# pyCancerSig: subclassifying human cancer with comprehensive single nucleotide, structural and microsatellite mutational signature deconstruction from whole genome sequencing

**DOI:** 10.1101/785410

**Authors:** Jessada Thutkawkorapin, Jesper Eisfeldt, Emma Tham, Daniel Nilsson

## Abstract

**Background:** DNA damage accumulates over the course of cancer development. The often-substantial amount of somatic mutations in cancer poses a challenge to traditional methods to characterize tumors based on driver mutations. However, advances in machine learning technology can take advantage of this substantial amount of data.

**Results:** We developed a command line interface python package, pyCancerSig, to perform sample profiling by integrating single nucleotide variation (SNV), structural variation (SV) and microsatellite instability (MSI) profiles into a unified profile. It also provides a command to decipher underlying cancer processes, employing an unsupervised learning technique, Non-negative Matrix Factorization, and a command to visualize the results. The package accepts common standard file formats (vcf, bam). The program was evaluated using a cohort of breast- and colorectal cancer from The Cancer Genome Atlas project (TCGA). The result showed that by integrating multiple mutations modes, the tool can correctly identify cases with known clear mutational signatures and can strengthen signatures in cases with unclear signal from an SNV-only profile.

**Conclusions:** pyCancerSig has demonstrated its capability in identifying known and unknown cancer processes, and at the same time, illuminates the association within and between the mutation modes.

## Introduction

Cancer is a genomic disorder, involving different kinds of DNA damage. DNA damage and imperfect repair occurs frequently in human cells. Changes accumulate over time, starting with our first cell, the fertilized egg, and progressively over the course of cell division [1]. Cellular proteins pertaining to replication, damage sensing and repair are important in limiting the damage. Most induced DNA variations will have no effect on the cell (so called passenger variants). However, over time there is a risk that accumulating DNA damage can result in tumorigenesis and later, metastasis, regardless of whether the initial driver mutation stems from inheritance, induced damage or imperfections in the replication of the hereditary material. The continued accumulation of DNA changes is caused by a combination of exposure to sources of DNA damage, both endogenous and exogenous and a broken DNA damage response mechanism. Each of these have their own respective profile with regard to the type of damage inflicted and bias in the repair mechanisms including double-strand DNA breaks [2], single-strand DNA breaks [3], and microsatellite instability [4].

Conventional methods to characterize tumor behavior are based on mutations in functional domains of cancer driver genes. Identifying the driver mutation is difficult, since the majority of the somatic events are passenger mutations [5]. An alternative is to classify tumors based on tumor behavior directly, by observing the pattern of somatic mutations. With an exploding amount of whole exome and whole genome sequencing data, together with advances in machine learning technology, a computational approach to characterize tumors based on their base-substitution profile was initiated, capturing mutation signatures [6]. These pioneering results were extended, showing e.g. how the mutational signatures associated with the presence of particular somatic driver mutations [7] and how a mutational signature enables classification of germline missense mutation [8]. However, base substitution is only one result from DNA damage. The others include copy number variation, structural rearrangement and microsatellite instability. Sensing both single nucleotide and structural variation (SV) has been shown efficient in predicting *BRCA1/BRCA*2 null mutation [9]. Tumor microsatellite instability (MSI) profiling has been proven useful clinically to inform testing for mutation in mismatch repair genes (MMR) [10] and more recently, to identify tumors eligible for immunotherapy [11, 12]. Tools to classify MSI status from tumor genome short read sequencing data [13] have been successfully developed.

Integrating several types of genetic aberrations from different types of DNA damage may not only reveal patterns and connections between these aberrations, but may also reveal novel combinations of cancer processes and possibly allow novel clinically actionable distinctions between tumors and inform interpretation of larger classes of hereditary variants of uncertain significance.

Most tools available for mutational signatures are implemented in R or as web interfaces, and require custom file formats [14]. Tools otherwise used in large scale processing of short read sequencing data in production environments are nearly exclusively command line based, and operate on a limited set of standardized file formats.

In this article, we describe pyCancerSig: an open source, command line interface python package for deciphering cancer signatures; integrating SNV, SV and MSI profiles in signatures decomposed using non-negative matrix factorization; and producing pdf reports.

## Materials and Methods

### Workflow processes

The workflow consists of 4 steps, data pre-processing, profiling, deciphering and visualizing (Figure 1, 2).

**Figure 1:**
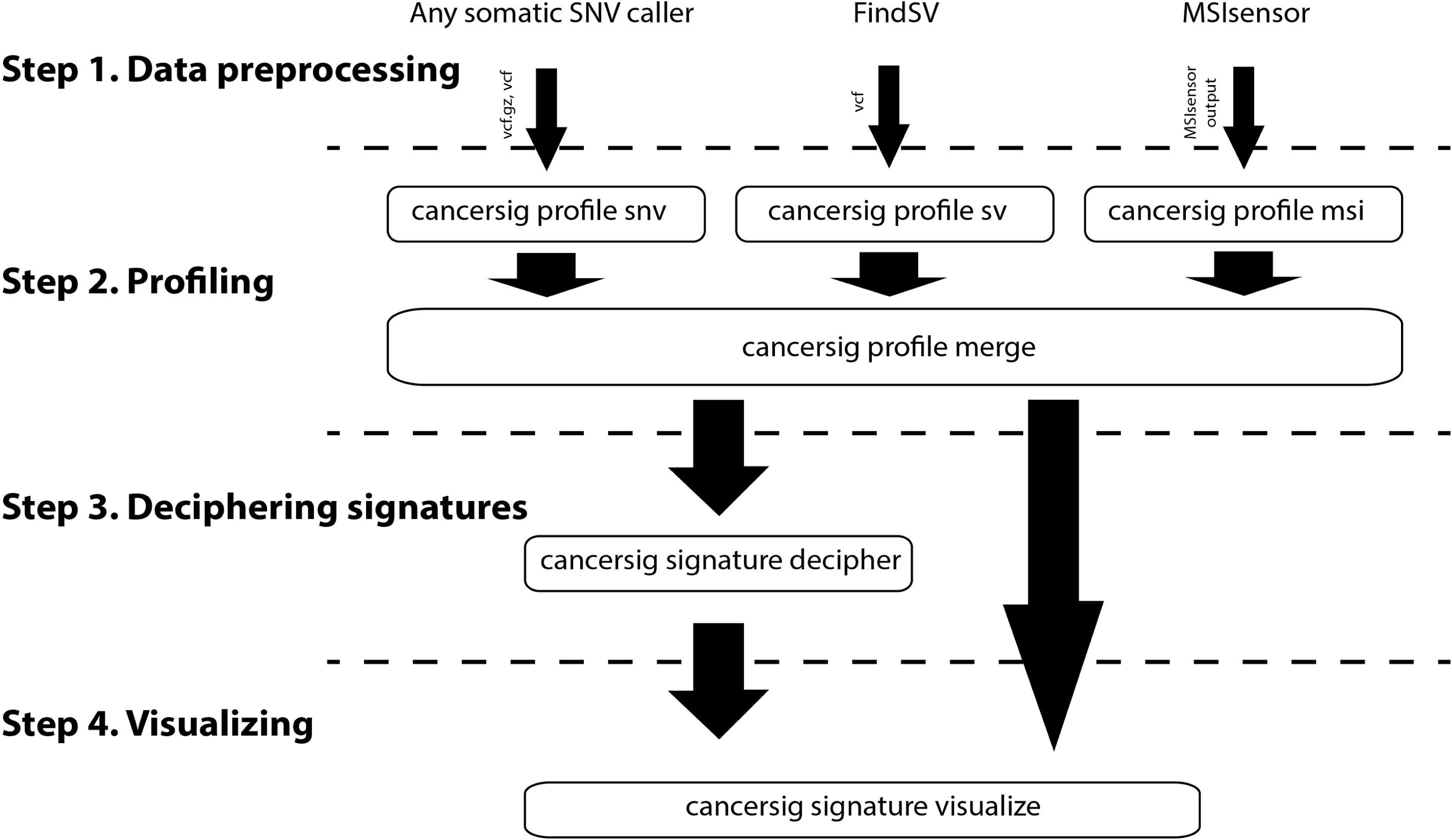
pyCancerSig workflow diagram. The workflow consists of 4 steps

1. Data preprocessing - The purpose of this step is to generate a list of variants. This step has to be performed by third party software.
  - Single nucleotide variant (SNV) - recommending MuTect2, otherwise Muse, VarScan2, or SomaticSniper.
  - Structural variant (SV) - dependency on FindSV
  - Microsatellite instability (MSI) - dependency on MSIsensor
2. Profiling (Feature extraction) - ‘cancersig profile’ - The purpose of this step is to turn information generated in the first step into matrix features usable by the model in the next step. The output of this stage has similar format as xshttps://cancer.sanger.ac.uk/cancergenome/assets/signatures_probabilities.txt, which consists of at least 3 columns.
  - Column 1, Variant type (Substitution Type in COSMIC)
  - Column 2, Variant subgroup (Trinucleotide in COSMIC)
  - Column 3, Feature ID (Somatic Mutation Type in COSMIC)
  - From column 4 onward, each column represents one sample
  There are subcommand to be used for each type of genetic variation
  -‘cancersig feature snv’ is for extraction single nucleotide variant feature
  - ‘cancersig feature sv’ is for extraction structural variant feature
  - ‘cancersig feature msi’ is for extraction microsatellite instability feature
  -‘cancersig feature merge’ is for merging all feature profiles into one single profile ready to be used by the next step
3. Deciphering mutational signatures - ‘cancersig signature decipher’ - The purpose of this step is to use unsupervised learning model to find mutational signature components in the tumors.
4. Visualizing profiles - ‘cancersig signature visualize’ - The purpose of this step is to visualize mutational signature component for each tumor.

**Figure 2:**
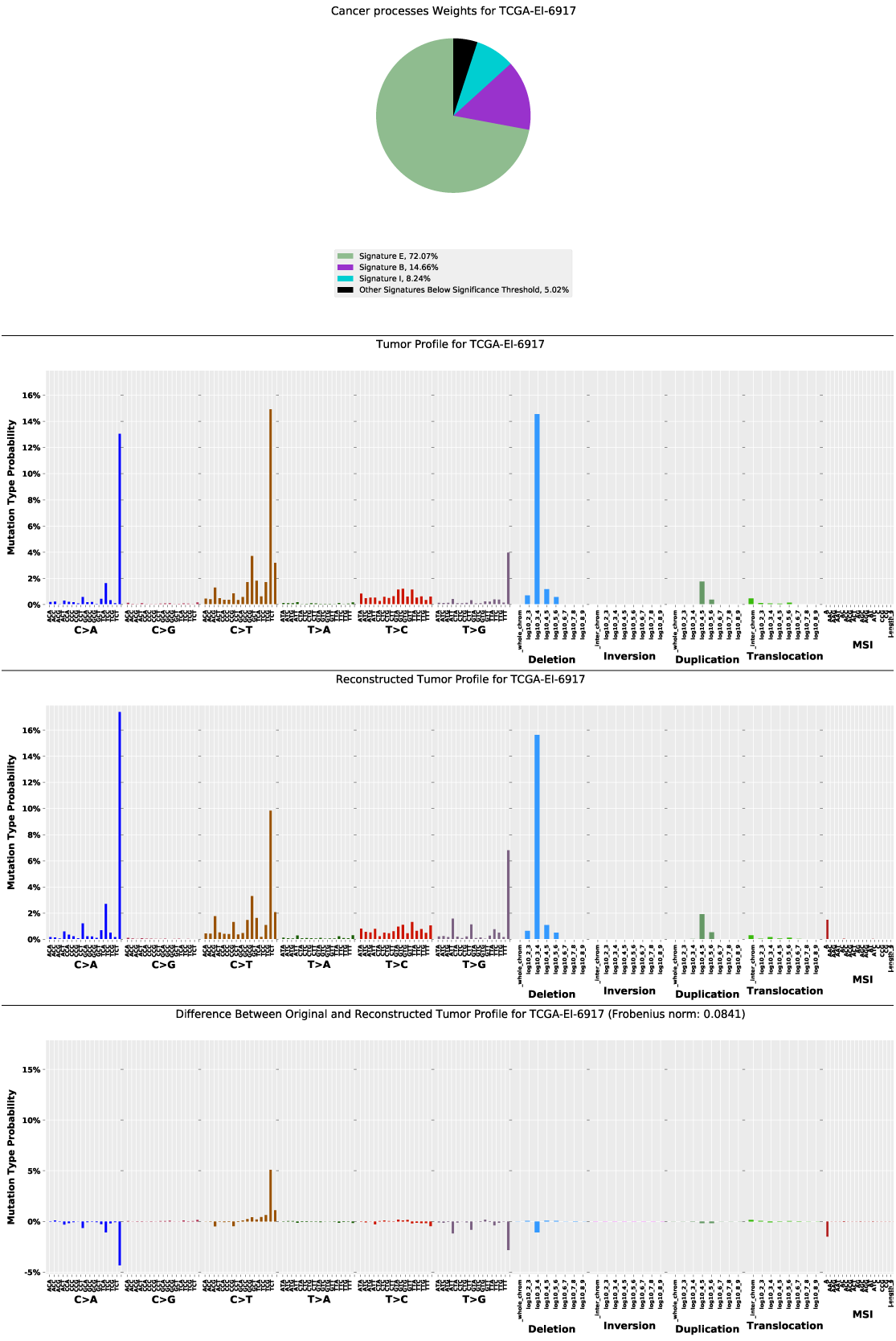
Example of a visualized tumor profile. The visualized profiles are generated in pdf format consisting of 4 pages. The first page is a pie chart representing mutational signature components in the profile. The second page displays the actual tumor profile. The third page displays the profile reconstructed from the predicted mutational signatures component in the first page. The last page displays the prediction error, which is the difference between the actual profile and the reconstructed profile.

#### Data pre-processing

The purpose of this step is to generate a list of variants. This step has to be performed by third party software. For SNVs, any somatic callers that produce variant calls in VCF format can be used. The same is true for SV calls, but even though the VCF standard has support for SVs, callers may not always be fully interchangeable. We have tested with, and recommend FindSV [15]. For MSI, the MSI profiling in the stage currently has a dependency on MSIsensor [13].

#### Profiling

The purpose of this step is to quantify information generated in the first step into matrix features usable by the deciphering process. There are three types of profiles, SNV, SV and MSI. SNV profiling scans the VCF file and quantifies six types of base substitution C:G > A:T, C:G > G:C, C:G > T:A, T:A > A:T, T:A > C:G, and T:A > G:C. Then, the changes are further subclassified by the sequence context, 5’ and 3’, of the mutation. In total, there are 96 mutation types (6 types of substitution x 4 types of 5’ base x 4 types of 3’ base).

SV profiling scans the VCF record for the INFO field SVTYPE to classify the mutation into structural duplication, structural deletion, inversion, and translocation. Then it checks another INFO field, SVLEN, for the length of the event, in order to subclassify the event based on its approximated size in log10 scale (size between 100-1Kbps, 1K-10K, 10K-100K, 100K-1M, 1M-10M, 10M-100M, 100M-1000M, and whole chromosome). In total, there were 32 mutation types (4 types of variation x 8 length groups).

MSI profiling quantifies all possible repeat patterns of a repeat unit with size between 1-3 nucleotides (A, C, AC, AG, AT, CG, AAC, AAG, AAT, ACC, ACG, ACT, AGC, AGG, ATC, CCG). For repeat unit sizes of 4 and 5 basepairs, it performs quantification without checking repeat patterns. The profile merger merges all sample profiles into one ready to be used by the deciphering- and visualization processes.

#### Deciphering

The purpose of this step is to use an unsupervised learning model to find mutational signature components in the tumors. The process was developed based on a well-established framework [6] and uses non-negative matrix factorization (NMF) to decipher signature probabilities, matrix P, from given input mutation profiles, matrix M, where M ≈ P × E. Matrix M represents fraction of each mutation type in each sample, each column for one sample and each row for one mutation type. Matrix P represents fraction of mutation type in each cancer process, each column for one cancer signature process and each row for one mutation type. Matrix E represents exposure, fraction of cancer signature process in each sample, each column for one sample and each row for one cancer signature process.

#### Visualizing

The purpose of this step is to decompose the tumor mutational signature components and visualize them together with mutation profiles of the tumor.

### Evaluation

#### Benchmarking data

A cohort of 92 breast cancer cases, TCGA-BRCA, and 38 colorectal cancer cases, TCGA-COAD and TCGA-READ, from The Cancer Genome Atlas project (TCGA) was used to evaluate the tool. The data consists of tumor-normal pair whole genome sequence in BAM format and SNV exome sequencing data called by MuTect2 [16] in VCF format (See Additional File 1: Benchmarking samples).

#### Preprocessing of the benchmarking data

For SNVs, the downloaded somatic VCF files called by MuTect2 were in a format ready to be used for SNV profiling in the next stage. For SVs, a singularity container implementation of FindSV was used with downloaded aligned bam files. Briefly, balanced structural rearrangements were called by TIDDIT [17]. Copy number variations were called by TIDDIT [17] and CNVnator [18]. Somatic aberrations were called by subtracting normal tissue events from tumor tissue events using SVDB [17]. For MSI, the instability loci were called by msisensor [13]from aligned bam files.

#### Sample labeling

Clinical information on the cases was provided by TCGA. Annotation of somatic SNVs in genes of interest (See Additional File 1: Pathway and related genes) was done using VEP [19]. ClinVar is a clinical database used for variant interpretation [20].

#### Profile types and evaluation processes

We evaluated the tool using three profile types: SNV-only profile, SNV+SV+MSI profile and SV-only profile. The purpose of the SNV-only profile was to verify the SNV profiling- and the visualization processes by comparing the results with [21], and to setup a baseline for the other two comparisons. The combined, SNV+SV+MSI, profile was used to evaluate the efficiency of the profile when integrating several groups of mutations. The SV-only profile was used to evaluate the use of structural variation as the sole input compared to the combined profile.

#### Signature probabilities and signature IDs

There are three groups of mutational signatures described in this article, one for each profile type. The signature probabilities used in the evaluation of SNV-only profile were downloaded from [22]. Signature IDs are represented as running numbers, “signature 1”, “signature 2”, “signature 3”, etc., the same as the ones in the downloaded signature probabilities file and the ones in [21]. For the combined profile, the signature probabilities were generated from the deciphering process. The signature IDs are represented using italic upper case English alphabets, “signature *A*”, “signature *B*”, “signature *C*”, etc. For the SV-only profile, the signature probabilities were also generated from the deciphering process. The signature IDs are represented using italic lower case English alphabets, “signature *a*”, “signature *b*”, “signature *c*”, etc.

### Sample-signature association

If a sample shows a decomposed fraction of more than 30% of a particular signature cancer process, it is defined as having an association between the signature and the sample.

### Tumor mutation burden (TMB)

TMB was quantified as the total number of base substitutions in the SNV profile divided by 30 to give an estimate of SNVs per Mbps sequence.

### Structural variation burden (SVB)

SVB was quantified as the total number of somatic structural variations in the sample, as determined from the SV profile.

## Results

We used pyCancerSig to profile the benchmark data on three profile types, SNV-only, SNV+SV+MSI (combined), and SV-only. The SNV-only signatures used in the package verification were from COSMIC Mutational Signatures (v2 - March 2015) [23]. pyCancerSig was used to decipher the combined- and the SV-only profile, resulting in nine (See Additional File 2: Figure 1) and twelve signatures (See Additional File 2: Figure 2) respectively. Tumor profiles were visualized for each profile type, together with their mutational signature components (Additional File 3-5).

### Verification using the SNV-only profile

#### *POLE* gene

Using SNV profiling by pyCancerSig, five cases were found to have signature 10, which has altered activity of the mutated *POLE* as proposed etiology. All of these cases were found to have somatic mutation in *POLE* and *POLD1* (Additional File 6). Moreover, these cases were found to have hypermutation with a median of 881 SNVs/Mbp (range 667-1686/Mbp), compared to the median 36/Mbp in the total data set and 22/Mbp in the tumors without *POLE/D1* variants. (Figure 3), another property of tumors with bi-allelic loss-of-function variants in *POLE* [24].

**Figure 3:**
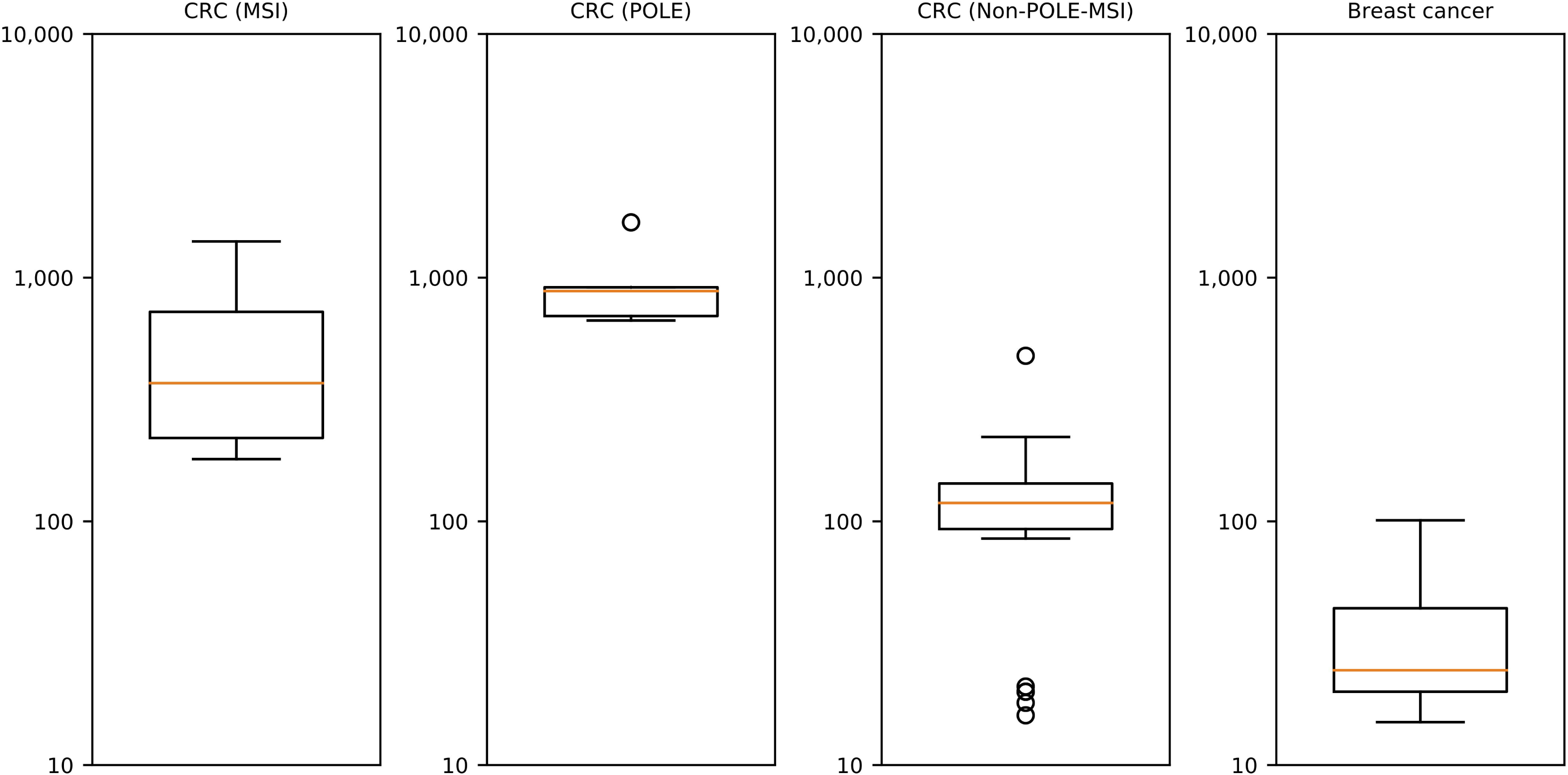
Boxplot of tumor mutation burden of 4 sub-populations evaluated in this article. The y-axis represents tumor mutation burden (TMB), which is quantified as the total number of base substitutions in the SNV profile divided by 30 to give an estimate of SNVs per Mbps sequence. The box represents values from the lower to the upper quartile. The yellow line represents the median.

#### DNA mismatch repair (MMR) genes

Signatures 6, 15, 20 and 26 were described to be associated with defective DNA mismatch repair. None of the cases in our cohort were associated with signature 15, 20 or 26. Using SNV profiling by pyCancerSig, two cases: TCGA-A6-6781 and TCGA-AA-A01R, were found to be associated with signature 6. Both of them were identified to be MSI-positive by MSIsensor. Both cases harbored pathogenic germline (and in one case also somatic) truncating variants in *MSH2* or *MSH6* (Additional File 6).

#### Double strand break pathway

Signature 3 was described to be associated with failure of DNA double-strand break (DSB) repair by homologous recombination. Using SNV profiling by pyCancerSig, 78 breast cancer– and 10 colorectal cancer cases were found to be associated with signature 3. Among these, eleven breast cancer cases and one colorectal case had pathogenic truncating variants in DSB-pathway genes. Four cases have truncating variants in *BRCA1*, p.Val1734Ter, p.Glu23ValfsTer17 (both germline) and p.Glu720Ter, c.4739-1G>A (both somatic). One case has a somatic truncating variant in *BRCA2*, c.7805+1G>A. One breast cancer case had bi-allelic truncating variants in *BARD1*. The rest had truncating variants in *BLM, ATM* and *ATRX*. In addition, 30 other cases (27 breast cancer and 3 CRC) had in-frame or missense variants in 22 DSB genes (Additional File 6).

### Evaluation using the combined profile

The optimal number of mutational signatures for the combined profile suggested by pyCancersig was nine (See Additional File 2: Figure 1). Signatures *E* and *I* were only associated with colorectal cancer cases, while signatures *C, D, G* and *H* were only associated with breast cancer cases and signatures. *A, B* and *F* were found to be associated with both cancer types (Additional File 6).

#### *POLE* and *POLD1* genes

Signature E exhibited a pattern of TCT > TAT, TCG > TTG and TTT > TGT, similar to those of signature 10, together with a pattern of structural deletion with size between 1K-10K. Signature E was associated with five cases and four of them were associated with signature 10. Three of these had known mutations in *POLE* and the fourth had a somatic truncating variant in *POLE* and a high TMB of 1686/Mbps. The fifth case with signature E had 26% signature 10, just below the 30% threshold applied in this paper and had a known mutation in *POLE* (p.Ser297Phe) as well as a high TMB (881/Mbps).

#### MMR genes

Signature *I* was the only signature with a pattern of MSI identified by msisensor [13]. It exhibited specific dominant characteristics in all mutation modes. SNVs were dominated by C > T substitutions, particularly on <u>C</u>G > <u>T</u>G backgrounds. The majority of SV events were structural deletion of 1K-10K. Signature *I* was associated with four colorectal cancer cases, two of which were the two cases associated with signature 6.

However, there were 2 other cases, TCGA-AD-6964 and TCGA-AZ-6601 with signature *I*. TCGA-AZ-6601 had both a germline truncating variant in *MSH6* (p.Lys537Ilefs*33) and a somatic in frame change in *MSH6* suggesting a mismatch repair defect, while no pathogenic variants were found in TCGA-AD-6964. (Additional File 6).

#### Double strand break pathway

From the aforementioned eleven breast cancer cases and one colorectal cancer case with signature 3 and pathogenic or likely-pathogenic variants in the DSB pathway, eleven were associated with seven different combined signatures (*A, B, C, D, F, G* and *H*) and one was not associated with any signatures. The median SV burden in these eleven cases (1004 events) did not differ significantly from the median of the entire cohort (Additional File 6).

#### Structural variation burden

The 10 cancer cases with highest SVB have a median of 10,016 events compared to the median 885 events in the total cohort. All ten were breast cancer samples and were divided between four signatures (*A, C, D* and *F*). (Additional File 6). To compare with the SNV-only profile, all of these cases, except TCGA-B6-A0IJ, were associated with signature 3. TCGA-B6-A0IJ was not associated with any signature using the SNV-only profile.

### Evaluation using the SV-only profile

The optimal number of mutational signatures for the combined profile suggested by pyCancerSig were 12 (See Additional File 2: Figure 2). Signatures *c, e, f, g* and *l* were associated with only breast cancer, while signatures *a, b* and *d* were associated with both cancer types.

#### *POLE, POLD1* and MMR genes

Signature *a* exhibited a very unique pattern. 96% of the mutation in this signature was structural deletion with size between 1K-10K. This pattern was found in signature *E* and *I*. Eleven breast cancer- and fifteen colorectal cancer cases were identified to have this signature. Among them, five cases were the ones with combined signature *E* and four cases were the ones with combined signature *I*.

#### Double strand break

Out of the aforementioned eleven breast cancer cases and one colorectal cancer case with signature 3 and pathogenic or likely-pathogenic variants in the DSB pathway, four were associated with SV-only signatures: *c, d* and *e*. The colorectal cancer case with an *ATM* mutation was associated with signature *a*. One thing in common between these four signatures was that they each represented one specific structural event (more than 95%), signature *a* for structural deletion 1K-10K, signature *c* for structural inversion of size 1K-10K, signature *d* for structural deletion of size 10K-100K and signature *e* for structural duplication of size 10K-100K. The rest were not associated with any signatures (Additional File 6).

#### Structural variation burden (SVB)

The 10 cancer cases with the highest SVB were all breast cancers and were mainly associated with two SV-only signatures: *c* and *d*. Both of these had one main SV event at high frequency. (Additional File 6).

## Discussion

In the present study, a command line tool was developed and tested on two different cancer types. We report a working tool, and study the use of a combined SV, SNV and MSI profile, contrasting to SNV or SV only profiles.

### Comparison between signatures generated by the SNV-only profile and signatures generated by the combined profile

This study has shown that combining SNV, SV and MSI into one single profile can correctly identify mutational signature processes caused by altered activity of the DNA polymerases *POLE* and *POLD1*: signature *E*, and loss-of-function in MMR genes: signature *I*. Moreover, beside backward compatibility to SNV-only signature 6, signature *I* generated by the combined profile has identified two additional cases, TCGA-AD-6964 and TCGA-AZ-6601, with limited association to signature 6 (23% and 18%) which were also MSI positive according to msisensor. In principle, it could be possible to reduce the required level of association to signature 6 in order to increase the sensitivity of the SNV-only profile. However, this also reduces the specificity. In total, 10 colorectal cancer and 20 breast cancer cases displayed a contribution from signature 6, ranging from weight 6% to 42% of the total mutational profile. Four of them had weights in the range 20-30%, including one MSI and three MSS cases. Integrating SNV, SV and MSI into one single profile significantly increased the strength of the signal (signature *I*) due to a specific SV pattern (structural deletion with size between 1K-10K) and MSI.

COSMIC signature 3 is generally described as associated to deficiency in homologous recombination repair of double stranded breaks [25]. SNV-only profiles categorized 78 of 92 (85%) breast cancer cases as having contributions from signature 3. Thus, this signature very likely appears in cancers with heterogeneous etiologies. The eleven breast cancer samples with signature 3 and a defect DSB pathway due to truncating mutations in DSB genes did not cluster together in any specific combined or SV-only signature, thus adding SV information did not identify a DSB-specific pattern. In fact, the four *BRCA1/2*-mutated samples displayed a nonspecific SV pattern and did not contain a high number of SV events. This is in contrast to previous work where *BRCA1/2* deficient structural variation-only profiles were detected as clustered events [9, 26, 27]. The present tool does not differentiate between clustered and non-clustered variants and indeed the SV callers used are not designed to detect such clusters, being essentially overlapping structural events.

### Comparing SV-only signatures to combined signatures

Whether to use a combined or SV-only signature depends on the problem at hand. Combining an informative profile with a non-informative one may weaken the signal, depending on the amount of training data available. On the other hand, combining two informative profiles of two different mutation modes will increase accuracy, as in the cases with MMR gene mutation.

When analyzing SVB, a clear difference between cases with high SVB and low SVB (See Additional File 7: Figure 1) can be seen. The high-SVB cases each have at least one dominant structural variant type, accounting for more than 30% (Additional File 7: Figure 2), which is never present in any of the low SVB cases. The ten high SVB samples clustered in two different SV-only profiles, each showing one single dominant structural variant type. However, when SNV data was added, the SV signal was weakened with two samples clustering in signatures that lacked SV specific patterns. In this scenario, interpretation is simplified by the SV-only model.

On the contrary, it would be disadvantageous to use an SV-only profile alone to identify tumors with *POLD1*/*E*-defect or loss of function in MMR gene. The analyzed colorectal cancer cases with pathogenic mutation in *POLE, POLD1* and *MSI* were all identified to have signature *a*, which was characterized by structural deletion with size between 1K-10K. However, signature *a* was not specific to only these cases. There were eleven breast cancer and six colorectal cancer cases with signature *a* which did not have any coding *POLD1/E* or MMR variants. This suggested a possible advantage of combining two informative mutational modes in increasing accuracy.

### Tumor mutation burden (TMB) as a profile component

Previous work has advised that TMB be used for tumor classification, genetic testing, and clinical trial design [28]. Including TMB as part of a molecular profile may strengthen accuracy. However, we have not included TMB for two reasons. First, different bioinformatic workflows can result in different TMB values [29]. A standardized workflow is needed to ensure that the same cut-off value can be used across platforms and bioinformatic workflows. Second, TMB analysis compares the number of mutations in one tumor to the others. But signature analysis compares relative incidences of mutation types within a tumor, resulting in a proportion of mutation types. Research on how TMB can be included in a form of a proportion is needed to ensure that TMB of hypermutated tumors does not overpower other mutation types.

### Signatures representing unknown biological mechanisms

Mathematically, NMF results represent sets of variables that capture most of the variation in the data. Hence, the signatures represent the underlying biological processes of cancer only when the captured variations are the result of the oncogenic mechanism in the analyzed tumor. The types of variation seen in dysfunction of one mechanism do not necessarily aid explanation of a dysfunction in another mechanism. Increasing sample numbers and number of cancer types under study does aid discovering more underlying signatures. As the study progressed to include also CRC samples, additional signatures were indeed discovered. Application to a larger and more diverse data set was, however, outside the scope of the current study.

It is worth noting that at least half of the COSMIC signatures have yet to be ascribed an underlying process. Adding sensors to detect variations caused by different cancer mechanisms will make it possible for the signatures to closer represent the underlying biological processes. Many different sensors of DNA change could be incorporated, and have indeed been used in different studies: multi nucleotide variants, small indels, clustered/non-clustered SVs, breakpoint characterization sensors, including microhomology and mutations clustered around the breakpoints. Having more quantitative information of omics data will strengthen the power of the model. Other omics data that can be included in the model include methylation data, RNA expression data and protein data. Depending on the amount of correlation between different sensor variables, signatures may be useful biomarkers even if the variant type repertoire is far from complete.

### Implementation

While it is being increasingly recognized that mutational signatures are useful for the analysis of tumors, they are not yet in use in many sequencing centers. We believe this is partly due to the lack of a modern, command line implementation of any such tools. Most available tools operate in the R environment, well suited for prototyping of mathematical methods, or by web interface. Many expect data in domain custom or rare formats. Sequencing facilities operate with command line interface tools that can be readily wrapped into pipelines, and a small number of standardized file formats. One notable exception is EMu [30] which also presents a command line interface. The EMu tool accepts copy number variation data, but relies on custom file formats for data input. It does not handle balanced structural variation or MSI. We opted for an implementation in python, with a command line interface accepting standard genomics file formats (vcf, bam) to be suitable for integration into standard pipelines. pyCancerSig also produces pdf reports. pyCancerSig requires whole genome sequencing input of both tumor and germline DNA in bam and VCF format. pyCancerSig optionally accepts COSMIC/WTSI format signatures. No particular constraint is made on SNV or SV VCF input, but SNV calls compatible with data used for Alexandrov et al [6] are accommodated, and the structural variant calling pipeline (FindSV) used in the present study is also documented for ease of use and reproduction.

### Choice of feature extraction model

Developing a feature extraction method that would reasonably represent the global mutation profile was challenging. The mathematical model, NMF, used in this study took a matrix M as an input and returned matrix P, where M ≈ P × E. The model treated every number in the matrix equally, regardless of the type of variation represented.

Relative differences in the number of mutation events in each mutation group played an important role in choosing the extraction method. It is known that mutation burden varies between different cancer types [31]. The number of somatic indels are significantly different between MSI and non-MSI cases [13]. In this study, we decided to limit the total percentage of features from each mutation group. For instance, the total fraction of base-substitution will always be 70% regardless of the relative difference between the number of base substitution events compared to copy number variation events. This prevented a very large number of mutations from one mutation group, like hypermutation due to defect MMR or the enormous amount of structural inversion events found in cases with signature *D* to overpower the others.

The numerical encoding of each type of structural event sensor variable also had an impact on the design. On the surface, the events might look equivalent. In detail, the quantitative values encode different molecular events. In structural deletion or structural duplication, the size of an event was the total number of nucleotides lost or gained. But for translocation events, the size of an event was a distance between the original position and the new position. If a piece of 100 bp DNA is translocated from a coordinate at 20 Mb to a coordinate at 25 Mb, in the same chromosome, the size of this event will be 5 Mb, instead of 100 bp. If the event is to a different chromosome, the size will be set to infinite. Moreover, there were also differences behind the quantitative value within each mutation type. For example, larger structural deletion events are not as numerous as their smaller counterparts. It is common to see 100 events of structural deletion with size between 100-1000bp but it is impossible to see the same number of events for whole chromosome deletions. In fact, having only 1 or 2 whole chromosome deletion events is a rare and pathogenic event. At first, we attempted a logarithmic scale for the quantitative values of structural events. However, in order to make SVs comparable with the base substitution profile and in order to keep the profile representation as a percentage (in line with previous publications [6], the present study does not use logarithmic values. This resulted in a risk that certain mutation types can overpower the others within the group, like in cases with signature *D*. Moreover, events that are very rare but pathologic were also underpowered by this choice.

### Usage of tumor profile

Cancer signatures can be applied in both supervised- and/or unsupervised learning models. Using cancer signatures in a supervised learning model can result in a high accuracy classifier, which is very efficient in answering questions with a limited set of answers. One such classifier is HRDetect [9], which can identify *BRCA1/BRCA2*-deficient tumors with 98.7% sensitivity.

However, the implementation in this study was in an unsupervised fashion. This means that the cases have not been labelled, including cancer type, cancer-related pathway, cancer-related genes or tumor heterogeneity prior to incorporation in the model. As a result, the model has brought out the patterns of profile that were commonly found in the cases, regardless of whether there were one or more cancer processes in the tumors, whether the processes are in known or unknown pathways, whether the processes are in known or unknown genes, or whether the damage was caused by one or more sources of DNA damage.

The unsupervised method aids interpretation by reducing the dimensionality of the data and presenting it in a way that humans can recognize. Therefore, this model can be used to support molecular classification of tumors; for instance, to compare a tumor to previously analyzed ones in order to delineate possible cancer processes. Given that the model has been trained with enough data, they can be used as supportive evidence in a clinic by associating a new unsolved case with signatures from solved cases, and perhaps in the future aid medical decisions regarding personalized therapy.

One way to make use of visualization of this clustering method is to use it as an intermediate result to infer the mechanism behind the tumor development and to suggest potential targets for therapy based on the suggested profile. For example, in MSI cases, if we profile a tumor and its profile is similar to sample TCGA-A6-6781 or TCGA-AA-A01R, the profile can be a supporting evidence suggesting a similar causative mechanism, in this case defect MMR genes. Also, since the profiles of the MSI cases suggest similar molecular mechanism caused by the mutation, this might be used as evidence that similar treatment on cases with similar profile is likely to have similar effect. Thus using combined mutation profiles may identify groups of cancers that may profit from specific therapies as has been previously suggested [32].

## Conclusions

This pattern-based study has shown that applying an unsupervised learning method on joint information from multiple variant types can identify the cases suggested by the SNV-only profile and demonstrated that integrating SV and MSI into the profile can strengthen previously unclear signals. This illuminated the association within and between the mutation modes. The tool presented, pyCancerSig, accepts common file formats, and is easily incorporated into a genome center pipeline.

## Availability and requirements

Project name: pyCancerSig

Project home page: https://github.com/jessada/pyCancerSig

Operating system(s): Platform independent.

Programming language: Python

Other requirements: Python 3.0 or higher

License: GNU v.3.0.

Any restrictions to use by non-academics: None.

## List of abbreviations

DSB: double-strand break
Mbp: mega basepair
MMR: mismatch repair genes
MSI: microsatellite instability
MSS: microsatellite Stable
NMF: non-negative matrix factorization
SNV: single nucleotide variation
SV: structural variation
SVB: structural variation burden
TCGA: The Cancer Genome Atlas project
TMB: tumor mutation burden

## Declaration

### Ethics approval and consent to participate

This study utilized previously obtained TCGA data, made available by dbGap after ethical review of our dbGap project #14084; and used under approval of this request [#55693-2]

### Consent for publication

Not applicable

### Availability of data and materials

The datasets used for evaluation of this software are available at https://portal.gdc.cancer.gov.

### Competing interests

The authors declare that they have no competing interests

### Funding

This study was supported by grants provided by the Stockholm County Council (ALF project) and Swedish cancer society.

### Authors’ contributions

JT applied for computational resources, gathered data, developed and evaluated the software, helped design the study and was a major contributor in writing the manuscript. JE was involved in structural variant analysis. ET performed pathogenicity interpretation of cancer-related genes and was involved in writing the manuscript. DN designed the study and was involved in designing software architecture and command line interface, and was involved in writing the manuscript. All authors approved of the final manuscript.

## Acknowledgements

We thank Annika Lindblom for help in applying for access to dbGaP TCGA data.

## Notes

https://portal.gdc.cancer.gov

