## Additional File 2 for "pyCancerSig: subclassifying human cancer with comprehensive single nucleotide, structural and microsatellite mutational signature deconstruction from whole genome sequencing"

### Figure 1: 9 signatures deciphered from the combined profiles

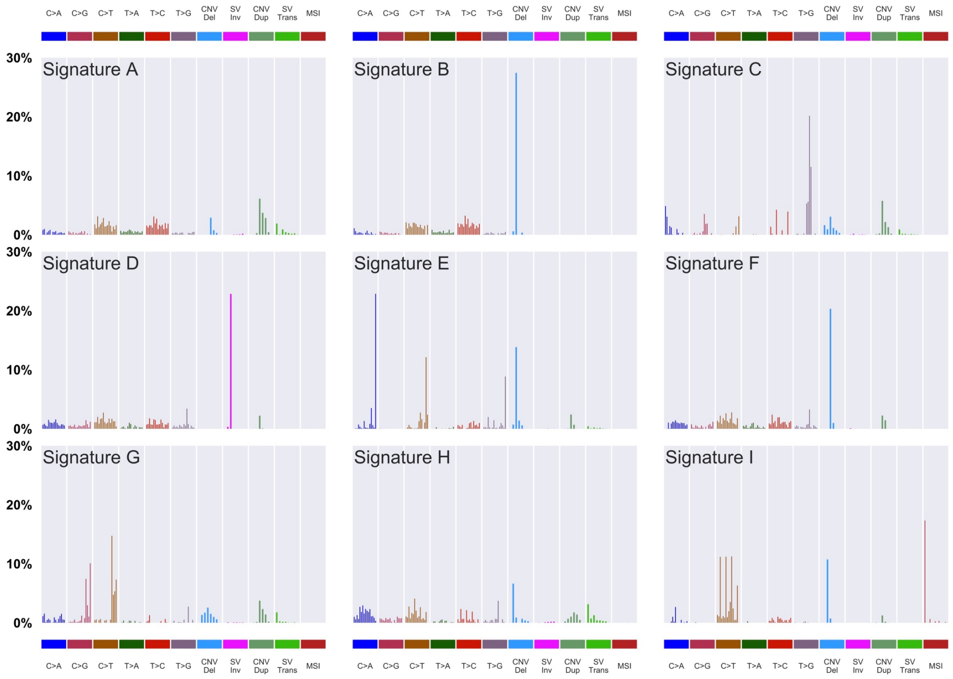


**Signature *A***

The majority of the base substitutions in this signature were C > T and T > C mutations. There was a high percentage of structural deletion with size between 100K-1M, structural duplication with size between 10K-10M, and interchromosomal translocation. 22 breast cancer and 25 colorectal cancer cases were associated with this signature (Additional File 6).

**Signature *B***

The most outstanding mutation type in this signature is structural deletion with size between 1K-10K. Five breast cancer and 10 colorectal cancer cases were associated with this signature (Additional File 6).

**Signature *C***

This signature exhibited a pattern of GT > GG mutations, especially at GTG > GGG and GTT > GGT context. This signature was associated only with breast cancer (28 cases) (Additional File 6).

**Signature *D***

The most salient mutation type in this signature is structural inversion with size between 1K-10K. Five breast cancer cases were associated with this signature (Additional File 6).

**Signature *E***

This signature exhibited a pattern of TCT > TAT, TCG > TTG, TTT > TGT and structural deletion with size between 1K-10K. It was associated only with colorectal cancer, five cases in total (Additional File 6).

**Signature *F***

Signature F was characterized by structural inversion with size between 10K-100K. Nine breast cancer and two colorectal cancer cases were associated with this signature (Additional File 6).

**Signature *G***

This signature exhibited a pattern of TC > TG and TC > TT mutations, particularly on TCA and TCT backgrounds. This signature was associated with five breast cancer cases (Additional File 6).

**Signature *H***

There was no single common mutation type in this signature. The distribution of SNVs was slightly biased toward C > A and C > T. Small structural deletions, size 100-1Kbps, and inter-chromosomal translocations dominated slightly compared to other structural events. Only breast cancers (29 cases) were associated with signature H (Additional File 6).

**Signature *I***

Signature I exhibited specific dominant characteristics in all mutation modes. In SNVs were dominated by C > T substitutions, particularly on CG > TG backgrounds. The majority of SV events were structural deletion of 1K-10K. And it was the only signature with MSI. This signature was associated with four colorectal cancer (Additional File 6).

### Figure2: 12 signatures deciphered from the SV-only profile

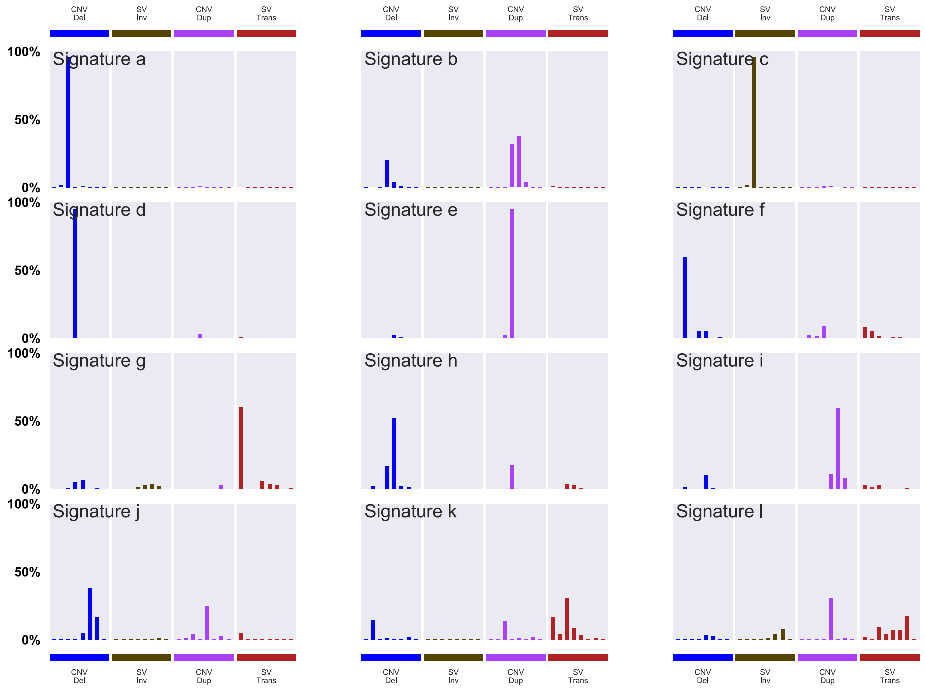


**Signature *a***

This signature almost only represented structural deletion of size 1K-10K. There were 11 breast cancer- and 15 colorectal cancer cases associated with this signature (Additional File 6).

**Signature *b***

This signature was dominated by structural duplication and structural deletion of size 10K-100K. There were 7 breast cancer- and 3 colorectal cancer cases associated with this signature (Additional File 6).

**Signature *c***

This signature almost only represented structural inversion of size 1K-10K. This signature was associated only with breast cancer, 7 cases in total (Additional File 6).

**Signature *d***

This signature almost only represented structural deletion of size 10K-100K. There were nine breast cancer and one colorectal cancer cases associated with this signature (Additional File 6).

**Signature *e***

This signature almost only represented structural duplication of size 10K-100K. There was only one breast cancer case, TCGA-A2-A04Q, associated with this signature (Additional File 6).

**Signature *f***

This signature was dominated by structural deletion, mainly with size 100-1Kbps. This signature was associated only with breast cancer, nine cases (Additional File 6).

**Signature *g***

Signature G exhibited inter-chromosomal translocation. There was only one breast cancer case, associated with this signature (Additional File 6).

**Signature *h***

This signature was dominated by structural duplication and structural deletion of size 10K-100K. None of the cases was associated with this signature (Additional File 6).

**Signature *i***

This signature exhibited structural duplication with size 1M-10M. None of the cases was associated with this signature (Additional File 6).

**Signature *j***

This signature was dominated by large structural deletion and structural duplication, size between 100K-10M. None of the cases was associated with this signature (Additional File 6).

**Signature *k***

This signature exhibited a combined pattern of structural deletion, structural duplication and inter- and intra-chromosomal translocation. None of the cases was associated with this signature (Additional File 6).

**Signature *l***

This signature was dominated by structural duplication of size 100K-1M and chromosomal translocation with distance 1K-100M. There was only one breast cancer case associated with this signature (Additional File 6).
