## Additional File 3 for "pyCancerSig: subclassifying human cancer with comprehensive single nucleotide, structural and microsatellite mutational signature deconstruction from whole genome sequencing"

### Cancer processes Weights for TCGA-E2-A14X

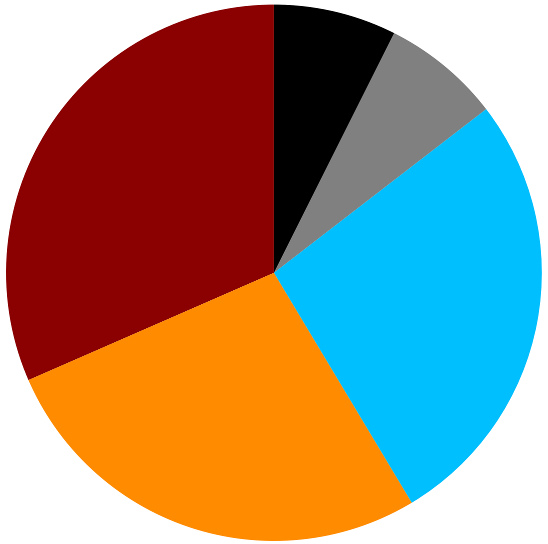

Signature 3, 31.55%
Signature 1, 27.05%
Signature 12, 26.85%
Signature 24, 7.15%
Other Signatures Below Significance Threshold, 7.4%

Tumor Profile for TCGA-E2-A14X

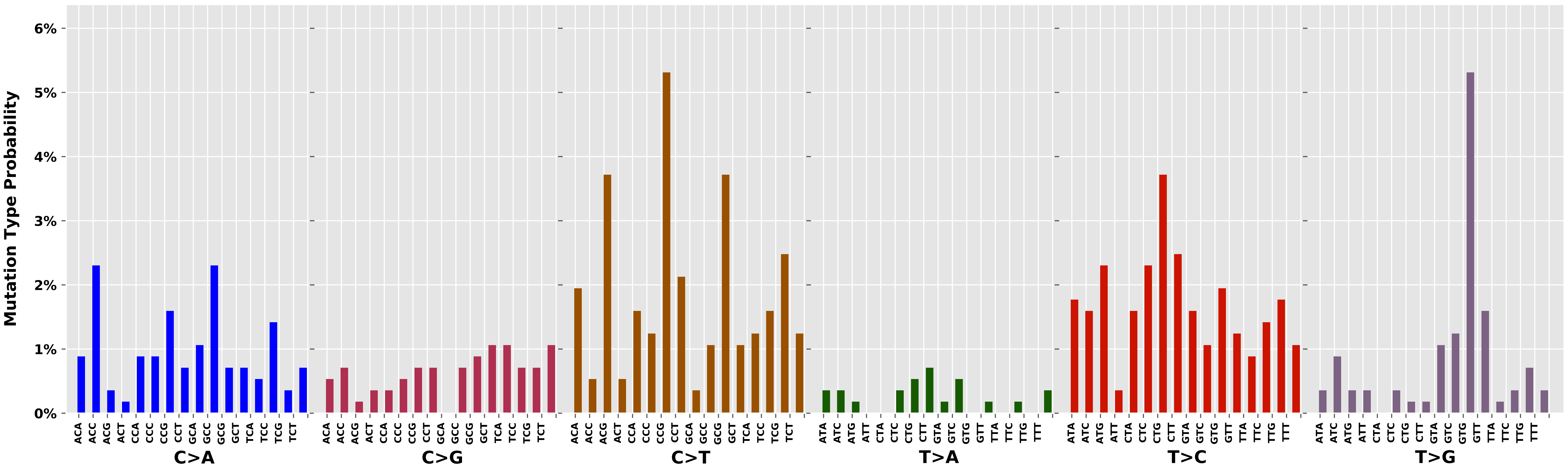

Cancer processes Weights for TCGA-A8-A08L

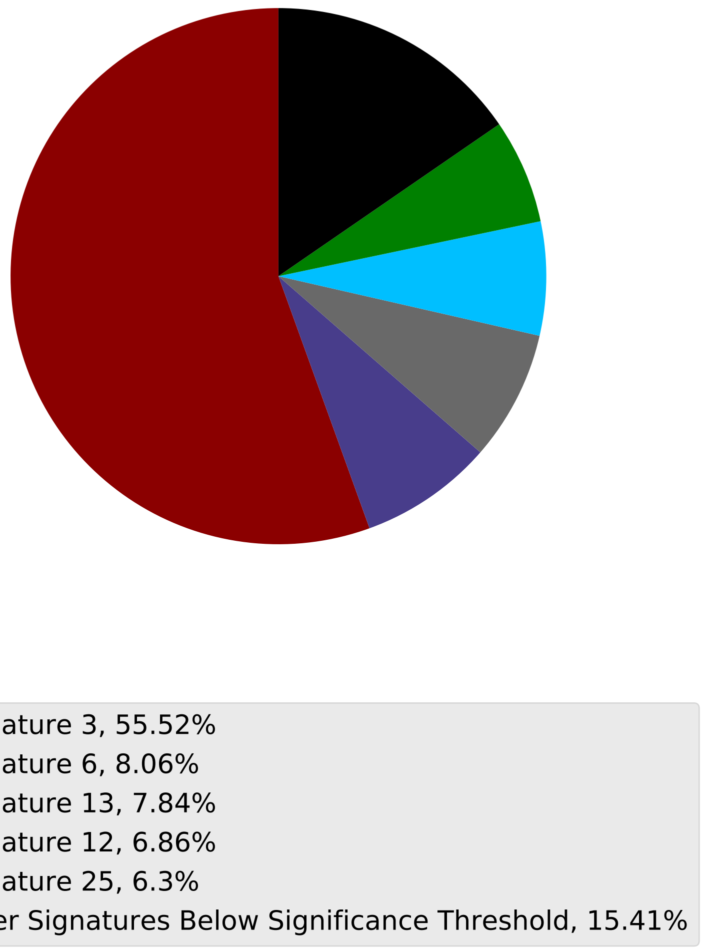

Tumor Profile for TCGA-A8-A08L

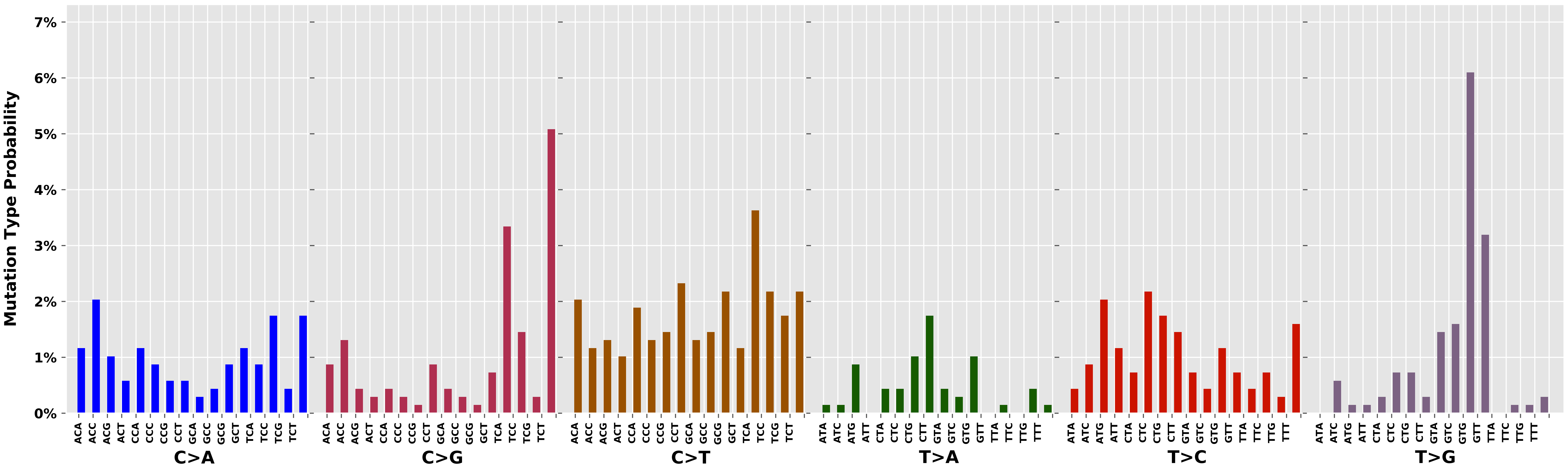

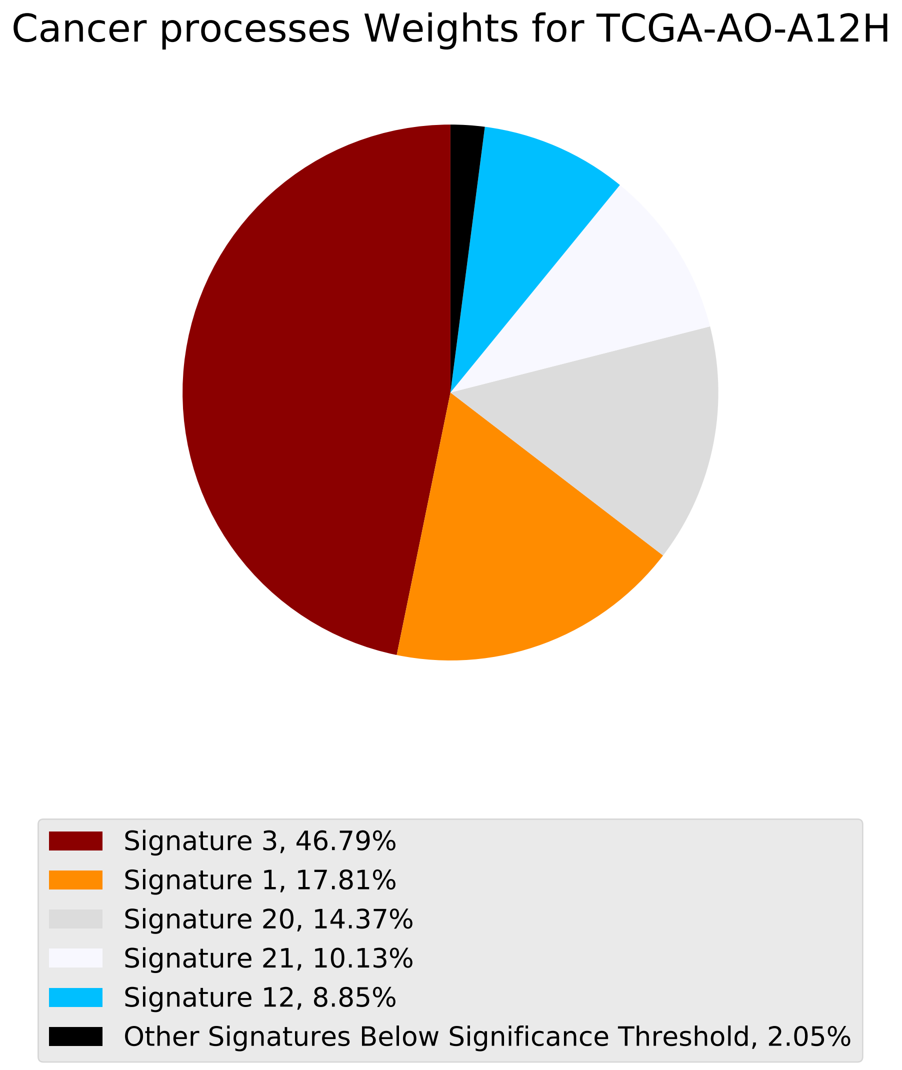
Cancer processes Weights for TCGA-AO-A12H
Signature 3, 46.79%
Signature 1, 17.81%
Signature 20, 14.37%
Signature 21, 10.13%
Signature 12, 8.85%
Other Signatures Below Significance Threshold, 2.05%

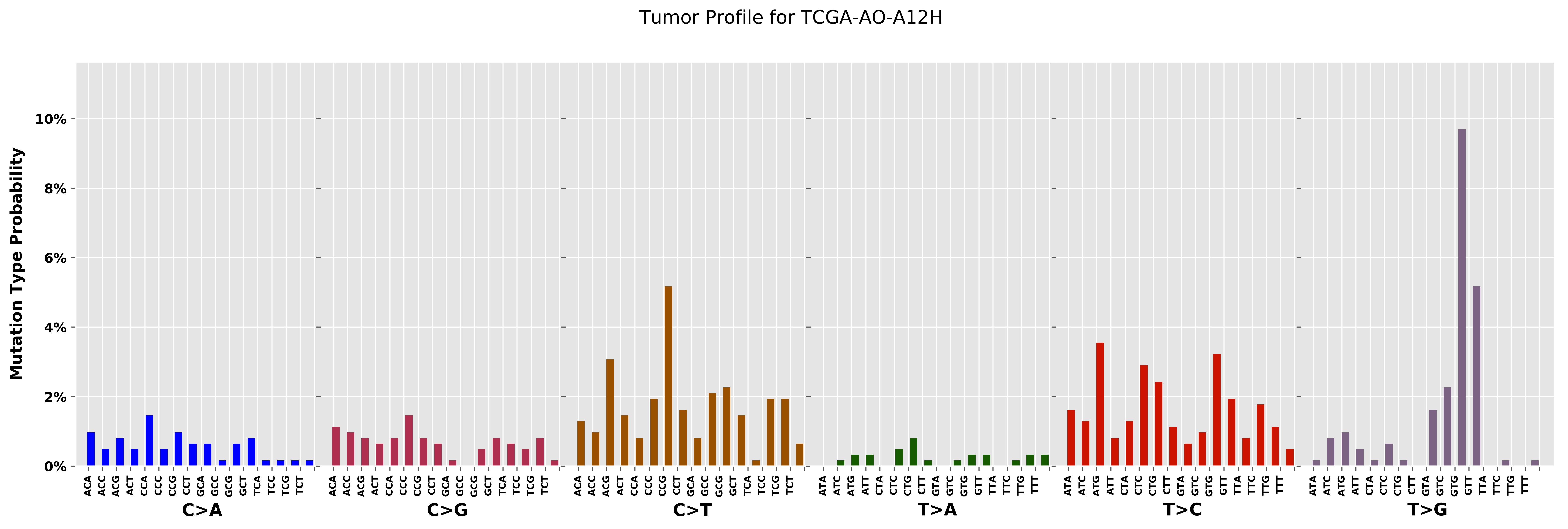

Tumor Profile for TCGA-AO-A12H
Mutation Type Probability
0%
2%
4%
6%
8%
10%
C>A
C>G
C>T
T>A
T>C
T>G
ACA ACC ACG ACT CCA CCC CCG CCT GCA GCC GCG GCT TCA TCC TCG TCT
ATA ATC ATG ATT CTA CTC CTG CTT GTA GTC GTG GTT TTA TTC TTG TTT

Cancer processes Weights for TCGA-EW-A3U0

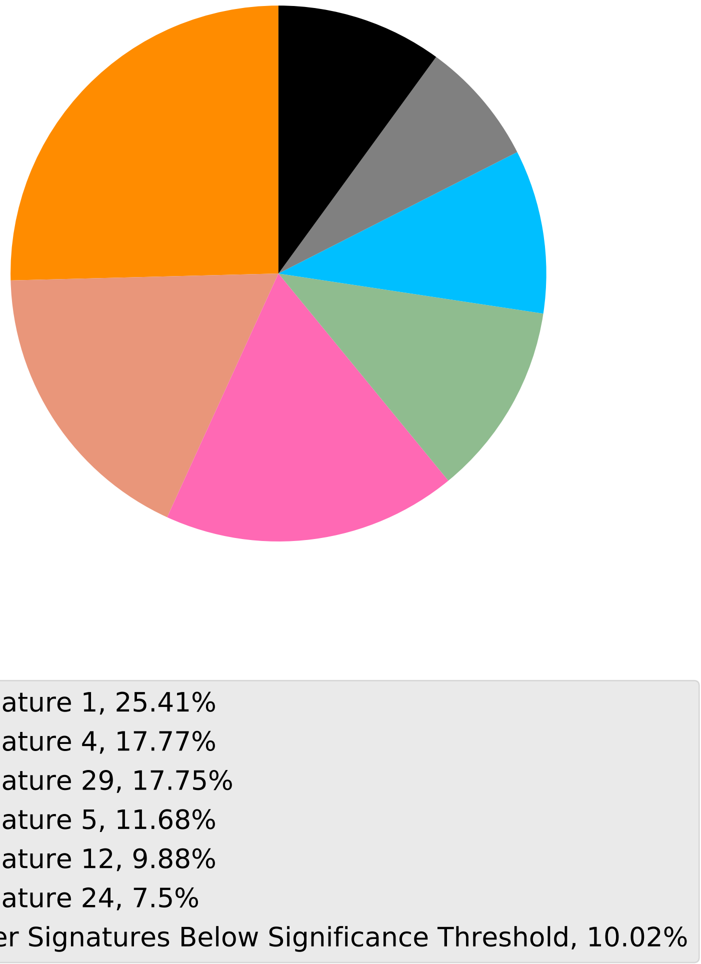

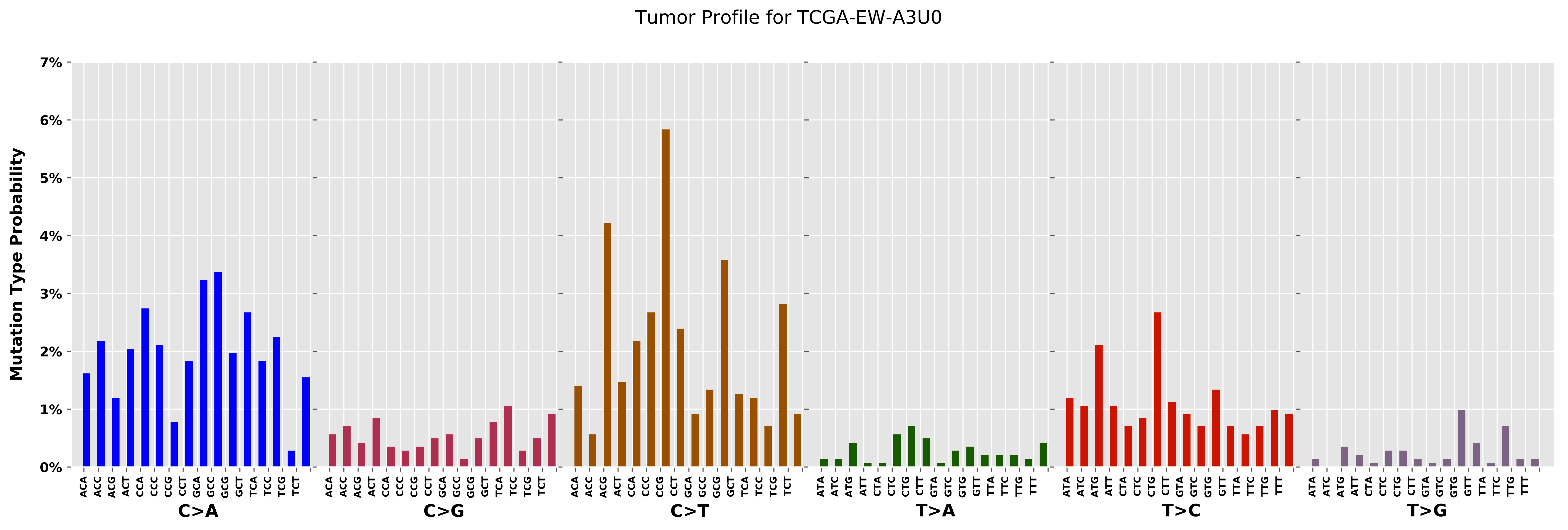
Tumor Profile for TCGA-EW-A3U0
Mutation Type Probability
0%
1%
2%
3%
4%
5%
6%
7%
C>A
C>G
C>T
T>A
T>C
T>G
ACA ACC ACG ACT CCA CCC CCG CCT GCA GCC GCG GCT TCA TCC TCG TCT
ATA ATC ATG ATT CTA CTC CTG CTT GTA GTC GTG GTT TTA TTC TTG TTT

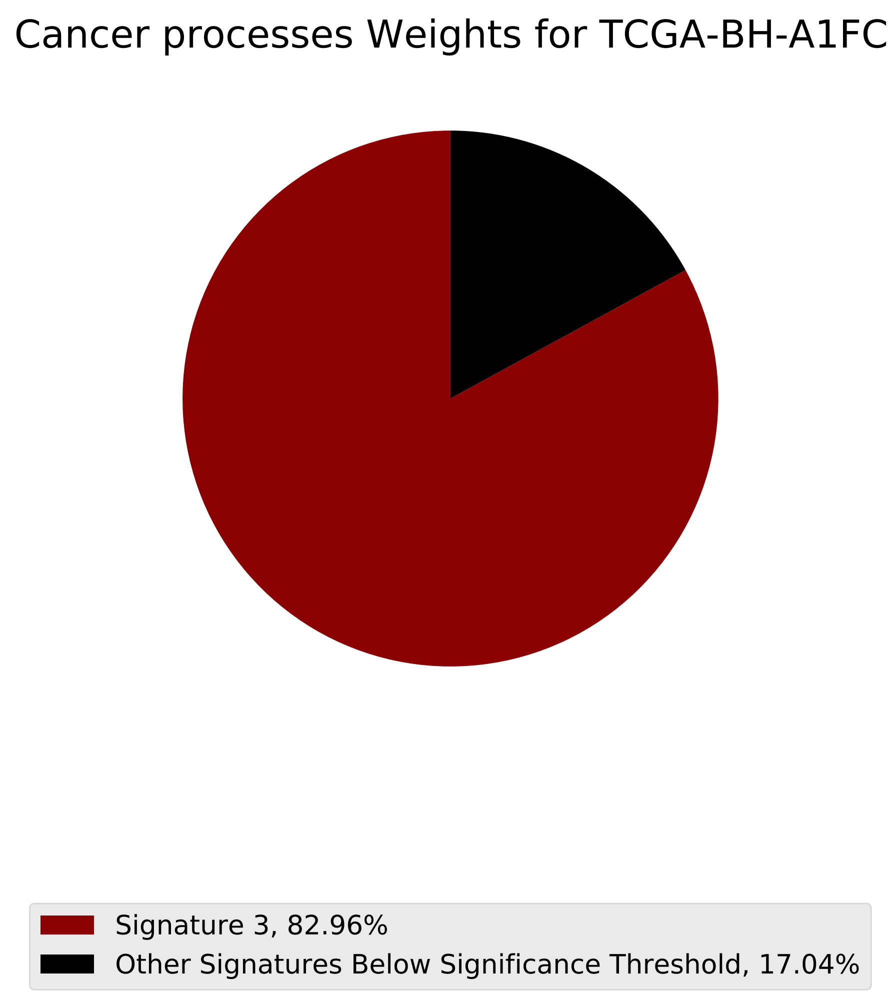
Cancer processes Weights for TCGA-BH-A1FC
Signature 3, 82.96%
Other Signatures Below Significance Threshold, 17.04%

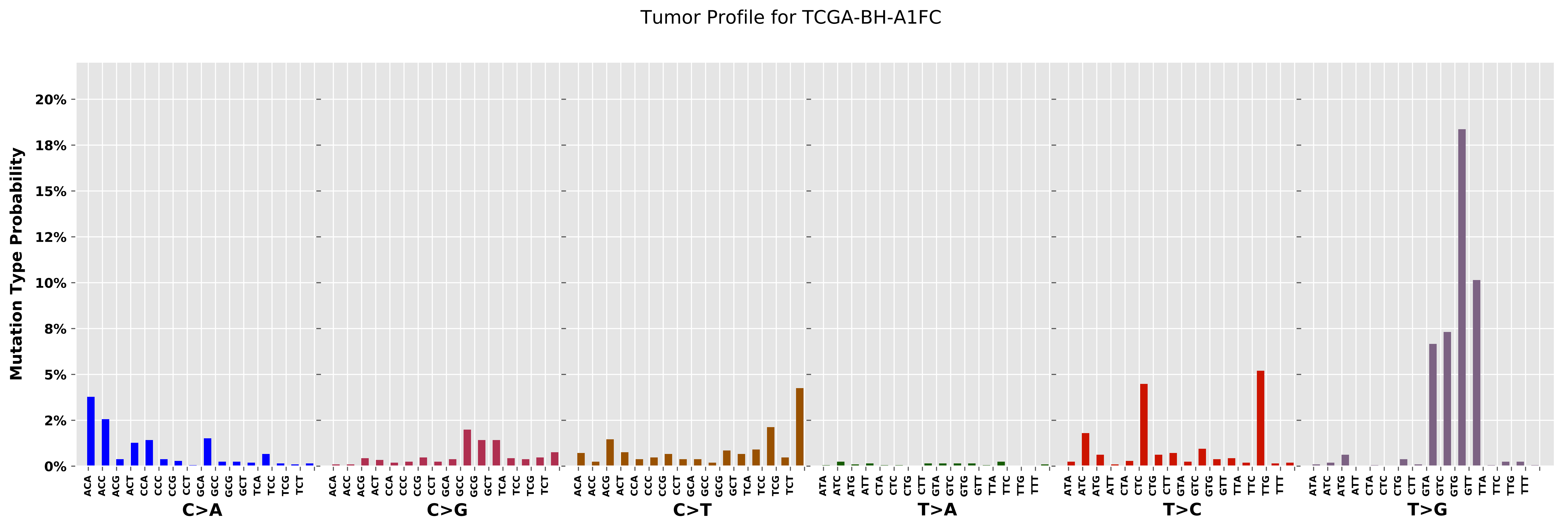

Tumor Profile for TCGA-BH-A1FC
Mutation Type Probability
20%
18%
15%
12%
10%
8%
5%
2%
0%
C>A
C>G
C>T
T>A
T>C
T>G

Cancer processes Weights for TCGA-AF-2691

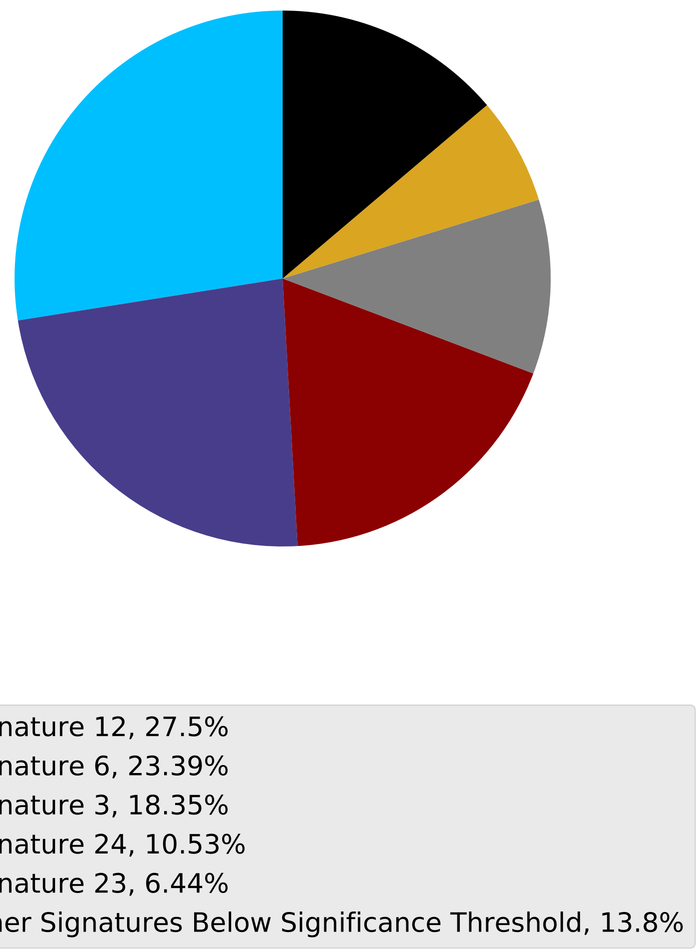

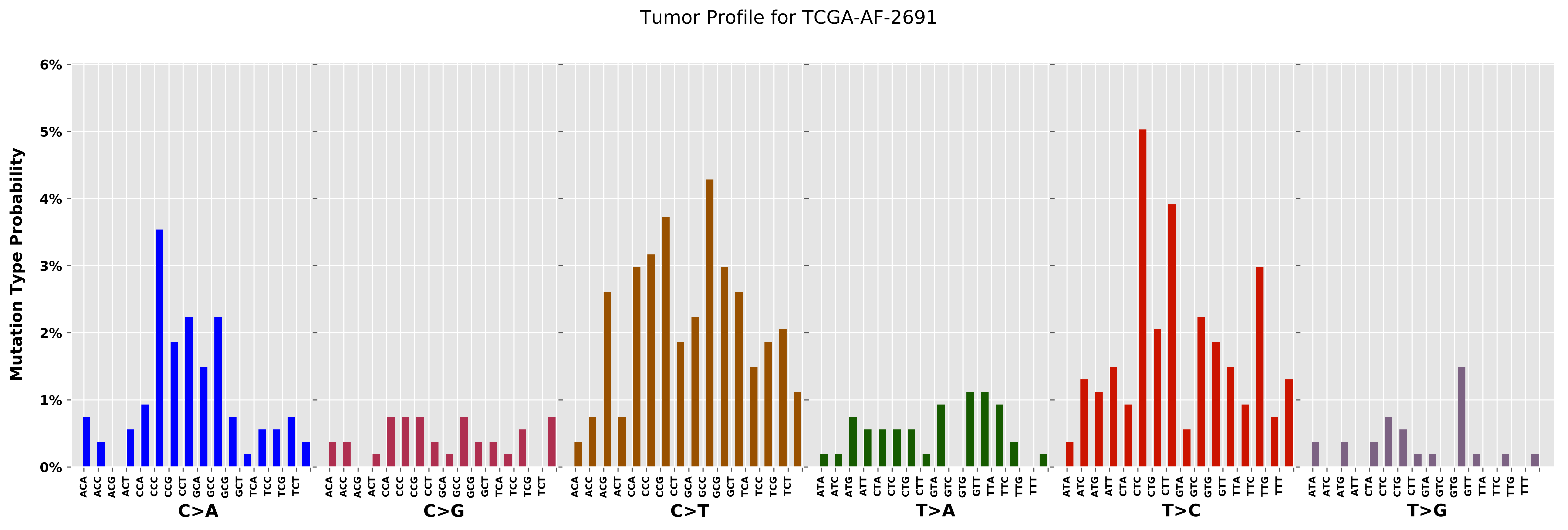
Tumor Profile for TCGA-AF-2691
Mutation Type Probability
6%
5%
4%
3%
2%
1%
0%
C>A
C>G
C>T
T>A
T>C
T>G
ACA ACC ACG ACT CCA CCC CCG CCT GCA GCC GCG GCT TCA TCC TCG TCT
ATA ATC ATG ATT CTA CTC CTG CTT GTA GTC GTG GTT TTA TTC TTG TTT

Cancer processes Weights for TCGA-AF-3913

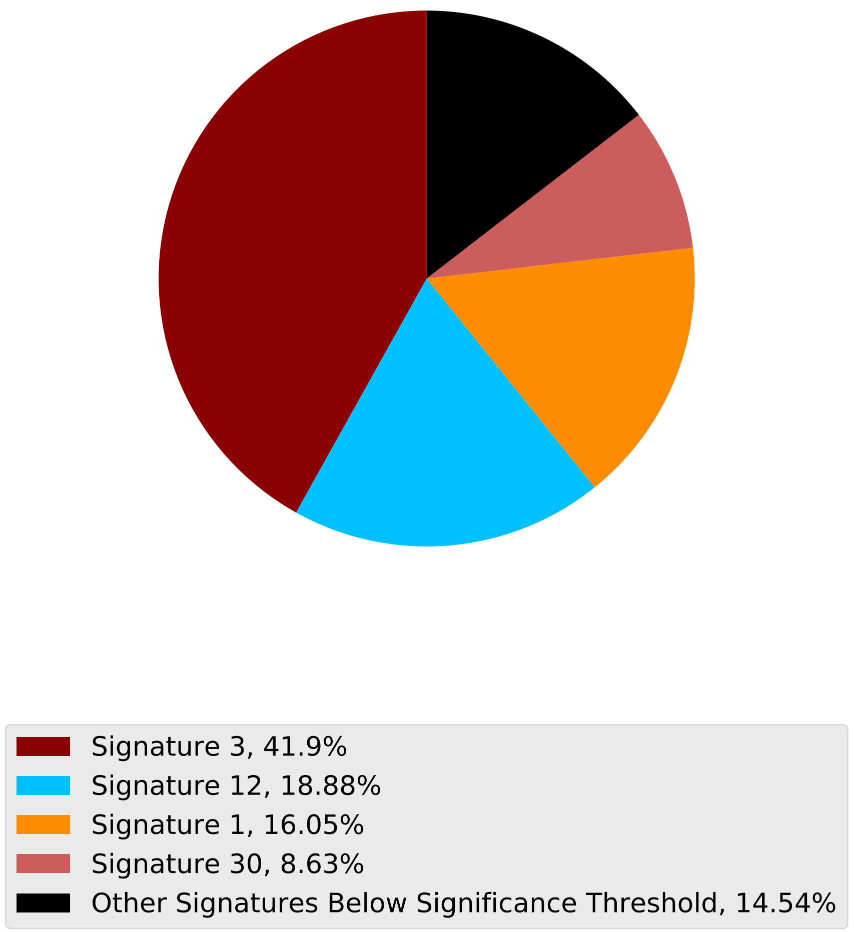

Tumor Profile for TCGA-AF-3913

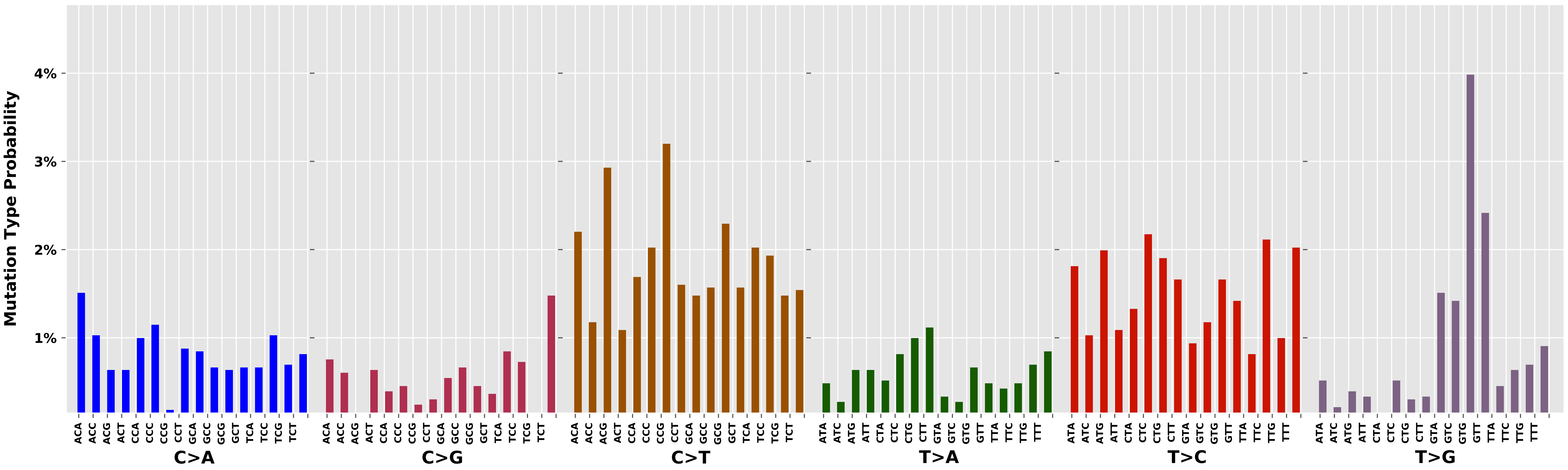

Cancer processes Weights for TCGA-AA-3977

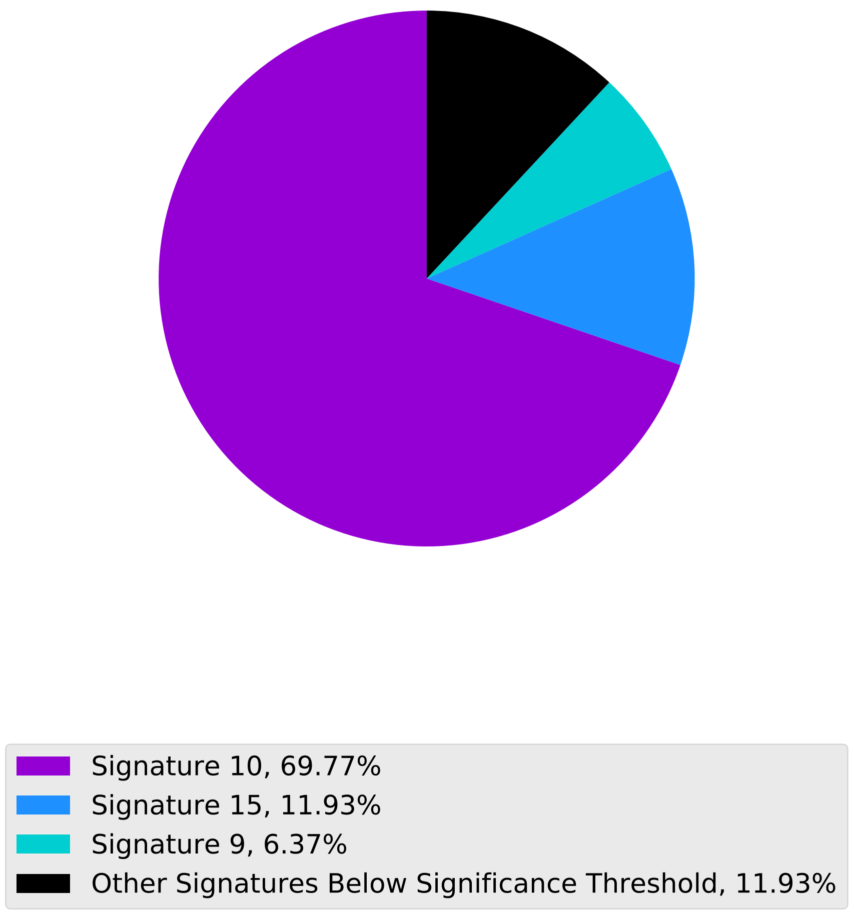

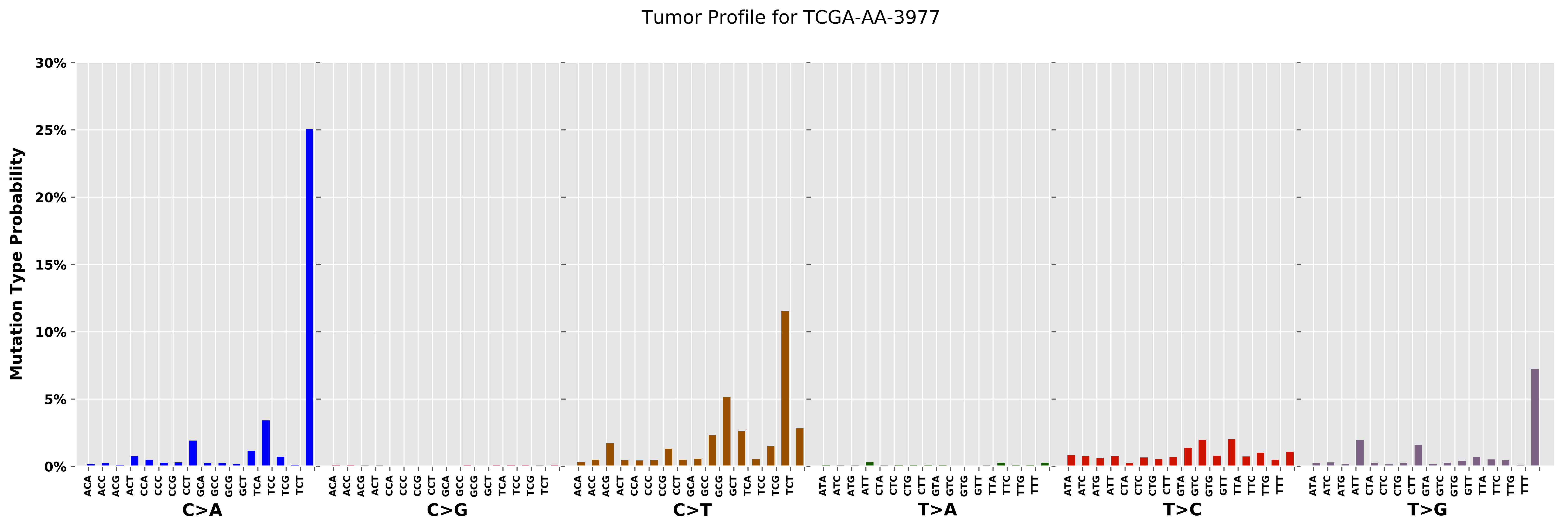
Tumor Profile for TCGA-AA-3977
Mutation Type Probability
30%
25%
20%
15%
10%
5%
0%
ACA ACC ACG ACT CCA CCC CCG CCT GCA GCC GCG GCT TCA TCC TCG TCT
C>A
C>G
C>T
ATA ATC ATG ATT CTA CTC CTG CTT GTA GTC GTG GTT TTA TTC TTG TTT
T>A
T>C
T>G

Cancer processes Weights for TCGA-A7-A0CE

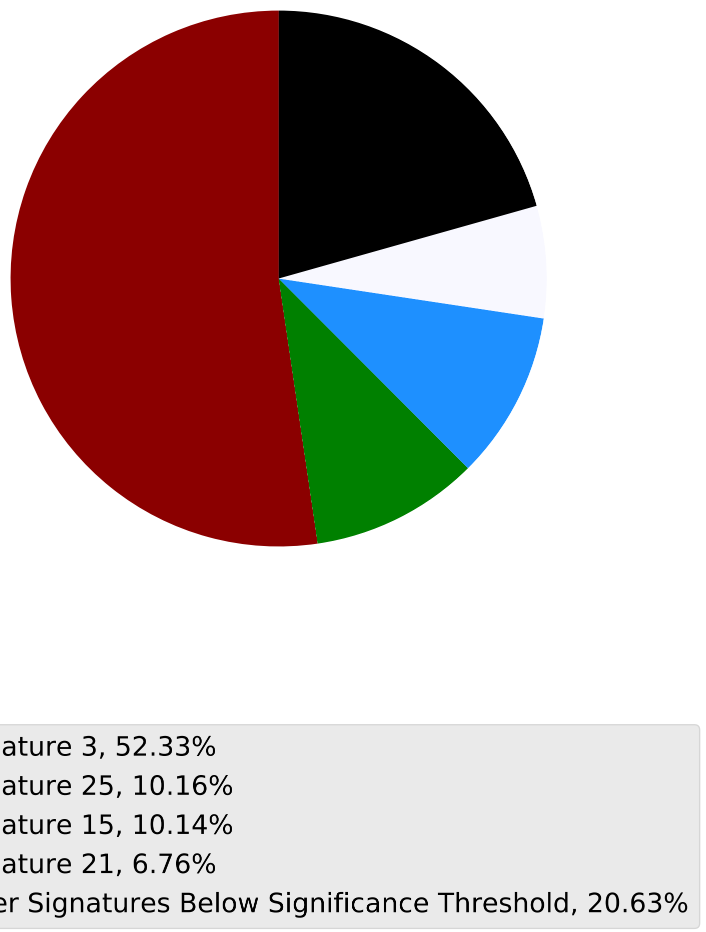

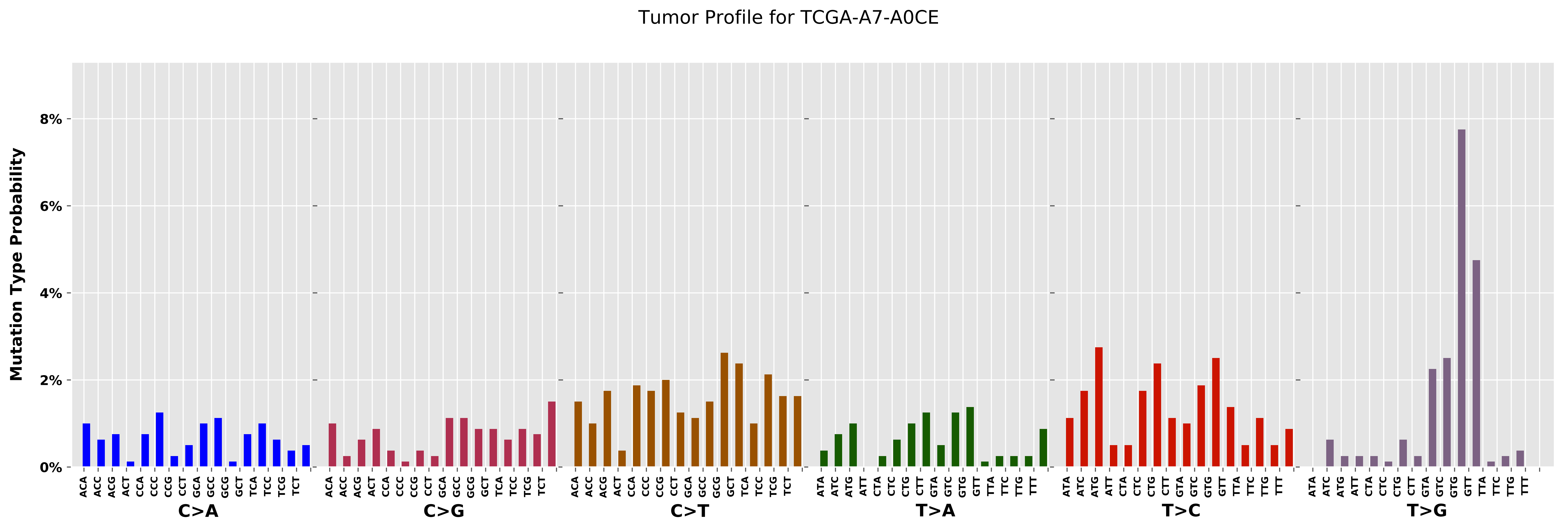

Tumor Profile for TCGA-A7-A0CE
Mutation Type Probability
0%
2%
4%
6%
8%
C>A
C>G
C>T
T>A
T>C
T>G

Cancer processes Weights for TCGA-AO-A0J4

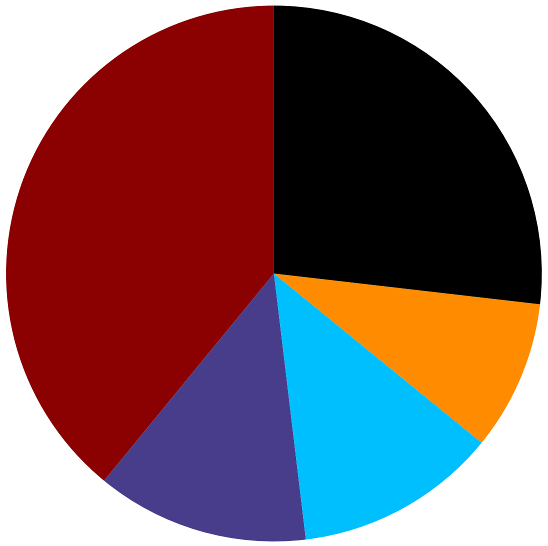

- Signature 3, 39.07%
- Signature 6, 12.84%
- Signature 12, 12.22%
- Signature 1, 9.05%
- Other Signatures Below Significance Threshold, 26.83%

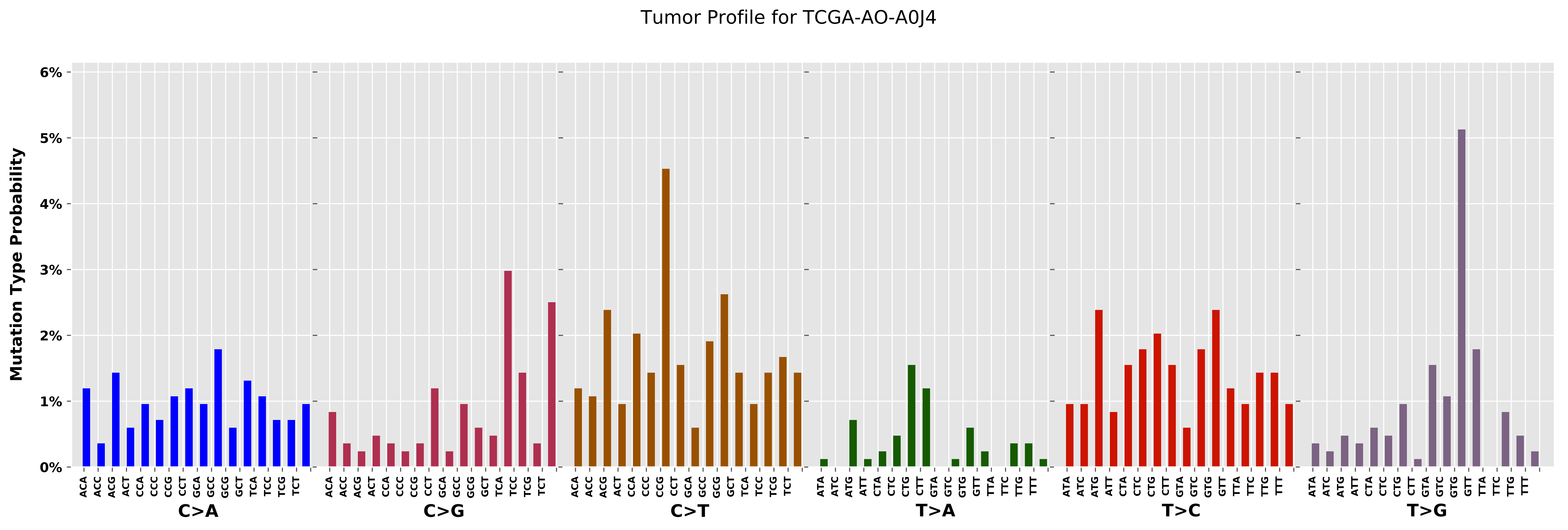

Tumor Profile for TCGA-AO-A0J4
Mutation Type Probability
6%
5%
4%
3%
2%
1%
0%
ACA
ACC
ACG
ACT
CCA
CCC
CCG
CCT
GCA
GCC
GCG
GCT
TCA
TCC
TCG
TCT
ATA
ATC
ATG
ATT
CTA
CTC
CTG
CTT
GTA
GTC
GTG
GTT
TTA
TTC
TTG
TTT
C>A
C>G
C>T
T>A
T>C
T>G

Cancer processes Weights for TCGA-AQ-A04J

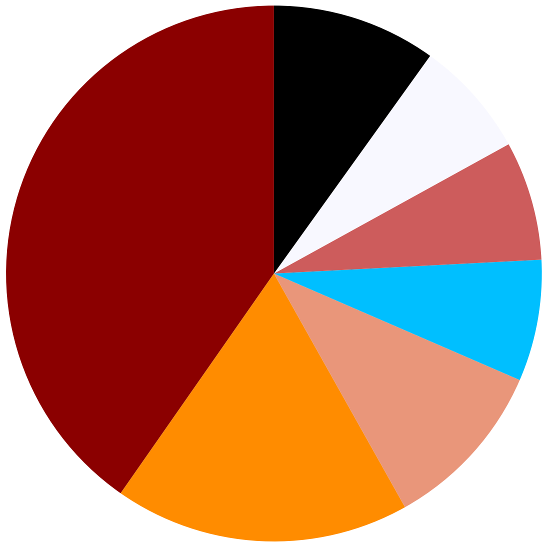

Signature 3, 40.3%
Signature 1, 17.84%
Signature 4, 10.35%
Signature 12, 7.33%
Signature 30, 7.19%
Signature 21, 7.08%
Other Signatures Below Significance Threshold, 9.91%

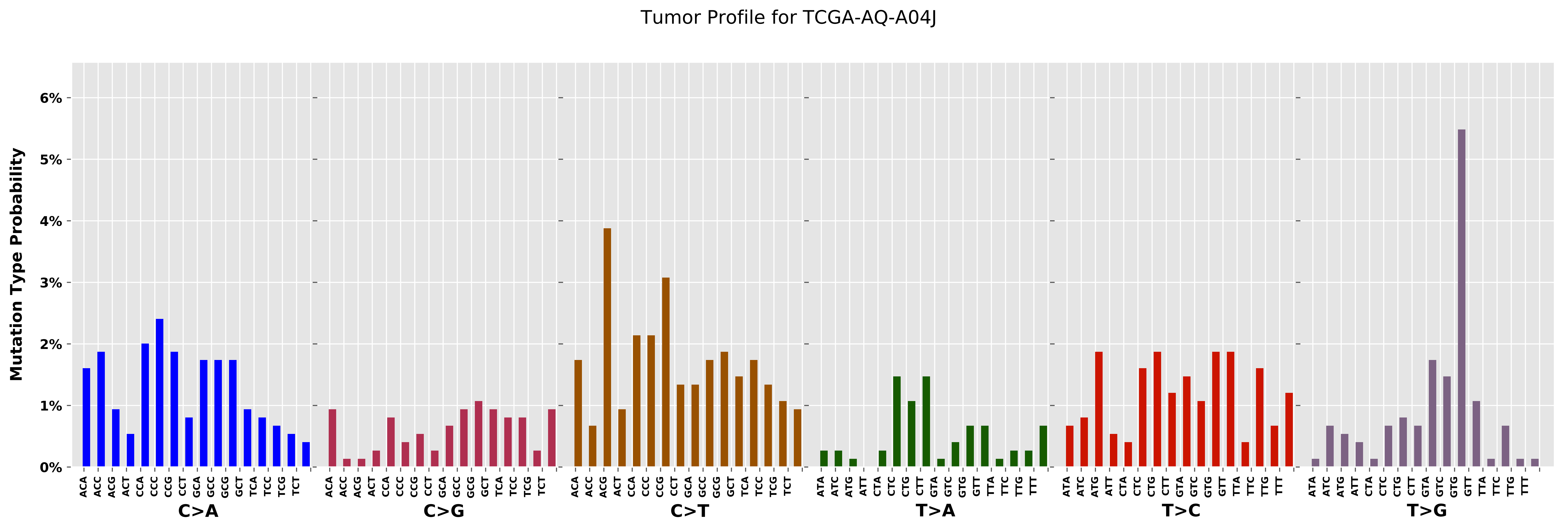

Tumor Profile for TCGA-AQ-A04J
Mutation Type Probability
0%
1%
2%
3%
4%
5%
6%
C>A
C>G
C>T
T>A
T>C
T>G
ACA ACC ACG ACT CCA CCC CCG CCT GCA GCC GCG GCT TCA TCC TCG TCT
ATA ATC ATG ATT CTA CTC CTG CTT GTA GTC GTG GTT TTA TTC TTG TTT

Cancer processes Weights for TCGA-AG-4007

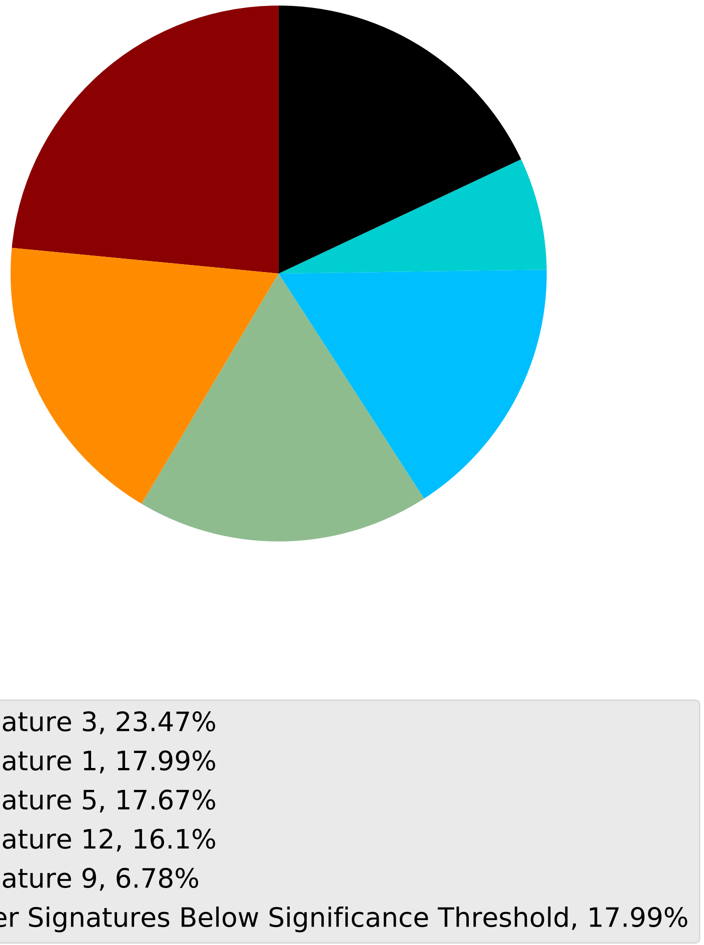

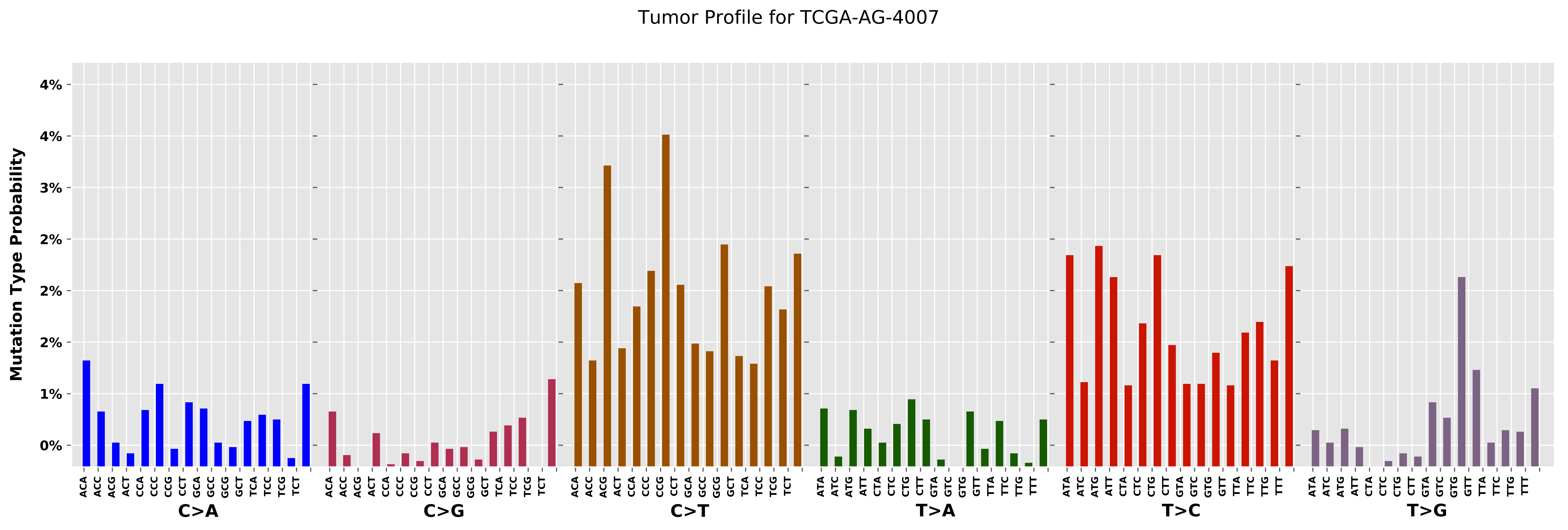

Tumor Profile for TCGA-AG-4007
Mutation Type Probability
4%
4%
3%
2%
2%
2%
1%
0%
C>A
C>G
C>T
T>A
T>C
T>G

Cancer processes Weights for TCGA-CA-6717

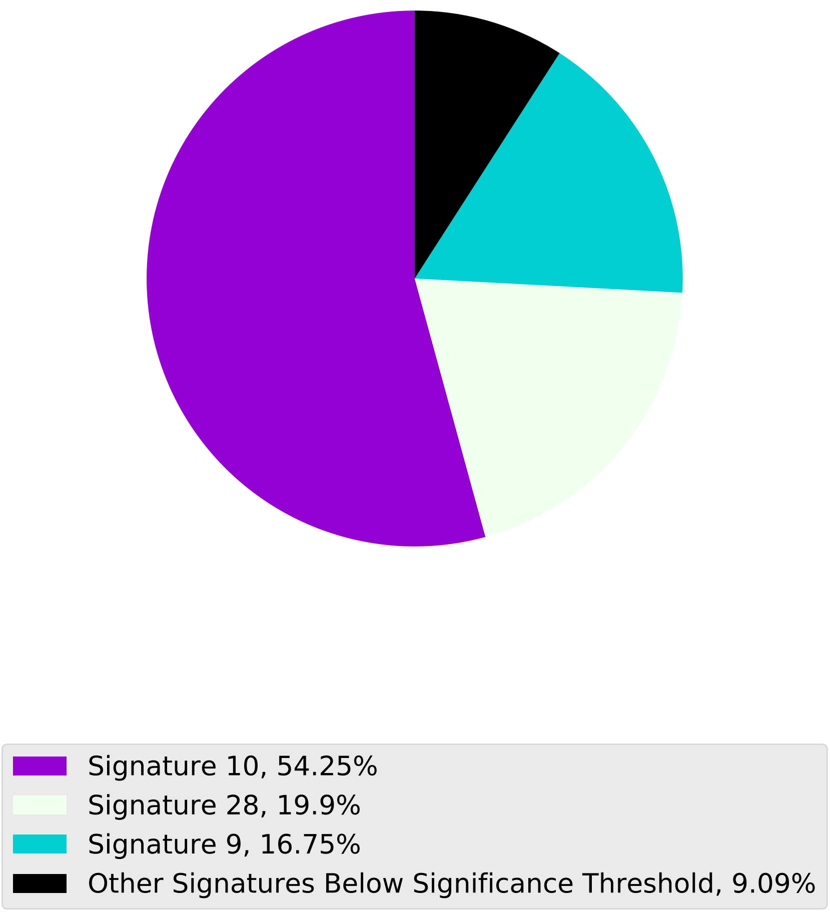

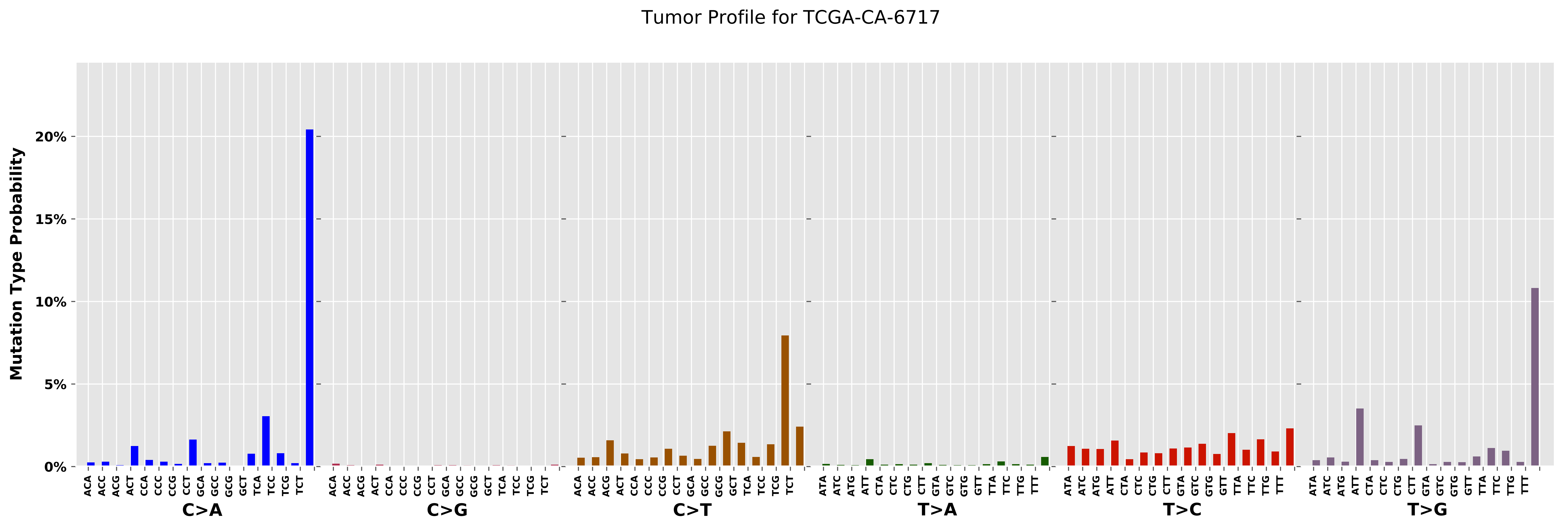

Tumor Profile for TCGA-CA-6717
Mutation Type Probability
20%
15%
10%
5%
0%
C>A
C>G
C>T
T>A
T>C
T>G
ACA ACC ACG ACT CCA CCC CCG CCT GCA GCC GCG GCT TCA TCC TCG TCT
ATA ATC ATG ATT CTA CTC CTG CTT GTA GTC GTG GTT TTA TTC TTG TTT

Cancer processes Weights for TCGA-E2-A14P

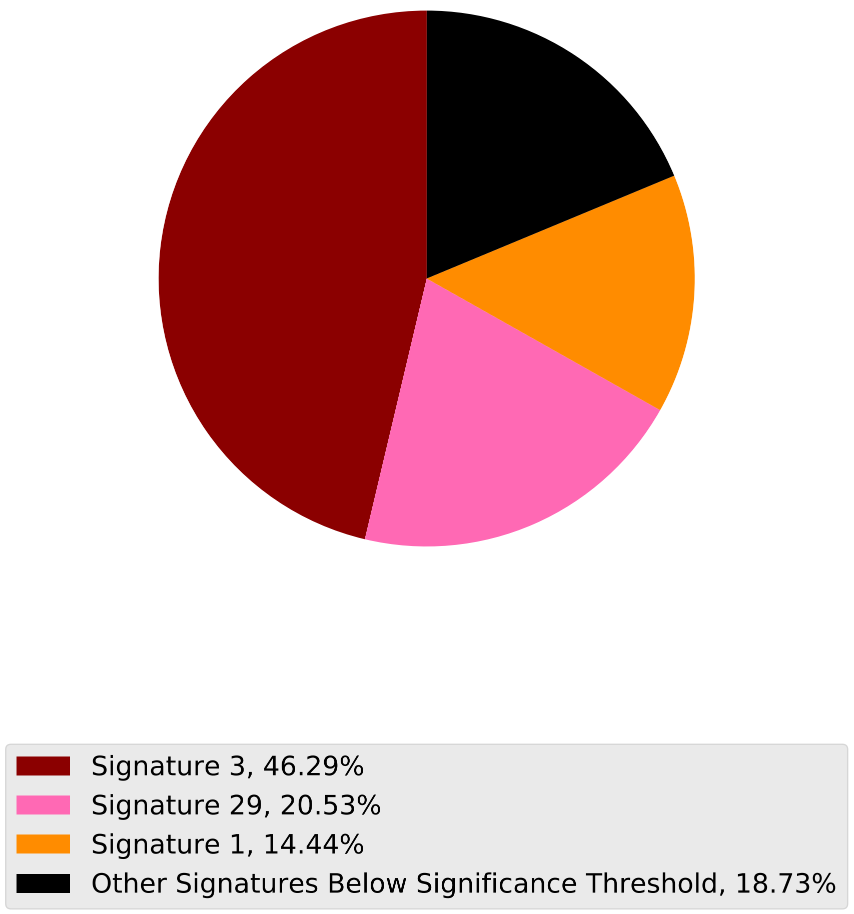

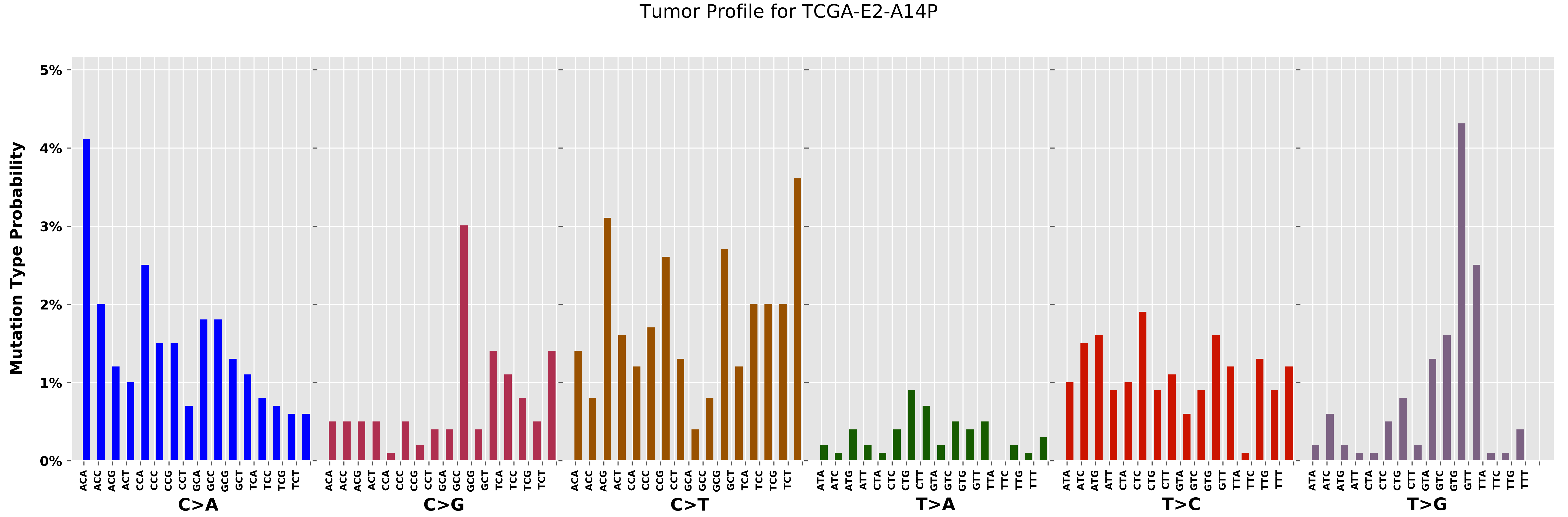
Tumor Profile for TCGA-E2-A14P
Mutation Type Probability
0%
1%
2%
3%
4%
5%
ACA ACC ACG ACT CCA CCC CCG CCT GCA GCC GCG GCT TCA TCC TCG TCT
ATA ATC ATG ATT CTA CTC CTG CTT GTA GTC GTG GTT TTA TTC TTG TTT
C>A
C>G
C>T
T>A
T>C
T>G

Cancer processes Weights for TCGA-AZ-6601

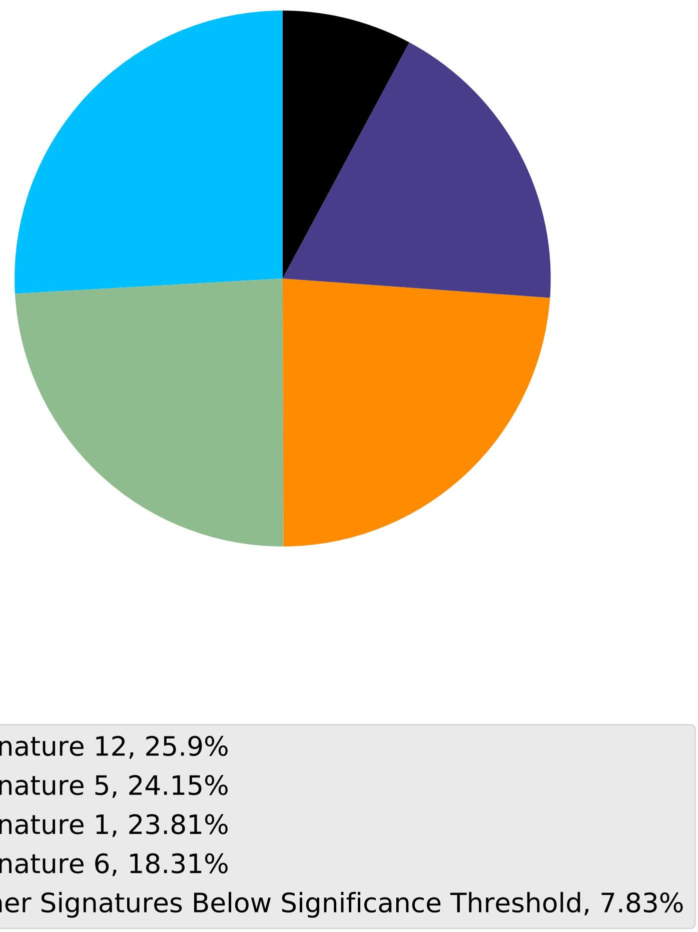

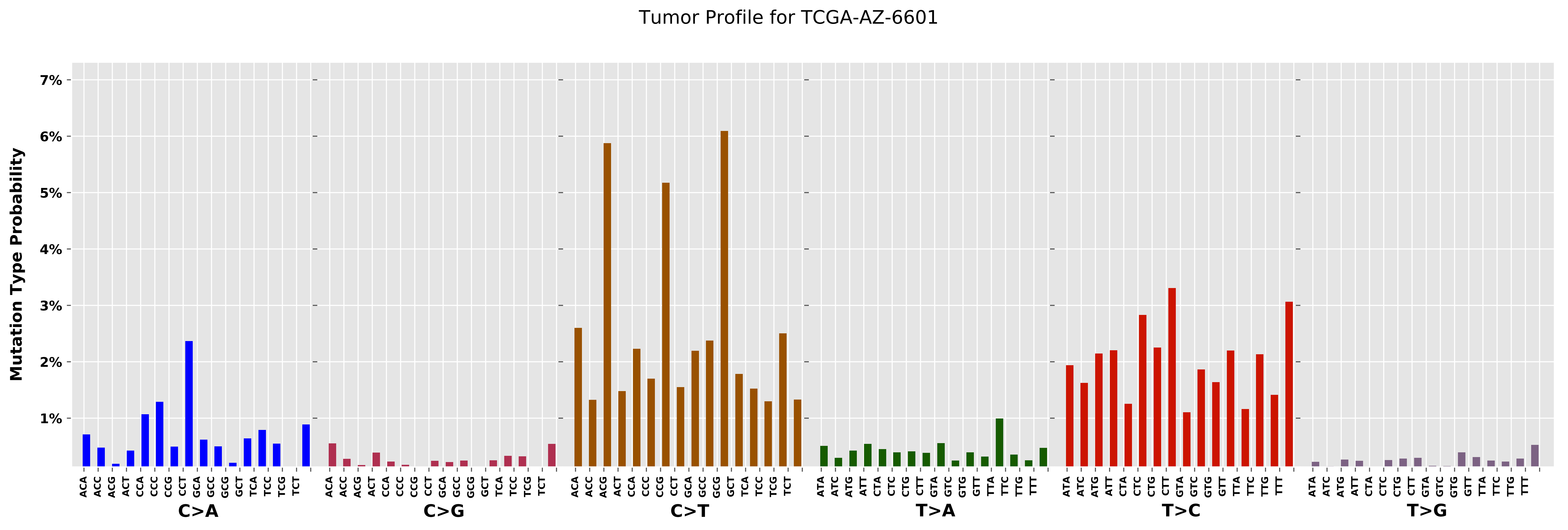

Tumor Profile for TCGA-AZ-6601
Mutation Type Probability
7%
6%
5%
4%
3%
2%
1%
ACA ACC ACG ACT CCA CCC CCG CCT GCA GCC GCG GCT TCA TCC TCG TCT
C>A
ACA ACC ACG ACT CCA CCC CCG CCT GCA GCC GCG GCT TCA TCC TCG TCT
C>G
ACA ACC ACG ACT CCA CCC CCG CCT GCA GCC GCG GCT TCA TCC TCG TCT
C>T
ATA ATC ATG ATT CTA CTC CTG CTT GTA GTC GTG GTT TTA TTC TTG TTT
T>A
ATA ATC ATG ATT CTA CTC CTG CTT GTA GTC GTG GTT TTA TTC TTG TTT
T>C
ATA ATC ATG ATT CTA CTC CTG CTT GTA GTC GTG GTT TTA TTC TTG TTT
T>G

Cancer processes Weights for TCGA-A6-6781

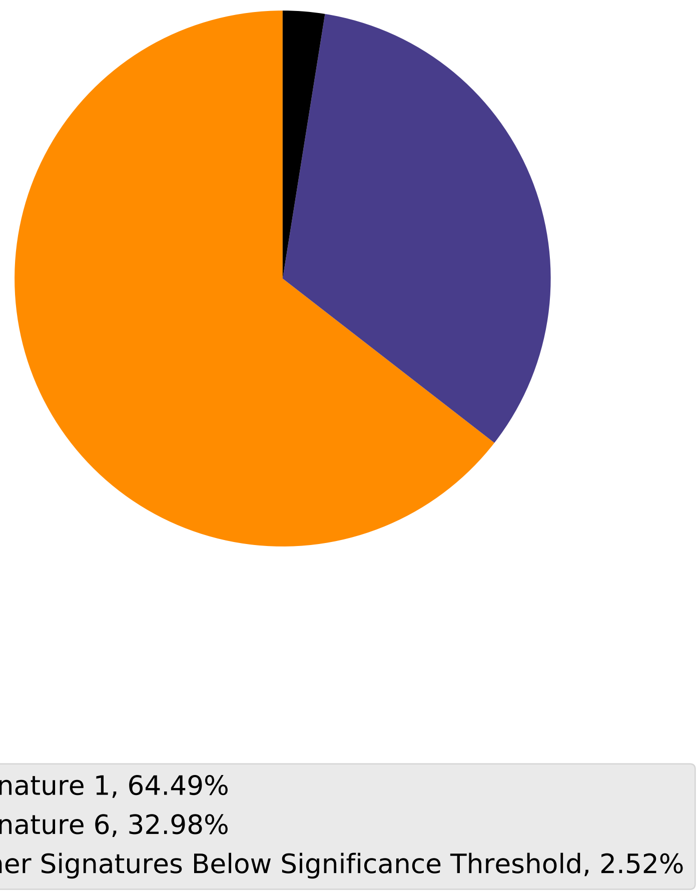

Tumor Profile for TCGA-A6-6781

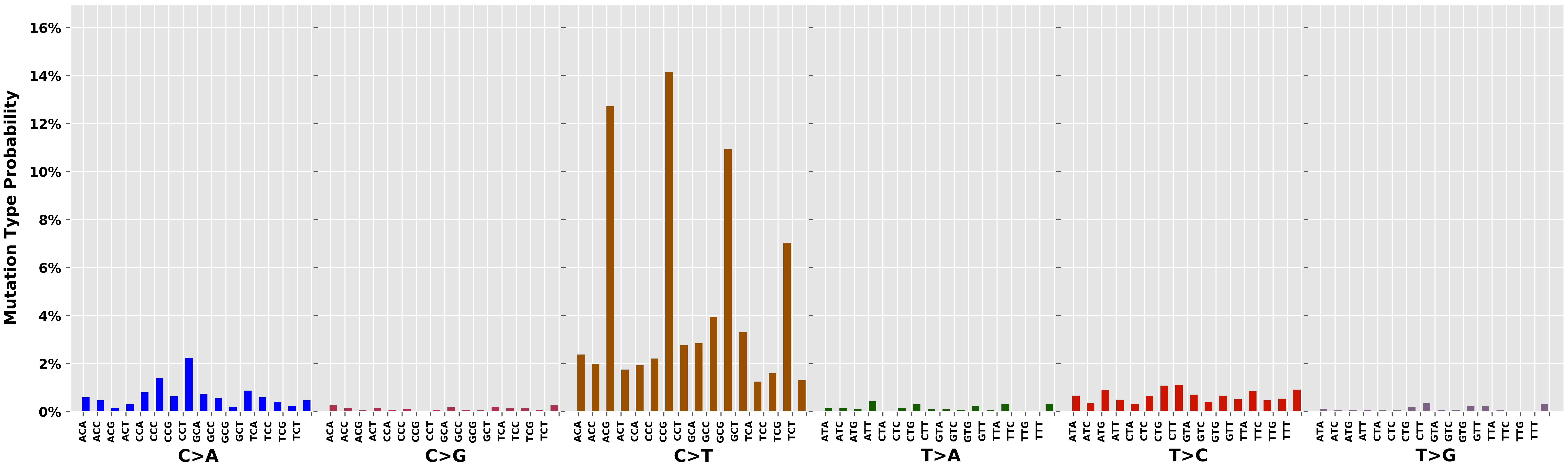

### Cancer processes Weights for TCGA-GM-A2DF

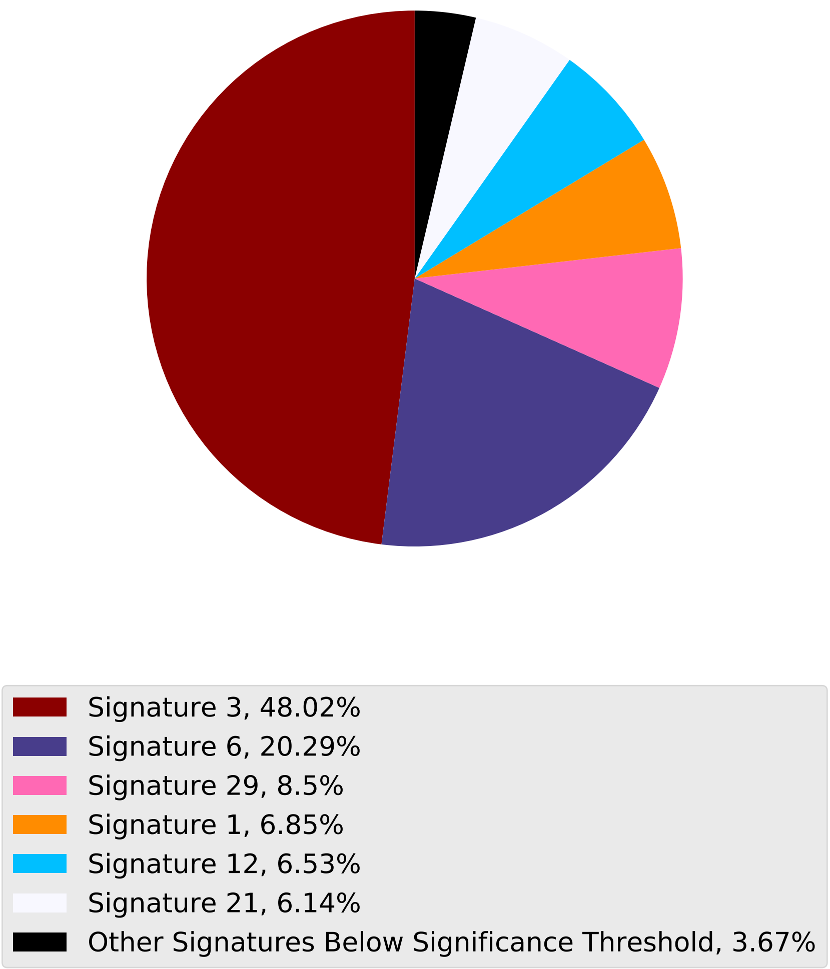

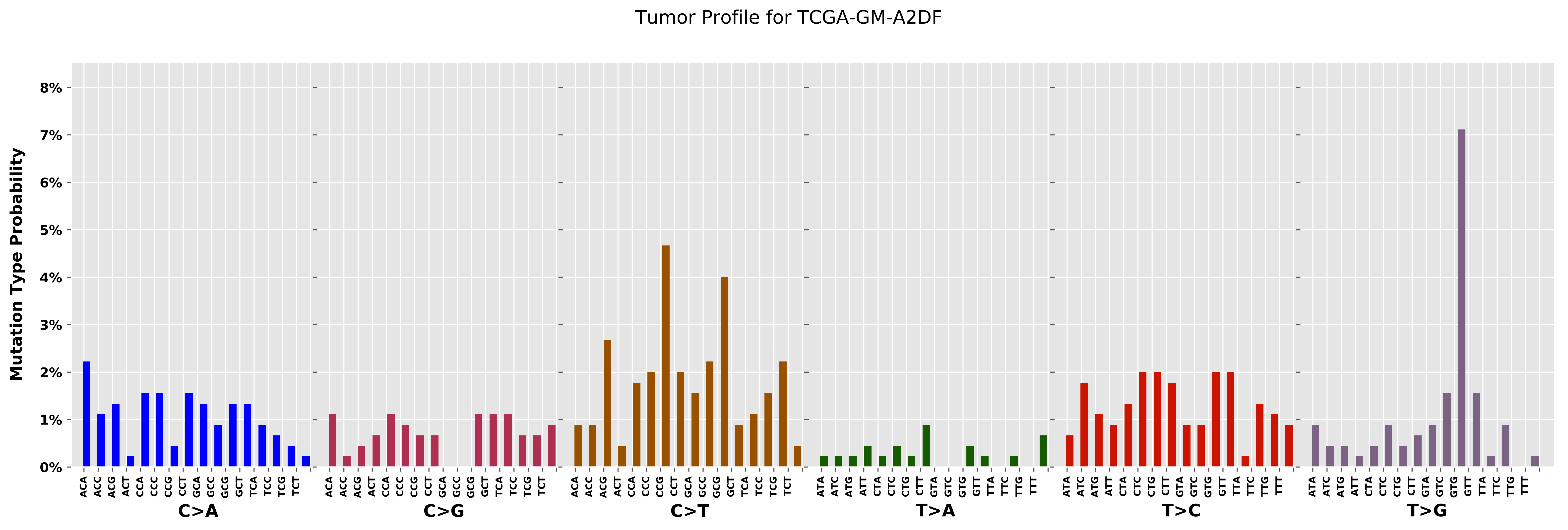

Tumor Profile for TCGA-GM-A2DF
Mutation Type Probability
0%
1%
2%
3%
4%
5%
6%
7%
8%
C>A
C>G
C>T
T>A
T>C
T>G
ACA ACC ACG ACT CCA CCC CCG CCT GCA GCC GCG GCT TCA TCC TCG TCT
ATA ATC ATG ATT CTA CTC CTG CTT GTA GTC GTG GTT TTA TTC TTG TTT

Cancer processes Weights for TCGA-C8-A12L

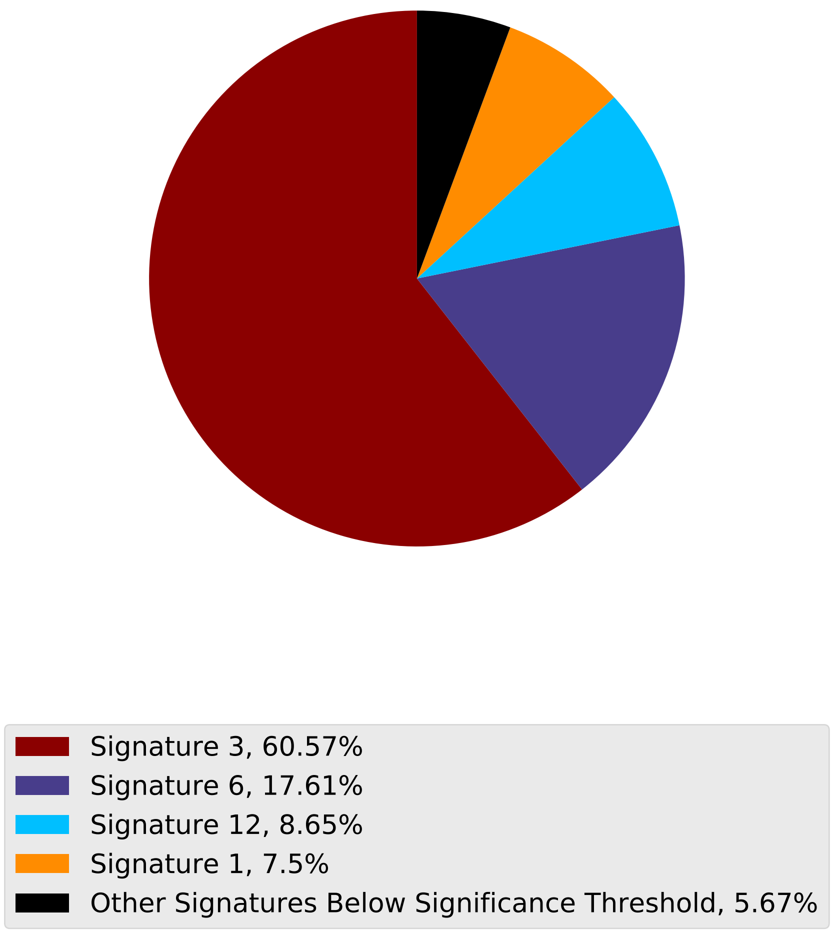

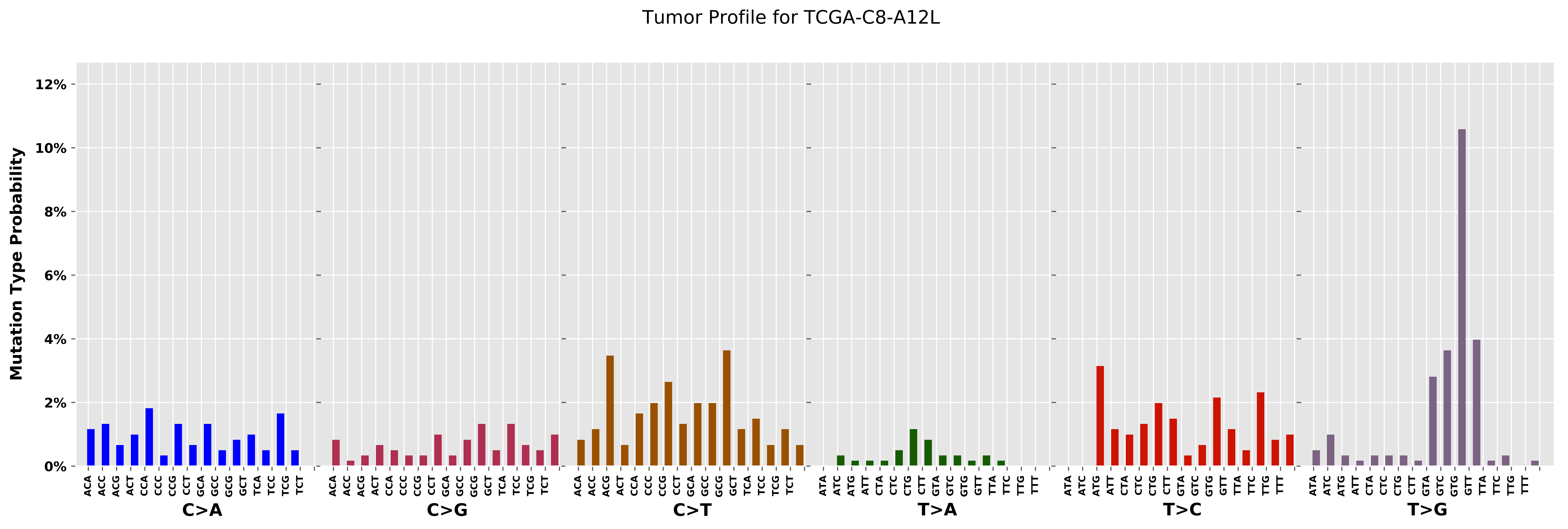
Tumor Profile for TCGA-C8-A12L
Mutation Type Probability
0%
2%
4%
6%
8%
10%
12%
C>A
C>G
C>T
T>A
T>C
T>G
ACA ACC ACG ACT CCA CCC CCG CCT GCA GCC GCG GCT TCA TCC TCG TCT
ATA ATC ATG ATT CTA CTC CTG CTT GTA GTC GTG GTT TTA TTC TTG TTT

#### Cancer processes Weights for TCGA-AG-3890

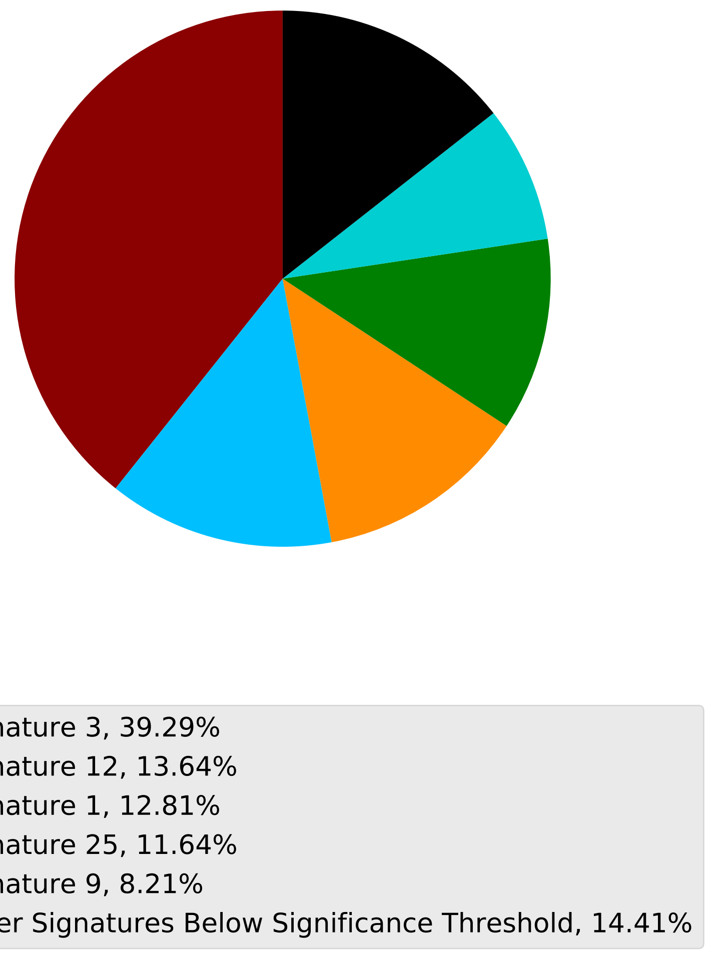

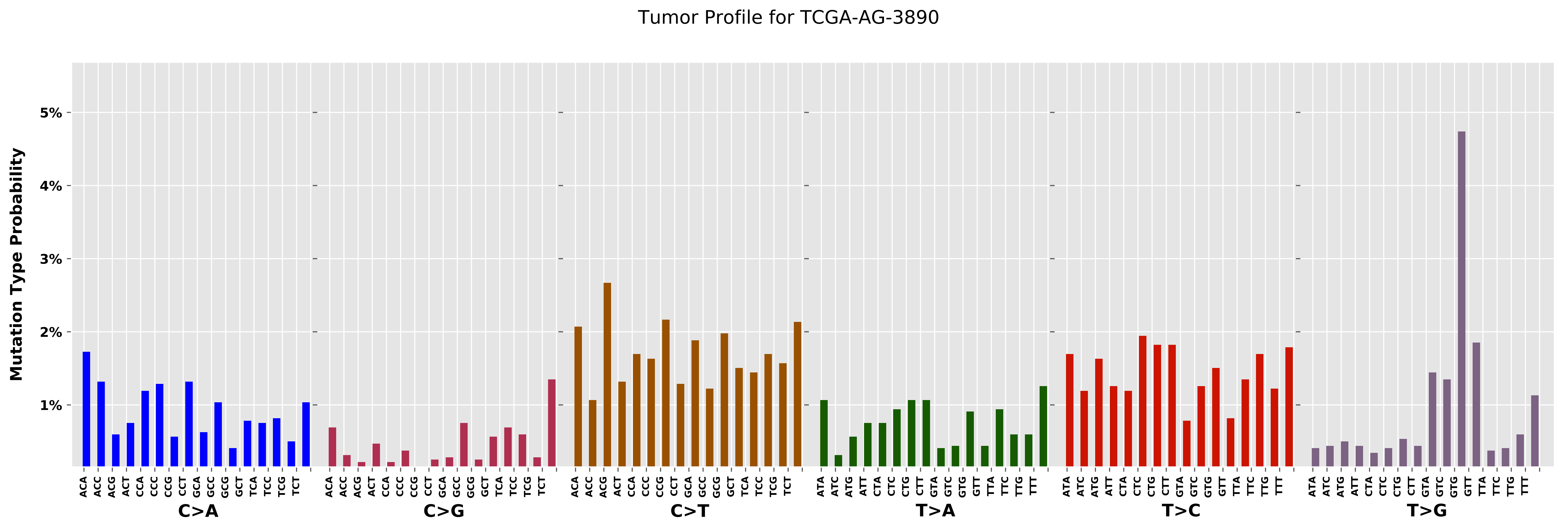

Tumor Profile for TCGA-AG-3890
Mutation Type Probability
5%
4%
3%
2%
1%
ACA ACC ACG ACT CCA CCC CCG CCT GCA GCC GCG GCT TCA TCC TCG TCT
C>A
C>G
C>T
ATA ATC ATG ATT CTA CTC CTG CTT GTA GTC GTG GTT TTA TTC TTG TTT
T>A
T>C
T>G

Cancer processes Weights for TCGA-E2-A1LK

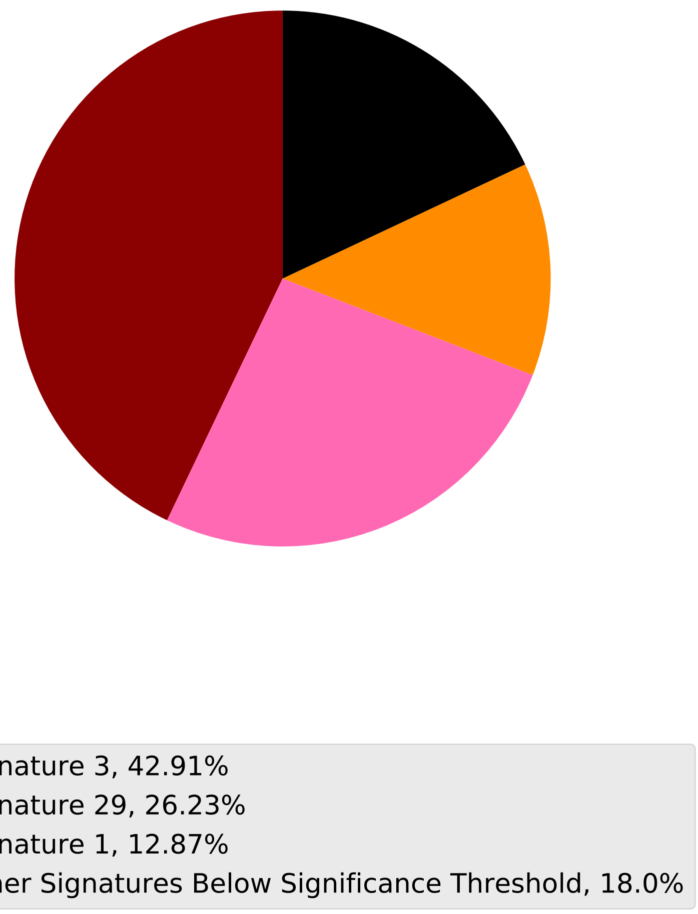

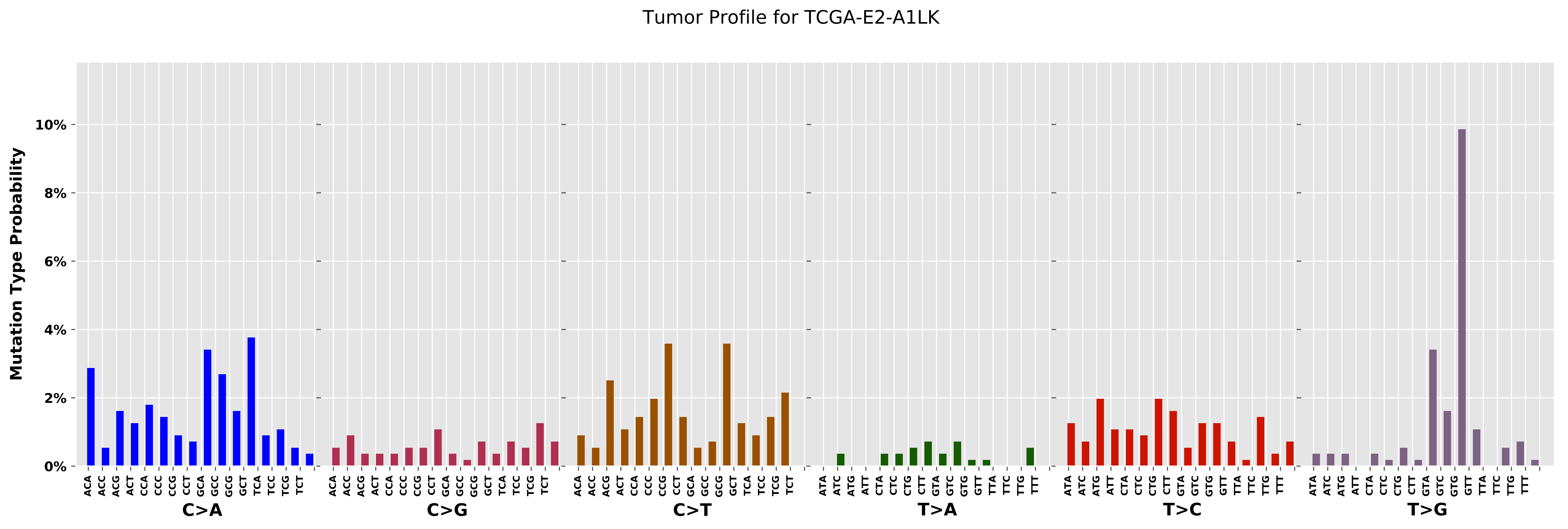
Tumor Profile for TCGA-E2-A1LK
Mutation Type Probability
0%
2%
4%
6%
8%
10%
C>A
C>G
C>T
T>A
T>C
T>G
ACA ACC ACG ACT CCA CCC CCG CCT GCA GCC GCG GCT TCA TCC TCG TCT
ATA ATC ATG ATT CTA CTC CTG CTT GTA GTC GTG GTT TTA TTC TTG TTT

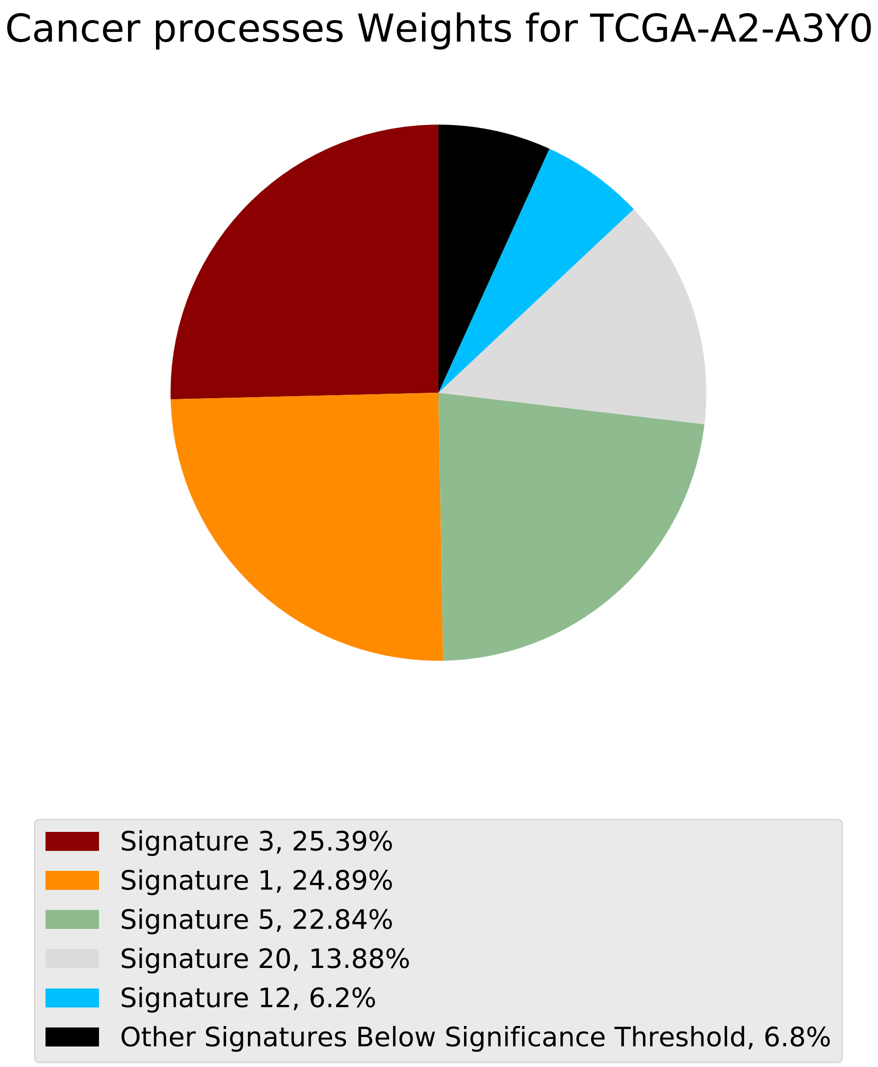

Cancer processes Weights for TCGA-A2-A3Y0
Signature 3, 25.39%
Signature 1, 24.89%
Signature 5, 22.84%
Signature 20, 13.88%
Signature 12, 6.2%
Other Signatures Below Significance Threshold, 6.8%

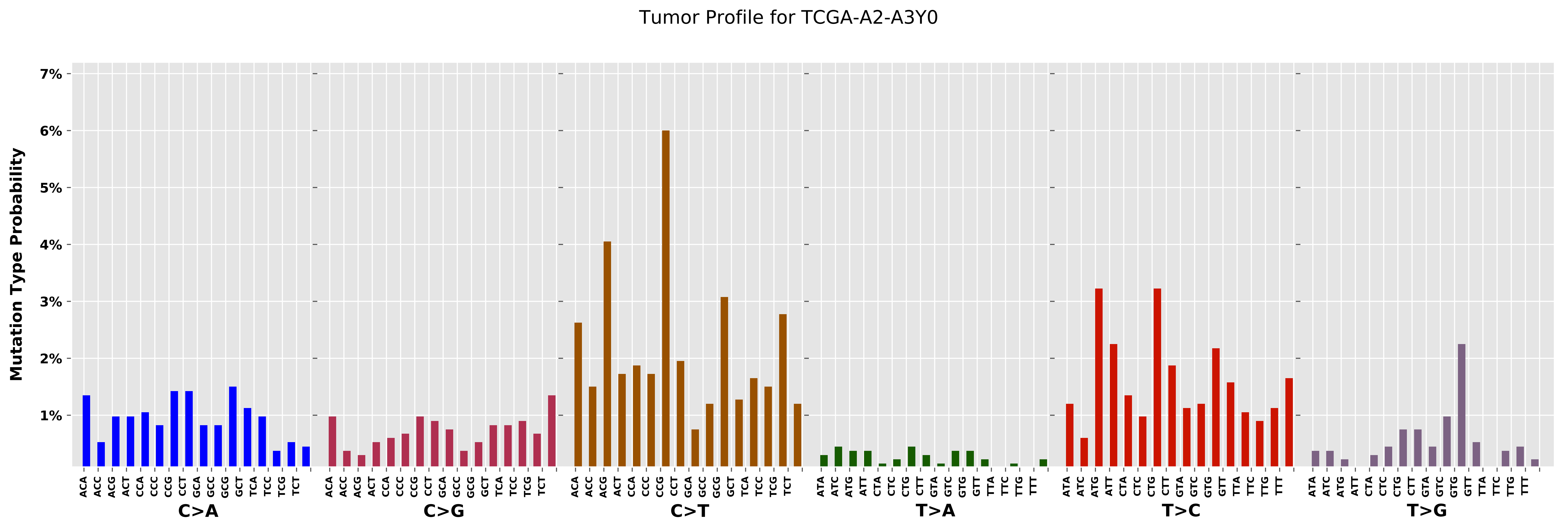

Tumor Profile for TCGA-A2-A3Y0
Mutation Type Probability
7%
6%
5%
4%
3%
2%
1%
C>A
C>G
C>T
T>A
T>C
T>G

#### Cancer processes Weights for TCGA-B6-A0I1

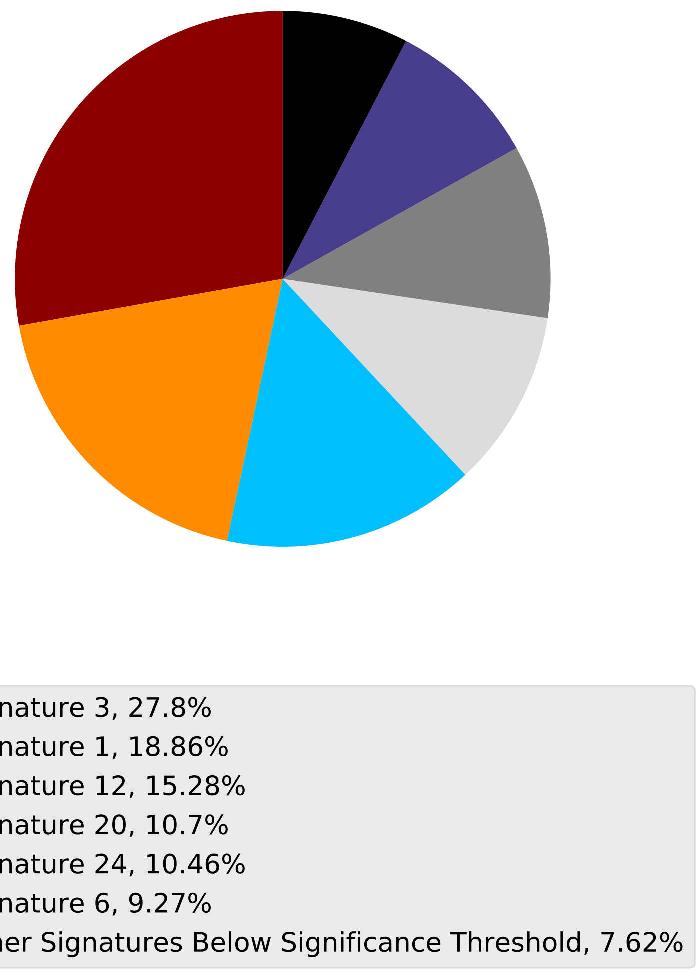

Tumor Profile for TCGA-B6-A0I1

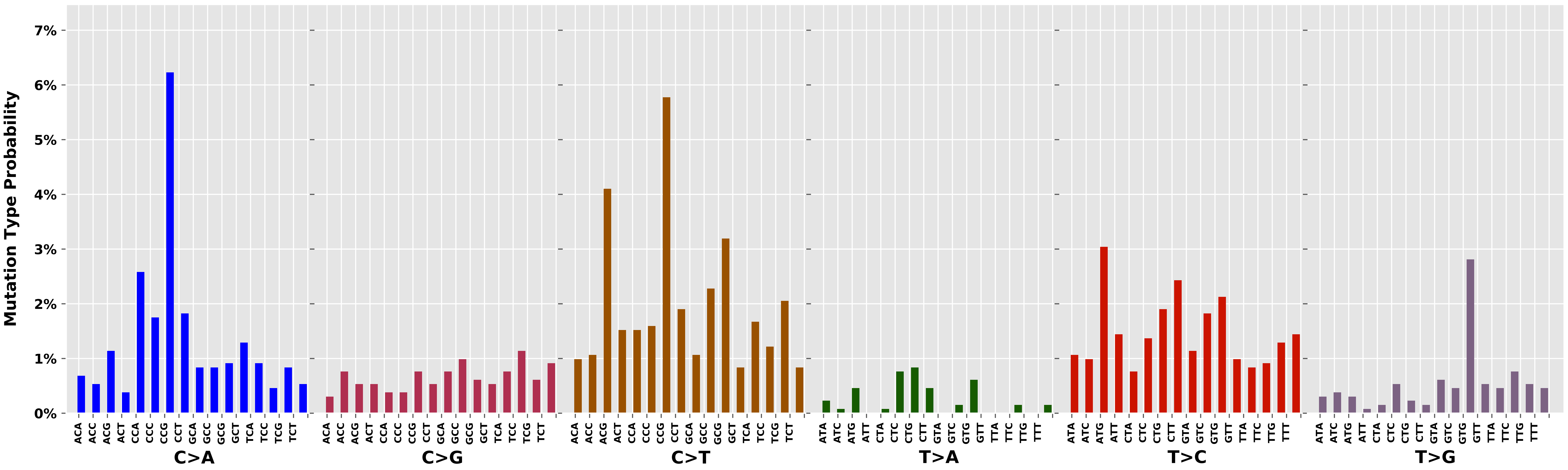

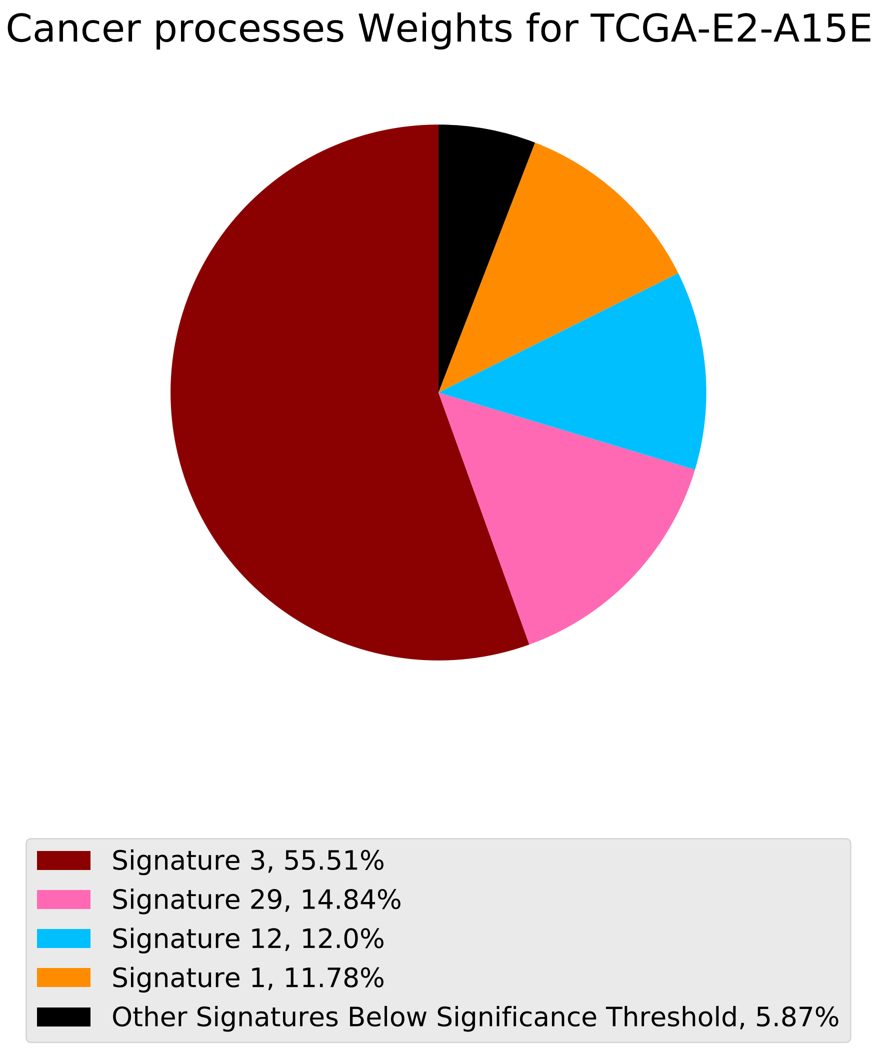
Cancer processes Weights for TCGA-E2-A15E
Signature 3, 55.51%
Signature 29, 14.84%
Signature 12, 12.0%
Signature 1, 11.78%
Other Signatures Below Significance Threshold, 5.87%

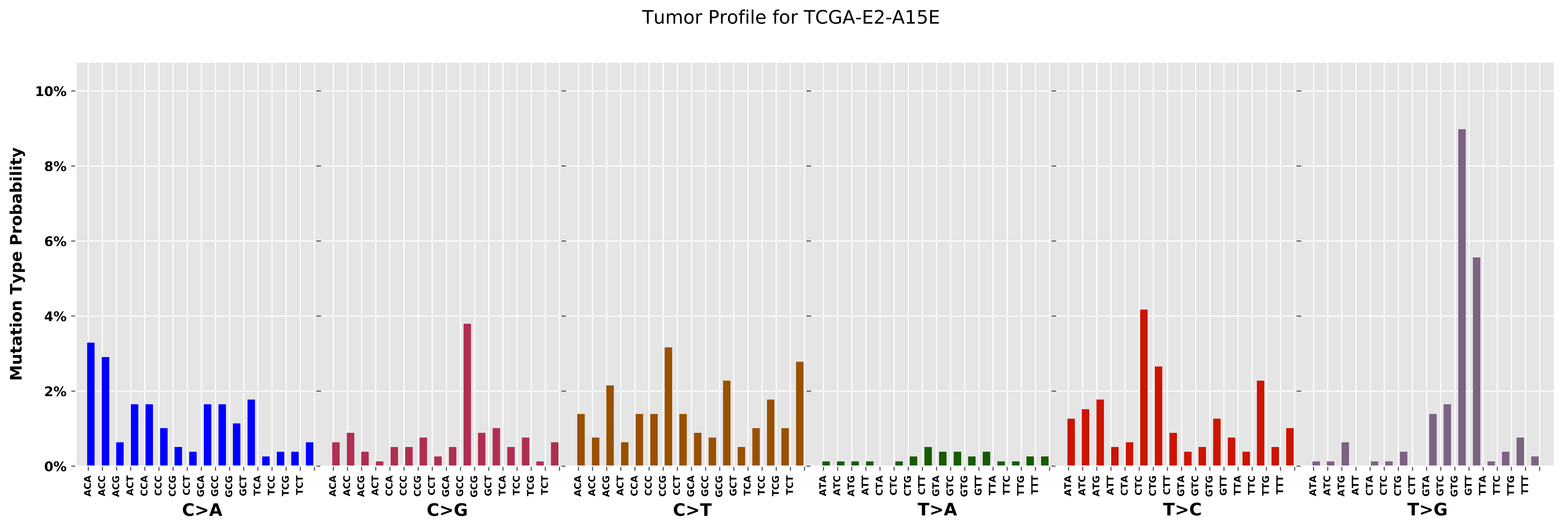
Tumor Profile for TCGA-E2-A15E
Mutation Type Probability
0%
2%
4%
6%
8%
10%
C>A
C>G
C>T
T>A
T>C
T>G

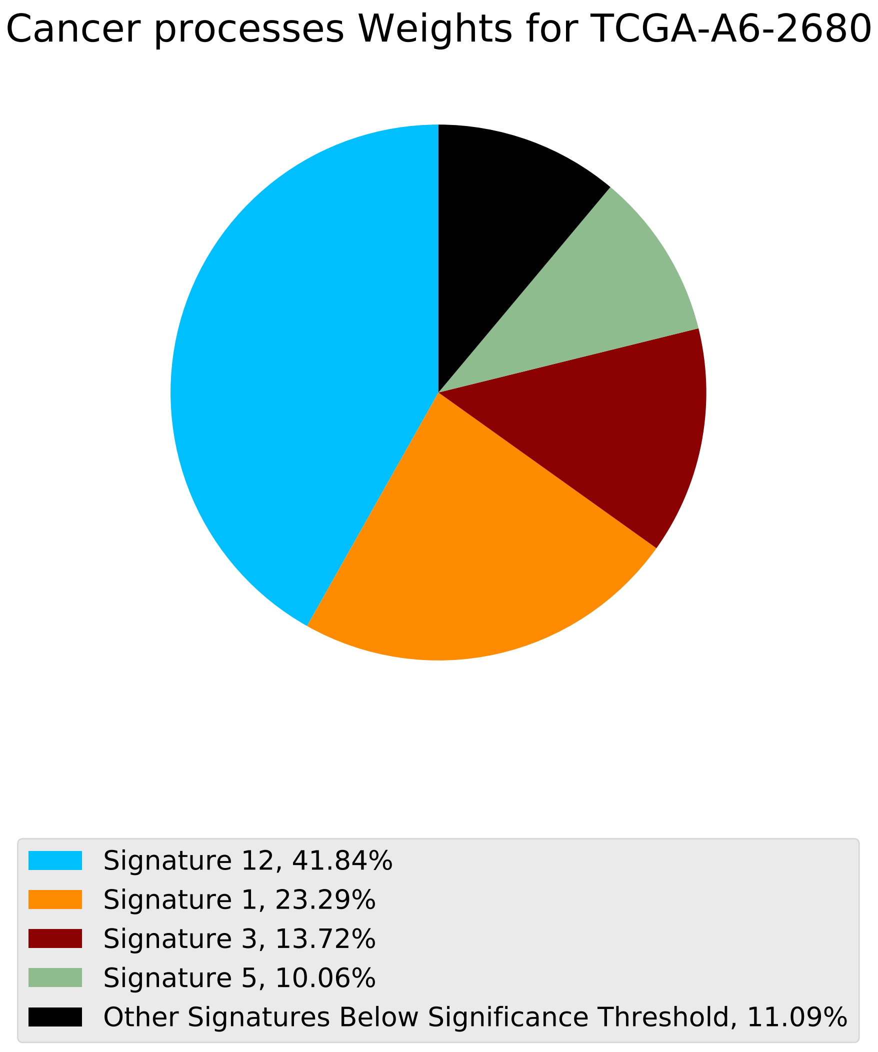
Cancer processes Weights for TCGA-A6-2680
Signature 12, 41.84%
Signature 1, 23.29%
Signature 3, 13.72%
Signature 5, 10.06%
Other Signatures Below Significance Threshold, 11.09%

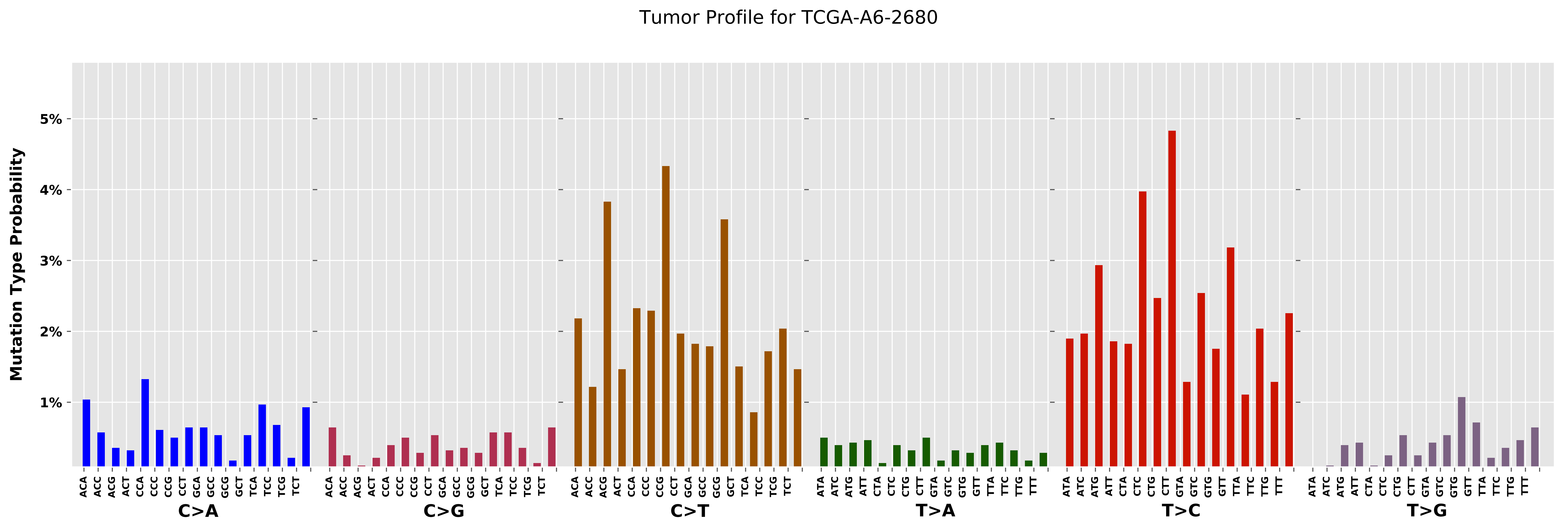
Tumor Profile for TCGA-A6-2680
Mutation Type Probability
5%
4%
3%
2%
1%
ACA ACC ACG ACT CCA CCC CCG CCT GCA GCC GCG GCT TCA TCC TCG TCT
C>A
C>G
C>T
ATA ATC ATG ATT CTA CTC CTG CTT GTA GTC GTG GTT TTA TTC TTG TTT
T>A
T>C
T>G

Cancer processes Weights for TCGA-A8-A07B

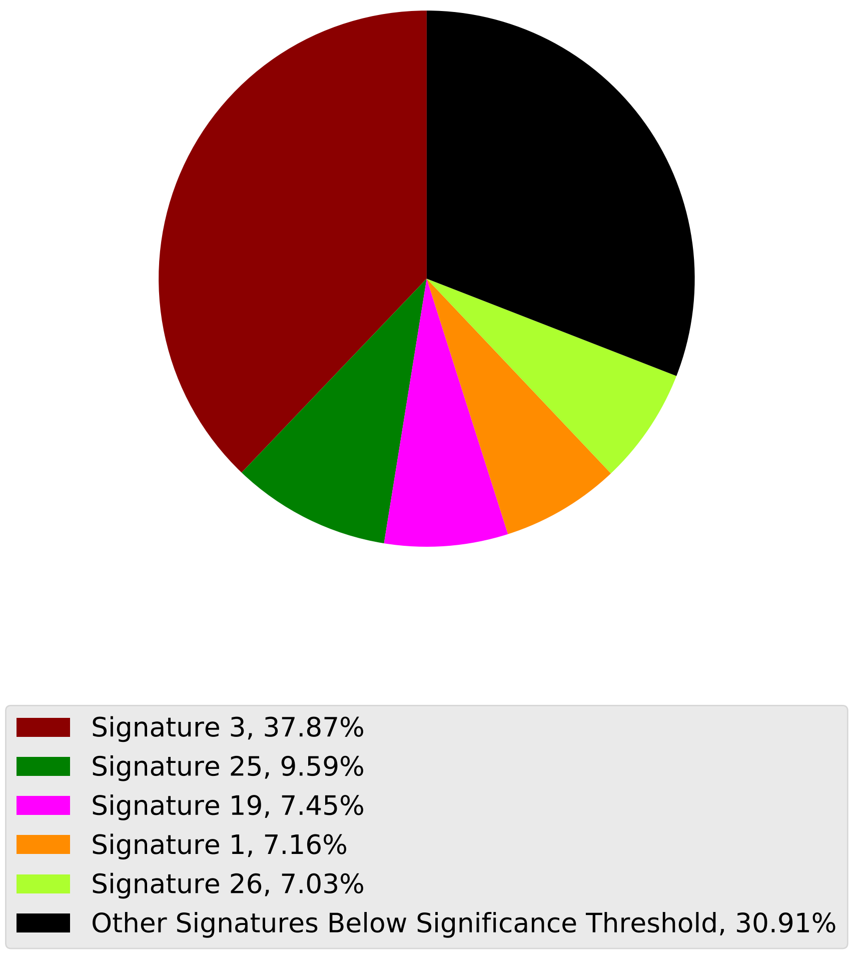

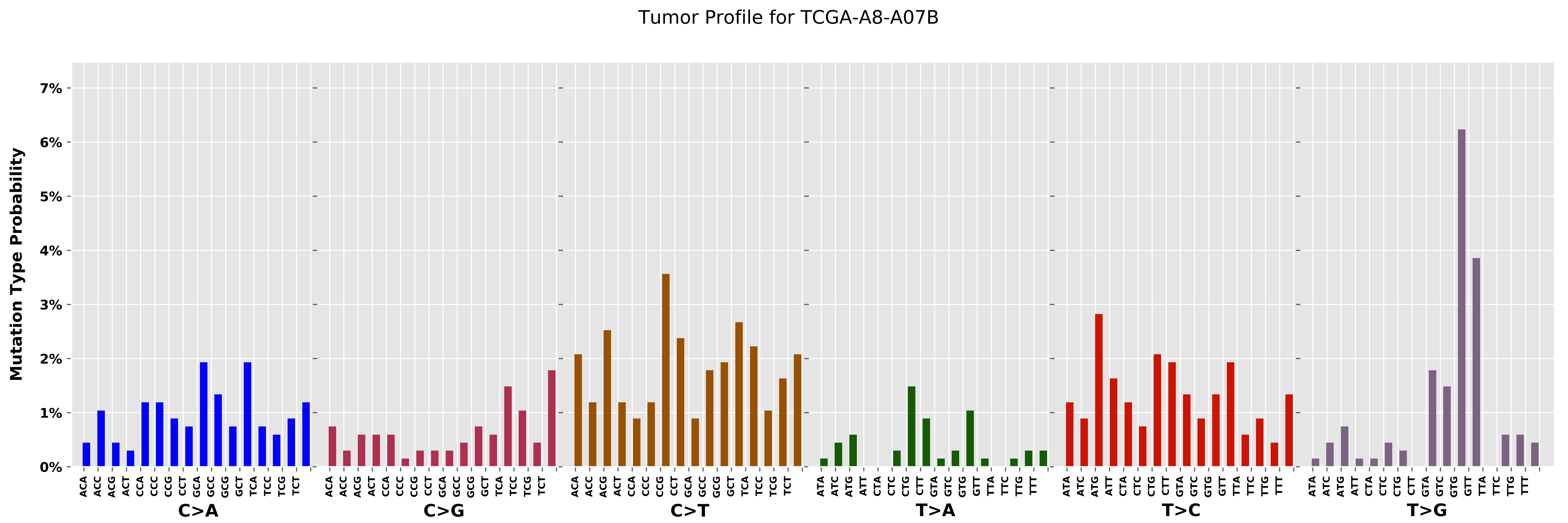
Tumor Profile for TCGA-A8-A07B
Mutation Type Probability
0%
1%
2%
3%
4%
5%
6%
7%
C>A
C>G
C>T
T>A
T>C
T>G

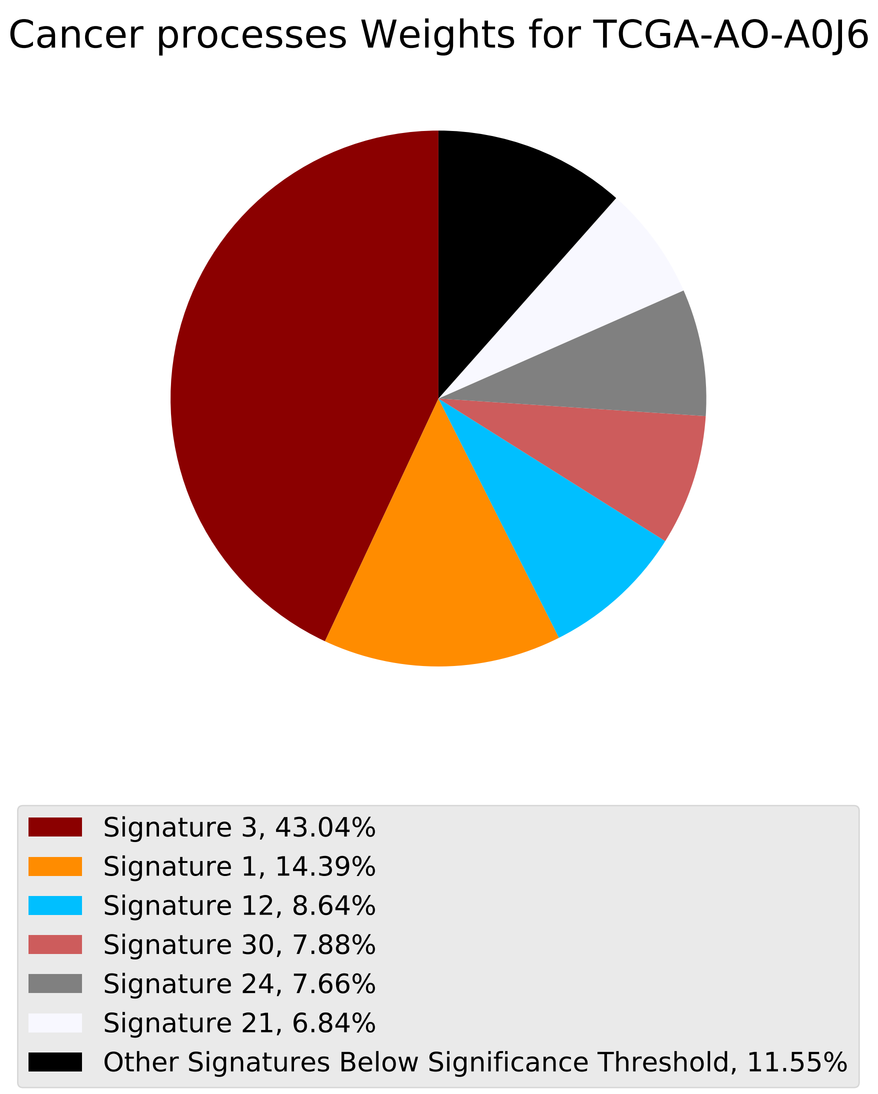
Cancer processes Weights for TCGA-AO-A0J6
Signature 3, 43.04%
Signature 1, 14.39%
Signature 12, 8.64%
Signature 30, 7.88%
Signature 24, 7.66%
Signature 21, 6.84%
Other Signatures Below Significance Threshold, 11.55%

### Tumor Profile for TCGA-AO-A0J6

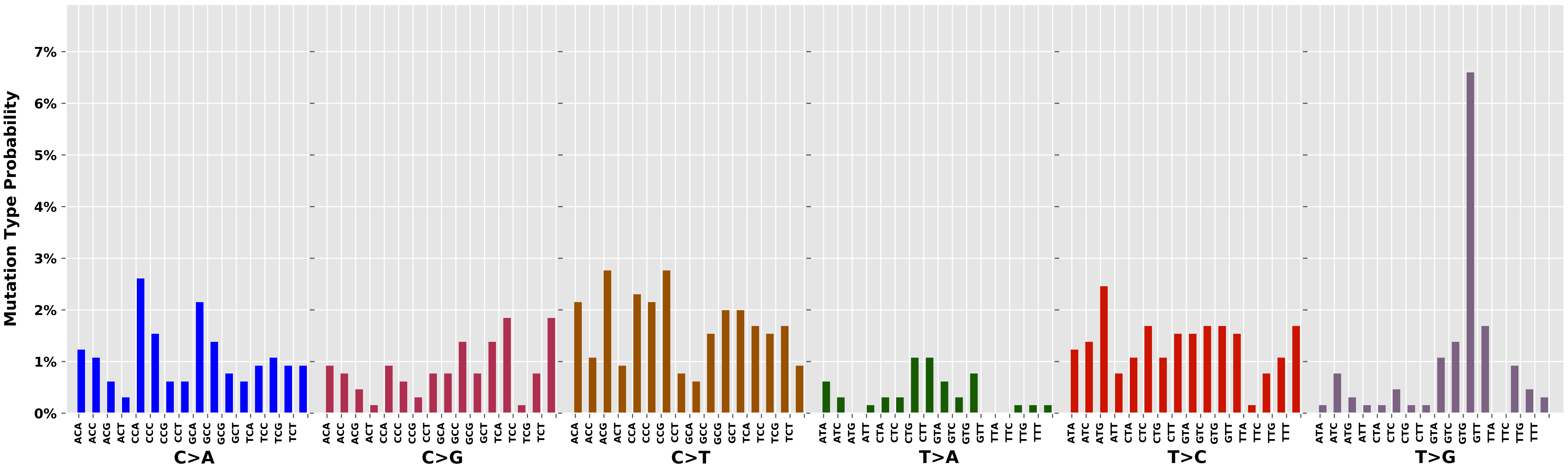

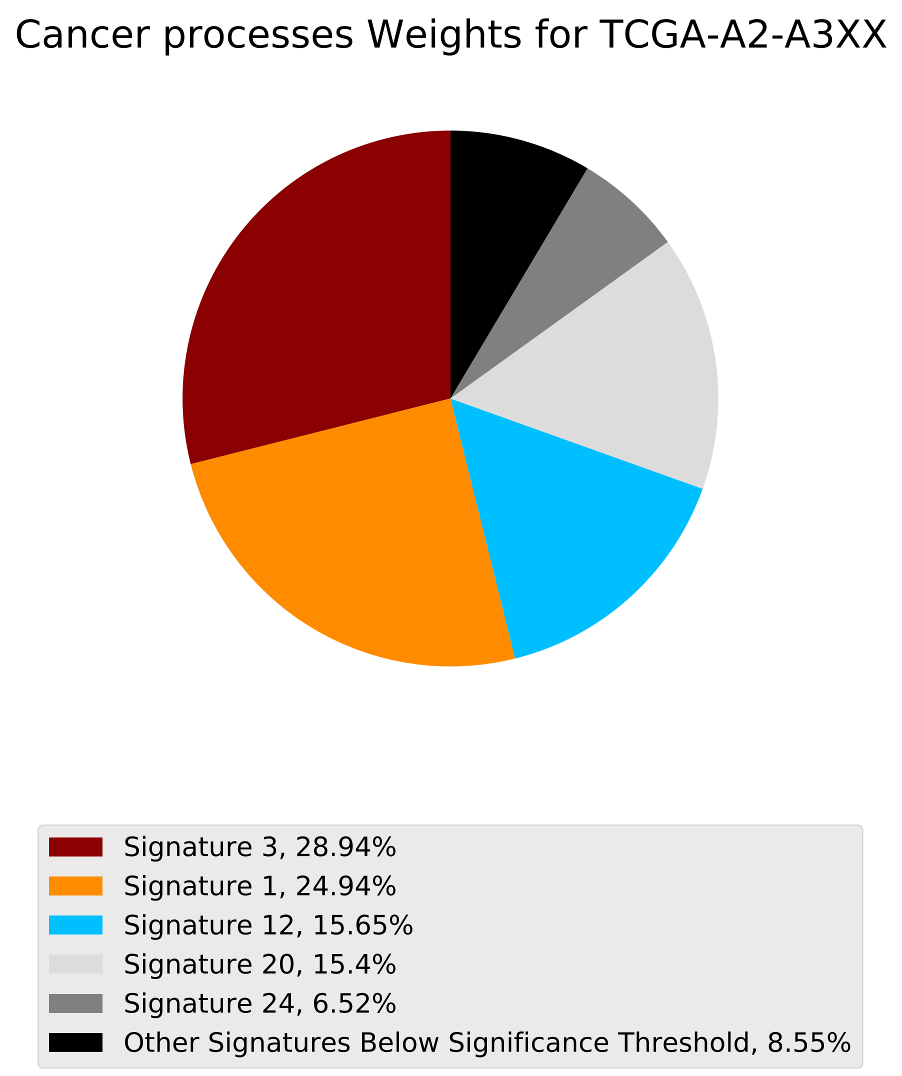
Cancer processes Weights for TCGA-A2-A3XX
Signature 3, 28.94%
Signature 1, 24.94%
Signature 12, 15.65%
Signature 20, 15.4%
Signature 24, 6.52%
Other Signatures Below Significance Threshold, 8.55%

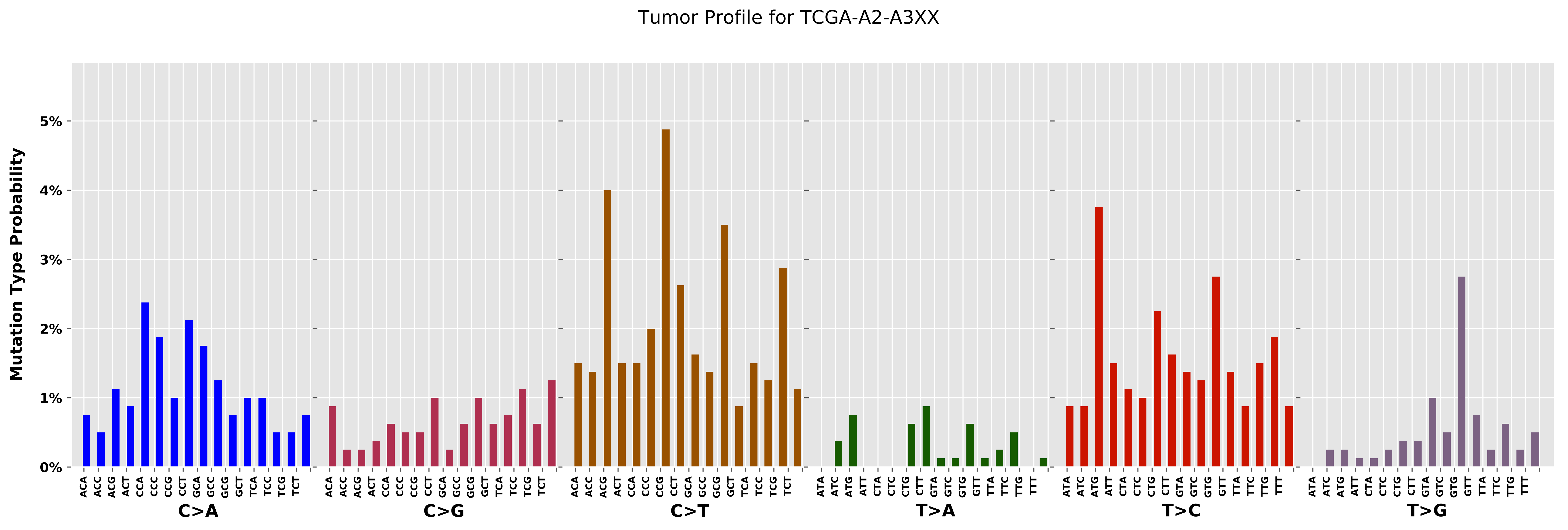
Tumor Profile for TCGA-A2-A3XX
Mutation Type Probability
0%
1%
2%
3%
4%
5%
C>A
C>G
C>T
T>A
T>C
T>G
ACA ACC ACG ACT CCA CCC CCG CCT GCA GCC GCG GCT TCA TCC TCG TCT
ATA ATC ATG ATT CTA CTC CTG CTT GTA GTC GTG GTT TTA TTC TTG TTT

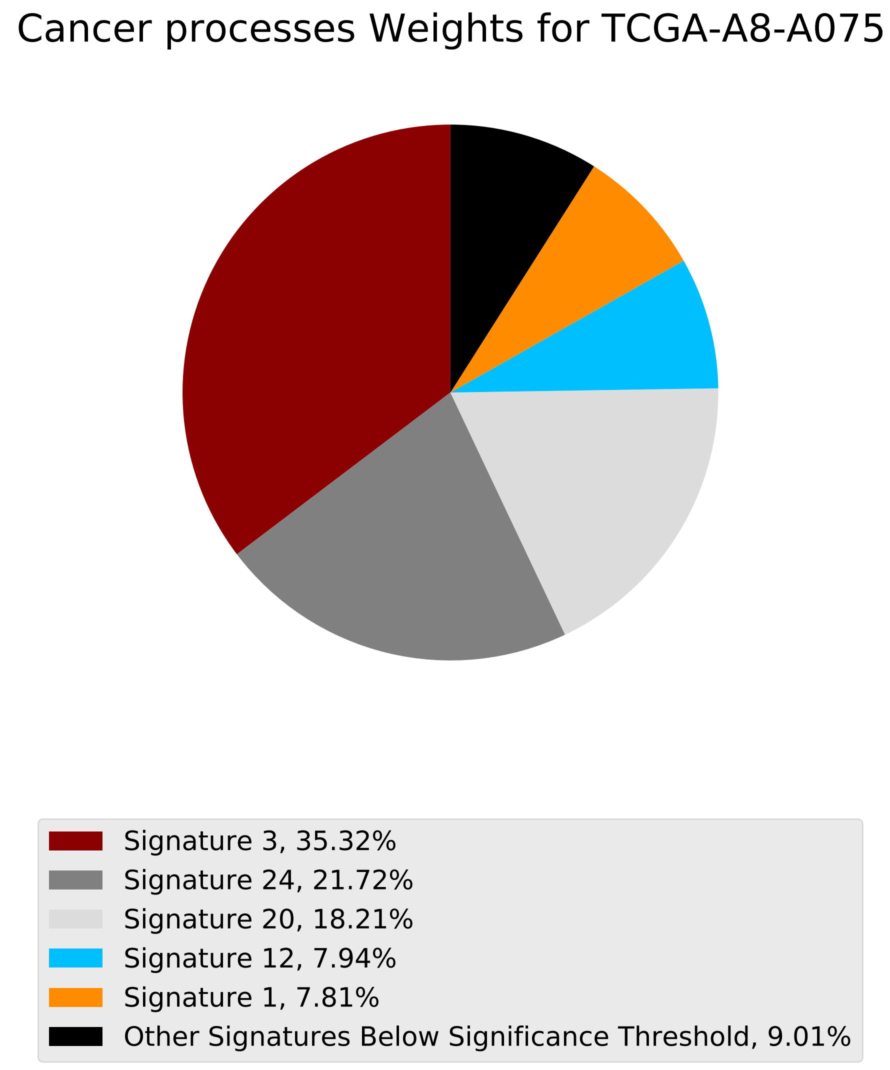

Cancer processes Weights for TCGA-A8-A075
Signature 3, 35.32%
Signature 24, 21.72%
Signature 20, 18.21%
Signature 12, 7.94%
Signature 1, 7.81%
Other Signatures Below Significance Threshold, 9.01%

Tumor Profile for TCGA-A8-A075

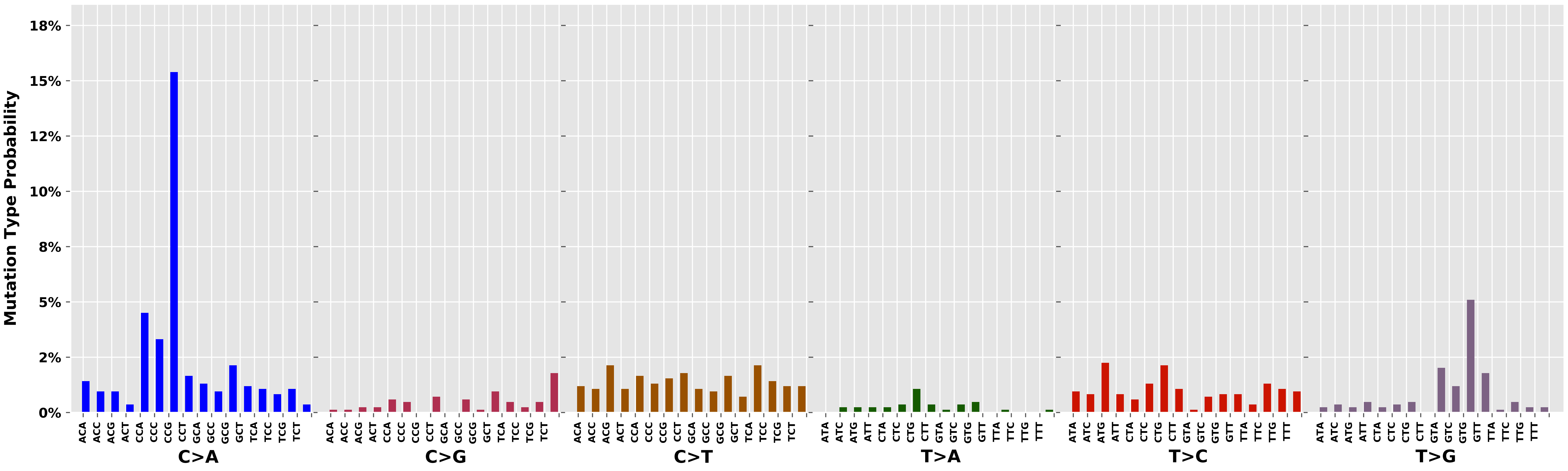

Cancer processes Weights for TCGA-AR-A1AY

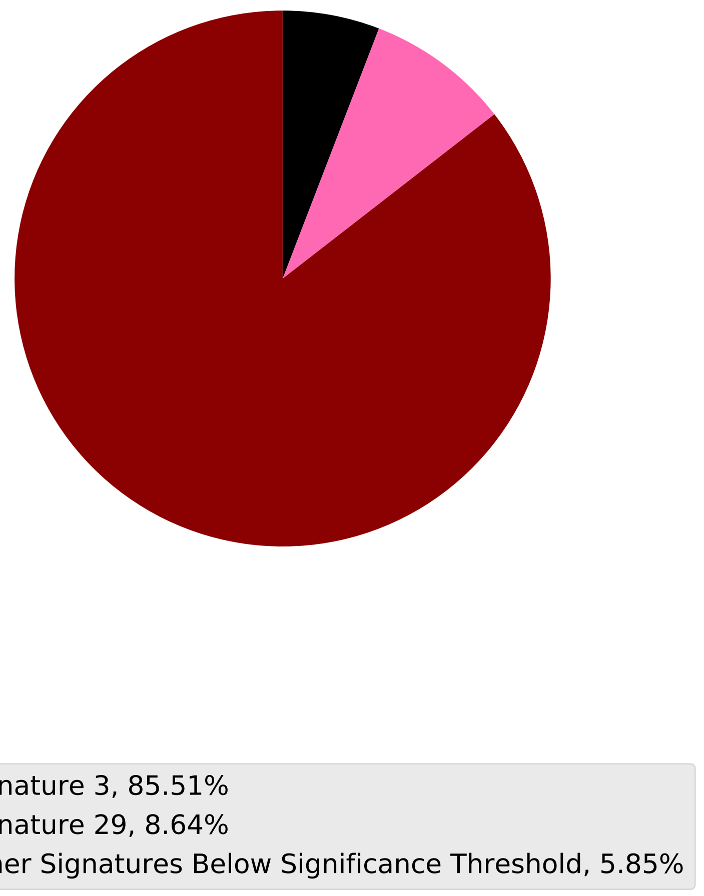

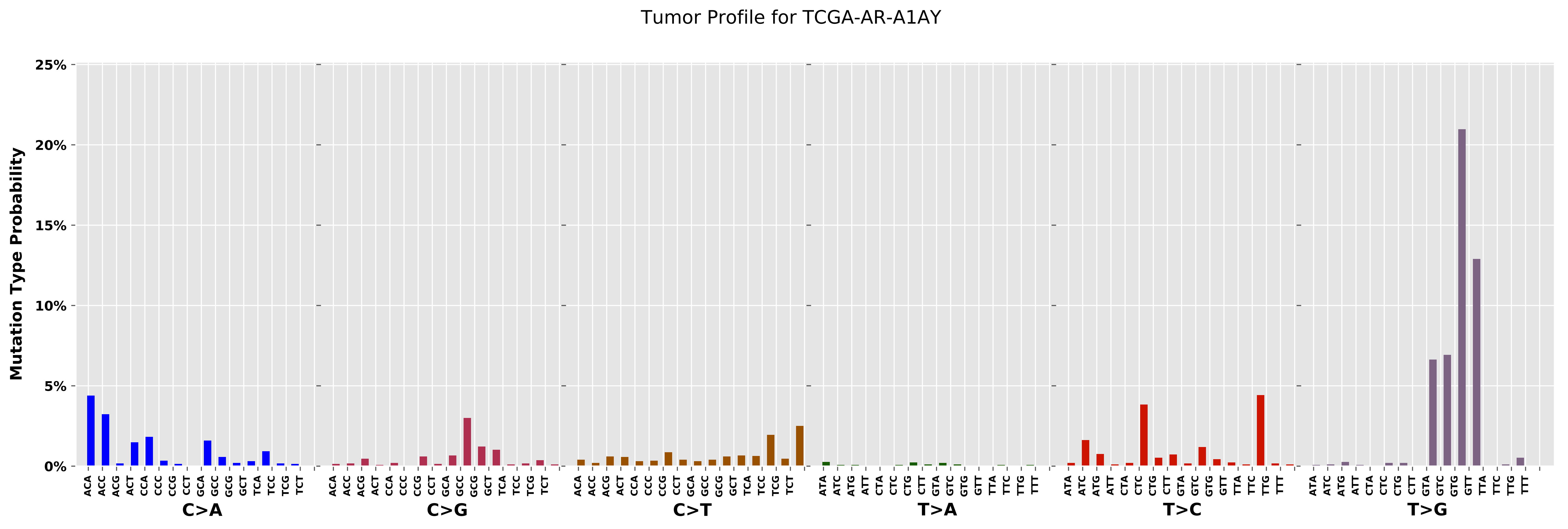
Tumor Profile for TCGA-AR-A1AY
Mutation Type Probability
25%
20%
15%
10%
5%
0%
ACA ACC ACG ACT CCA CCC CCG CCT GCA GCC GCG GCT TCA TCC TCG TCT
C>A
ACA ACC ACG ACT CCA CCC CCG CCT GCA GCC GCG GCT TCA TCC TCG TCT
C>G
ACA ACC ACG ACT CCA CCC CCG CCT GCA GCC GCG GCT TCA TCC TCG TCT
C>T
ATA ATC ATG ATT CTA CTC CTG CTT GTA GTC GTG GTT TTA TTC TTG TTT
T>A
ATA ATC ATG ATT CTA CTC CTG CTT GTA GTC GTG GTT TTA TTC TTG TTT
T>C
ATA ATC ATG ATT CTA CTC CTG CTT GTA GTC GTG GTT TTA TTC TTG TTT
T>G

Cancer processes Weights for TCGA-AG-3901

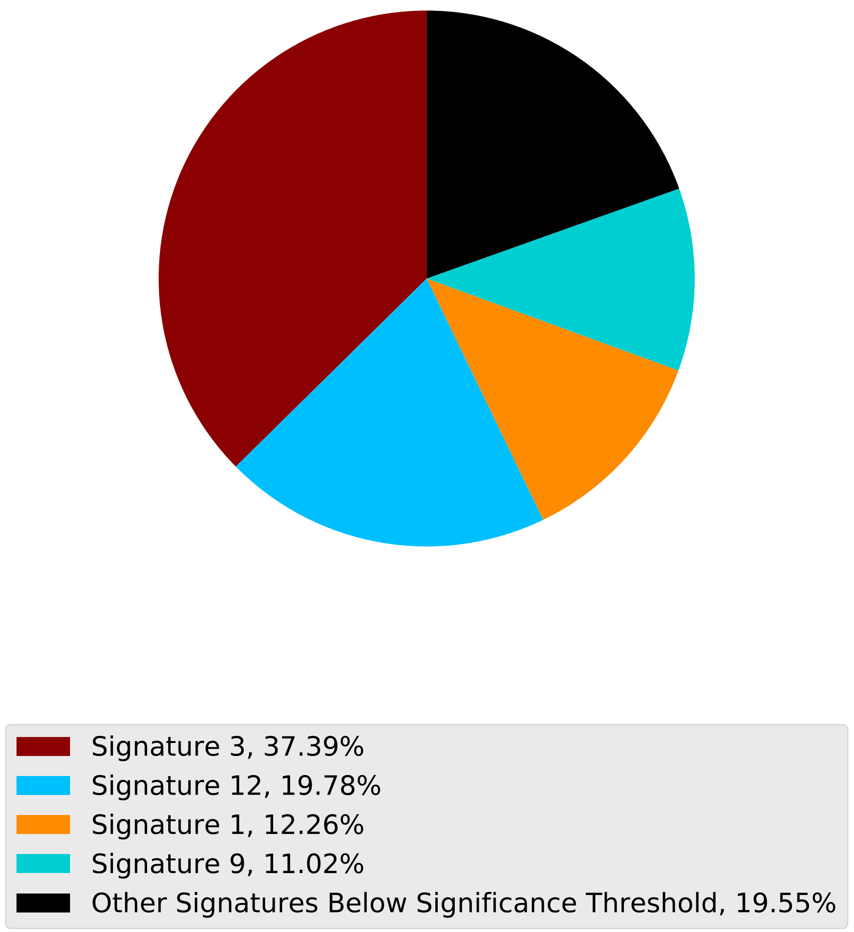

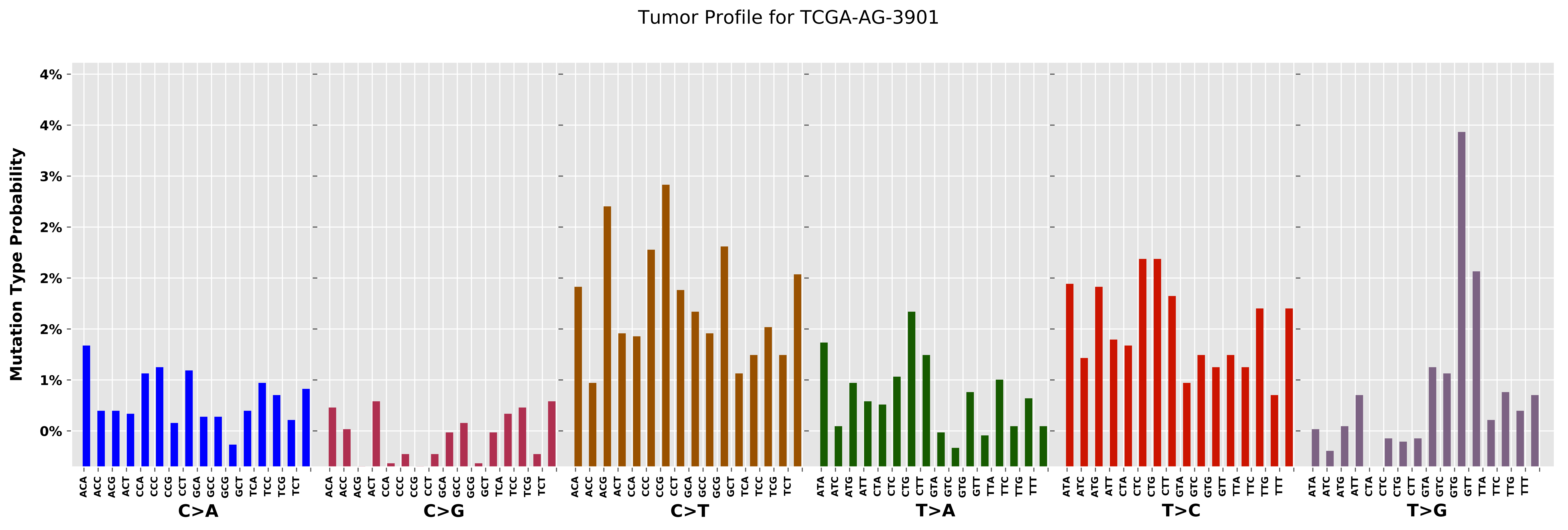

Tumor Profile for TCGA-AG-3901
Mutation Type Probability
4%
4%
3%
2%
2%
2%
1%
0%
C>A
C>G
C>T
T>A
T>C
T>G

Cancer processes Weights for TCGA-E2-A152

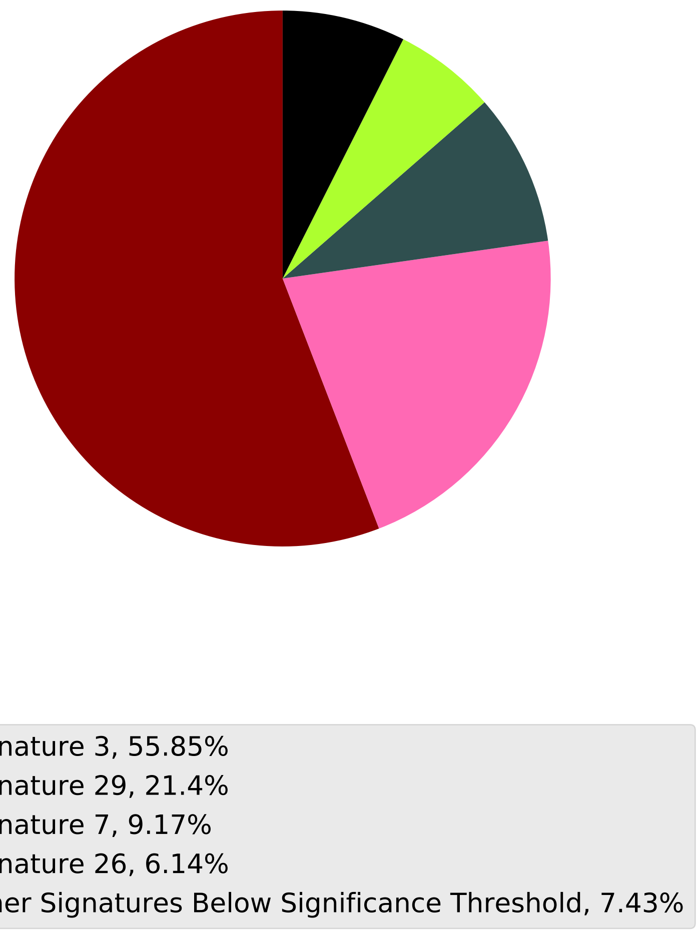

Tumor Profile for TCGA-E2-A152

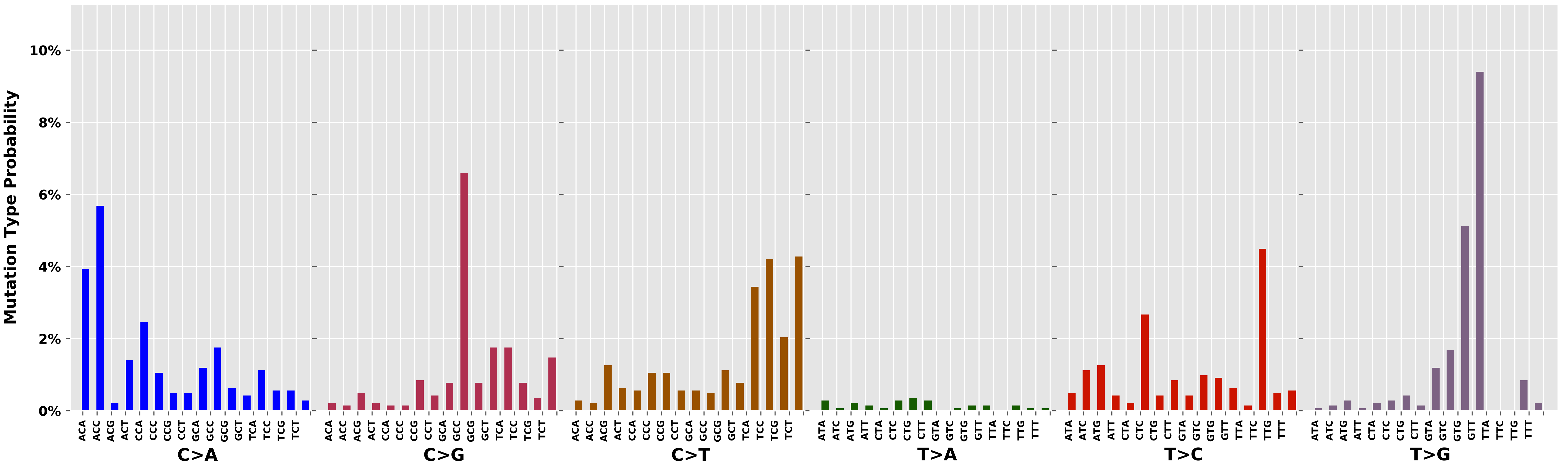

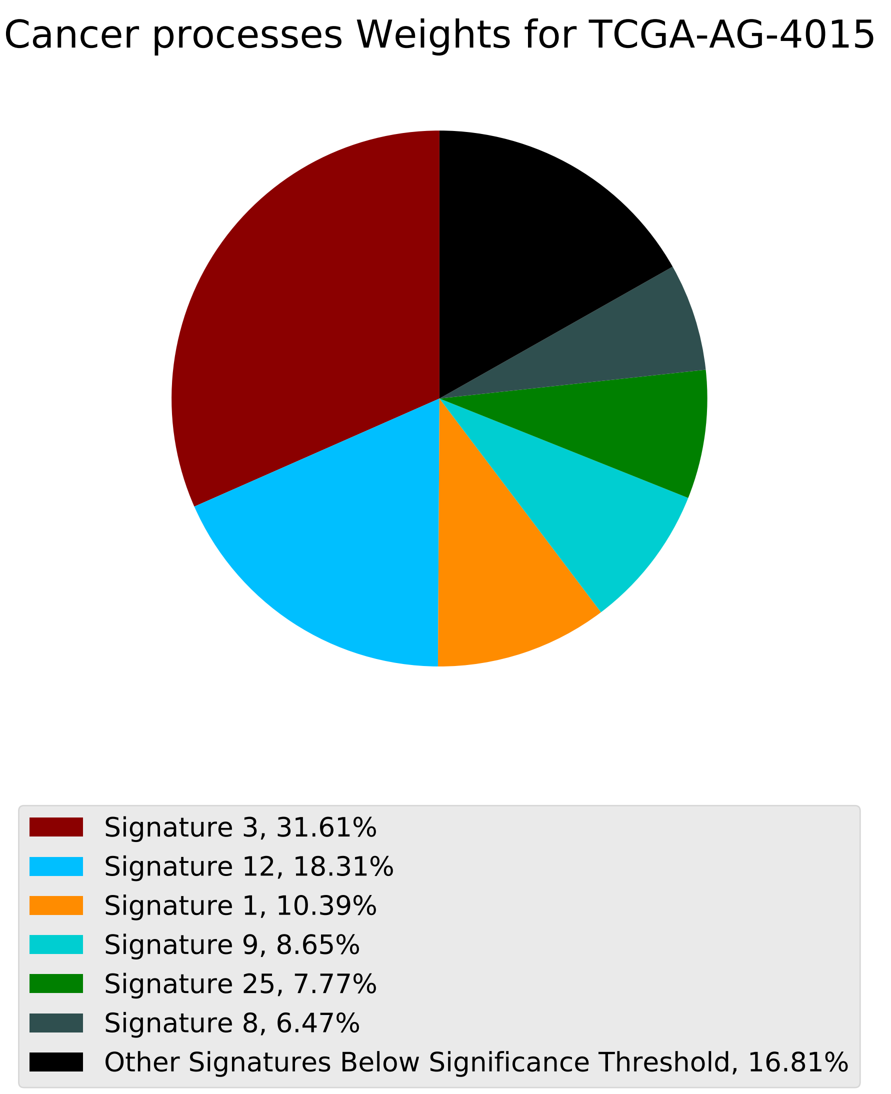
Cancer processes Weights for TCGA-AG-4015
Signature 3, 31.61%
Signature 12, 18.31%
Signature 1, 10.39%
Signature 9, 8.65%
Signature 25, 7.77%
Signature 8, 6.47%
Other Signatures Below Significance Threshold, 16.81%

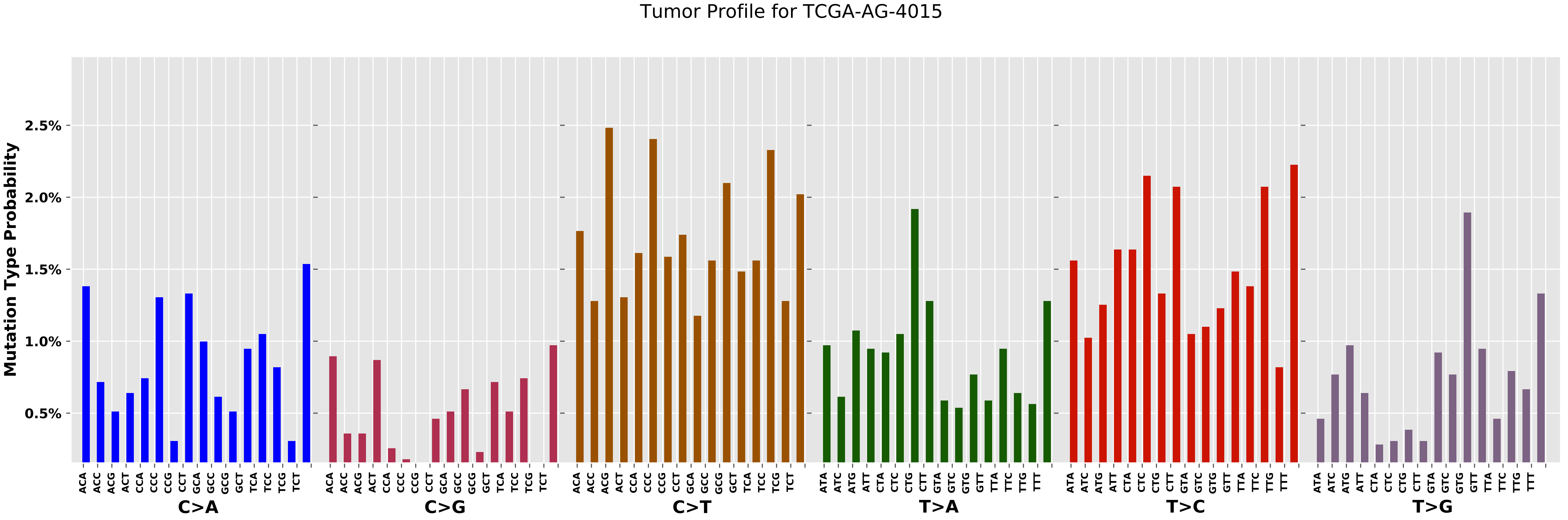
Tumor Profile for TCGA-AG-4015
Mutation Type Probability
2.5%
2.0%
1.5%
1.0%
0.5%
C>A
C>G
C>T
T>A
T>C
T>G
ACA ACC ACG ACT CCA CCC CCG CCT GCA GCC GCG GCT TCA TCC TCG TCT
ATA ATC ATG ATT CTA CTC CTG CTT GTA GTC GTG GTT TTA TTC TTG TTT

Cancer processes Weights for TCGA-BH-A0B9

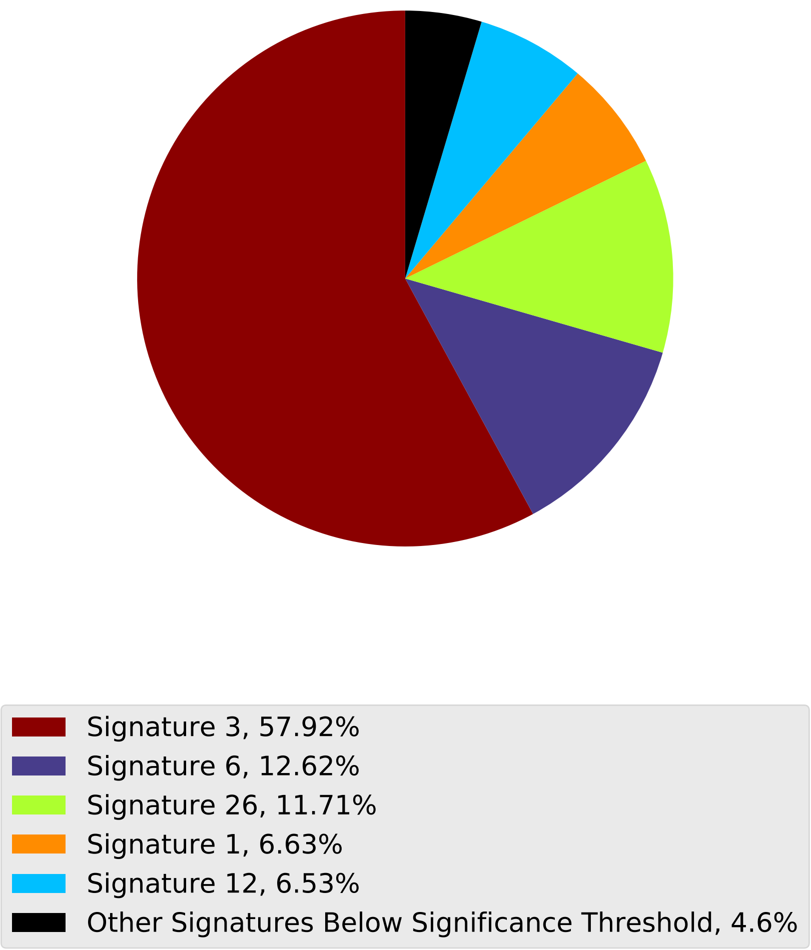

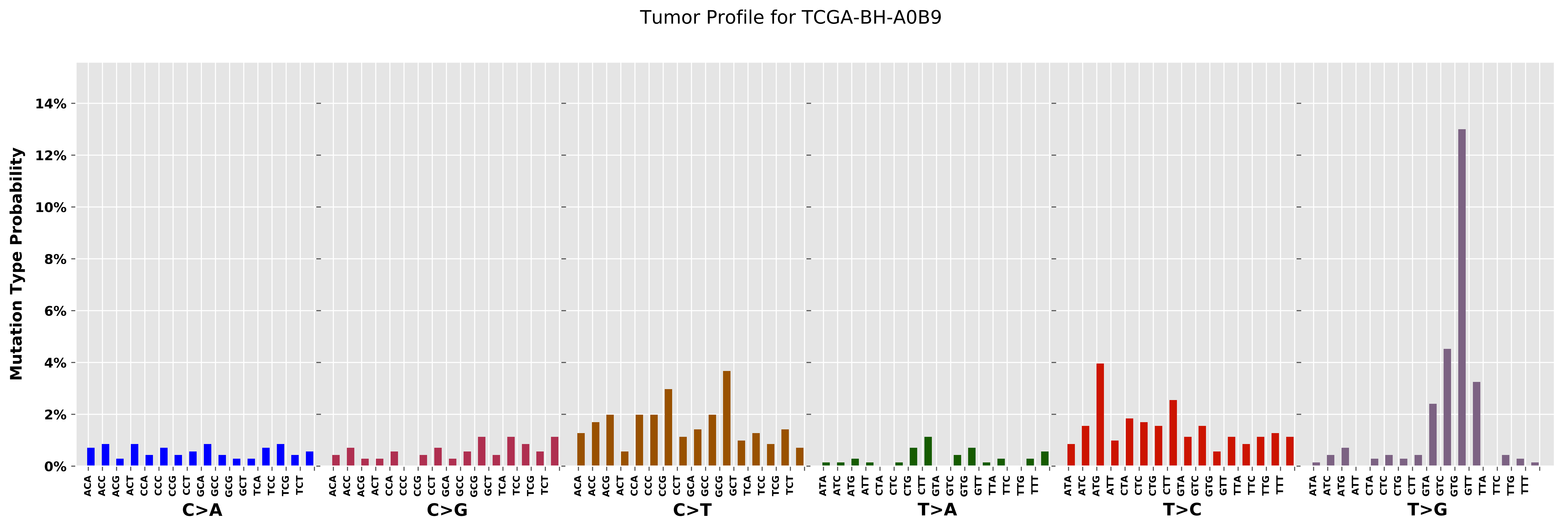

Tumor Profile for TCGA-BH-A0B9
Mutation Type Probability
0%
2%
4%
6%
8%
10%
12%
14%
ACA ACC ACG ACT CCA CCC CCG CCT GCA GCC GCG GCT TCA TCC TCG TCT
C>A
ACA ACC ACG ACT CCA CCC CCG CCT GCA GCC GCG GCT TCA TCC TCG TCT
C>G
ACA ACC ACG ACT CCA CCC CCG CCT GCA GCC GCG GCT TCA TCC TCG TCT
C>T
ATA ATC ATG ATT CTA CTC CTG CTT GTA GTC GTG GTT TTA TTC TTG TTT
T>A
ATA ATC ATG ATT CTA CTC CTG CTT GTA GTC GTG GTT TTA TTC TTG TTT
T>C
ATA ATC ATG ATT CTA CTC CTG CTT GTA GTC GTG GTT TTA TTC TTG TTT
T>G

Cancer processes Weights for TCGA-A2-A0D1

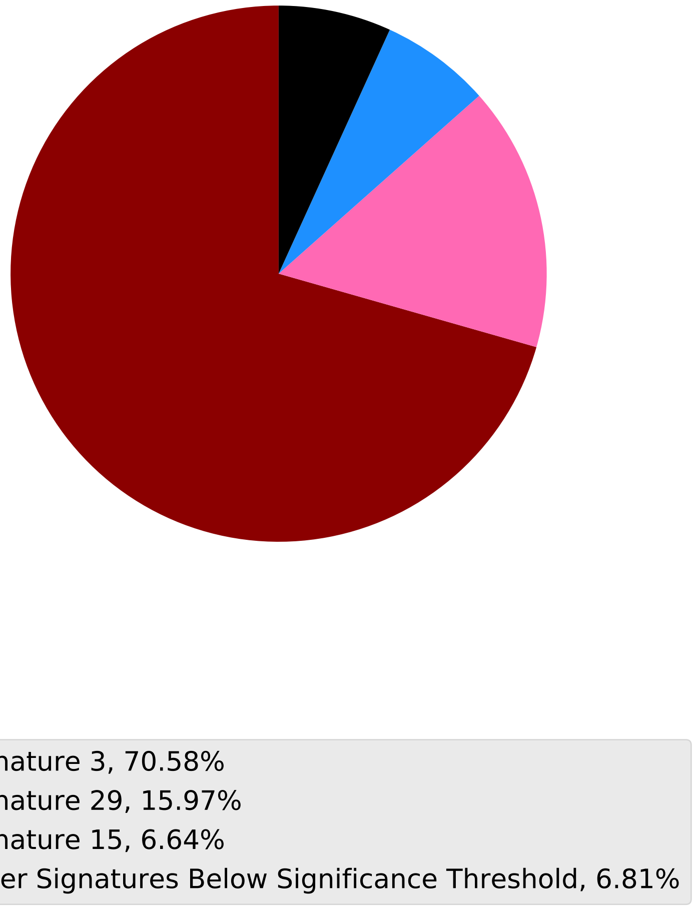

Tumor Profile for TCGA-A2-A0D1

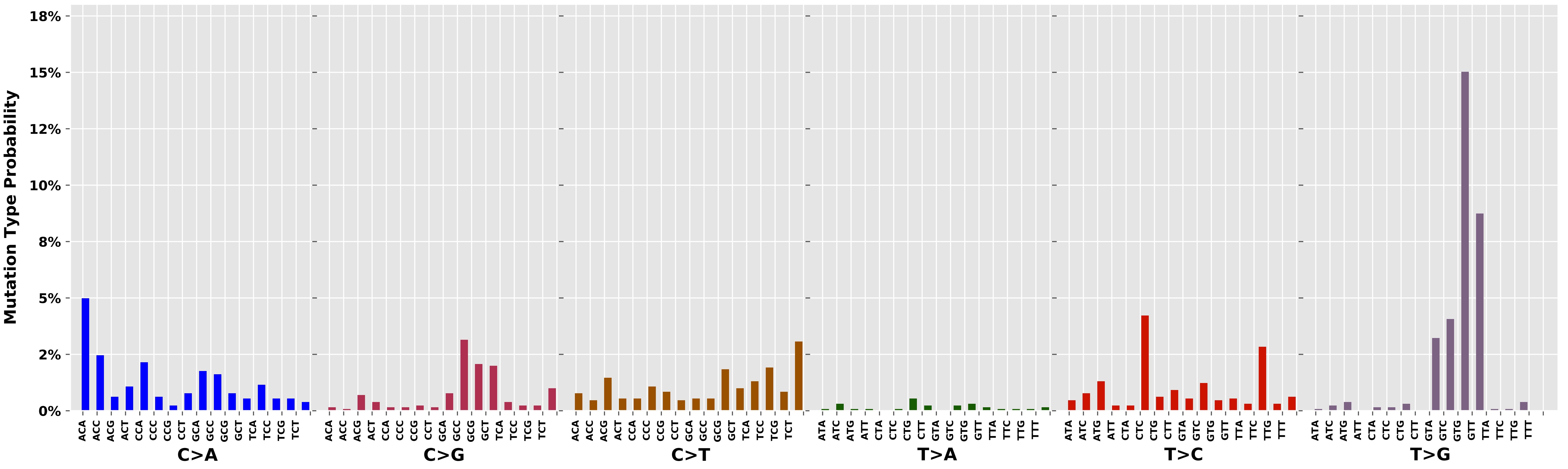

Cancer processes Weights for TCGA-AA-3529

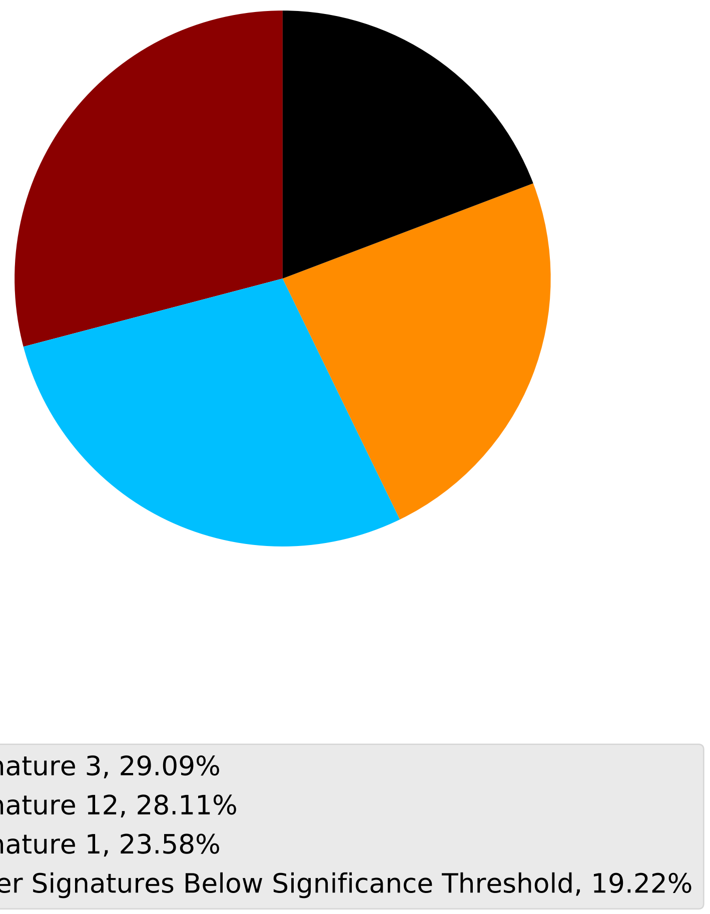

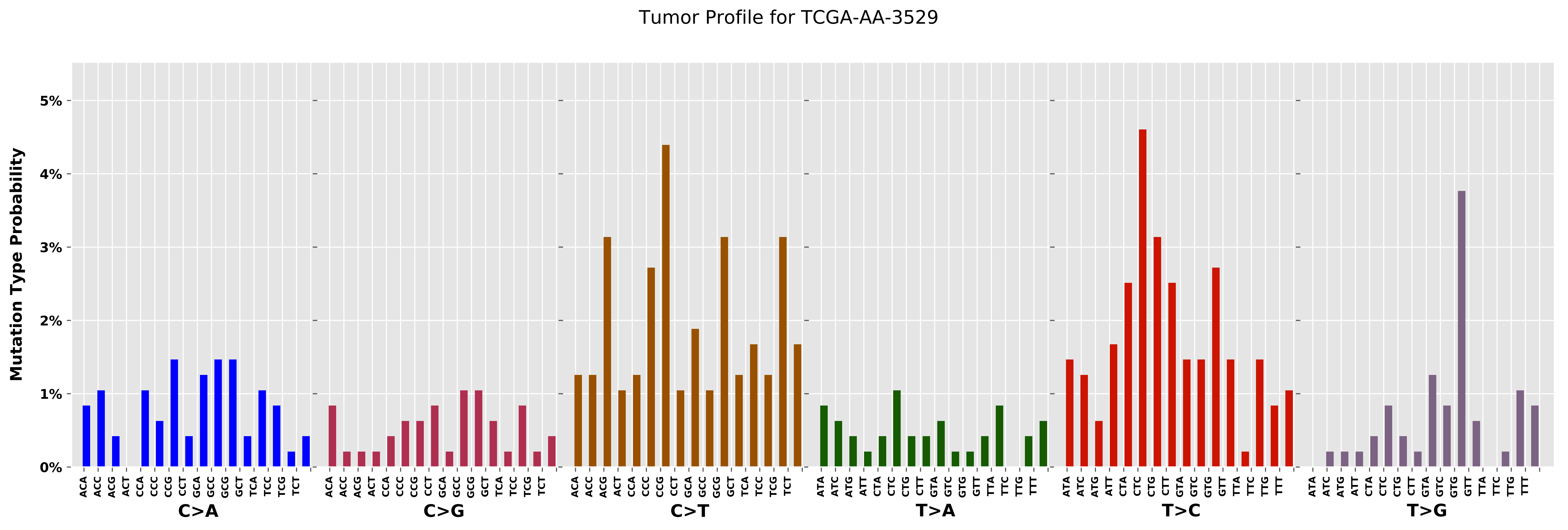
Tumor Profile for TCGA-AA-3529
Mutation Type Probability
0%
1%
2%
3%
4%
5%
C>A
C>G
C>T
T>A
T>C
T>G

Cancer processes Weights for TCGA-A6-3807

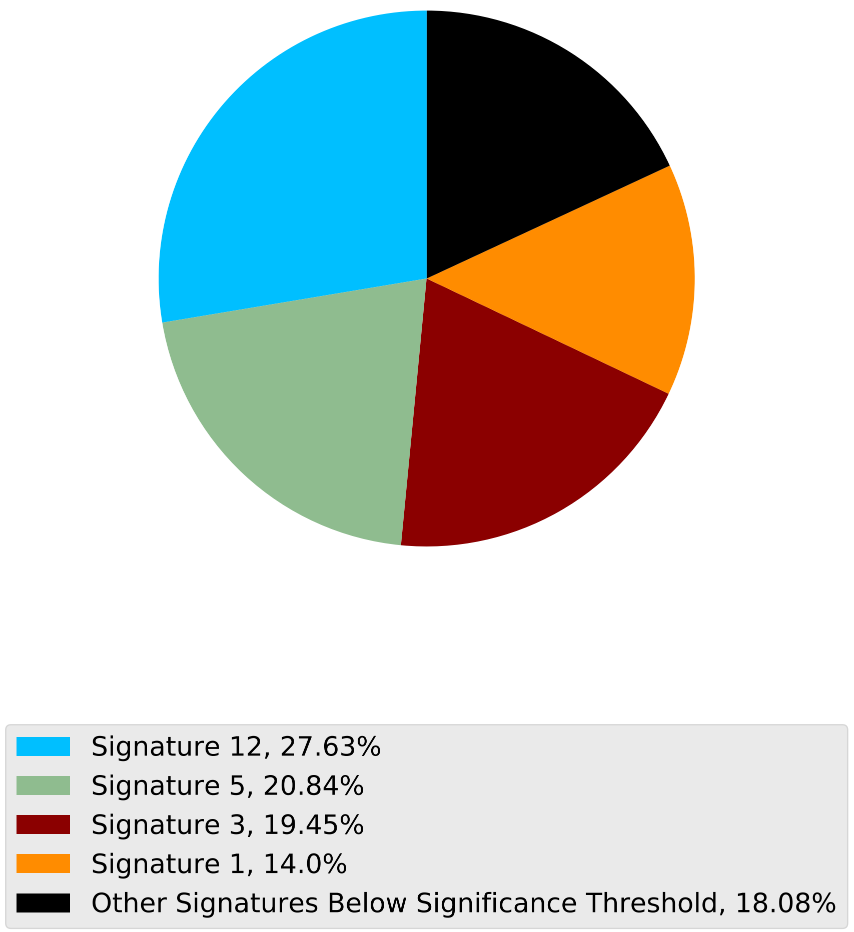

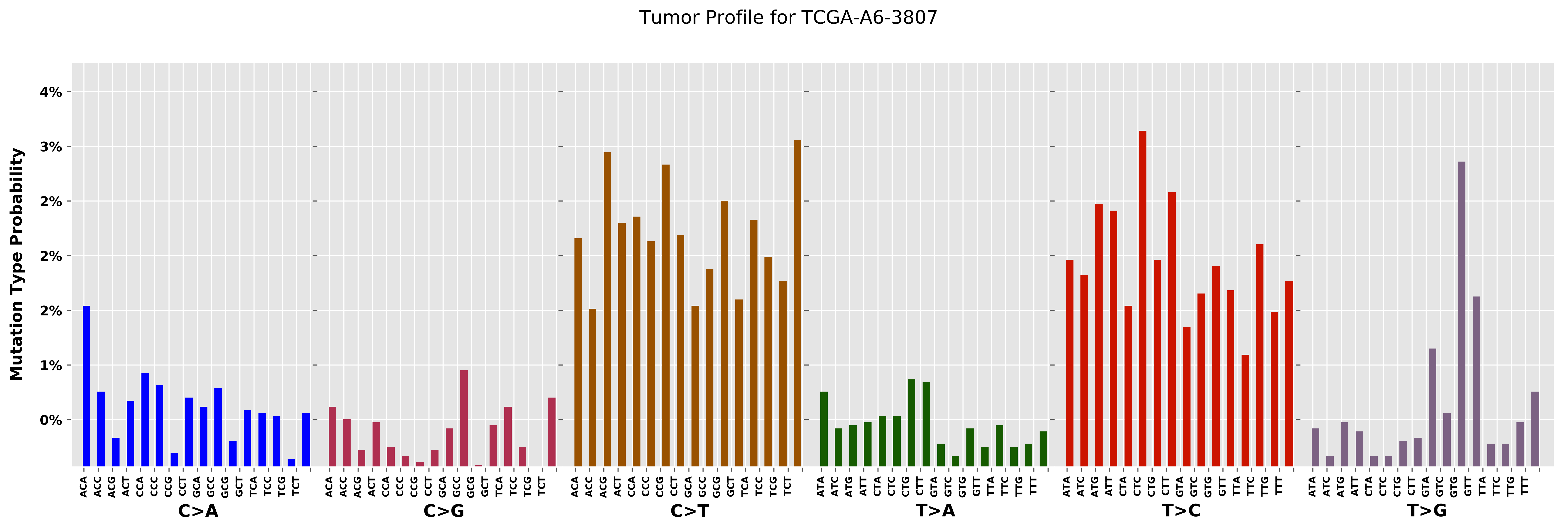

Tumor Profile for TCGA-A6-3807
Mutation Type Probability
4%
3%
2%
2%
2%
1%
0%
C>A
C>G
C>T
T>A
T>C
T>G
ACA ACC ACG ACT CCA CCC CCG CCT GCA GCC GCG GCT TCA TCC TCG TCT
ATA ATC ATG ATT CTA CTC CTG CTT GTA GTC GTG GTT TTA TTC TTG TTT

#### Cancer processes Weights for TCGA-A2-A0YG

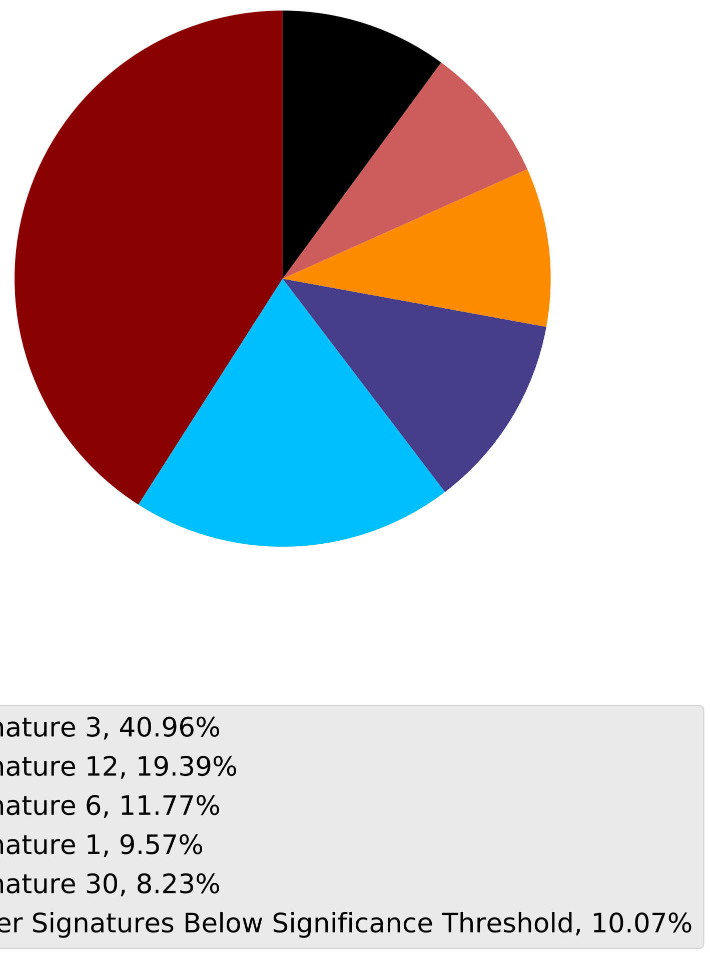

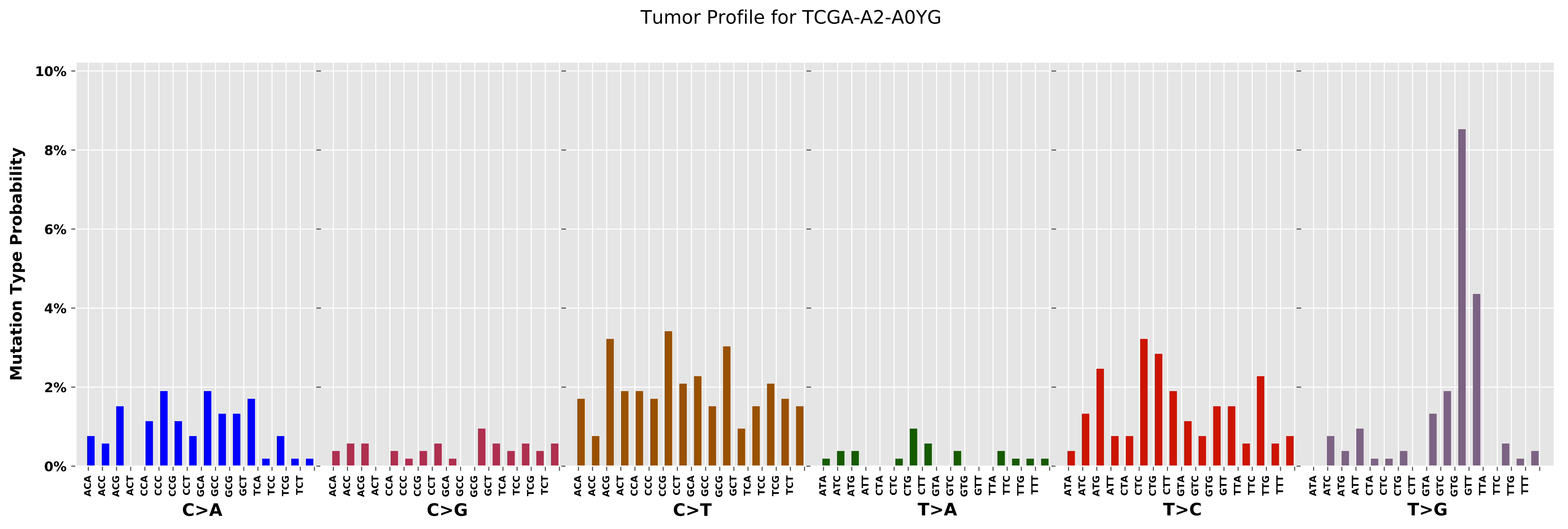

Tumor Profile for TCGA-A2-A0YG
Mutation Type Probability
10%
8%
6%
4%
2%
0%
C>A
C>G
C>T
T>A
T>C
T>G

Cancer processes Weights for TCGA-C8-A12Q

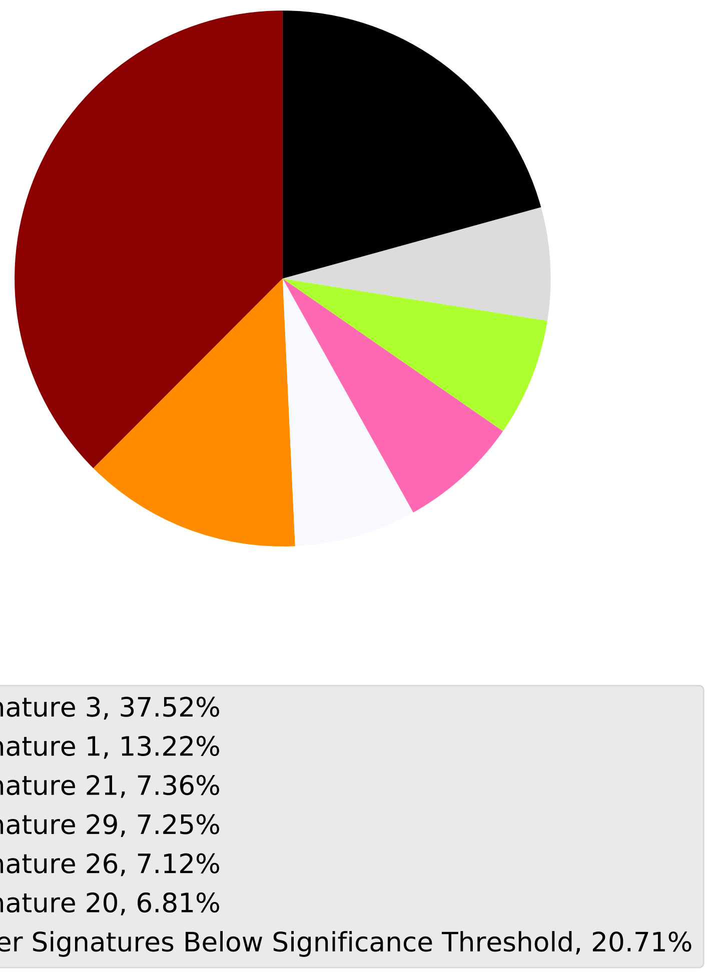

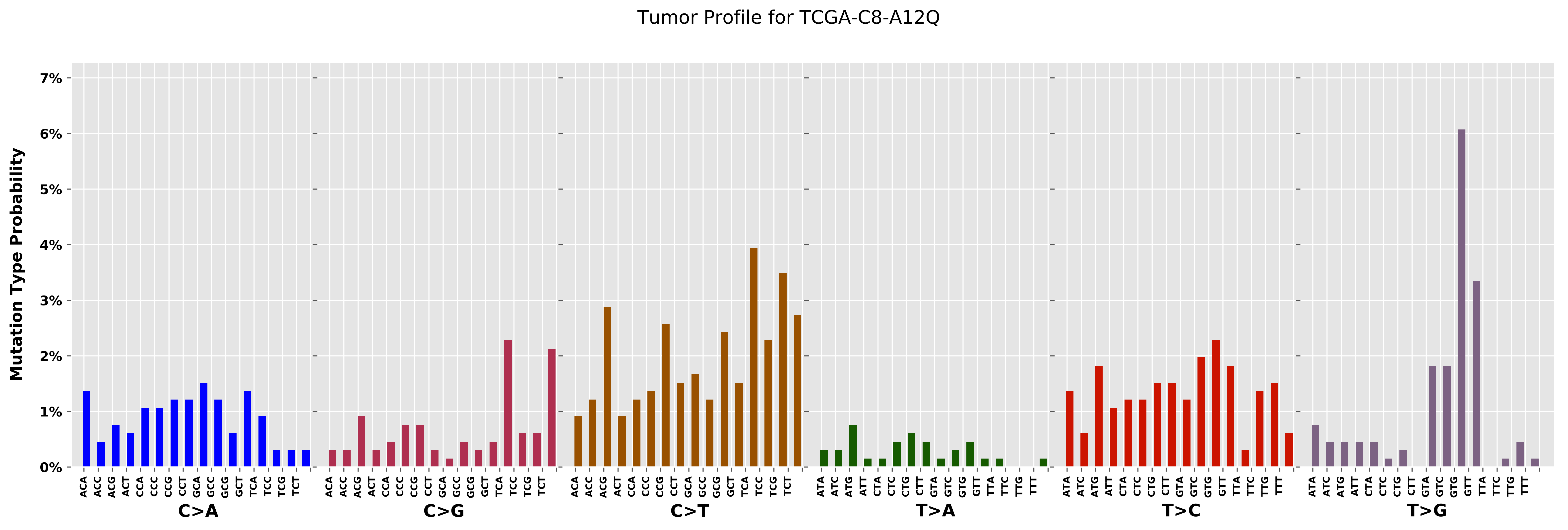
Tumor Profile for TCGA-C8-A12Q
Mutation Type Probability
0%
1%
2%
3%
4%
5%
6%
7%
C>A
C>G
C>T
T>A
T>C
T>G
ACA ACC ACG ACT CCA CCC CCG CCT GCA GCC GCG GCT TCA TCC TCG TCT
ATA ATC ATG ATT CTA CTC CTG CTT GTA GTC GTG GTT TTA TTC TTG TTT

Cancer processes Weights for TCGA-BH-A0BW

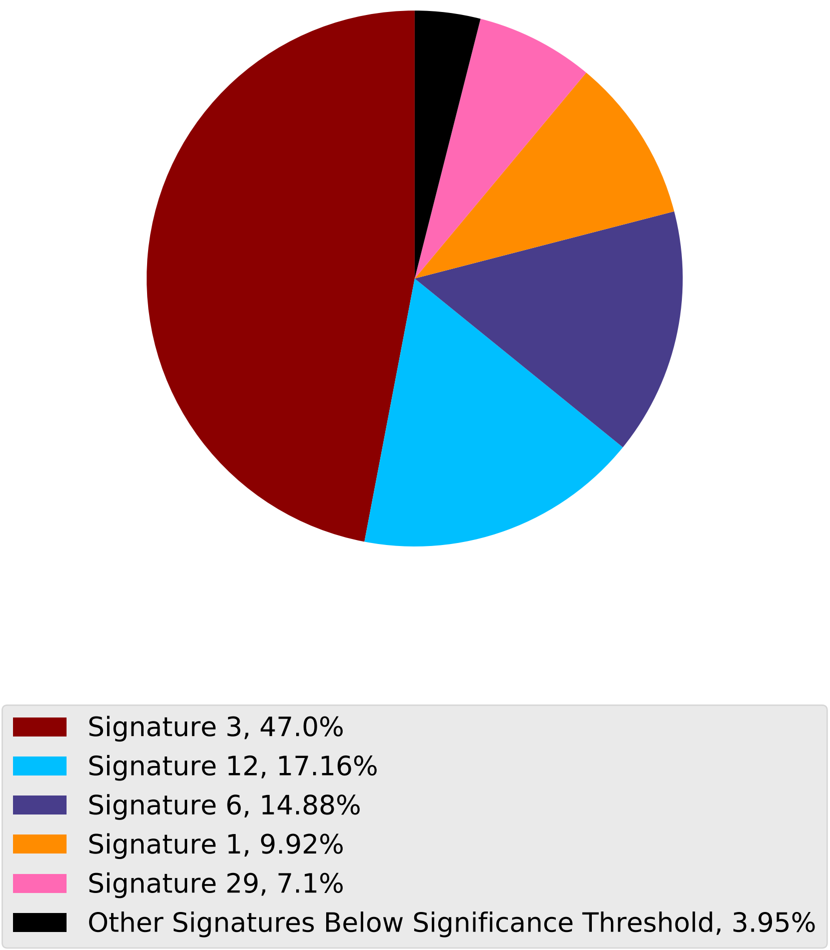

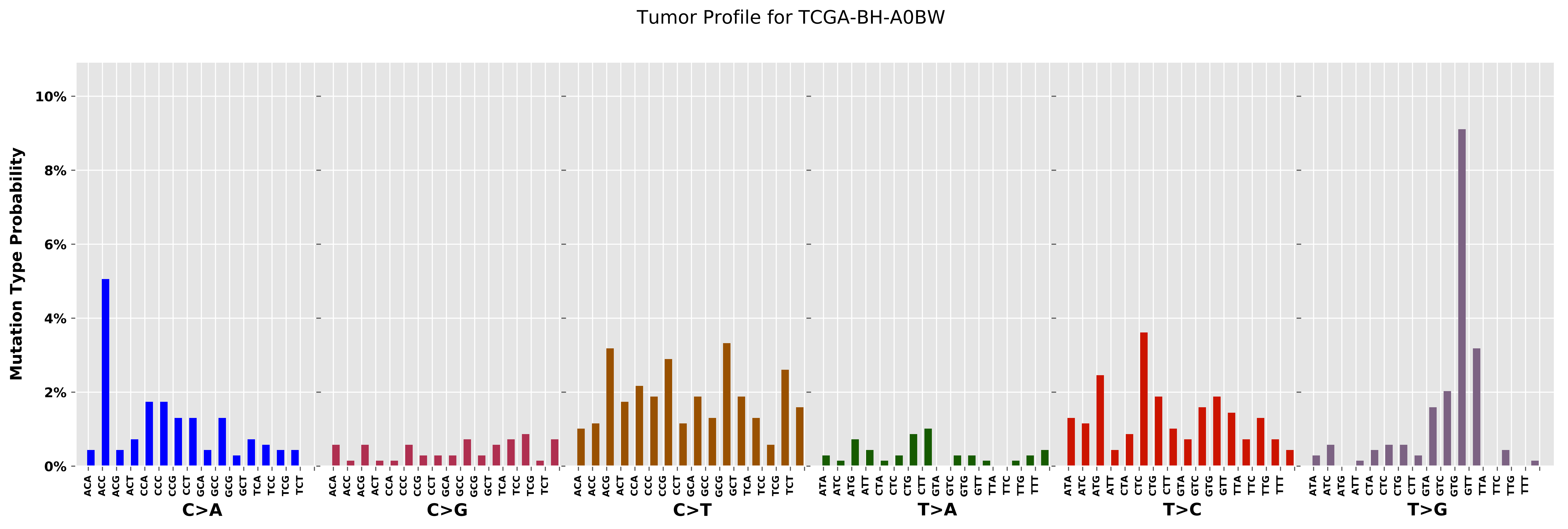
Tumor Profile for TCGA-BH-A0BW
Mutation Type Probability
10%
8%
6%
4%
2%
0%
ACA ACC ACG ACT CCA CCC CCG CCT GCA GCC GCG GCT TCA TCC TCG TCT
C>A
ACA ACC ACG ACT CCA CCC CCG CCT GCA GCC GCG GCT TCA TCC TCG TCT
C>G
ACA ACC ACG ACT CCA CCC CCG CCT GCA GCC GCG GCT TCA TCC TCG TCT
C>T
ATA ATC ATG ATT CTA CTC CTG CTT GTA GTC GTG GTT TTA TTC TTG TTT
T>A
ATA ATC ATG ATT CTA CTC CTG CTT GTA GTC GTG GTT TTA TTC TTG TTT
T>C
ATA ATC ATG ATT CTA CTC CTG CTT GTA GTC GTG GTT TTA TTC TTG TTT
T>G

Cancer processes Weights for TCGA-A2-A0EY

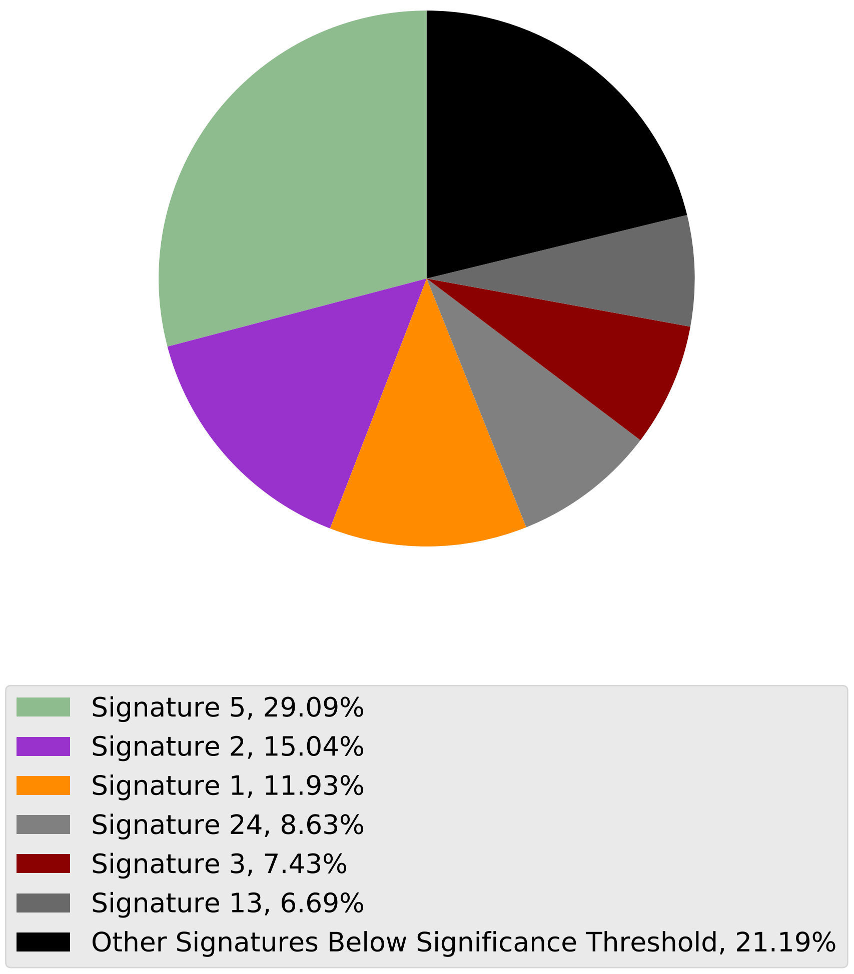

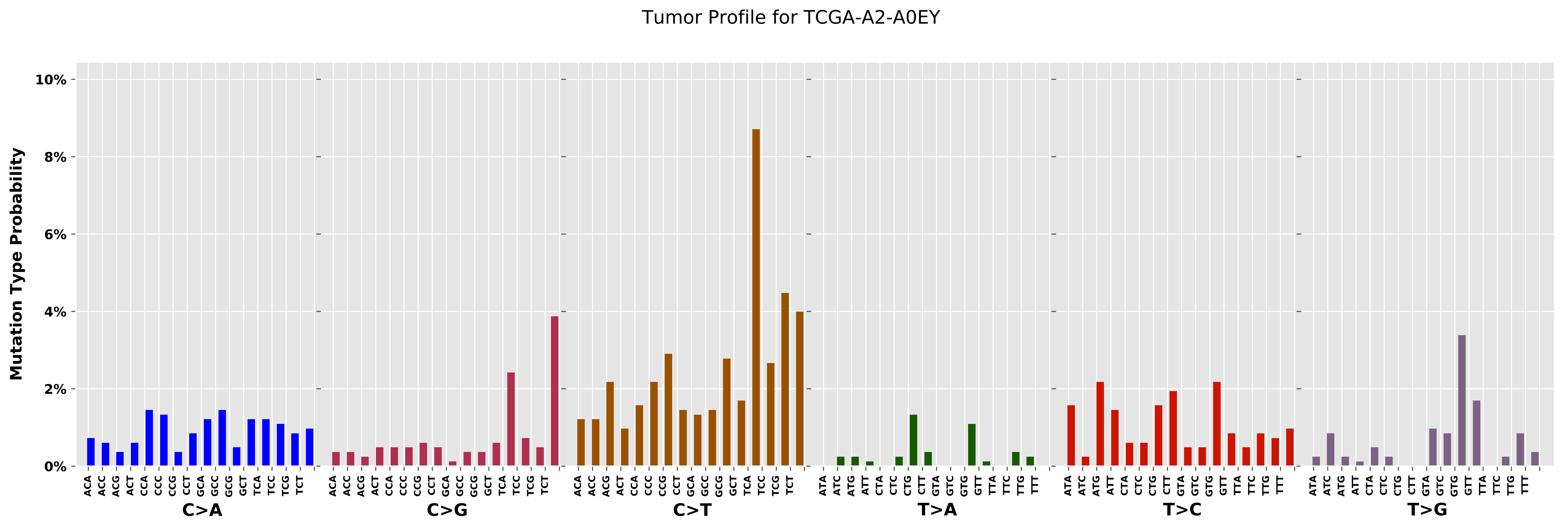

Tumor Profile for TCGA-A2-A0EY
Mutation Type Probability
0%
2%
4%
6%
8%
10%
ACA ACC ACG ACT CCA CCC CCG CCT GCA GCC GCG GCT TCA TCC TCG TCT
C>A
ACA ACC ACG ACT CCA CCC CCG CCT GCA GCC GCG GCT TCA TCC TCG TCT
C>G
ACA ACC ACG ACT CCA CCC CCG CCT GCA GCC GCG GCT TCA TCC TCG TCT
C>T
ATA ATC ATG ATT CTA CTC CTG CTT GTA GTC GTG GTT TTA TTC TTG TTT
T>A
ATA ATC ATG ATT CTA CTC CTG CTT GTA GTC GTG GTT TTA TTC TTG TTT
T>C
ATA ATC ATG ATT CTA CTC CTG CTT GTA GTC GTG GTT TTA TTC TTG TTT
T>G

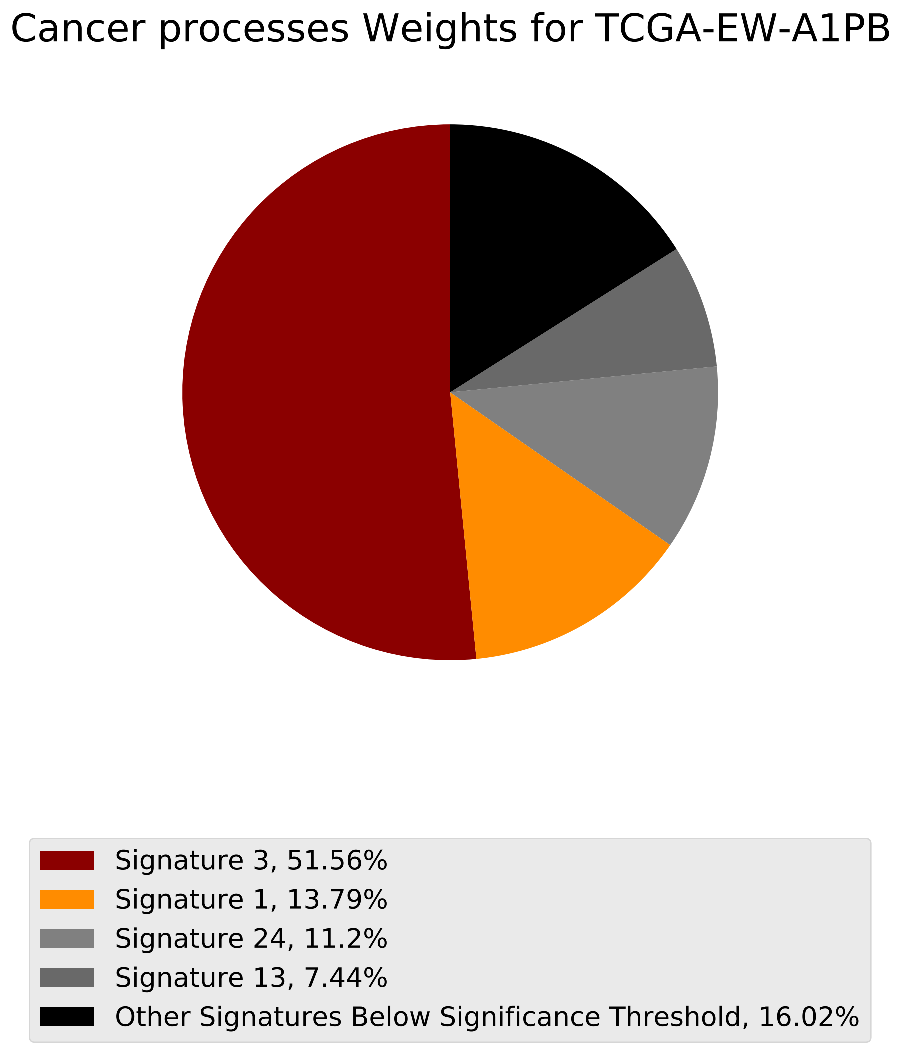
Cancer processes Weights for TCGA-EW-A1PB
Signature 3, 51.56%
Signature 1, 13.79%
Signature 24, 11.2%
Signature 13, 7.44%
Other Signatures Below Significance Threshold, 16.02%

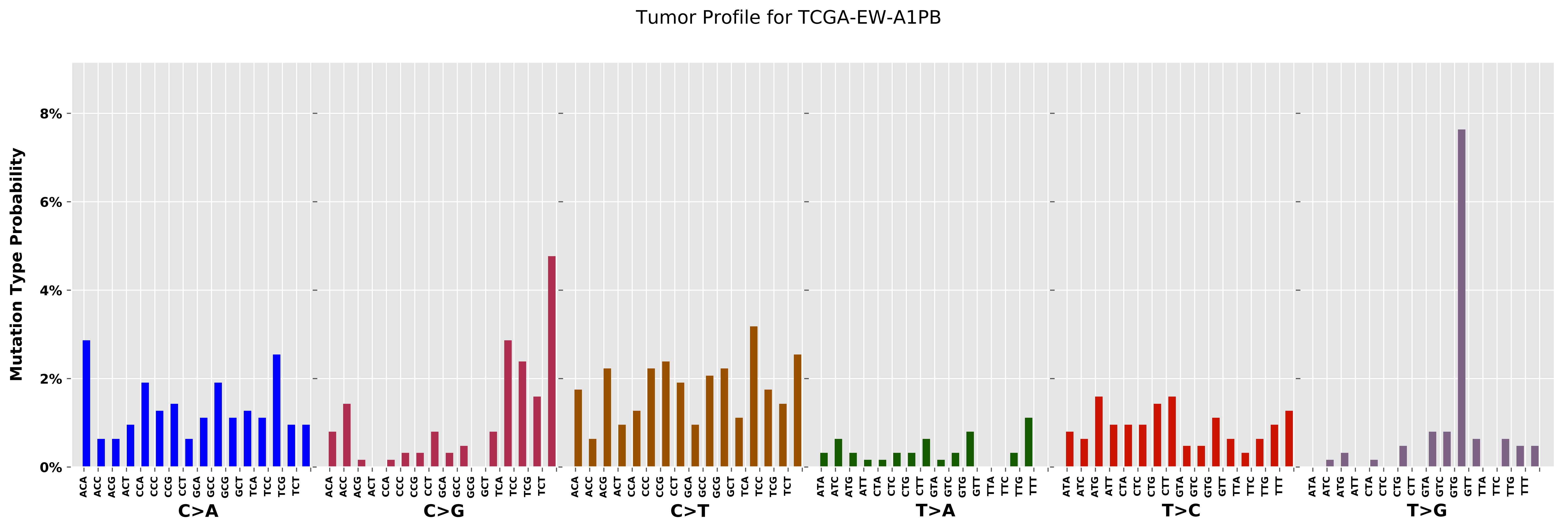

Tumor Profile for TCGA-EW-A1PB
Mutation Type Probability
8%
6%
4%
2%
0%
C>A
C>G
C>T
T>A
T>C
T>G

Cancer processes Weights for TCGA-AD-6964

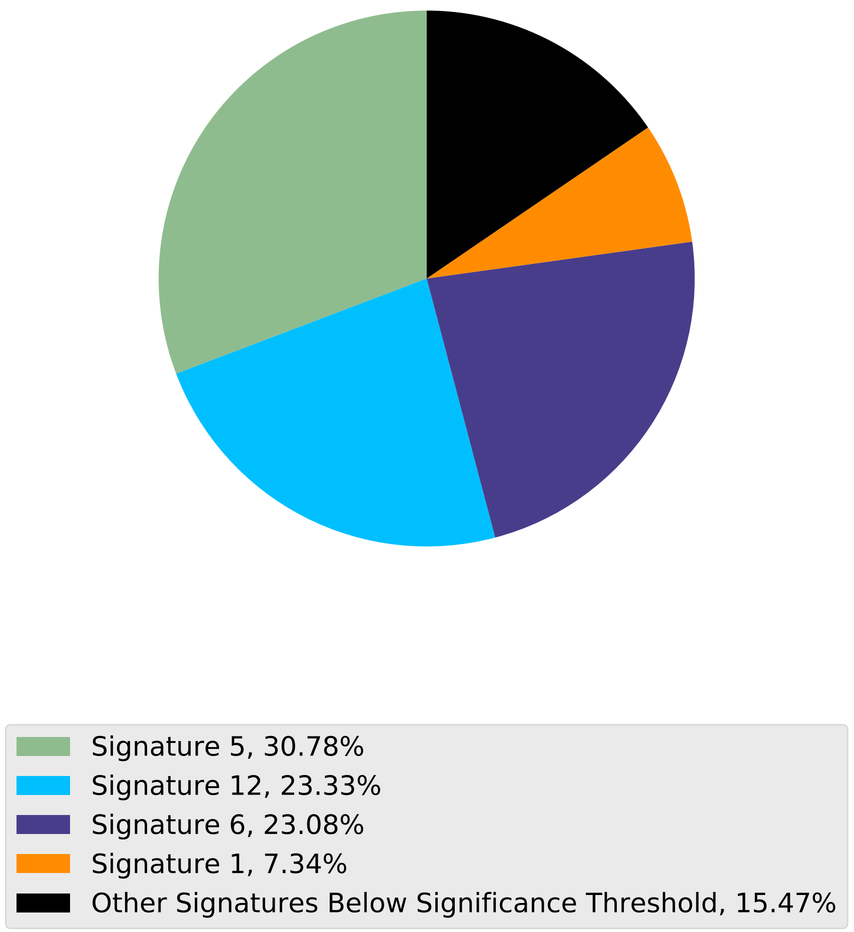

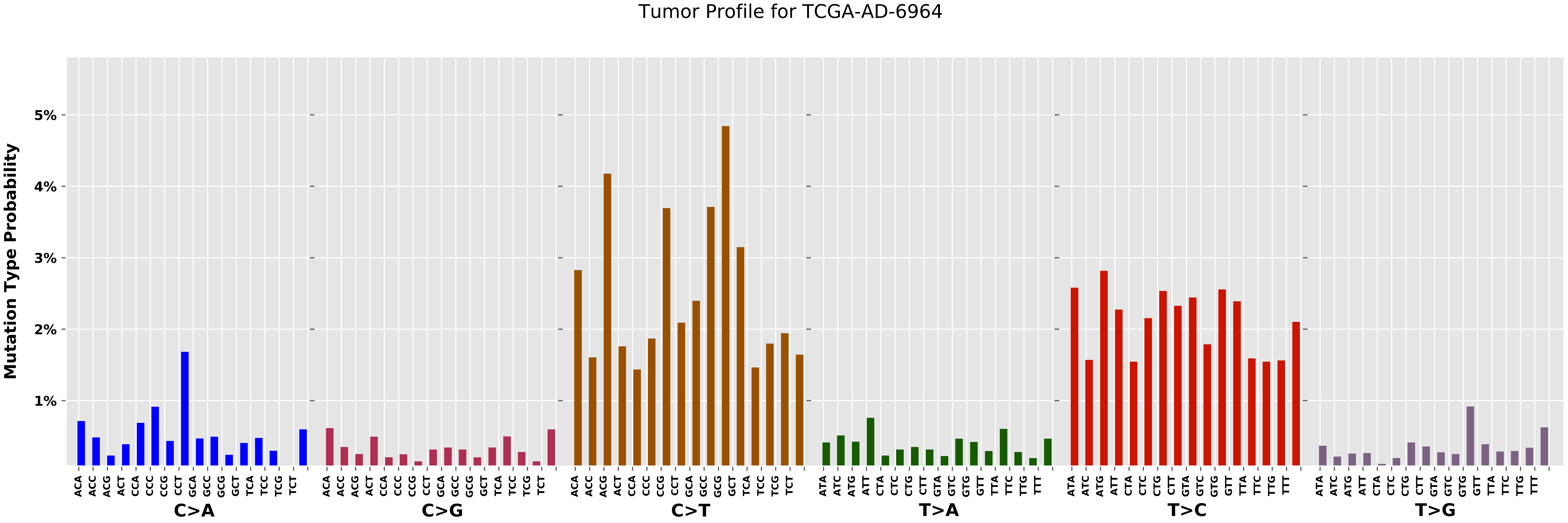
Tumor Profile for TCGA-AD-6964
Mutation Type Probability
5%
4%
3%
2%
1%
ACA ACC ACG ACT CCA CCC CCG CCT GCA GCC GCG GCT TCA TCC TCG TCT
C>A
C>G
C>T
ATA ATC ATG ATT CTA CTC CTG CTT GTA GTC GTG GTT TTA TTC TTG TTT
T>A
T>C
T>G

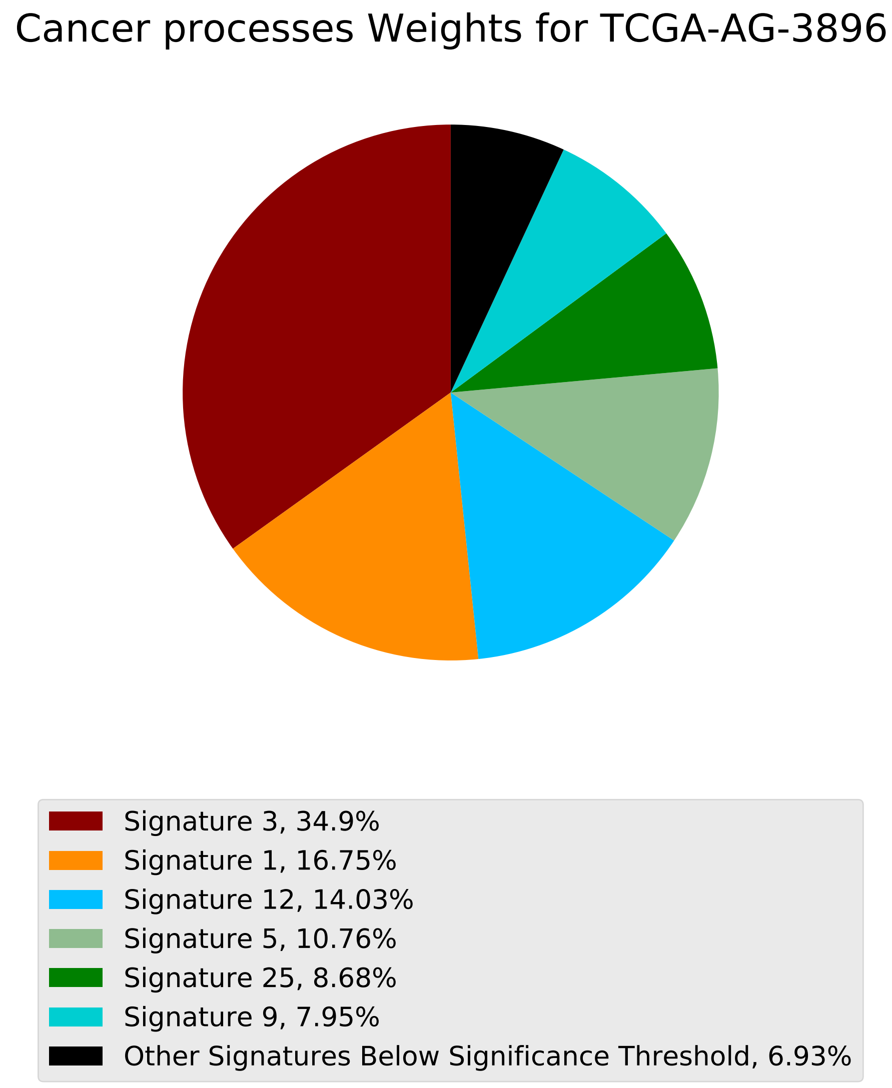
Cancer processes Weights for TCGA-AG-3896
Signature 3, 34.9%
Signature 1, 16.75%
Signature 12, 14.03%
Signature 5, 10.76%
Signature 25, 8.68%
Signature 9, 7.95%
Other Signatures Below Significance Threshold, 6.93%

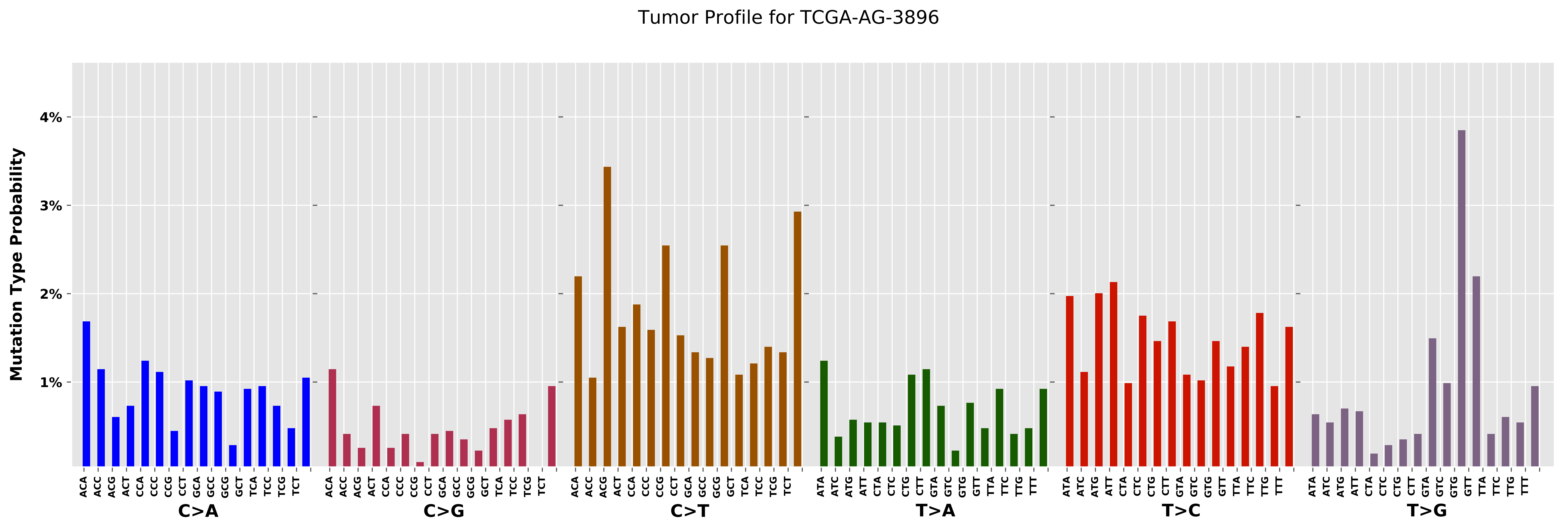
Tumor Profile for TCGA-AG-3896
Mutation Type Probability
4%
3%
2%
1%
C>A
C>G
C>T
T>A
T>C
T>G

Cancer processes Weights for TCGA-E2-A15K

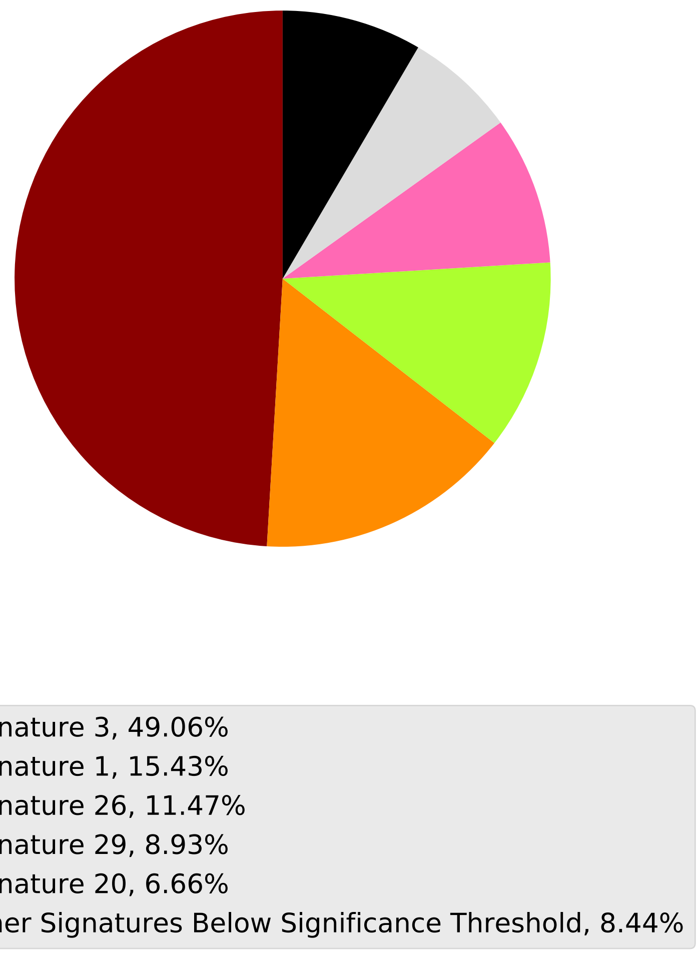

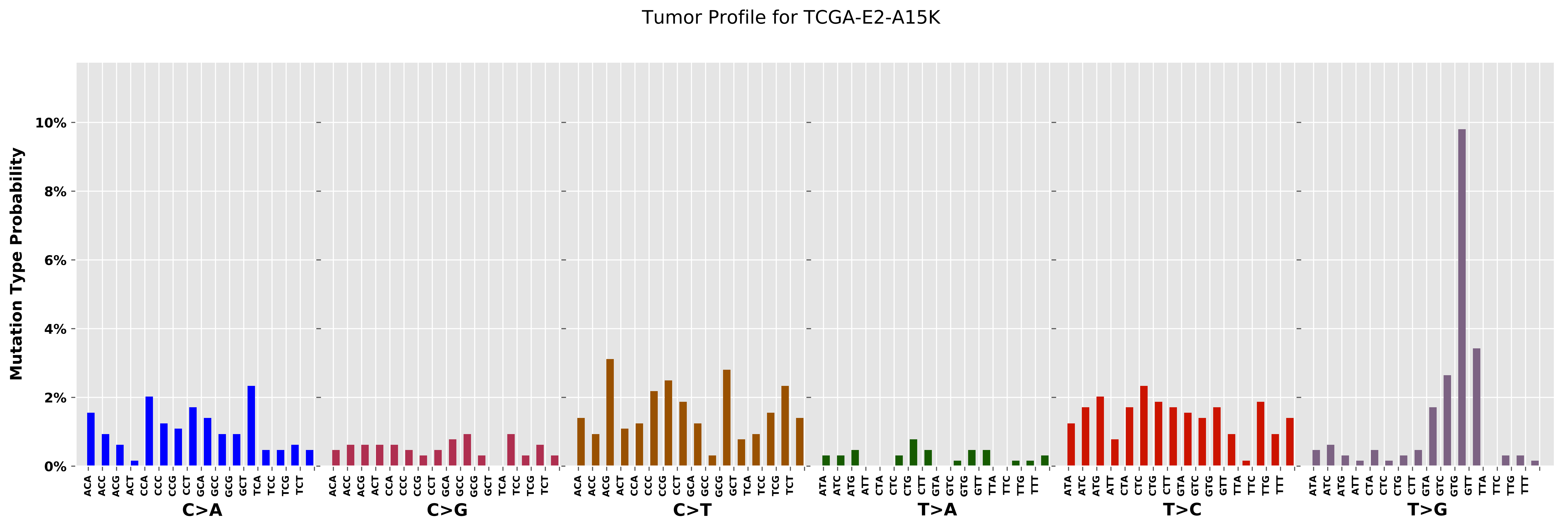

Tumor Profile for TCGA-E2-A15K
Mutation Type Probability
0%
2%
4%
6%
8%
10%
C>A
C>G
C>T
T>A
T>C
T>G

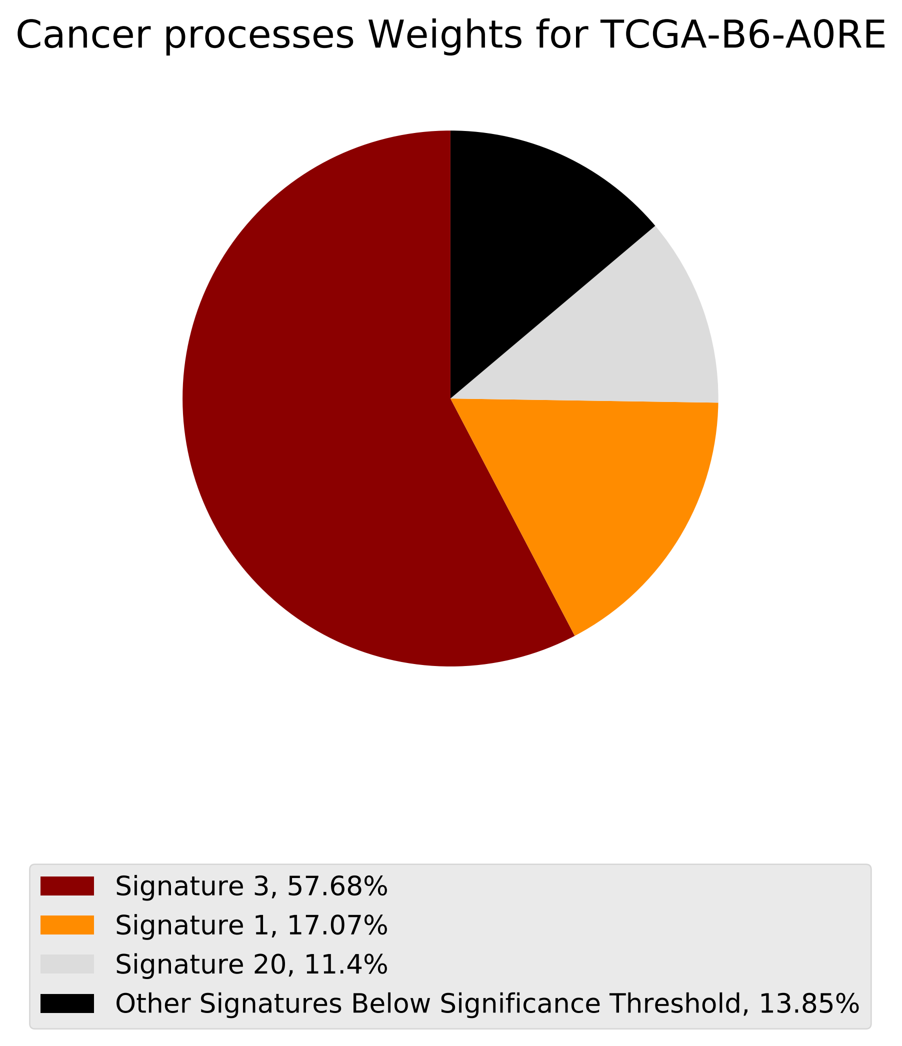
Cancer processes Weights for TCGA-B6-A0RE
Signature 3, 57.68%
Signature 1, 17.07%
Signature 20, 11.4%
Other Signatures Below Significance Threshold, 13.85%

Tumor Profile for TCGA-B6-A0RE

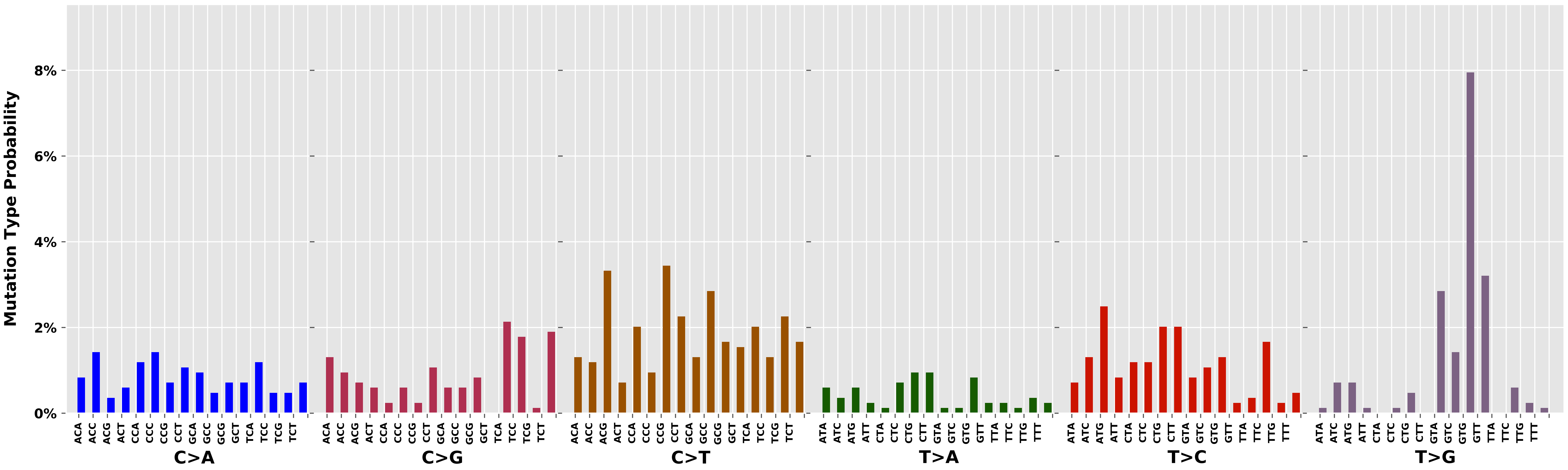

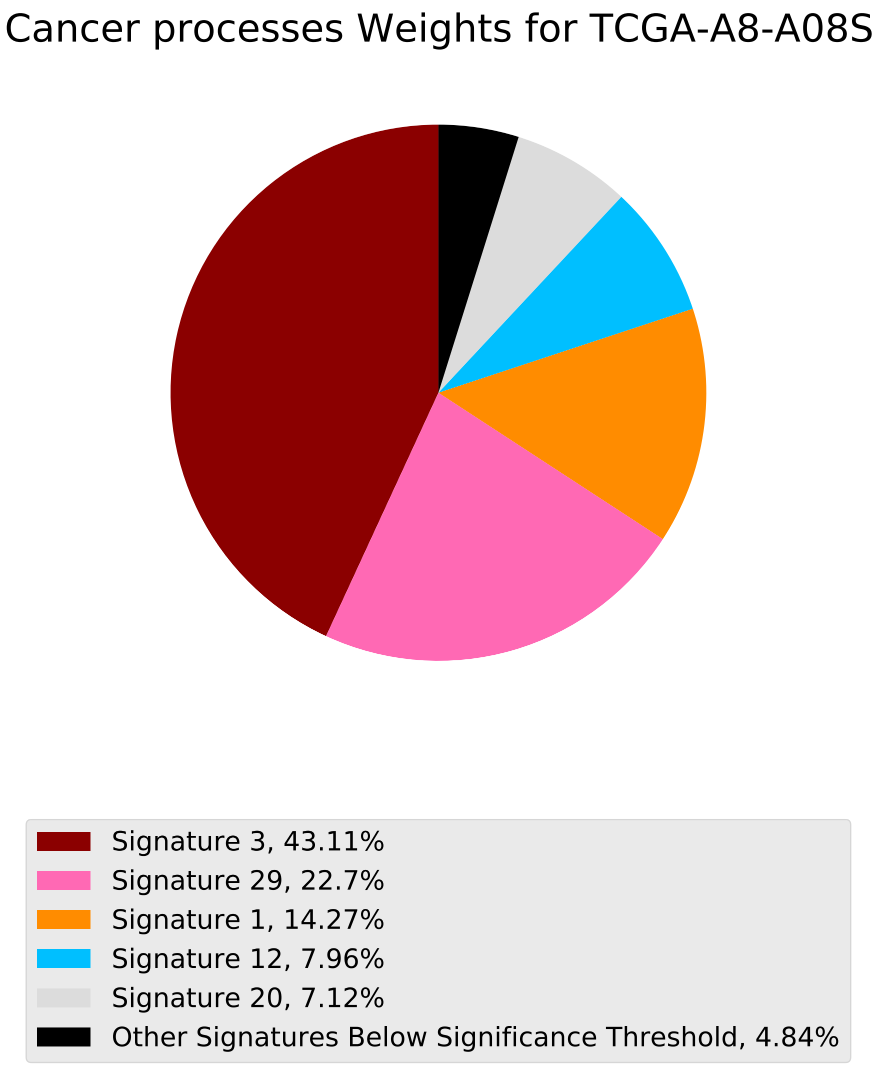
Cancer processes Weights for TCGA-A8-A08S
Signature 3, 43.11%
Signature 29, 22.7%
Signature 1, 14.27%
Signature 12, 7.96%
Signature 20, 7.12%
Other Signatures Below Significance Threshold, 4.84%

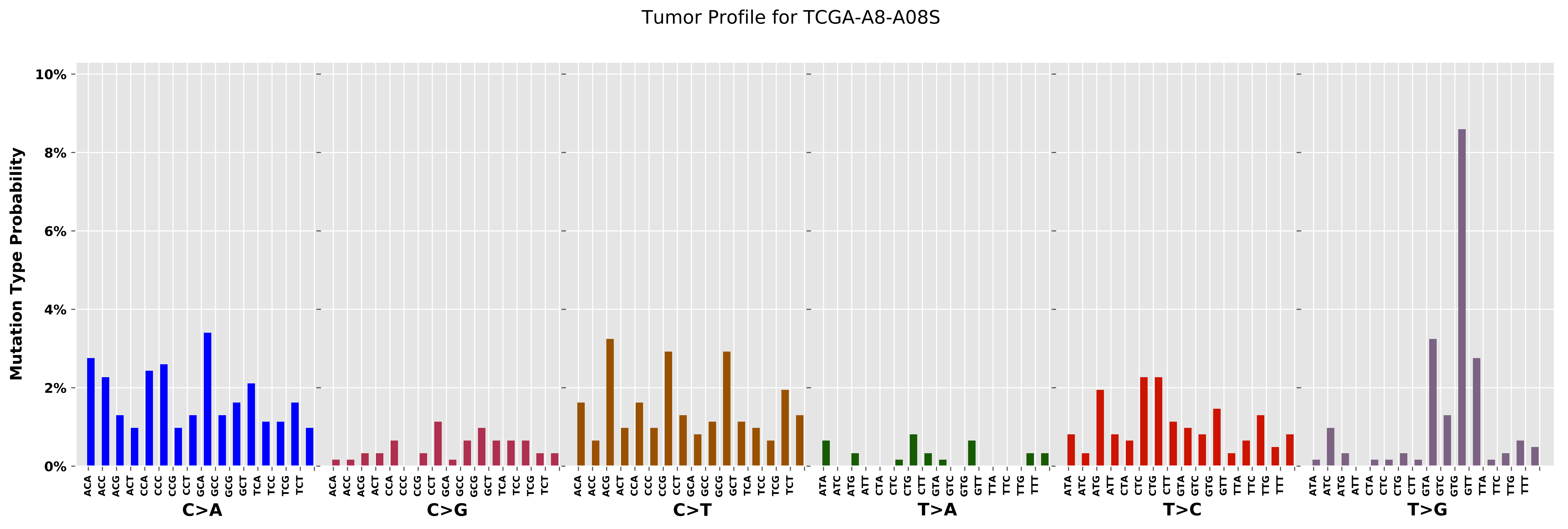
Tumor Profile for TCGA-A8-A08S
Mutation Type Probability
10%
8%
6%
4%
2%
0%
C>A
C>G
C>T
T>A
T>C
T>G
ACA ACC ACG ACT CCA CCC CCG CCT GCA GCC GCG GCT TCA TCC TCG TCT
ATA ATC ATG ATT CTA CTC CTG CTT GTA GTC GTG GTT TTA TTC TTG TTT

Cancer processes Weights for TCGA-AA-A03F

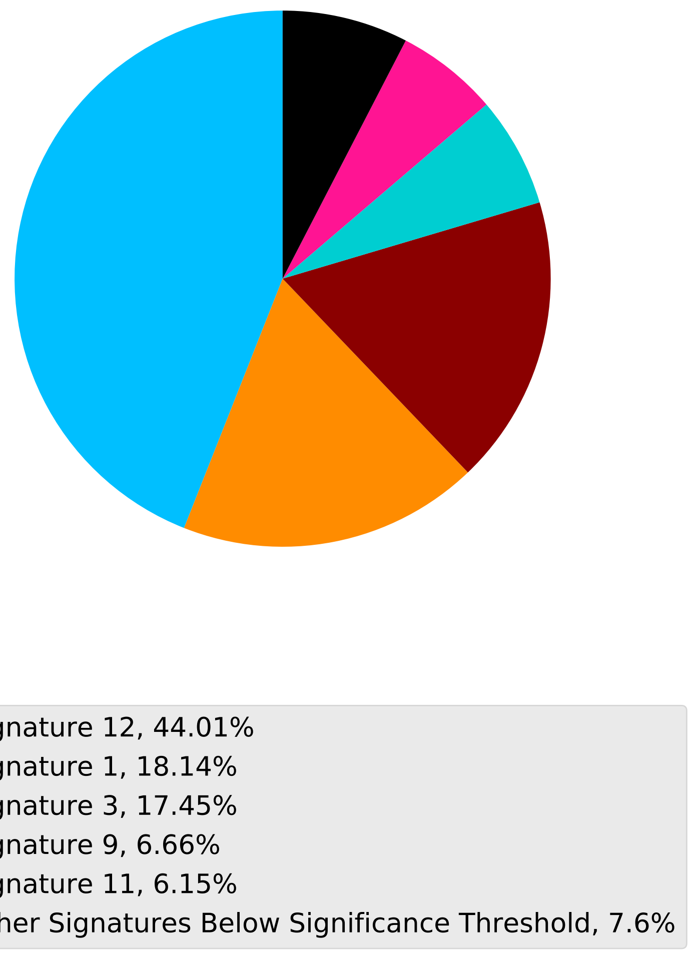

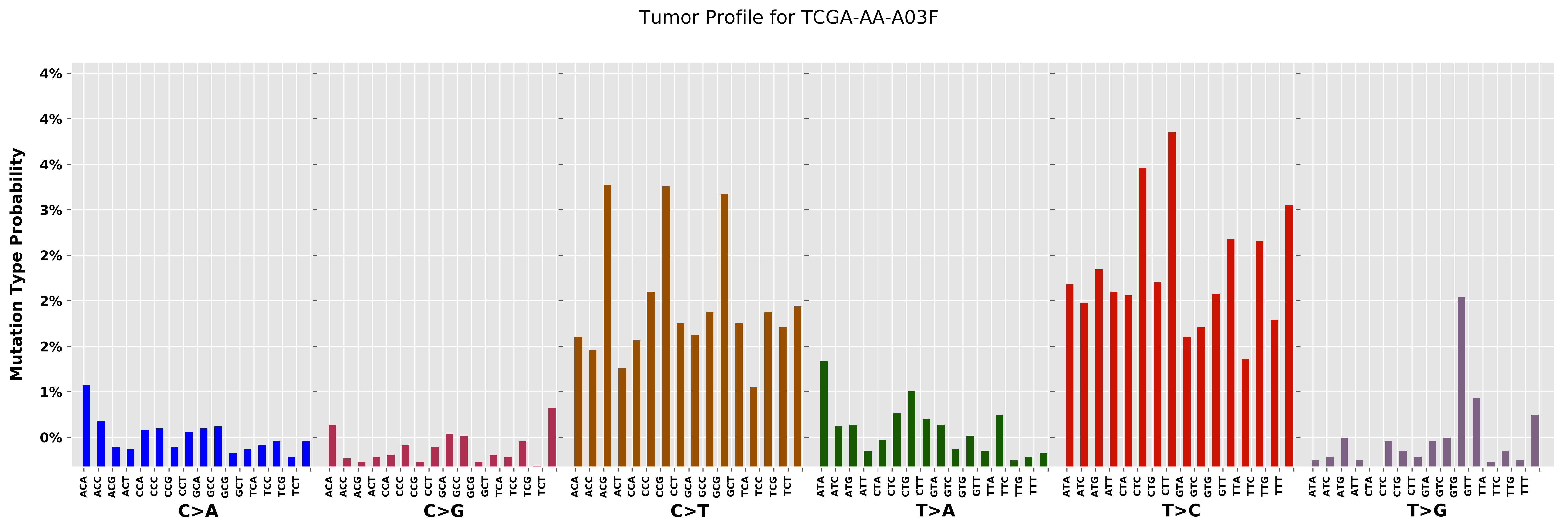

Tumor Profile for TCGA-AA-A03F
Mutation Type Probability
4%
4%
4%
3%
2%
2%
2%
1%
0%
C>A
C>G
C>T
T>A
T>C
T>G
ACA ACC ACG ACT CCA CCC CCG CCT GCA GCC GCG GCT TCA TCC TCG TCT
ATA ATC ATG ATT CTA CTC CTG CTT GTA GTC GTG GTT TTA TTC TTG TTT

Cancer processes Weights for TCGA-EW-A1PH

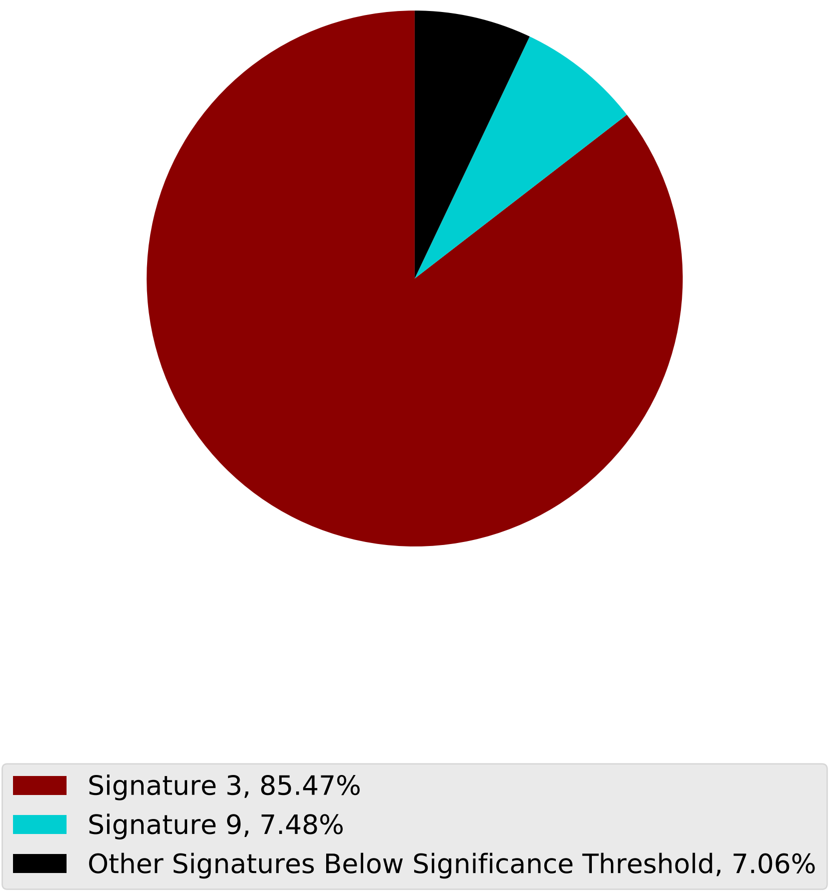

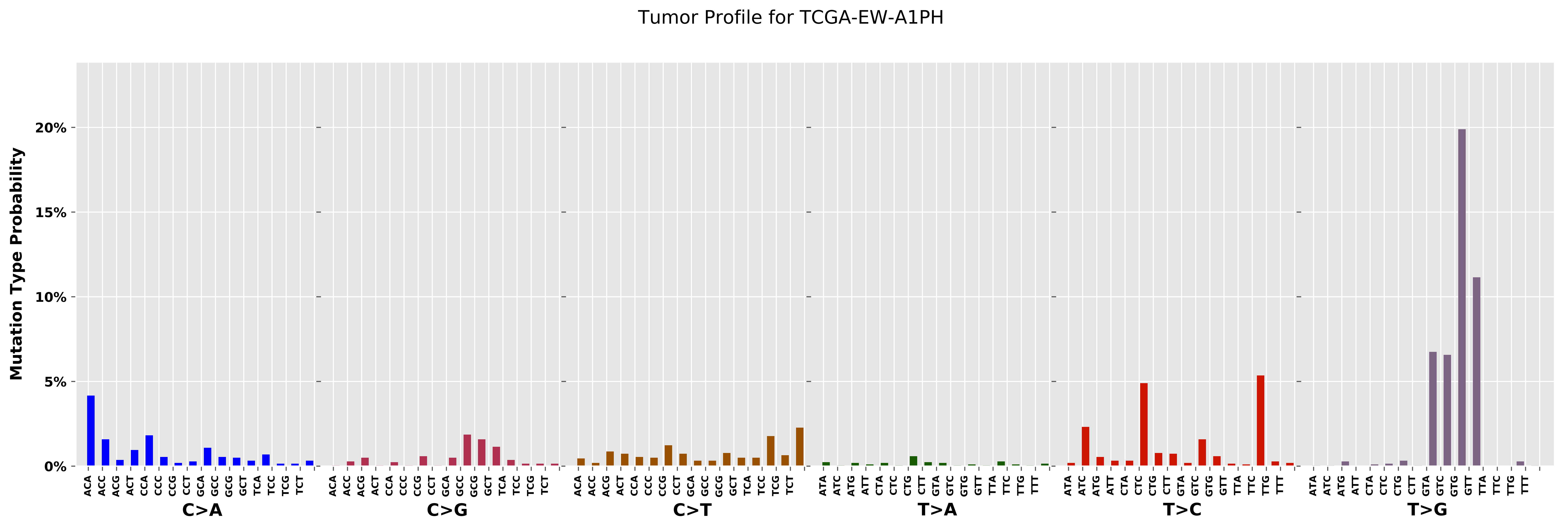

Tumor Profile for TCGA-EW-A1PH
Mutation Type Probability
0%
5%
10%
15%
20%
C>A
C>G
C>T
T>A
T>C
T>G

Cancer processes Weights for TCGA-E2-A1LG

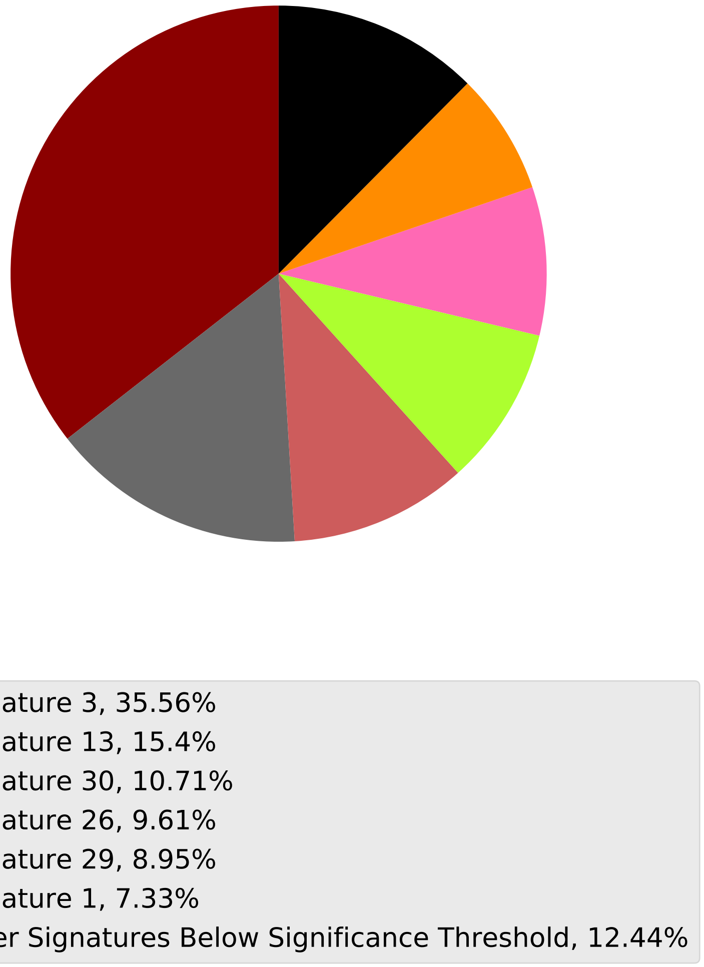

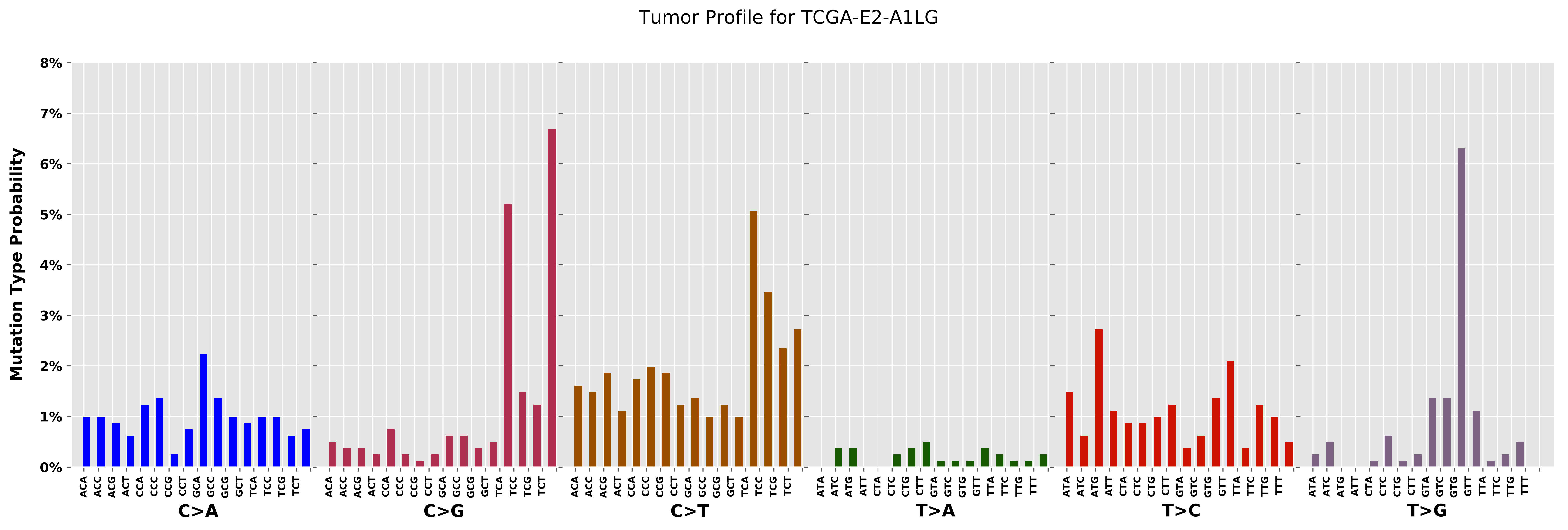

Tumor Profile for TCGA-E2-A1LG
Mutation Type Probability
8%
7%
6%
5%
4%
3%
2%
1%
0%
C>A
C>G
C>T
T>A
T>C
T>G
ACA ACC ACG ACT CCA CCC CCG CCT GCA GCC GCG GCT TCA TCC TCG TCT
ATA ATC ATG ATT CTA CTC CTG CTT GTA GTC GTG GTT TTA TTC TTG TTT

Cancer processes Weights for TCGA-A8-A08B

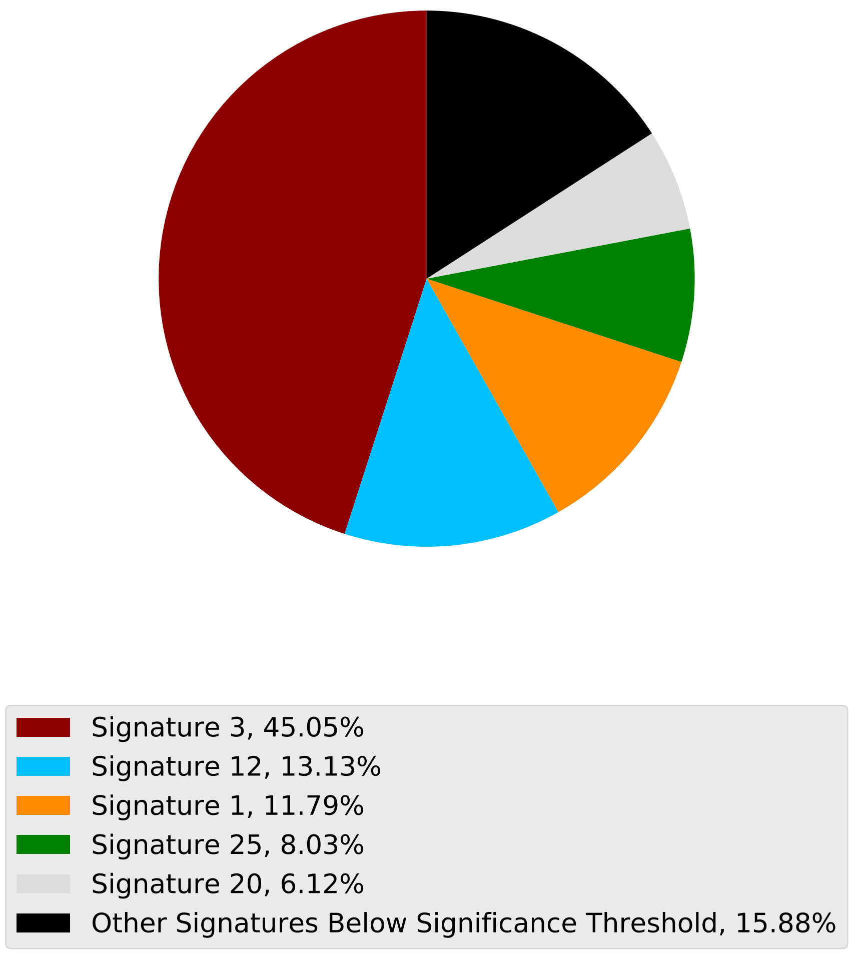

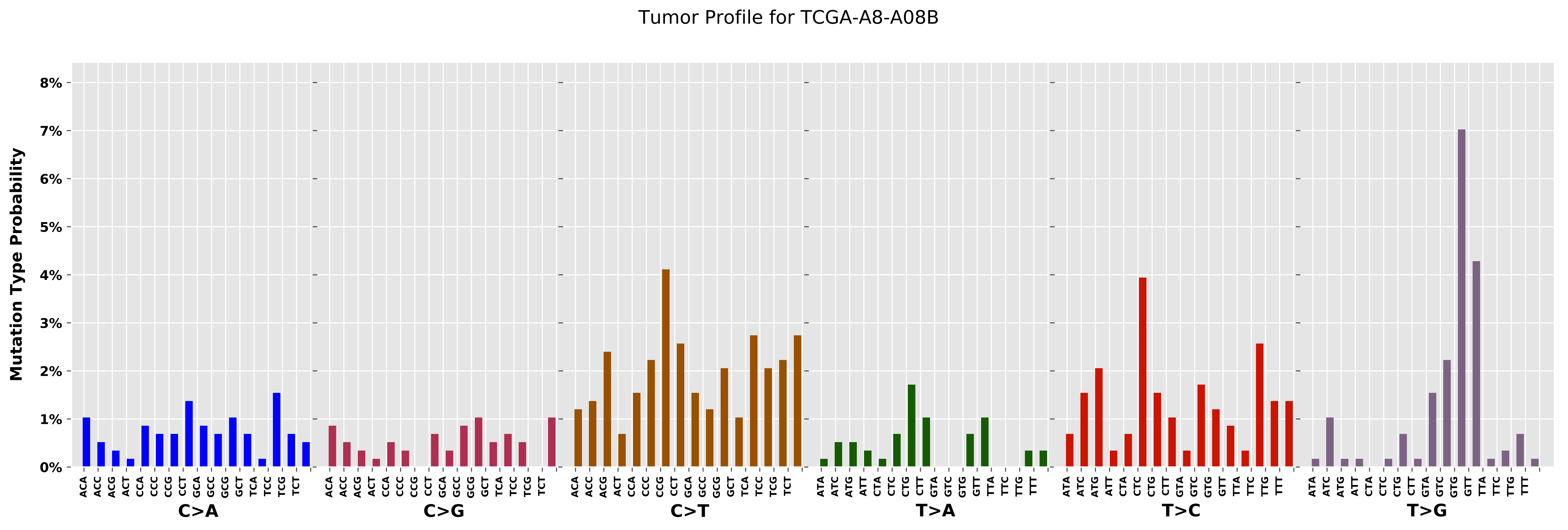
Tumor Profile for TCGA-A8-A08B
Mutation Type Probability
0%
1%
2%
3%
4%
5%
6%
7%
8%
C>A
C>G
C>T
T>A
T>C
T>G
ACA ACC ACG ACT CCA CCC CCG CCT GCA GCC GCG GCT TCA TCC TCG TCT
ATA ATC ATG ATT CTA CTC CTG CTT GTA GTC GTG GTT TTA TTC TTG TTT

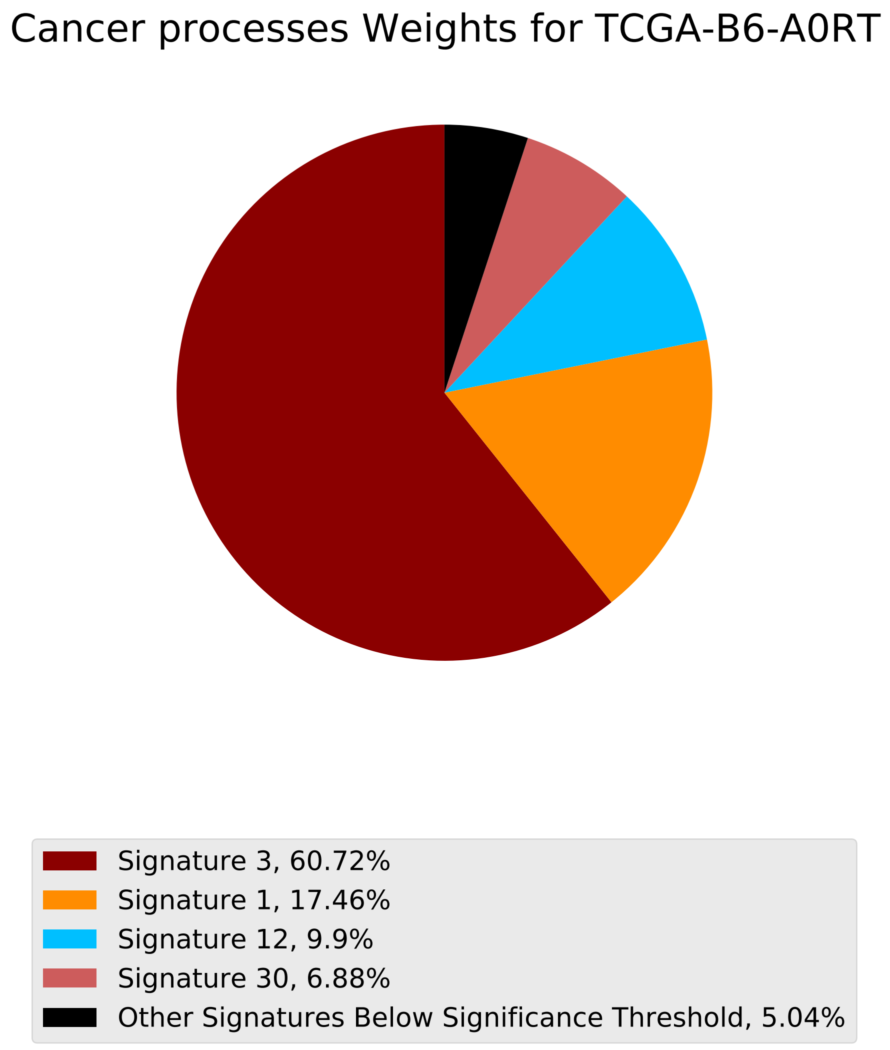
Cancer processes Weights for TCGA-B6-A0RT
Signature 3, 60.72%
Signature 1, 17.46%
Signature 12, 9.9%
Signature 30, 6.88%
Other Signatures Below Significance Threshold, 5.04%

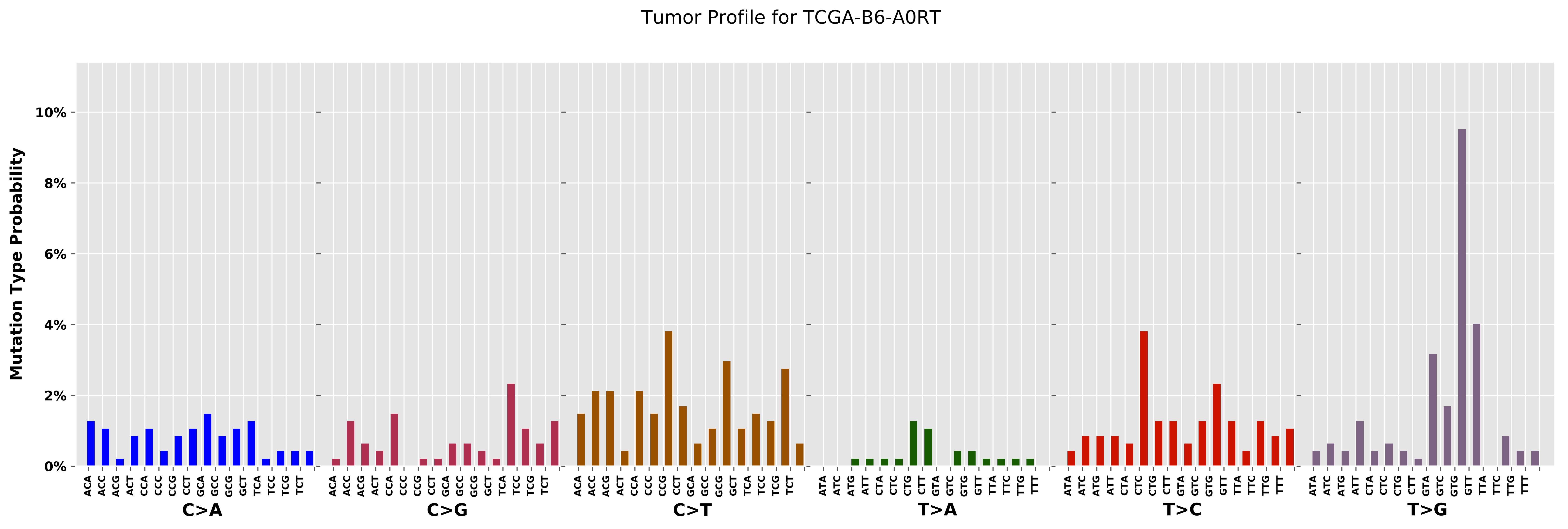

Tumor Profile for TCGA-B6-A0RT
Mutation Type Probability
10%
8%
6%
4%
2%
0%
C>A
C>G
C>T
T>A
T>C
T>G

Cancer processes Weights for TCGA-GM-A3XL

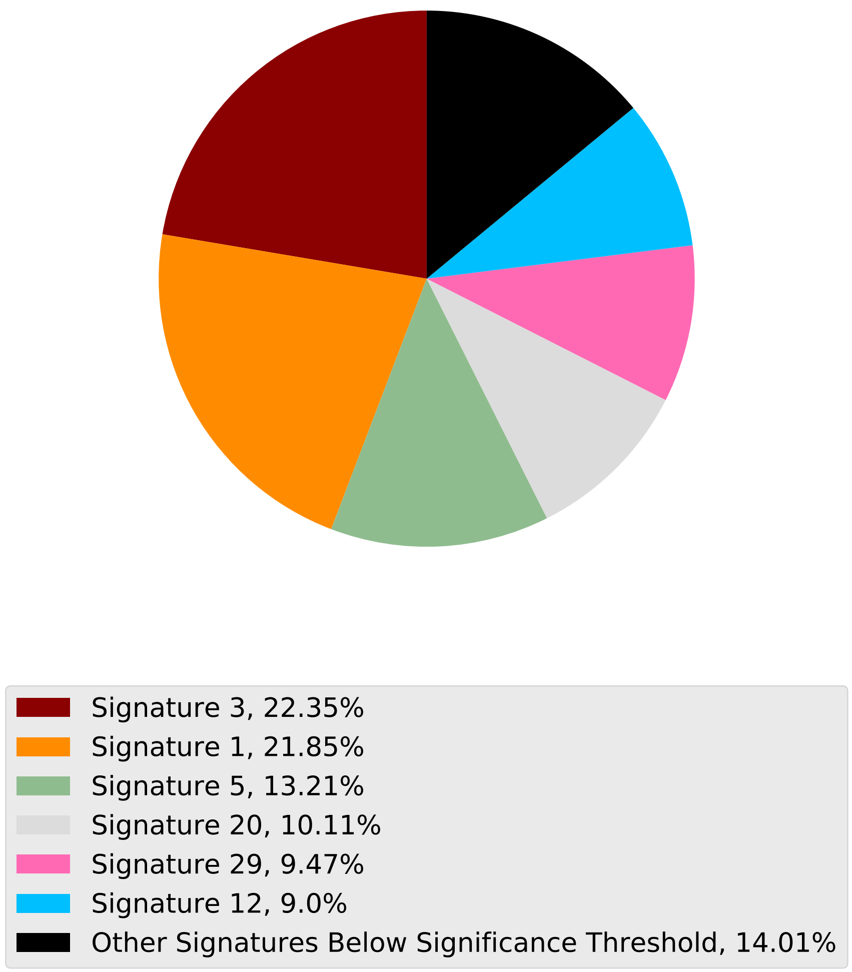

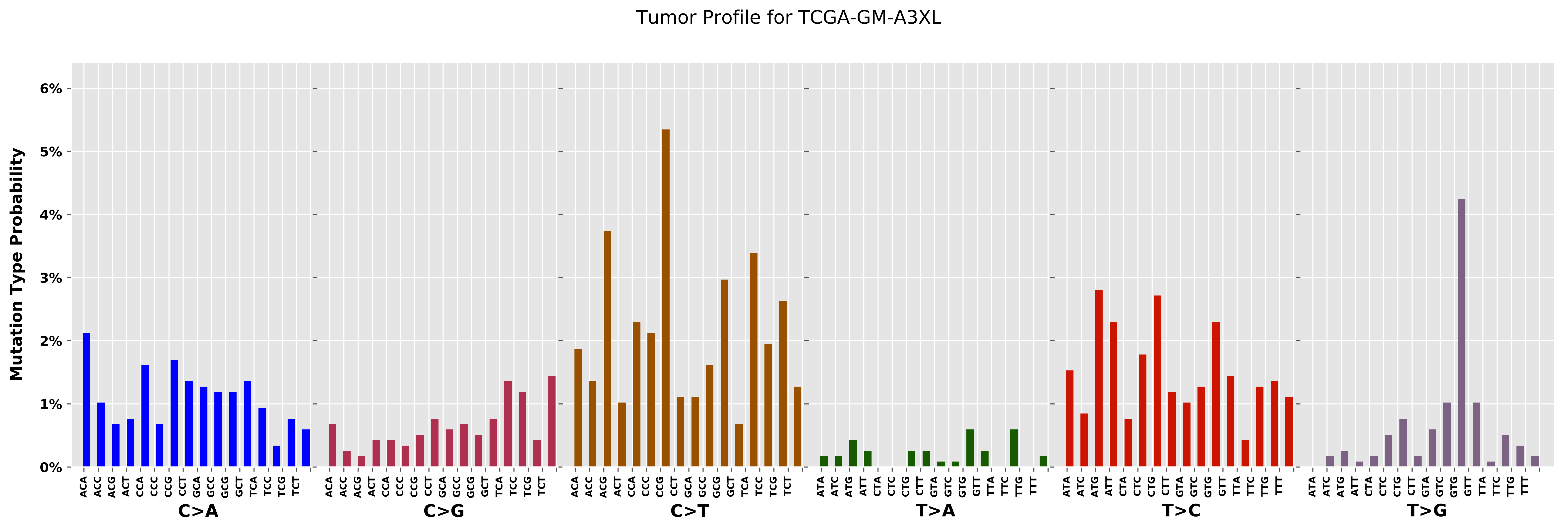

Tumor Profile for TCGA-GM-A3XL
Mutation Type Probability
0%
1%
2%
3%
4%
5%
6%
C>A
C>G
C>T
T>A
T>C
T>G
ACA
ACC
ACG
ACT
CCA
CCC
CCG
CCT
GCA
GCC
GCG
GCT
TCA
TCC
TCG
TCT
ATA
ATC
ATG
ATT
CTA
CTC
CTG
CTT
GTA
GTC
GTG
GTT
TTA
TTC
TTG
TTT

Cancer processes Weights for TCGA-AO-A0J2

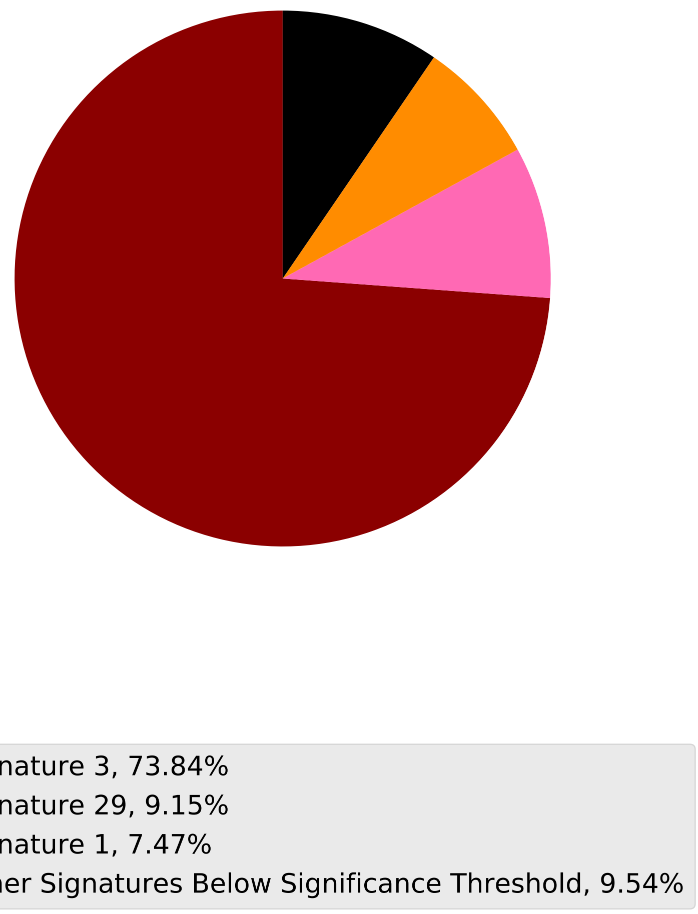

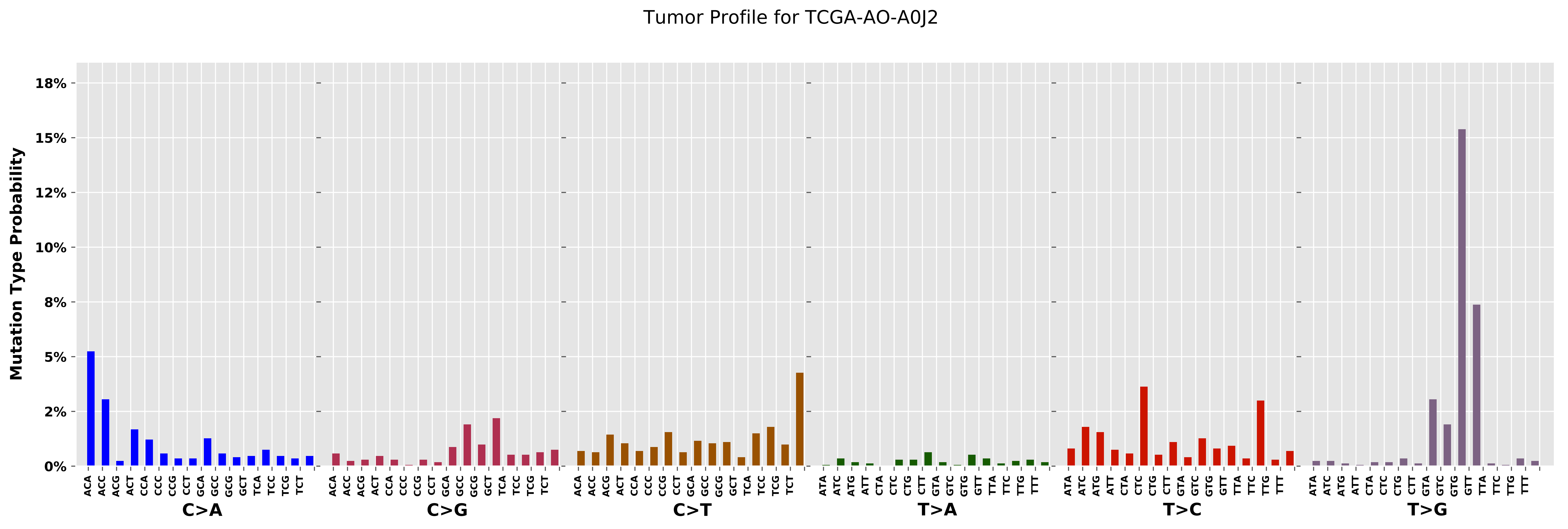

Tumor Profile for TCGA-AO-A0J2
Mutation Type Probability
18%
15%
12%
10%
8%
5%
2%
0%
C>A
C>G
C>T
T>A
T>C
T>G
ACA ACC ACG ACT CCA CCC CCG CCT GCA GCC GCG GCT TCA TCC TCG TCT
ATA ATC ATG ATT CTA CTC CTG CTT GTA GTC GTG GTT TTA TTC TTG TTT

Cancer processes Weights for TCGA-E2-A156

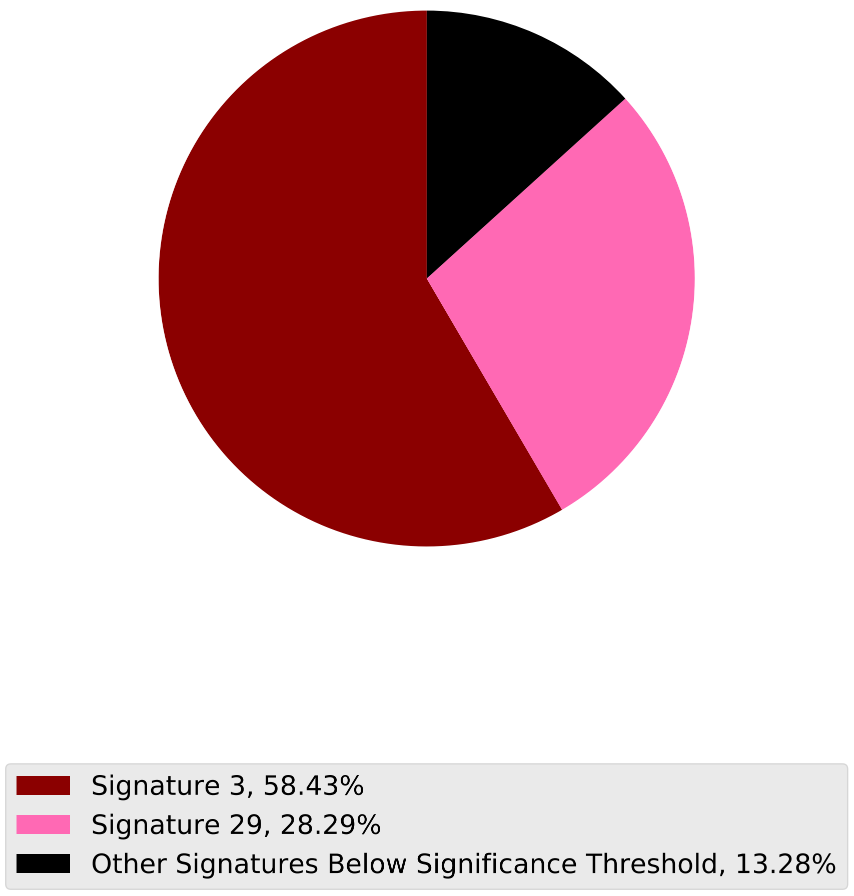

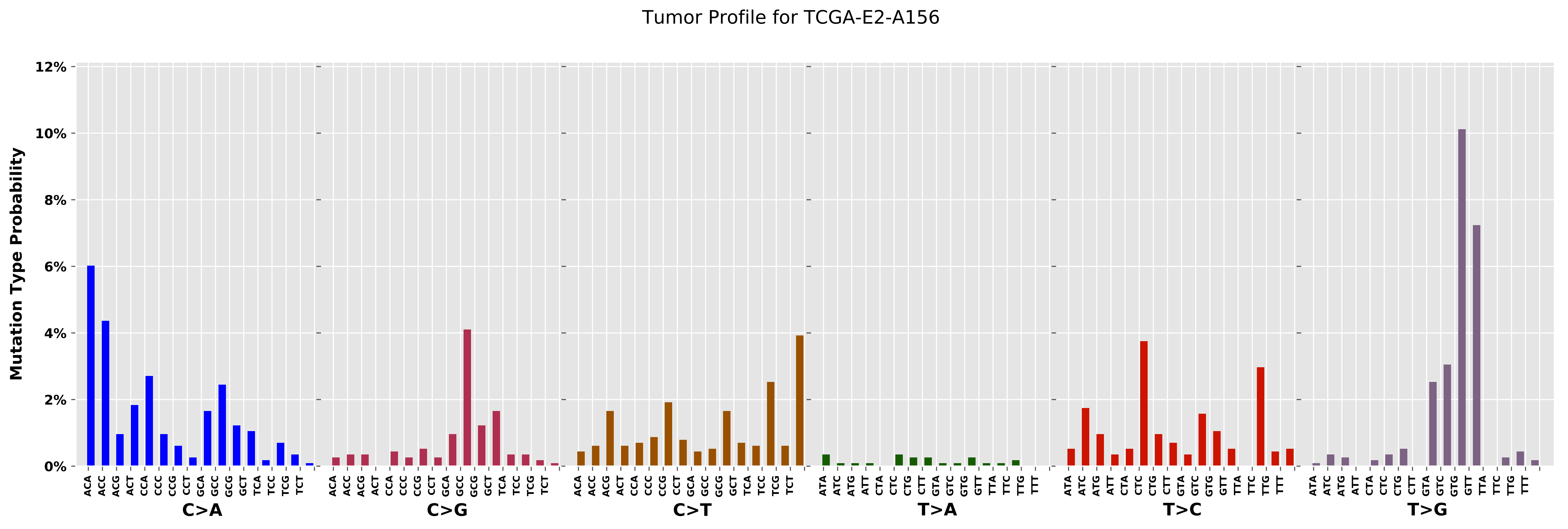

Tumor Profile for TCGA-E2-A156
Mutation Type Probability
12%
10%
8%
6%
4%
2%
0%
ACA ACC ACG ACT CCA CCC CCG CCT GCA GCC GCG GCT TCA TCC TCG TCT
C>A
ACA ACC ACG ACT CCA CCC CCG CCT GCA GCC GCG GCT TCA TCC TCG TCT
C>G
ACA ACC ACG ACT CCA CCC CCG CCT GCA GCC GCG GCT TCA TCC TCG TCT
C>T
ATA ATC ATG ATT CTA CTC CTG CTT GTA GTC GTG GTT TTA TTC TTG TTT
T>A
ATA ATC ATG ATT CTA CTC CTG CTT GTA GTC GTG GTT TTA TTC TTG TTT
T>C
ATA ATC ATG ATT CTA CTC CTG CTT GTA GTC GTG GTT TTA TTC TTG TTT
T>G

Cancer processes Weights for TCGA-BH-A18R

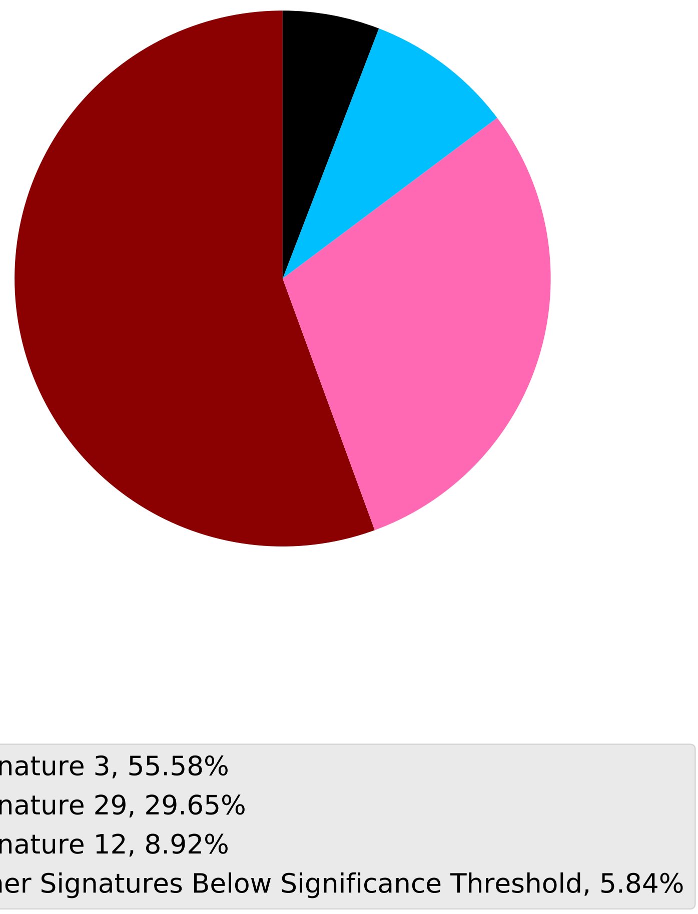

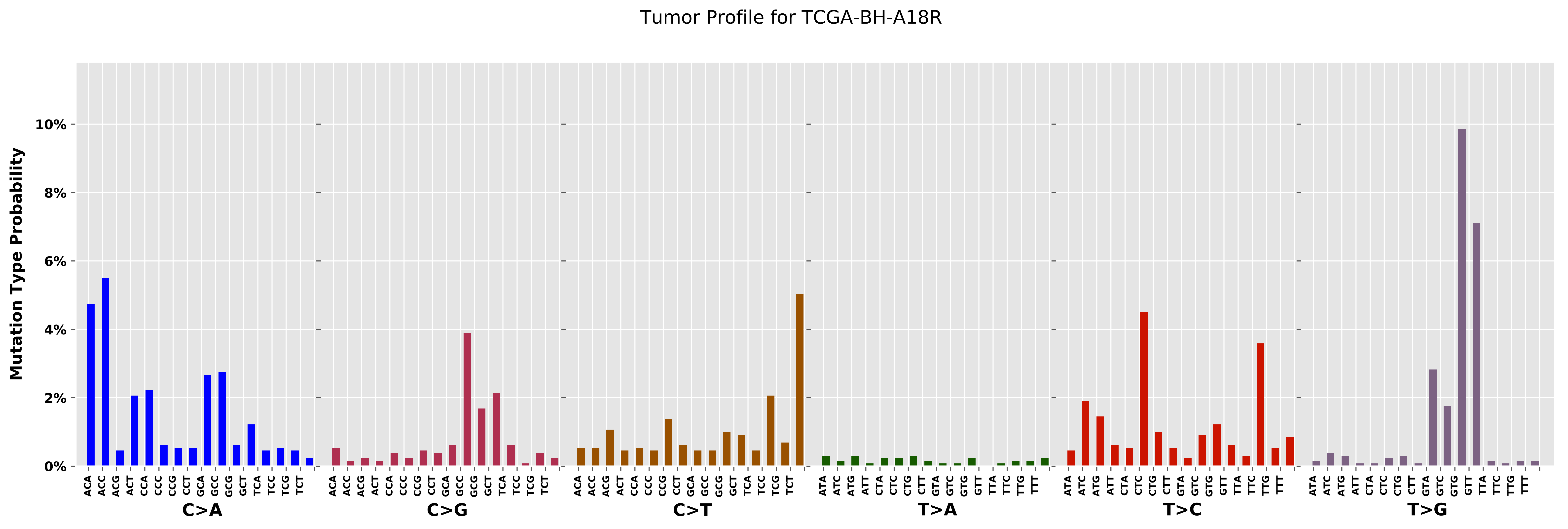

Tumor Profile for TCGA-BH-A18R
Mutation Type Probability
0%
2%
4%
6%
8%
10%
C>A
C>G
C>T
T>A
T>C
T>G
ACA
ACC
ACG
ACT
CCA
CCC
CCG
CCT
GCA
GCC
GCG
GCT
TCA
TCC
TCG
TCT
ATA
ATC
ATG
ATT
CTA
CTC
CTG
CTT
GTA
GTC
GTG
GTT
TTA
TTC
TTG
TTT

Cancer processes Weights for TCGA-AO-A0JM

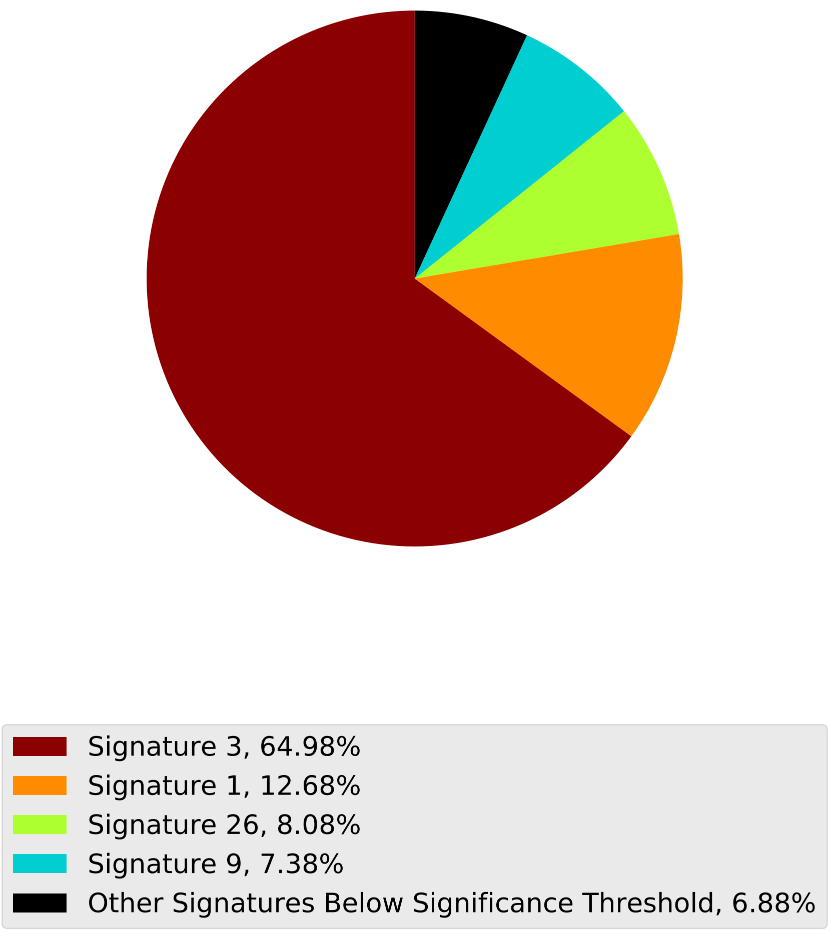

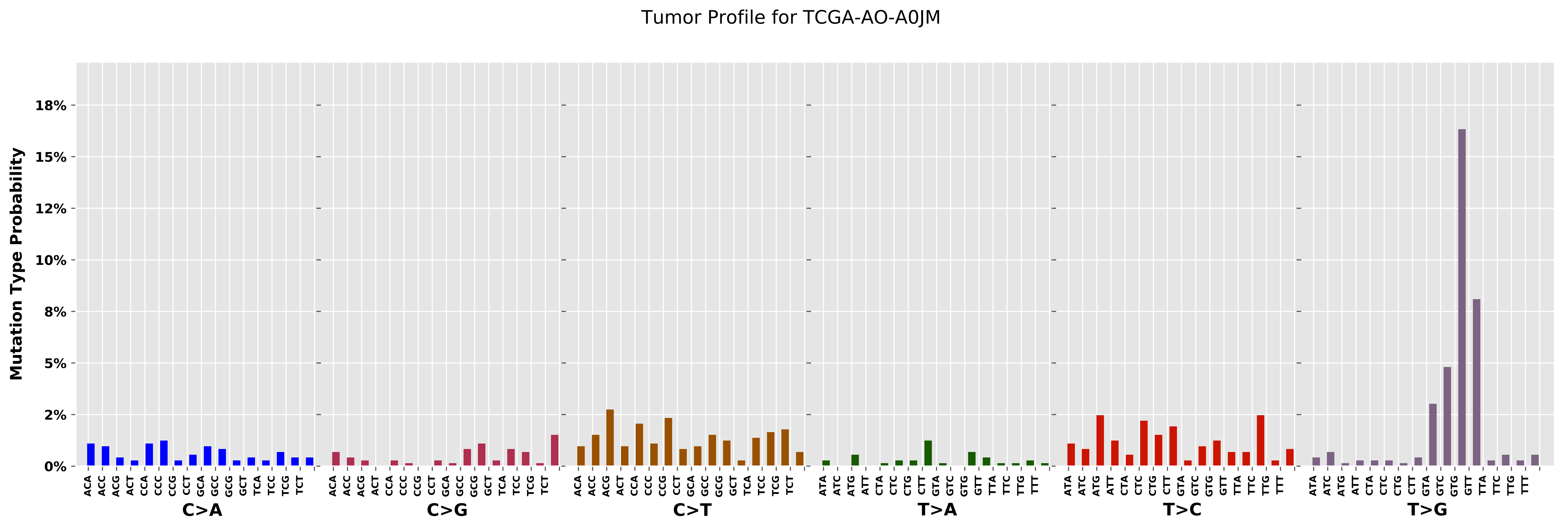

Tumor Profile for TCGA-AO-A0JM
Mutation Type Probability
18%
15%
12%
10%
8%
5%
2%
0%
C>A
C>G
C>T
T>A
T>C
T>G
ACA ACC ACG ACT CCA CCC CCG CCT GCA GCC GCG GCT TCA TCC TCG TCT
ATA ATC ATG ATT CTA CTC CTG CTT GTA GTC GTG GTT TTA TTC TTG TTT

Cancer processes Weights for TCGA-AA-A01S

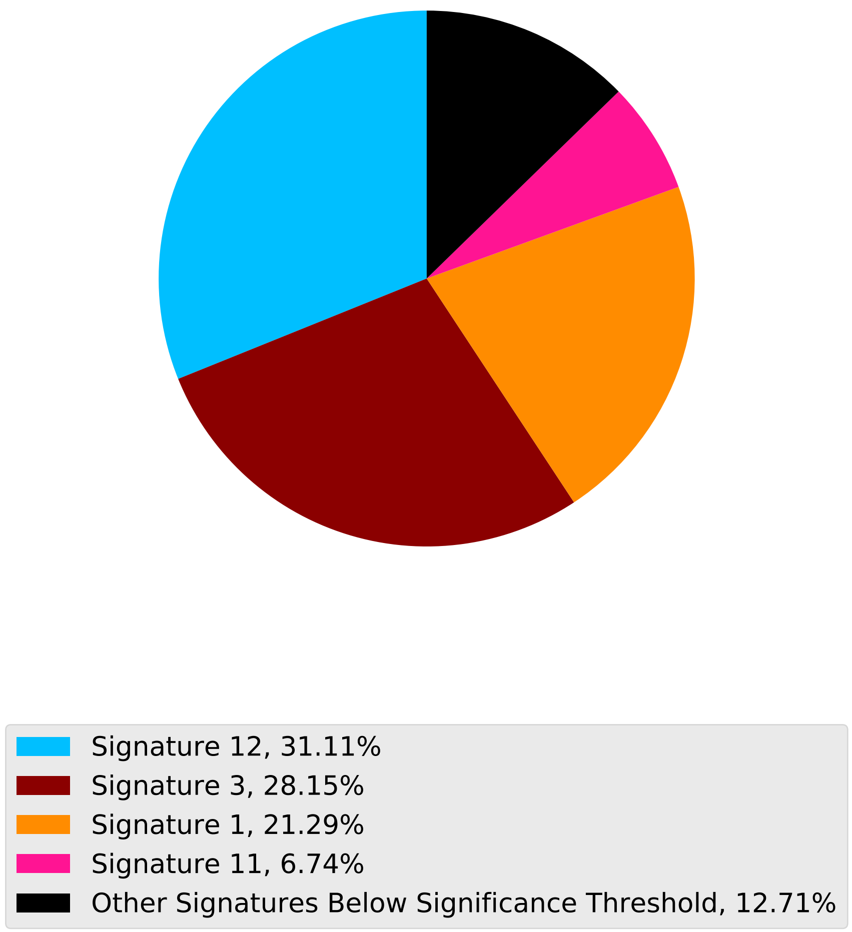

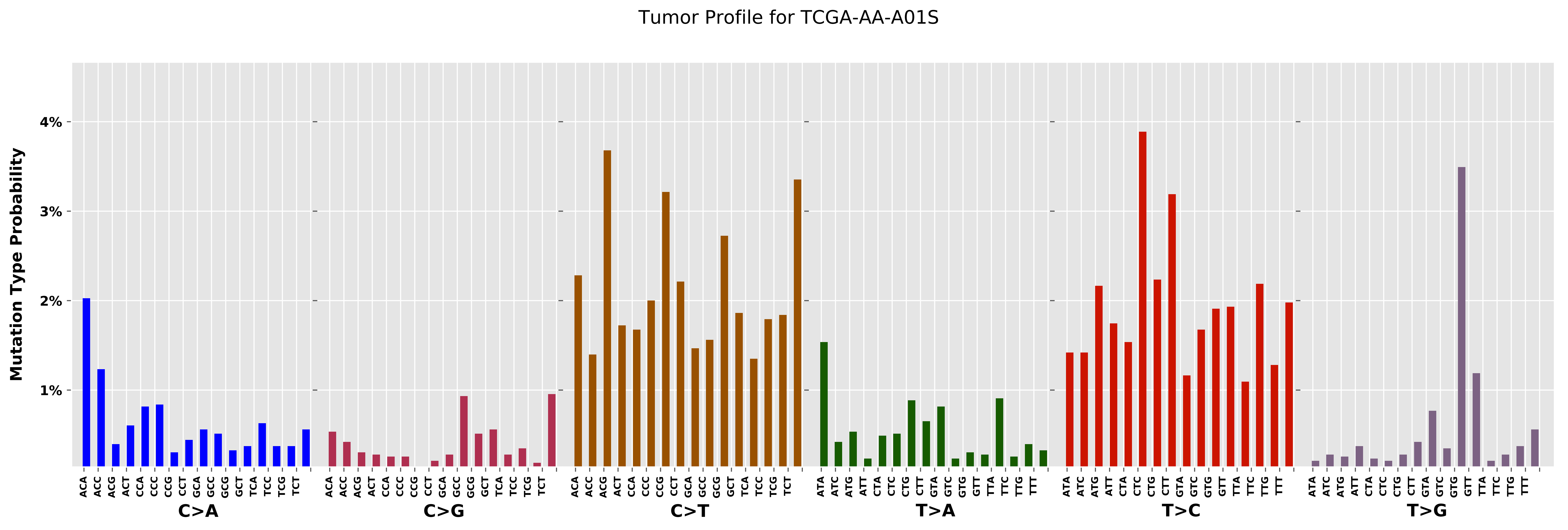

Tumor Profile for TCGA-AA-A01S
Mutation Type Probability
4%
3%
2%
1%
C>A
C>G
C>T
T>A
T>C
T>G
ACA ACC ACG ACT CCA CCC CCG CCT GCA GCC GCG GCT TCA TCC TCG TCT
ATA ATC ATG ATT CTA CTC CTG CTT GTA GTC GTG GTT TTA TTC TTG TTT

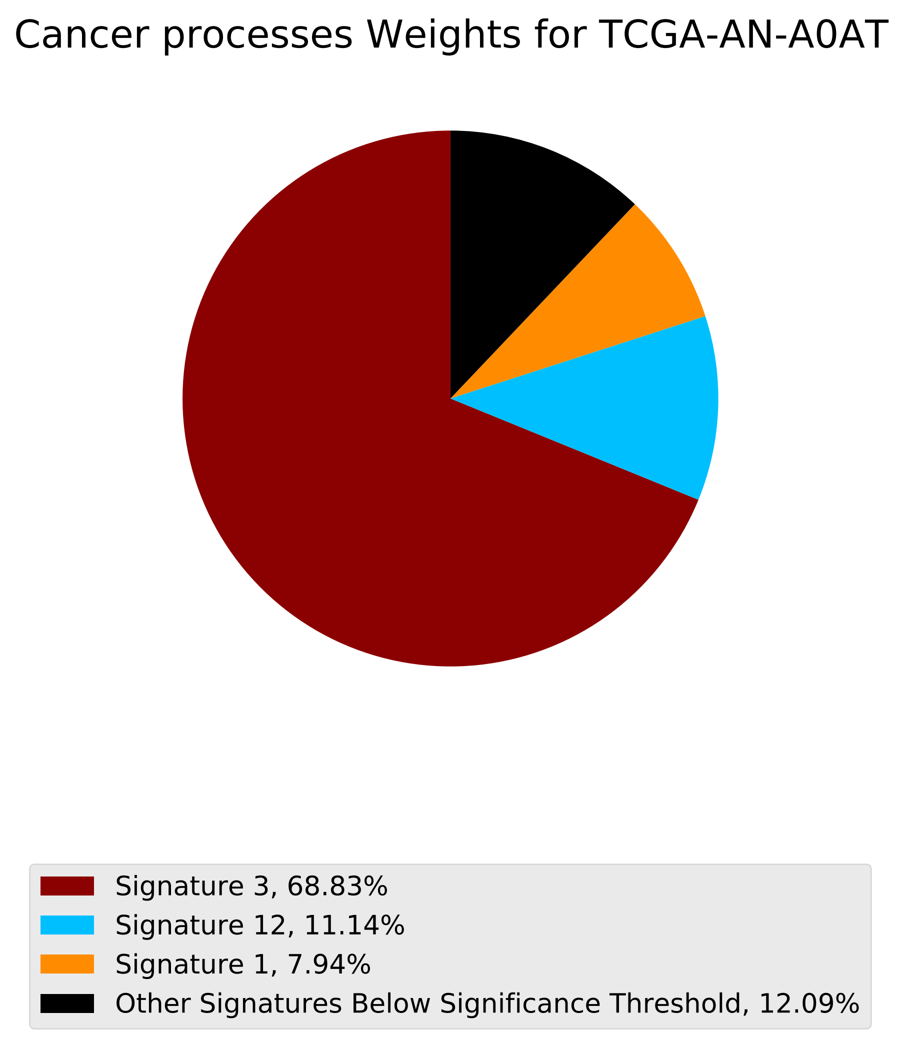
Cancer processes Weights for TCGA-AN-A0AT
Signature 3, 68.83%
Signature 12, 11.14%
Signature 1, 7.94%
Other Signatures Below Significance Threshold, 12.09%

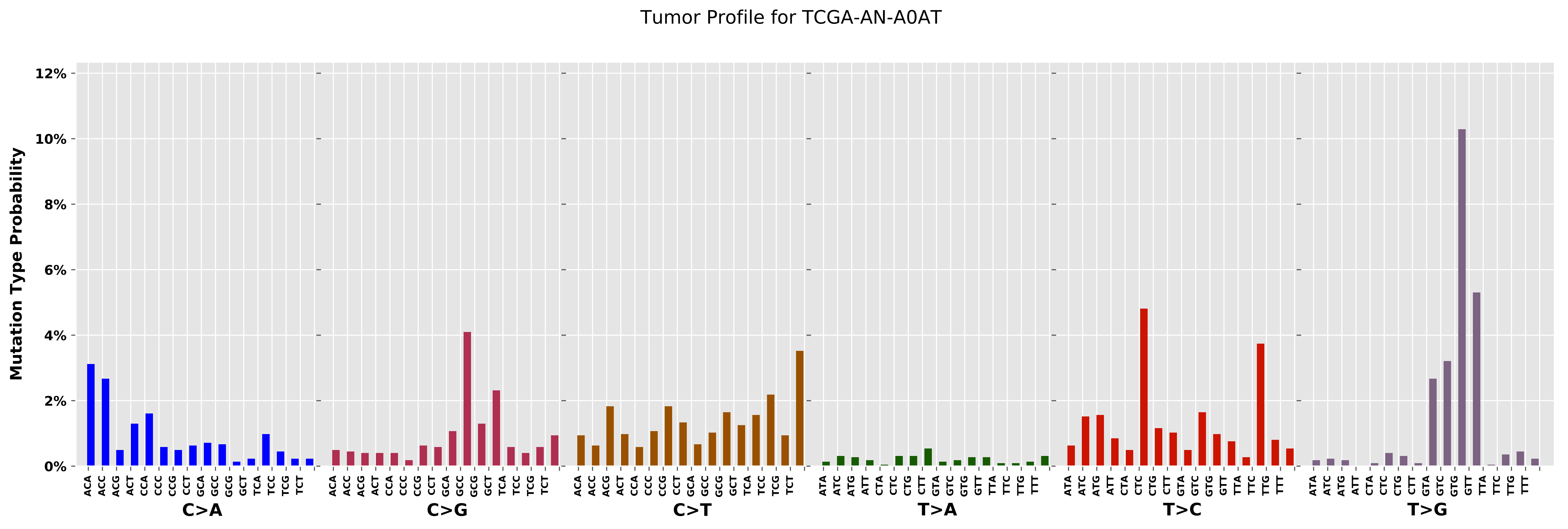
Tumor Profile for TCGA-AN-A0AT
Mutation Type Probability
12%
10%
8%
6%
4%
2%
0%
ACA ACC ACG ACT CCA CCC CCG CCT GCA GCC GCG GCT TCA TCC TCG TCT
C>A
ACA ACC ACG ACT CCA CCC CCG CCT GCA GCC GCG GCT TCA TCC TCG TCT
C>G
ACA ACC ACG ACT CCA CCC CCG CCT GCA GCC GCG GCT TCA TCC TCG TCT
C>T
ATA ATC ATG ATT CTA CTC CTG CTT GTA GTC GTG GTT TTA TTC TTG TTT
T>A
ATA ATC ATG ATT CTA CTC CTG CTT GTA GTC GTG GTT TTA TTC TTG TTT
T>C
ATA ATC ATG ATT CTA CTC CTG CTT GTA GTC GTG GTT TTA TTC TTG TTT
T>G

Cancer processes Weights for TCGA-B6-A0WX

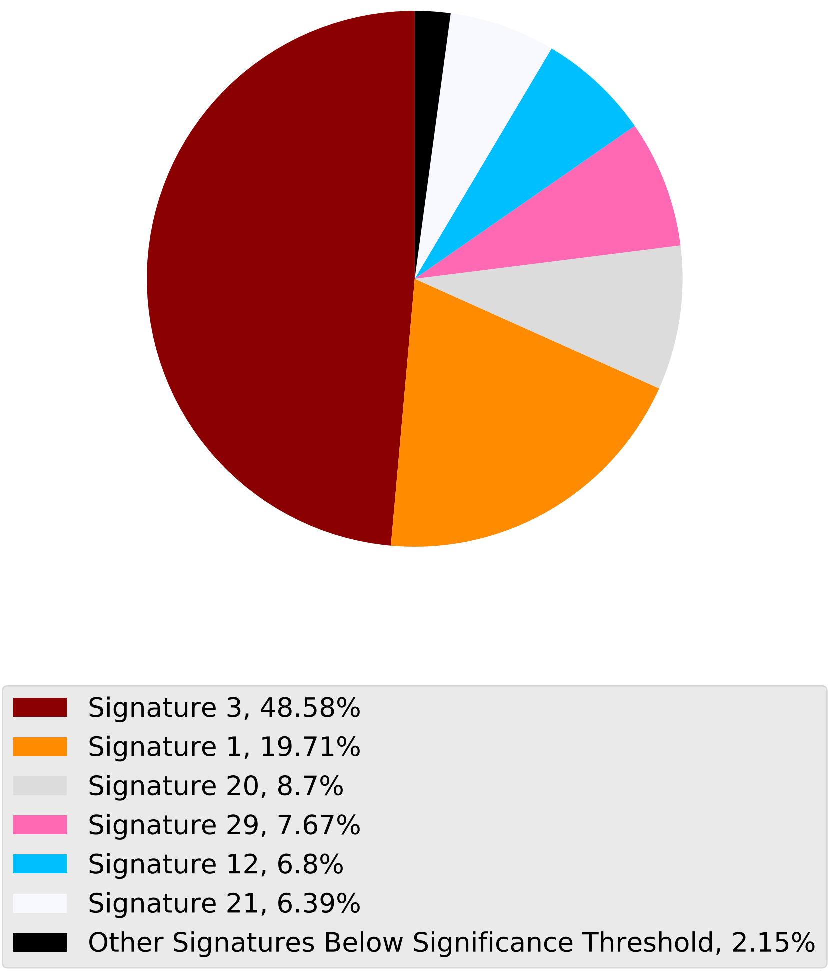

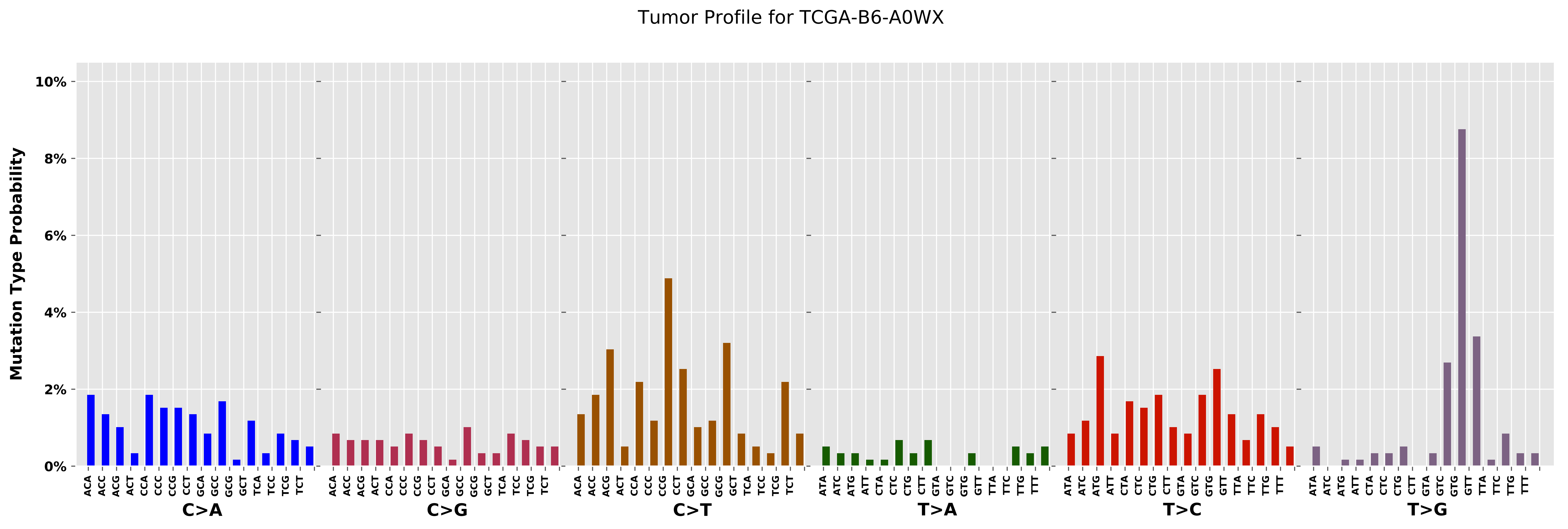

Tumor Profile for TCGA-B6-A0WX
Mutation Type Probability
10%
8%
6%
4%
2%
0%
C>A
C>G
C>T
T>A
T>C
T>G

### Cancer processes Weights for TCGA-BH-A0WA

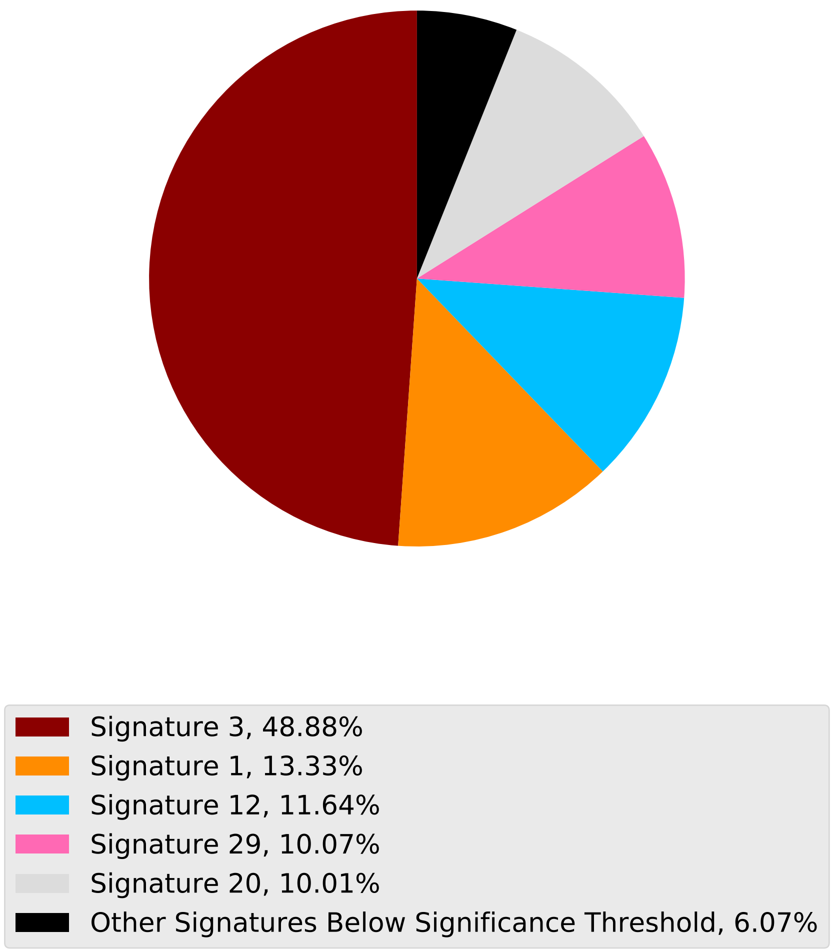

Tumor Profile for TCGA-BH-A0WA

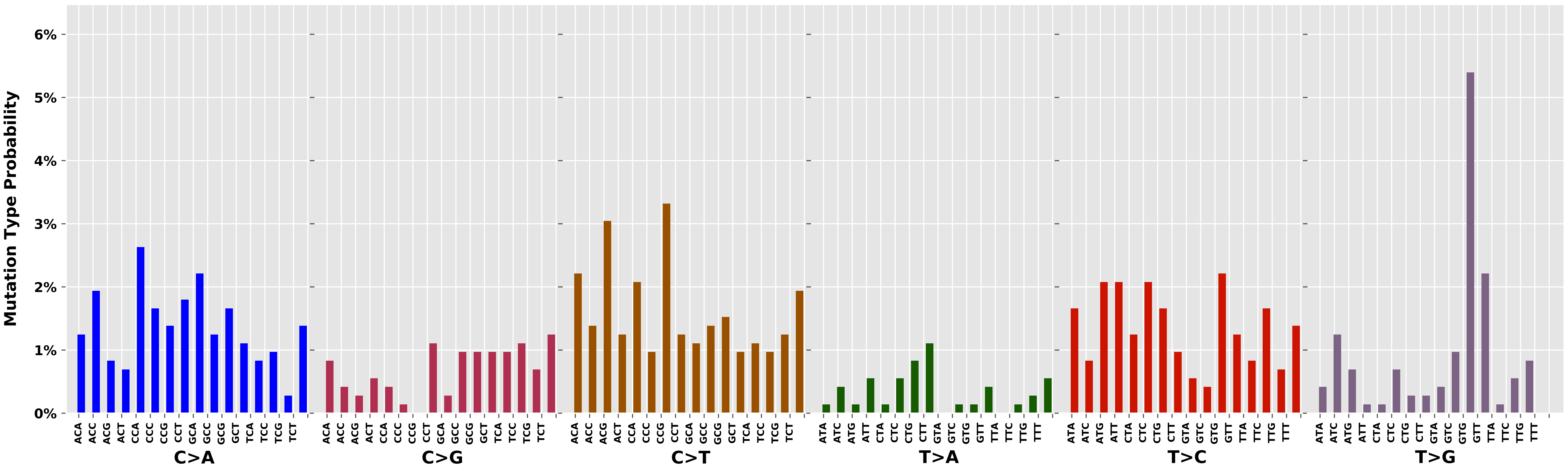

Cancer processes Weights for TCGA-A7-A13D

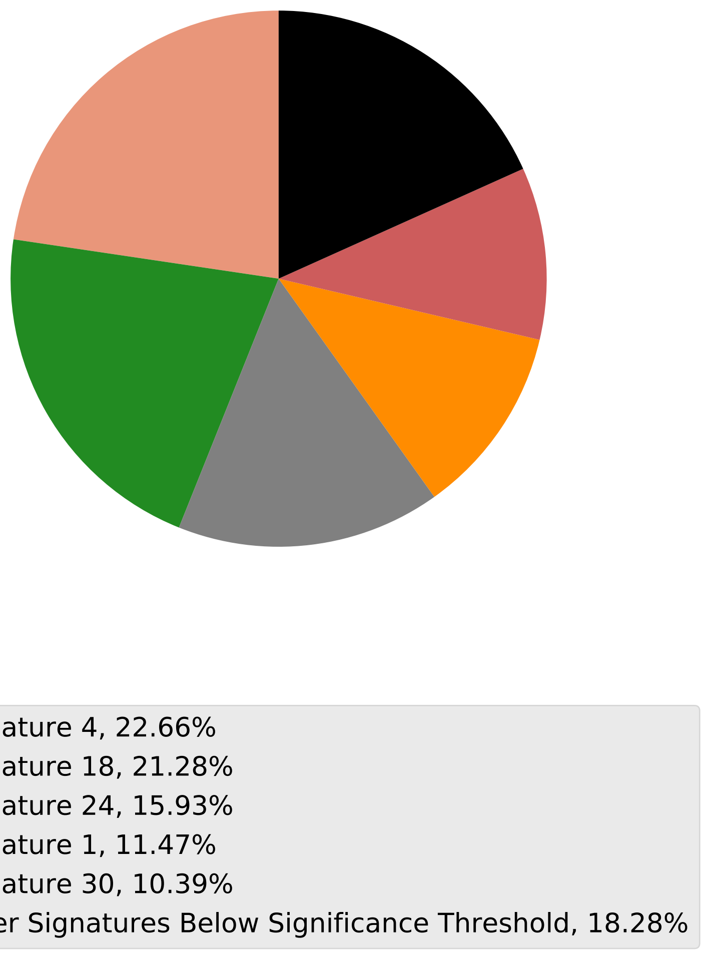

Tumor Profile for TCGA-A7-A13D

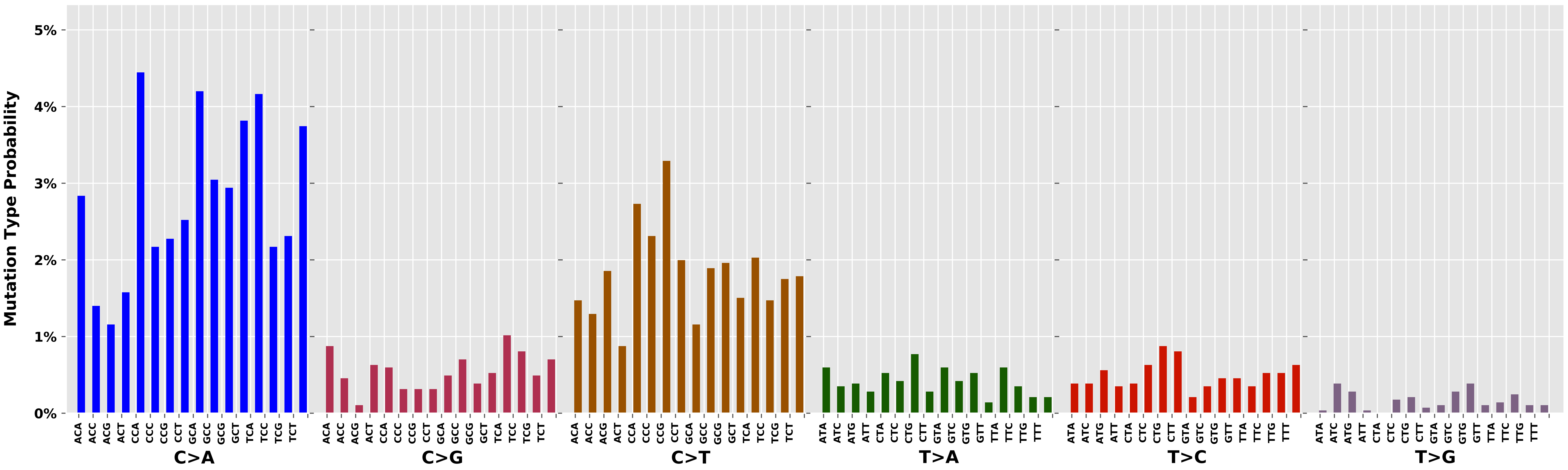

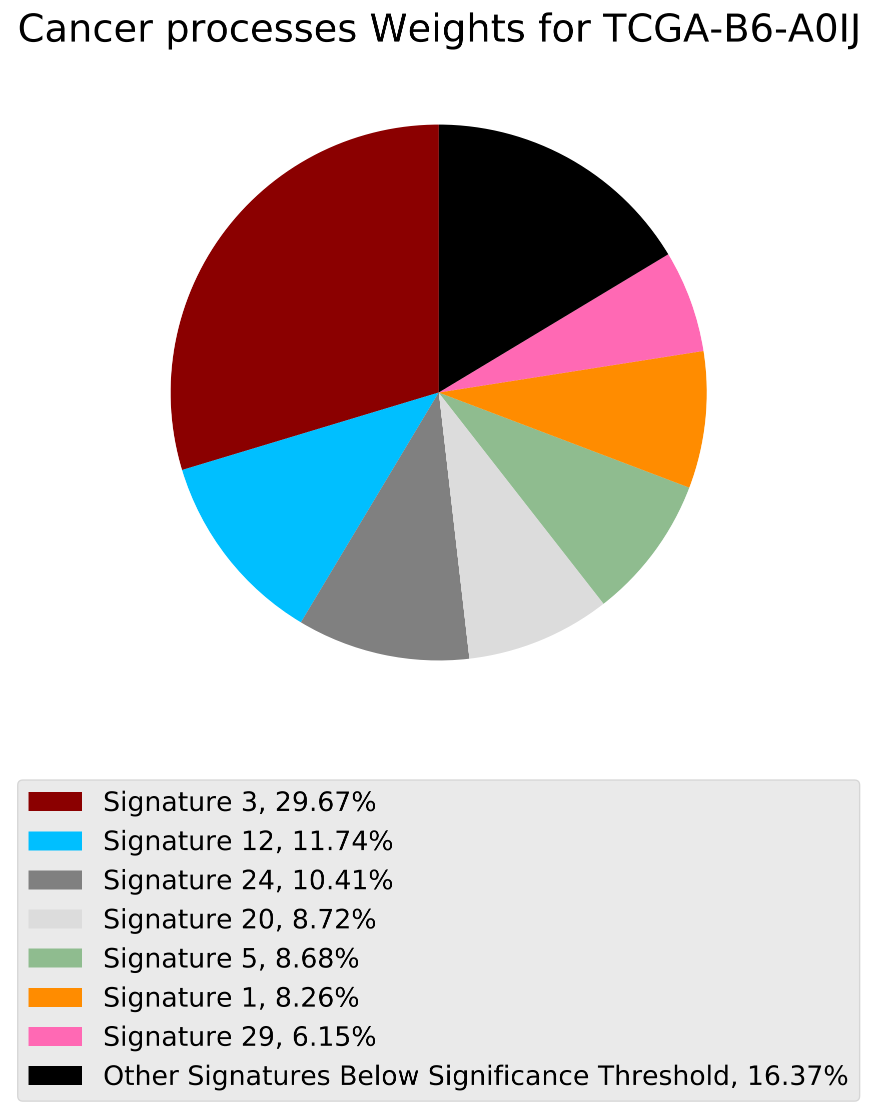
Cancer processes Weights for TCGA-B6-A0IJ
Signature 3, 29.67%
Signature 12, 11.74%
Signature 24, 10.41%
Signature 20, 8.72%
Signature 5, 8.68%
Signature 1, 8.26%
Signature 29, 6.15%
Other Signatures Below Significance Threshold, 16.37%

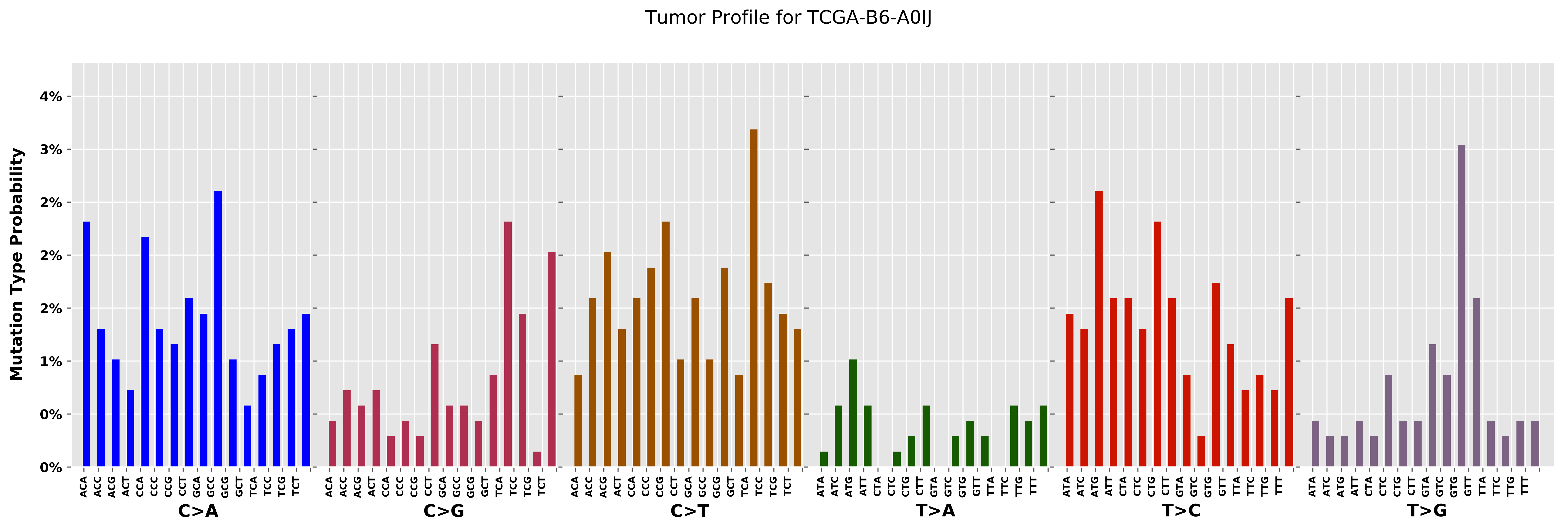

Tumor Profile for TCGA-B6-A0IJ
Mutation Type Probability
4%
3%
2%
2%
2%
1%
0%
0%
C>A
C>G
C>T
T>A
T>C
T>G
ACA
ACC
ACG
ACT
CCA
CCC
CCG
CCT
GCA
GCC
GCG
GCT
TCA
TCC
TCG
TCT
ATA
ATC
ATG
ATT
CTA
CTC
CTG
CTT
GTA
GTC
GTG
GTT
TTA
TTC
TTG
TTT

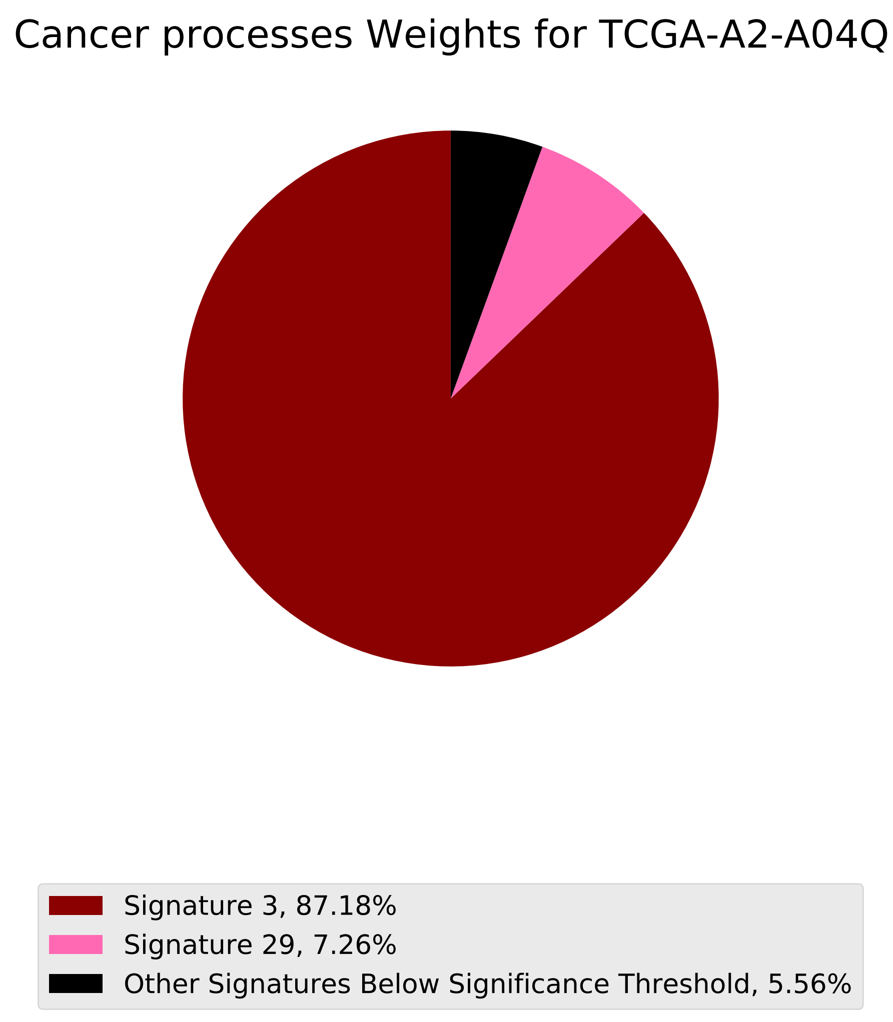
Cancer processes Weights for TCGA-A2-A04Q
Signature 3, 87.18%
Signature 29, 7.26%
Other Signatures Below Significance Threshold, 5.56%

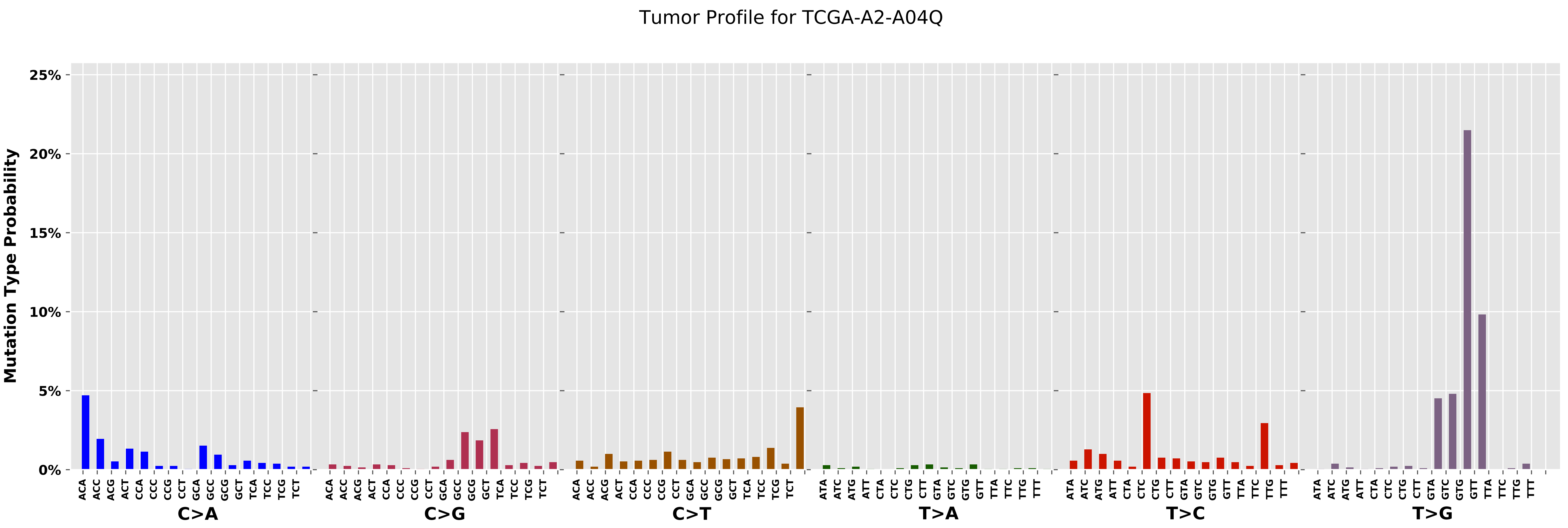
Tumor Profile for TCGA-A2-A04Q
Mutation Type Probability
25%
20%
15%
10%
5%
0%
C>A
C>G
C>T
T>A
T>C
T>G

Cancer processes Weights for TCGA-BH-A0B3

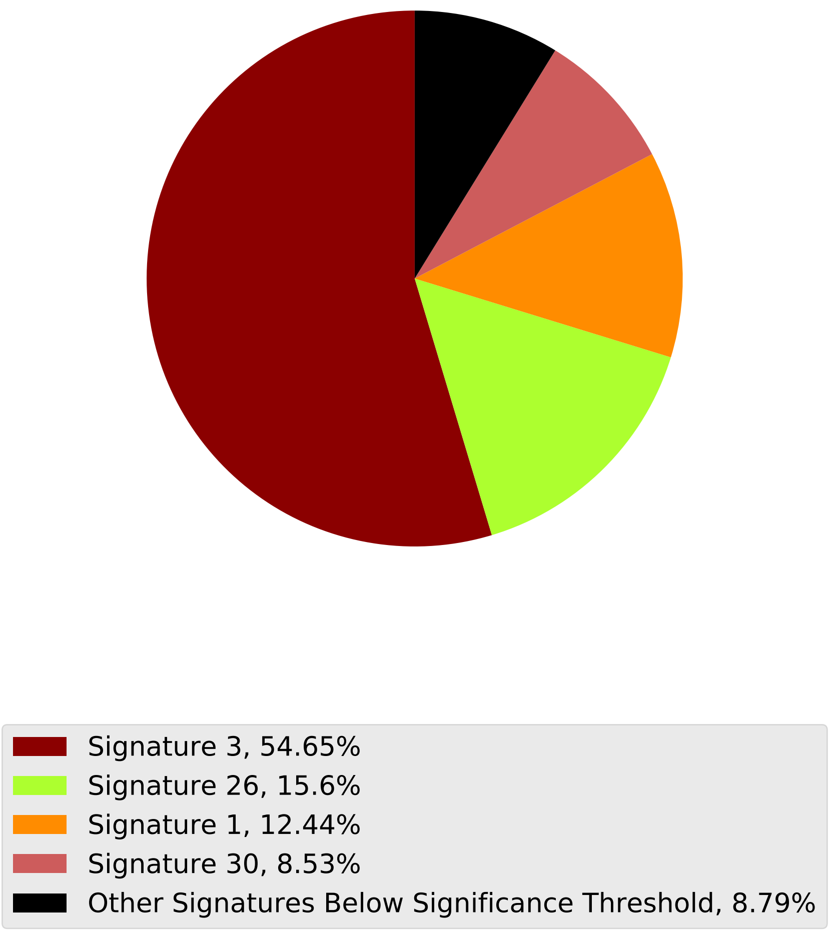

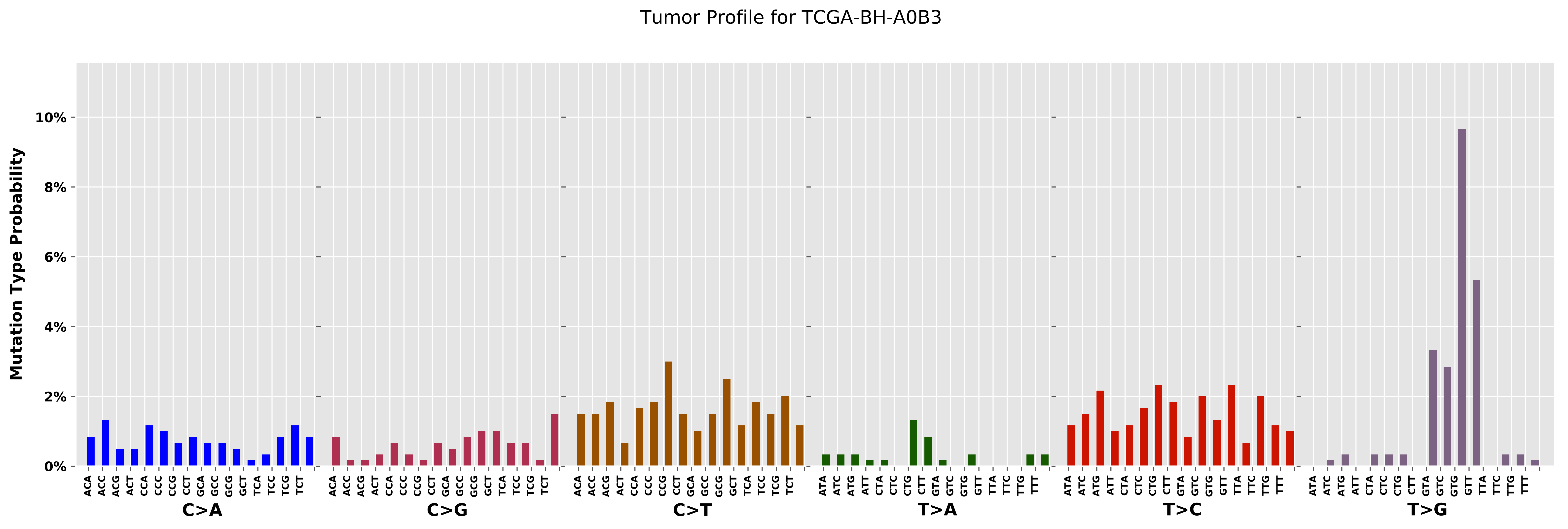

Tumor Profile for TCGA-BH-A0B3
Mutation Type Probability
10%
8%
6%
4%
2%
0%
C>A
C>G
C>T
T>A
T>C
T>G

Cancer processes Weights for TCGA-BH-A18U

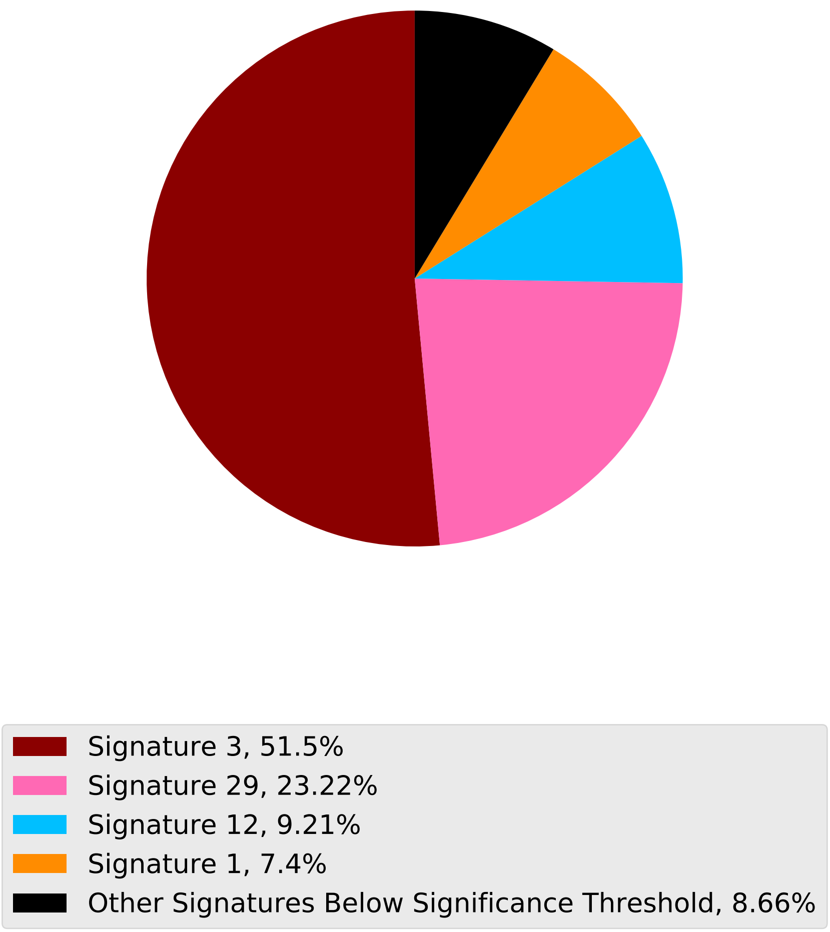

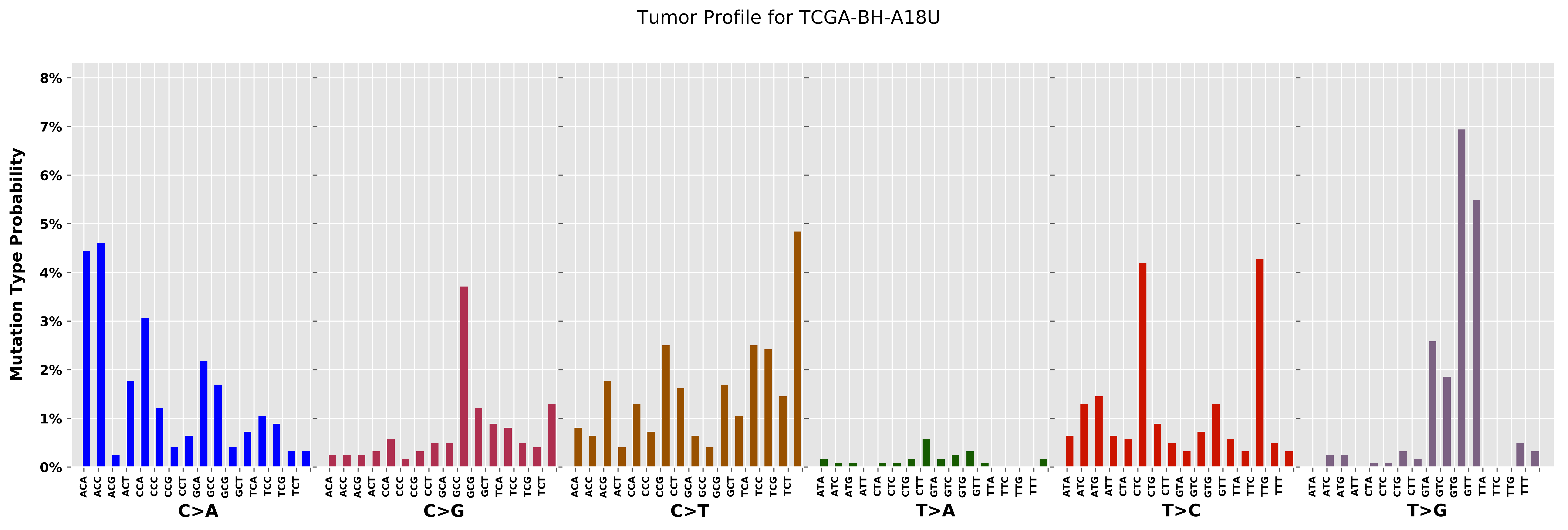

Tumor Profile for TCGA-BH-A18U
Mutation Type Probability
8%
7%
6%
5%
4%
3%
2%
1%
0%
C>A
C>G
C>T
T>A
T>C
T>G

Cancer processes Weights for TCGA-GI-A2C9

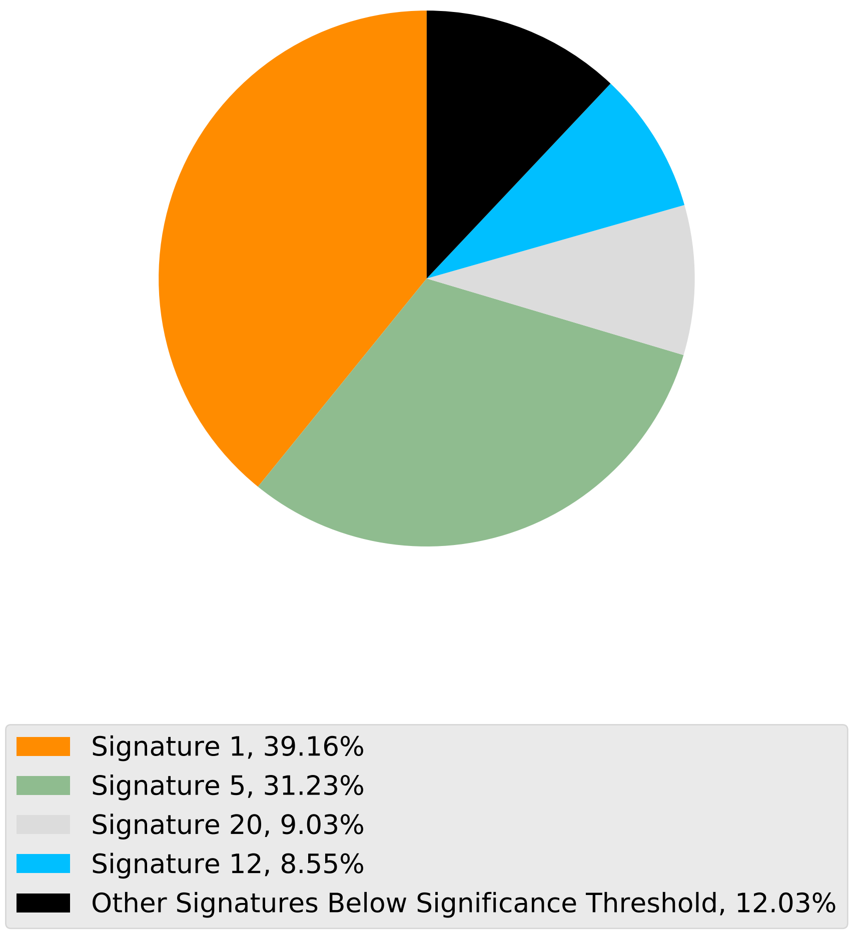

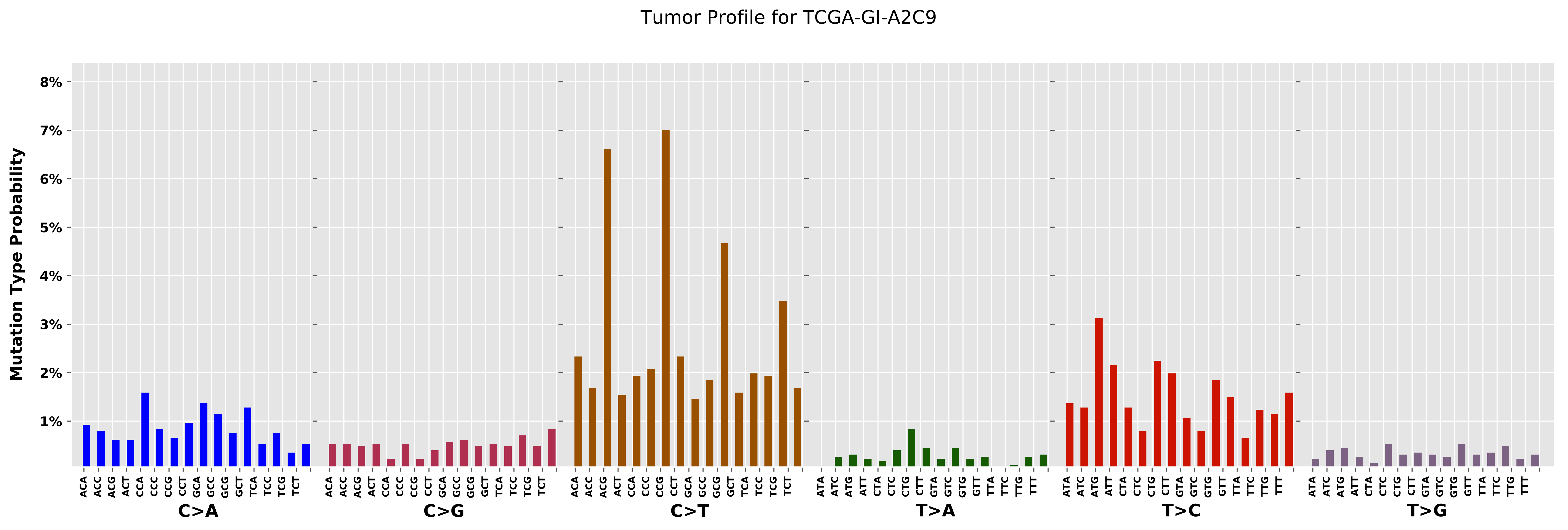

Tumor Profile for TCGA-GI-A2C9
Mutation Type Probability
8%
7%
6%
5%
4%
3%
2%
1%
ACA ACC ACG ACT CCA CCC CCG CCT GCA GCC GCG GCT TCA TCC TCG TCT
ATA ATC ATG ATT CTA CTC CTG CTT GTA GTC GTG GTT TTA TTC TTG TTT
C>A
C>G
C>T
T>A
T>C
T>G

Cancer processes Weights for TCGA-BH-A0DT

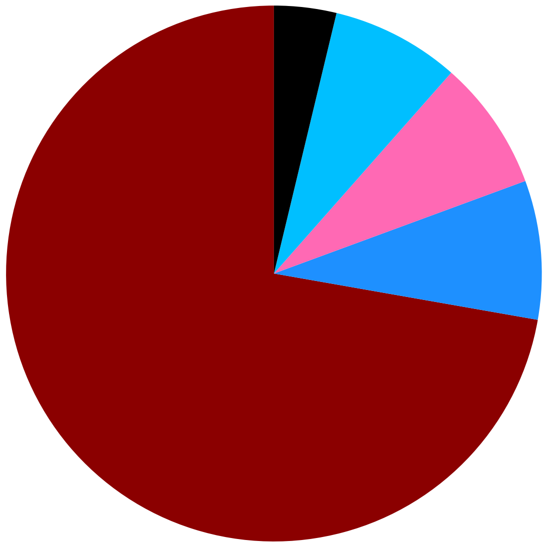

Signature 3, 72.23%
Signature 15, 8.38%
Signature 29, 7.87%
Signature 12, 7.76%
Other Signatures Below Significance Threshold, 3.76%

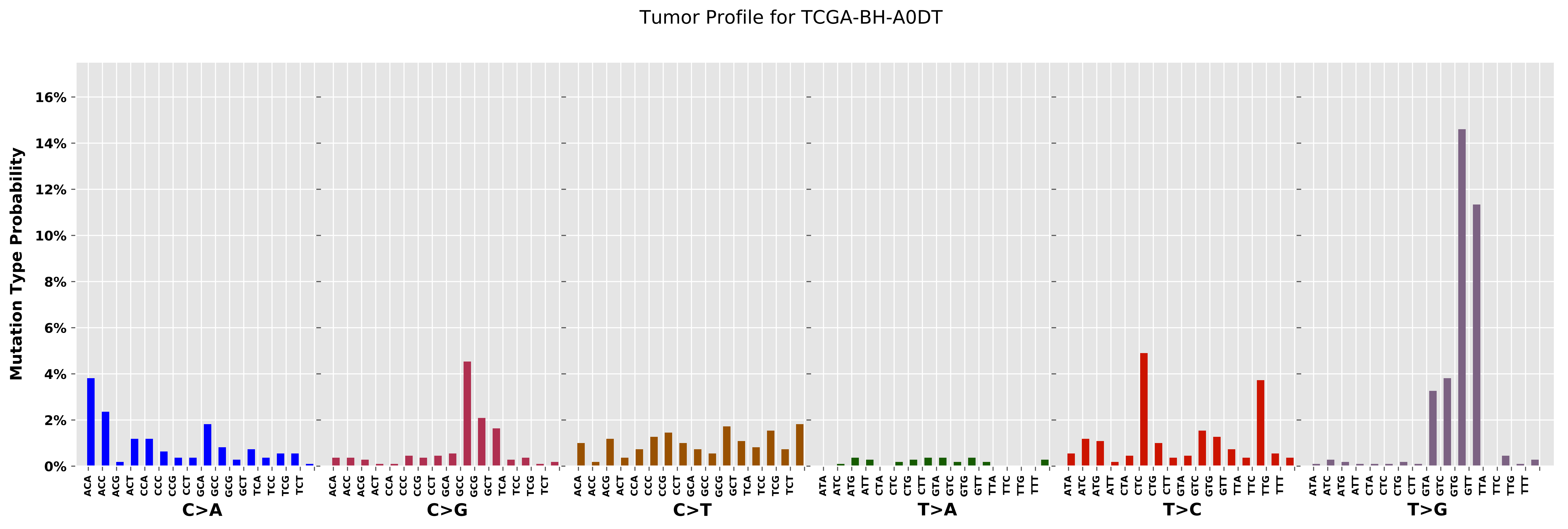

Tumor Profile for TCGA-BH-A0DT
Mutation Type Probability
16%
14%
12%
10%
8%
6%
4%
2%
0%
C>A
C>G
C>T
T>A
T>C
T>G
ACA ACC ACG ACT CCA CCC CCG CCT GCA GCC GCG GCT TCA TCC TCG TCT
ATA ATC ATG ATT CTA CTC CTG CTT GTA GTC GTG GTT TTA TTC TTG TTT

Cancer processes Weights for TCGA-A2-A0D4

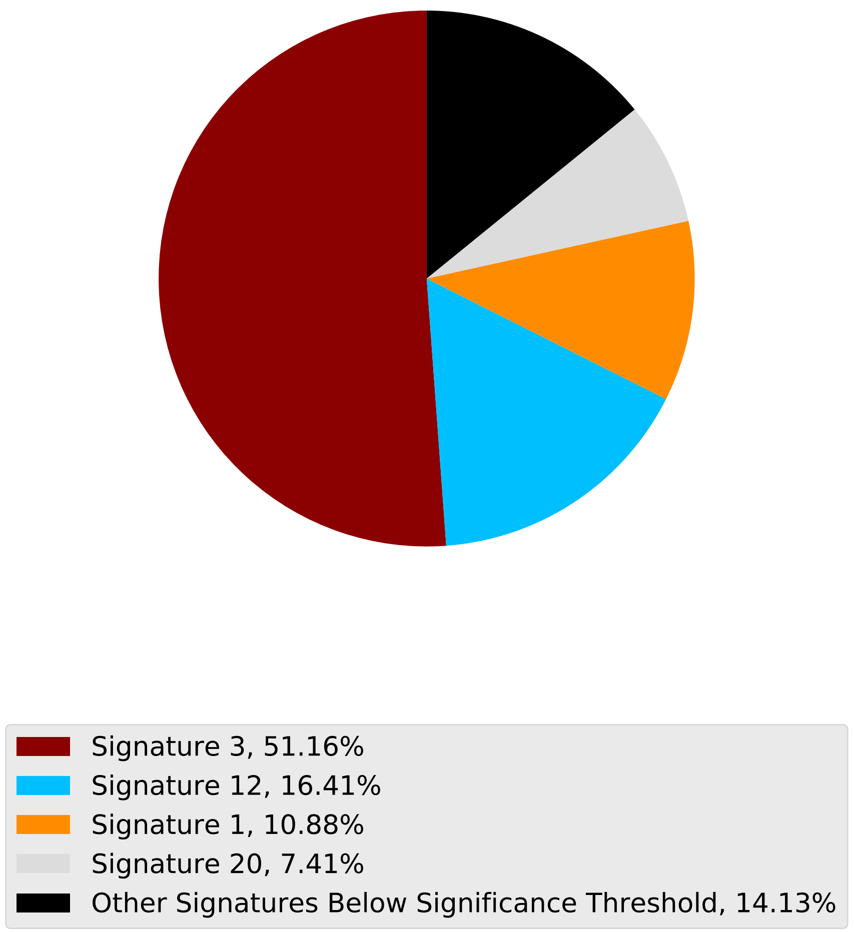

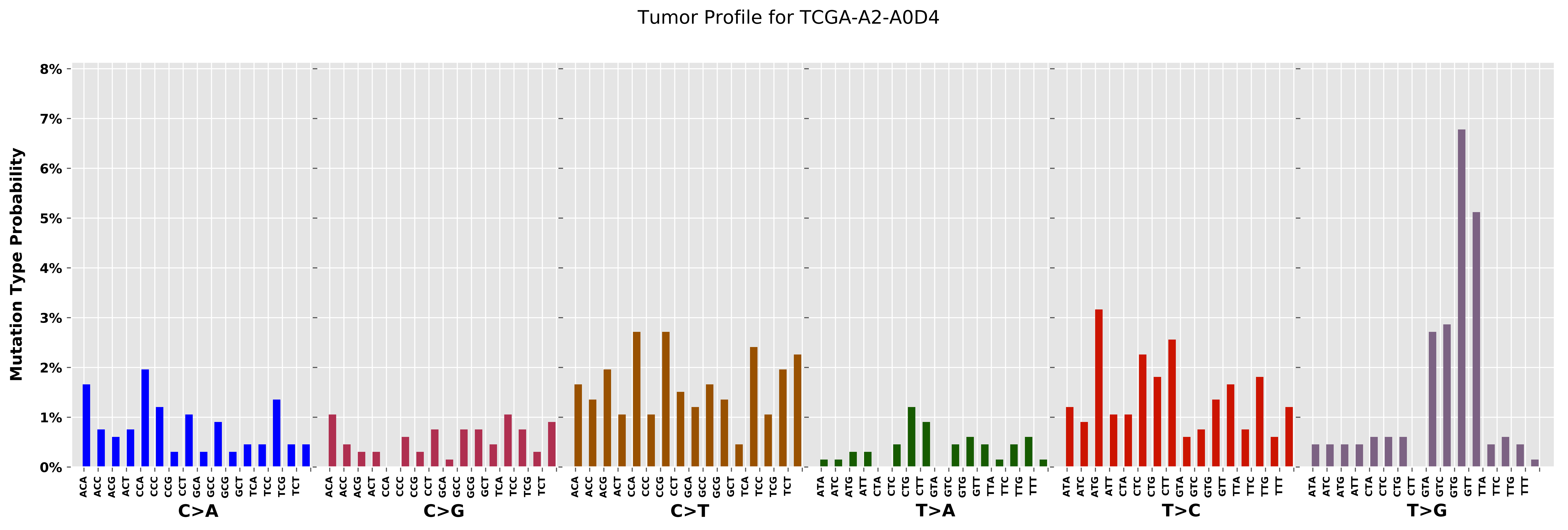
Tumor Profile for TCGA-A2-A0D4
Mutation Type Probability
0%
1%
2%
3%
4%
5%
6%
7%
8%
C>A
C>G
C>T
T>A
T>C
T>G
ACA ACC ACG ACT CCA CCC CCG CCT GCA GCC GCG GCT TCA TCC TCG TCT
ATA ATC ATG ATT CTA CTC CTG CTT GTA GTC GTG GTT TTA TTC TTG TTT

Cancer processes Weights for TCGA-EW-A1P8

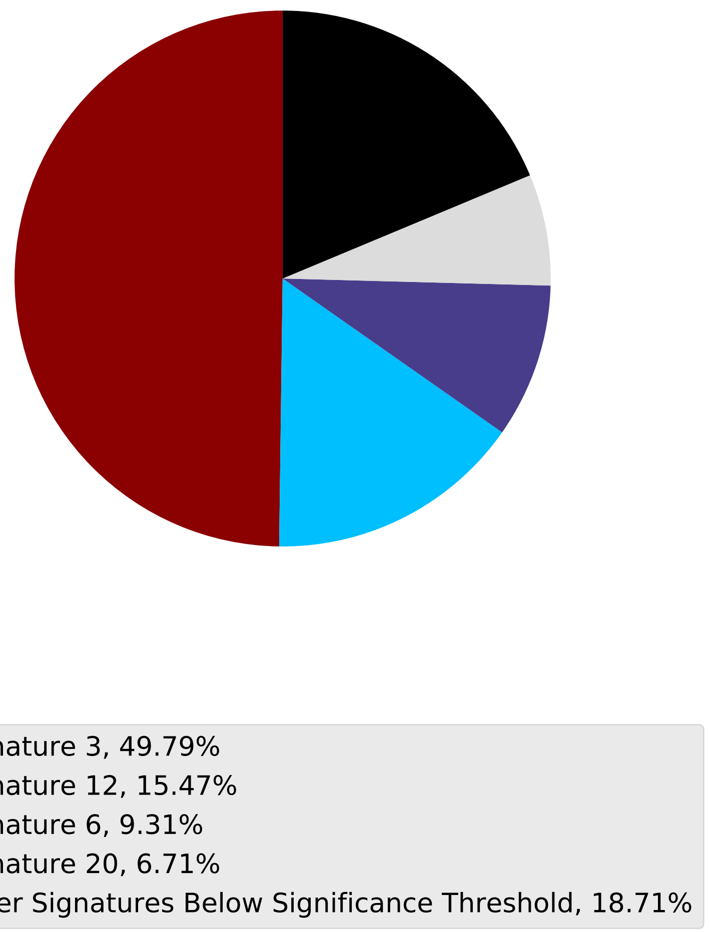

Tumor Profile for TCGA-EW-A1P8

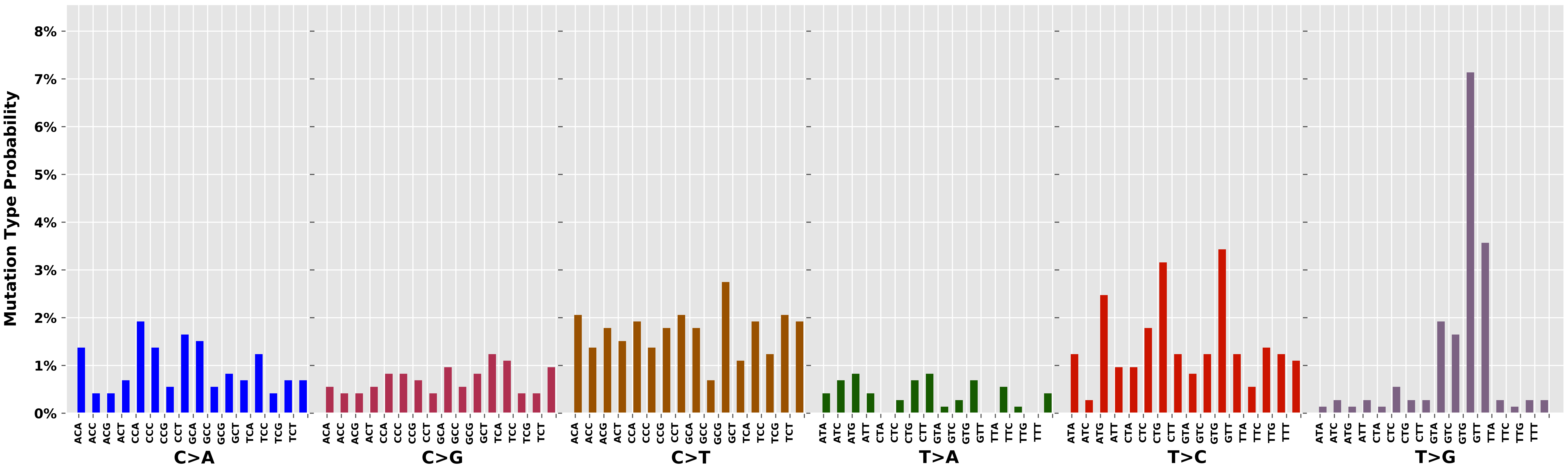

Cancer processes Weights for TCGA-A6-2683

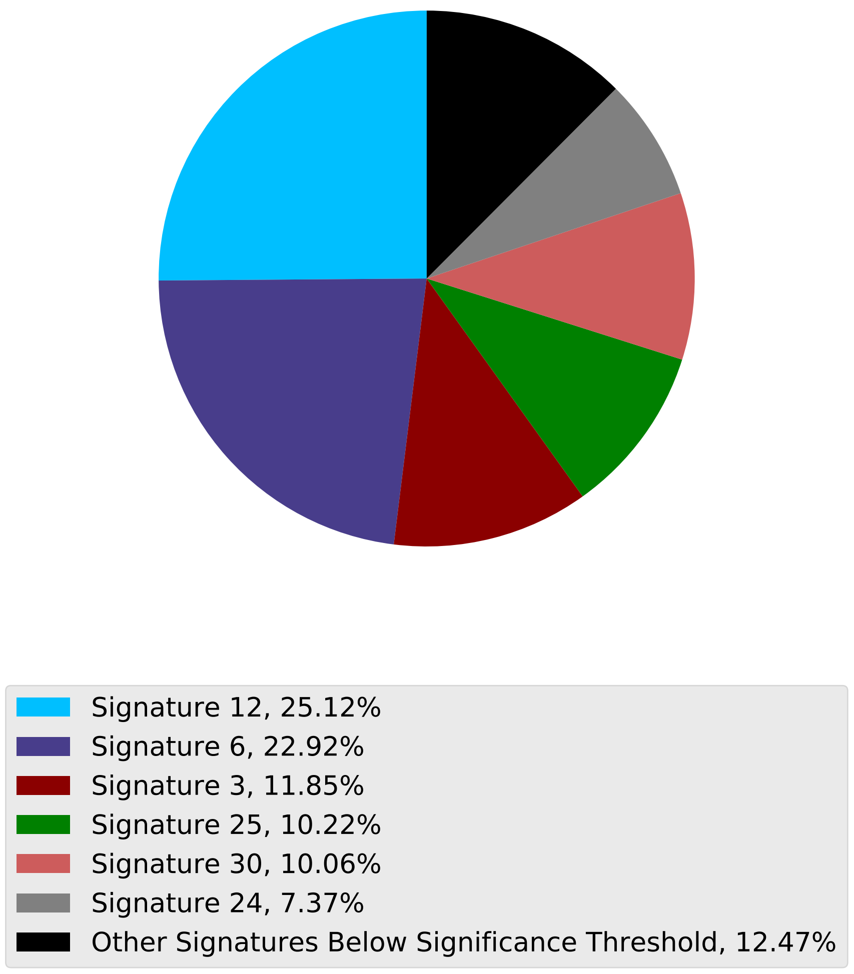

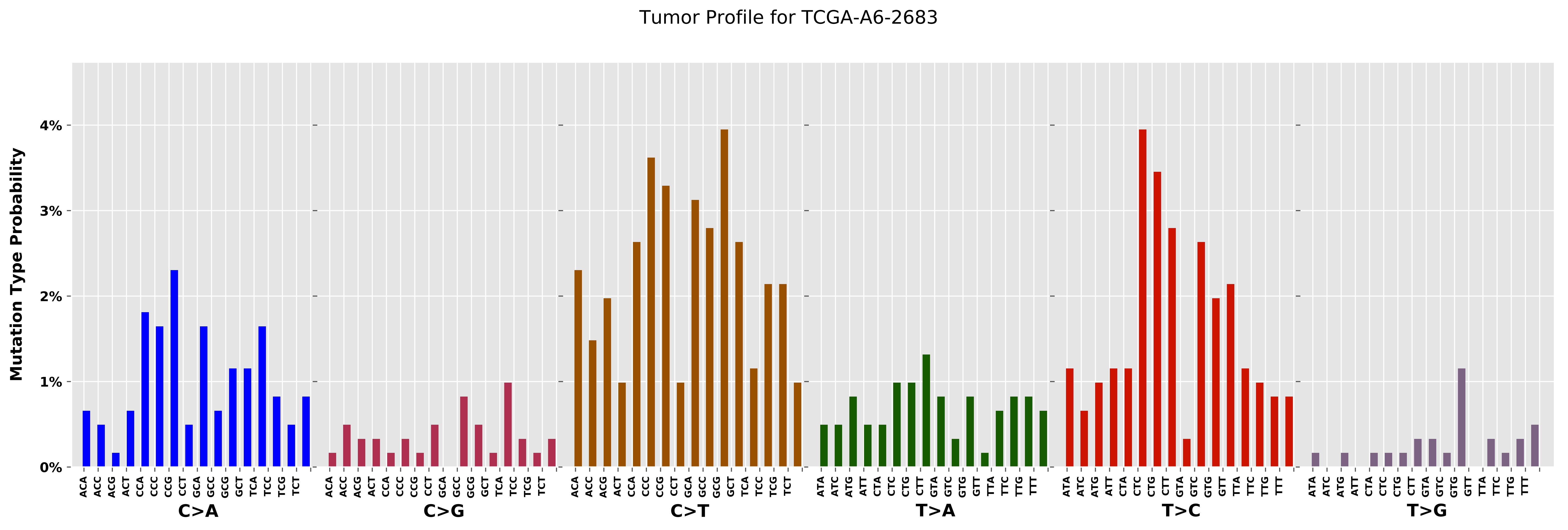

Tumor Profile for TCGA-A6-2683
Mutation Type Probability
0%
1%
2%
3%
4%
C>A
C>G
C>T
T>A
T>C
T>G

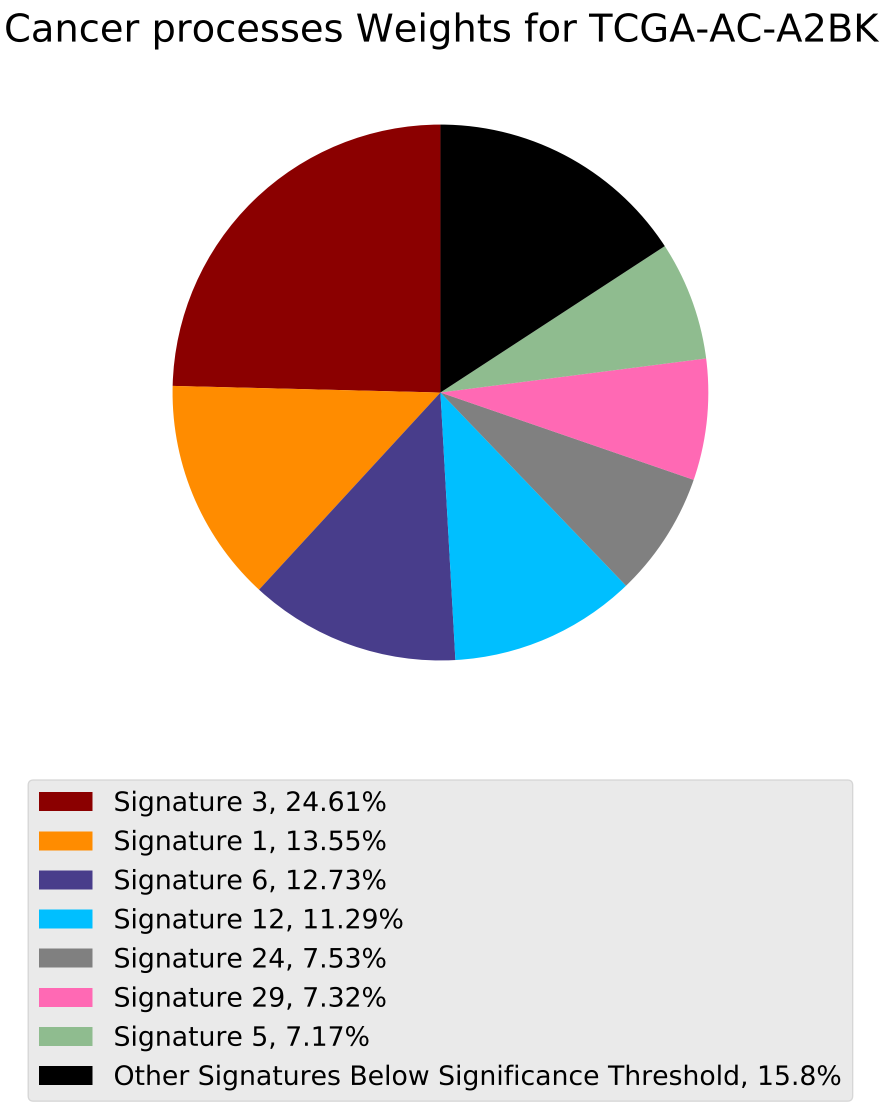

Cancer processes Weights for TCGA-AC-A2BK
Signature 3, 24.61%
Signature 1, 13.55%
Signature 6, 12.73%
Signature 12, 11.29%
Signature 24, 7.53%
Signature 29, 7.32%
Signature 5, 7.17%
Other Signatures Below Significance Threshold, 15.8%

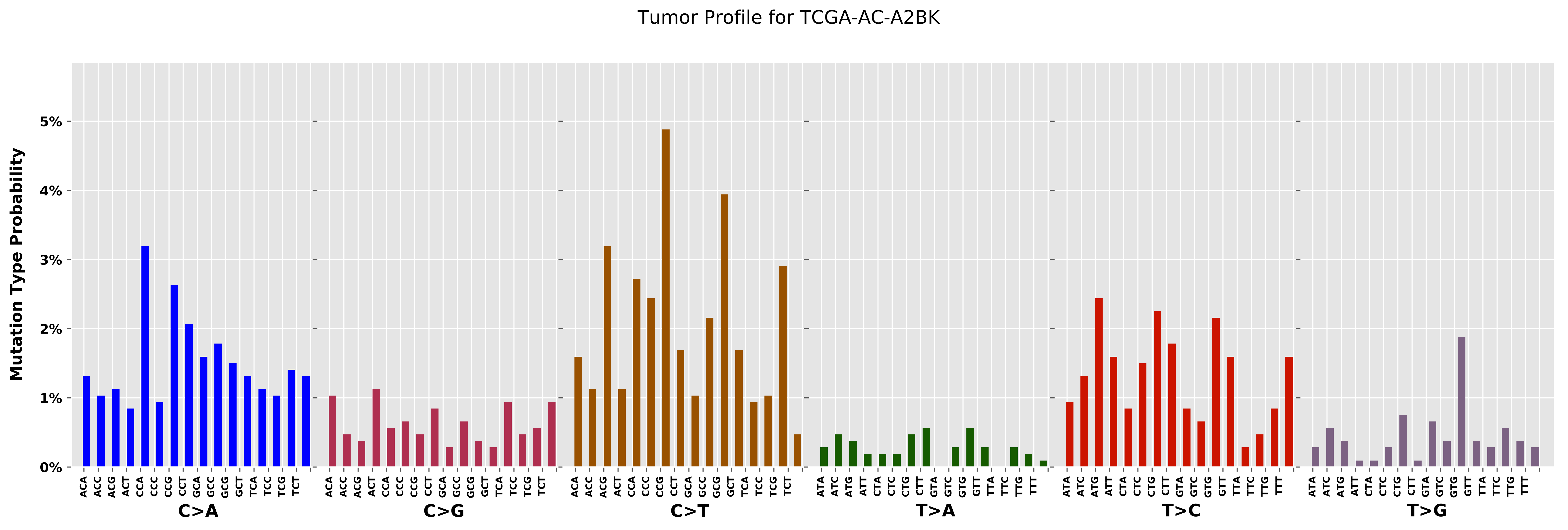

Tumor Profile for TCGA-AC-A2BK
Mutation Type Probability
0%
1%
2%
3%
4%
5%
C>A
C>G
C>T
T>A
T>C
T>G
ACA
ACC
ACG
ACT
CCA
CCC
CCG
CCT
GCA
GCC
GCG
GCT
TCA
TCC
TCG
TCT
ATA
ATC
ATG
ATT
CTA
CTC
CTG
CTT
GTA
GTC
GTG
GTT
TTA
TTC
TTG
TTT

Cancer processes Weights for TCGA-AG-A032

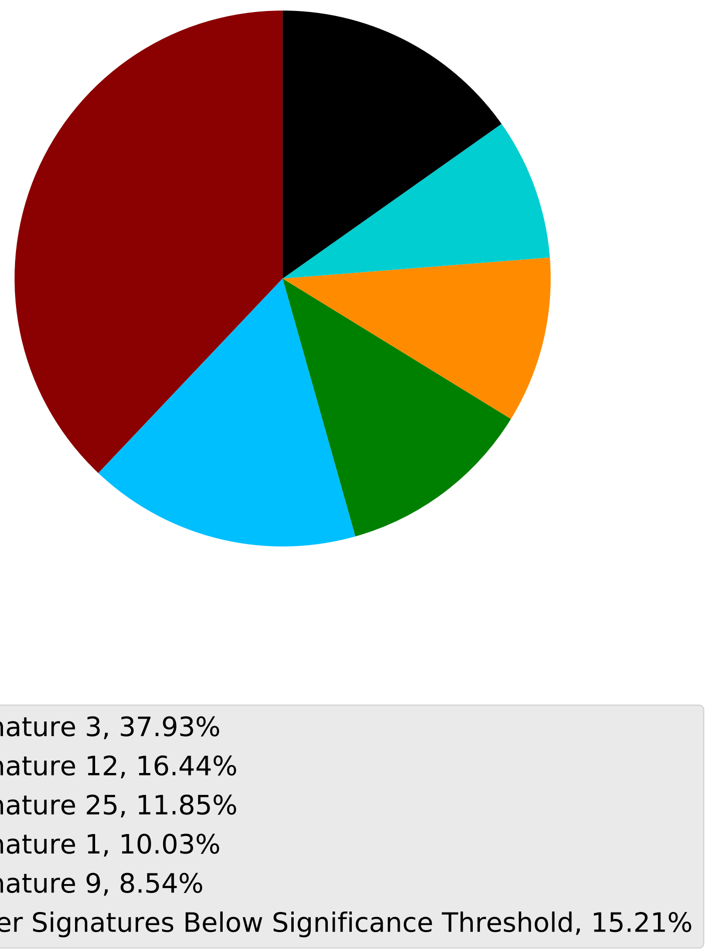

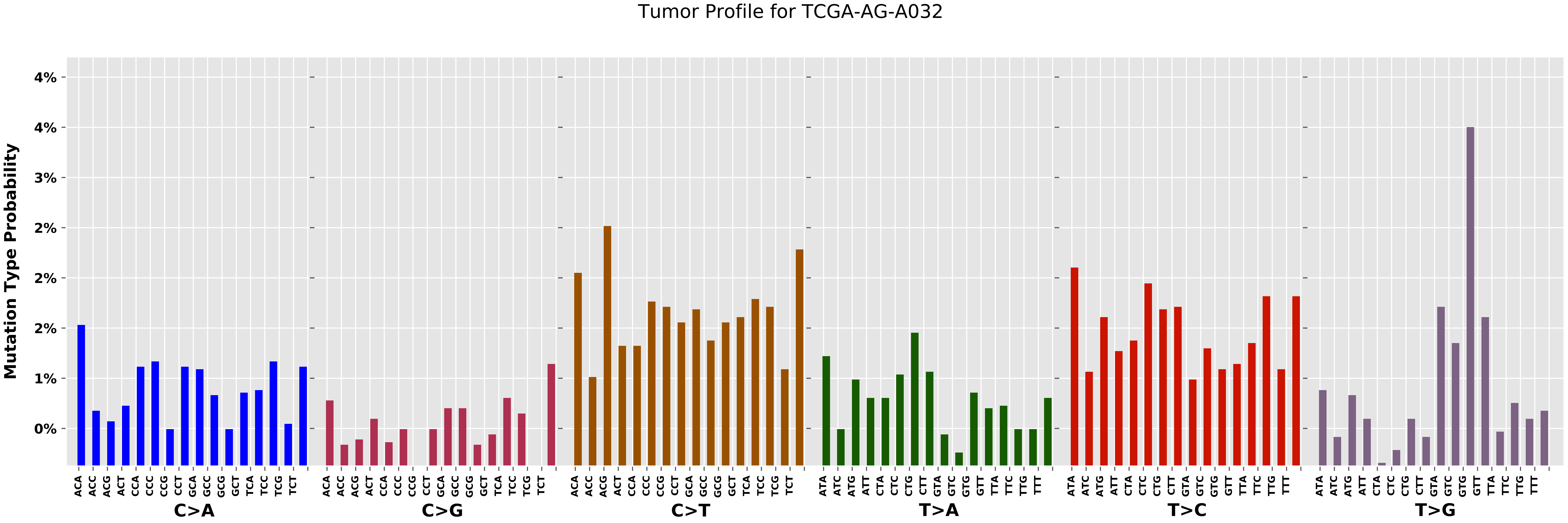

Tumor Profile for TCGA-AG-A032
Mutation Type Probability
4%
4%
3%
2%
2%
2%
1%
0%
ACA
ACC
ACG
ACT
CCA
CCC
CCG
CCT
GCA
GCC
GCG
GCT
TCA
TCC
TCG
TCT
ATA
ATC
ATG
ATT
CTA
CTC
CTG
CTT
GTA
GTC
GTG
GTT
TTA
TTC
TTG
TTT
C>A
C>G
C>T
T>A
T>C
T>G

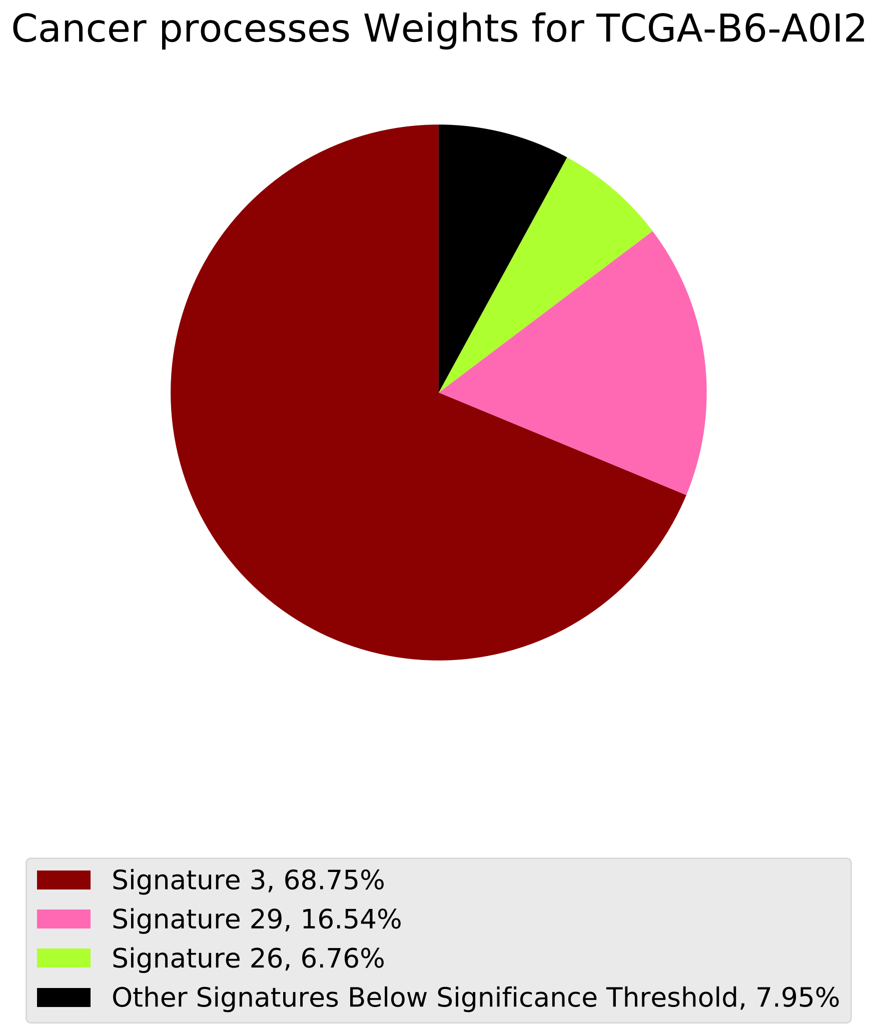
Cancer processes Weights for TCGA-B6-A0I2
Signature 3, 68.75%
Signature 29, 16.54%
Signature 26, 6.76%
Other Signatures Below Significance Threshold, 7.95%

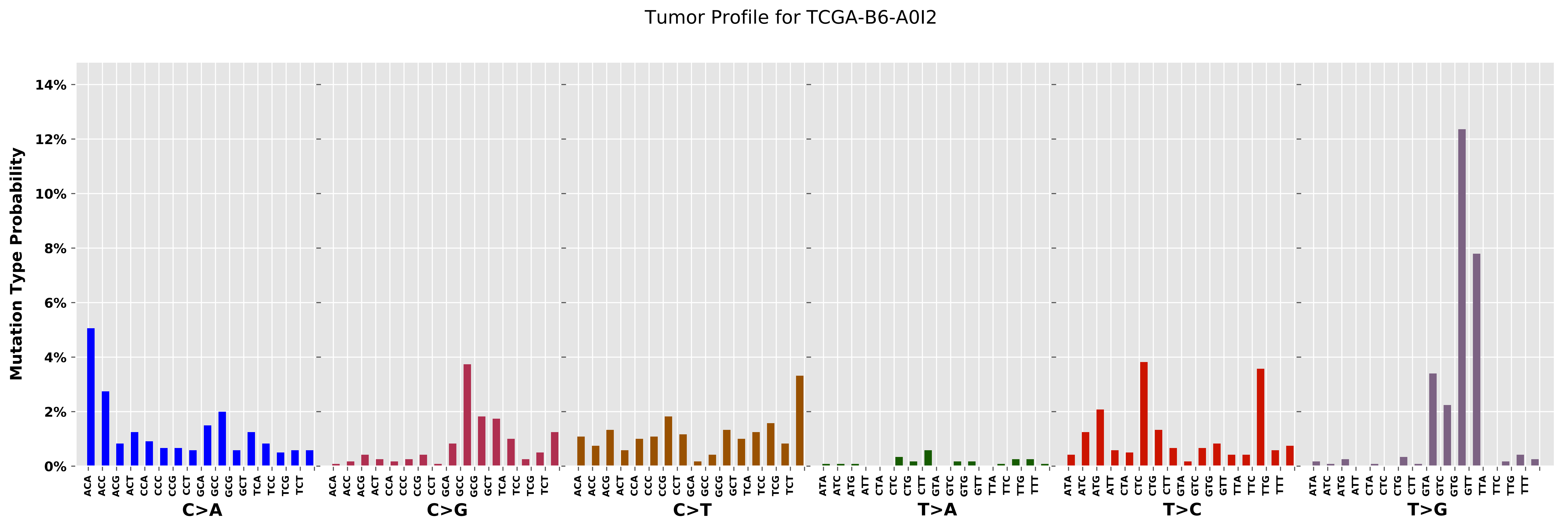

Tumor Profile for TCGA-B6-A0I2
Mutation Type Probability
14%
12%
10%
8%
6%
4%
2%
0%
C>A
C>G
C>T
T>A
T>C
T>G
ACA ACC ACG ACT CCA CCC CCG CCT GCA GCC GCG GCT TCA TCC TCG TCT
ATA ATC ATG ATT CTA CTC CTG CTT GTA GTC GTG GTT TTA TTC TTG TTT

Cancer processes Weights for TCGA-A8-A094

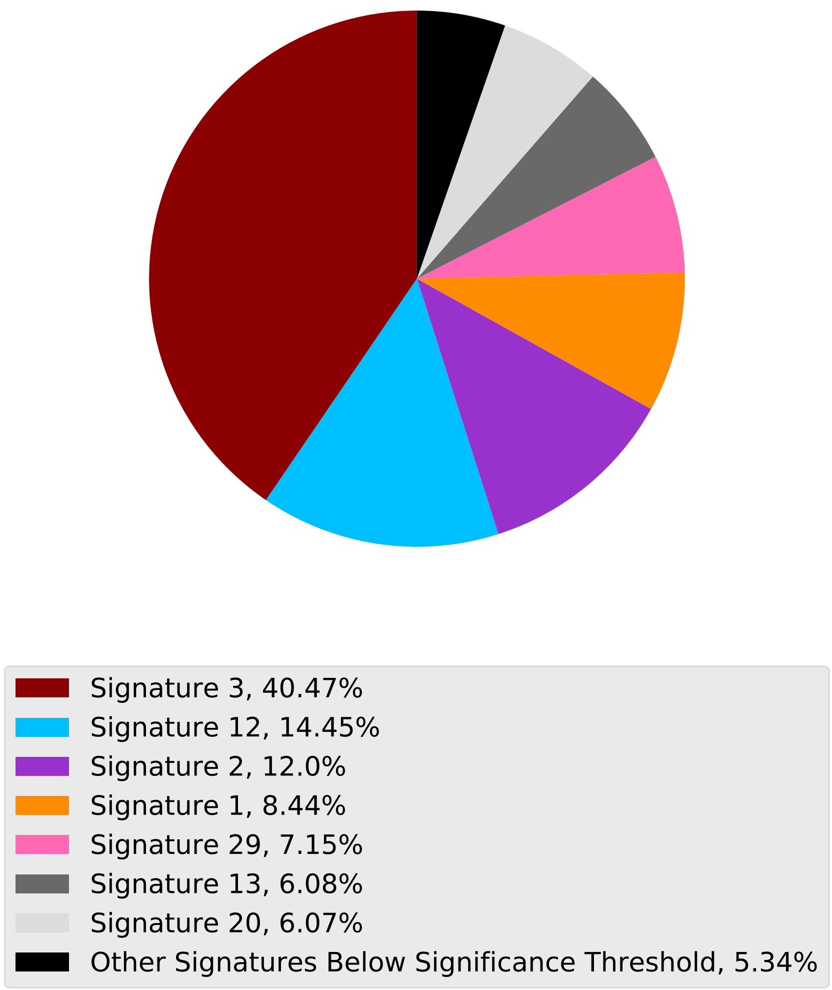

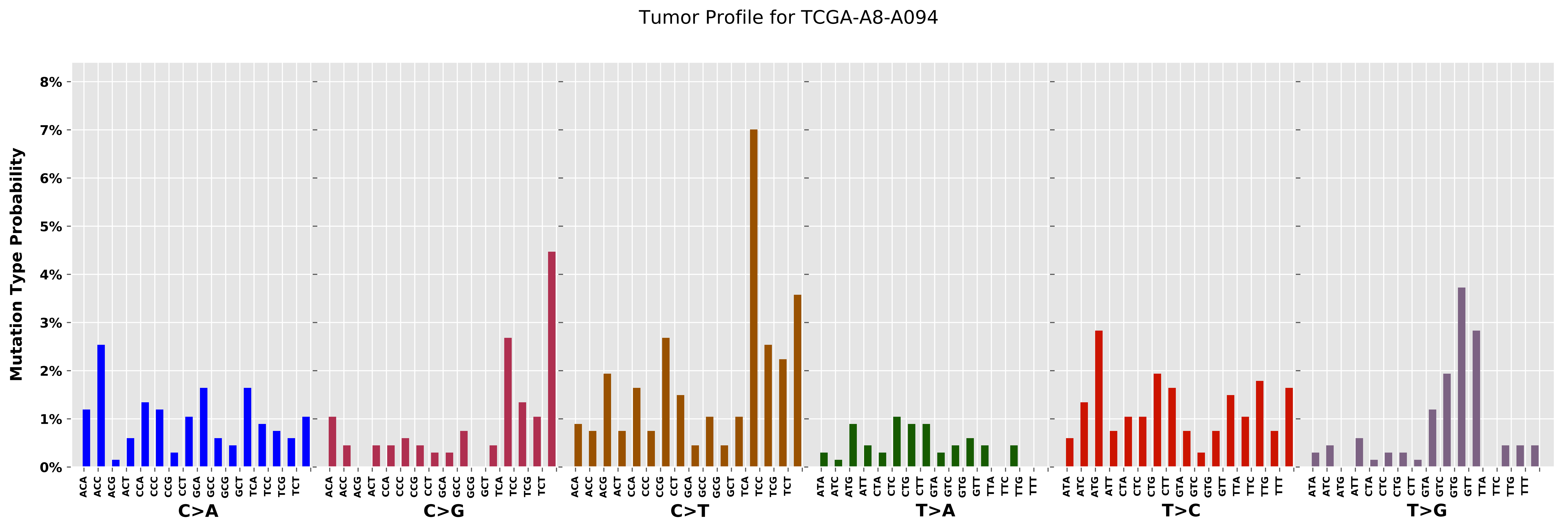

Tumor Profile for TCGA-A8-A094
Mutation Type Probability
0%
1%
2%
3%
4%
5%
6%
7%
8%
C>A
C>G
C>T
T>A
T>C
T>G

### Cancer processes Weights for TCGA-AA-3994

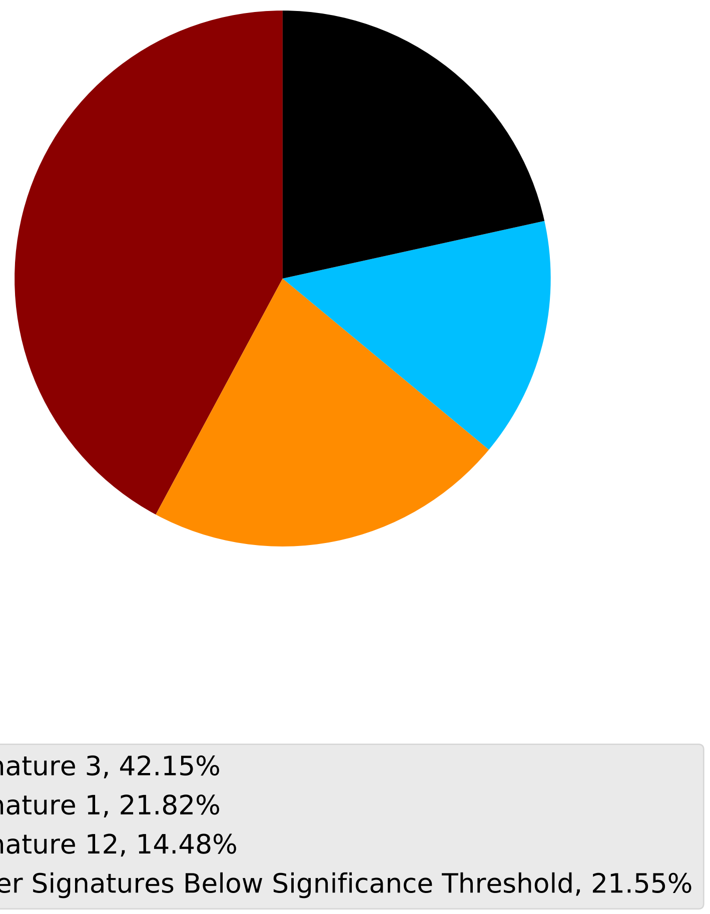

Tumor Profile for TCGA-AA-3994

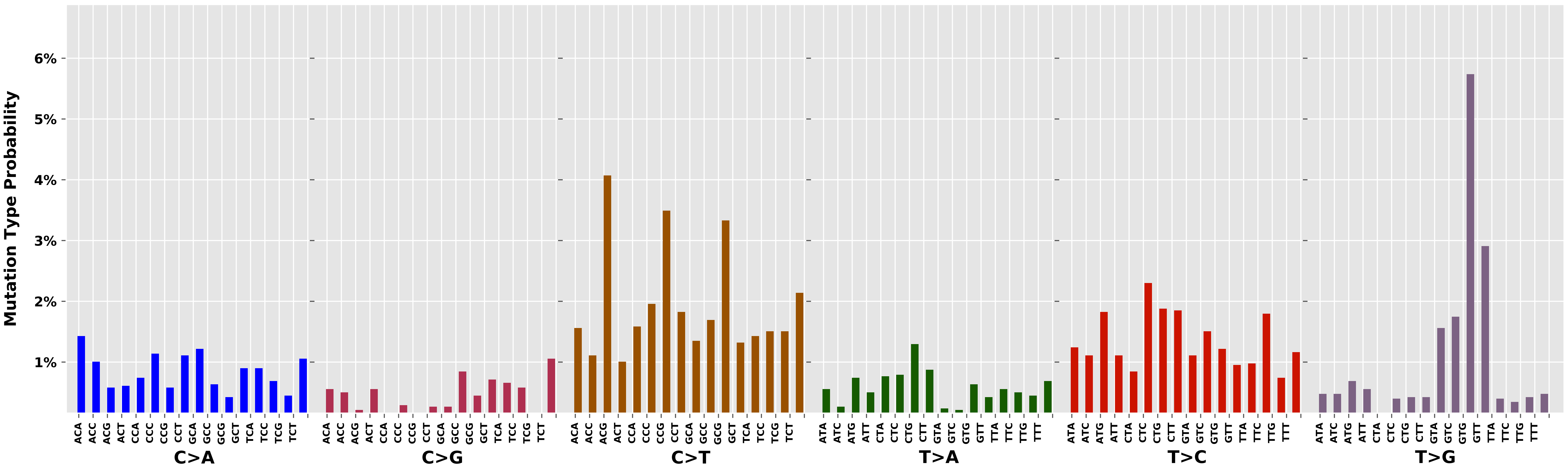

Cancer processes Weights for TCGA-AG-3574

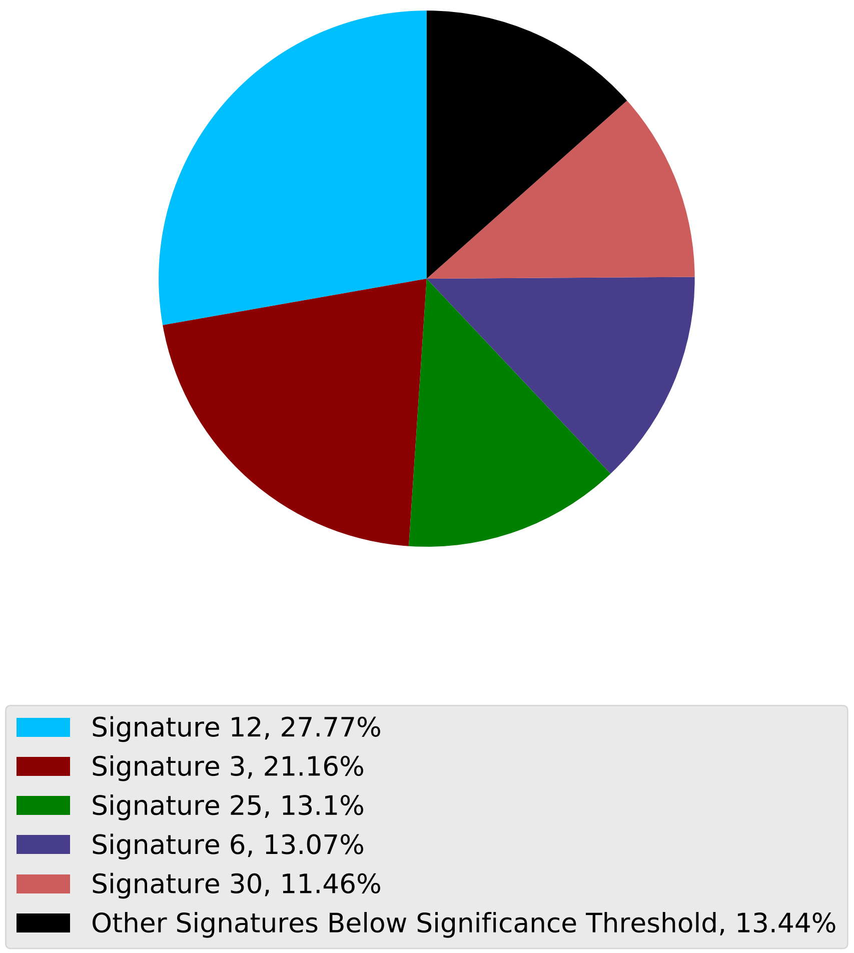

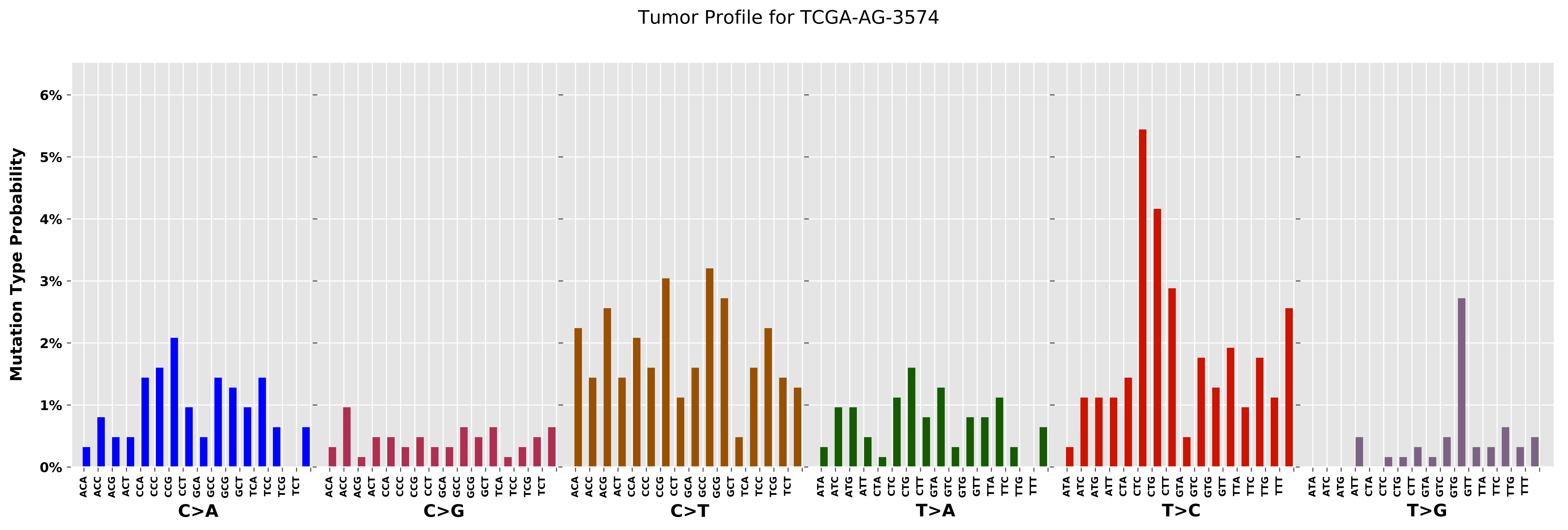

Tumor Profile for TCGA-AG-3574
Mutation Type Probability
0%
1%
2%
3%
4%
5%
6%
C>A
C>G
C>T
T>A
T>C
T>G

Cancer processes Weights for TCGA-AO-A03L

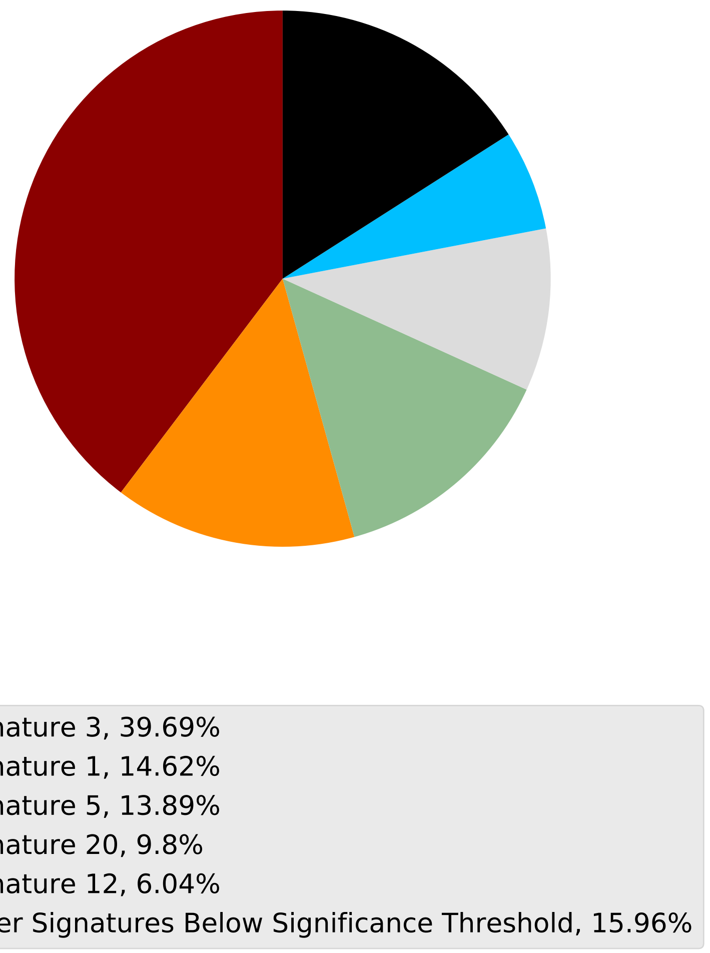

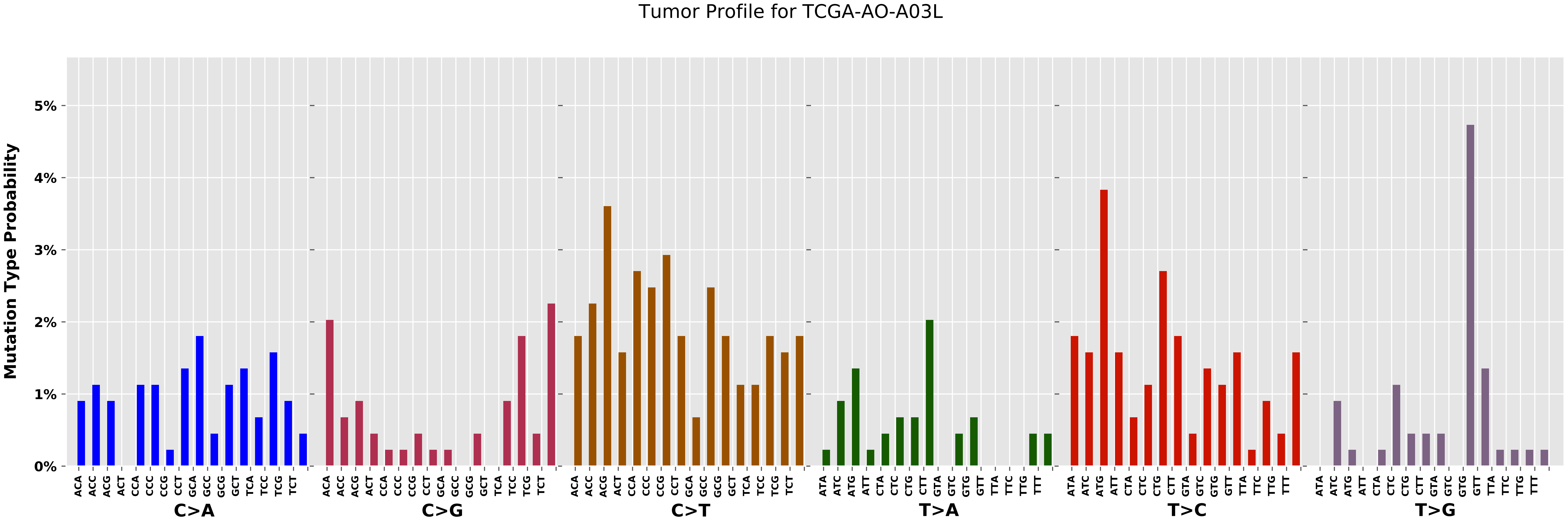
Tumor Profile for TCGA-AO-A03L
Mutation Type Probability
0%
1%
2%
3%
4%
5%
C>A
C>G
C>T
T>A
T>C
T>G
ACA
ACC
ACG
ACT
CCA
CCC
CCG
CCT
GCA
GCC
GCG
GCT
TCA
TCC
TCG
TCT
ATA
ATC
ATG
ATT
CTA
CTC
CTG
CTT
GTA
GTC
GTG
GTT
TTA
TTC
TTG
TTT

Cancer processes Weights for TCGA-A6-6141

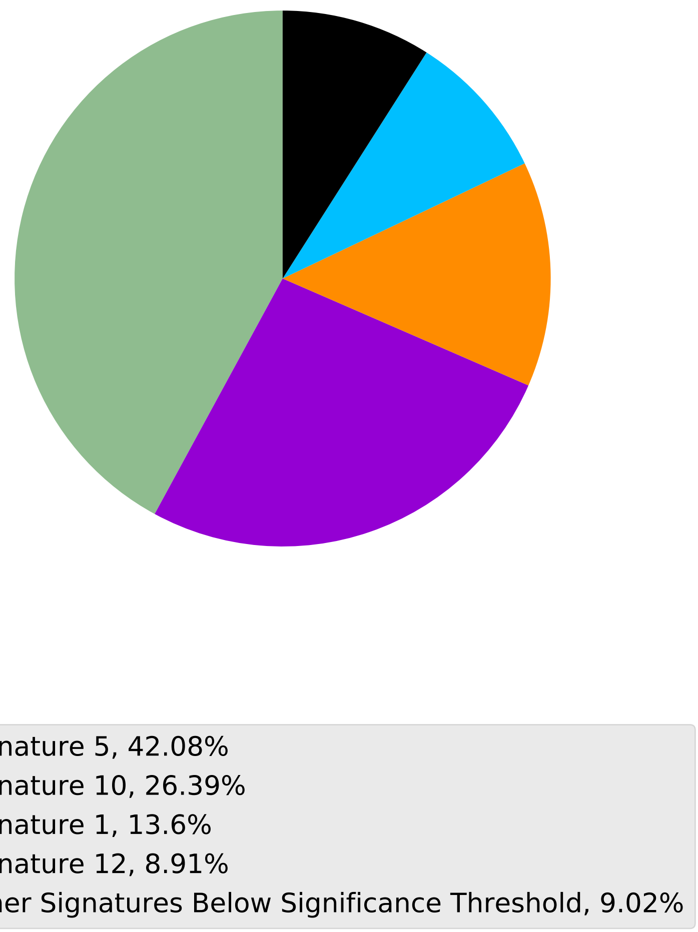

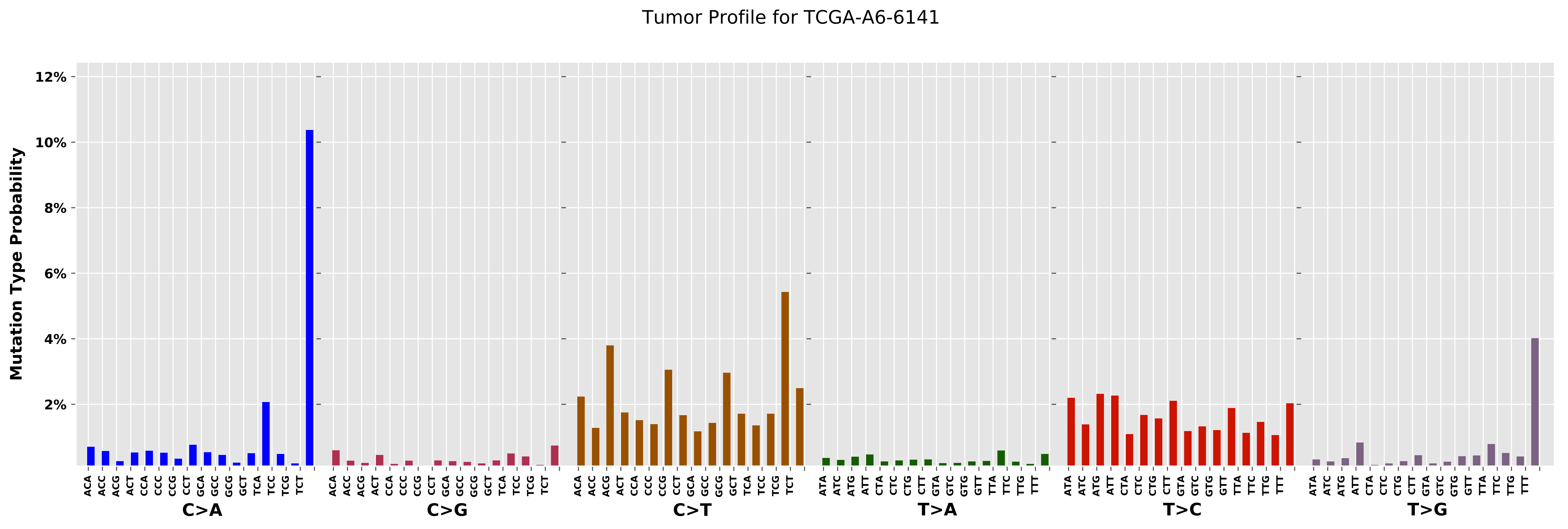

Tumor Profile for TCGA-A6-6141
Mutation Type Probability
12%
10%
8%
6%
4%
2%
ACA ACC ACG ACT CCA CCC CCG CCT GCA GCC GCG GCT TCA TCC TCG TCT
C>A
ACA ACC ACG ACT CCA CCC CCG CCT GCA GCC GCG GCT TCA TCC TCG TCT
C>G
ACA ACC ACG ACT CCA CCC CCG CCT GCA GCC GCG GCT TCA TCC TCG TCT
C>T
ATA ATC ATG ATT CTA CTC CTG CTT GTA GTC GTG GTT TTA TTC TTG TTT
T>A
ATA ATC ATG ATT CTA CTC CTG CTT GTA GTC GTG GTT TTA TTC TTG TTT
T>C
ATA ATC ATG ATT CTA CTC CTG CTT GTA GTC GTG GTT TTA TTC TTG TTT
T>G

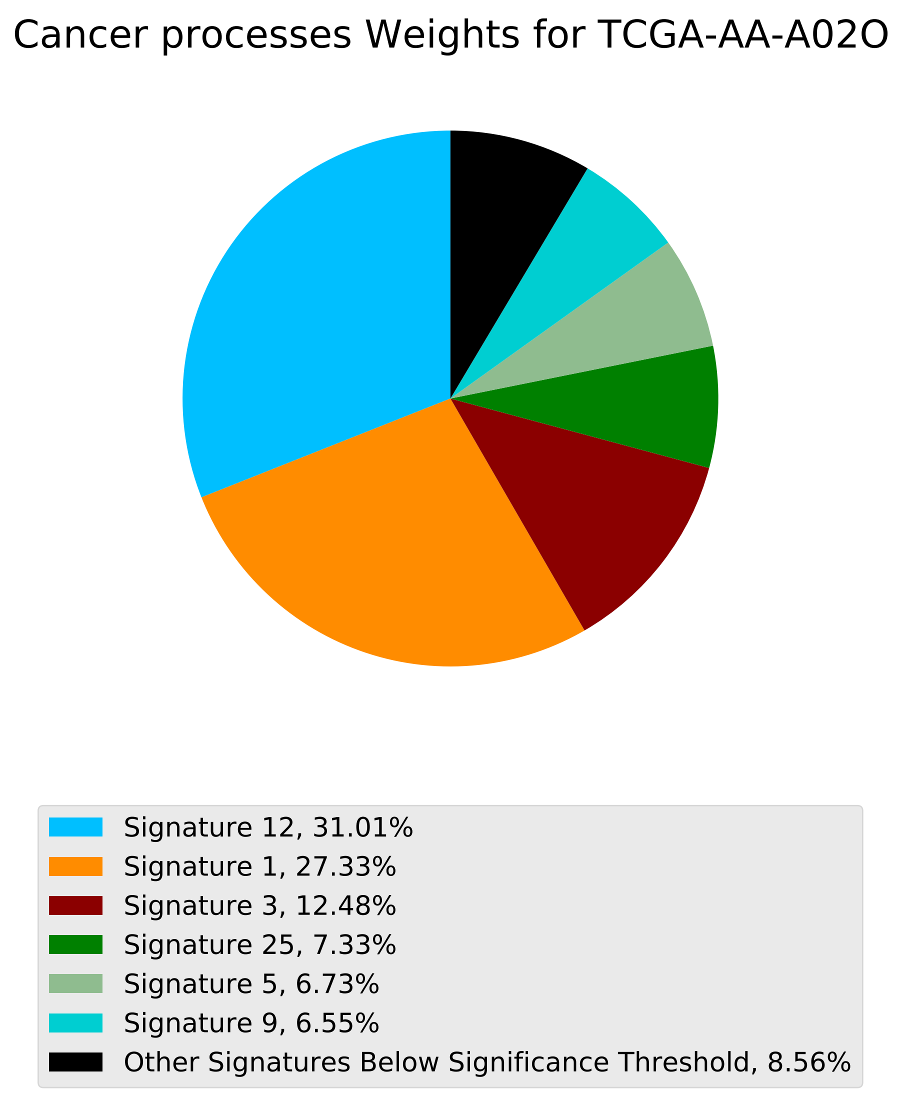

Cancer processes Weights for TCGA-AA-A02O
Signature 12, 31.01%
Signature 1, 27.33%
Signature 3, 12.48%
Signature 25, 7.33%
Signature 5, 6.73%
Signature 9, 6.55%
Other Signatures Below Significance Threshold, 8.56%

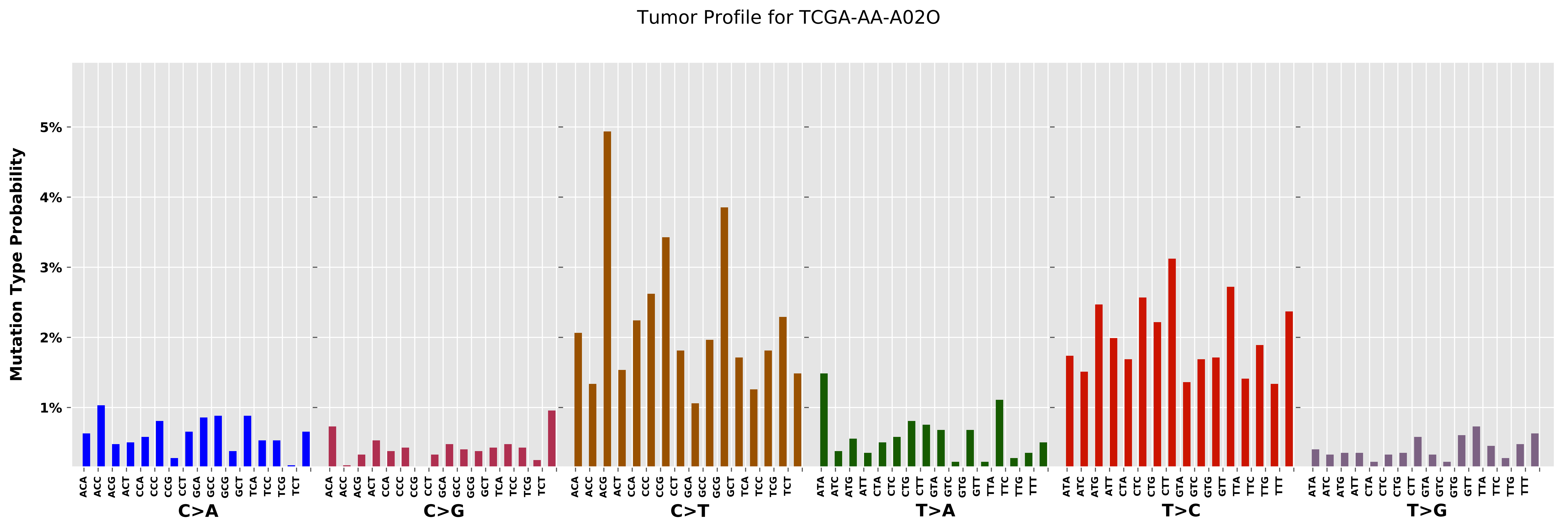

Tumor Profile for TCGA-AA-A02O
Mutation Type Probability
5%
4%
3%
2%
1%
ACA
ACC
ACG
ACT
CCA
CCC
CCG
CCT
GCA
GCC
GCG
GCT
TCA
TCC
TCG
TCT
C>A
C>G
C>T
ATA
ATC
ATG
ATT
CTA
CTC
CTG
CTT
GTA
GTC
GTG
GTT
TTA
TTC
TTG
TTT
T>A
T>C
T>G

Cancer processes Weights for TCGA-AG-3885

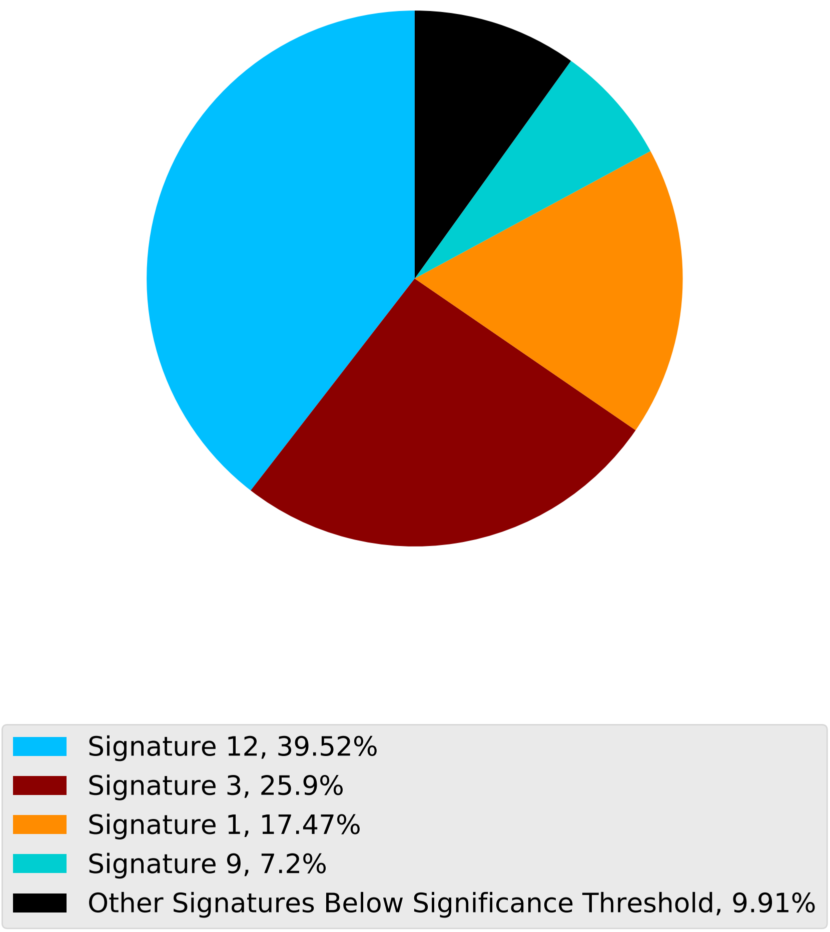

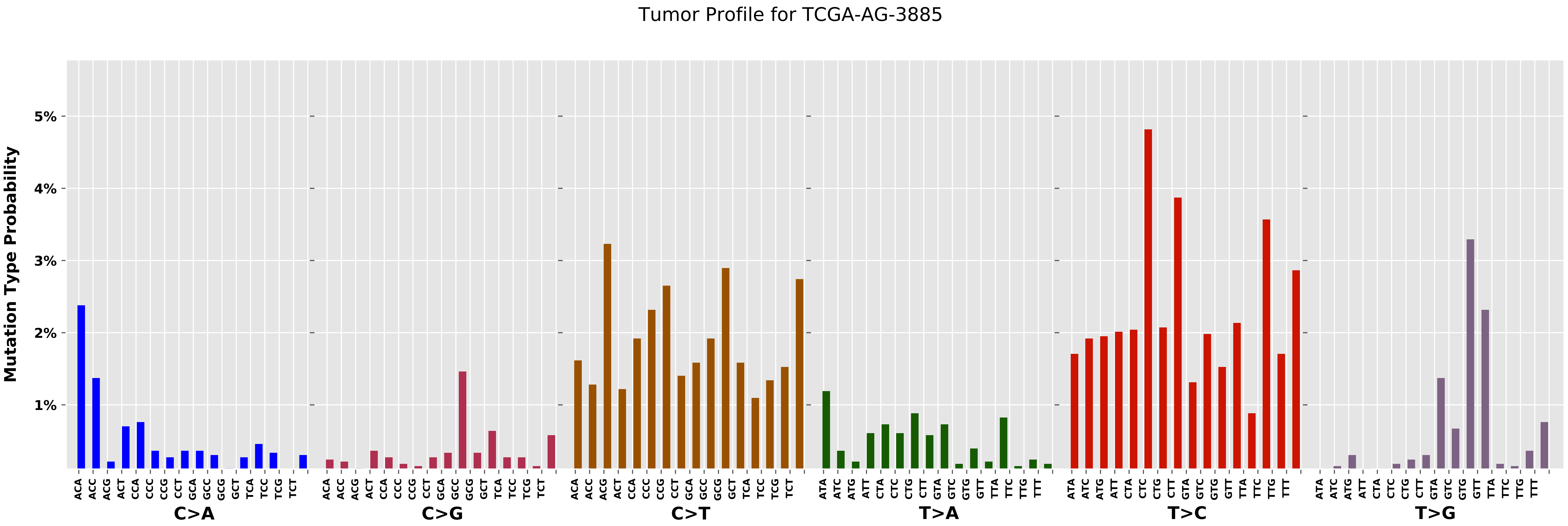
Tumor Profile for TCGA-AG-3885
Mutation Type Probability
5%
4%
3%
2%
1%
C>A
C>G
C>T
T>A
T>C
T>G

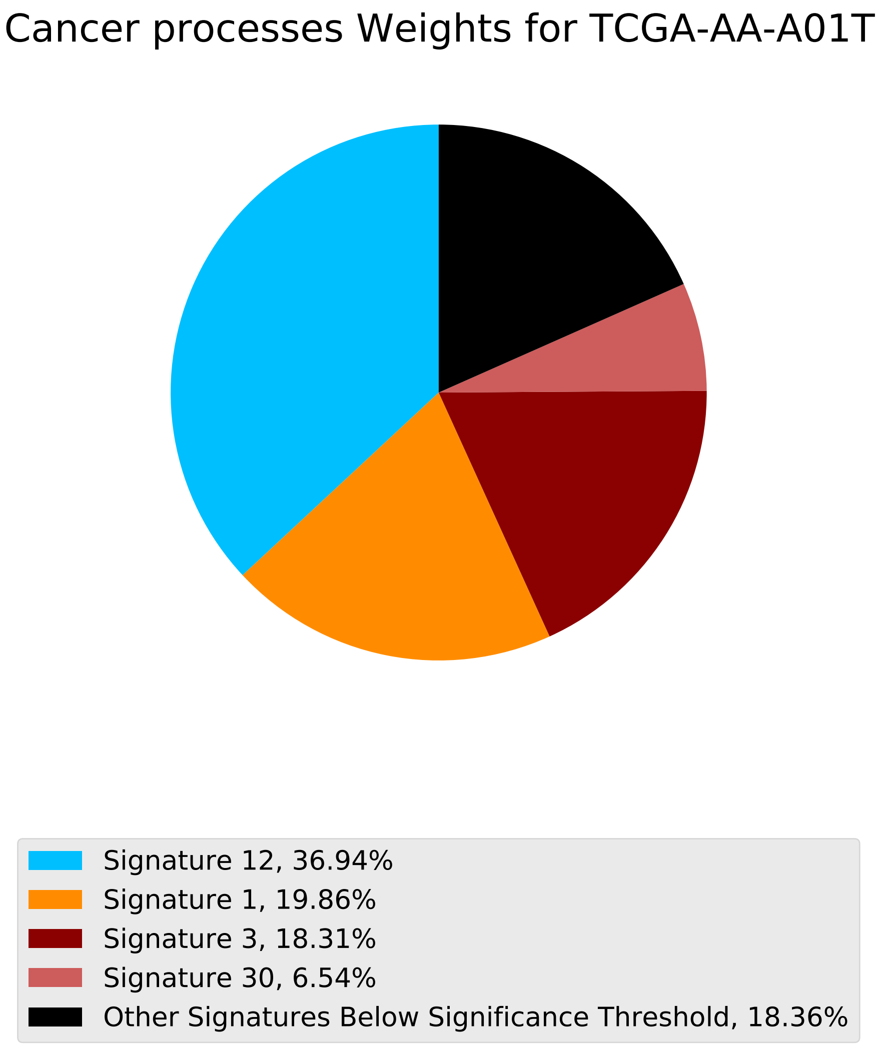
Cancer processes Weights for TCGA-AA-A01T
Signature 12, 36.94%
Signature 1, 19.86%
Signature 3, 18.31%
Signature 30, 6.54%
Other Signatures Below Significance Threshold, 18.36%

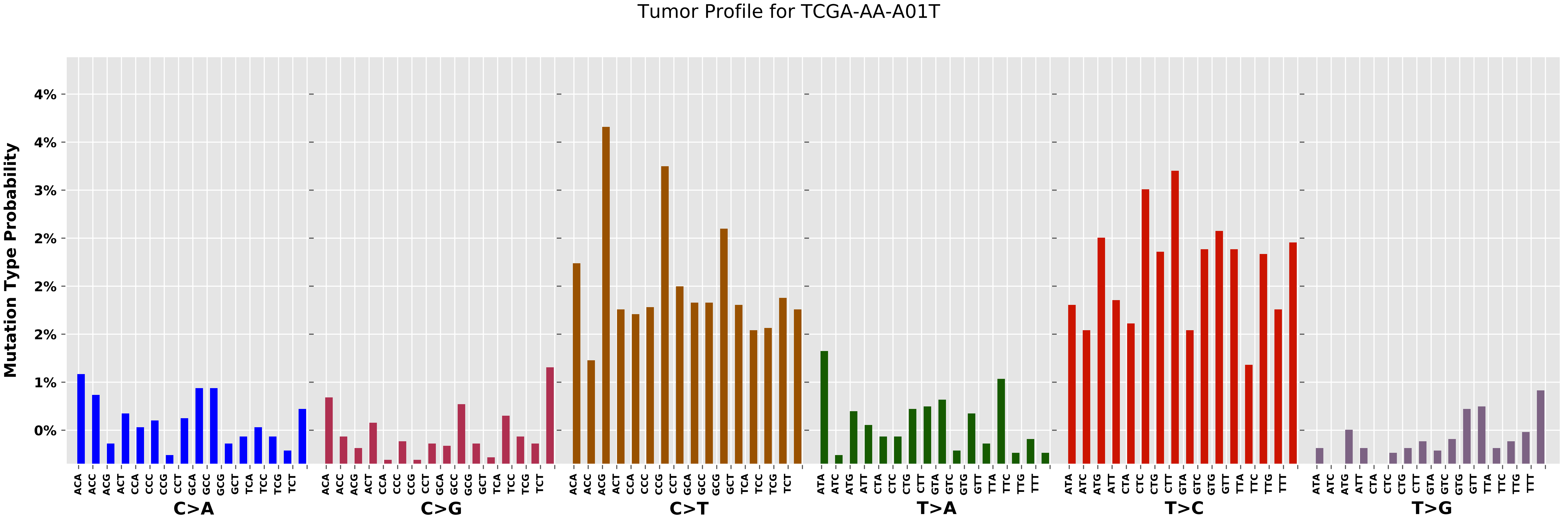

Tumor Profile for TCGA-AA-A01T
Mutation Type Probability
4%
4%
3%
2%
2%
2%
1%
0%
C>A
C>G
C>T
T>A
T>C
T>G
ACA ACC ACG ACT CCA CCC CCG CCT GCA GCC GCG GCT TCA TCC TCG TCT
ATA ATC ATG ATT CTA CTC CTG CTT GTA GTC GTG GTT TTA TTC TTG TTT

Cancer processes Weights for TCGA-A8-A07I

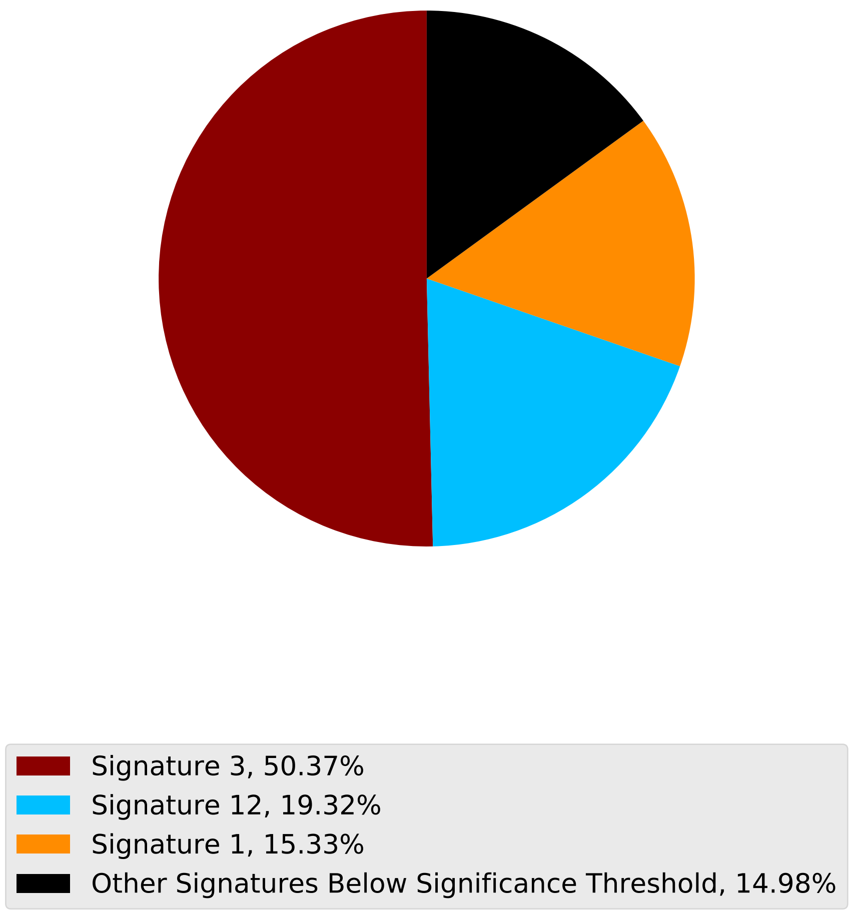

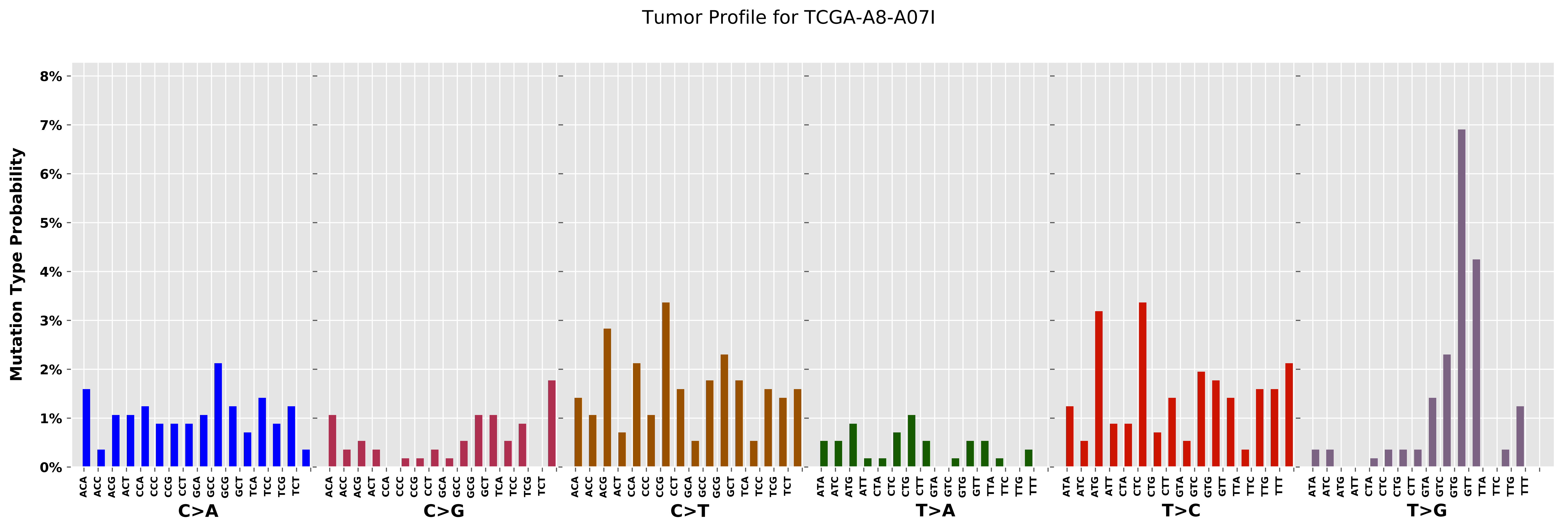

Tumor Profile for TCGA-A8-A07I
Mutation Type Probability
0%
1%
2%
3%
4%
5%
6%
7%
8%
C>A
C>G
C>T
T>A
T>C
T>G
ACA ACC ACG ACT CCA CCC CCG CCT GCA GCC GCG GCT TCA TCC TCG TCT
ATA ATC ATG ATT CTA CTC CTG CTT GTA GTC GTG GTT TTA TTC TTG TTT

Cancer processes Weights for TCGA-A6-2681

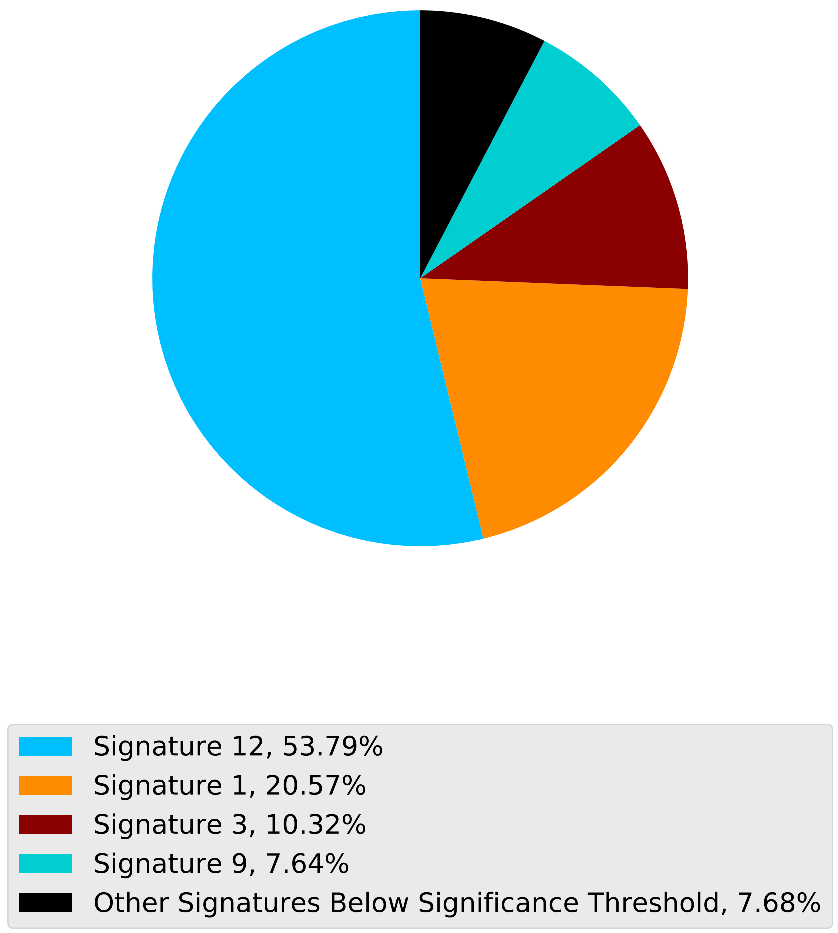

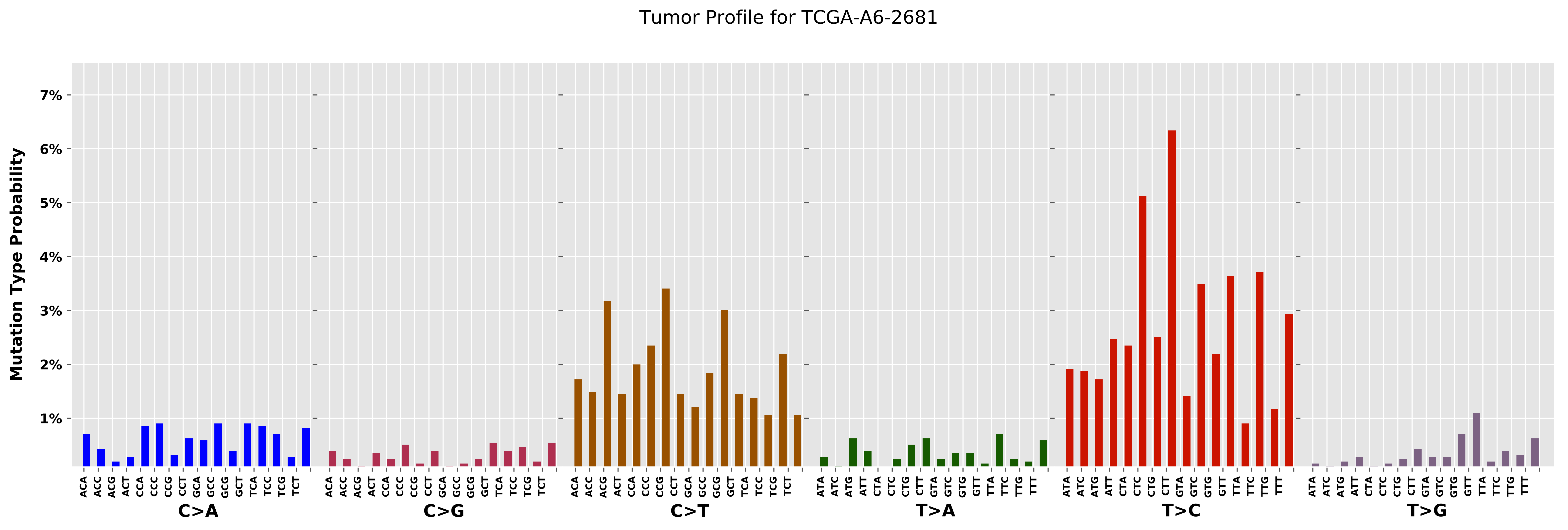

Tumor Profile for TCGA-A6-2681
Mutation Type Probability
7%
6%
5%
4%
3%
2%
1%
ACA
ACC
ACG
ACT
CCA
CCC
CCG
CCT
GCA
GCC
GCG
GCT
TCA
TCC
TCG
TCT
C>A
C>G
C>T
ATA
ATC
ATG
ATT
CTA
CTC
CTG
CTT
GTA
GTC
GTG
GTT
TTA
TTC
TTG
TTT
T>A
T>C
T>G

Cancer processes Weights for TCGA-AO-A124

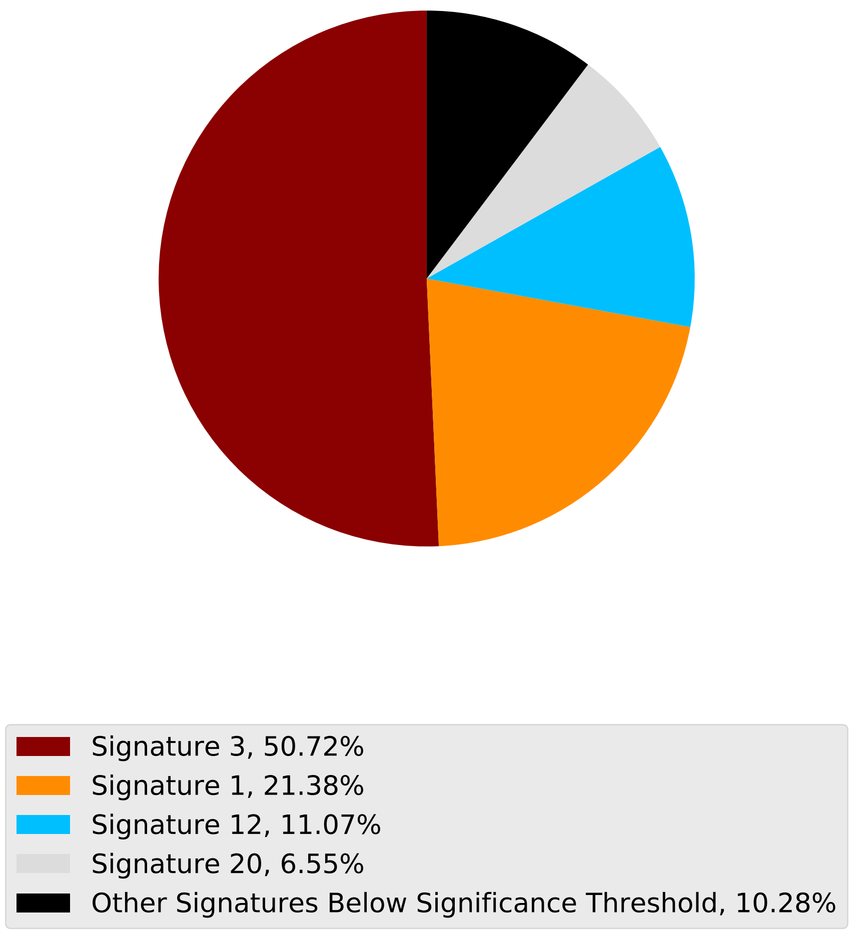

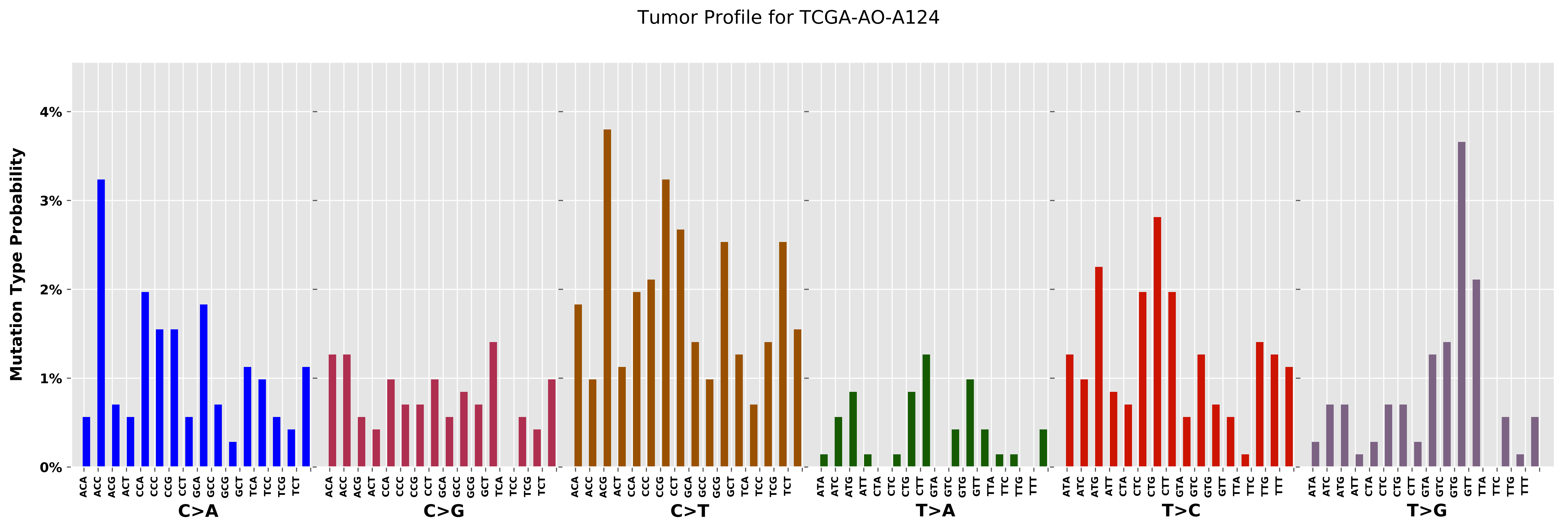
Tumor Profile for TCGA-AO-A124
Mutation Type Probability
0%
1%
2%
3%
4%
C>A
C>G
C>T
T>A
T>C
T>G

Cancer processes Weights for TCGA-A2-A259

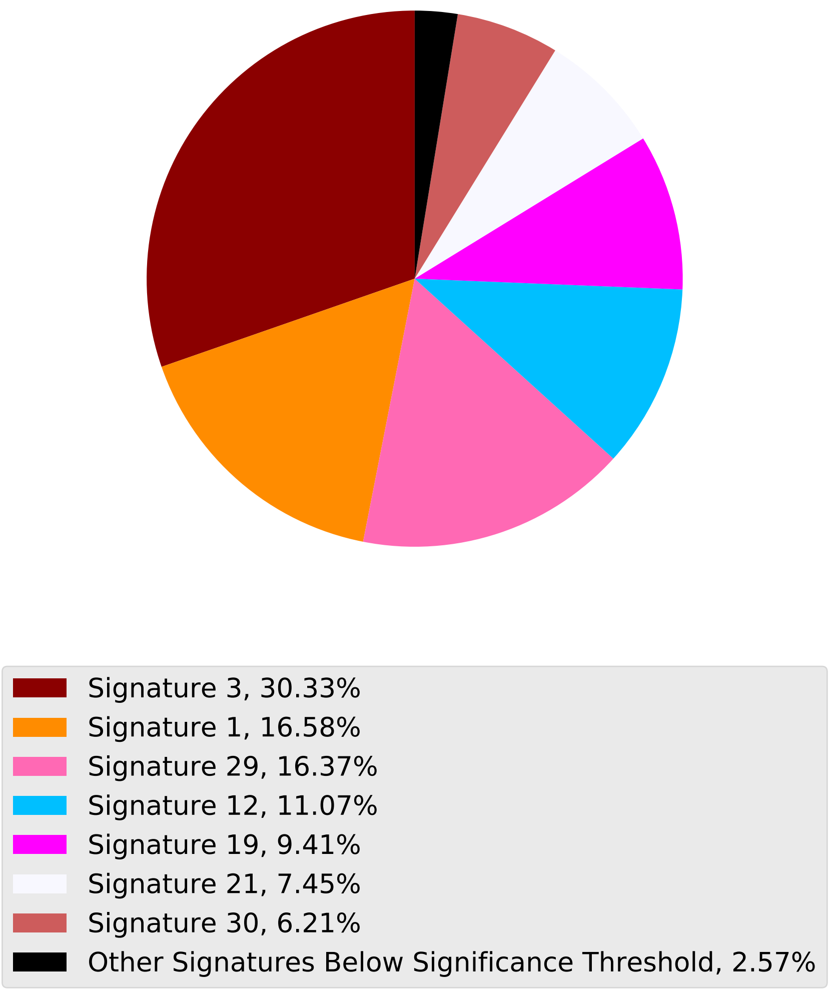

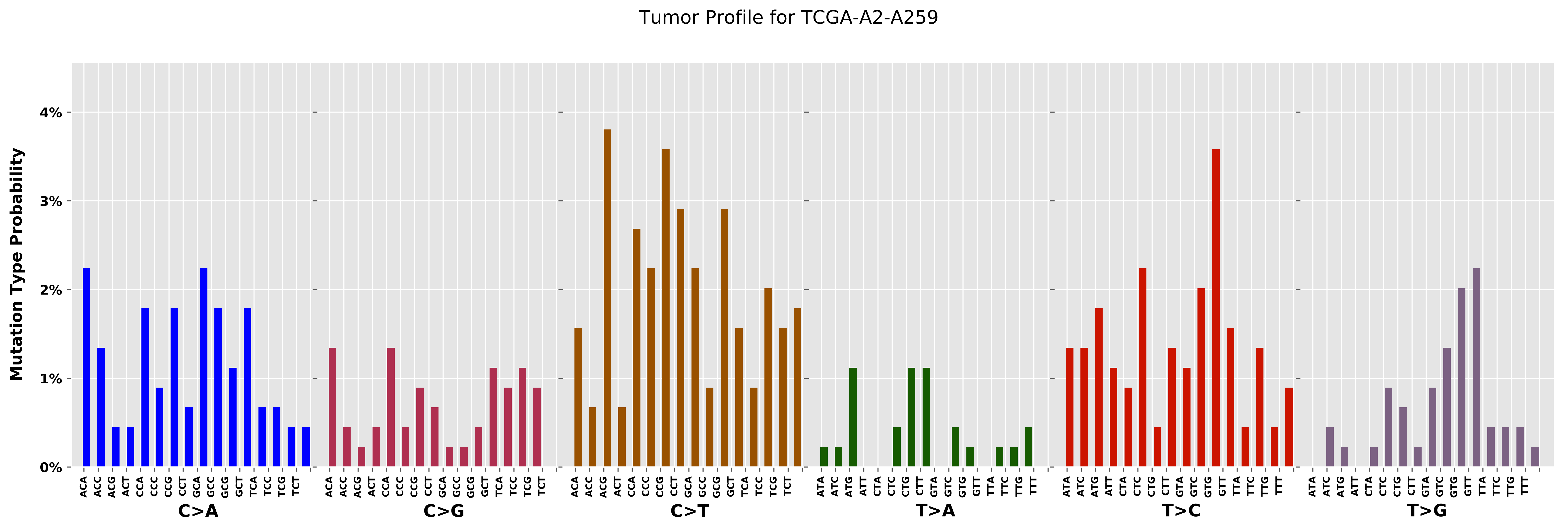
Tumor Profile for TCGA-A2-A259
Mutation Type Probability
0%
1%
2%
3%
4%
C>A
C>G
C>T
T>A
T>C
T>G

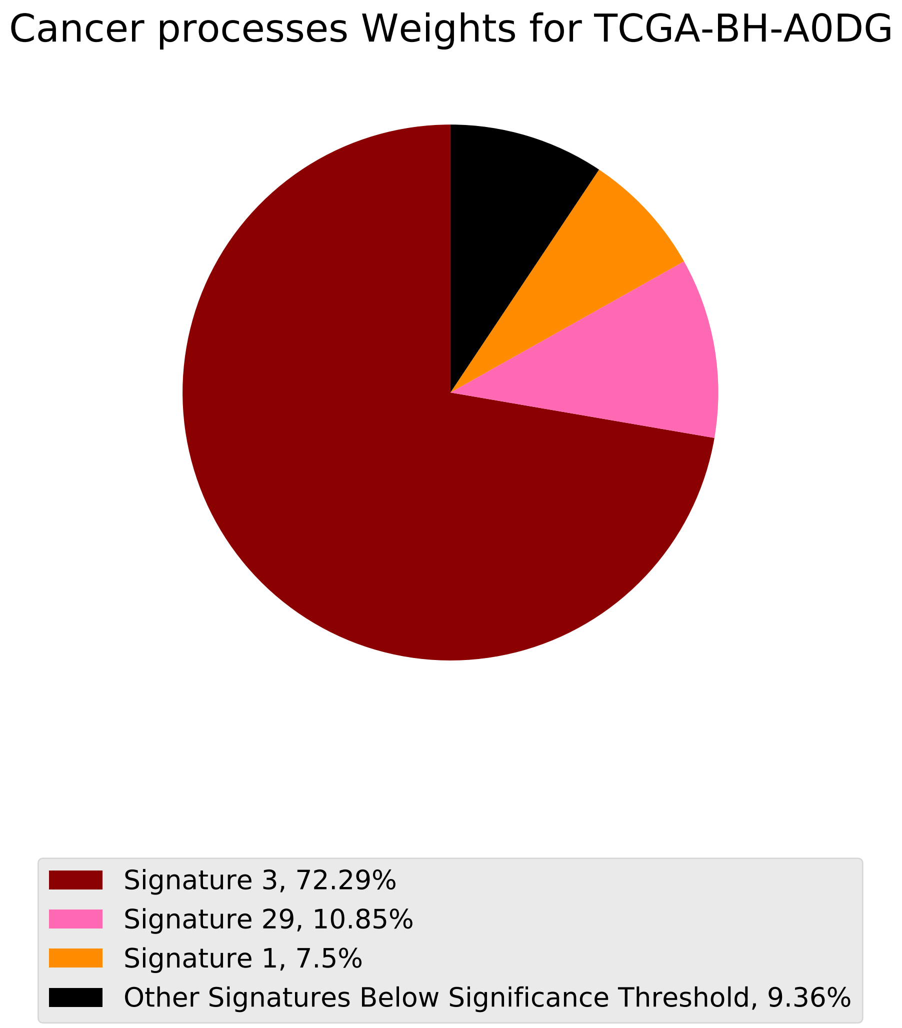
Cancer processes Weights for TCGA-BH-A0DG
Signature 3, 72.29%
Signature 29, 10.85%
Signature 1, 7.5%
Other Signatures Below Significance Threshold, 9.36%

Tumor Profile for TCGA-BH-A0DG

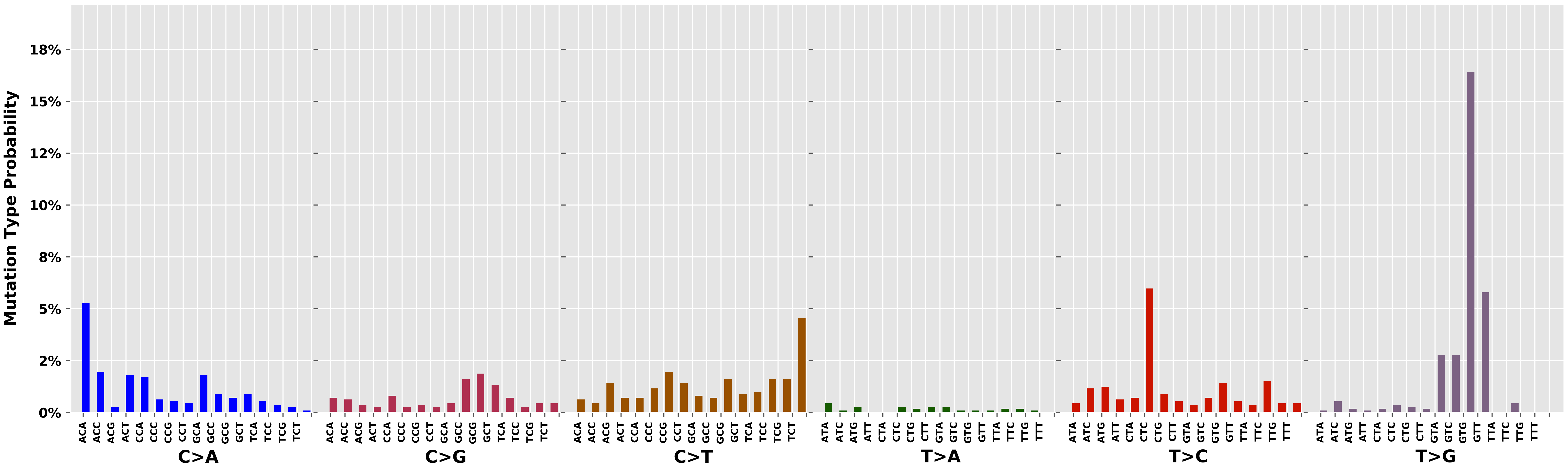

Cancer processes Weights for TCGA-A2-A3KC

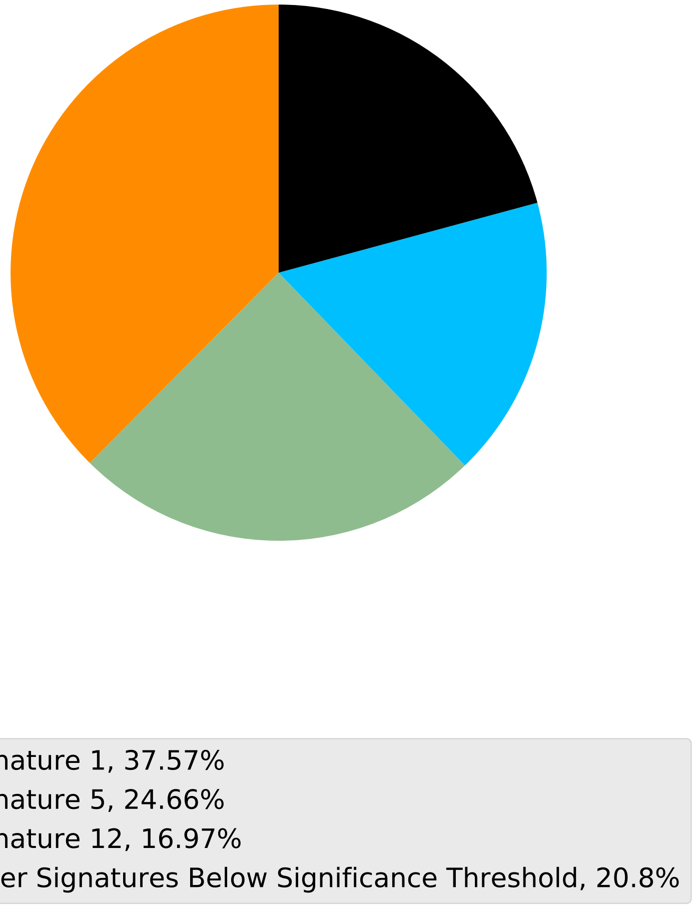

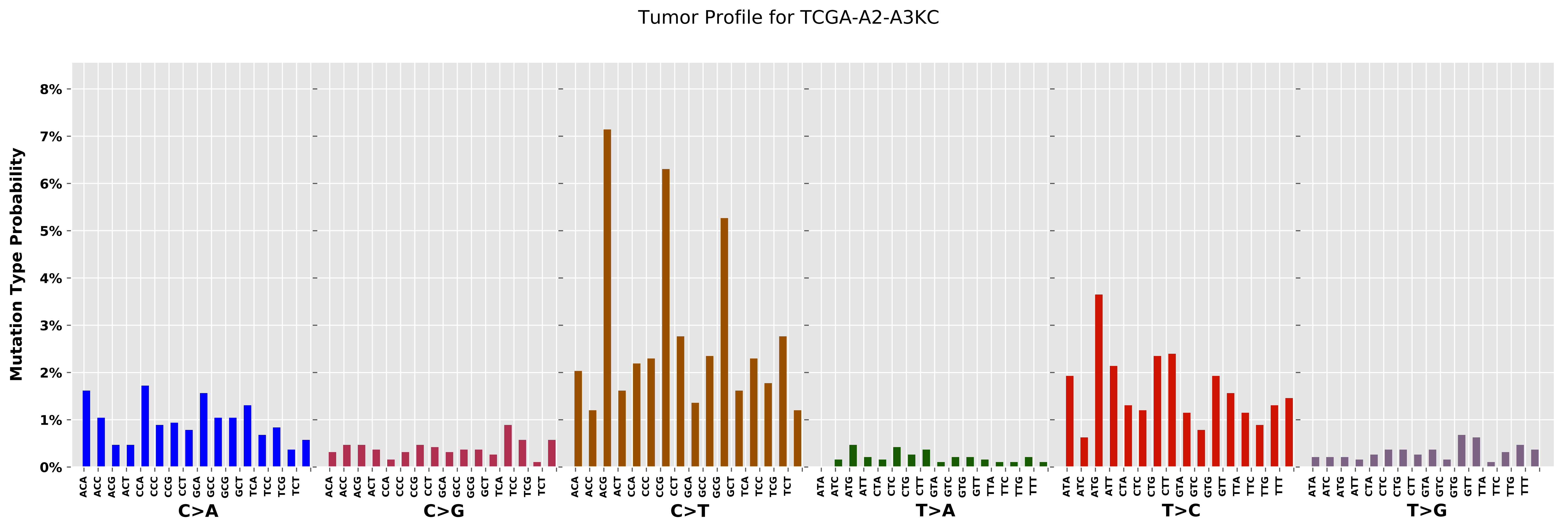
Tumor Profile for TCGA-A2-A3KC
Mutation Type Probability
0%
1%
2%
3%
4%
5%
6%
7%
8%
C>A
C>G
C>T
T>A
T>C
T>G
ACA
ACC
ACG
ACT
CCA
CCC
CCG
CCT
GCA
GCC
GCG
GCT
TCA
TCC
TCG
TCT
ATA
ATC
ATG
ATT
CTA
CTC
CTG
CTT
GTA
GTC
GTG
GTT
TTA
TTC
TTG
TTT

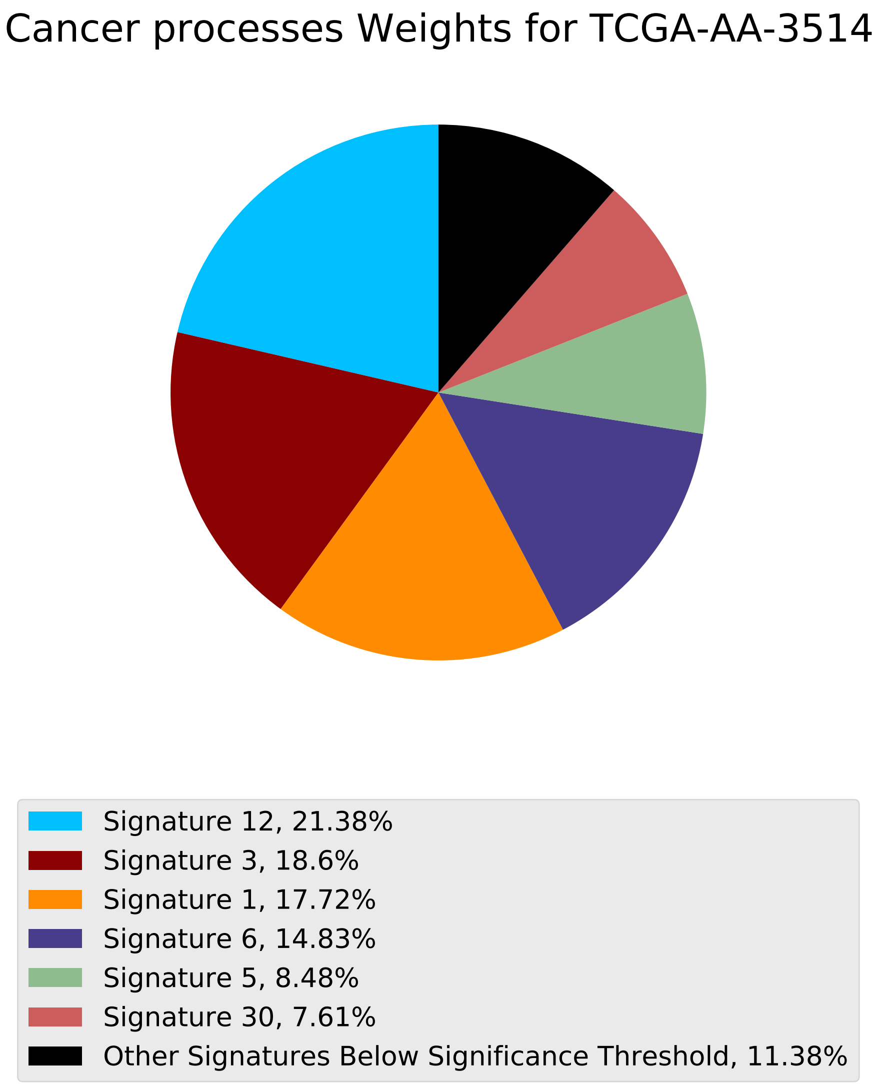
Cancer processes Weights for TCGA-AA-3514
Signature 12, 21.38%
Signature 3, 18.6%
Signature 1, 17.72%
Signature 6, 14.83%
Signature 5, 8.48%
Signature 30, 7.61%
Other Signatures Below Significance Threshold, 11.38%

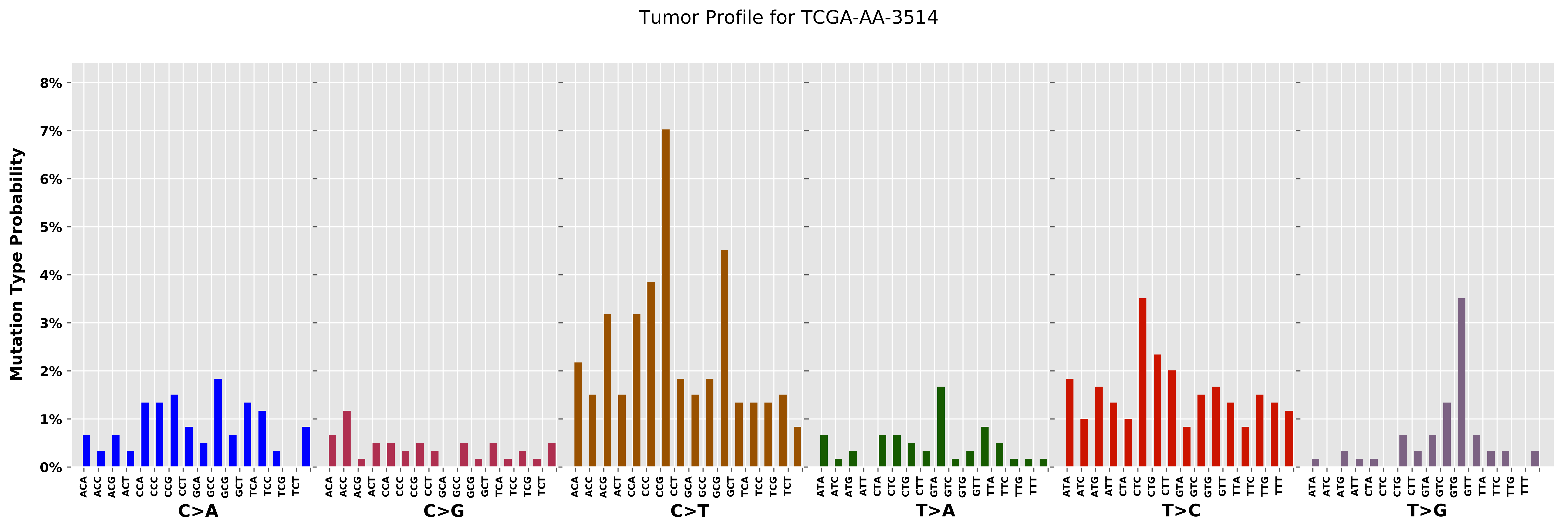

Tumor Profile for TCGA-AA-3514
Mutation Type Probability
0%
1%
2%
3%
4%
5%
6%
7%
8%
C>A
C>G
C>T
T>A
T>C
T>G
ACA ACC ACG ACT CCA CCC CCG CCT GCA GCC GCG GCT TCA TCC TCG TCT
ATA ATC ATG ATT CTA CTC CTG CTT GTA GTC GTG GTT TTA TTC TTG TTT

Cancer processes Weights for TCGA-AN-A0XR

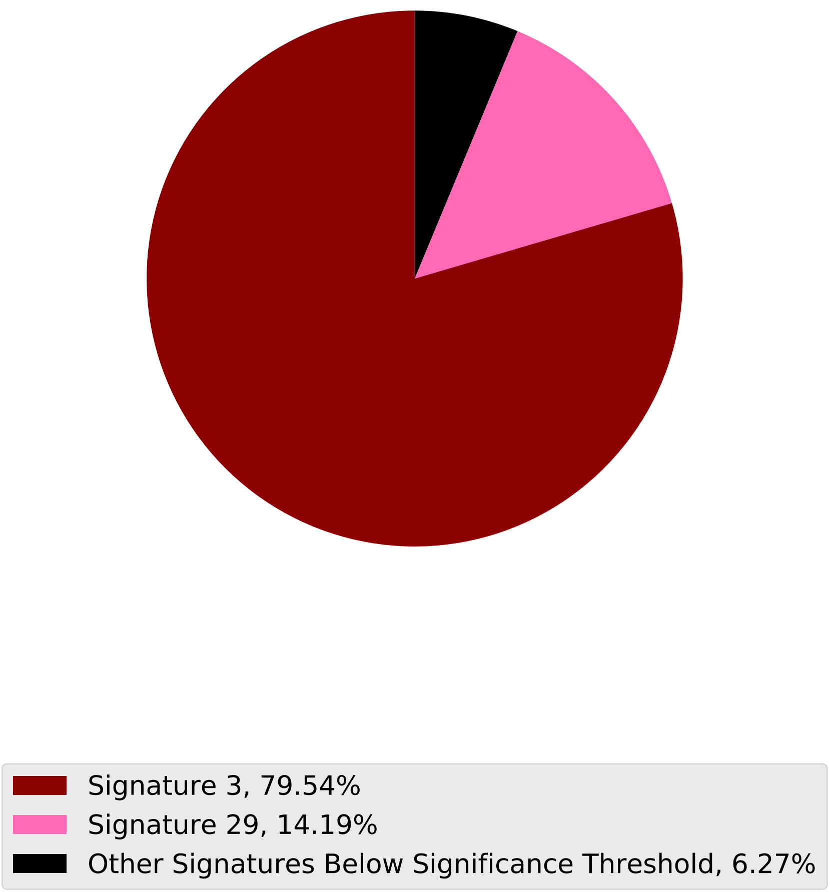

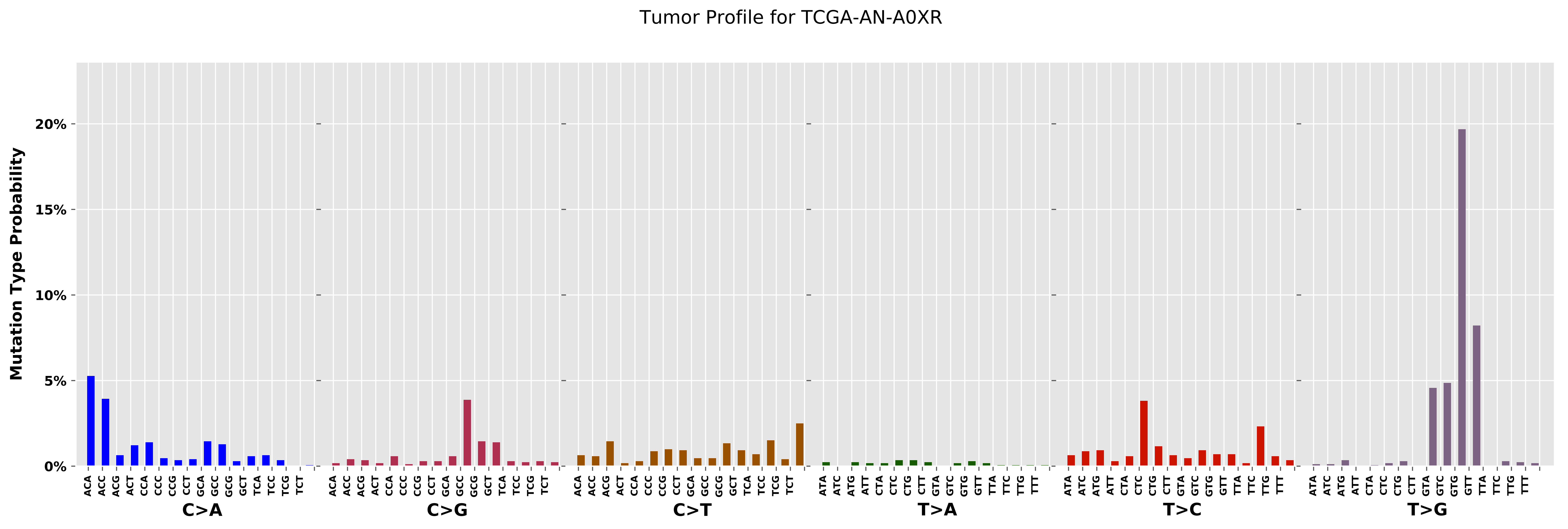
Tumor Profile for TCGA-AN-A0XR
Mutation Type Probability
20%
15%
10%
5%
0%
C>A
C>G
C>T
T>A
T>C
T>G

Cancer processes Weights for TCGA-BH-A0EA

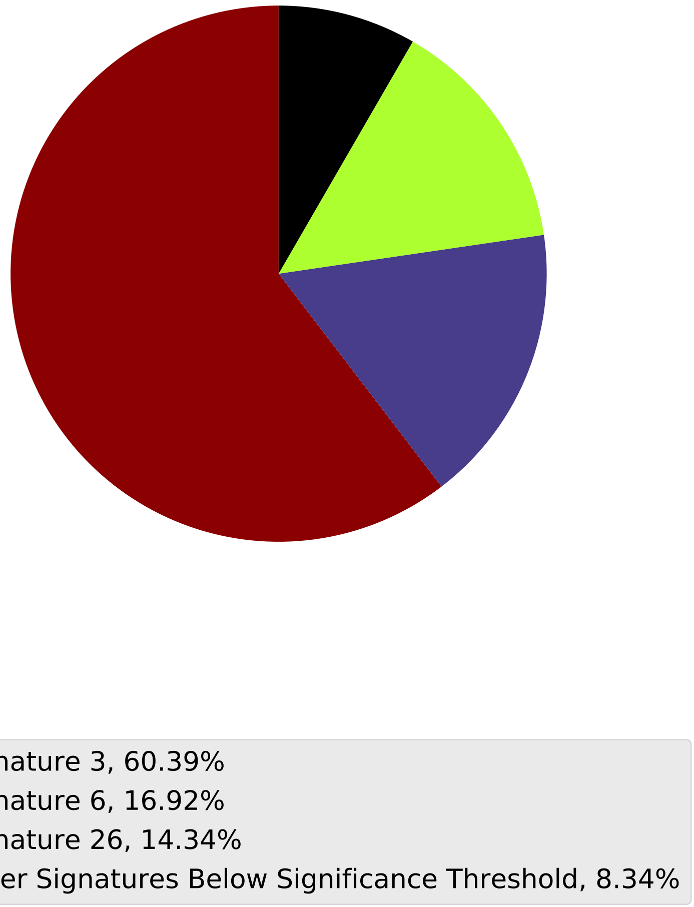

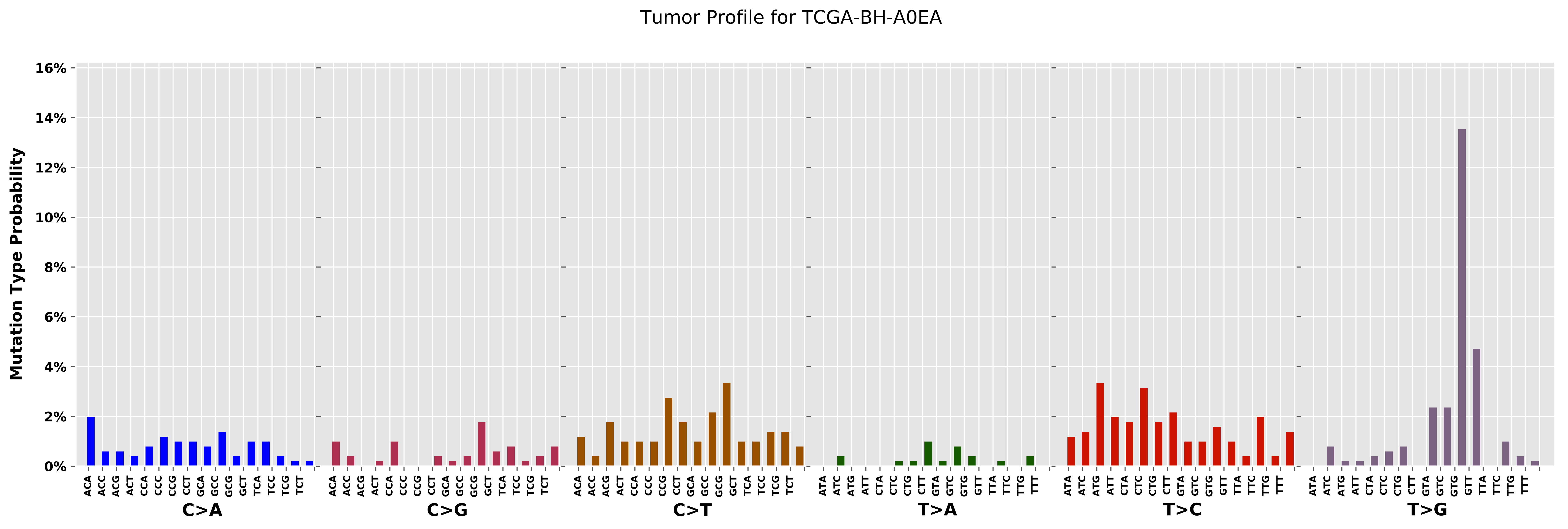

Tumor Profile for TCGA-BH-A0EA
Mutation Type Probability
16%
14%
12%
10%
8%
6%
4%
2%
0%
C>A
C>G
C>T
T>A
T>C
T>G

Cancer processes Weights for TCGA-AR-A256

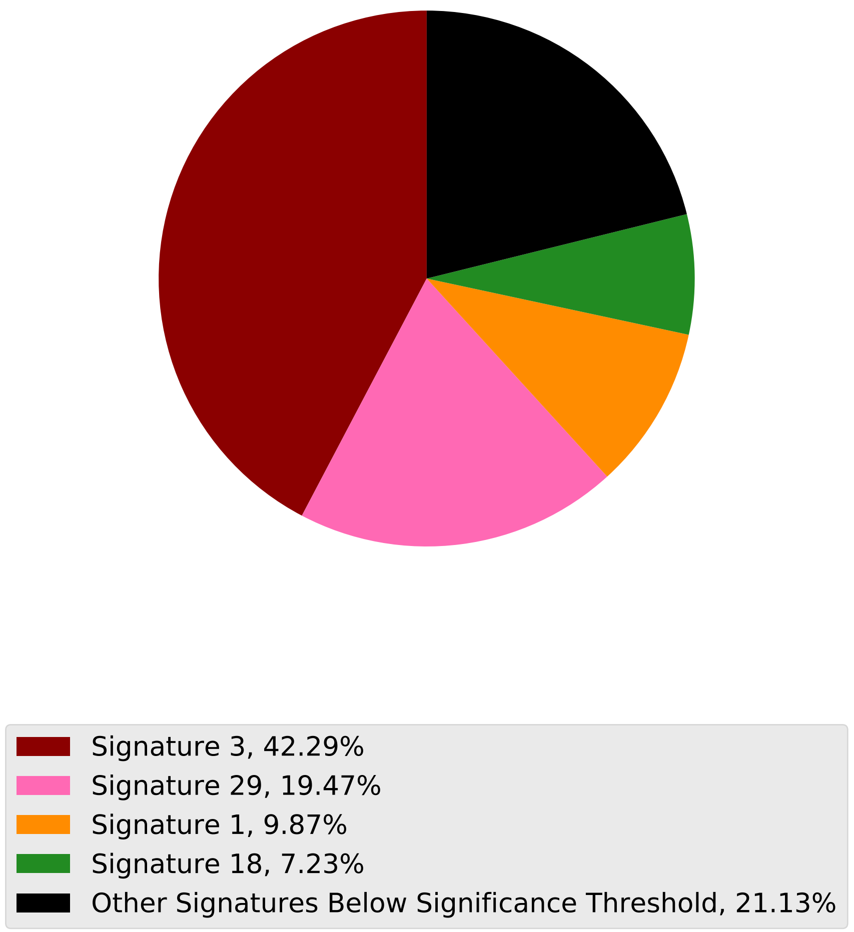

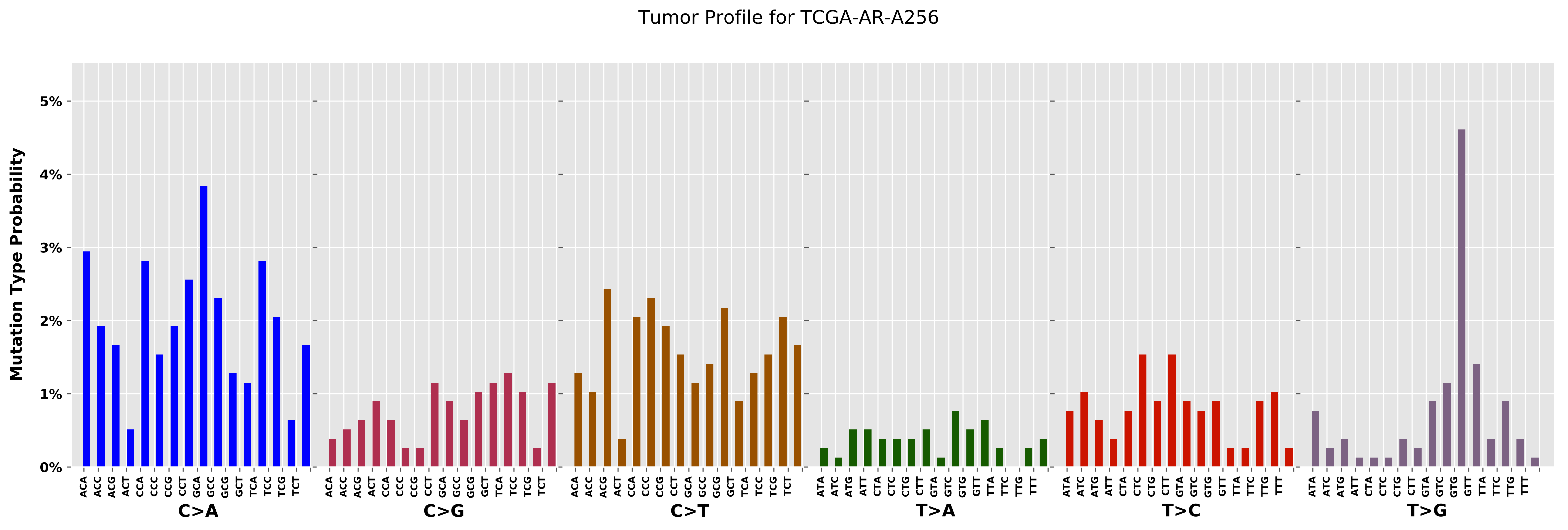

Tumor Profile for TCGA-AR-A256
Mutation Type Probability
0%
1%
2%
3%
4%
5%
C>A
C>G
C>T
T>A
T>C
T>G
ACA ACC ACG ACT CCA CCC CCG CCT GCA GCC GCG GCT TCA TCC TCG TCT
ATA ATC ATG ATT CTA CTC CTG CTT GTA GTC GTG GTT TTA TTC TTG TTT

Cancer processes Weights for TCGA-A8-A09X

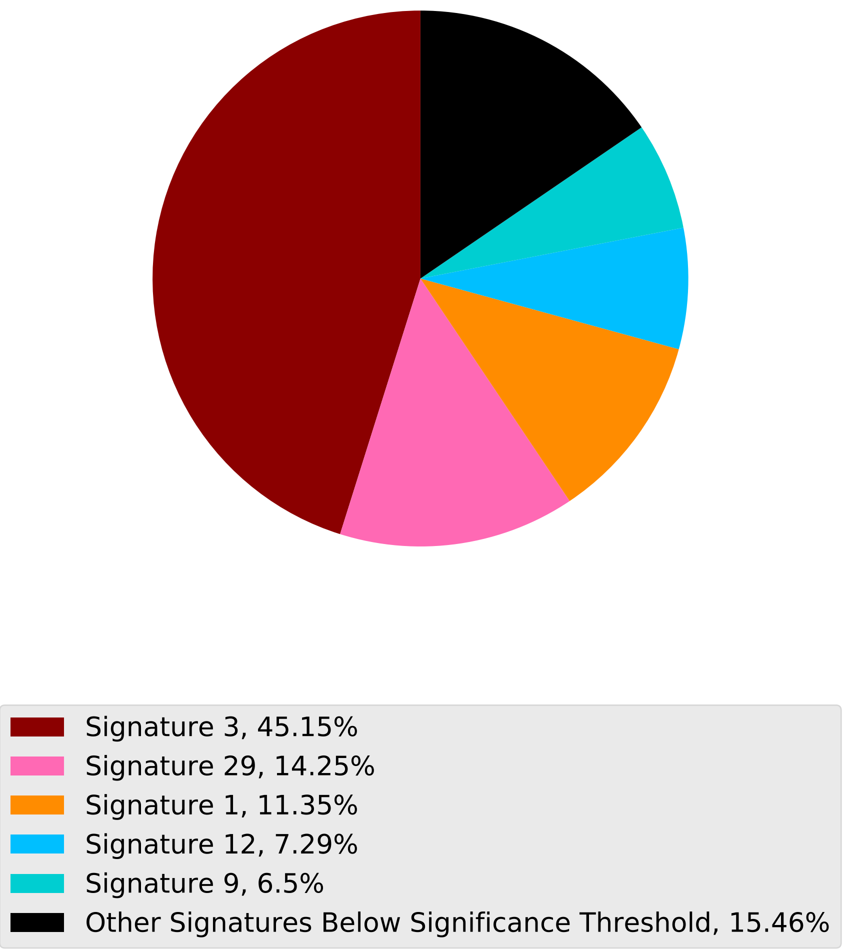

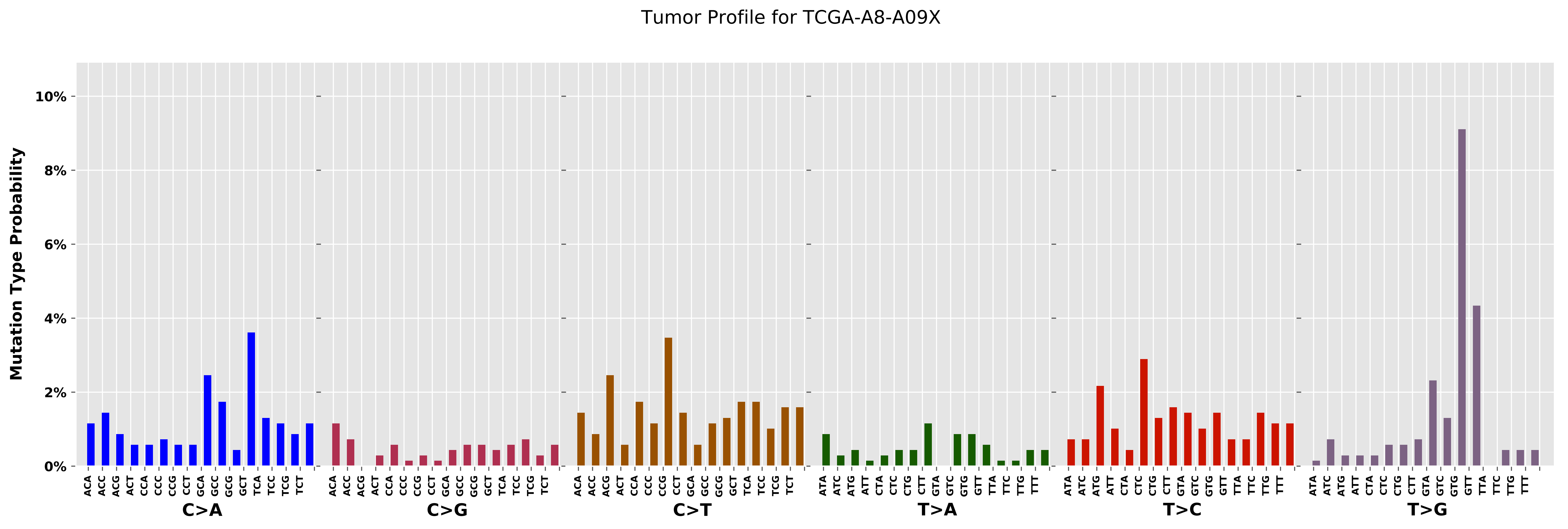
Tumor Profile for TCGA-A8-A09X
Mutation Type Probability
0%
2%
4%
6%
8%
10%
ACA ACC ACG ACT CCA CCC CCG CCT GCA GCC GCG GCT TCA TCC TCG TCT
C>A
ACA ACC ACG ACT CCA CCC CCG CCT GCA GCC GCG GCT TCA TCC TCG TCT
C>G
ACA ACC ACG ACT CCA CCC CCG CCT GCA GCC GCG GCT TCA TCC TCG TCT
C>T
ATA ATC ATG ATT CTA CTC CTG CTT GTA GTC GTG GTT TTA TTC TTG TTT
T>A
ATA ATC ATG ATT CTA CTC CTG CTT GTA GTC GTG GTT TTA TTC TTG TTT
T>C
ATA ATC ATG ATT CTA CTC CTG CTT GTA GTC GTG GTT TTA TTC TTG TTT
T>G

### Cancer processes Weights for TCGA-AA-A01V

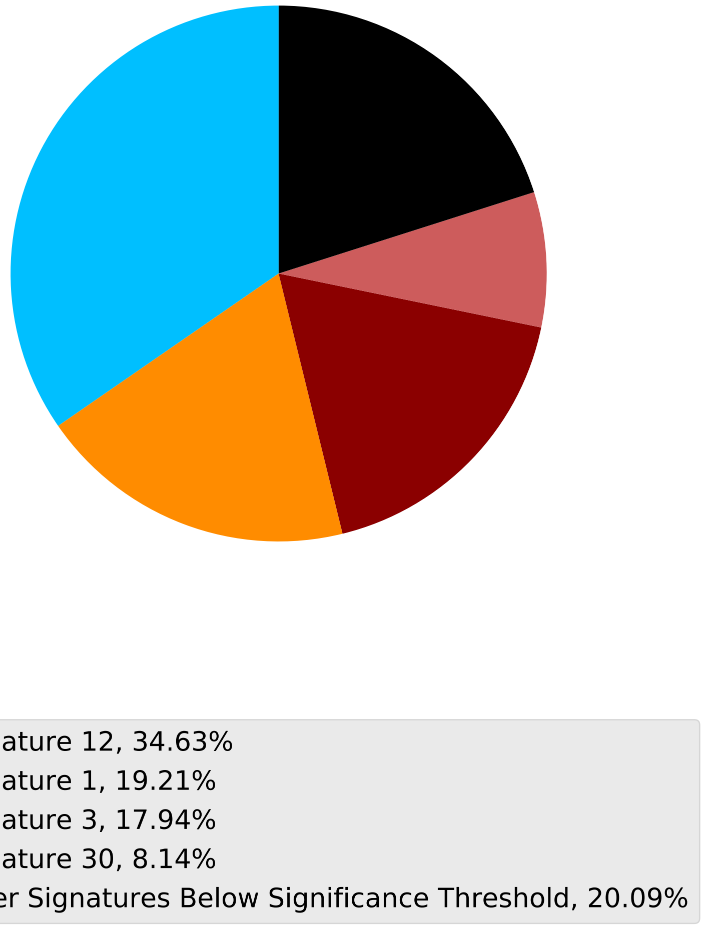

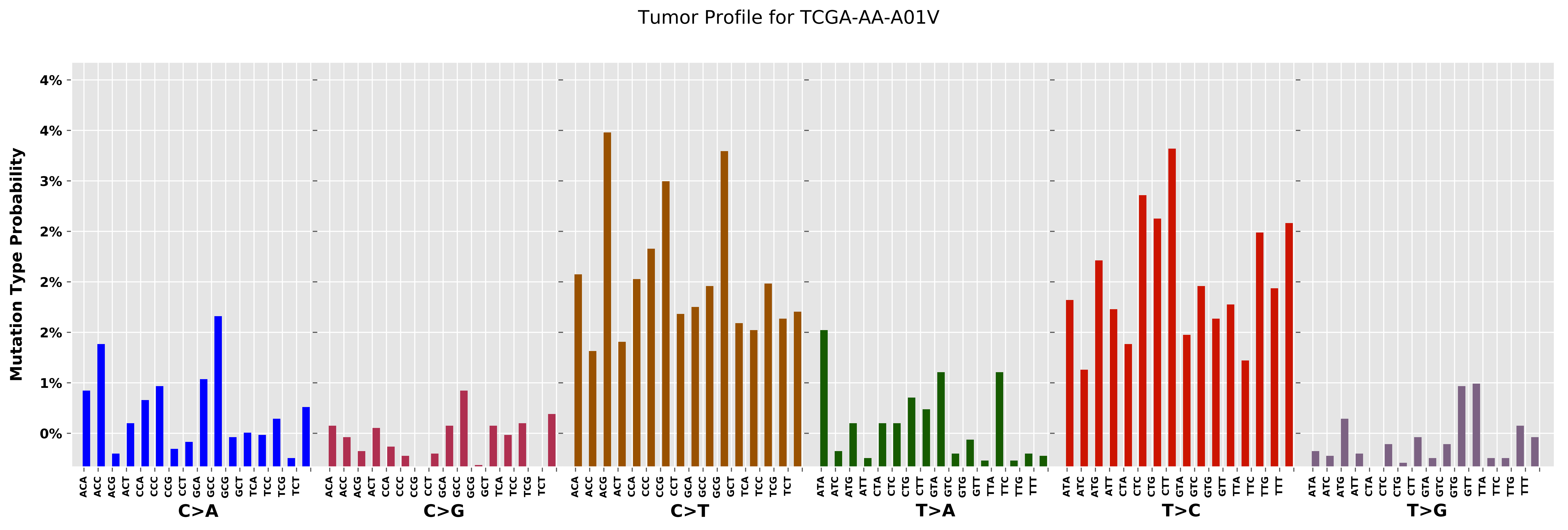
Tumor Profile for TCGA-AA-A01V
Mutation Type Probability
4%
4%
3%
2%
2%
2%
1%
0%
C>A
C>G
C>T
T>A
T>C
T>G
ACA ACC ACG ACT CCA CCC CCG CCT GCA GCC GCG GCT TCA TCC TCG TCT
ATA ATC ATG ATT CTA CTC CTG CTT GTA GTC GTG GTT TTA TTC TTG TTT

#### Cancer processes Weights for TCGA-A2-A04T

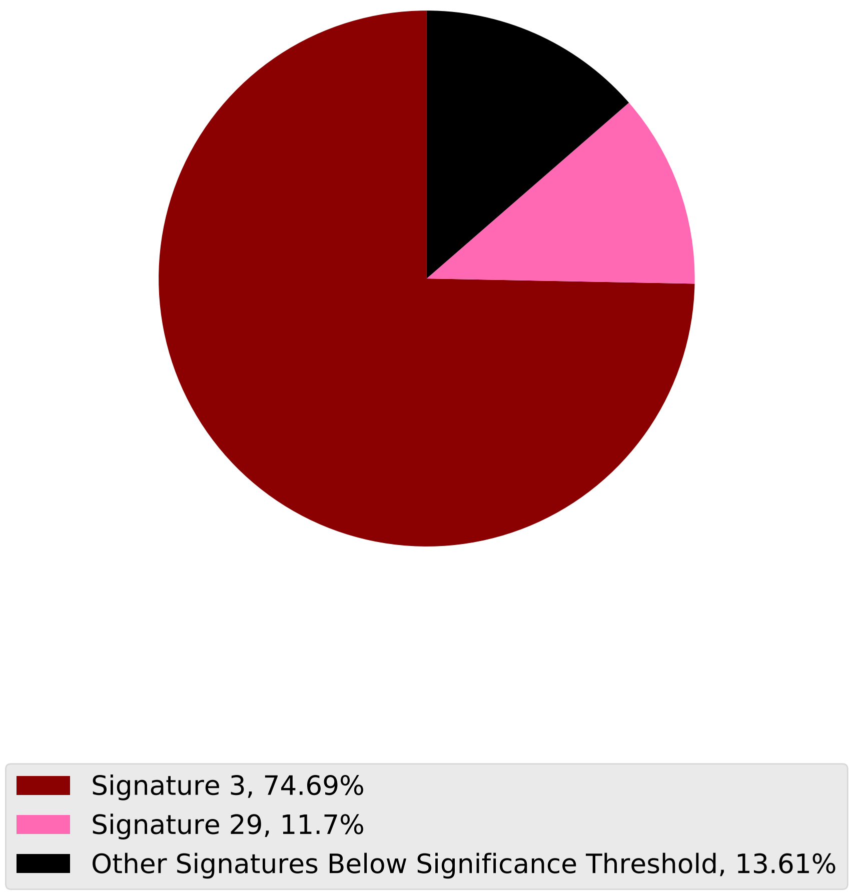

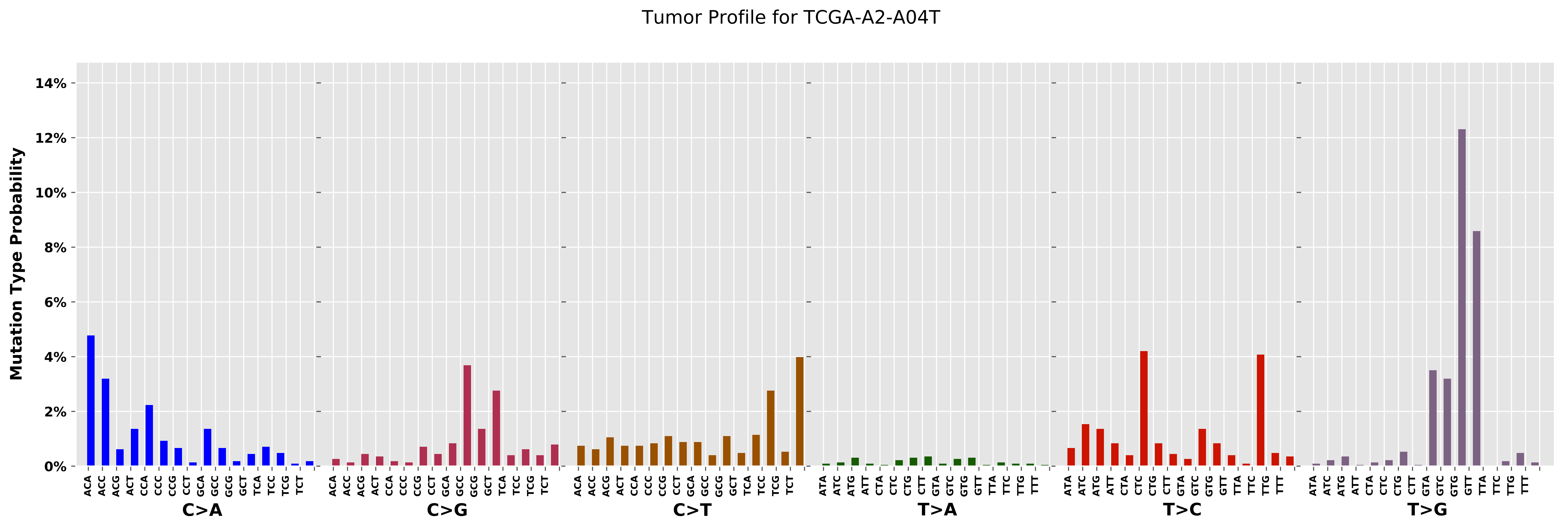

Tumor Profile for TCGA-A2-A04T
Mutation Type Probability
0%
2%
4%
6%
8%
10%
12%
14%
C>A
C>G
C>T
T>A
T>C
T>G
ACA ACC ACG ACT CCA CCC CCG CCT GCA GCC GCG GCT TCA TCC TCG TCT
ATA ATC ATG ATT CTA CTC CTG CTT GTA GTC GTG GTT TTA TTC TTG TTT

Cancer processes Weights for TCGA-BH-A0H6

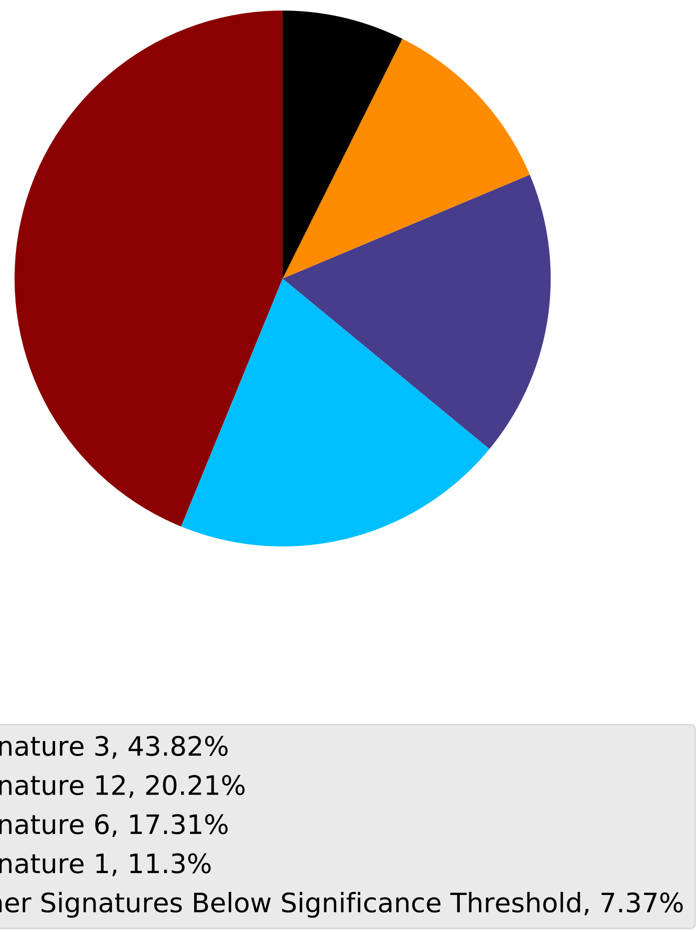

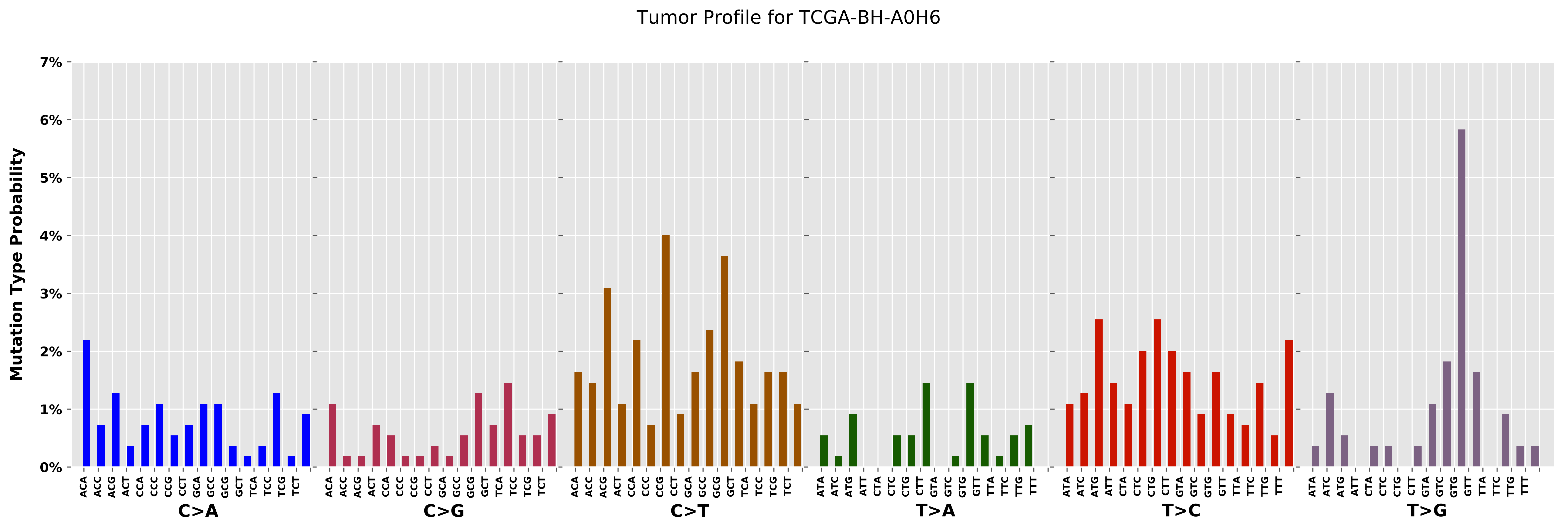
Tumor Profile for TCGA-BH-A0H6
Mutation Type Probability
0%
1%
2%
3%
4%
5%
6%
7%
C>A
C>G
C>T
T>A
T>C
T>G
ACA ACC ACG ACT CCA CCC CCG CCT GCA GCC GCG GCT TCA TCC TCG TCT
ATA ATC ATG ATT CTA CTC CTG CTT GTA GTC GTG GTT TTA TTC TTG TTT

Cancer processes Weights for TCGA-D8-A27F

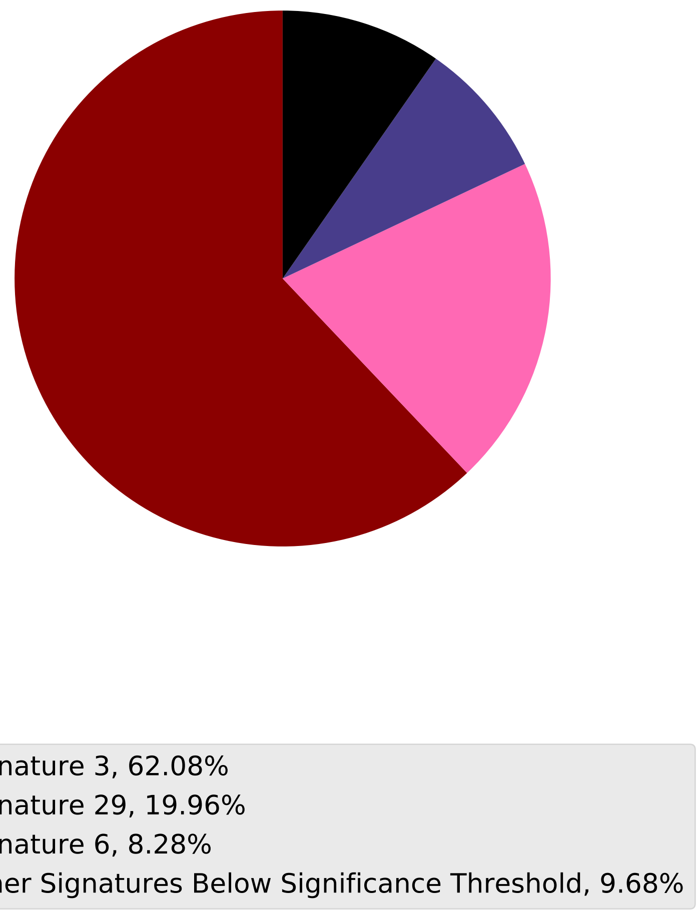

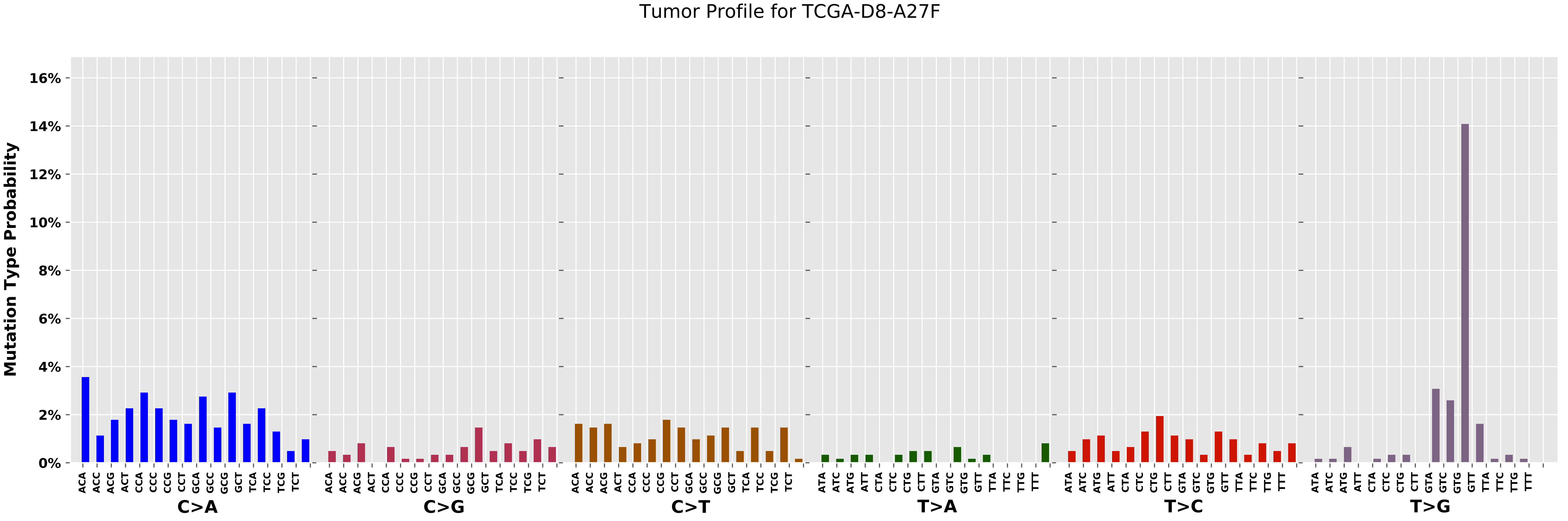
Tumor Profile for TCGA-D8-A27F
Mutation Type Probability
0%
2%
4%
6%
8%
10%
12%
14%
16%
C>A
C>G
C>T
T>A
T>C
T>G
ACA ACC ACG ACT CCA CCC CCG CCT GCA GCC GCG GCT TCA TCC TCG TCT
ATA ATC ATG ATT CTA CTC CTG CTT GTA GTC GTG GTT TTA TTC TTG TTT

Cancer processes Weights for TCGA-AG-3727

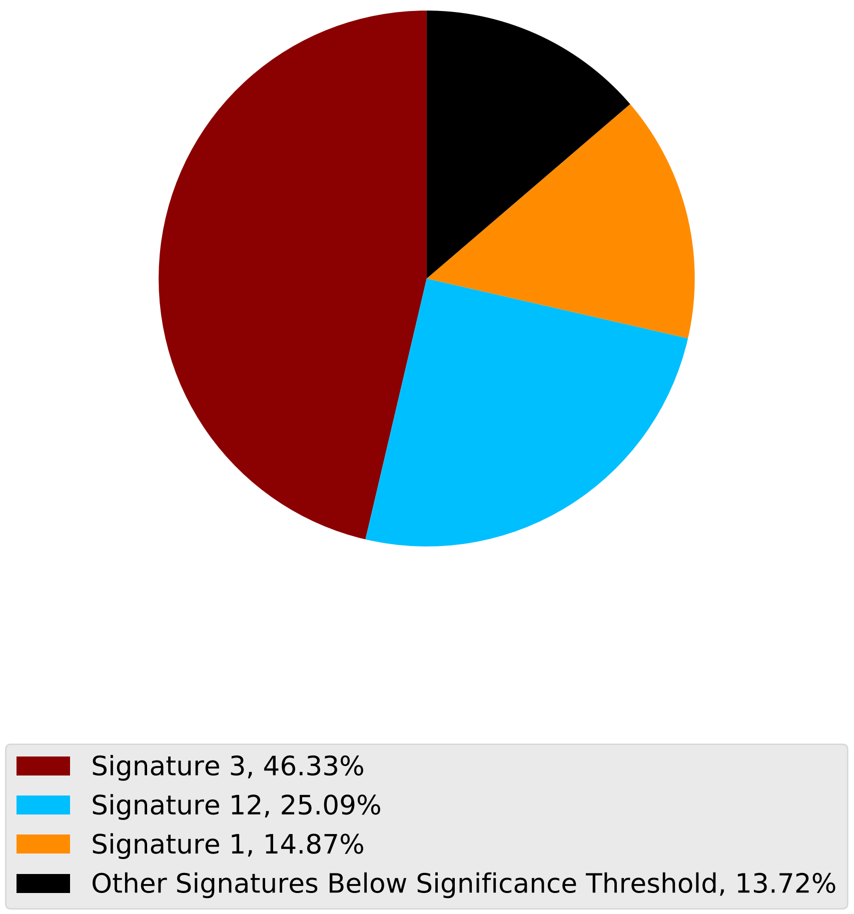

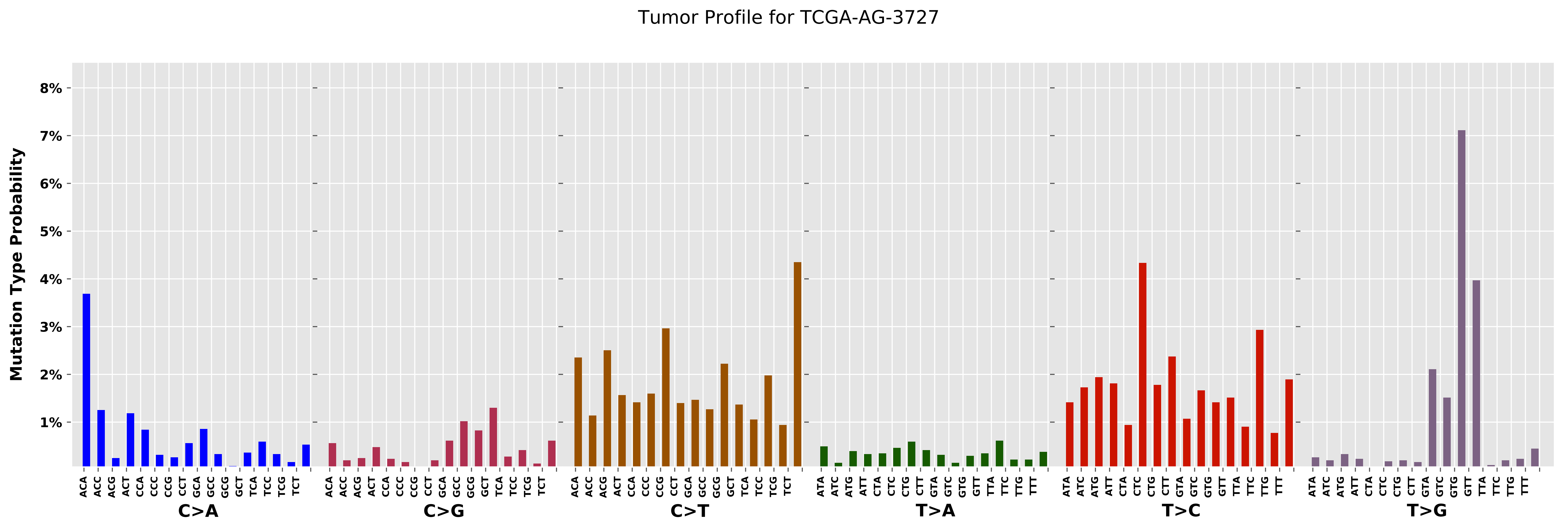

Tumor Profile for TCGA-AG-3727
Mutation Type Probability
8%
7%
6%
5%
4%
3%
2%
1%
C>A
C>G
C>T
T>A
T>C
T>G
ACA ACC ACG ACT CCA CCC CCG CCT GCA GCC GCG GCT TCA TCC TCG TCT
ATA ATC ATG ATT CTA CTC CTG CTT GTA GTC GTG GTT TTA TTC TTG TTT

Cancer processes Weights for TCGA-A8-A09I

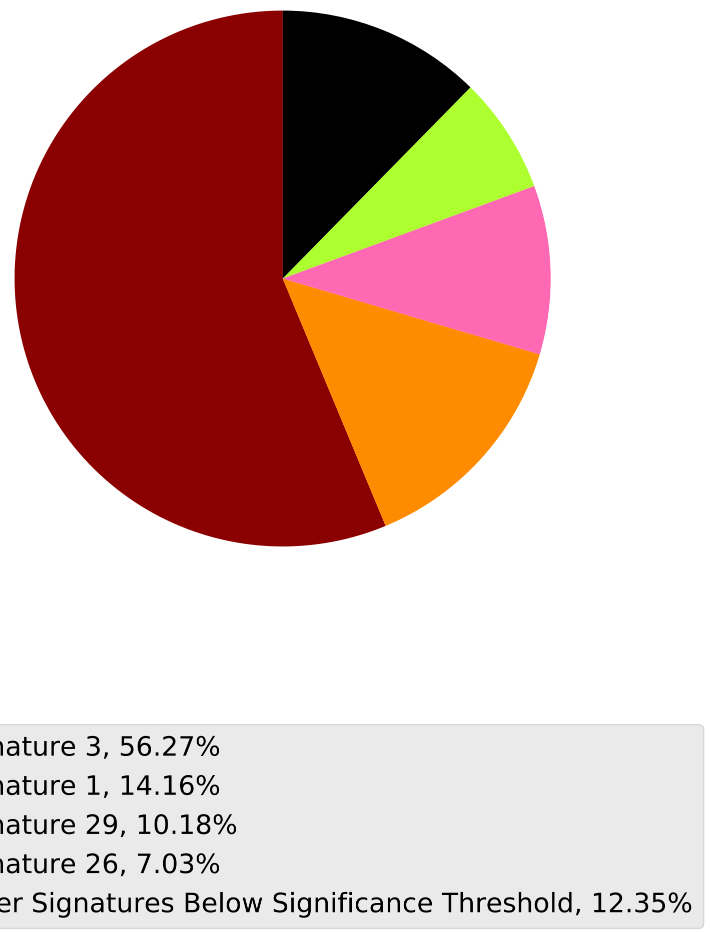

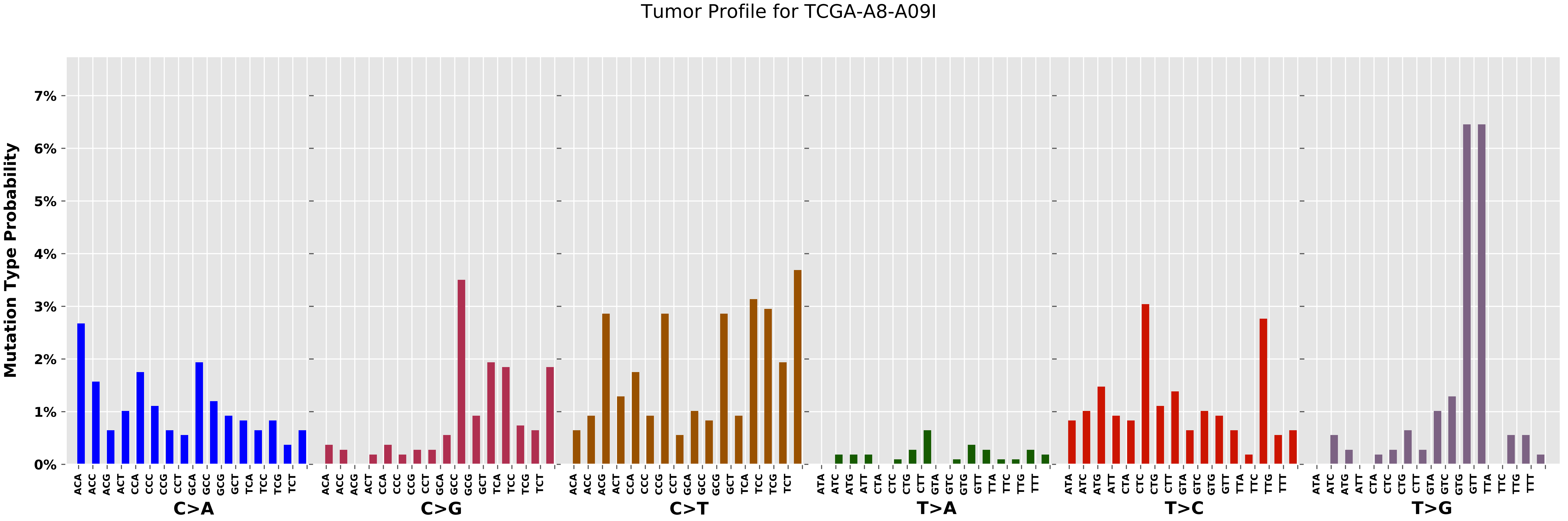

Tumor Profile for TCGA-A8-A09I
Mutation Type Probability
0%
1%
2%
3%
4%
5%
6%
7%
C>A
C>G
C>T
T>A
T>C
T>G
ACA ACC ACG ACT CCA CCC CCG CCT GCA GCC GCG GCT TCA TCC TCG TCT
ATA ATC ATG ATT CTA CTC CTG CTT GTA GTC GTG GTT TTA TTC TTG TTT

Cancer processes Weights for TCGA-AA-3664

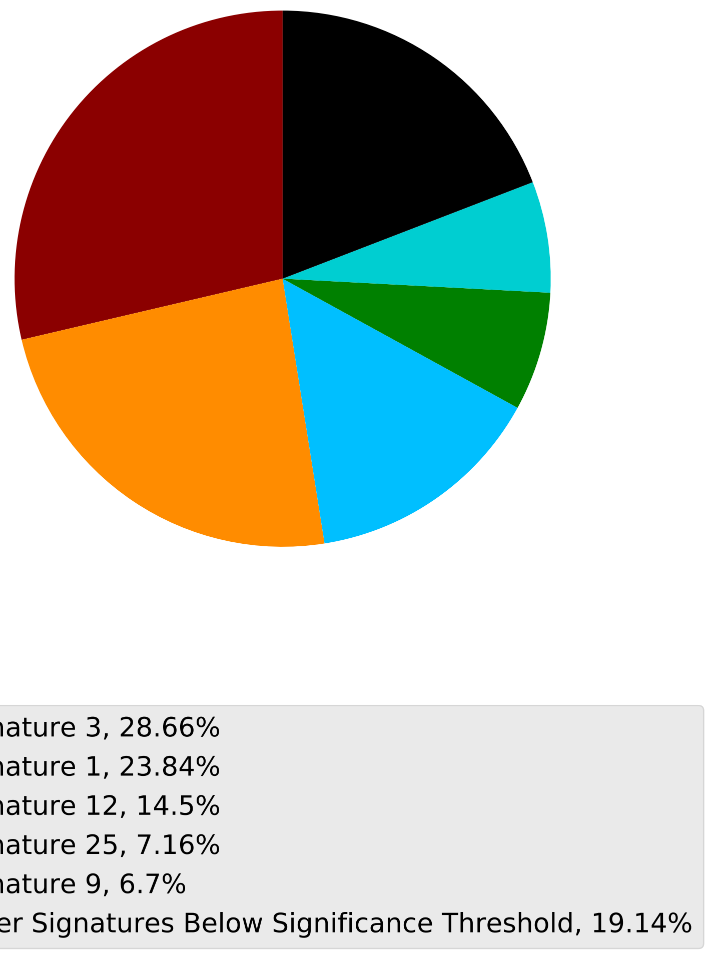

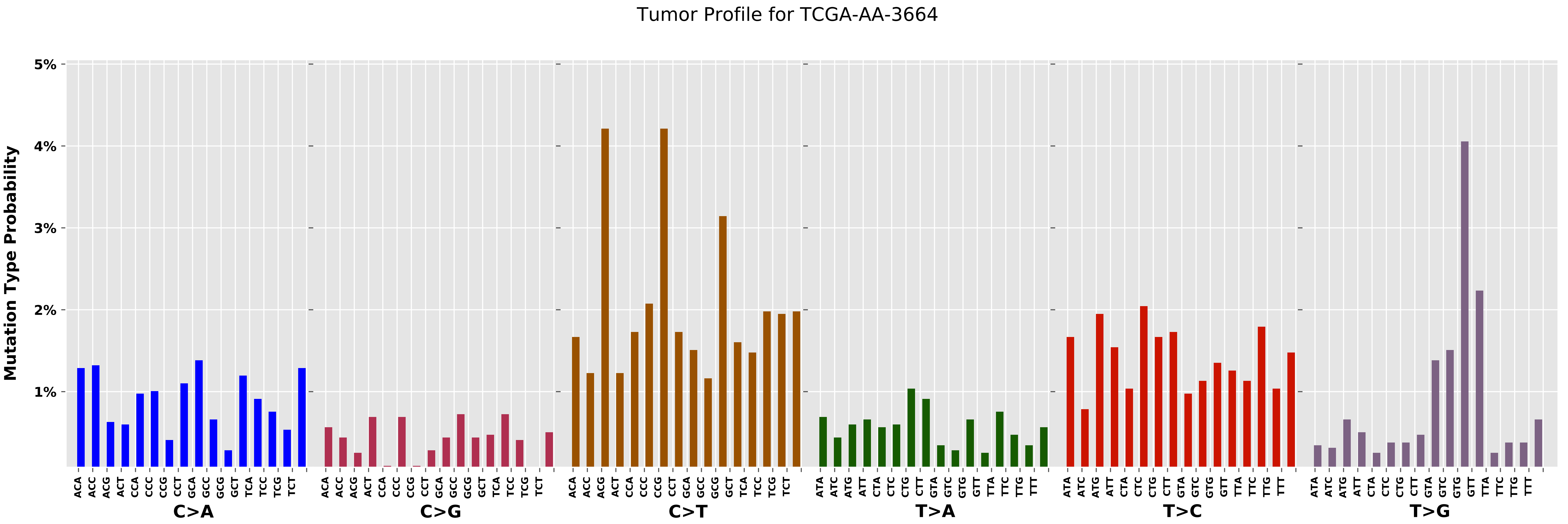

Tumor Profile for TCGA-AA-3664
Mutation Type Probability
5%
4%
3%
2%
1%
C>A
C>G
C>T
T>A
T>C
T>G
ACA ACC ACG ACT CCA CCC CCG CCT GCA GCC GCG GCT TCA TCC TCG TCT
ATA ATC ATG ATT CTA CTC CTG CTT GTA GTC GTG GTT TTA TTC TTG TTT

#### Cancer processes Weights for TCGA-E2-A1LL

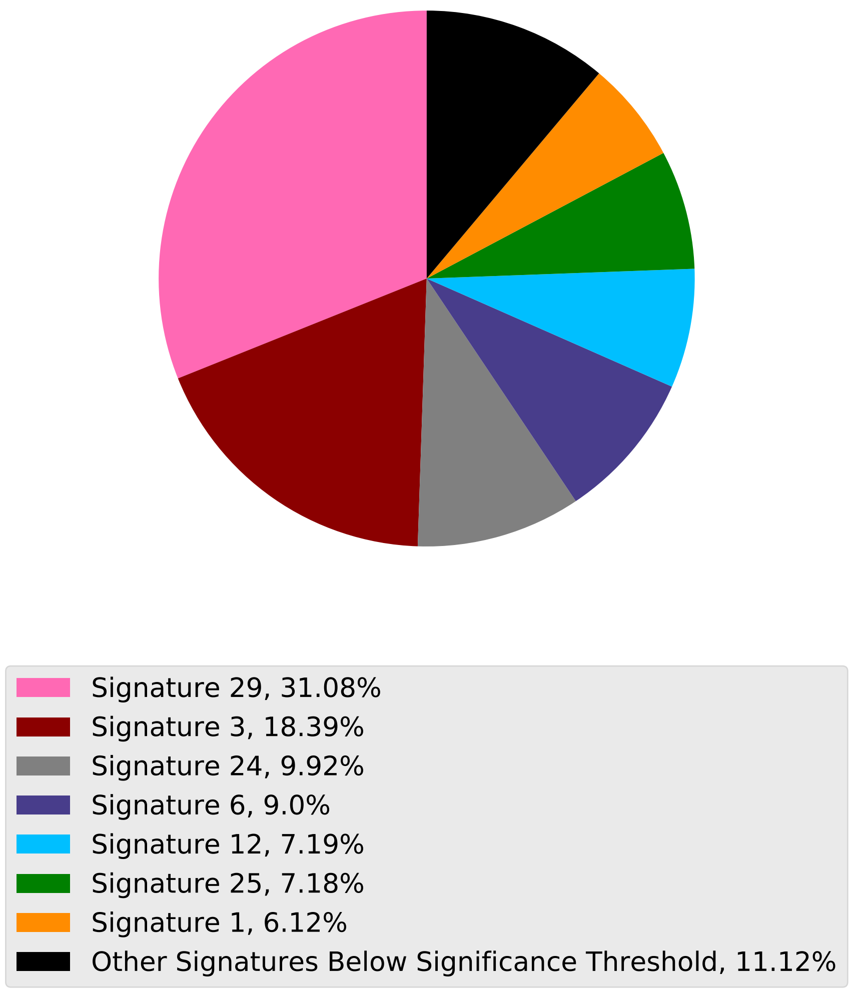

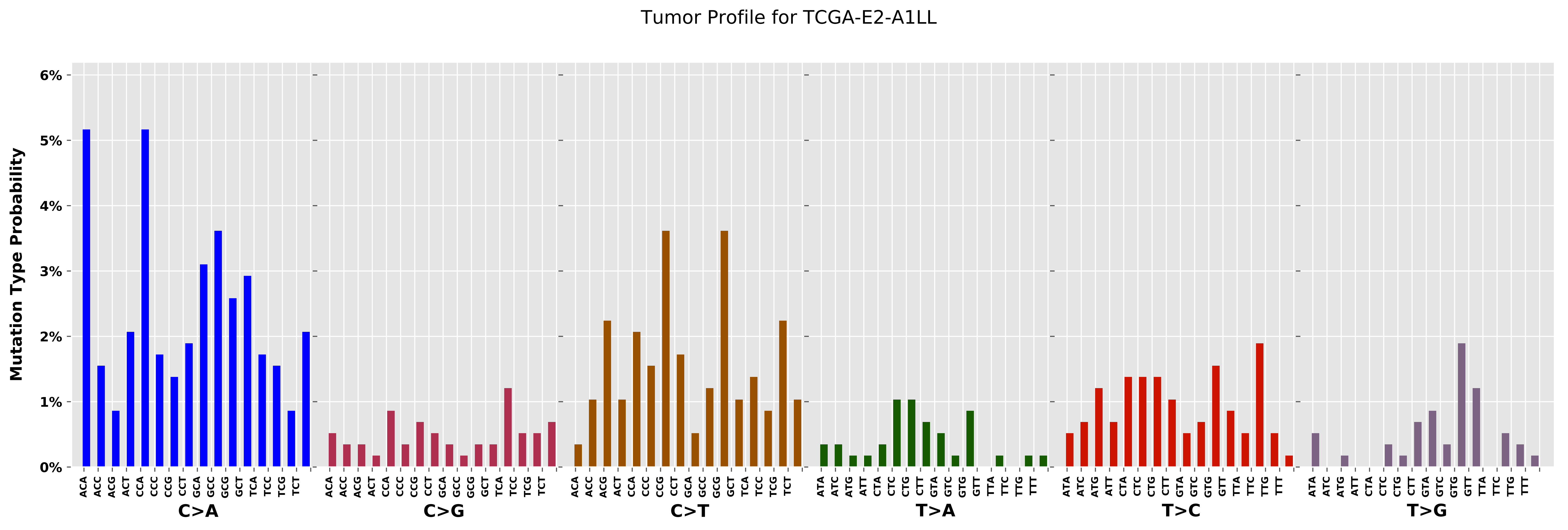
Tumor Profile for TCGA-E2-A1LL
Mutation Type Probability
0%
1%
2%
3%
4%
5%
6%
C>A
C>G
C>T
T>A
T>C
T>G
ACA ACC ACG ACT CCA CCC CCG CCT GCA GCC GCG GCT TCA TCC TCG TCT
ATA ATC ATG ATT CTA CTC CTG CTT GTA GTC GTG GTT TTA TTC TTG TTT

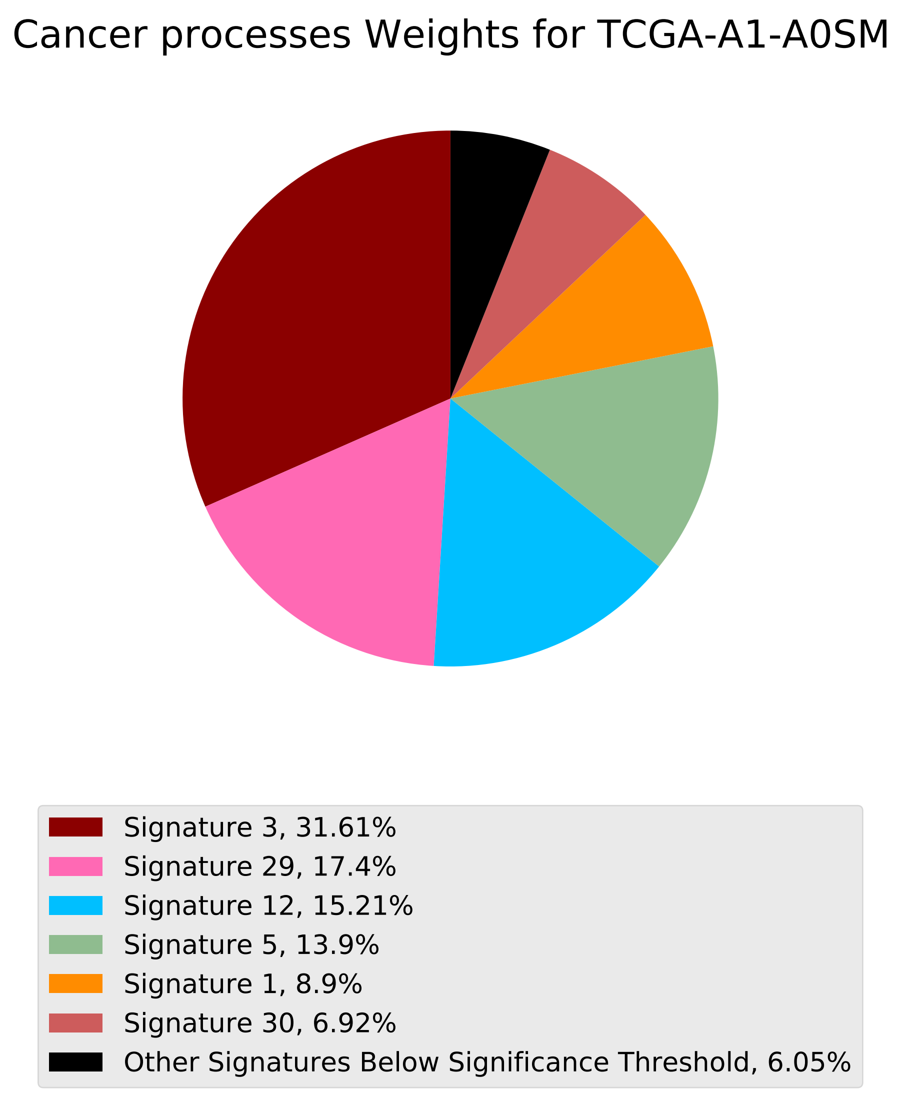
Cancer processes Weights for TCGA-A1-A0SM
Signature 3, 31.61%
Signature 29, 17.4%
Signature 12, 15.21%
Signature 5, 13.9%
Signature 1, 8.9%
Signature 30, 6.92%
Other Signatures Below Significance Threshold, 6.05%

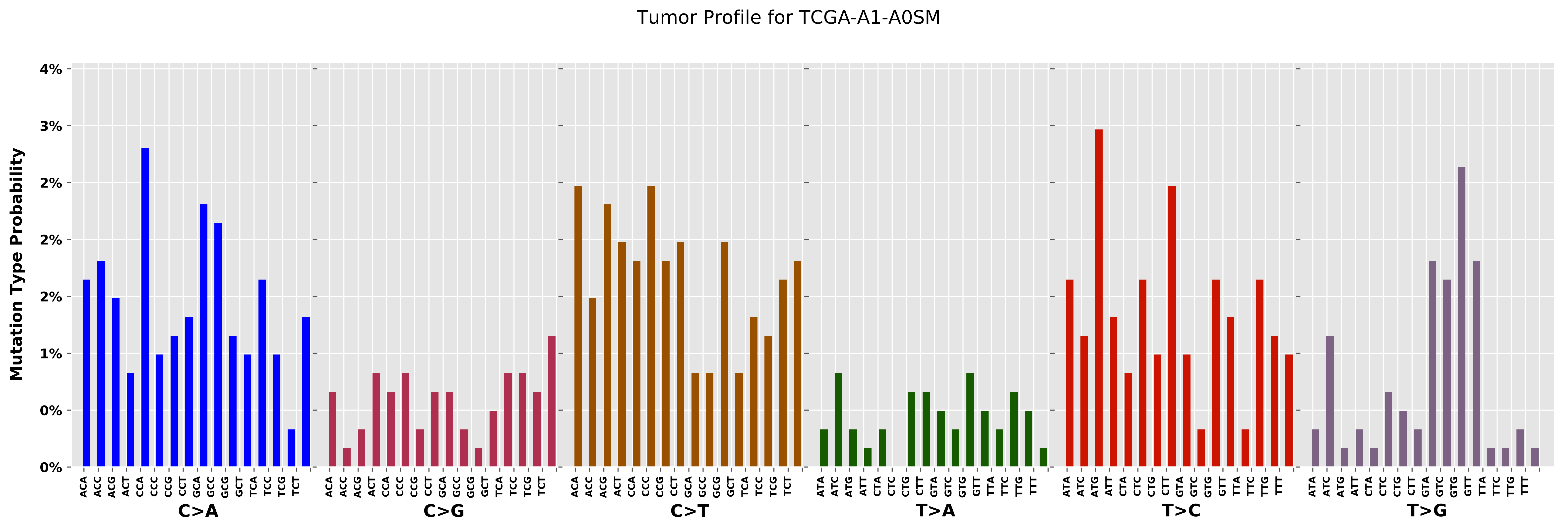
Tumor Profile for TCGA-A1-A0SM
Mutation Type Probability
4%
3%
2%
2%
2%
1%
0%
0%
C>A
C>G
C>T
T>A
T>C
T>G
ACA
ACC
ACG
ACT
CCA
CCC
CCG
CCT
GCA
GCC
GCG
GCT
TCA
TCC
TCG
TCT
ATA
ATC
ATG
ATT
CTA
CTC
CTG
CTT
GTA
GTC
GTG
GTT
TTA
TTC
TTG
TTT

Cancer processes Weights for TCGA-C8-A130

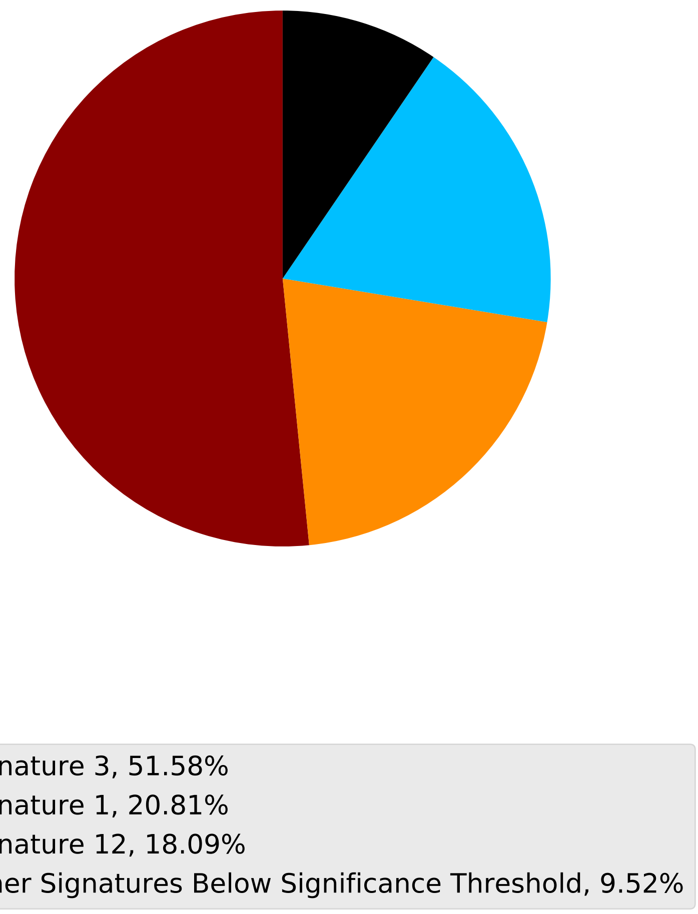

Tumor Profile for TCGA-C8-A130

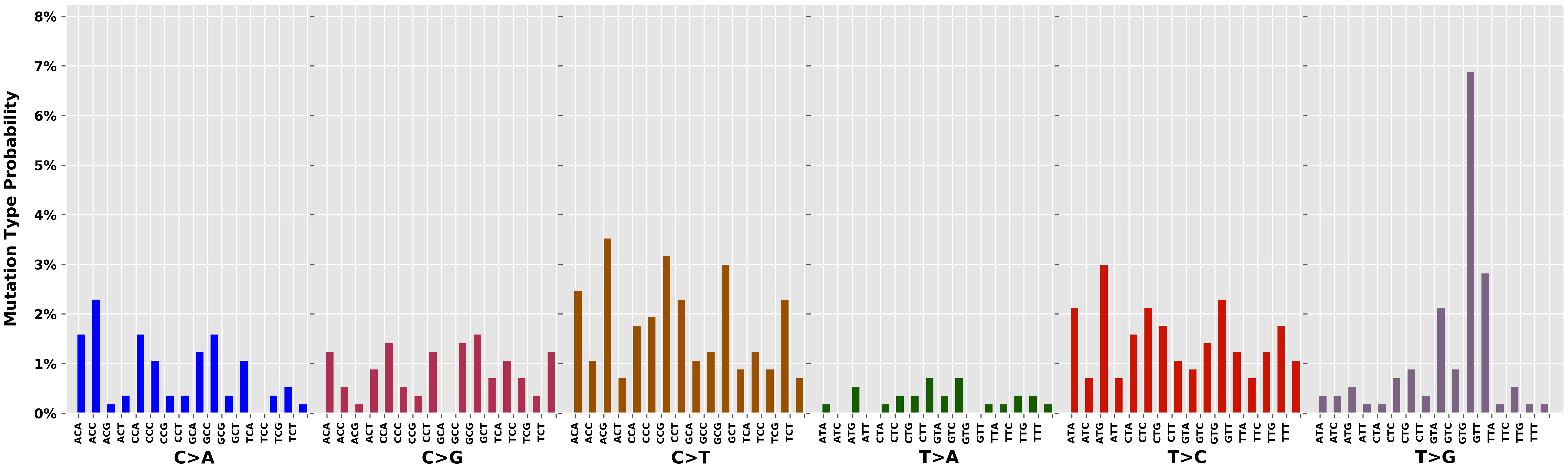

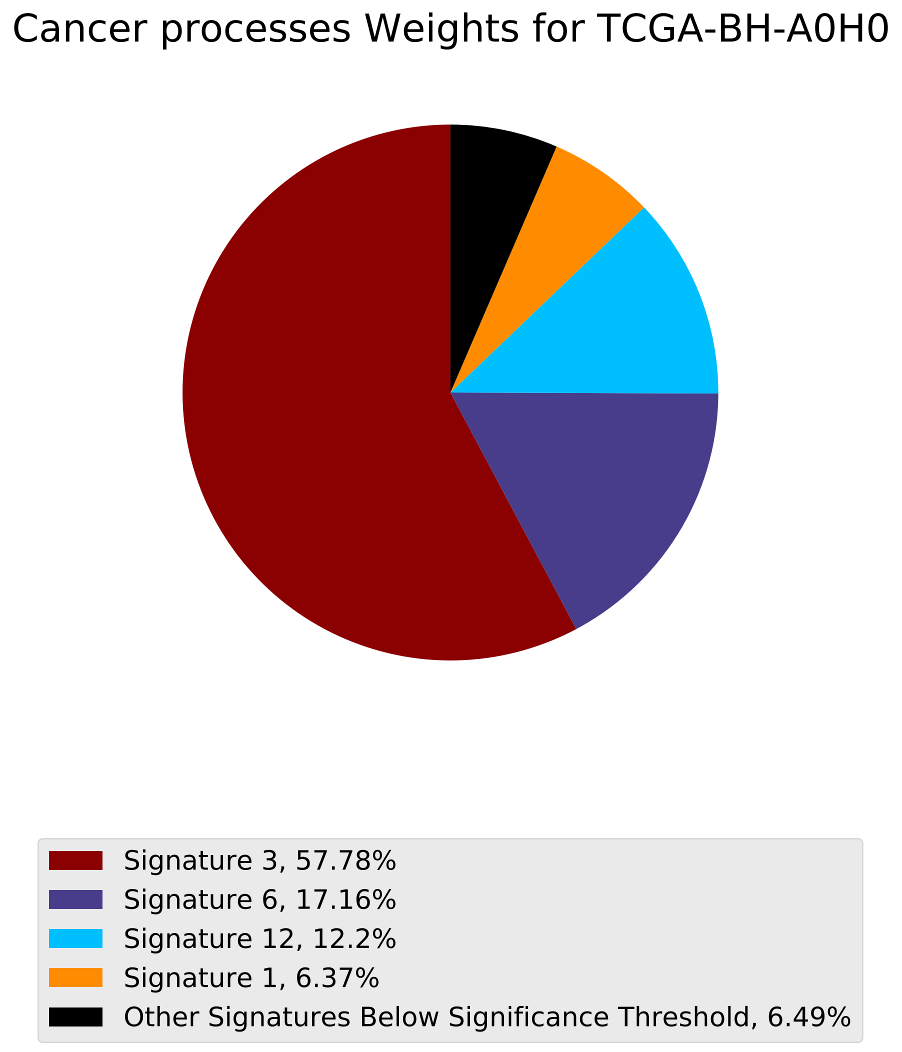
Cancer processes Weights for TCGA-BH-A0H0
Signature 3, 57.78%
Signature 6, 17.16%
Signature 12, 12.2%
Signature 1, 6.37%
Other Signatures Below Significance Threshold, 6.49%

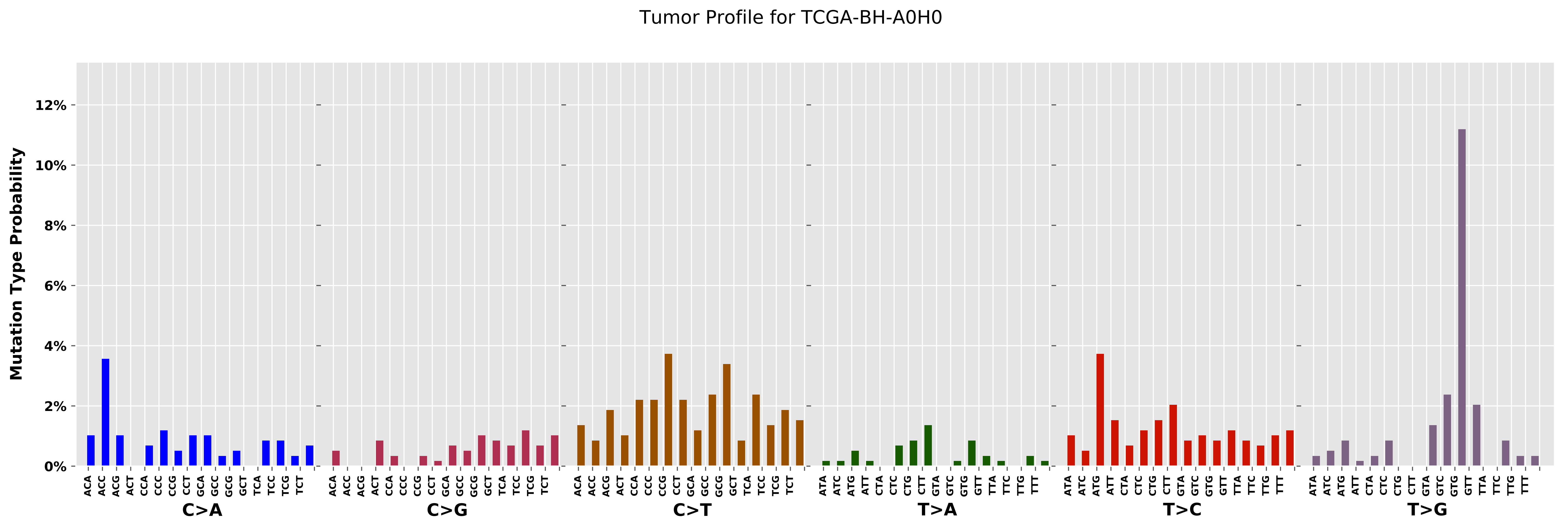

Tumor Profile for TCGA-BH-A0H0
Mutation Type Probability
0%
2%
4%
6%
8%
10%
12%
C>A
C>G
C>T
T>A
T>C
T>G

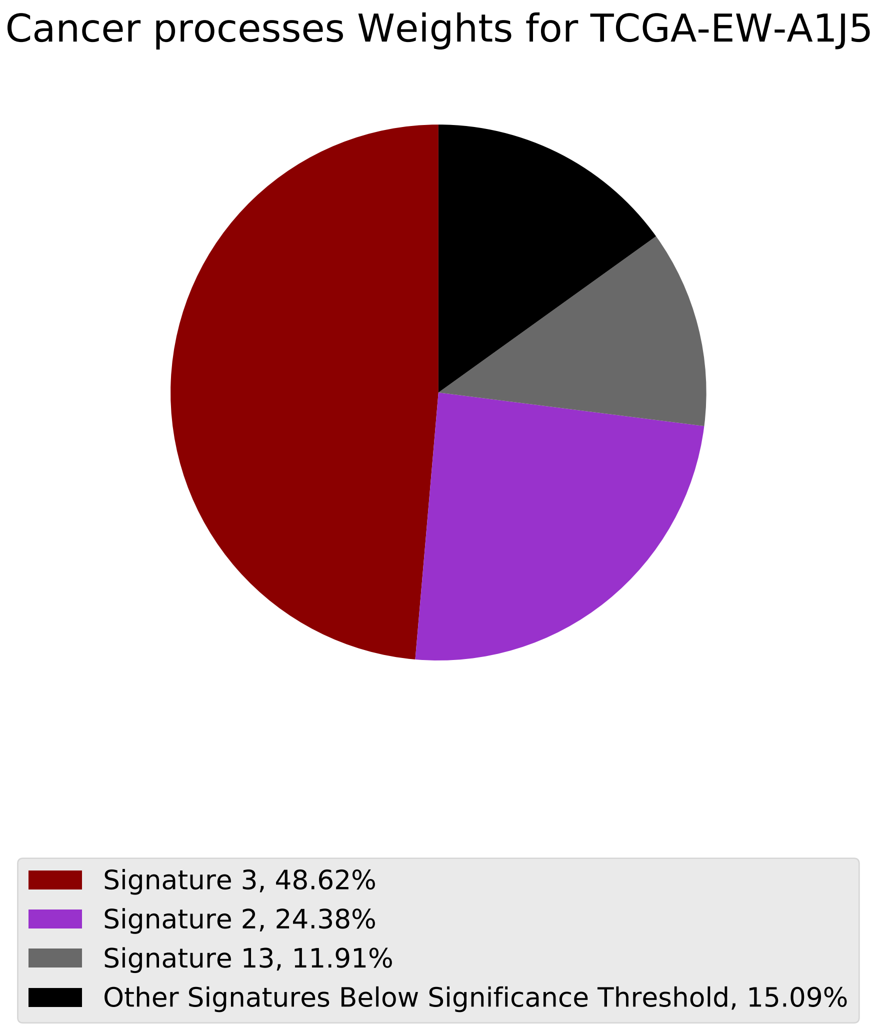

Cancer processes Weights for TCGA-EW-A1J5
Signature 3, 48.62%
Signature 2, 24.38%
Signature 13, 11.91%
Other Signatures Below Significance Threshold, 15.09%

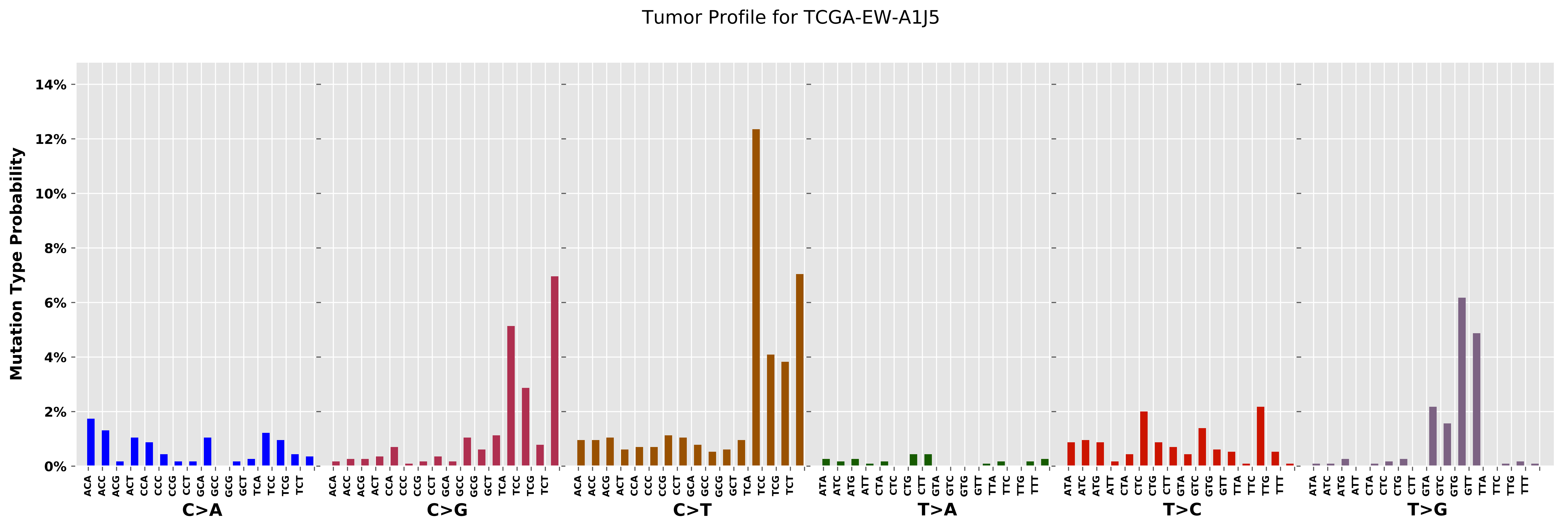

Tumor Profile for TCGA-EW-A1J5
Mutation Type Probability
14%
12%
10%
8%
6%
4%
2%
0%
ACA ACC ACG ACT CCA CCC CCG CCT GCA GCC GCG GCT TCA TCC TCG TCT
C>A
ACA ACC ACG ACT CCA CCC CCG CCT GCA GCC GCG GCT TCA TCC TCG TCT
C>G
ACA ACC ACG ACT CCA CCC CCG CCT GCA GCC GCG GCT TCA TCC TCG TCT
C>T
ATA ATC ATG ATT CTA CTC CTG CTT GTA GTC GTG GTT TTA TTC TTG TTT
T>A
ATA ATC ATG ATT CTA CTC CTG CTT GTA GTC GTG GTT TTA TTC TTG TTT
T>C
ATA ATC ATG ATT CTA CTC CTG CTT GTA GTC GTG GTT TTA TTC TTG TTT
T>G

#### Cancer processes Weights for TCGA-EW-A1PC

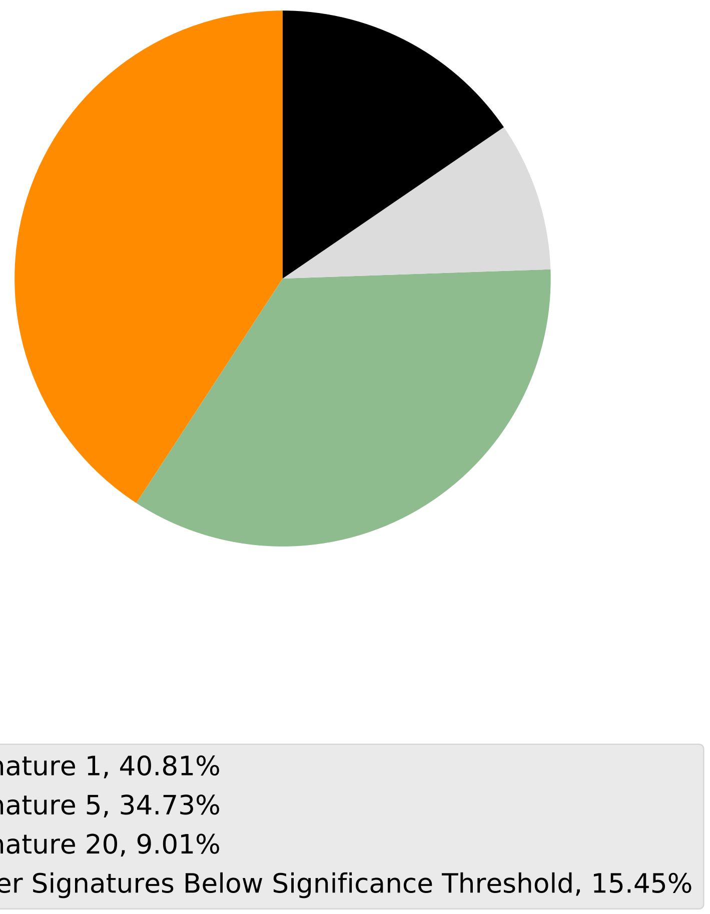

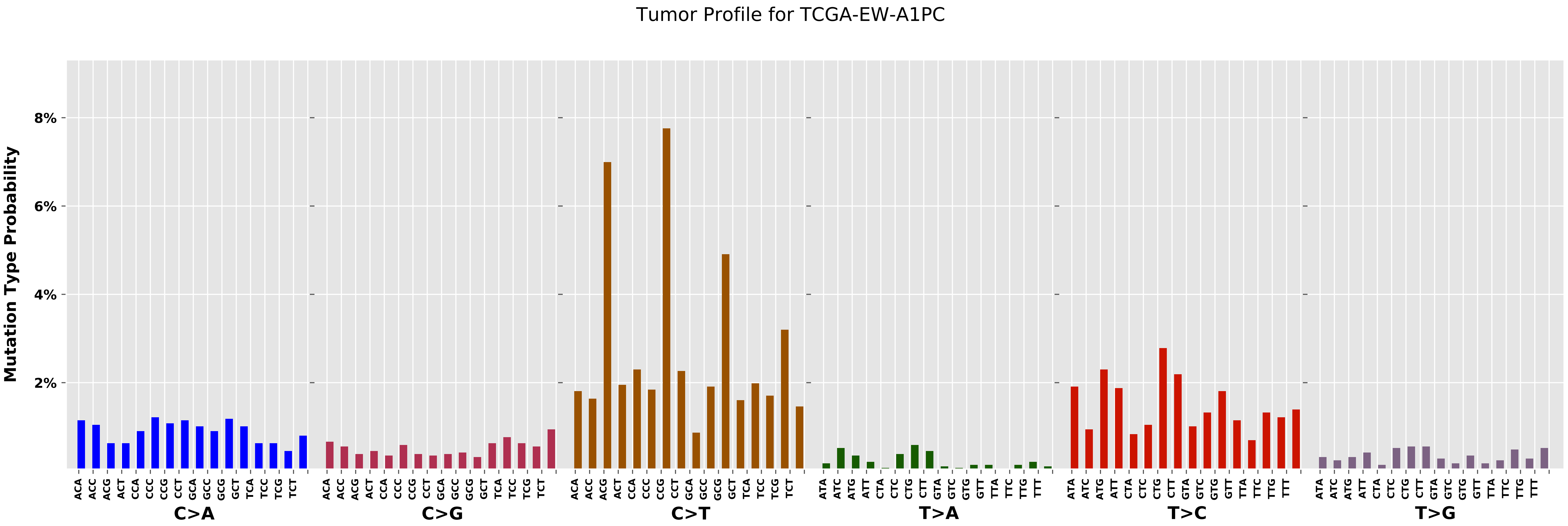
Tumor Profile for TCGA-EW-A1PC
Mutation Type Probability
8%
6%
4%
2%
ACA ACC ACG ACT CCA CCC CCG CCT GCA GCC GCG GCT TCA TCC TCG TCT
ATA ATC ATG ATT CTA CTC CTG CTT GTA GTC GTG GTT TTA TTC TTG TTT
C>A
C>G
C>T
T>A
T>C
T>G

Cancer processes Weights for TCGA-BH-A0E0

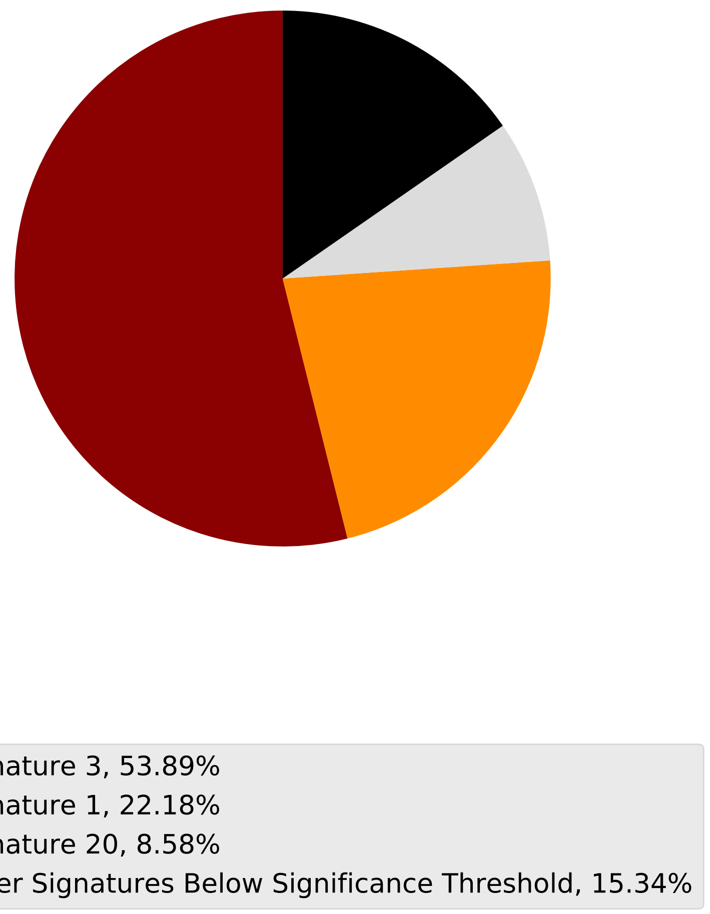

Tumor Profile for TCGA-BH-A0E0

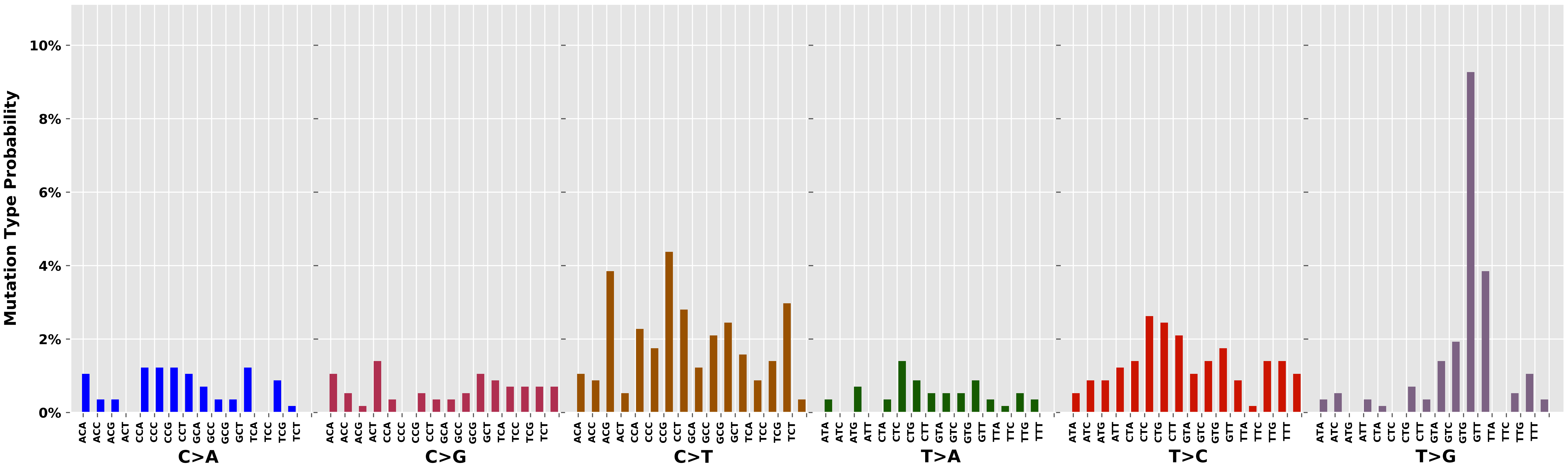

Cancer processes Weights for TCGA-A2-A0D0

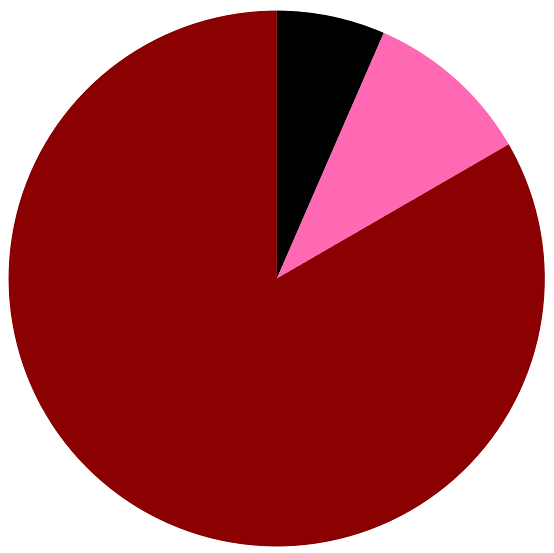

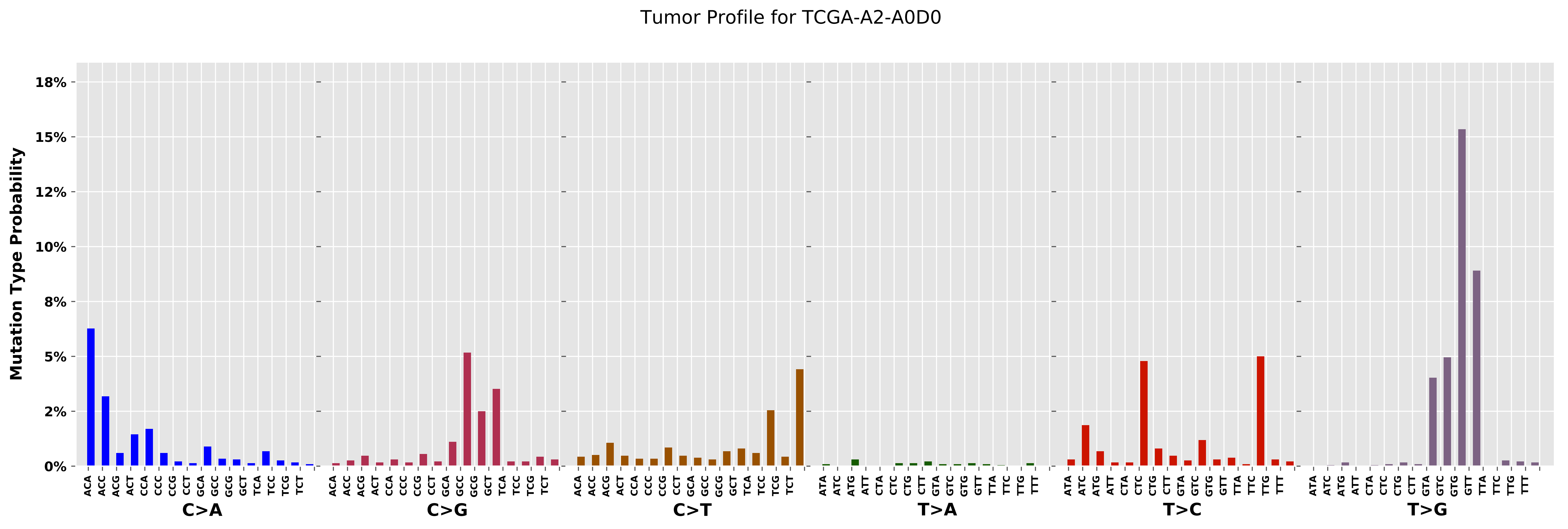

Tumor Profile for TCGA-A2-A0D0
Mutation Type Probability
0%
2%
5%
8%
10%
12%
15%
18%
C>A
C>G
C>T
T>A
T>C
T>G

Cancer processes Weights for TCGA-AA-3956

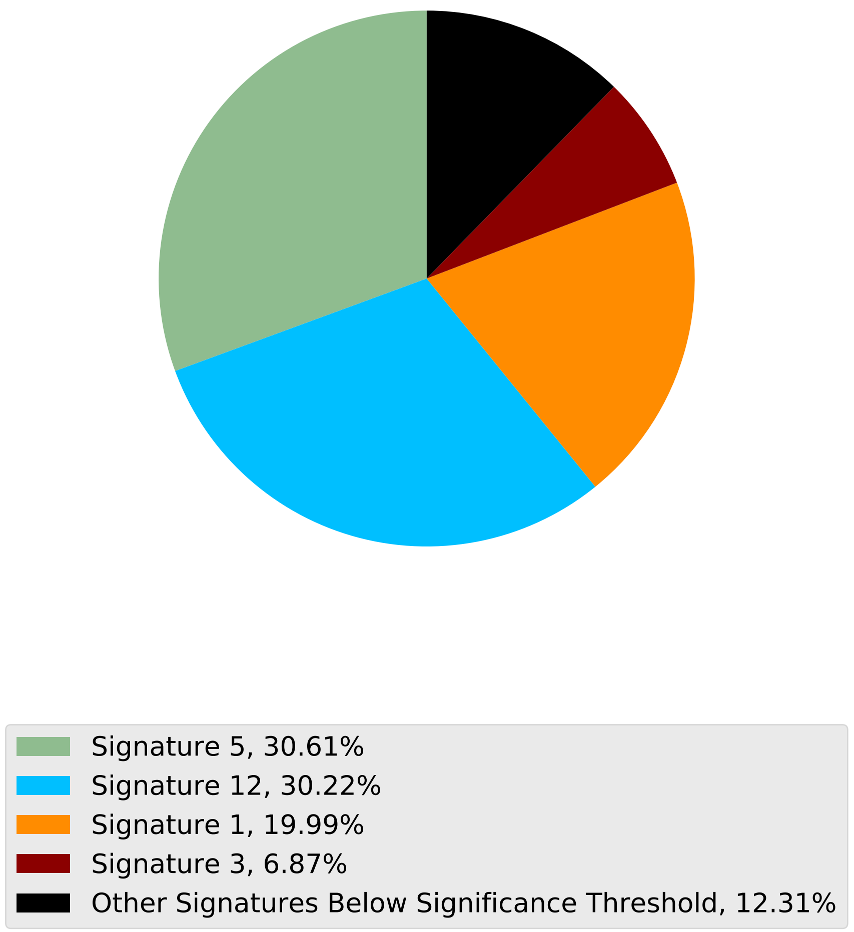

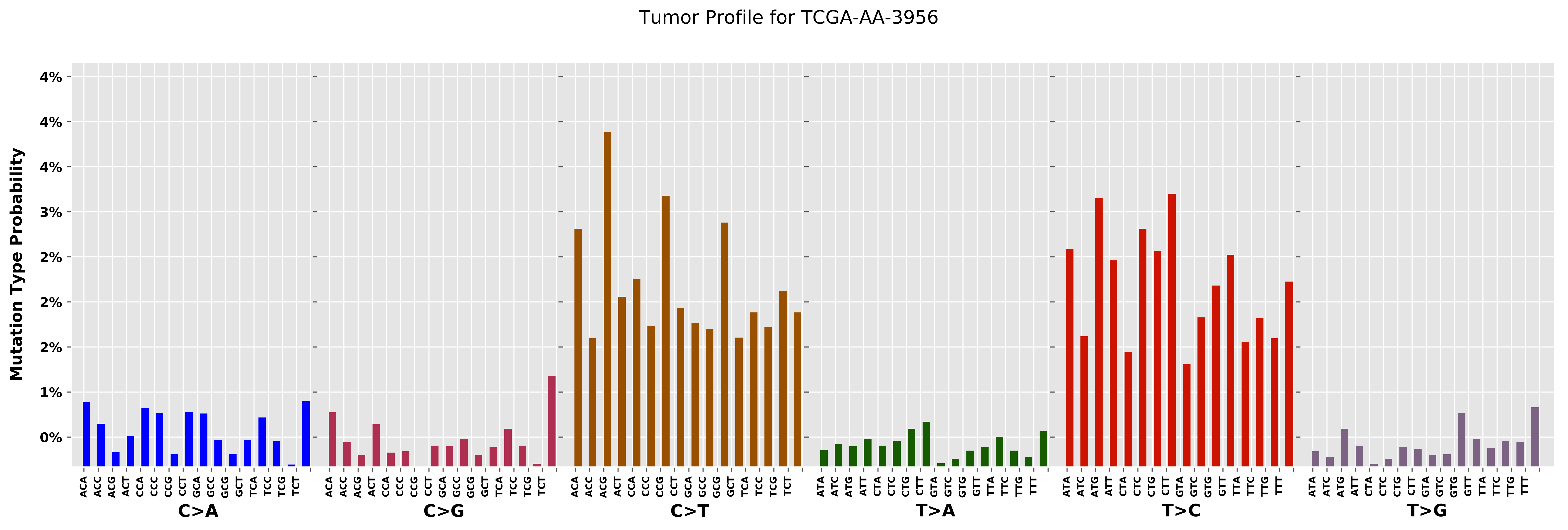
Tumor Profile for TCGA-AA-3956
Mutation Type Probability
4%
4%
4%
3%
2%
2%
2%
1%
0%
C>A
C>G
C>T
T>A
T>C
T>G

Cancer processes Weights for TCGA-AR-A0TX

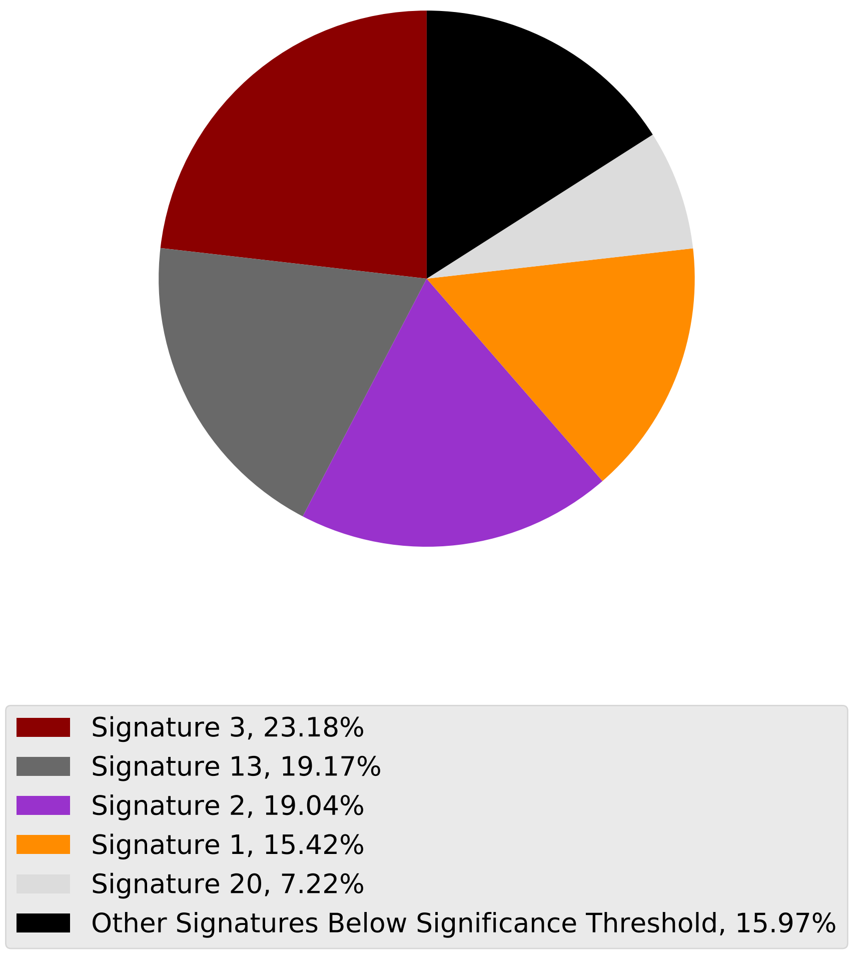

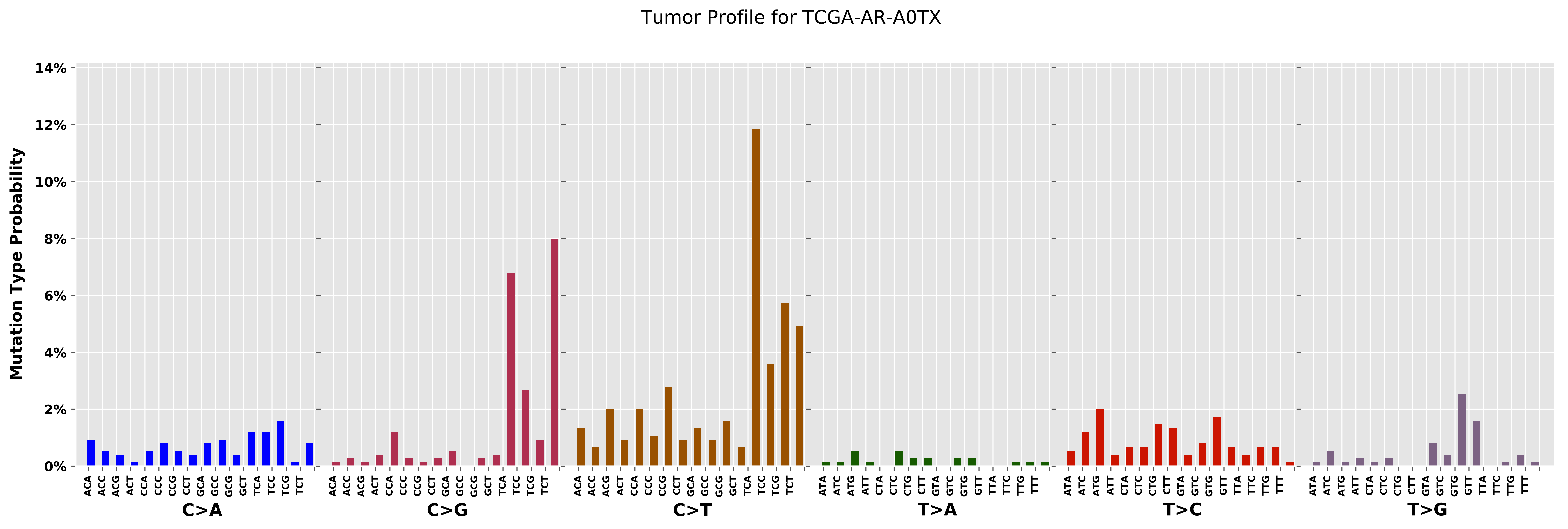
Tumor Profile for TCGA-AR-A0TX
Mutation Type Probability
0%
2%
4%
6%
8%
10%
12%
14%
ACA ACC ACG ACT CCA CCC CCG CCT GCA GCC GCG GCT TCA TCC TCG TCT
C>A
ACA ACC ACG ACT CCA CCC CCG CCT GCA GCC GCG GCT TCA TCC TCG TCT
C>G
ACA ACC ACG ACT CCA CCC CCG CCT GCA GCC GCG GCT TCA TCC TCG TCT
C>T
ATA ATC ATG ATT CTA CTC CTG CTT GTA GTC GTG GTT TTA TTC TTG TTT
T>A
ATA ATC ATG ATT CTA CTC CTG CTT GTA GTC GTG GTT TTA TTC TTG TTT
T>C
ATA ATC ATG ATT CTA CTC CTG CTT GTA GTC GTG GTT TTA TTC TTG TTT
T>G

Cancer processes Weights for TCGA-EI-6917

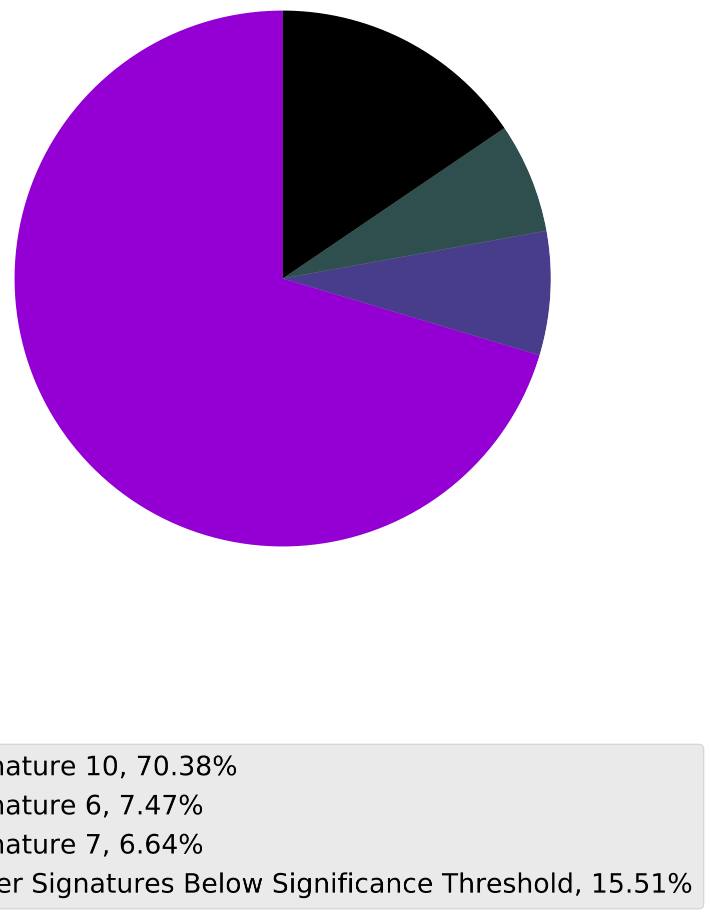

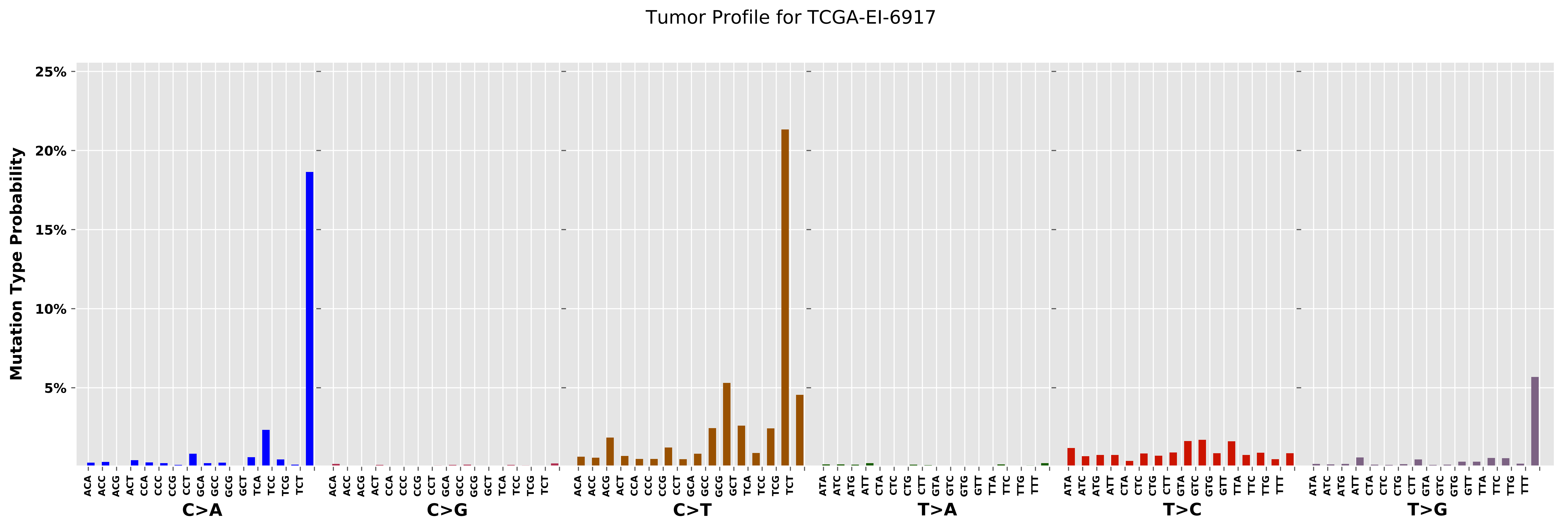

Tumor Profile for TCGA-EI-6917
Mutation Type Probability
25%
20%
15%
10%
5%
C>A
C>G
C>T
T>A
T>C
T>G
ACA
ACC
ACG
ACT
CCA
CCC
CCG
CCT
GCA
GCC
GCG
GCT
TCA
TCC
TCG
TCT
ATA
ATC
ATG
ATT
CTA
CTC
CTG
CTT
GTA
GTC
GTG
GTT
TTA
TTC
TTG
TTT

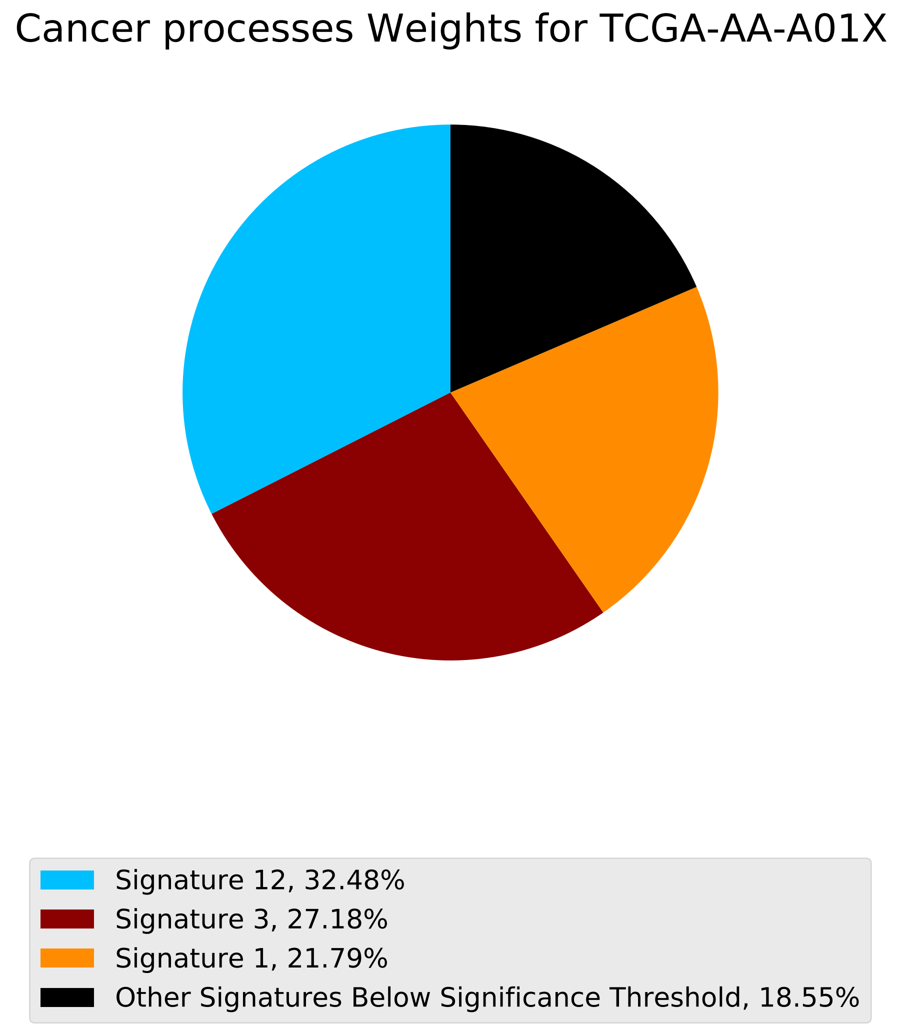

Cancer processes Weights for TCGA-AA-A01X
Signature 12, 32.48%
Signature 3, 27.18%
Signature 1, 21.79%
Other Signatures Below Significance Threshold, 18.55%

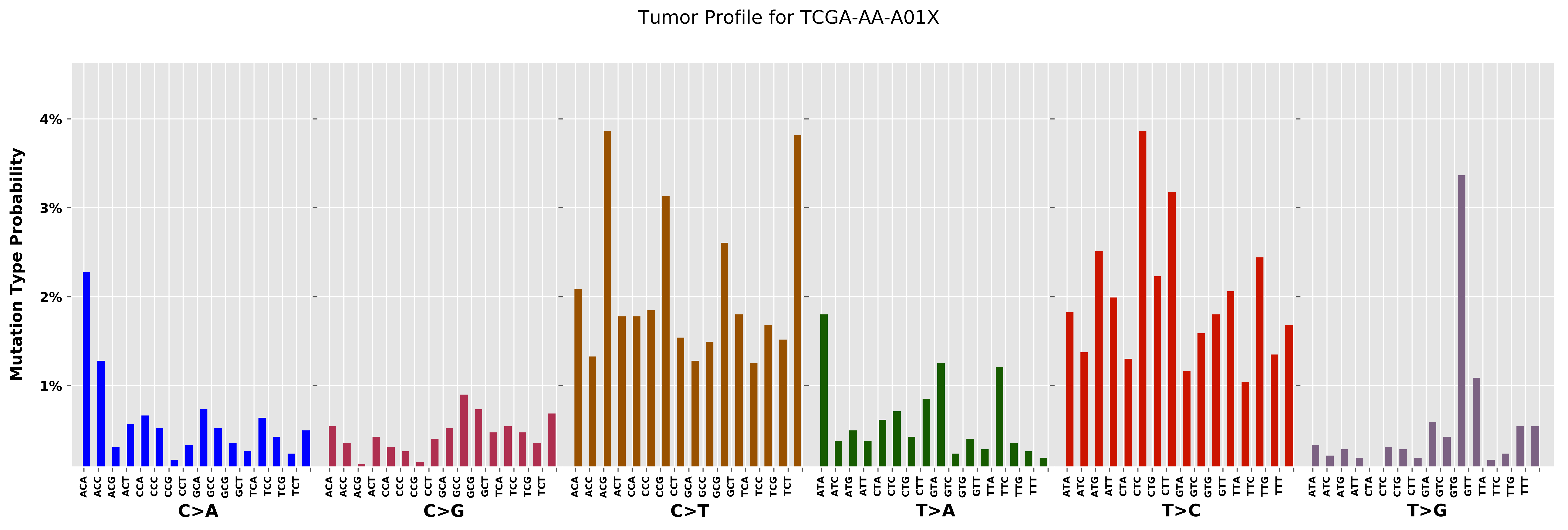
Tumor Profile for TCGA-AA-A01X
Mutation Type Probability
4%
3%
2%
1%
ACA ACC ACG ACT CCA CCC CCG CCT GCA GCC GCG GCT TCA TCC TCG TCT
C>A
ACA ACC ACG ACT CCA CCC CCG CCT GCA GCC GCG GCT TCA TCC TCG TCT
C>G
ACA ACC ACG ACT CCA CCC CCG CCT GCA GCC GCG GCT TCA TCC TCG TCT
C>T
ATA ATC ATG ATT CTA CTC CTG CTT GTA GTC GTG GTT TTA TTC TTG TTT
T>A
ATA ATC ATG ATT CTA CTC CTG CTT GTA GTC GTG GTT TTA TTC TTG TTT
T>C
ATA ATC ATG ATT CTA CTC CTG CTT GTA GTC GTG GTT TTA TTC TTG TTT
T>G

Cancer processes Weights for TCGA-E2-A15H

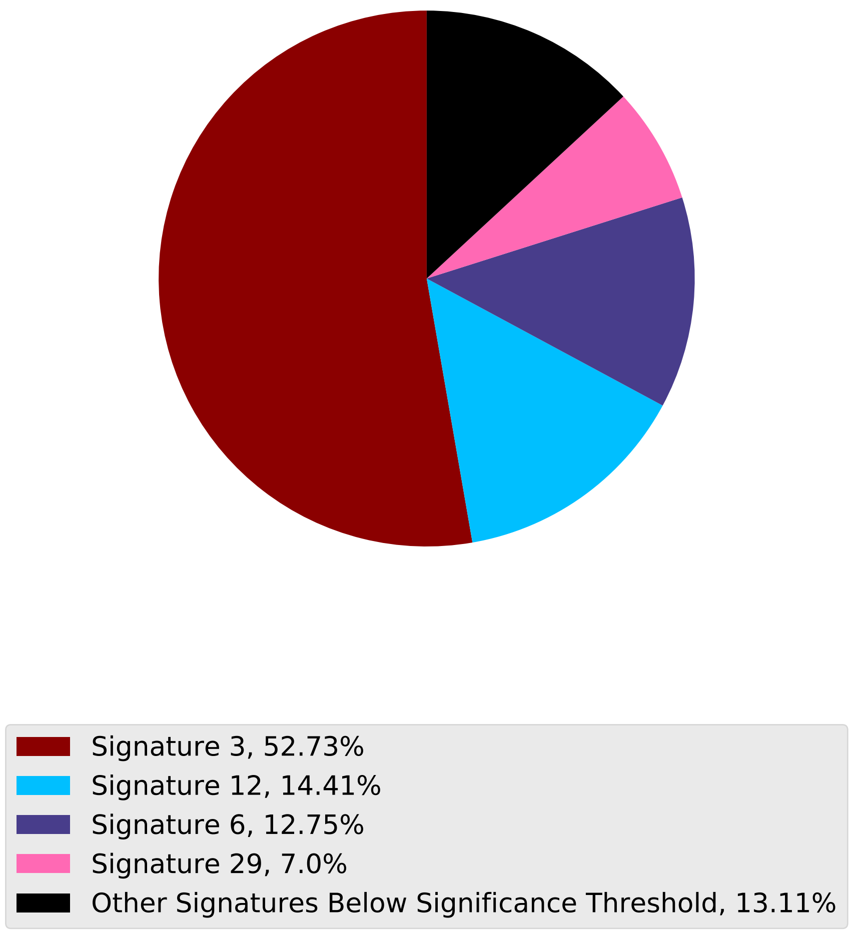

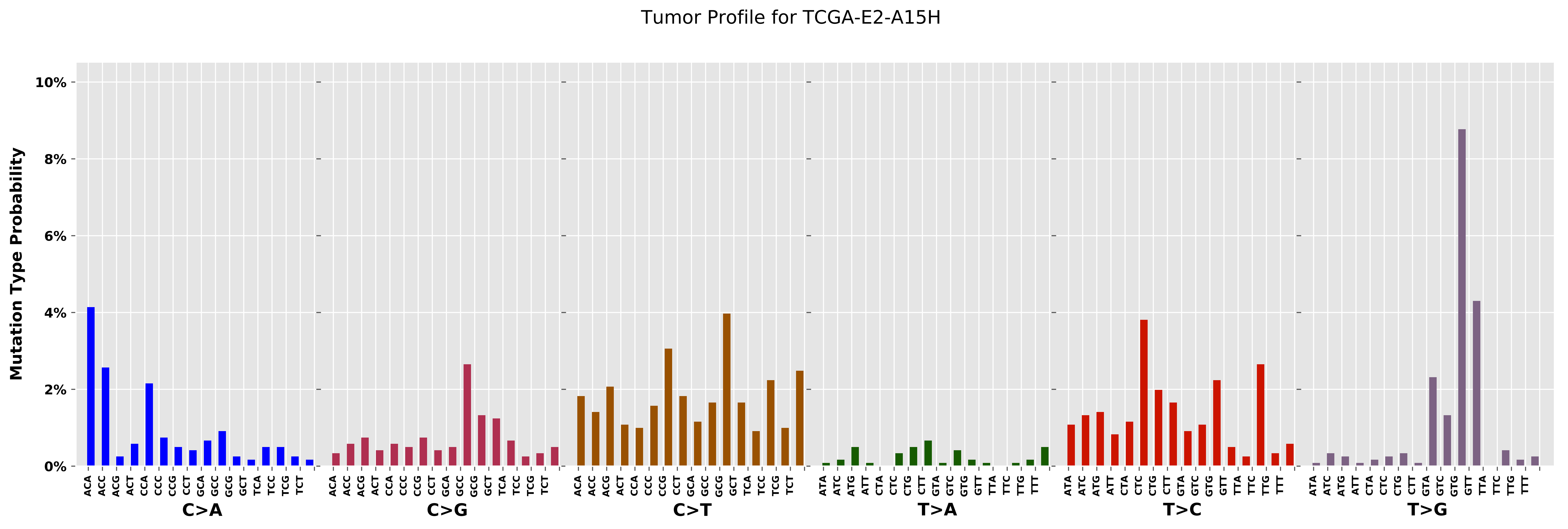
Tumor Profile for TCGA-E2-A15H
Mutation Type Probability
10%
8%
6%
4%
2%
0%
C>A
C>G
C>T
T>A
T>C
T>G
ACA ACC ACG ACT CCA CCC CCG CCT GCA GCC GCG GCT TCA TCC TCG TCT
ATA ATC ATG ATT CTA CTC CTG CTT GTA GTC GTG GTT TTA TTC TTG TTT

Cancer processes Weights for TCGA-E9-A1NH

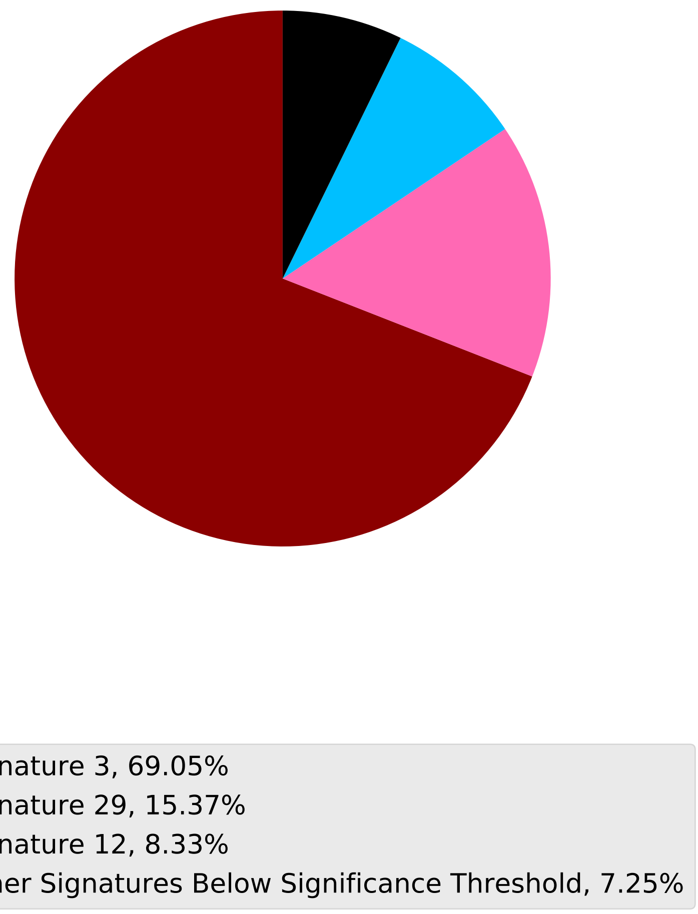

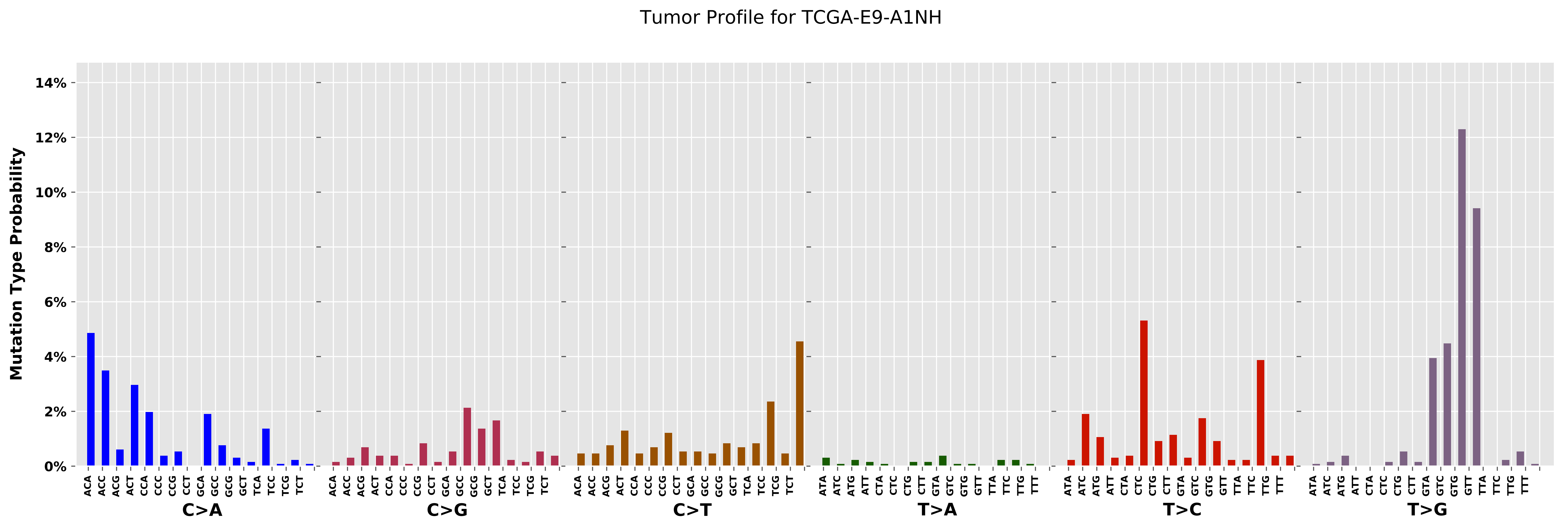

Tumor Profile for TCGA-E9-A1NH
Mutation Type Probability
14%
12%
10%
8%
6%
4%
2%
0%
C>A
C>G
C>T
T>A
T>C
T>G

Cancer processes Weights for TCGA-AA-A01R

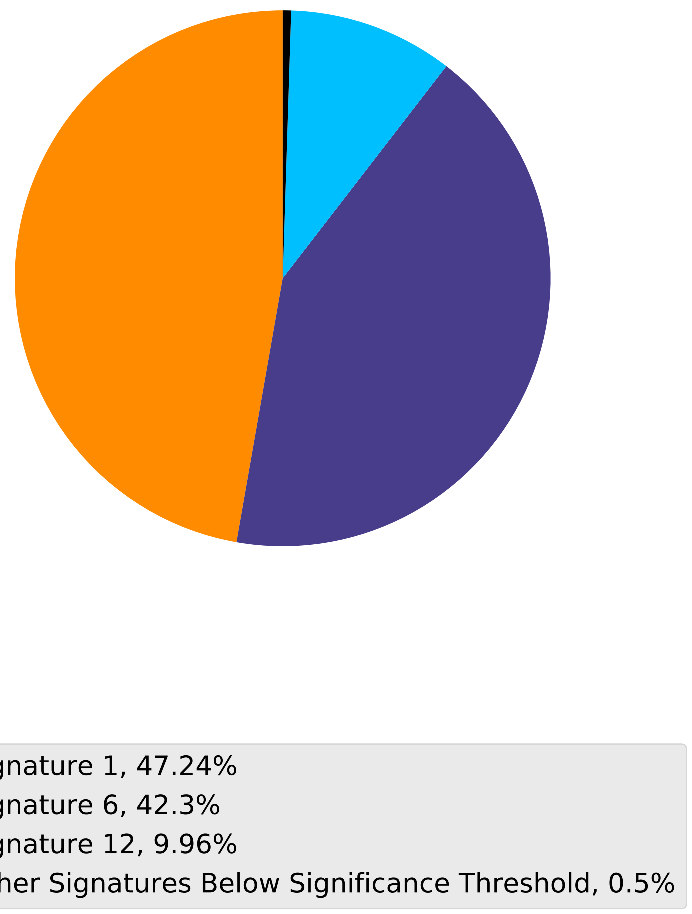

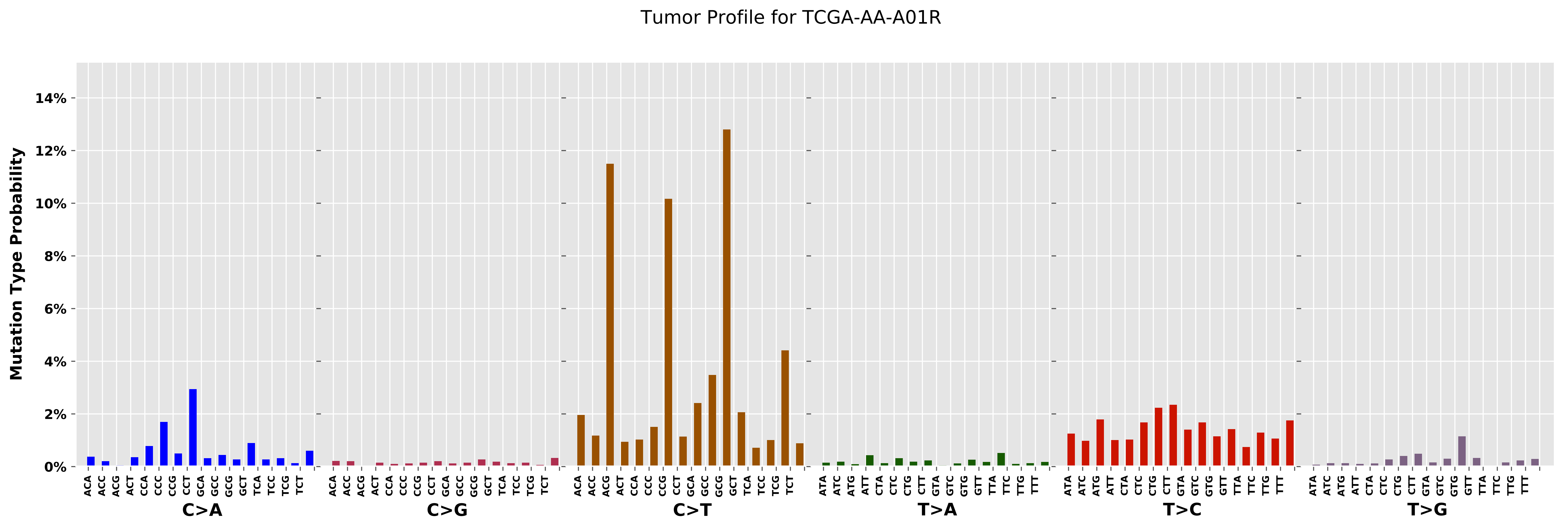

Tumor Profile for TCGA-AA-A01R
Mutation Type Probability
0%
2%
4%
6%
8%
10%
12%
14%
C>A
C>G
C>T
T>A
T>C
T>G
ACA ACC ACG ACT CCA CCC CCG CCT GCA GCC GCG GCT TCA TCC TCG TCT
ATA ATC ATG ATT CTA CTC CTG CTT GTA GTC GTG GTT TTA TTC TTG TTT

Cancer processes Weights for TCGA-AA-3666

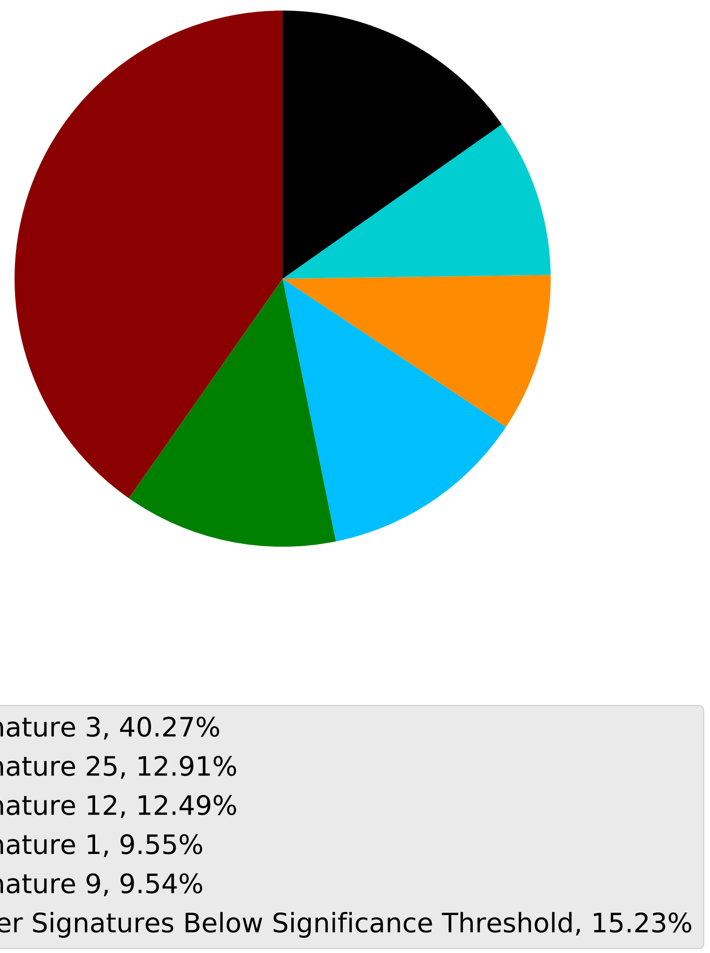

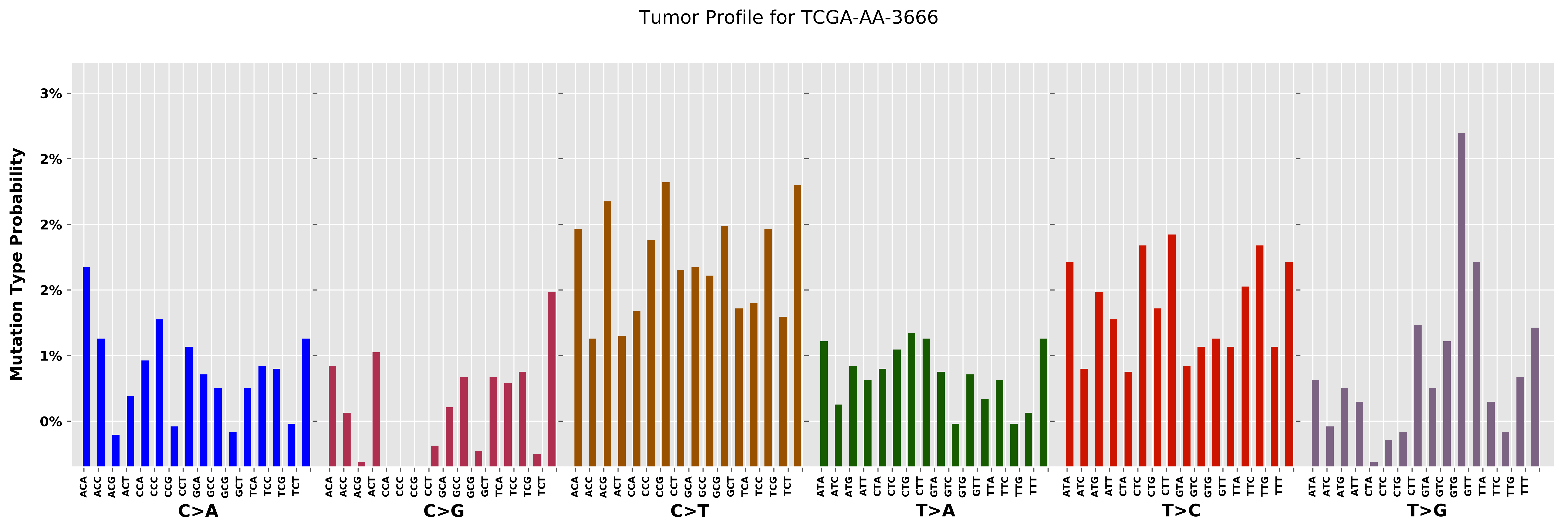
Tumor Profile for TCGA-AA-3666
Mutation Type Probability
3%
2%
2%
2%
2%
1%
0%
C>A
C>G
C>T
T>A
T>C
T>G
ACA
ACC
ACG
ACT
CCA
CCC
CCG
CCT
GCA
GCC
GCG
GCT
TCA
TCC
TCG
TCT
ATA
ATC
ATG
ATT
CTA
CTC
CTG
CTT
GTA
GTC
GTG
GTT
TTA
TTC
TTG
TTT

Cancer processes Weights for TCGA-AR-A2LK

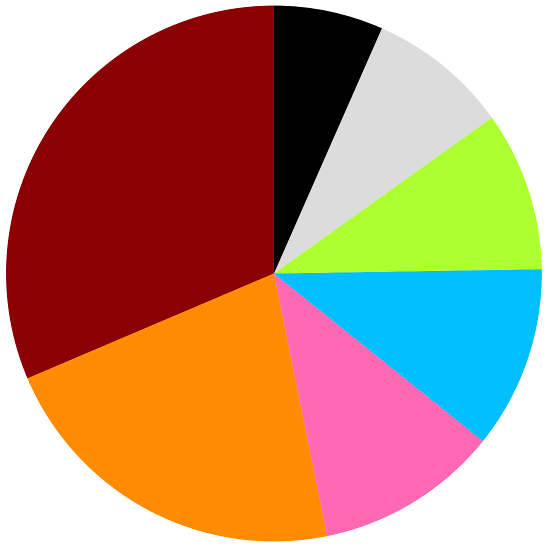

Signature 3, 31.4%
Signature 1, 21.78%
Signature 29, 11.05%
Signature 12, 11.02%
Signature 26, 9.62%
Signature 20, 8.52%
Other Signatures Below Significance Threshold, 6.6%

Tumor Profile for TCGA-AR-A2LK

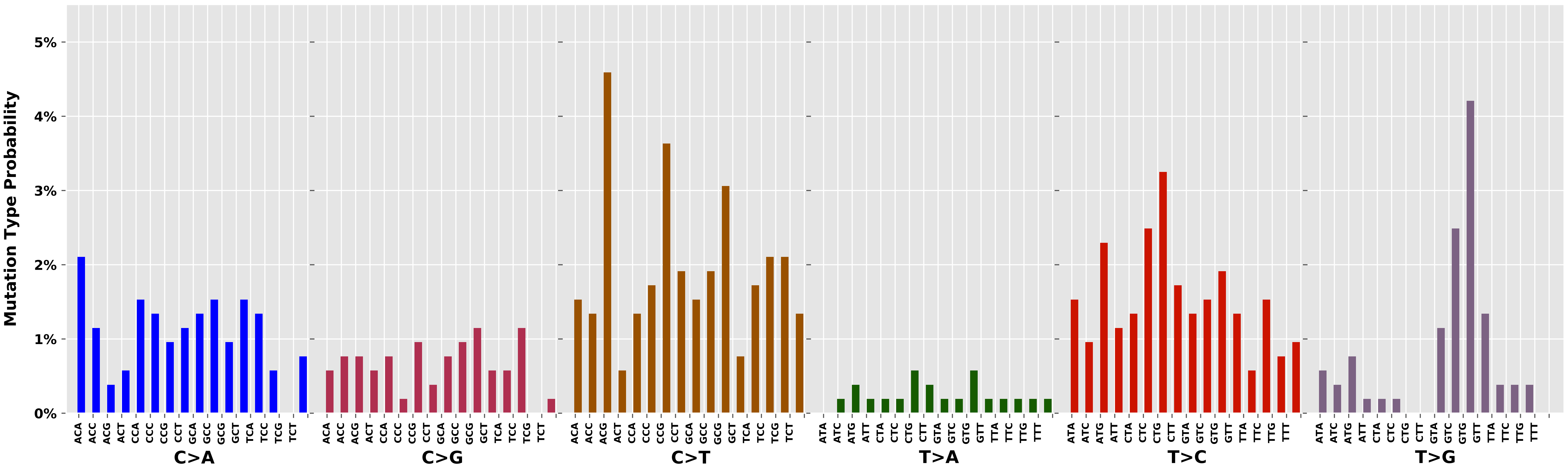

Cancer processes Weights for TCGA-AA-3534

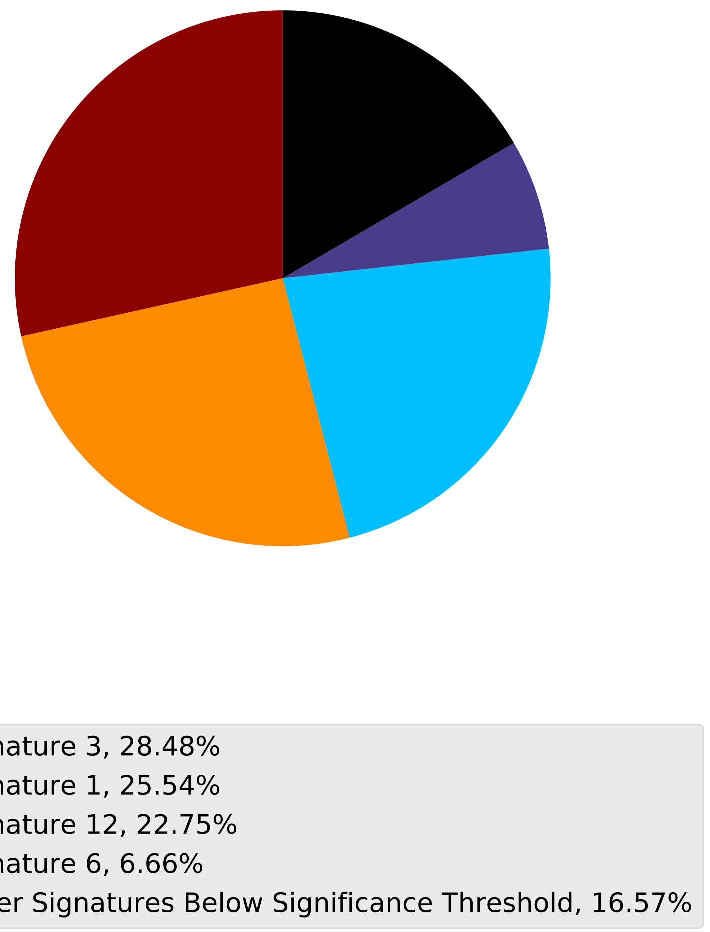

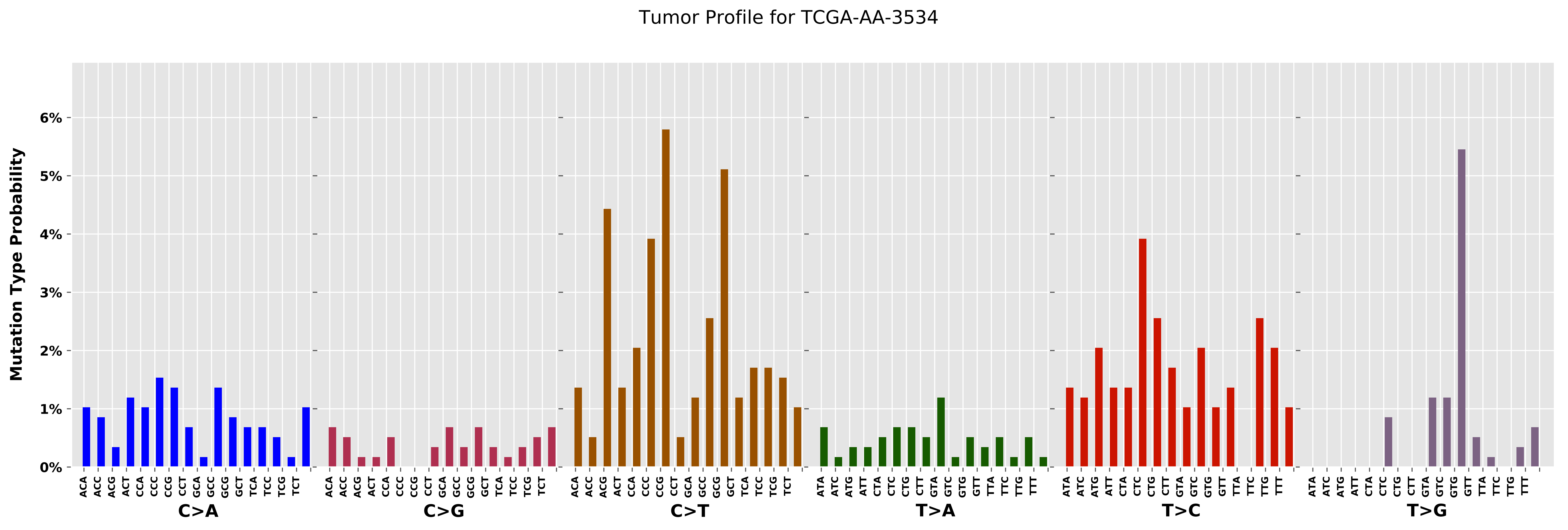

Tumor Profile for TCGA-AA-3534
Mutation Type Probability
0%
1%
2%
3%
4%
5%
6%
ACA ACC ACG ACT CCA CCC CCG CCT GCA GCC GCG GCT TCA TCC TCG TCT
C>A
C>G
C>T
ATA ATC ATG ATT CTA CTC CTG CTT GTA GTC GTG GTT TTA TTC TTG TTT
T>A
T>C
T>G

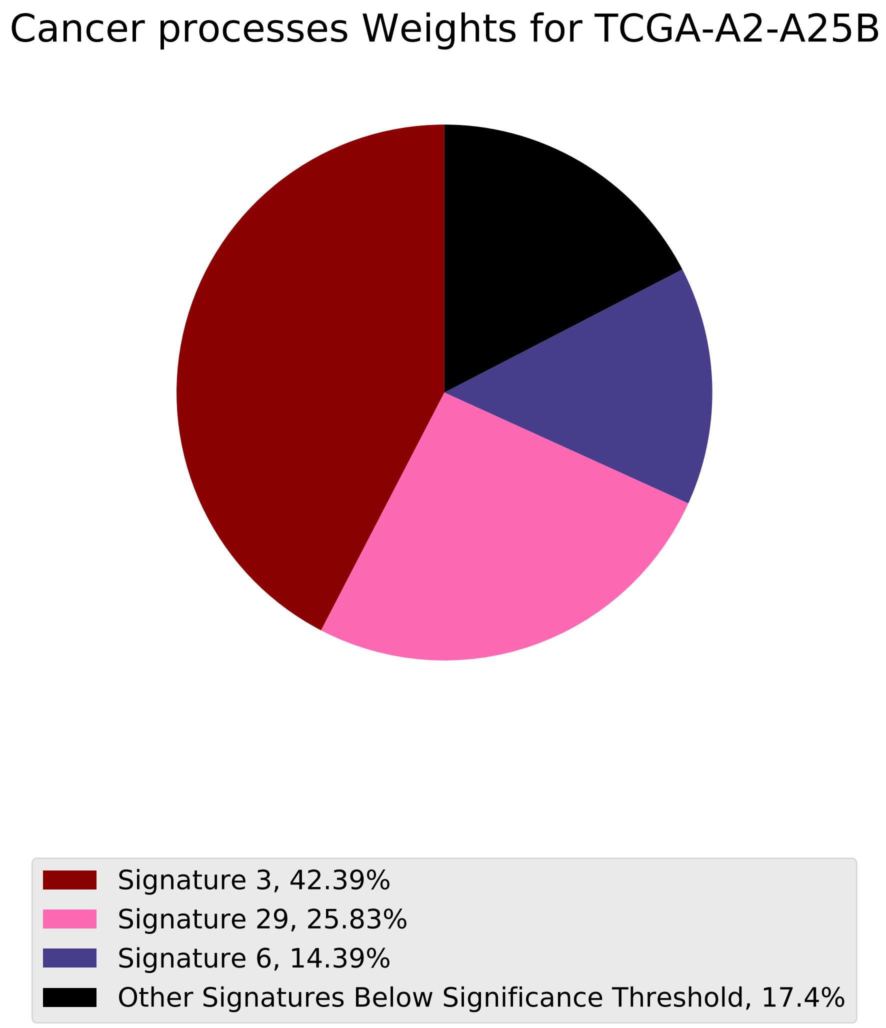
Cancer processes Weights for TCGA-A2-A25B
Signature 3, 42.39%
Signature 29, 25.83%
Signature 6, 14.39%
Other Signatures Below Significance Threshold, 17.4%

Tumor Profile for TCGA-A2-A25B

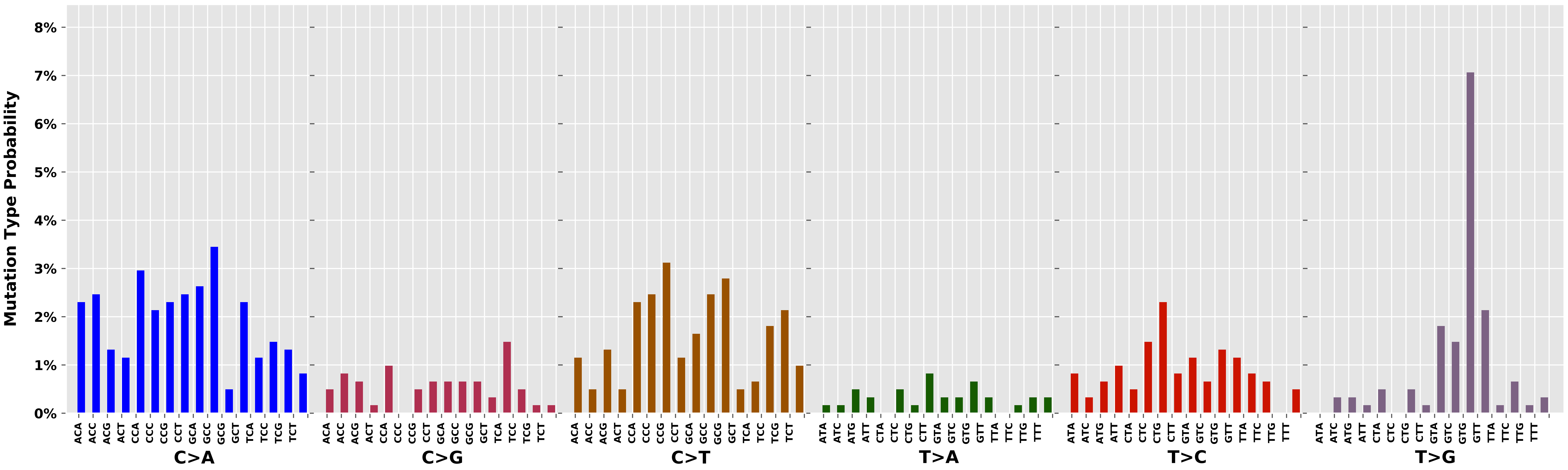

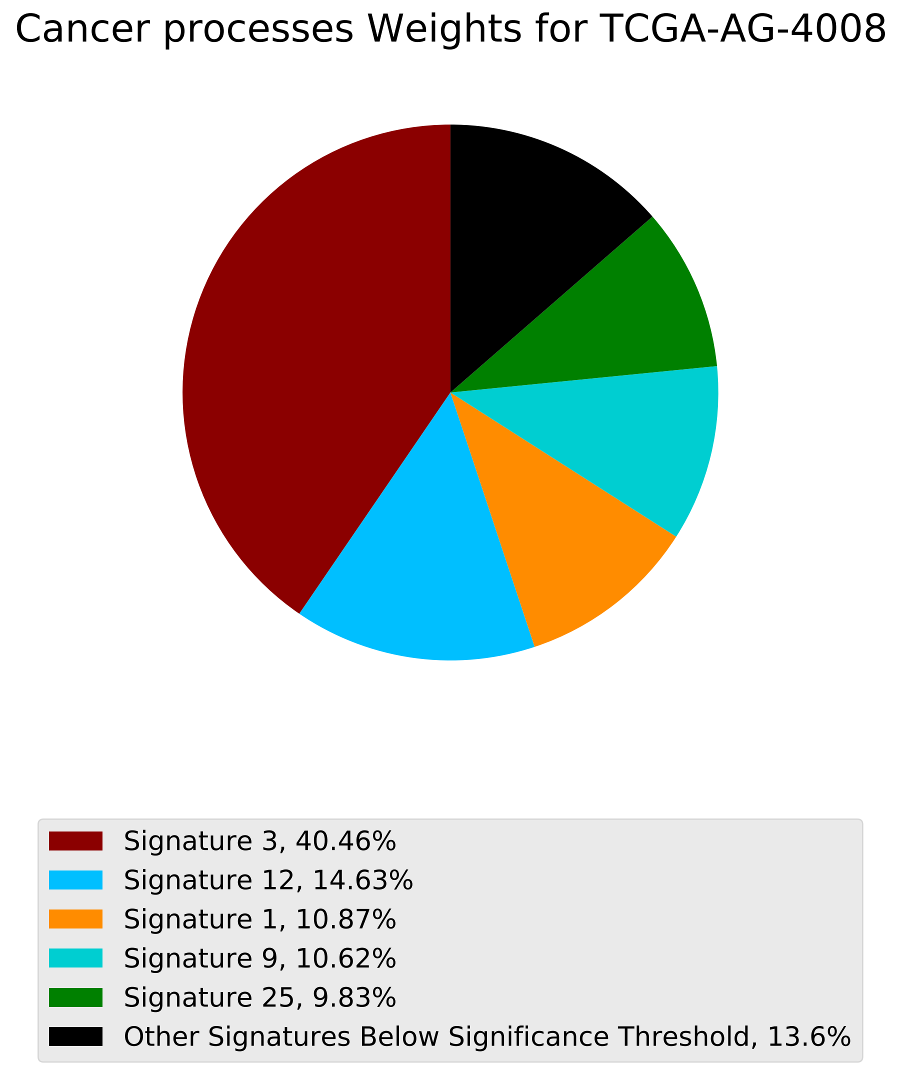
Cancer processes Weights for TCGA-AG-4008
Signature 3, 40.46%
Signature 12, 14.63%
Signature 1, 10.87%
Signature 9, 10.62%
Signature 25, 9.83%
Other Signatures Below Significance Threshold, 13.6%

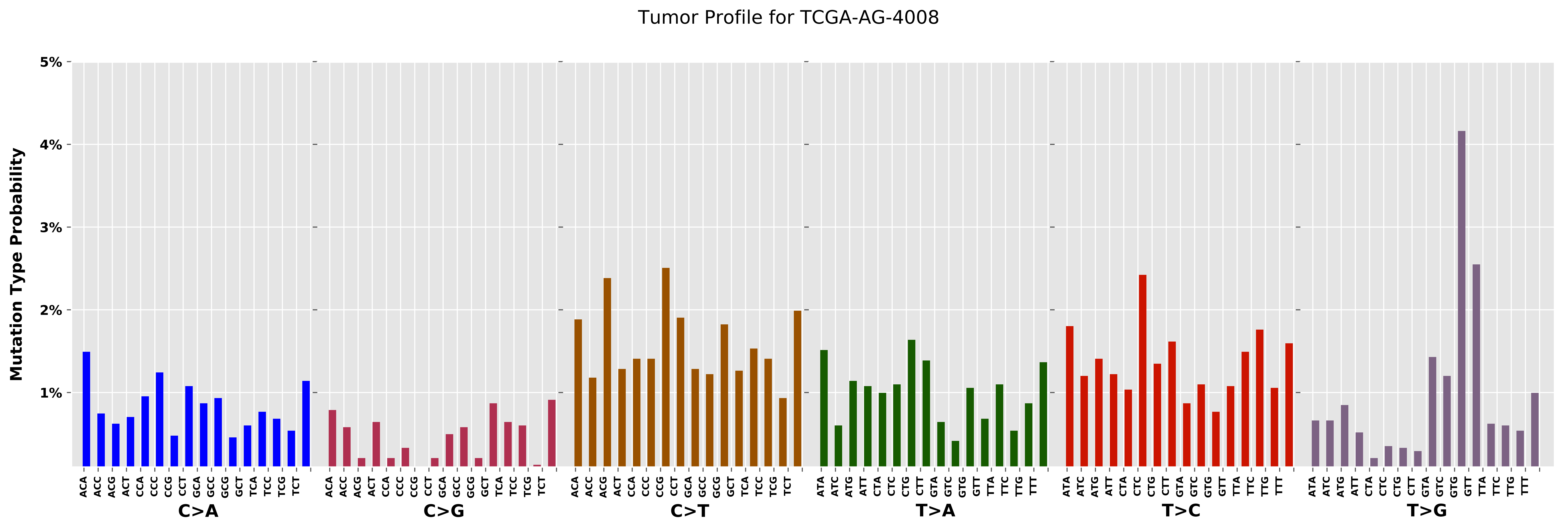

Tumor Profile for TCGA-AG-4008
Mutation Type Probability
5%
4%
3%
2%
1%
ACA ACC ACG ACT CCA CCC CCG CCT GCA GCC GCG GCT TCA TCC TCG TCT
C>A
C>G
C>T
ATA ATC ATG ATT CTA CTC CTG CTT GTA GTC GTG GTT TTA TTC TTG TTT
T>A
T>C
T>G

Cancer processes Weights for TCGA-A7-A26G

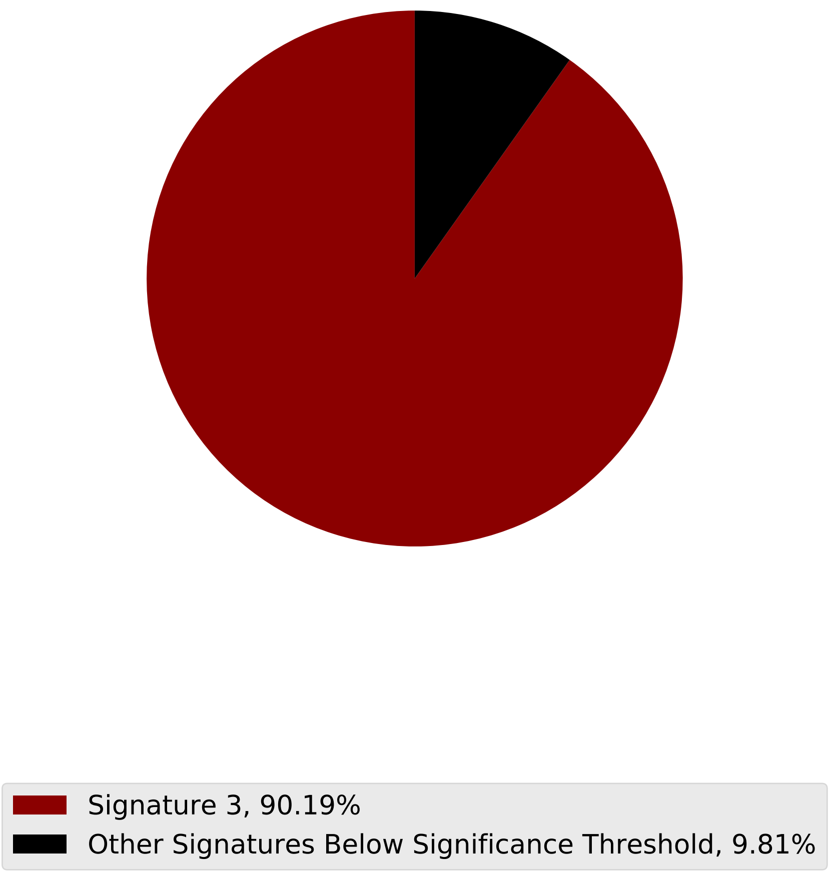

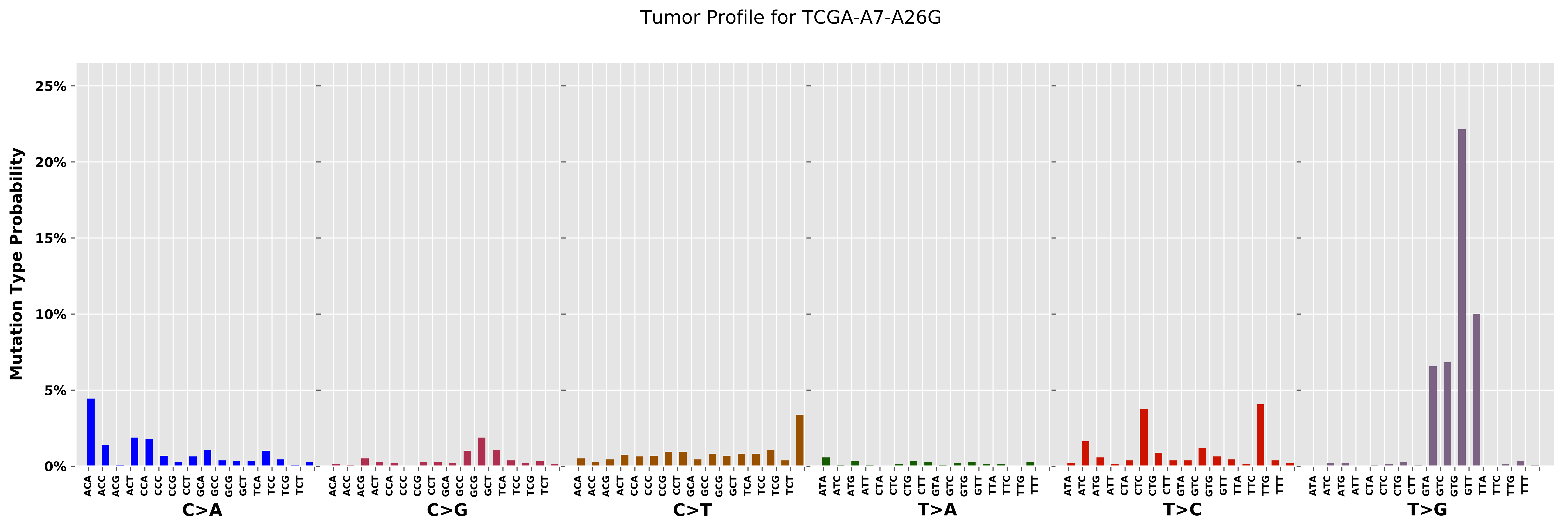

Tumor Profile for TCGA-A7-A26G
Mutation Type Probability
25%
20%
15%
10%
5%
0%
C>A
C>G
C>T
T>A
T>C
T>G
ACA ACC ACG ACT CCA CCC CCG CCT GCA GCC GCG GCT TCA TCC TCG TCT
ATA ATC ATG ATT CTA CTC CTG CTT GTA GTC GTG GTT TTA TTC TTG TTT

Cancer processes Weights for TCGA-AR-A24Z

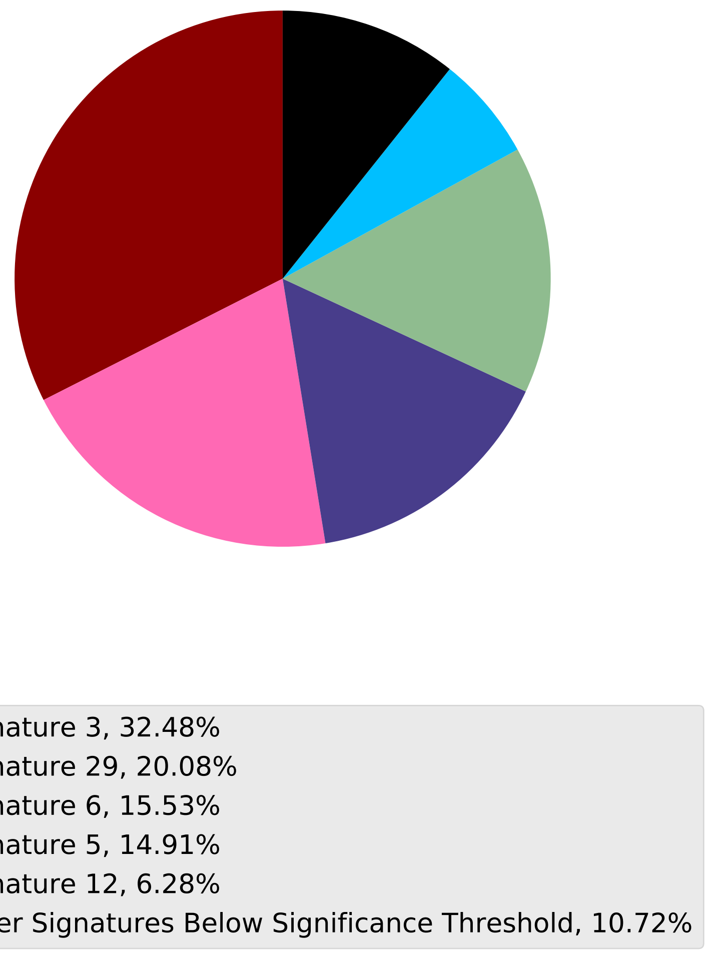

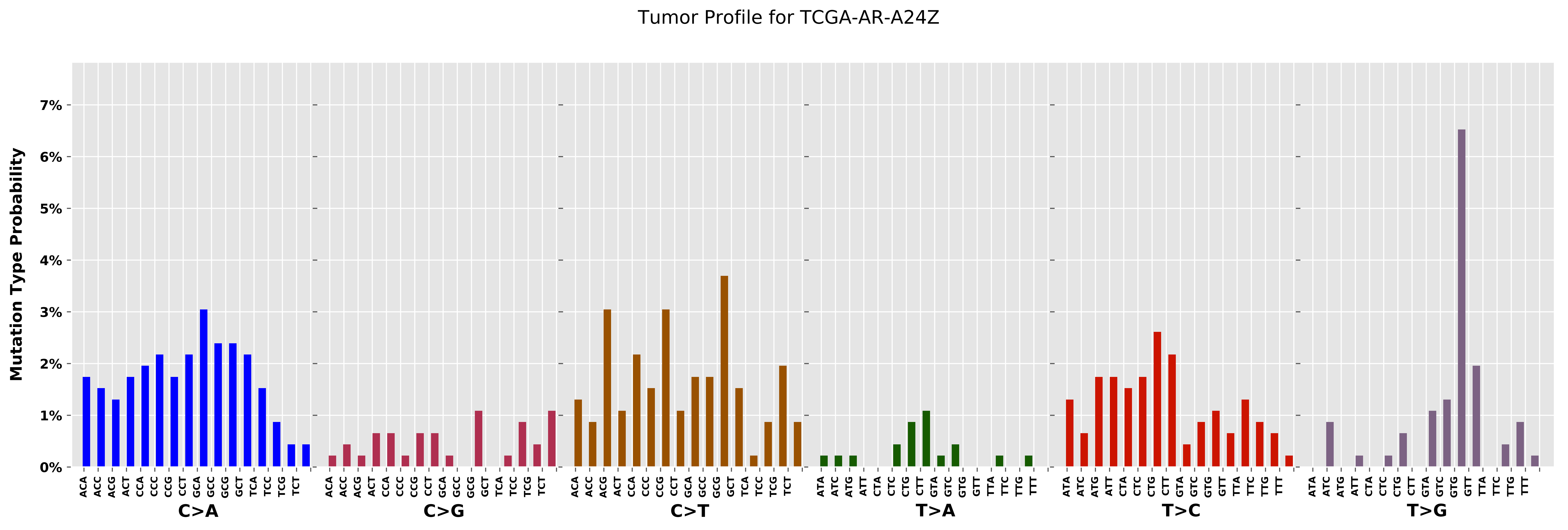
Tumor Profile for TCGA-AR-A24Z
Mutation Type Probability
0%
1%
2%
3%
4%
5%
6%
7%
C>A
C>G
C>T
T>A
T>C
T>G

Cancer processes Weights for TCGA-A2-A04P

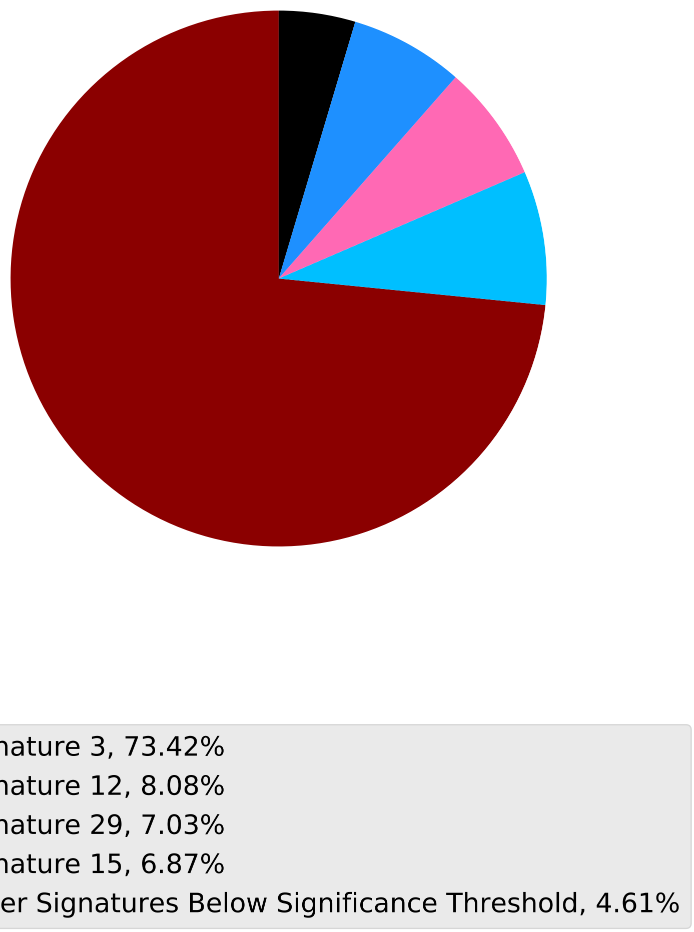

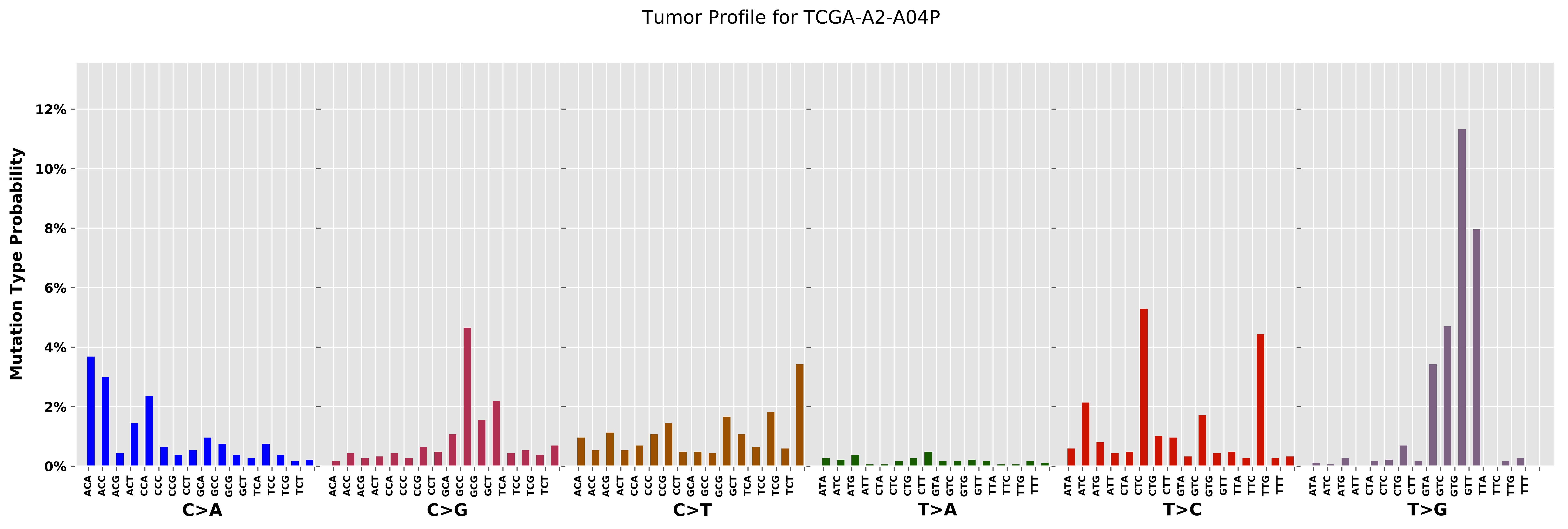
Tumor Profile for TCGA-A2-A04P
Mutation Type Probability
0%
2%
4%
6%
8%
10%
12%
C>A
C>G
C>T
T>A
T>C
T>G
ACA ACC ACG ACT CCA CCC CCG CCT GCA GCC GCG GCT TCA TCC TCG TCT
ATA ATC ATG ATT CTA CTC CTG CTT GTA GTC GTG GTT TTA TTC TTG TTT

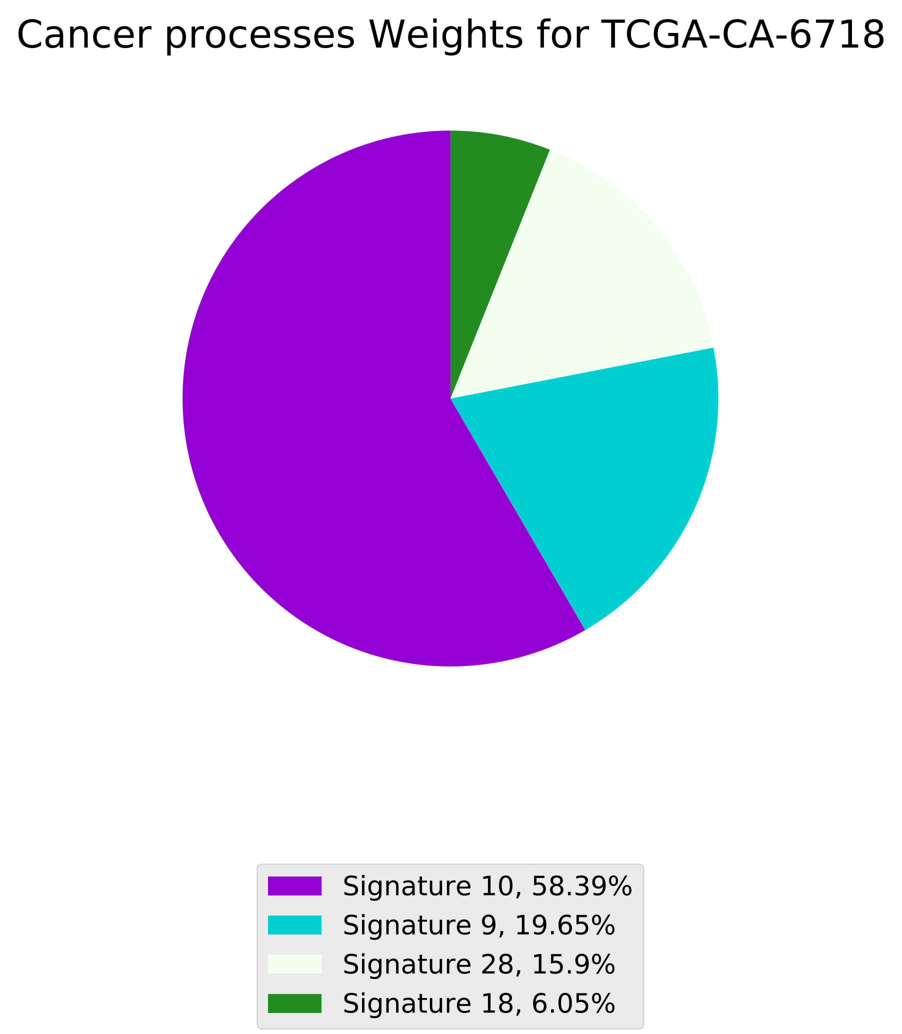
Cancer processes Weights for TCGA-CA-6718
Signature 10, 58.39%
Signature 9, 19.65%
Signature 28, 15.9%
Signature 18, 6.05%

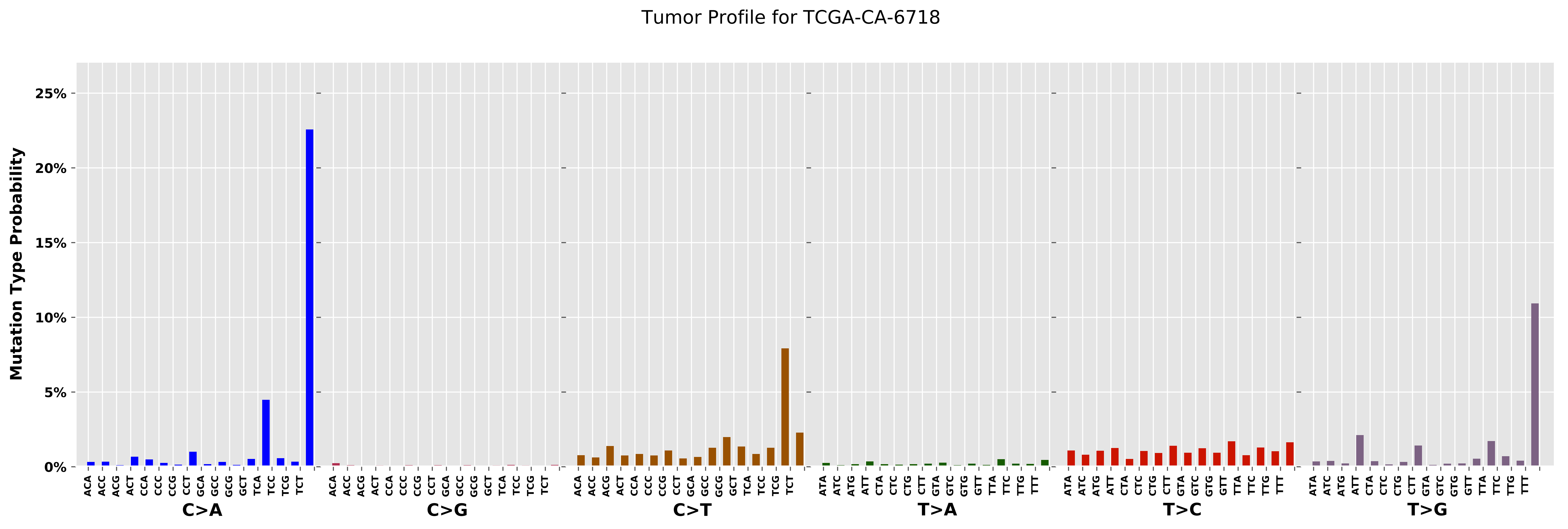
Tumor Profile for TCGA-CA-6718
Mutation Type Probability
25%
20%
15%
10%
5%
0%
ACA ACC ACG ACT CCA CCC CCG CCT GCA GCC GCG GCT TCA TCC TCG TCT
C>A
ACA ACC ACG ACT CCA CCC CCG CCT GCA GCC GCG GCT TCA TCC TCG TCT
C>G
ACA ACC ACG ACT CCA CCC CCG CCT GCA GCC GCG GCT TCA TCC TCG TCT
C>T
ATA ATC ATG ATT CTA CTC CTG CTT GTA GTC GTG GTT TTA TTC TTG TTT
T>A
ATA ATC ATG ATT CTA CTC CTG CTT GTA GTC GTG GTT TTA TTC TTG TTT
T>C
ATA ATC ATG ATT CTA CTC CTG CTT GTA GTC GTG GTT TTA TTC TTG TTT
T>G

Cancer processes Weights for TCGA-BH-A0AV

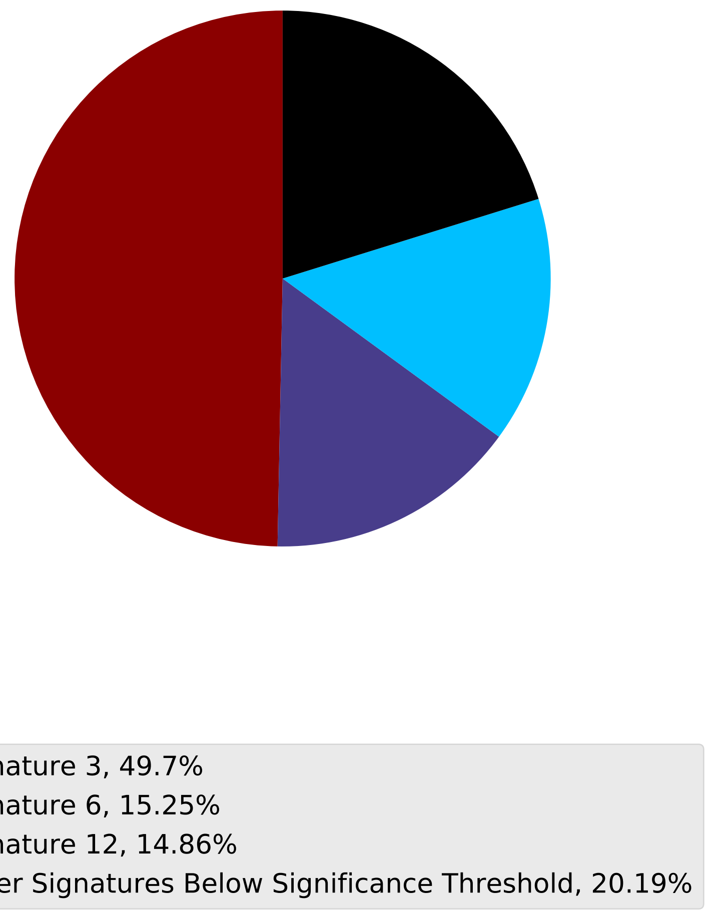

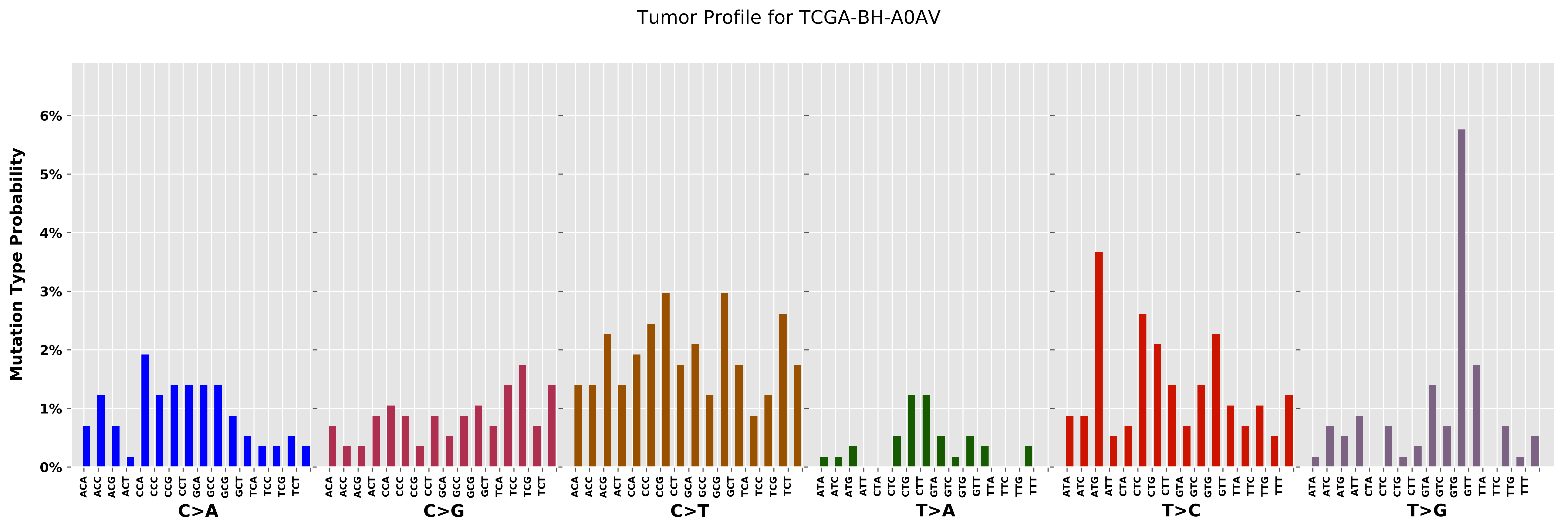

Tumor Profile for TCGA-BH-A0AV
Mutation Type Probability
0%
1%
2%
3%
4%
5%
6%
C>A
C>G
C>T
T>A
T>C
T>G
ACA ACC ACG ACT CCA CCC CCG CCT GCA GCC GCG GCT TCA TCC TCG TCT
ATA ATC ATG ATT CTA CTC CTG CTT GTA GTC GTG GTT TTA TTC TTG TTT

Cancer processes Weights for TCGA-A2-A0CM

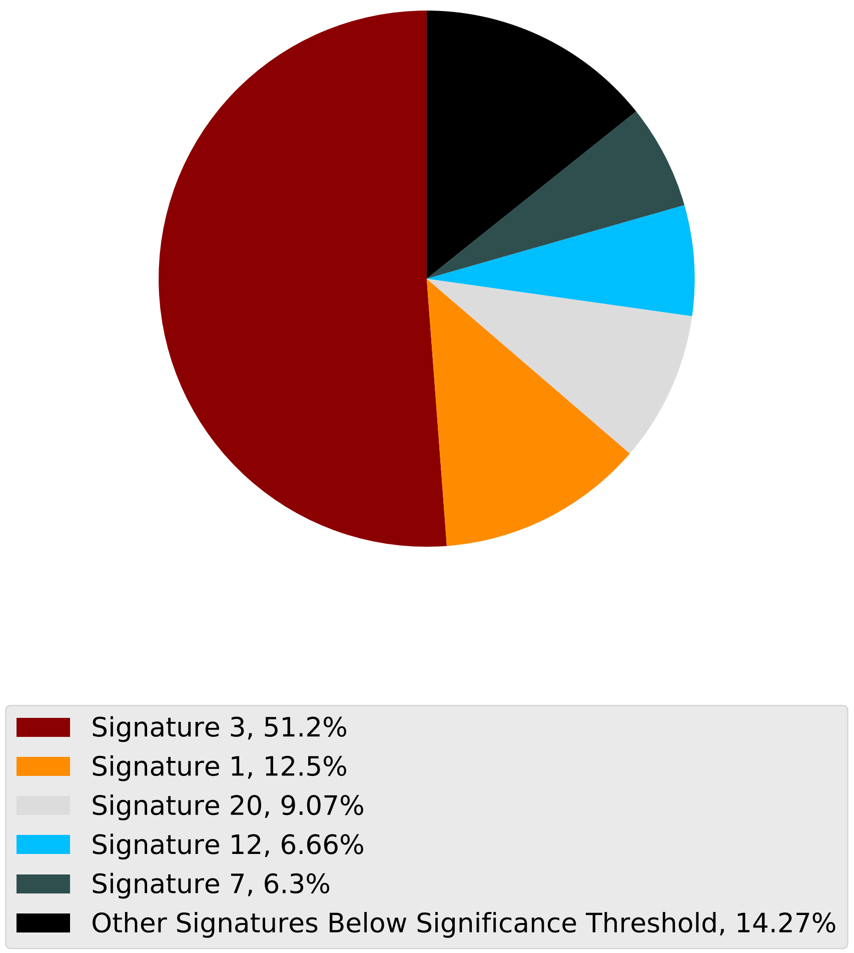

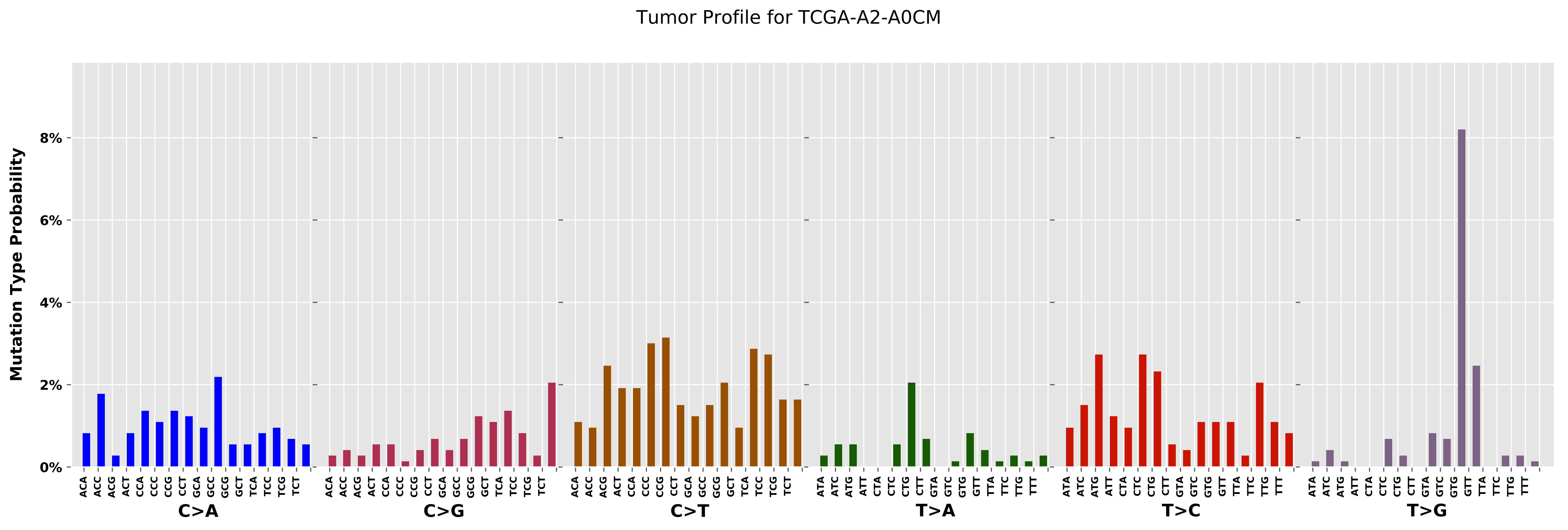

Tumor Profile for TCGA-A2-A0CM
Mutation Type Probability
0%
2%
4%
6%
8%
C>A
C>G
C>T
T>A
T>C
T>G

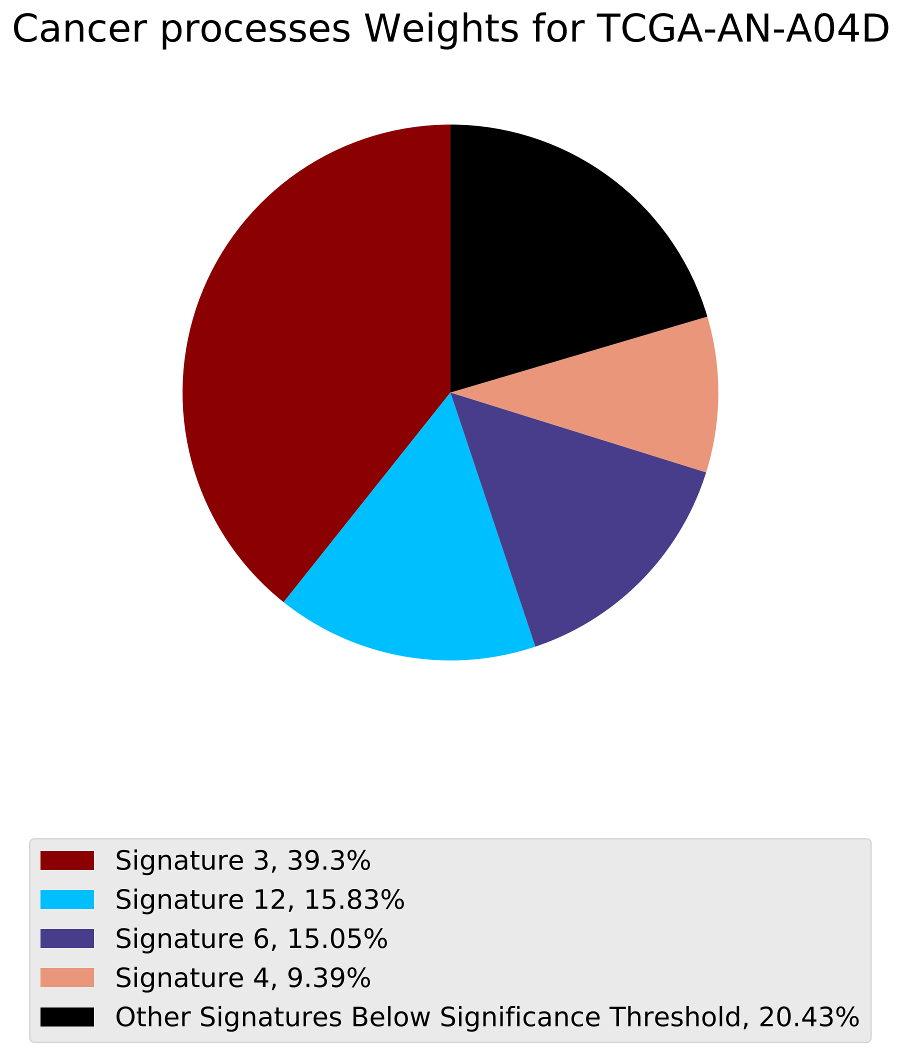
Cancer processes Weights for TCGA-AN-A04D
Signature 3, 39.3%
Signature 12, 15.83%
Signature 6, 15.05%
Signature 4, 9.39%
Other Signatures Below Significance Threshold, 20.43%

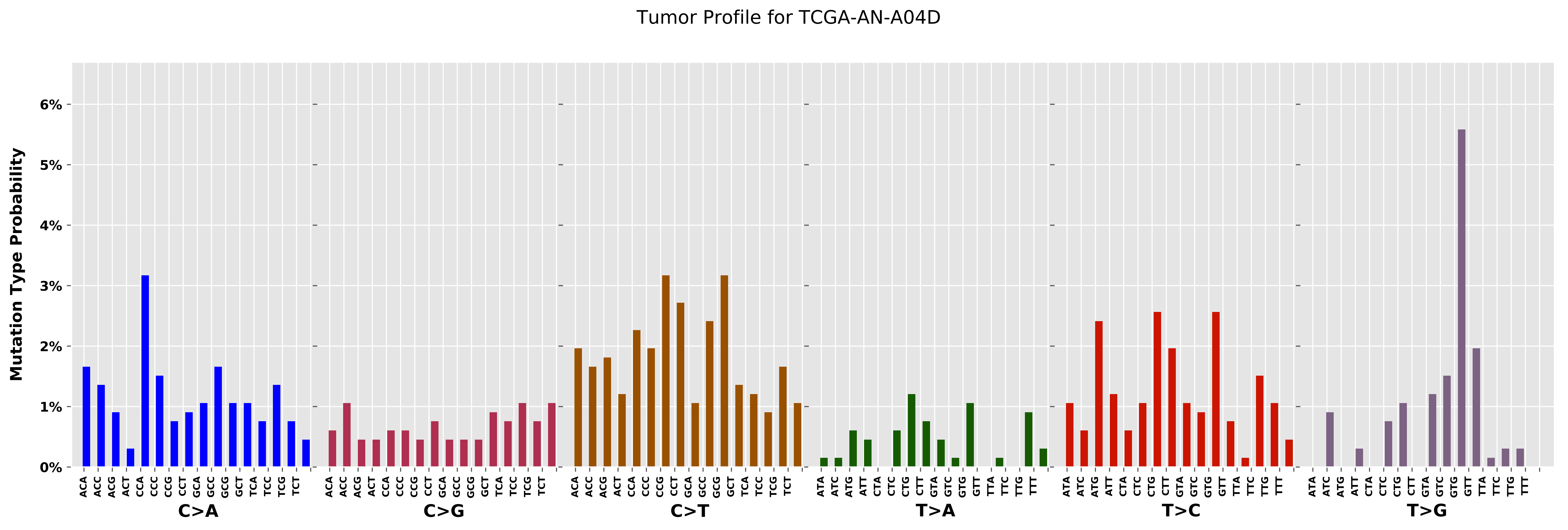

Tumor Profile for TCGA-AN-A04D
Mutation Type Probability
0%
1%
2%
3%
4%
5%
6%
C>A
C>G
C>T
T>A
T>C
T>G

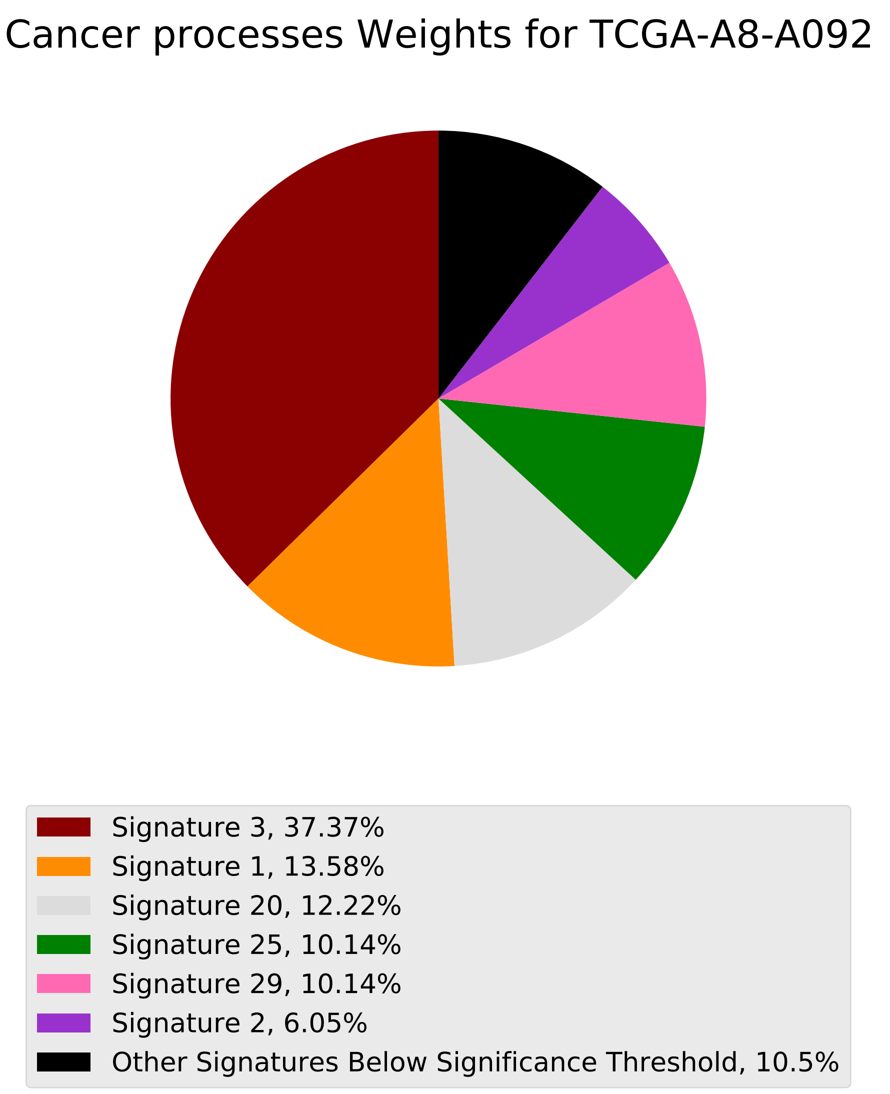
Cancer processes Weights for TCGA-A8-A092
Signature 3, 37.37%
Signature 1, 13.58%
Signature 20, 12.22%
Signature 25, 10.14%
Signature 29, 10.14%
Signature 2, 6.05%
Other Signatures Below Significance Threshold, 10.5%

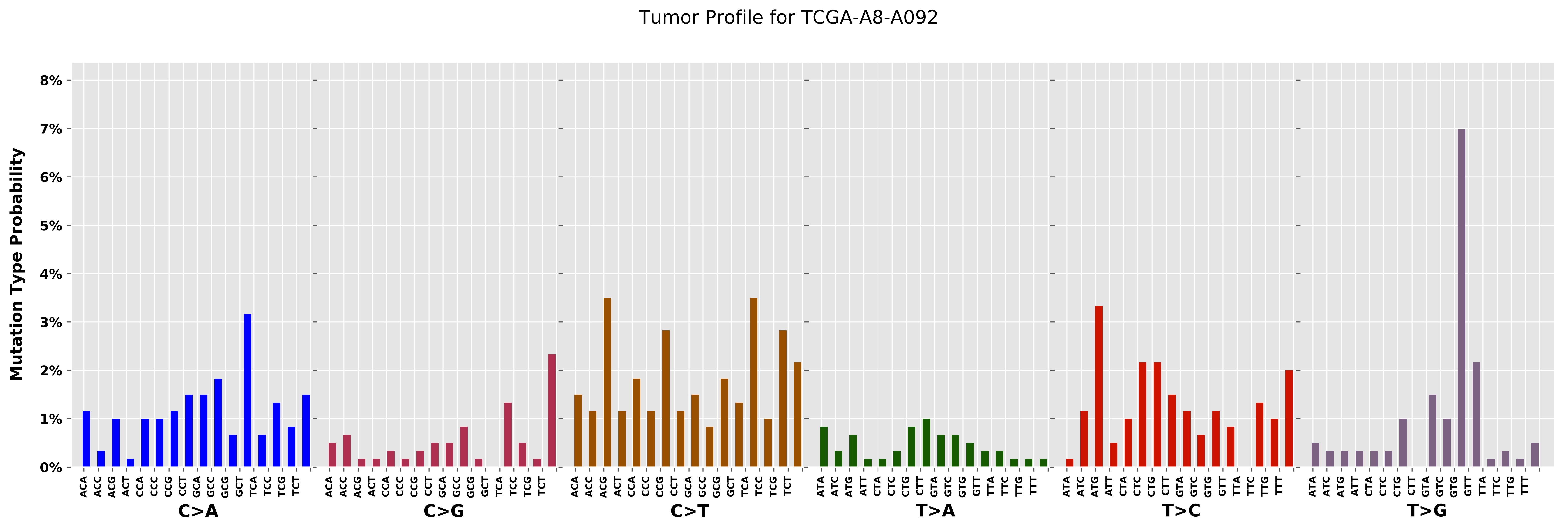
Tumor Profile for TCGA-A8-A092
Mutation Type Probability
8%
7%
6%
5%
4%
3%
2%
1%
0%
ACA ACC ACG ACT CCA CCC CCG CCT GCA GCC GCG GCT TCA TCC TCG TCT
C>A
C>G
C>T
ATA ATC ATG ATT CTA CTC CTG CTT GTA GTC GTG GTT TTA TTC TTG TTT
T>A
T>C
T>G

Cancer processes Weights for TCGA-A2-A04X

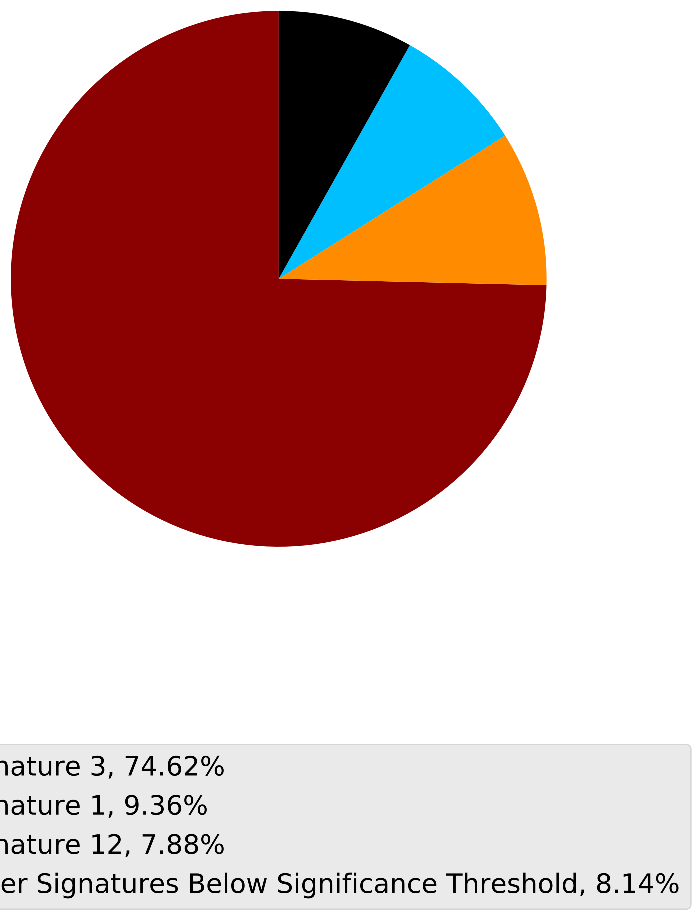

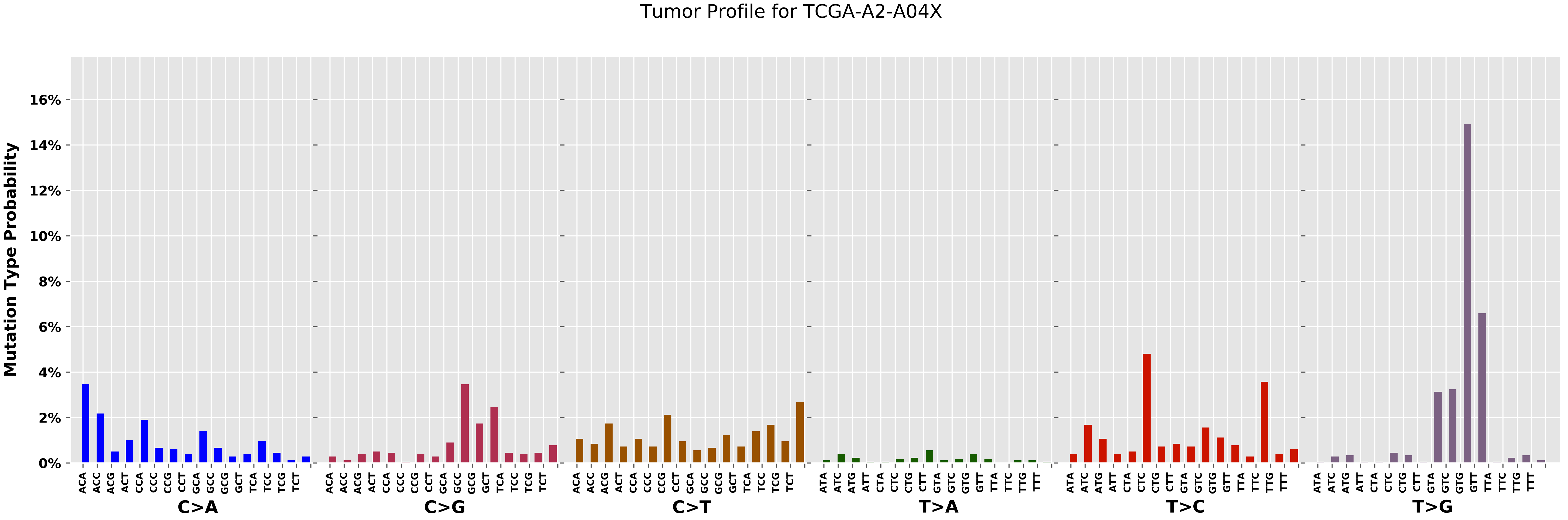

Tumor Profile for TCGA-A2-A04X
Mutation Type Probability
0%
2%
4%
6%
8%
10%
12%
14%
16%
C>A
C>G
C>T
T>A
T>C
T>G

Cancer processes Weights for TCGA-A2-A0D2

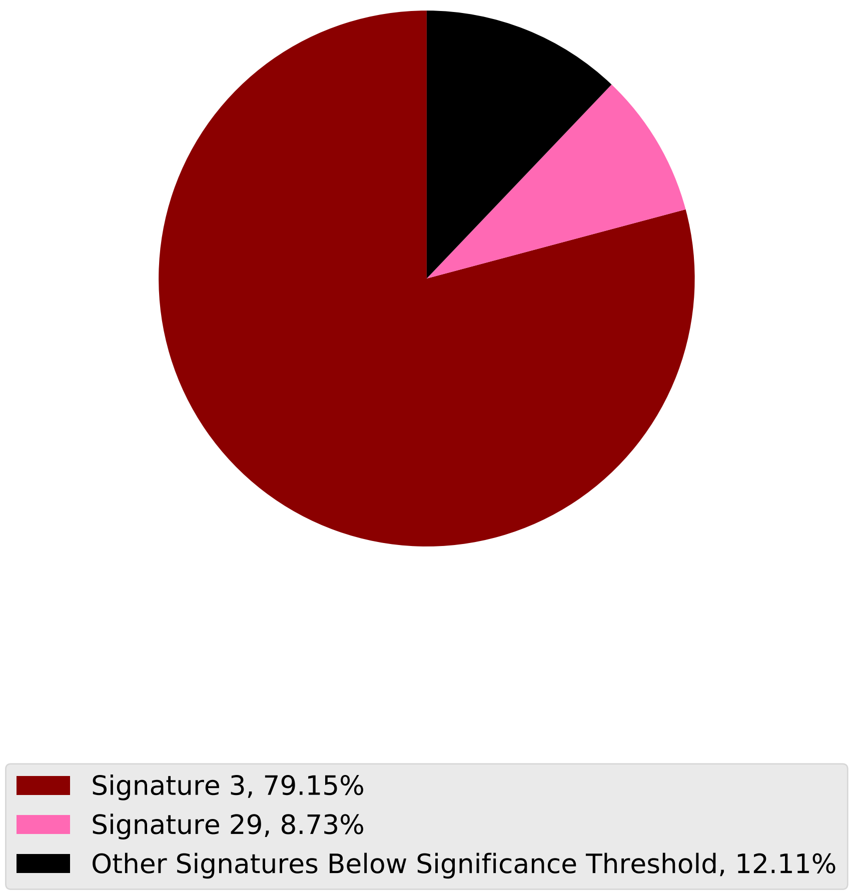

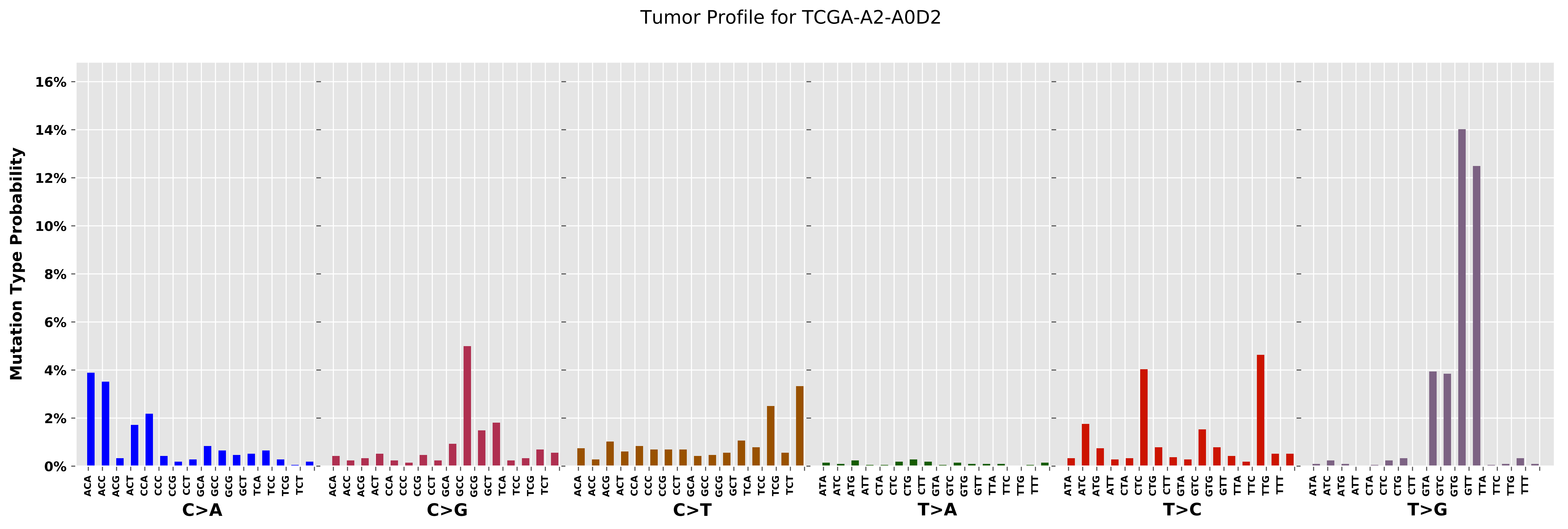

Tumor Profile for TCGA-A2-A0D2
Mutation Type Probability
0%
2%
4%
6%
8%
10%
12%
14%
16%
C>A
C>G
C>T
T>A
T>C
T>G
ACA ACC ACG ACT CCA CCC CCG CCT GCA GCC GCG GCT TCA TCC TCG TCT
ATA ATC ATG ATT CTA CTC CTG CTT GTA GTC GTG GTT TTA TTC TTG TTT

Cancer processes Weights for TCGA-B6-A0RU

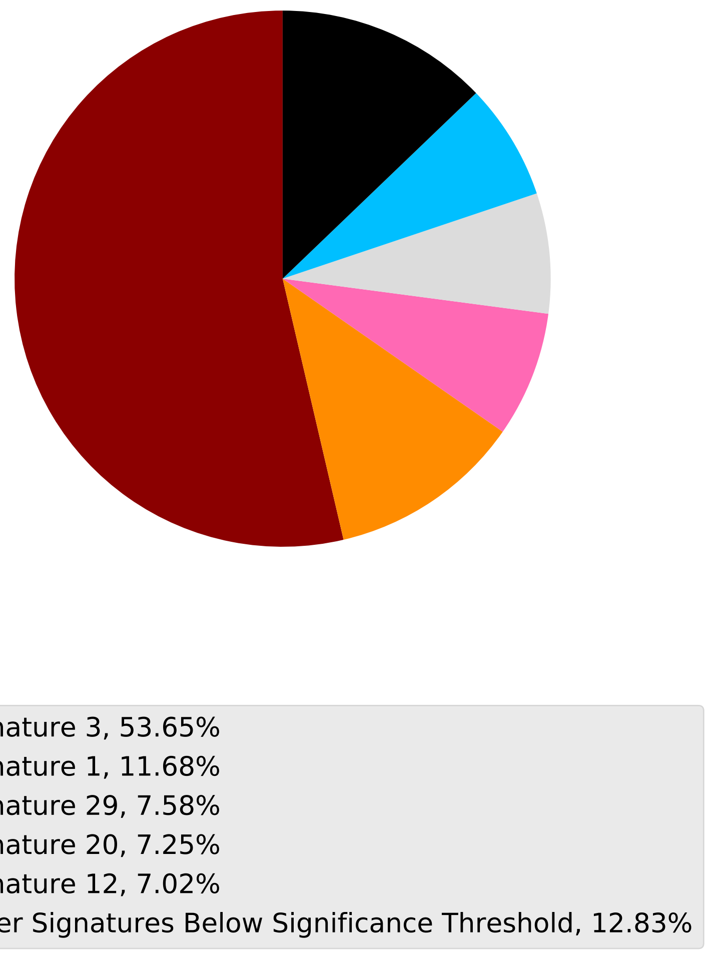

Tumor Profile for TCGA-B6-A0RU

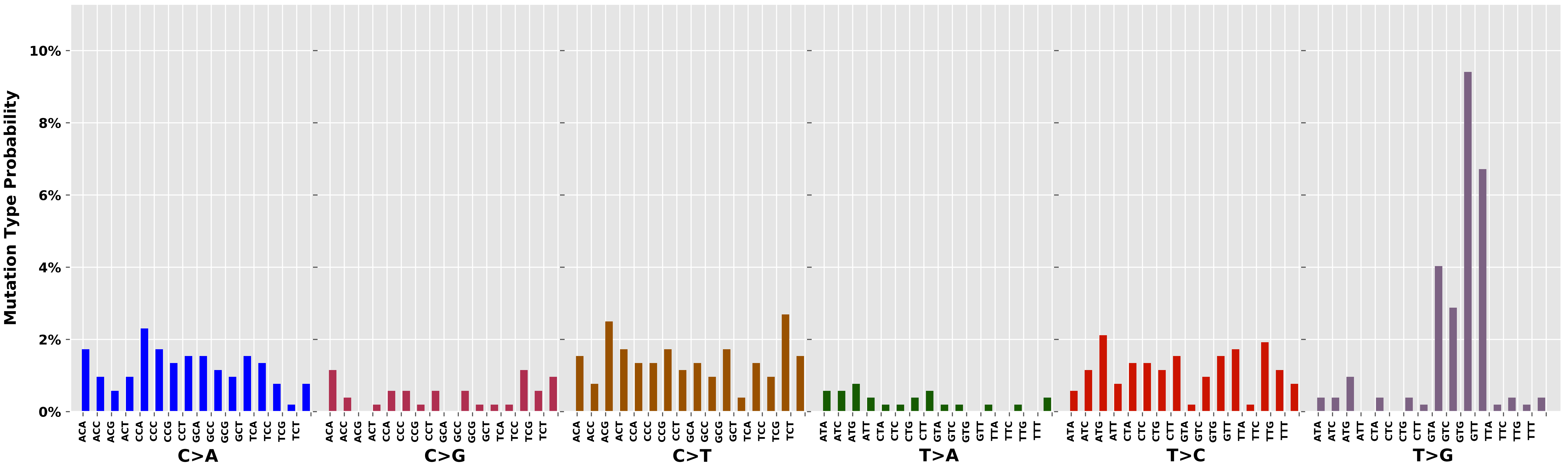
