## Additional File 4 for "pyCancerSig: subclassifying human cancer with comprehensive single nucleotide, structural and microsatellite mutational signature deconstruction from whole genome sequencing"

Cancer processes Weights for TCGA-E2-A14X

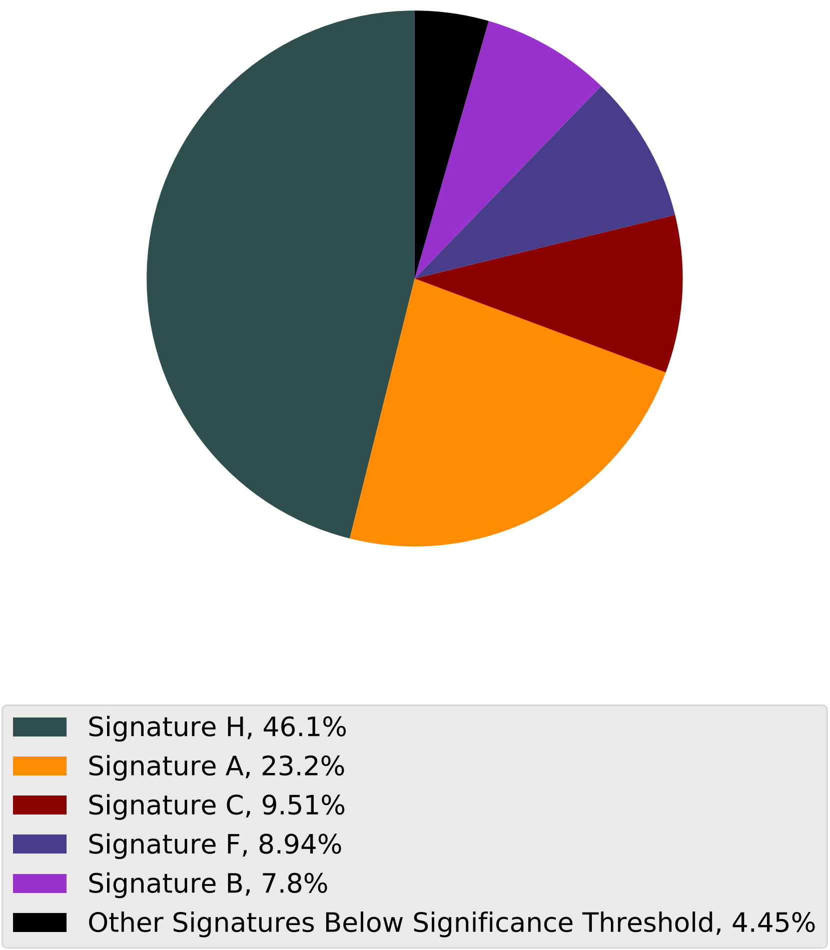

### Tumor Profile for TCGA-E2-A14X

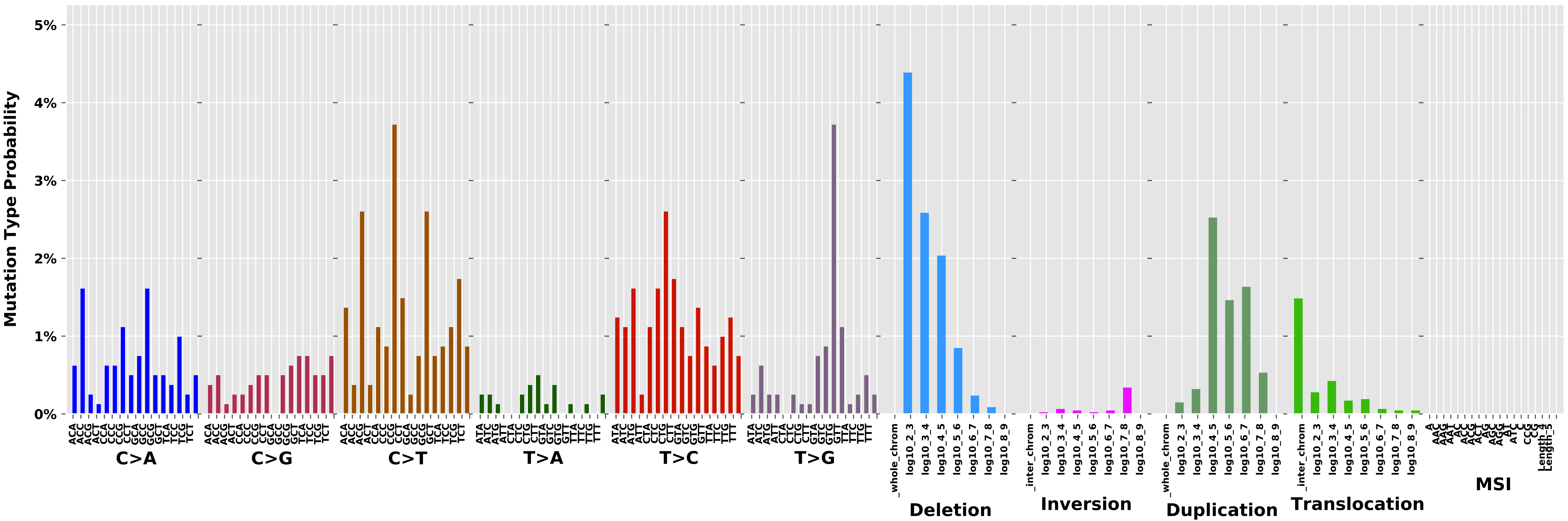

Cancer processes Weights for TCGA-A8-A08L

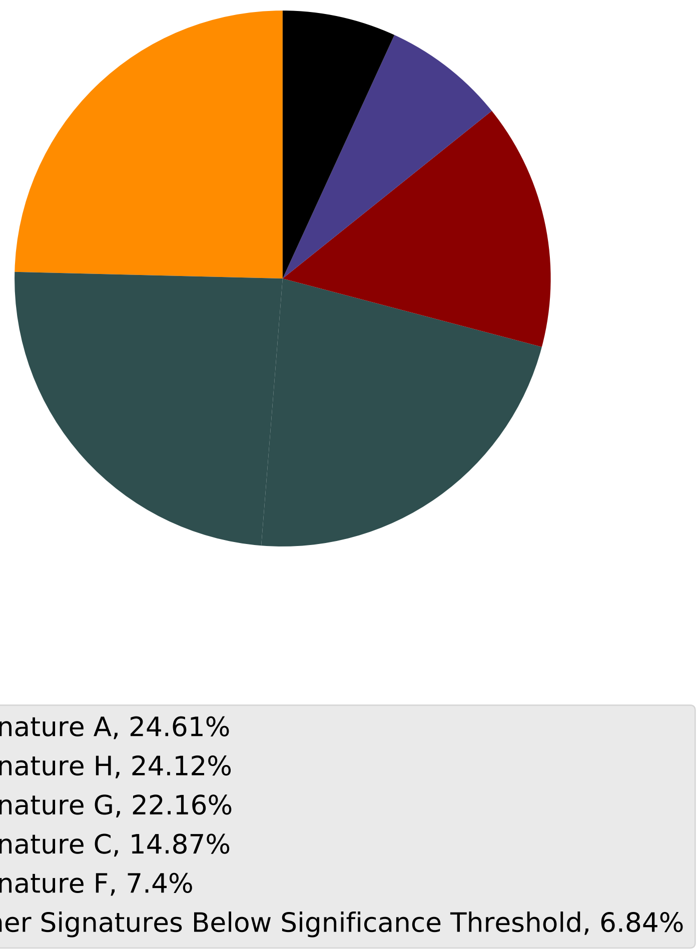

Tumor Profile for TCGA-A8-A08L

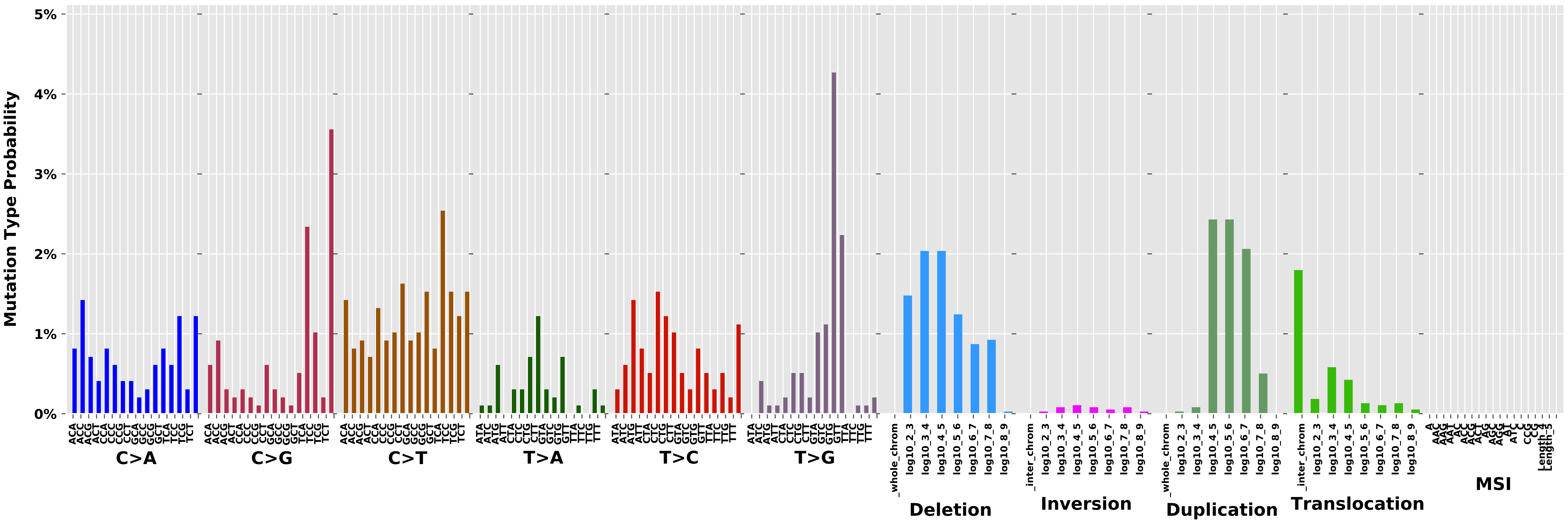

Cancer processes Weights for TCGA-AO-A12H

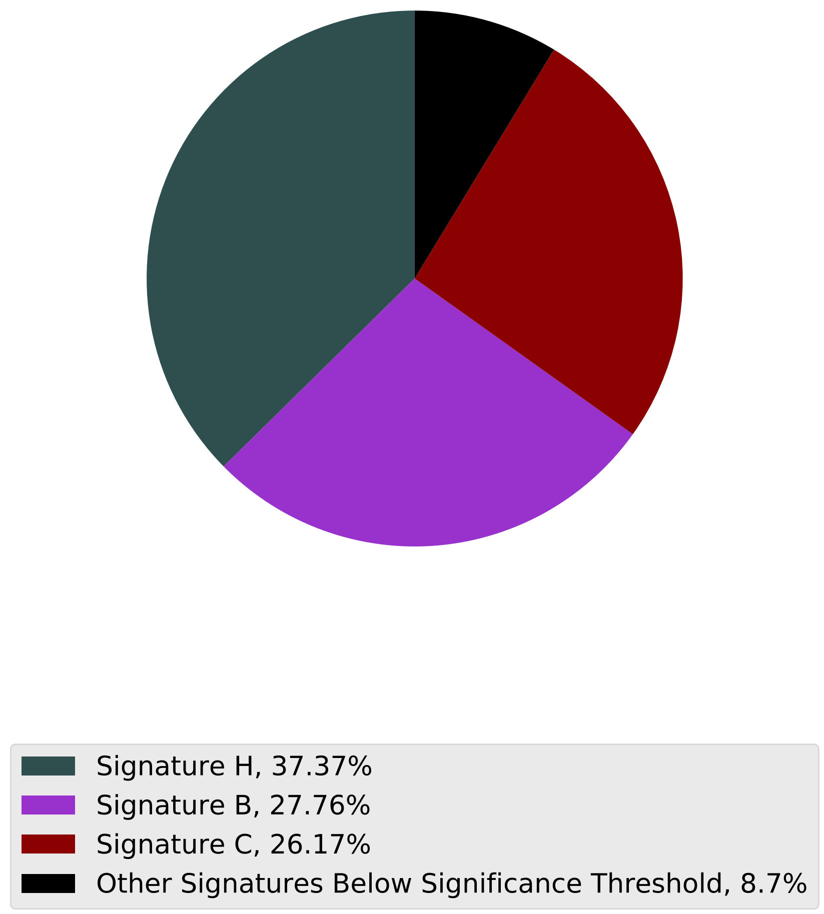

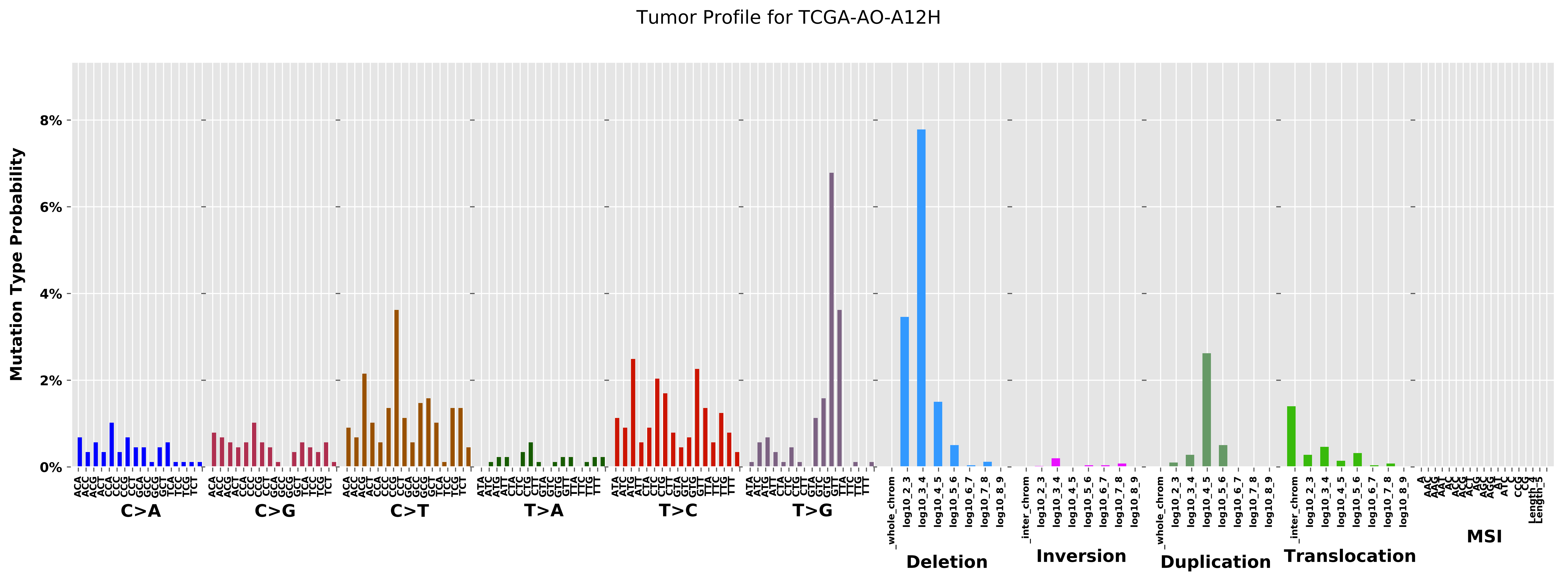

Tumor Profile for TCGA-AO-A12H
Mutation Type Probability
8%
6%
4%
2%
0%
C>A
C>G
C>T
T>A
T>C
T>G
Deletion
Inversion
Duplication
Translocation
MSI

#### Cancer processes Weights for TCGA-EW-A3U0

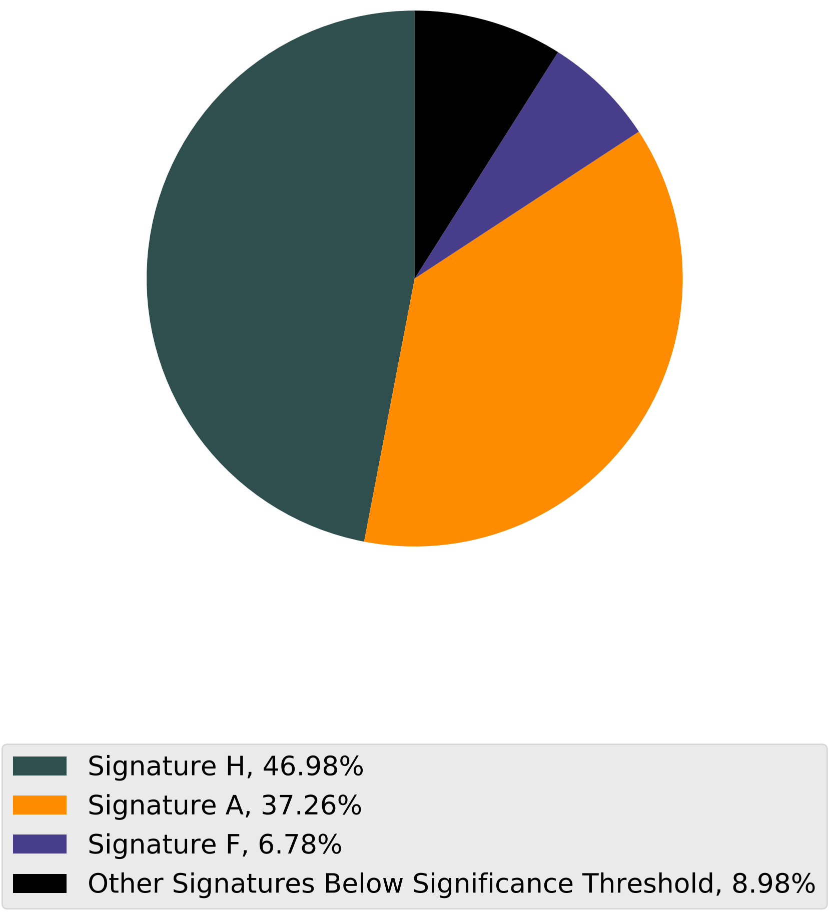

Tumor Profile for TCGA-EW-A3U0

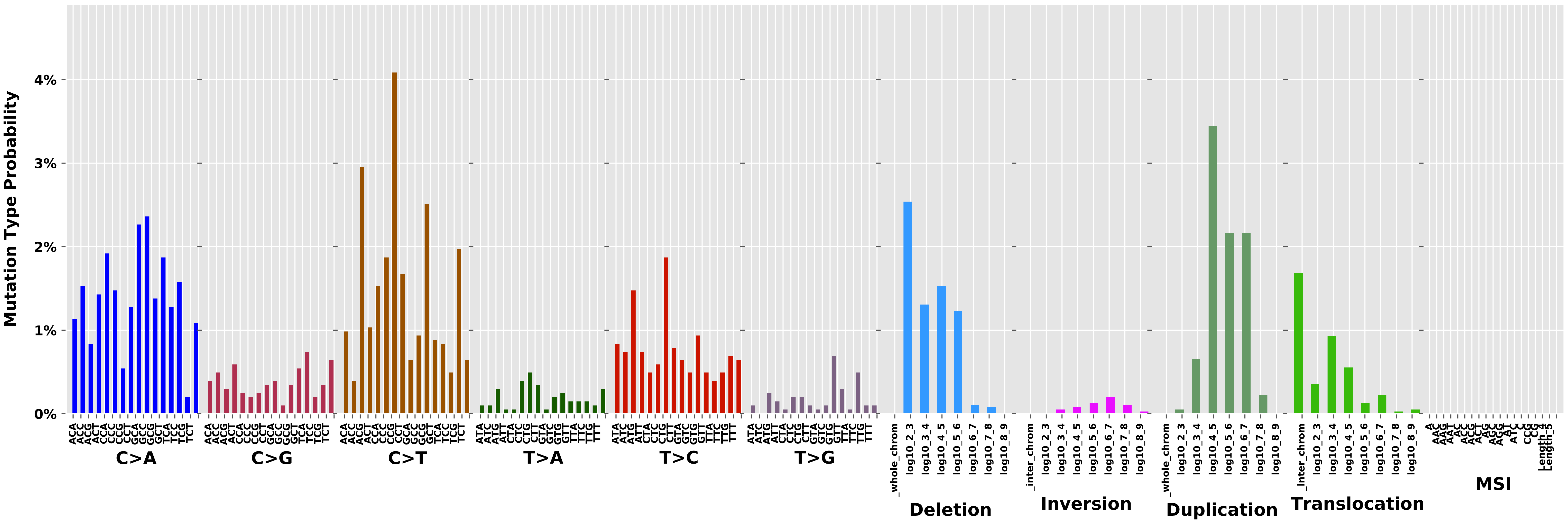

Cancer processes Weights for TCGA-BH-A1FC

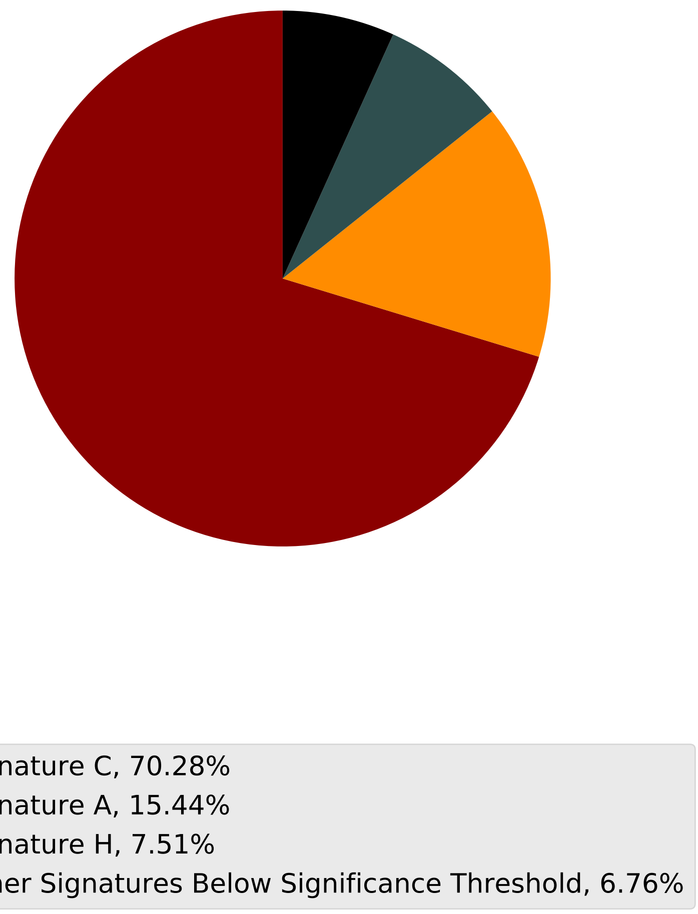

Tumor Profile for TCGA-BH-A1FC

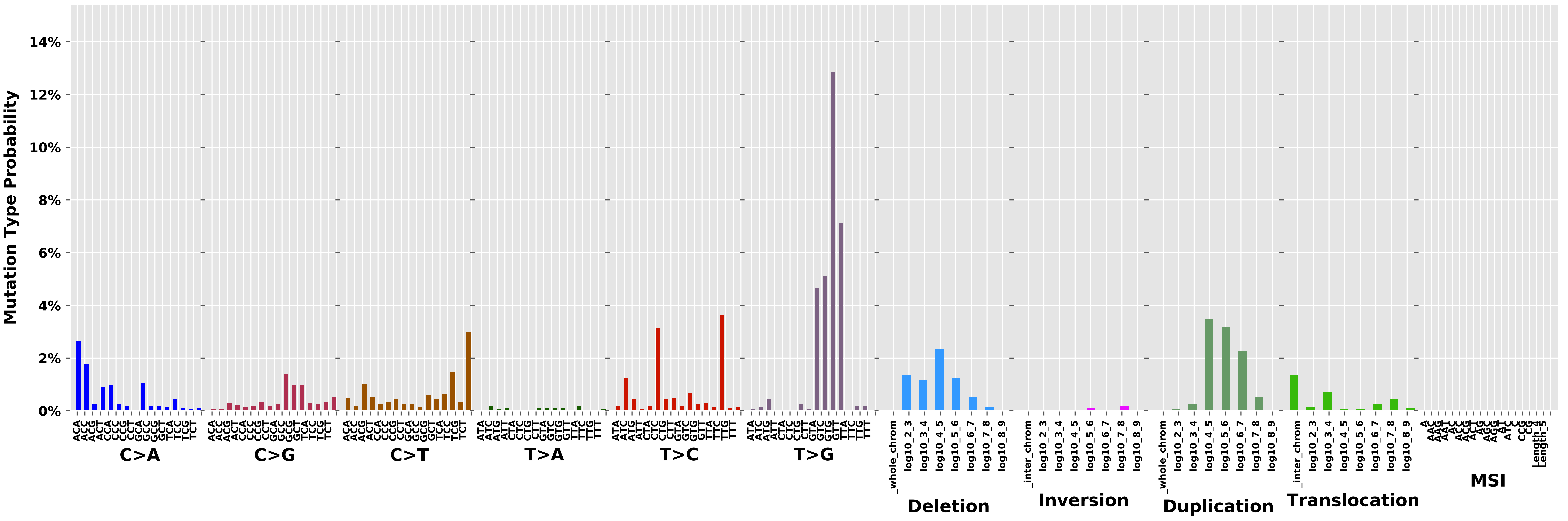

Cancer processes Weights for TCGA-AF-2691

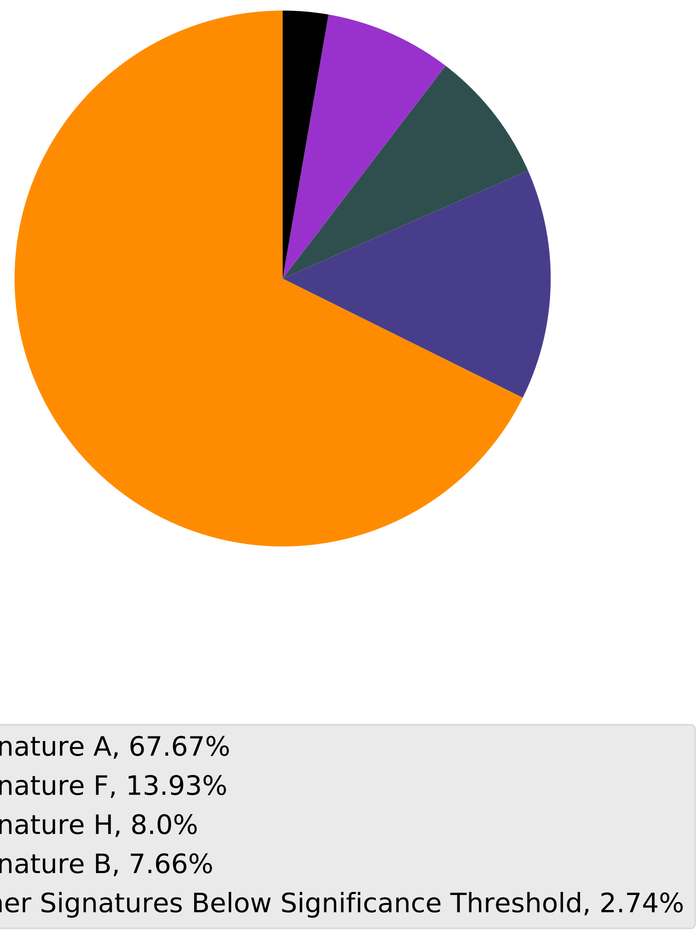

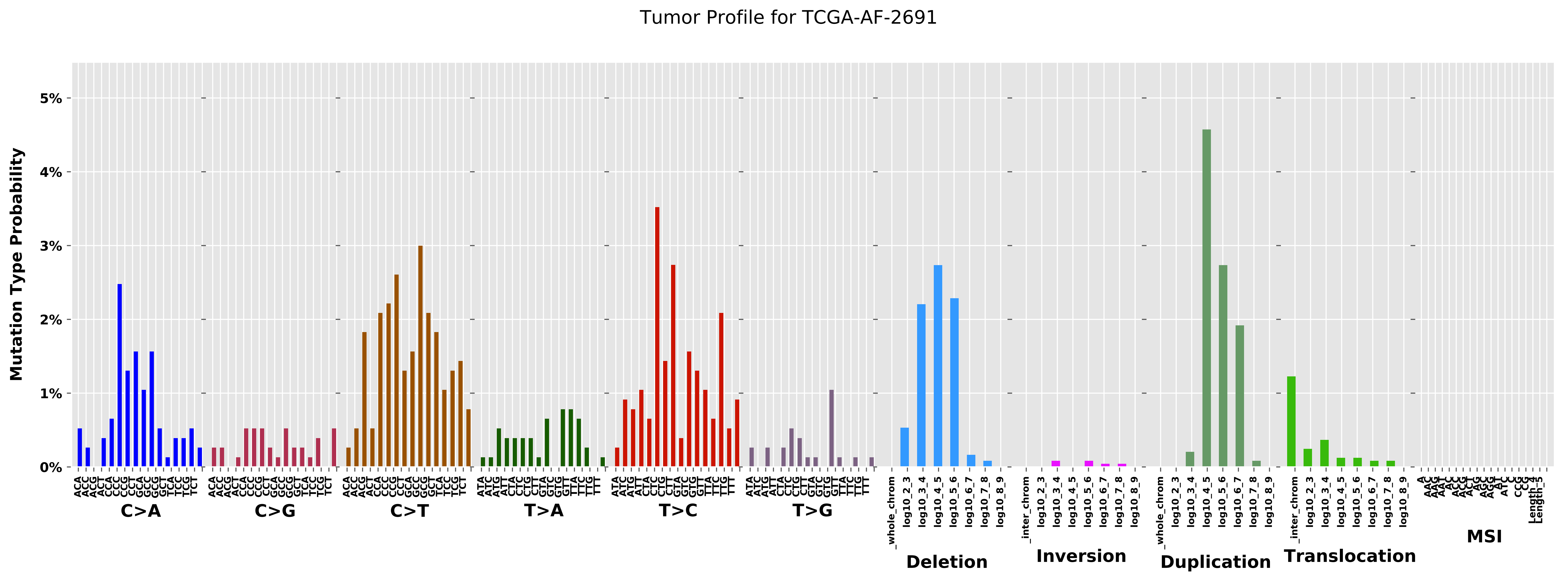
Tumor Profile for TCGA-AF-2691
Mutation Type Probability
0%
1%
2%
3%
4%
5%
C>A
C>G
C>T
T>A
T>C
T>G
Deletion
Inversion
Duplication
Translocation
MSI

Cancer processes Weights for TCGA-AF-3913

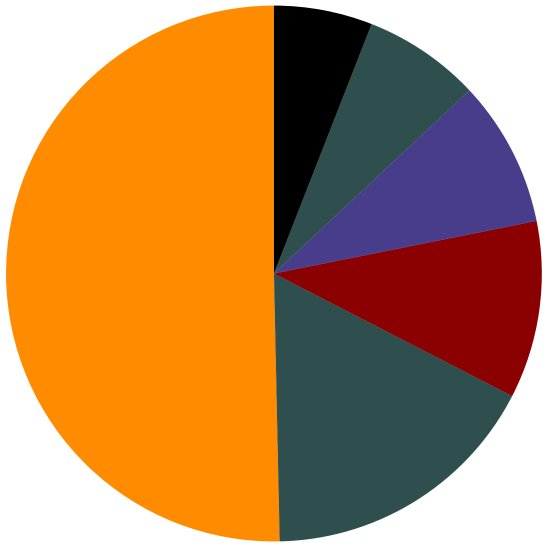

- Signature A, 50.34%
- Signature H, 17.09%
- Signature C, 10.71%
- Signature F, 8.82%
- Signature G, 7.09%
- Other Signatures Below Significance Threshold, 5.94%

Tumor Profile for TCGA-AF-3913

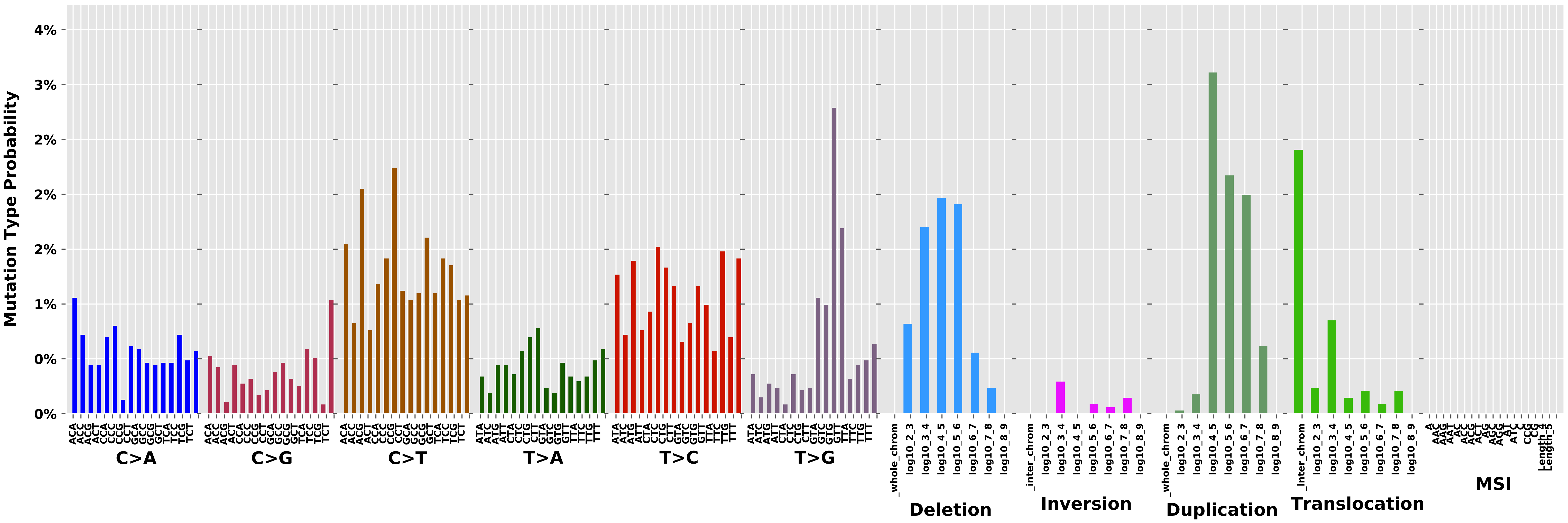

Cancer processes Weights for TCGA-AA-3977

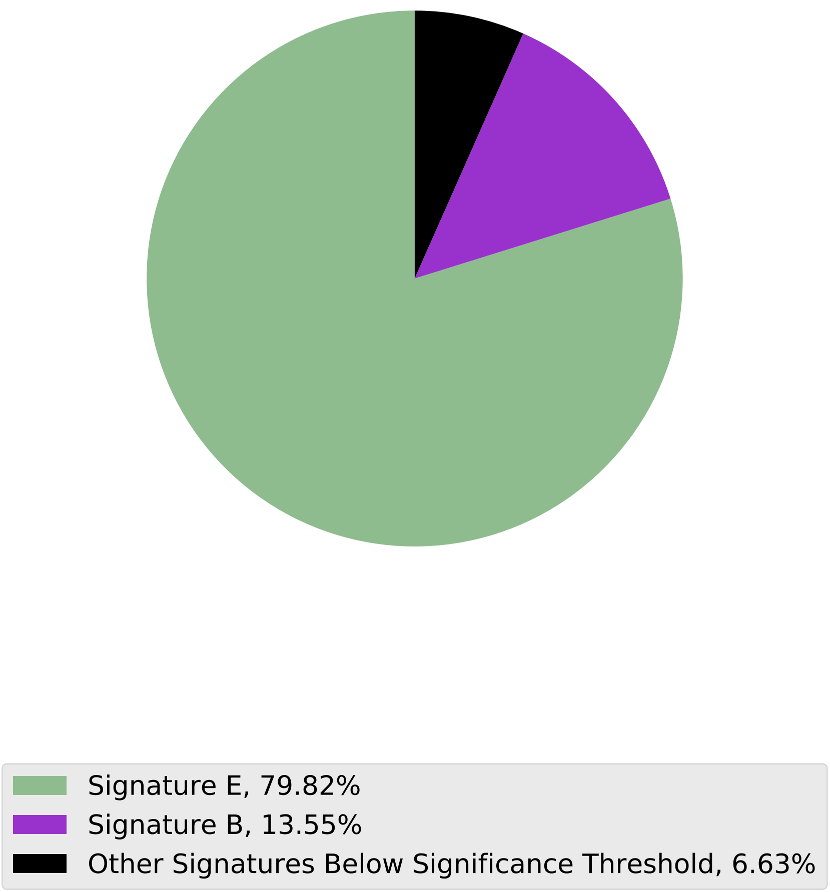

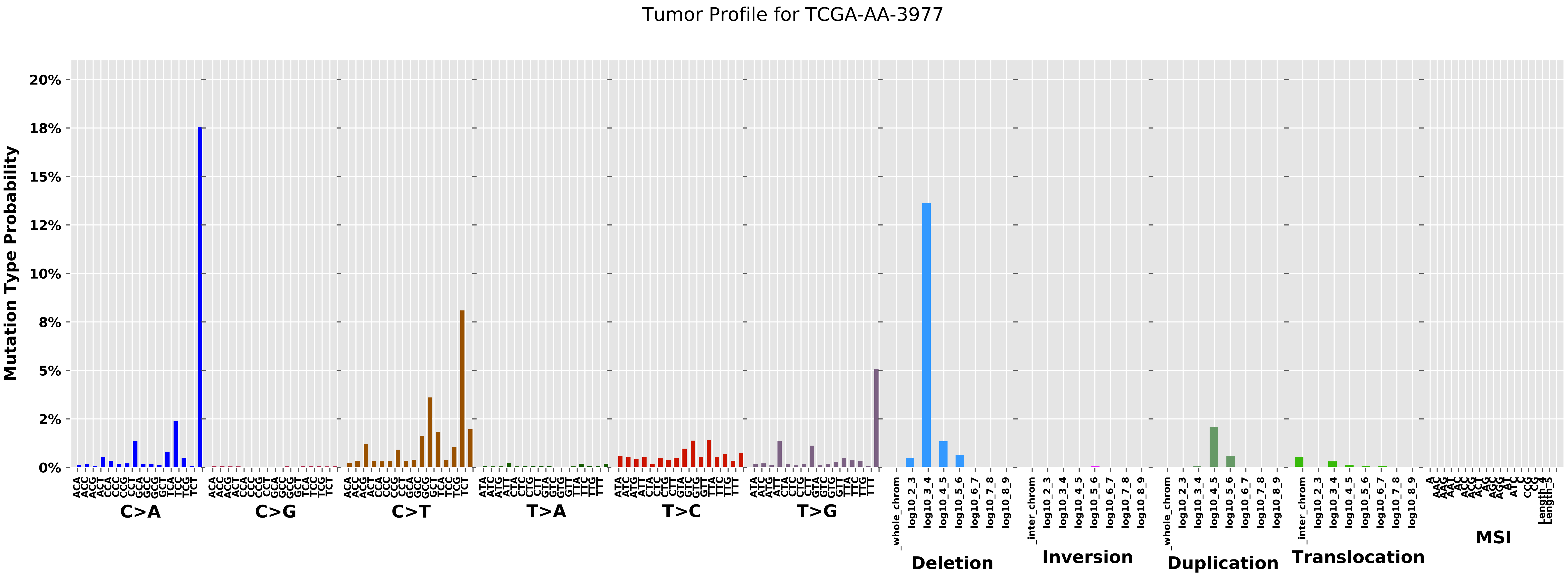

Tumor Profile for TCGA-AA-3977
Mutation Type Probability
20%
18%
15%
12%
10%
8%
5%
2%
0%
C>A
C>G
C>T
T>A
T>C
T>G
Deletion
Inversion
Duplication
Translocation
MSI
_whole_chrom
_inter_chrom
log10_2_3
log10_3_4
log10_4_5
log10_5_6
log10_6_7
log10_7_8
log10_8_9
Length_4
Length_5

Cancer processes Weights for TCGA-A7-A0CE

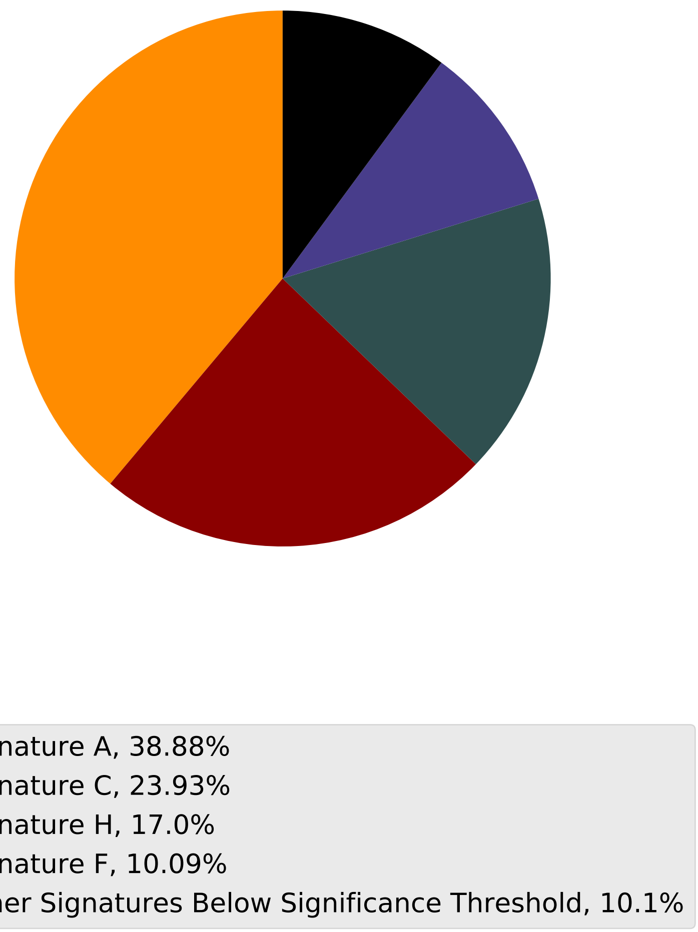

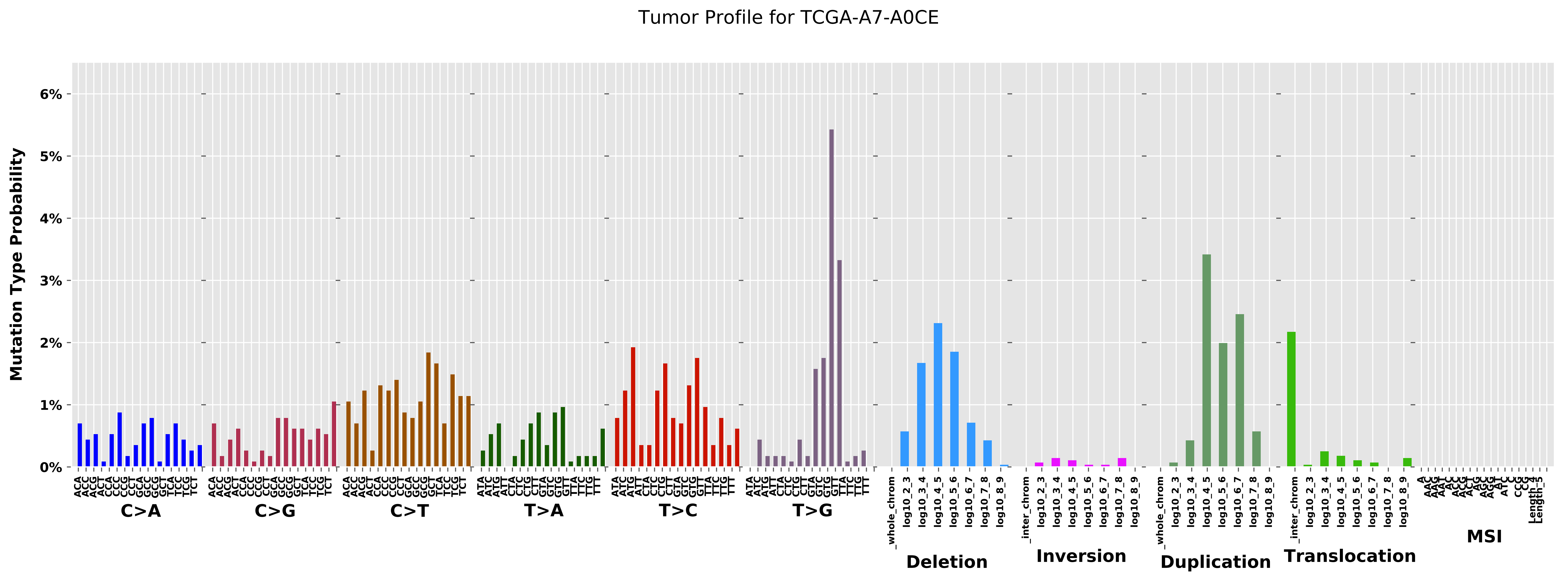

Tumor Profile for TCGA-A7-A0CE
Mutation Type Probability
0%
1%
2%
3%
4%
5%
6%
C>A
C>G
C>T
T>A
T>C
T>G
Deletion
Inversion
Duplication
Translocation
MSI

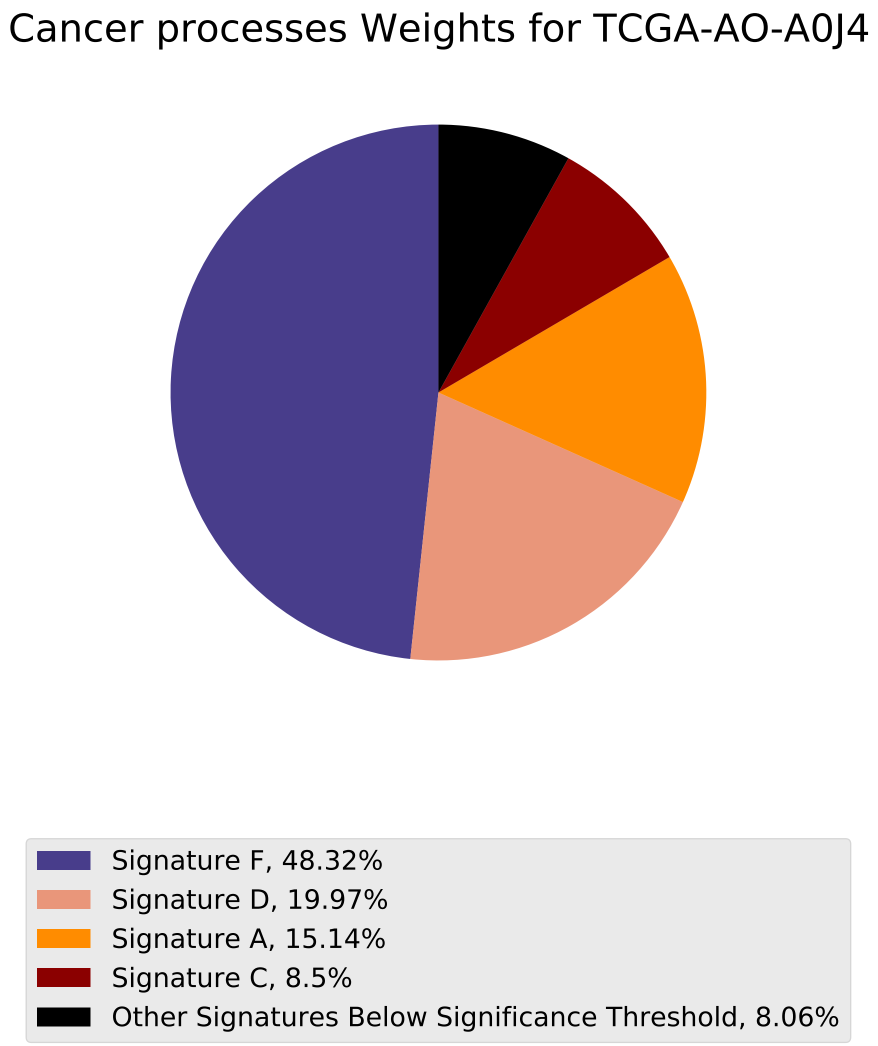
Cancer processes Weights for TCGA-AO-A0J4
Signature F, 48.32%
Signature D, 19.97%
Signature A, 15.14%
Signature C, 8.5%
Other Signatures Below Significance Threshold, 8.06%

Tumor Profile for TCGA-AO-A0J4

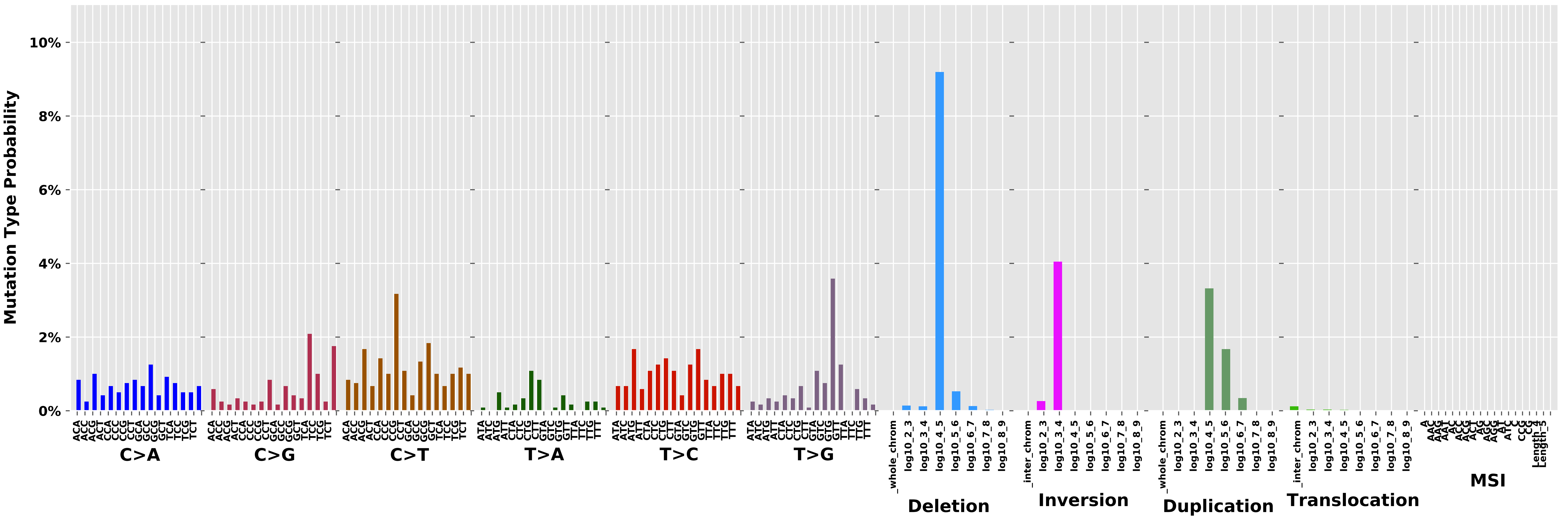

Cancer processes Weights for TCGA-AQ-A04J

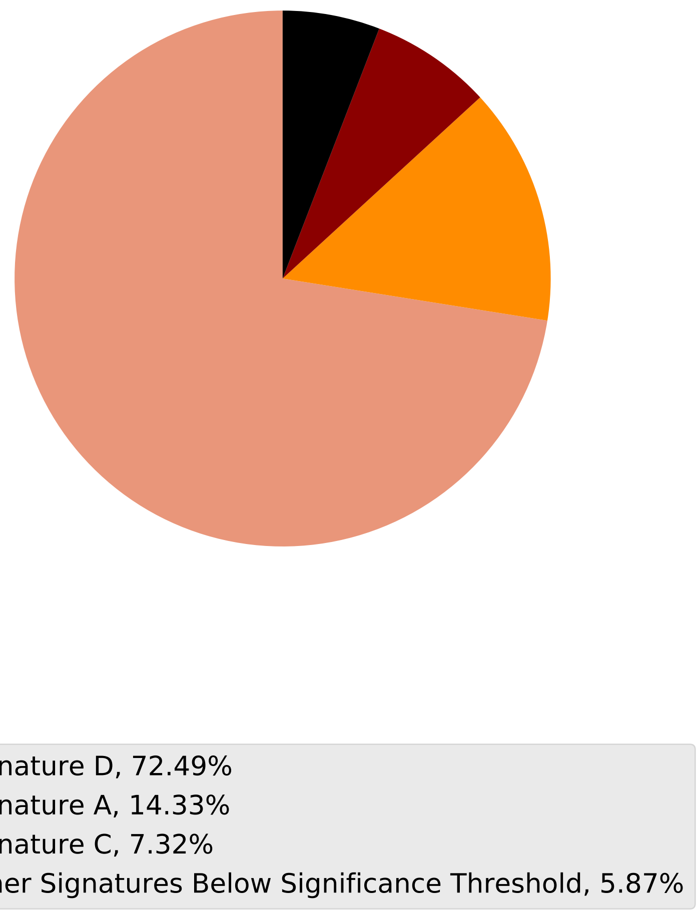

Tumor Profile for TCGA-AQ-A04J

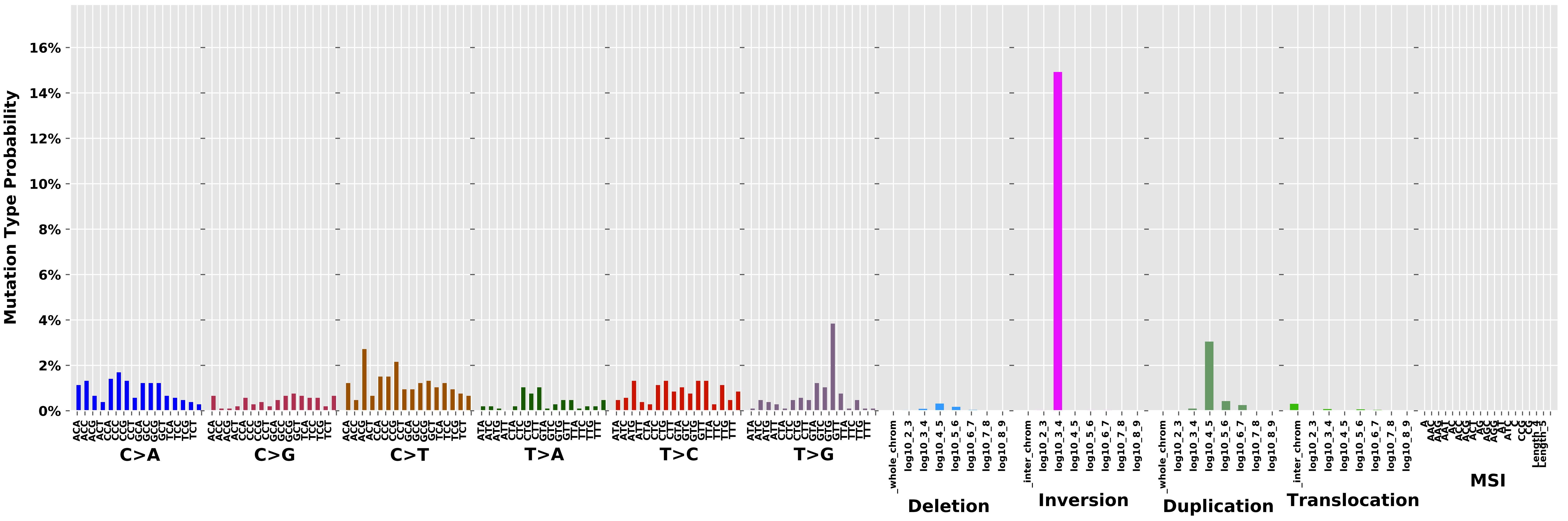

Cancer processes Weights for TCGA-AG-4007

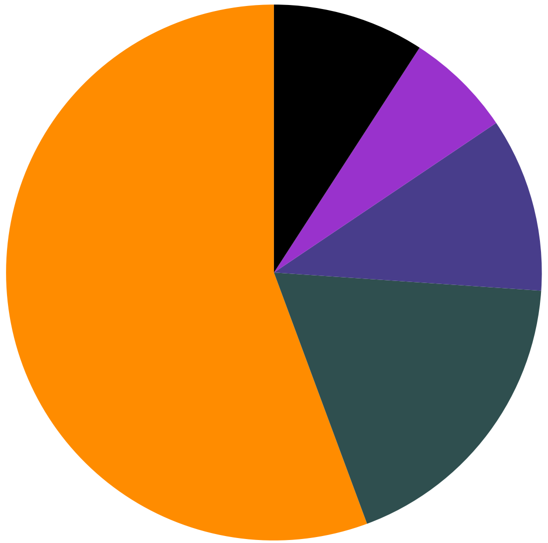

Signature A, 55.65%
Signature H, 18.27%
Signature F, 10.51%
Signature B, 6.42%
Other Signatures Below Significance Threshold, 9.15%

Tumor Profile for TCGA-AG-4007

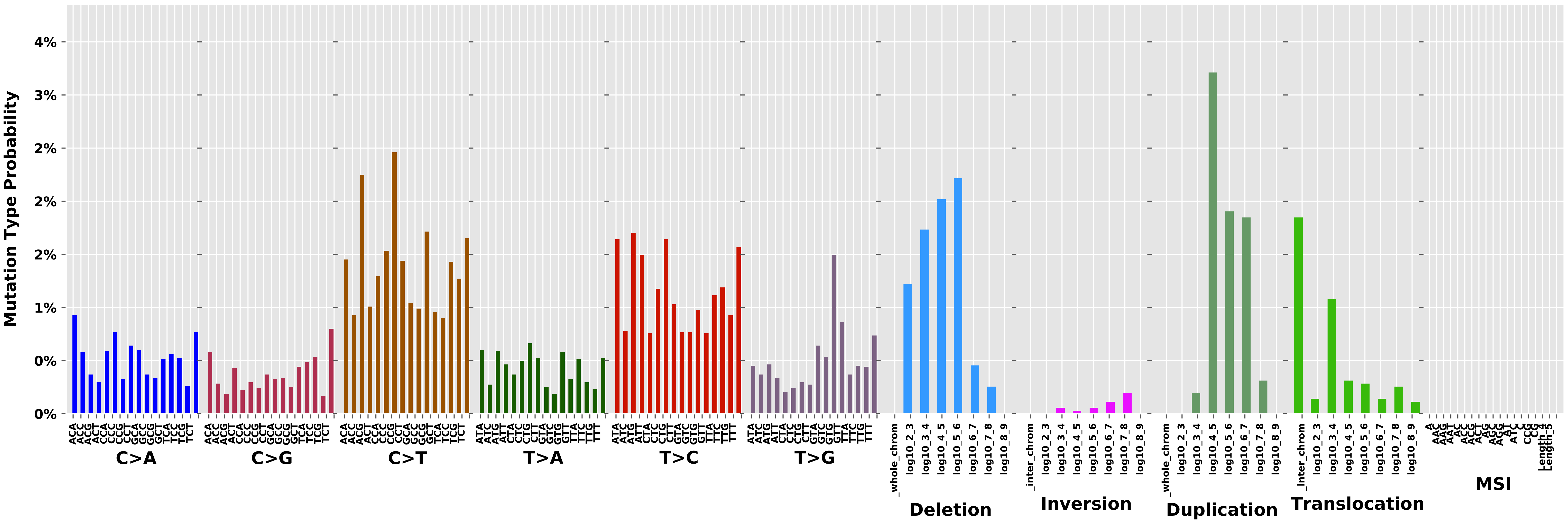

Cancer processes Weights for TCGA-CA-6717

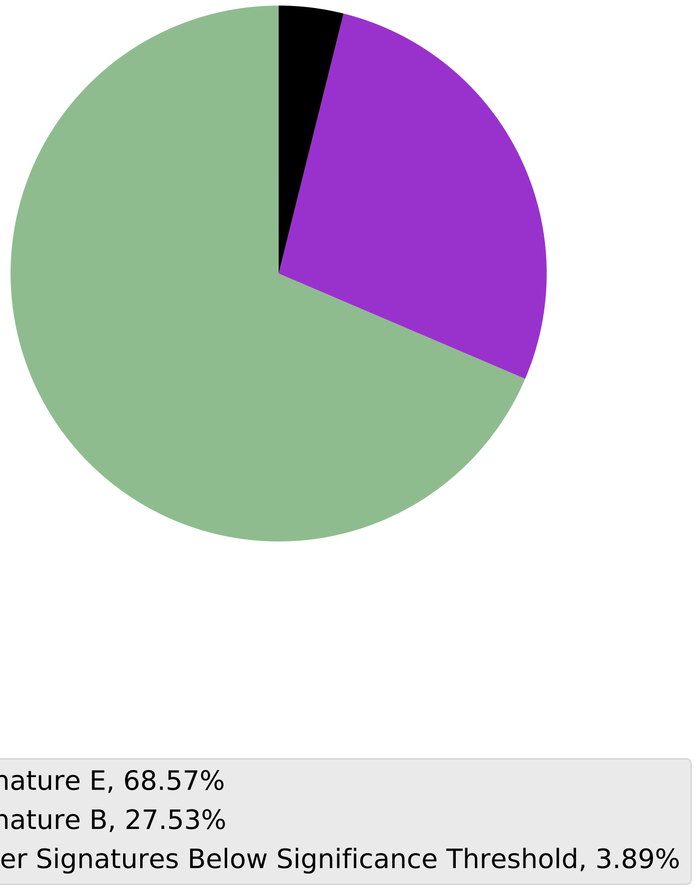

Tumor Profile for TCGA-CA-6717

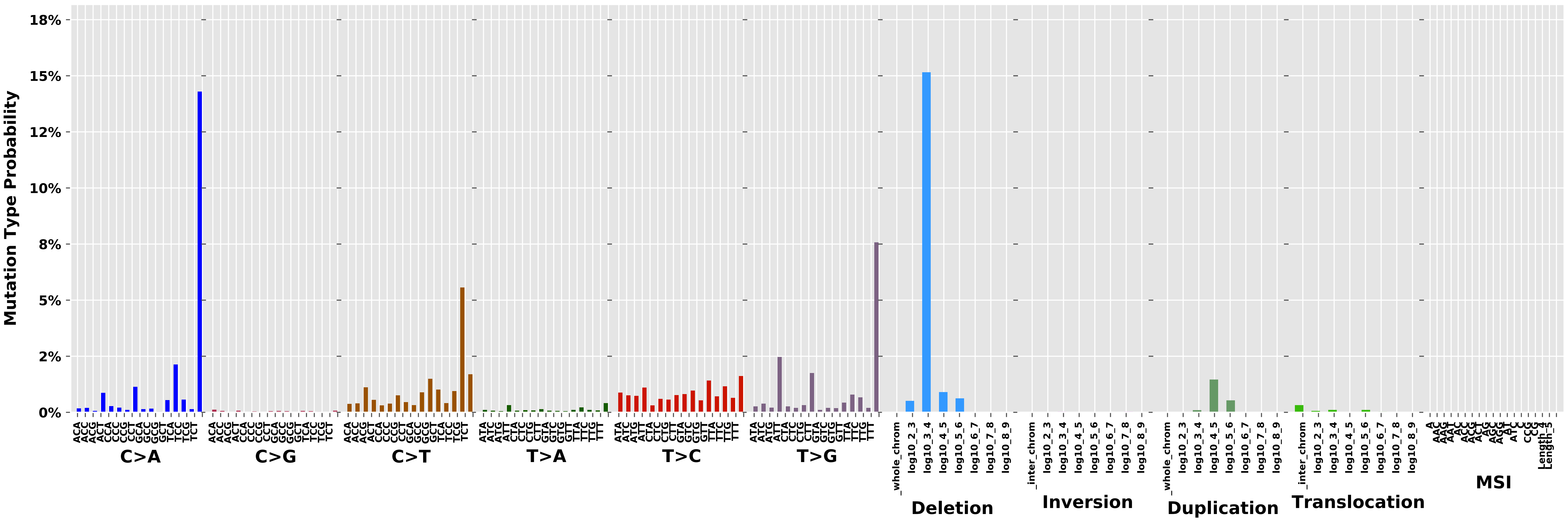

Cancer processes Weights for TCGA-E2-A14P

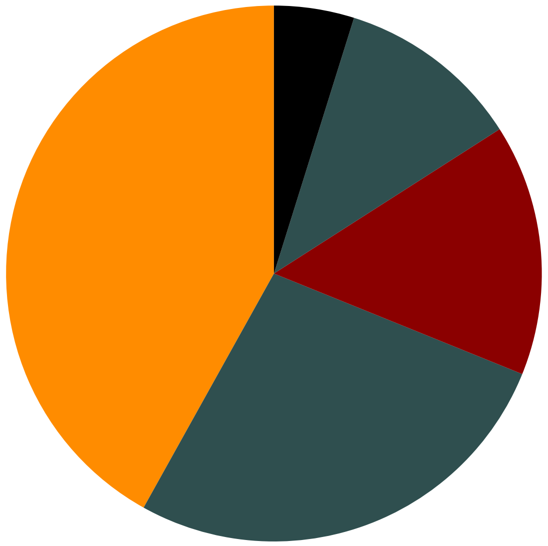

- Signature A, 41.91%
- Signature H, 26.97%
- Signature C, 15.18%
- Signature G, 11.11%
- Other Signatures Below Significance Threshold, 4.83%

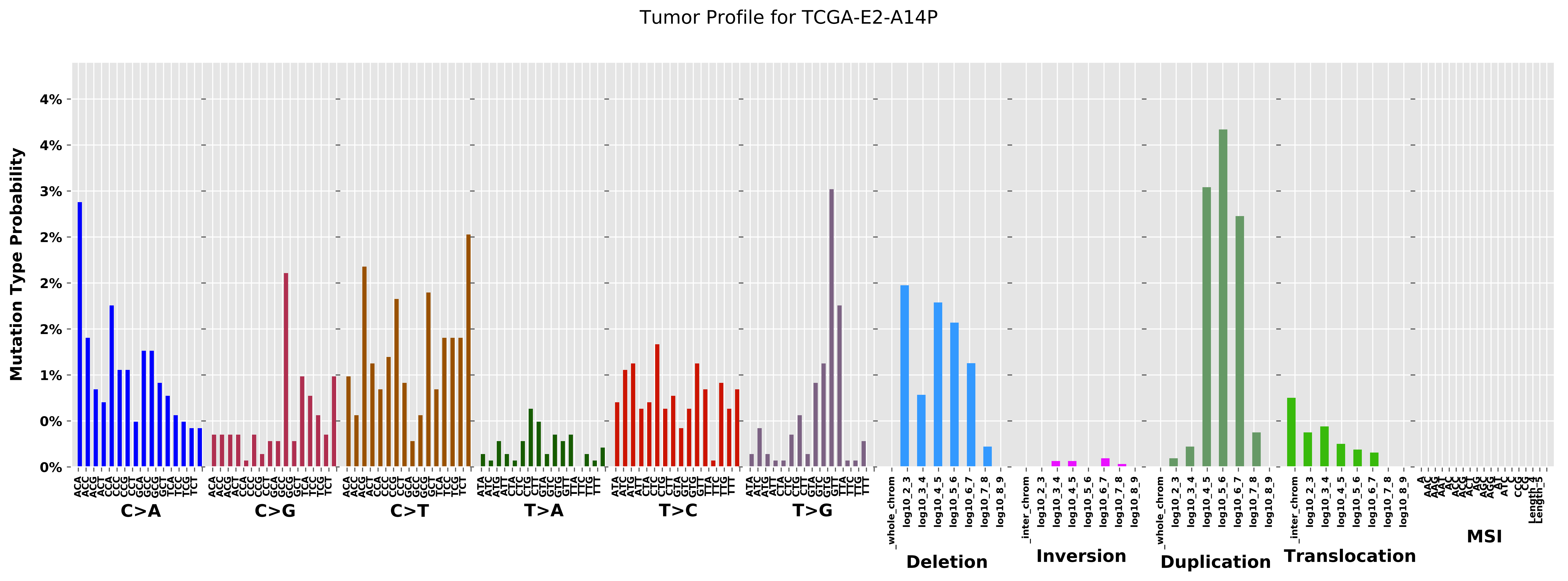
Tumor Profile for TCGA-E2-A14P
Mutation Type Probability
C>A
C>G
C>T
T>A
T>C
T>G
Deletion
Inversion
Duplication
Translocation
MSI

Cancer processes Weights for TCGA-AZ-6601

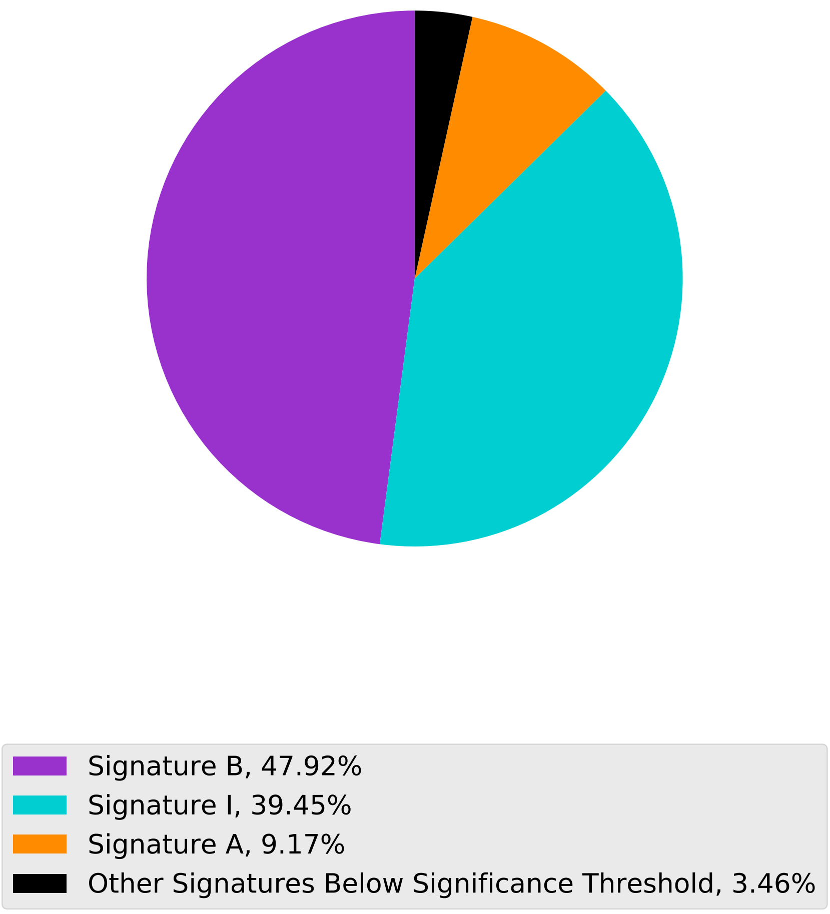

Tumor Profile for TCGA-AZ-6601

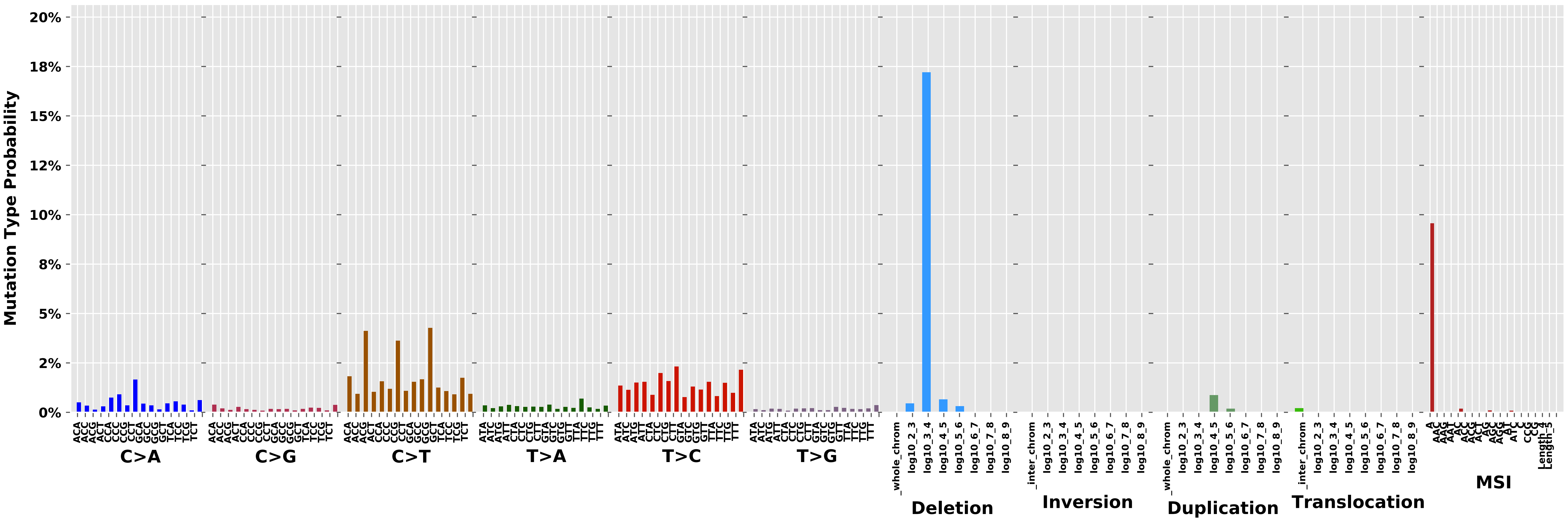

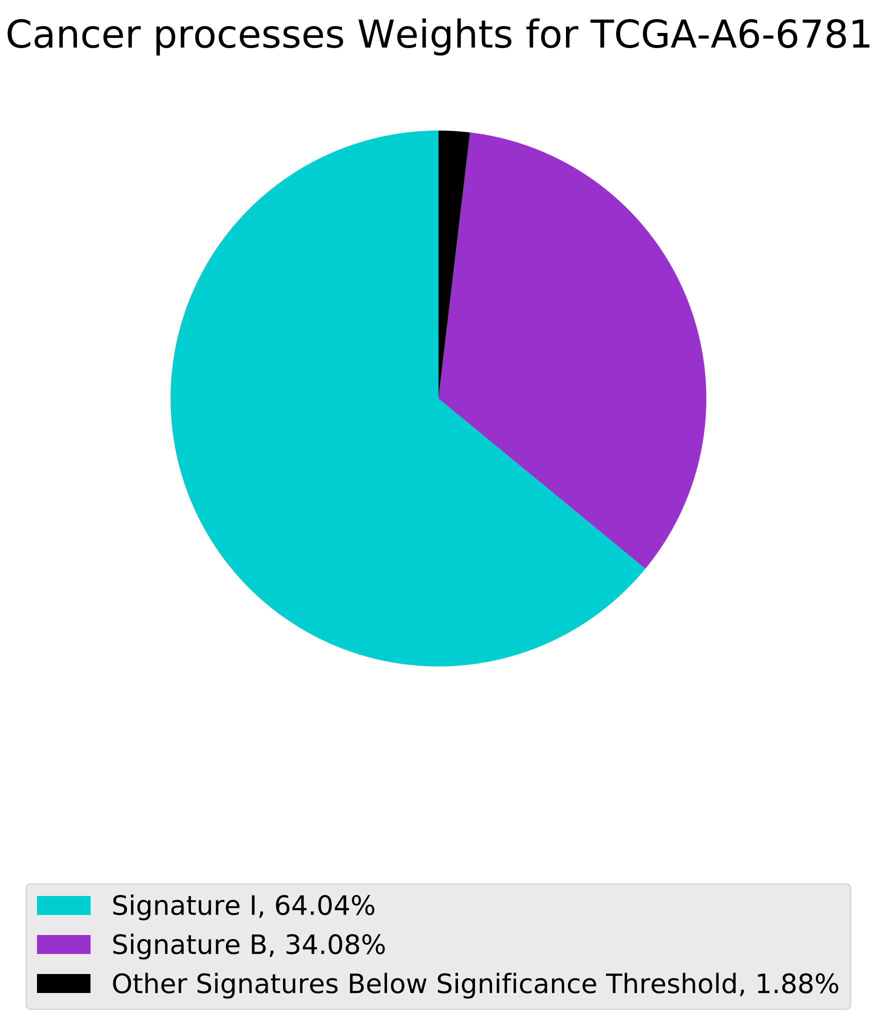

Cancer processes Weights for TCGA-A6-6781
Signature I, 64.04%
Signature B, 34.08%
Other Signatures Below Significance Threshold, 1.88%

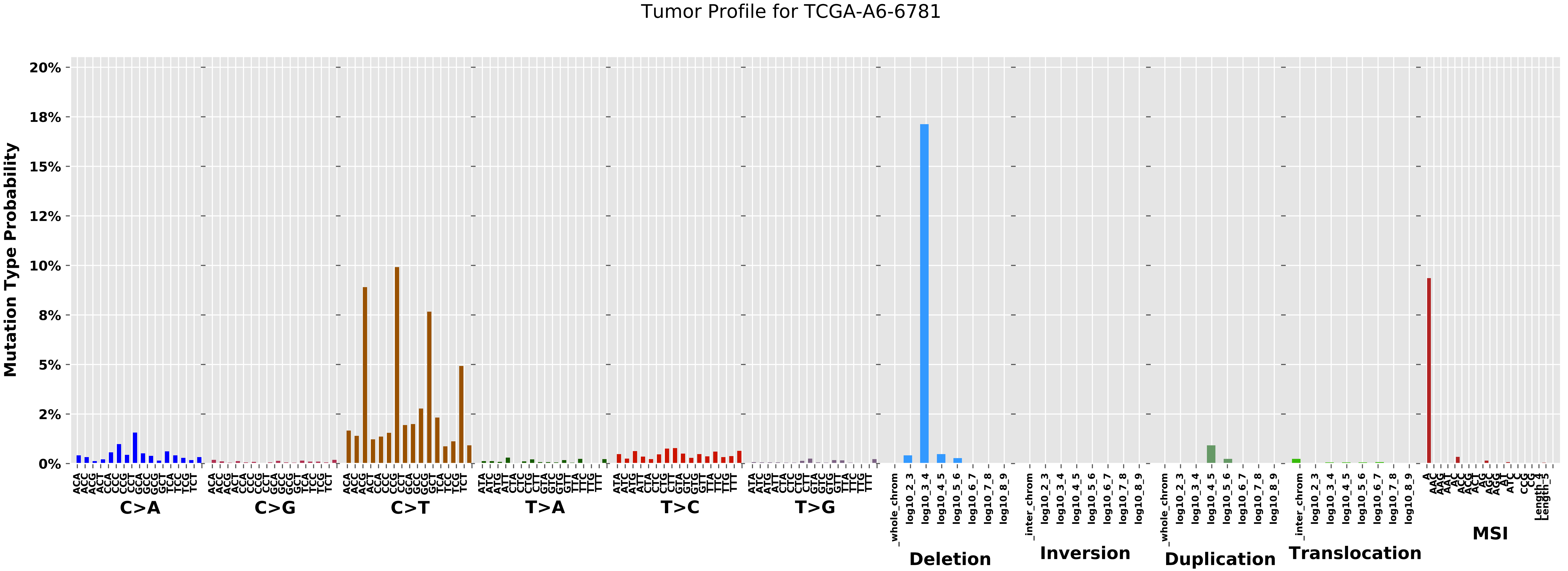
Tumor Profile for TCGA-A6-6781
Mutation Type Probability
20%
18%
15%
12%
10%
8%
5%
2%
0%
C>A
C>G
C>T
T>A
T>C
T>G
Deletion
Inversion
Duplication
Translocation
MSI
_whole_chrom
_inter_chrom
log10_2_3
log10_3_4
log10_4_5
log10_5_6
log10_6_7
log10_7_8
log10_8_9

Cancer processes Weights for TCGA-GM-A2DF

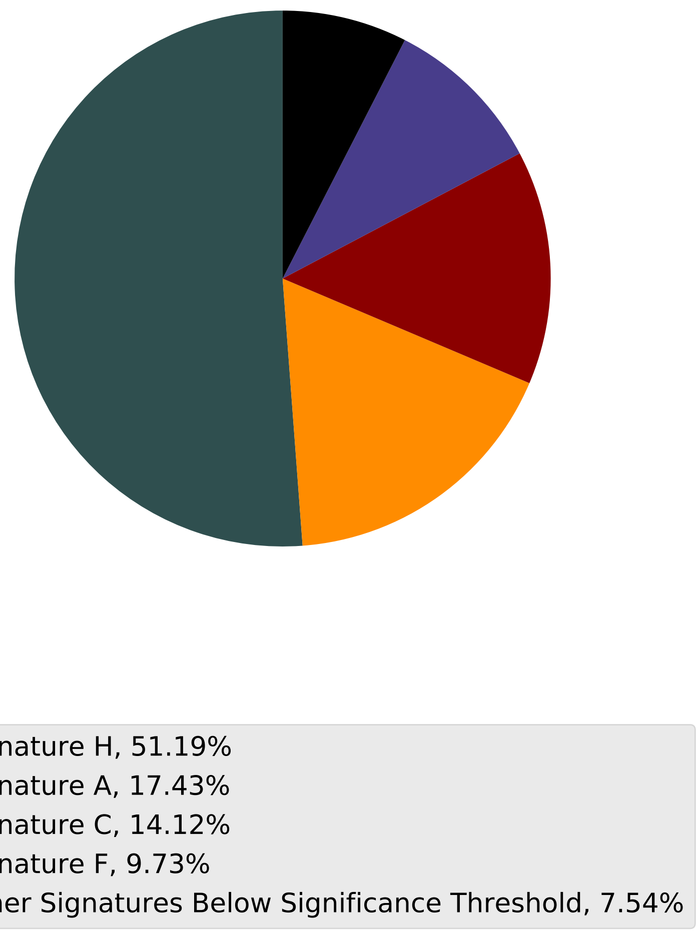

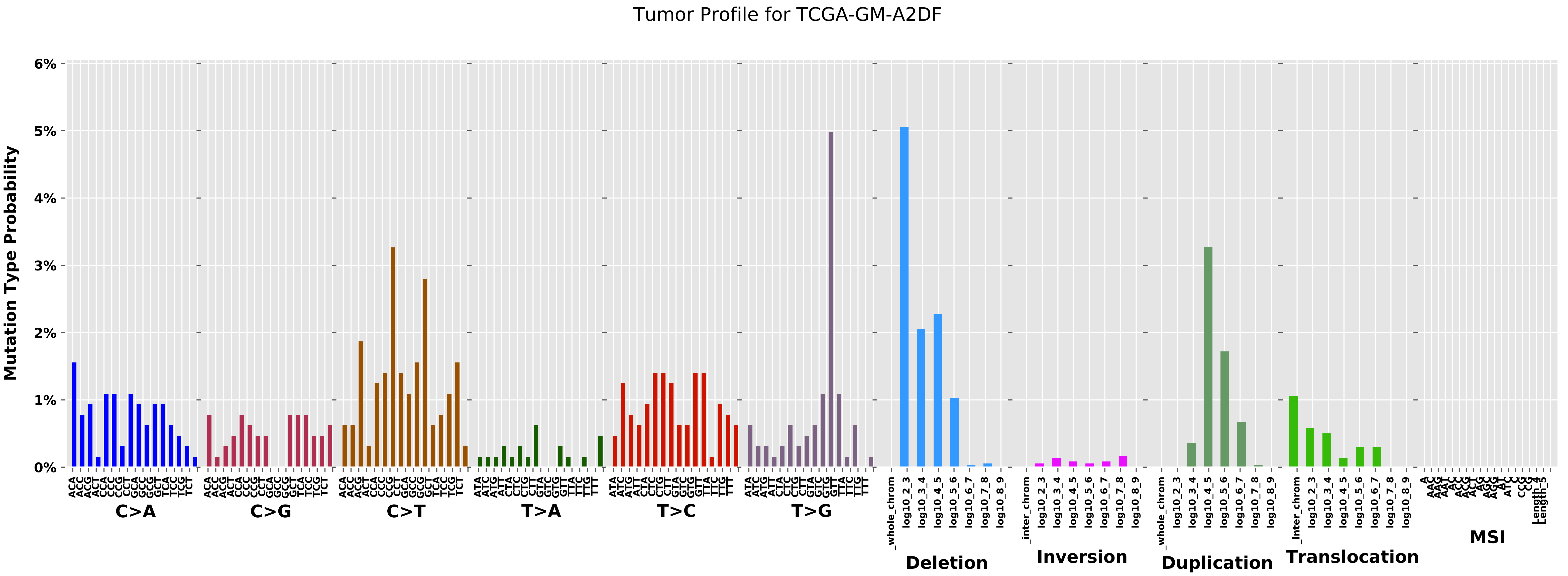
Tumor Profile for TCGA-GM-A2DF
Mutation Type Probability
0%
1%
2%
3%
4%
5%
6%
C>A
C>G
C>T
T>A
T>C
T>G
Deletion
Inversion
Duplication
Translocation
MSI

Cancer processes Weights for TCGA-C8-A12L

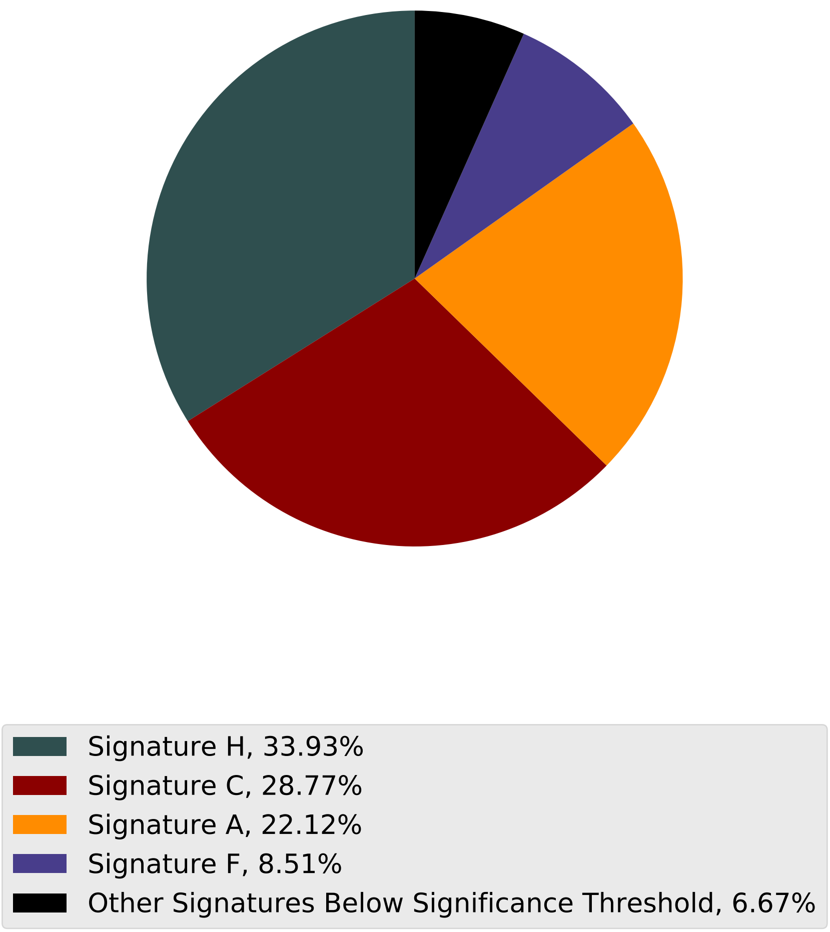

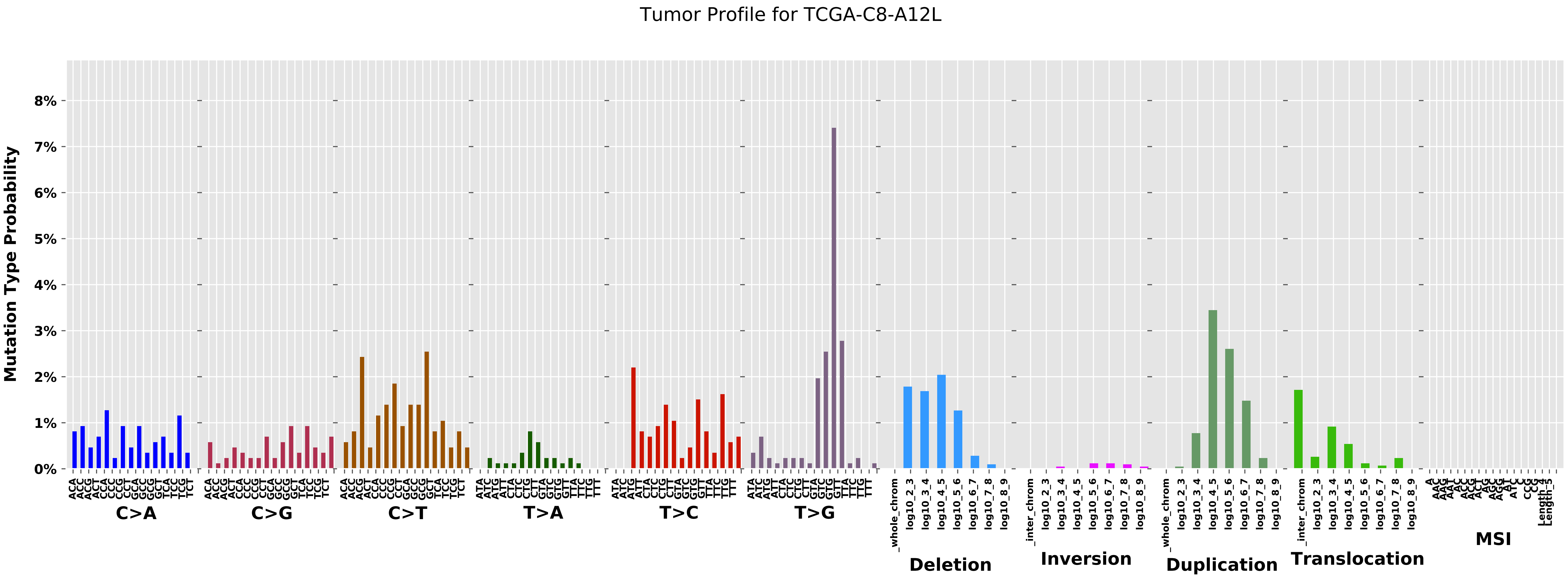
Tumor Profile for TCGA-C8-A12L
Mutation Type Probability
0%
1%
2%
3%
4%
5%
6%
7%
8%
C>A
C>G
C>T
T>A
T>C
T>G
Deletion
Inversion
Duplication
Translocation
MSI
_whole_chrom
_inter_chrom
log10_2_3
log10_3_4
log10_4_5
log10_5_6
log10_6_7
log10_7_8
log10_8_9

Cancer processes Weights for TCGA-AG-3890

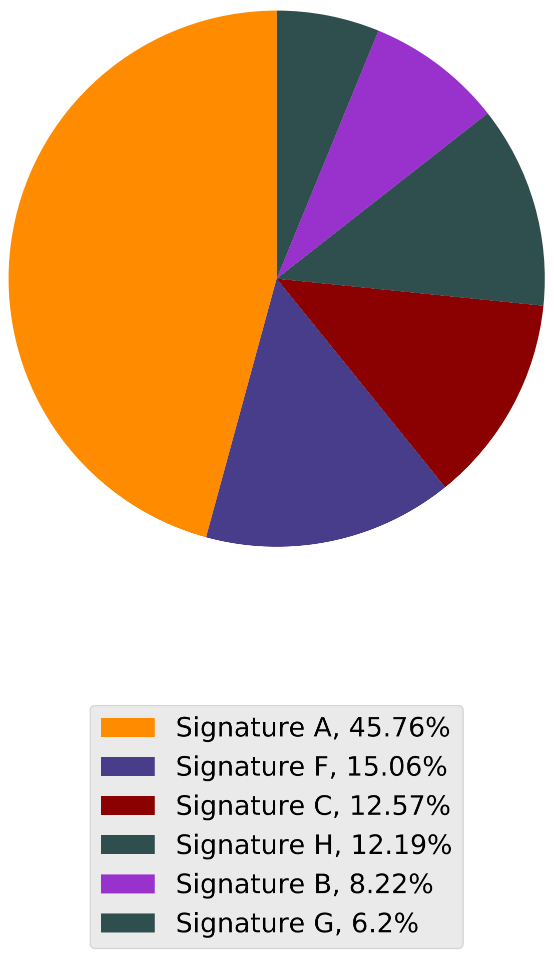

Tumor Profile for TCGA-AG-3890

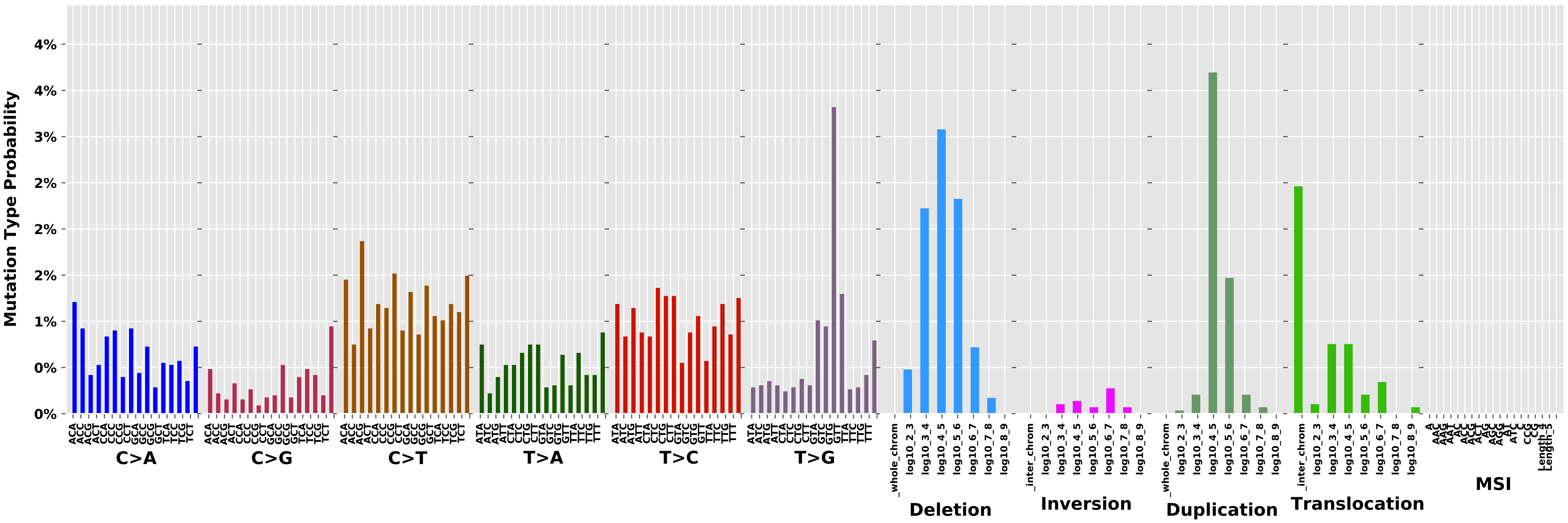

Cancer processes Weights for TCGA-E2-A1LK

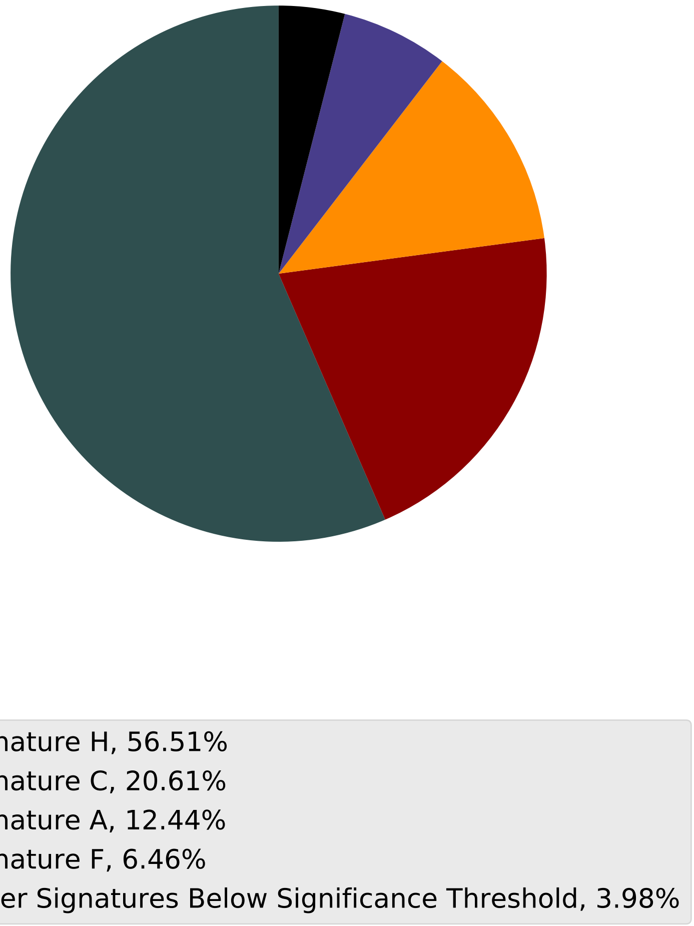

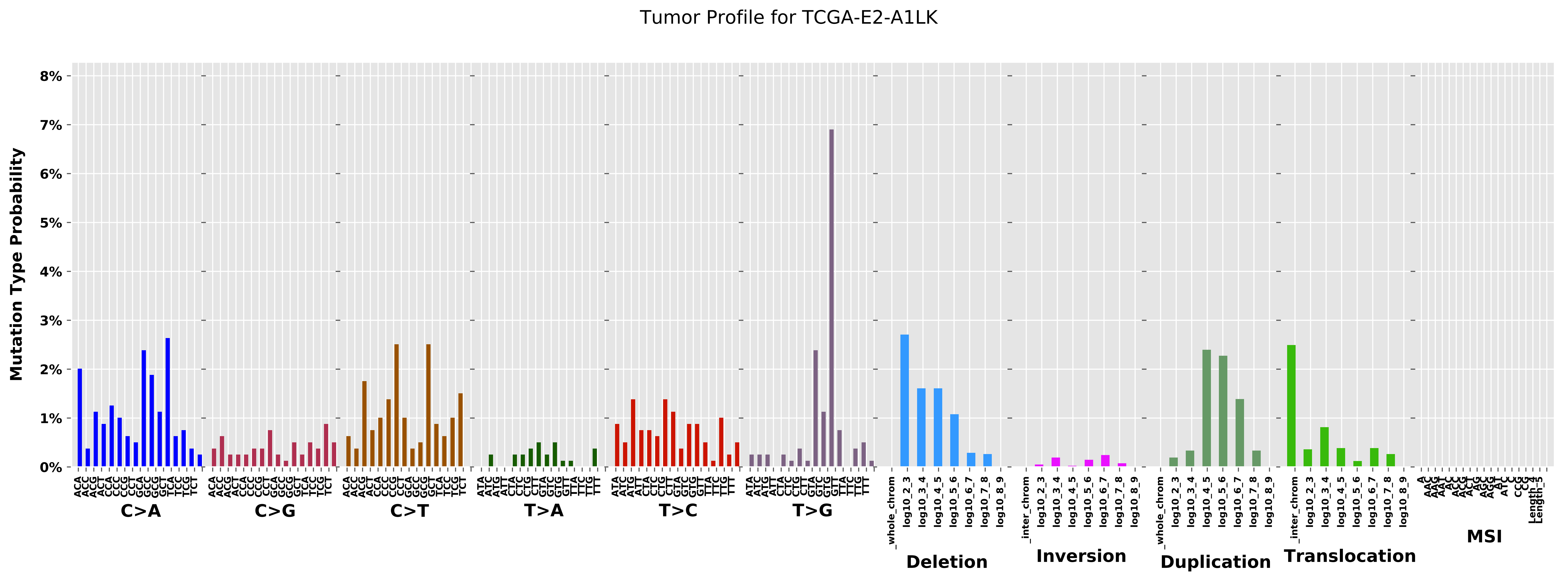

Tumor Profile for TCGA-E2-A1LK
Mutation Type Probability
0%
1%
2%
3%
4%
5%
6%
7%
8%
C>A
C>G
C>T
T>A
T>C
T>G
Deletion
Inversion
Duplication
Translocation
MSI

Cancer processes Weights for TCGA-A2-A3Y0

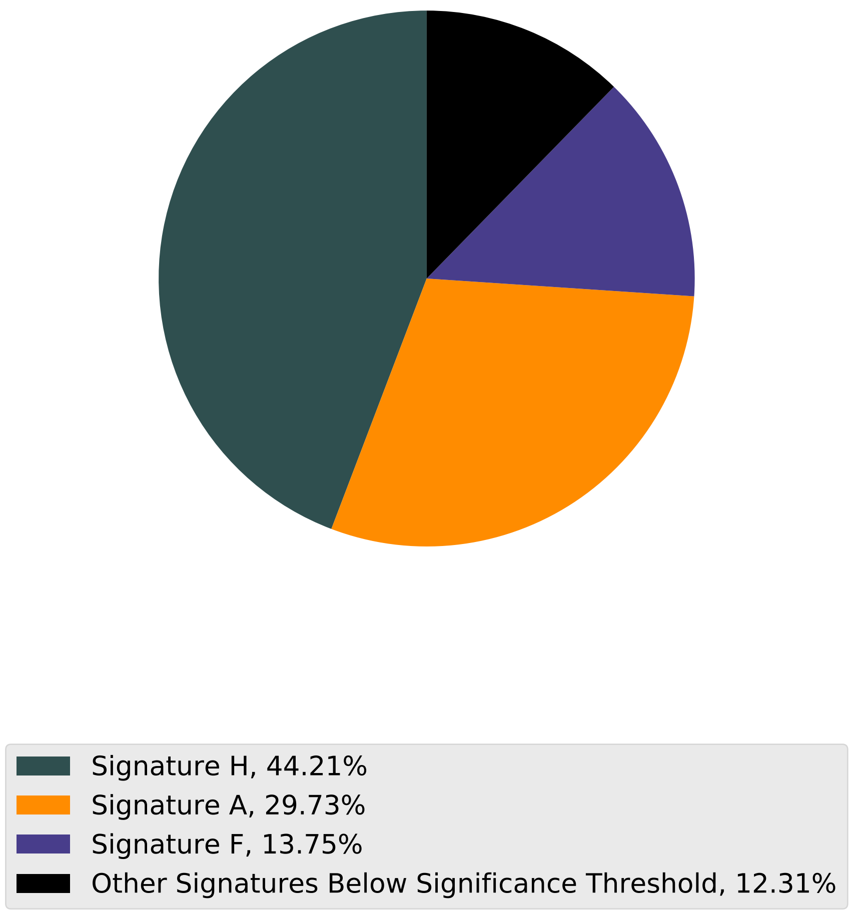

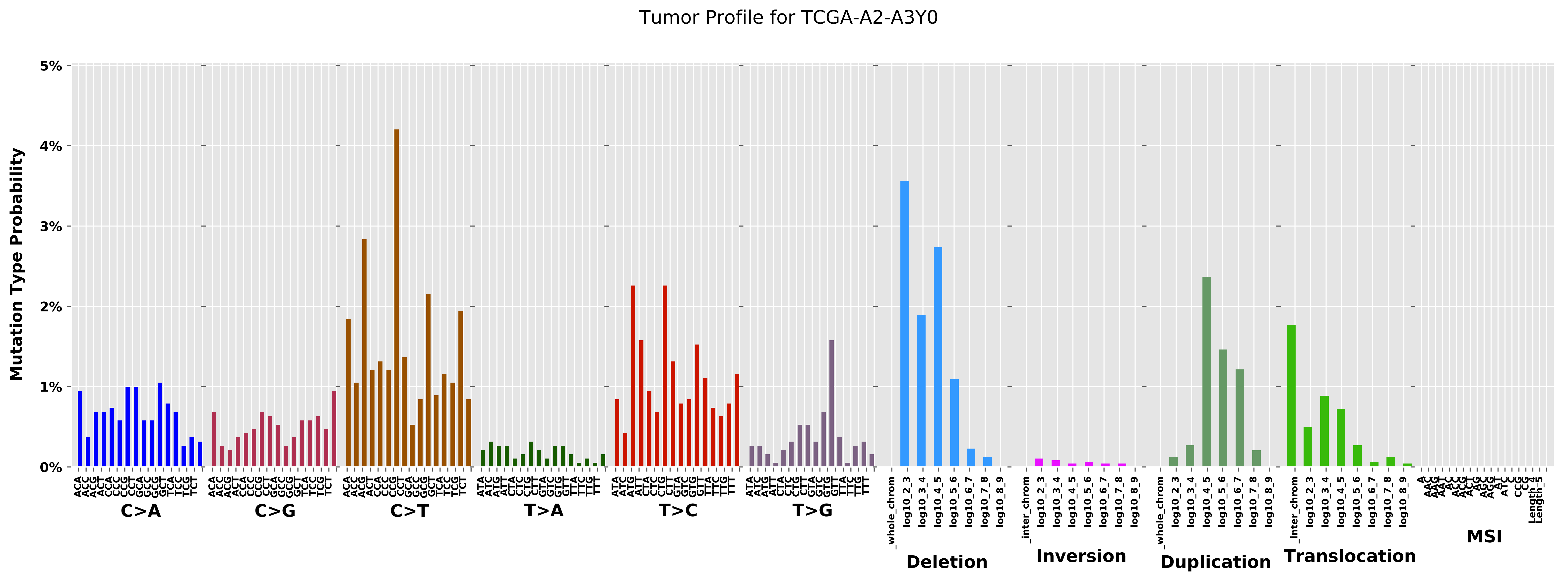

Tumor Profile for TCGA-A2-A3Y0
Mutation Type Probability
0%
1%
2%
3%
4%
5%
C>A
C>G
C>T
T>A
T>C
T>G
Deletion
Inversion
Duplication
Translocation
MSI
_whole_chrom
_inter_chrom
log10_2_3
log10_3_4
log10_4_5
log10_5_6
log10_6_7
log10_7_8
log10_8_9

Cancer processes Weights for TCGA-B6-A0I1

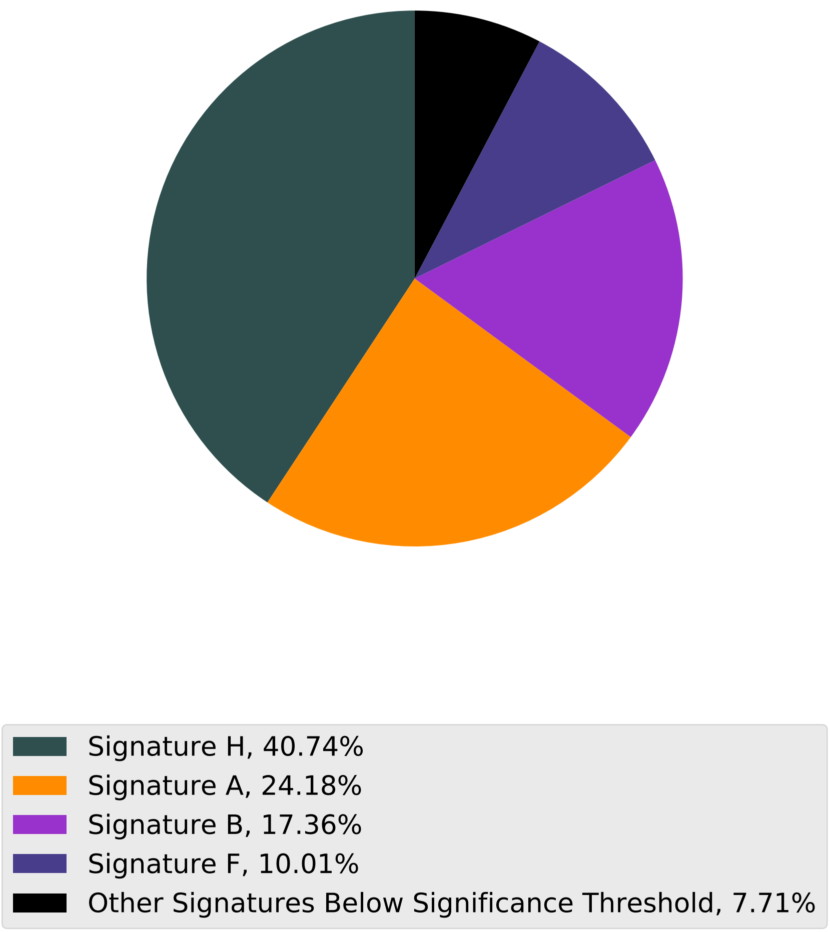

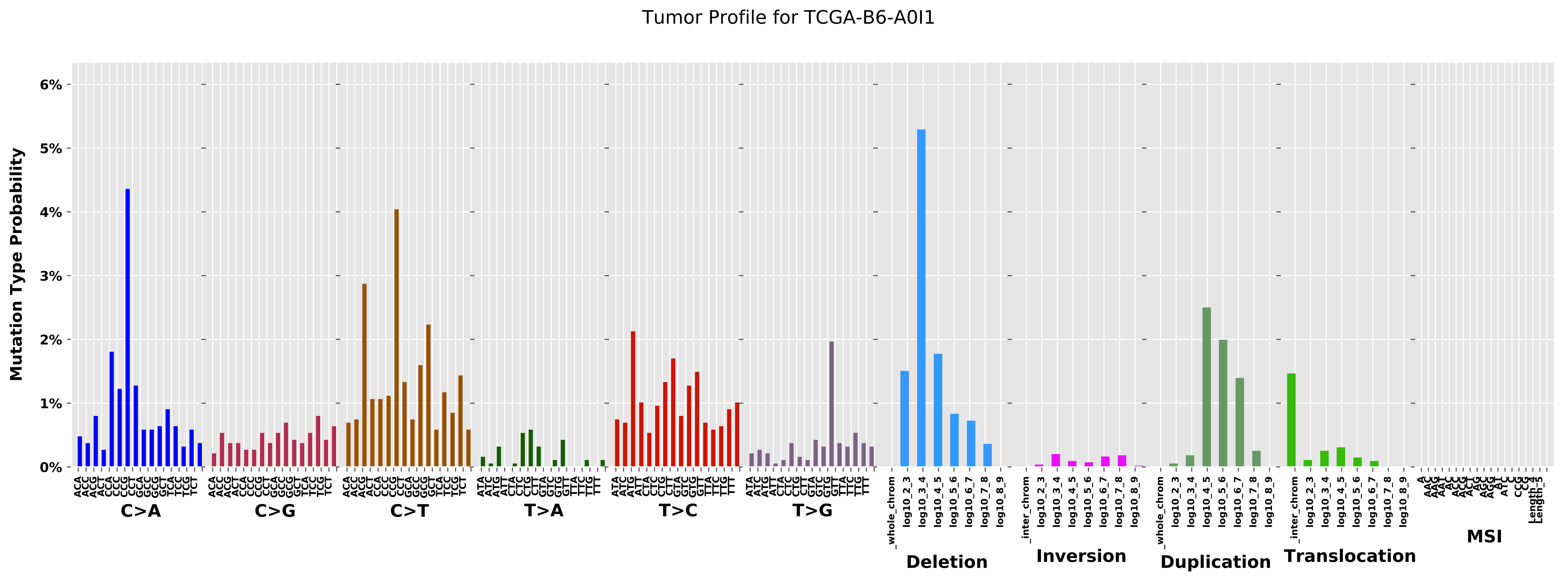
Tumor Profile for TCGA-B6-A0I1
Mutation Type Probability
0%
1%
2%
3%
4%
5%
6%
C>A
C>G
C>T
T>A
T>C
T>G
Deletion
Inversion
Duplication
Translocation
MSI

Cancer processes Weights for TCGA-E2-A15E

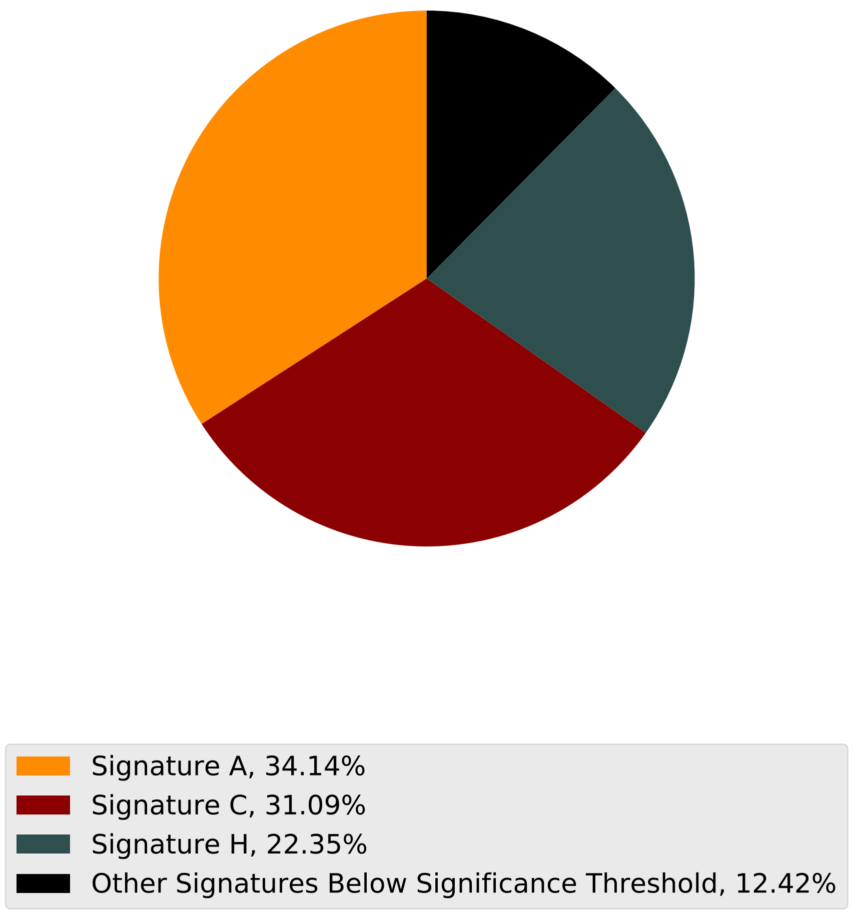

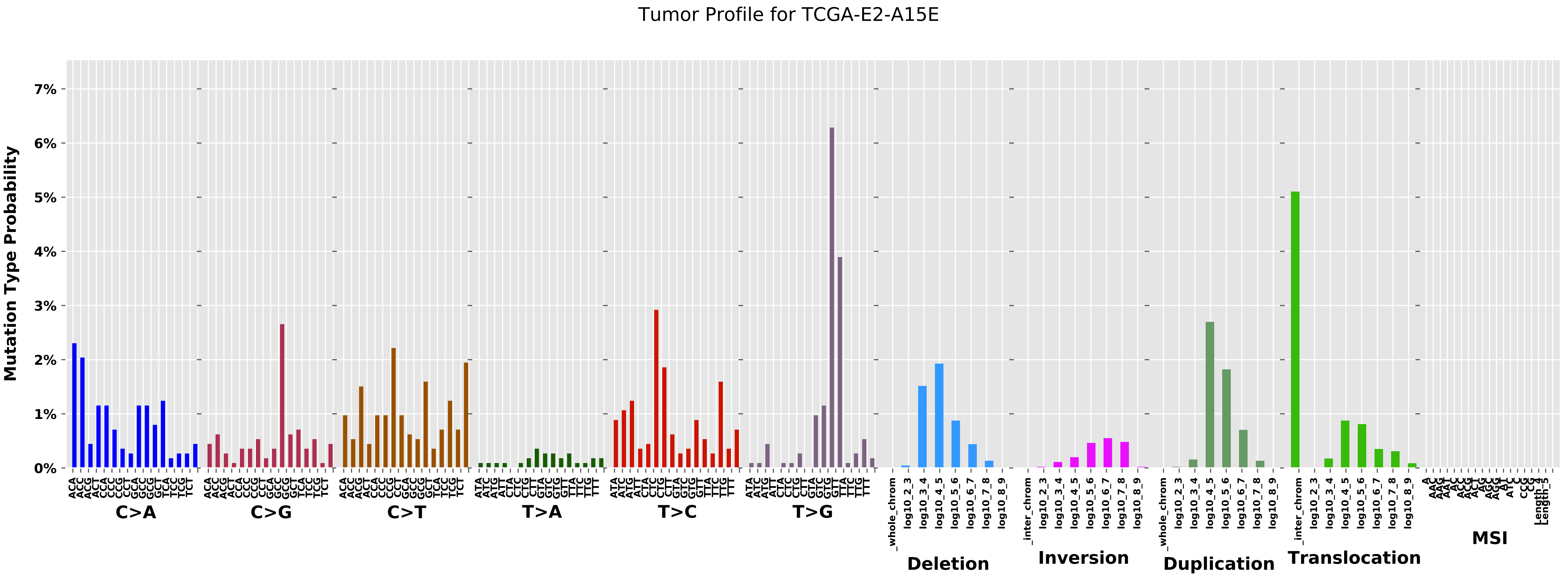

Tumor Profile for TCGA-E2-A15E
Mutation Type Probability
0%
1%
2%
3%
4%
5%
6%
7%
C>A
C>G
C>T
T>A
T>C
T>G
Deletion
Inversion
Duplication
Translocation
MSI

Cancer processes Weights for TCGA-A6-2680

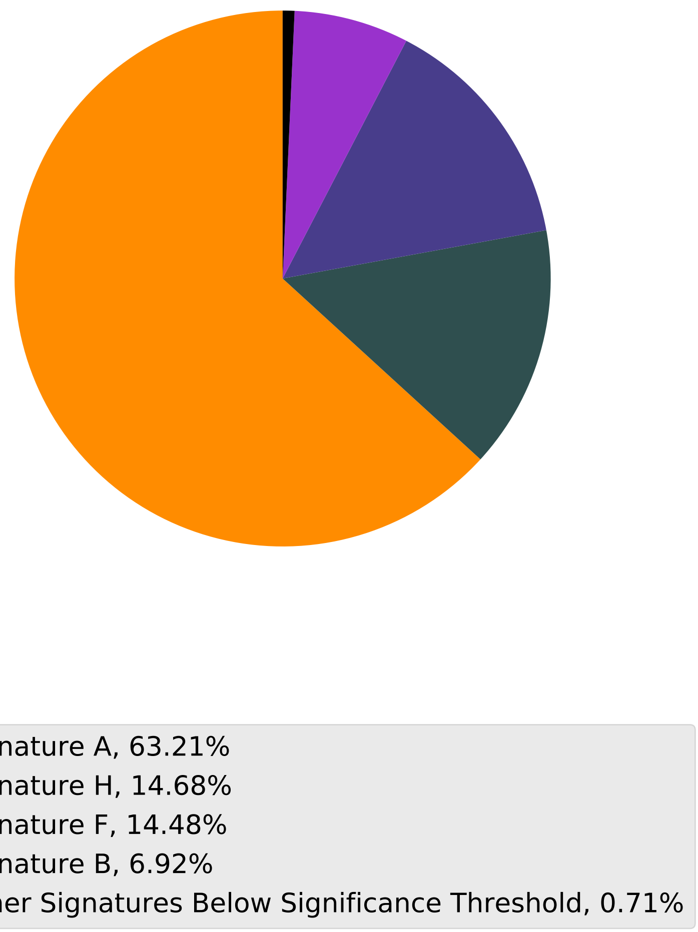

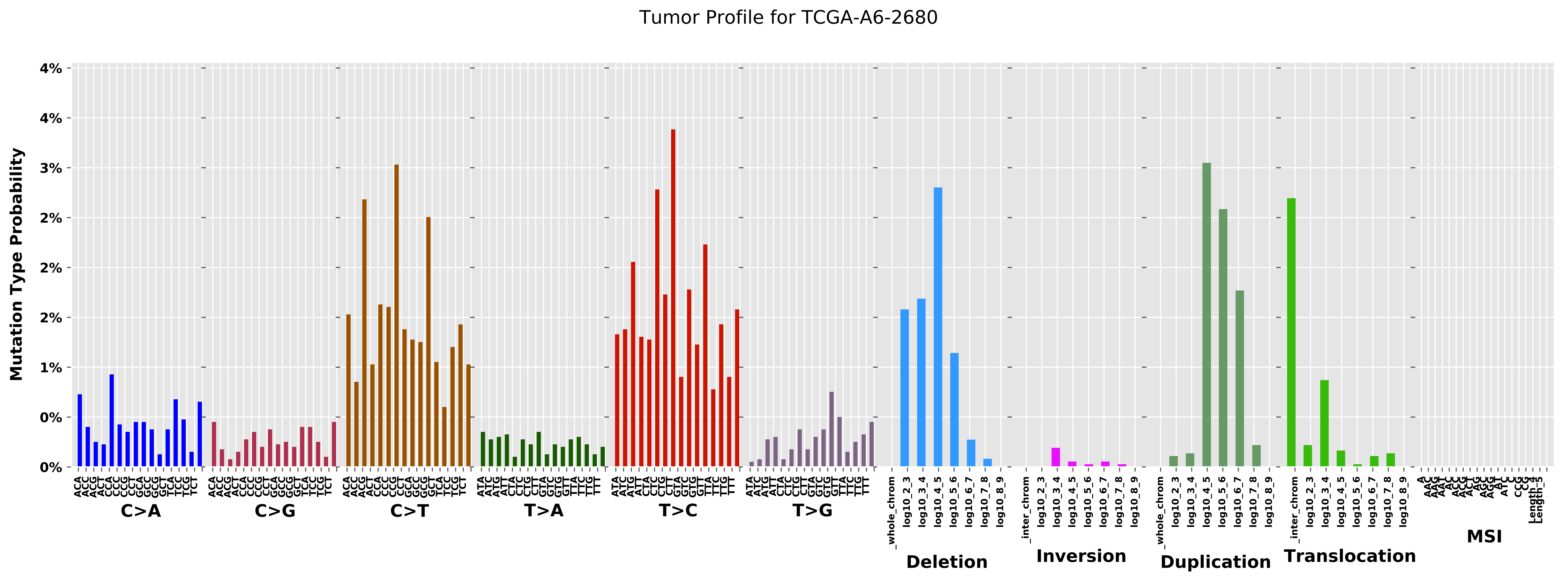

Tumor Profile for TCGA-A6-2680
Mutation Type Probability
0%
1%
2%
3%
4%
C>A
C>G
C>T
T>A
T>C
T>G
Deletion
Inversion
Duplication
Translocation
MSI

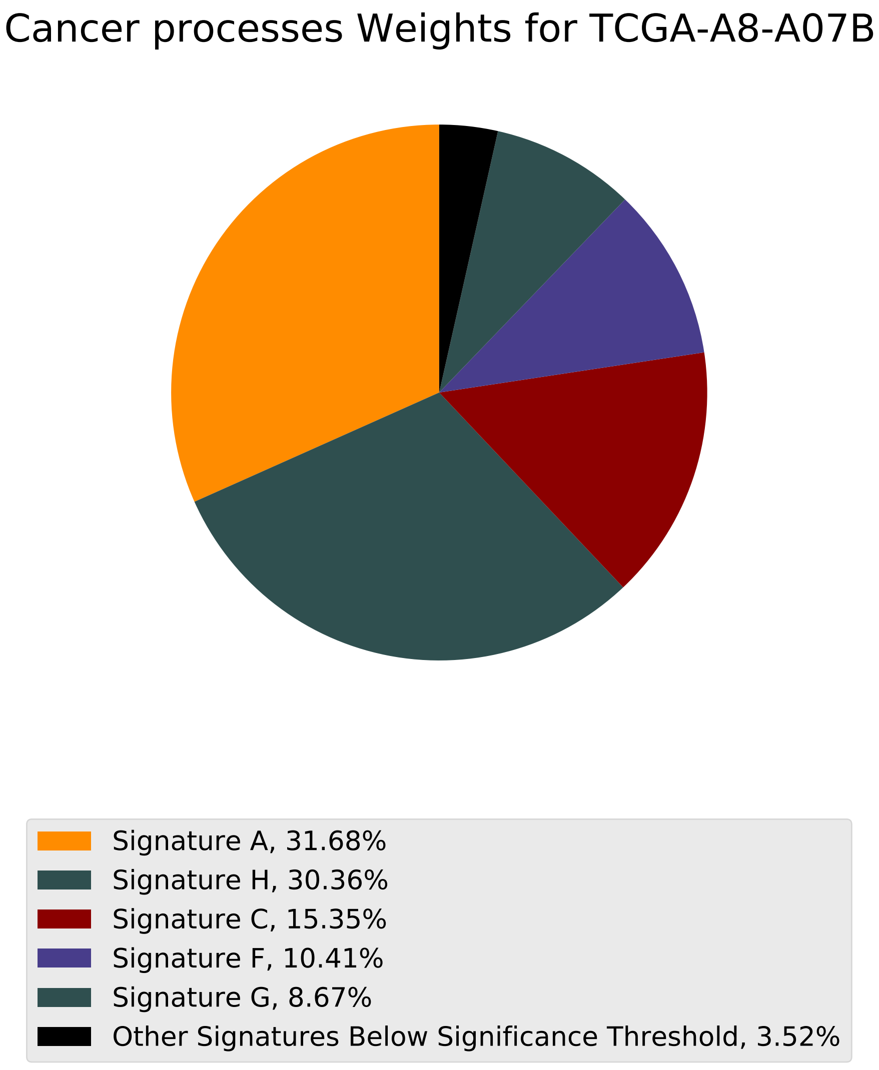
Cancer processes Weights for TCGA-A8-A07B
Signature A, 31.68%
Signature H, 30.36%
Signature C, 15.35%
Signature F, 10.41%
Signature G, 8.67%
Other Signatures Below Significance Threshold, 3.52%

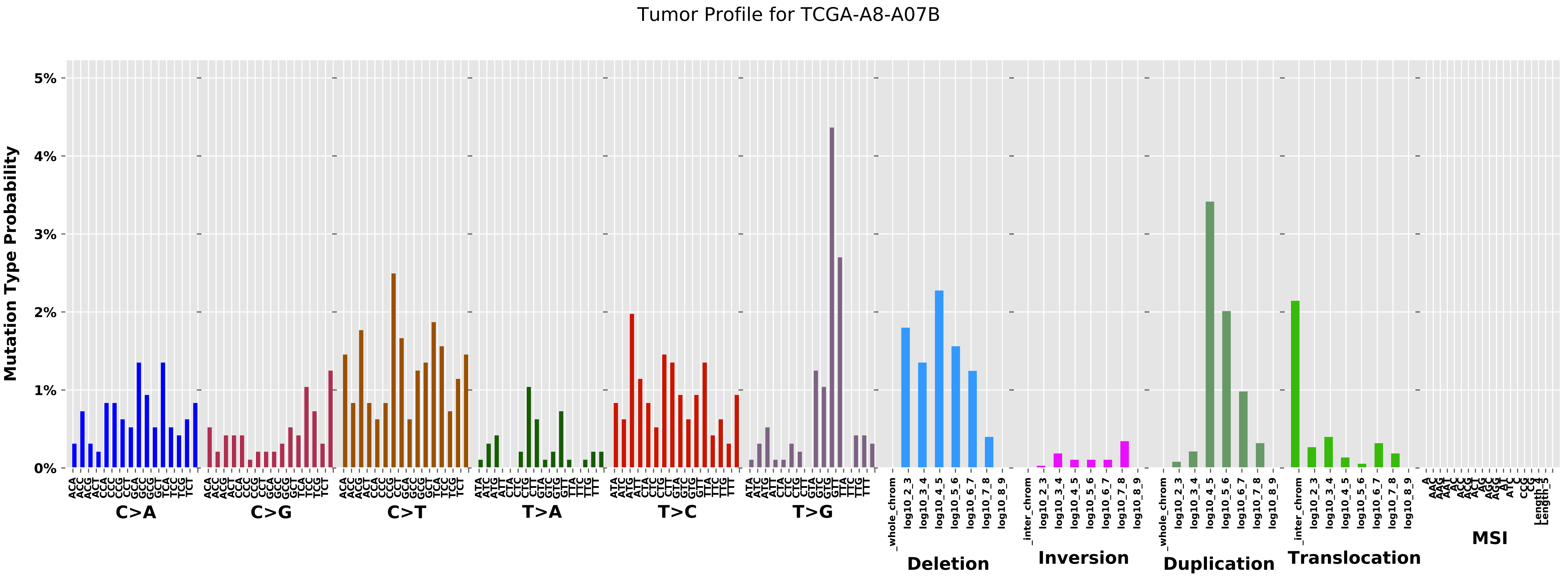

Tumor Profile for TCGA-A8-A07B
Mutation Type Probability
0%
1%
2%
3%
4%
5%
C>A
C>G
C>T
T>A
T>C
T>G
Deletion
Inversion
Duplication
Translocation
MSI

Cancer processes Weights for TCGA-AO-A0J6

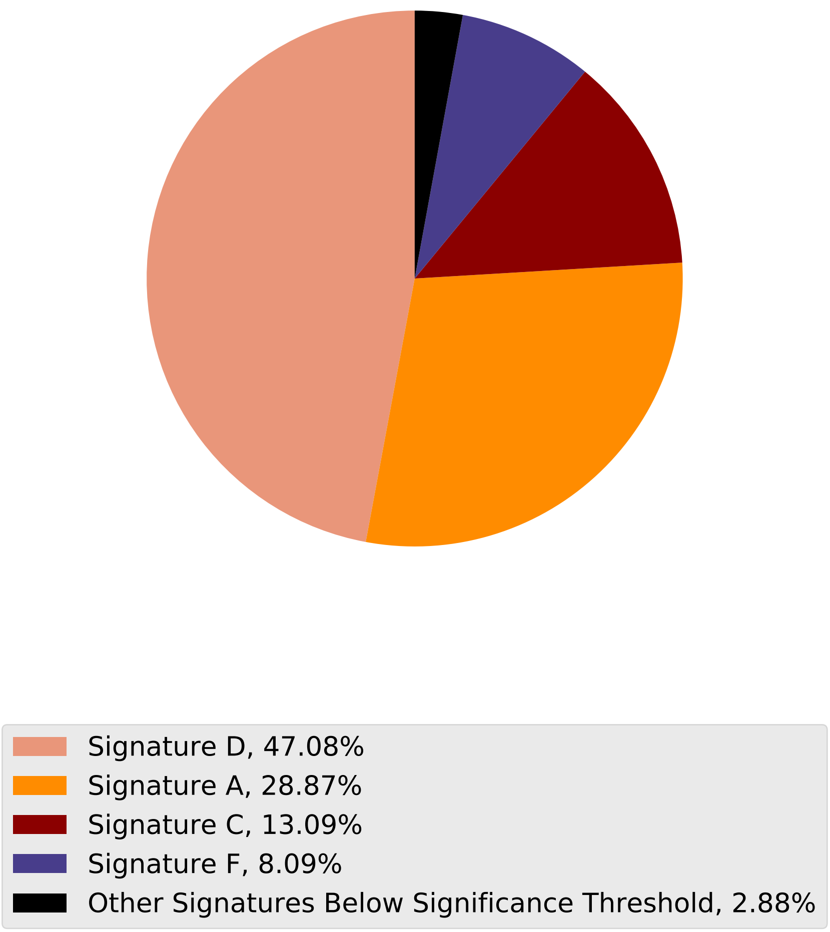

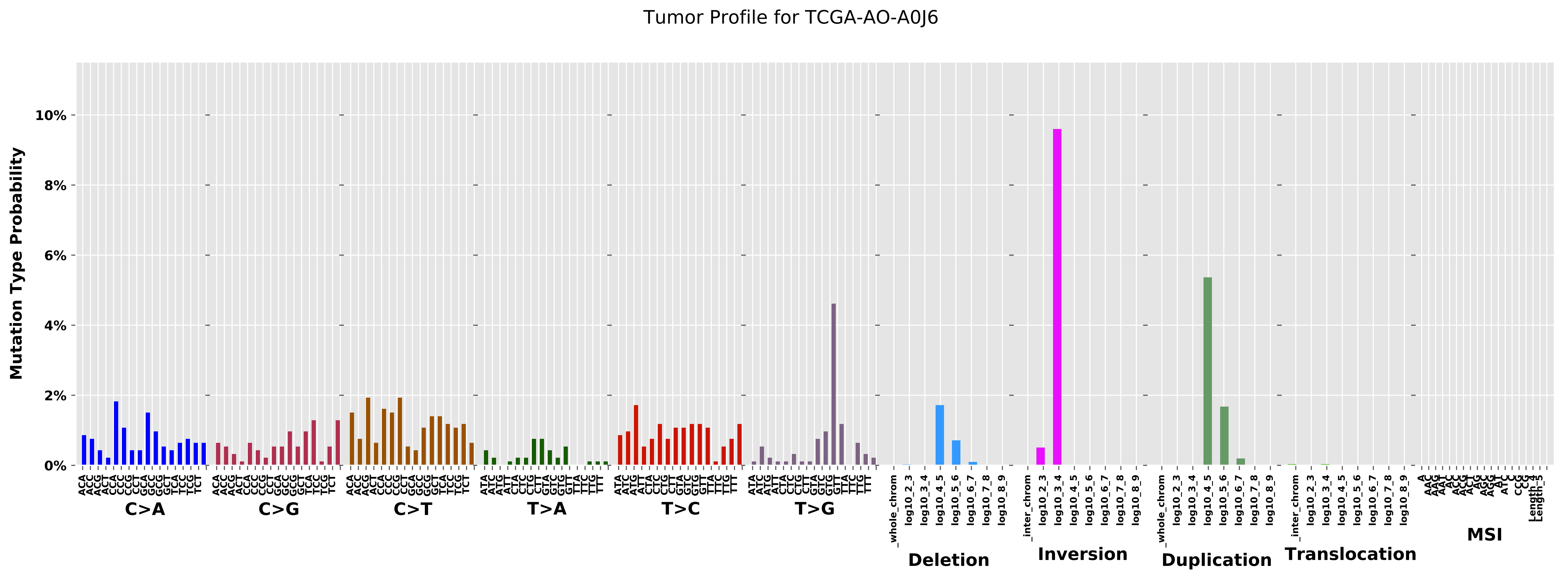

Tumor Profile for TCGA-AO-A0J6
Mutation Type Probability
0%
2%
4%
6%
8%
10%
C>A
C>G
C>T
T>A
T>C
T>G
Deletion
Inversion
Duplication
Translocation
MSI
_whole_chrom
_inter_chrom
log10_2_3
log10_3_4
log10_4_5
log10_5_6
log10_6_7
log10_7_8
log10_8_9

Cancer processes Weights for TCGA-A2-A3XX

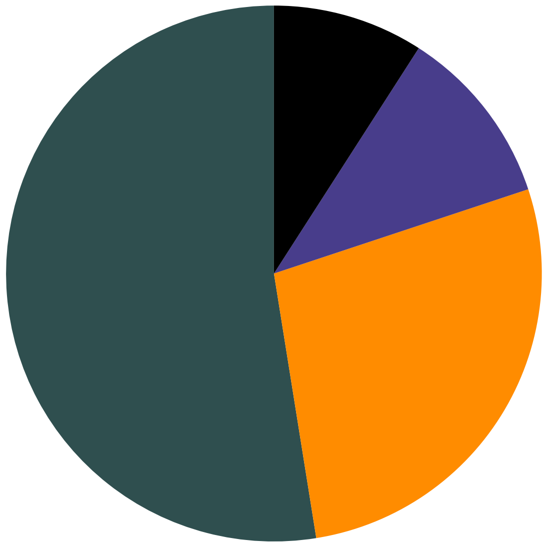

- Signature H, 52.53%
- Signature A, 27.56%
- Signature F, 10.81%
- Other Signatures Below Significance Threshold, 9.09%

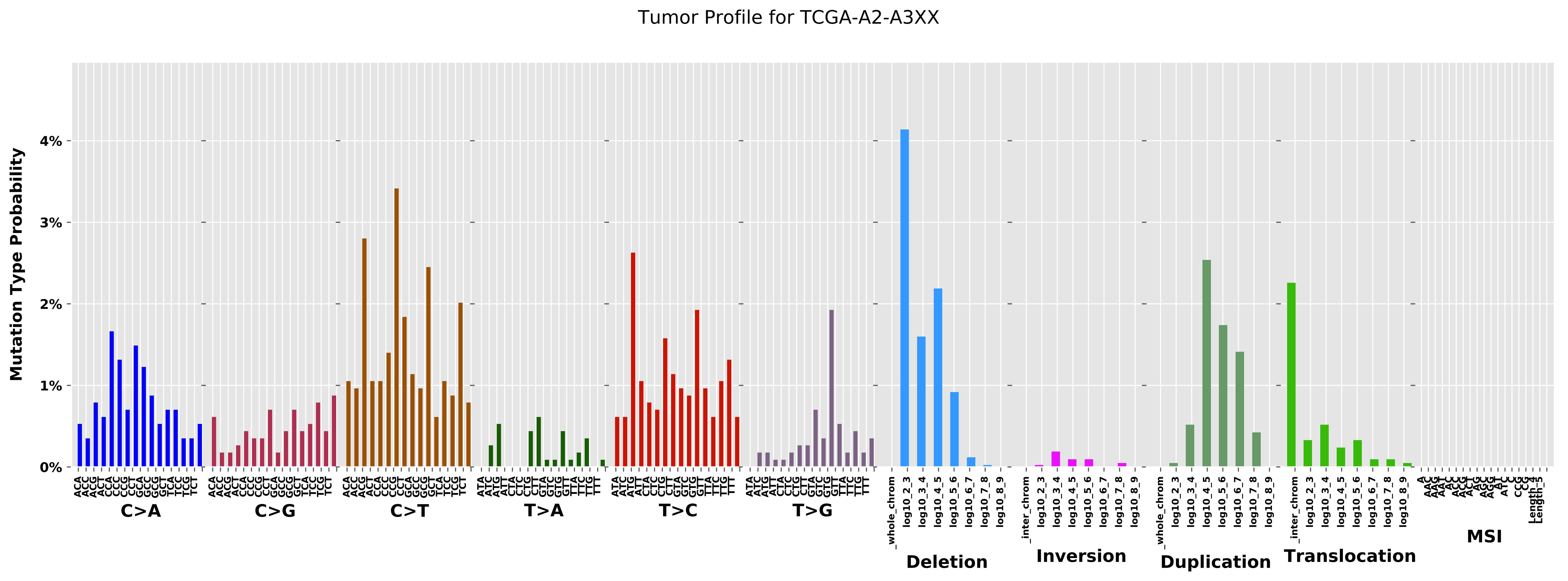

Tumor Profile for TCGA-A2-A3XX
Mutation Type Probability
0%
1%
2%
3%
4%
C>A
C>G
C>T
T>A
T>C
T>G
Deletion
Inversion
Duplication
Translocation
MSI

#### Cancer processes Weights for TCGA-A8-A075

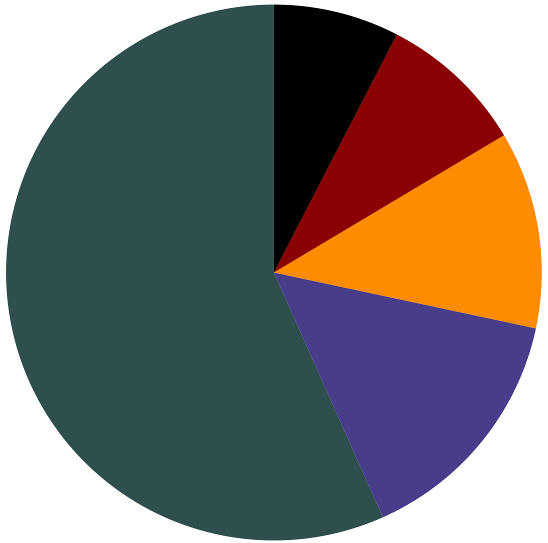

Signature H, 56.67%
Signature F, 14.99%
Signature A, 11.91%
Signature C, 8.85%
Other Signatures Below Significance Threshold, 7.59%

Tumor Profile for TCGA-A8-A075

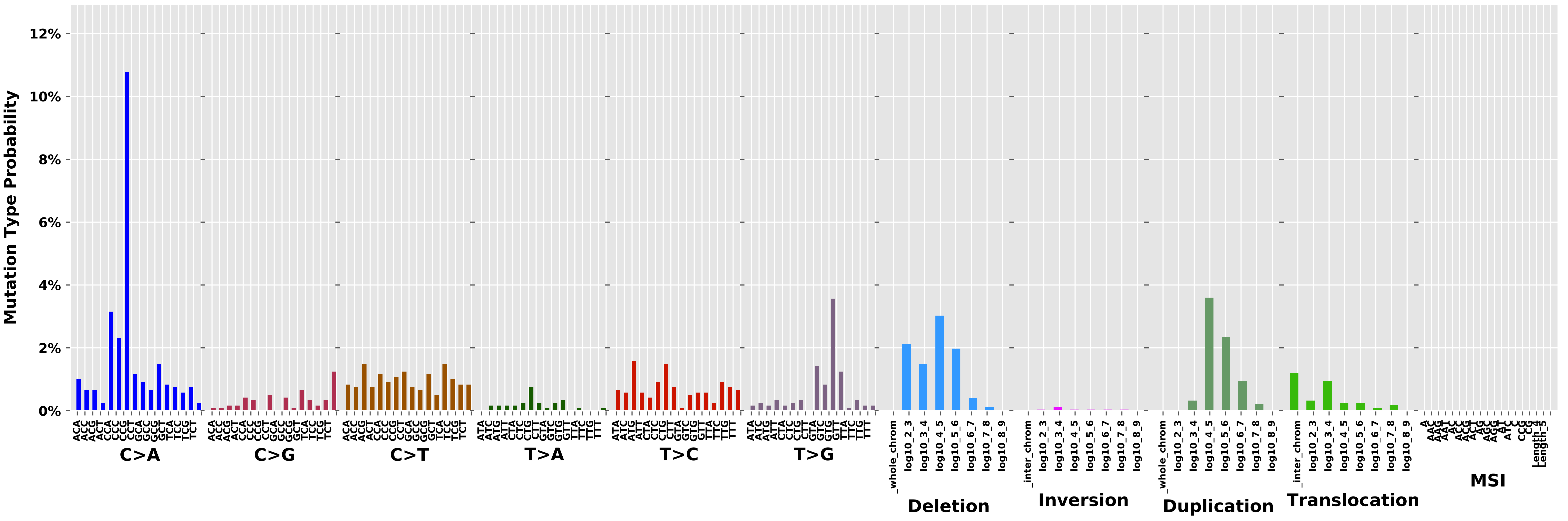

Cancer processes Weights for TCGA-AR-A1AY

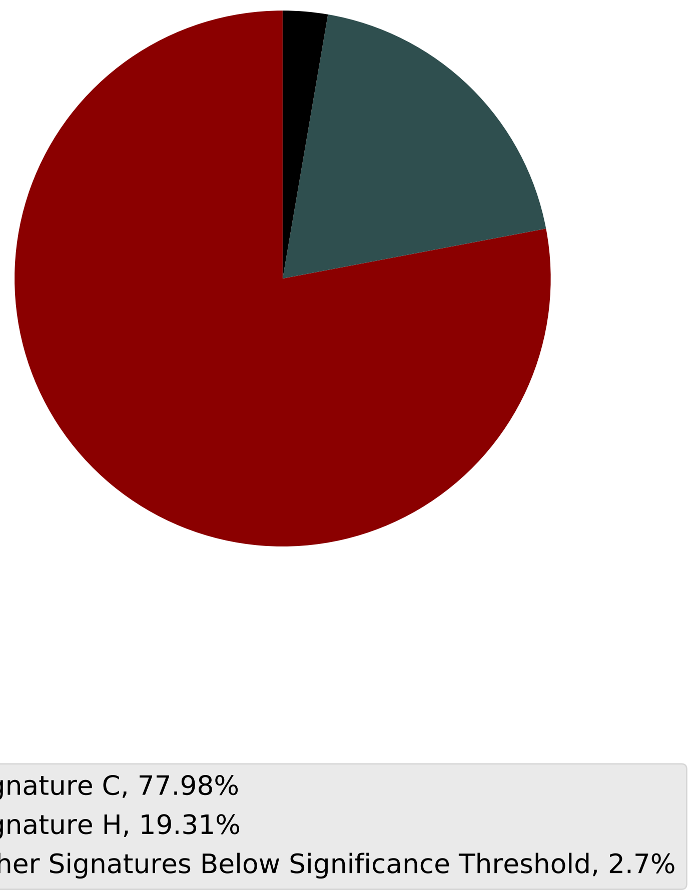

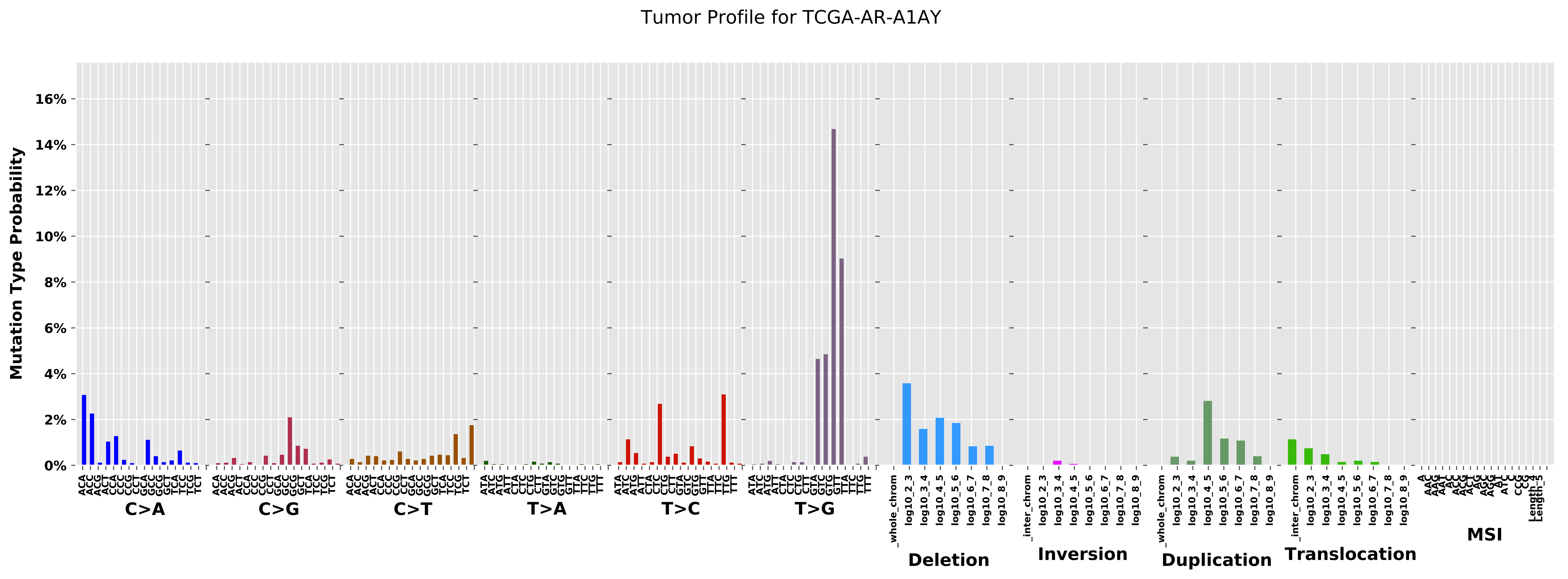

Tumor Profile for TCGA-AR-A1AY
Mutation Type Probability
0%
2%
4%
6%
8%
10%
12%
14%
16%
ACA ACC ACG ACT CCA CCC CCG CCT GCA GCC GCG GCT TCA TCC TCG TCT
C>A
ACA ACC ACG ACT CCA CCC CCG CCT GCA GCC GCG GCT TCA TCC TCG TCT
C>G
ACA ACC ACG ACT CCA CCC CCG CCT GCA GCC GCG GCT TCA TCC TCG TCT
C>T
ATA ATC ATG ATT CTA CTC CTG CTT GTA GTC GTG GTT TTA TTC TTG TTT
T>A
ATA ATC ATG ATT CTA CTC CTG CTT GTA GTC GTG GTT TTA TTC TTG TTT
T>C
ATA ATC ATG ATT CTA CTC CTG CTT GTA GTC GTG GTT TTA TTC TTG TTT
T>G
_whole_chrom log10_2_3 log10_3_4 log10_4_5 log10_5_6 log10_6_7 log10_7_8 log10_8_9
Deletion
_inter_chrom log10_2_3 log10_3_4 log10_4_5 log10_5_6 log10_6_7 log10_7_8 log10_8_9
Inversion
_whole_chrom log10_2_3 log10_3_4 log10_4_5 log10_5_6 log10_6_7 log10_7_8 log10_8_9
Duplication
_inter_chrom log10_2_3 log10_3_4 log10_4_5 log10_5_6 log10_6_7 log10_7_8 log10_8_9
Translocation
MSI

Cancer processes Weights for TCGA-AG-3901

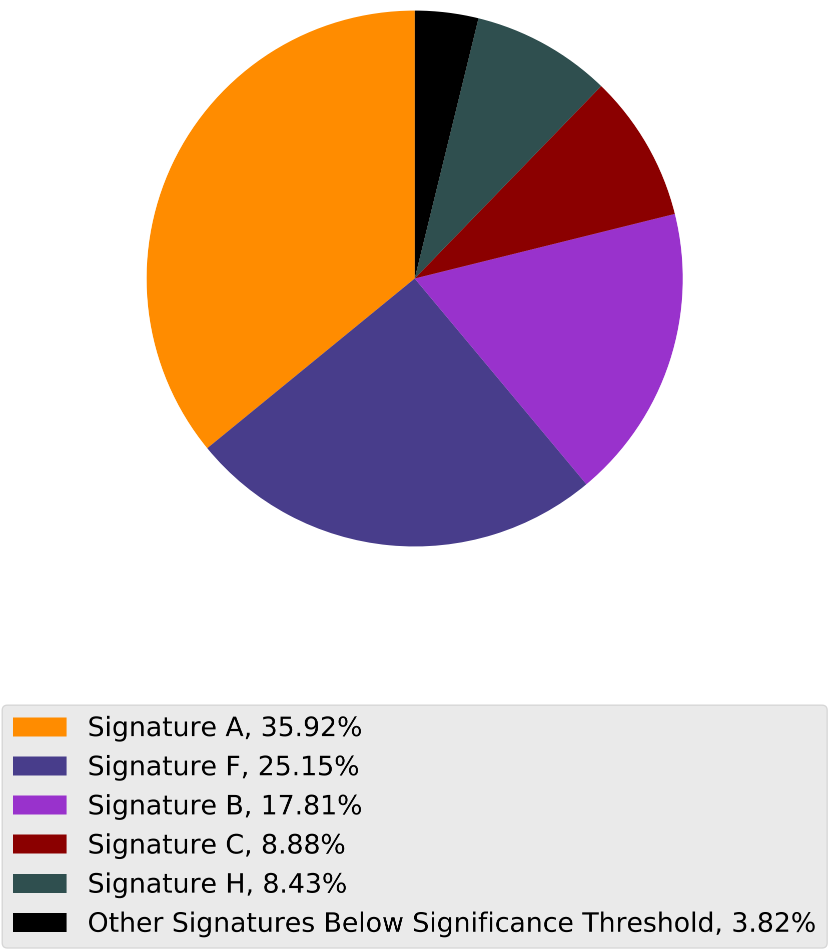

Tumor Profile for TCGA-AG-3901

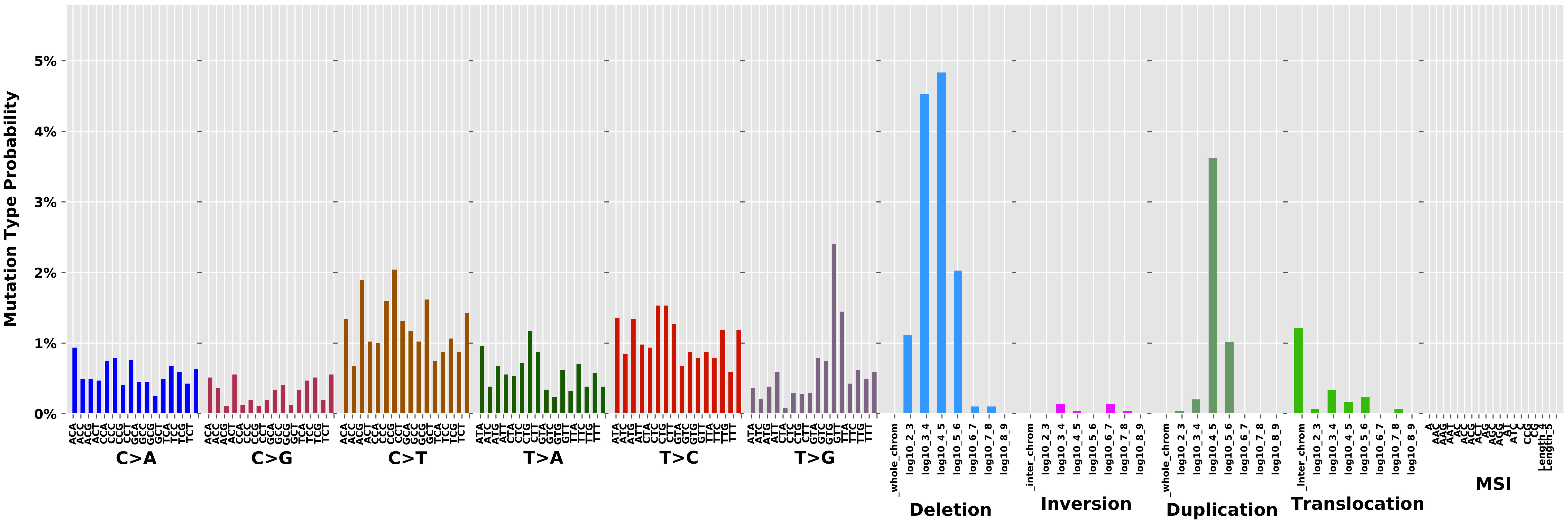

Cancer processes Weights for TCGA-E2-A152

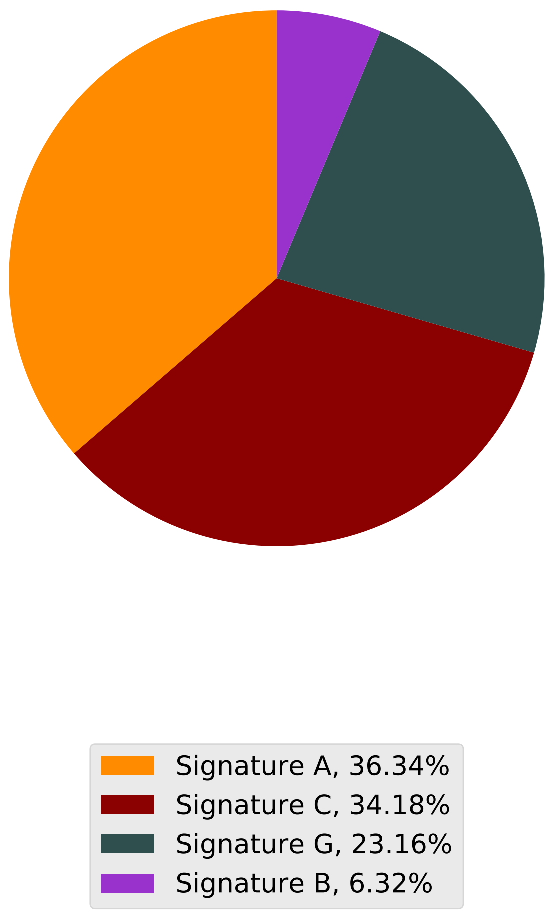

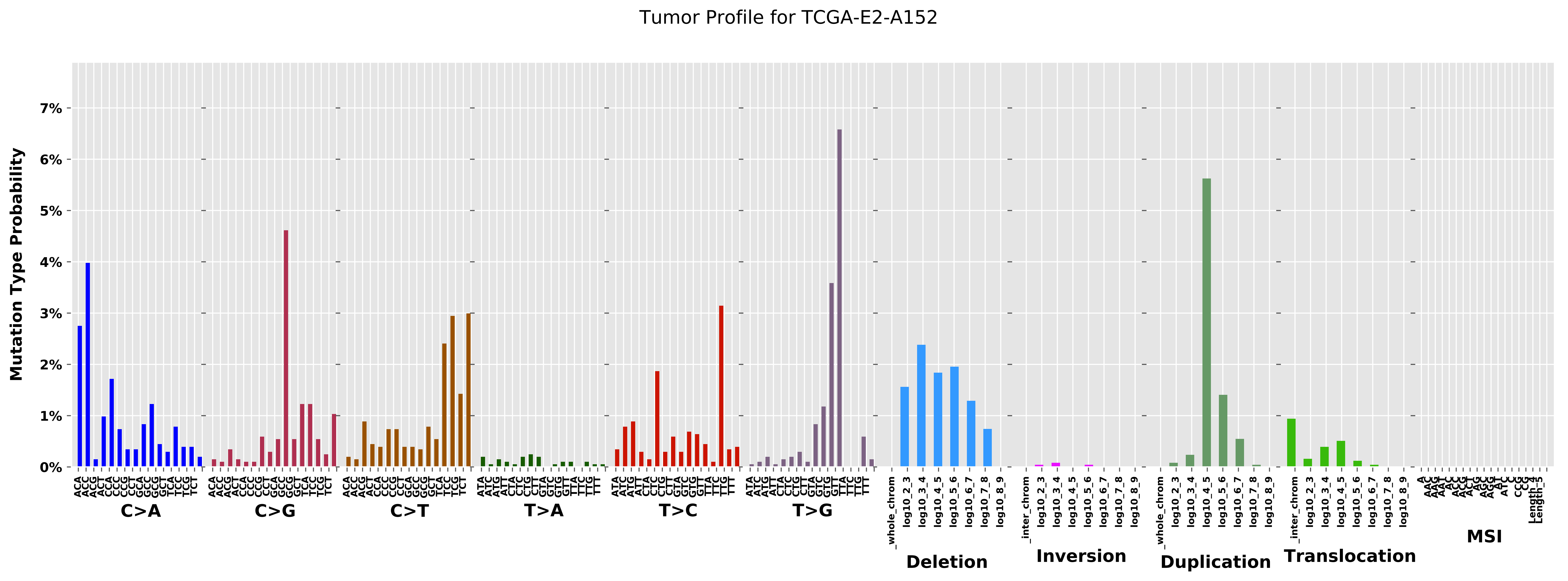
Tumor Profile for TCGA-E2-A152
Mutation Type Probability
C>A
C>G
C>T
T>A
T>C
T>G
Deletion
Inversion
Duplication
Translocation
MSI

Cancer processes Weights for TCGA-AG-4015

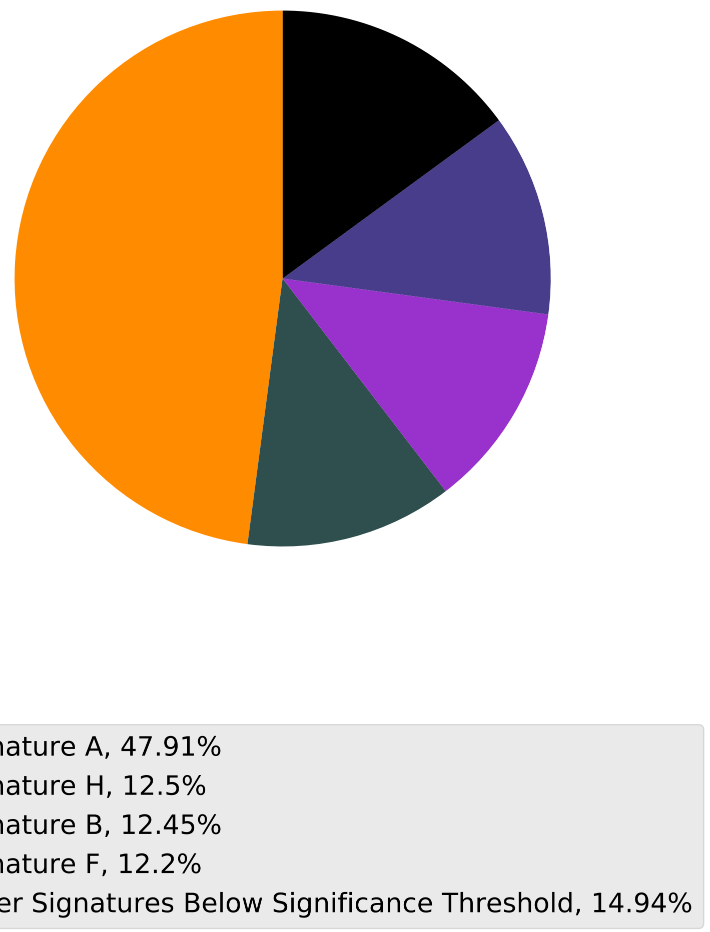

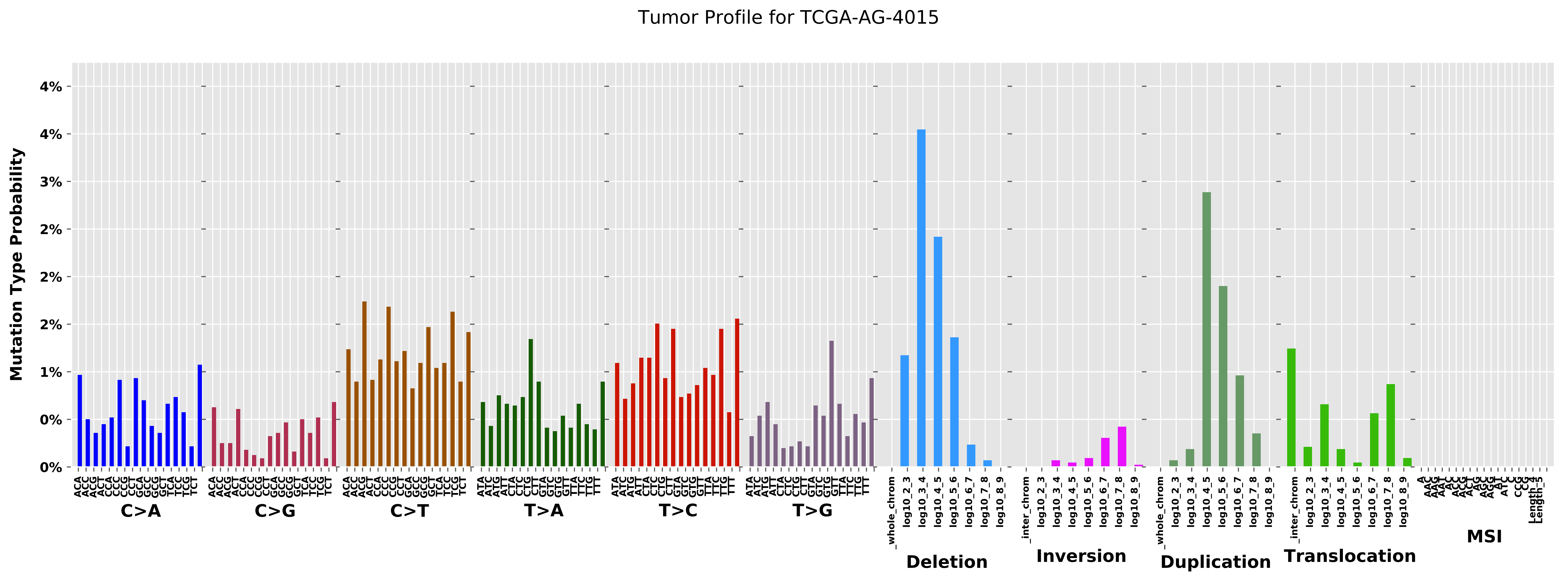

Tumor Profile for TCGA-AG-4015
Mutation Type Probability
0%
0%
1%
2%
2%
3%
4%
4%
C>A
C>G
C>T
T>A
T>C
T>G
_whole_chrom
log10_2_3
log10_3_4
log10_4_5
log10_5_6
log10_6_7
log10_7_8
log10_8_9
Deletion
_inter_chrom
Inversion
Duplication
Translocation
MSI

#### Cancer processes Weights for TCGA-BH-A0B9

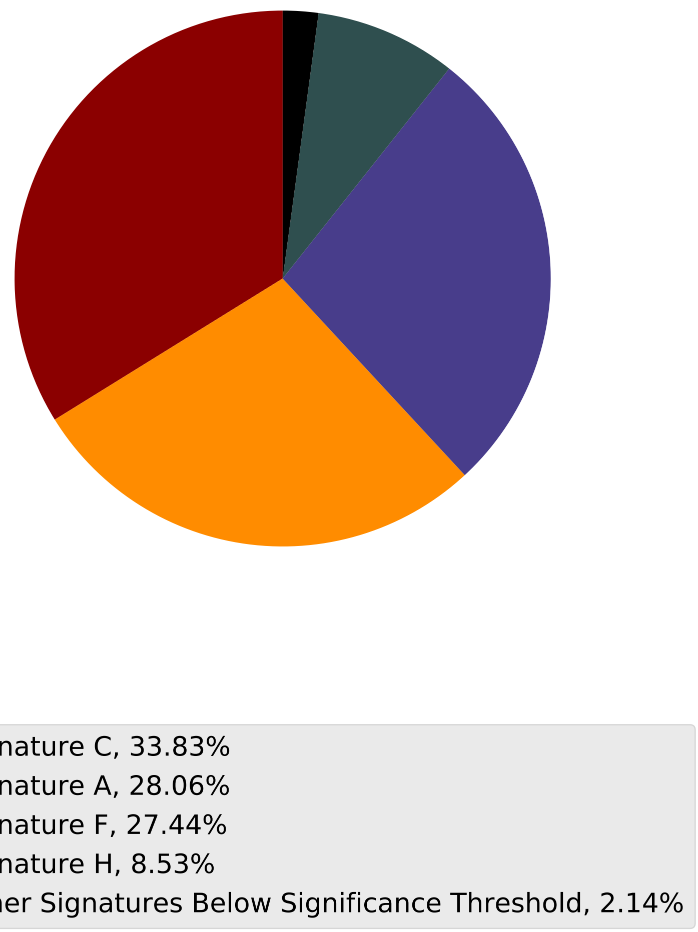

Tumor Profile for TCGA-BH-A0B9

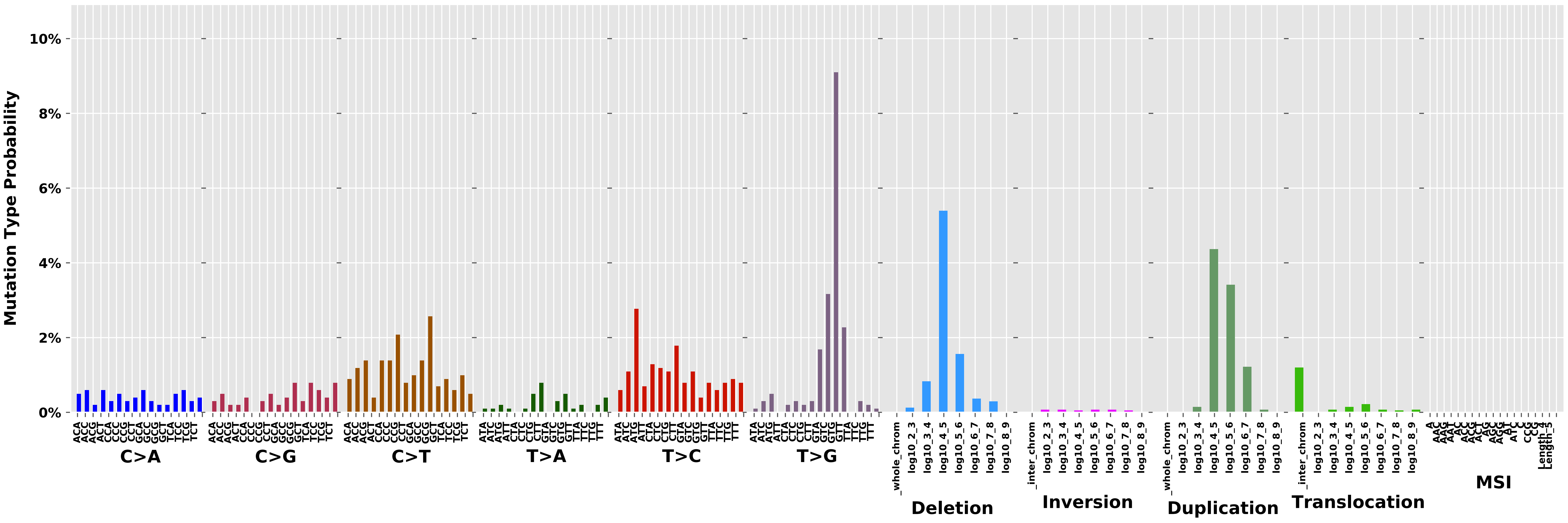

Cancer processes Weights for TCGA-A2-A0D1

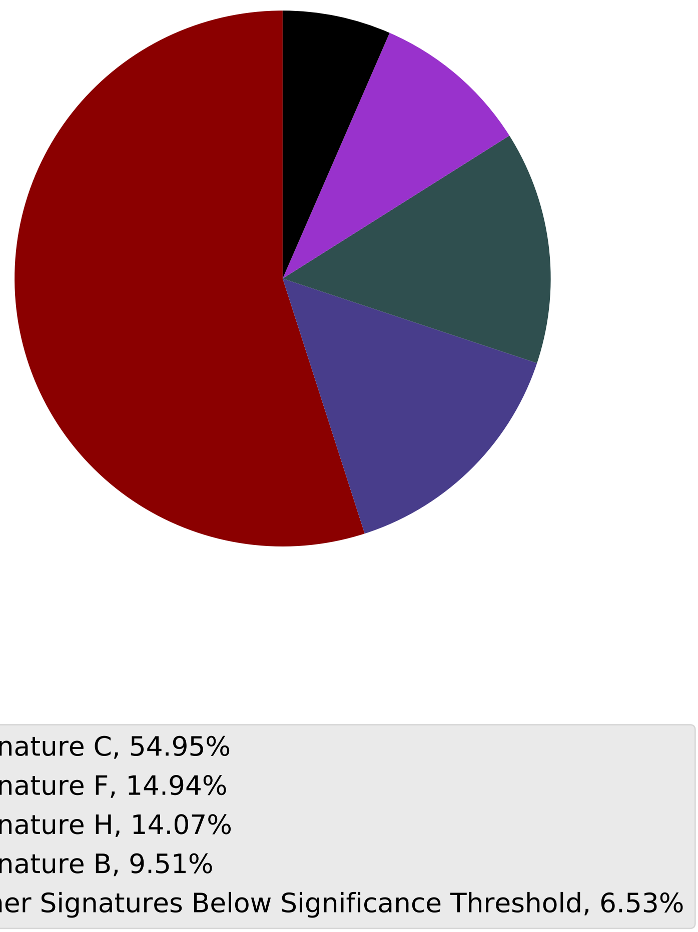

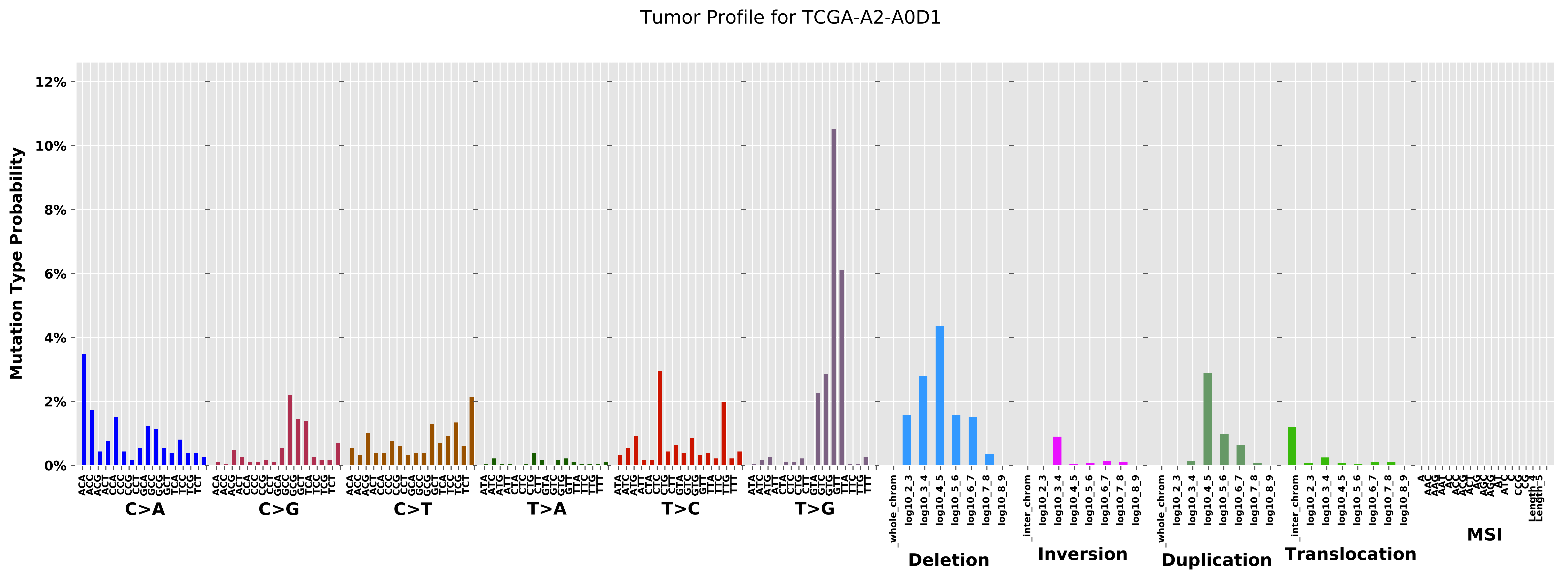
Tumor Profile for TCGA-A2-A0D1
Mutation Type Probability
0%
2%
4%
6%
8%
10%
12%
C>A
C>G
C>T
T>A
T>C
T>G
Deletion
Inversion
Duplication
Translocation
MSI
_whole_chrom
_inter_chrom
log10_2_3
log10_3_4
log10_4_5
log10_5_6
log10_6_7
log10_7_8
log10_8_9

Cancer processes Weights for TCGA-AA-3529

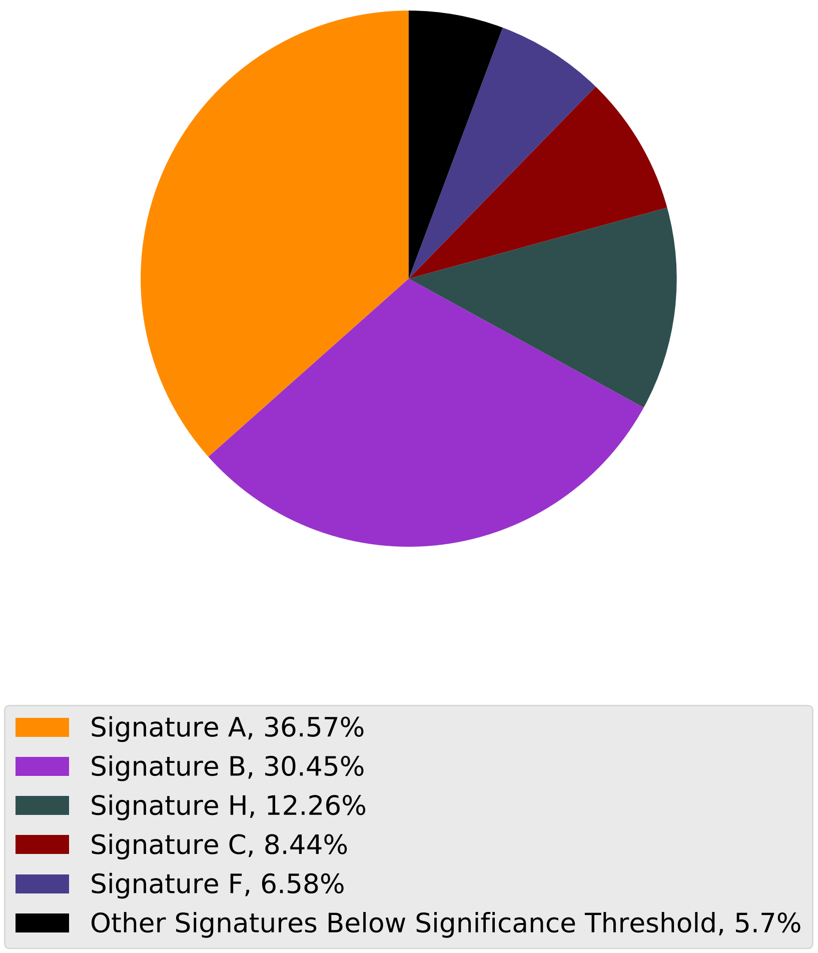

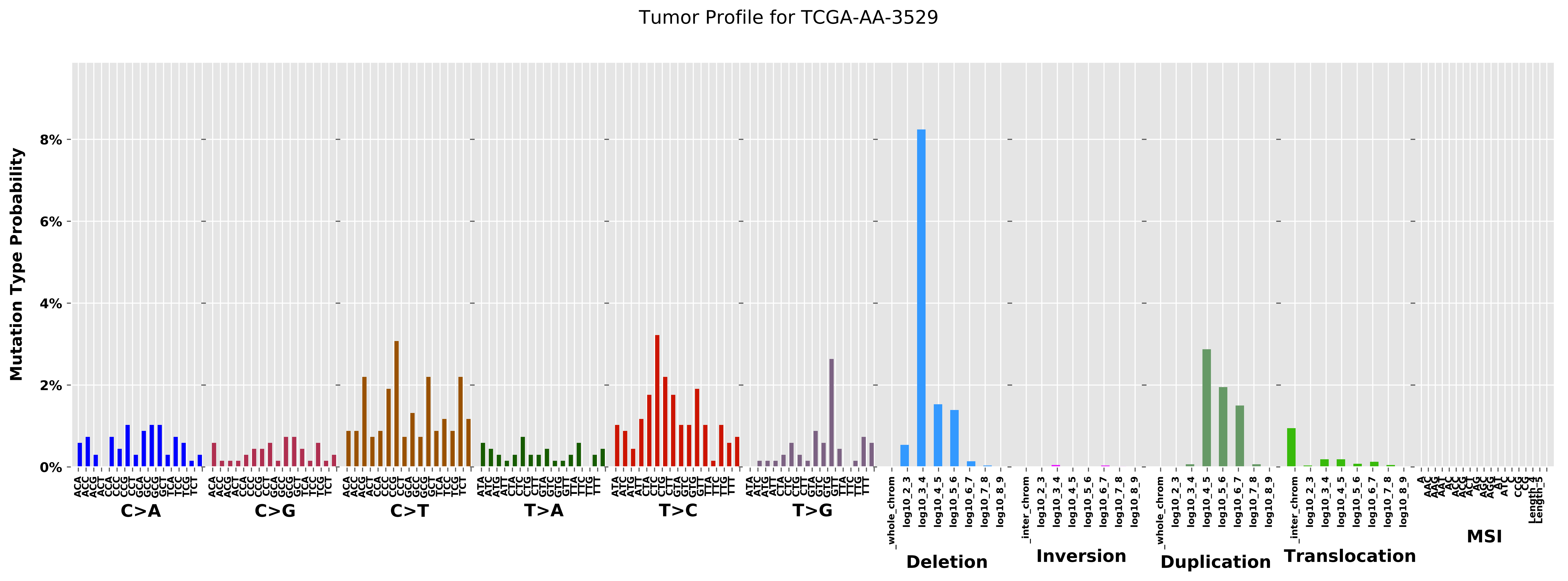
Tumor Profile for TCGA-AA-3529
Mutation Type Probability
0%
2%
4%
6%
8%
C>A
C>G
C>T
T>A
T>C
T>G
Deletion
Inversion
Duplication
Translocation
MSI

Cancer processes Weights for TCGA-A6-3807

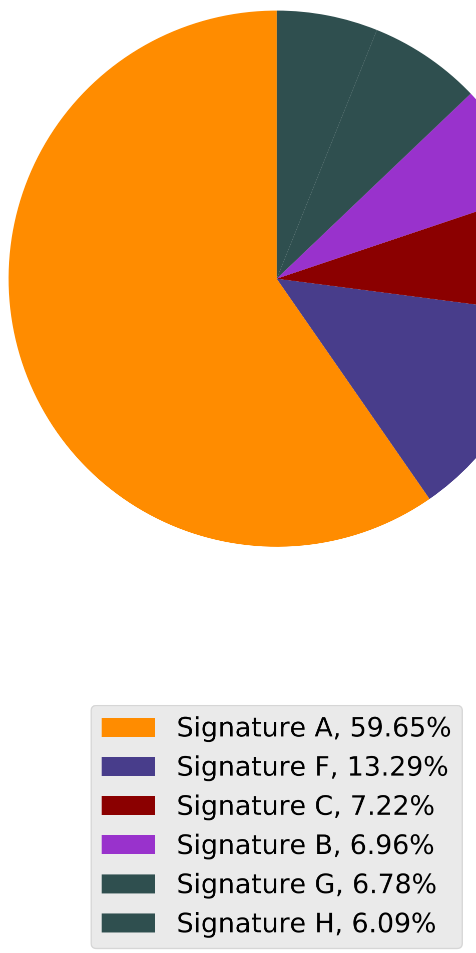

Tumor Profile for TCGA-A6-3807

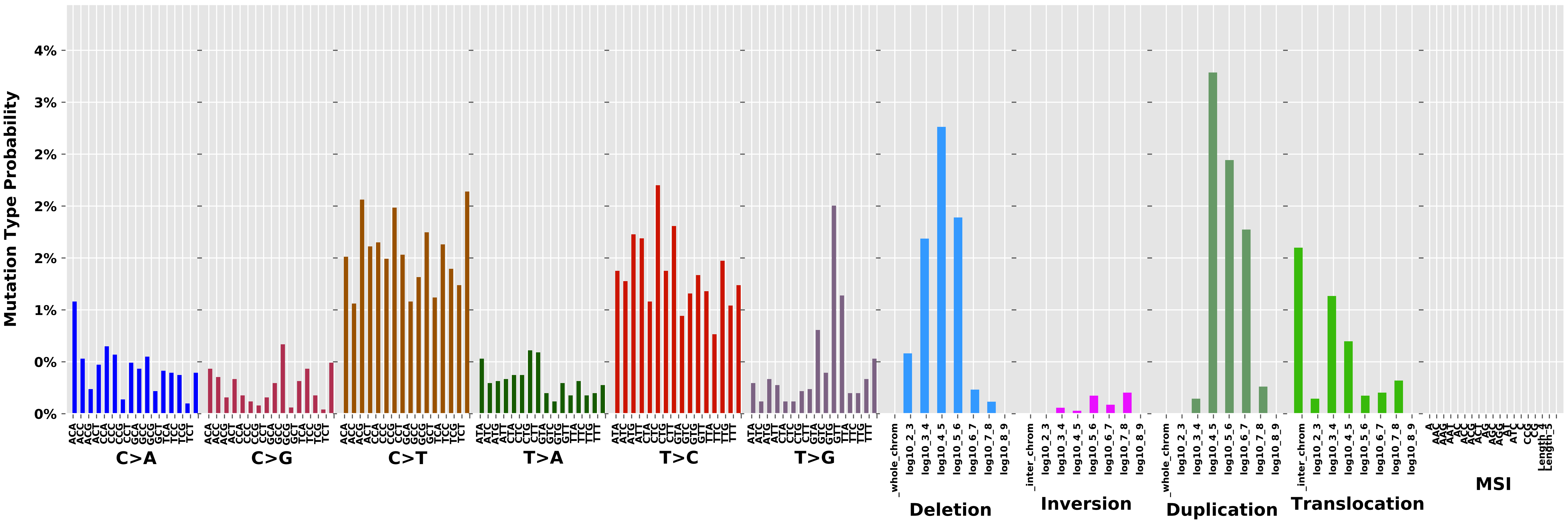

Cancer processes Weights for TCGA-A2-A0YG

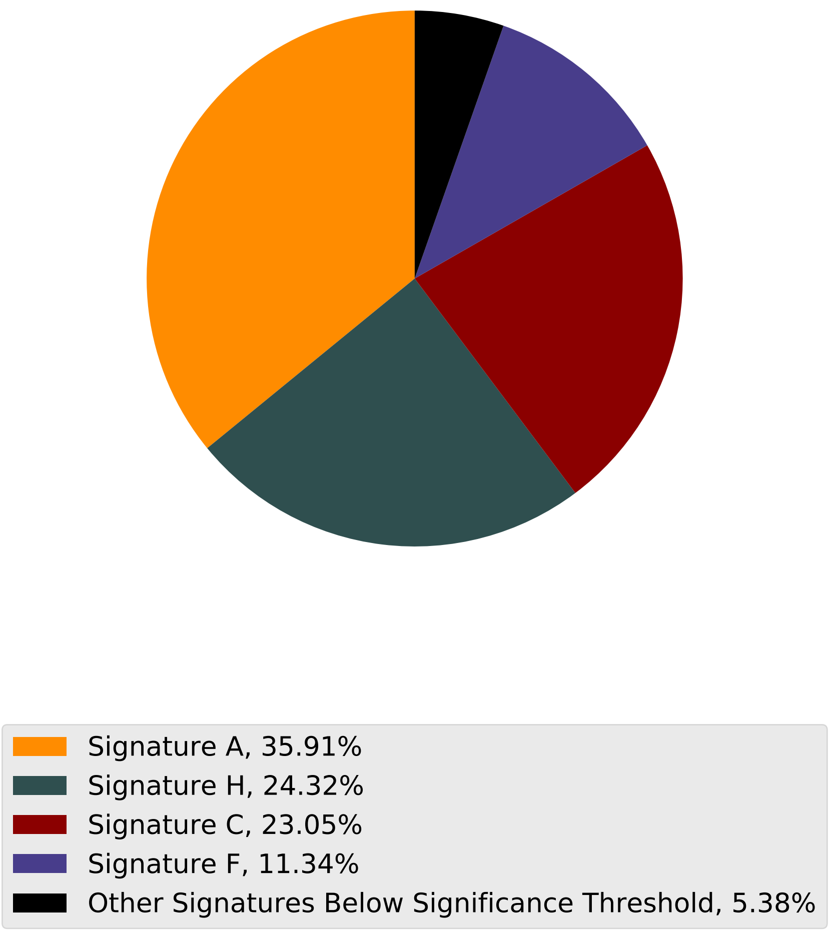

Tumor Profile for TCGA-A2-A0YG

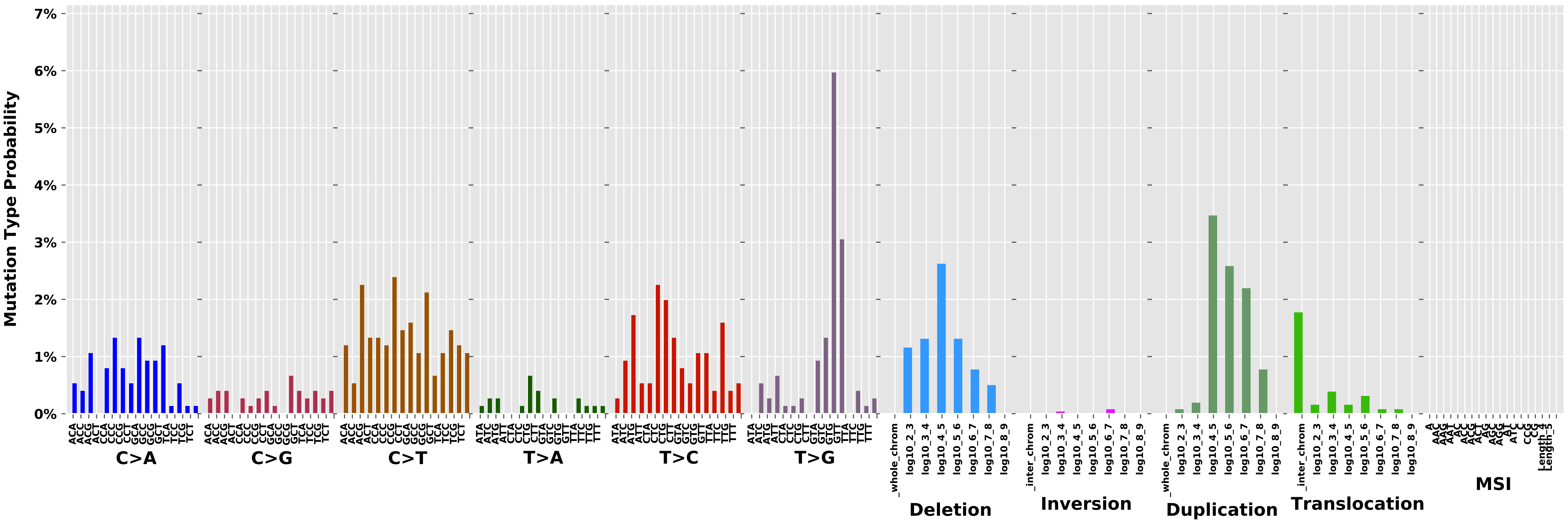

#### Cancer processes Weights for TCGA-C8-A12Q

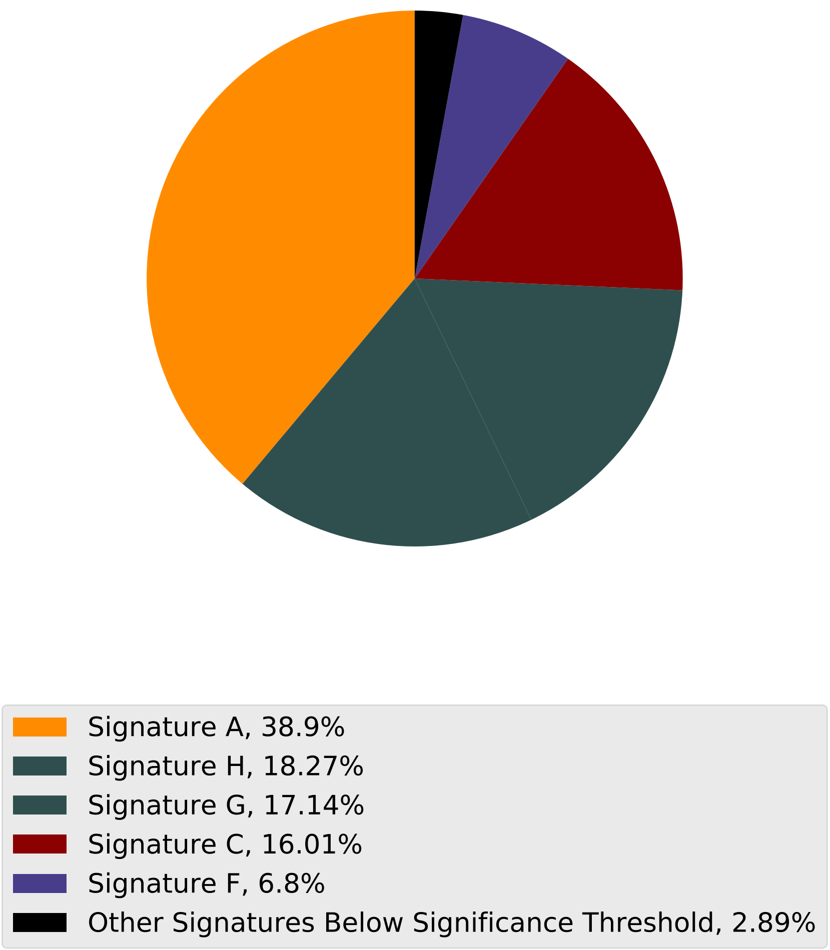

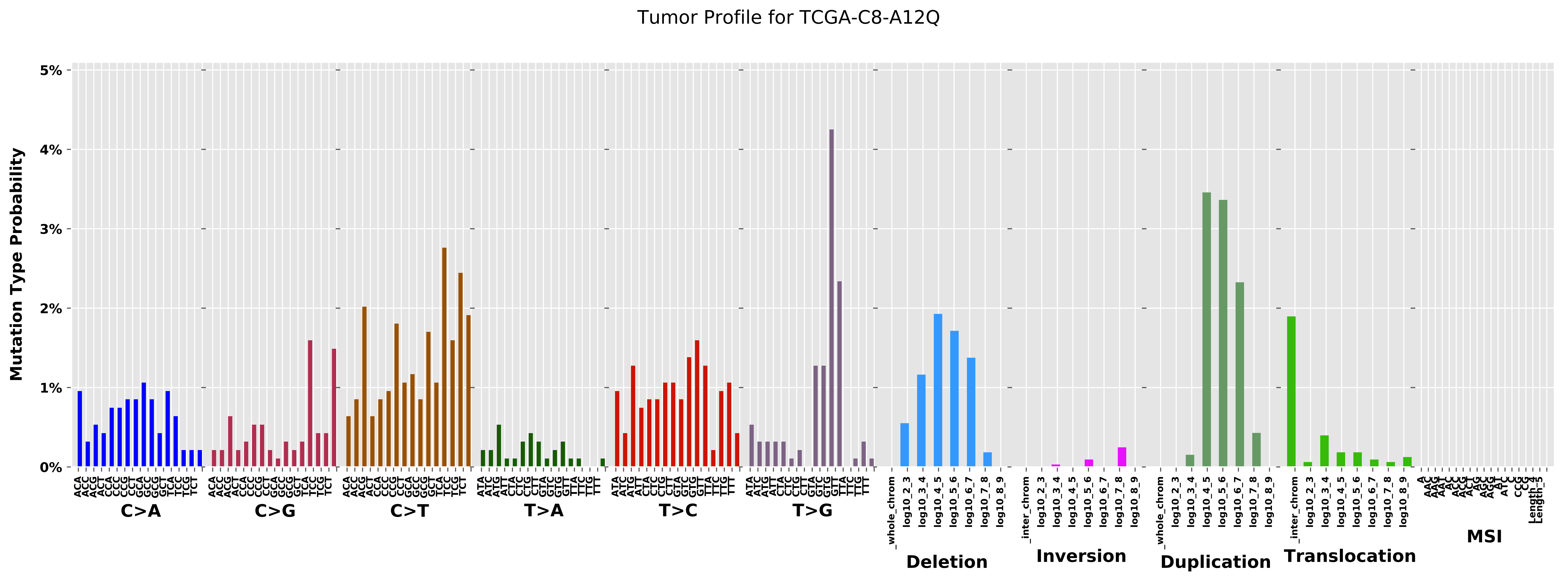
Tumor Profile for TCGA-C8-A12Q
Mutation Type Probability
0%
1%
2%
3%
4%
5%
C>A
C>G
C>T
T>A
T>C
T>G
Deletion
Inversion
Duplication
Translocation
MSI
_whole_chrom
_inter_chrom
log10_2_3
log10_3_4
log10_4_5
log10_5_6
log10_6_7
log10_7_8
log10_8_9

Cancer processes Weights for TCGA-BH-A0BW

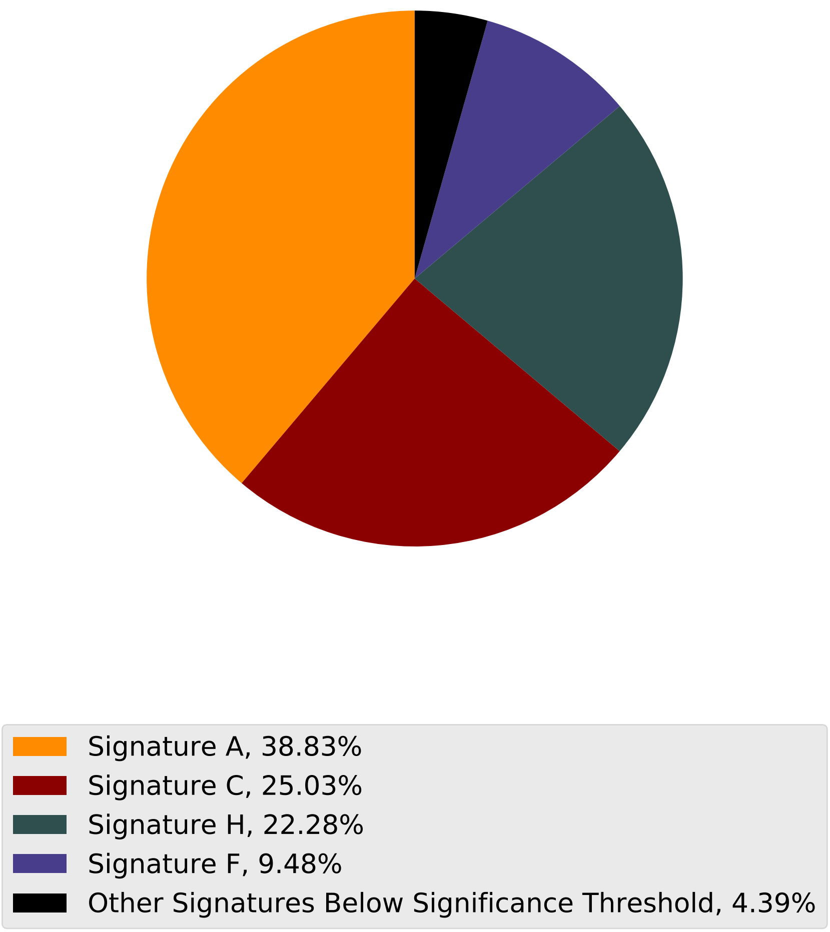

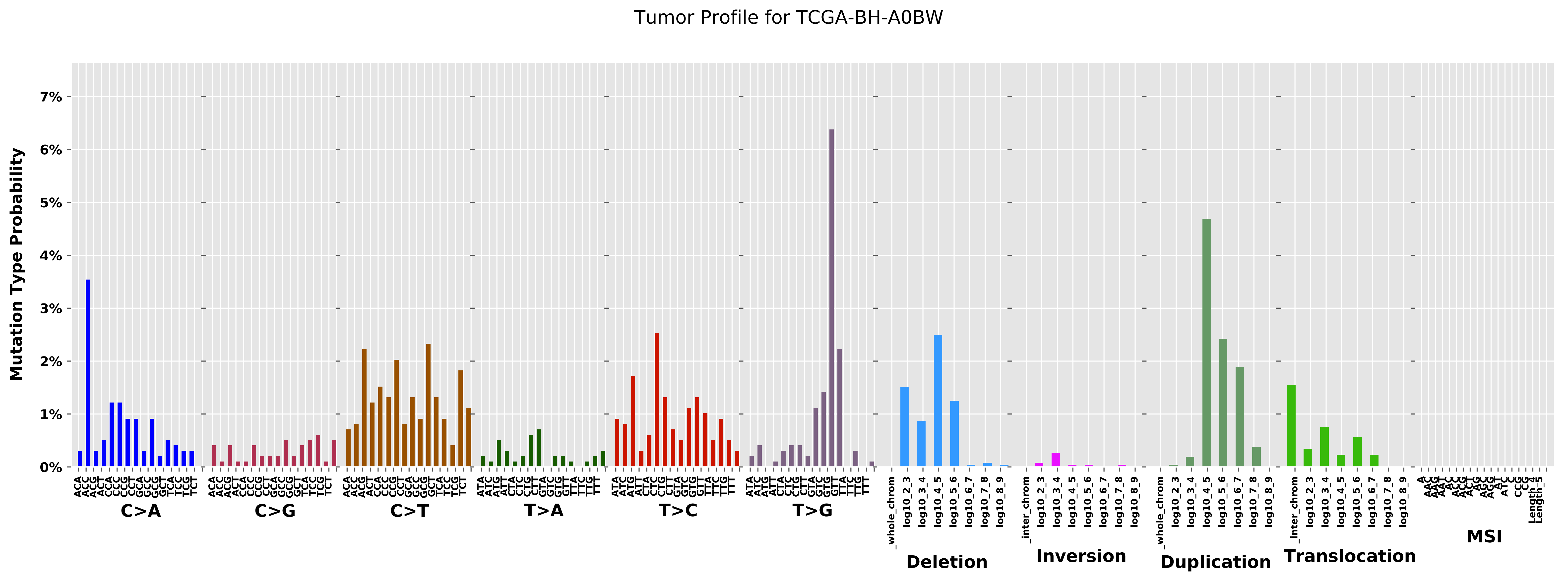

Tumor Profile for TCGA-BH-A0BW
Mutation Type Probability
7%
6%
5%
4%
3%
2%
1%
0%
C>A
C>G
C>T
T>A
T>C
T>G
Deletion
Inversion
Duplication
Translocation
MSI
_whole_chrom
_inter_chrom
log10_2_3
log10_3_4
log10_4_5
log10_5_6
log10_6_7
log10_7_8
log10_8_9

Cancer processes Weights for TCGA-A2-A0EY

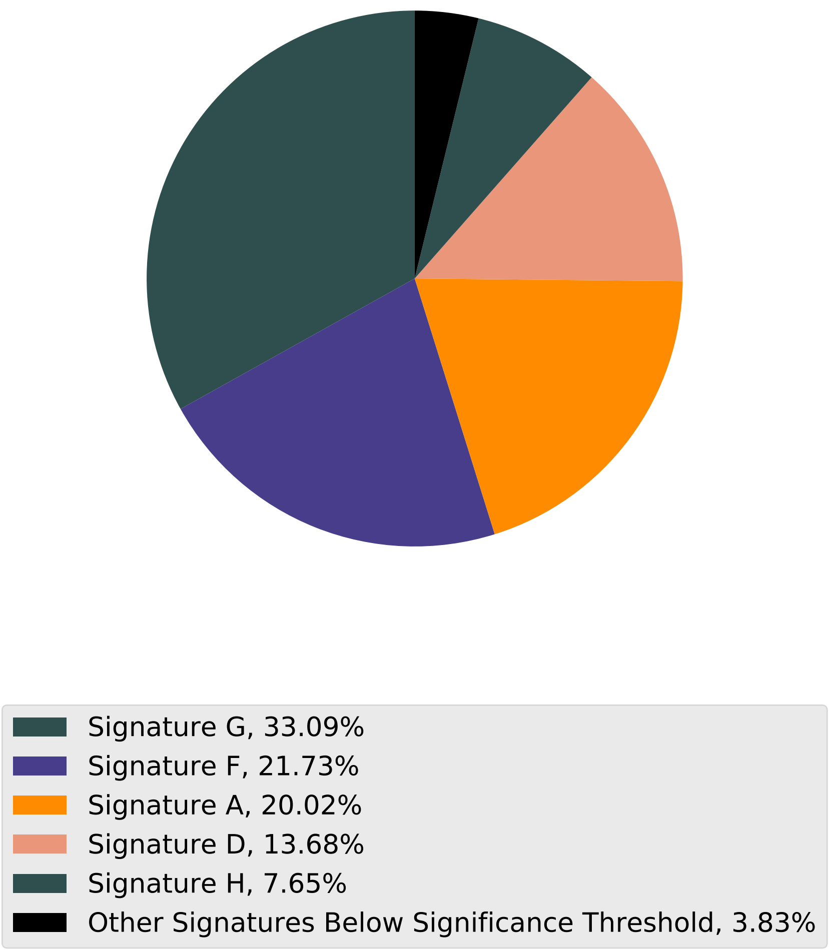

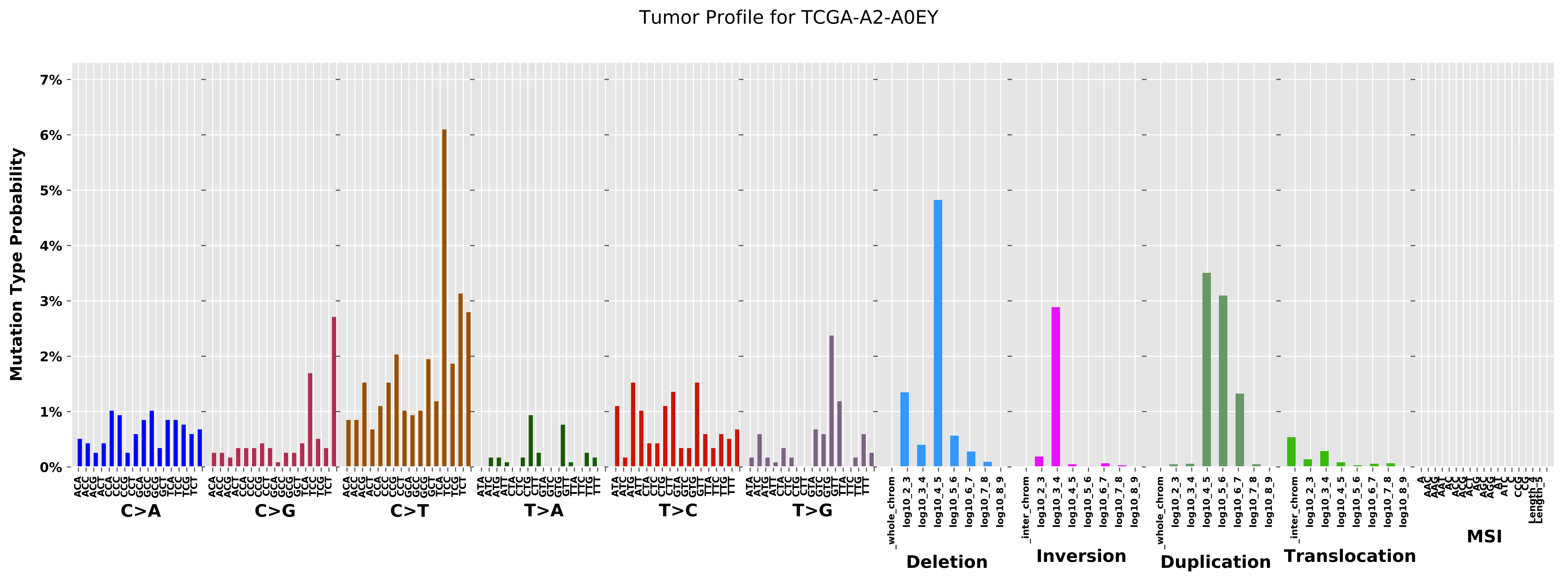

Tumor Profile for TCGA-A2-A0EY
Mutation Type Probability
0%
1%
2%
3%
4%
5%
6%
7%
C>A
C>G
C>T
T>A
T>C
T>G
Deletion
Inversion
Duplication
Translocation
MSI

Cancer processes Weights for TCGA-EW-A1PB

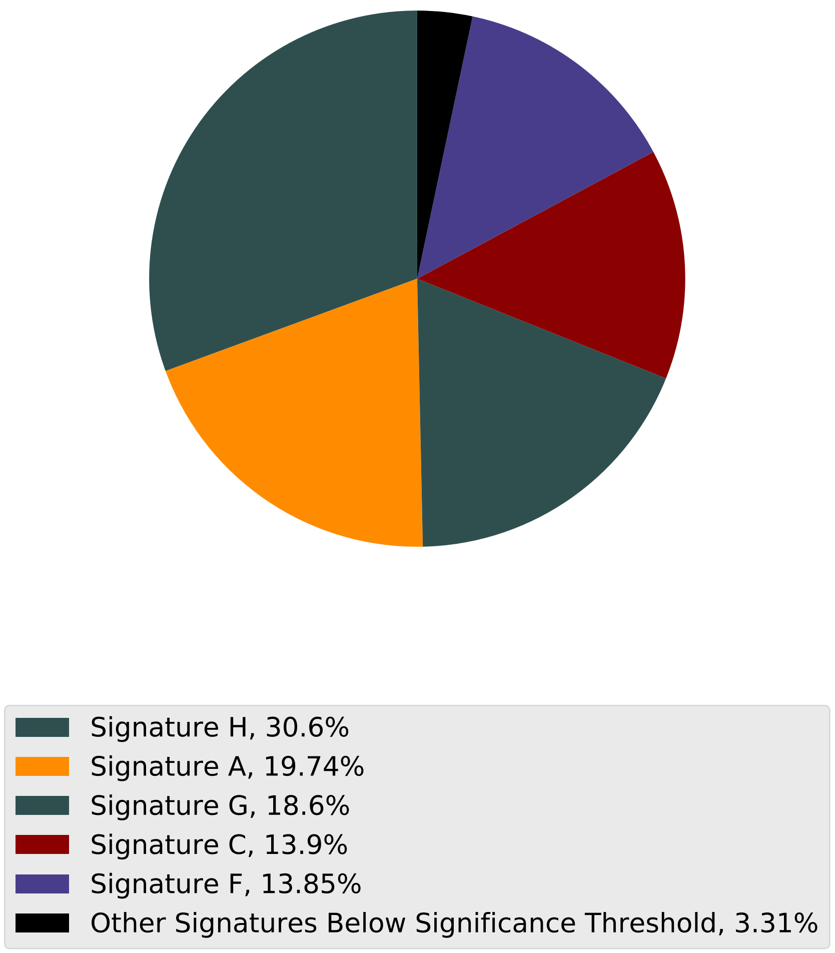

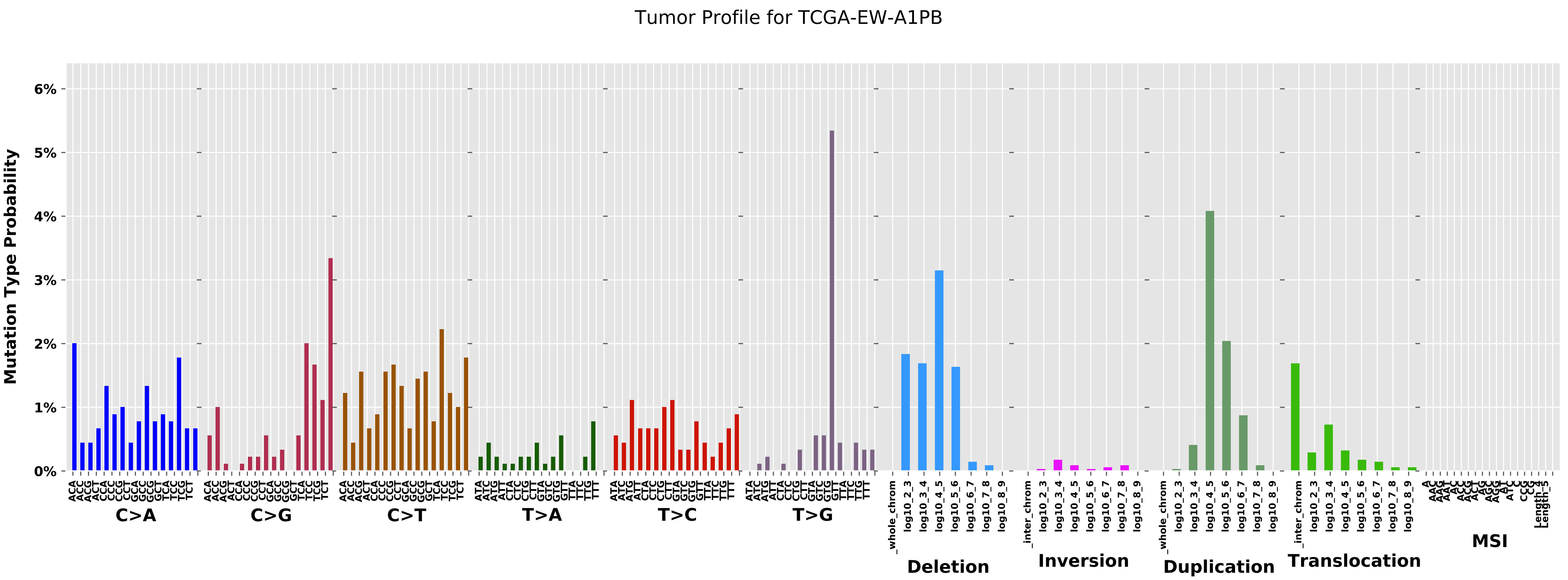

Tumor Profile for TCGA-EW-A1PB
Mutation Type Probability
0%
1%
2%
3%
4%
5%
6%
C>A
C>G
C>T
T>A
T>C
T>G
Deletion
Inversion
Duplication
Translocation
MSI

Cancer processes Weights for TCGA-AD-6964

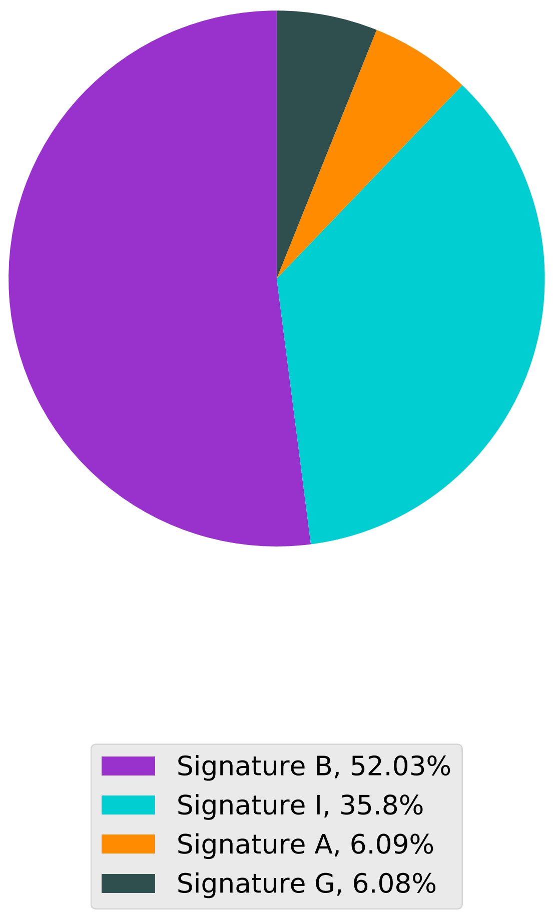

Tumor Profile for TCGA-AD-6964

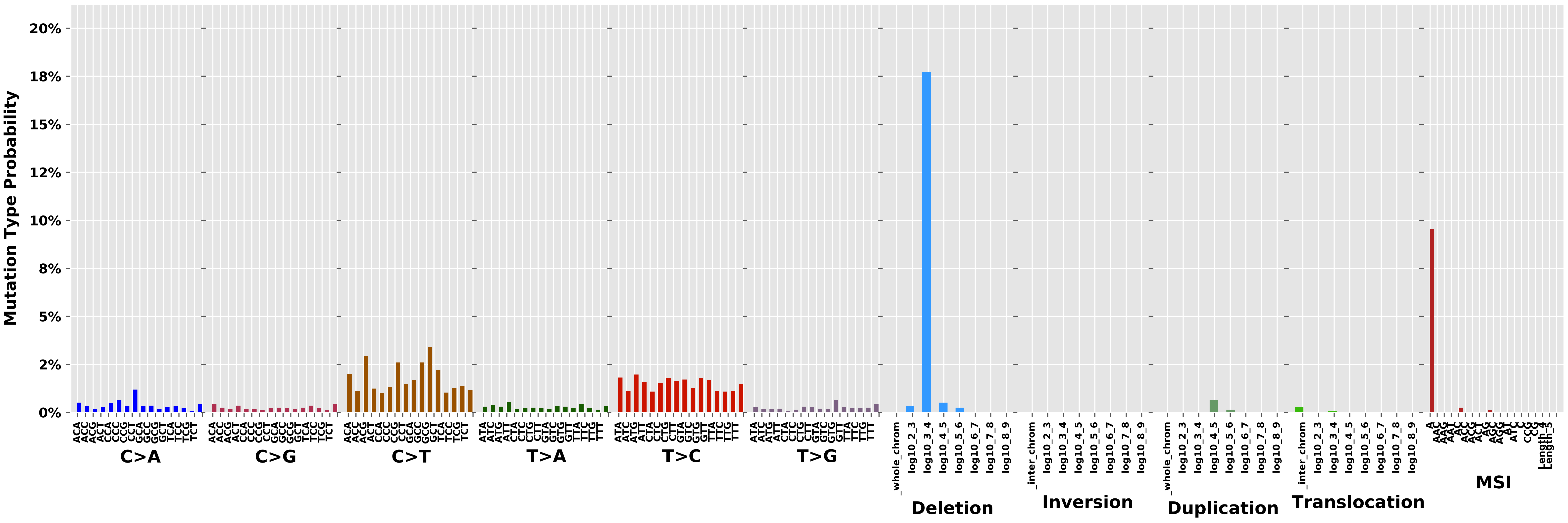

Cancer processes Weights for TCGA-AG-3896

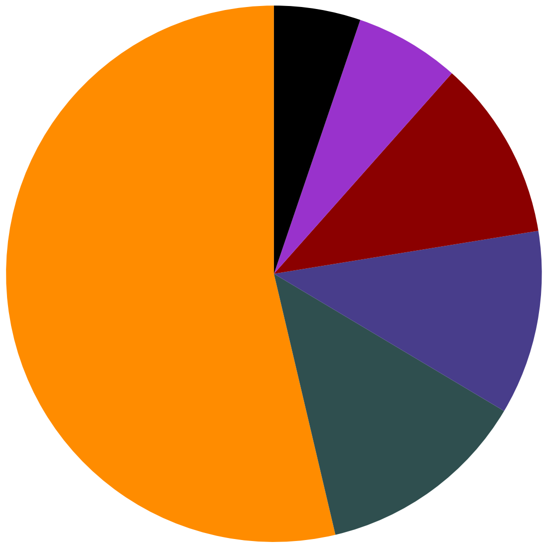

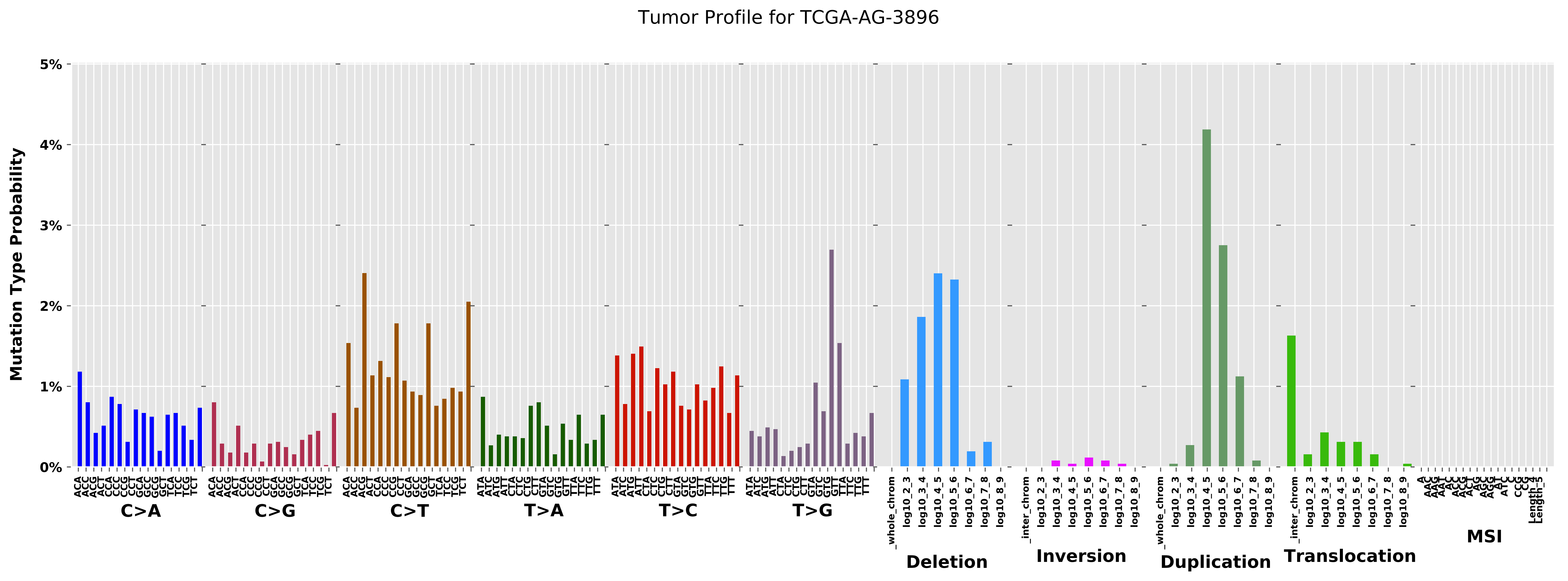

Tumor Profile for TCGA-AG-3896
Mutation Type Probability
0%
1%
2%
3%
4%
5%
C>A
C>G
C>T
T>A
T>C
T>G
Deletion
Inversion
Duplication
Translocation
MSI

Cancer processes Weights for TCGA-E2-A15K

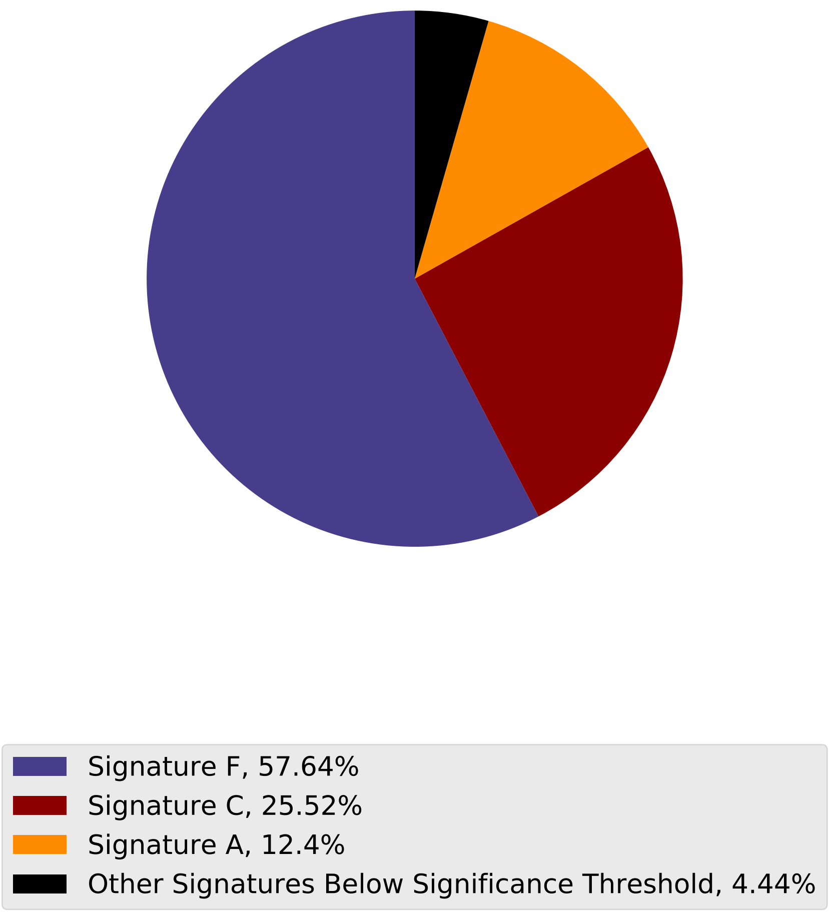

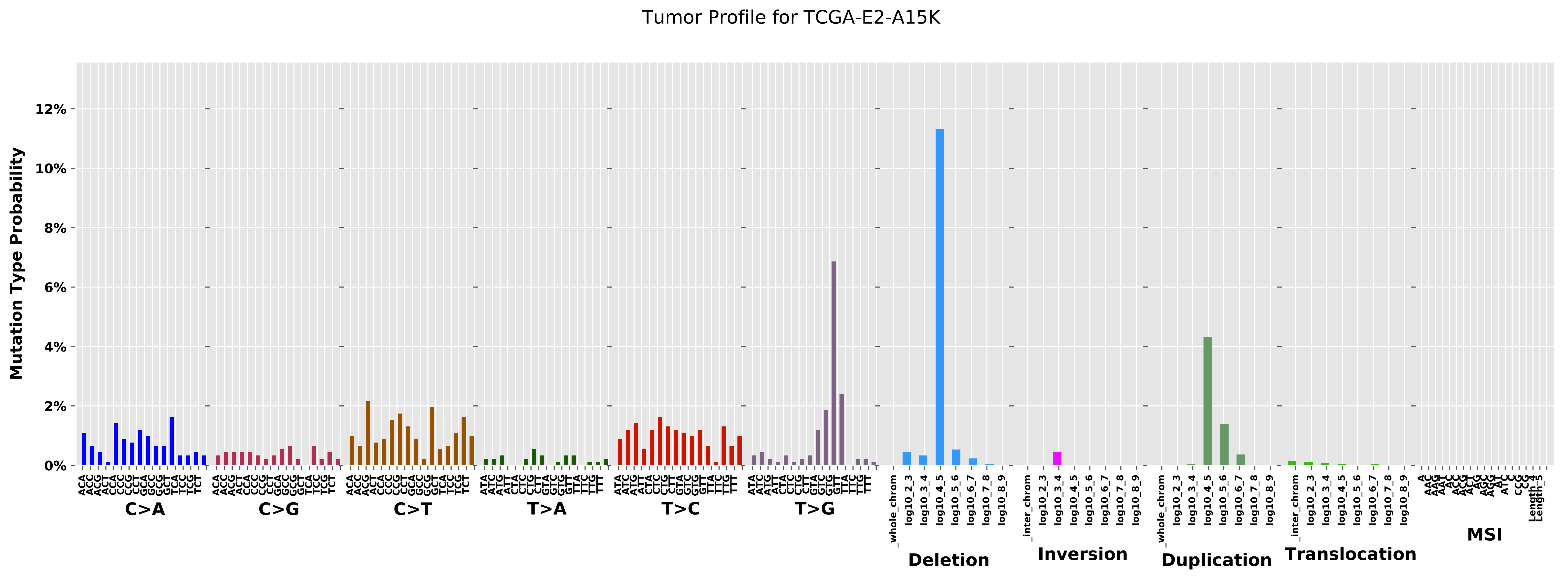
Tumor Profile for TCGA-E2-A15K
Mutation Type Probability
12%
10%
8%
6%
4%
2%
0%
C>A
C>G
C>T
T>A
T>C
T>G
Deletion
Inversion
Duplication
Translocation
MSI
_whole_chrom
_inter_chrom
log10_2_3
log10_3_4
log10_4_5
log10_5_6
log10_6_7
log10_7_8
log10_8_9

Cancer processes Weights for TCGA-B6-A0RE

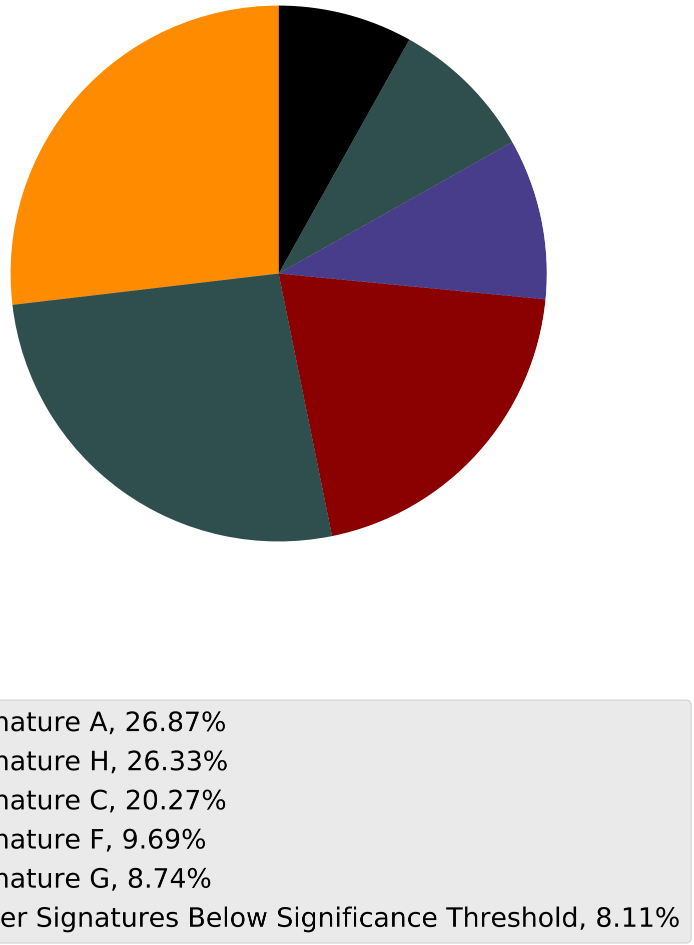

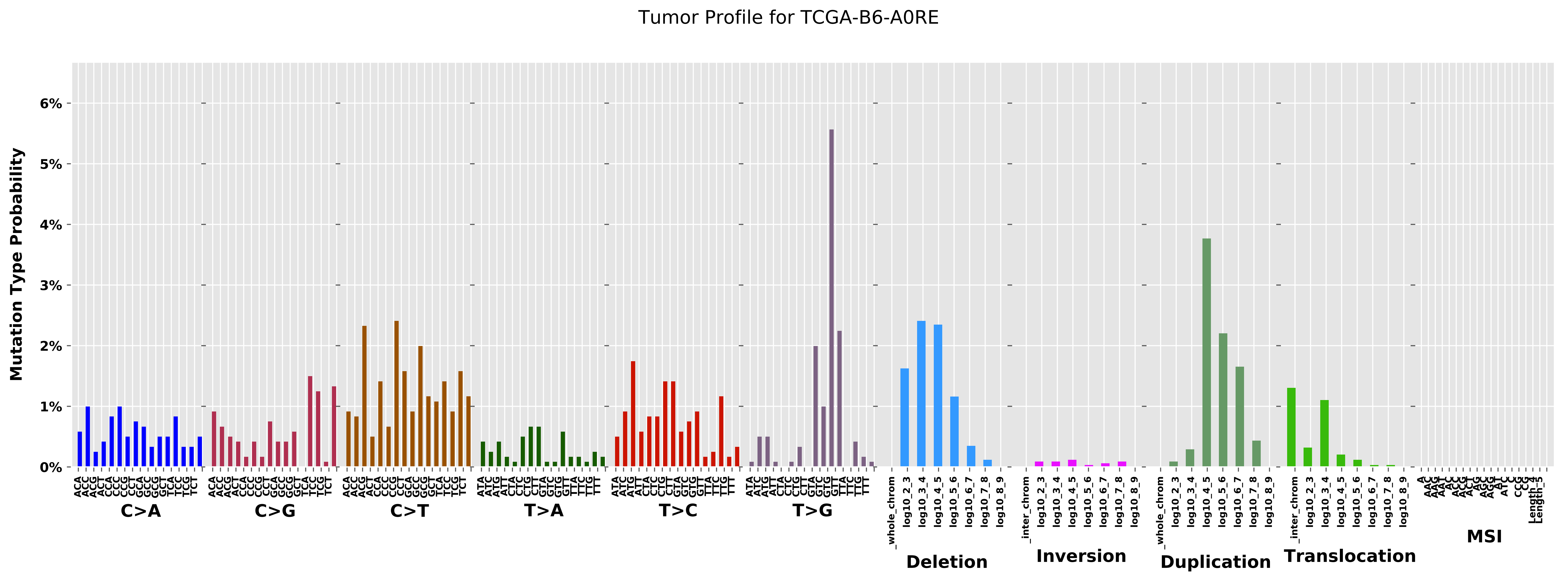
Tumor Profile for TCGA-B6-A0RE
Mutation Type Probability
6%
5%
4%
3%
2%
1%
0%
C>A
C>G
C>T
T>A
T>C
T>G
Deletion
Inversion
Duplication
Translocation
MSI

Cancer processes Weights for TCGA-A8-A08S

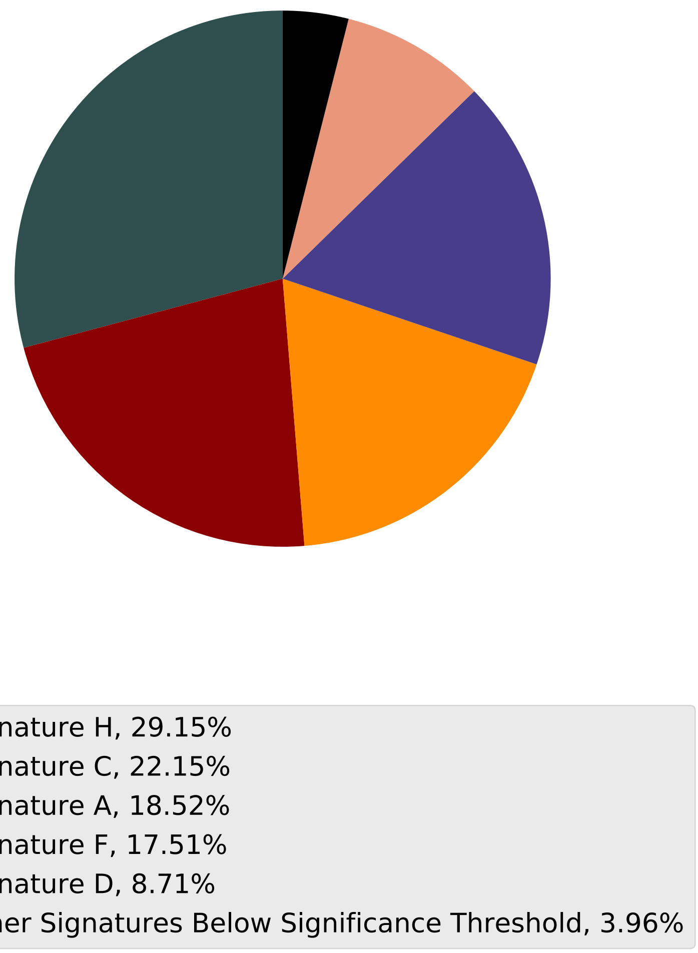

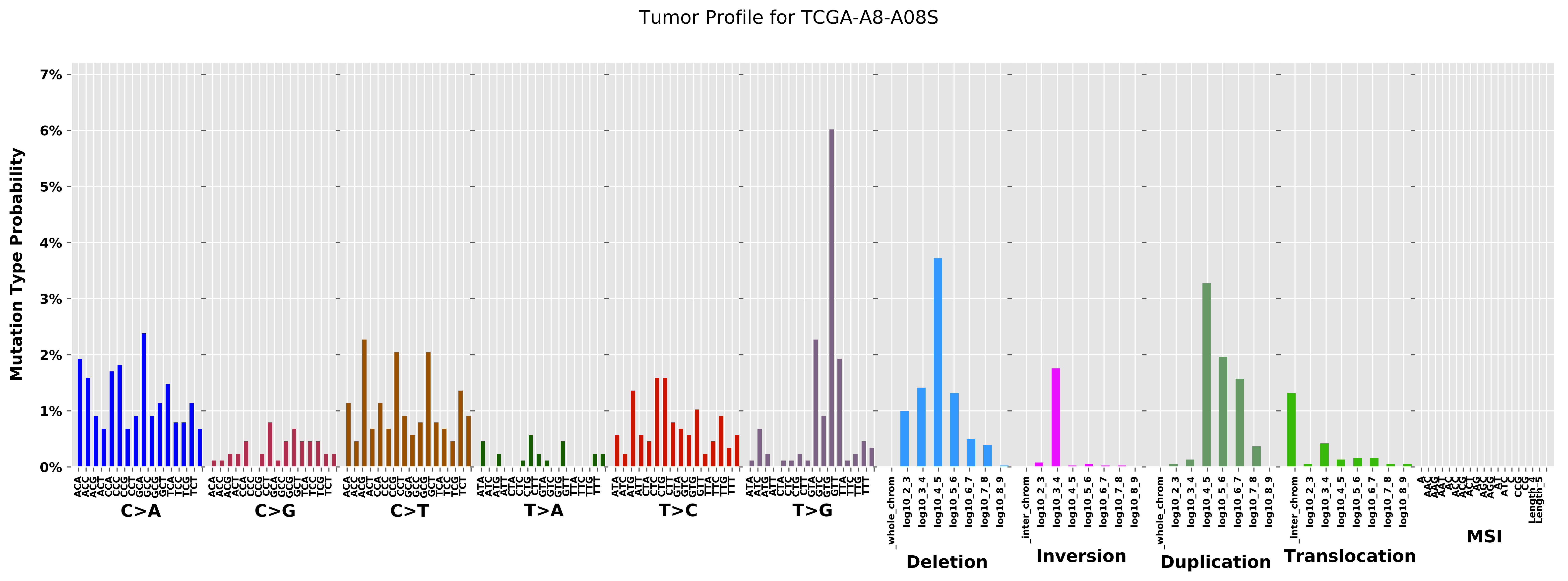
Tumor Profile for TCGA-A8-A08S
Mutation Type Probability
0%
1%
2%
3%
4%
5%
6%
7%
C>A
C>G
C>T
T>A
T>C
T>G
Deletion
Inversion
Duplication
Translocation
MSI
_whole_chrom
_inter_chrom
log10_2_3
log10_3_4
log10_4_5
log10_5_6
log10_6_7
log10_7_8
log10_8_9

Cancer processes Weights for TCGA-AA-A03F

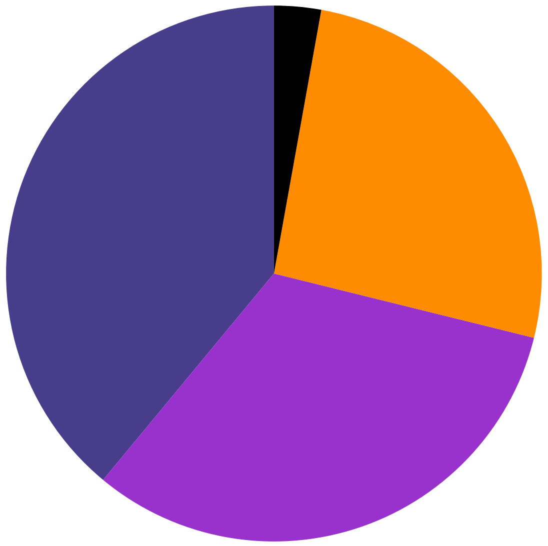

Signature F, 38.99%
Signature B, 32.15%
Signature A, 26.02%
Other Signatures Below Significance Threshold, 2.83%

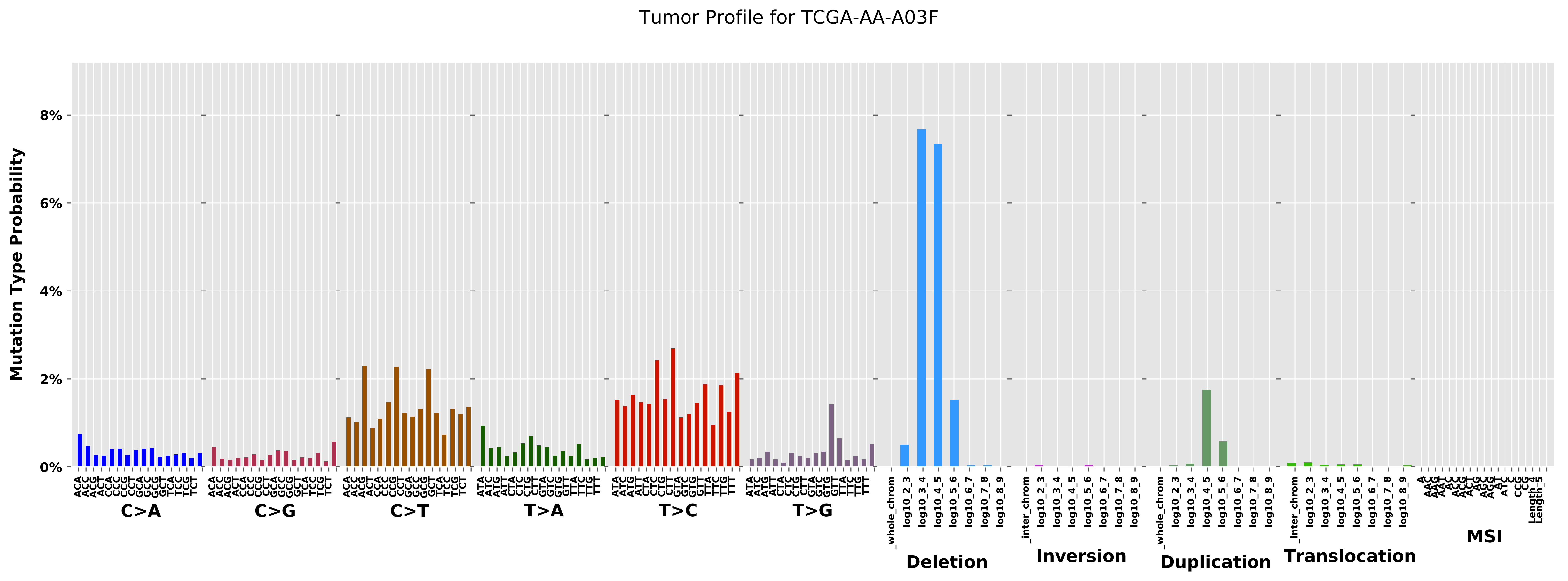
Tumor Profile for TCGA-AA-A03F
Mutation Type Probability
8%
6%
4%
2%
0%
C>A
C>G
C>T
T>A
T>C
T>G
Deletion
Inversion
Duplication
Translocation
MSI

Cancer processes Weights for TCGA-EW-A1PH

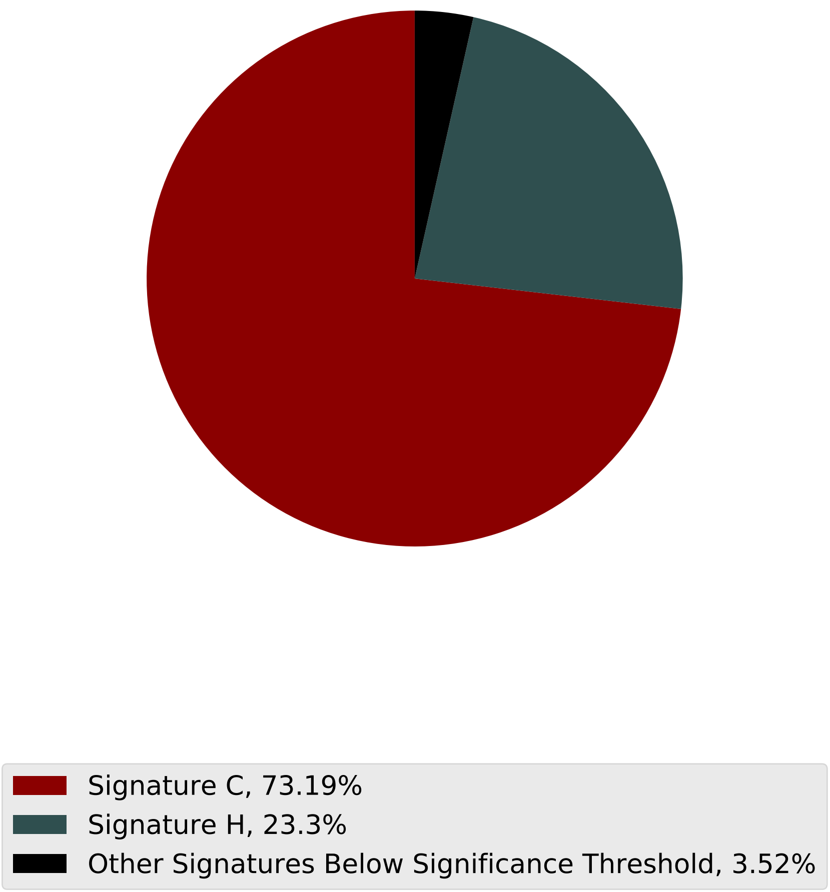

Tumor Profile for TCGA-EW-A1PH

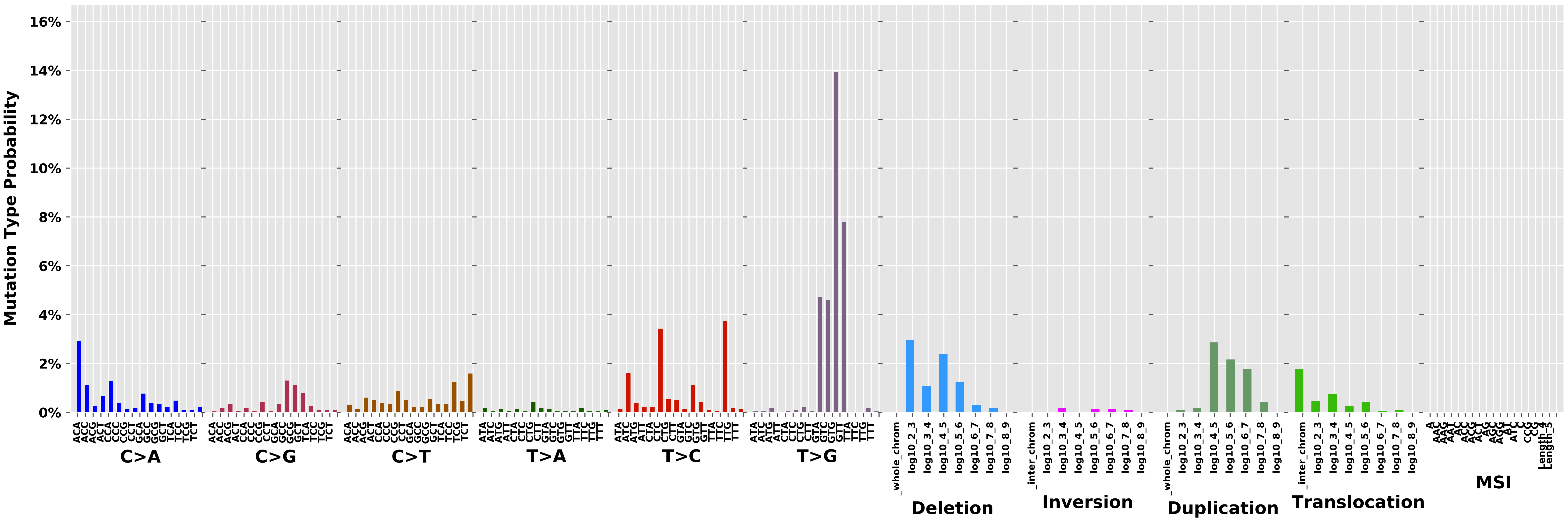

Cancer processes Weights for TCGA-E2-A1LG

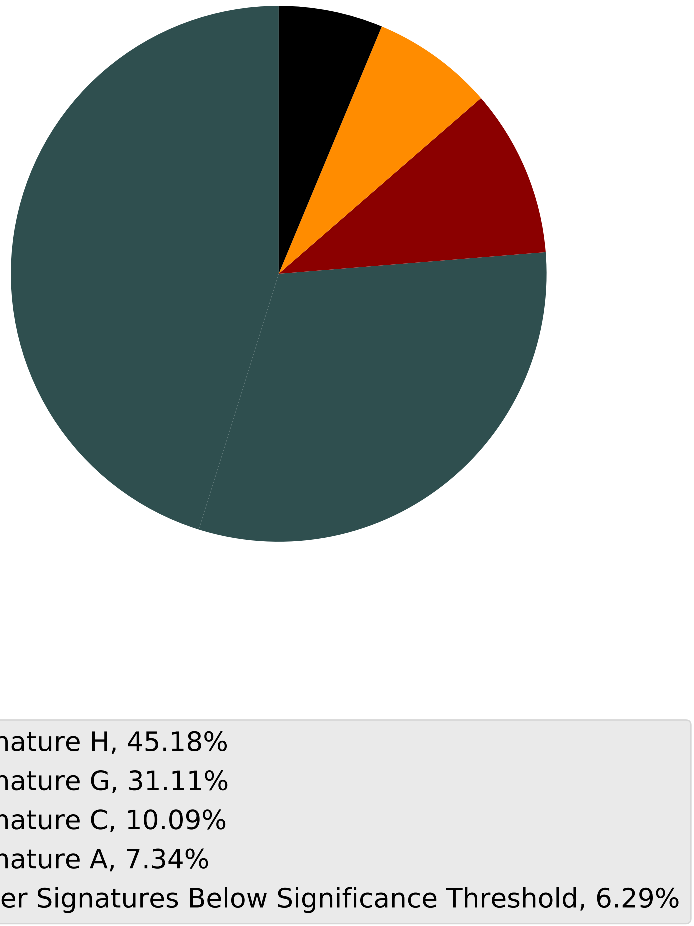

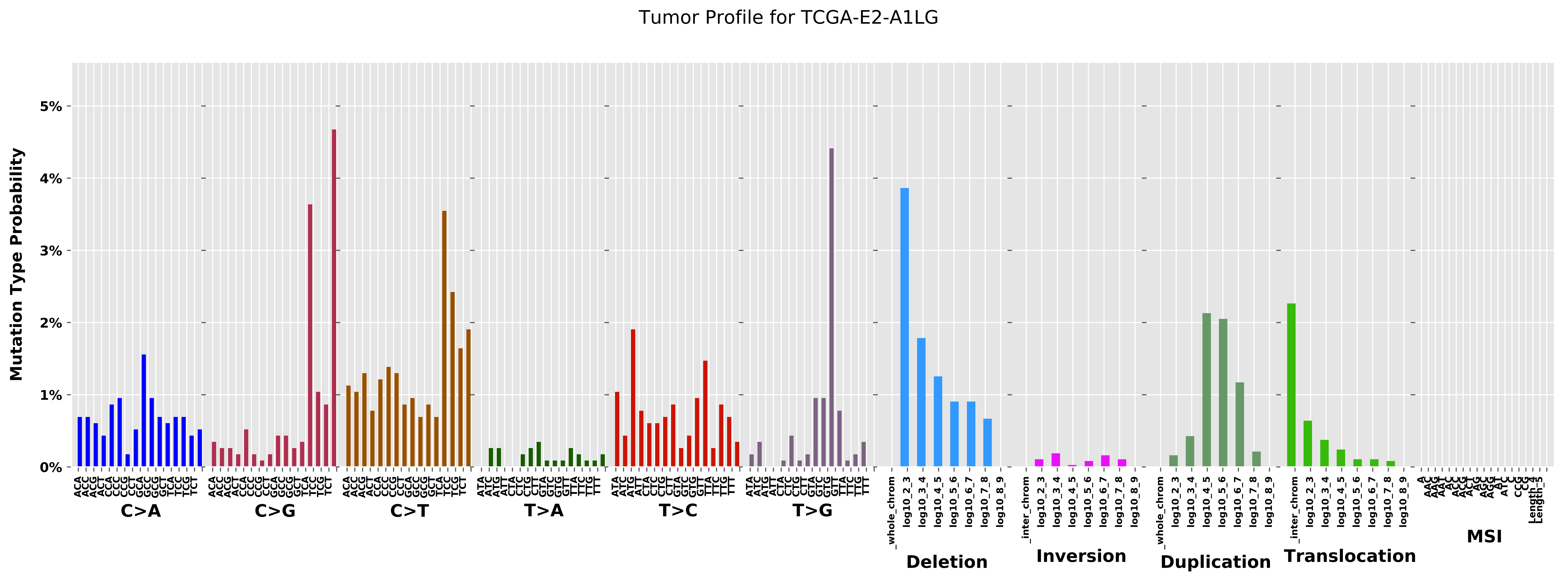

Tumor Profile for TCGA-E2-A1LG
Mutation Type Probability
0%
1%
2%
3%
4%
5%
C>A
C>G
C>T
T>A
T>C
T>G
Deletion
Inversion
Duplication
Translocation
MSI

Cancer processes Weights for TCGA-A8-A08B

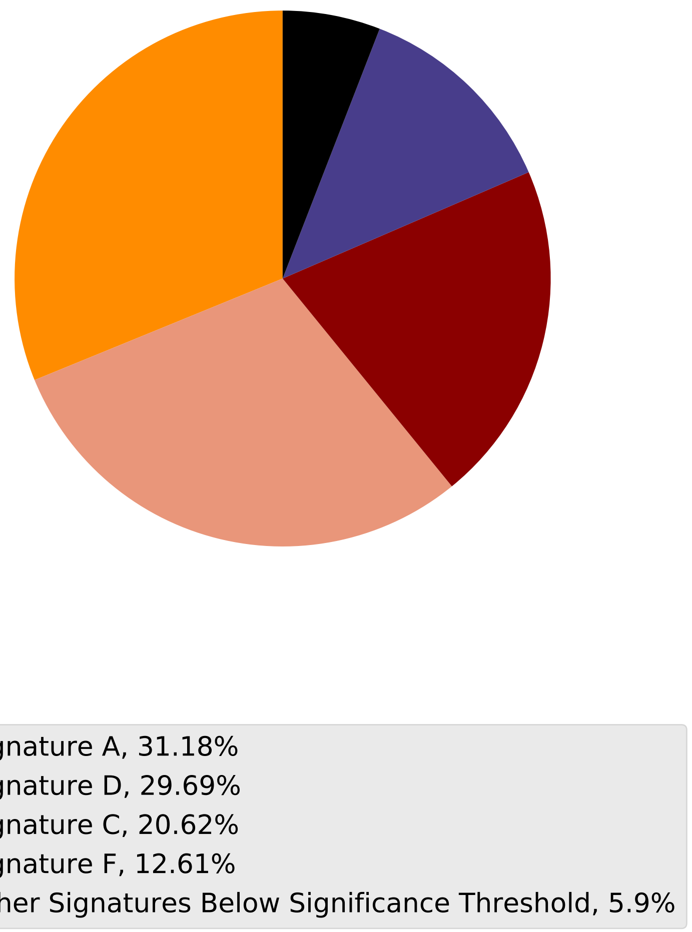

Tumor Profile for TCGA-A8-A08B

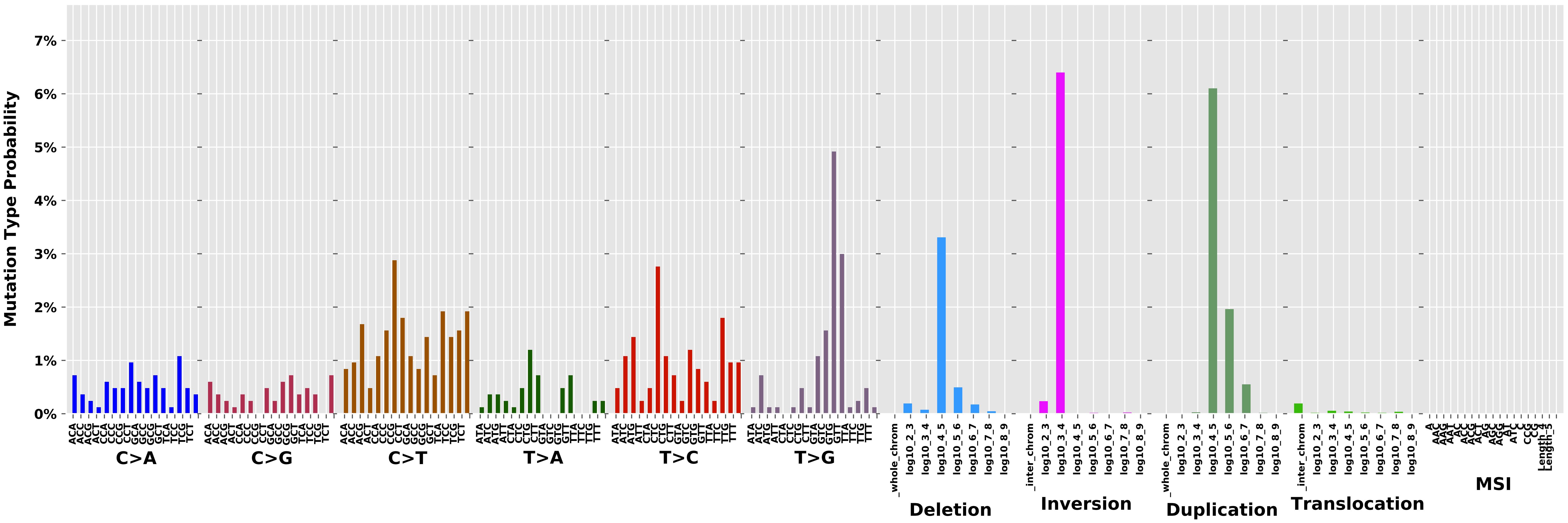

Cancer processes Weights for TCGA-B6-A0RT

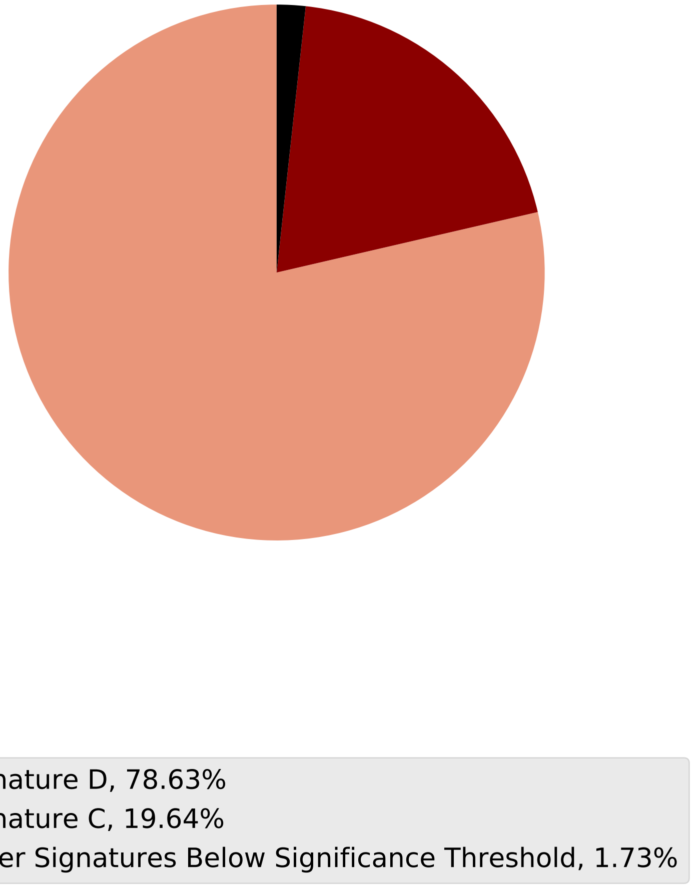

Tumor Profile for TCGA-B6-A0RT

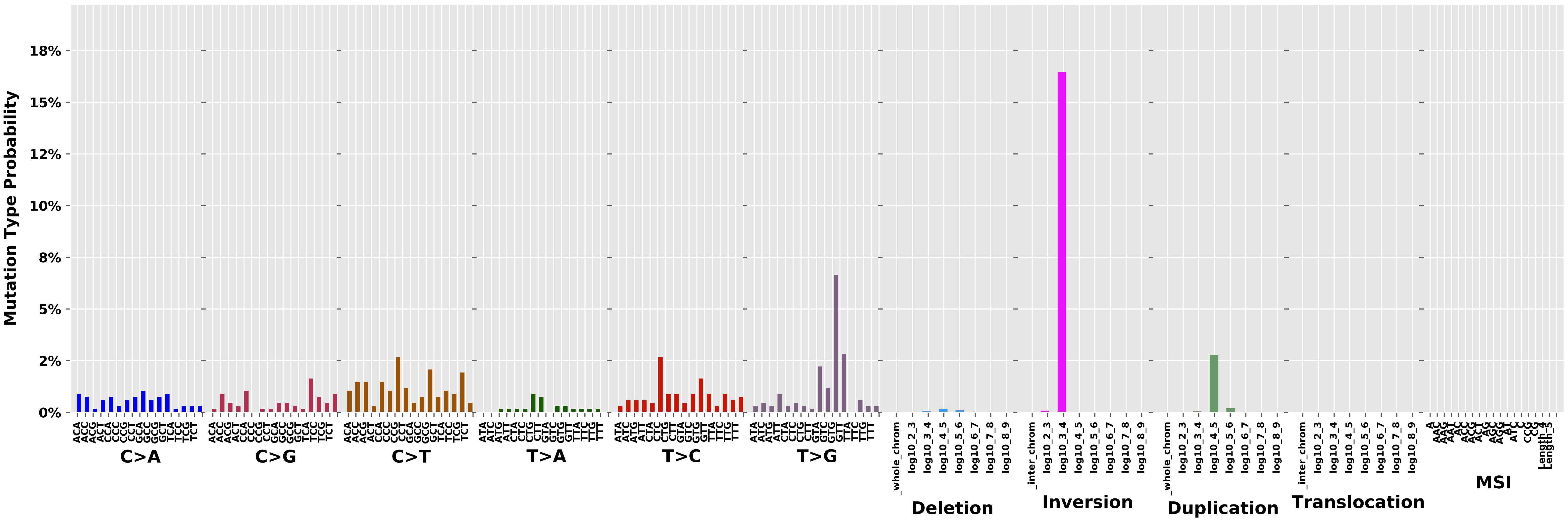

Cancer processes Weights for TCGA-GM-A3XL

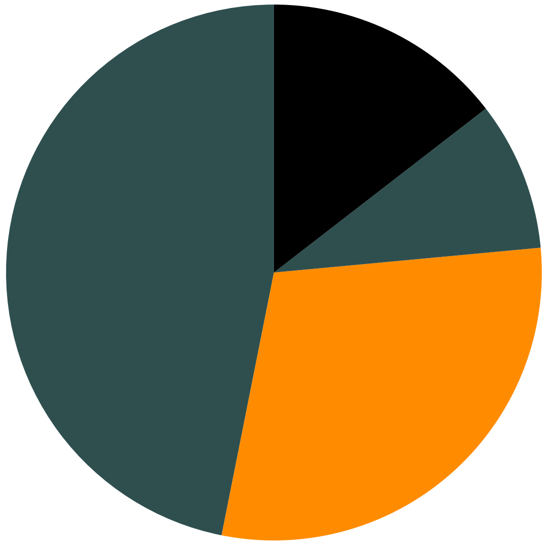

Signature H, 46.87%
Signature A, 29.61%
Signature G, 9.0%
Other Signatures Below Significance Threshold, 14.52%

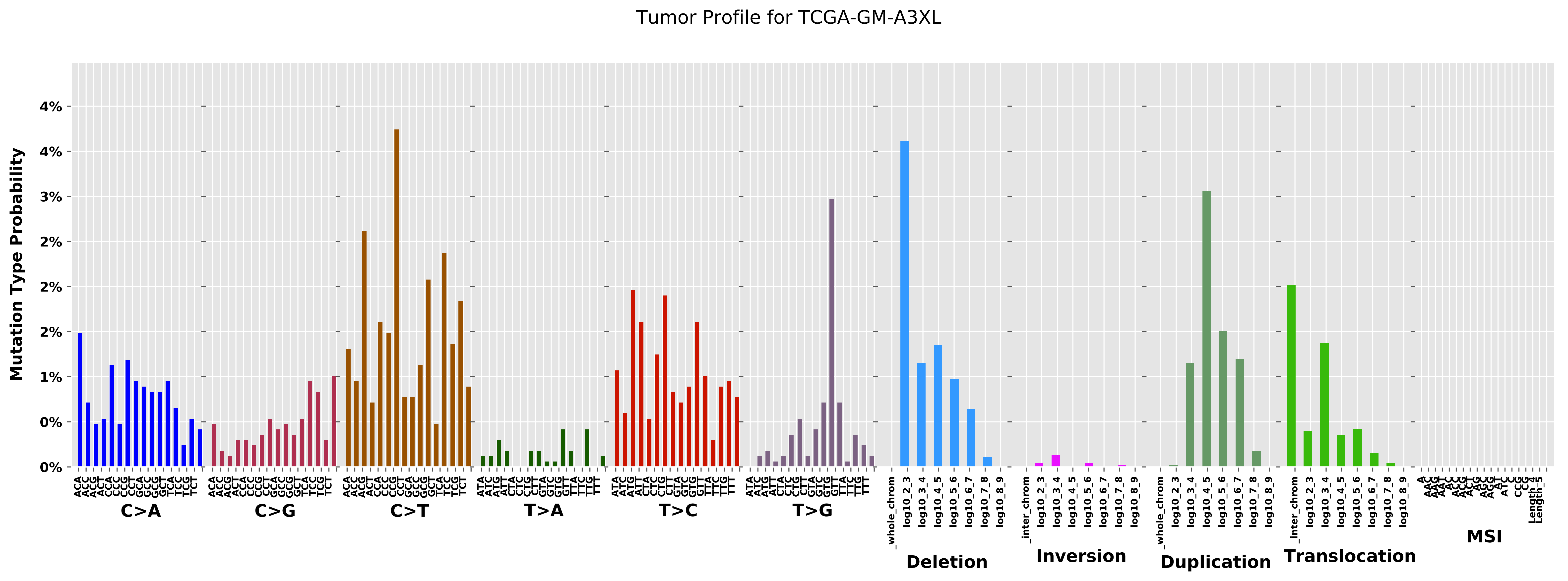
Tumor Profile for TCGA-GM-A3XL
Mutation Type Probability
C>A
C>G
C>T
T>A
T>C
T>G
Deletion
Inversion
Duplication
Translocation
MSI

Cancer processes Weights for TCGA-AO-A0J2

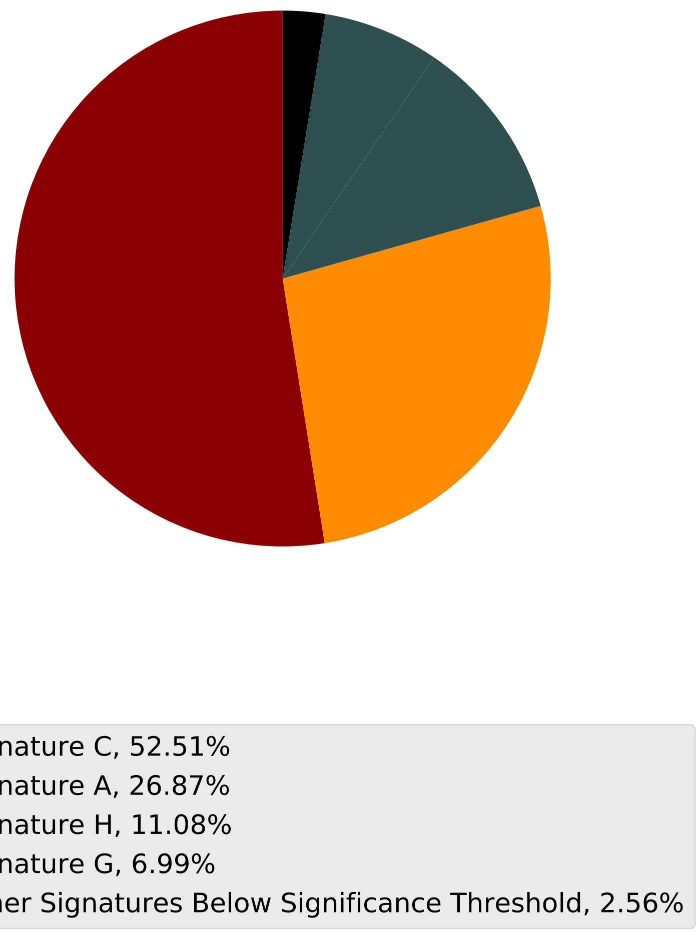

Tumor Profile for TCGA-AO-A0J2

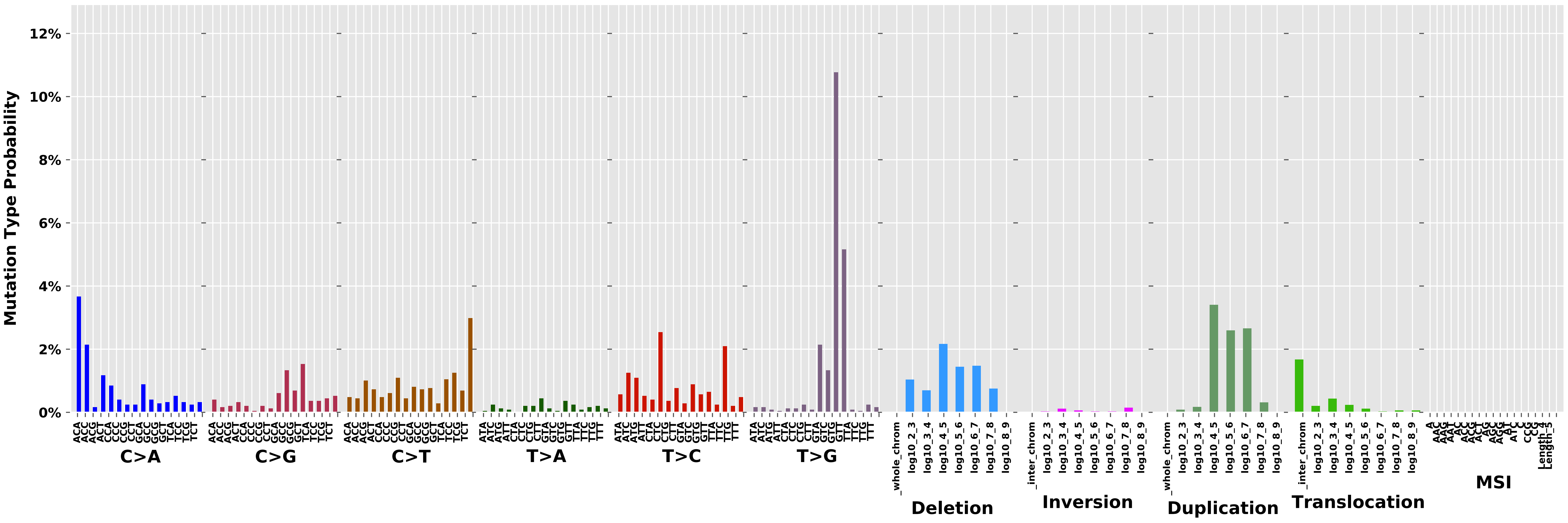

Cancer processes Weights for TCGA-E2-A156

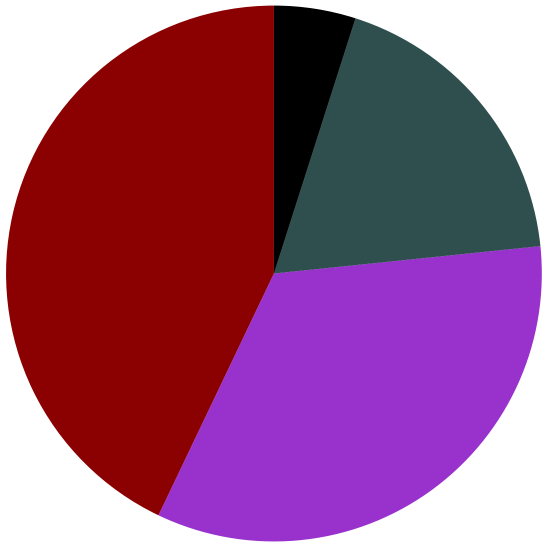

Signature C, 42.91%
Signature B, 33.72%
Signature H, 18.45%
Other Signatures Below Significance Threshold, 4.92%

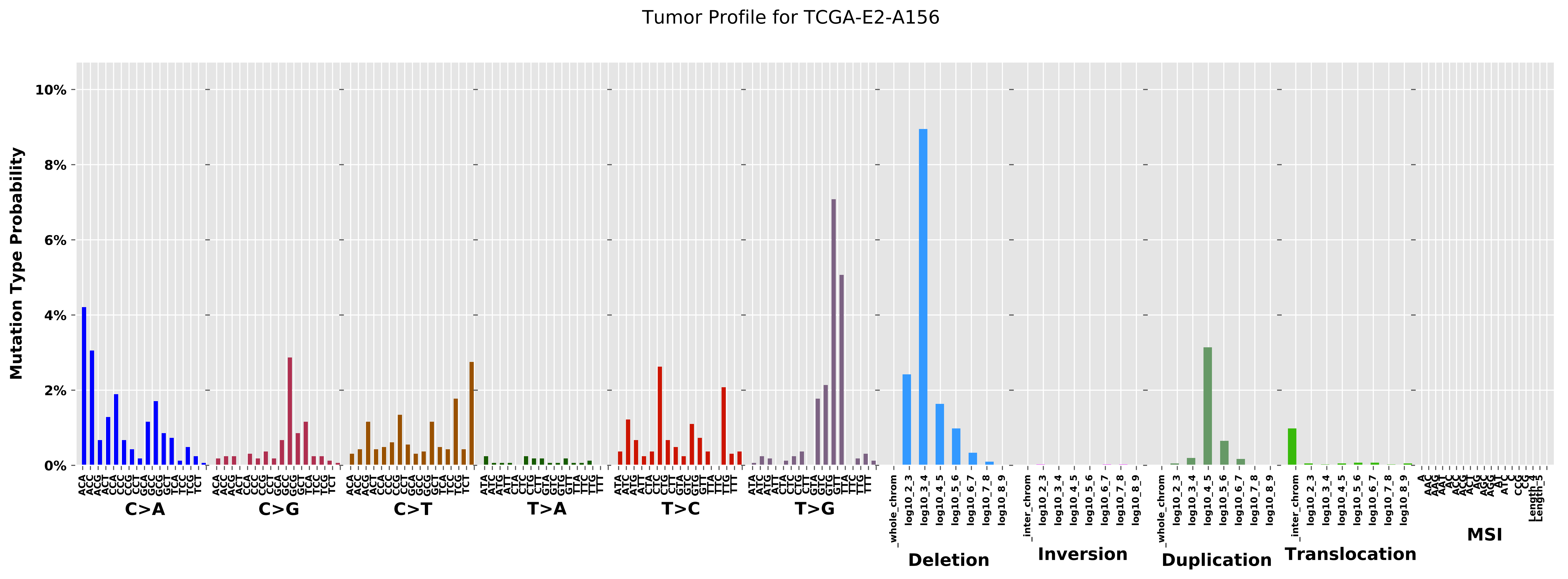

Tumor Profile for TCGA-E2-A156
Mutation Type Probability
10%
8%
6%
4%
2%
0%
C>A
C>G
C>T
T>A
T>C
T>G
Deletion
Inversion
Duplication
Translocation
MSI

Cancer processes Weights for TCGA-BH-A18R

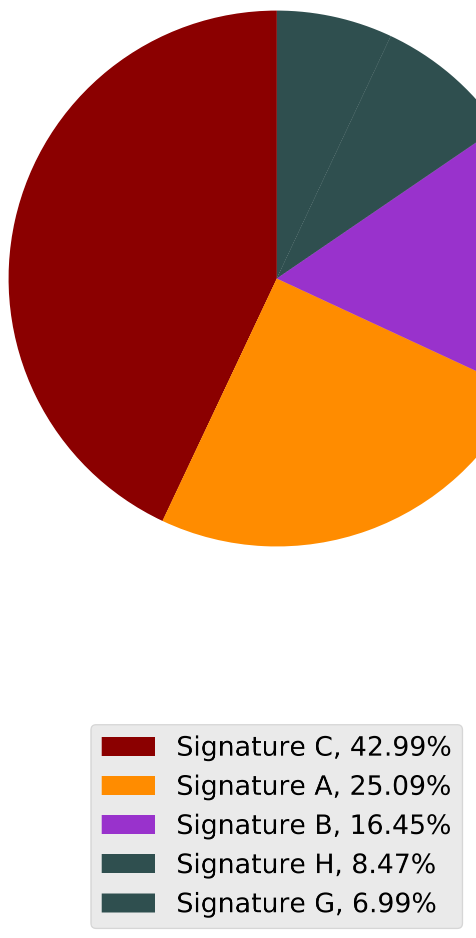

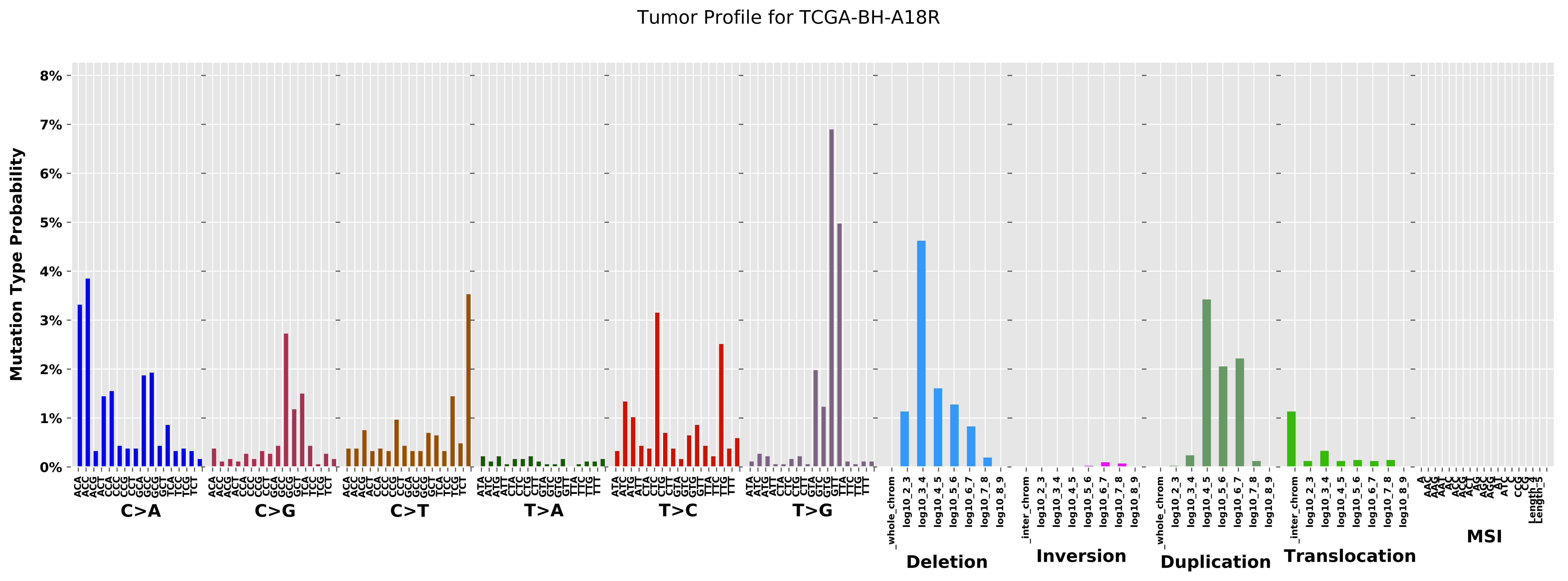

Tumor Profile for TCGA-BH-A18R
Mutation Type Probability
0%
1%
2%
3%
4%
5%
6%
7%
8%
C>A
C>G
C>T
T>A
T>C
T>G
Deletion
Inversion
Duplication
Translocation
MSI

Cancer processes Weights for TCGA-AO-A0JM

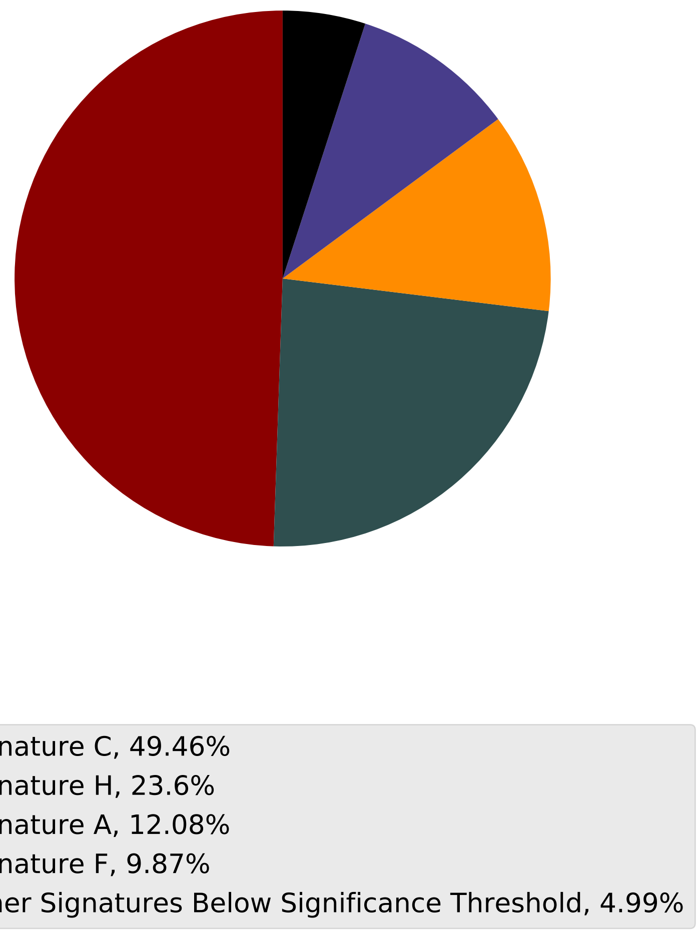

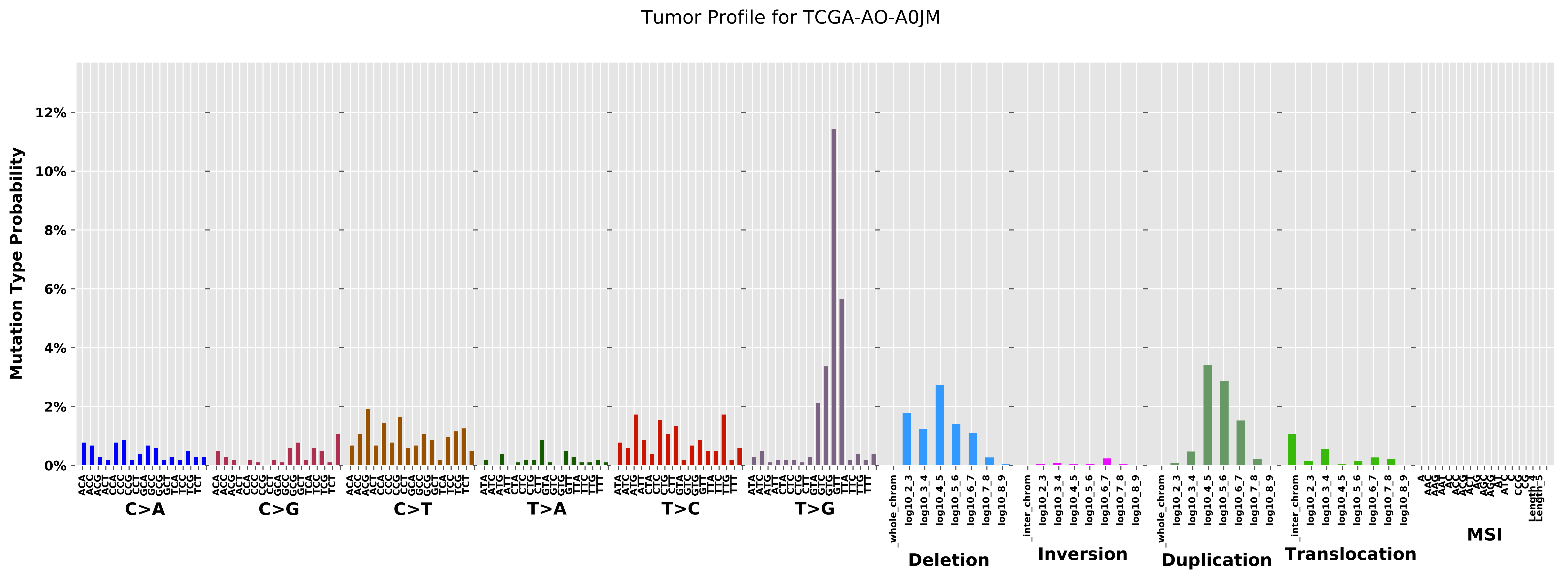
Tumor Profile for TCGA-AO-A0JM
Mutation Type Probability
0%
2%
4%
6%
8%
10%
12%
C>A
C>G
C>T
T>A
T>C
T>G
_whole_chrom
log10_2_3
log10_3_4
log10_4_5
log10_5_6
log10_6_7
log10_7_8
log10_8_9
Deletion
_inter_chrom
Inversion
Duplication
Translocation
MSI

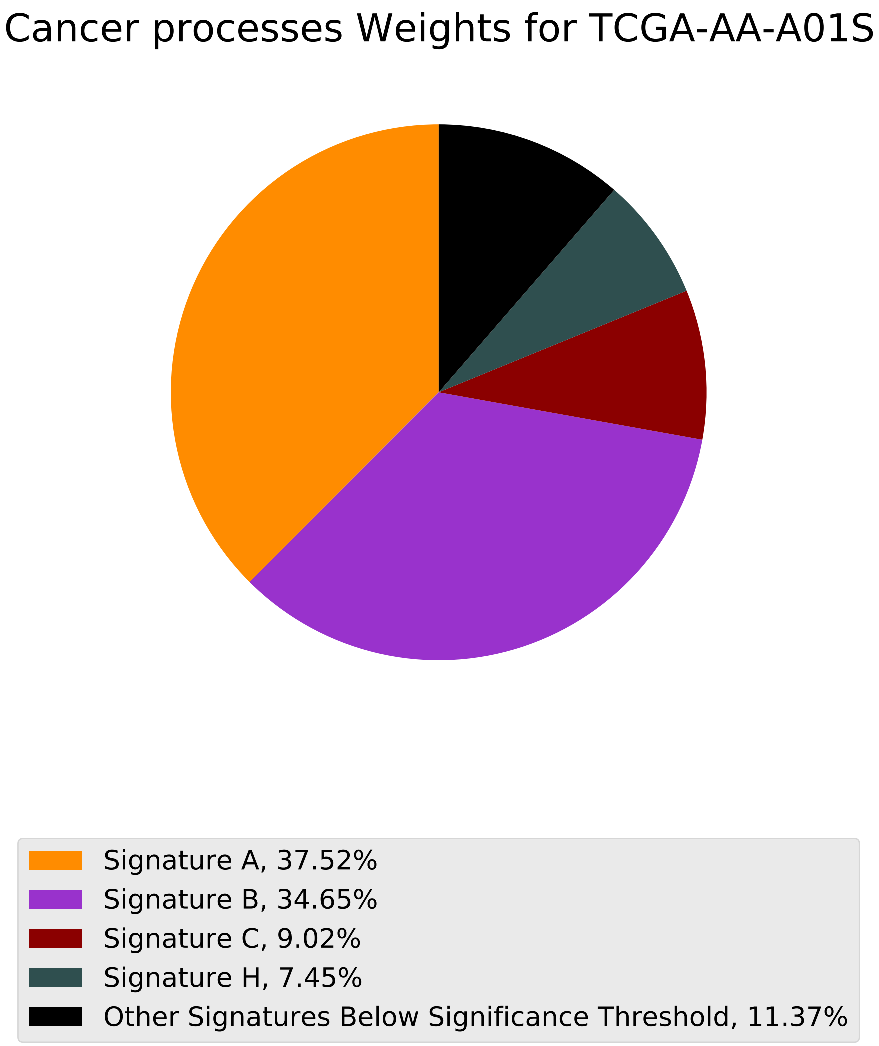
Cancer processes Weights for TCGA-AA-A01S
Signature A, 37.52%
Signature B, 34.65%
Signature C, 9.02%
Signature H, 7.45%
Other Signatures Below Significance Threshold, 11.37%

Tumor Profile for TCGA-AA-A01S

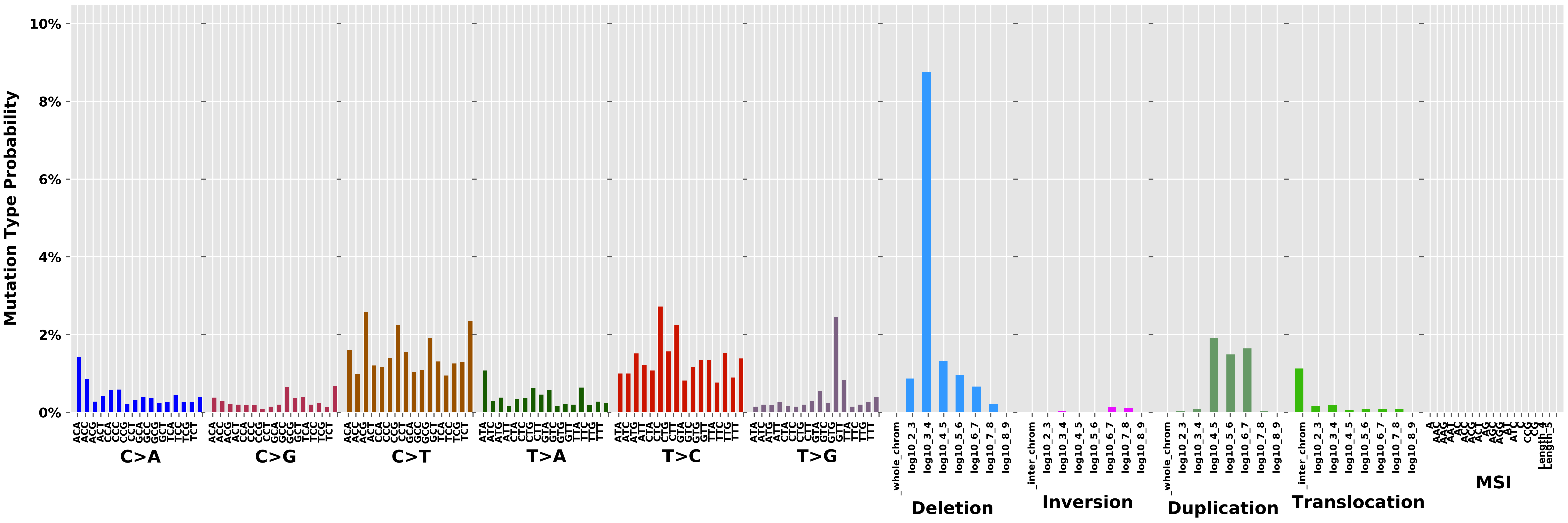

#### Cancer processes Weights for TCGA-AN-A0AT

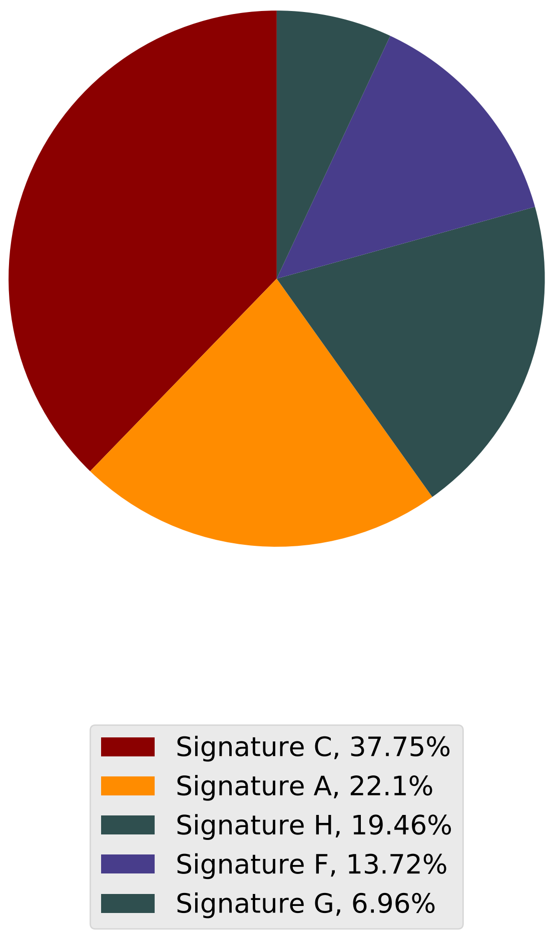

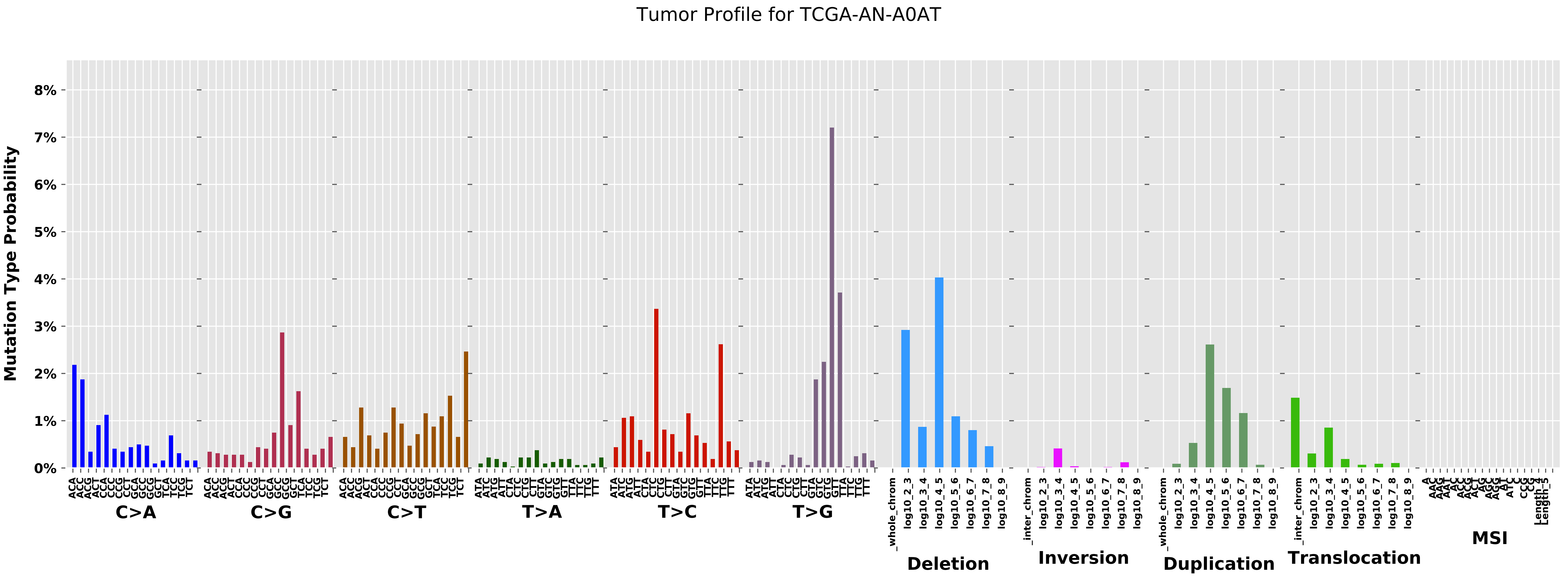
Tumor Profile for TCGA-AN-A0AT
Mutation Type Probability
0%
1%
2%
3%
4%
5%
6%
7%
8%
C>A
C>G
C>T
T>A
T>C
T>G
Deletion
Inversion
Duplication
Translocation
MSI

### Cancer processes Weights for TCGA-B6-A0WX

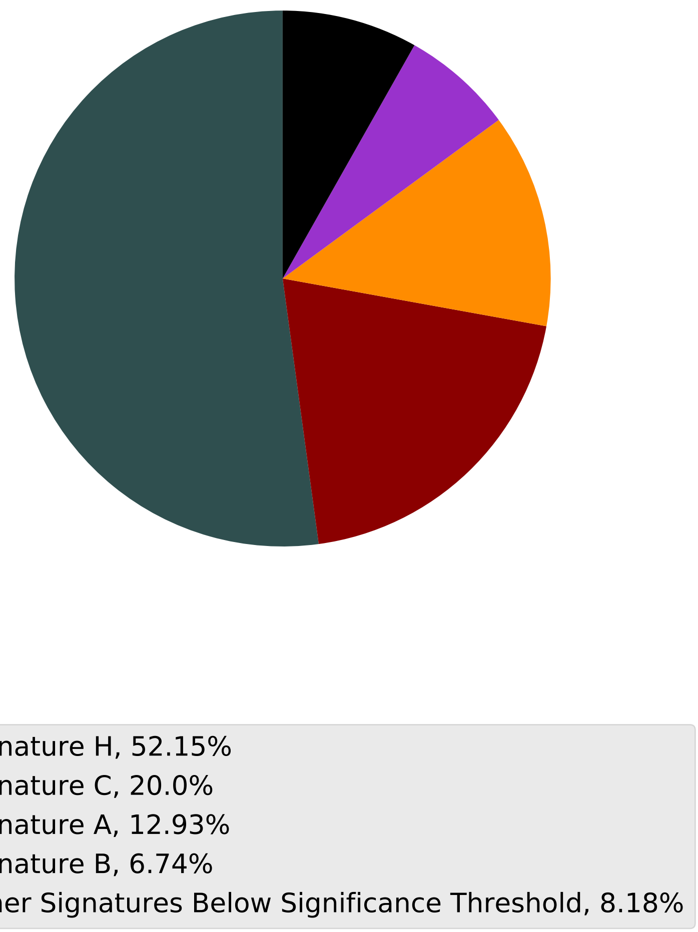

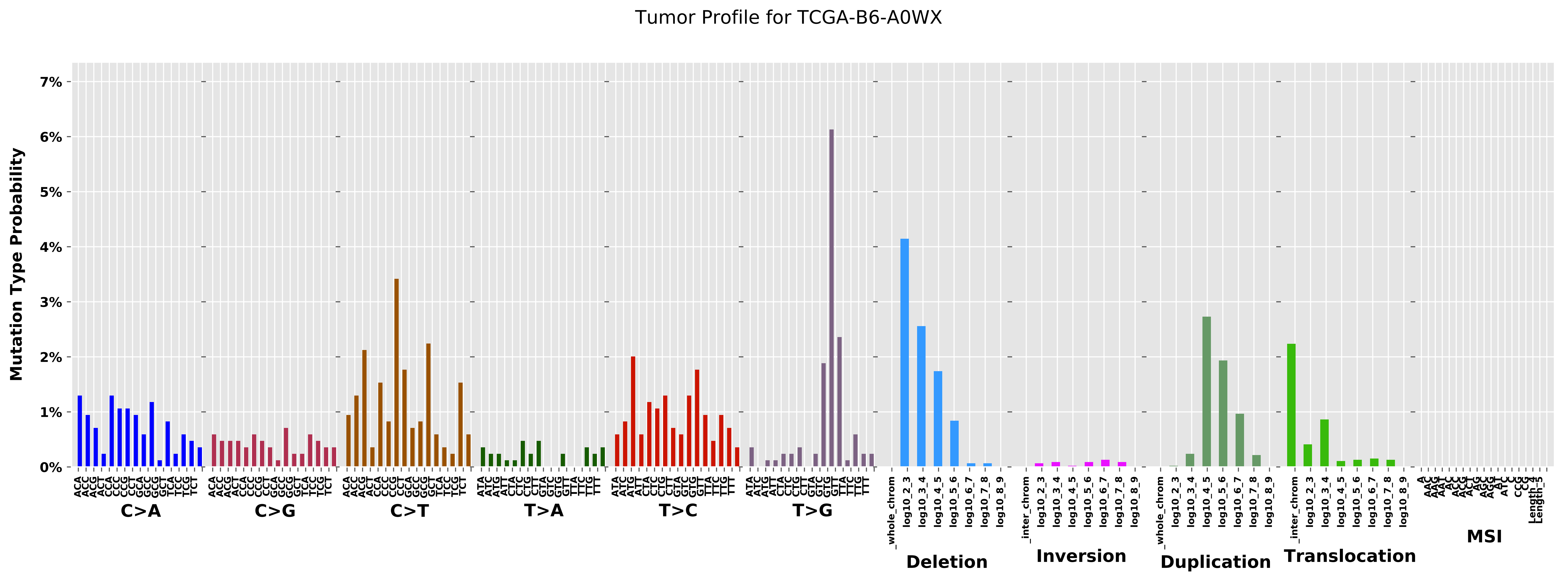

Tumor Profile for TCGA-B6-A0WX
Mutation Type Probability
7%
6%
5%
4%
3%
2%
1%
0%
C>A
C>G
C>T
T>A
T>C
T>G
Deletion
Inversion
Duplication
Translocation
MSI

Cancer processes Weights for TCGA-BH-A0WA

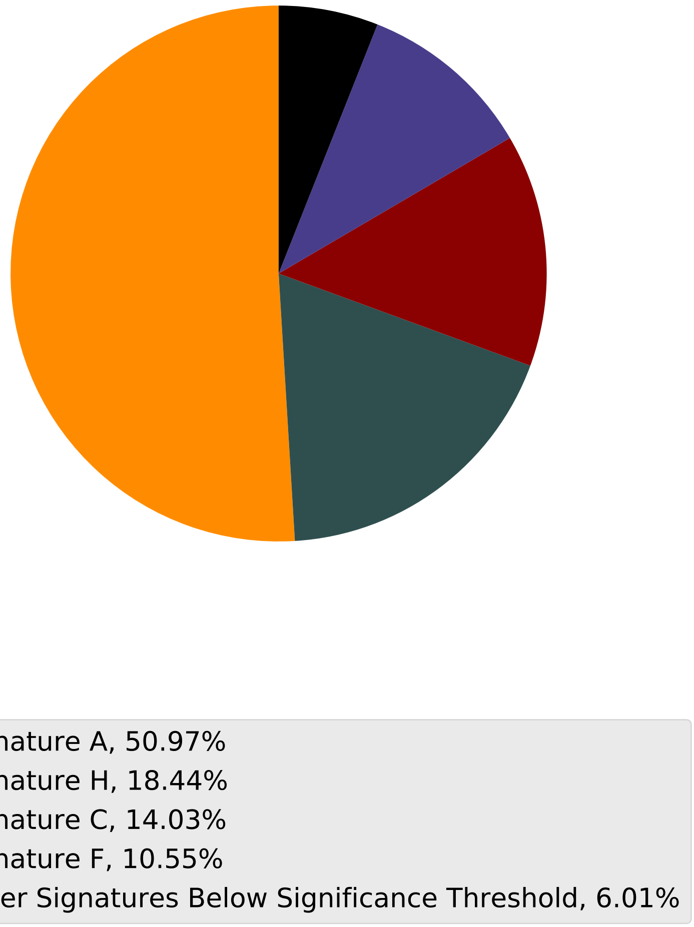

Tumor Profile for TCGA-BH-A0WA

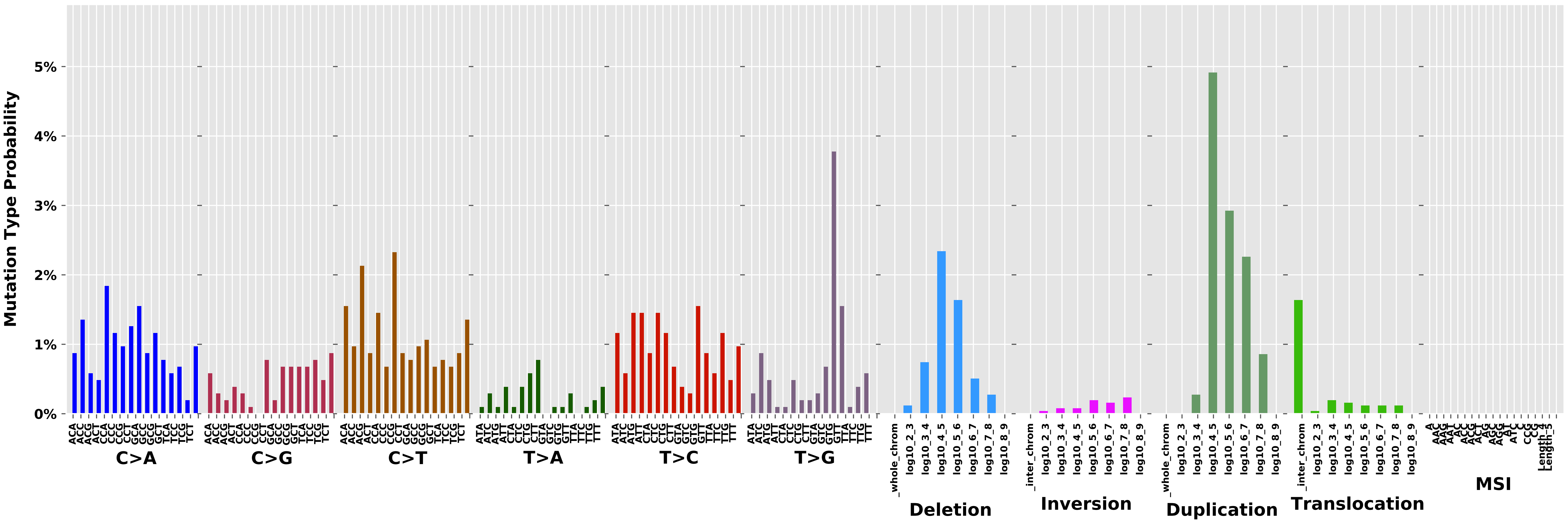

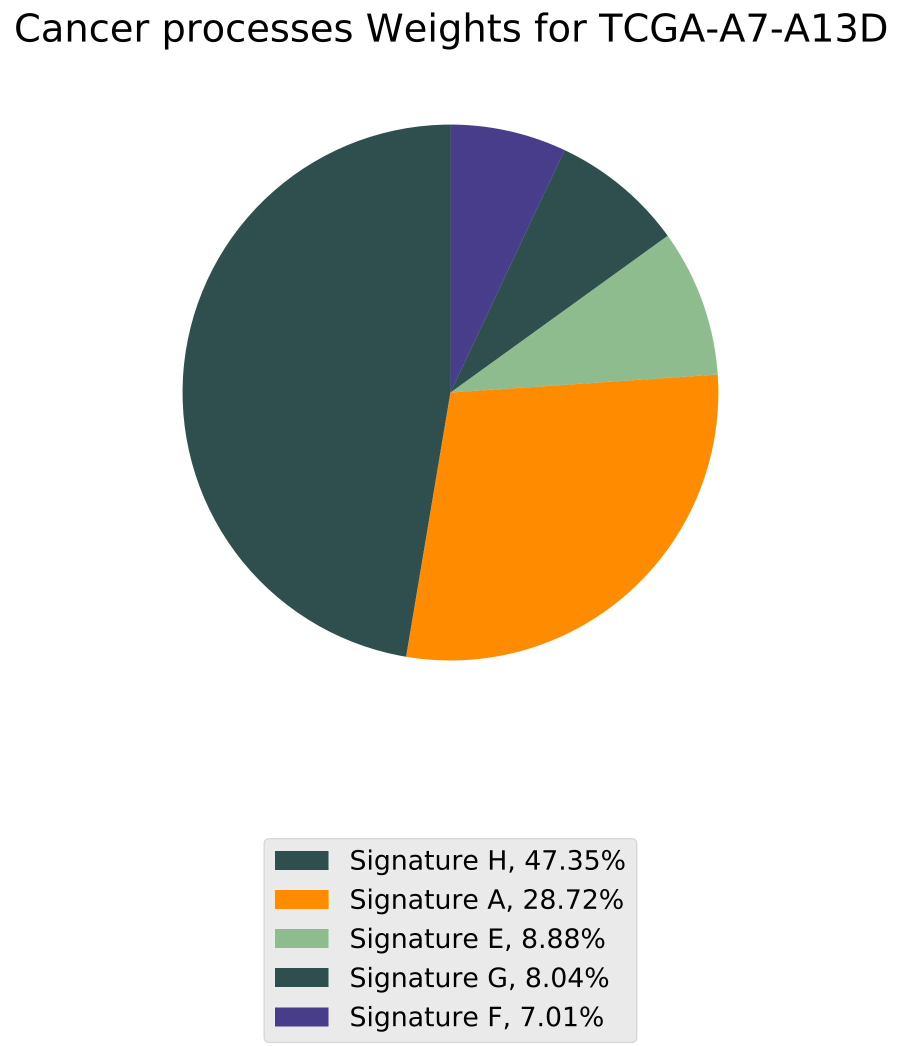
Cancer processes Weights for TCGA-A7-A13D
Signature H, 47.35%
Signature A, 28.72%
Signature E, 8.88%
Signature G, 8.04%
Signature F, 7.01%

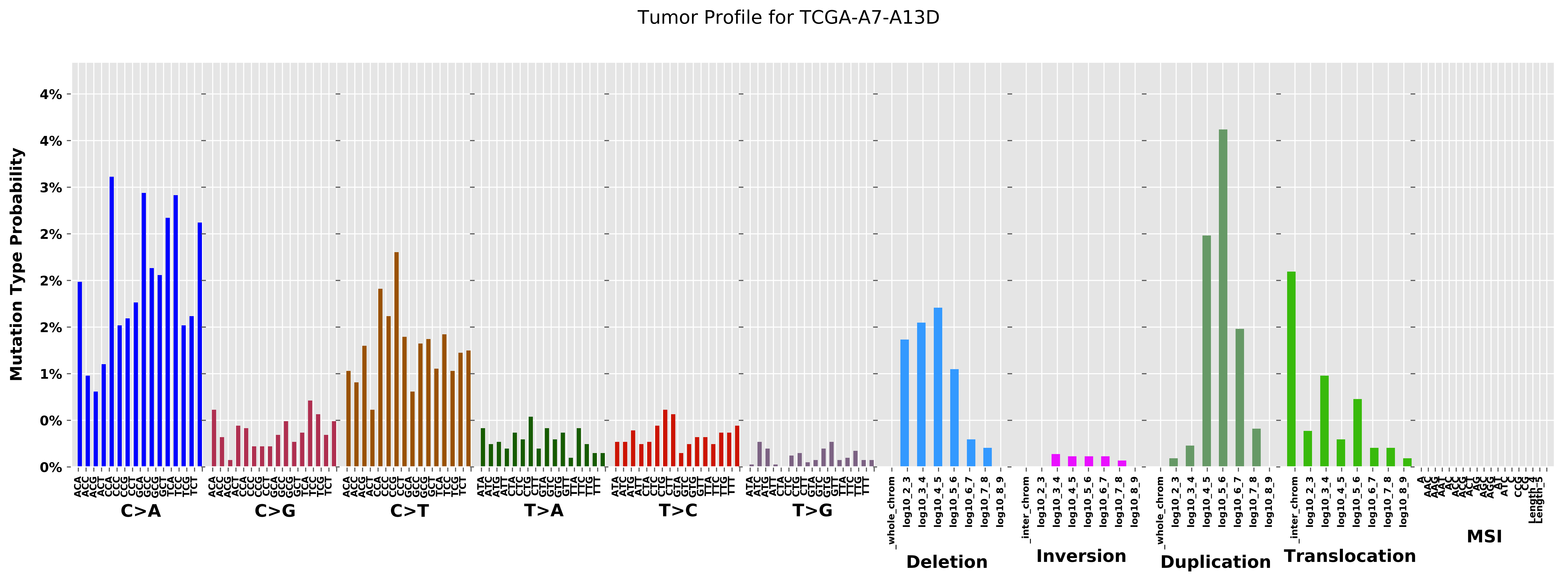

Tumor Profile for TCGA-A7-A13D
Mutation Type Probability
C>A
C>G
C>T
T>A
T>C
T>G
Deletion
Inversion
Duplication
Translocation
MSI

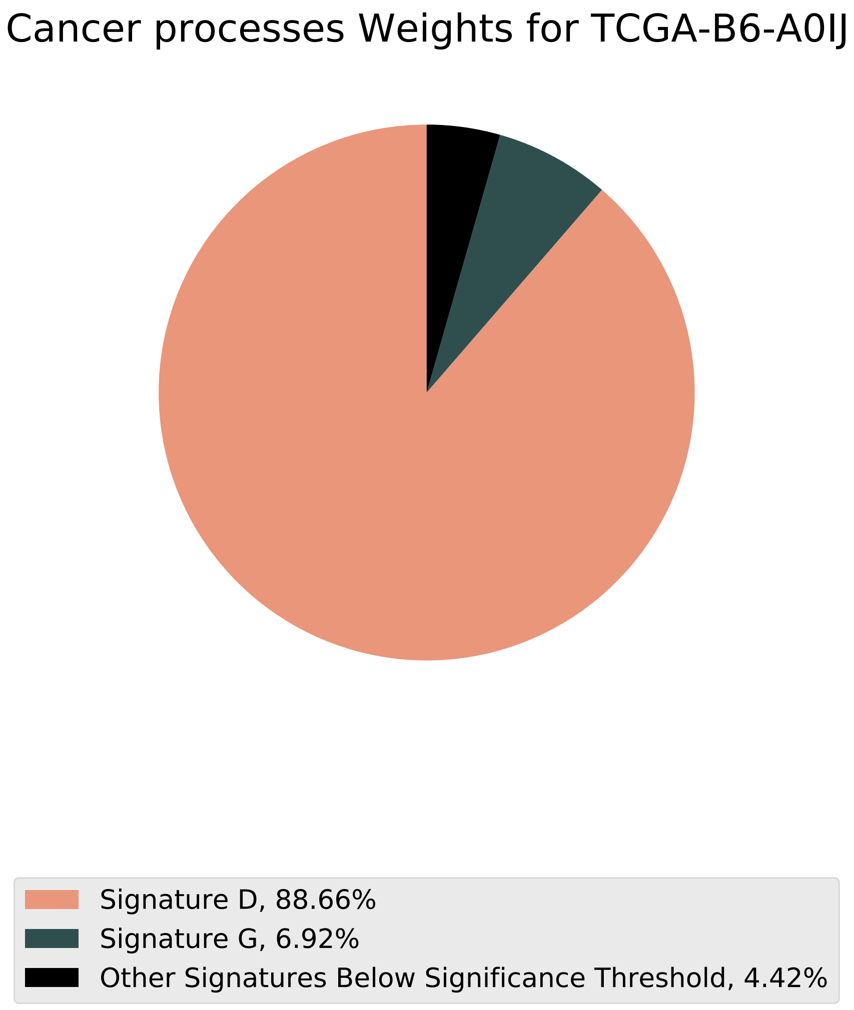
Cancer processes Weights for TCGA-B6-A0IJ
Signature D, 88.66%
Signature G, 6.92%
Other Signatures Below Significance Threshold, 4.42%

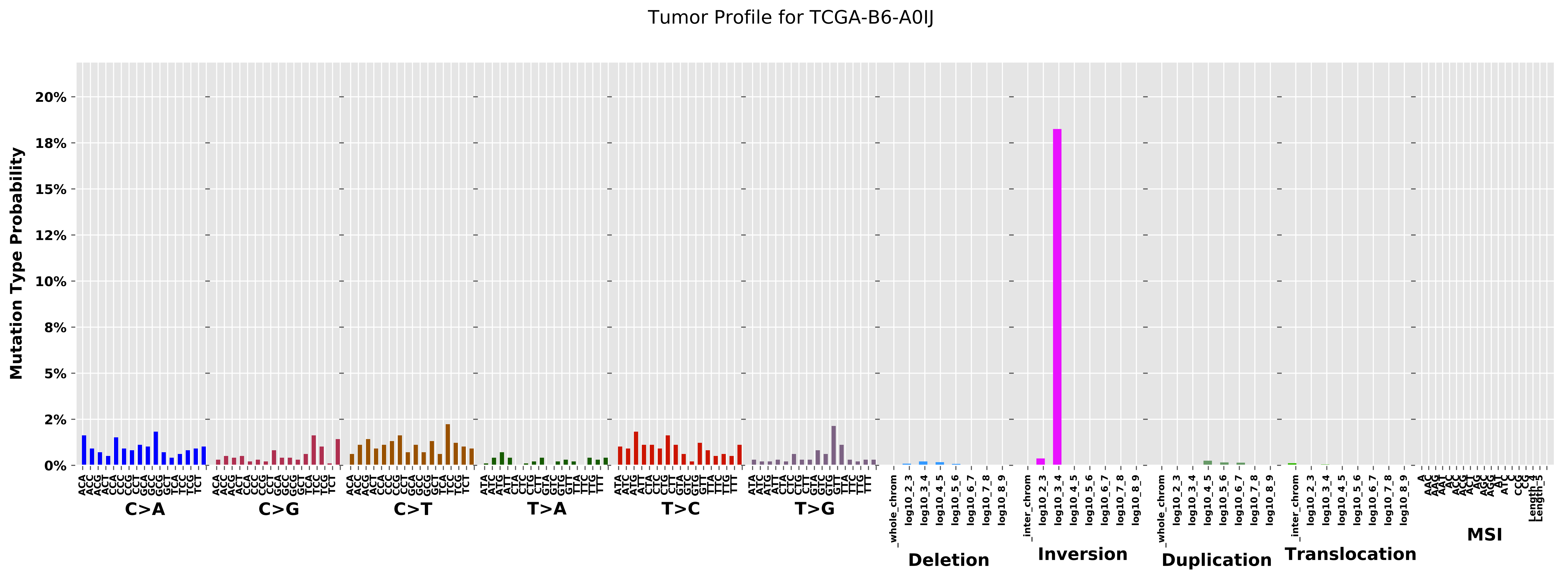

Tumor Profile for TCGA-B6-A0IJ
Mutation Type Probability
0%
2%
5%
8%
10%
12%
15%
18%
20%
C>A
C>G
C>T
T>A
T>C
T>G
Deletion
Inversion
Duplication
Translocation
MSI

Cancer processes Weights for TCGA-A2-A04Q

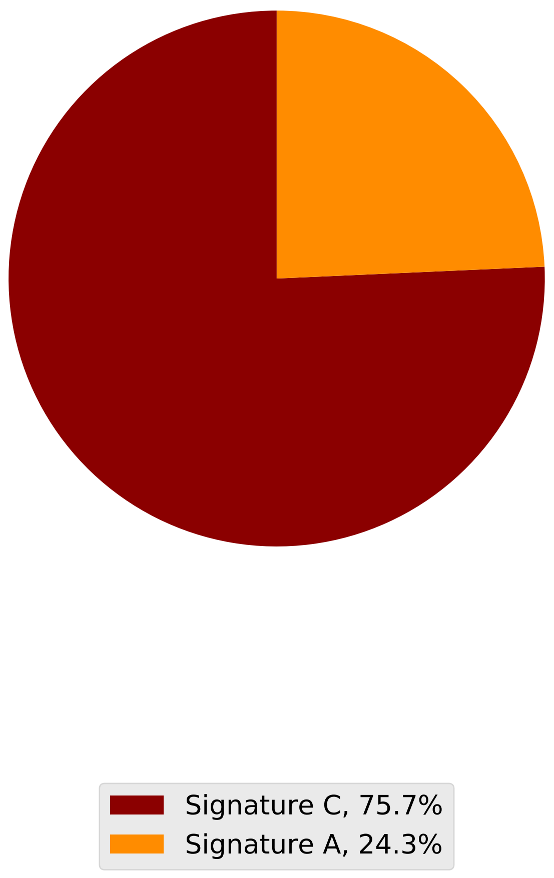

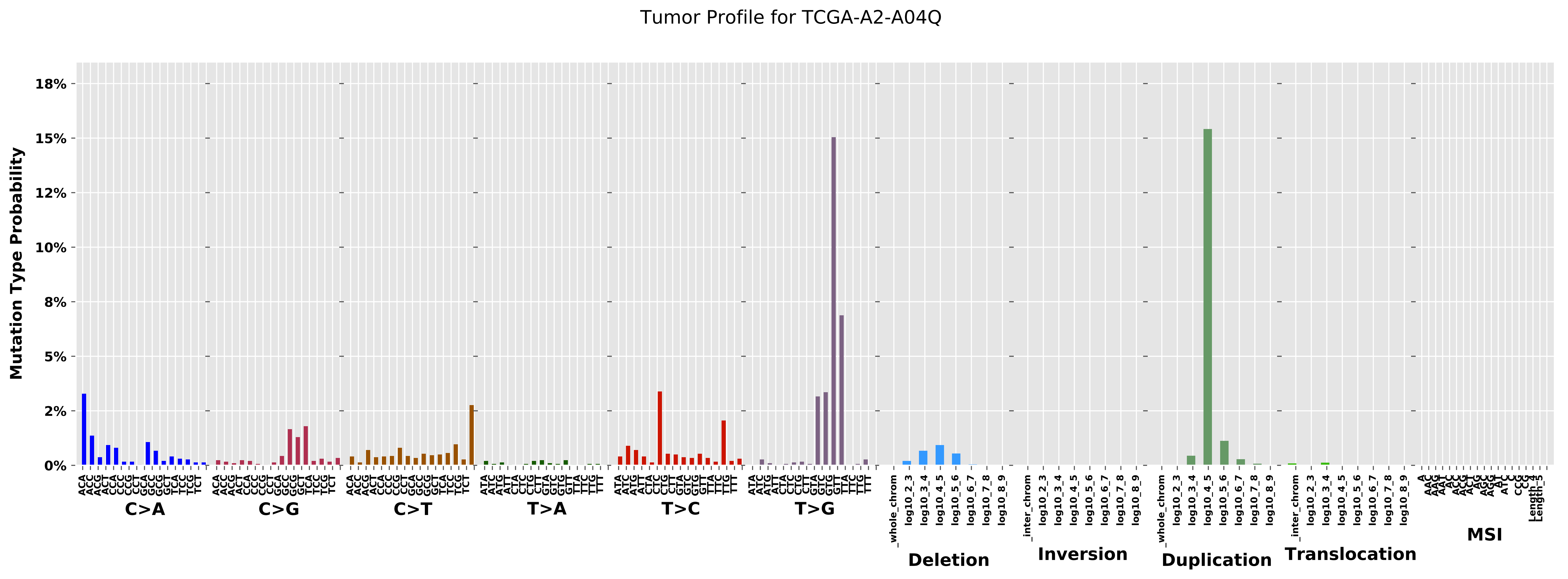

Tumor Profile for TCGA-A2-A04Q
Mutation Type Probability
18%
15%
12%
10%
8%
5%
2%
0%
C>A
C>G
C>T
T>A
T>C
T>G
Deletion
Inversion
Duplication
Translocation
MSI

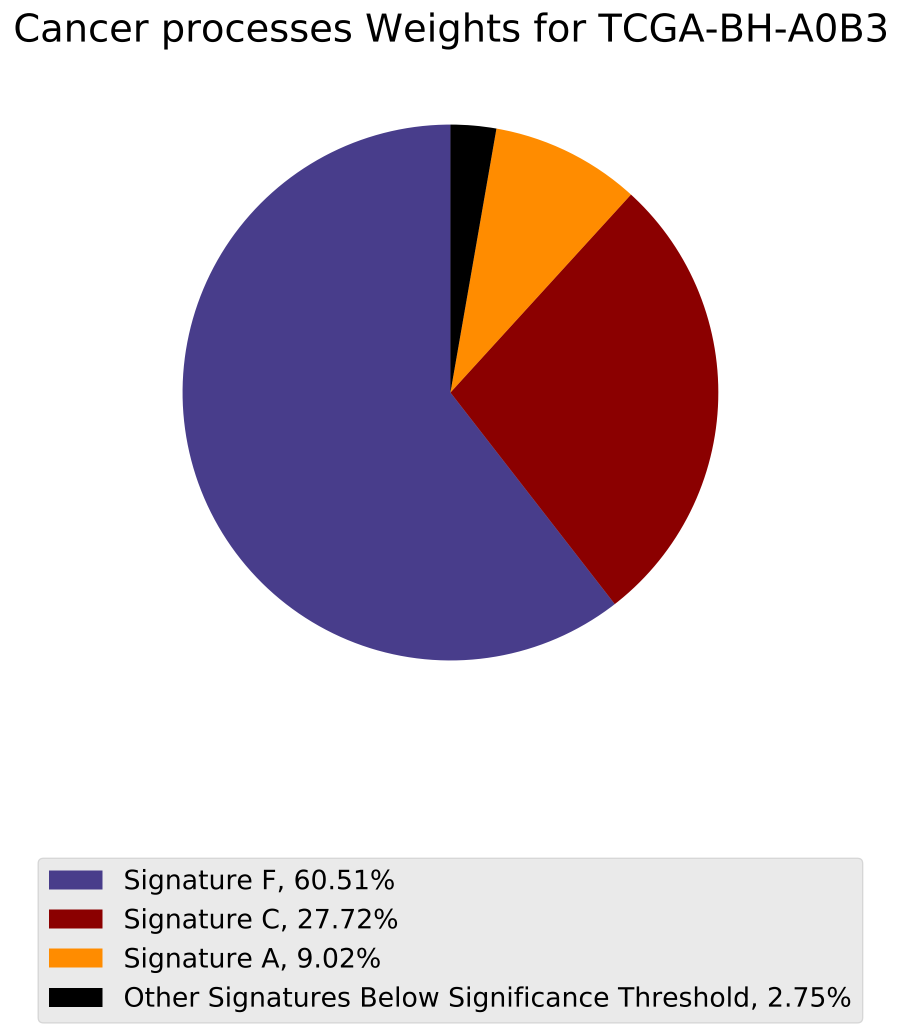
Cancer processes Weights for TCGA-BH-A0B3
Signature F, 60.51%
Signature C, 27.72%
Signature A, 9.02%
Other Signatures Below Significance Threshold, 2.75%

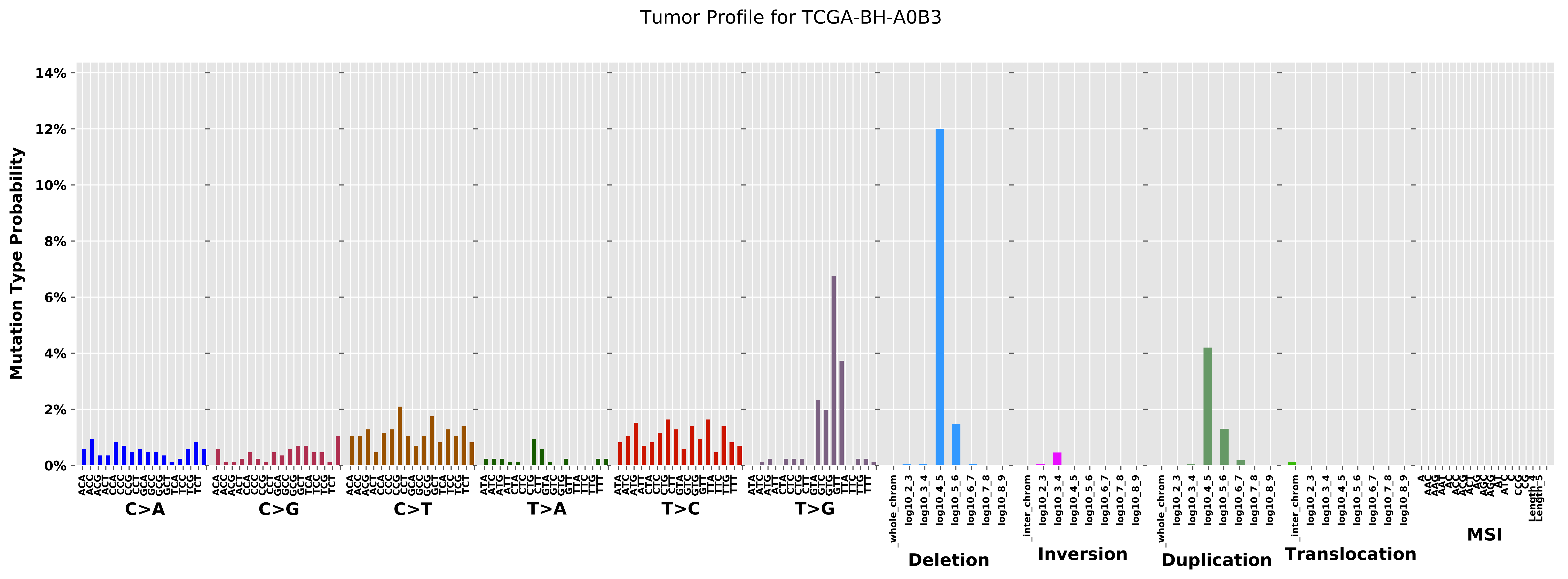

Tumor Profile for TCGA-BH-A0B3
Mutation Type Probability
0%
2%
4%
6%
8%
10%
12%
14%
C>A
C>G
C>T
T>A
T>C
T>G
Deletion
Inversion
Duplication
Translocation
MSI

Cancer processes Weights for TCGA-BH-A18U

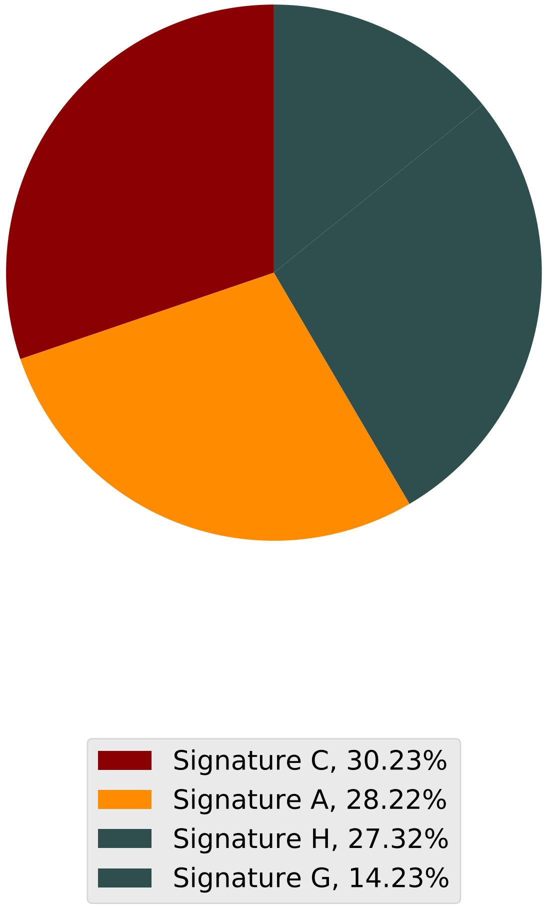

Tumor Profile for TCGA-BH-A18U

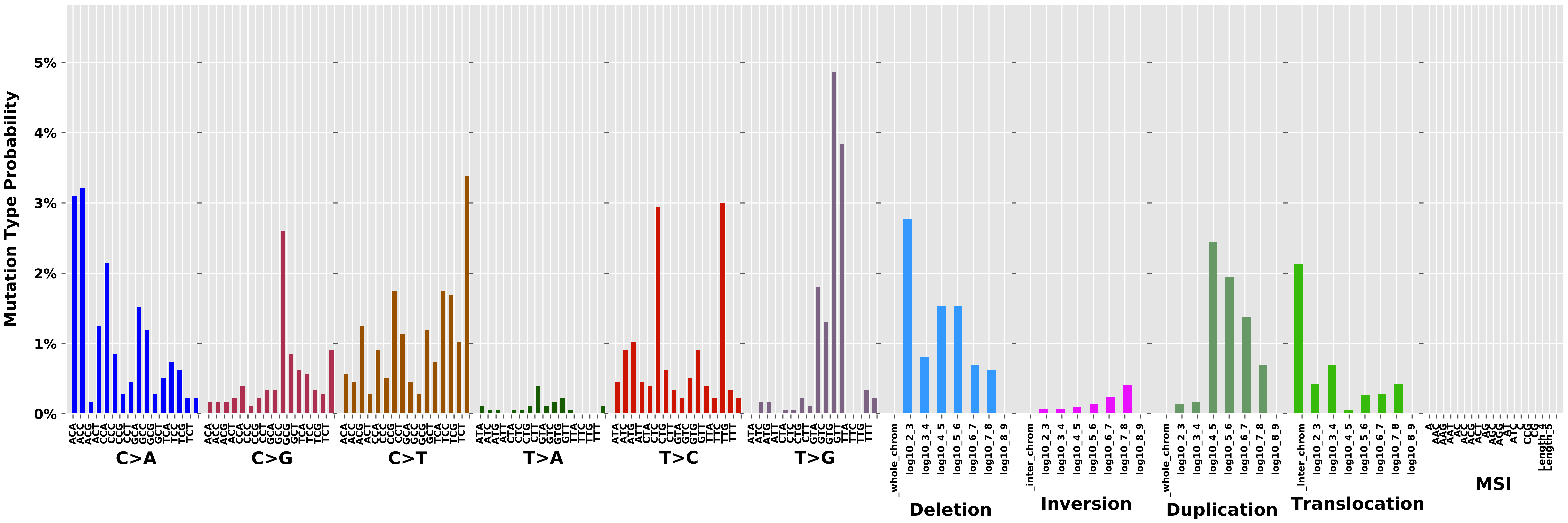

Cancer processes Weights for TCGA-GI-A2C9

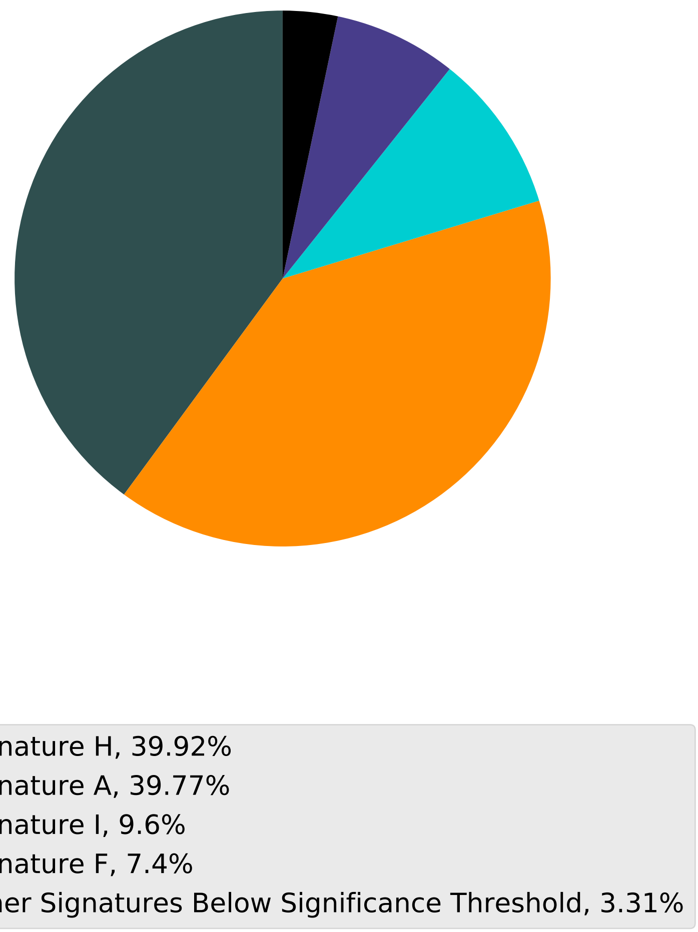

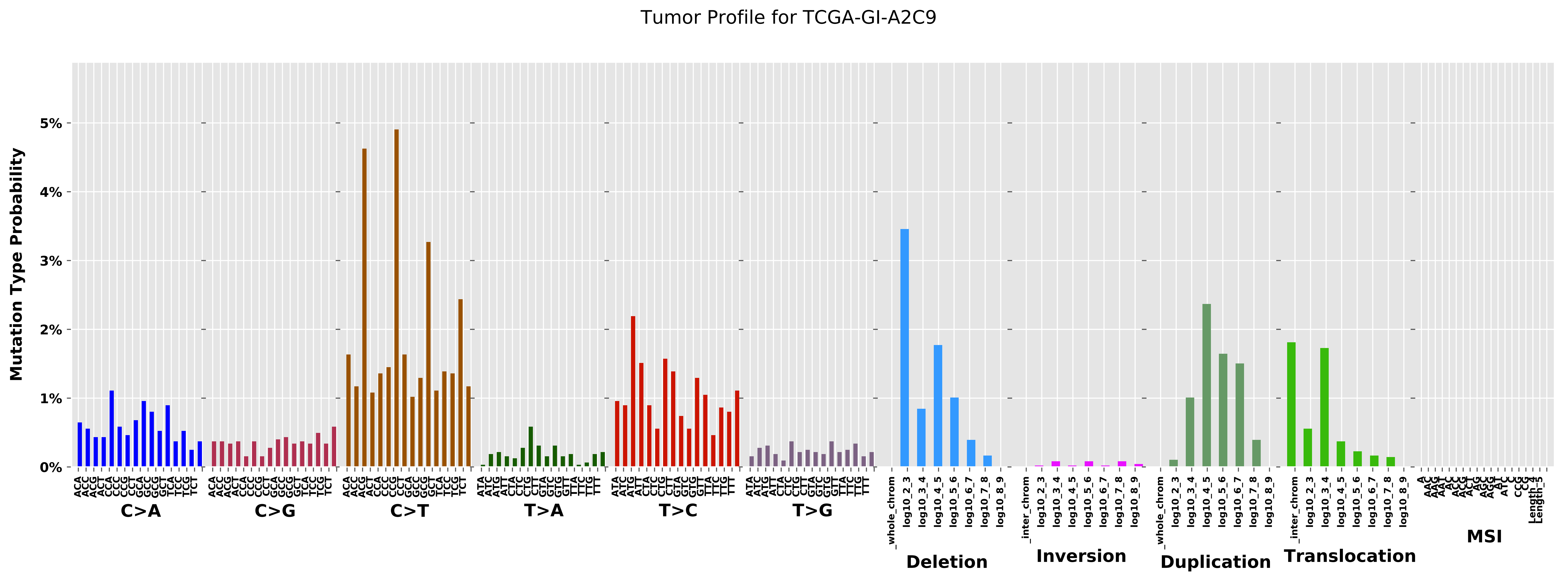

Tumor Profile for TCGA-GI-A2C9
Mutation Type Probability
0%
1%
2%
3%
4%
5%
C>A
C>G
C>T
T>A
T>C
T>G
Deletion
Inversion
Duplication
Translocation
MSI

Cancer processes Weights for TCGA-BH-A0DT

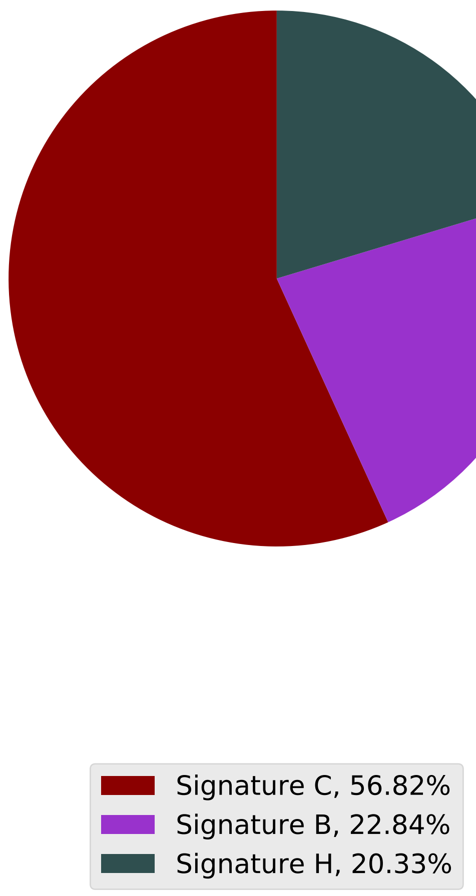

Tumor Profile for TCGA-BH-A0DT

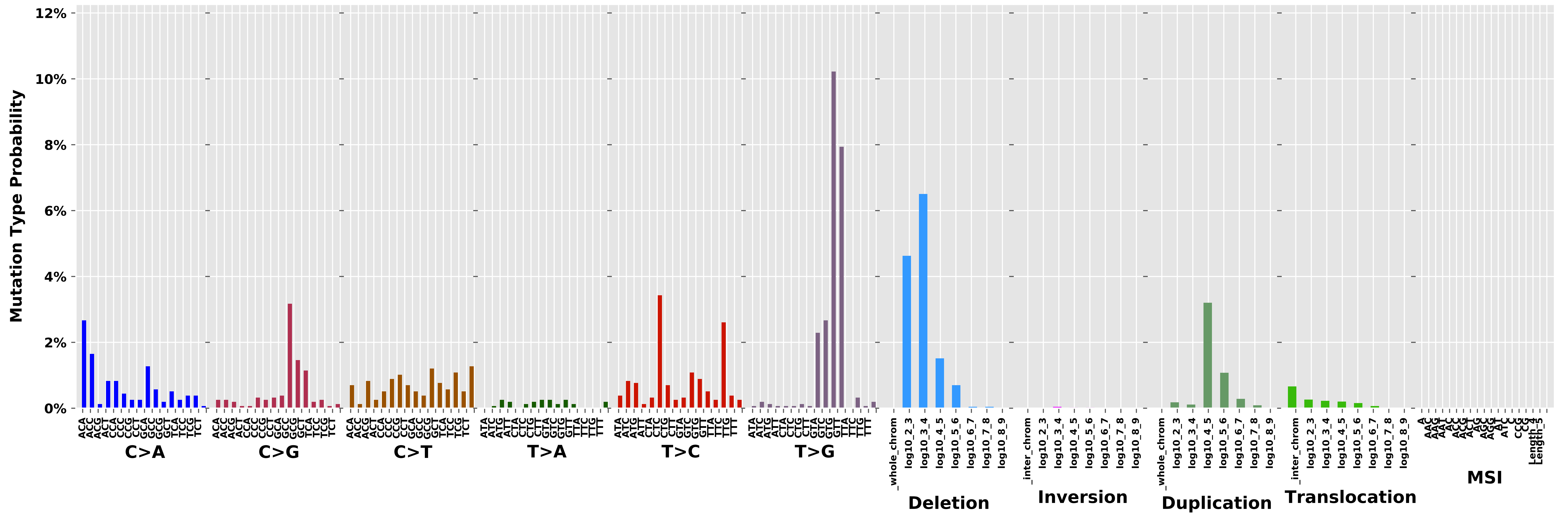

Cancer processes Weights for TCGA-A2-A0D4

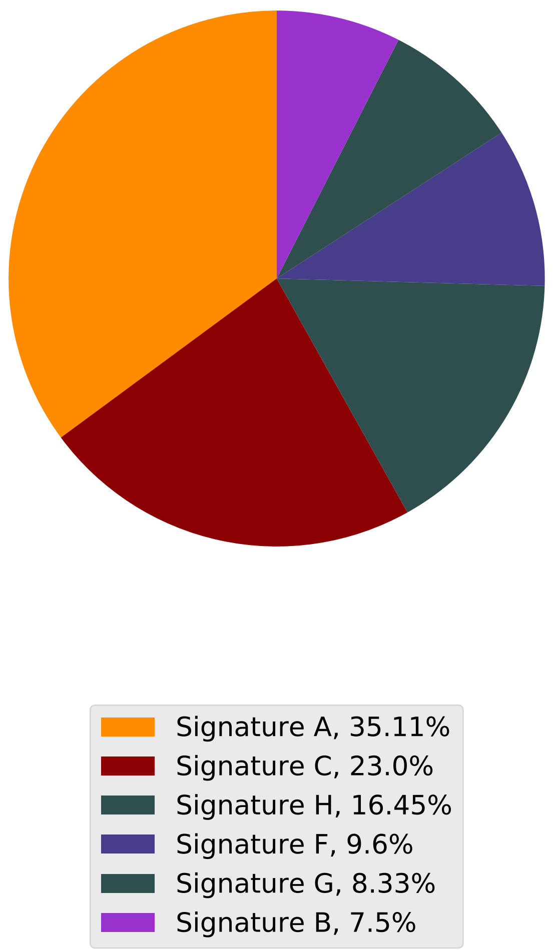

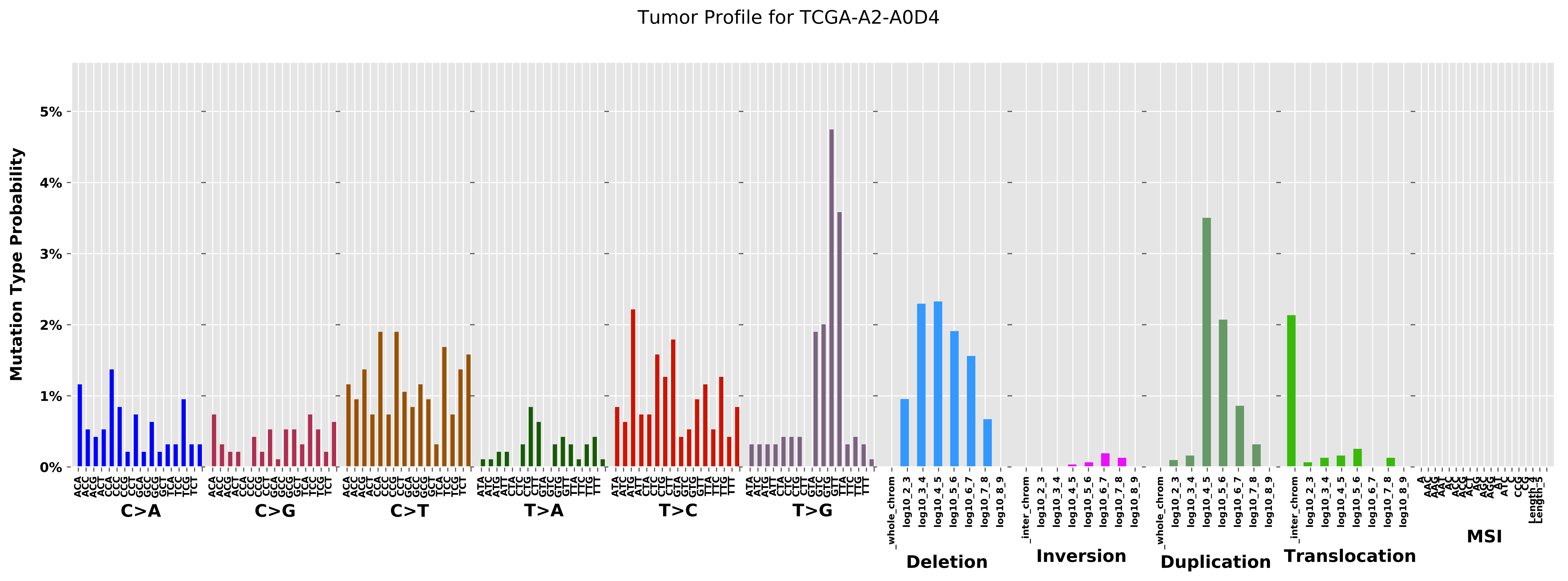

Tumor Profile for TCGA-A2-A0D4
Mutation Type Probability
0%
1%
2%
3%
4%
5%
C>A
C>G
C>T
T>A
T>C
T>G
Deletion
Inversion
Duplication
Translocation
MSI

#### Cancer processes Weights for TCGA-EW-A1P8

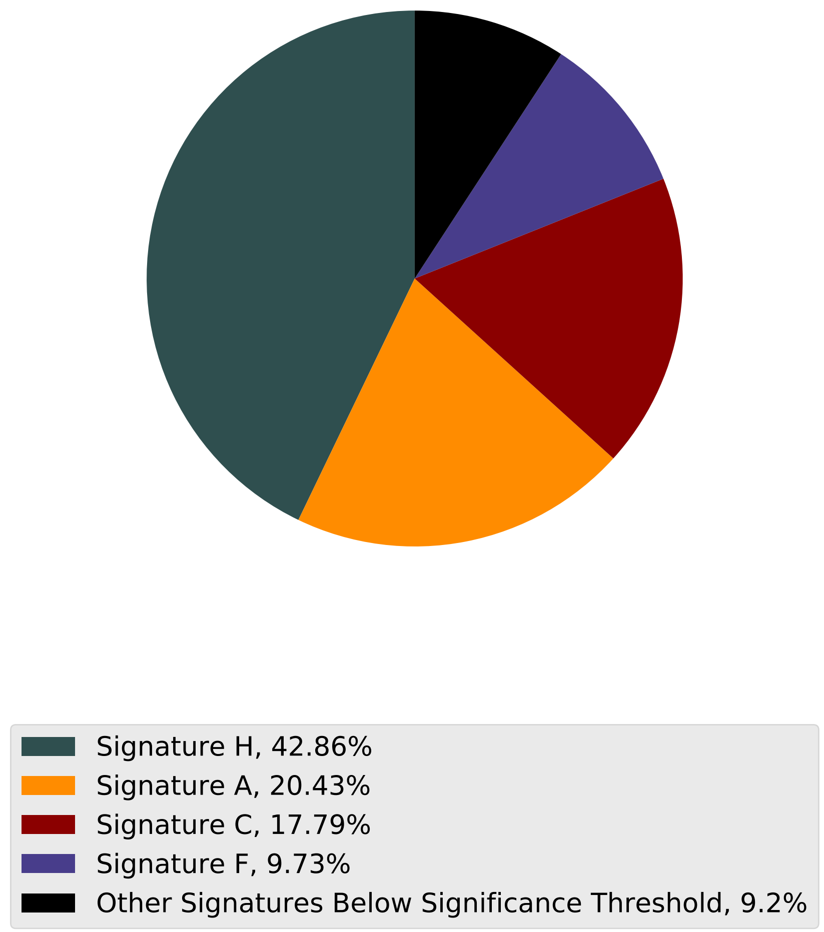

Tumor Profile for TCGA-EW-A1P8

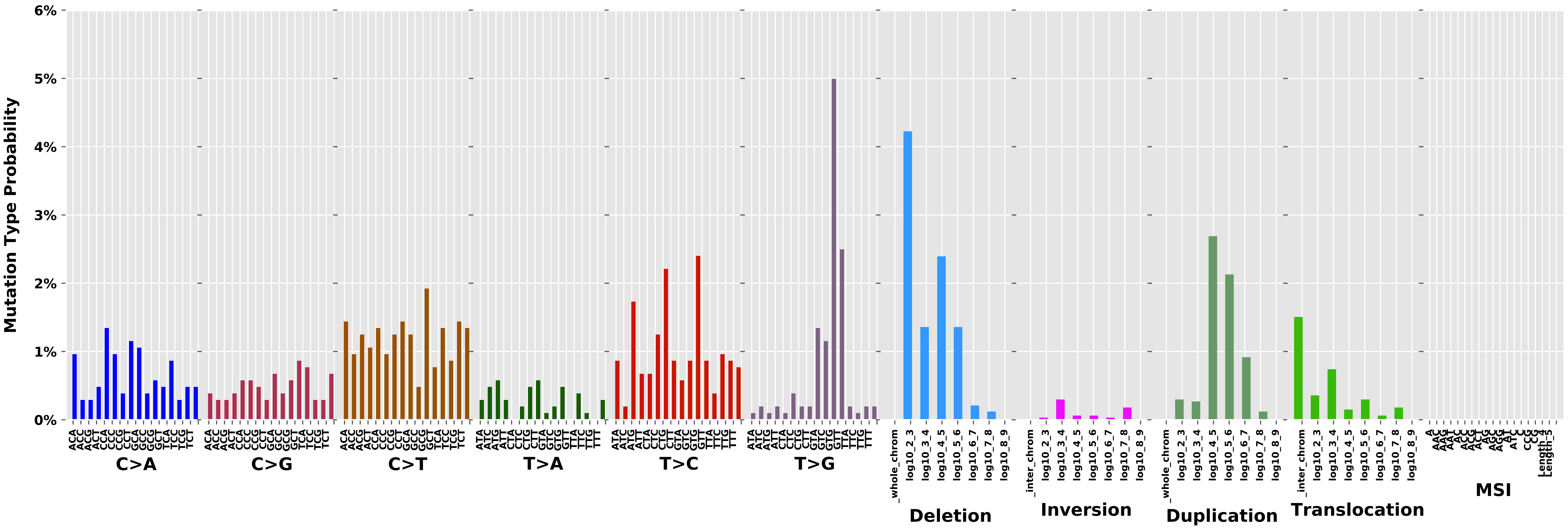

Cancer processes Weights for TCGA-A6-2683

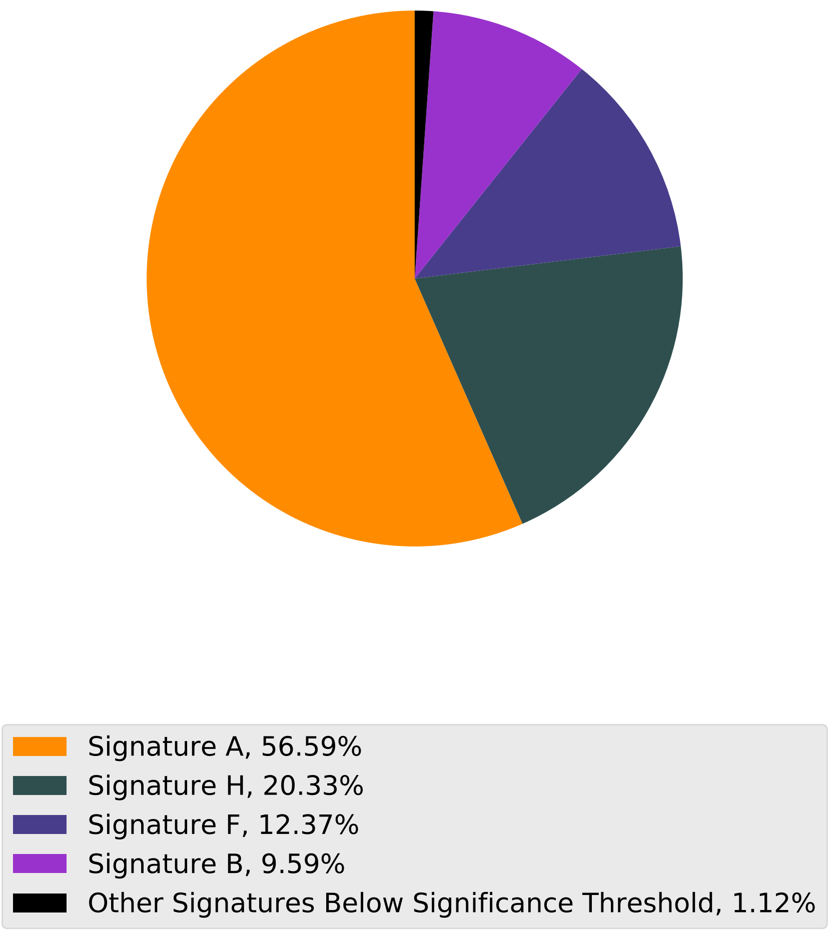

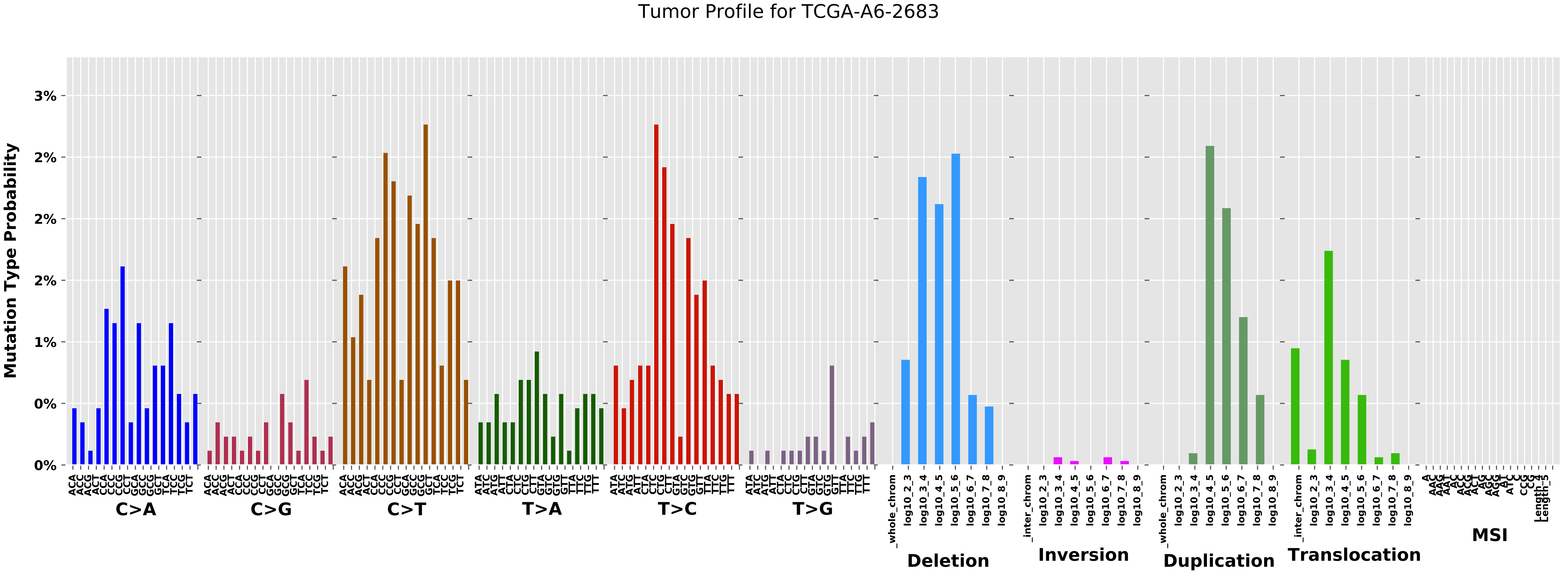
Tumor Profile for TCGA-A6-2683
Mutation Type Probability
3%
2%
2%
2%
1%
0%
0%
C>A
C>G
C>T
T>A
T>C
T>G
Deletion
Inversion
Duplication
Translocation
MSI

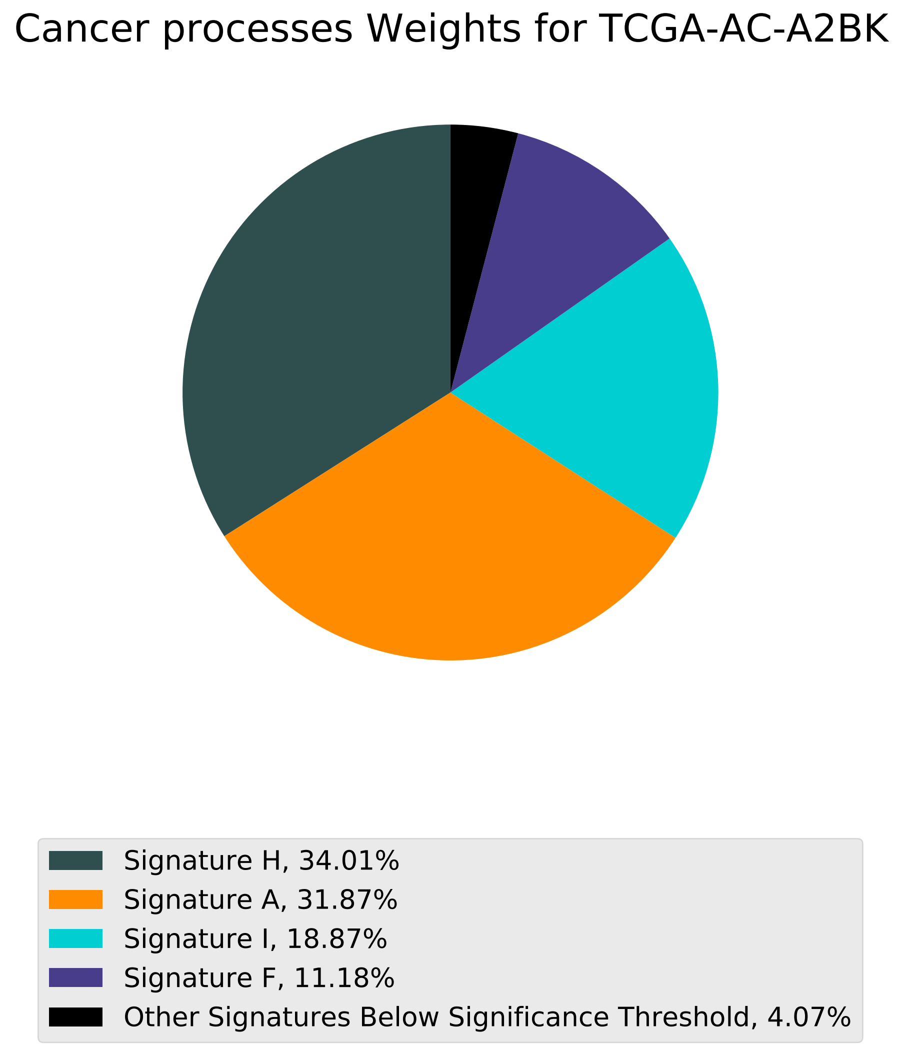
Cancer processes Weights for TCGA-AC-A2BK
Signature H, 34.01%
Signature A, 31.87%
Signature I, 18.87%
Signature F, 11.18%
Other Signatures Below Significance Threshold, 4.07%

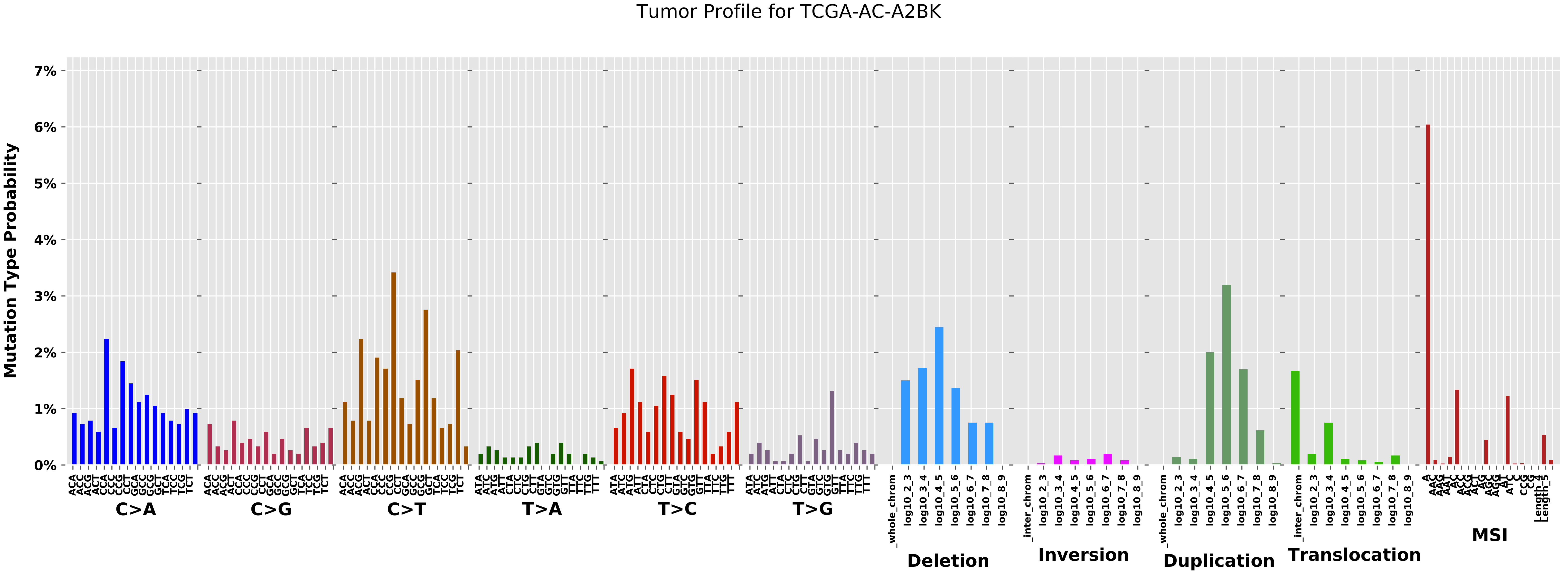

Tumor Profile for TCGA-AC-A2BK
Mutation Type Probability
7%
6%
5%
4%
3%
2%
1%
0%
C>A
C>G
C>T
T>A
T>C
T>G
Deletion
Inversion
Duplication
Translocation
MSI

Cancer processes Weights for TCGA-AG-A032

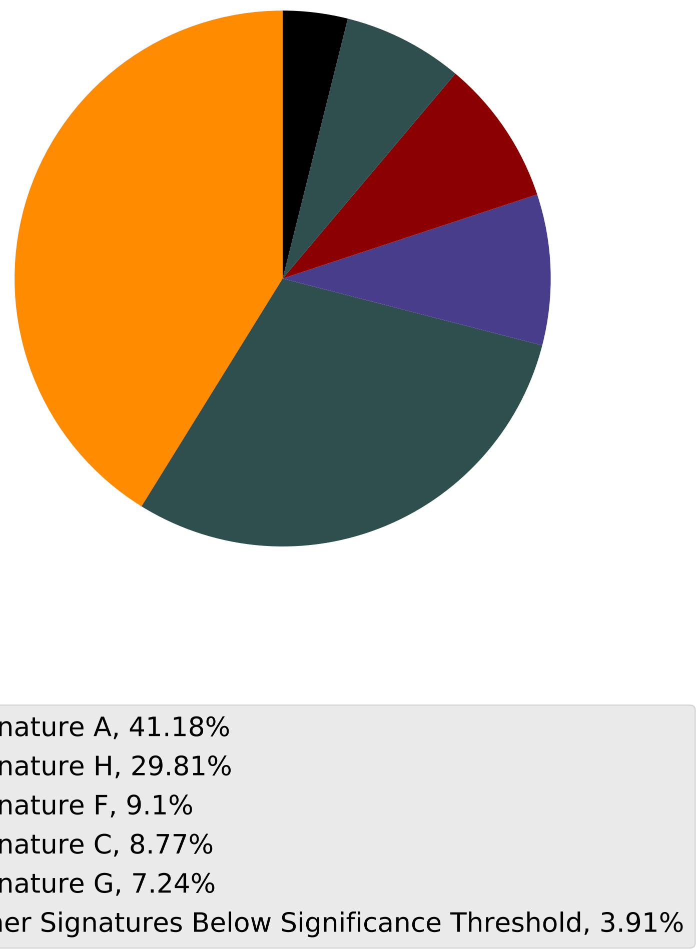

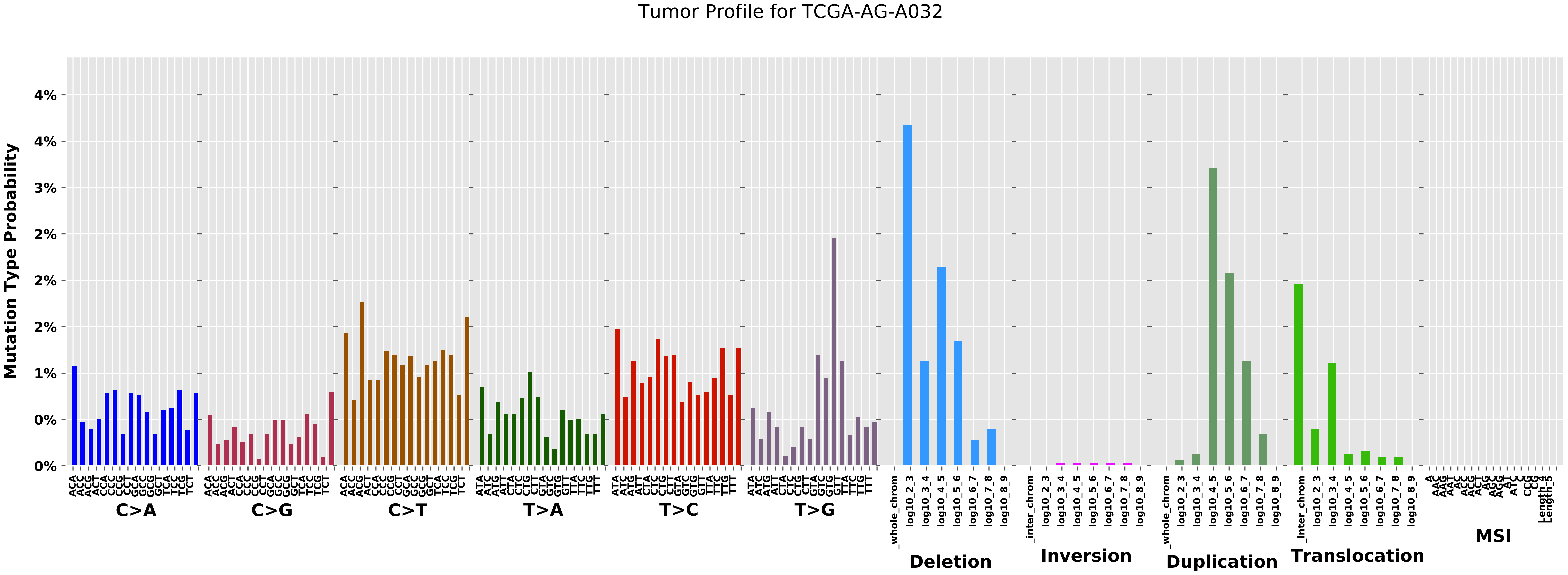

Tumor Profile for TCGA-AG-A032
Mutation Type Probability
4%
4%
3%
2%
2%
2%
1%
0%
0%
C>A
C>G
C>T
T>A
T>C
T>G
Deletion
Inversion
Duplication
Translocation
MSI

Cancer processes Weights for TCGA-B6-A0I2

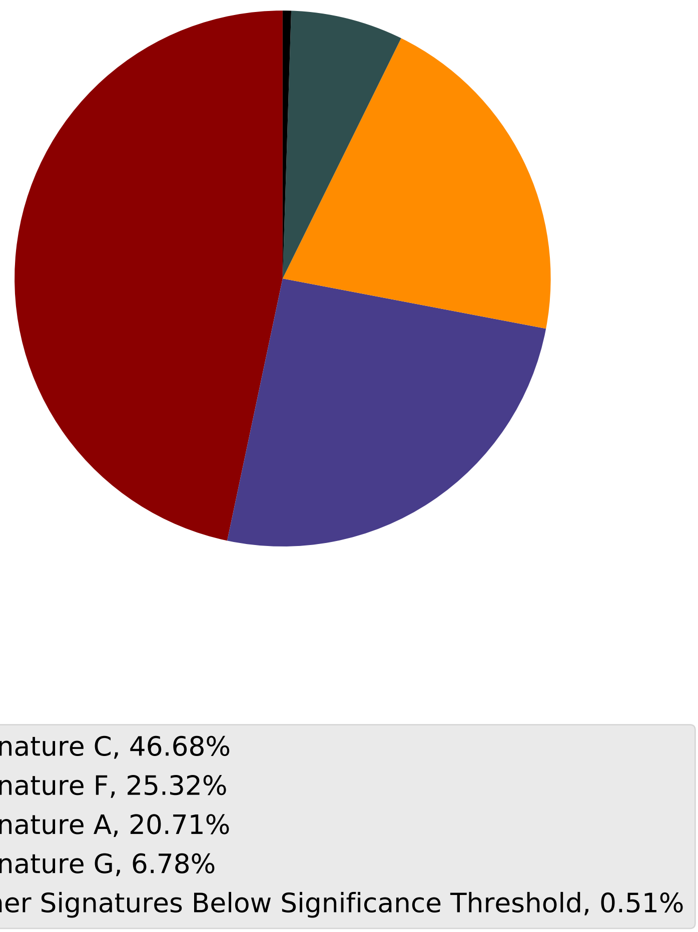

Tumor Profile for TCGA-B6-A0I2

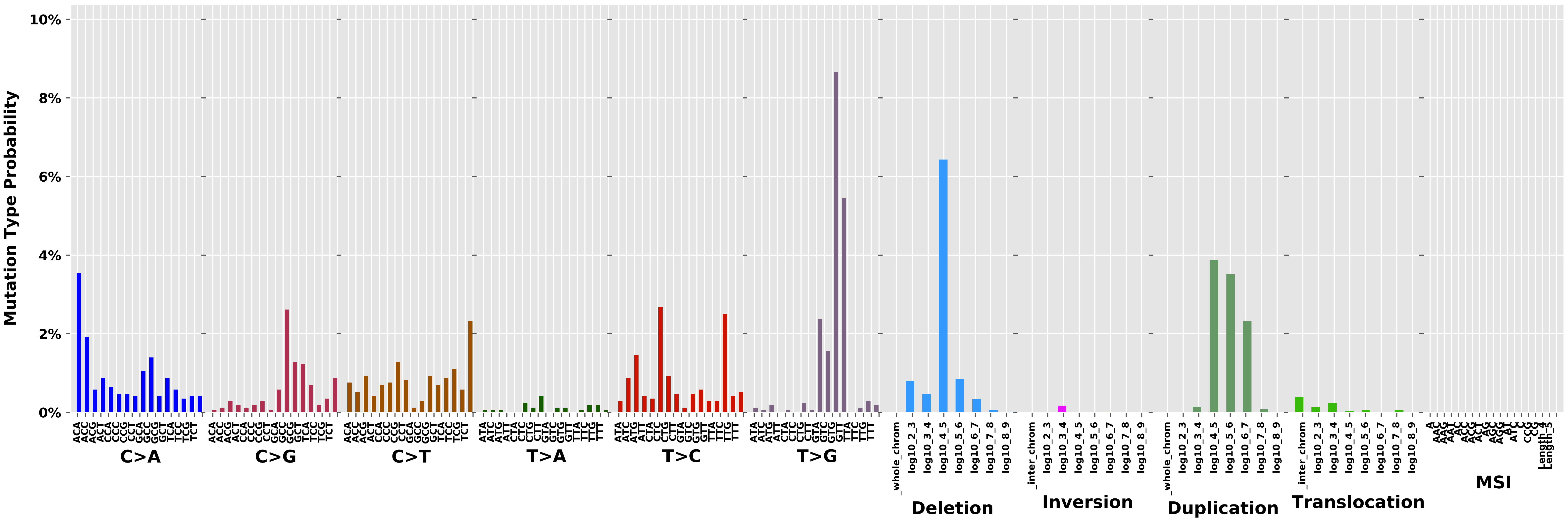

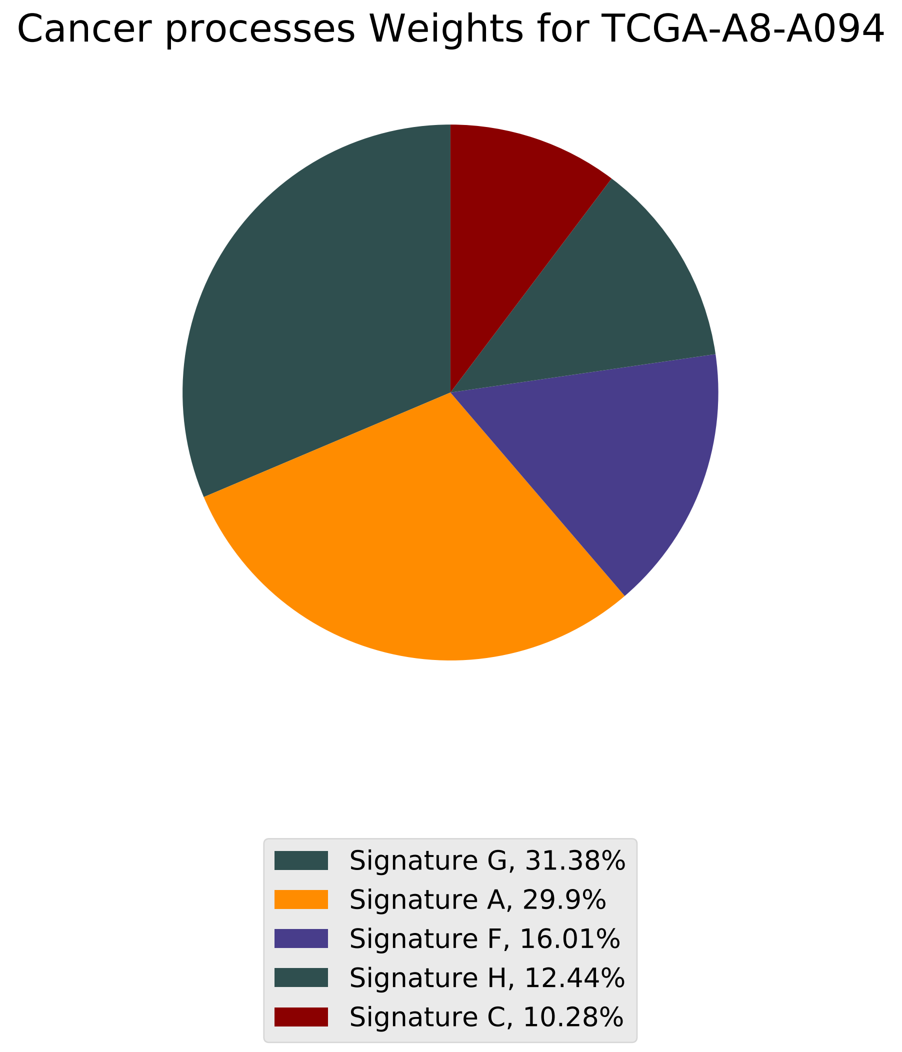
Cancer processes Weights for TCGA-A8-A094
Signature G, 31.38%
Signature A, 29.9%
Signature F, 16.01%
Signature H, 12.44%
Signature C, 10.28%

Tumor Profile for TCGA-A8-A094

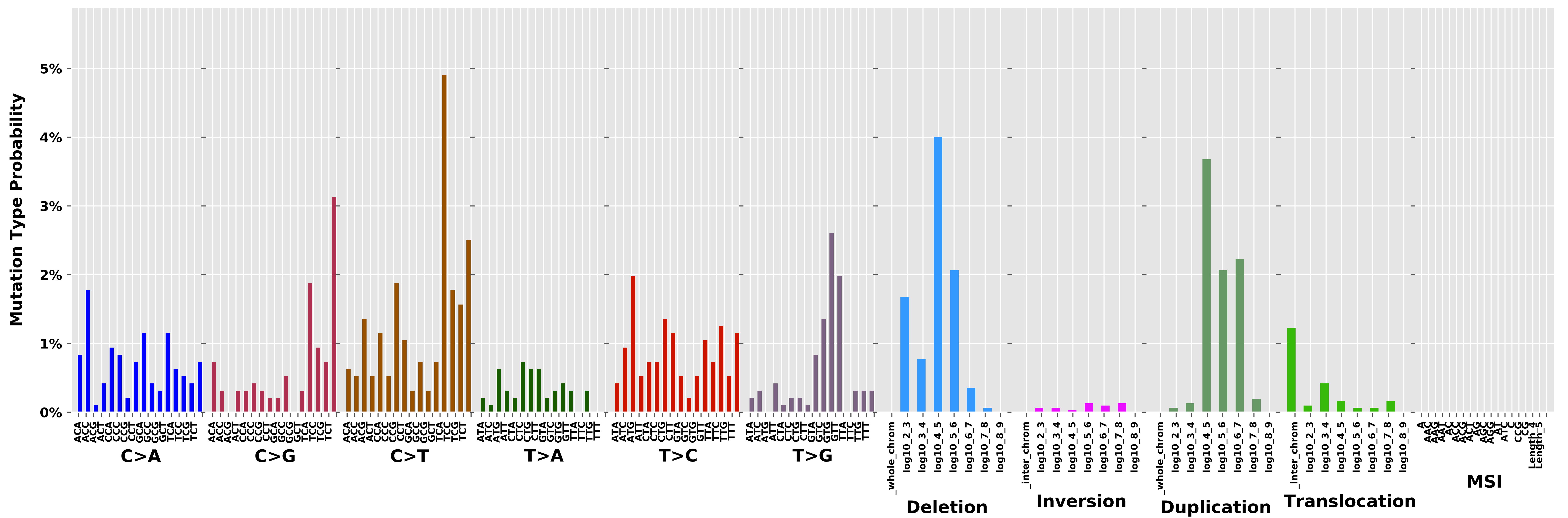

#### Cancer processes Weights for TCGA-AA-3994

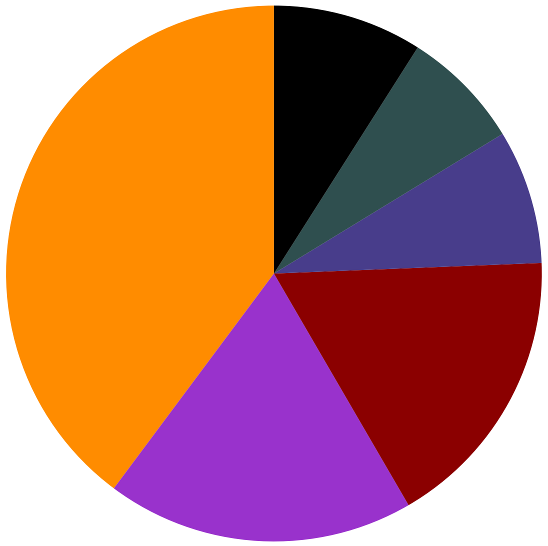

- Signature A, 39.81%
- Signature B, 18.58%
- Signature C, 17.25%
- Signature F, 8.09%
- Signature H, 7.26%
- Other Signatures Below Significance Threshold, 9.02%

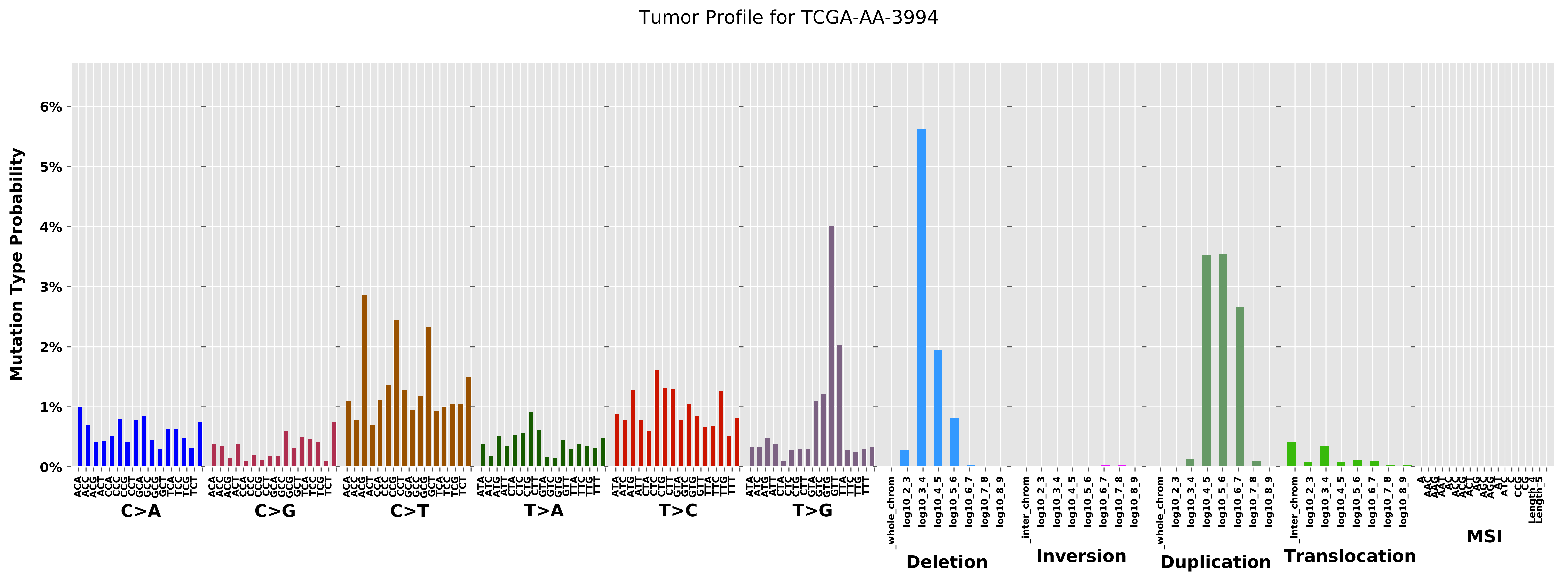
Tumor Profile for TCGA-AA-3994
Mutation Type Probability
6%
5%
4%
3%
2%
1%
0%
ACA ACC ACG ACT CCA CCC CCG CCT GCA GCC GCG GCT TCA TCC TCG TCT
C>A
ACA ACC ACG ACT CCA CCC CCG CCT GCA GCC GCG GCT TCA TCC TCG TCT
C>G
ACA ACC ACG ACT CCA CCC CCG CCT GCA GCC GCG GCT TCA TCC TCG TCT
C>T
ATA ATC ATG ATT CTA CTC CTG CTT GTA GTC GTG GTT TTA TTC TTG TTT
T>A
ATA ATC ATG ATT CTA CTC CTG CTT GTA GTC GTG GTT TTA TTC TTG TTT
T>C
ATA ATC ATG ATT CTA CTC CTG CTT GTA GTC GTG GTT TTA TTC TTG TTT
T>G
_whole_chrom log10_2_3 log10_3_4 log10_4_5 log10_5_6 log10_6_7 log10_7_8 log10_8_9
Deletion
_inter_chrom log10_2_3 log10_3_4 log10_4_5 log10_5_6 log10_6_7 log10_7_8 log10_8_9
Inversion
_whole_chrom log10_2_3 log10_3_4 log10_4_5 log10_5_6 log10_6_7 log10_7_8 log10_8_9
Duplication
_inter_chrom log10_2_3 log10_3_4 log10_4_5 log10_5_6 log10_6_7 log10_7_8 log10_8_9
Translocation
MSI

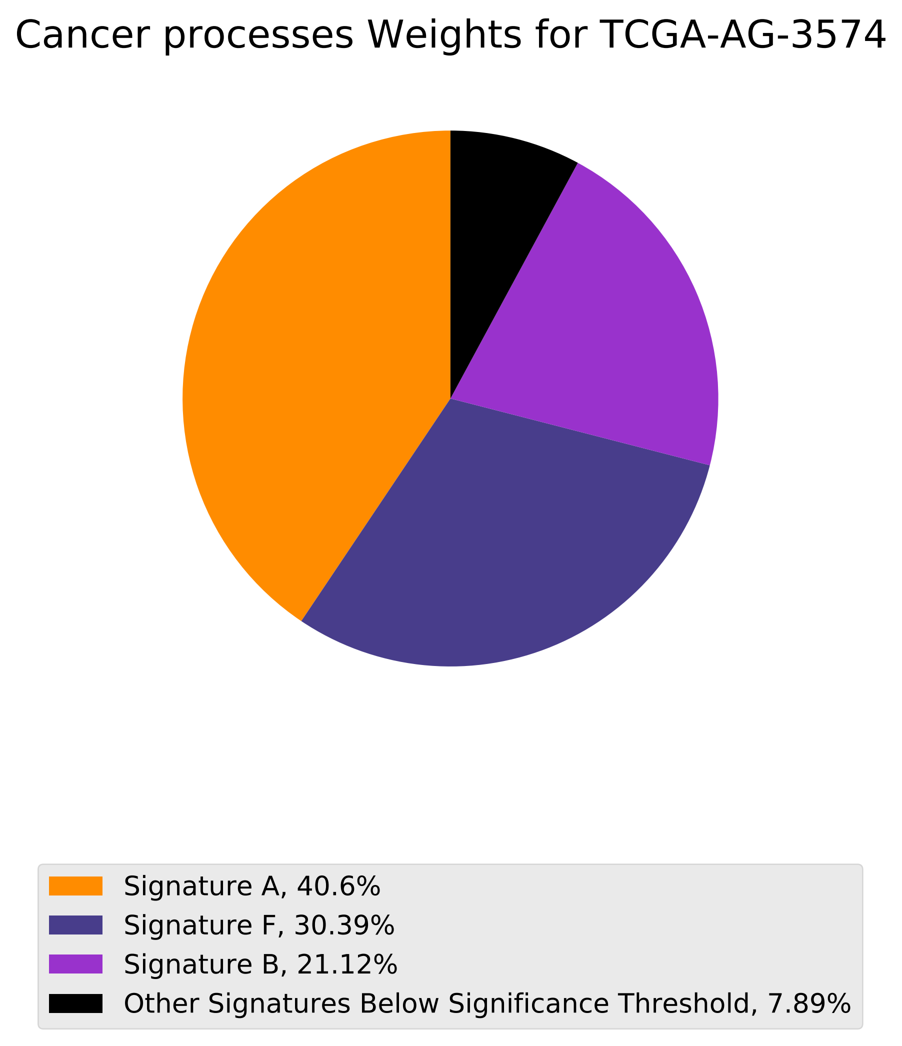
Cancer processes Weights for TCGA-AG-3574
Signature A, 40.6%
Signature F, 30.39%
Signature B, 21.12%
Other Signatures Below Significance Threshold, 7.89%

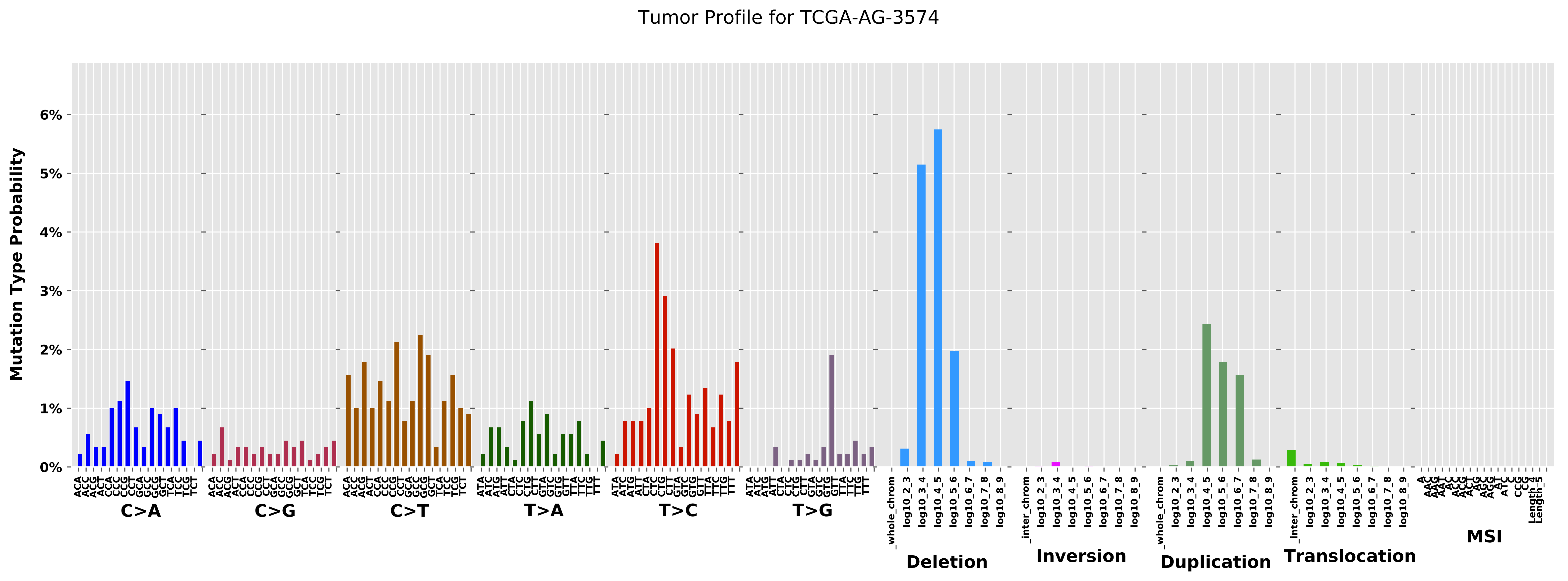

Tumor Profile for TCGA-AG-3574
Mutation Type Probability
0%
1%
2%
3%
4%
5%
6%
C>A
C>G
C>T
T>A
T>C
T>G
Deletion
Inversion
Duplication
Translocation
MSI

Cancer processes Weights for TCGA-AO-A03L

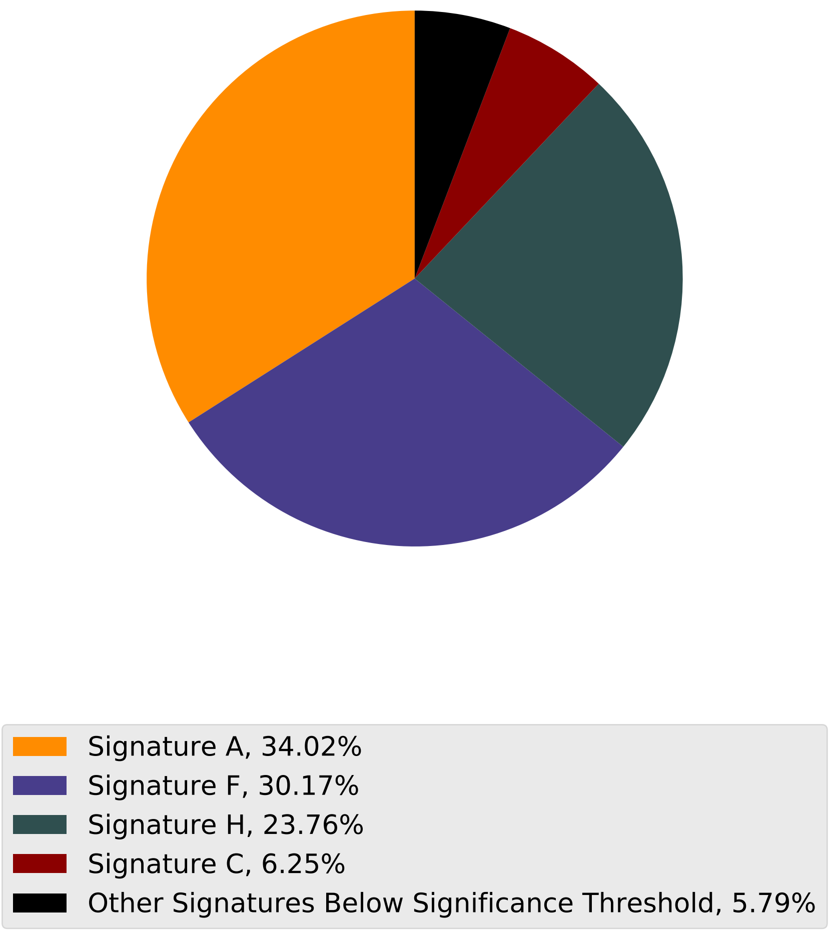

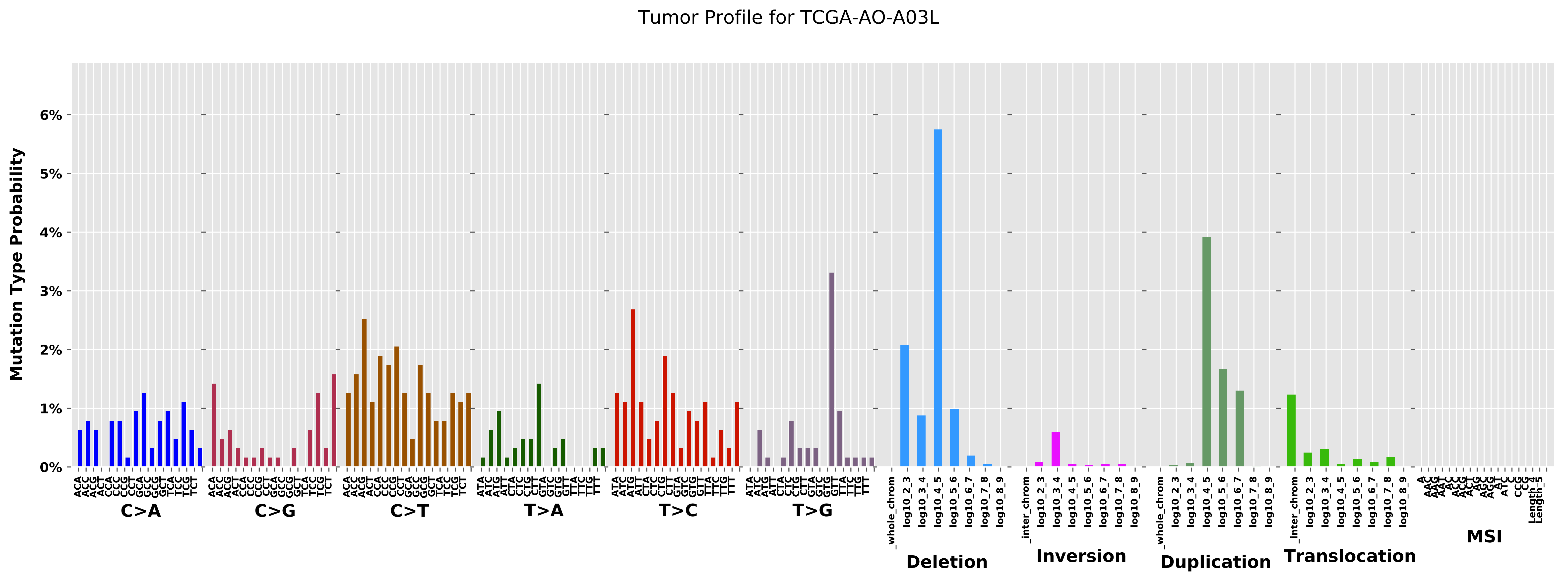
Tumor Profile for TCGA-AO-A03L
Mutation Type Probability
0%
1%
2%
3%
4%
5%
6%
C>A
C>G
C>T
T>A
T>C
T>G
Deletion
Inversion
Duplication
Translocation
MSI

Cancer processes Weights for TCGA-A6-6141

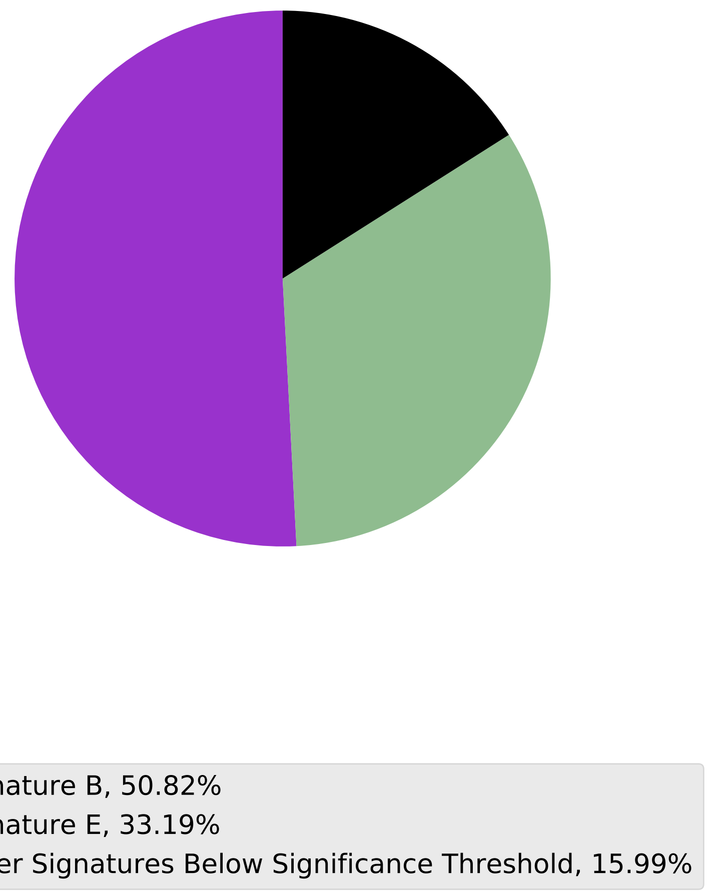

Tumor Profile for TCGA-A6-6141

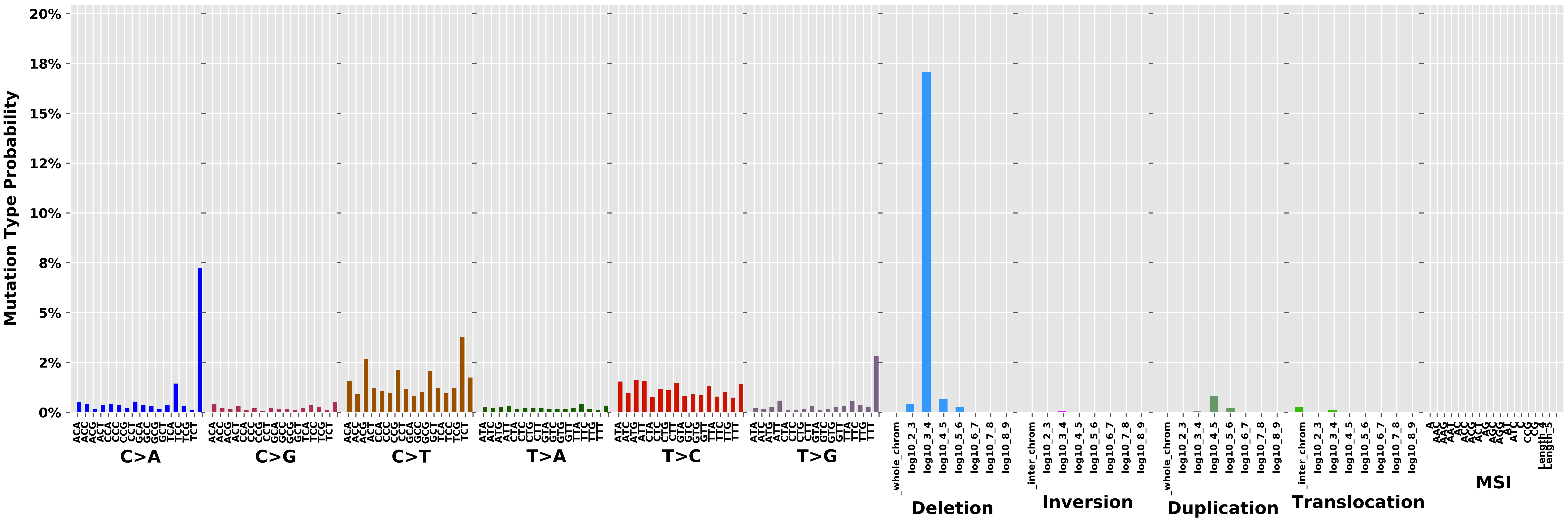

Cancer processes Weights for TCGA-AA-A02O

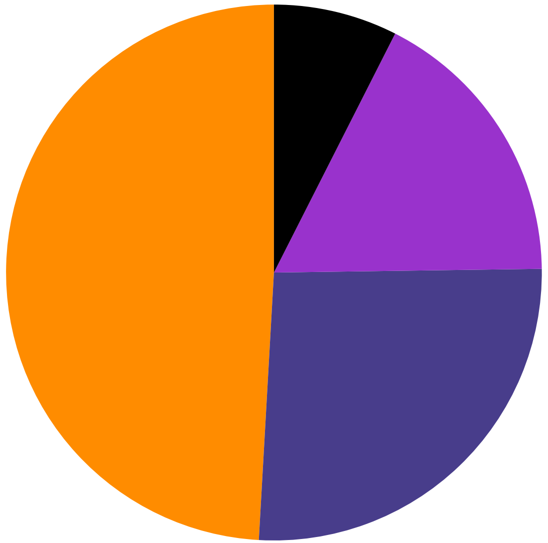

Signature A, 49.1%
Signature F, 26.12%
Signature B, 17.31%
Other Signatures Below Significance Threshold, 7.48%

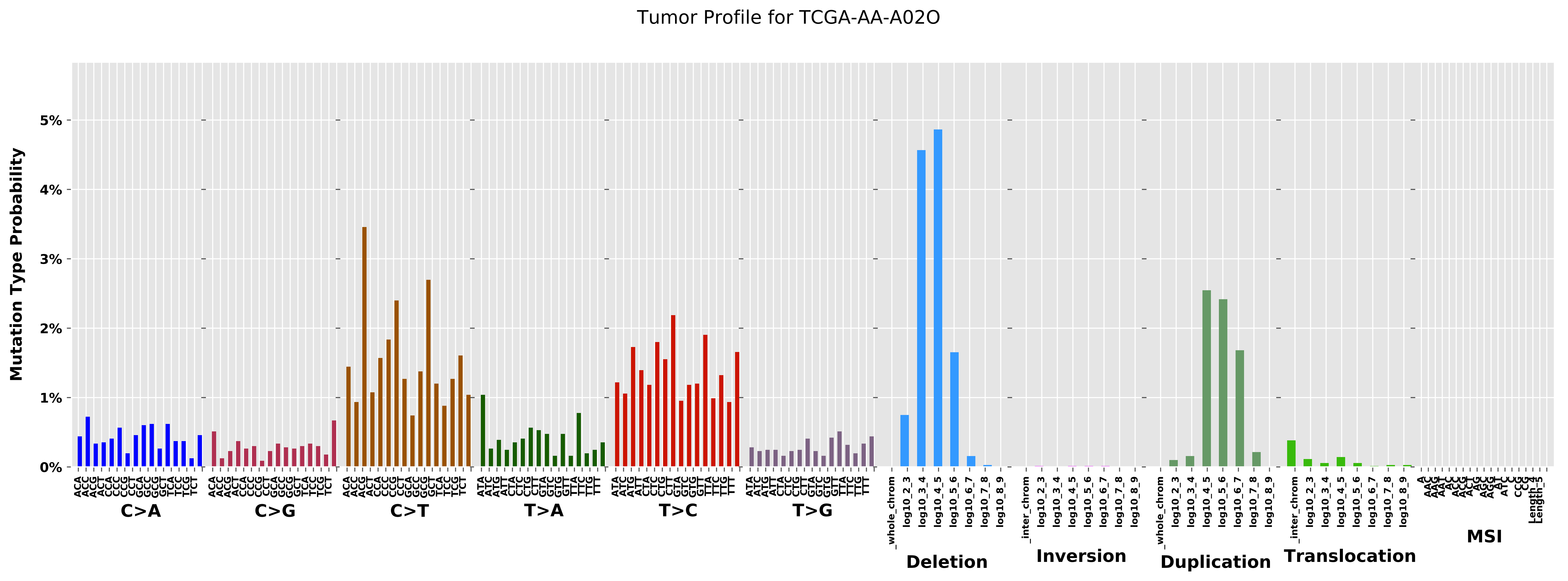

Tumor Profile for TCGA-AA-A02O
Mutation Type Probability
5%
4%
3%
2%
1%
0%
C>A
C>G
C>T
T>A
T>C
T>G
Deletion
Inversion
Duplication
Translocation
MSI

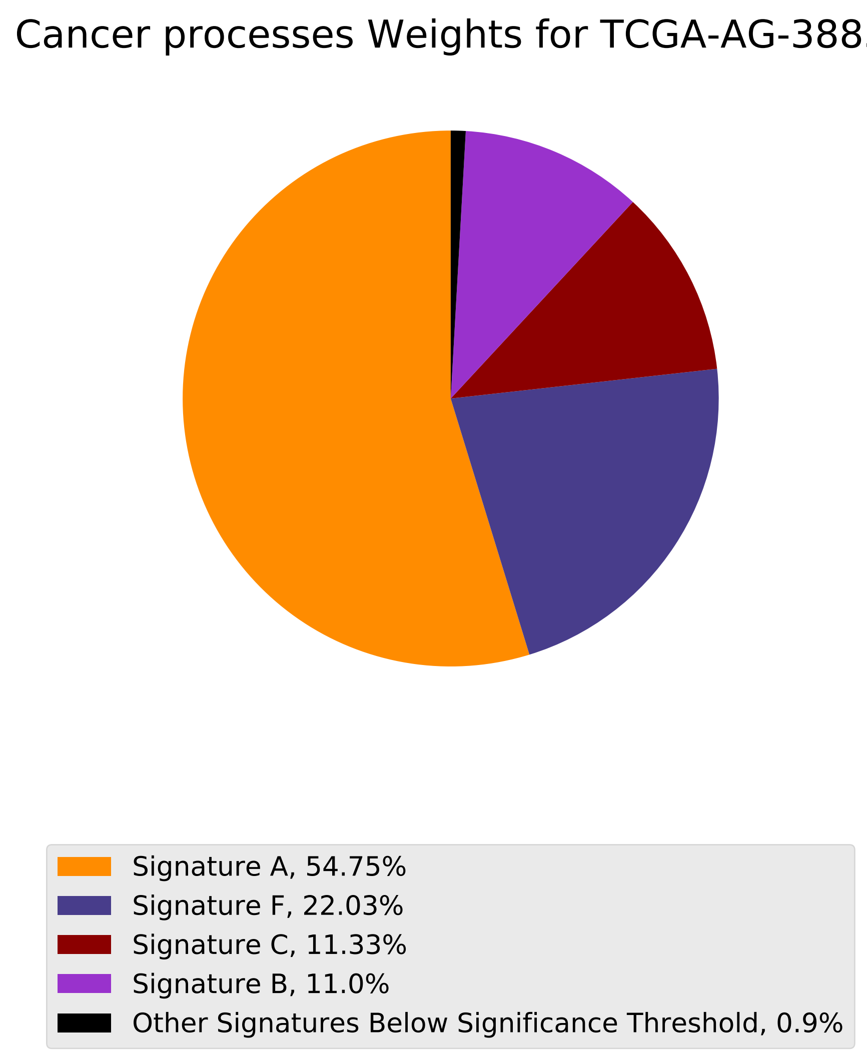
Cancer processes Weights for TCGA-AG-3885
Signature A, 54.75%
Signature F, 22.03%
Signature C, 11.33%
Signature B, 11.0%
Other Signatures Below Significance Threshold, 0.9%

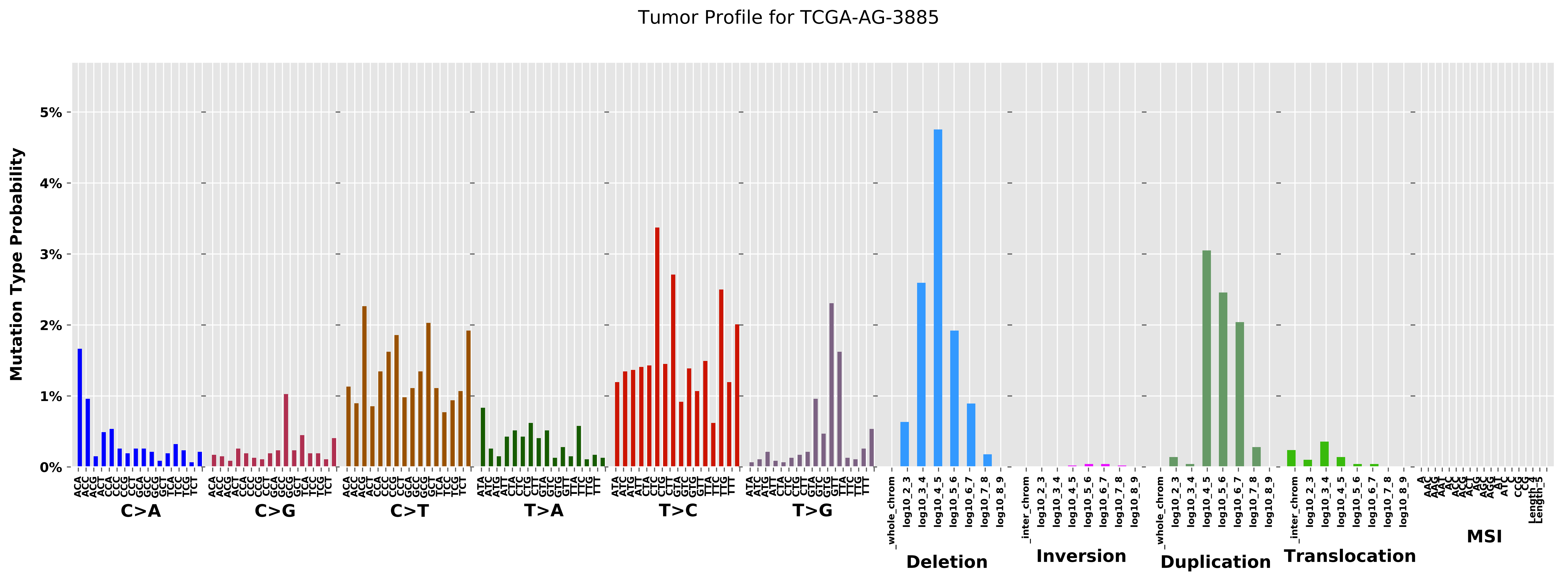

Tumor Profile for TCGA-AG-3885
Mutation Type Probability
0%
1%
2%
3%
4%
5%
C>A
C>G
C>T
T>A
T>C
T>G
_whole_chrom
_inter_chrom
log10_2_3
log10_3_4
log10_4_5
log10_5_6
log10_6_7
log10_7_8
log10_8_9
Deletion
Inversion
Duplication
Translocation
MSI

Cancer processes Weights for TCGA-AA-A01T

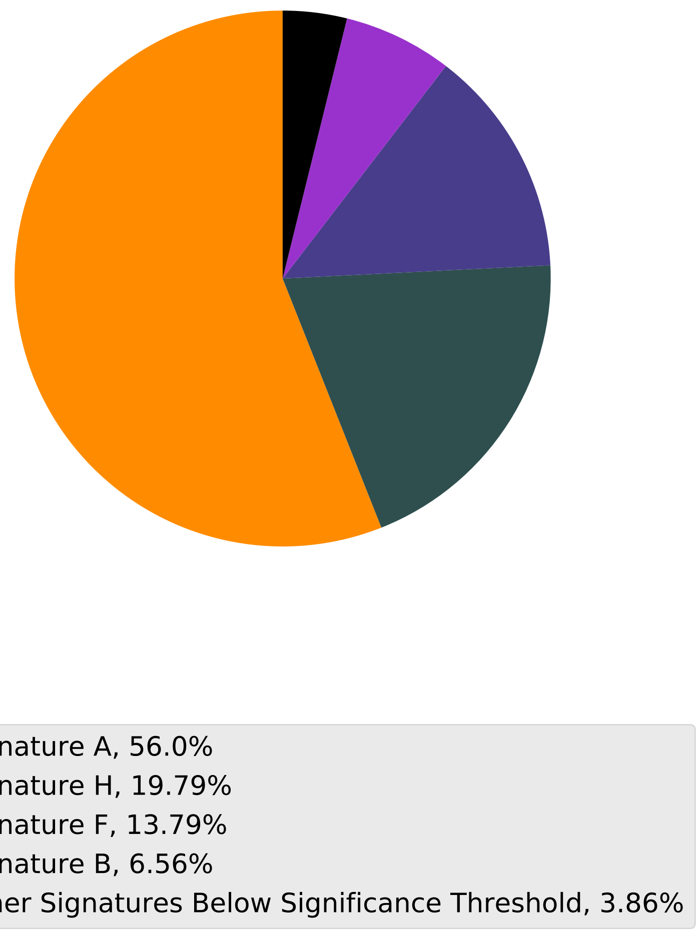

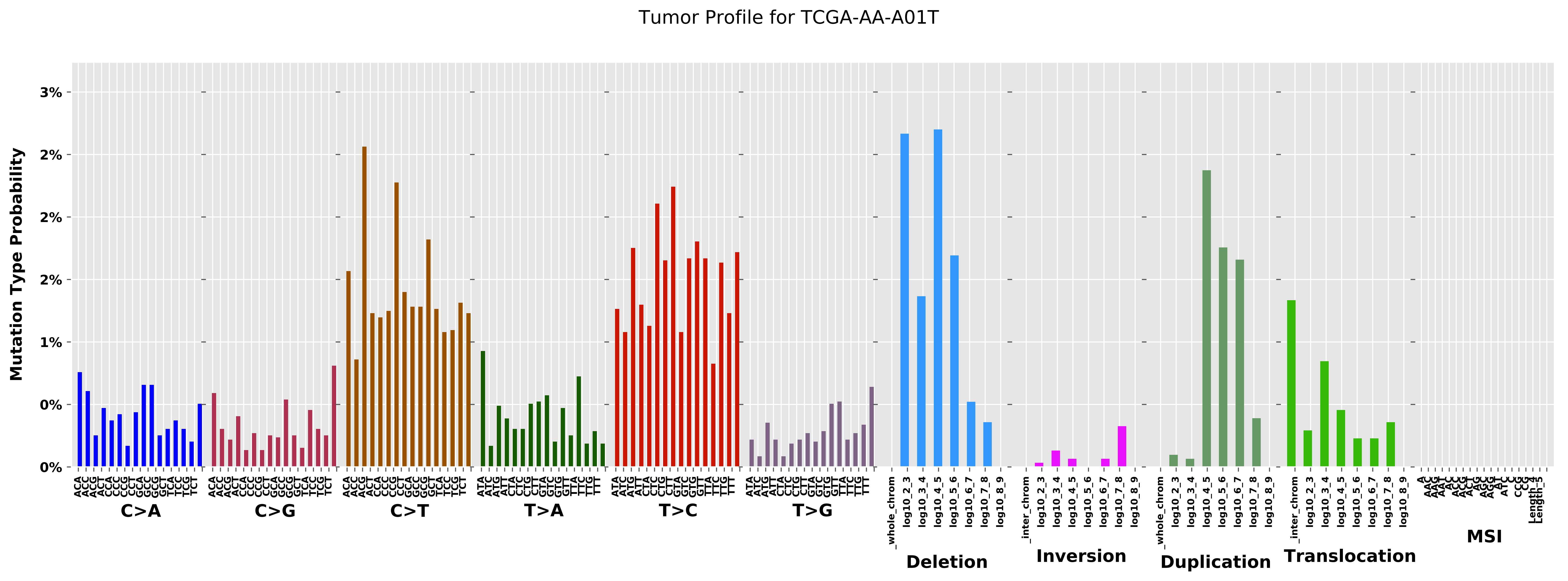

Tumor Profile for TCGA-AA-A01T
Mutation Type Probability
3%
2%
2%
2%
1%
0%
0%
C>A
C>G
C>T
T>A
T>C
T>G
_whole_chrom
log10_2_3
log10_3_4
log10_4_5
log10_5_6
log10_6_7
log10_7_8
log10_8_9
Deletion
_inter_chrom
Inversion
Duplication
Translocation
MSI

#### Cancer processes Weights for TCGA-A8-A07I

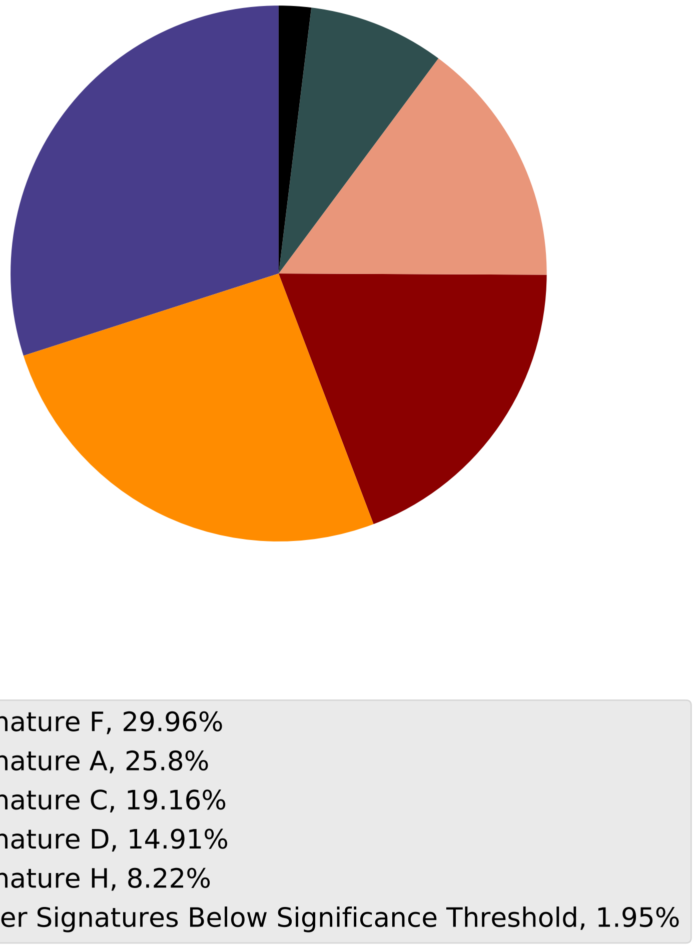

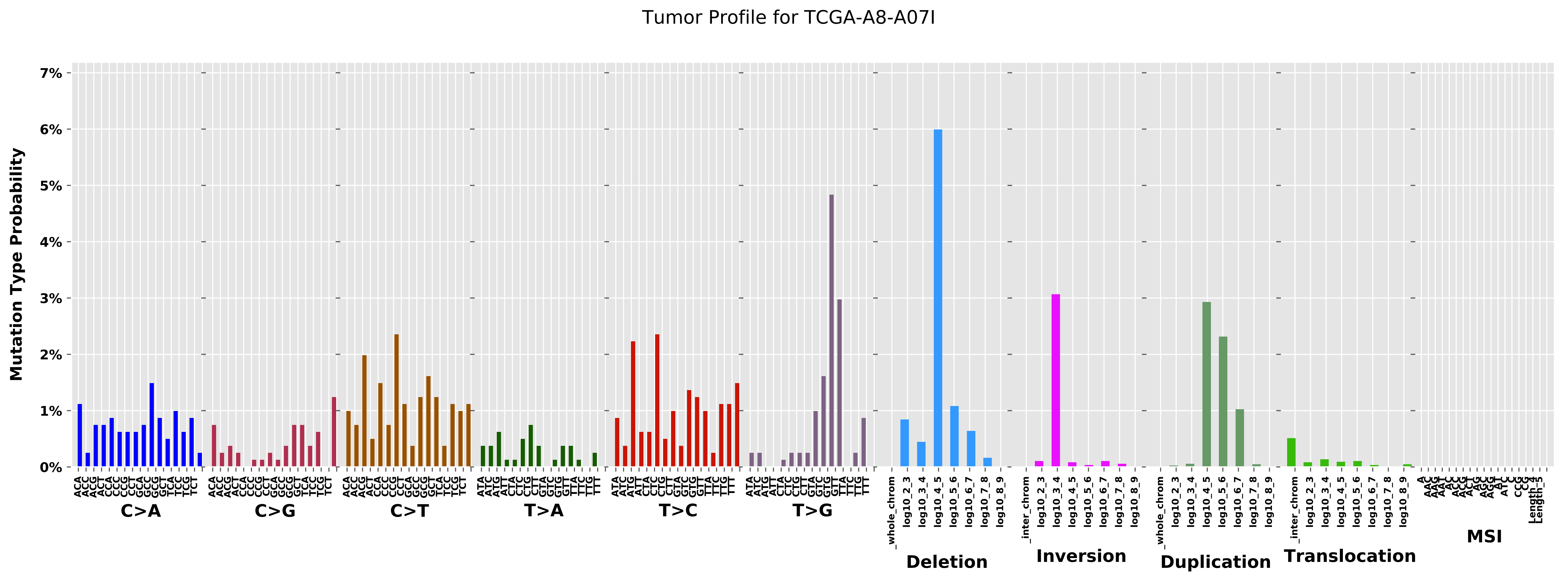
Tumor Profile for TCGA-A8-A07I
Mutation Type Probability
0%
1%
2%
3%
4%
5%
6%
7%
C>A
C>G
C>T
T>A
T>C
T>G
_whole_chrom
log10_2_3
log10_3_4
log10_4_5
log10_5_6
log10_6_7
log10_7_8
log10_8_9
Deletion
_inter_chrom
Inversion
Duplication
Translocation
MSI

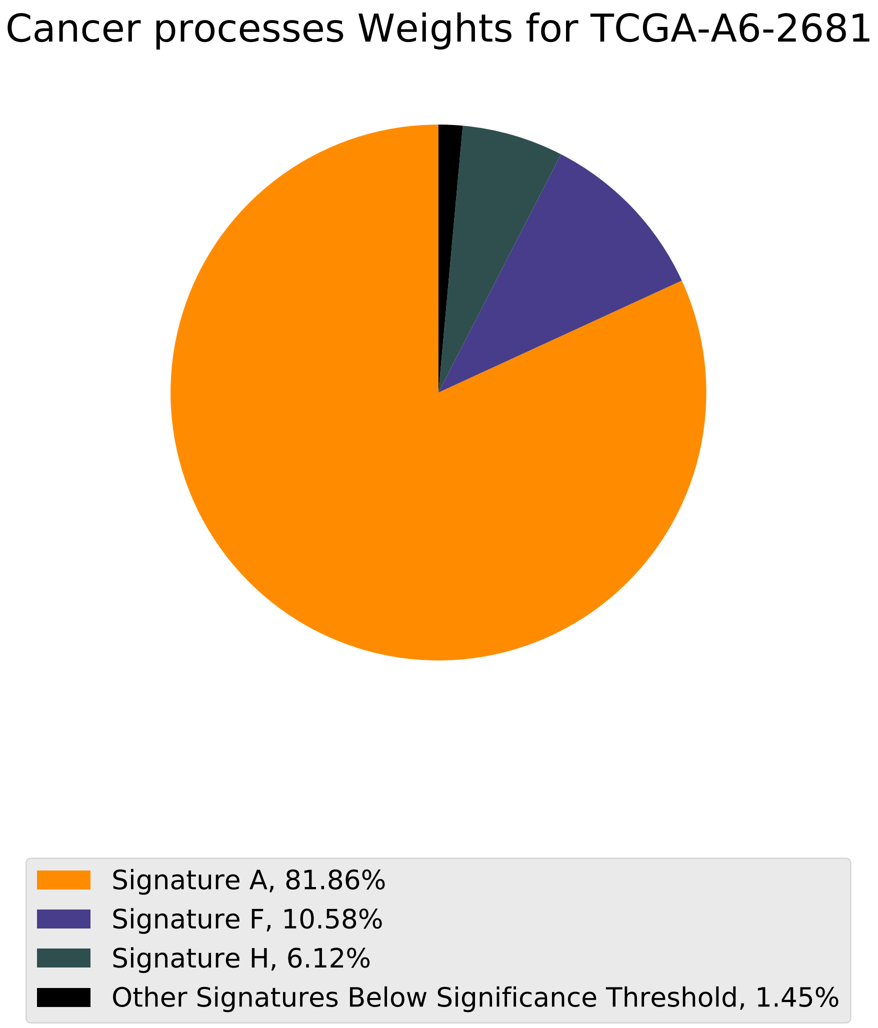
Cancer processes Weights for TCGA-A6-2681
Signature A, 81.86%
Signature F, 10.58%
Signature H, 6.12%
Other Signatures Below Significance Threshold, 1.45%

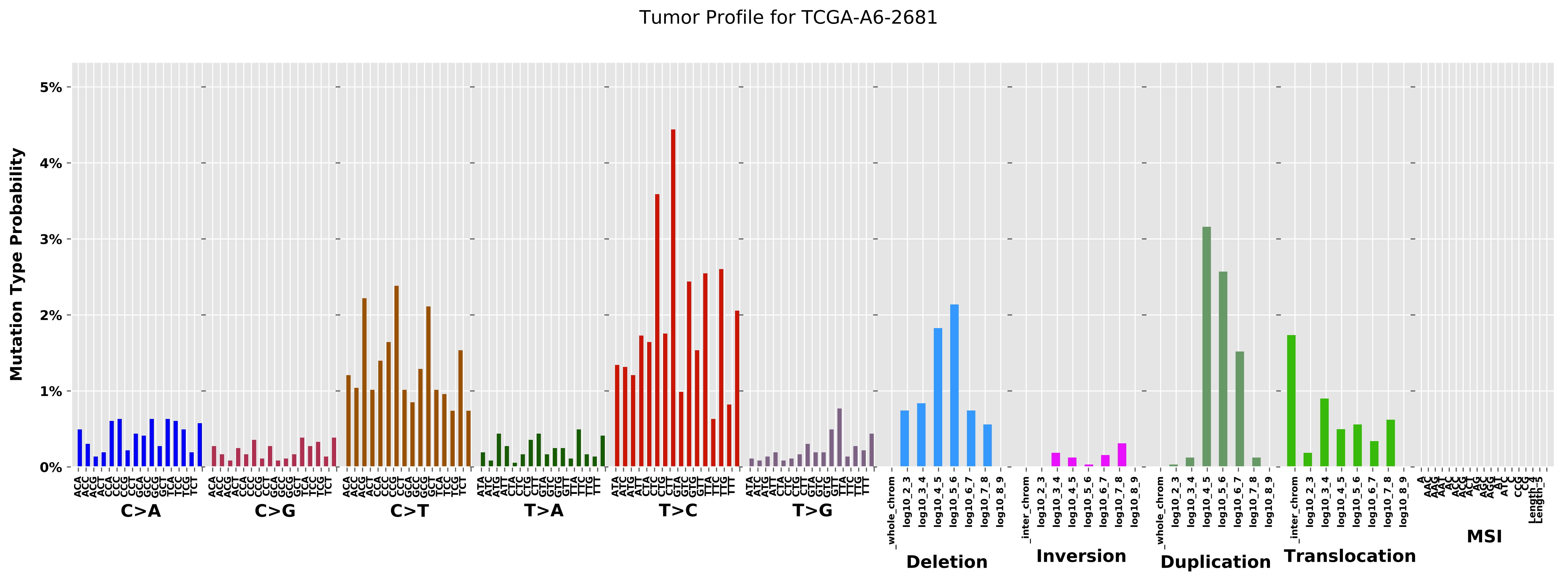

Tumor Profile for TCGA-A6-2681
Mutation Type Probability
0%
1%
2%
3%
4%
5%
C>A
C>G
C>T
T>A
T>C
T>G
Deletion
Inversion
Duplication
Translocation
MSI
_whole_chrom
_inter_chrom
log10_2_3
log10_3_4
log10_4_5
log10_5_6
log10_6_7
log10_7_8
log10_8_9

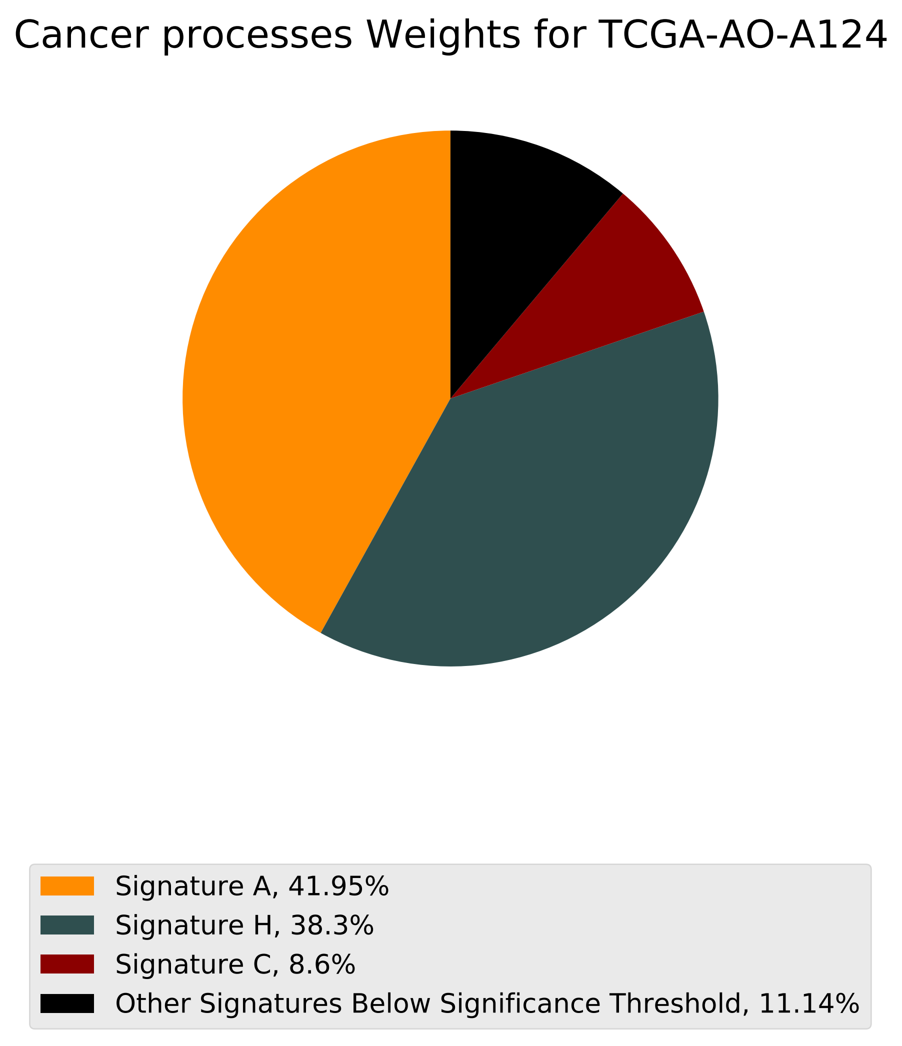
Cancer processes Weights for TCGA-AO-A124
Signature A, 41.95%
Signature H, 38.3%
Signature C, 8.6%
Other Signatures Below Significance Threshold, 11.14%

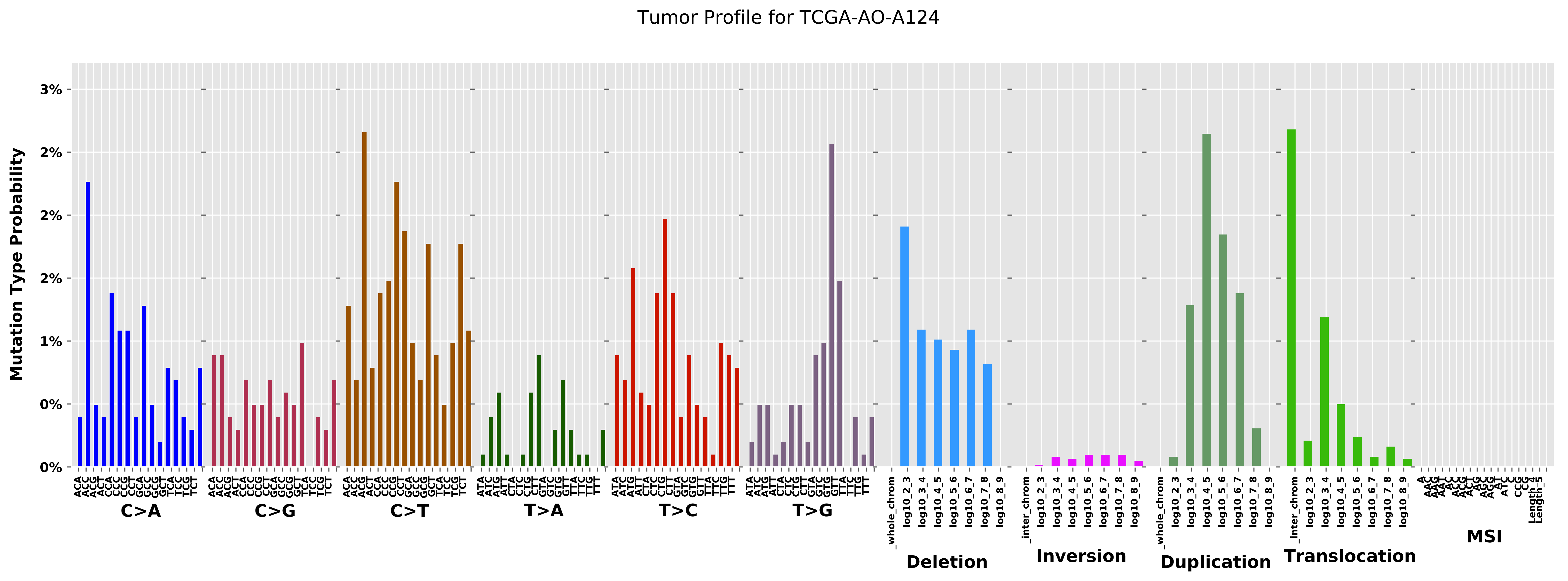

Tumor Profile for TCGA-AO-A124
Mutation Type Probability
3%
2%
2%
2%
1%
0%
0%
C>A
C>G
C>T
T>A
T>C
T>G
Deletion
Inversion
Duplication
Translocation
MSI
_whole_chrom
_inter_chrom
log10_2_3
log10_3_4
log10_4_5
log10_5_6
log10_6_7
log10_7_8
log10_8_9

Cancer processes Weights for TCGA-A2-A259

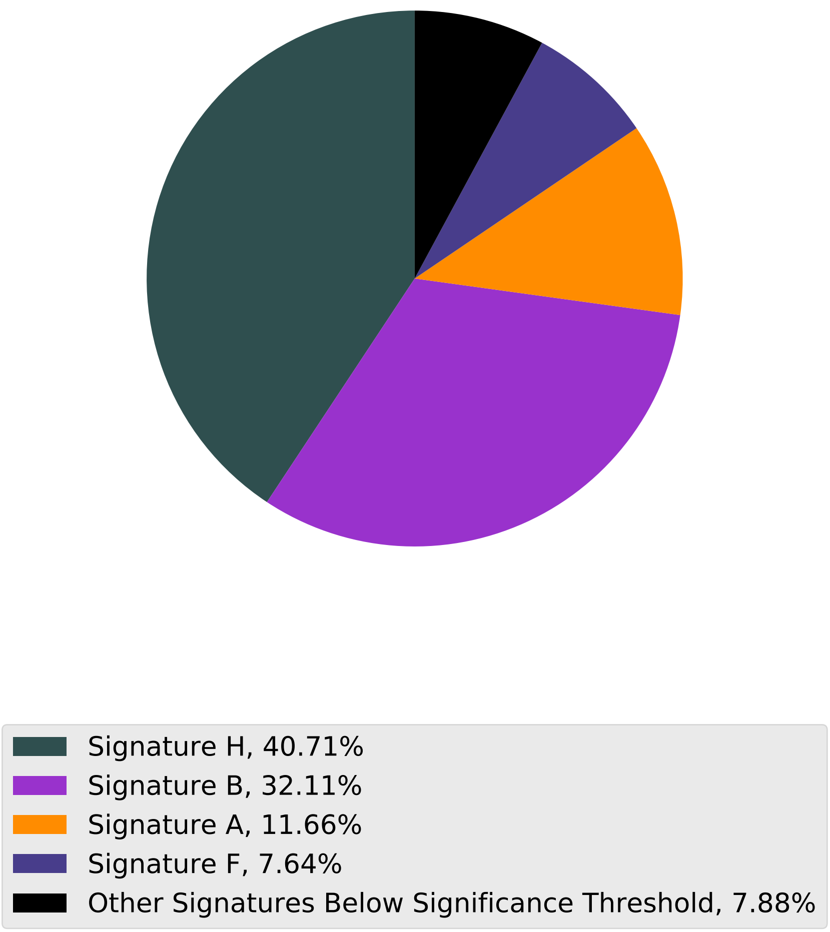

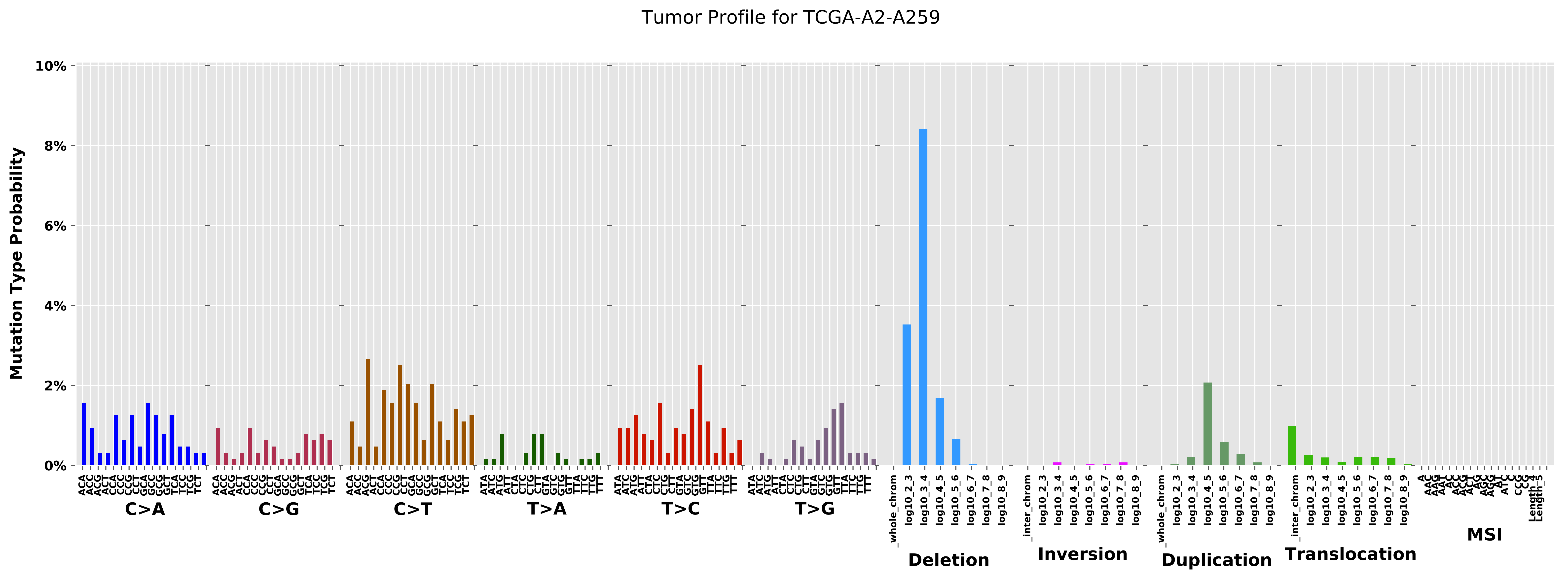
Tumor Profile for TCGA-A2-A259
Mutation Type Probability
10%
8%
6%
4%
2%
0%
C>A
C>G
C>T
T>A
T>C
T>G
Deletion
Inversion
Duplication
Translocation
MSI
_whole_chrom
_inter_chrom
log10_2_3
log10_3_4
log10_4_5
log10_5_6
log10_6_7
log10_7_8
log10_8_9

Cancer processes Weights for TCGA-BH-A0DG

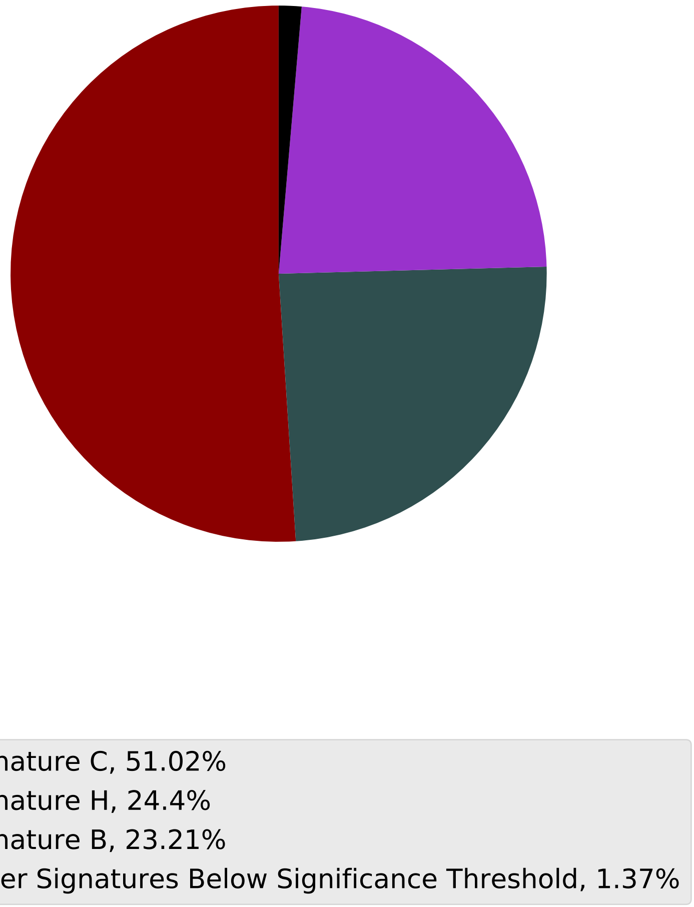

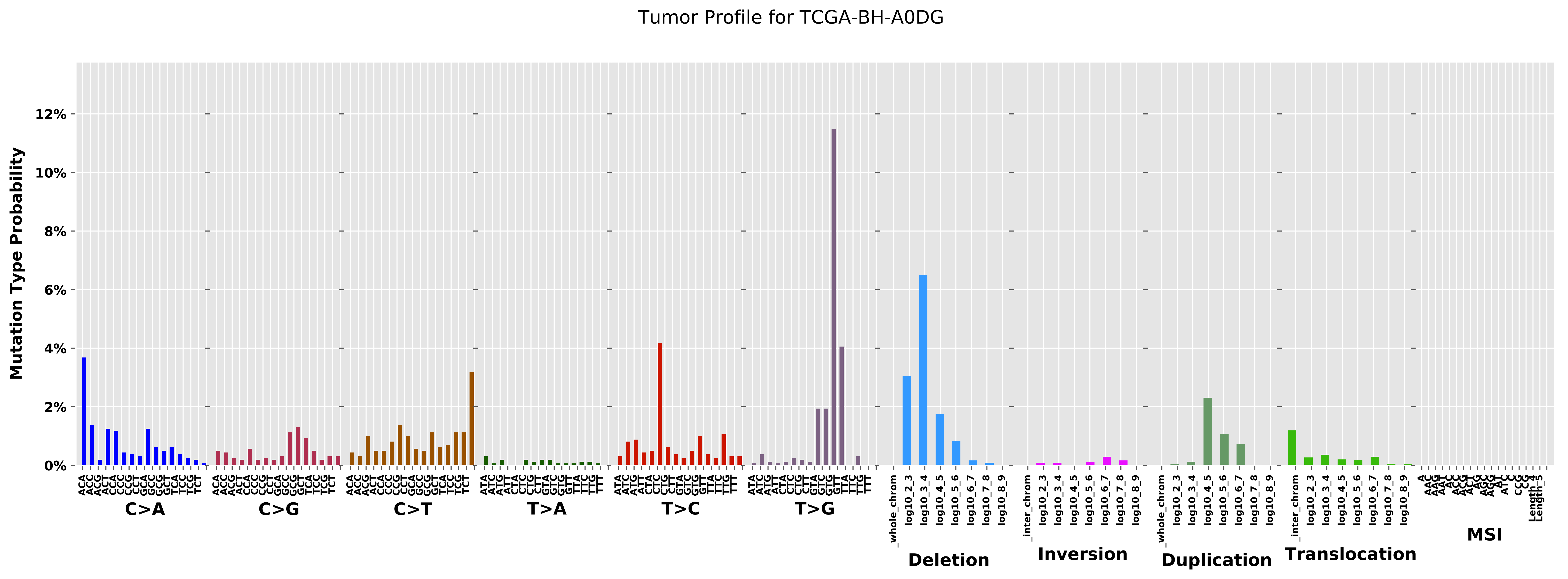

Tumor Profile for TCGA-BH-A0DG
Mutation Type Probability
0%
2%
4%
6%
8%
10%
12%
C>A
C>G
C>T
T>A
T>C
T>G
Deletion
Inversion
Duplication
Translocation
MSI
_whole_chrom
_inter_chrom
log10_2_3
log10_3_4
log10_4_5
log10_5_6
log10_6_7
log10_7_8
log10_8_9

Cancer processes Weights for TCGA-A2-A3KC

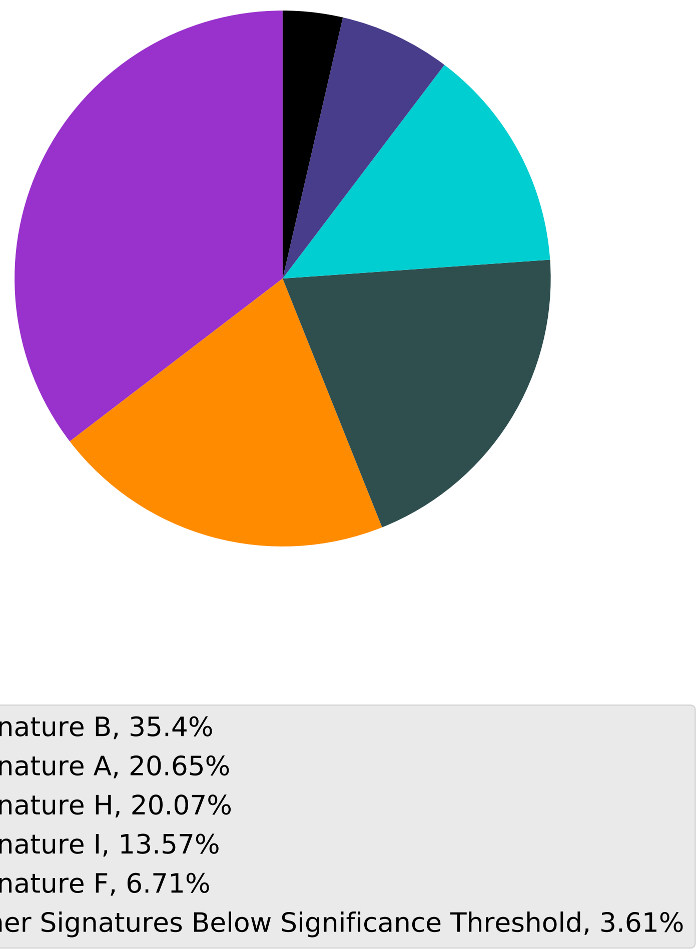

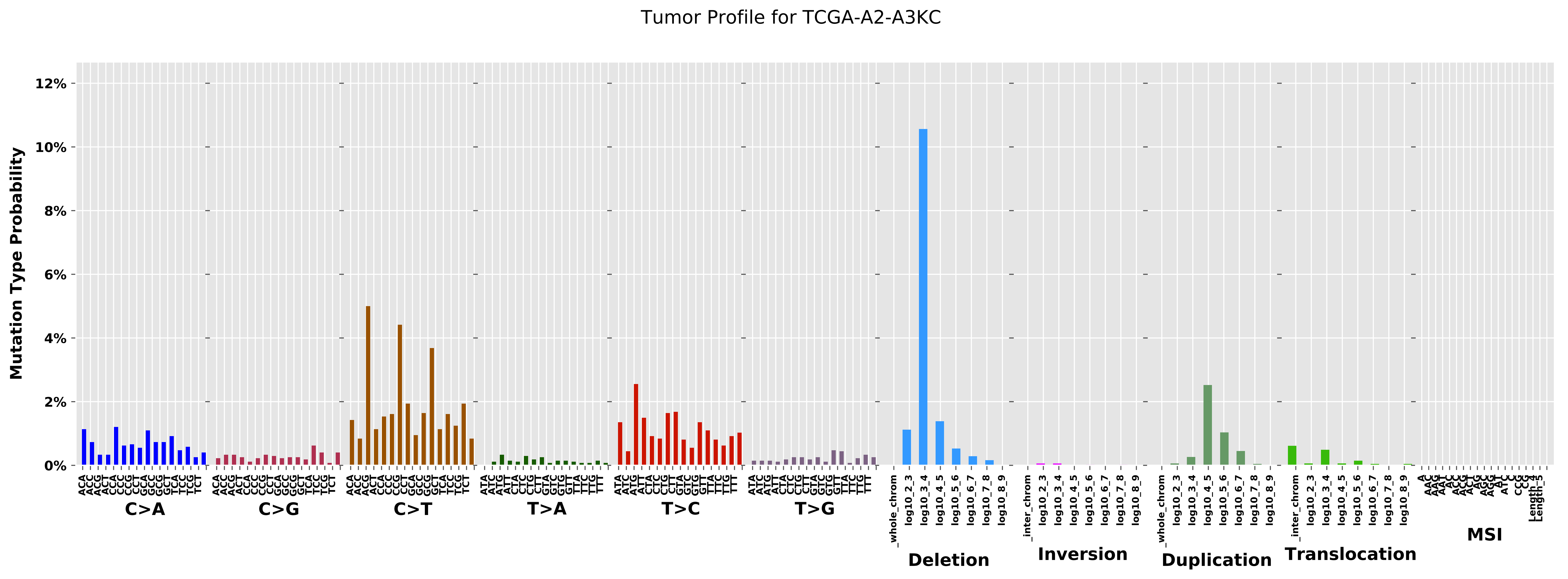

Tumor Profile for TCGA-A2-A3KC
Mutation Type Probability
0%
2%
4%
6%
8%
10%
12%
C>A
C>G
C>T
T>A
T>C
T>G
_whole_chrom
log10_2_3
log10_3_4
log10_4_5
log10_5_6
log10_6_7
log10_7_8
log10_8_9
Deletion
_inter_chrom
Inversion
Duplication
Translocation
MSI

Cancer processes Weights for TCGA-AA-3514

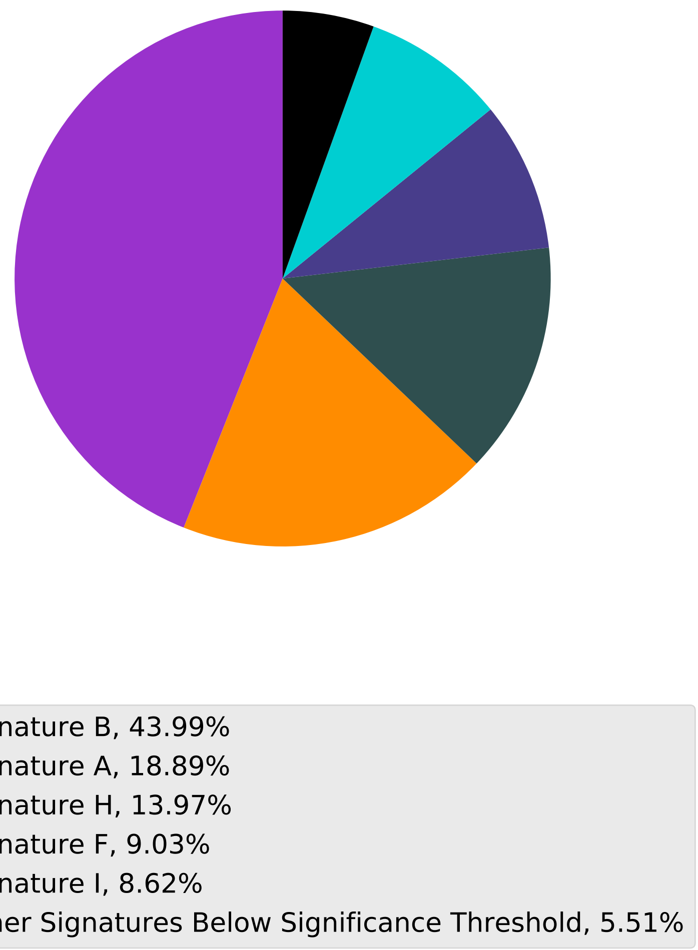

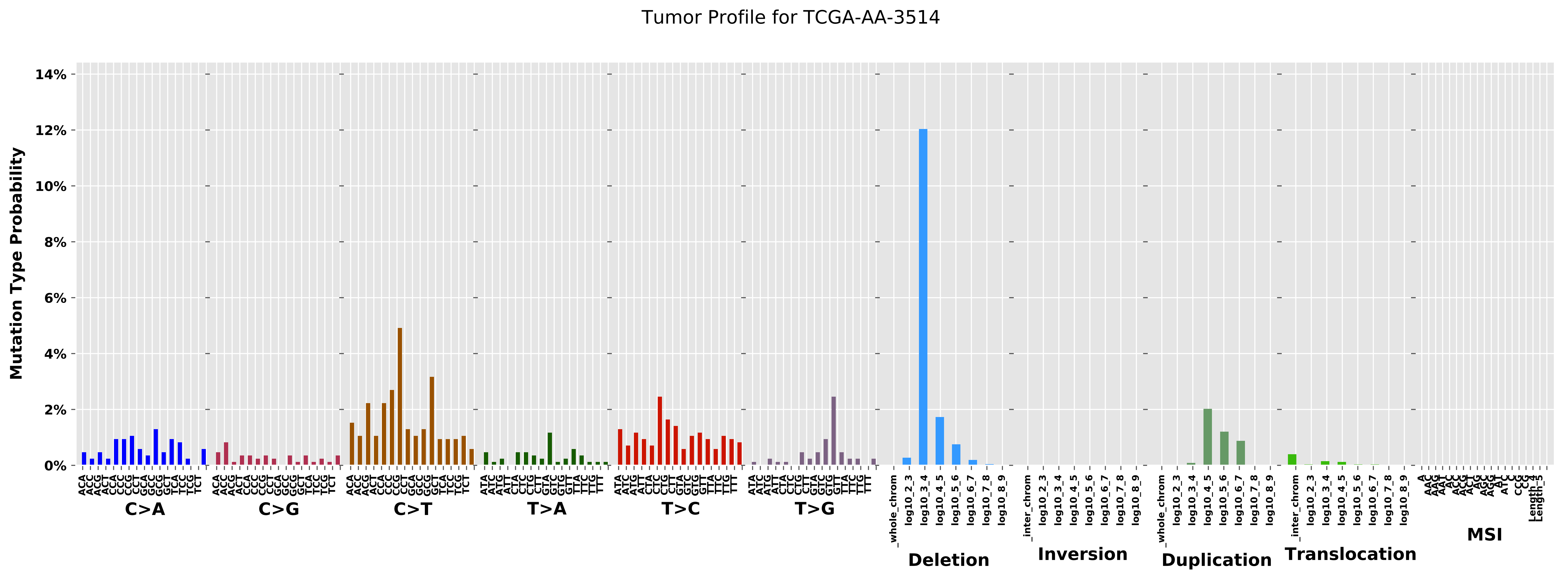
Tumor Profile for TCGA-AA-3514
Mutation Type Probability
0%
2%
4%
6%
8%
10%
12%
14%
C>A
C>G
C>T
T>A
T>C
T>G
Deletion
Inversion
Duplication
Translocation
MSI
_whole_chrom
_inter_chrom
log10_2_3
log10_3_4
log10_4_5
log10_5_6
log10_6_7
log10_7_8
log10_8_9

Cancer processes Weights for TCGA-AN-A0XR

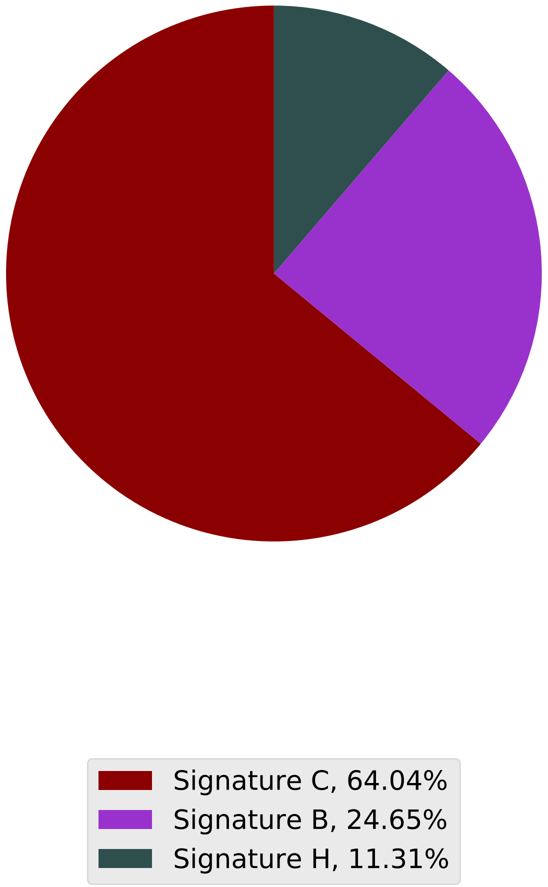

Tumor Profile for TCGA-AN-A0XR

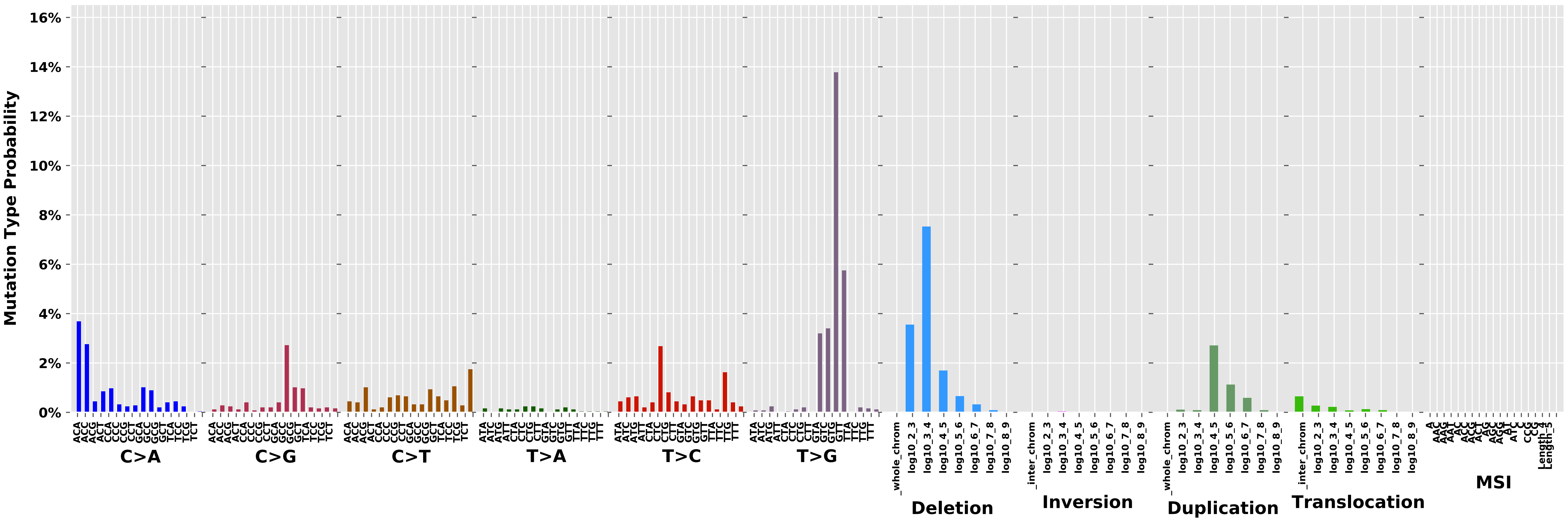

Cancer processes Weights for TCGA-BH-A0EA

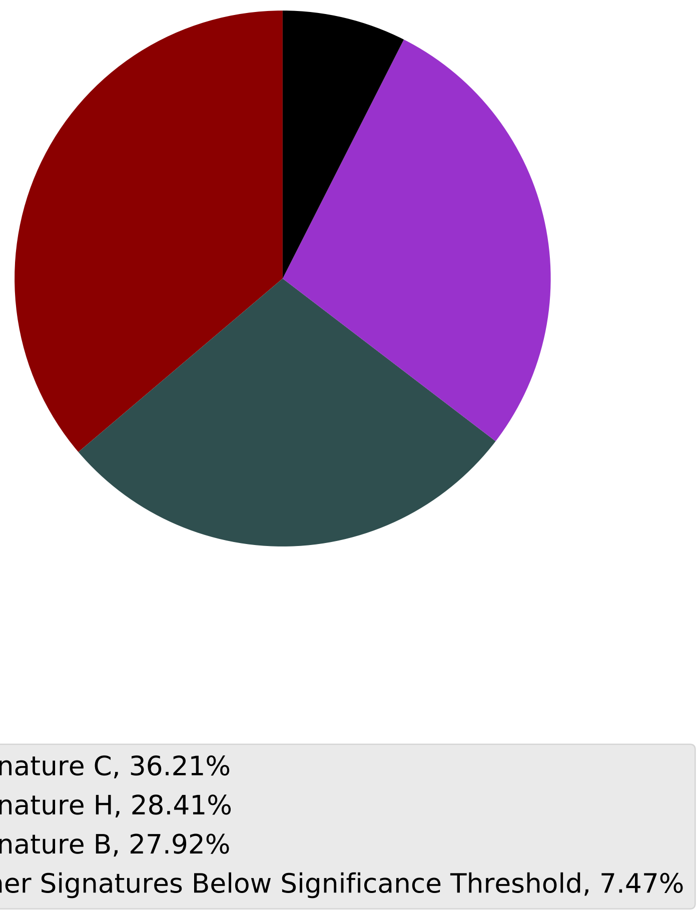

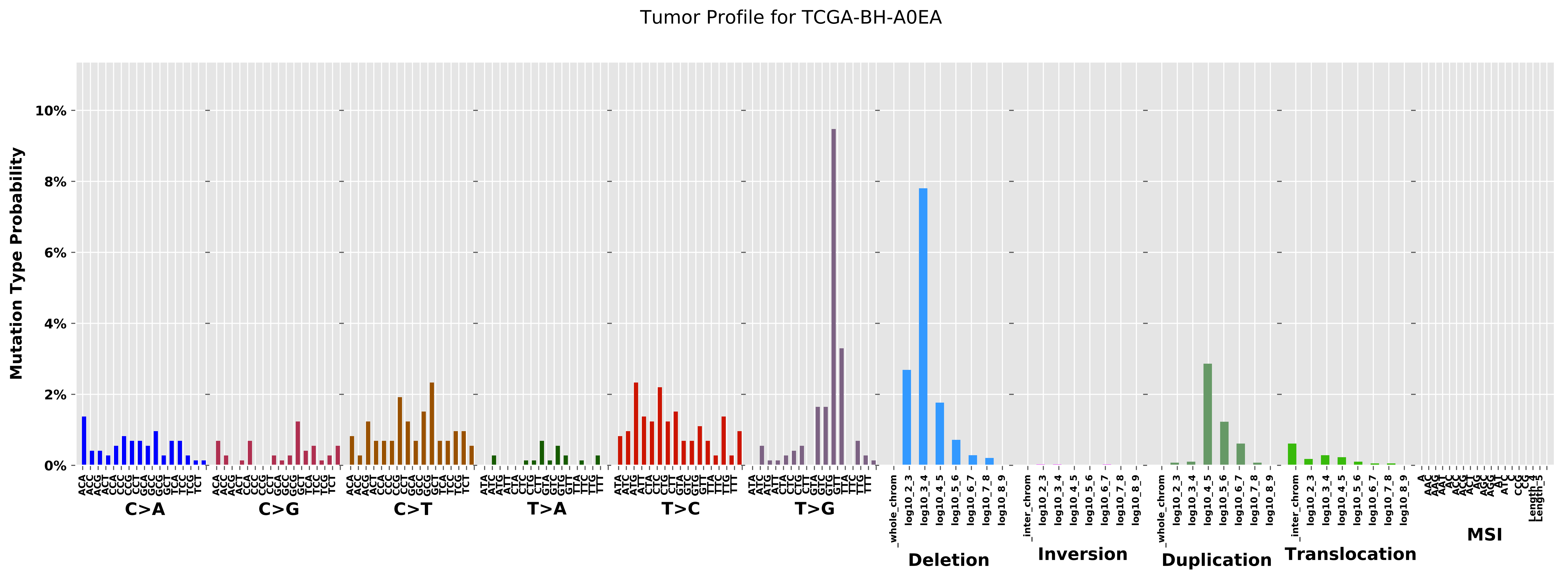
Tumor Profile for TCGA-BH-A0EA
Mutation Type Probability
10%
8%
6%
4%
2%
0%
C>A
C>G
C>T
T>A
T>C
T>G
Deletion
Inversion
Duplication
Translocation
MSI
_whole_chrom
_inter_chrom
log10_2_3
log10_3_4
log10_4_5
log10_5_6
log10_6_7
log10_7_8
log10_8_9

Cancer processes Weights for TCGA-AR-A256

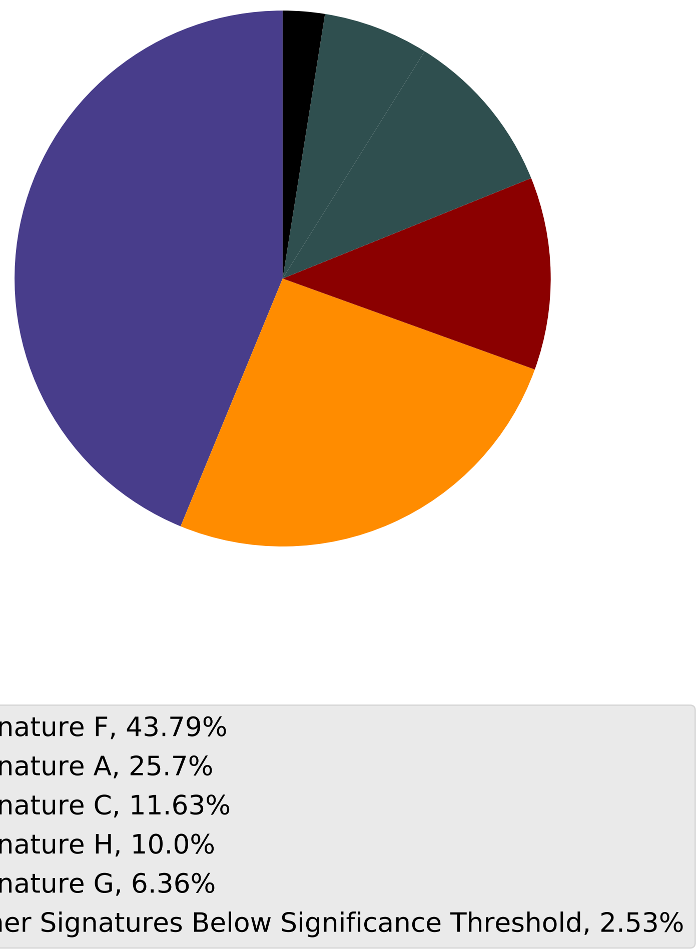

Tumor Profile for TCGA-AR-A256

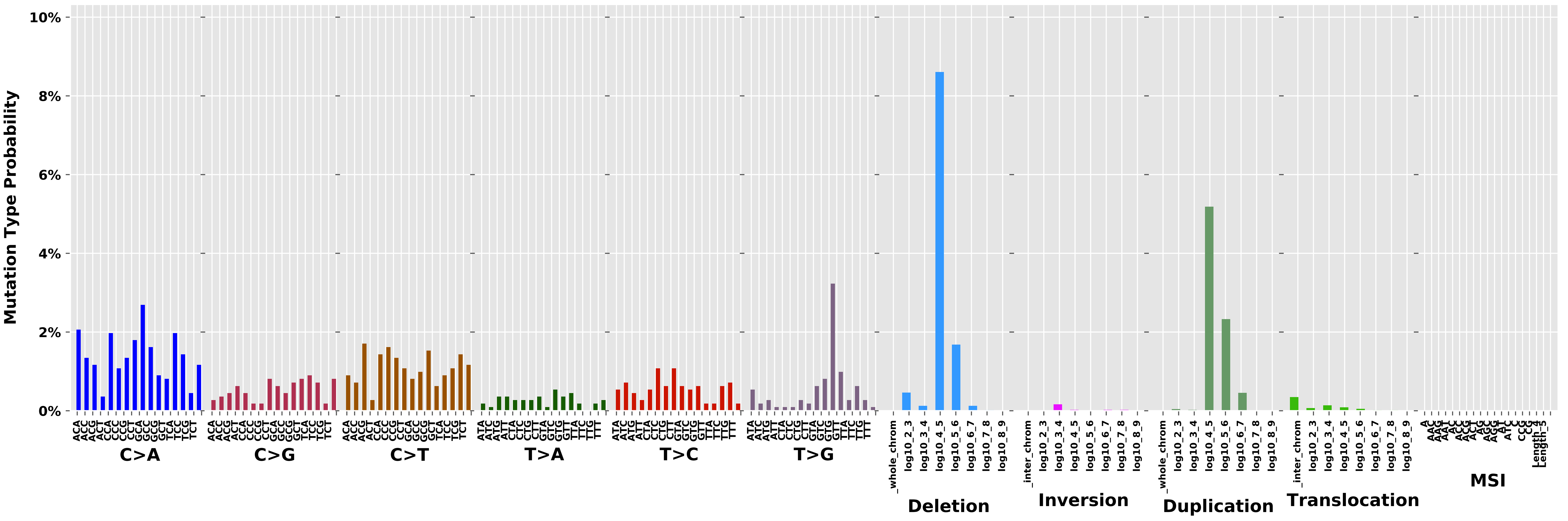

Cancer processes Weights for TCGA-A8-A09X

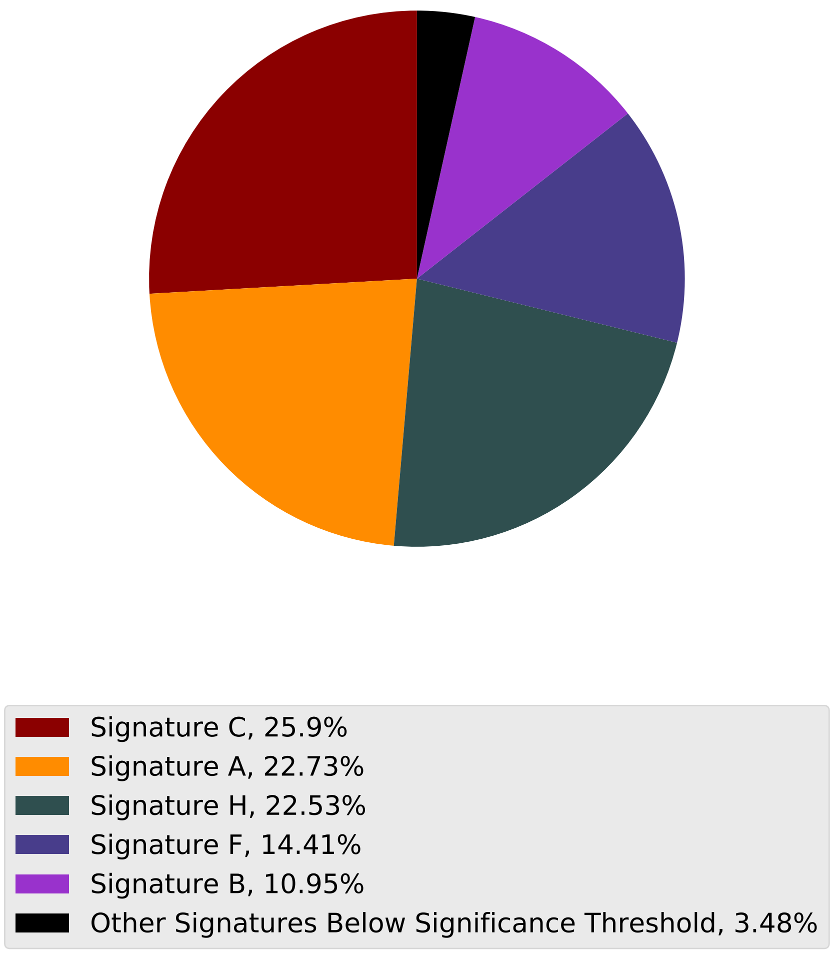

Tumor Profile for TCGA-A8-A09X

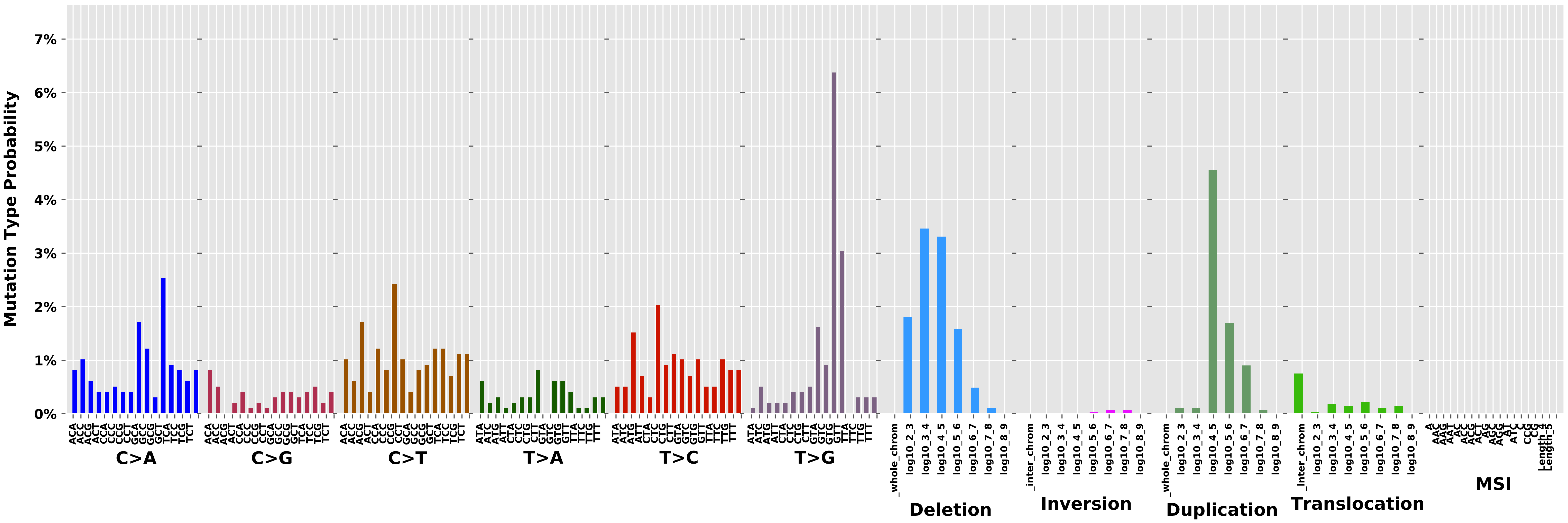

Cancer processes Weights for TCGA-AA-A01V

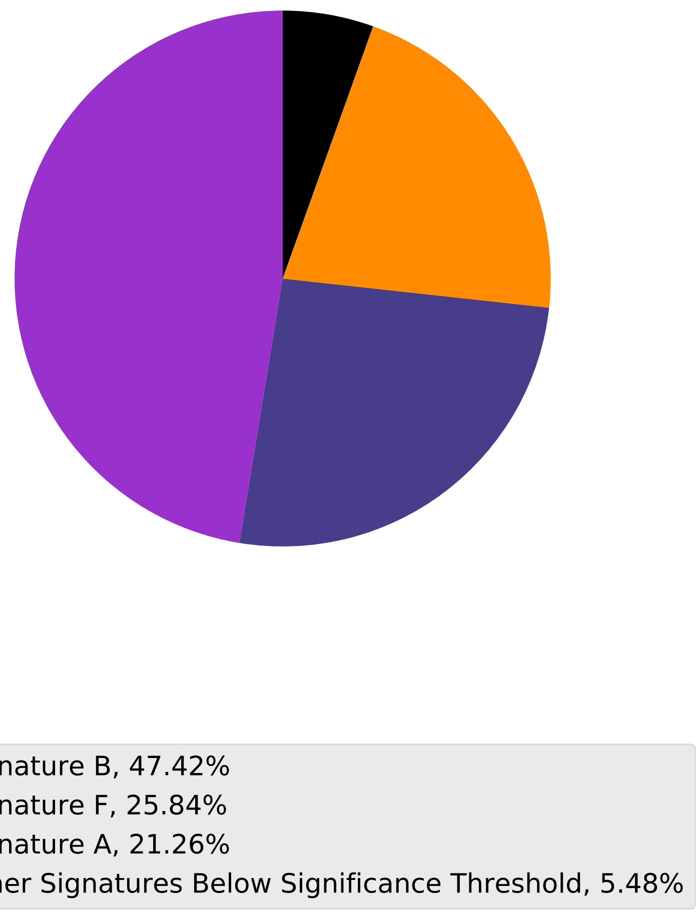

Tumor Profile for TCGA-AA-A01V

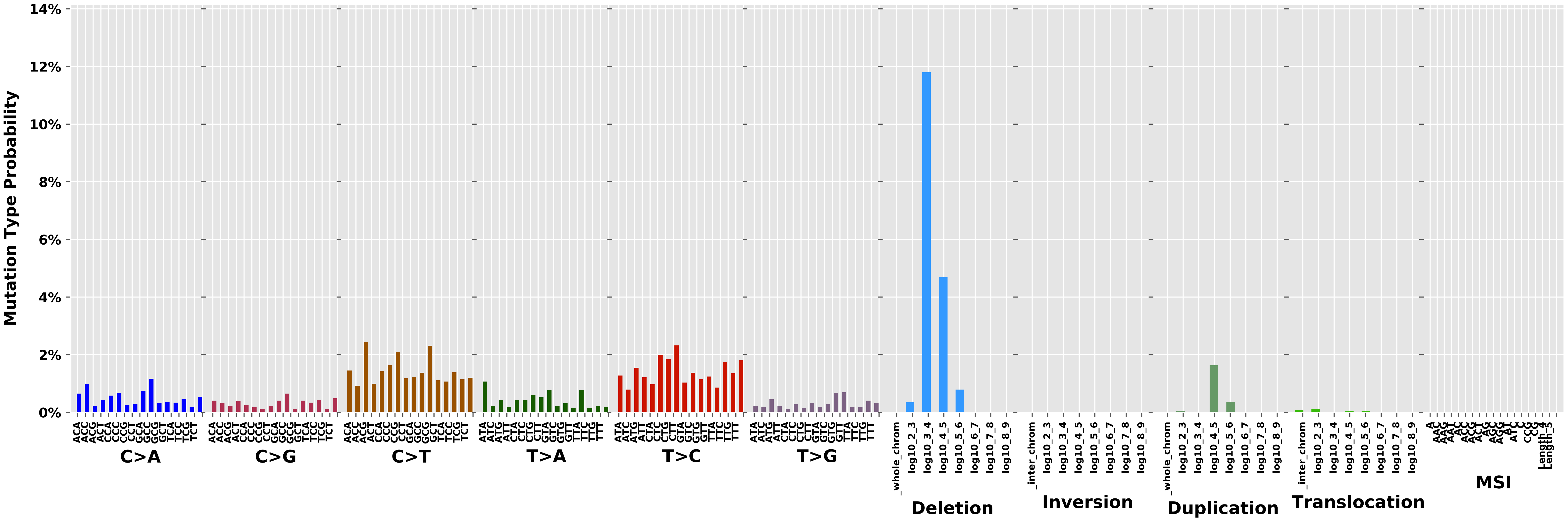

Cancer processes Weights for TCGA-A2-A04T

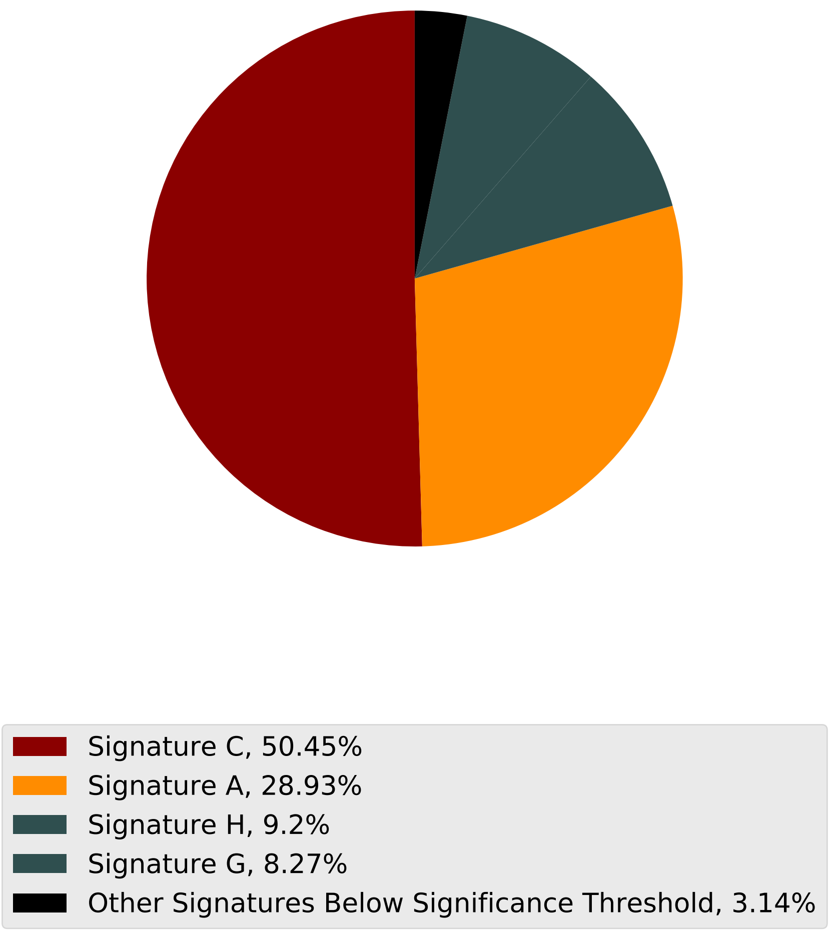

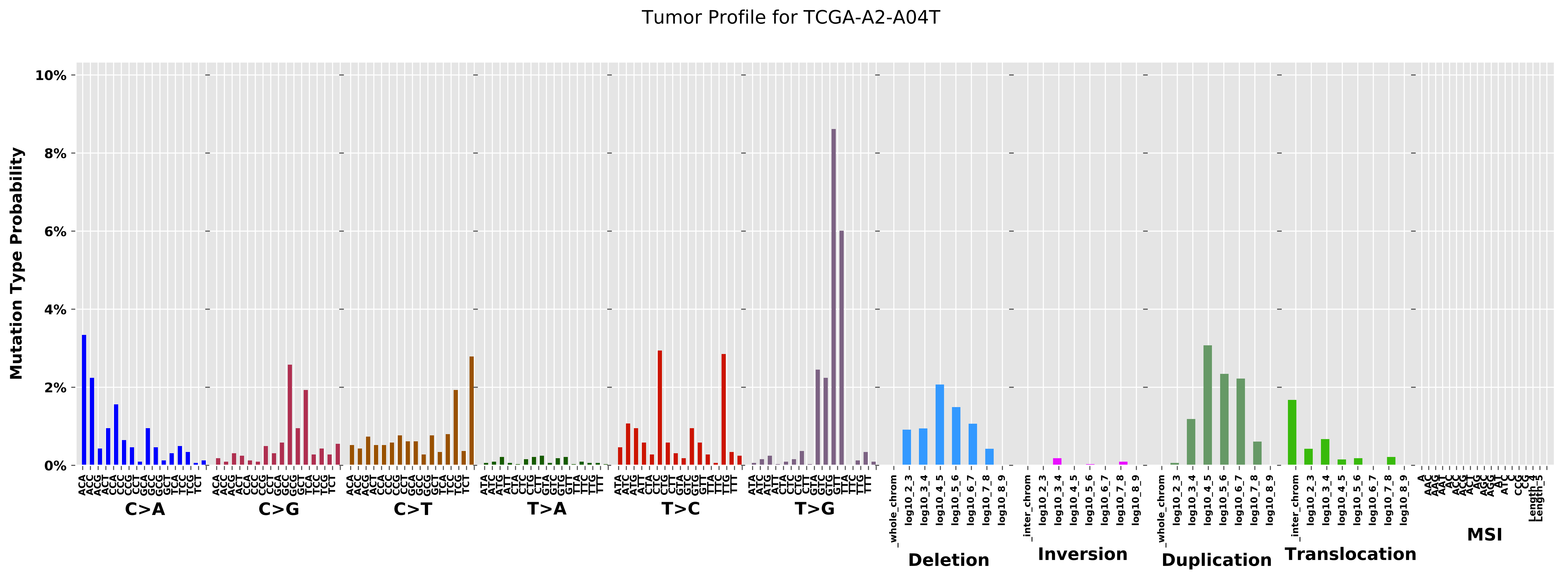
Tumor Profile for TCGA-A2-A04T
Mutation Type Probability
10%
8%
6%
4%
2%
0%
C>A
C>G
C>T
T>A
T>C
T>G
Deletion
Inversion
Duplication
Translocation
MSI
_whole_chrom
log10_2_3
log10_3_4
log10_4_5
log10_5_6
log10_6_7
log10_7_8
log10_8_9
_inter_chrom
Length_4
Length_5

Cancer processes Weights for TCGA-BH-A0H6

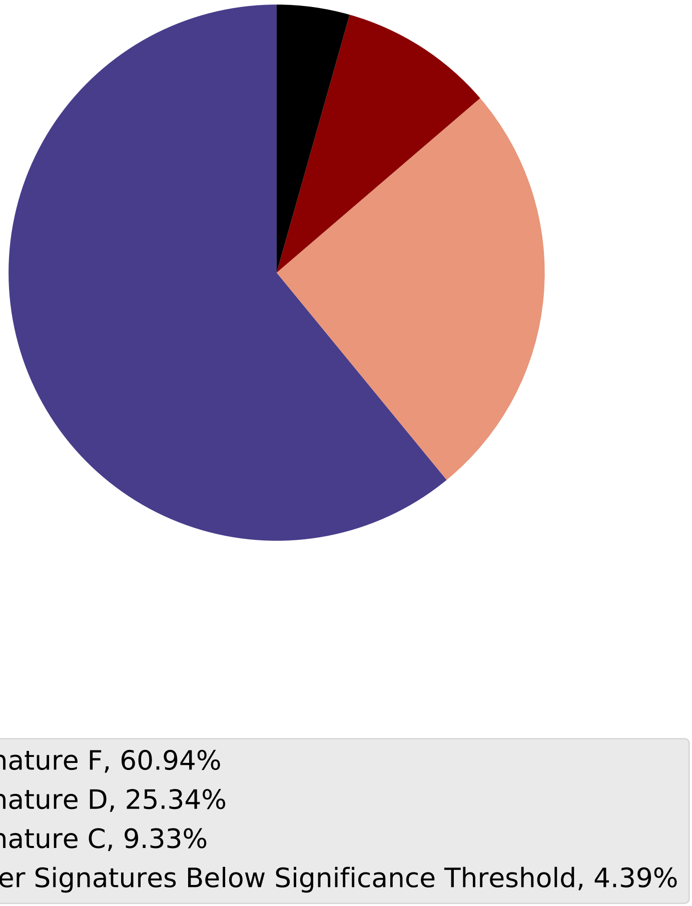

Tumor Profile for TCGA-BH-A0H6

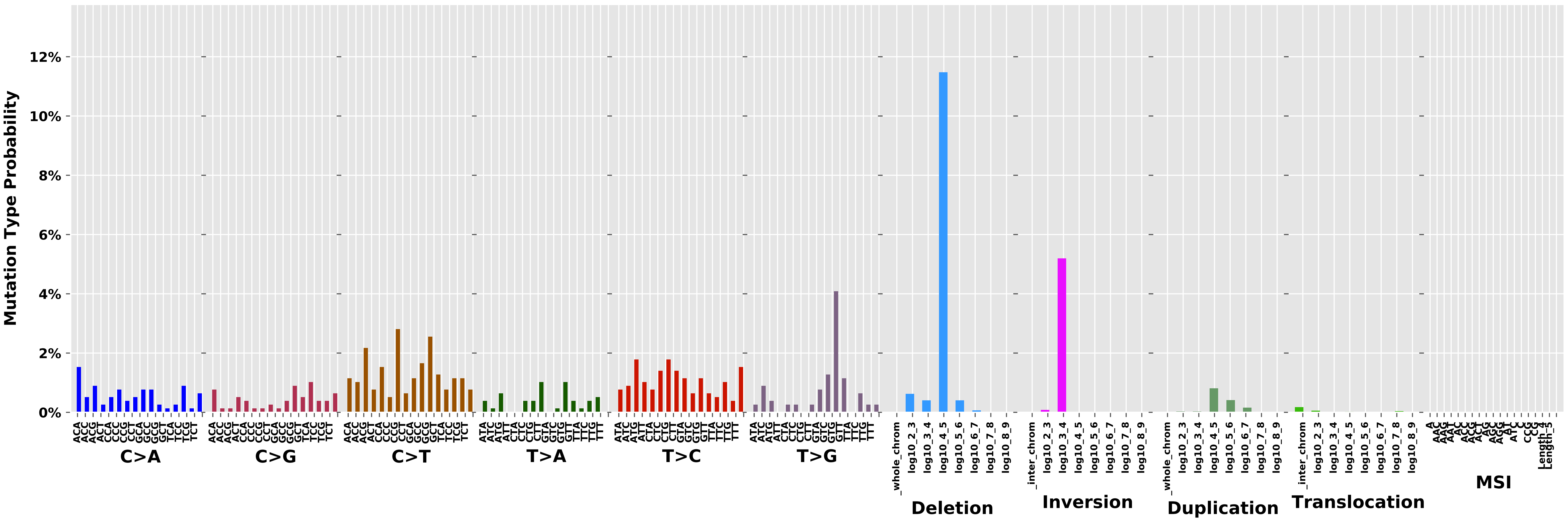

#### Cancer processes Weights for TCGA-D8-A27F

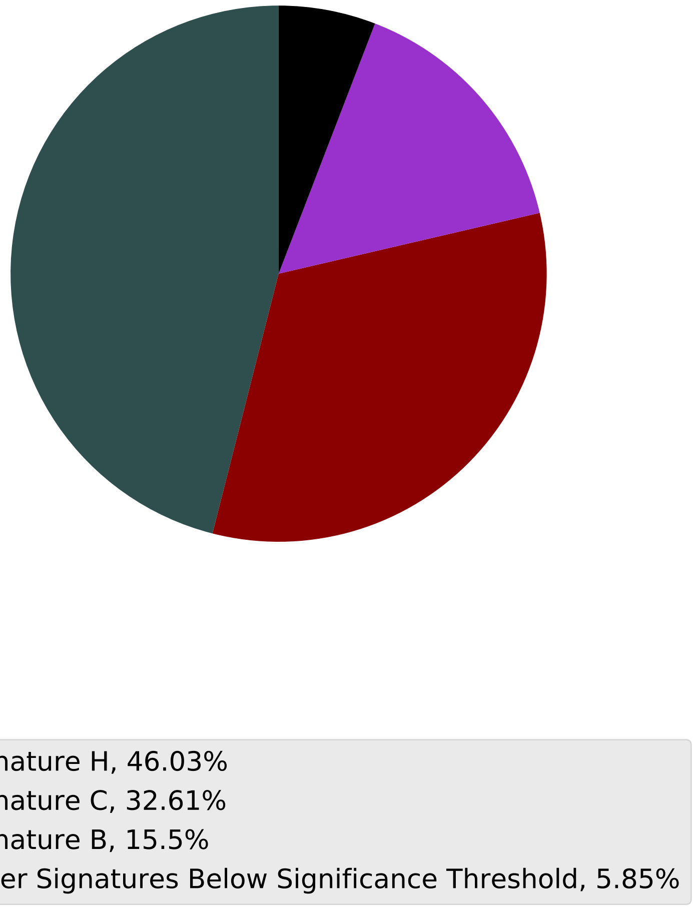

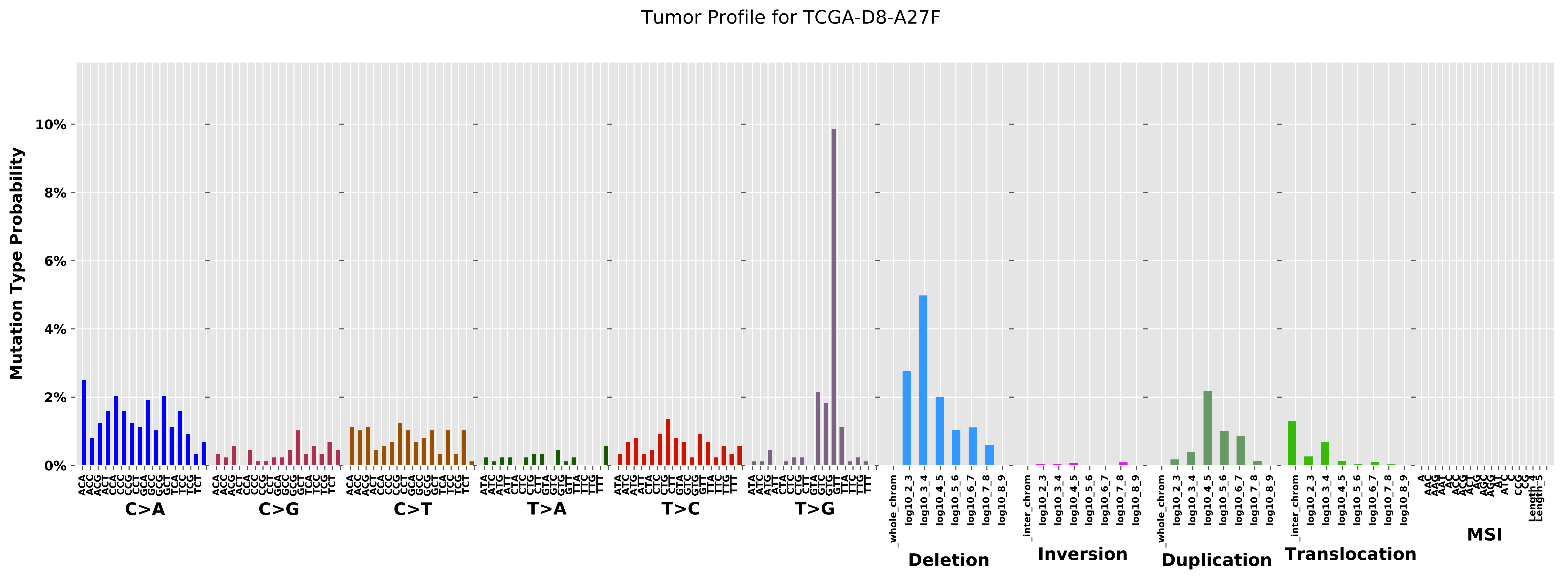

Tumor Profile for TCGA-D8-A27F
Mutation Type Probability
0%
2%
4%
6%
8%
10%
C>A
C>G
C>T
T>A
T>C
T>G
Deletion
Inversion
Duplication
Translocation
MSI
_whole_chrom
_inter_chrom
log10_2_3
log10_3_4
log10_4_5
log10_5_6
log10_6_7
log10_7_8
log10_8_9

Cancer processes Weights for TCGA-AG-3727

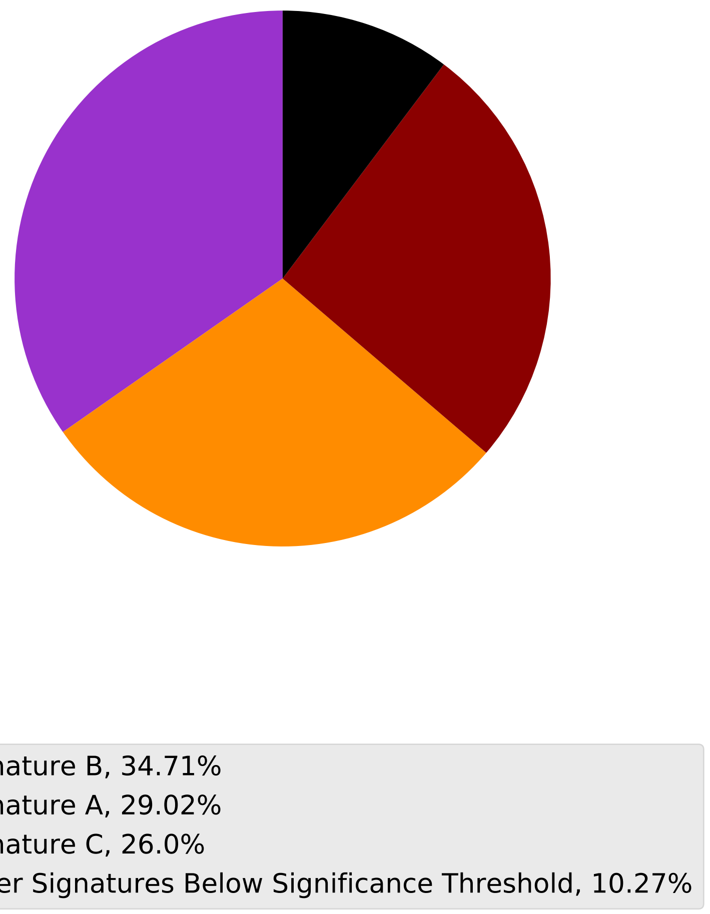

Tumor Profile for TCGA-AG-3727

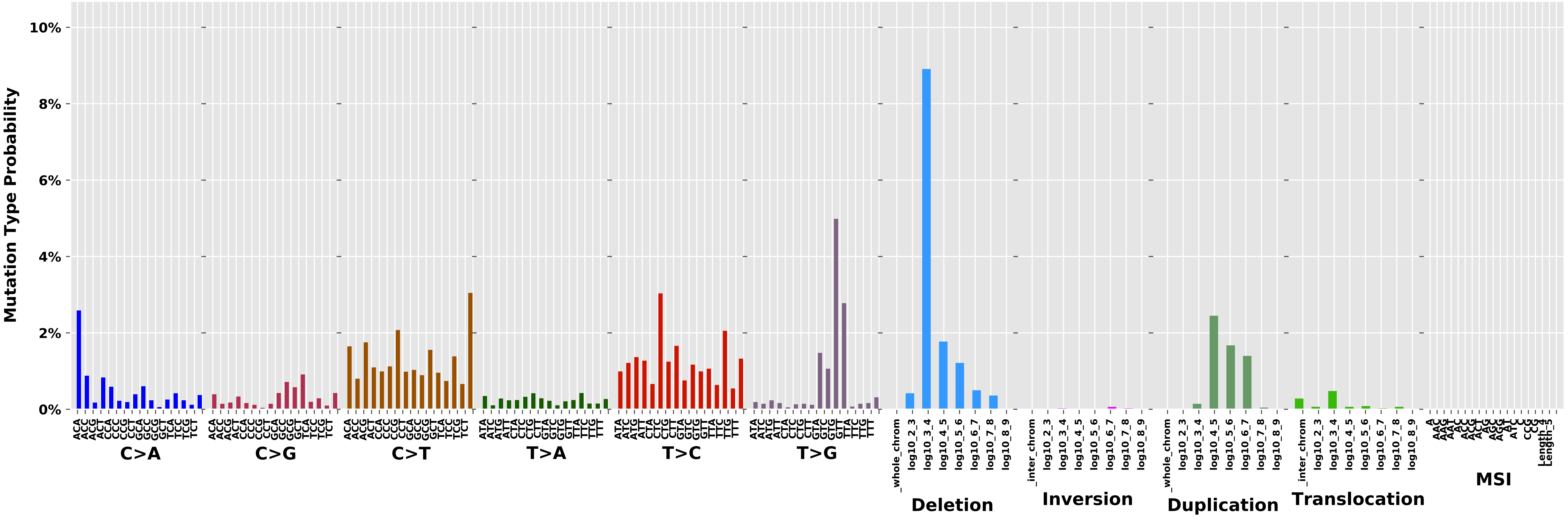

Cancer processes Weights for TCGA-A8-A09I

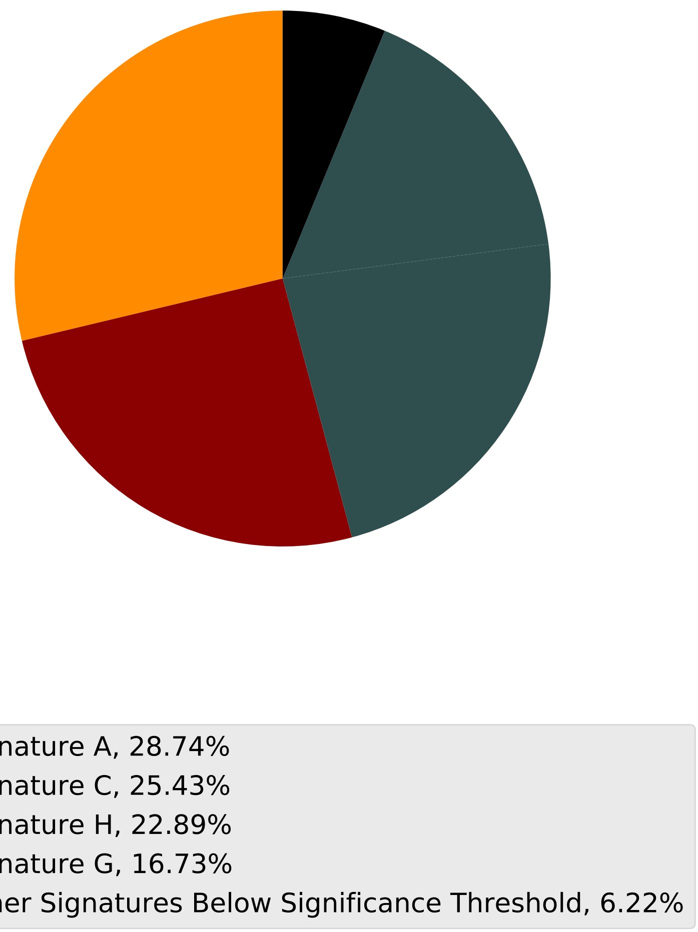

Tumor Profile for TCGA-A8-A09I

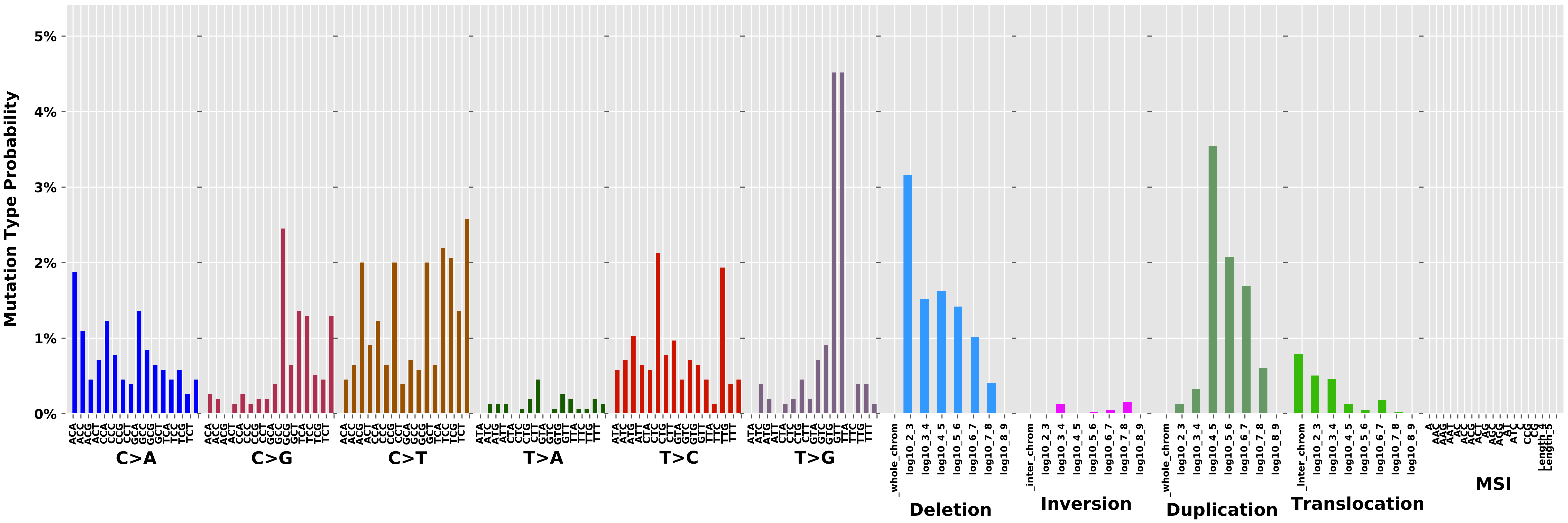

Cancer processes Weights for TCGA-AA-3664

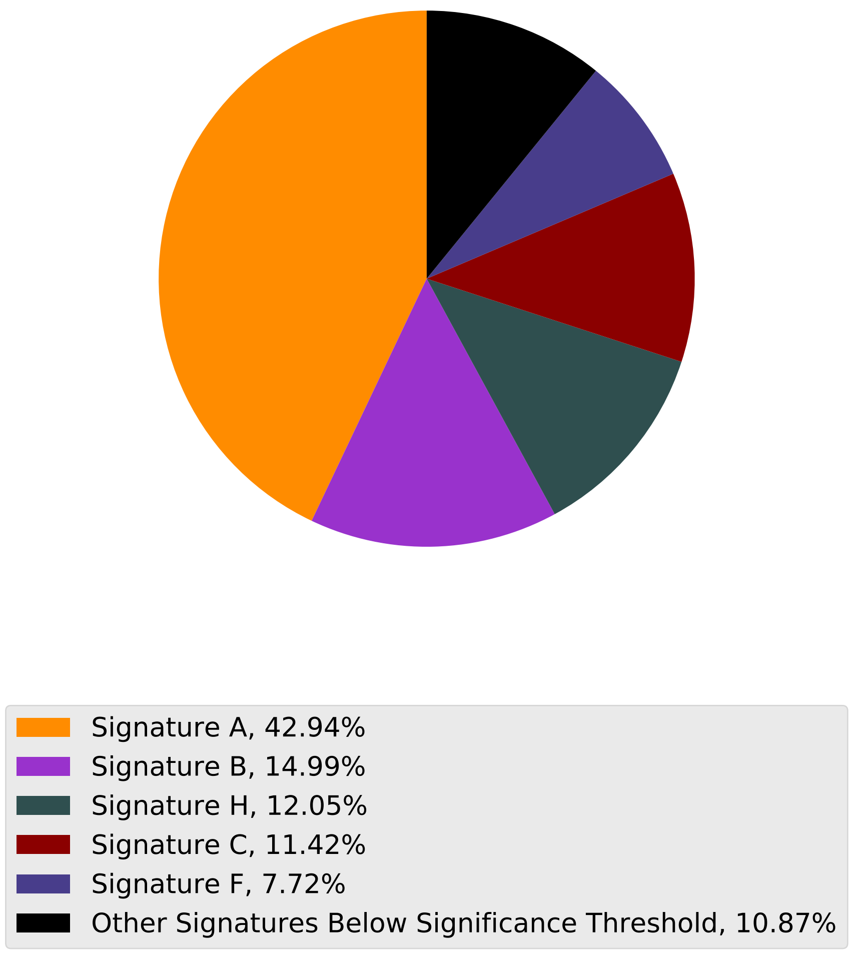

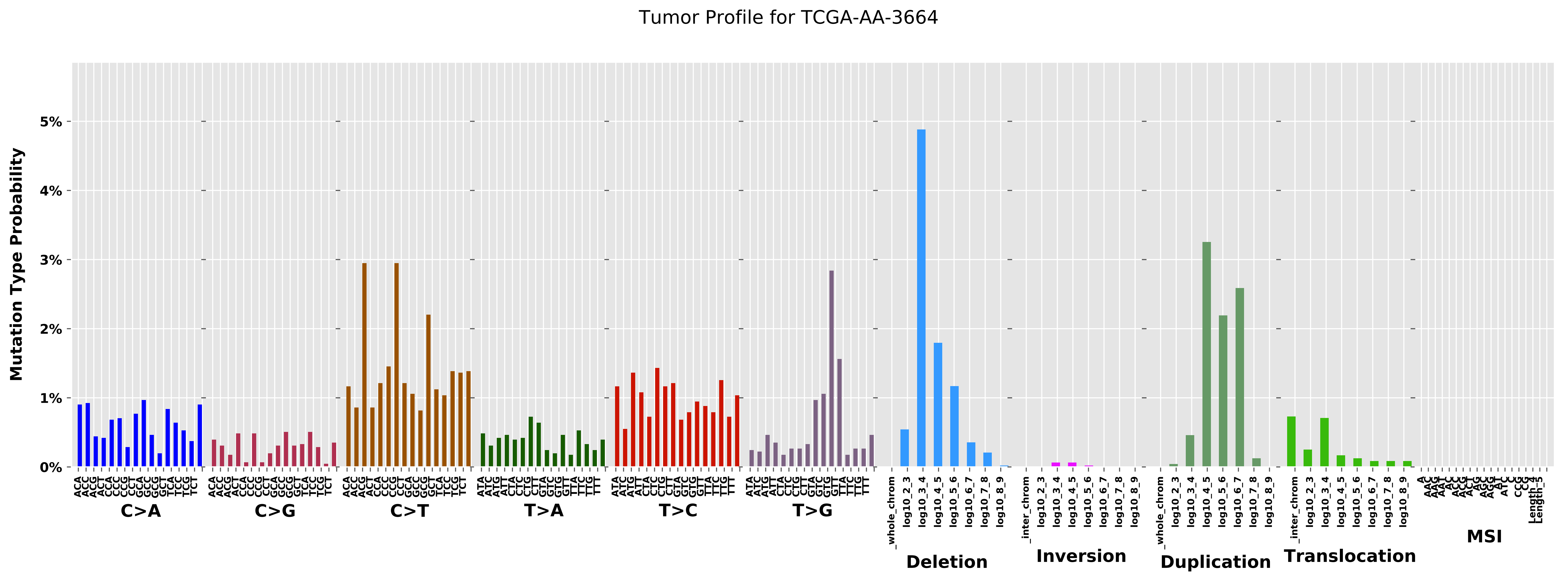
Tumor Profile for TCGA-AA-3664
Mutation Type Probability
0%
1%
2%
3%
4%
5%
C>A
C>G
C>T
T>A
T>C
T>G
Deletion
Inversion
Duplication
Translocation
MSI

Cancer processes Weights for TCGA-E2-A1LL

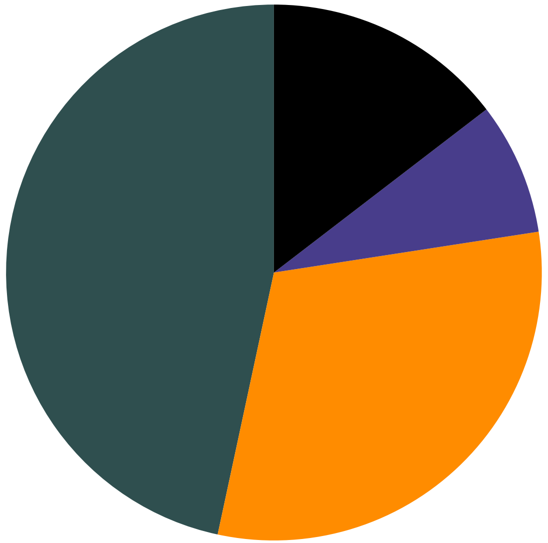

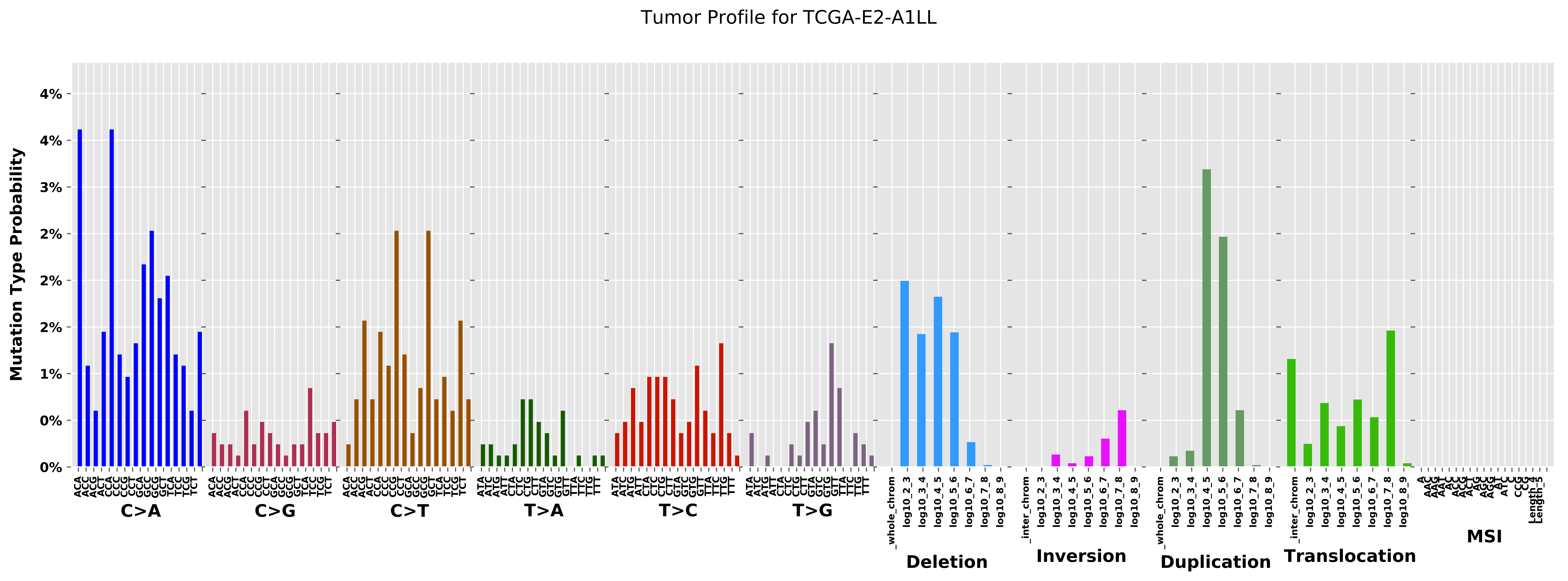
Tumor Profile for TCGA-E2-A1LL
Mutation Type Probability
0%
1%
2%
3%
4%
C>A
C>G
C>T
T>A
T>C
T>G
Deletion
Inversion
Duplication
Translocation
MSI

Cancer processes Weights for TCGA-A1-A0SM

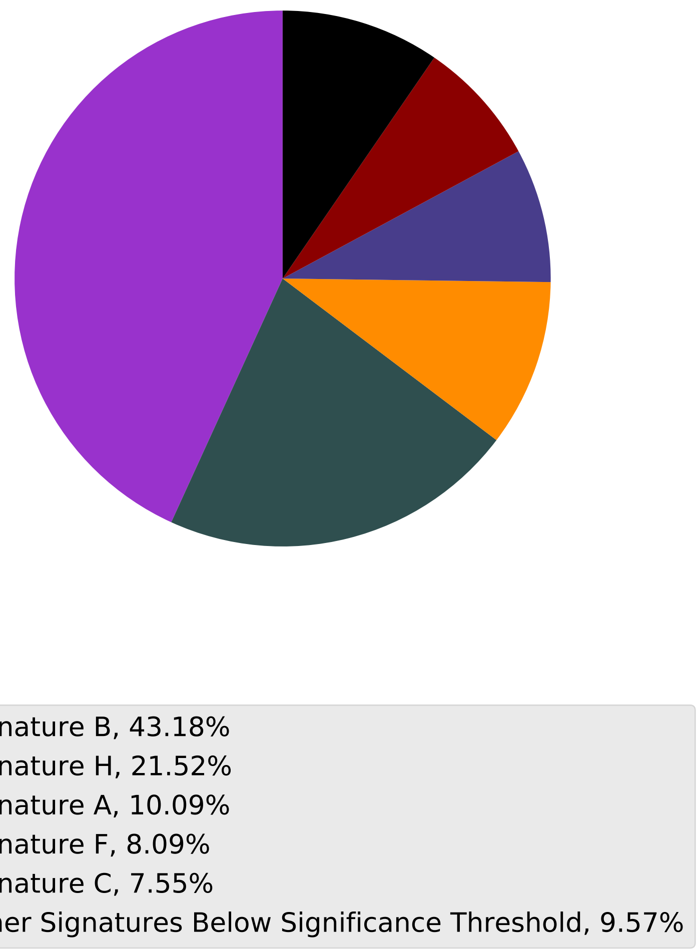

Tumor Profile for TCGA-A1-A0SM

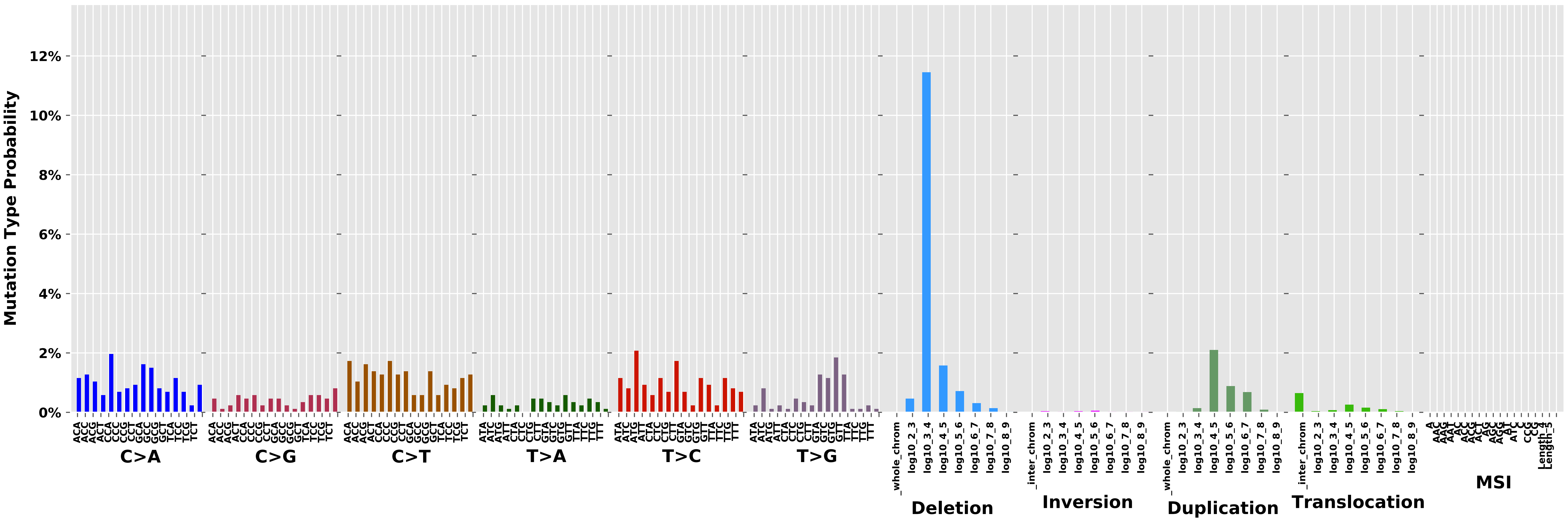

Cancer processes Weights for TCGA-C8-A130

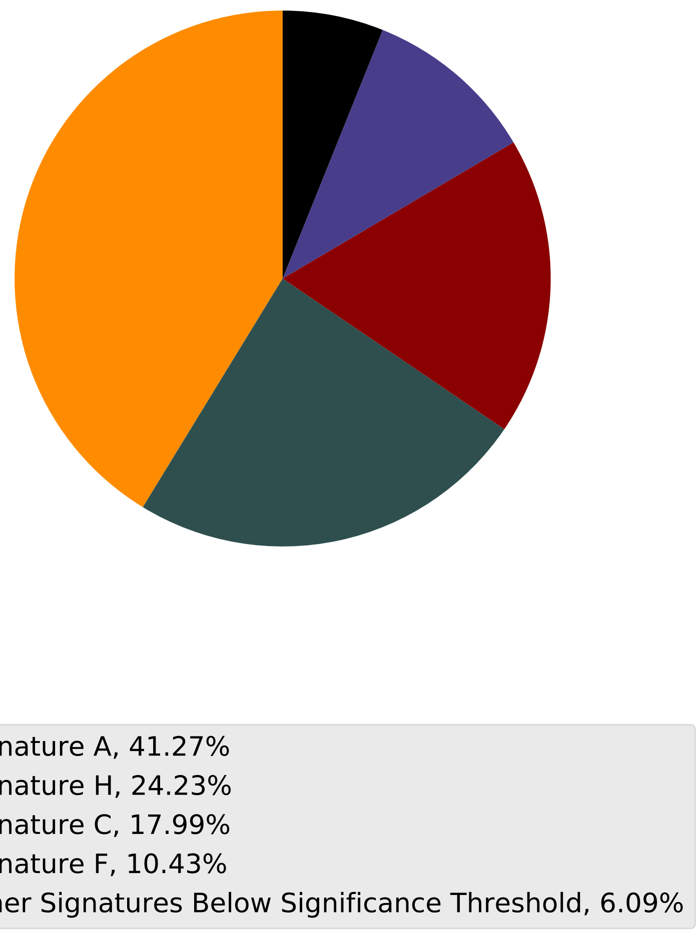

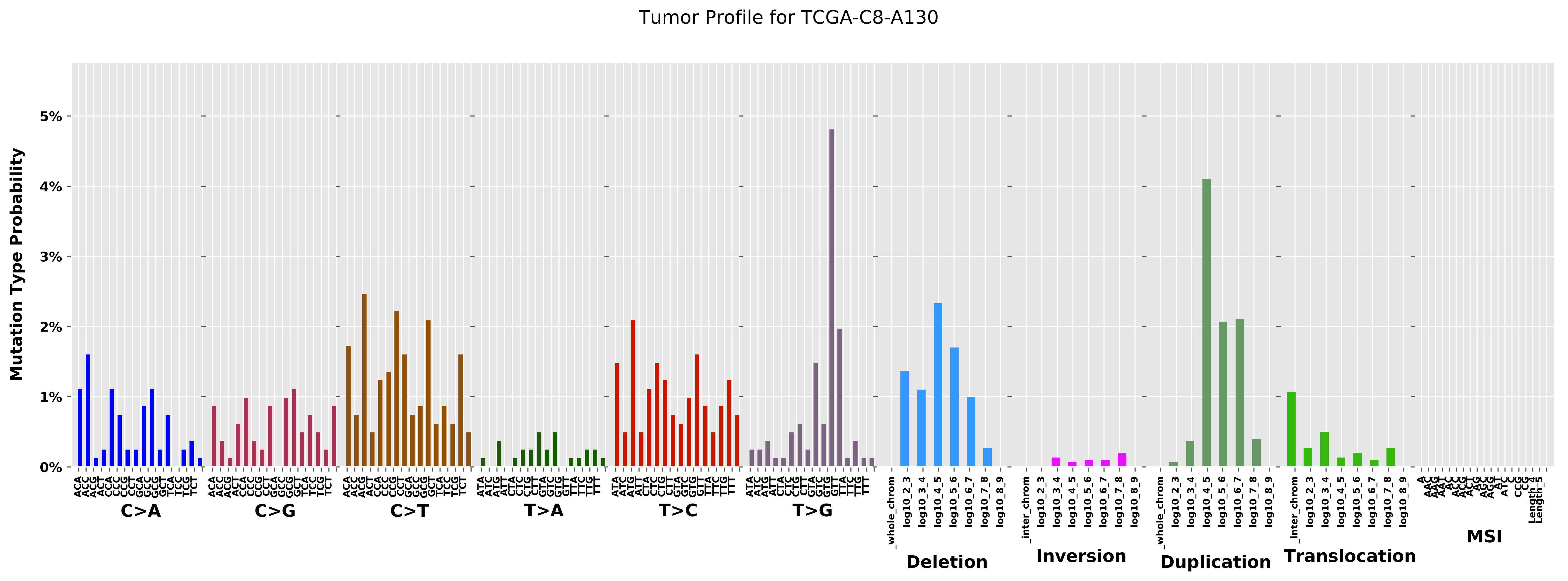

Tumor Profile for TCGA-C8-A130
Mutation Type Probability
0%
1%
2%
3%
4%
5%
C>A
C>G
C>T
T>A
T>C
T>G
Deletion
Inversion
Duplication
Translocation
MSI
_whole_chrom
_inter_chrom
log10_2_3
log10_3_4
log10_4_5
log10_5_6
log10_6_7
log10_7_8
log10_8_9

Cancer processes Weights for TCGA-BH-A0H0

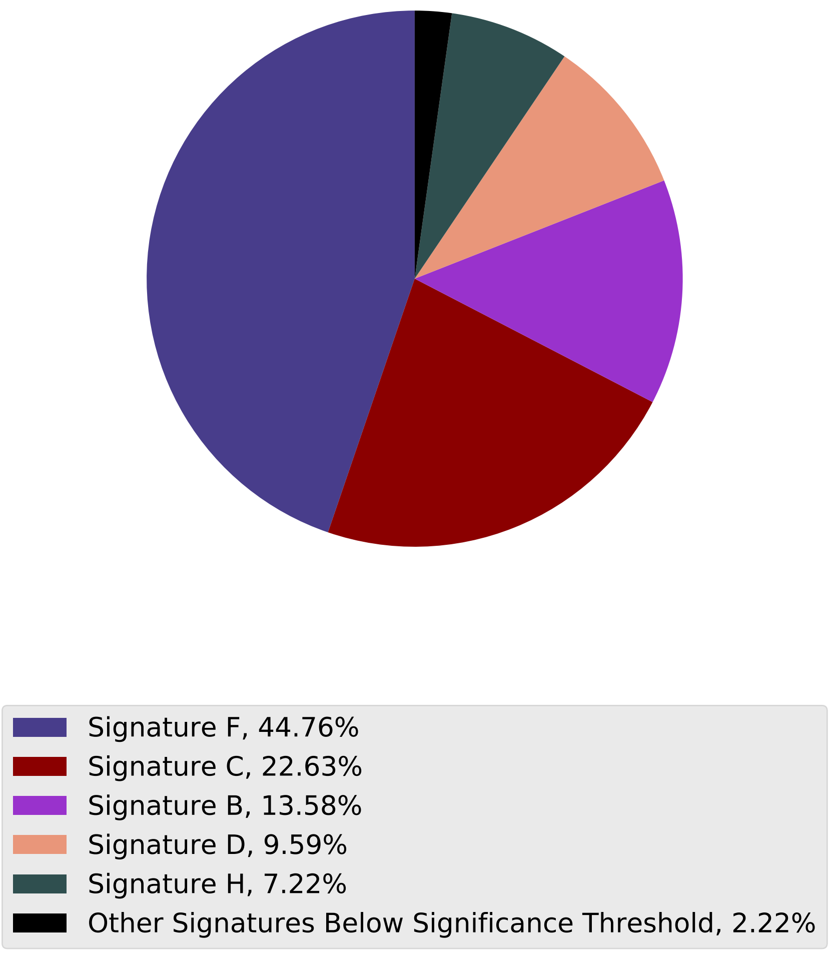

Tumor Profile for TCGA-BH-A0H0

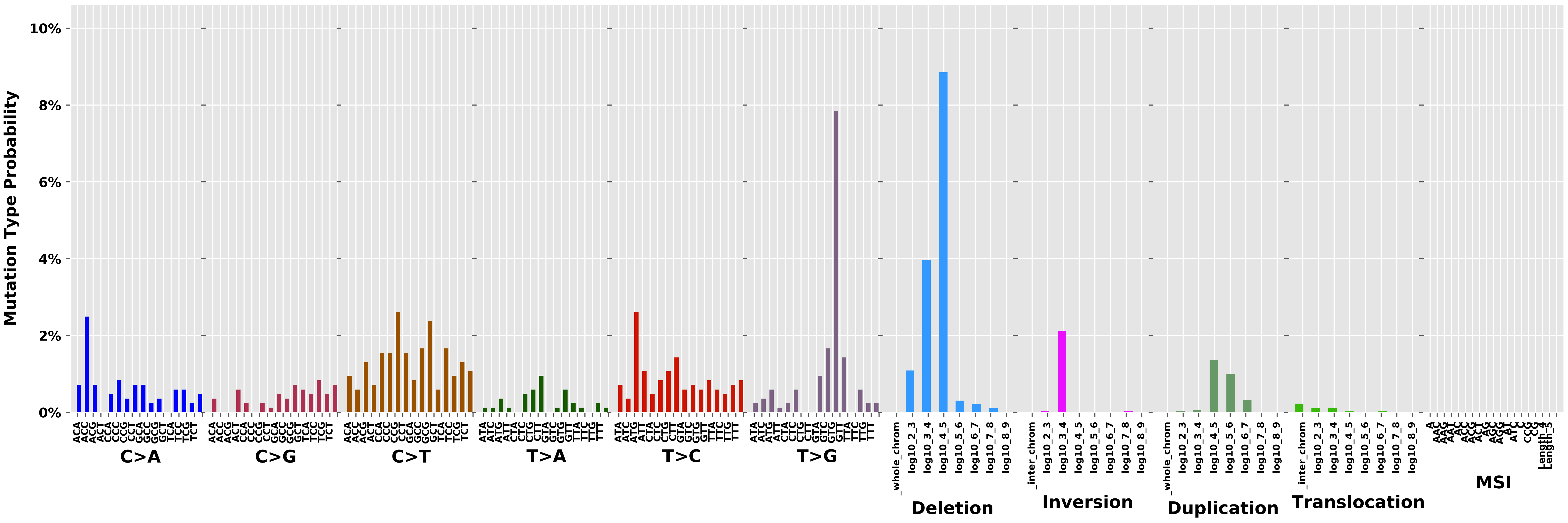

### Cancer processes Weights for TCGA-EW-A1J5

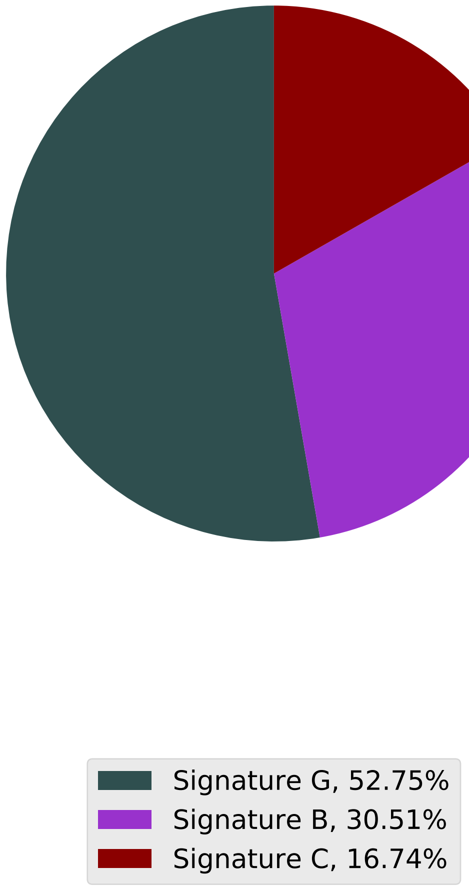

Tumor Profile for TCGA-EW-A1J5

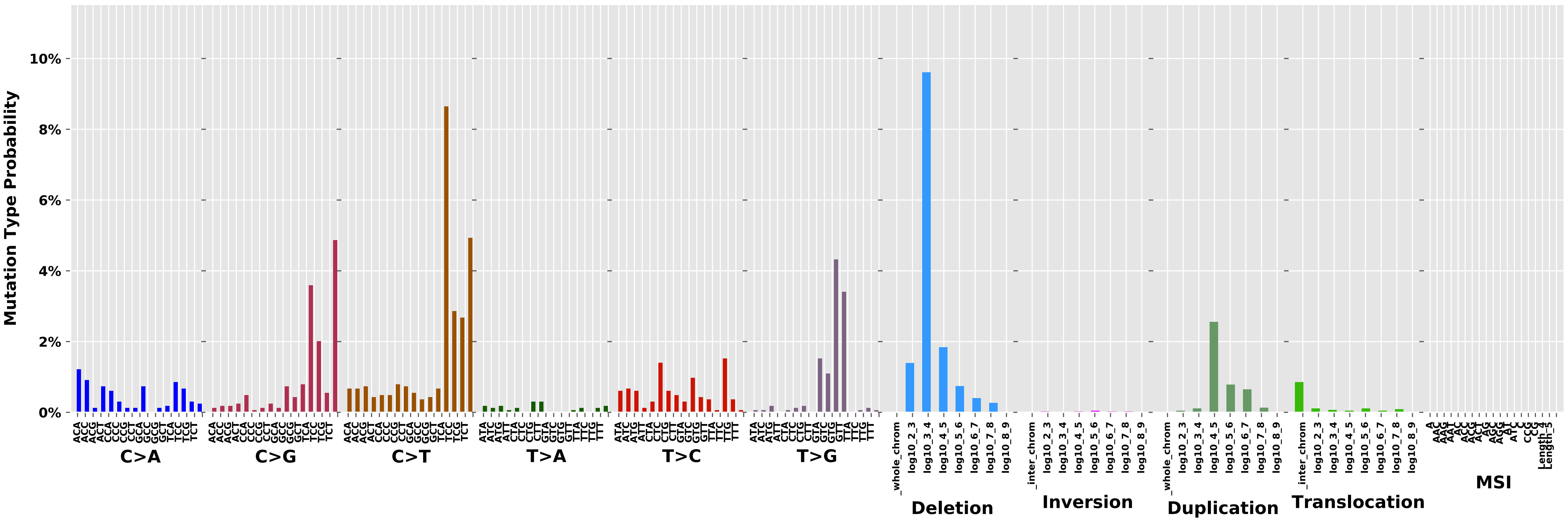

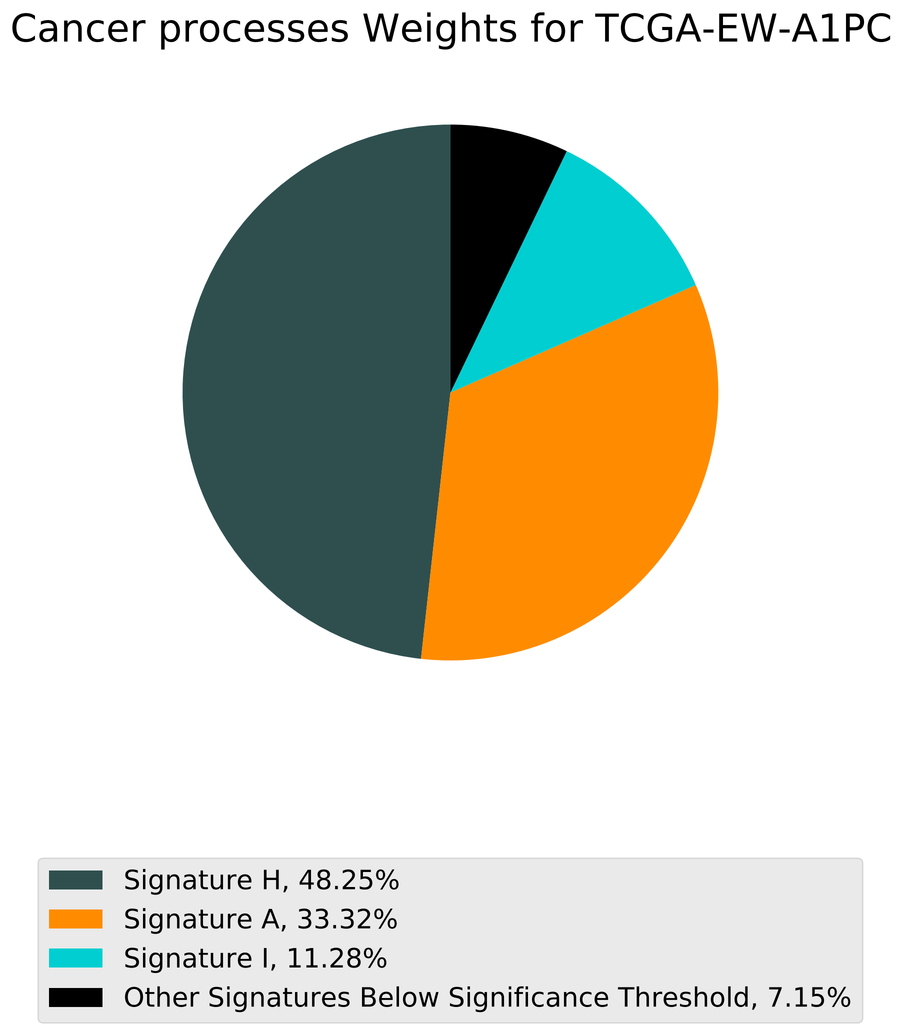
Cancer processes Weights for TCGA-EW-A1PC
Signature H, 48.25%
Signature A, 33.32%
Signature I, 11.28%
Other Signatures Below Significance Threshold, 7.15%

Tumor Profile for TCGA-EW-A1PC

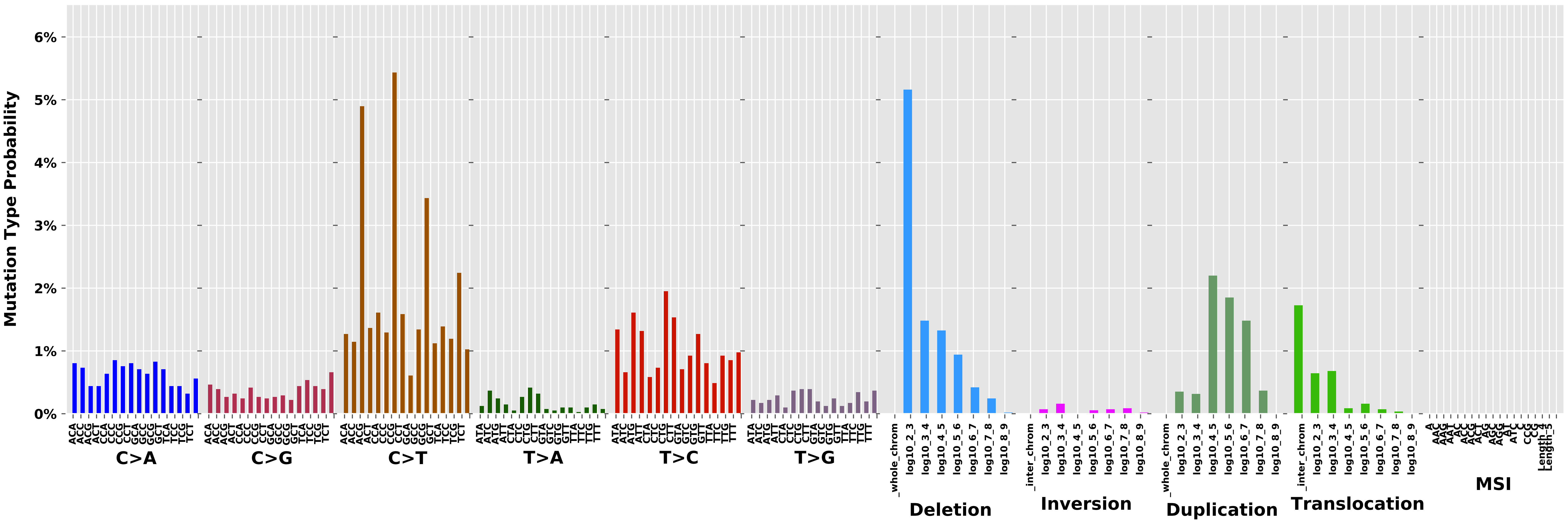

### Cancer processes Weights for TCGA-BH-A0E0

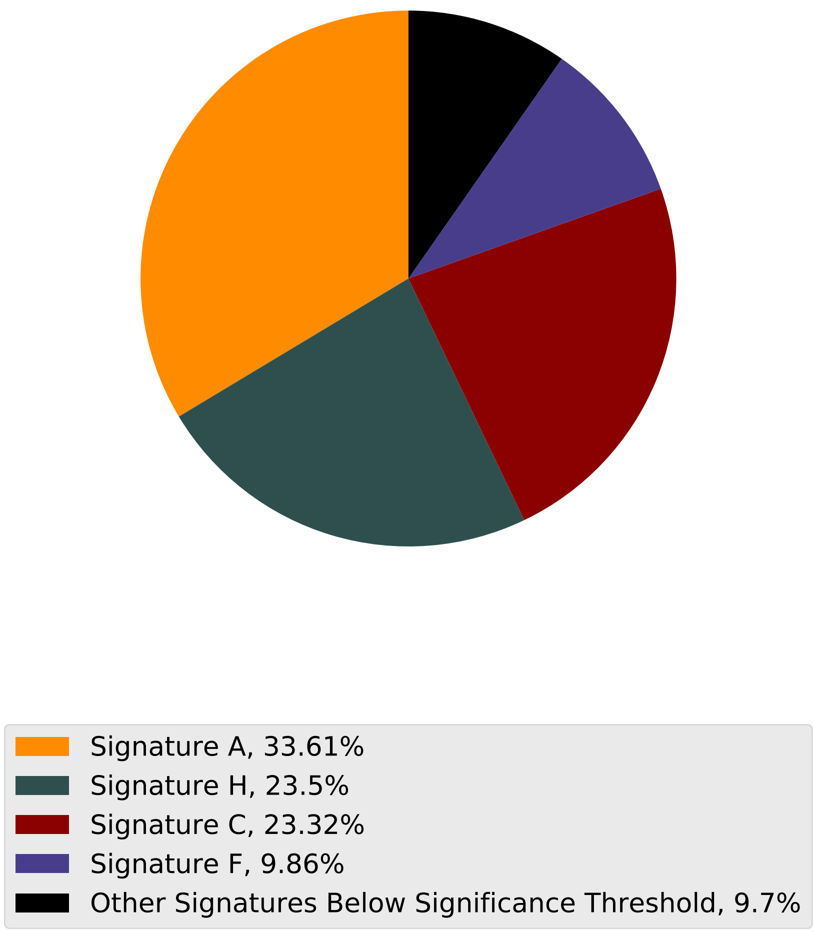

Tumor Profile for TCGA-BH-A0E0

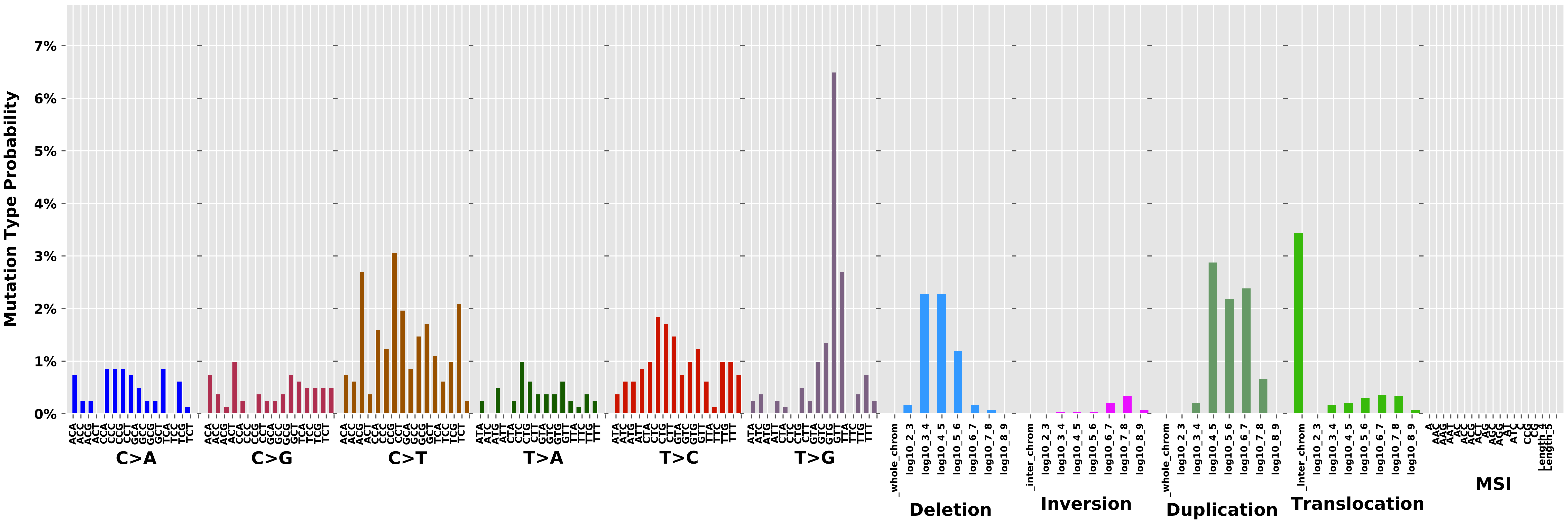

Cancer processes Weights for TCGA-A2-A0D0

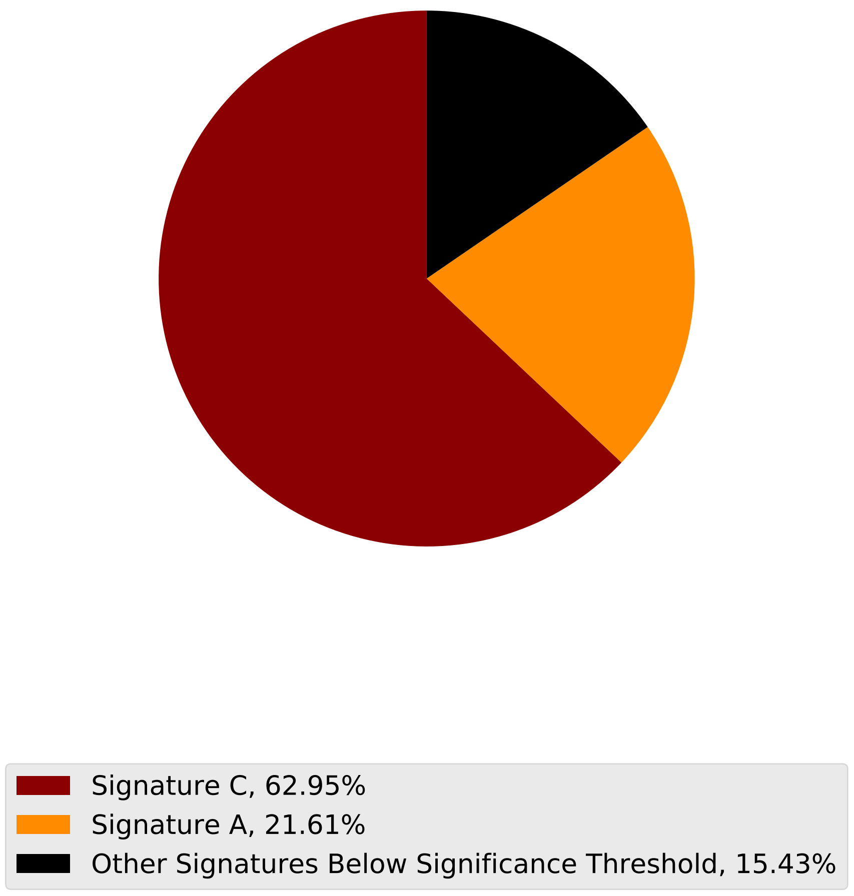

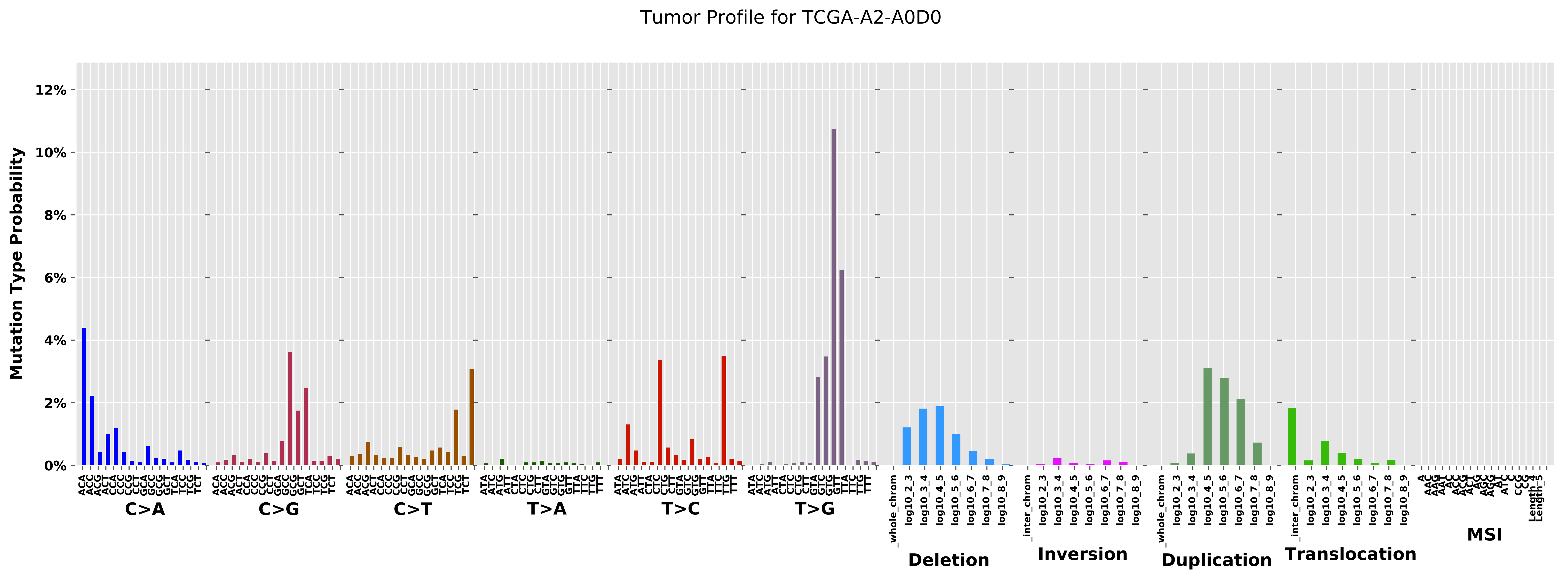
Tumor Profile for TCGA-A2-A0D0
Mutation Type Probability
0%
2%
4%
6%
8%
10%
12%
C>A
C>G
C>T
T>A
T>C
T>G
Deletion
Inversion
Duplication
Translocation
MSI
_whole_chrom
_inter_chrom
log10_2_3
log10_3_4
log10_4_5
log10_5_6
log10_6_7
log10_7_8
log10_8_9

Cancer processes Weights for TCGA-AA-3956

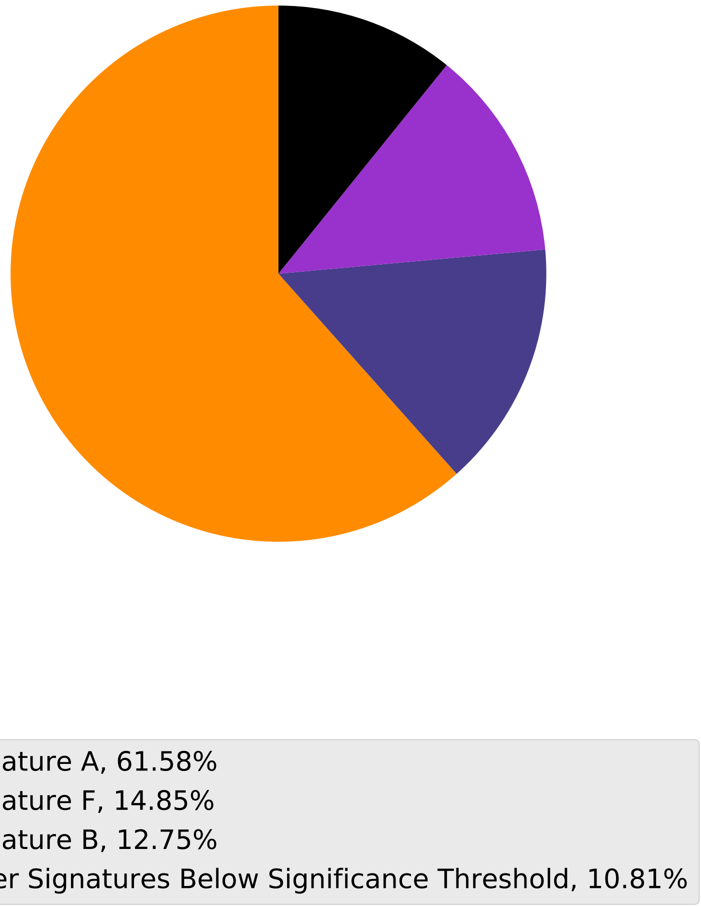

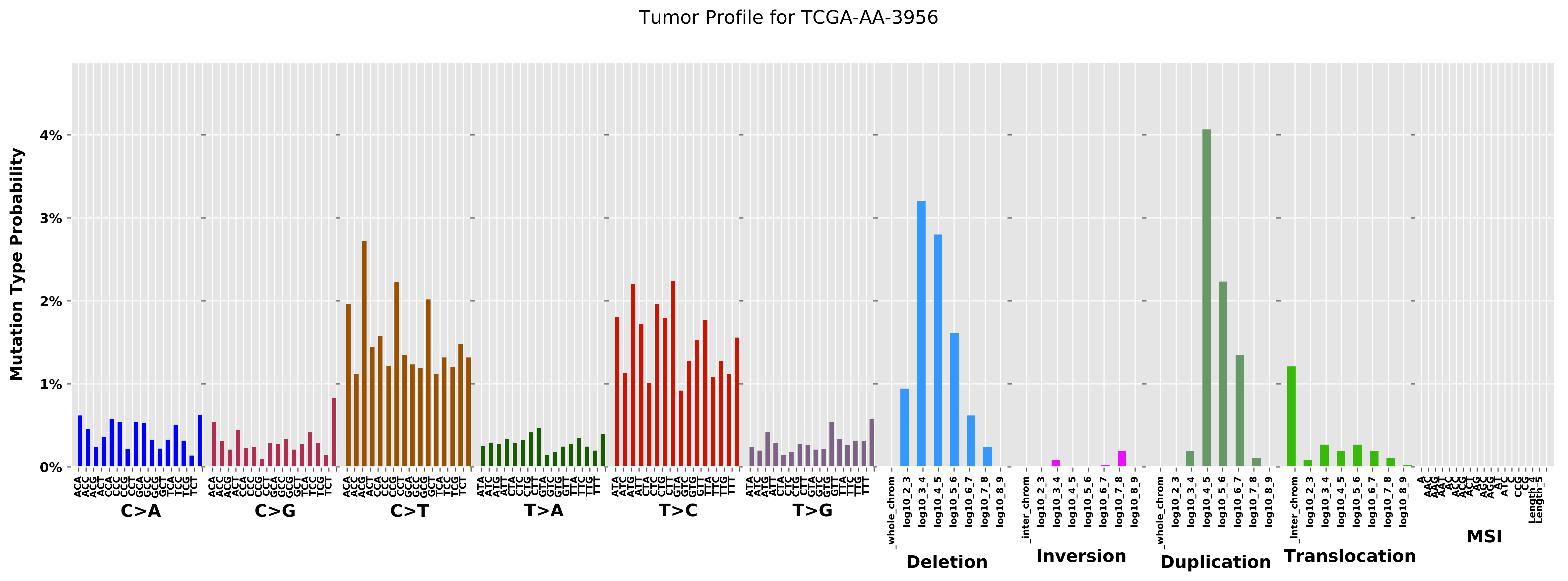
Tumor Profile for TCGA-AA-3956
Mutation Type Probability
0%
1%
2%
3%
4%
C>A
C>G
C>T
T>A
T>C
T>G
Deletion
Inversion
Duplication
Translocation
MSI

Cancer processes Weights for TCGA-AR-A0TX

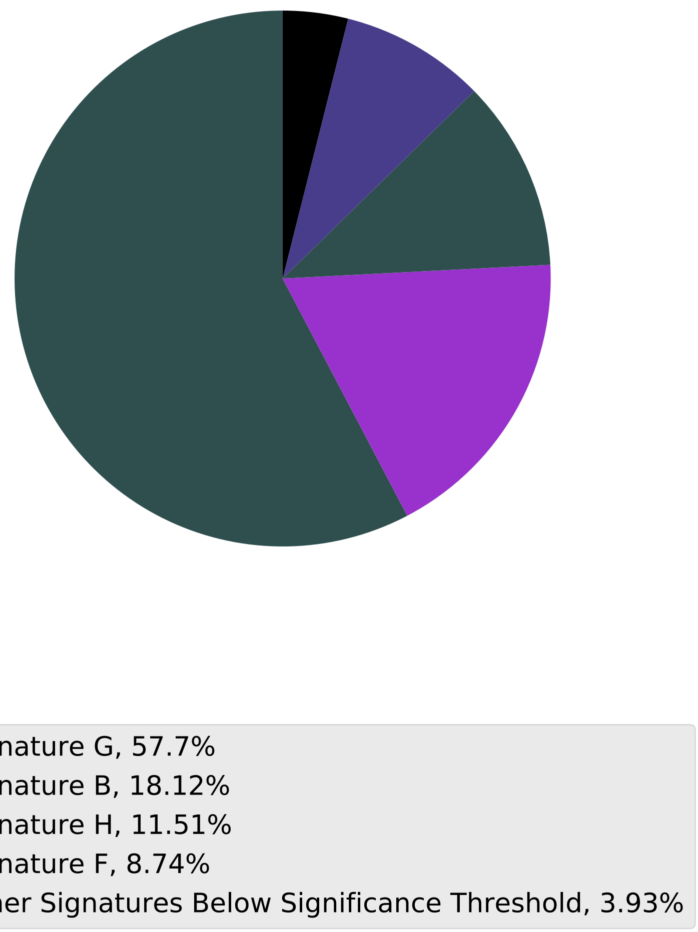

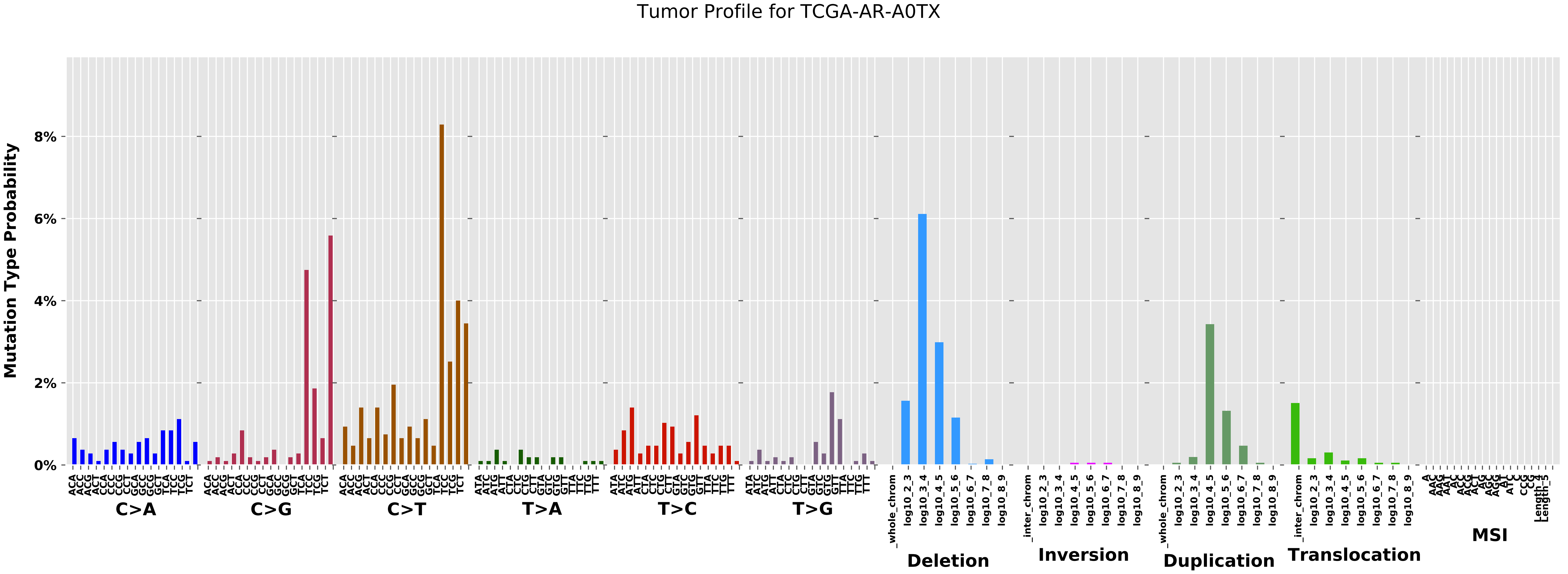

Tumor Profile for TCGA-AR-A0TX
Mutation Type Probability
0%
2%
4%
6%
8%
C>A
C>G
C>T
T>A
T>C
T>G
Deletion
Inversion
Duplication
Translocation
MSI

Cancer processes Weights for TCGA-EI-6917

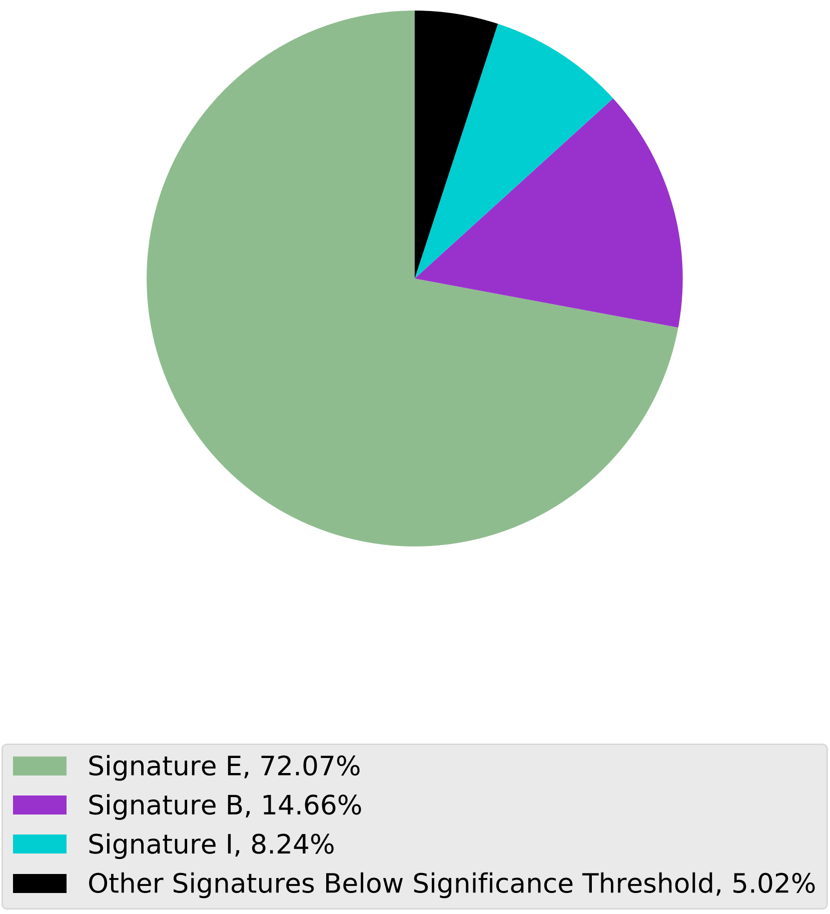

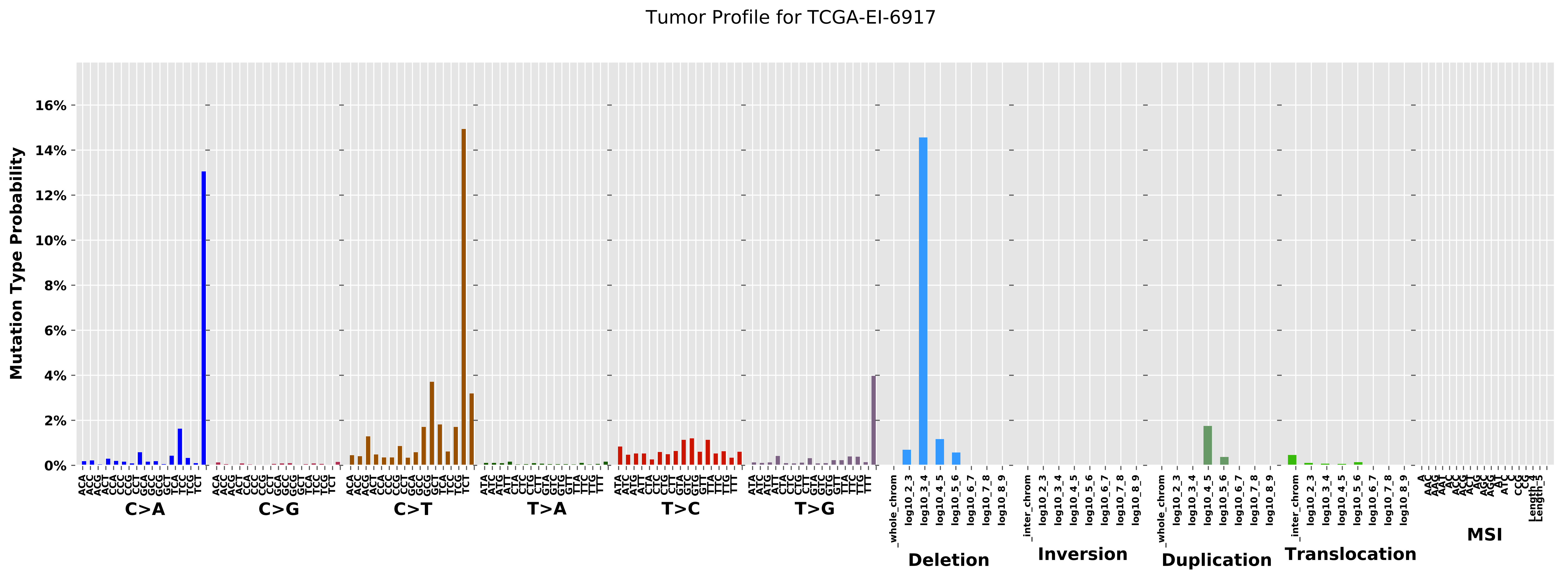

Tumor Profile for TCGA-EI-6917
Mutation Type Probability
0%
2%
4%
6%
8%
10%
12%
14%
16%
C>A
C>G
C>T
T>A
T>C
T>G
Deletion
Inversion
Duplication
Translocation
MSI

### Cancer processes Weights for TCGA-AA-A01X

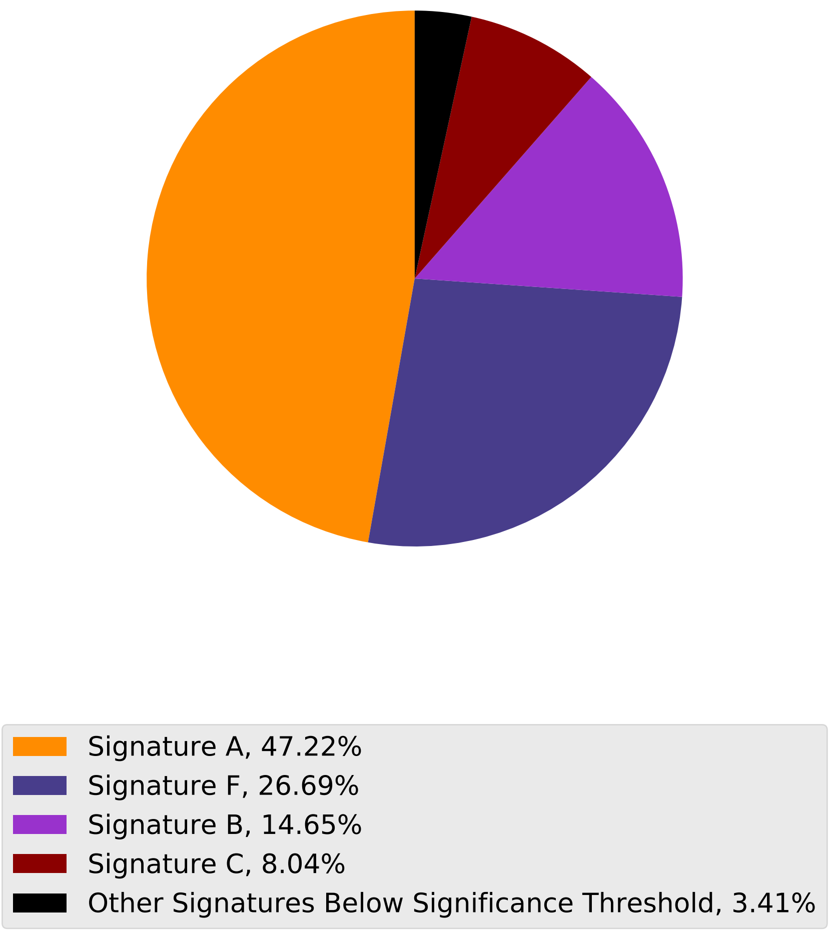

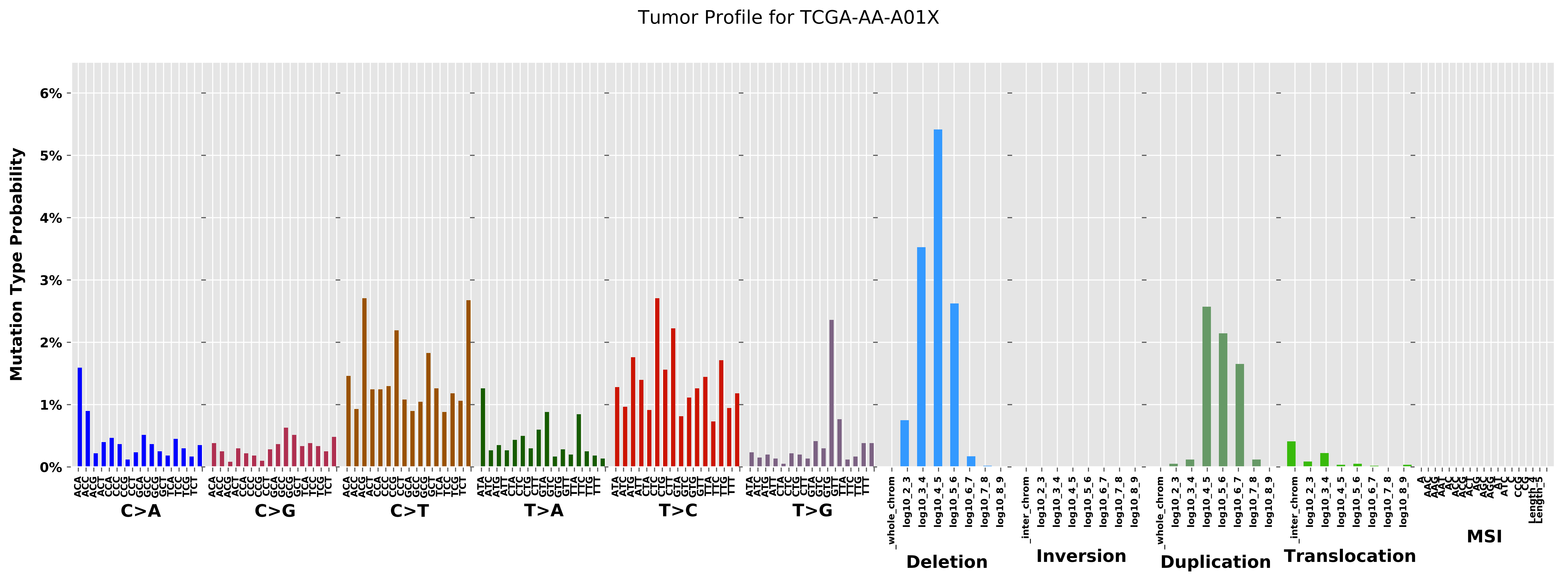
Tumor Profile for TCGA-AA-A01X
Mutation Type Probability
0%
1%
2%
3%
4%
5%
6%
C>A
C>G
C>T
T>A
T>C
T>G
Deletion
Inversion
Duplication
Translocation
MSI
_whole_chrom
_inter_chrom
log10_2_3
log10_3_4
log10_4_5
log10_5_6
log10_6_7
log10_7_8
log10_8_9

Cancer processes Weights for TCGA-E2-A15H

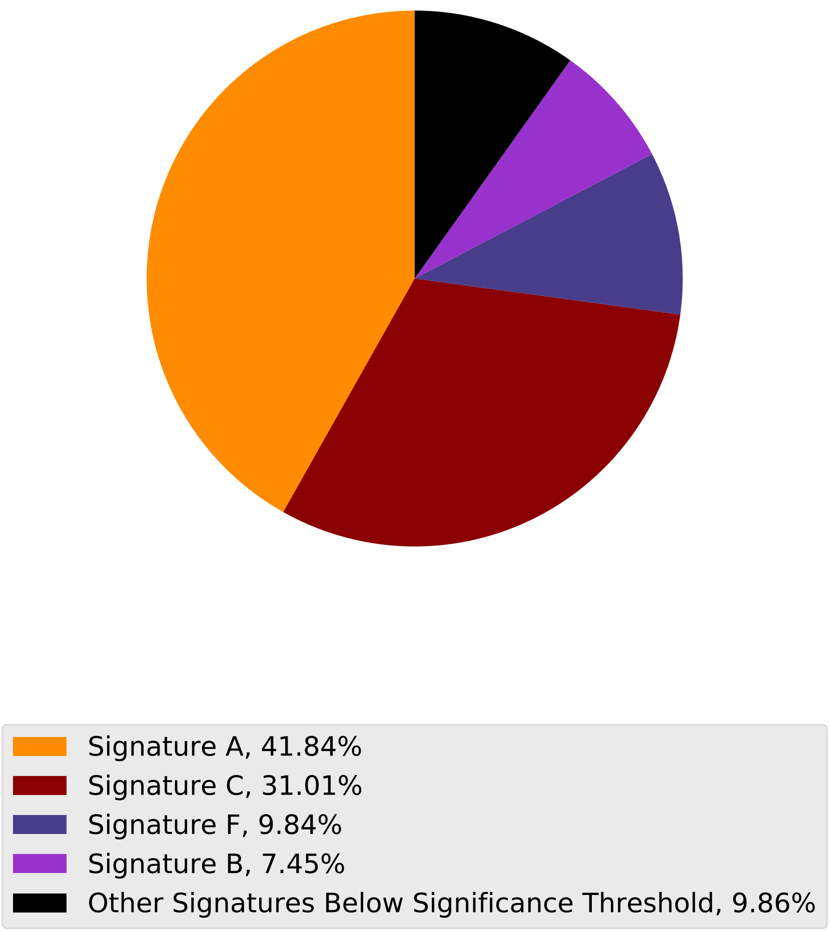

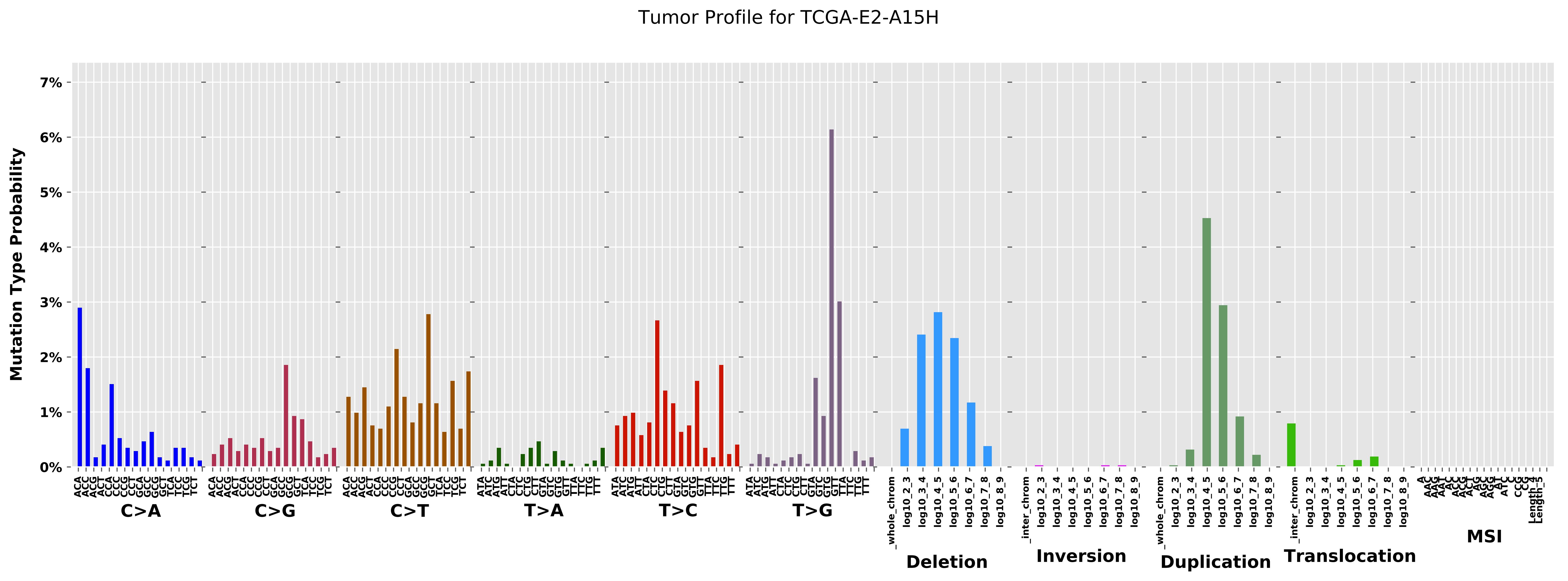
Tumor Profile for TCGA-E2-A15H
Mutation Type Probability
0%
1%
2%
3%
4%
5%
6%
7%
C>A
C>G
C>T
T>A
T>C
T>G
Deletion
Inversion
Duplication
Translocation
MSI
_whole_chrom
_inter_chrom
log10_2_3
log10_3_4
log10_4_5
log10_5_6
log10_6_7
log10_7_8
log10_8_9

Cancer processes Weights for TCGA-E9-A1NH

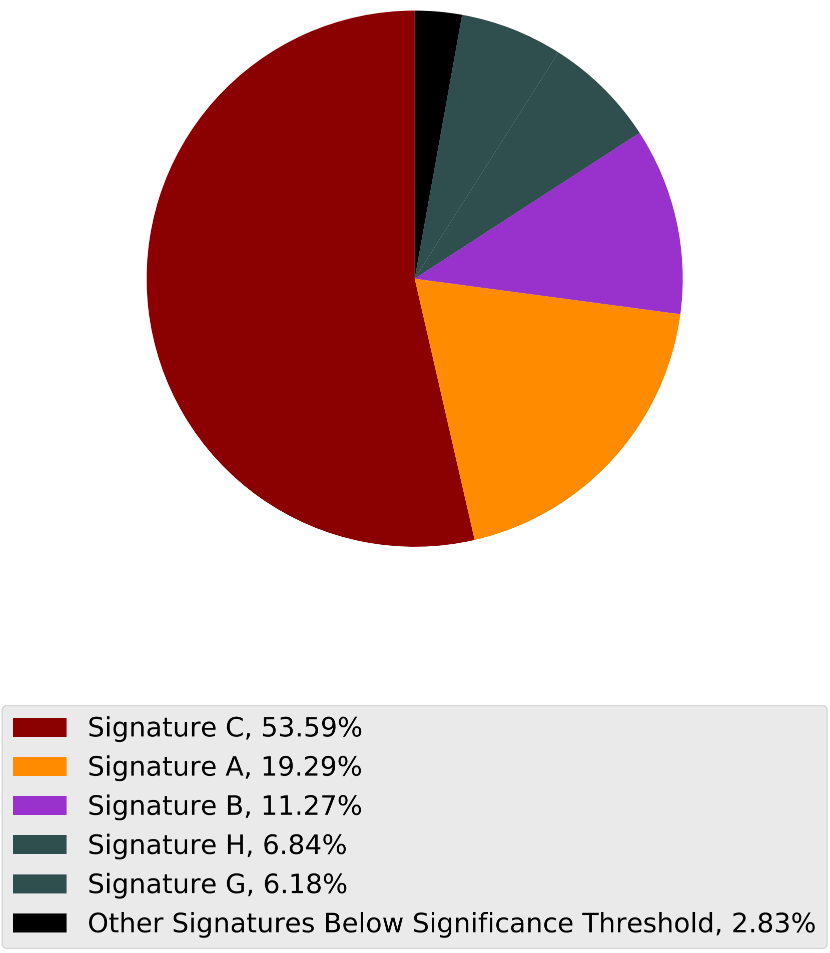

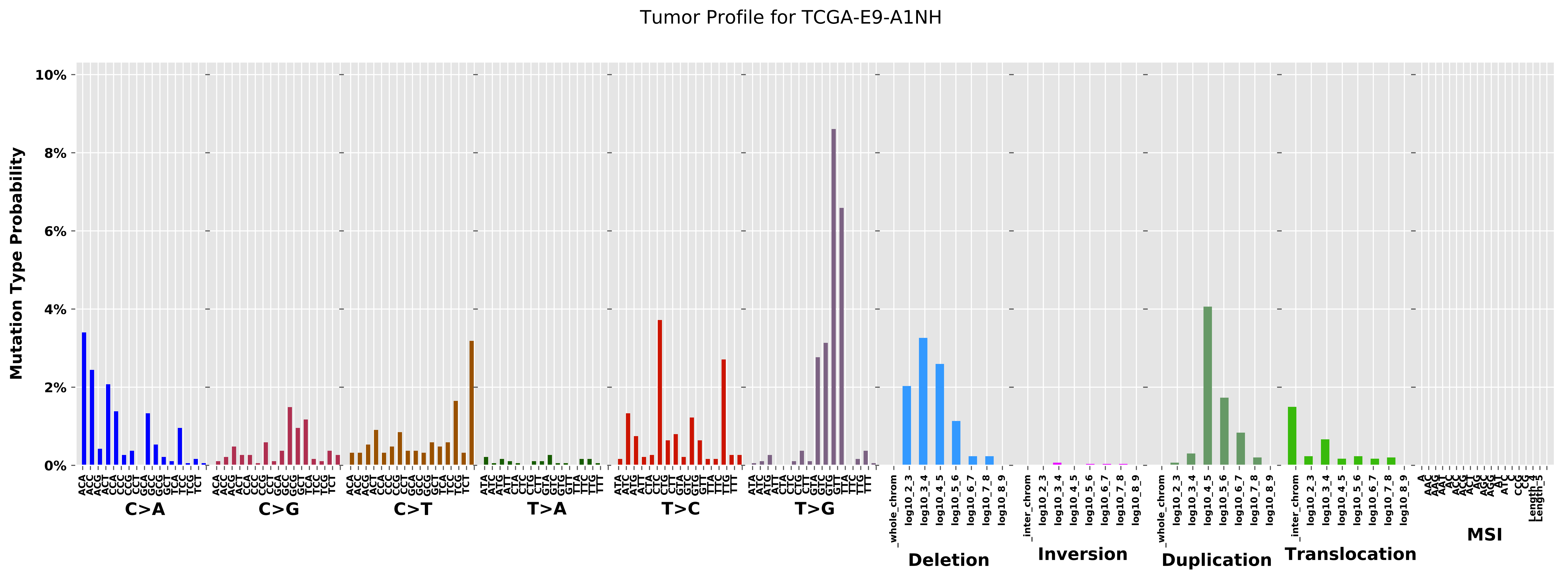

Tumor Profile for TCGA-E9-A1NH
Mutation Type Probability
10%
8%
6%
4%
2%
0%
C>A
C>G
C>T
T>A
T>C
T>G
Deletion
Inversion
Duplication
Translocation
MSI
_whole_chrom
_inter_chrom
log10_2_3
log10_3_4
log10_4_5
log10_5_6
log10_6_7
log10_7_8
log10_8_9

Cancer processes Weights for TCGA-AA-A01R

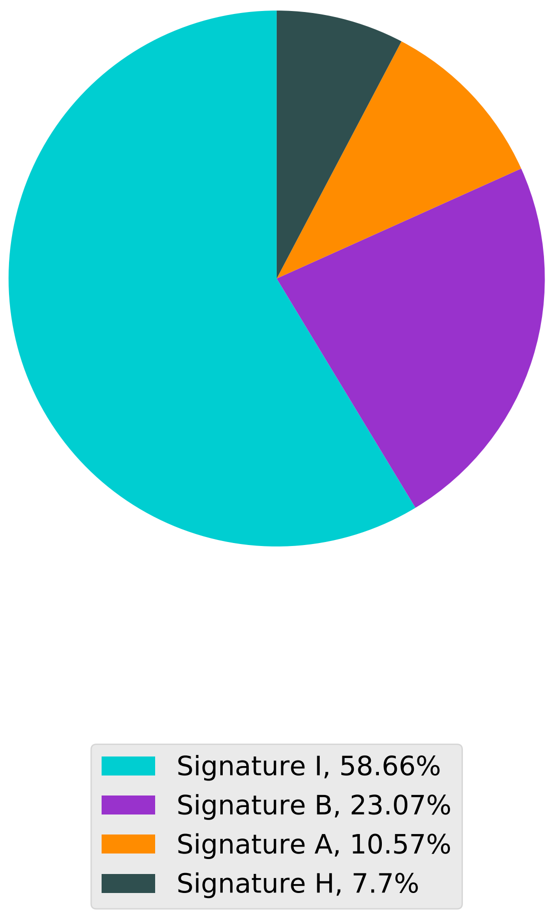

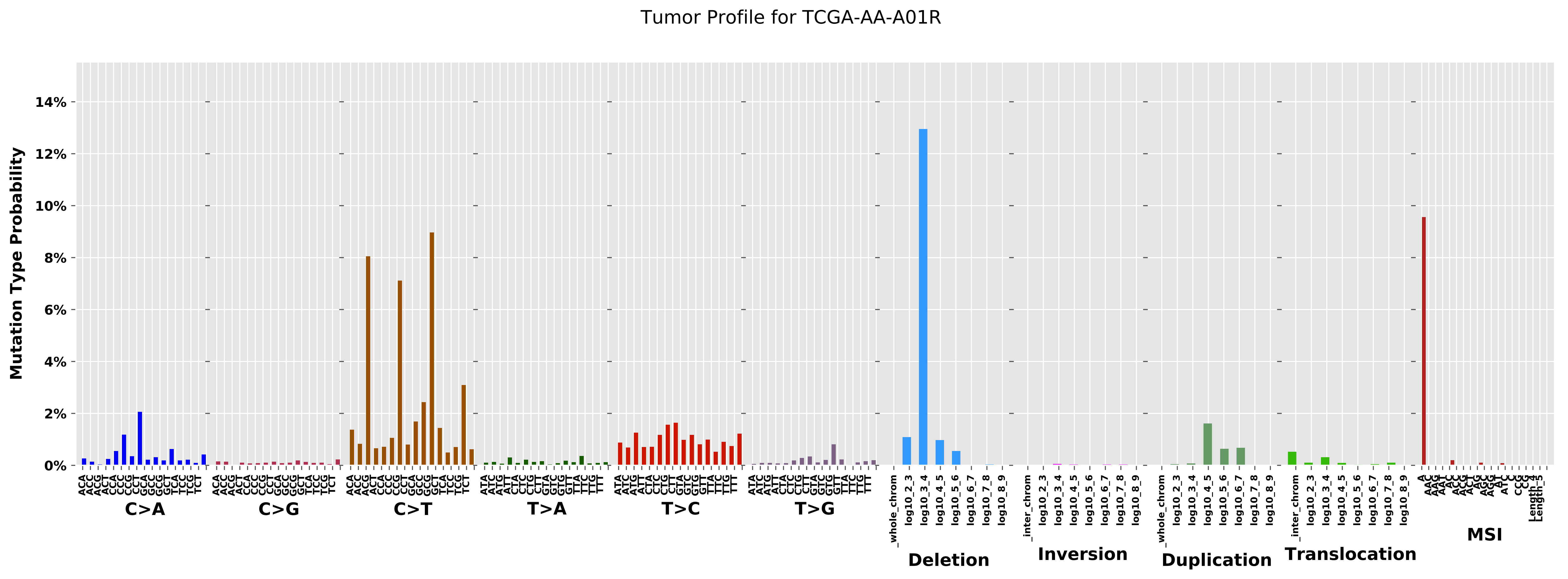

Tumor Profile for TCGA-AA-A01R
Mutation Type Probability
0%
2%
4%
6%
8%
10%
12%
14%
C>A
C>G
C>T
T>A
T>C
T>G
Deletion
Inversion
Duplication
Translocation
MSI

Cancer processes Weights for TCGA-AA-3666

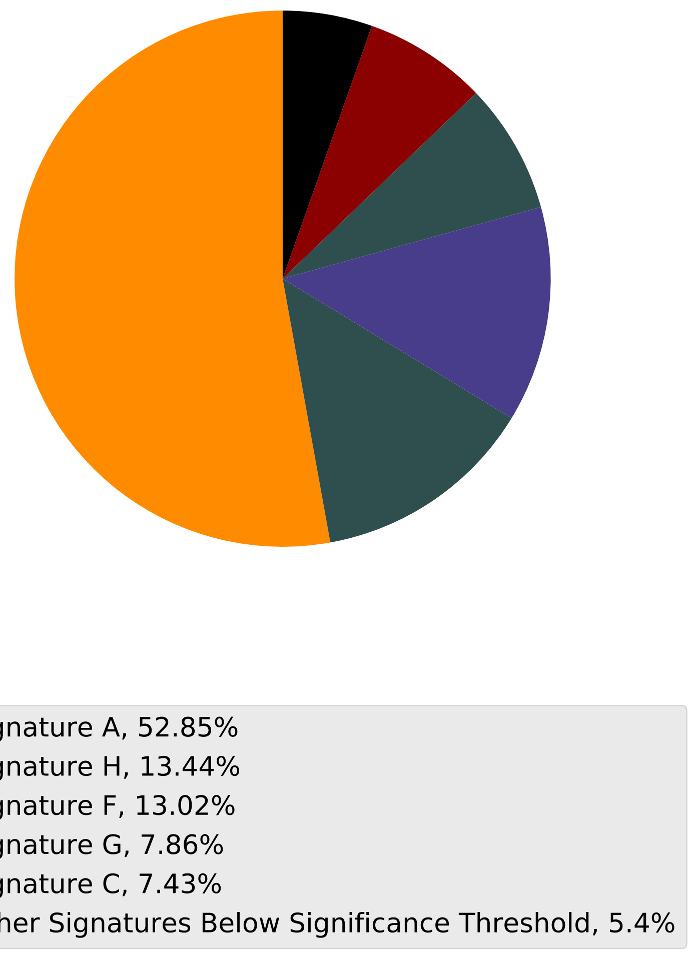

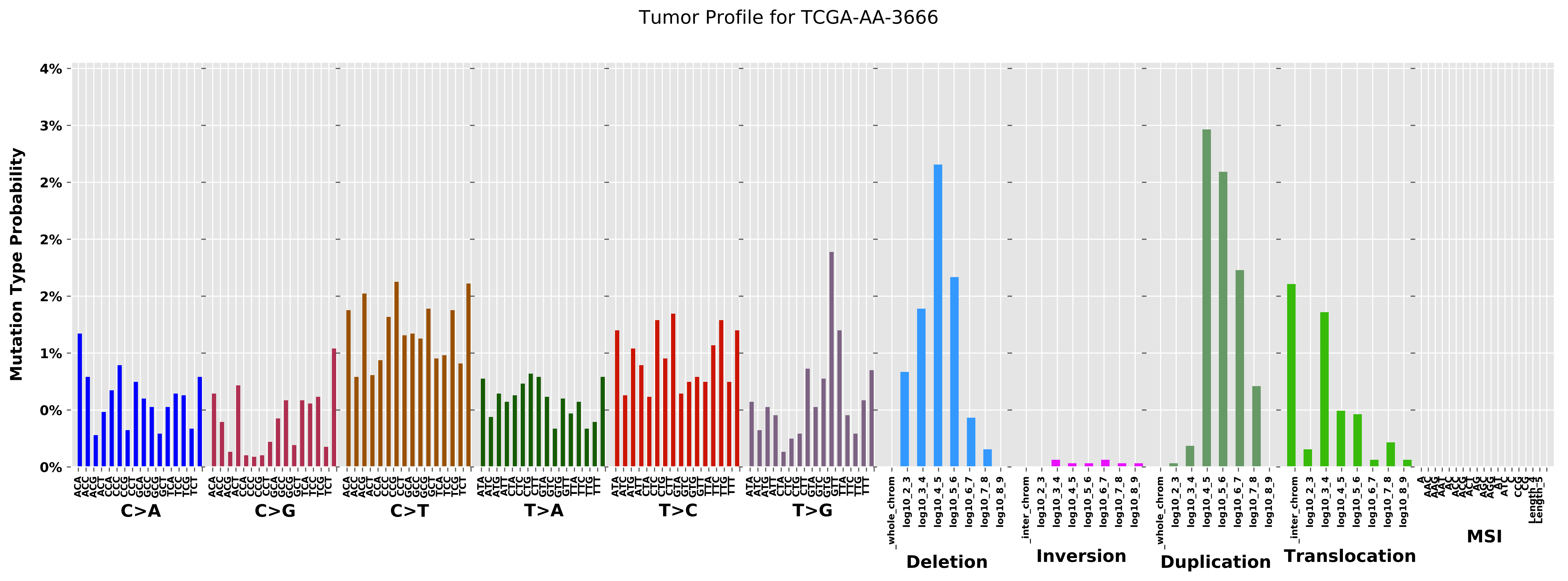
Tumor Profile for TCGA-AA-3666
Mutation Type Probability
4%
3%
2%
2%
1%
1%
0%
0%
C>A
C>G
C>T
T>A
T>C
T>G
Deletion
Inversion
Duplication
Translocation
MSI

Cancer processes Weights for TCGA-AR-A2LK

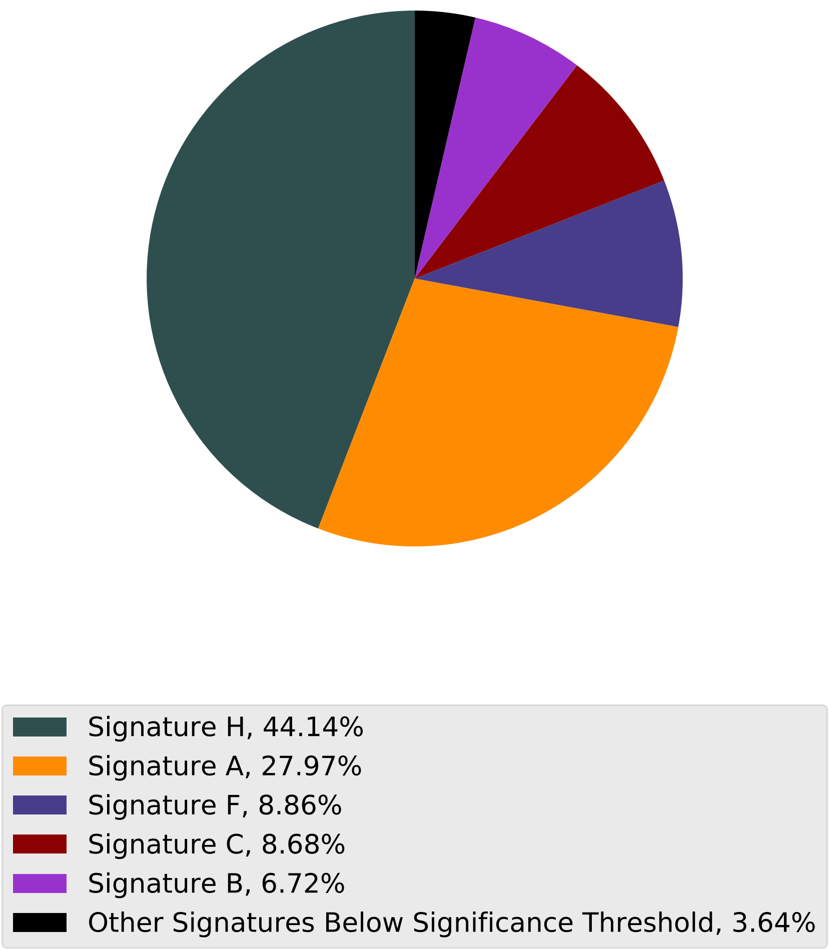

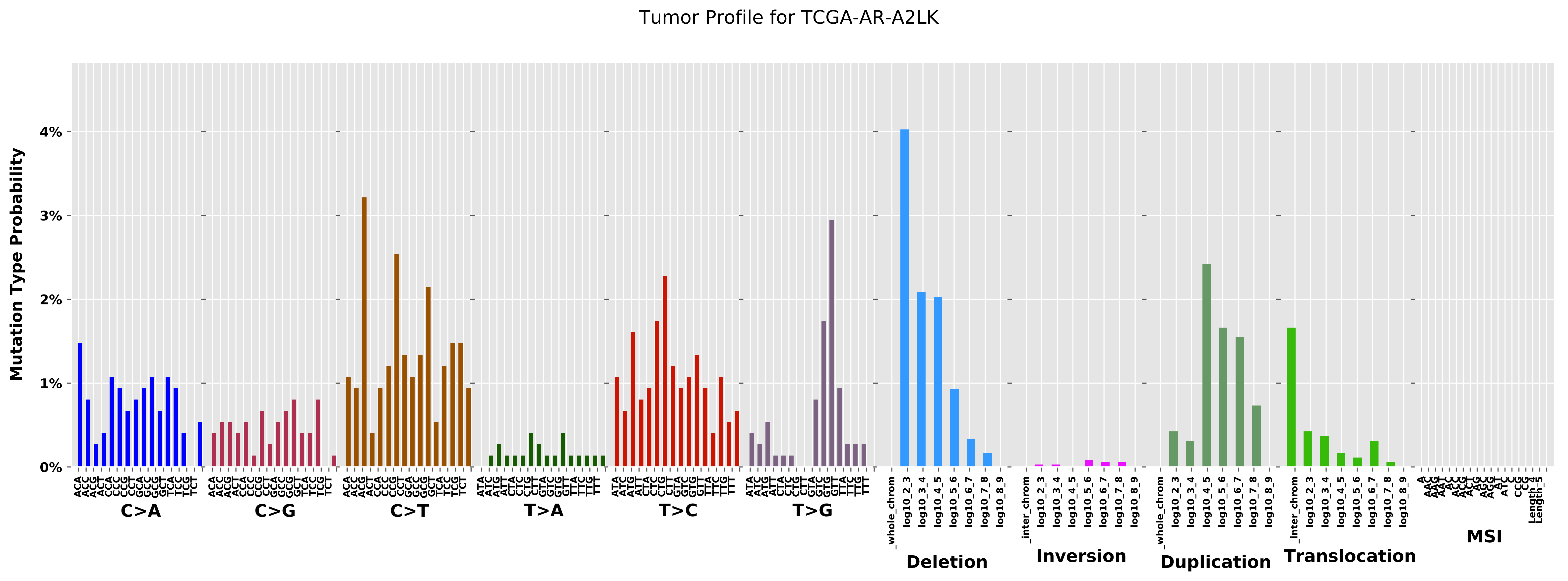

Tumor Profile for TCGA-AR-A2LK
Mutation Type Probability
0%
1%
2%
3%
4%
C>A
C>G
C>T
T>A
T>C
T>G
Deletion
Inversion
Duplication
Translocation
MSI

Cancer processes Weights for TCGA-AA-3534

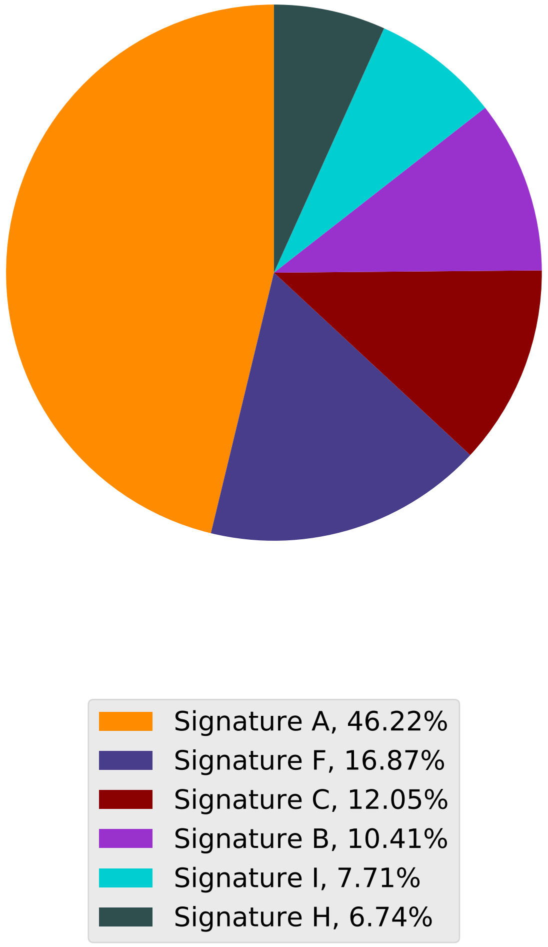

Tumor Profile for TCGA-AA-3534

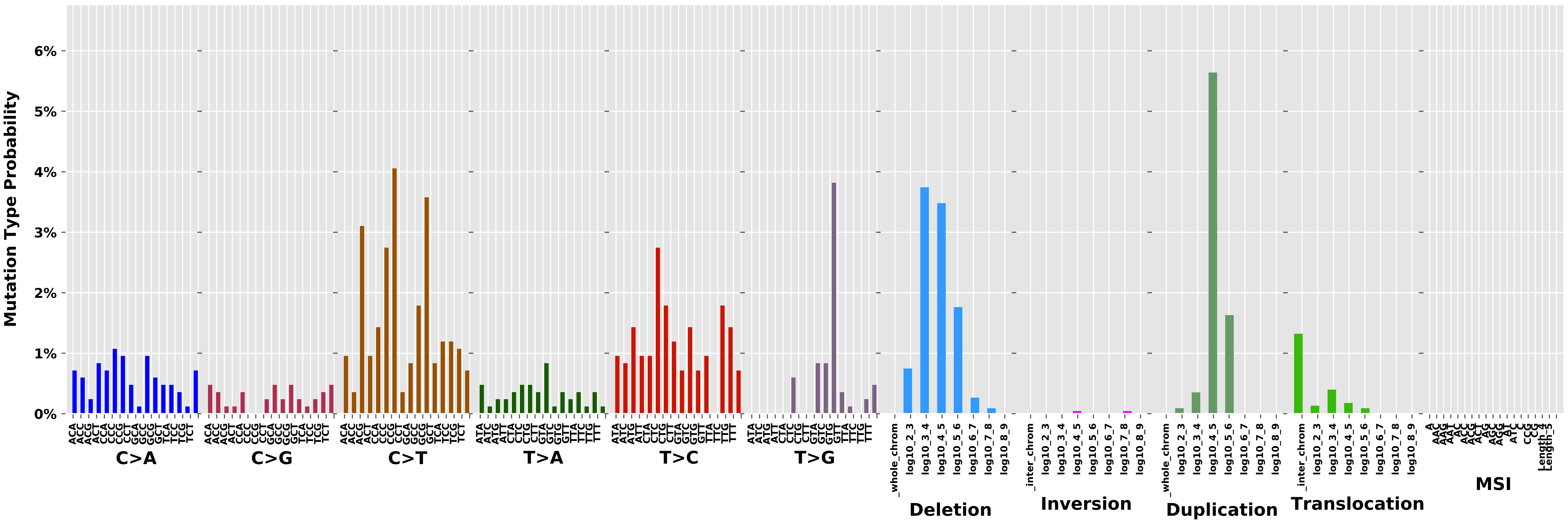

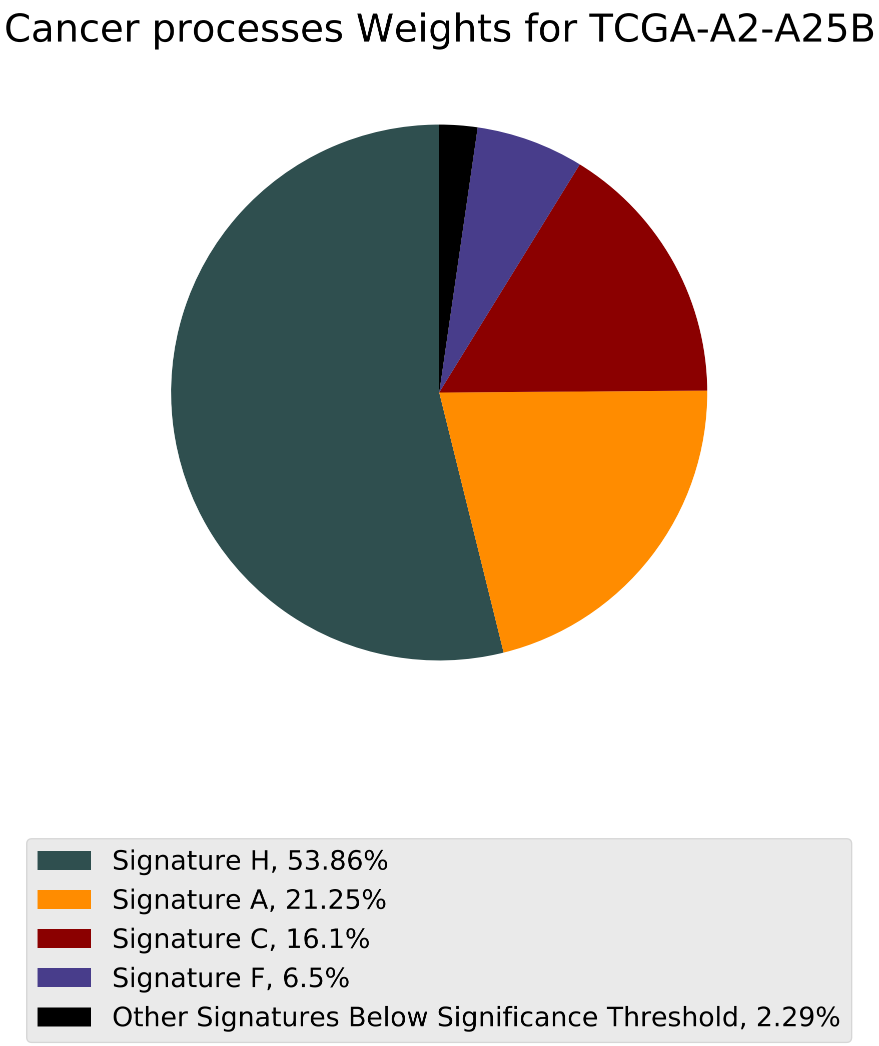
Cancer processes Weights for TCGA-A2-A25B
Signature H, 53.86%
Signature A, 21.25%
Signature C, 16.1%
Signature F, 6.5%
Other Signatures Below Significance Threshold, 2.29%

Tumor Profile for TCGA-A2-A25B

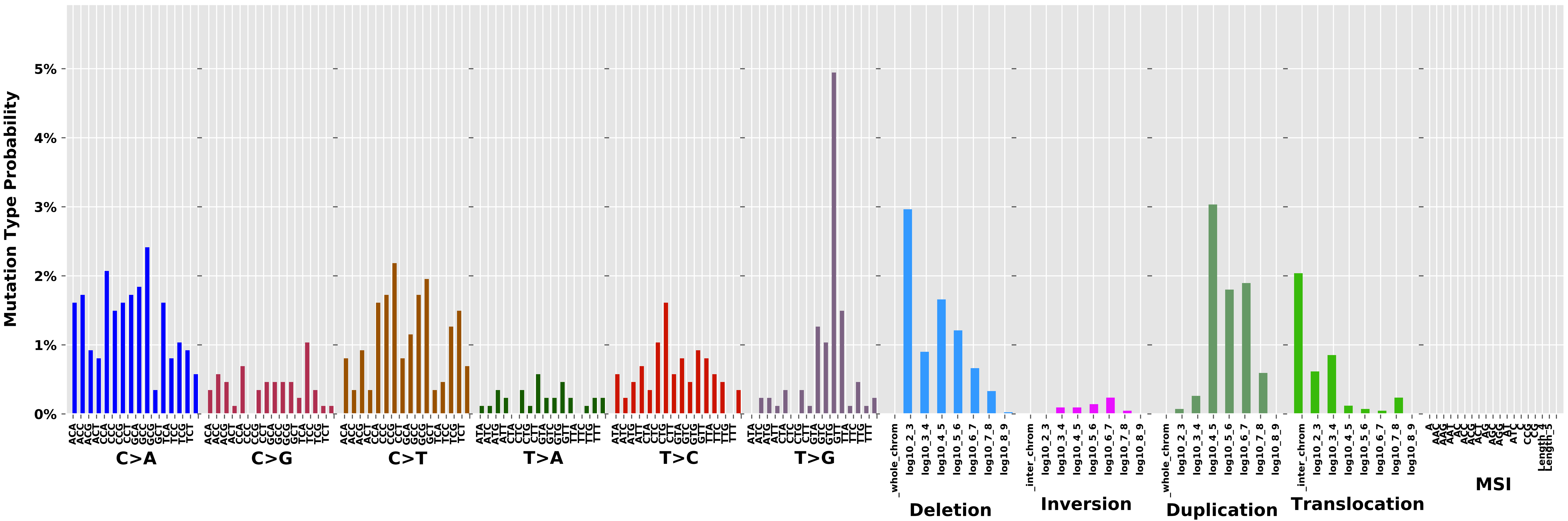

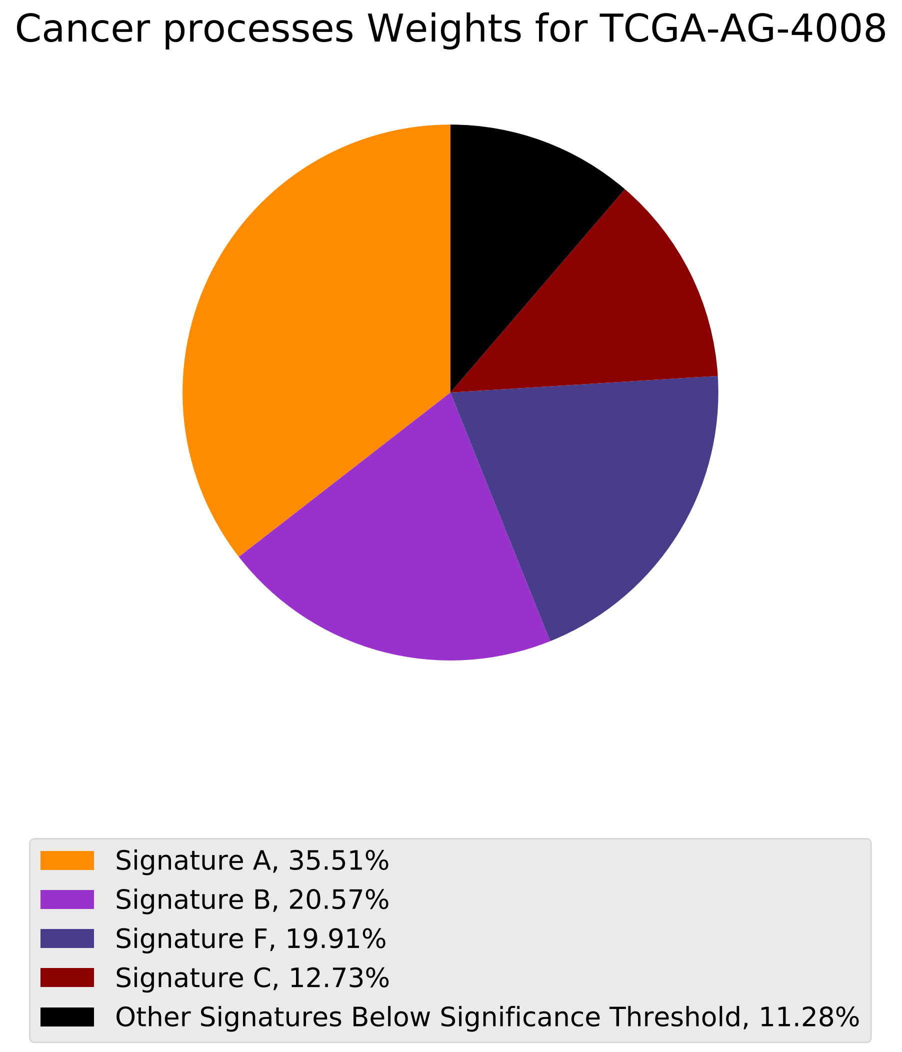
Cancer processes Weights for TCGA-AG-4008
Signature A, 35.51%
Signature B, 20.57%
Signature F, 19.91%
Signature C, 12.73%
Other Signatures Below Significance Threshold, 11.28%

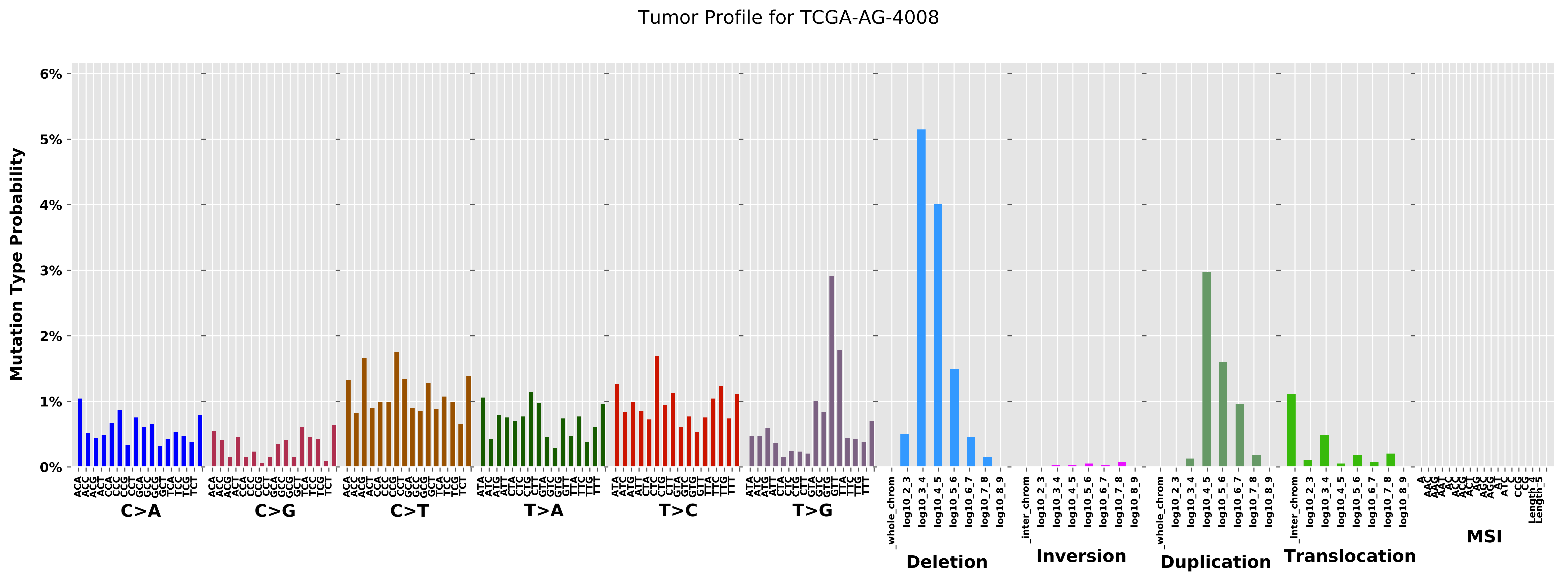
Tumor Profile for TCGA-AG-4008
Mutation Type Probability
0%
1%
2%
3%
4%
5%
6%
C>A
C>G
C>T
T>A
T>C
T>G
Deletion
Inversion
Duplication
Translocation
MSI
_whole_chrom
_inter_chrom
log10_2_3
log10_3_4
log10_4_5
log10_5_6
log10_6_7
log10_7_8
log10_8_9

Cancer processes Weights for TCGA-A7-A26G

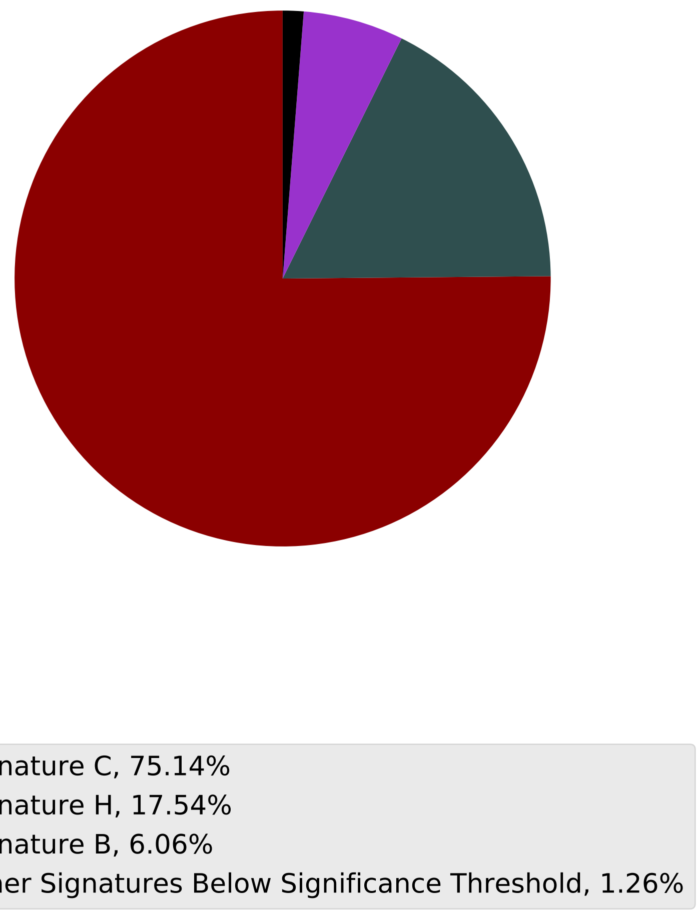

Tumor Profile for TCGA-A7-A26G

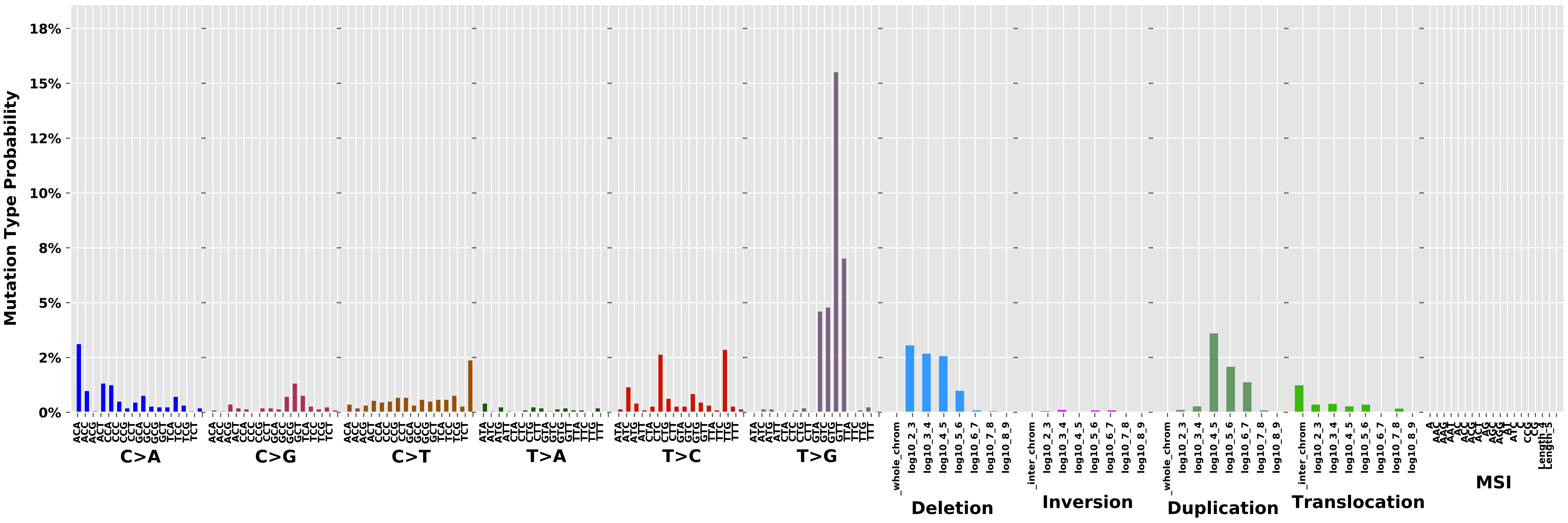

### Cancer processes Weights for TCGA-AR-A24Z

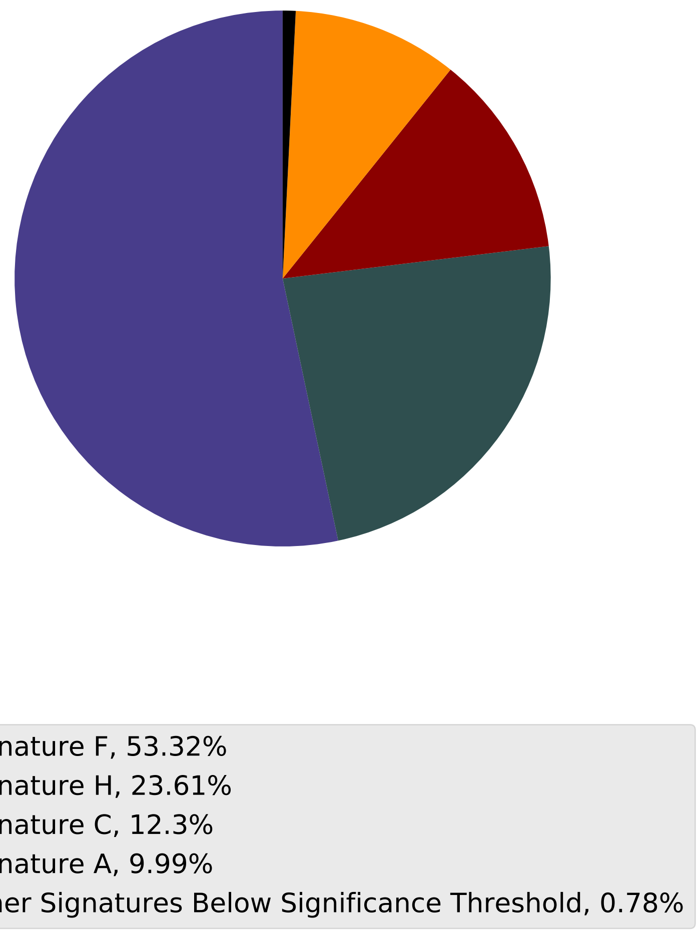

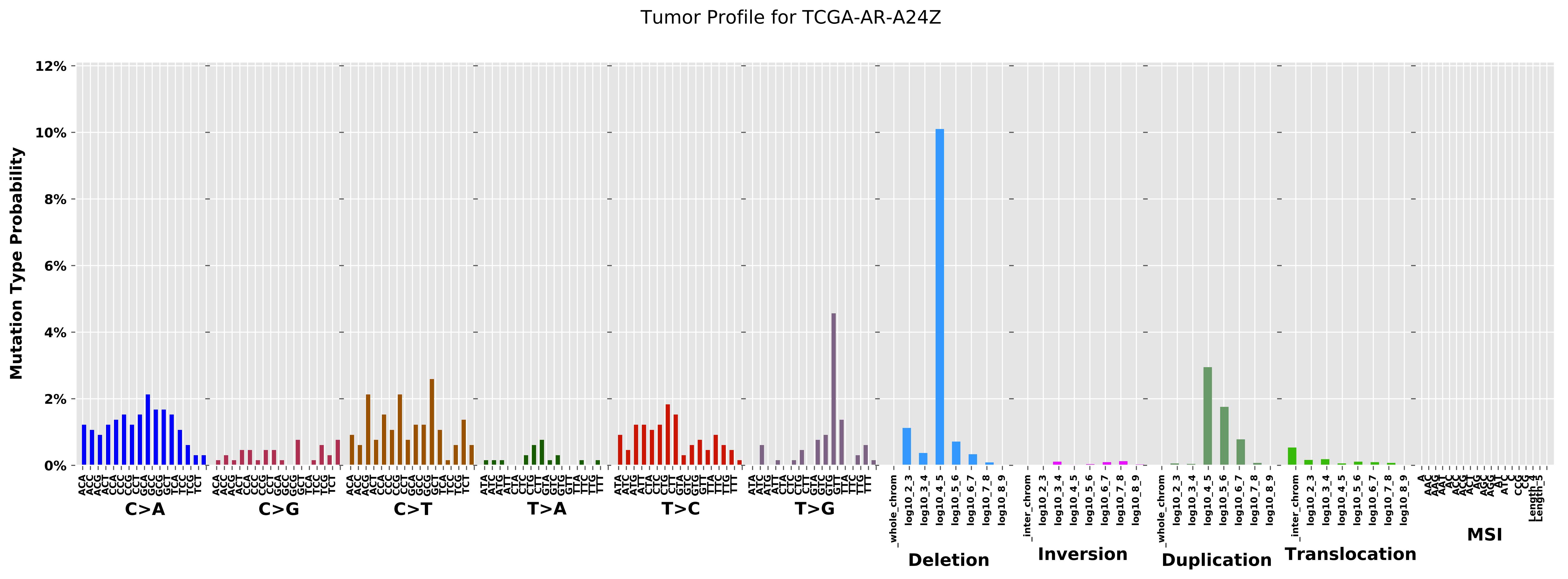

Tumor Profile for TCGA-AR-A24Z
Mutation Type Probability
12%
10%
8%
6%
4%
2%
0%
C>A
C>G
C>T
T>A
T>C
T>G
Deletion
Inversion
Duplication
Translocation
MSI

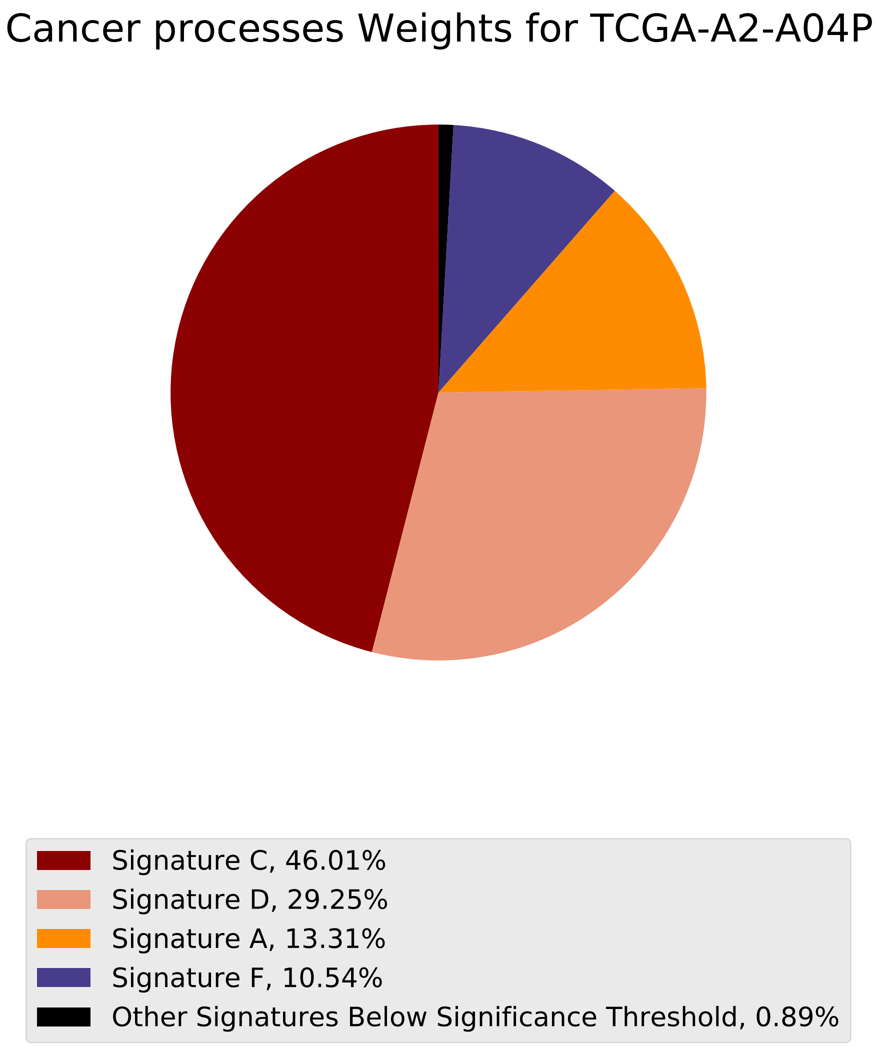
Cancer processes Weights for TCGA-A2-A04P
Signature C, 46.01%
Signature D, 29.25%
Signature A, 13.31%
Signature F, 10.54%
Other Signatures Below Significance Threshold, 0.89%

Tumor Profile for TCGA-A2-A04P

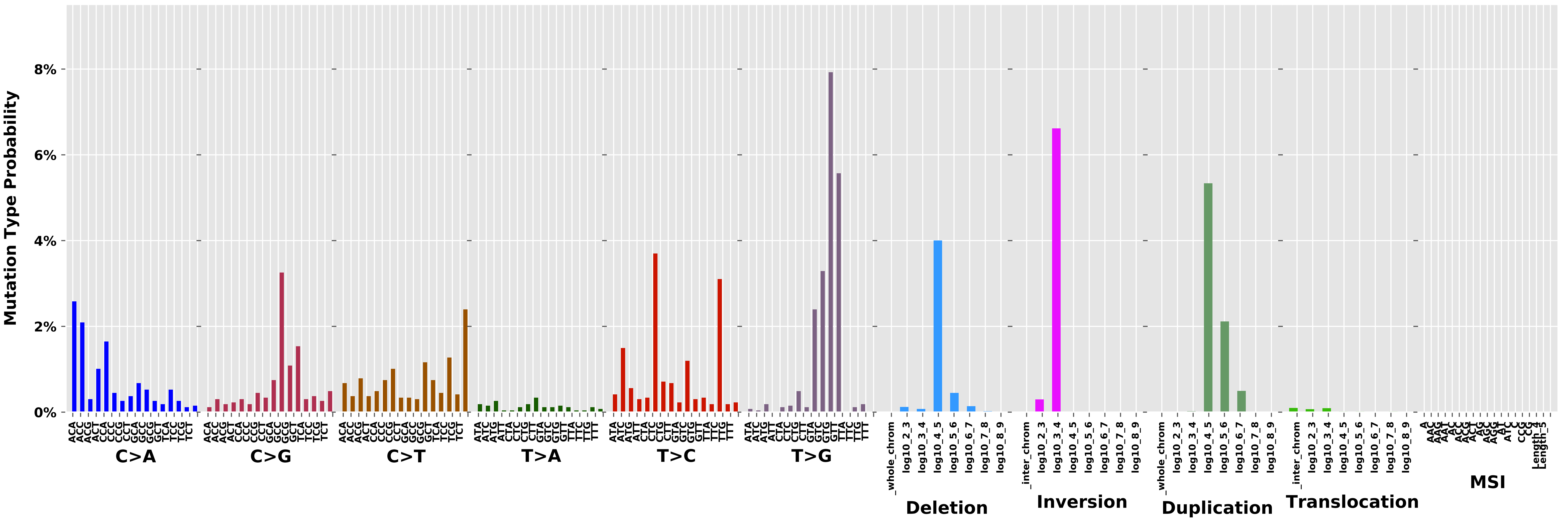

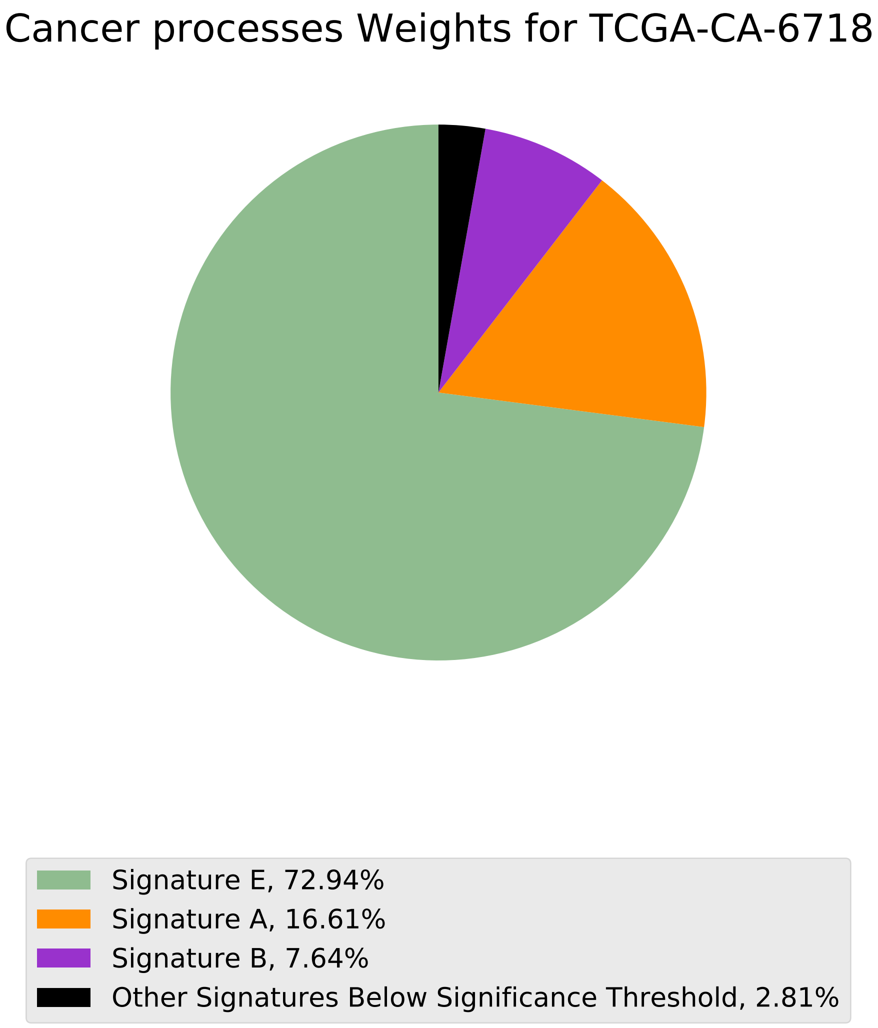
Cancer processes Weights for TCGA-CA-6718
Signature E, 72.94%
Signature A, 16.61%
Signature B, 7.64%
Other Signatures Below Significance Threshold, 2.81%

Tumor Profile for TCGA-CA-6718

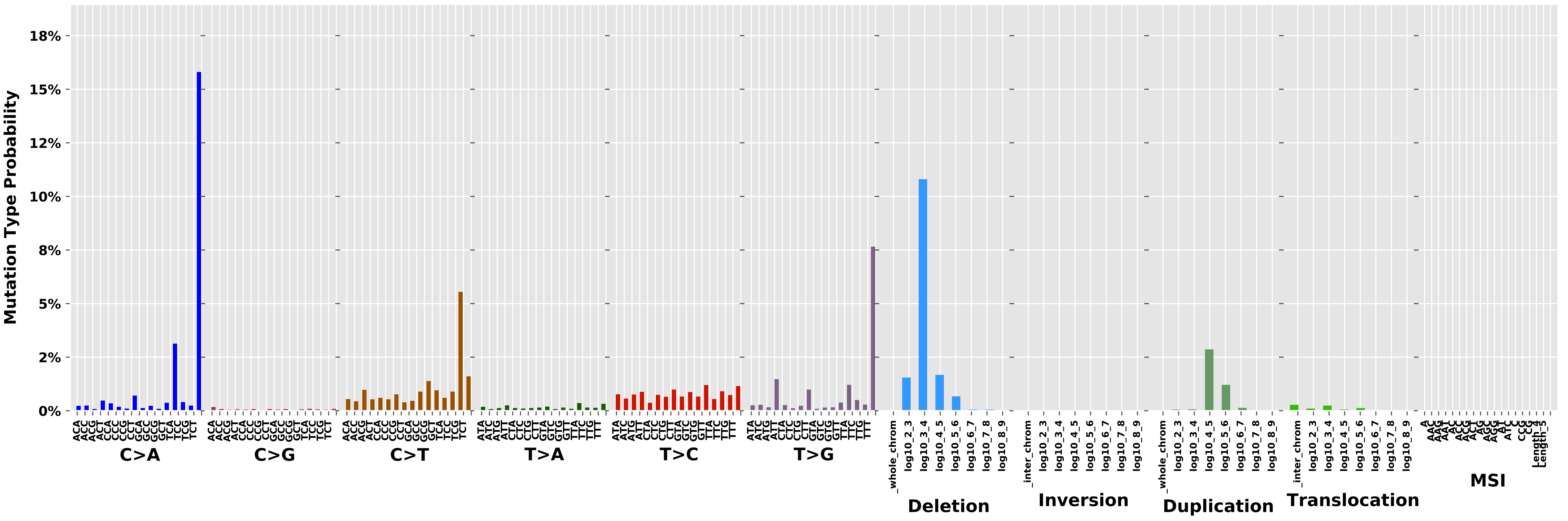

Cancer processes Weights for TCGA-BH-A0AV

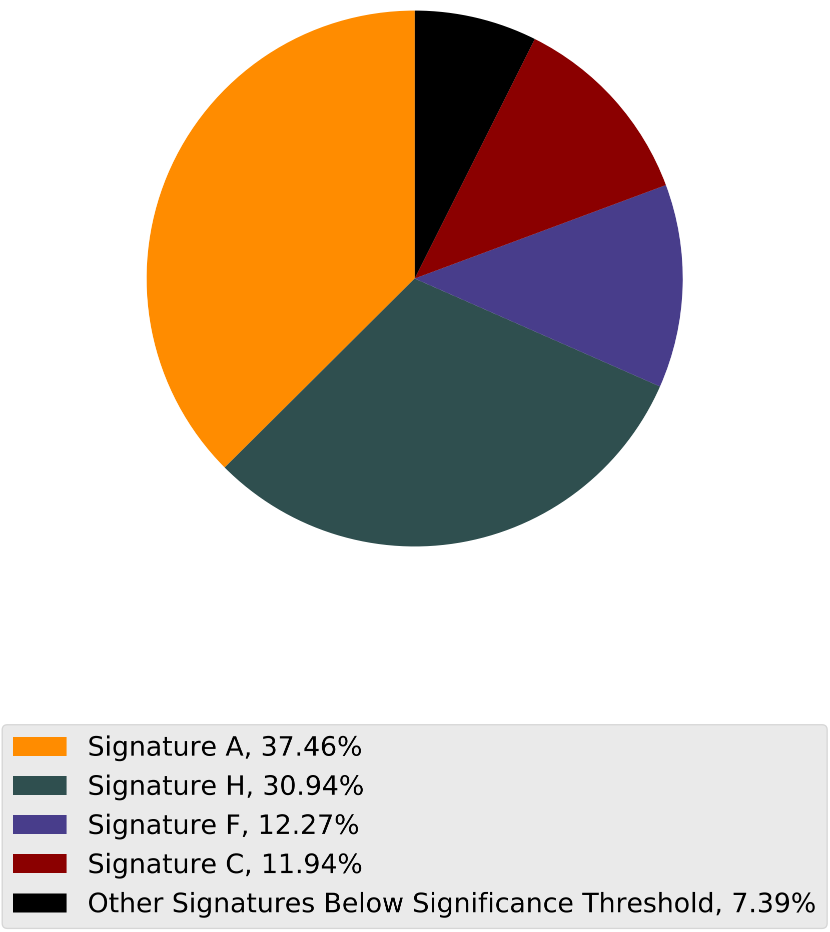

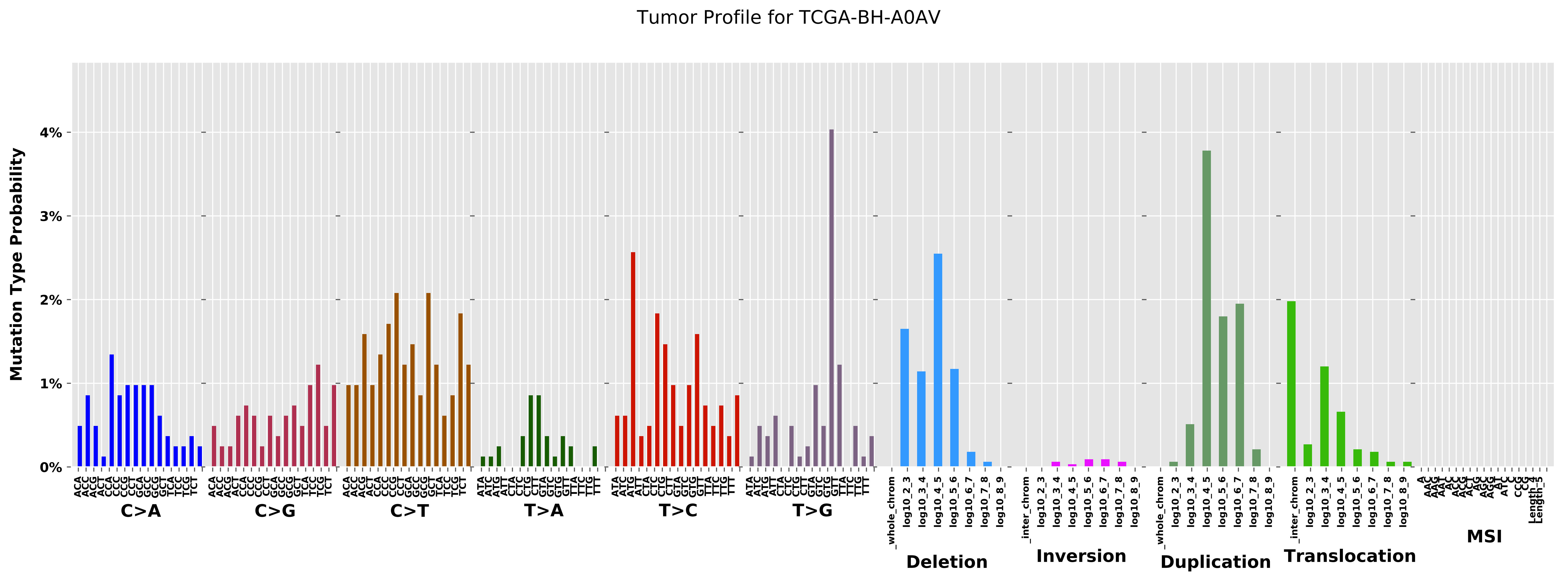

Tumor Profile for TCGA-BH-A0AV
Mutation Type Probability
0%
1%
2%
3%
4%
C>A
C>G
C>T
T>A
T>C
T>G
Deletion
Inversion
Duplication
Translocation
MSI

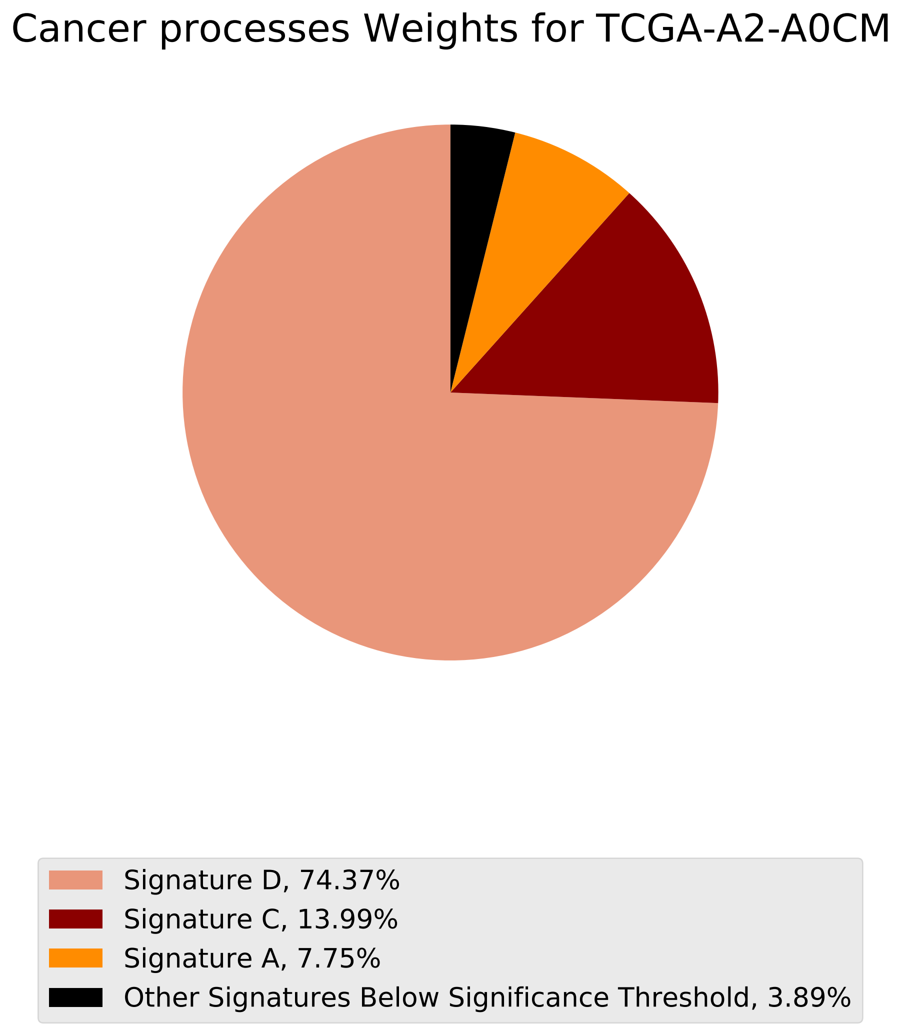
Cancer processes Weights for TCGA-A2-A0CM
Signature D, 74.37%
Signature C, 13.99%
Signature A, 7.75%
Other Signatures Below Significance Threshold, 3.89%

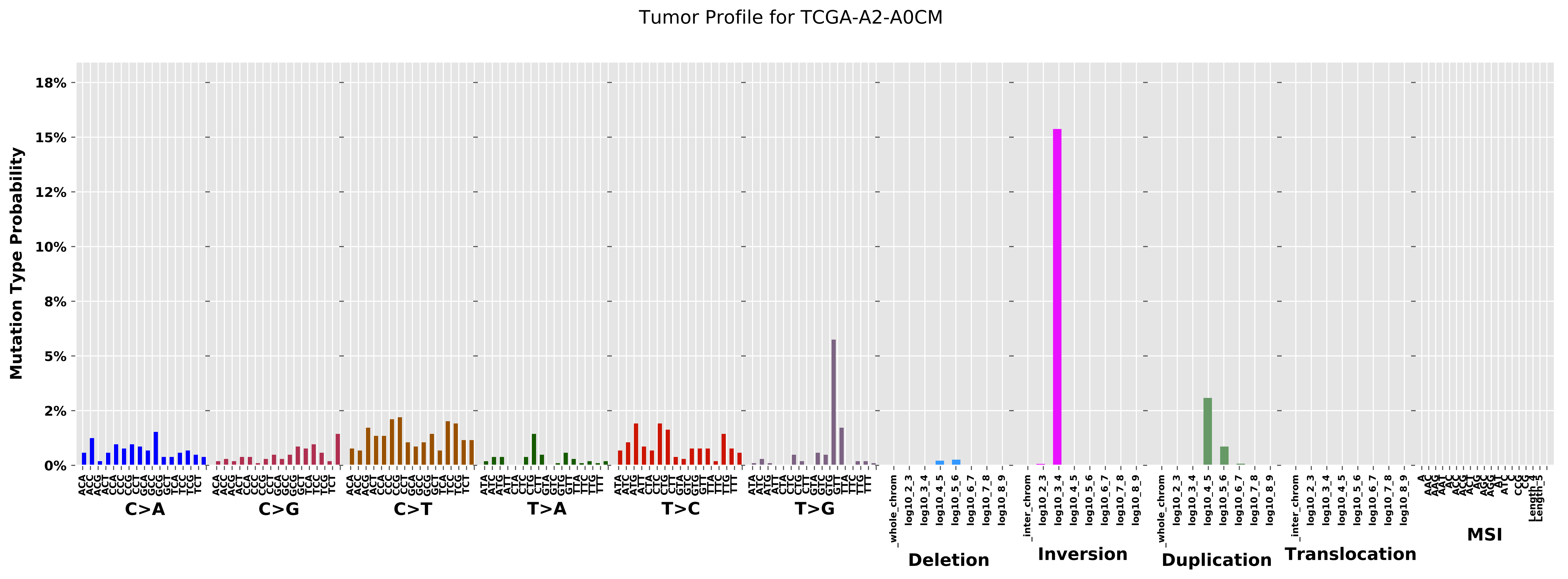

Tumor Profile for TCGA-A2-A0CM
Mutation Type Probability
18%
15%
12%
10%
8%
5%
2%
0%
C>A
C>G
C>T
T>A
T>C
T>G
Deletion
Inversion
Duplication
Translocation
MSI

### Cancer processes Weights for TCGA-AN-A04D

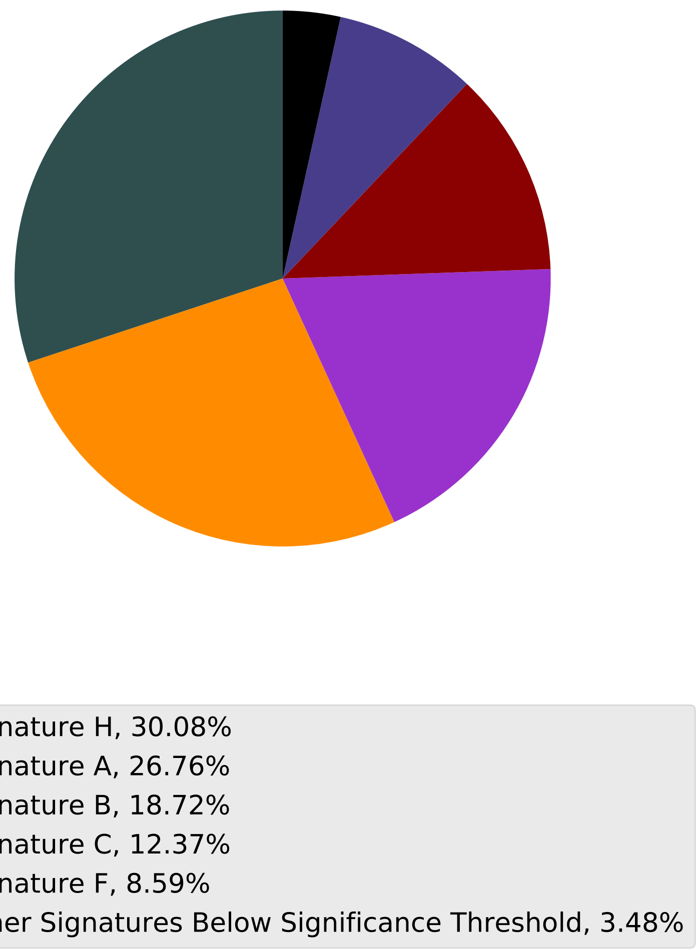

Tumor Profile for TCGA-AN-A04D

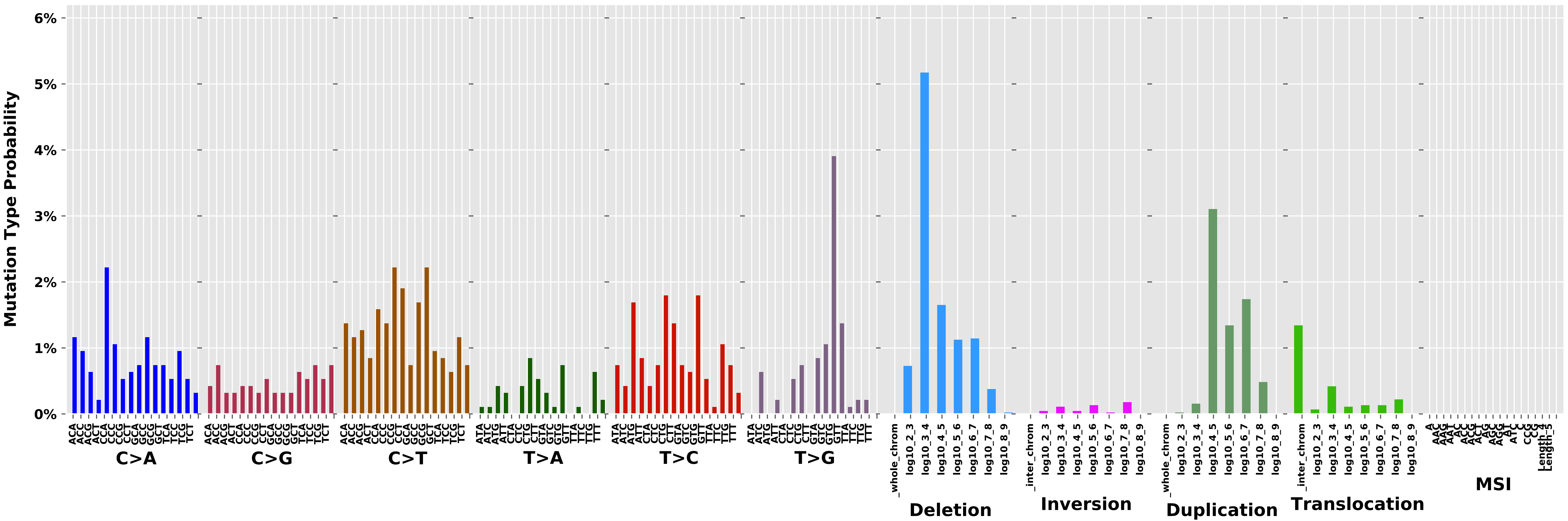

Cancer processes Weights for TCGA-A8-A092

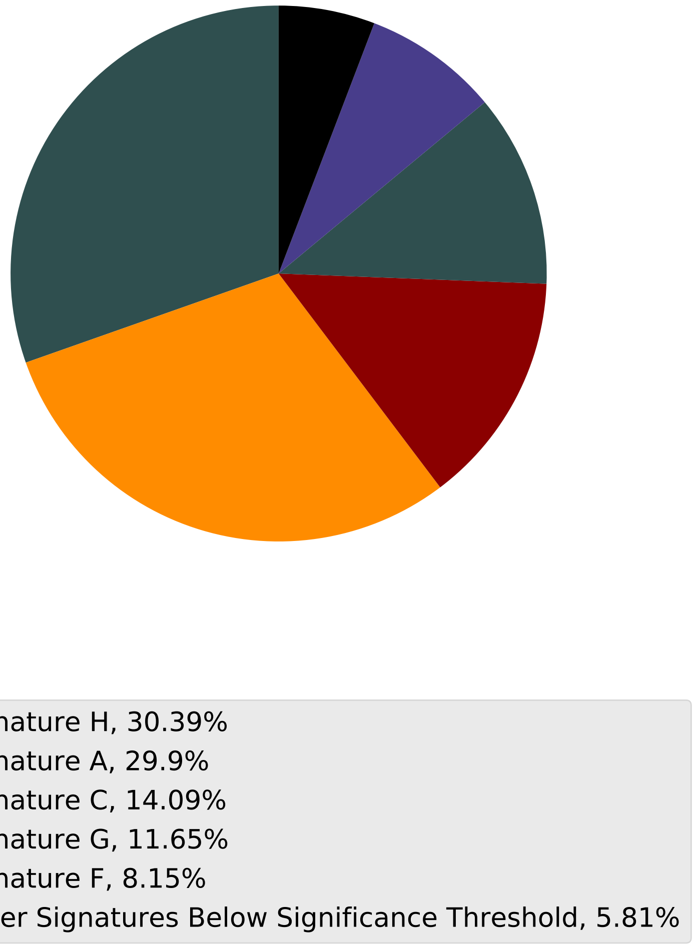

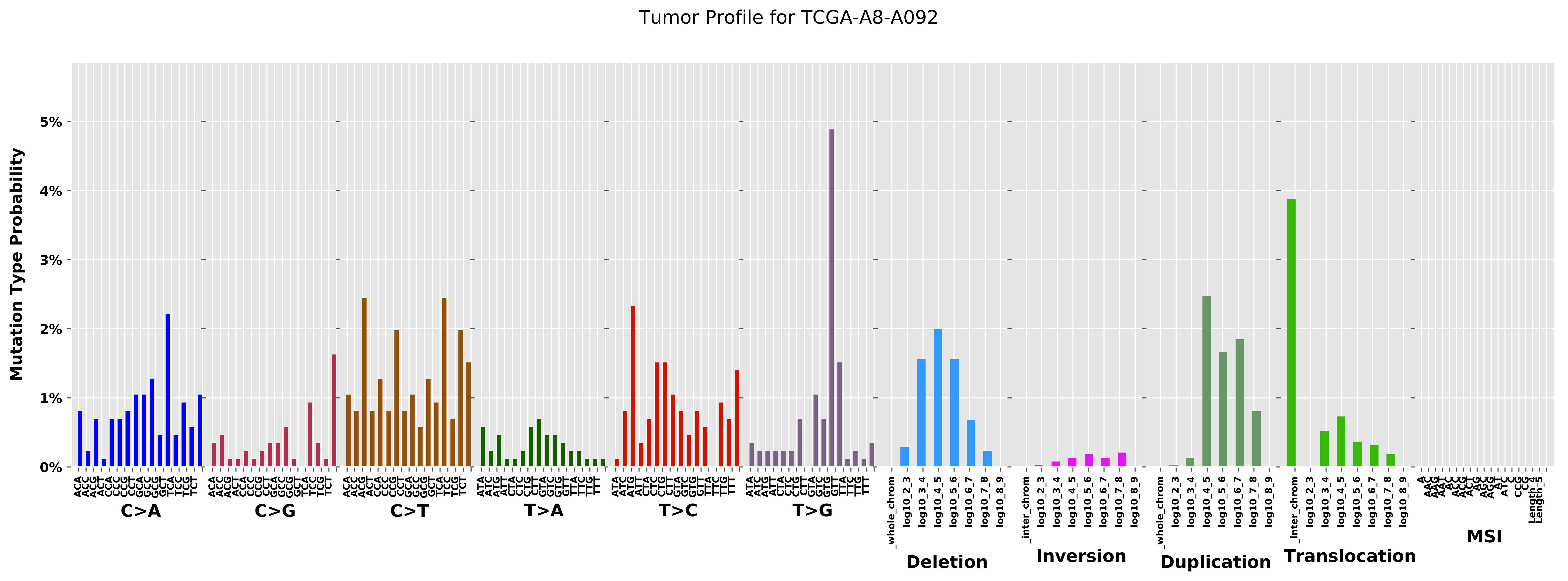
Tumor Profile for TCGA-A8-A092
Mutation Type Probability
0%
1%
2%
3%
4%
5%
C>A
C>G
C>T
T>A
T>C
T>G
Deletion
Inversion
Duplication
Translocation
MSI

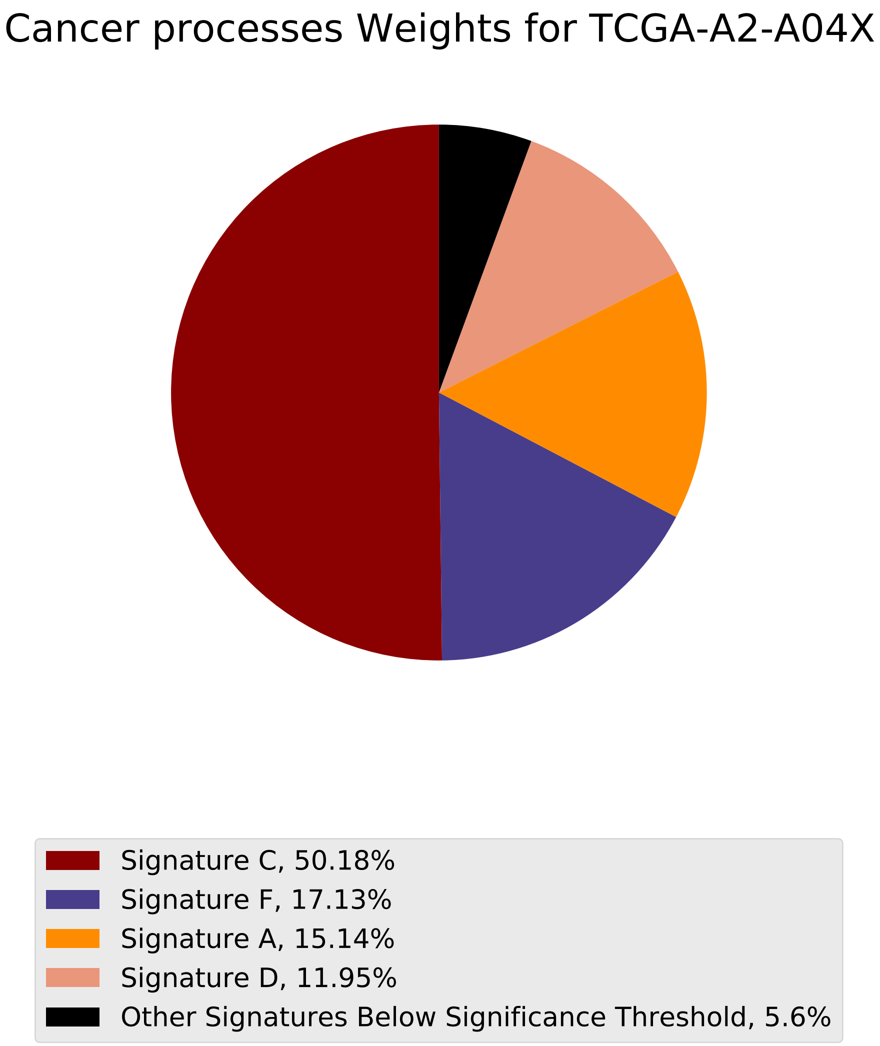
Cancer processes Weights for TCGA-A2-A04X
Signature C, 50.18%
Signature F, 17.13%
Signature A, 15.14%
Signature D, 11.95%
Other Signatures Below Significance Threshold, 5.6%

Tumor Profile for TCGA-A2-A04X

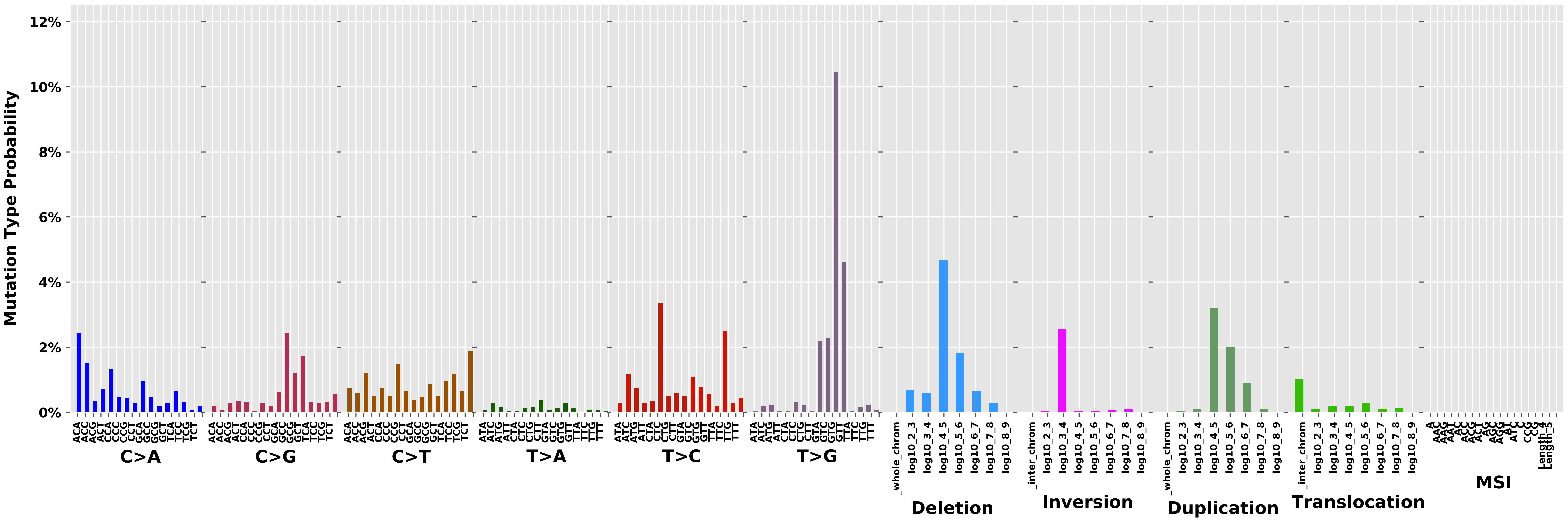

Cancer processes Weights for TCGA-A2-A0D2

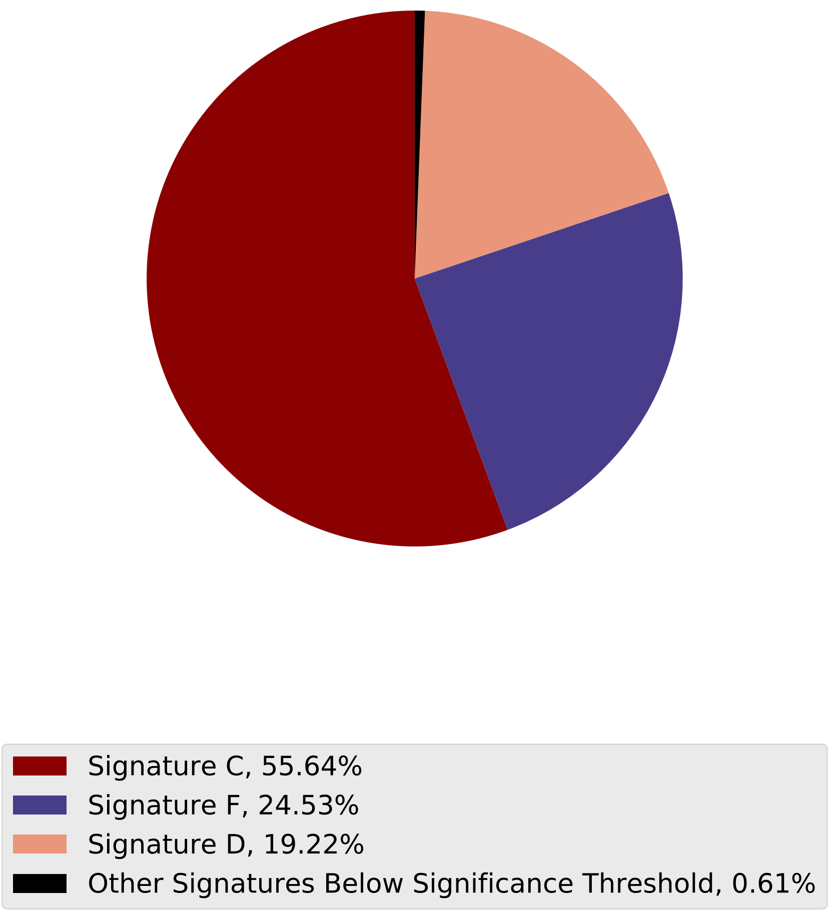

Tumor Profile for TCGA-A2-A0D2

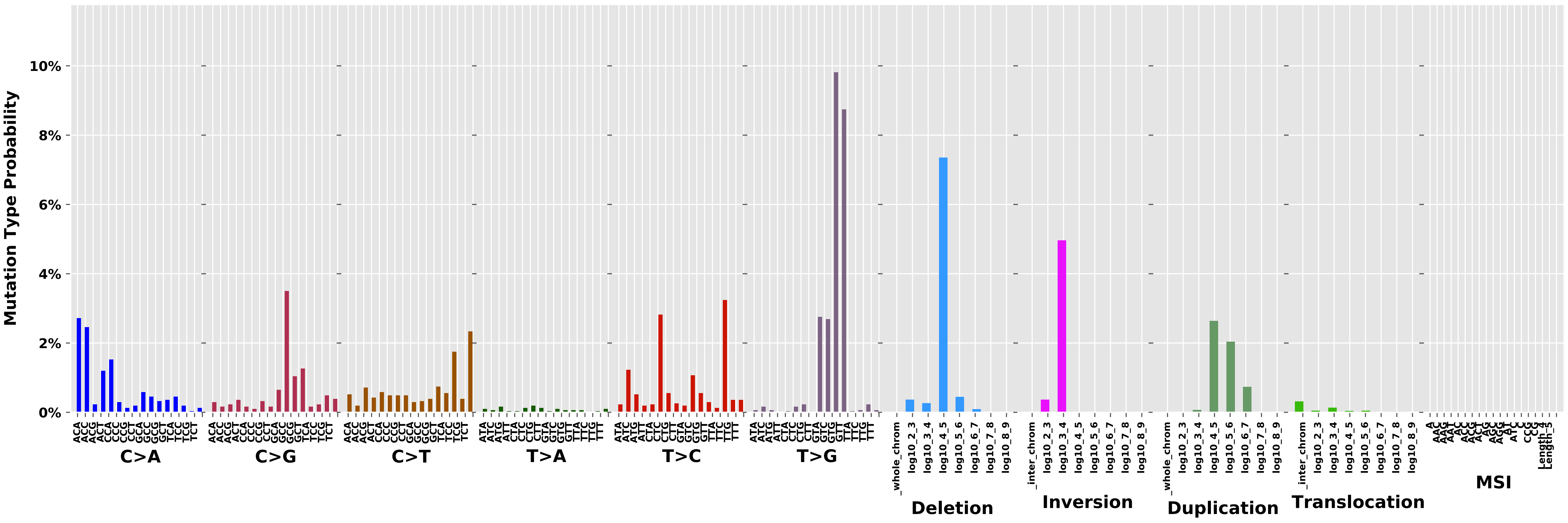

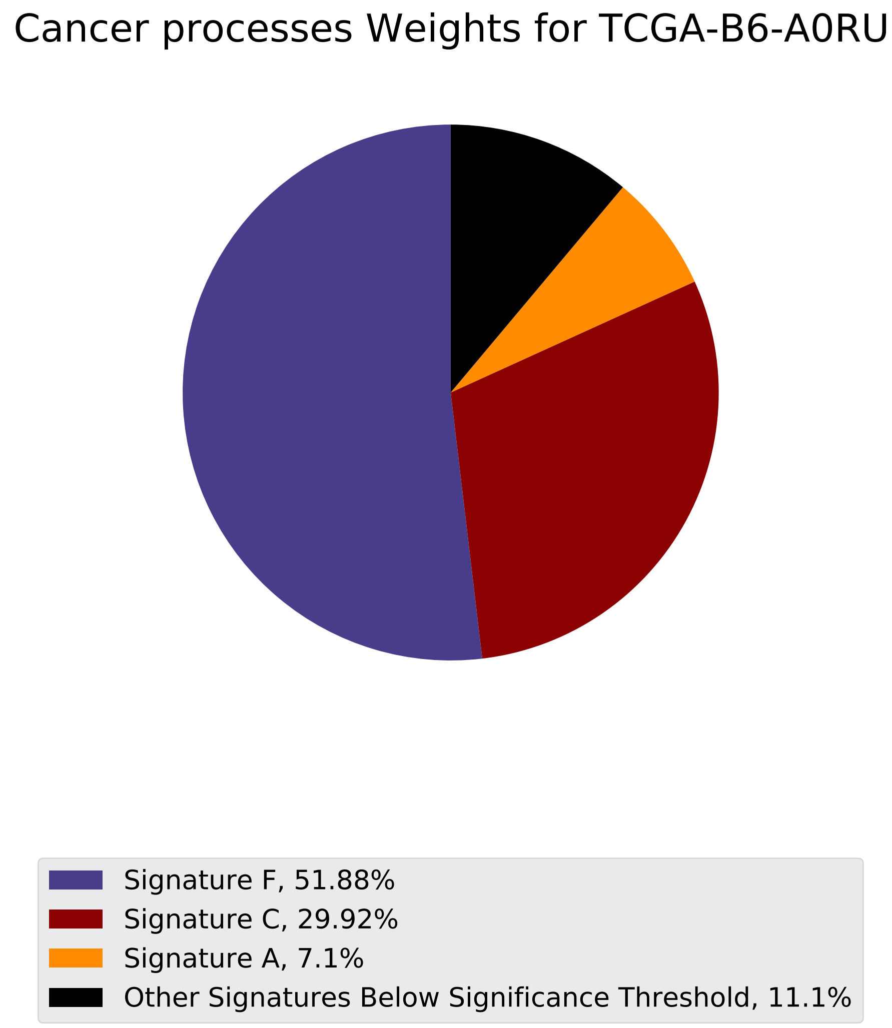
Cancer processes Weights for TCGA-B6-A0RU
Signature F, 51.88%
Signature C, 29.92%
Signature A, 7.1%
Other Signatures Below Significance Threshold, 11.1%

### Tumor Profile for TCGA-B6-A0RU

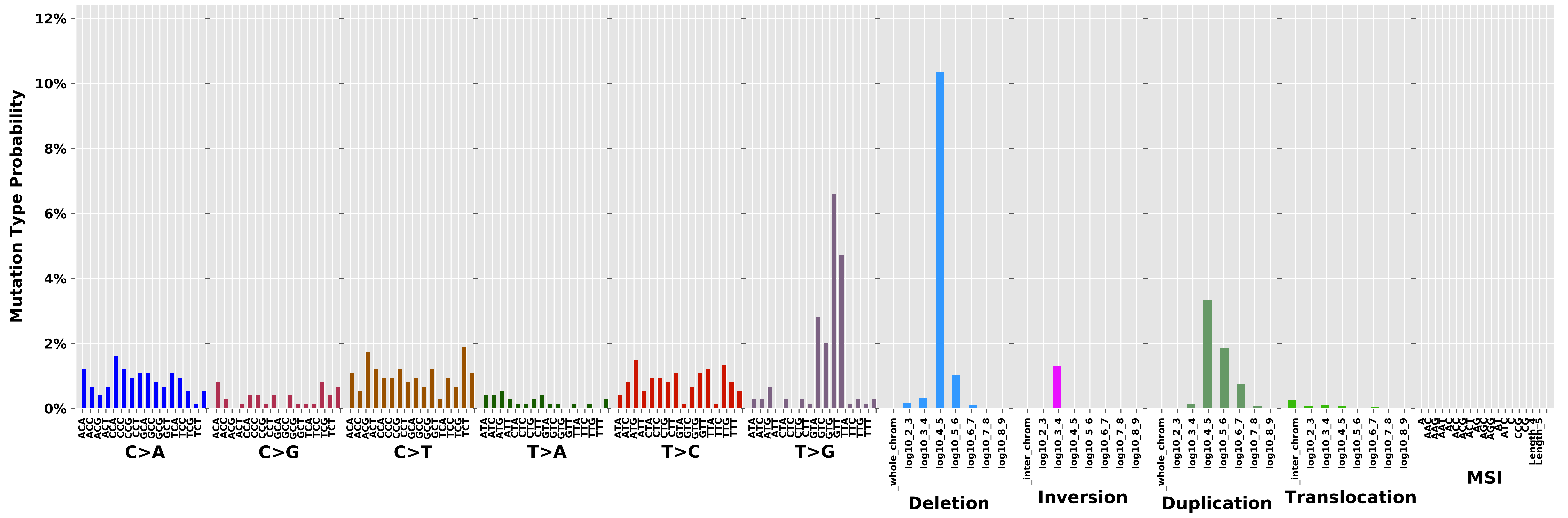
