## Additional File 5 for "pyCancerSig: subclassifying human cancer with comprehensive single nucleotide, structural and microsatellite mutational signature deconstruction from whole genome sequencing"

Cancer processes Weights for TCGA-E2-A14X

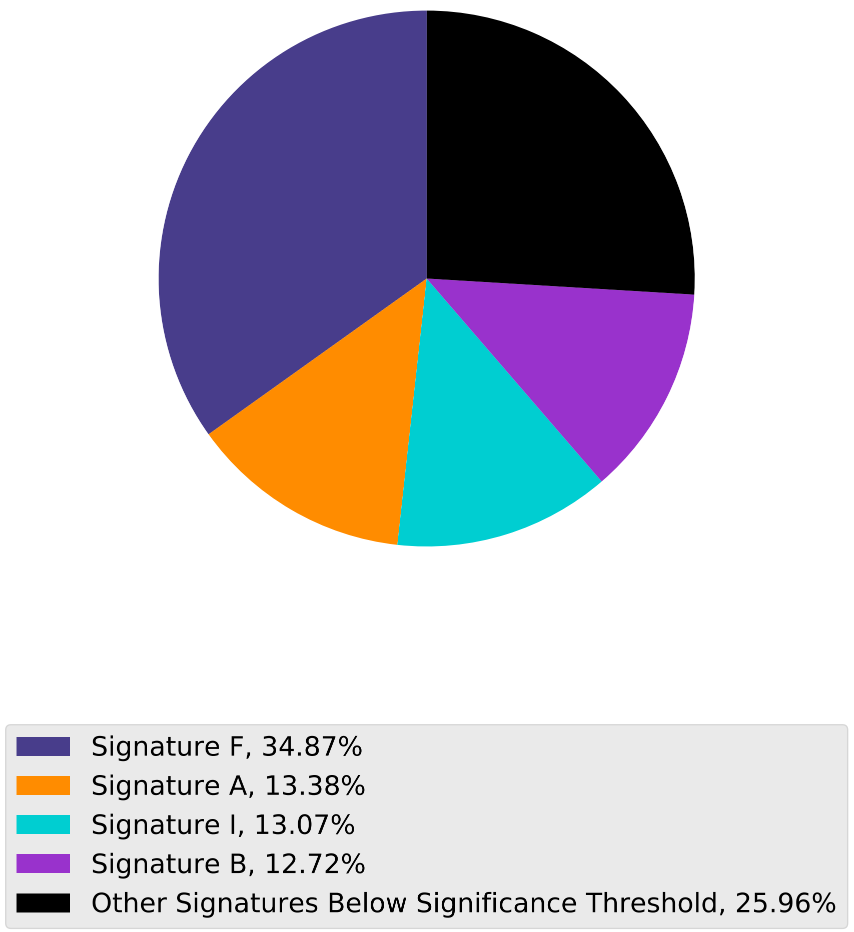

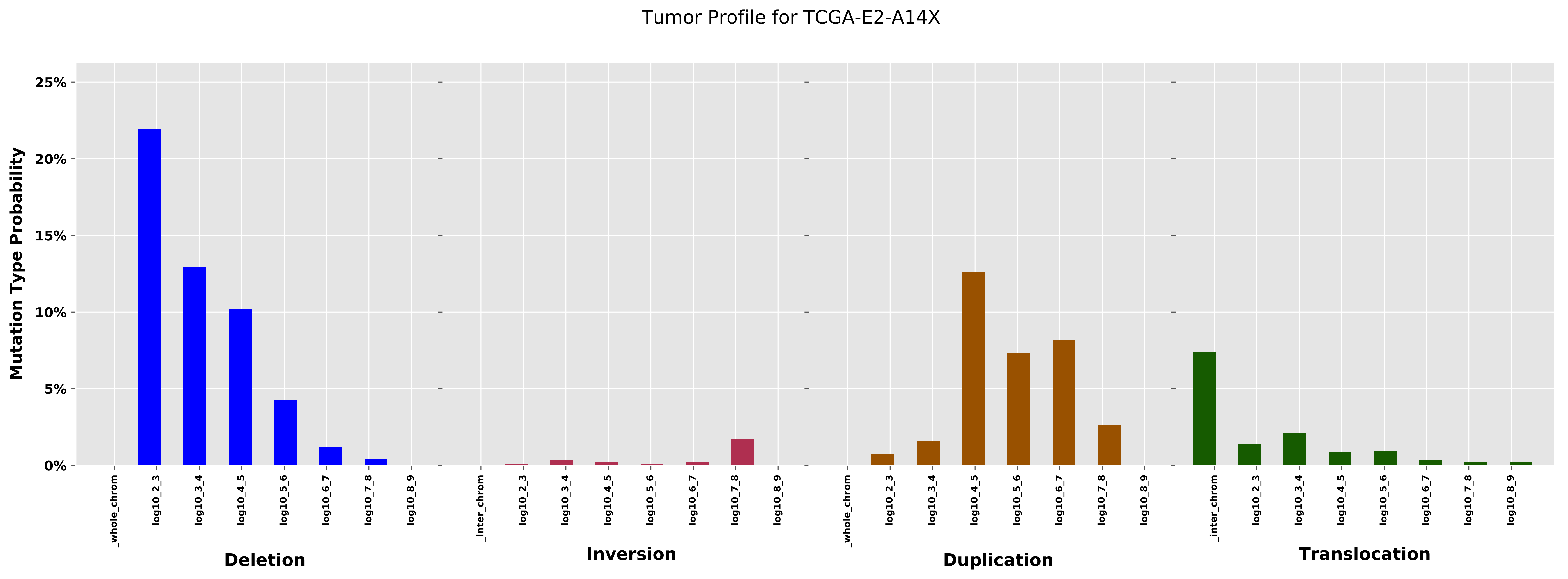
Tumor Profile for TCGA-E2-A14X
Mutation Type Probability
0%
5%
10%
15%
20%
25%
_whole_chrom
log10_2_3
log10_3_4
log10_4_5
log10_5_6
log10_6_7
log10_7_8
log10_8_9
_inter_chrom
Deletion
Inversion
Duplication
Translocation

Cancer processes Weights for TCGA-A8-A08L

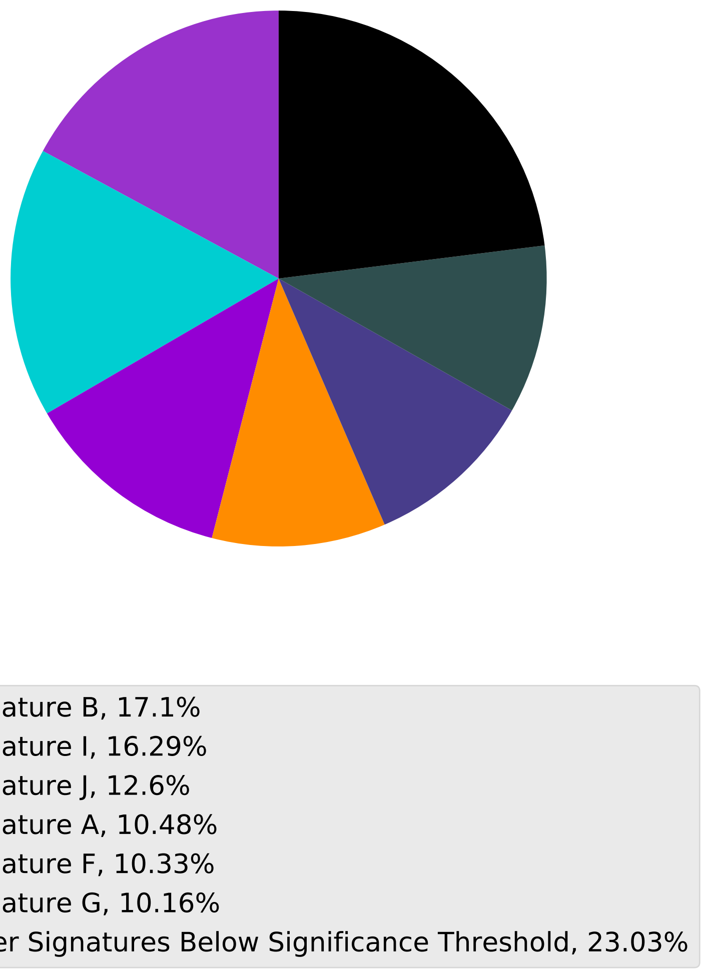

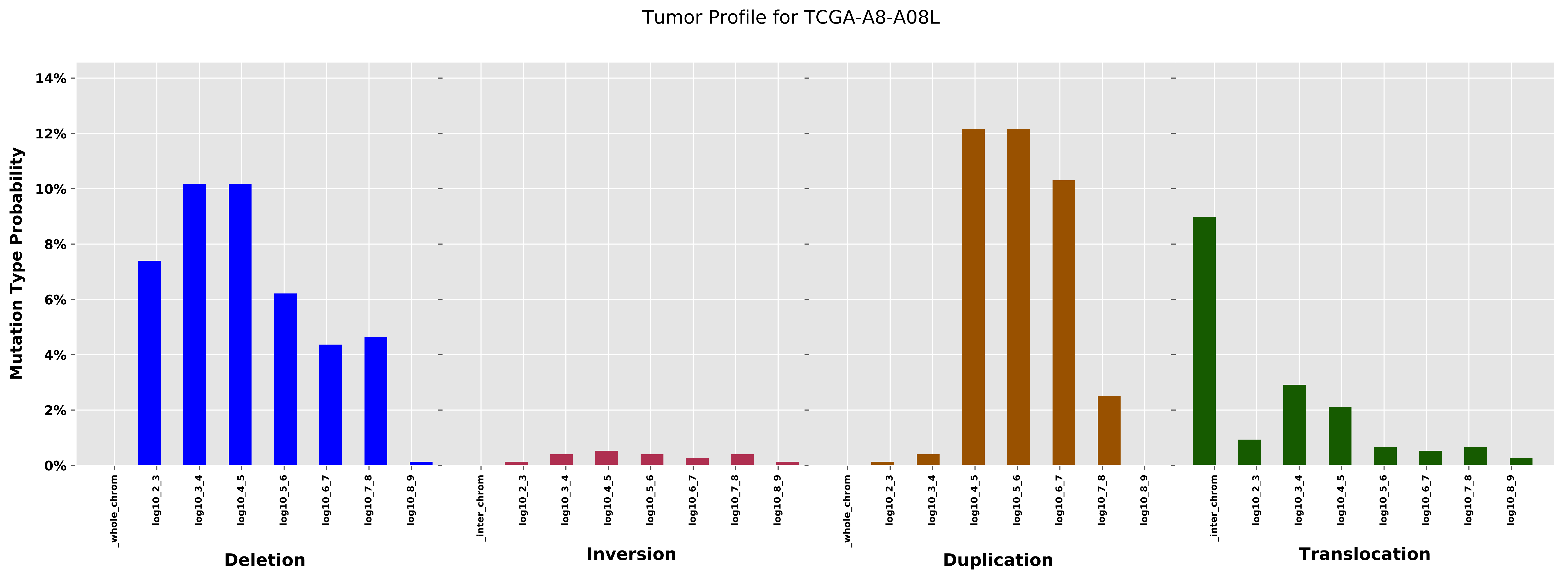

Tumor Profile for TCGA-A8-A08L
Mutation Type Probability
14%
12%
10%
8%
6%
4%
2%
0%
_whole_chrom
log10_2_3
log10_3_4
log10_4_5
log10_5_6
log10_6_7
log10_7_8
log10_8_9
_inter_chrom
Deletion
Inversion
Duplication
Translocation

Cancer processes Weights for TCGA-AO-A12H

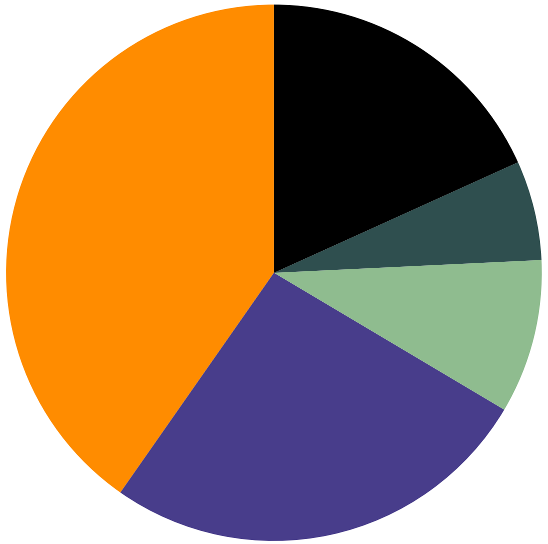

- Signature A, 40.28%
- Signature F, 26.19%
- Signature E, 9.29%
- Signature G, 6.0%
- Other Signatures Below Significance Threshold, 18.24%

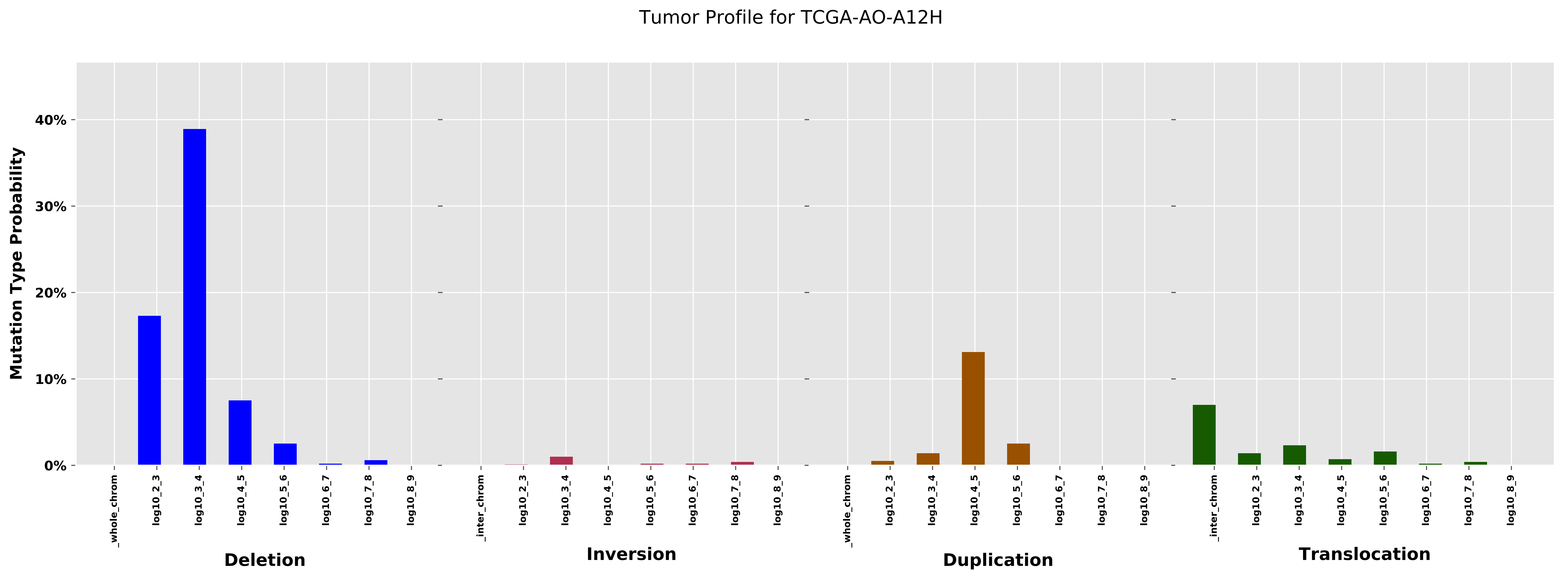
Tumor Profile for TCGA-AO-A12H
Mutation Type Probability
40%
30%
20%
10%
0%
_whole_chrom
log10_2_3
log10_3_4
log10_4_5
log10_5_6
log10_6_7
log10_7_8
log10_8_9
Deletion
_inter_chrom
log10_2_3
log10_3_4
log10_4_5
log10_5_6
log10_6_7
log10_7_8
log10_8_9
Inversion
_whole_chrom
log10_2_3
log10_3_4
log10_4_5
log10_5_6
log10_6_7
log10_7_8
log10_8_9
Duplication
_inter_chrom
log10_2_3
log10_3_4
log10_4_5
log10_5_6
log10_6_7
log10_7_8
log10_8_9
Translocation

#### Cancer processes Weights for TCGA-EW-A3U0

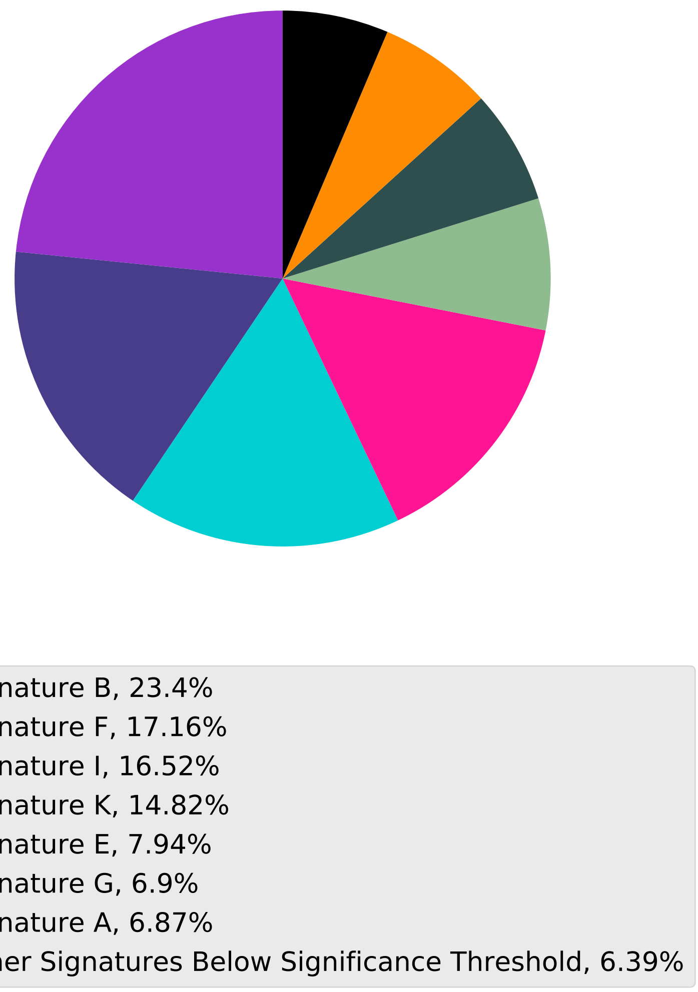

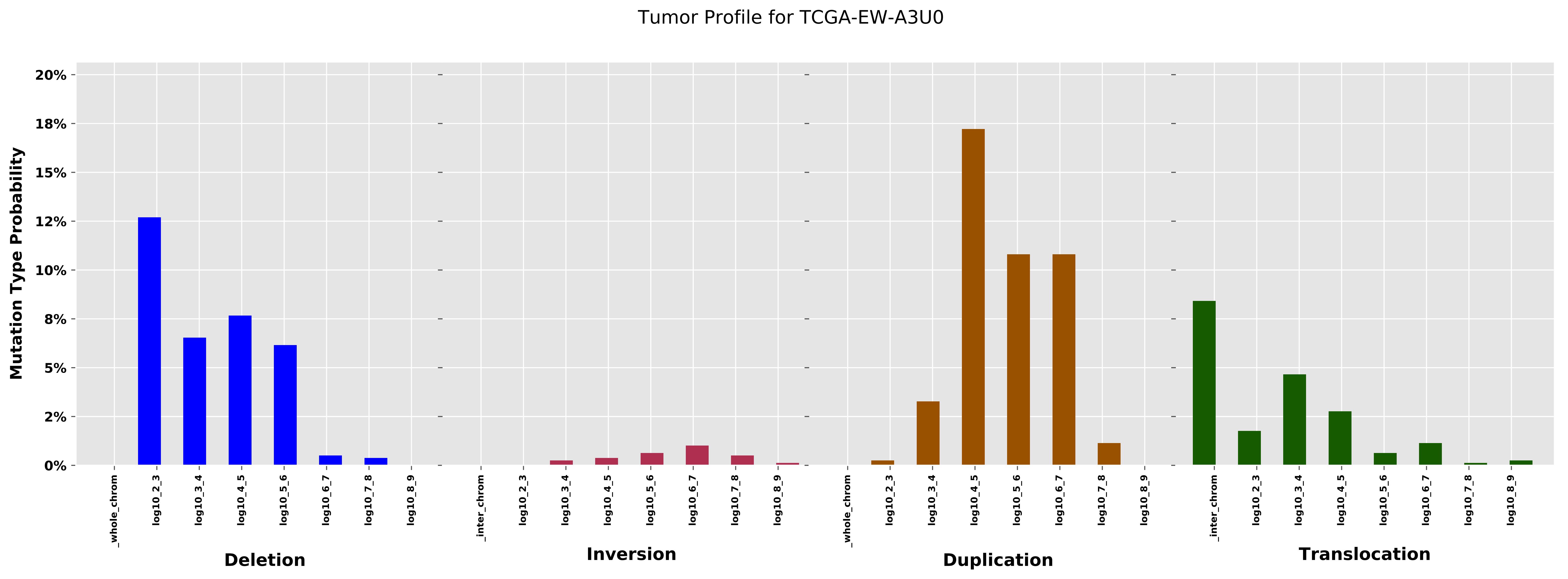
Tumor Profile for TCGA-EW-A3U0
Mutation Type Probability
20%
18%
15%
12%
10%
8%
5%
2%
0%
_whole_chrom
log10_2_3
log10_3_4
log10_4_5
log10_5_6
log10_6_7
log10_7_8
log10_8_9
_inter_chrom
Deletion
Inversion
Duplication
Translocation

Cancer processes Weights for TCGA-BH-A1FC

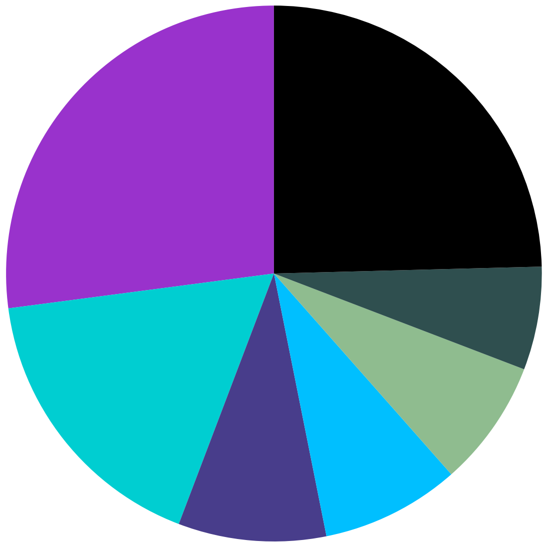

- Signature B, 27.08%
- Signature I, 17.14%
- Signature F, 8.92%
- Signature L, 8.37%
- Signature E, 7.66%
- Signature G, 6.23%
- Other Signatures Below Significance Threshold, 24.59%

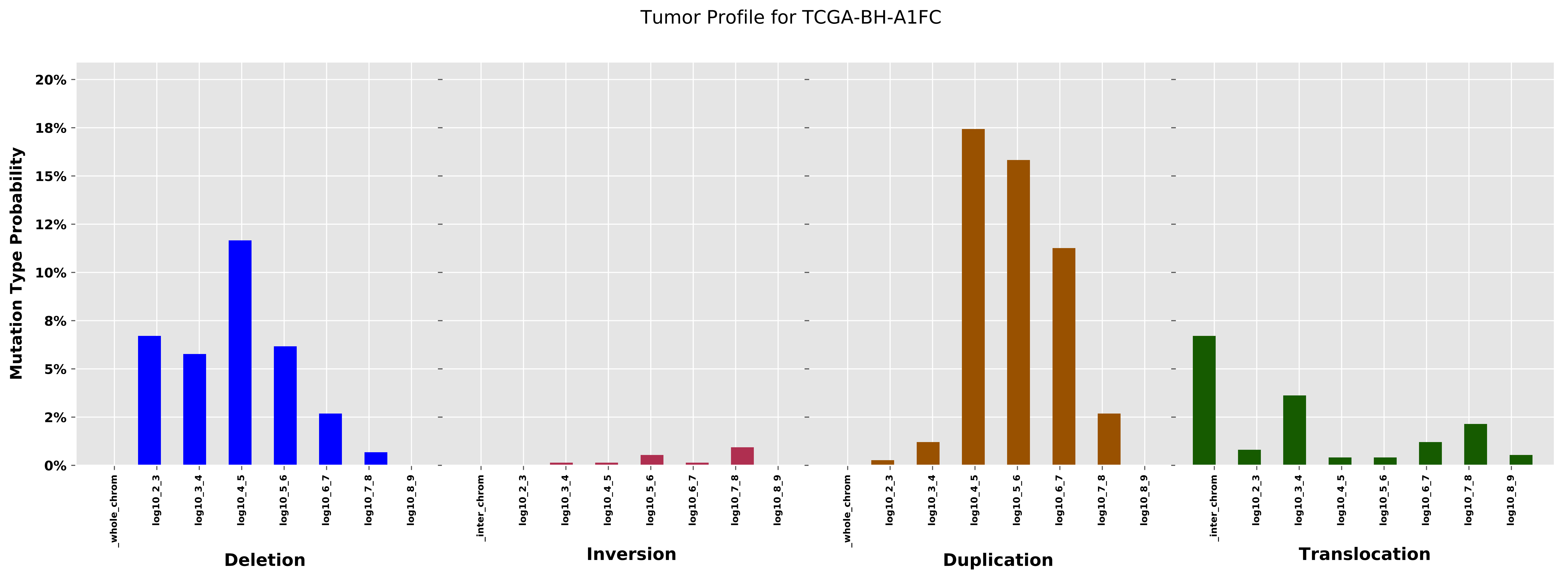

Tumor Profile for TCGA-BH-A1FC
Mutation Type Probability
20%
18%
15%
12%
10%
8%
5%
2%
0%
_whole_chrom
log10_2_3
log10_3_4
log10_4_5
log10_5_6
log10_6_7
log10_7_8
log10_8_9
_inter_chrom
Deletion
Inversion
Duplication
Translocation

Cancer processes Weights for TCGA-AF-2691

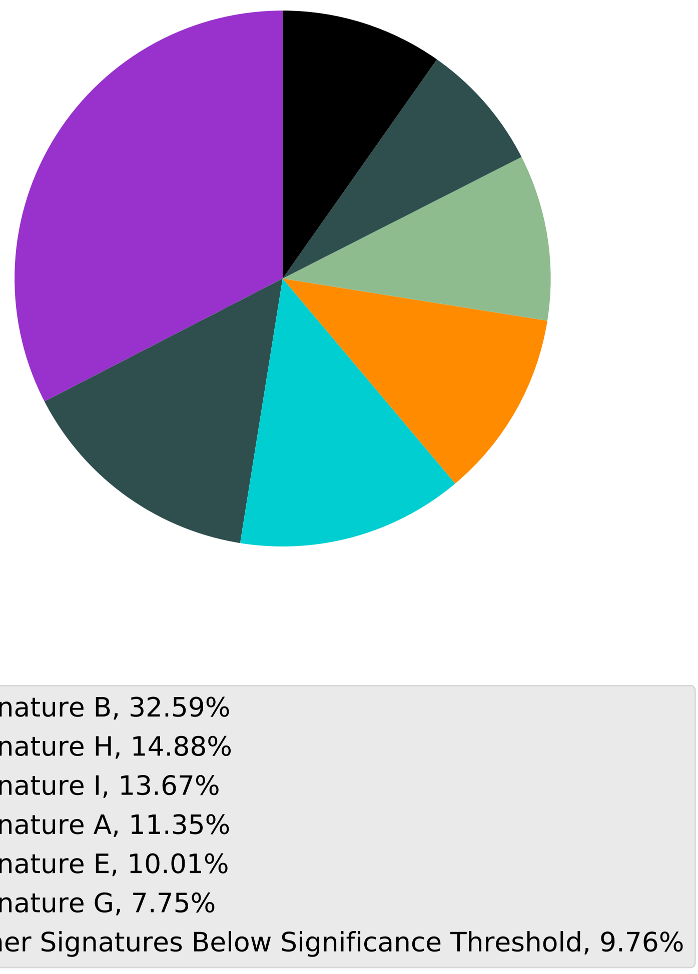

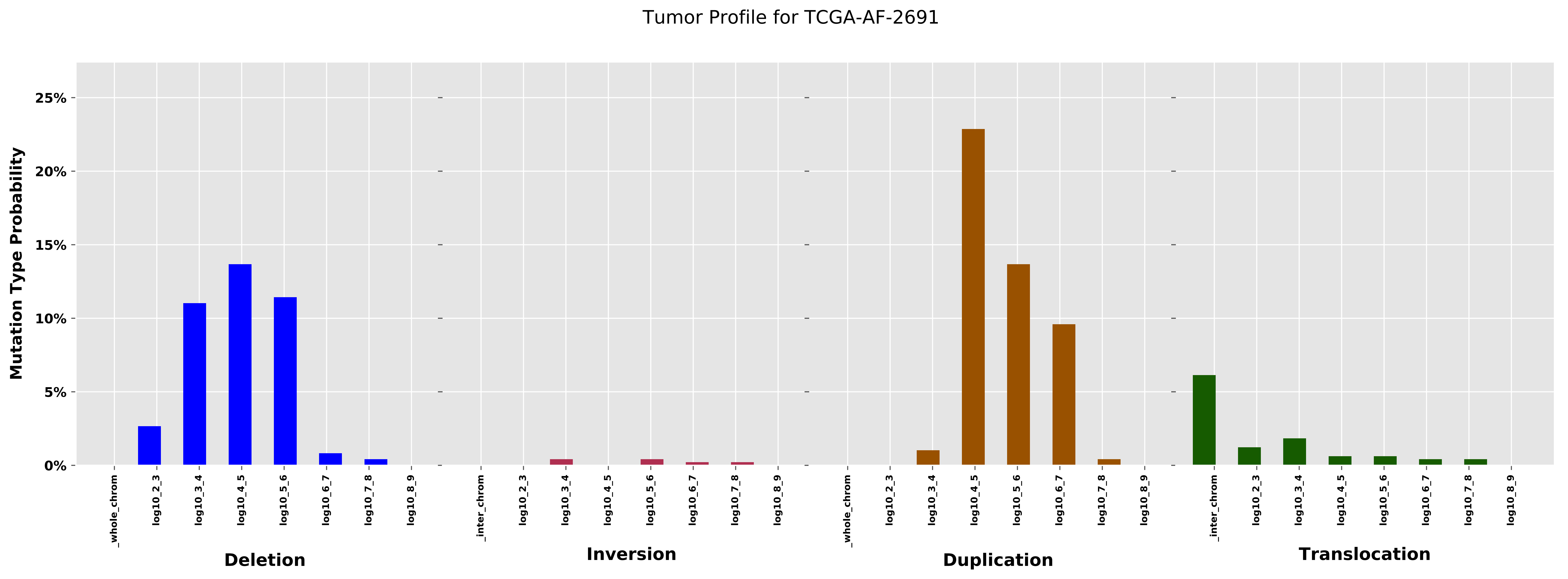

Tumor Profile for TCGA-AF-2691
Mutation Type Probability
25%
20%
15%
10%
5%
0%
_whole_chrom
log10_2_3
log10_3_4
log10_4_5
log10_5_6
log10_6_7
log10_7_8
log10_8_9
Deletion
_inter_chrom
log10_2_3
log10_3_4
log10_4_5
log10_5_6
log10_6_7
log10_7_8
log10_8_9
Inversion
_whole_chrom
log10_2_3
log10_3_4
log10_4_5
log10_5_6
log10_6_7
log10_7_8
log10_8_9
Duplication
_inter_chrom
log10_2_3
log10_3_4
log10_4_5
log10_5_6
log10_6_7
log10_7_8
log10_8_9
Translocation

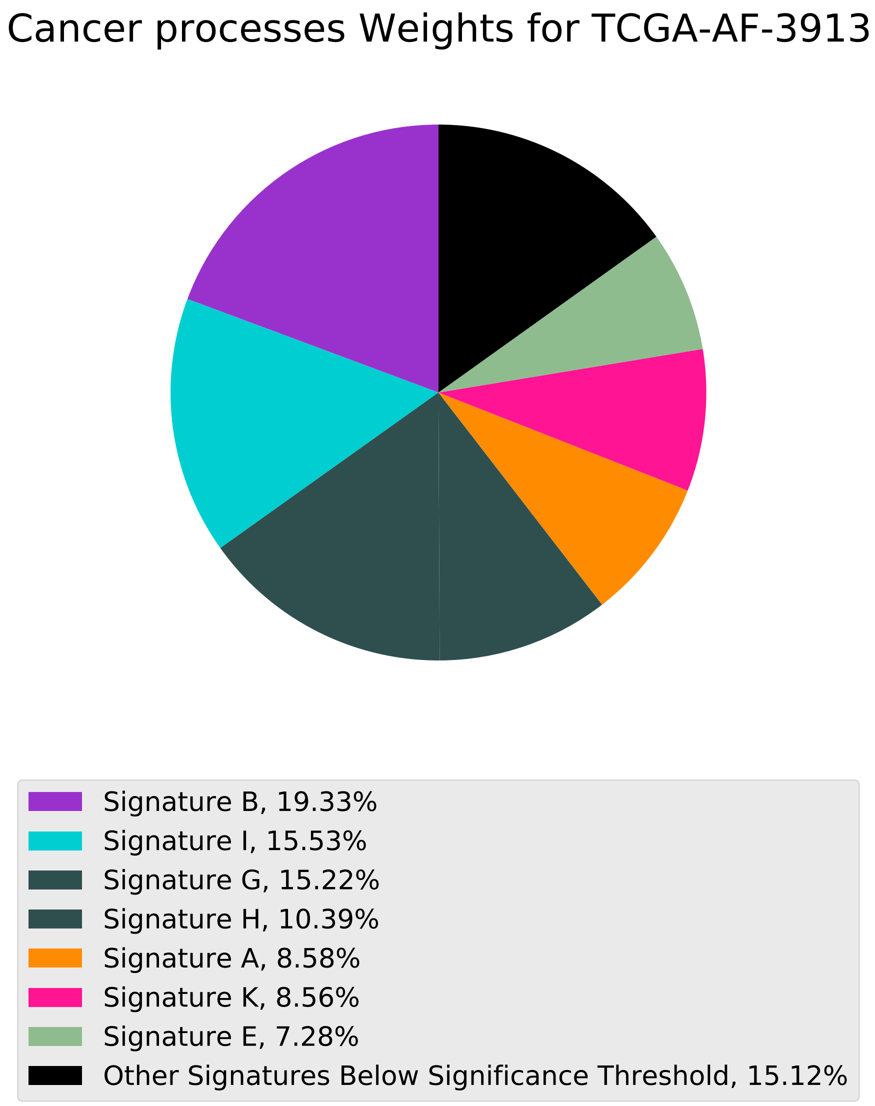
Cancer processes Weights for TCGA-AF-3913
Signature B, 19.33%
Signature I, 15.53%
Signature G, 15.22%
Signature H, 10.39%
Signature A, 8.58%
Signature K, 8.56%
Signature E, 7.28%
Other Signatures Below Significance Threshold, 15.12%

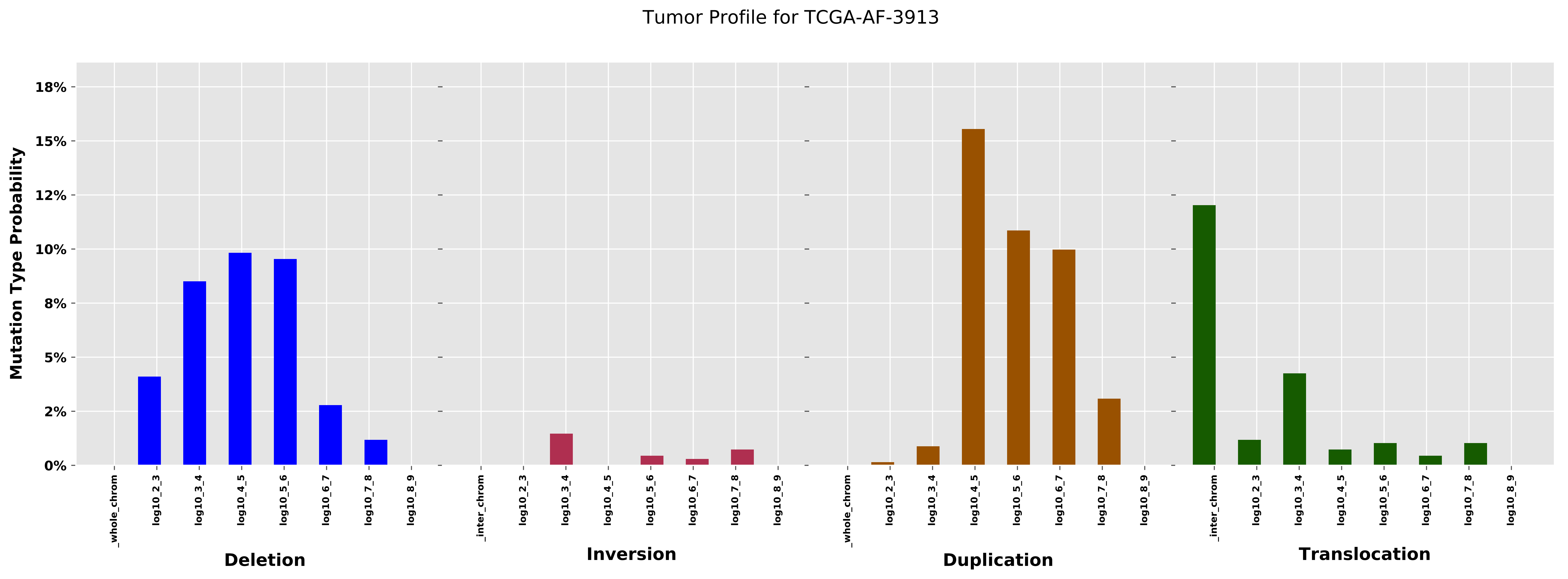
Tumor Profile for TCGA-AF-3913
Mutation Type Probability
18%
15%
12%
10%
8%
5%
2%
0%
_whole_chrom
_inter_chrom
log10_2_3
log10_3_4
log10_4_5
log10_5_6
log10_6_7
log10_7_8
log10_8_9
Deletion
Inversion
Duplication
Translocation

#### Cancer processes Weights for TCGA-AA-3977

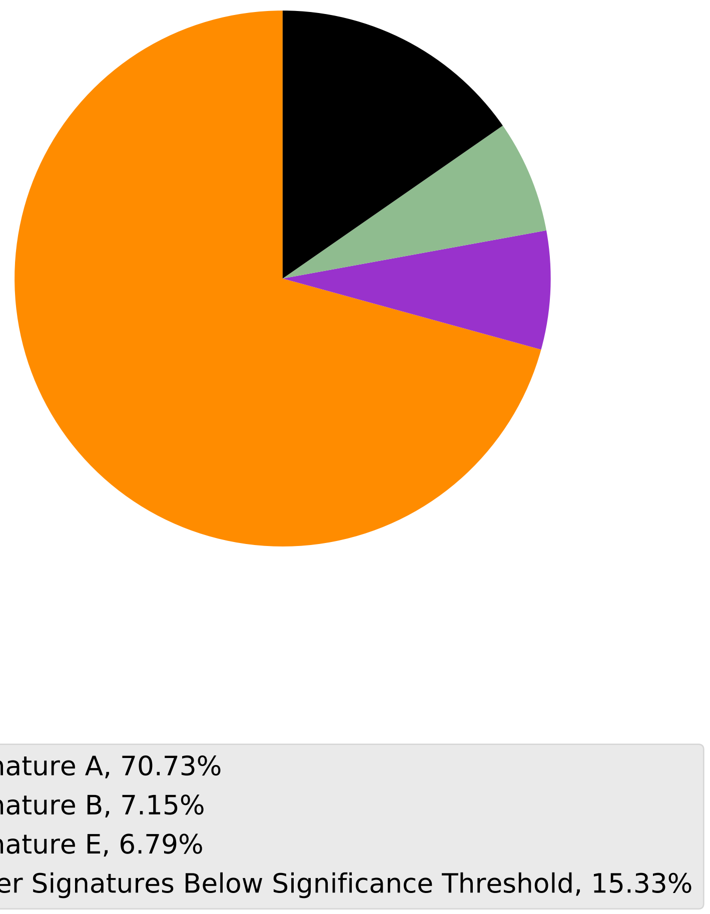

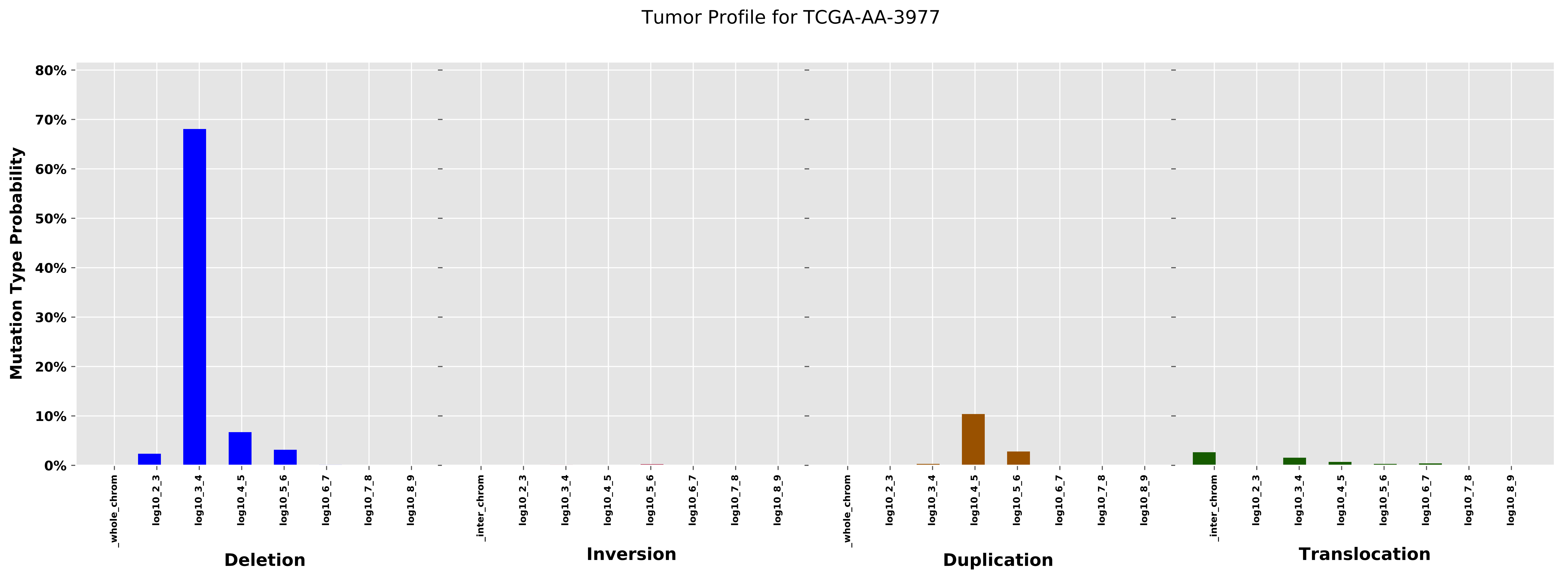
Tumor Profile for TCGA-AA-3977
Mutation Type Probability
80%
70%
60%
50%
40%
30%
20%
10%
0%
_whole_chrom
log10_2_3
log10_3_4
log10_4_5
log10_5_6
log10_6_7
log10_7_8
log10_8_9
_inter_chrom
Deletion
Inversion
Duplication
Translocation

Cancer processes Weights for TCGA-A7-A0CE

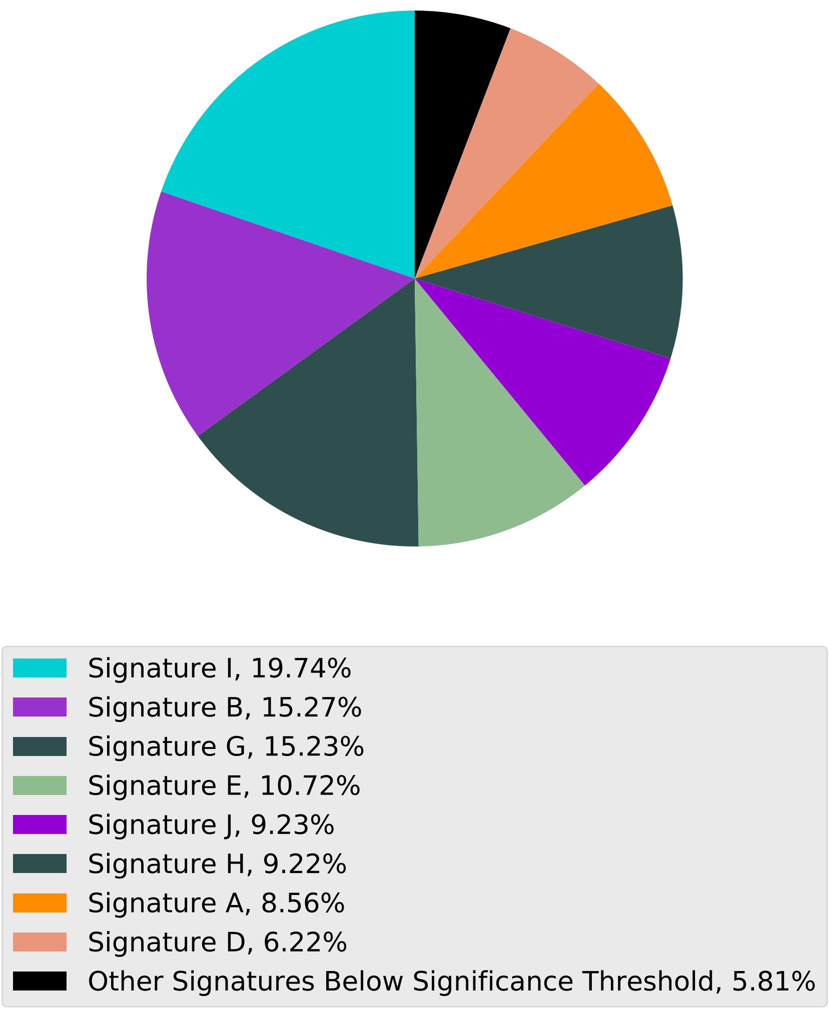

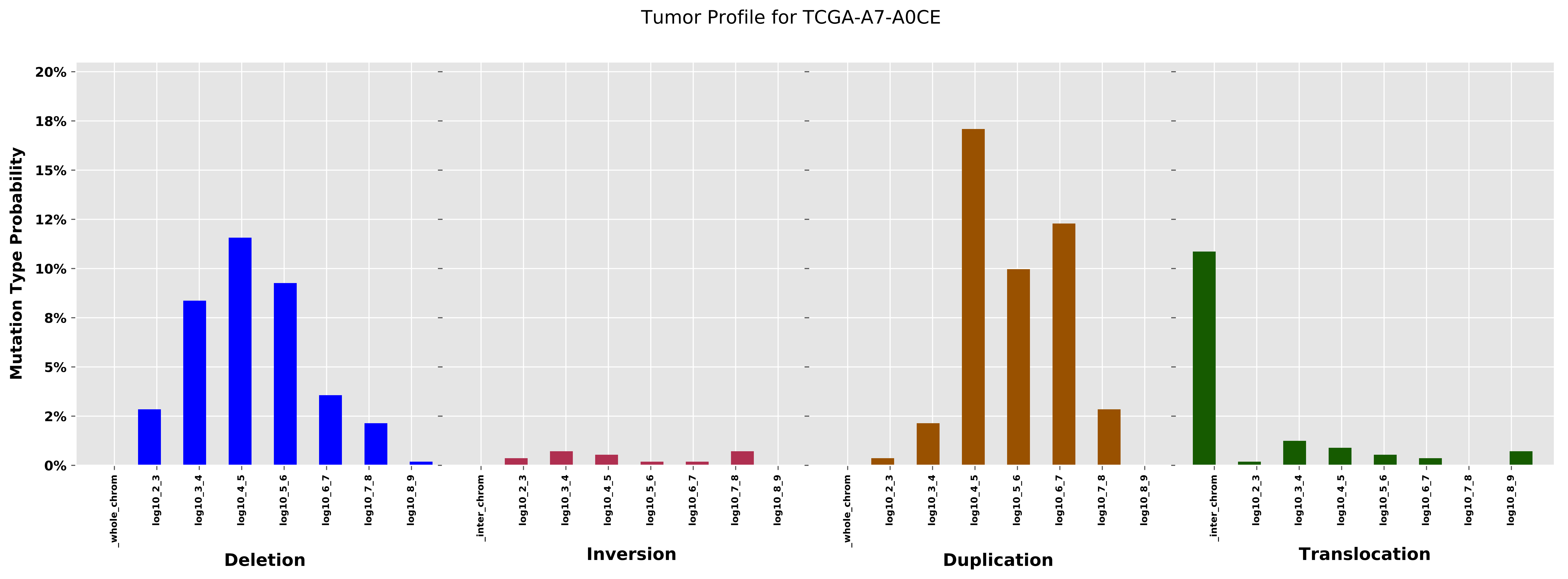

Tumor Profile for TCGA-A7-A0CE
Mutation Type Probability
20%
18%
15%
12%
10%
8%
5%
2%
0%
_whole_chrom
log10_2_3
log10_3_4
log10_4_5
log10_5_6
log10_6_7
log10_7_8
log10_8_9
_inter_chrom
Deletion
Inversion
Duplication
Translocation

Cancer processes Weights for TCGA-AO-A0J4

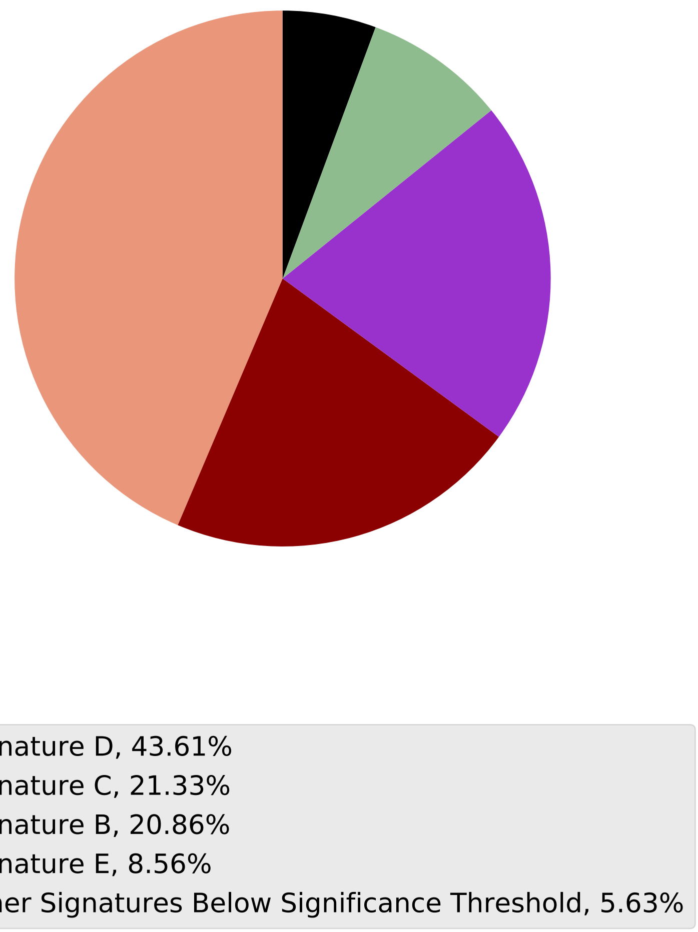

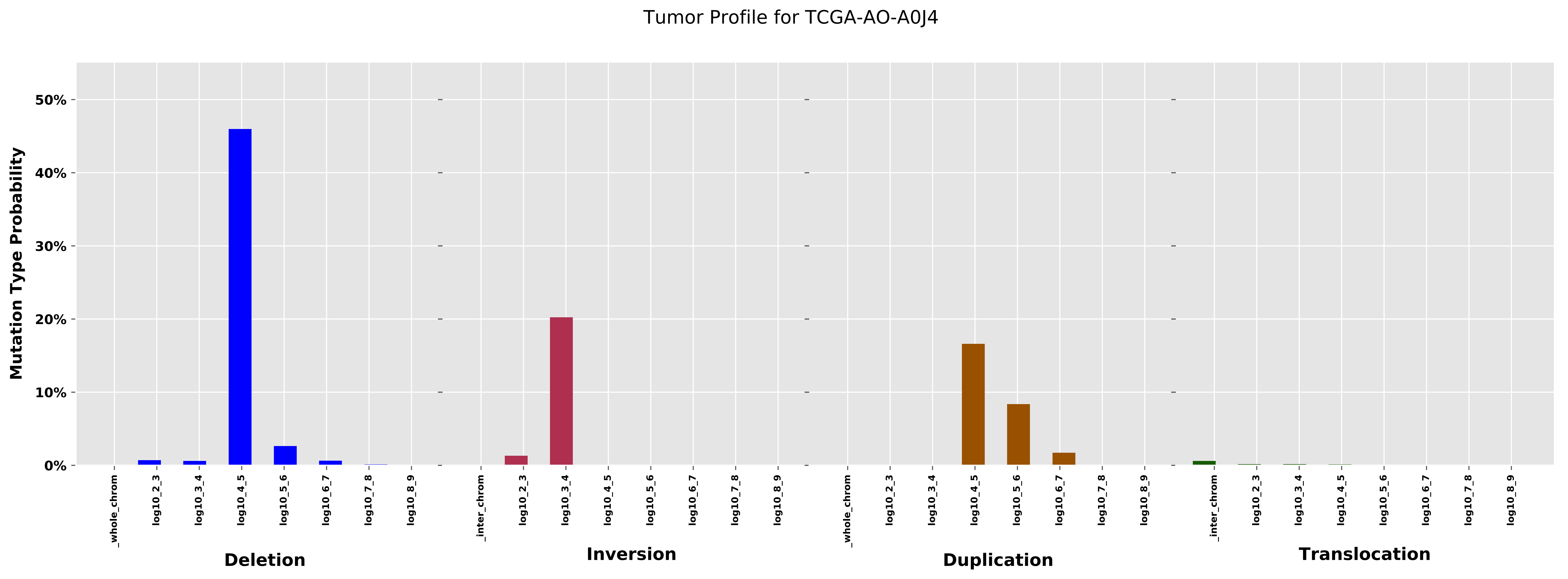
Tumor Profile for TCGA-AO-A0J4
Mutation Type Probability
50%
40%
30%
20%
10%
0%
_whole_chrom
log10_2_3
log10_3_4
log10_4_5
log10_5_6
log10_6_7
log10_7_8
log10_8_9
_inter_chrom
Deletion
Inversion
Duplication
Translocation

Cancer processes Weights for TCGA-AQ-A04J

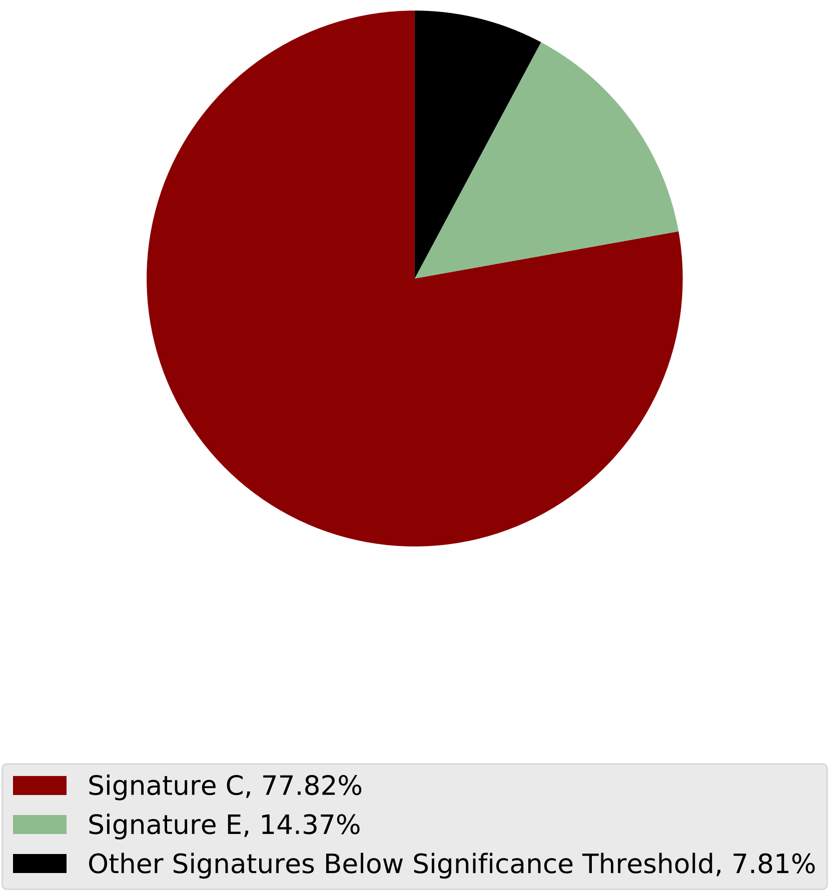

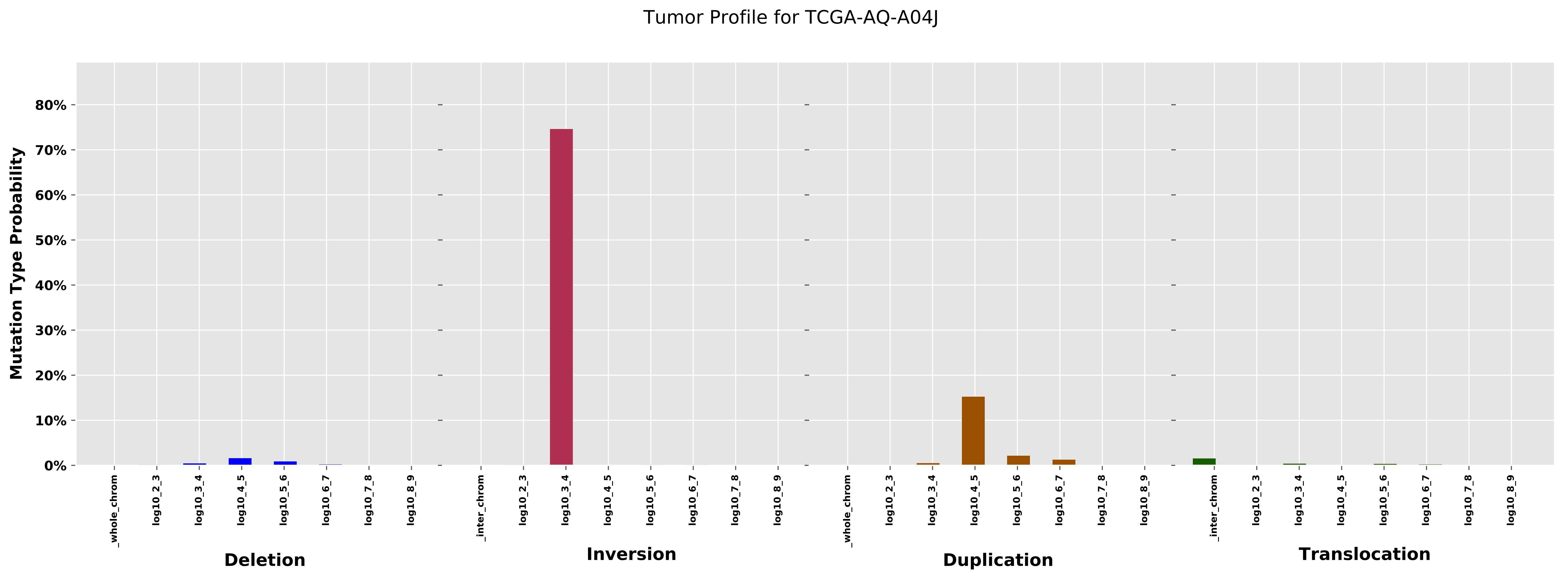

Tumor Profile for TCGA-AQ-A04J
Mutation Type Probability
80%
70%
60%
50%
40%
30%
20%
10%
0%
_whole_chrom
log10_2_3
log10_3_4
log10_4_5
log10_5_6
log10_6_7
log10_7_8
log10_8_9
_inter_chrom
Deletion
Inversion
Duplication
Translocation

#### Cancer processes Weights for TCGA-AG-4007

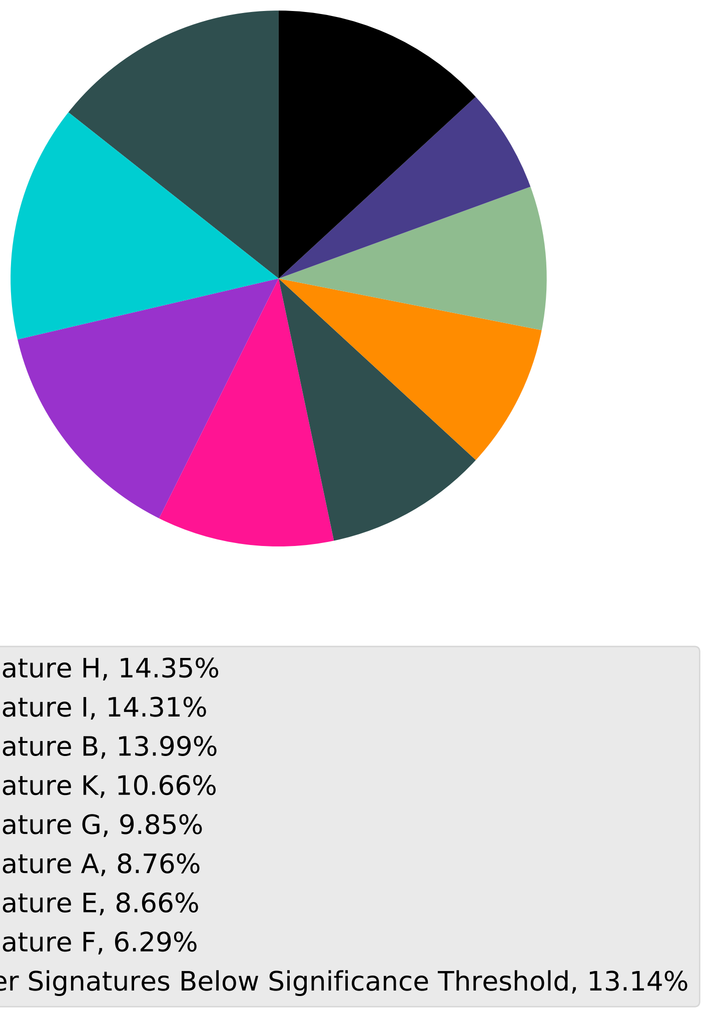

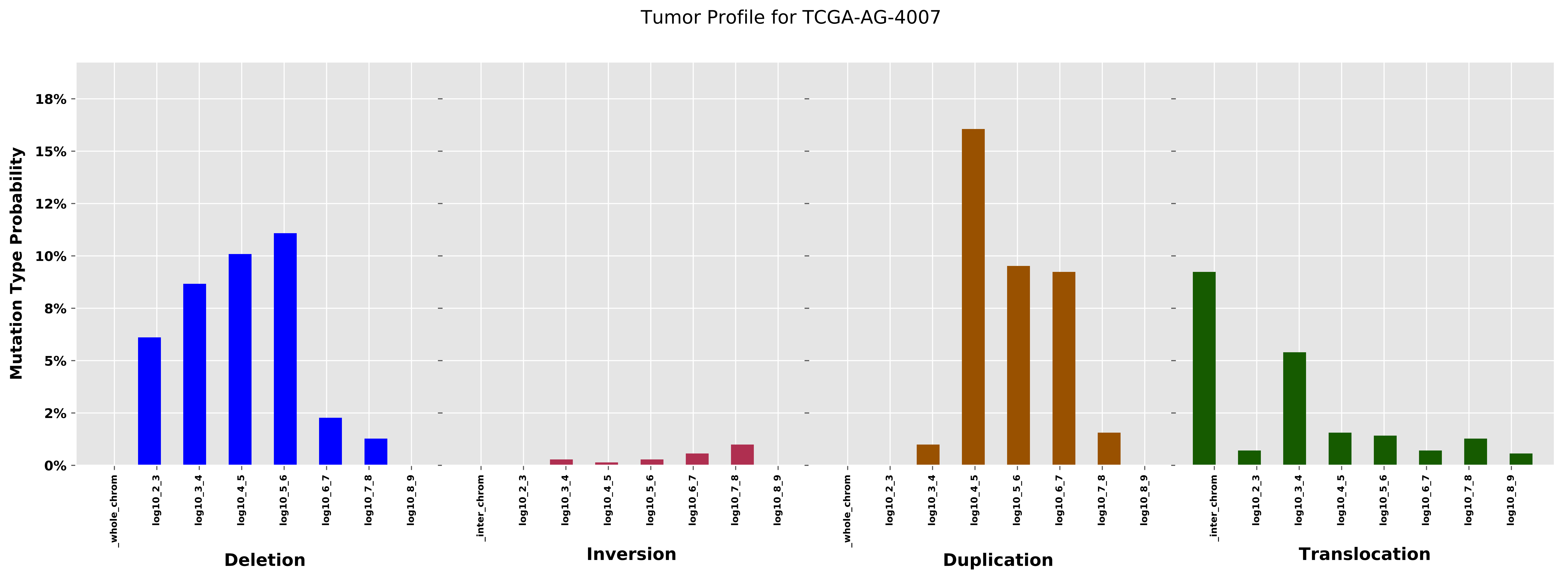

Tumor Profile for TCGA-AG-4007
Mutation Type Probability
18%
15%
12%
10%
8%
5%
2%
0%
_whole_chrom
log10_2_3
log10_3_4
log10_4_5
log10_5_6
log10_6_7
log10_7_8
log10_8_9
_inter_chrom
Deletion
Inversion
Duplication
Translocation

Cancer processes Weights for TCGA-CA-6717

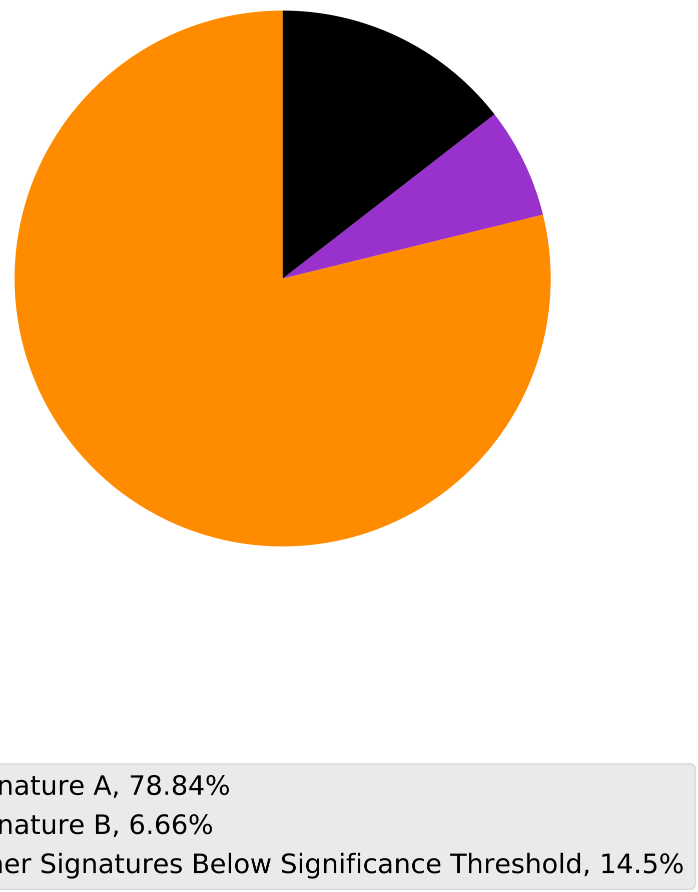

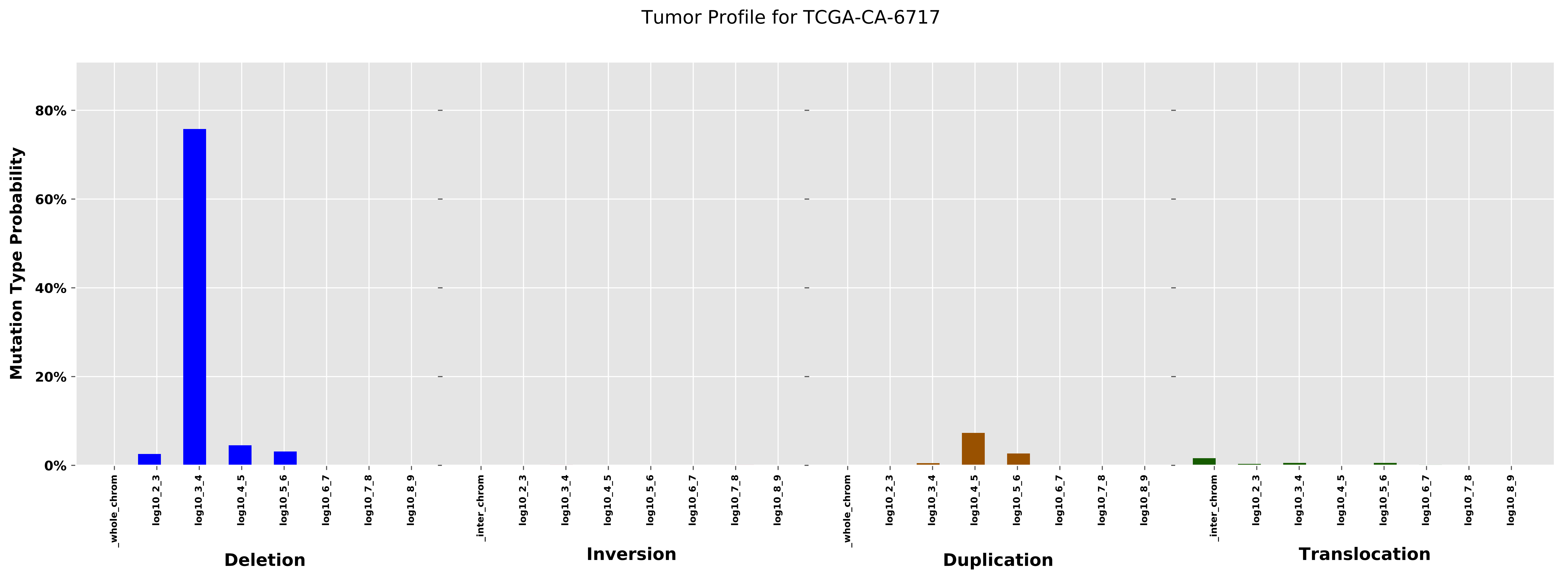
Tumor Profile for TCGA-CA-6717
Mutation Type Probability
80%
60%
40%
20%
0%
_whole_chrom
log10_2_3
log10_3_4
log10_4_5
log10_5_6
log10_6_7
log10_7_8
log10_8_9
Deletion
_inter_chrom
log10_2_3
log10_3_4
log10_4_5
log10_5_6
log10_6_7
log10_7_8
log10_8_9
Inversion
_whole_chrom
log10_2_3
log10_3_4
log10_4_5
log10_5_6
log10_6_7
log10_7_8
log10_8_9
Duplication
_inter_chrom
log10_2_3
log10_3_4
log10_4_5
log10_5_6
log10_6_7
log10_7_8
log10_8_9
Translocation

#### Cancer processes Weights for TCGA-E2-A14P

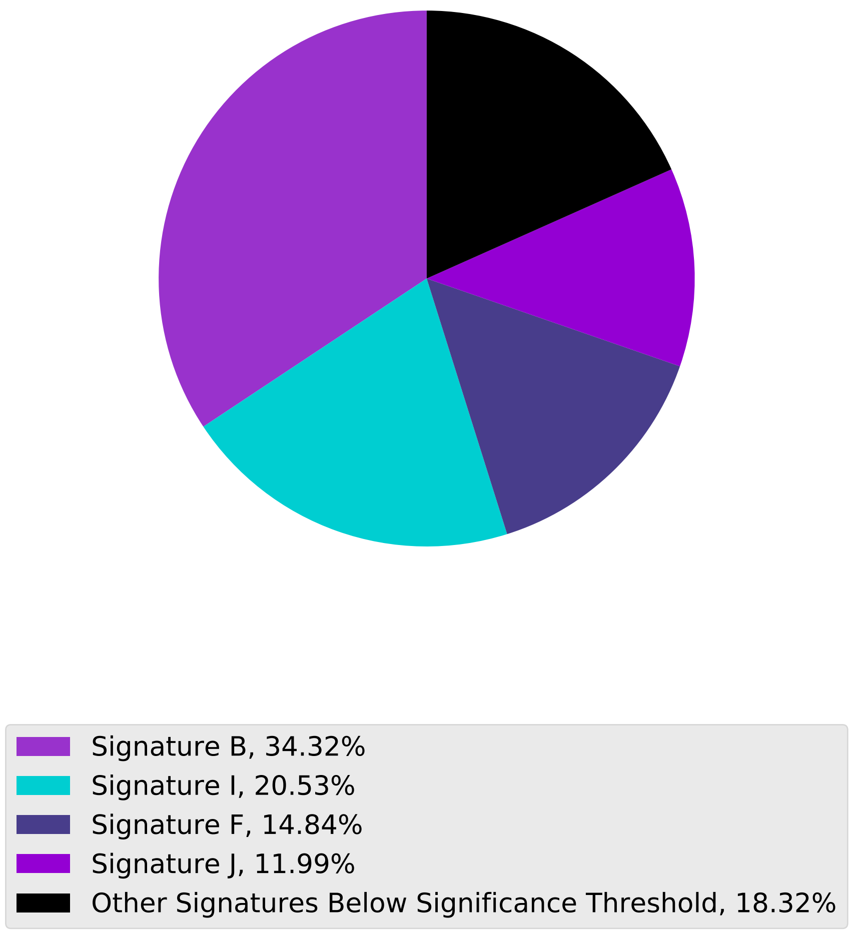

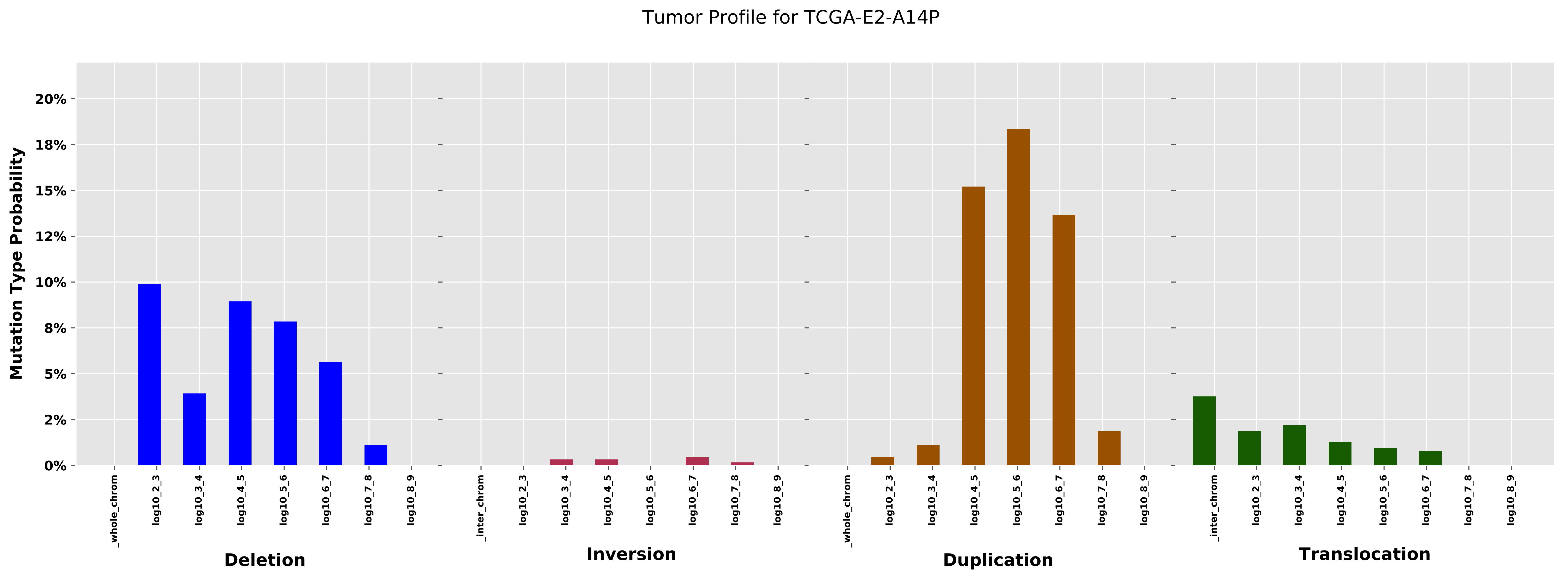
Tumor Profile for TCGA-E2-A14P
Mutation Type Probability
20%
18%
15%
12%
10%
8%
5%
2%
0%
_whole_chrom
log10_2_3
log10_3_4
log10_4_5
log10_5_6
log10_6_7
log10_7_8
log10_8_9
_inter_chrom
Deletion
Inversion
Duplication
Translocation

Cancer processes Weights for TCGA-AZ-6601

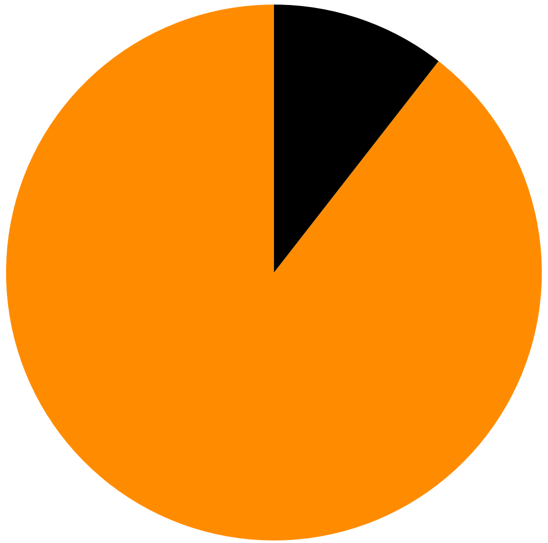

Signature A, 89.47%
Other Signatures Below Significance Threshold, 10.53%

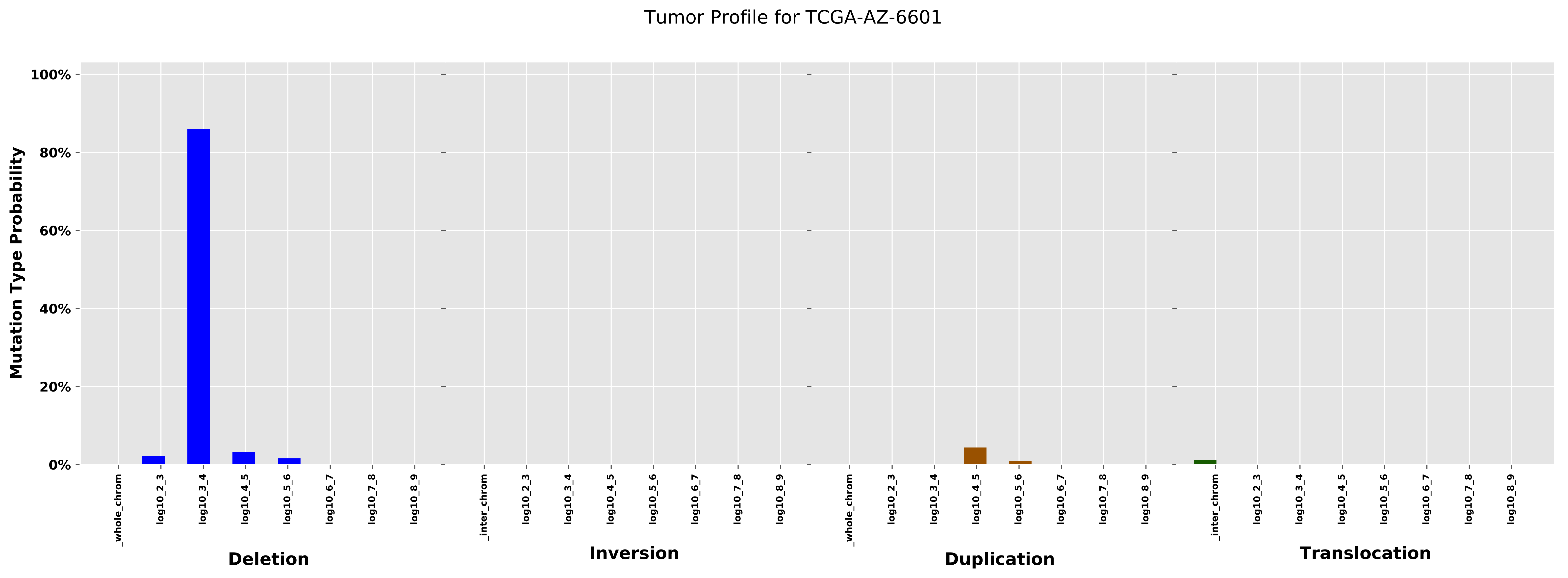
Tumor Profile for TCGA-AZ-6601
Mutation Type Probability
100%
80%
60%
40%
20%
0%
_whole_chrom
log10_2_3
log10_3_4
log10_4_5
log10_5_6
log10_6_7
log10_7_8
log10_8_9
_inter_chrom
Deletion
Inversion
Duplication
Translocation

Cancer processes Weights for TCGA-A6-6781

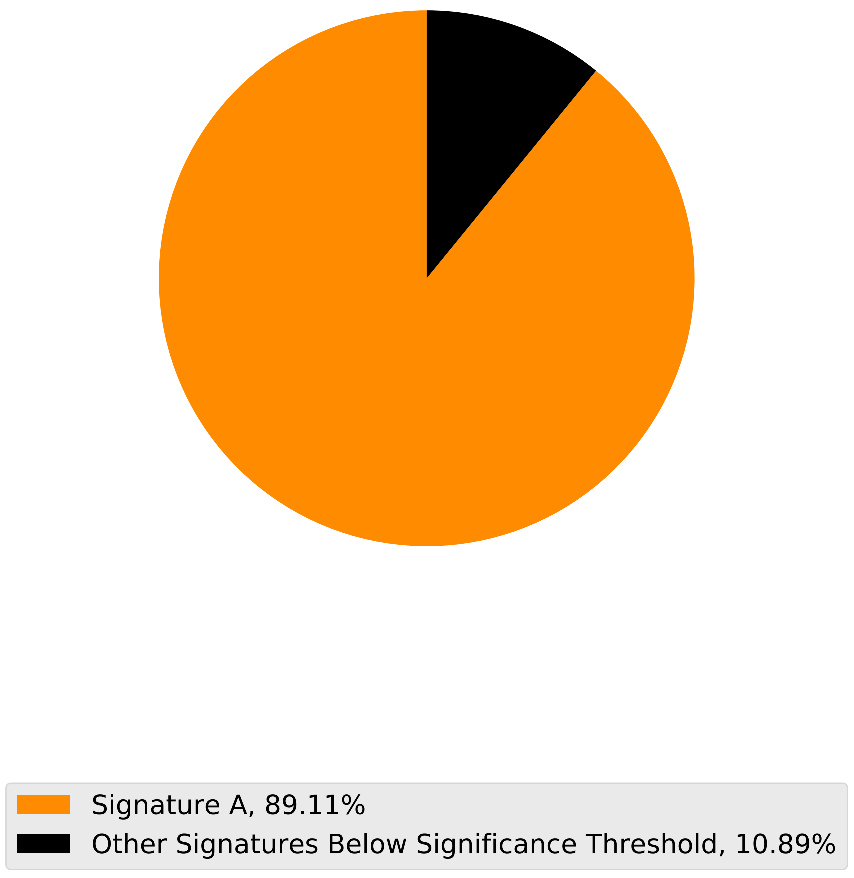

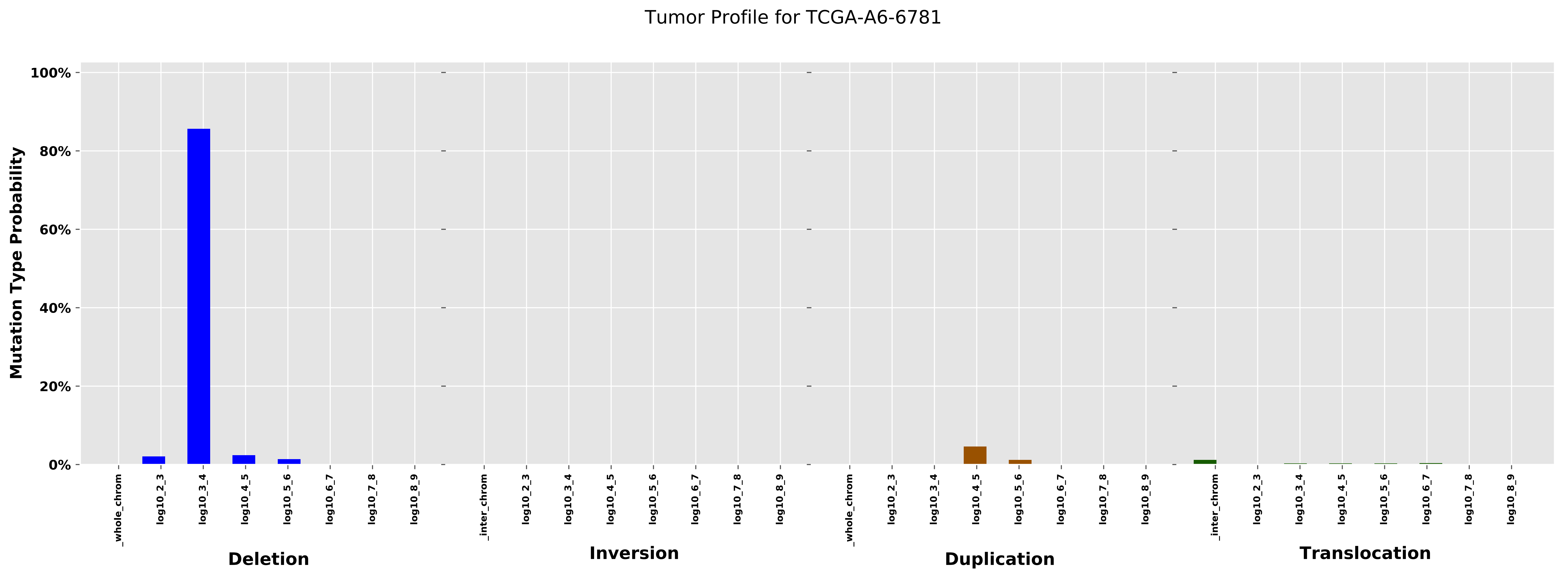
Tumor Profile for TCGA-A6-6781
Mutation Type Probability
100%
80%
60%
40%
20%
0%
_whole_chrom
log10_2_3
log10_3_4
log10_4_5
log10_5_6
log10_6_7
log10_7_8
log10_8_9
_inter_chrom
Deletion
Inversion
Duplication
Translocation

Cancer processes Weights for TCGA-GM-A2DF

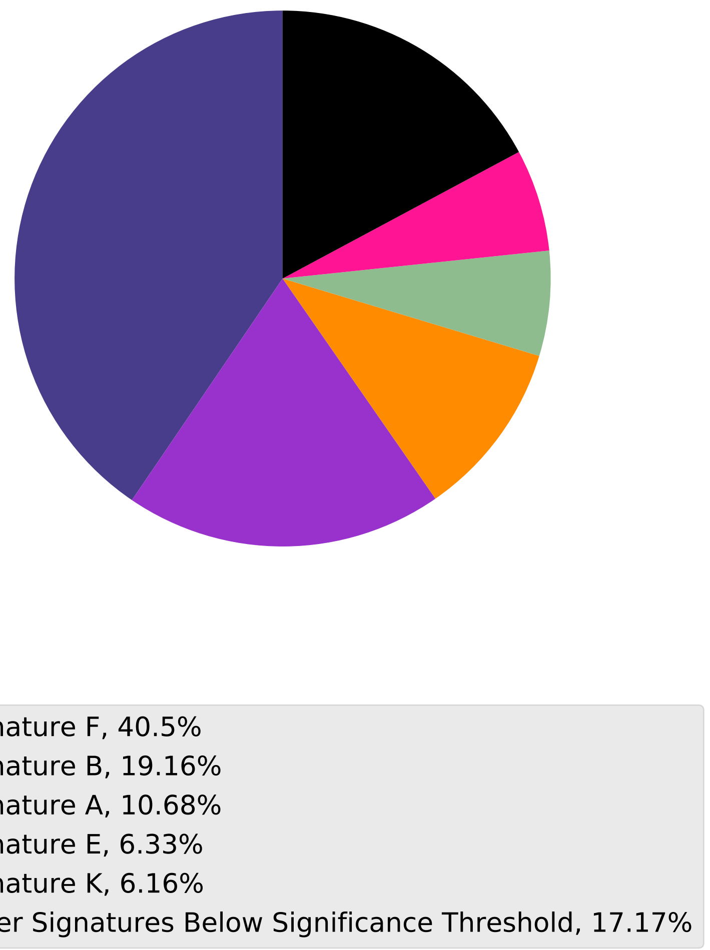

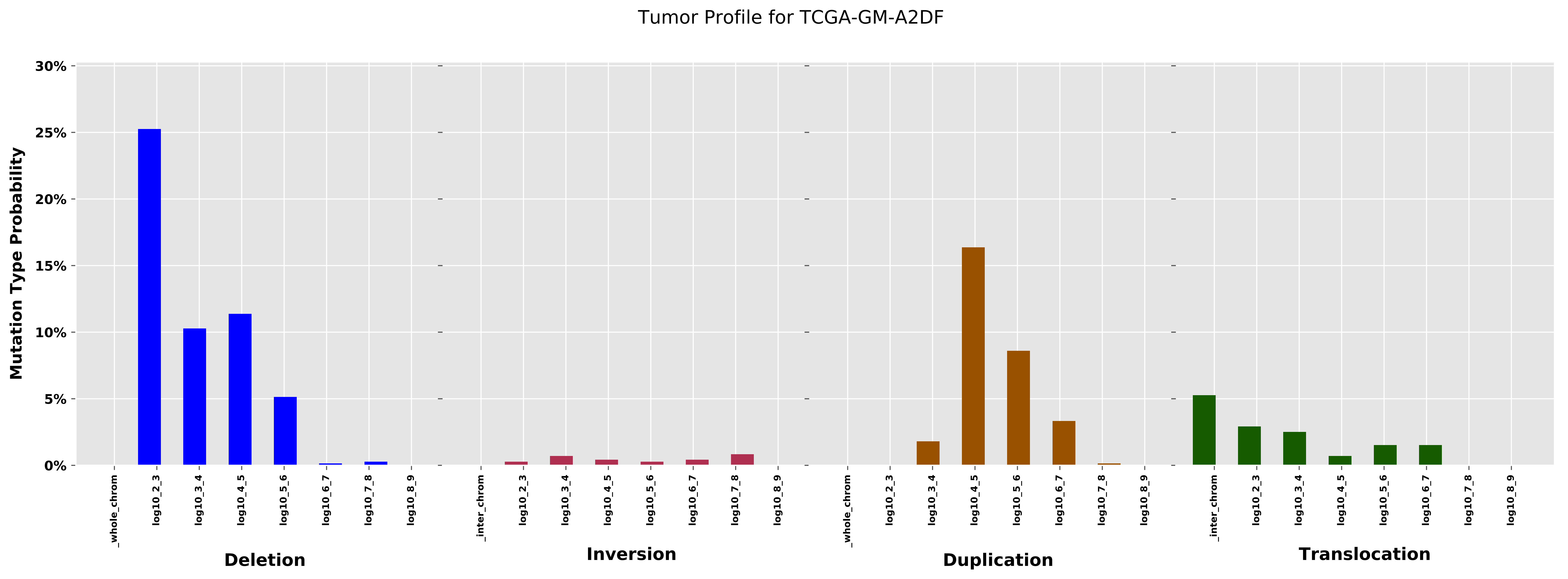

Tumor Profile for TCGA-GM-A2DF
Mutation Type Probability
0%
5%
10%
15%
20%
25%
30%
_whole_chrom
log10_2_3
log10_3_4
log10_4_5
log10_5_6
log10_6_7
log10_7_8
log10_8_9
Deletion
_inter_chrom
Inversion
Duplication
Translocation

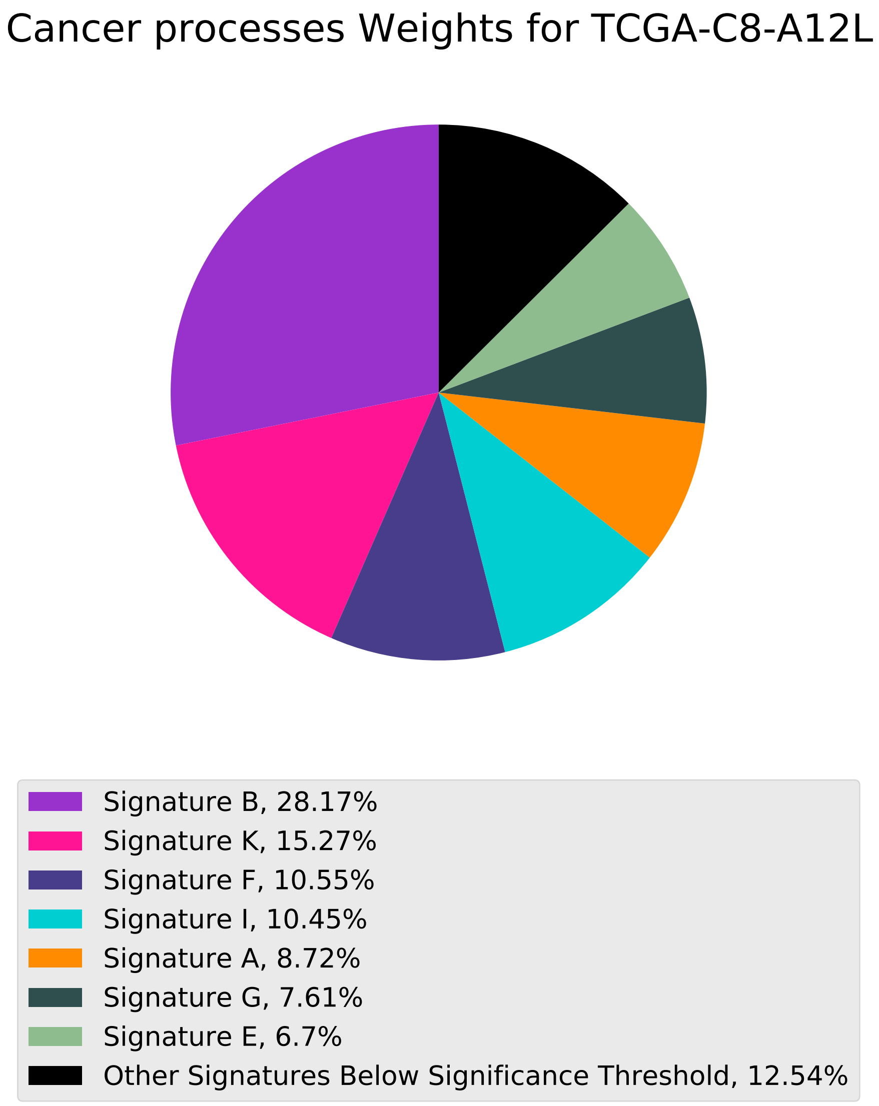
Cancer processes Weights for TCGA-C8-A12L
Signature B, 28.17%
Signature K, 15.27%
Signature F, 10.55%
Signature I, 10.45%
Signature A, 8.72%
Signature G, 7.61%
Signature E, 6.7%
Other Signatures Below Significance Threshold, 12.54%

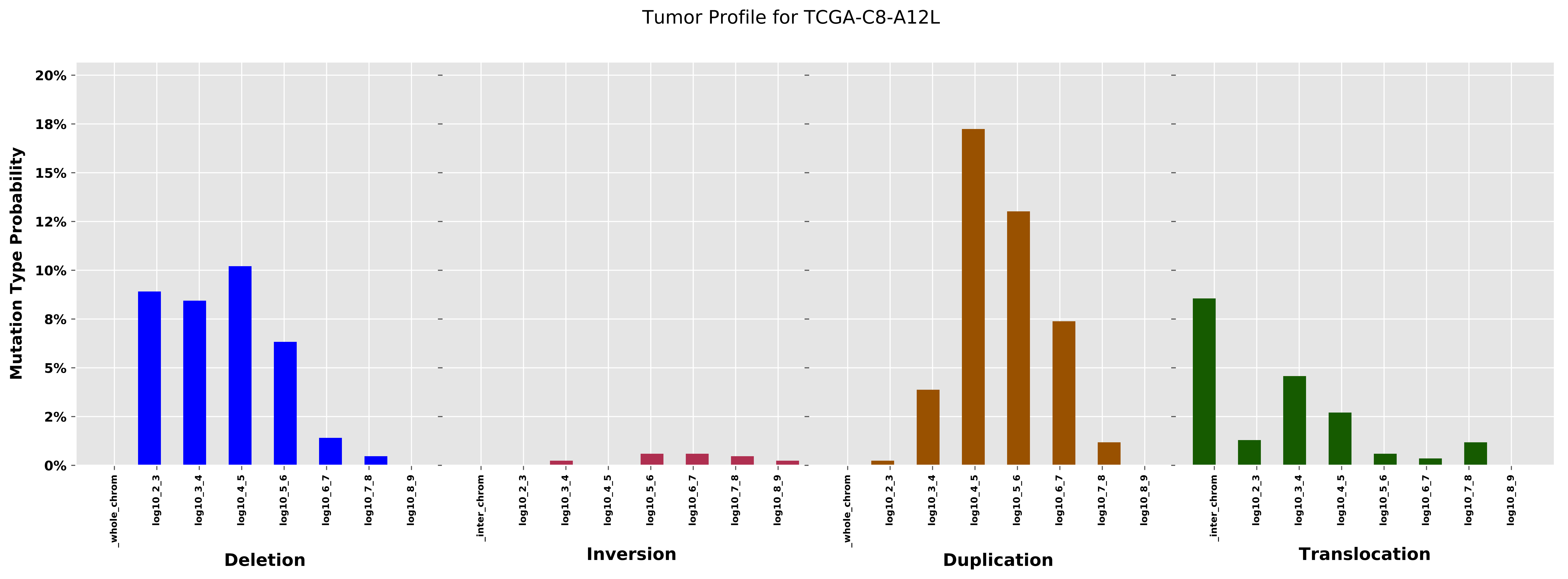

Tumor Profile for TCGA-C8-A12L
Mutation Type Probability
20%
18%
15%
12%
10%
8%
5%
2%
0%
_whole_chrom
log10_2_3
log10_3_4
log10_4_5
log10_5_6
log10_6_7
log10_7_8
log10_8_9
Deletion
_inter_chrom
Inversion
Duplication
Translocation

Cancer processes Weights for TCGA-AG-3890

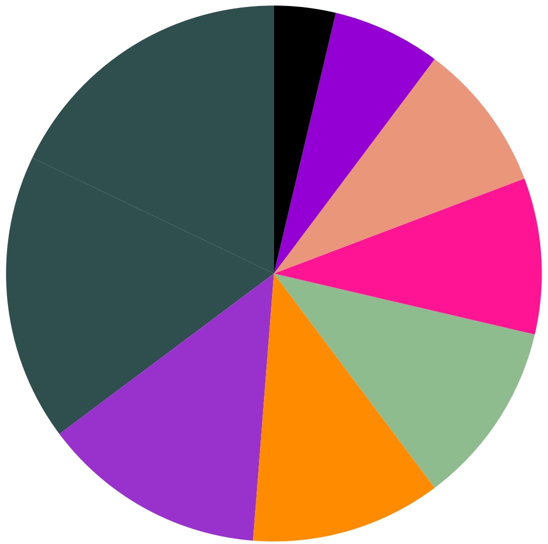

Signature H, 17.9%
Signature G, 17.32%
Signature B, 13.53%
Signature A, 11.49%
Signature E, 11.1%
Signature K, 9.42%
Signature D, 8.99%
Signature J, 6.52%
Other Signatures Below Significance Threshold, 3.71%

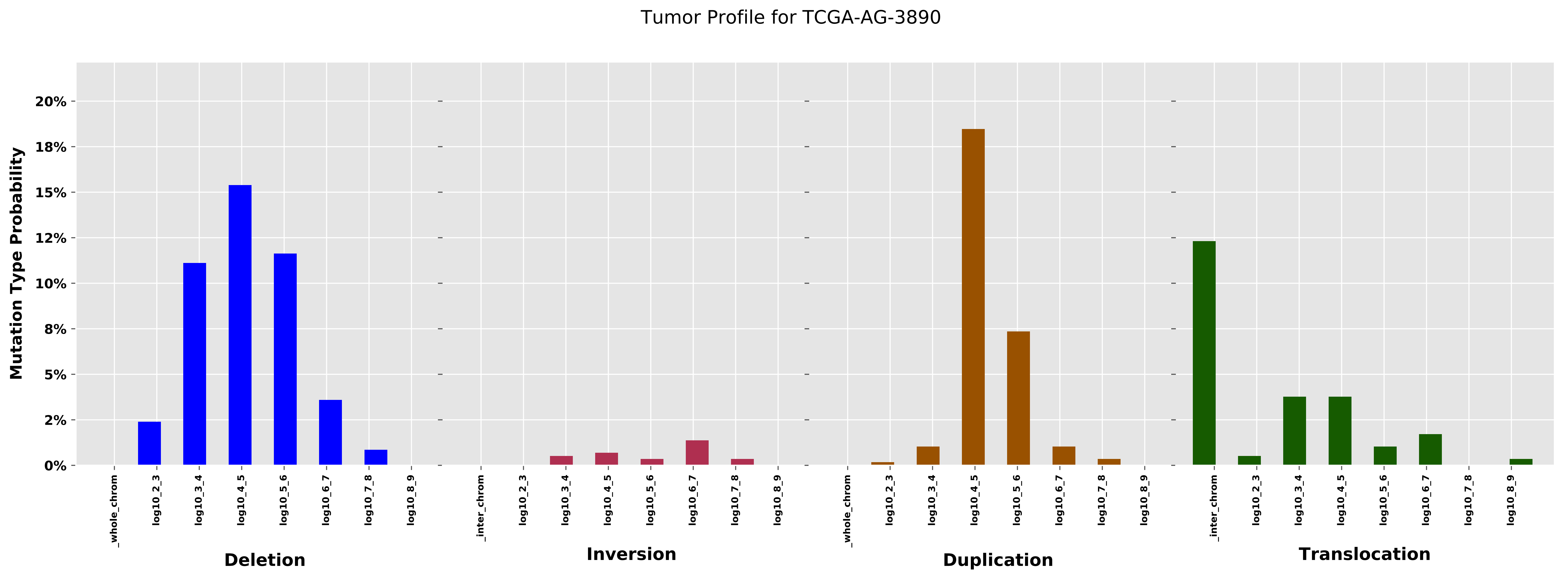

Tumor Profile for TCGA-AG-3890
Mutation Type Probability
20%
18%
15%
12%
10%
8%
5%
2%
0%
_whole_chrom
log10_2_3
log10_3_4
log10_4_5
log10_5_6
log10_6_7
log10_7_8
log10_8_9
_inter_chrom
Deletion
Inversion
Duplication
Translocation

Cancer processes Weights for TCGA-E2-A1LK

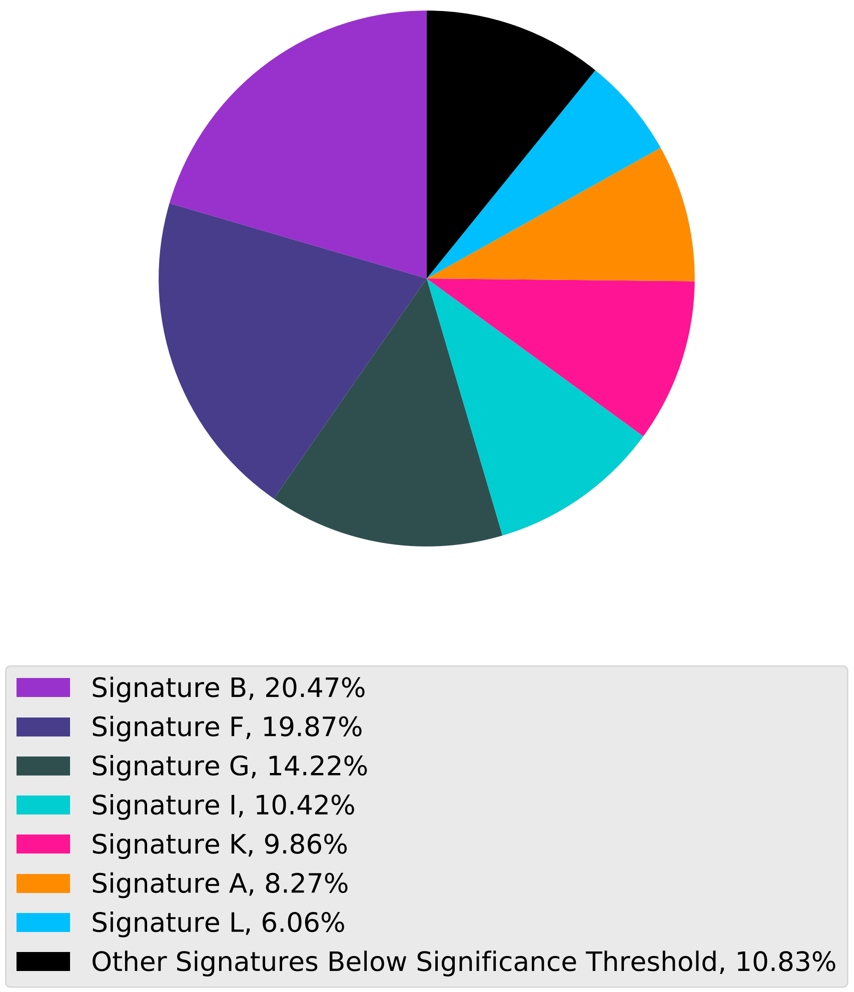

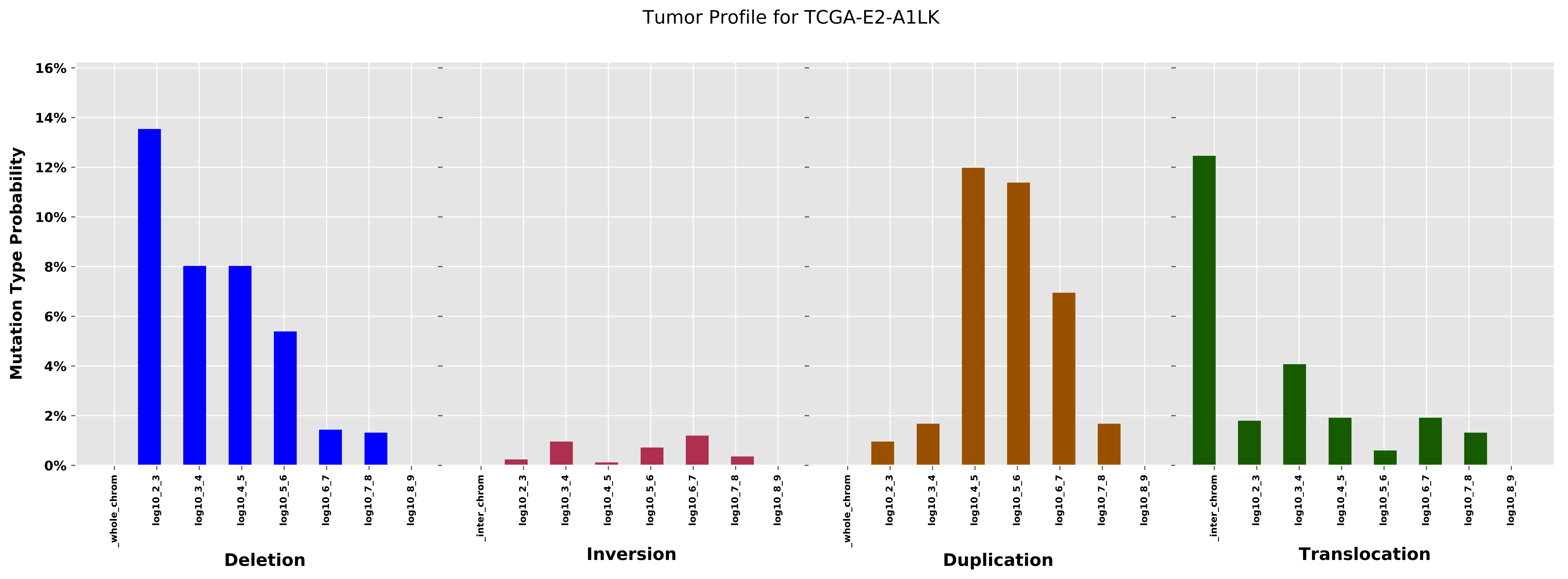

Tumor Profile for TCGA-E2-A1LK
Mutation Type Probability
16%
14%
12%
10%
8%
6%
4%
2%
0%
_whole_chrom
log10_2_3
log10_3_4
log10_4_5
log10_5_6
log10_6_7
log10_7_8
log10_8_9
Deletion
_inter_chrom
log10_2_3
log10_3_4
log10_4_5
log10_5_6
log10_6_7
log10_7_8
log10_8_9
Inversion
_whole_chrom
log10_2_3
log10_3_4
log10_4_5
log10_5_6
log10_6_7
log10_7_8
log10_8_9
Duplication
_inter_chrom
log10_2_3
log10_3_4
log10_4_5
log10_5_6
log10_6_7
log10_7_8
log10_8_9
Translocation

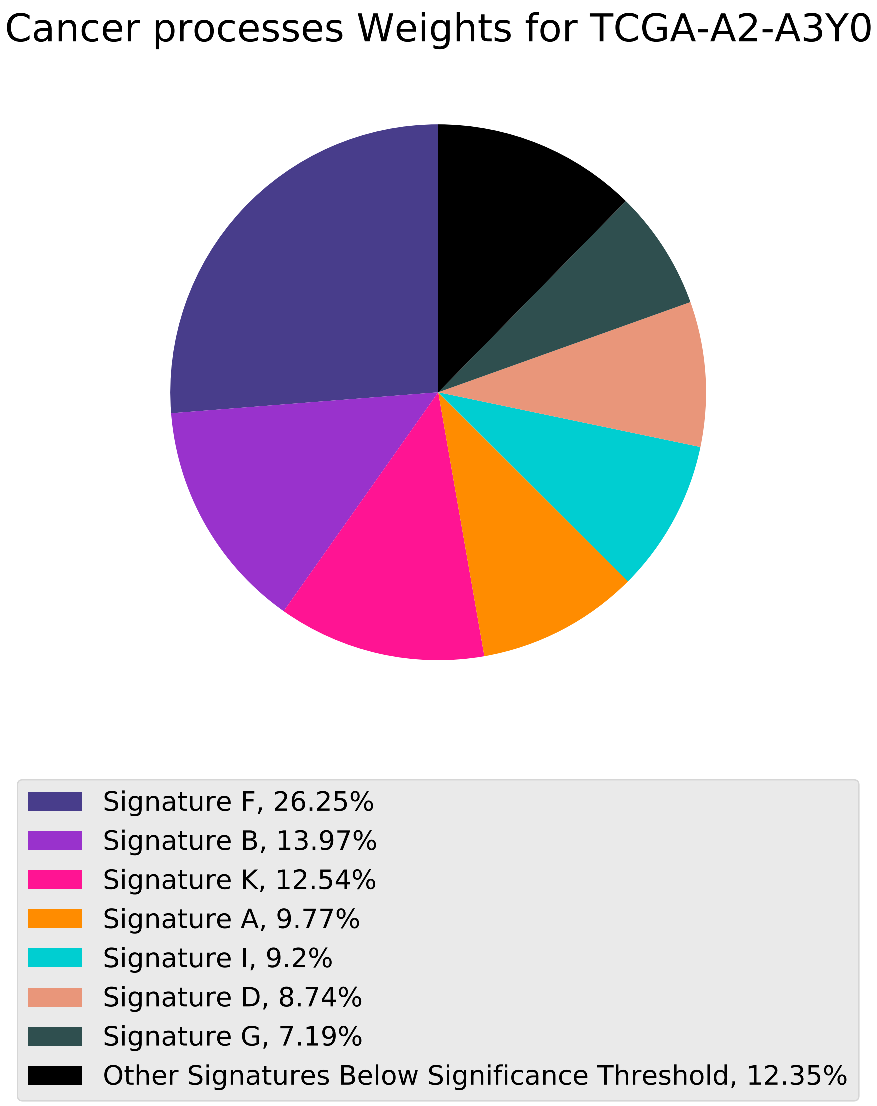
Cancer processes Weights for TCGA-A2-A3Y0
Signature F, 26.25%
Signature B, 13.97%
Signature K, 12.54%
Signature A, 9.77%
Signature I, 9.2%
Signature D, 8.74%
Signature G, 7.19%
Other Signatures Below Significance Threshold, 12.35%

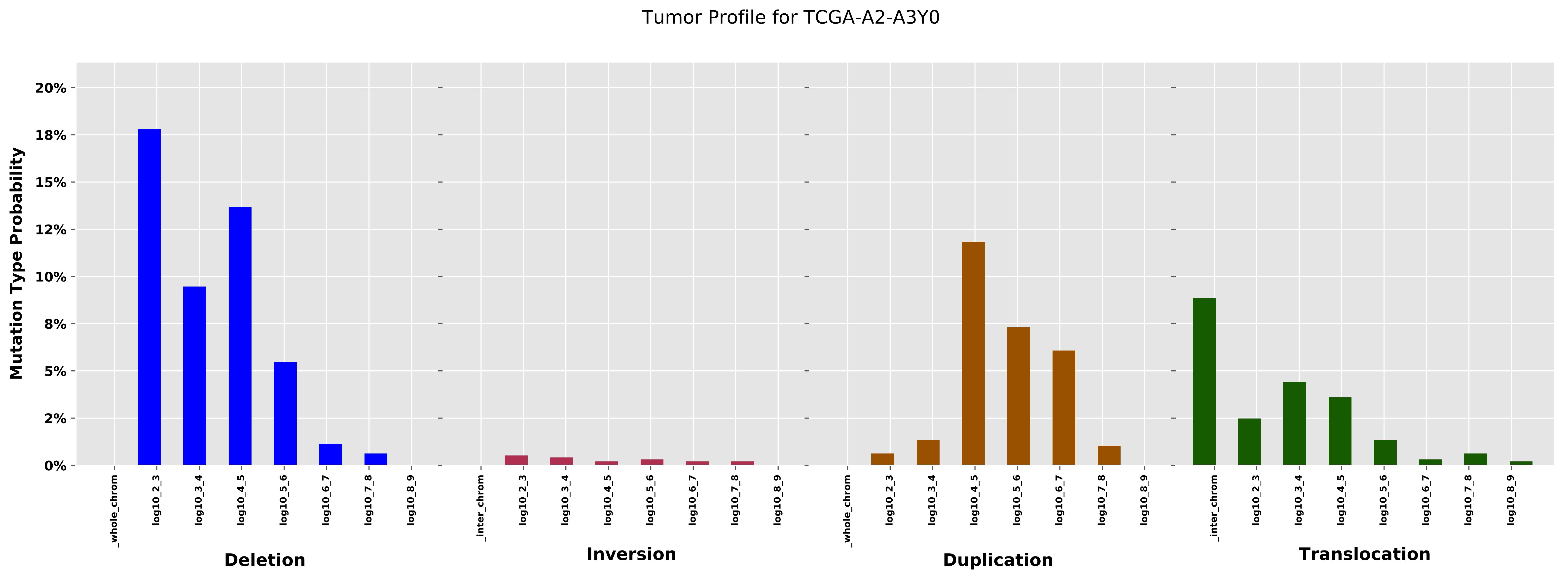
Tumor Profile for TCGA-A2-A3Y0
Mutation Type Probability
20%
18%
15%
12%
10%
8%
5%
2%
0%
_whole_chrom
log10_2_3
log10_3_4
log10_4_5
log10_5_6
log10_6_7
log10_7_8
log10_8_9
_inter_chrom
Deletion
Inversion
Duplication
Translocation

#### Cancer processes Weights for TCGA-B6-A0I1

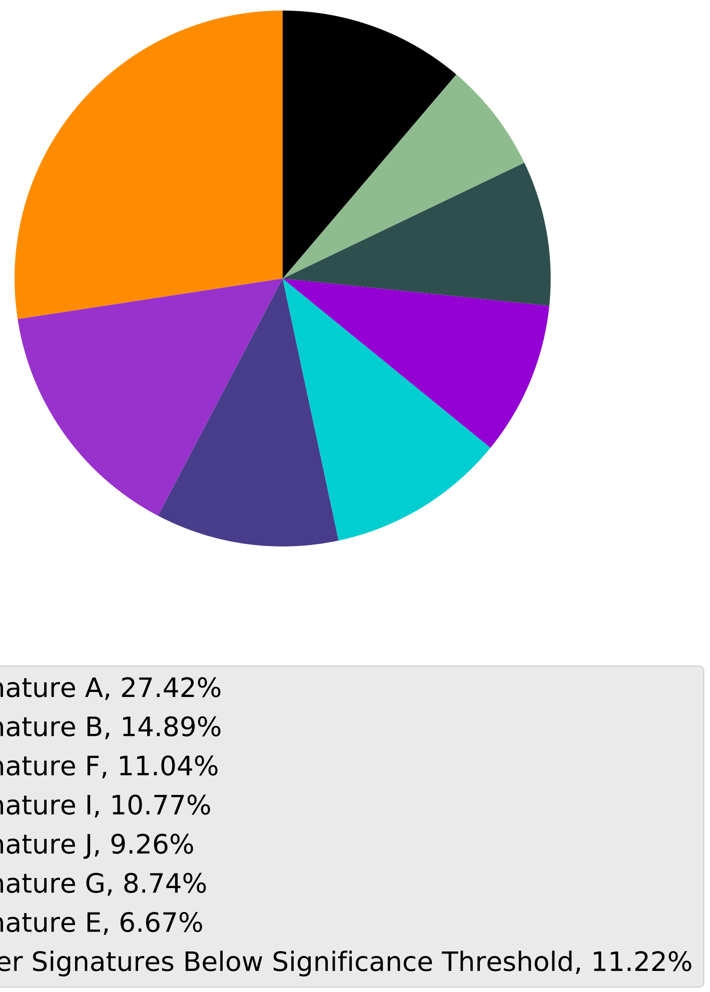

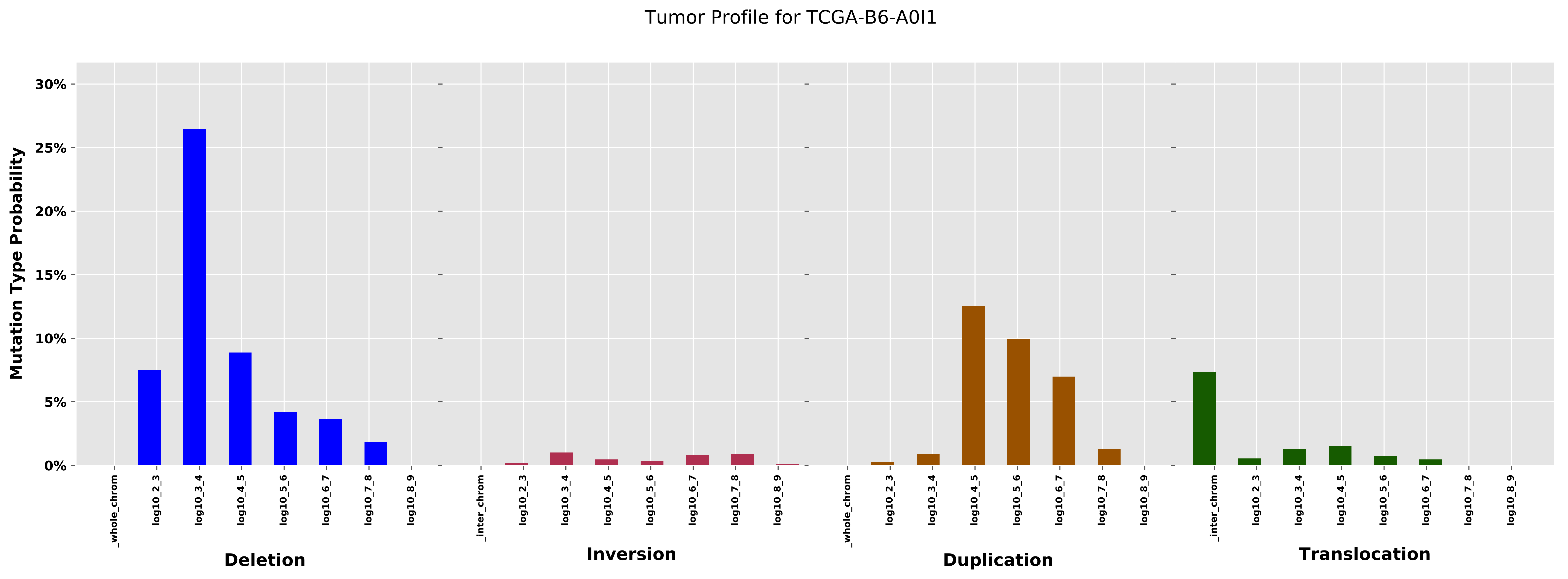
Tumor Profile for TCGA-B6-A0I1
Mutation Type Probability
0%
5%
10%
15%
20%
25%
30%
_whole_chrom
log10_2_3
log10_3_4
log10_4_5
log10_5_6
log10_6_7
log10_7_8
log10_8_9
_inter_chrom
Deletion
Inversion
Duplication
Translocation

Cancer processes Weights for TCGA-E2-A15E

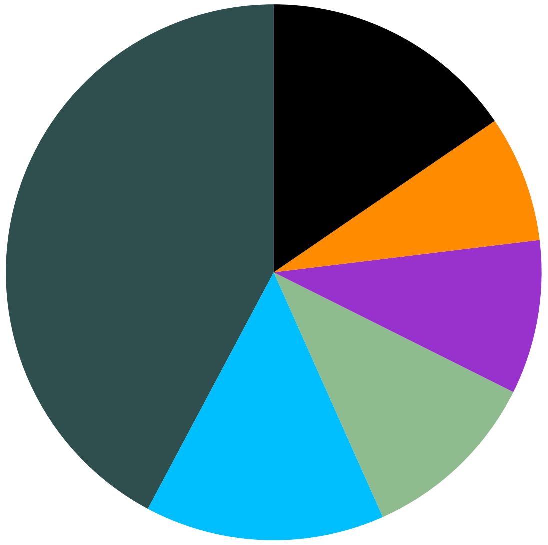

Signature G, 42.21%
Signature L, 14.47%
Signature E, 10.95%
Signature B, 9.3%
Signature A, 7.63%
Other Signatures Below Significance Threshold, 15.44%

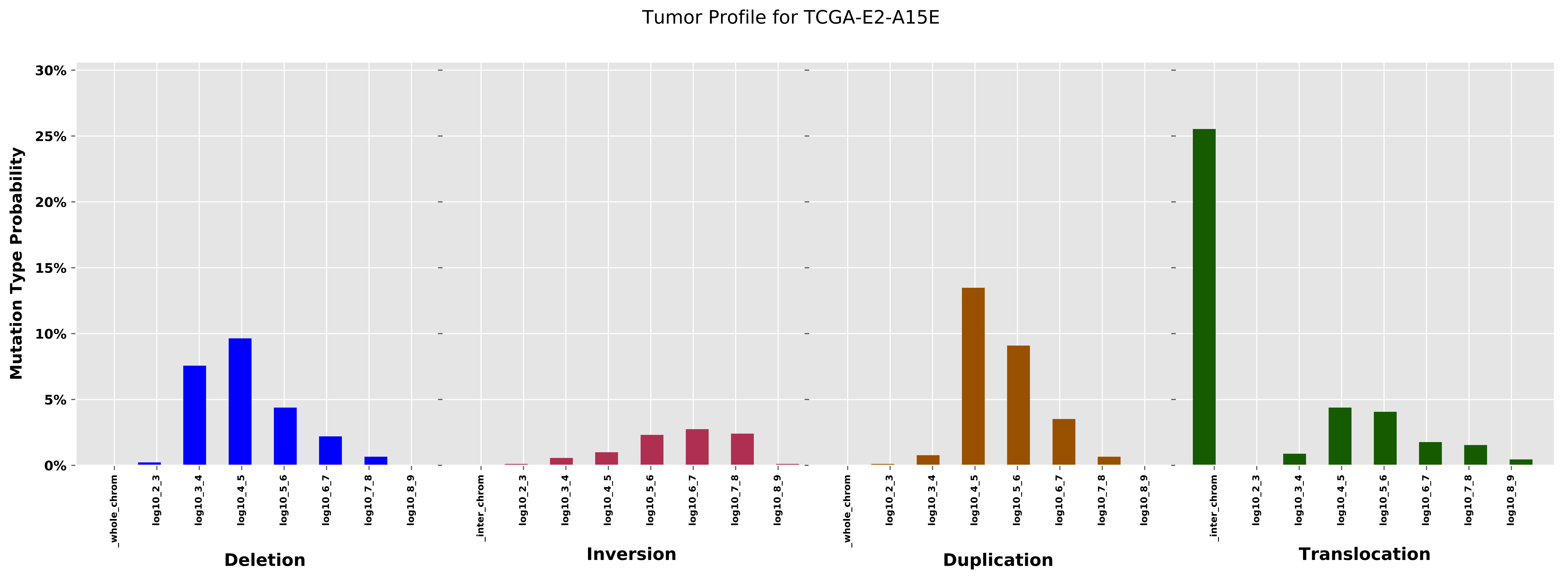

Tumor Profile for TCGA-E2-A15E
Mutation Type Probability
30%
25%
20%
15%
10%
5%
0%
_whole_chrom
log10_2_3
log10_3_4
log10_4_5
log10_5_6
log10_6_7
log10_7_8
log10_8_9
_inter_chrom
Deletion
Inversion
Duplication
Translocation

Cancer processes Weights for TCGA-A6-2680

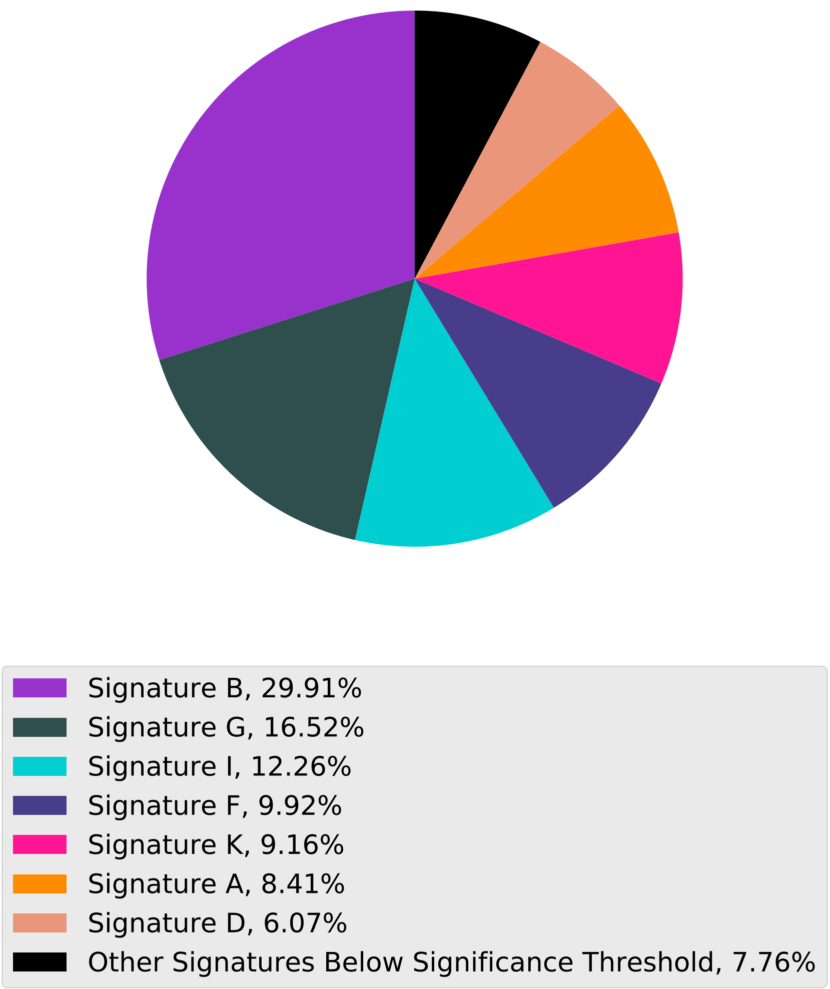

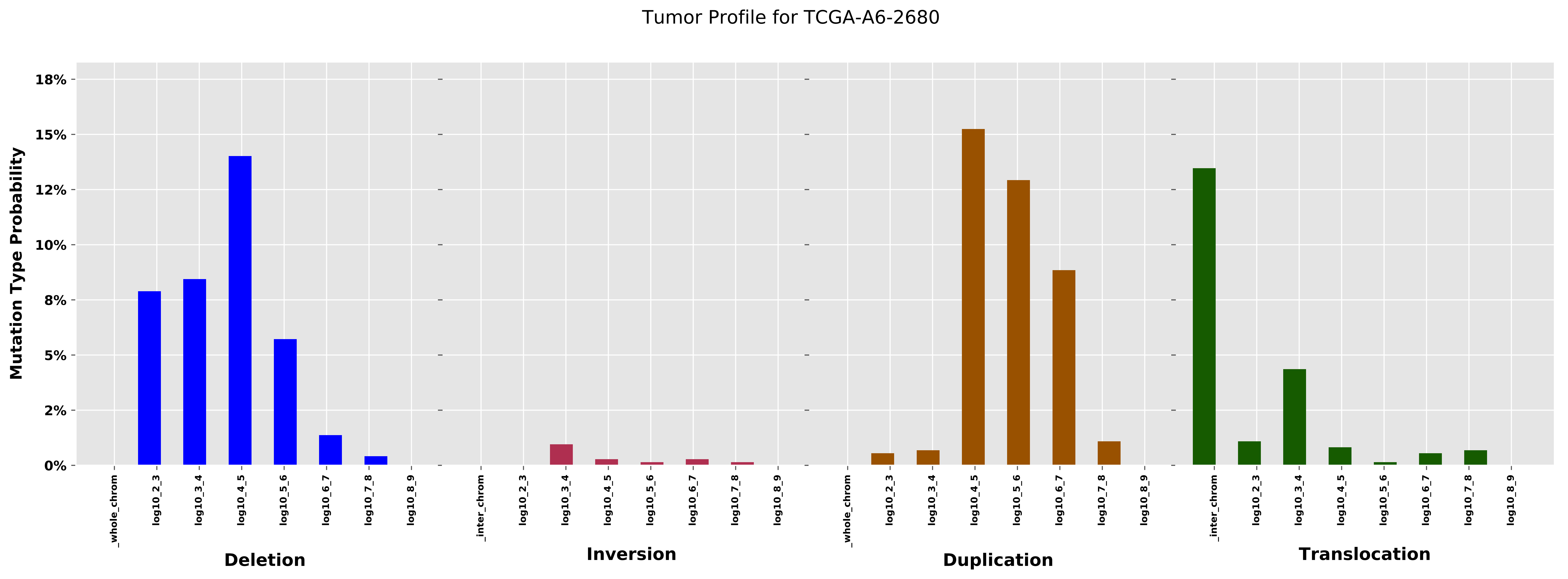

Tumor Profile for TCGA-A6-2680
Mutation Type Probability
18%
15%
12%
10%
8%
5%
2%
0%
_whole_chrom
log10_2_3
log10_3_4
log10_4_5
log10_5_6
log10_6_7
log10_7_8
log10_8_9
_inter_chrom
Deletion
Inversion
Duplication
Translocation

Cancer processes Weights for TCGA-A8-A07B

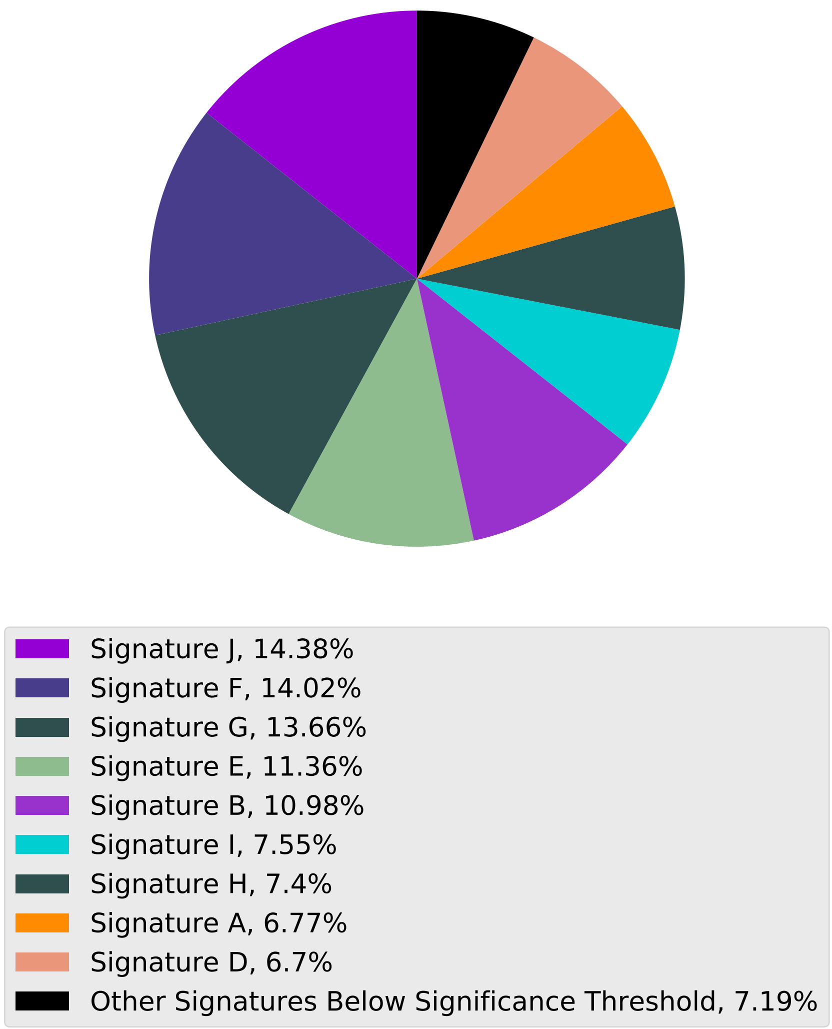

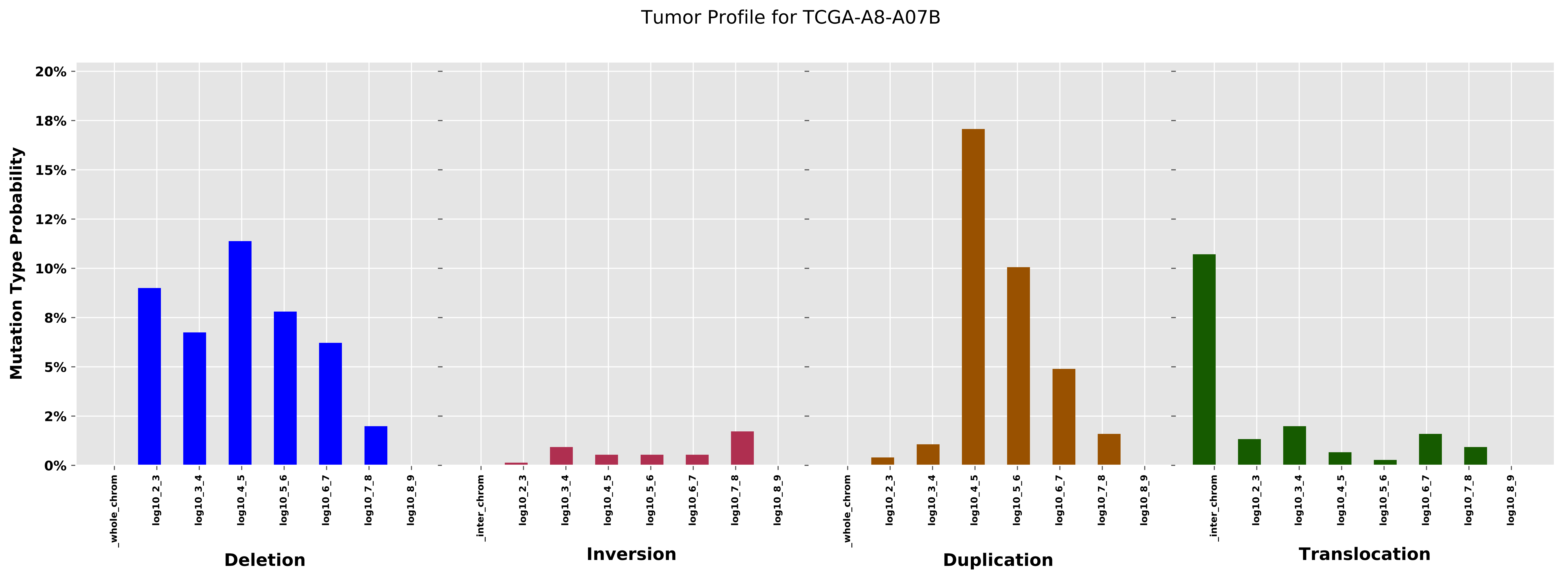
Tumor Profile for TCGA-A8-A07B
Mutation Type Probability
20%
18%
15%
12%
10%
8%
5%
2%
0%
_whole_chrom
log10_2_3
log10_3_4
log10_4_5
log10_5_6
log10_6_7
log10_7_8
log10_8_9
_inter_chrom
Deletion
Inversion
Duplication
Translocation

Cancer processes Weights for TCGA-AO-A0J6

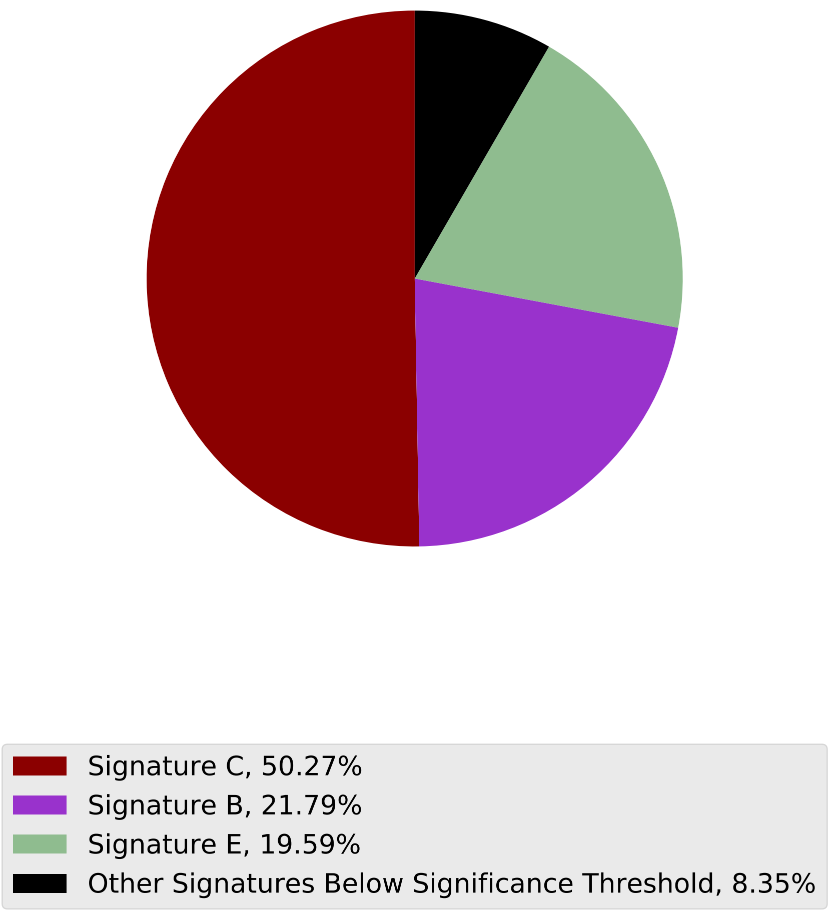

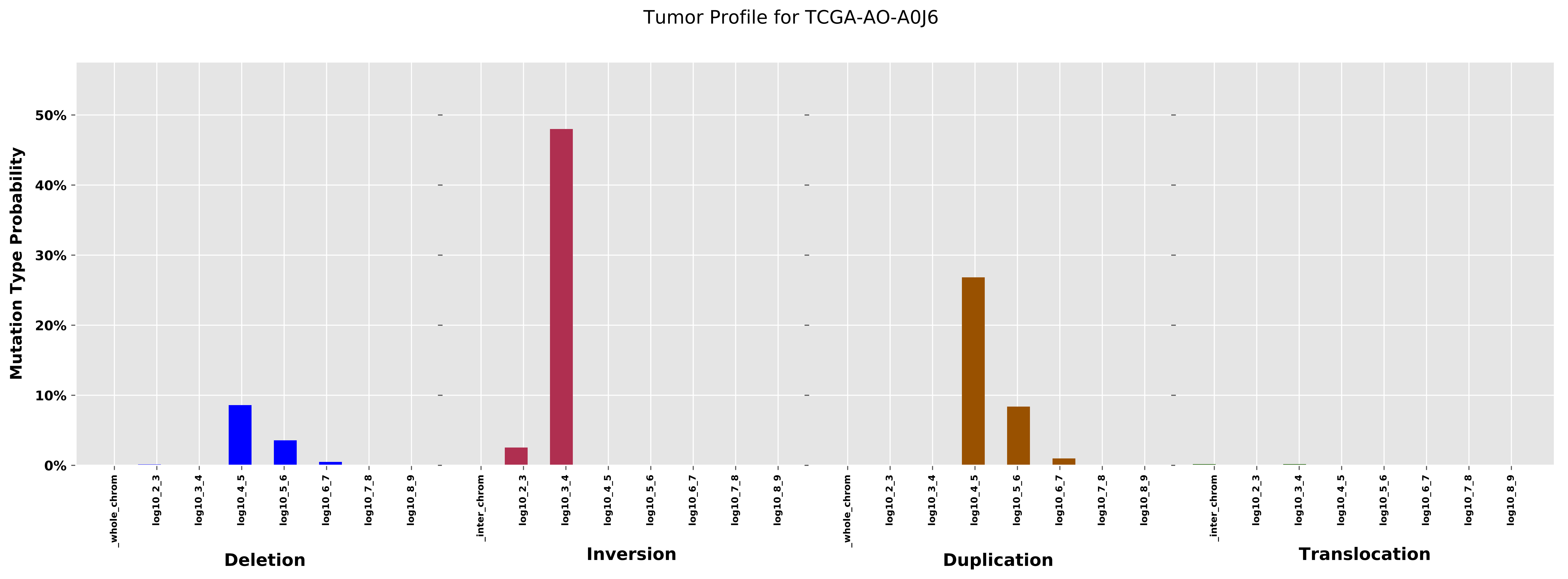

Tumor Profile for TCGA-AO-A0J6
Mutation Type Probability
50%
40%
30%
20%
10%
0%
_whole_chrom
log10_2_3
log10_3_4
log10_4_5
log10_5_6
log10_6_7
log10_7_8
log10_8_9
_inter_chrom
Deletion
Inversion
Duplication
Translocation

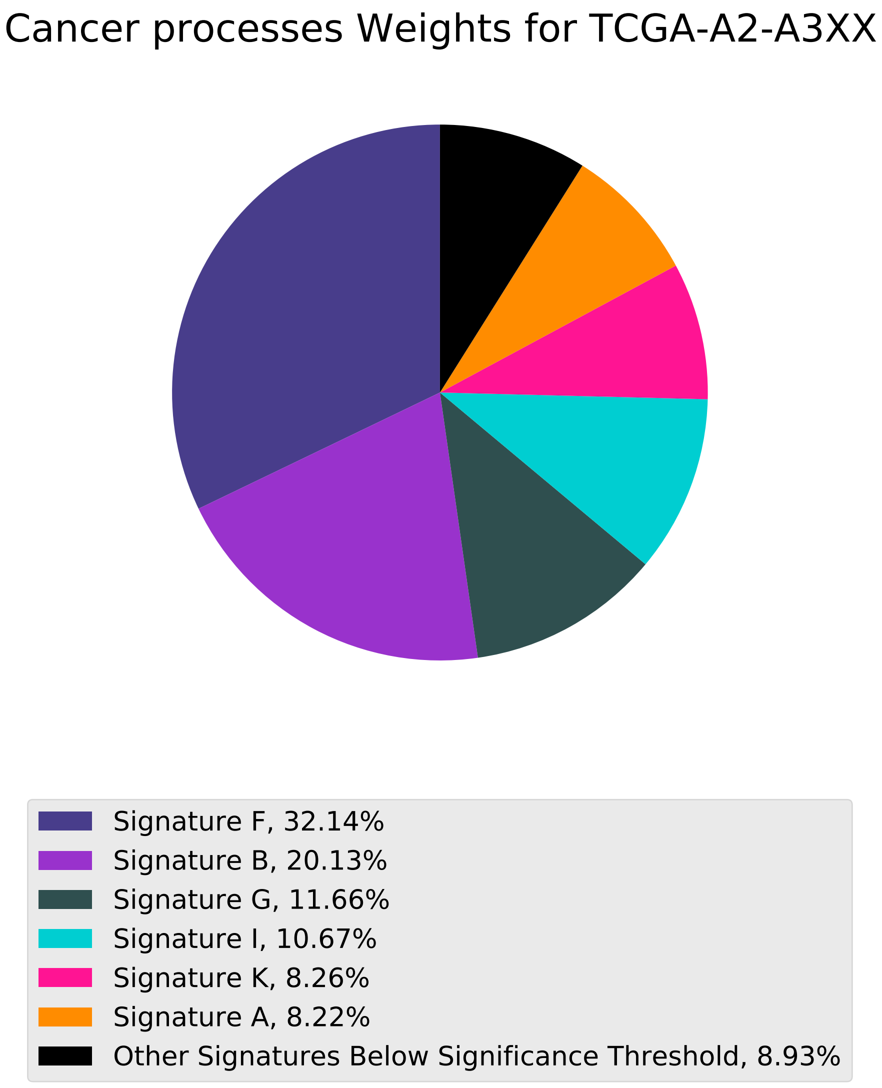
Cancer processes Weights for TCGA-A2-A3XX
Signature F, 32.14%
Signature B, 20.13%
Signature G, 11.66%
Signature I, 10.67%
Signature K, 8.26%
Signature A, 8.22%
Other Signatures Below Significance Threshold, 8.93%

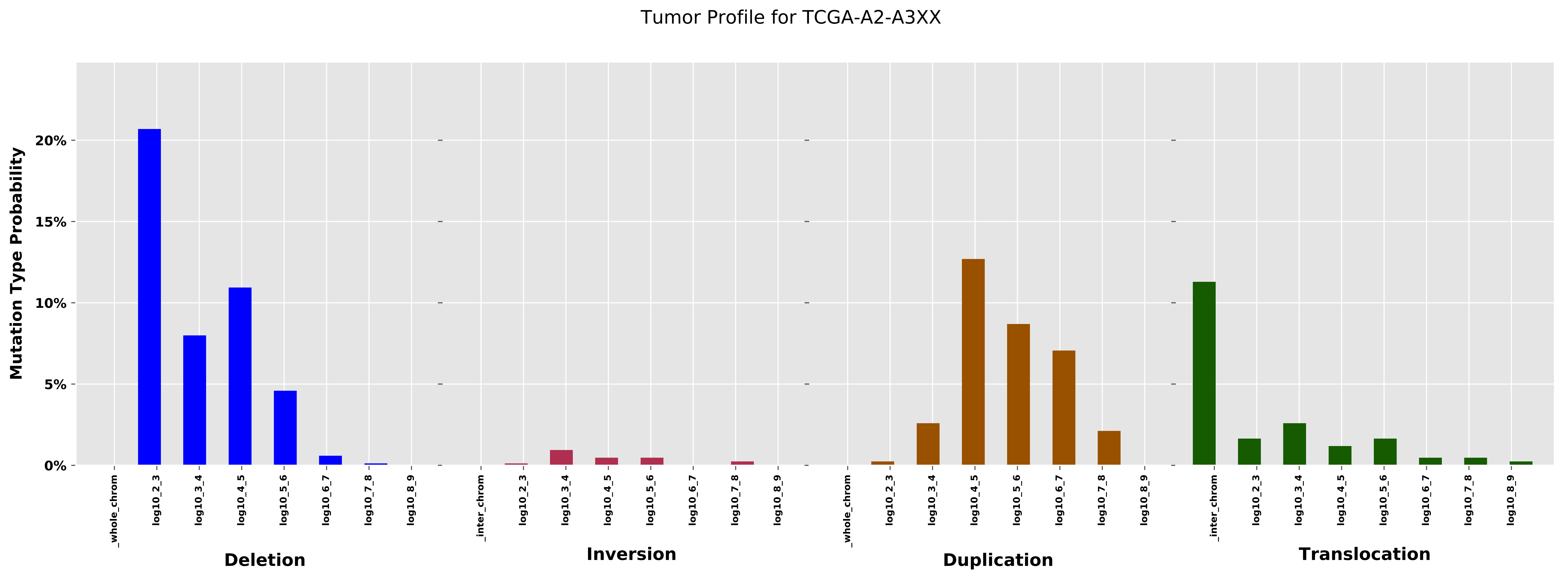

Tumor Profile for TCGA-A2-A3XX
Mutation Type Probability
0%
5%
10%
15%
20%
_whole_chrom
log10_2_3
log10_3_4
log10_4_5
log10_5_6
log10_6_7
log10_7_8
log10_8_9
Deletion
_inter_chrom
log10_2_3
log10_3_4
log10_4_5
log10_5_6
log10_6_7
log10_7_8
log10_8_9
Inversion
_whole_chrom
log10_2_3
log10_3_4
log10_4_5
log10_5_6
log10_6_7
log10_7_8
log10_8_9
Duplication
_inter_chrom
log10_2_3
log10_3_4
log10_4_5
log10_5_6
log10_6_7
log10_7_8
log10_8_9
Translocation

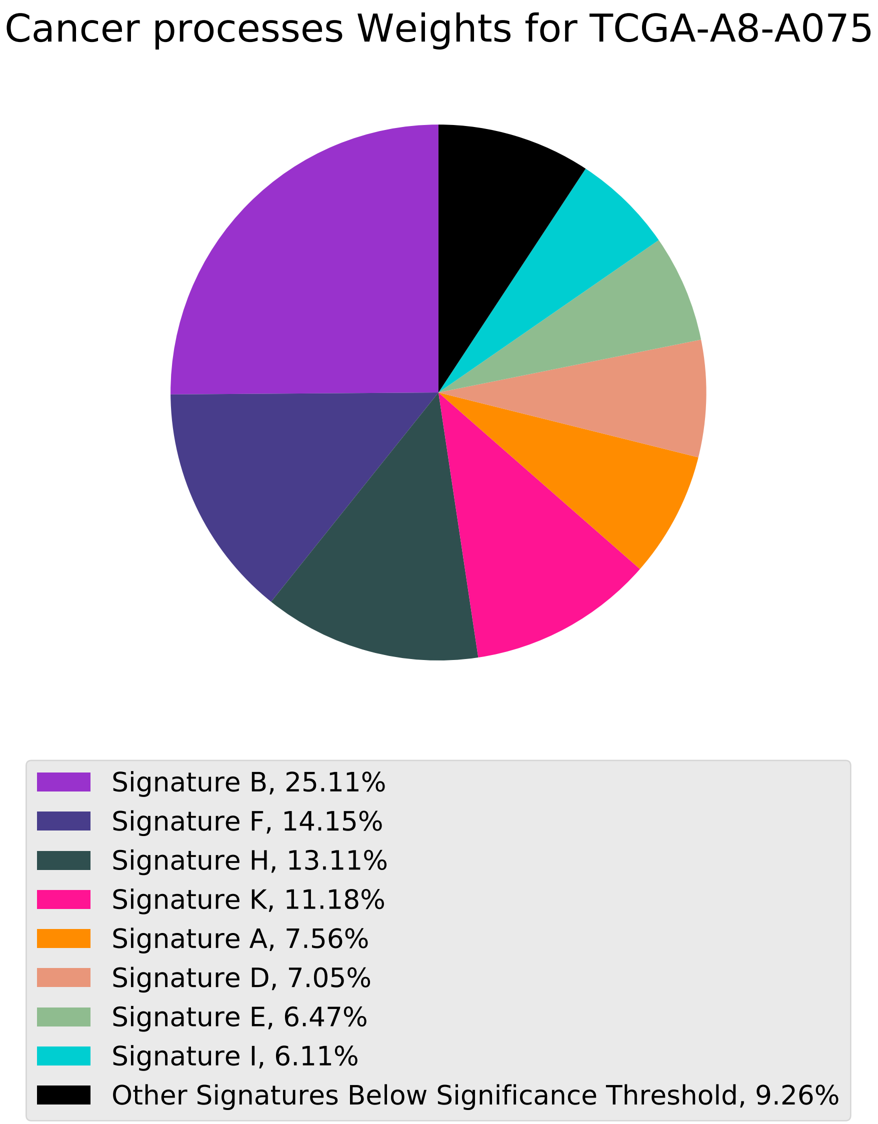

Cancer processes Weights for TCGA-A8-A075
Signature B, 25.11%
Signature F, 14.15%
Signature H, 13.11%
Signature K, 11.18%
Signature A, 7.56%
Signature D, 7.05%
Signature E, 6.47%
Signature I, 6.11%
Other Signatures Below Significance Threshold, 9.26%

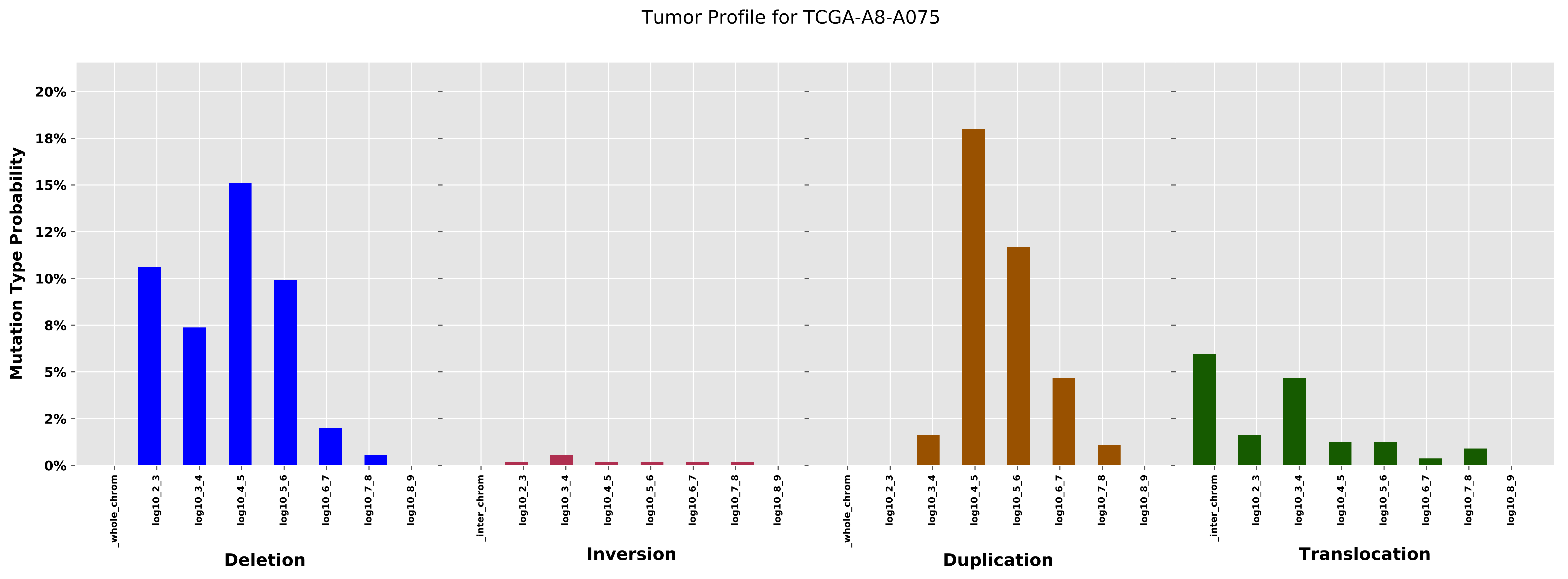

Tumor Profile for TCGA-A8-A075
Mutation Type Probability
0%
2%
5%
8%
10%
12%
15%
18%
20%
_whole_chrom
log10_2_3
log10_3_4
log10_4_5
log10_5_6
log10_6_7
log10_7_8
log10_8_9
_inter_chrom
Deletion
Inversion
Duplication
Translocation

Cancer processes Weights for TCGA-AR-A1AY

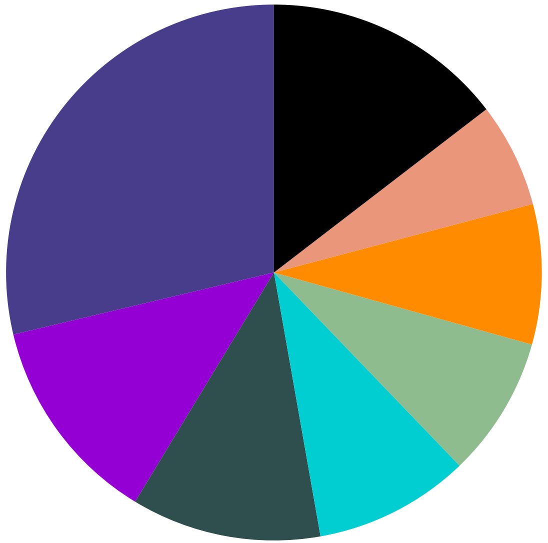

| |
|---|
| Signature F, 28.72% |
| Signature J, 12.61% |
| Signature H, 11.45% |
| Signature I, 9.4% |
| Signature E, 8.48% |
| Signature A, 8.45% |
| Signature D, 6.31% |
| Other Signatures Below Significance Threshold, 14.57% |

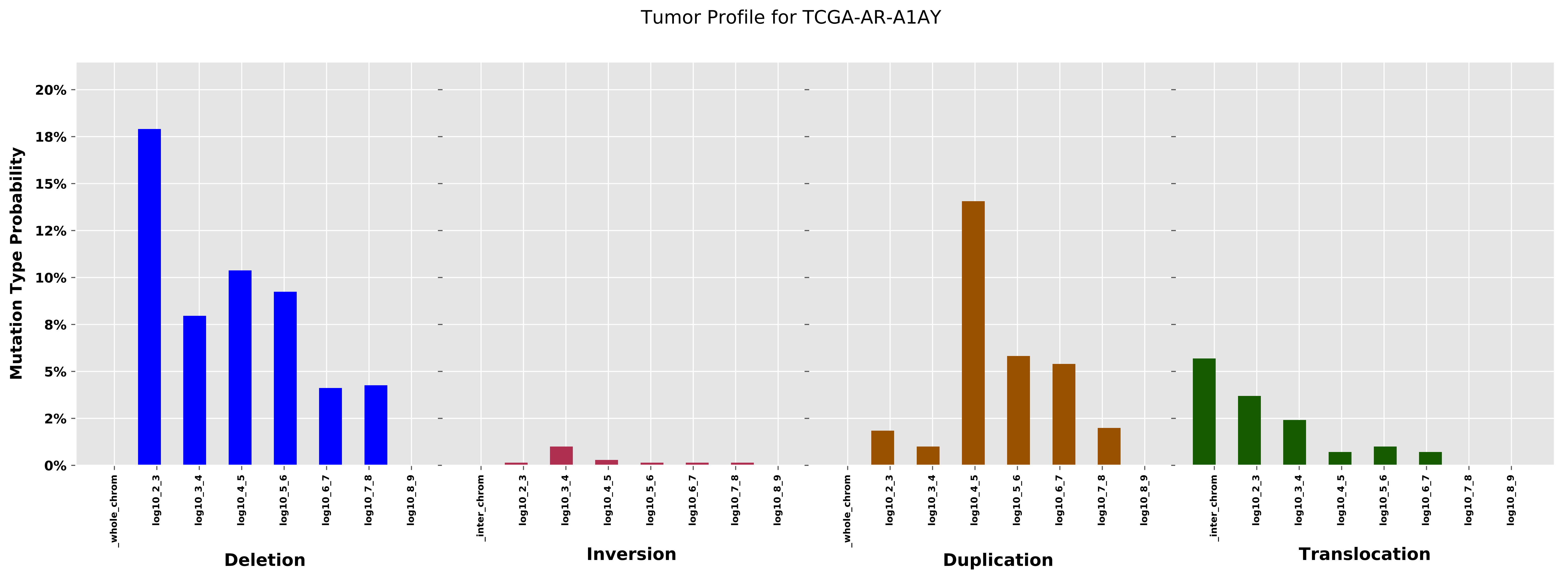

Tumor Profile for TCGA-AR-A1AY
Mutation Type Probability
20%
18%
15%
12%
10%
8%
5%
2%
0%
_whole_chrom
log10_2_3
log10_3_4
log10_4_5
log10_5_6
log10_6_7
log10_7_8
log10_8_9
_inter_chrom
Deletion
Inversion
Duplication
Translocation

Cancer processes Weights for TCGA-AG-3901

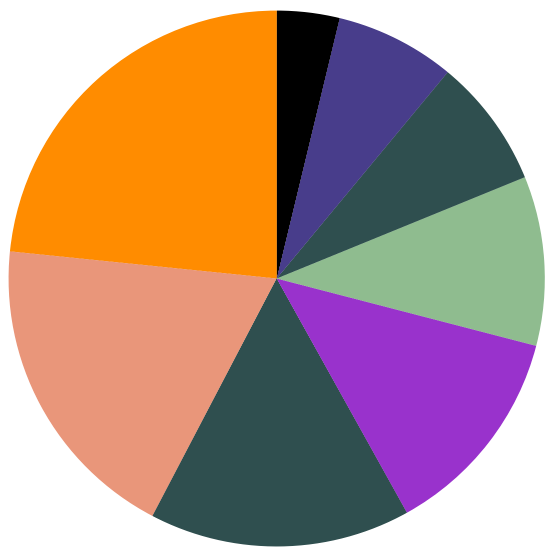

Signature A, 23.39%
Signature D, 18.95%
Signature H, 15.75%
Signature B, 12.88%
Signature E, 10.22%
Signature G, 7.81%
Signature F, 7.24%
Other Signatures Below Significance Threshold, 3.77%

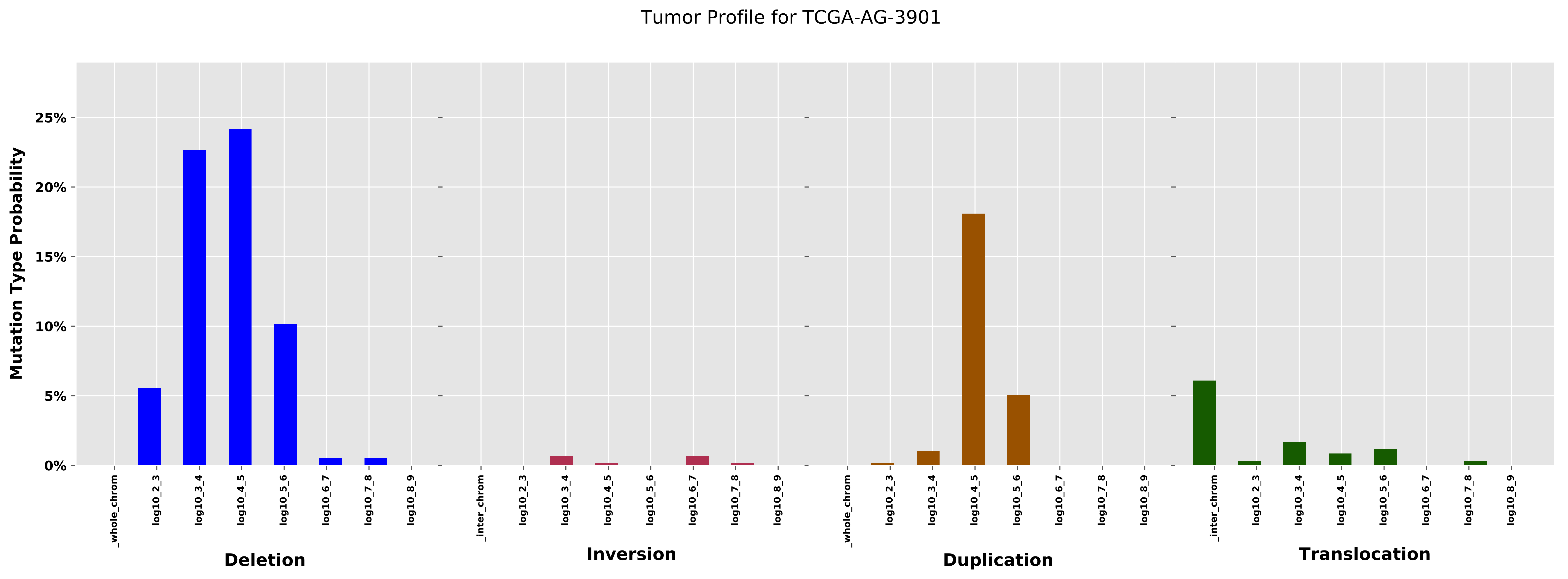

Tumor Profile for TCGA-AG-3901
Mutation Type Probability
0%
5%
10%
15%
20%
25%
_whole_chrom
log10_2_3
log10_3_4
log10_4_5
log10_5_6
log10_6_7
log10_7_8
log10_8_9
Deletion
_inter_chrom
log10_2_3
log10_3_4
log10_4_5
log10_5_6
log10_6_7
log10_7_8
log10_8_9
Inversion
_whole_chrom
log10_2_3
log10_3_4
log10_4_5
log10_5_6
log10_6_7
log10_7_8
log10_8_9
Duplication
_inter_chrom
log10_2_3
log10_3_4
log10_4_5
log10_5_6
log10_6_7
log10_7_8
log10_8_9
Translocation

#### Cancer processes Weights for TCGA-E2-A152

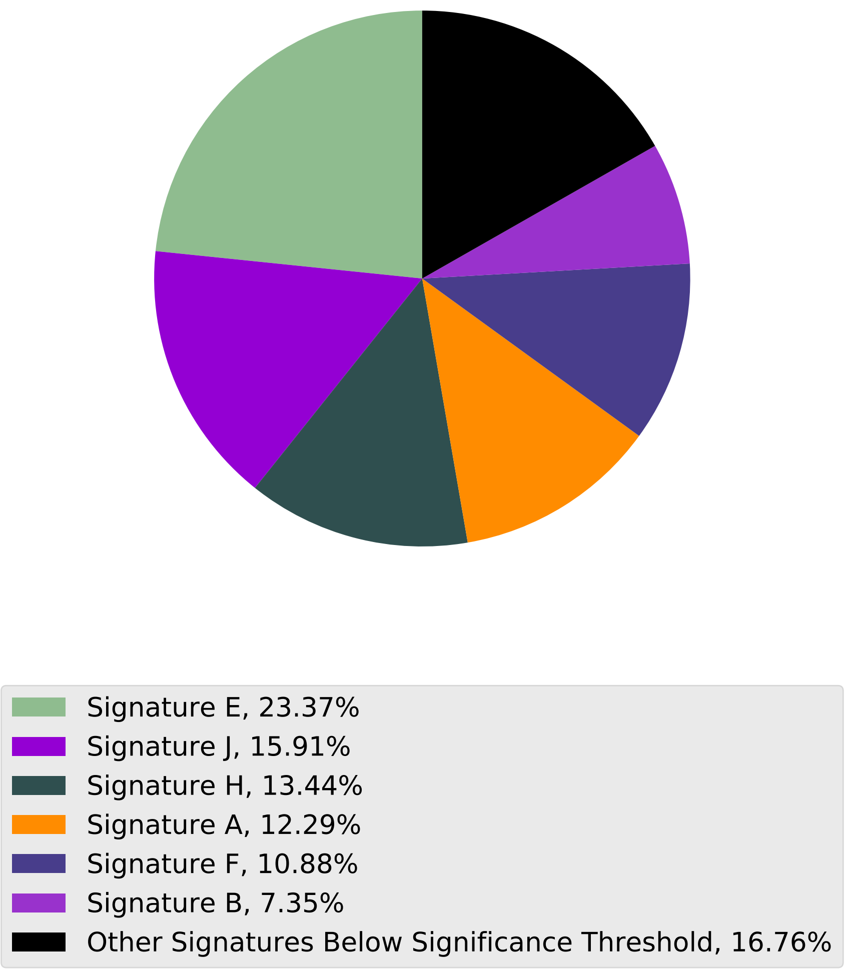

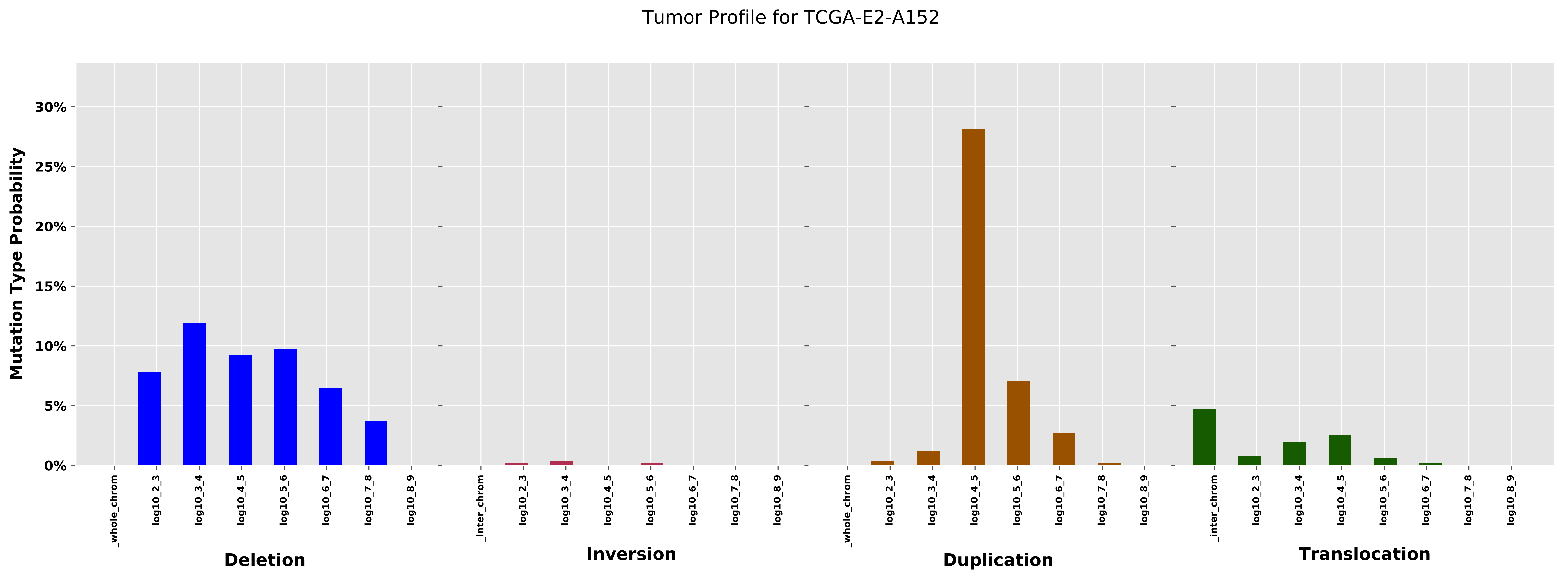

Tumor Profile for TCGA-E2-A152
Mutation Type Probability
0%
5%
10%
15%
20%
25%
30%
_whole_chrom
log10_2_3
log10_3_4
log10_4_5
log10_5_6
log10_6_7
log10_7_8
log10_8_9
_inter_chrom
Deletion
Inversion
Duplication
Translocation

Cancer processes Weights for TCGA-AG-4015

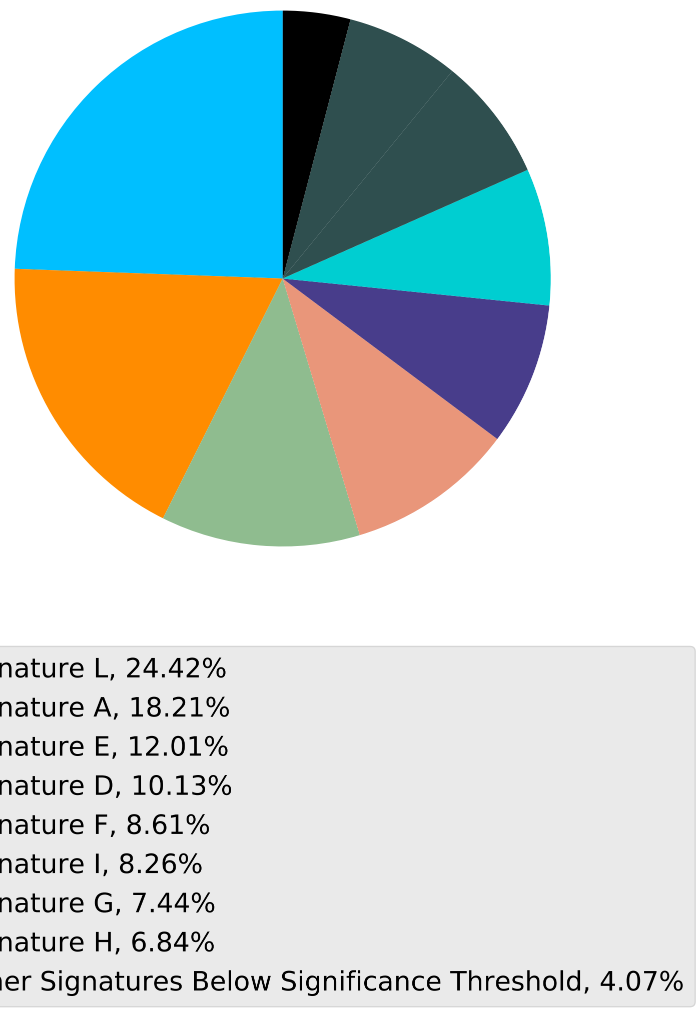

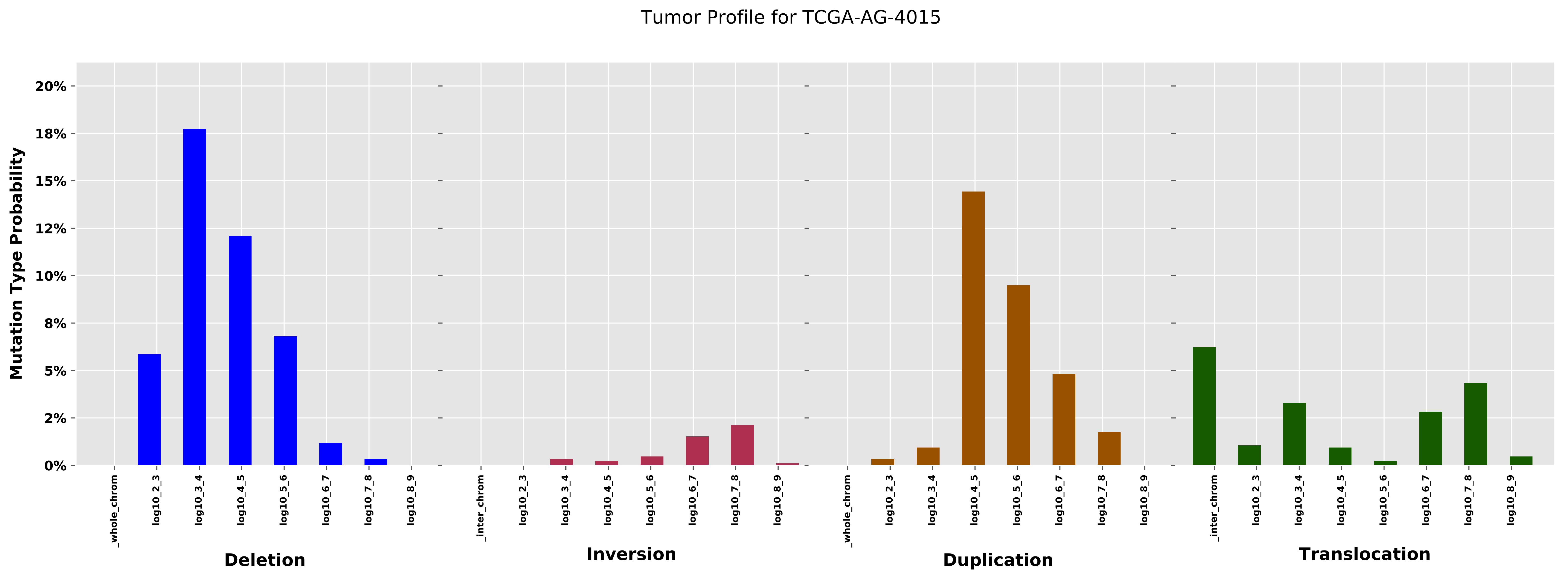
Tumor Profile for TCGA-AG-4015
Mutation Type Probability
20%
18%
15%
12%
10%
8%
5%
2%
0%
_whole_chrom
log10_2_3
log10_3_4
log10_4_5
log10_5_6
log10_6_7
log10_7_8
log10_8_9
_inter_chrom
Deletion
Inversion
Duplication
Translocation

Cancer processes Weights for TCGA-BH-A0B9

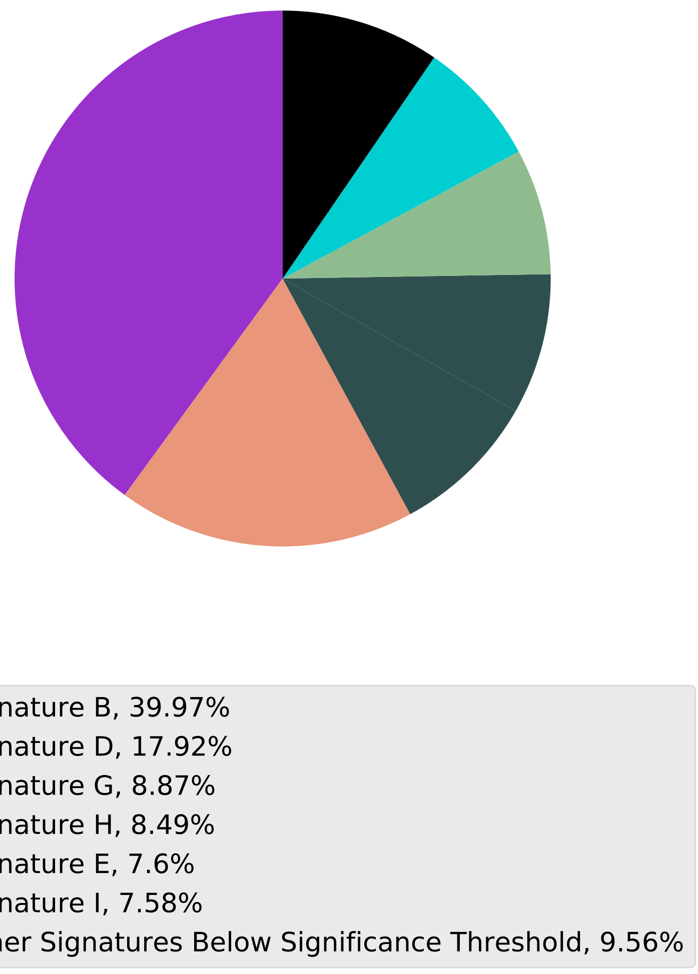

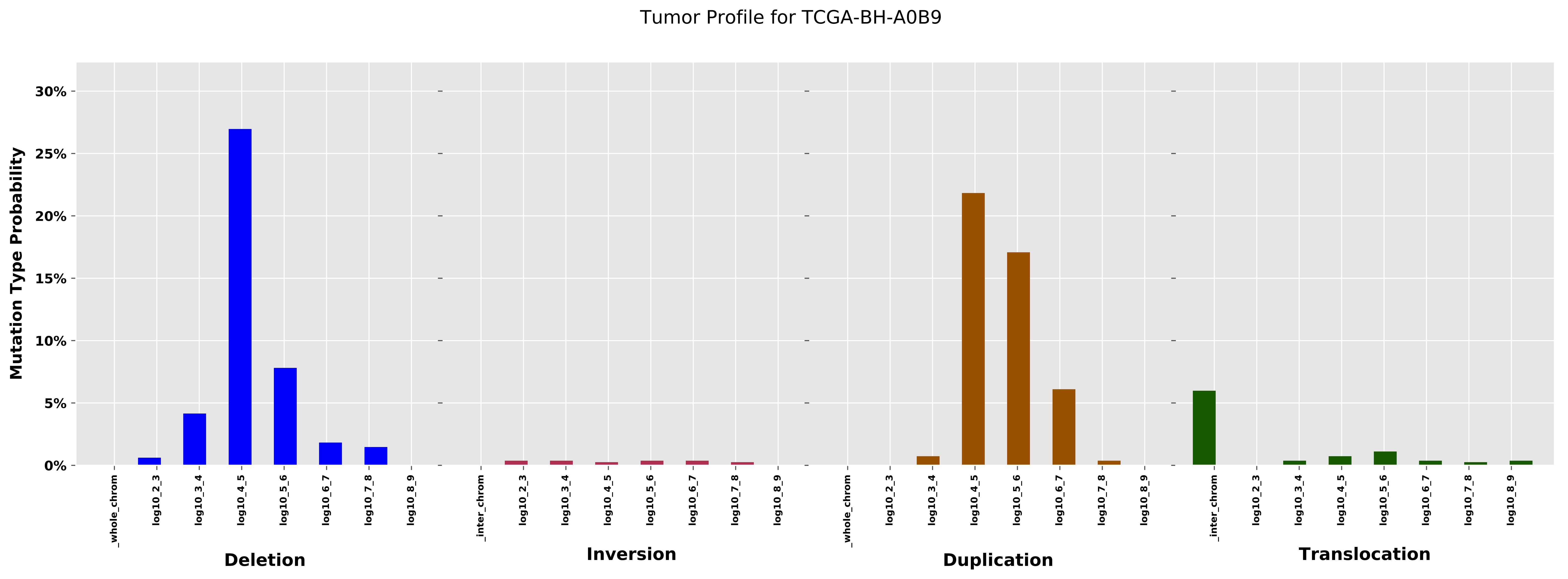

Tumor Profile for TCGA-BH-A0B9
Mutation Type Probability
0%
5%
10%
15%
20%
25%
30%
_whole_chrom
log10_2_3
log10_3_4
log10_4_5
log10_5_6
log10_6_7
log10_7_8
log10_8_9
_inter_chrom
Deletion
Inversion
Duplication
Translocation

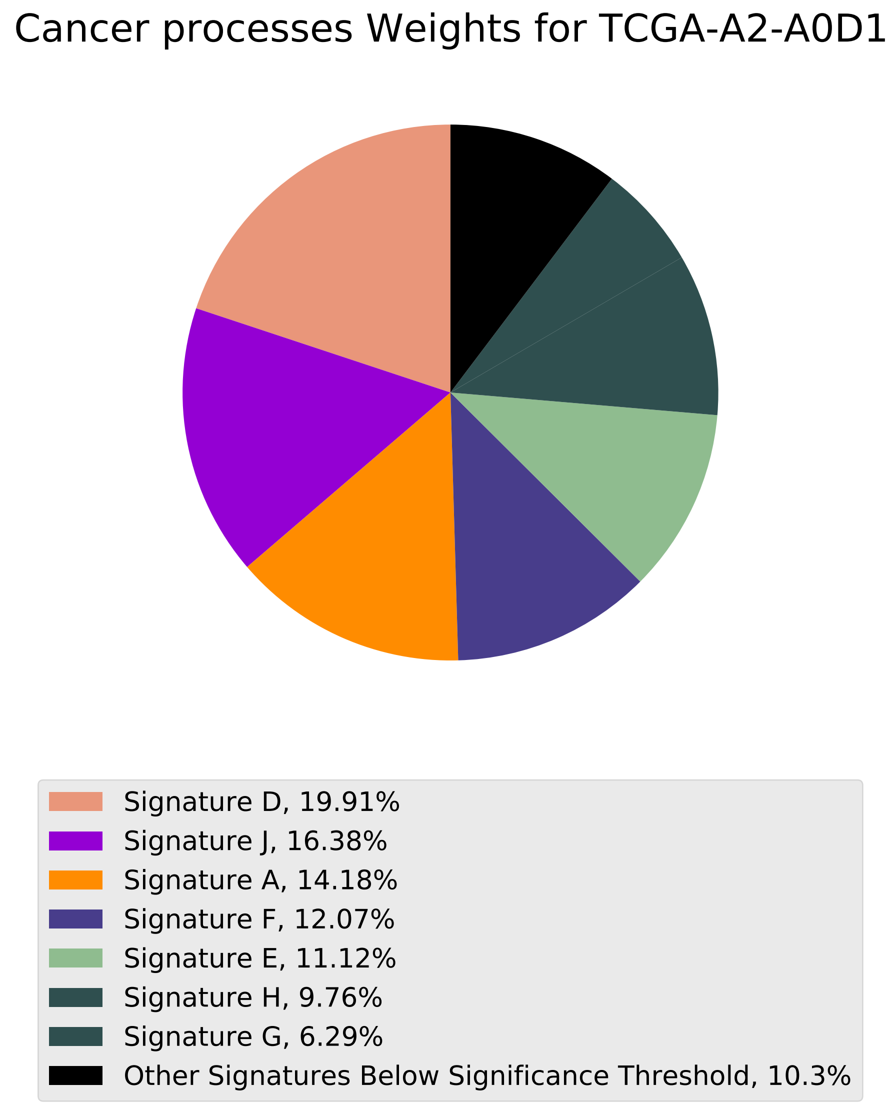

Cancer processes Weights for TCGA-A2-A0D1
Signature D, 19.91%
Signature J, 16.38%
Signature A, 14.18%
Signature F, 12.07%
Signature E, 11.12%
Signature H, 9.76%
Signature G, 6.29%
Other Signatures Below Significance Threshold, 10.3%

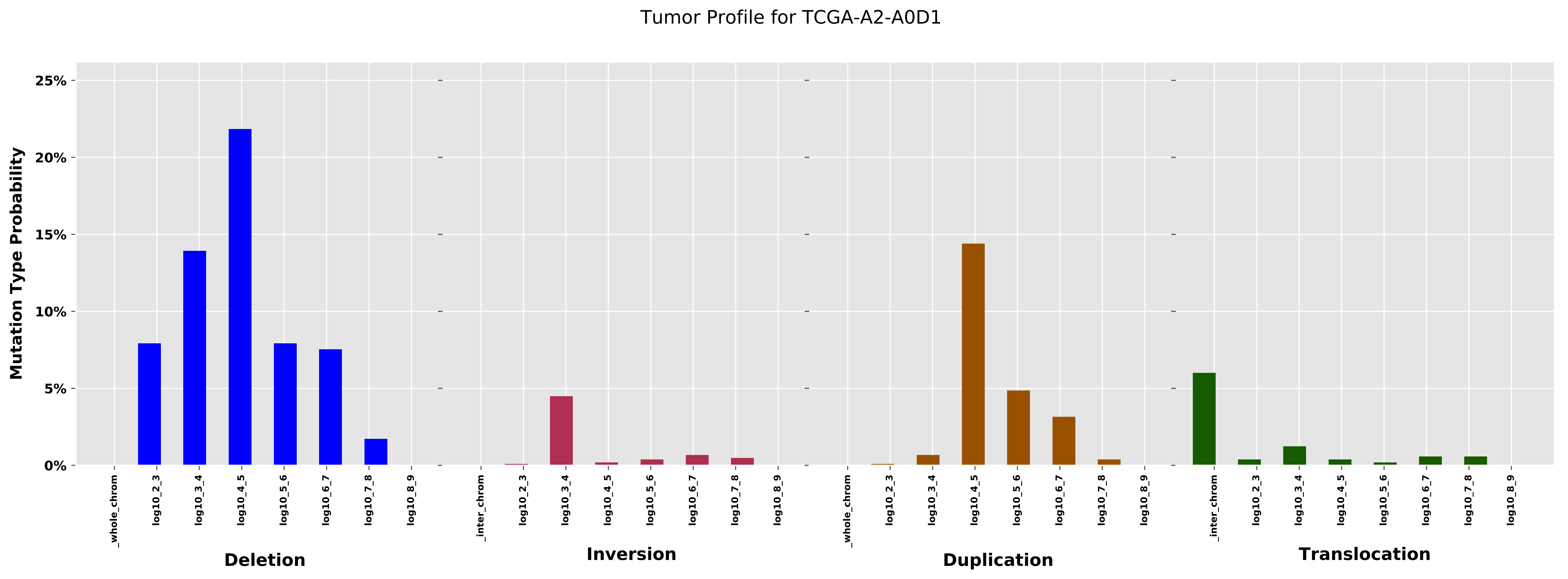
Tumor Profile for TCGA-A2-A0D1
Mutation Type Probability
25%
20%
15%
10%
5%
0%
_whole_chrom
log10_2_3
log10_3_4
log10_4_5
log10_5_6
log10_6_7
log10_7_8
log10_8_9
Deletion
_inter_chrom
log10_2_3
log10_3_4
log10_4_5
log10_5_6
log10_6_7
log10_7_8
log10_8_9
Inversion
_whole_chrom
log10_2_3
log10_3_4
log10_4_5
log10_5_6
log10_6_7
log10_7_8
log10_8_9
Duplication
_inter_chrom
log10_2_3
log10_3_4
log10_4_5
log10_5_6
log10_6_7
log10_7_8
log10_8_9
Translocation

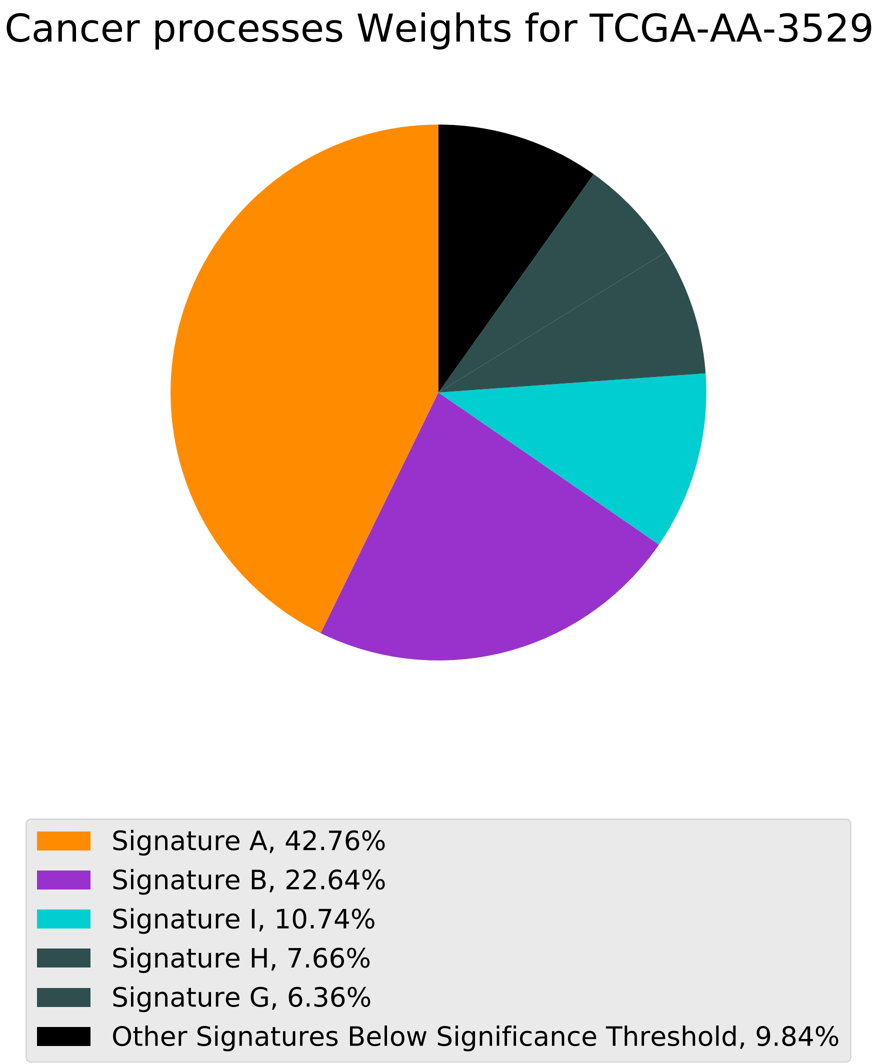
Cancer processes Weights for TCGA-AA-3529
Signature A, 42.76%
Signature B, 22.64%
Signature I, 10.74%
Signature H, 7.66%
Signature G, 6.36%
Other Signatures Below Significance Threshold, 9.84%

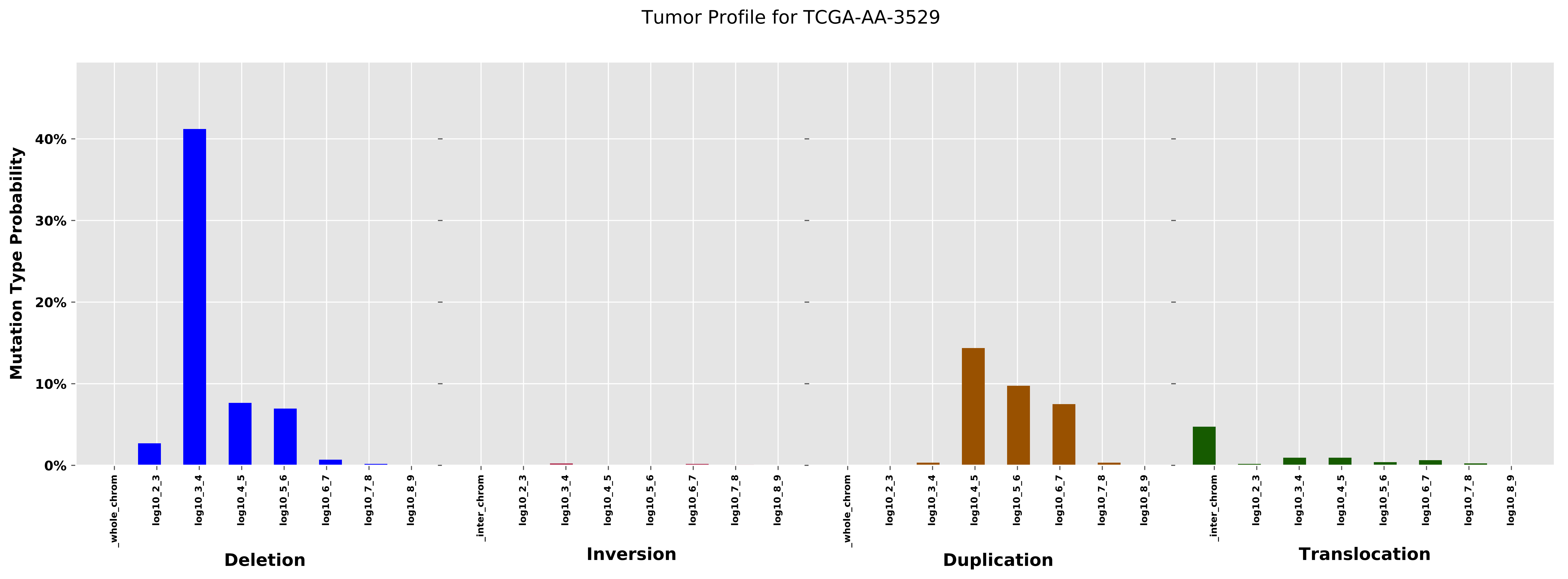
Tumor Profile for TCGA-AA-3529
Mutation Type Probability
40%
30%
20%
10%
0%
_whole_chrom
log10_2_3
log10_3_4
log10_4_5
log10_5_6
log10_6_7
log10_7_8
log10_8_9
_inter_chrom
Deletion
Inversion
Duplication
Translocation

Cancer processes Weights for TCGA-A6-3807

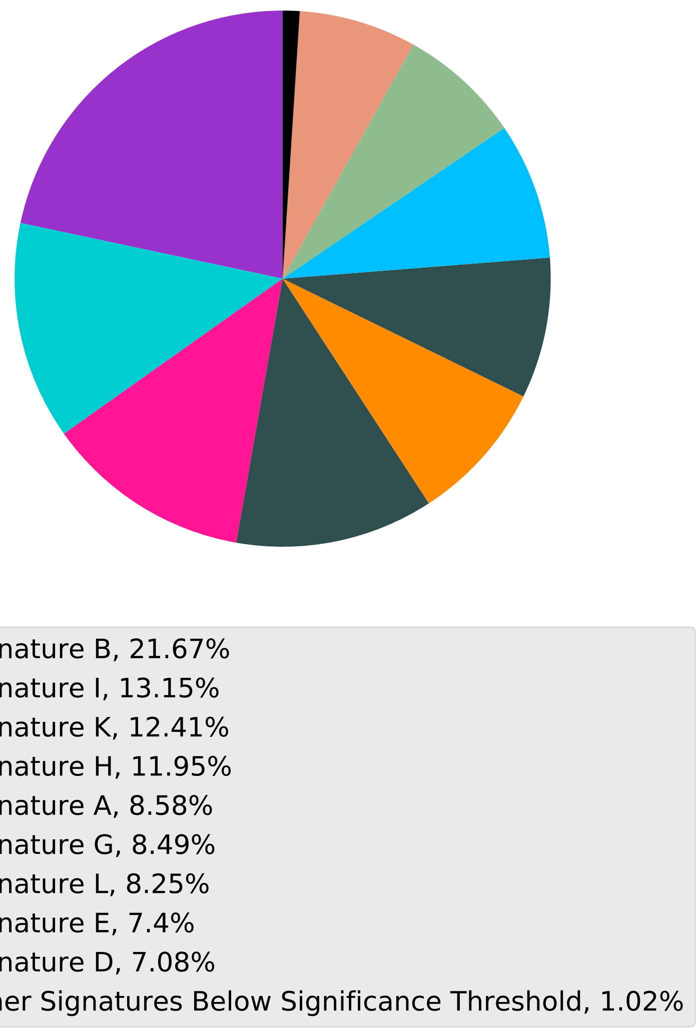

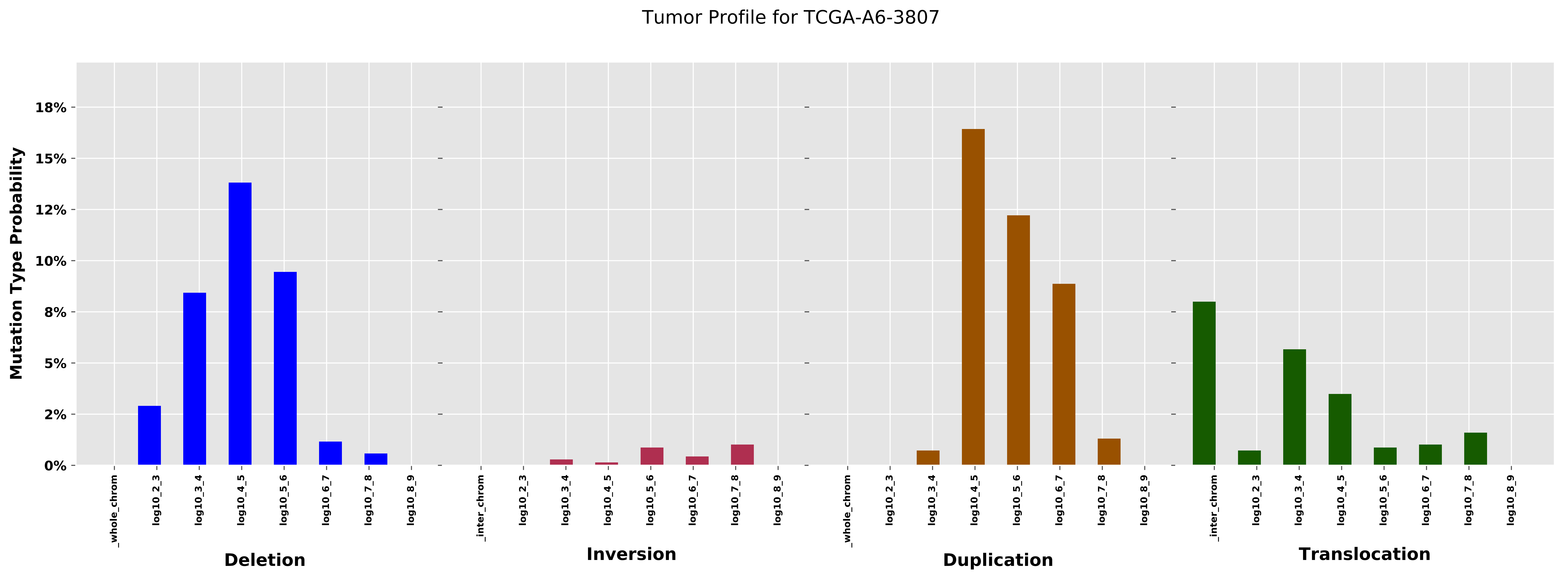

Tumor Profile for TCGA-A6-3807
Mutation Type Probability
18%
15%
12%
10%
8%
5%
2%
0%
_whole_chrom
log10_2_3
log10_3_4
log10_4_5
log10_5_6
log10_6_7
log10_7_8
log10_8_9
_inter_chrom
Deletion
Inversion
Duplication
Translocation

Cancer processes Weights for TCGA-B6-A0IQ

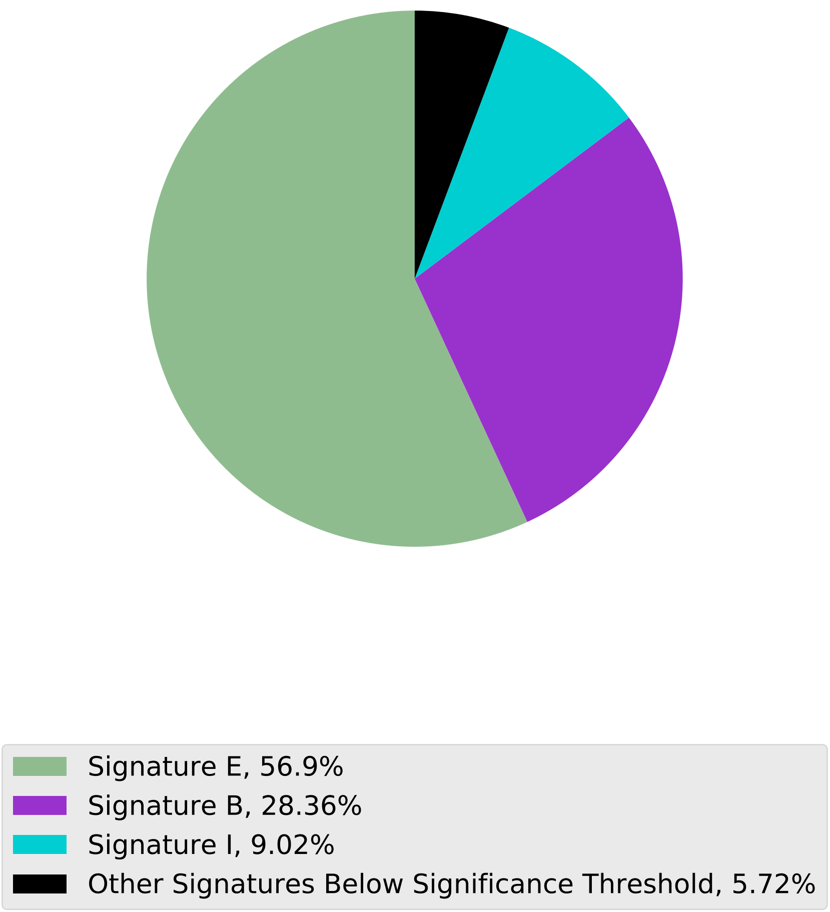

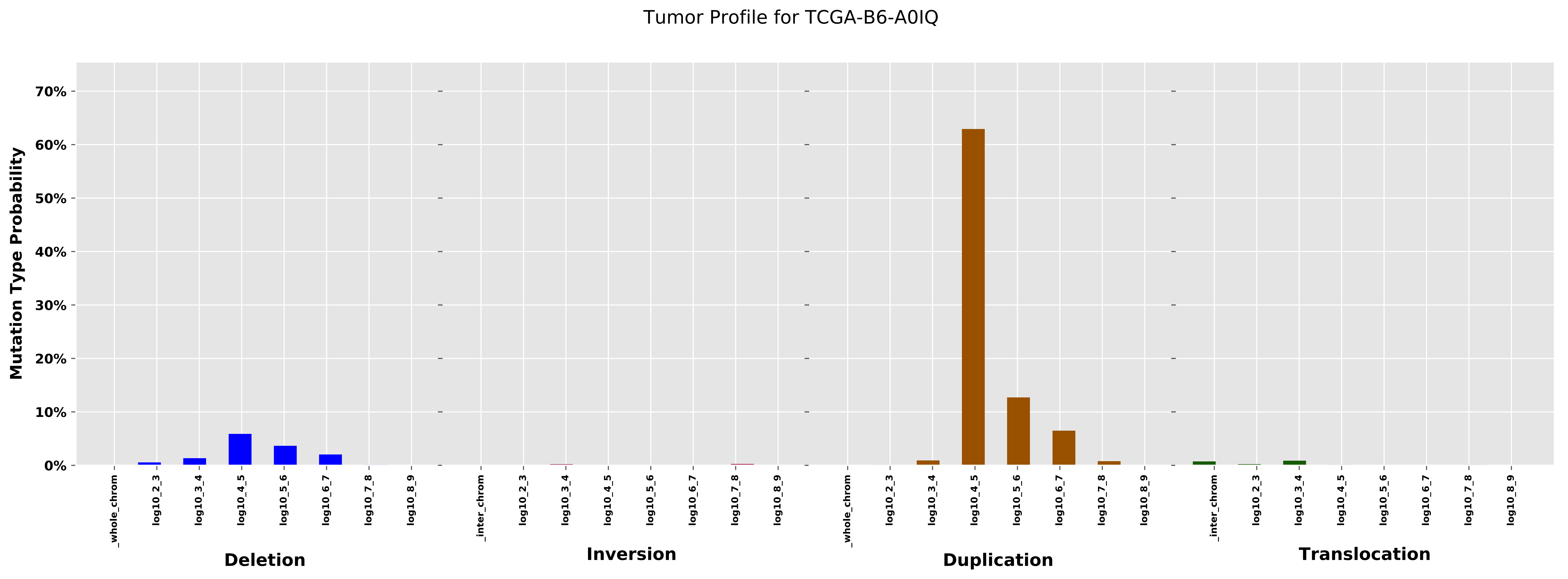

Tumor Profile for TCGA-B6-A0IQ
Mutation Type Probability
70%
60%
50%
40%
30%
20%
10%
0%
_whole_chrom
log10_2_3
log10_3_4
log10_4_5
log10_5_6
log10_6_7
log10_7_8
log10_8_9
_inter_chrom
log10_2_3
log10_3_4
log10_4_5
log10_5_6
log10_6_7
log10_7_8
log10_8_9
_whole_chrom
log10_2_3
log10_3_4
log10_4_5
log10_5_6
log10_6_7
log10_7_8
log10_8_9
_inter_chrom
log10_2_3
log10_3_4
log10_4_5
log10_5_6
log10_6_7
log10_7_8
log10_8_9
Deletion
Inversion
Duplication
Translocation

#### Cancer processes Weights for TCGA-A2-A0YG

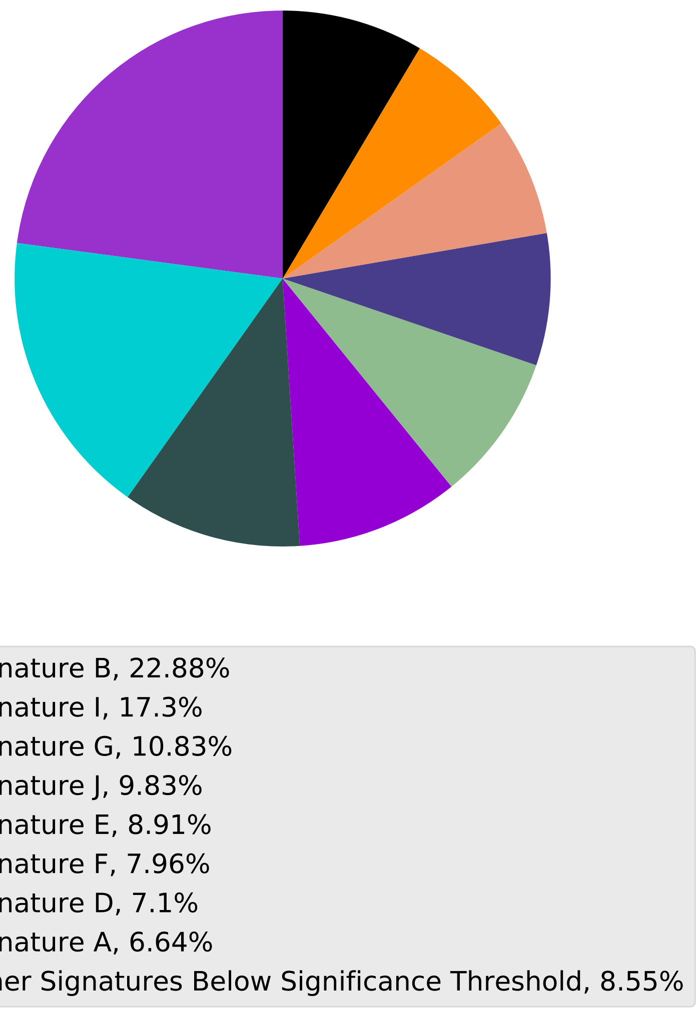

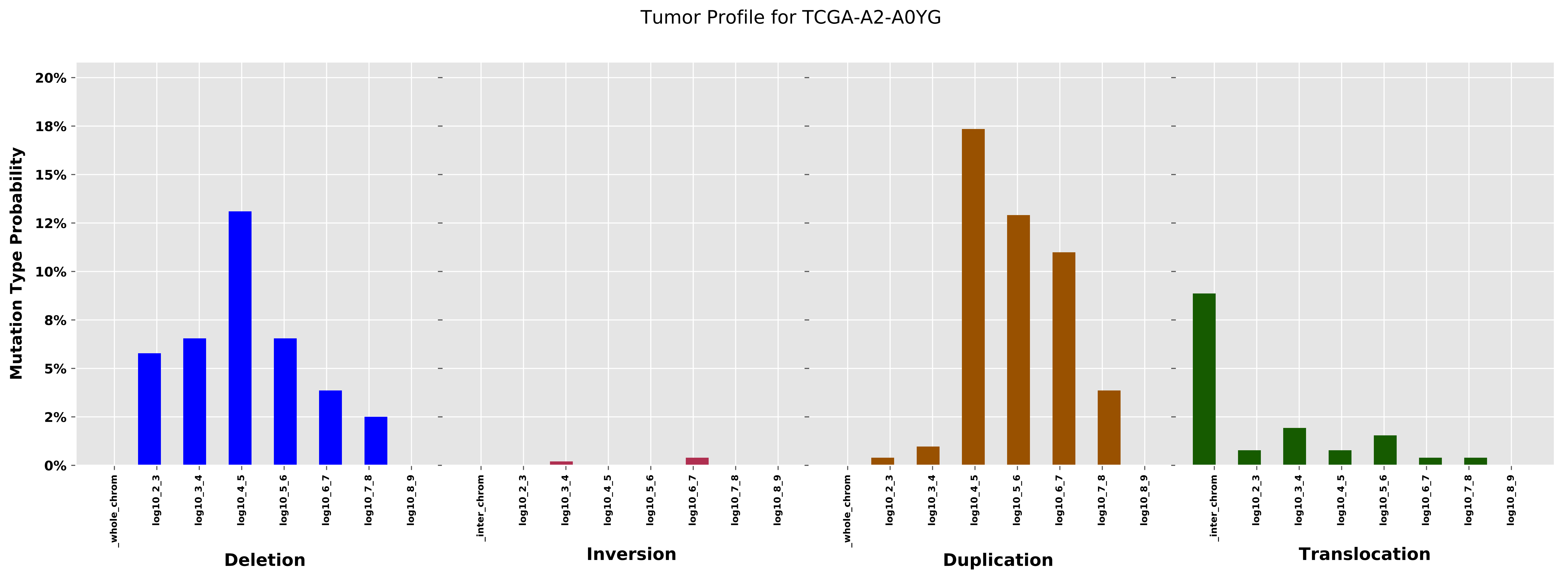

Tumor Profile for TCGA-A2-A0YG
Mutation Type Probability
20%
18%
15%
12%
10%
8%
5%
2%
0%
_whole_chrom
log10_2_3
log10_3_4
log10_4_5
log10_5_6
log10_6_7
log10_7_8
log10_8_9
_inter_chrom
Deletion
Inversion
Duplication
Translocation

#### Cancer processes Weights for TCGA-C8-A12Q

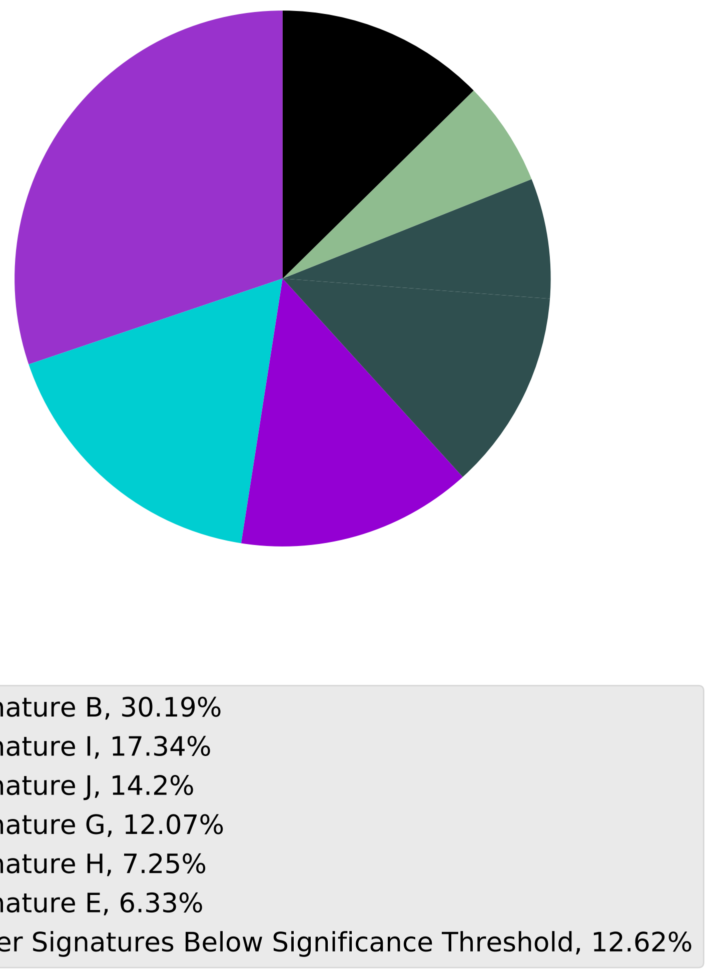

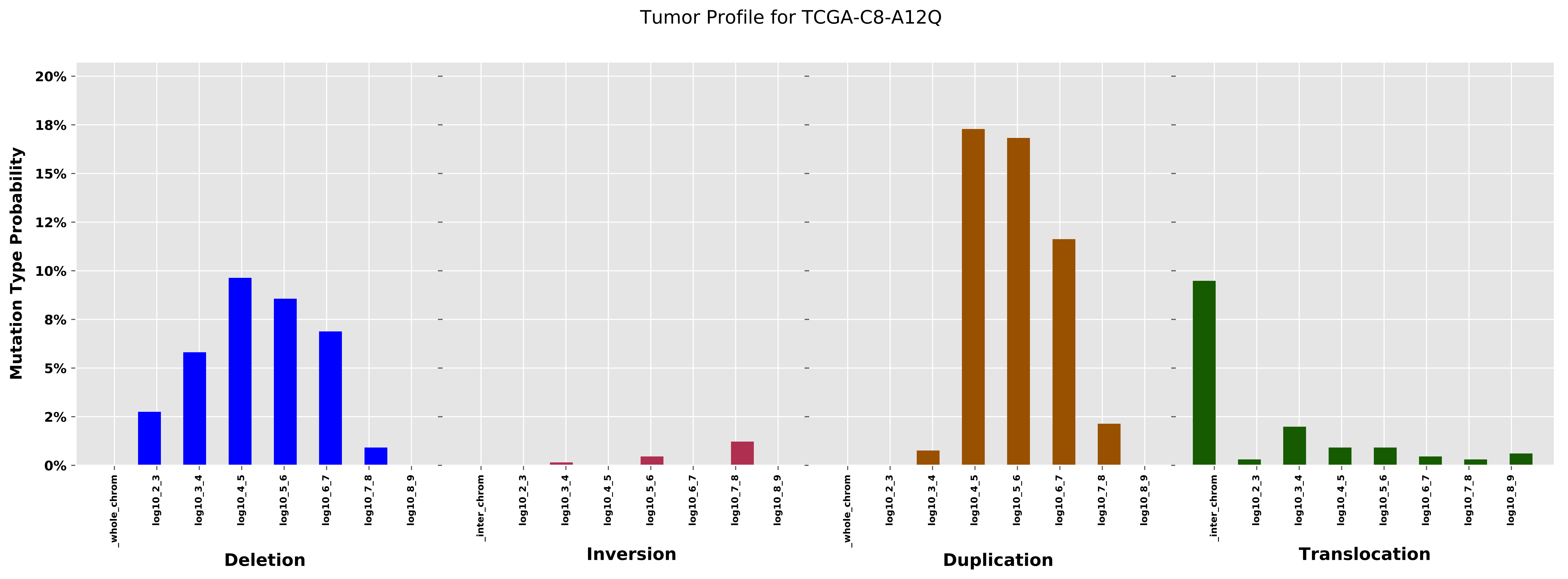

Tumor Profile for TCGA-C8-A12Q
Mutation Type Probability
20%
18%
15%
12%
10%
8%
5%
2%
0%
_whole_chrom
log10_2_3
log10_3_4
log10_4_5
log10_5_6
log10_6_7
log10_7_8
log10_8_9
_inter_chrom
Deletion
Inversion
Duplication
Translocation

Cancer processes Weights for TCGA-BH-A0BW

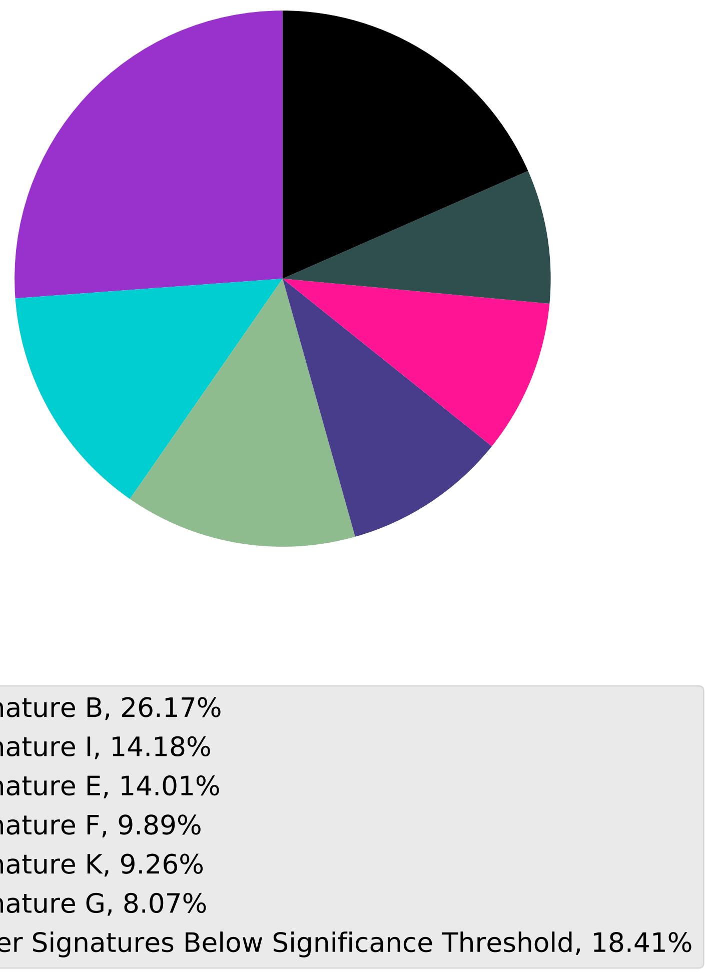

Tumor Profile for TCGA-BH-A0BW

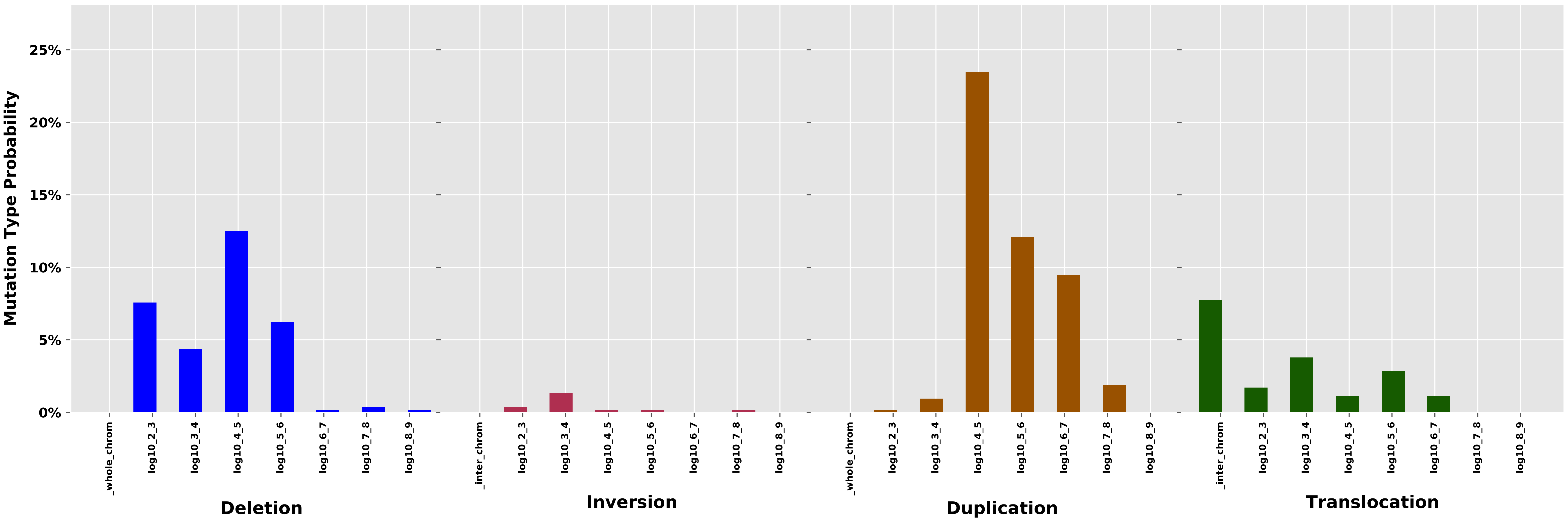

Cancer processes Weights for TCGA-A2-A0EY

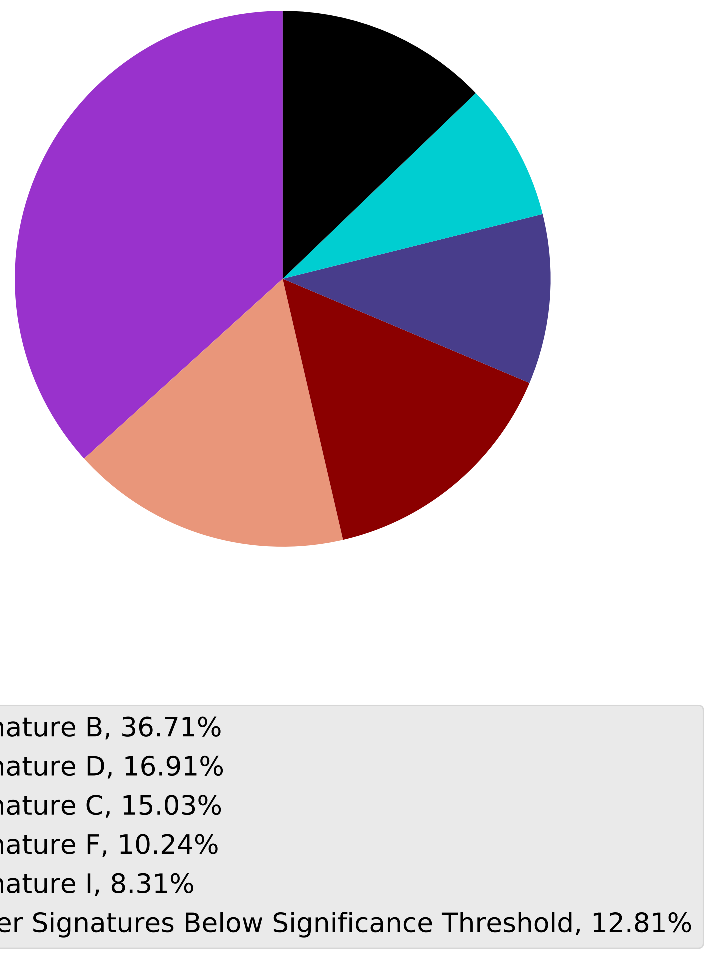

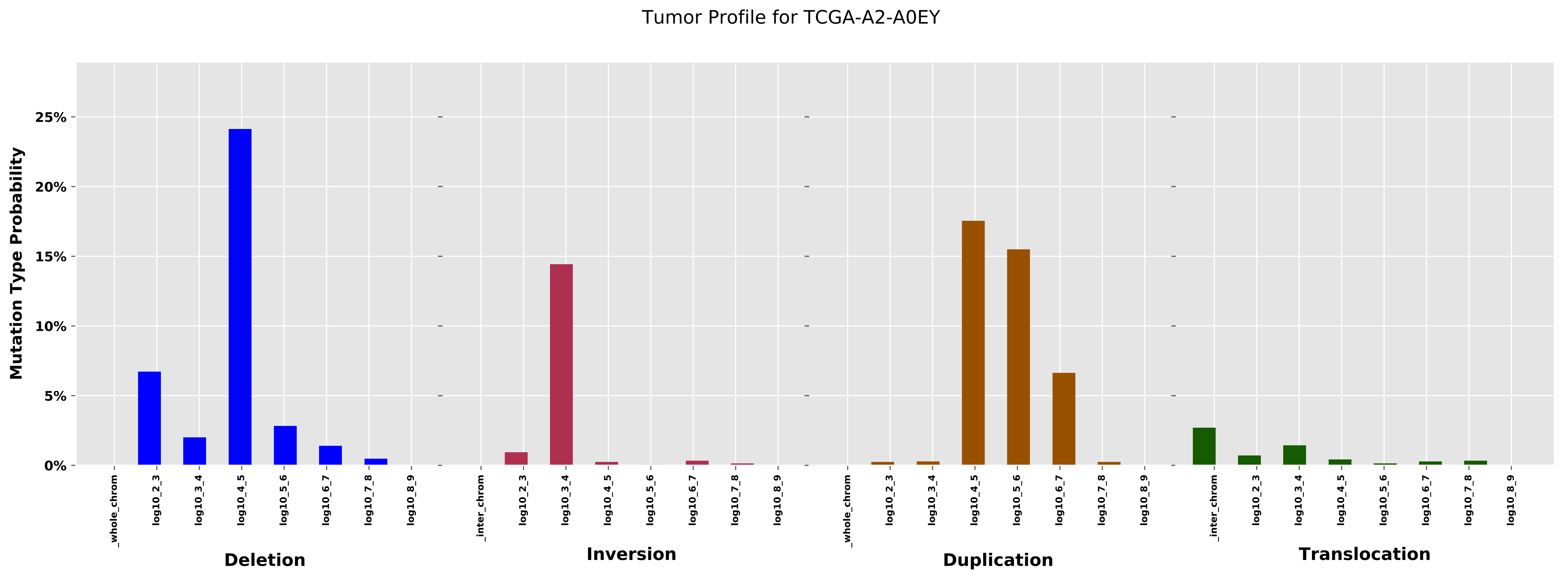

Tumor Profile for TCGA-A2-A0EY
Mutation Type Probability
25%
20%
15%
10%
5%
0%
_whole_chrom
log10_2_3
log10_3_4
log10_4_5
log10_5_6
log10_6_7
log10_7_8
log10_8_9
Deletion
_inter_chrom
log10_2_3
log10_3_4
log10_4_5
log10_5_6
log10_6_7
log10_7_8
log10_8_9
Inversion
_whole_chrom
log10_2_3
log10_3_4
log10_4_5
log10_5_6
log10_6_7
log10_7_8
log10_8_9
Duplication
_inter_chrom
log10_2_3
log10_3_4
log10_4_5
log10_5_6
log10_6_7
log10_7_8
log10_8_9
Translocation

Cancer processes Weights for TCGA-EW-A1PB

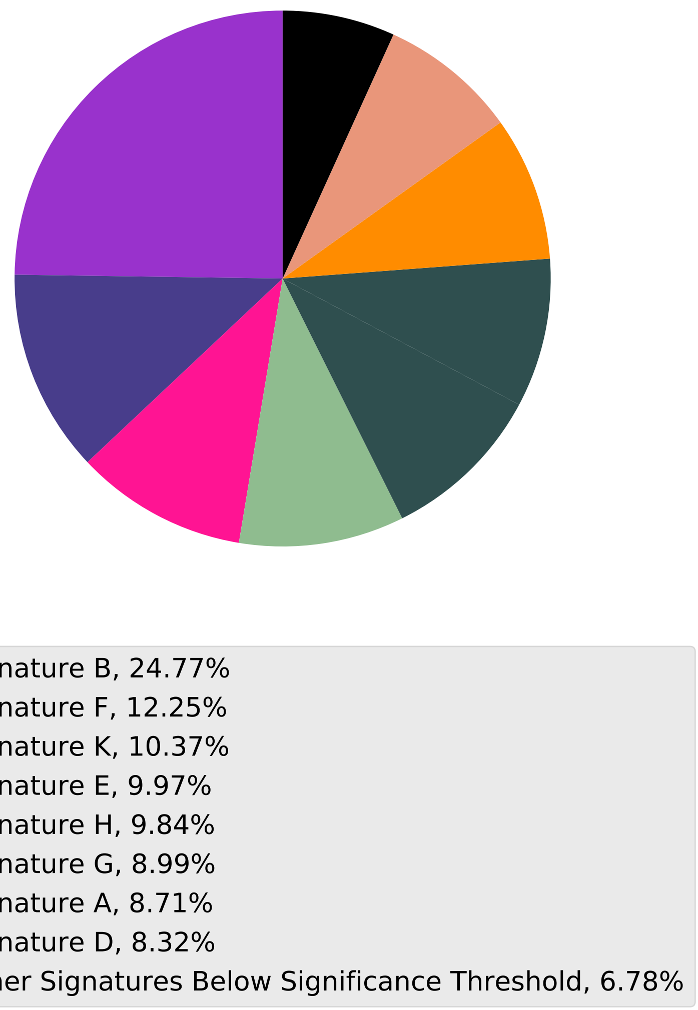

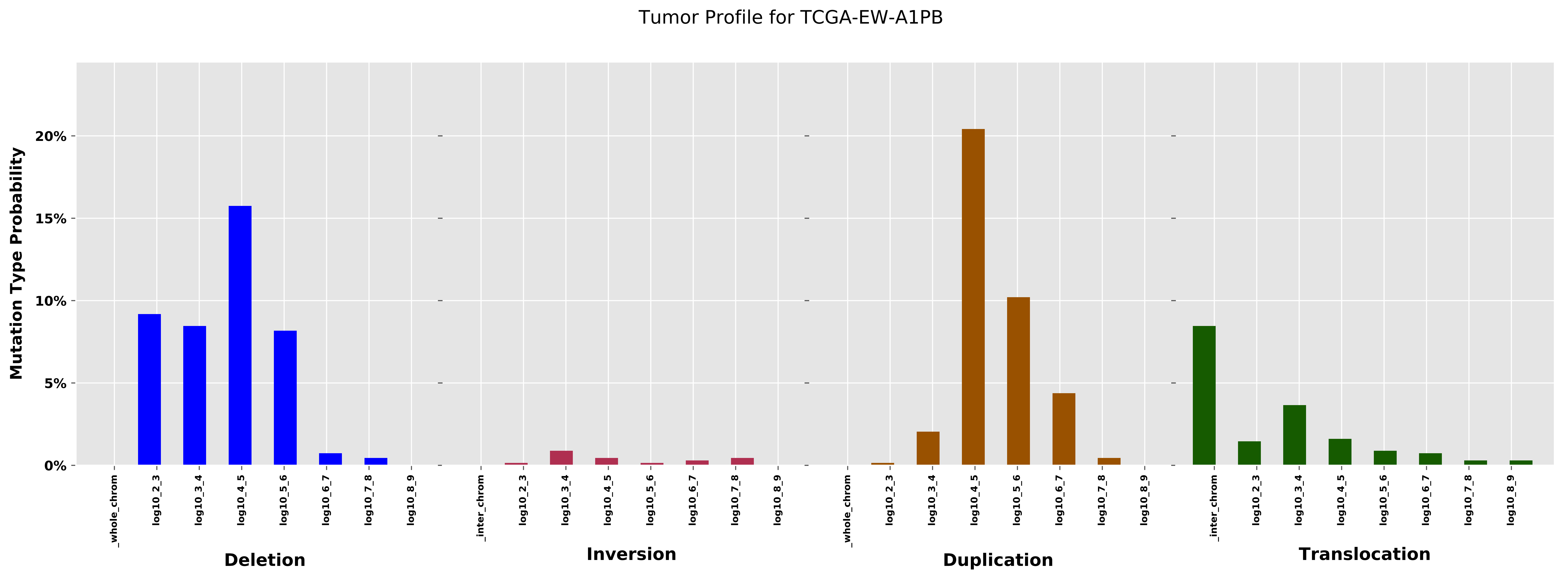

Tumor Profile for TCGA-EW-A1PB
Mutation Type Probability
0%
5%
10%
15%
20%
_whole_chrom
log10_2_3
log10_3_4
log10_4_5
log10_5_6
log10_6_7
log10_7_8
log10_8_9
_inter_chrom
Deletion
Inversion
Duplication
Translocation

Cancer processes Weights for TCGA-AD-6964

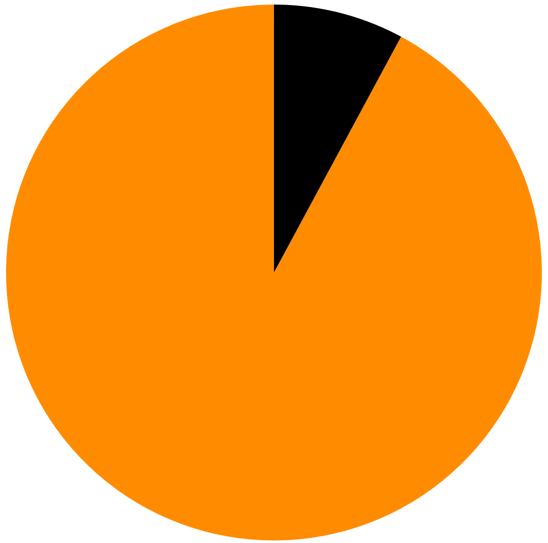

Signature A, 92.13%
Other Signatures Below Significance Threshold, 7.87%

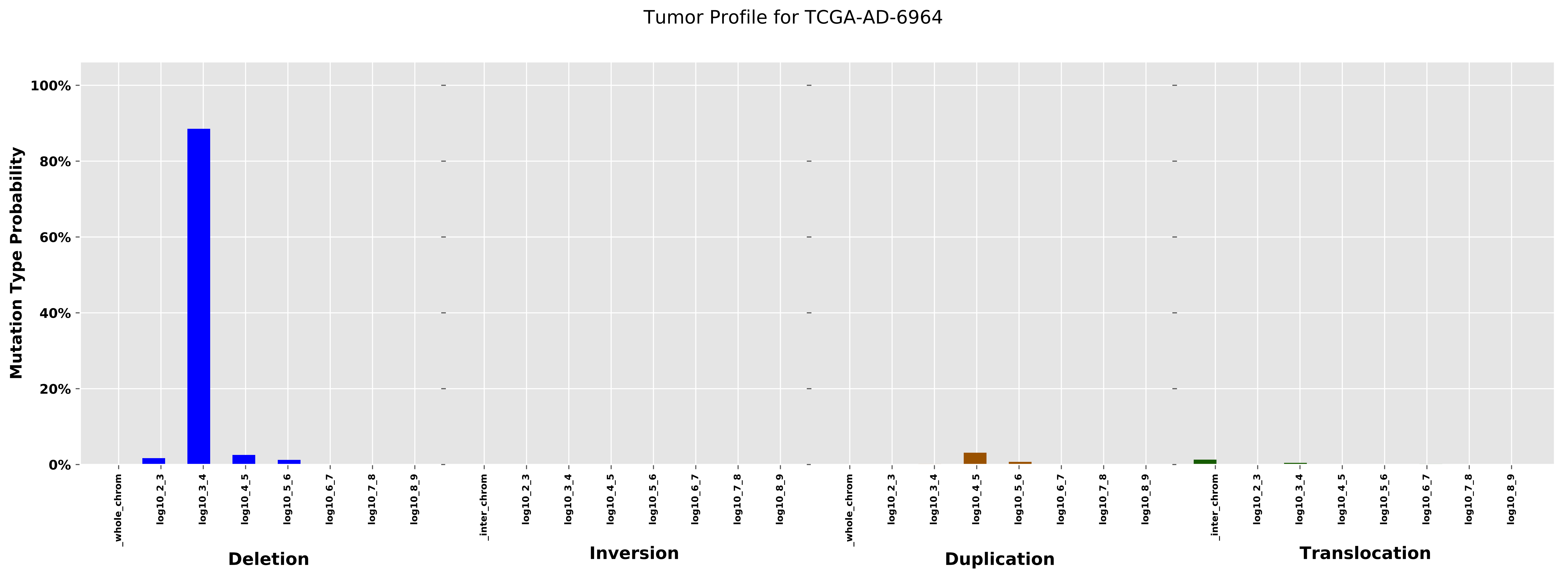

Tumor Profile for TCGA-AD-6964
Mutation Type Probability
100%
80%
60%
40%
20%
0%
_whole_chrom
log10_2_3
log10_3_4
log10_4_5
log10_5_6
log10_6_7
log10_7_8
log10_8_9
Deletion
_inter_chrom
Inversion
Duplication
Translocation

Cancer processes Weights for TCGA-AG-3896

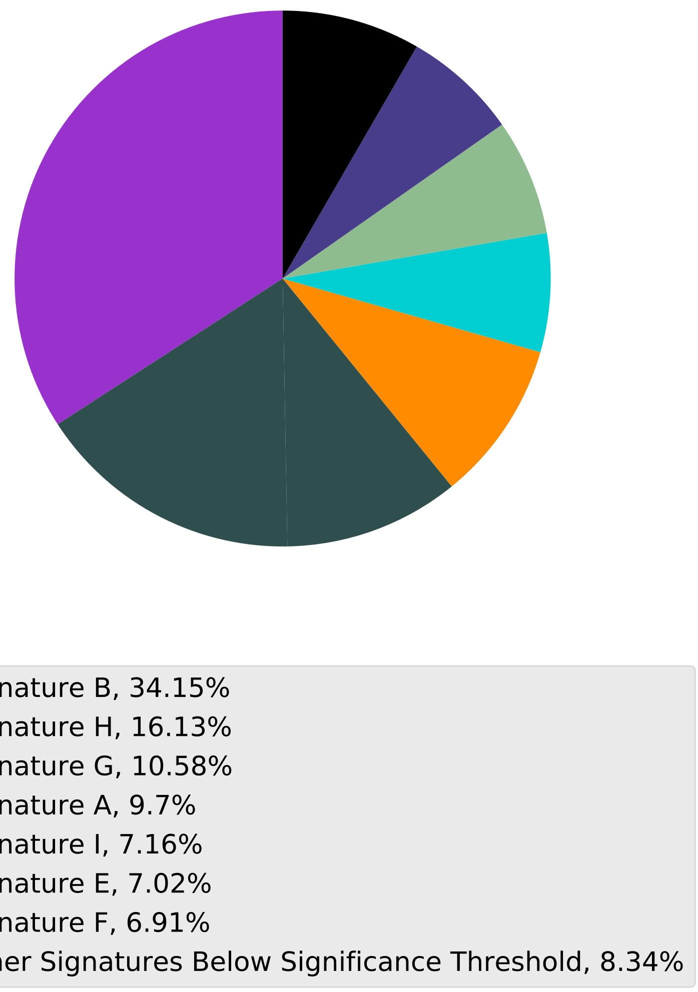

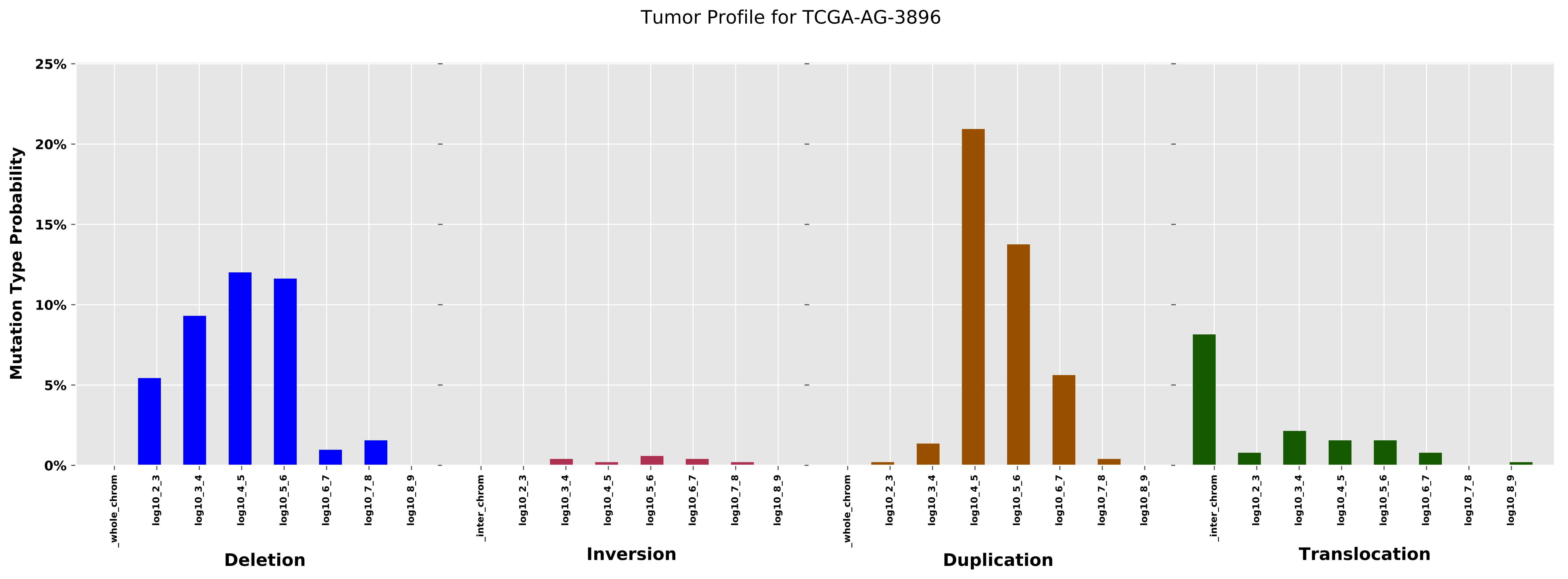
Tumor Profile for TCGA-AG-3896
Mutation Type Probability
25%
20%
15%
10%
5%
0%
_whole_chrom
log10_2_3
log10_3_4
log10_4_5
log10_5_6
log10_6_7
log10_7_8
log10_8_9
Deletion
_inter_chrom
log10_2_3
log10_3_4
log10_4_5
log10_5_6
log10_6_7
log10_7_8
log10_8_9
Inversion
_whole_chrom
log10_2_3
log10_3_4
log10_4_5
log10_5_6
log10_6_7
log10_7_8
log10_8_9
Duplication
_inter_chrom
log10_2_3
log10_3_4
log10_4_5
log10_5_6
log10_6_7
log10_7_8
log10_8_9
Translocation

Cancer processes Weights for TCGA-E2-A15K

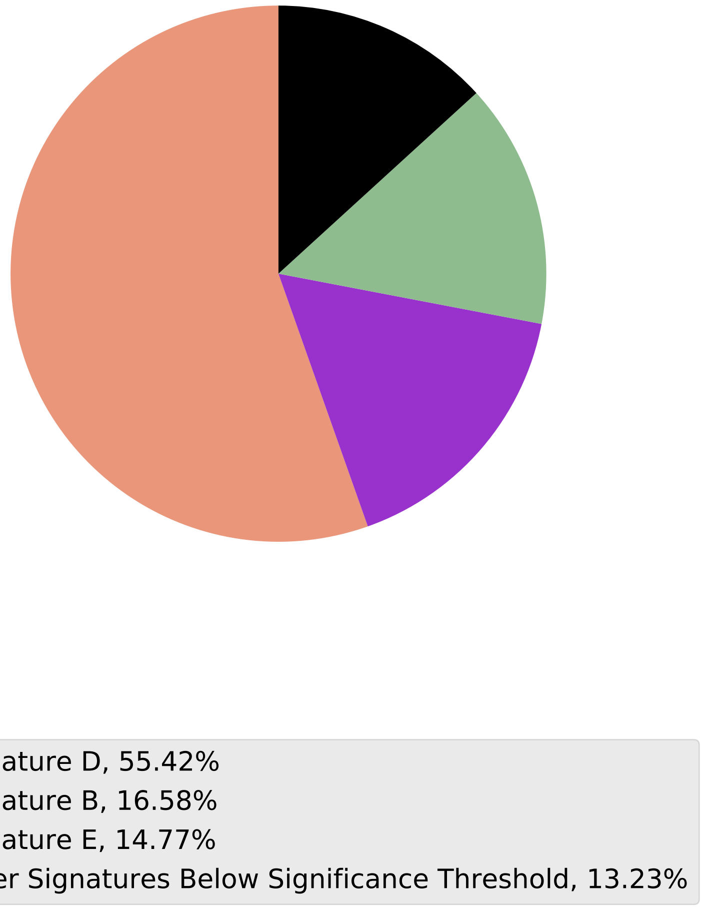

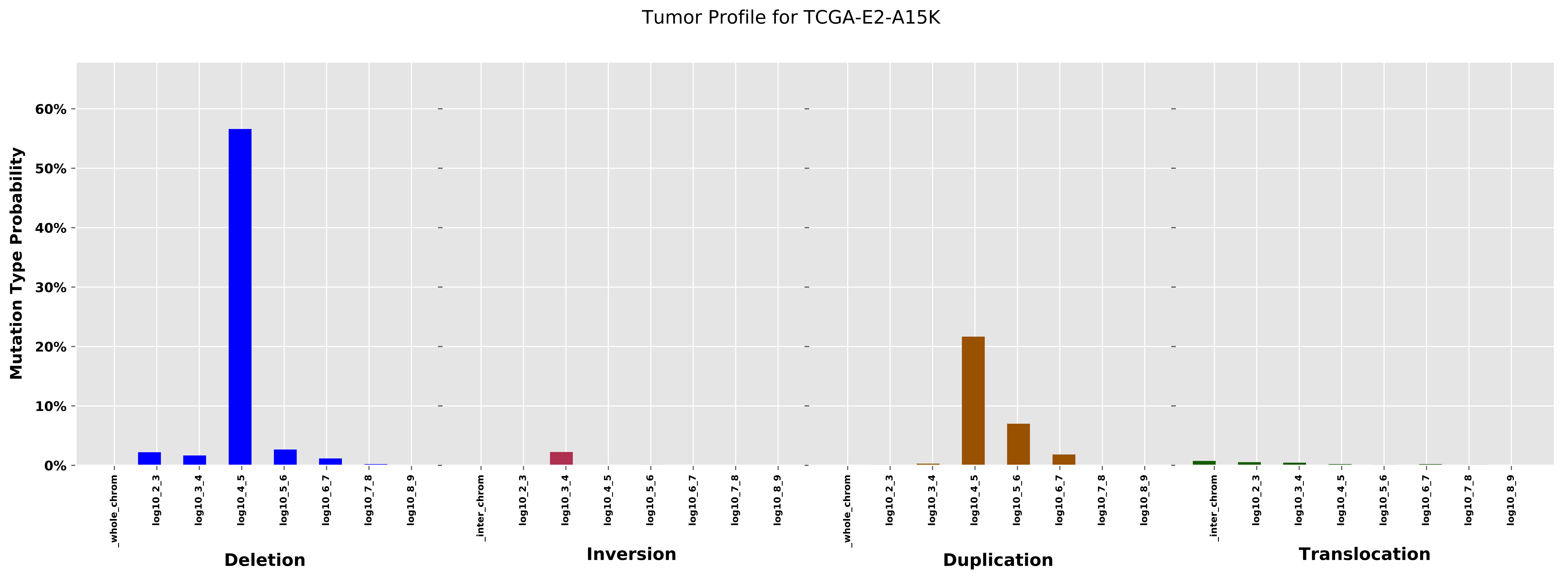

Tumor Profile for TCGA-E2-A15K
Mutation Type Probability
60%
50%
40%
30%
20%
10%
0%
_whole_chrom
log10_2_3
log10_3_4
log10_4_5
log10_5_6
log10_6_7
log10_7_8
log10_8_9
_inter_chrom
Deletion
Inversion
Duplication
Translocation

Cancer processes Weights for TCGA-B6-A0RE

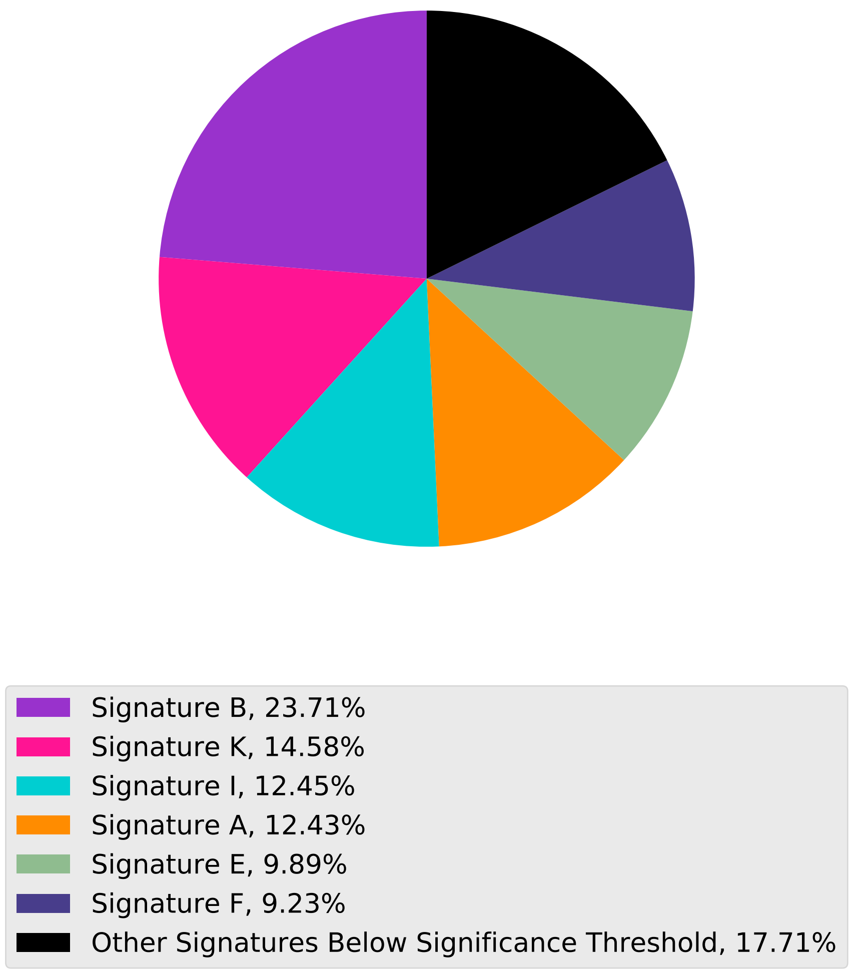

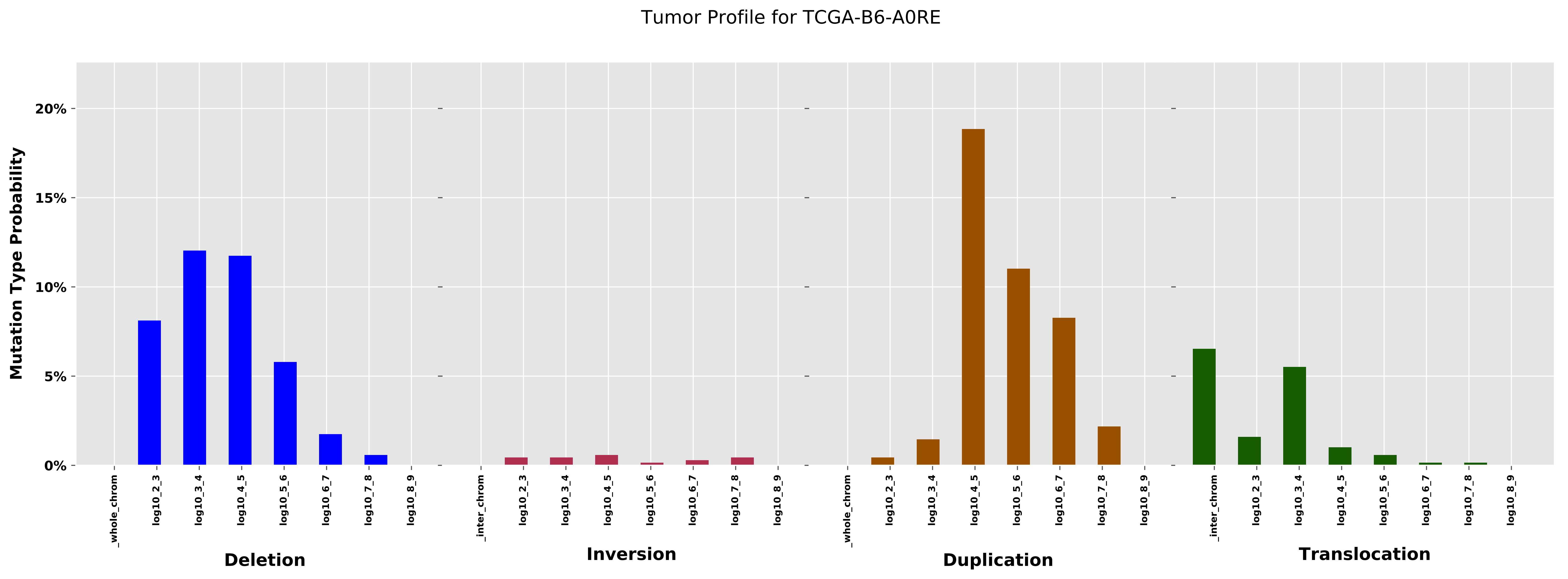

Tumor Profile for TCGA-B6-A0RE
Mutation Type Probability
0%
5%
10%
15%
20%
_whole_chrom
log10_2_3
log10_3_4
log10_4_5
log10_5_6
log10_6_7
log10_7_8
log10_8_9
_inter_chrom
Deletion
Inversion
Duplication
Translocation

Cancer processes Weights for TCGA-A8-A08S

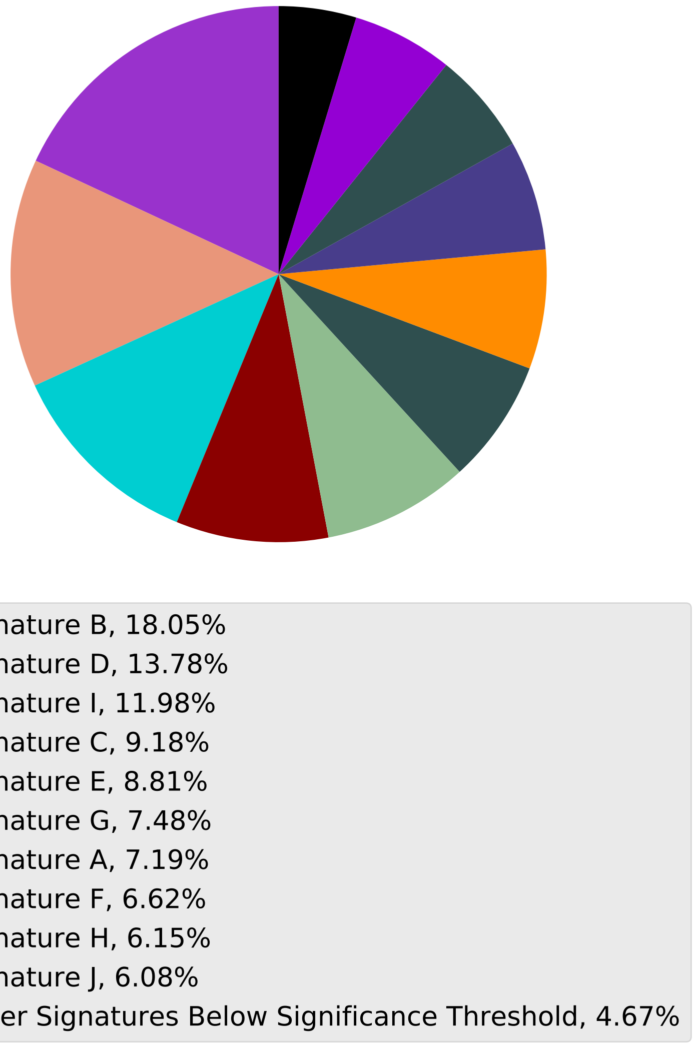

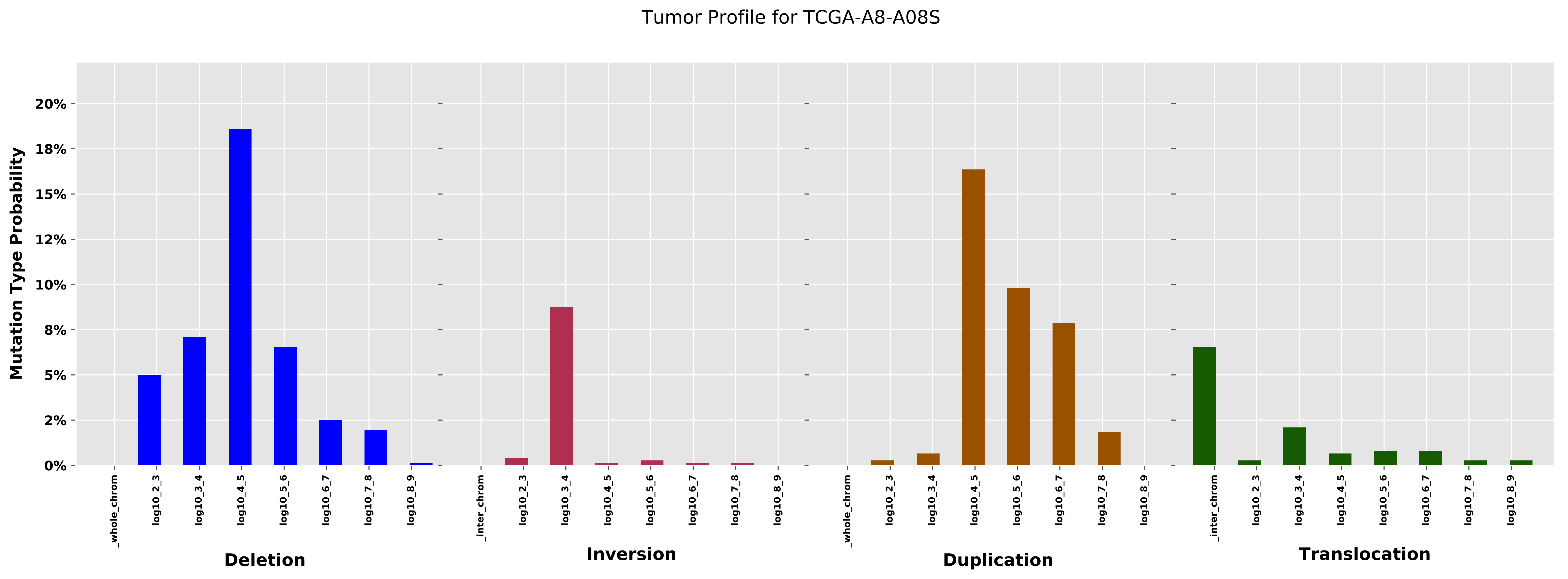
Tumor Profile for TCGA-A8-A08S
Mutation Type Probability
0%
2%
5%
8%
10%
12%
15%
18%
20%
_whole_chrom
log10_2_3
log10_3_4
log10_4_5
log10_5_6
log10_6_7
log10_7_8
log10_8_9
Deletion
_inter_chrom
Inversion
Duplication
Translocation

Cancer processes Weights for TCGA-AA-A03F

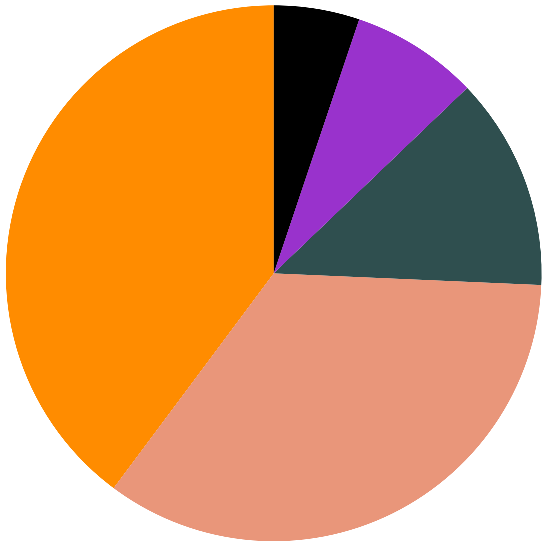

Signature A, 39.8%
Signature D, 34.5%
Signature H, 12.85%
Signature B, 7.68%
Other Signatures Below Significance Threshold, 5.17%

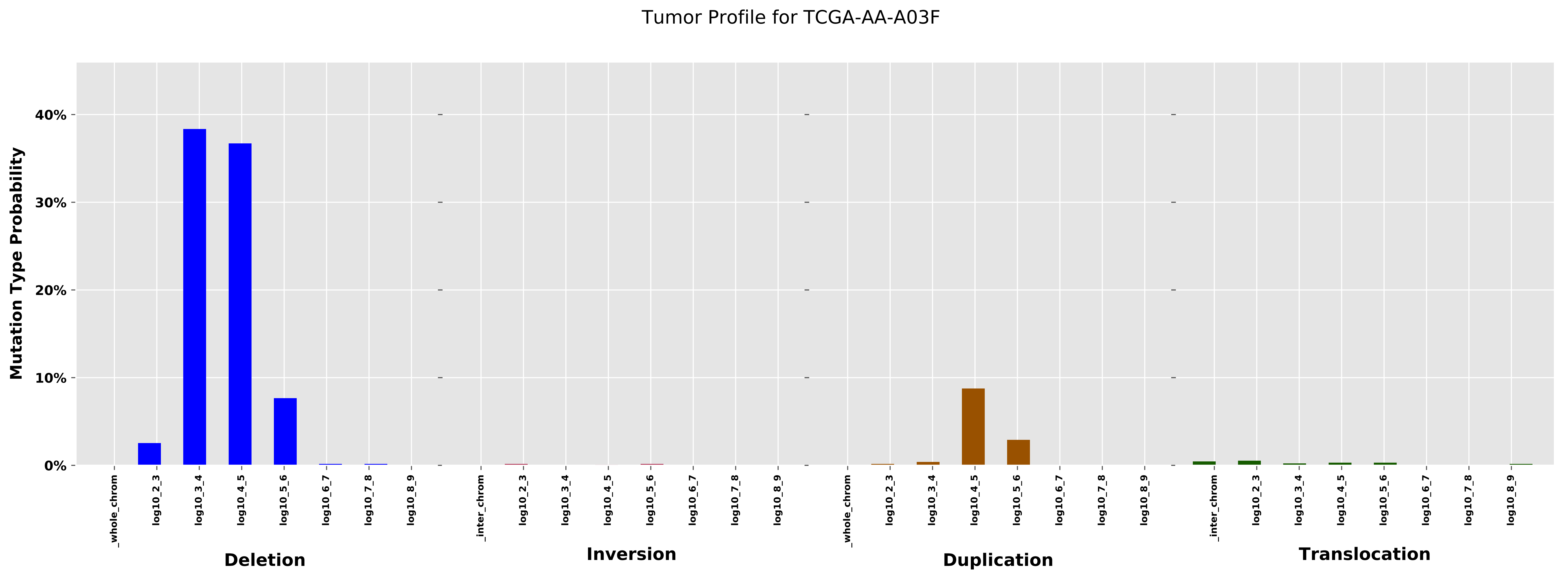
Tumor Profile for TCGA-AA-A03F
Mutation Type Probability
40%
30%
20%
10%
0%
_whole_chrom
log10_2_3
log10_3_4
log10_4_5
log10_5_6
log10_6_7
log10_7_8
log10_8_9
Deletion
_inter_chrom
log10_2_3
log10_3_4
log10_4_5
log10_5_6
log10_6_7
log10_7_8
log10_8_9
Inversion
_whole_chrom
log10_2_3
log10_3_4
log10_4_5
log10_5_6
log10_6_7
log10_7_8
log10_8_9
Duplication
_inter_chrom
log10_2_3
log10_3_4
log10_4_5
log10_5_6
log10_6_7
log10_7_8
log10_8_9
Translocation

Cancer processes Weights for TCGA-EW-A1PH

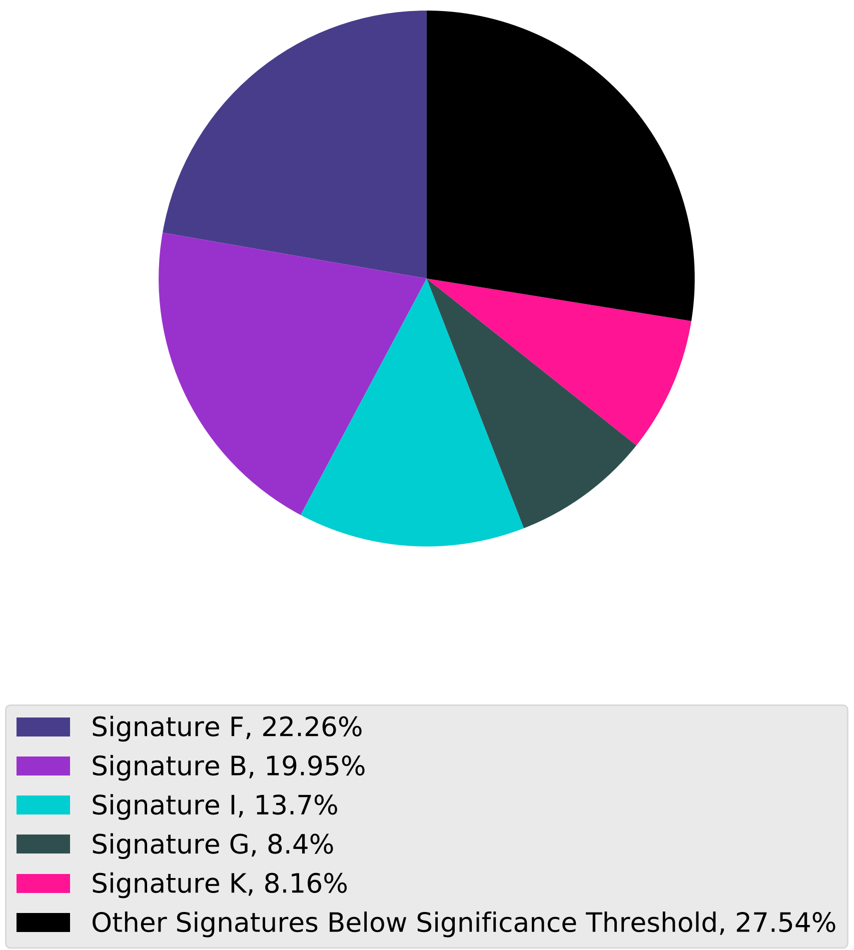

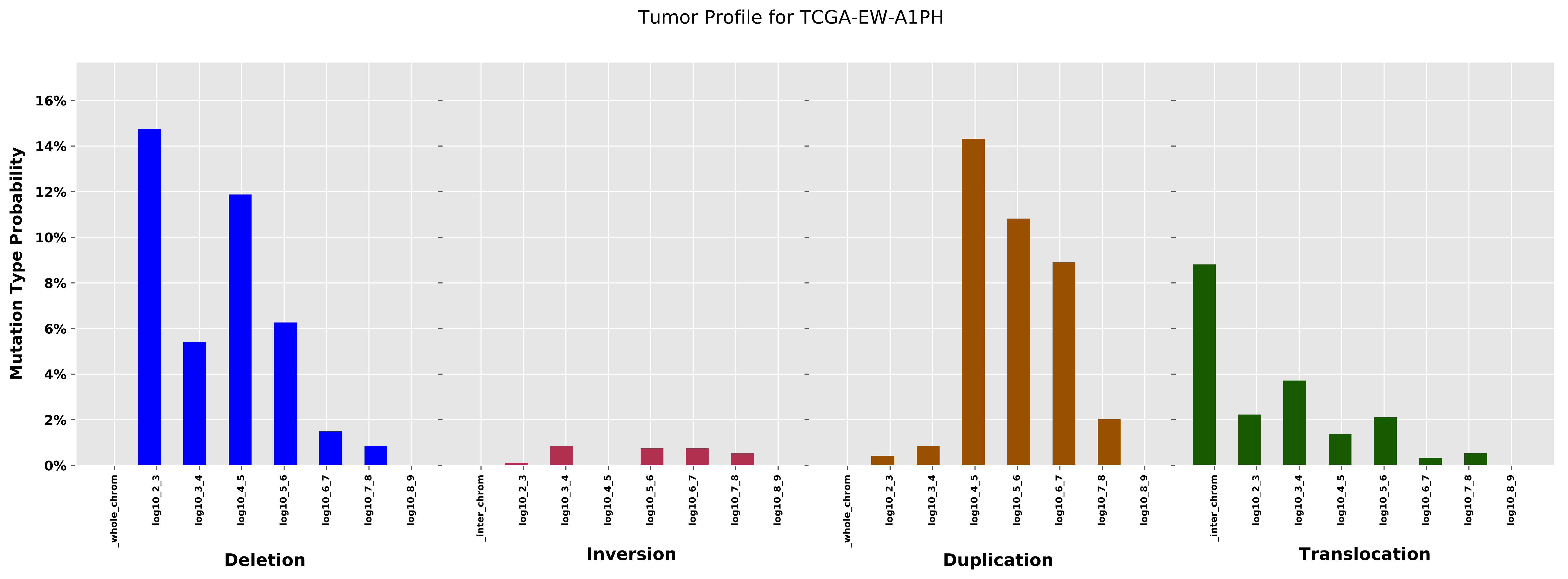
Tumor Profile for TCGA-EW-A1PH
Mutation Type Probability
0%
2%
4%
6%
8%
10%
12%
14%
16%
_whole_chrom
log10_2_3
log10_3_4
log10_4_5
log10_5_6
log10_6_7
log10_7_8
log10_8_9
_inter_chrom
Deletion
Inversion
Duplication
Translocation

Cancer processes Weights for TCGA-E2-A1LG

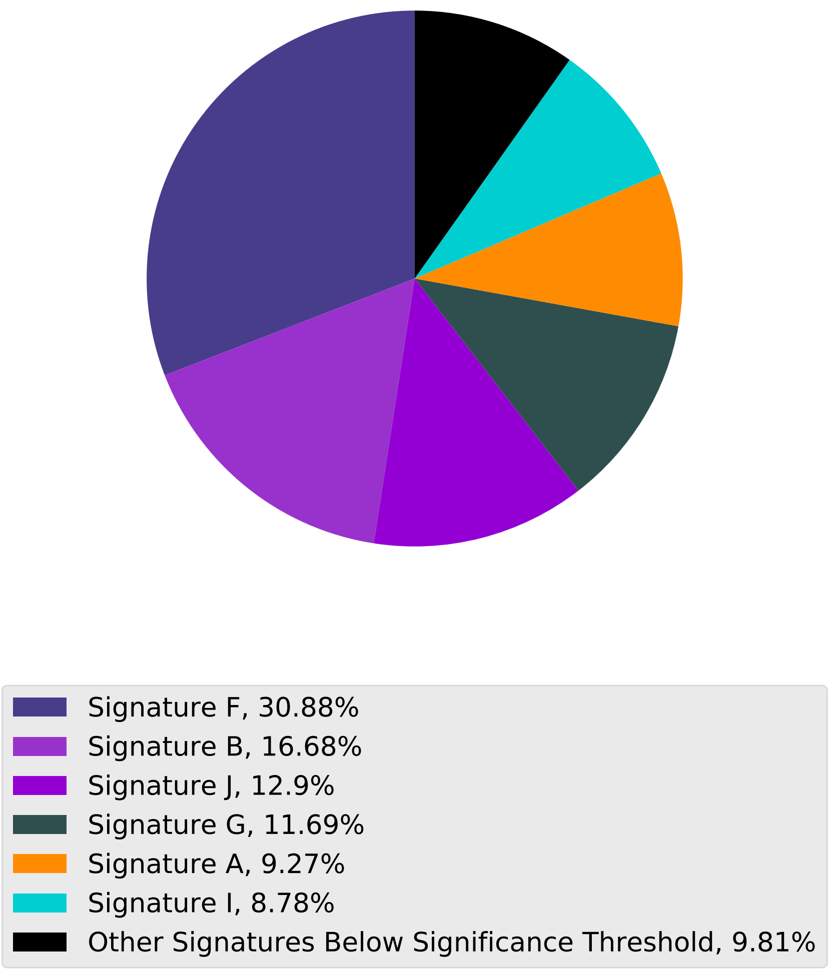

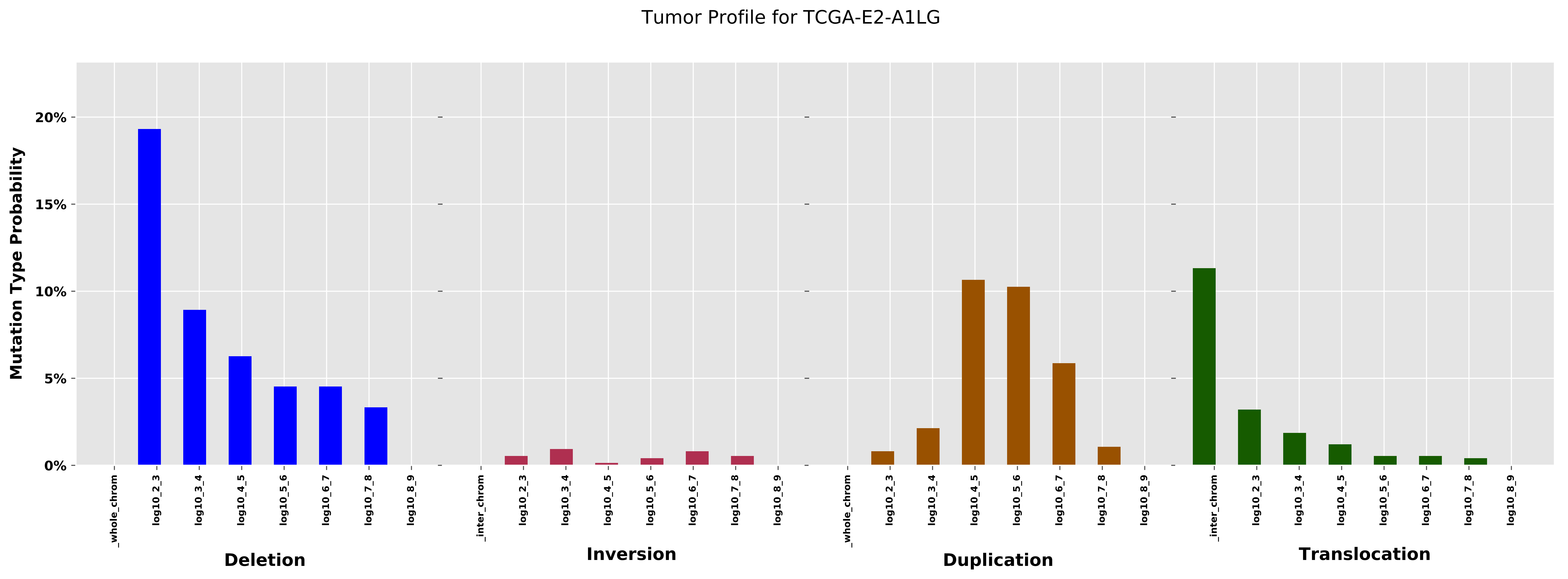

Tumor Profile for TCGA-E2-A1LG
Mutation Type Probability
0%
5%
10%
15%
20%
_whole_chrom
log10_2_3
log10_3_4
log10_4_5
log10_5_6
log10_6_7
log10_7_8
log10_8_9
_inter_chrom
Deletion
Inversion
Duplication
Translocation

Cancer processes Weights for TCGA-A8-A08B

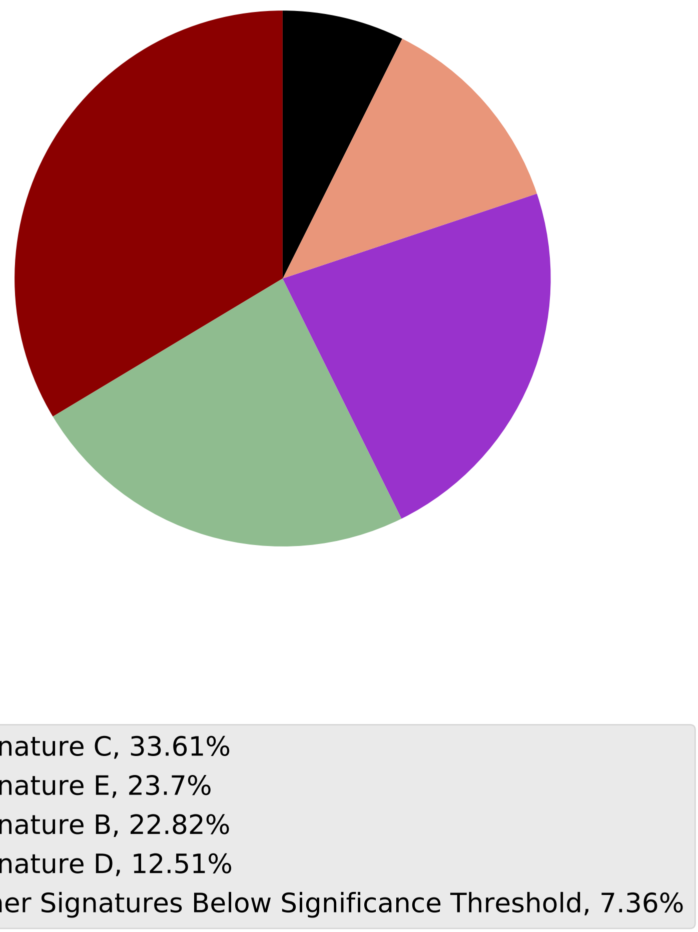

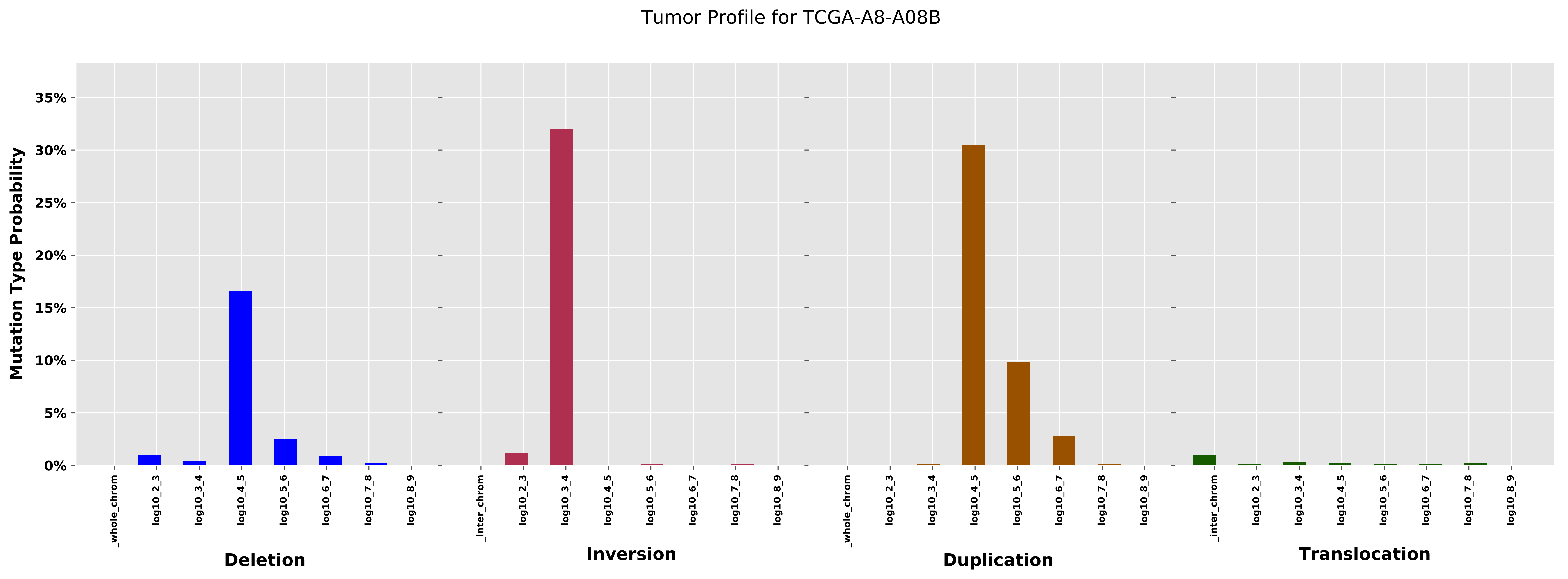

Tumor Profile for TCGA-A8-A08B
Mutation Type Probability
0%
5%
10%
15%
20%
25%
30%
35%
_whole_chrom
log10_2_3
log10_3_4
log10_4_5
log10_5_6
log10_6_7
log10_7_8
log10_8_9
_inter_chrom
Deletion
Inversion
Duplication
Translocation

Cancer processes Weights for TCGA-B6-A0RT

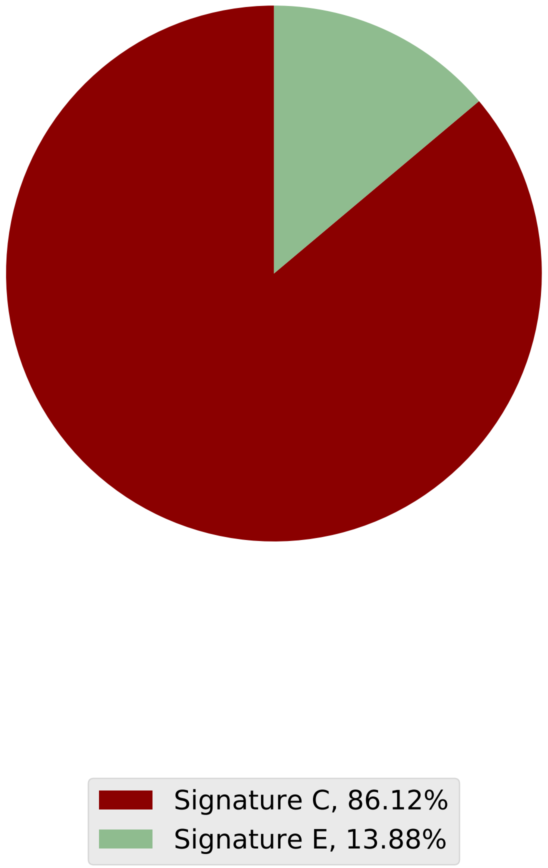

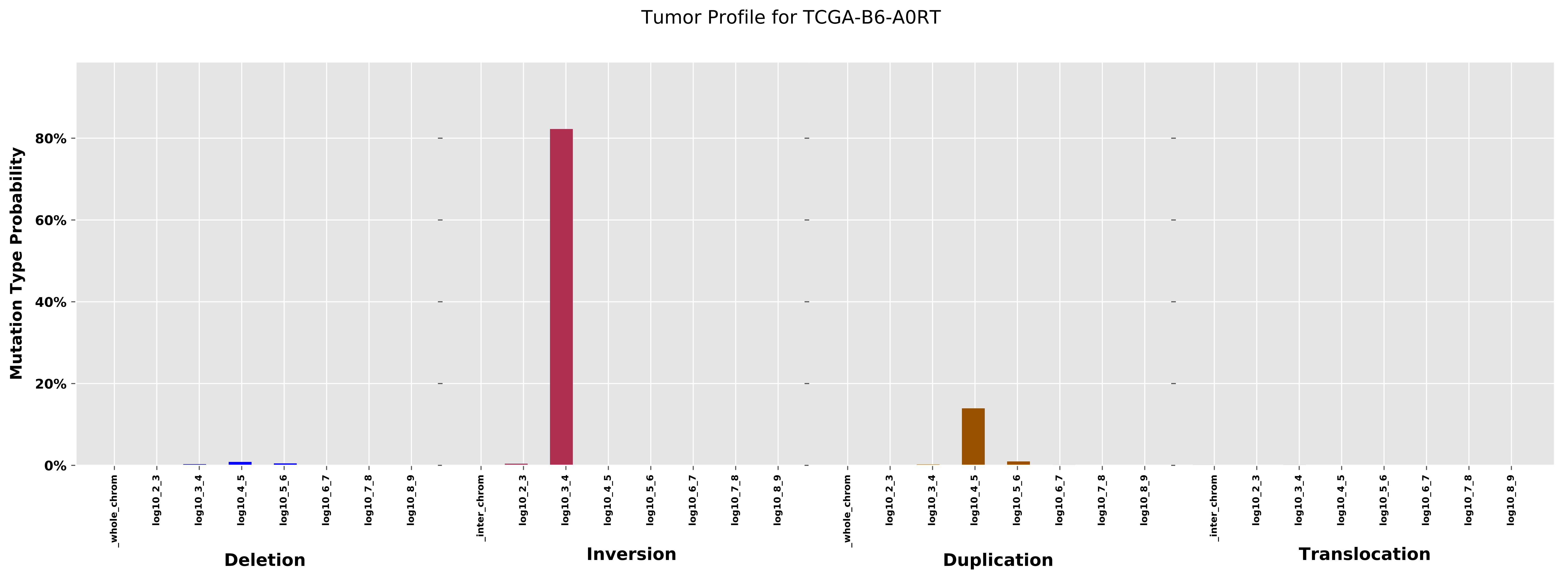
Tumor Profile for TCGA-B6-A0RT
Mutation Type Probability
80%
60%
40%
20%
0%
_whole_chrom
log10_2_3
log10_3_4
log10_4_5
log10_5_6
log10_6_7
log10_7_8
log10_8_9
_inter_chrom
Deletion
Inversion
Duplication
Translocation

Cancer processes Weights for TCGA-GM-A3XL

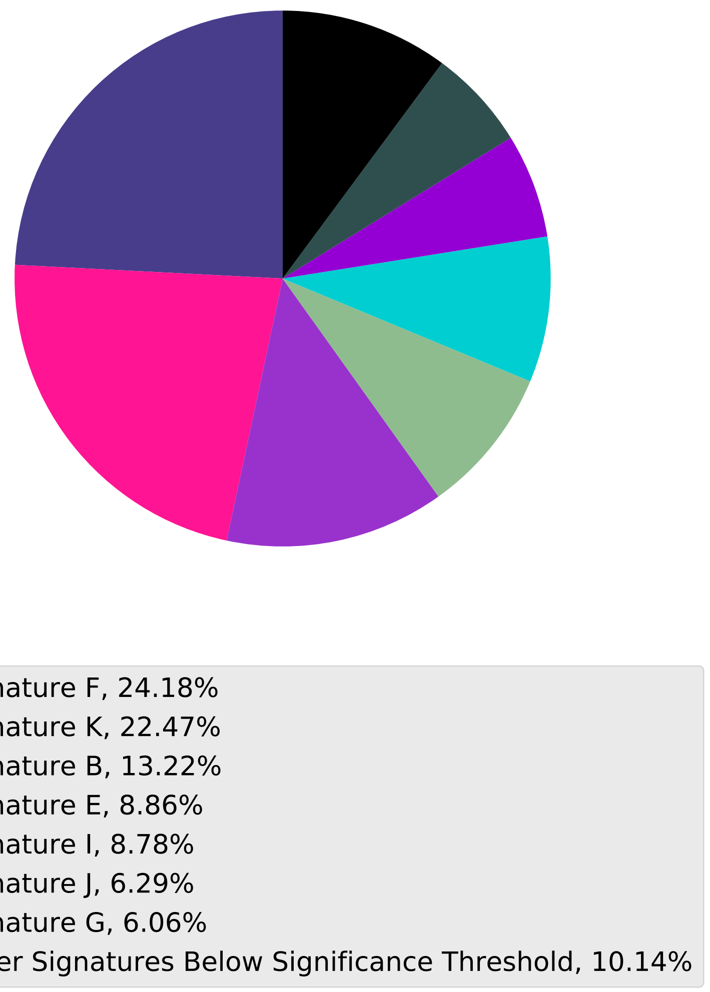

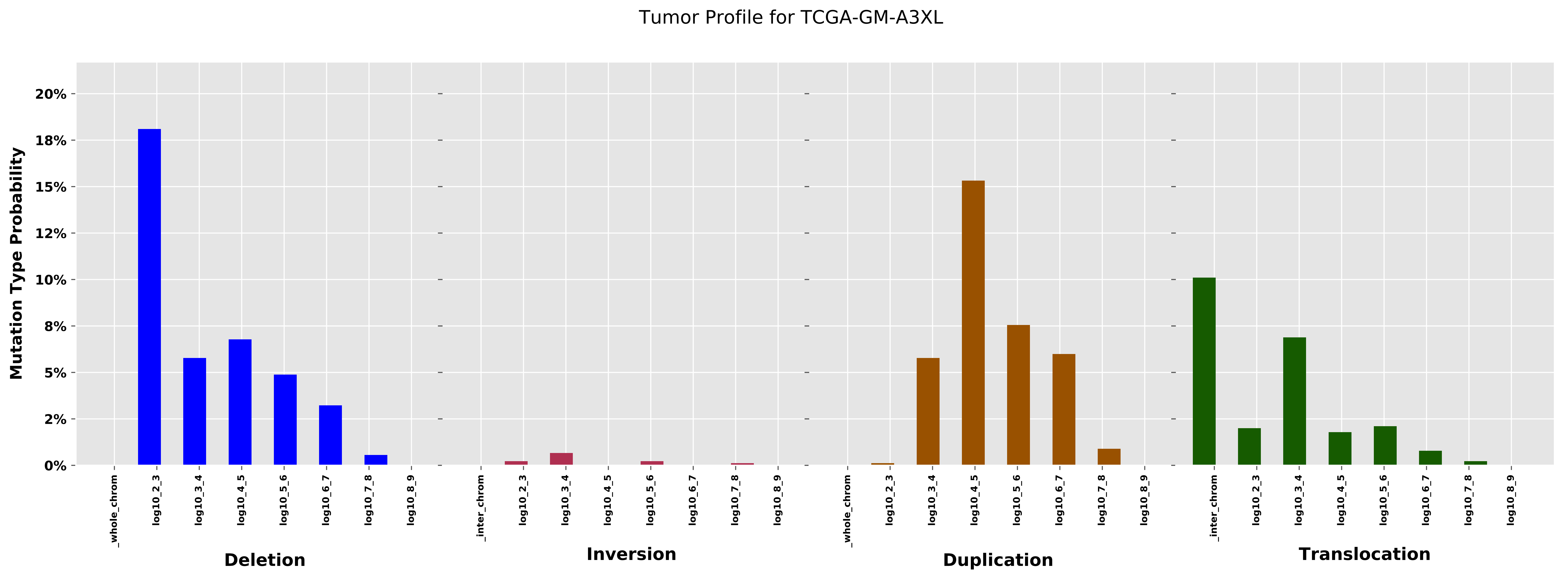

Tumor Profile for TCGA-GM-A3XL
Mutation Type Probability
0%
2%
5%
8%
10%
12%
15%
18%
20%
_whole_chrom
log10_2_3
log10_3_4
log10_4_5
log10_5_6
log10_6_7
log10_7_8
log10_8_9
_inter_chrom
Deletion
Inversion
Duplication
Translocation

Cancer processes Weights for TCGA-AO-A0J2

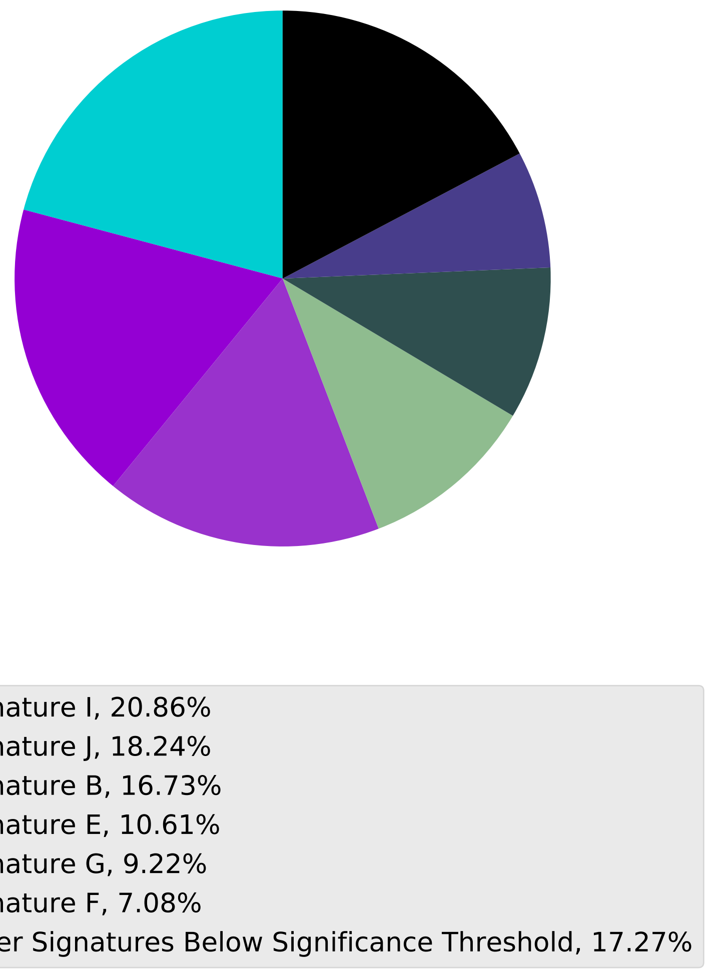

Tumor Profile for TCGA-AO-A0J2

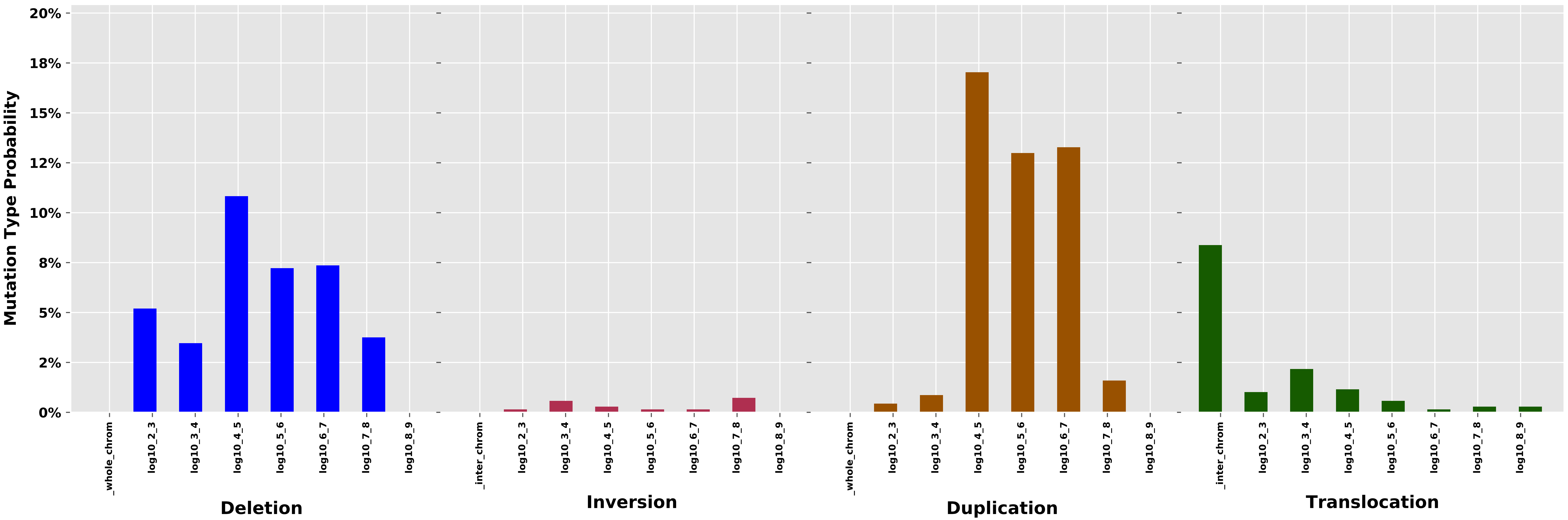

#### Cancer processes Weights for TCGA-E2-A156

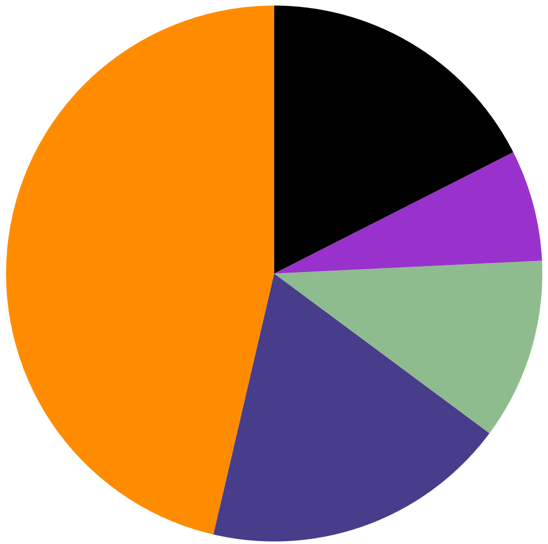

- Signature A, 46.36%
- Signature F, 18.47%
- Signature E, 10.93%
- Signature B, 6.72%
- Other Signatures Below Significance Threshold, 17.52%

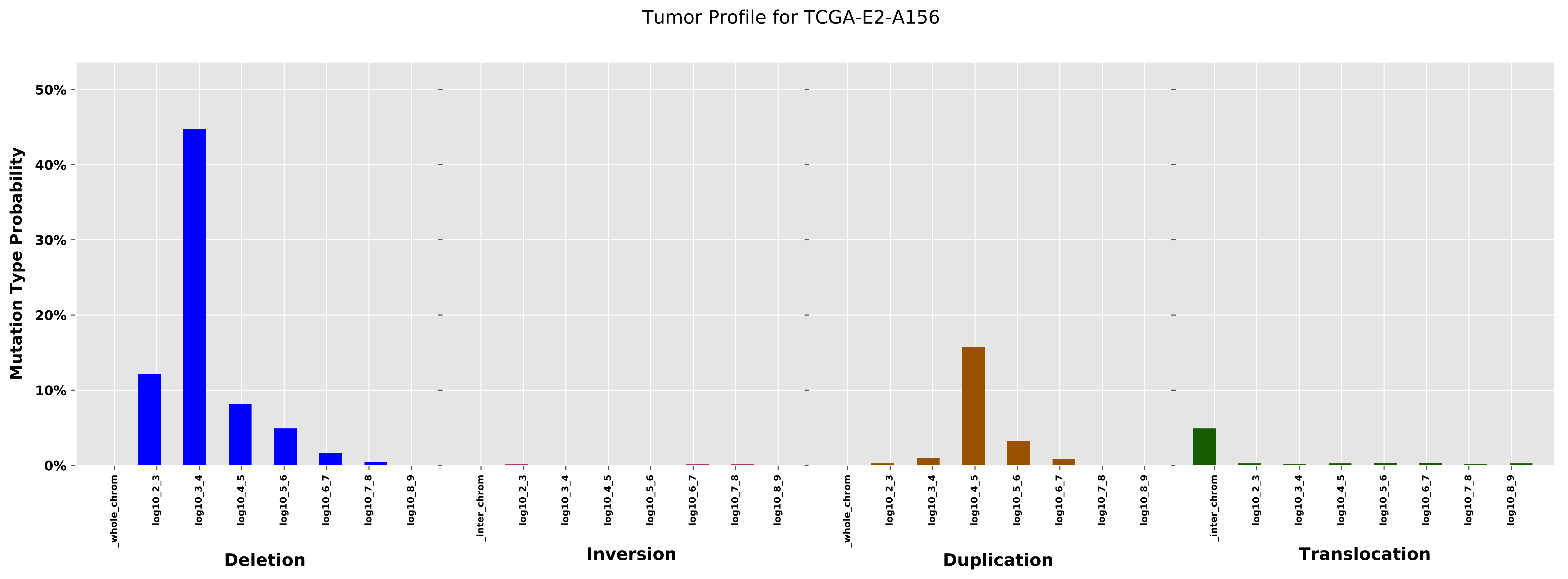

Tumor Profile for TCGA-E2-A156
Mutation Type Probability
50%
40%
30%
20%
10%
0%
_whole_chrom
log10_2_3
log10_3_4
log10_4_5
log10_5_6
log10_6_7
log10_7_8
log10_8_9
_inter_chrom
Deletion
Inversion
Duplication
Translocation

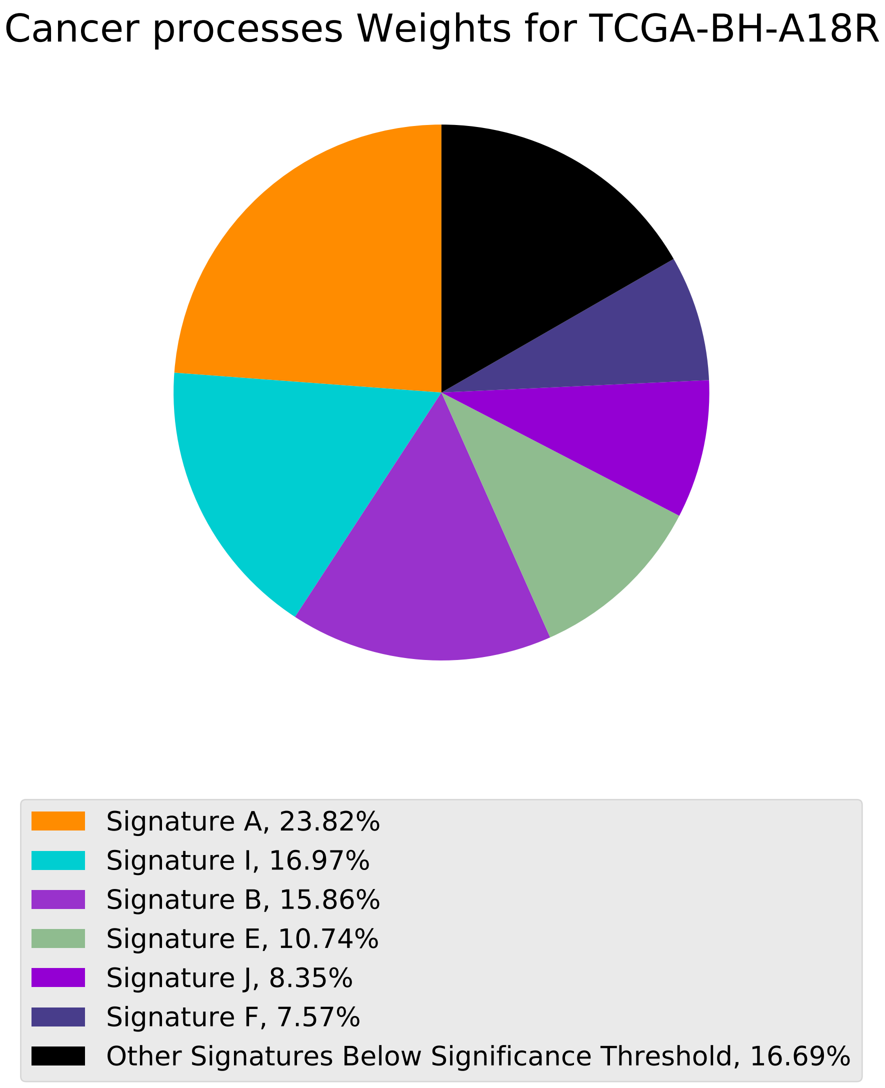
Cancer processes Weights for TCGA-BH-A18R
Signature A, 23.82%
Signature I, 16.97%
Signature B, 15.86%
Signature E, 10.74%
Signature J, 8.35%
Signature F, 7.57%
Other Signatures Below Significance Threshold, 16.69%

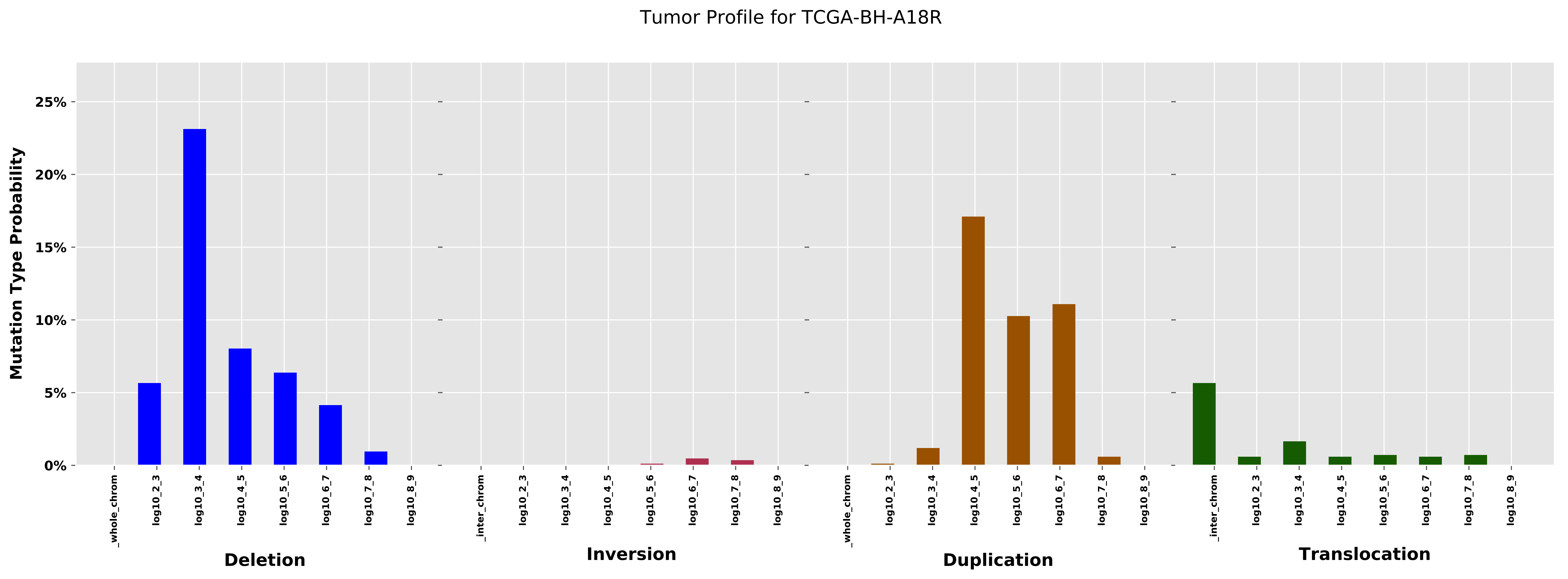
Tumor Profile for TCGA-BH-A18R
Mutation Type Probability
25%
20%
15%
10%
5%
0%
_whole_chrom
log10_2_3
log10_3_4
log10_4_5
log10_5_6
log10_6_7
log10_7_8
log10_8_9
_inter_chrom
Deletion
Inversion
Duplication
Translocation

Cancer processes Weights for TCGA-AO-A0JM

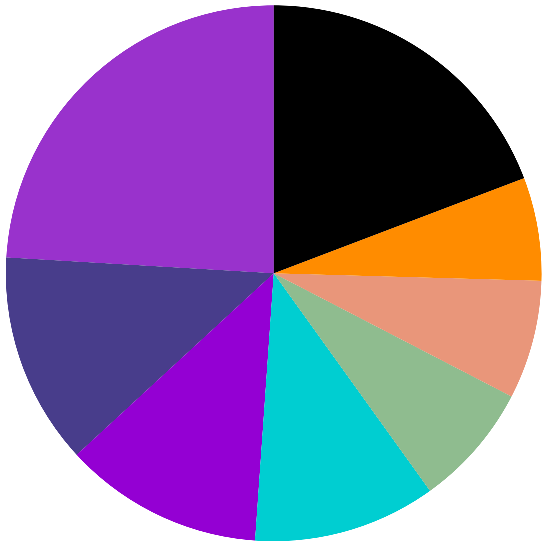

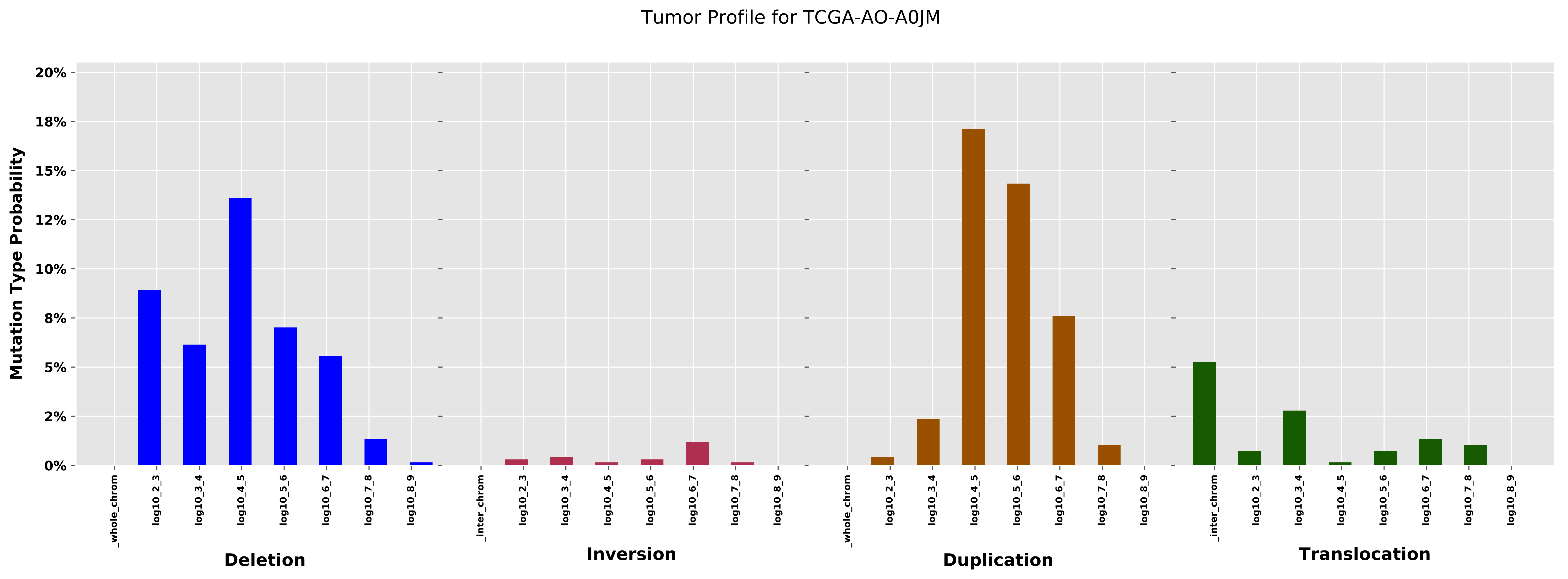

Tumor Profile for TCGA-AO-A0JM
Mutation Type Probability
20%
18%
15%
12%
10%
8%
5%
2%
0%
_whole_chrom
log10_2_3
log10_3_4
log10_4_5
log10_5_6
log10_6_7
log10_7_8
log10_8_9
_inter_chrom
Deletion
Inversion
Duplication
Translocation

Cancer processes Weights for TCGA-AA-A01S

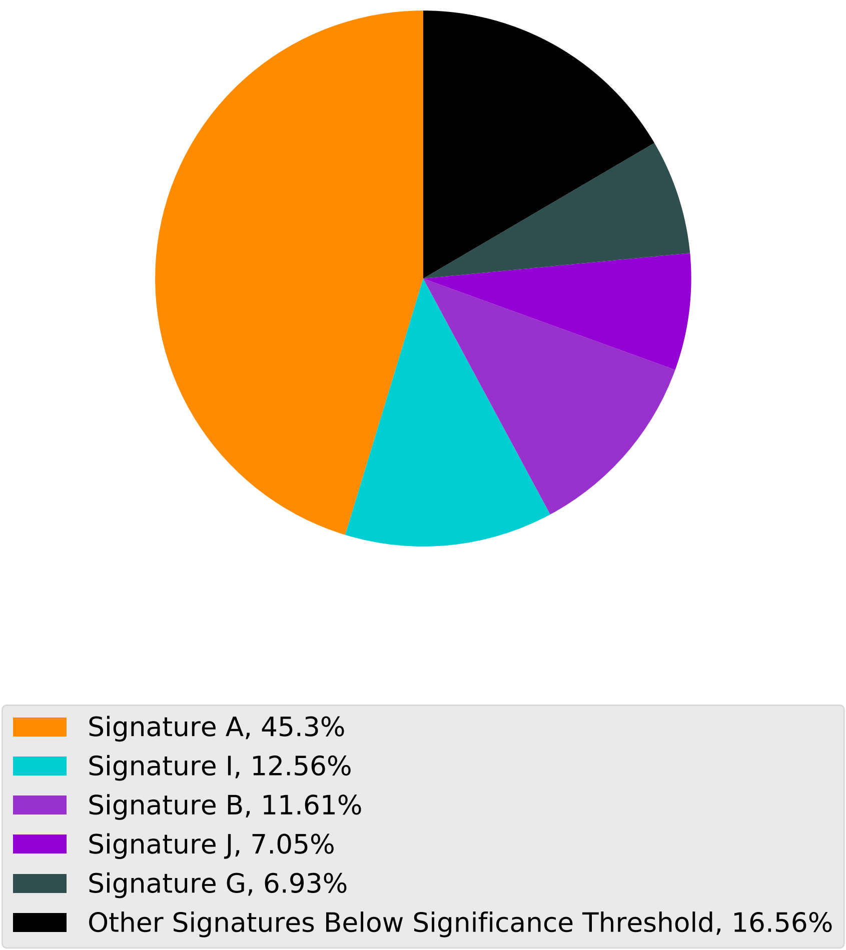

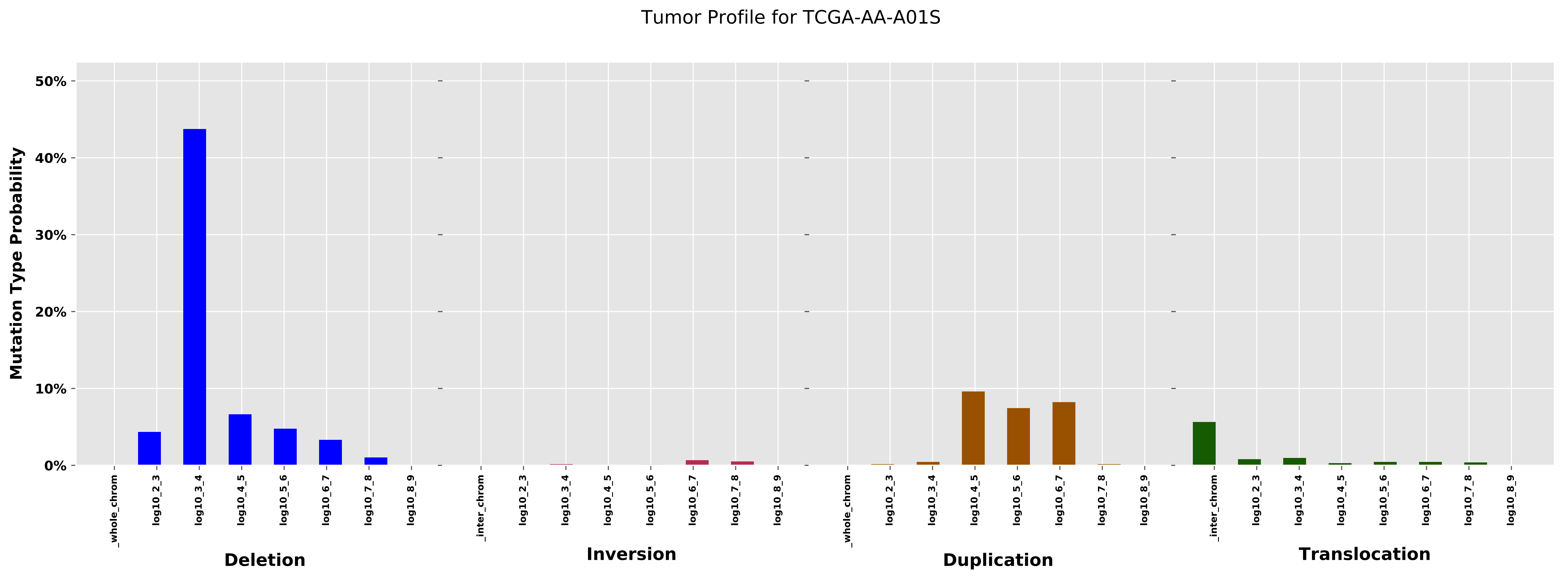

Tumor Profile for TCGA-AA-A01S
Mutation Type Probability
50%
40%
30%
20%
10%
0%
_whole_chrom
log10_2_3
log10_3_4
log10_4_5
log10_5_6
log10_6_7
log10_7_8
log10_8_9
_inter_chrom
Deletion
Inversion
Duplication
Translocation

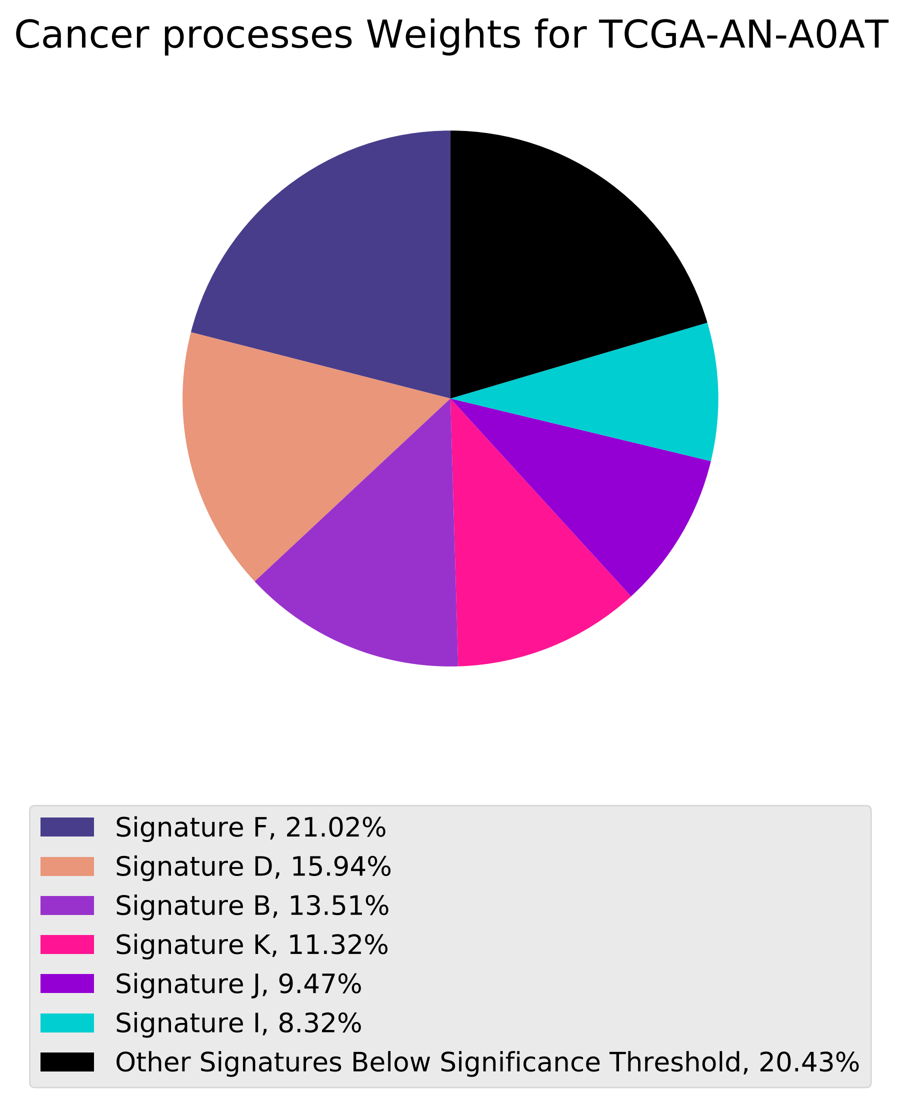

Cancer processes Weights for TCGA-AN-A0AT
Signature F, 21.02%
Signature D, 15.94%
Signature B, 13.51%
Signature K, 11.32%
Signature J, 9.47%
Signature I, 8.32%
Other Signatures Below Significance Threshold, 20.43%

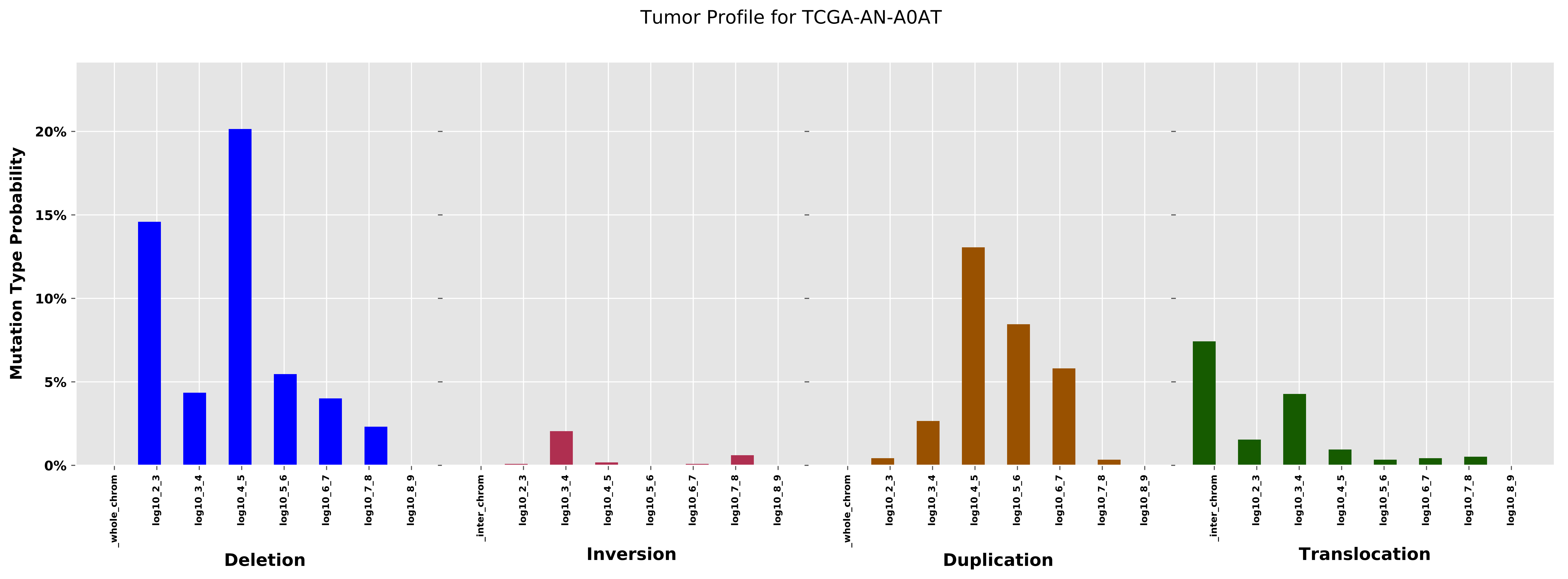

Tumor Profile for TCGA-AN-A0AT
Mutation Type Probability
0%
5%
10%
15%
20%
_whole_chrom
log10_2_3
log10_3_4
log10_4_5
log10_5_6
log10_6_7
log10_7_8
log10_8_9
Deletion
_inter_chrom
Inversion
Duplication
Translocation

Cancer processes Weights for TCGA-B6-A0WX

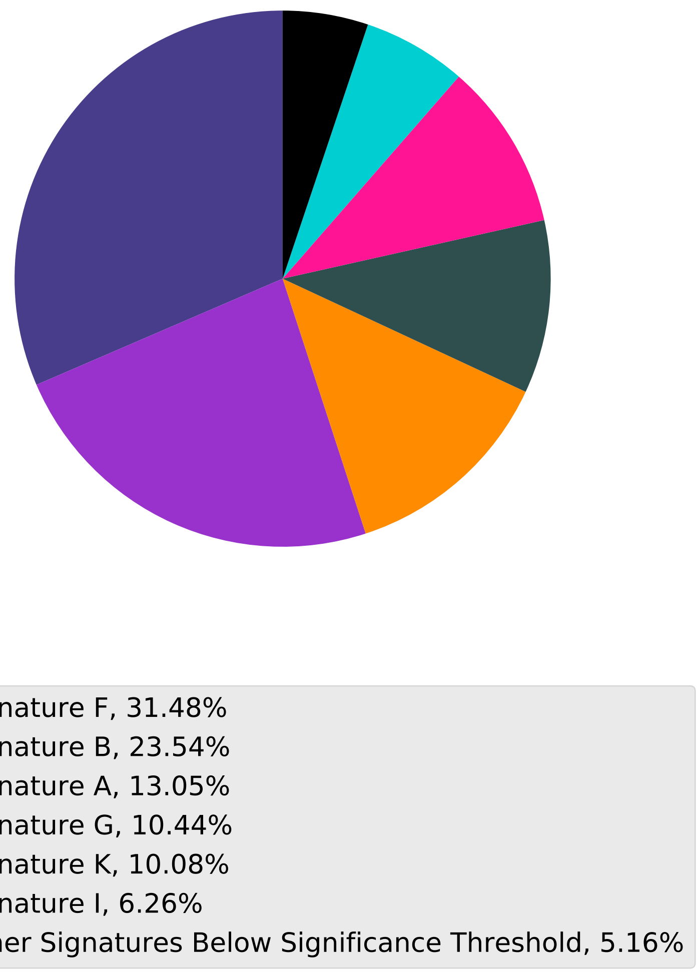

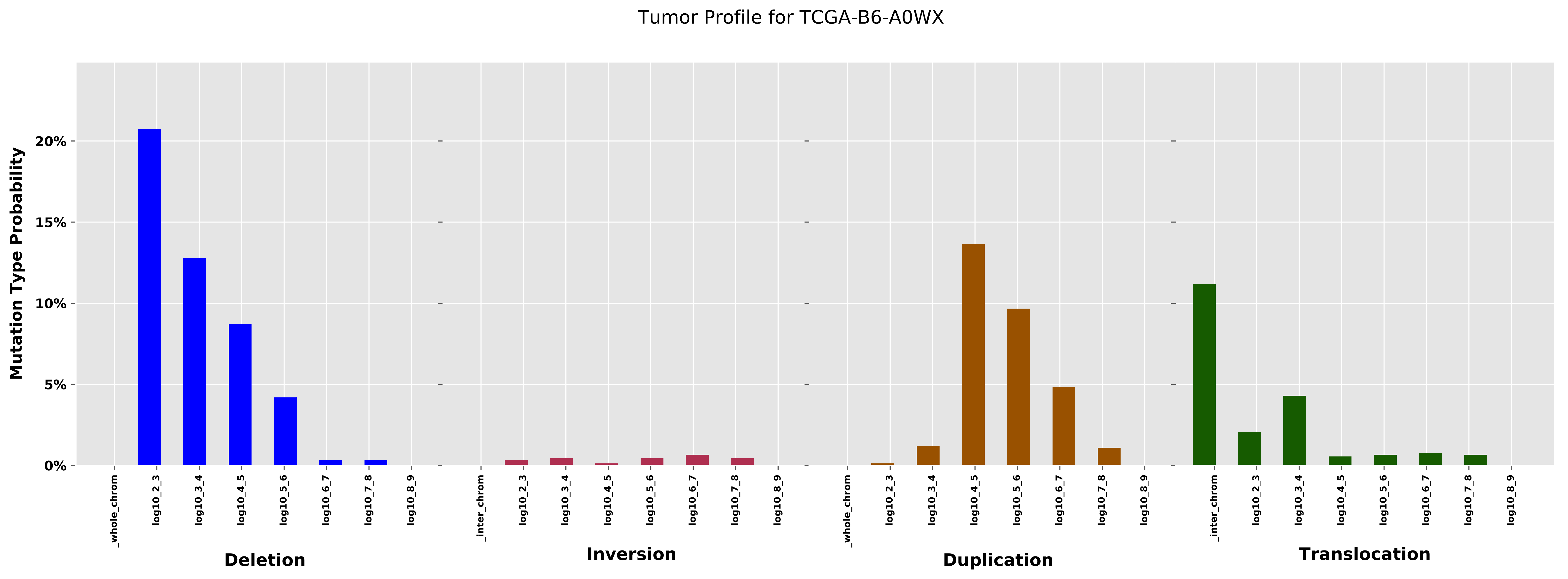
Tumor Profile for TCGA-B6-A0WX
Mutation Type Probability
0%
5%
10%
15%
20%
_whole_chrom
log10_2_3
log10_3_4
log10_4_5
log10_5_6
log10_6_7
log10_7_8
log10_8_9
Deletion
_inter_chrom
log10_2_3
log10_3_4
log10_4_5
log10_5_6
log10_6_7
log10_7_8
log10_8_9
Inversion
_whole_chrom
log10_2_3
log10_3_4
log10_4_5
log10_5_6
log10_6_7
log10_7_8
log10_8_9
Duplication
_inter_chrom
log10_2_3
log10_3_4
log10_4_5
log10_5_6
log10_6_7
log10_7_8
log10_8_9
Translocation

Cancer processes Weights for TCGA-BH-A0WA

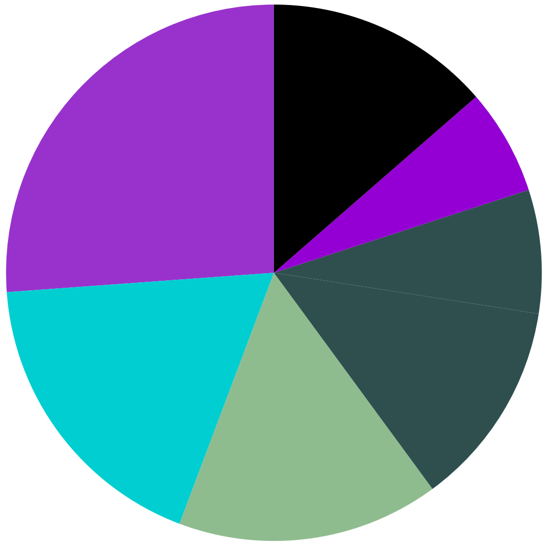

- Signature B, 26.15%
- Signature I, 18.12%
- Signature E, 15.81%
- Signature G, 12.47%
- Signature H, 7.44%
- Signature J, 6.39%
- Other Signatures Below Significance Threshold, 13.61%

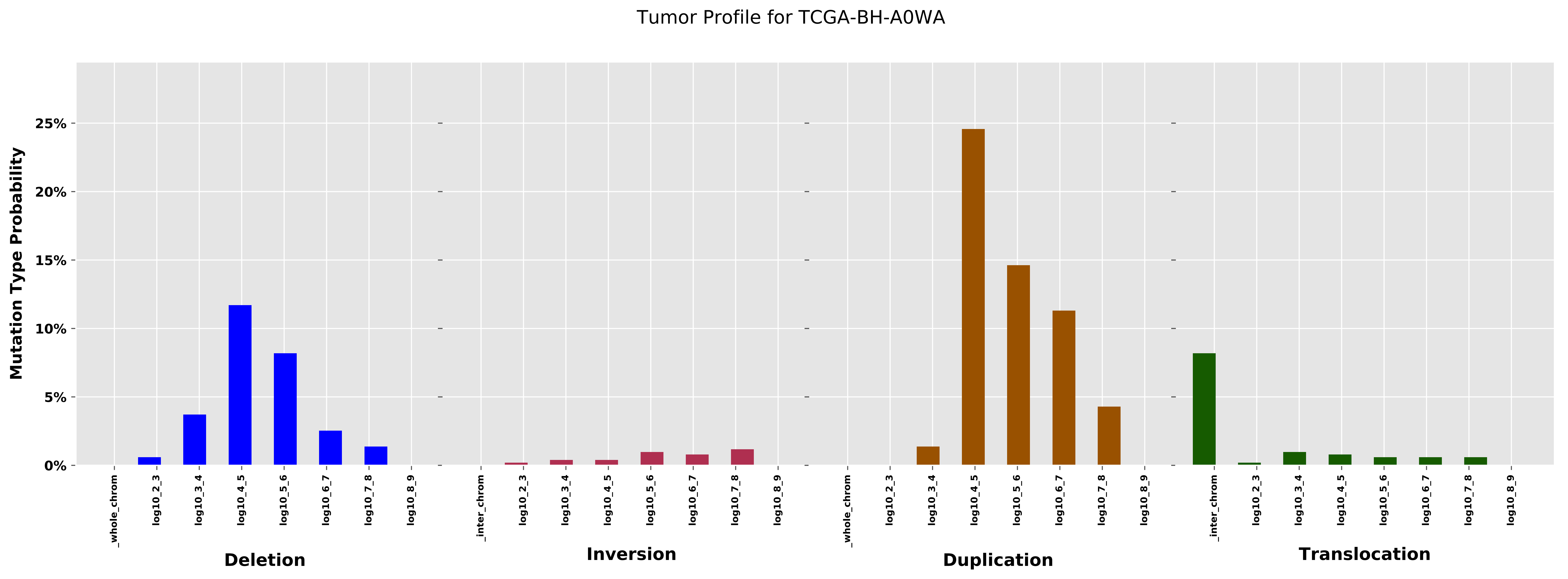

Tumor Profile for TCGA-BH-A0WA
Mutation Type Probability
25%
20%
15%
10%
5%
0%
_whole_chrom
log10_2_3
log10_3_4
log10_4_5
log10_5_6
log10_6_7
log10_7_8
log10_8_9
_inter_chrom
Deletion
Inversion
Duplication
Translocation

Cancer processes Weights for TCGA-A7-A13D

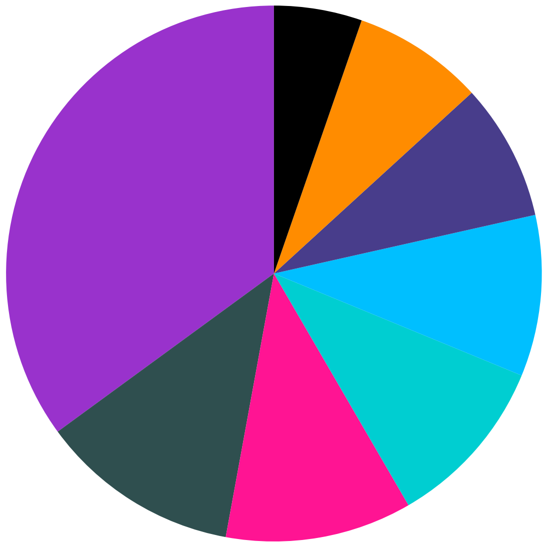

| |
|---|
| Signature B, 35.05% |
| Signature G, 12.07% |
| Signature K, 11.26% |
| Signature I, 10.41% |
| Signature L, 9.73% |
| Signature F, 8.27% |
| Signature A, 7.91% |
| Other Signatures Below Significance Threshold, 5.31% |

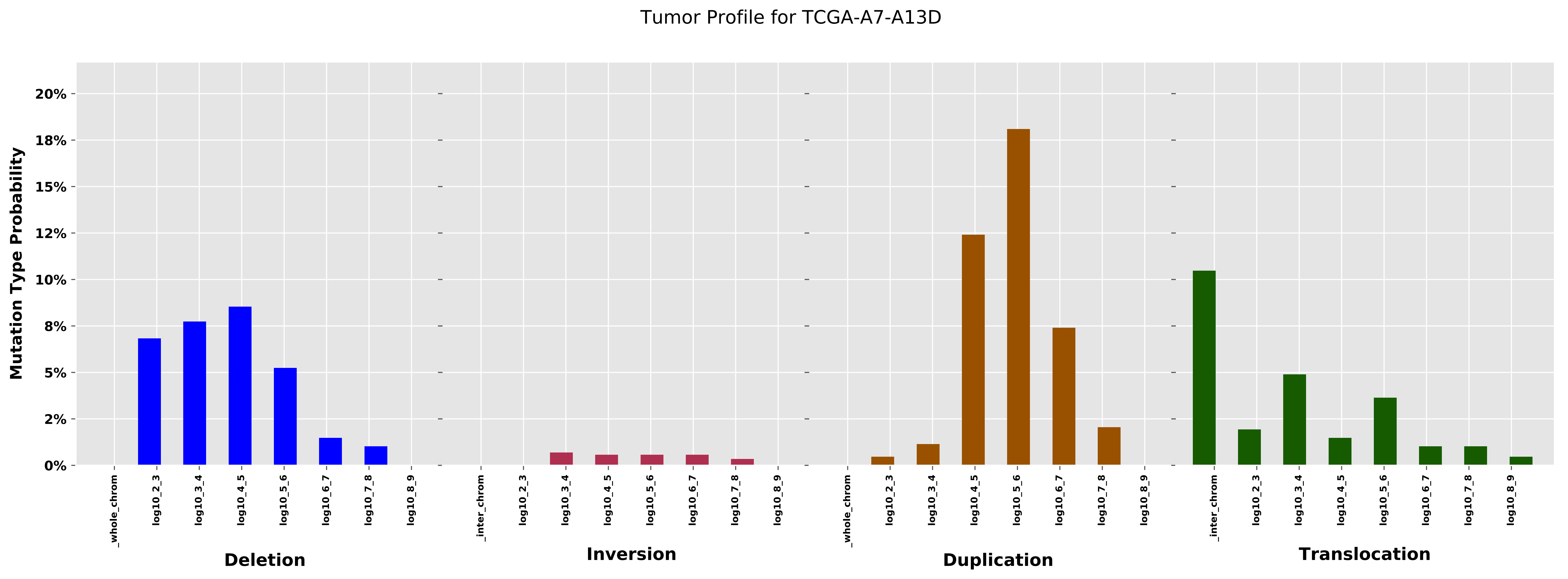

Tumor Profile for TCGA-A7-A13D
Mutation Type Probability
20%
18%
15%
12%
10%
8%
5%
2%
0%
_whole_chrom
log10_2_3
log10_3_4
log10_4_5
log10_5_6
log10_6_7
log10_7_8
log10_8_9
Deletion
_inter_chrom
log10_2_3
log10_3_4
log10_4_5
log10_5_6
log10_6_7
log10_7_8
log10_8_9
Inversion
_whole_chrom
log10_2_3
log10_3_4
log10_4_5
log10_5_6
log10_6_7
log10_7_8
log10_8_9
Duplication
_inter_chrom
log10_2_3
log10_3_4
log10_4_5
log10_5_6
log10_6_7
log10_7_8
log10_8_9
Translocation

Cancer processes Weights for TCGA-B6-A0IJ

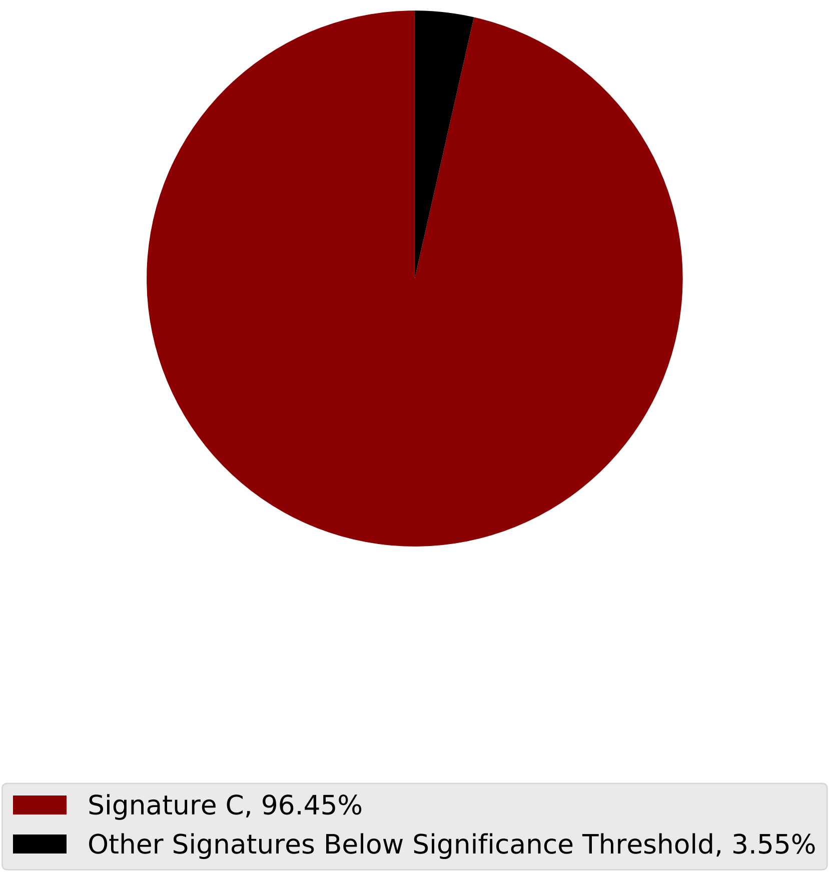

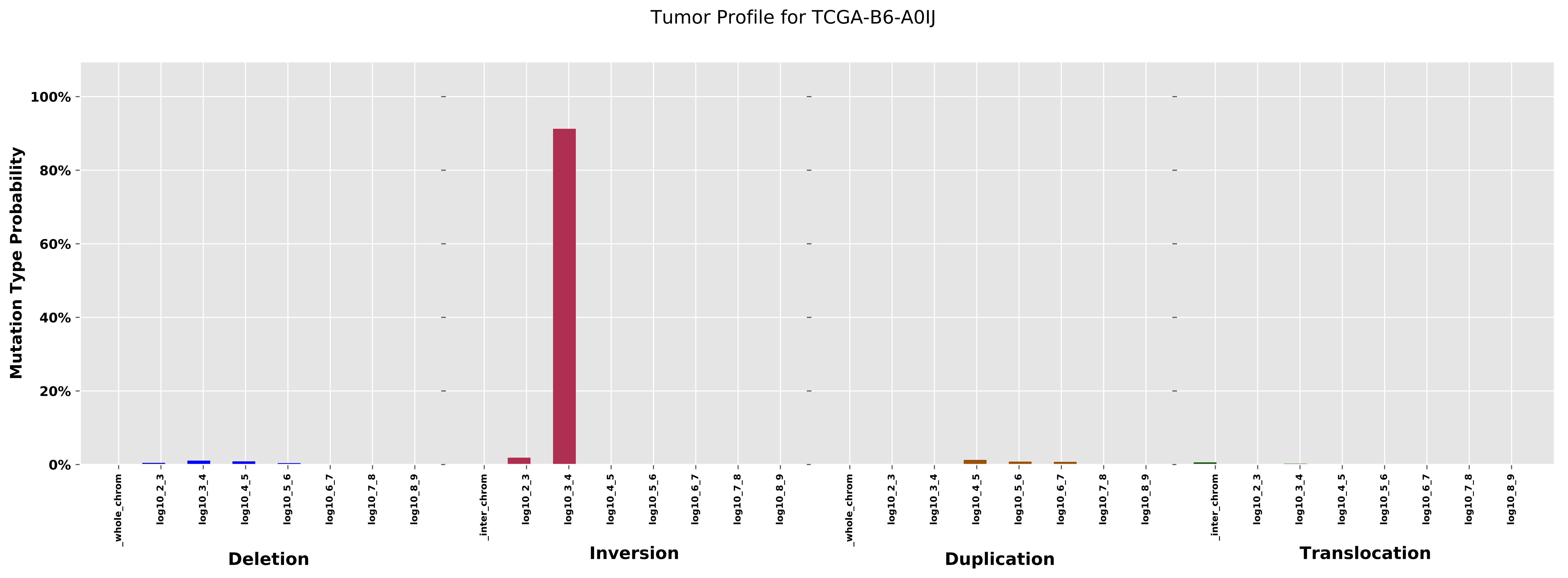
Tumor Profile for TCGA-B6-A0IJ
Mutation Type Probability
100%
80%
60%
40%
20%
0%
_whole_chrom
log10_2_3
log10_3_4
log10_4_5
log10_5_6
log10_6_7
log10_7_8
log10_8_9
_inter_chrom
Deletion
Inversion
Duplication
Translocation

#### Cancer processes Weights for TCGA-A2-A04Q

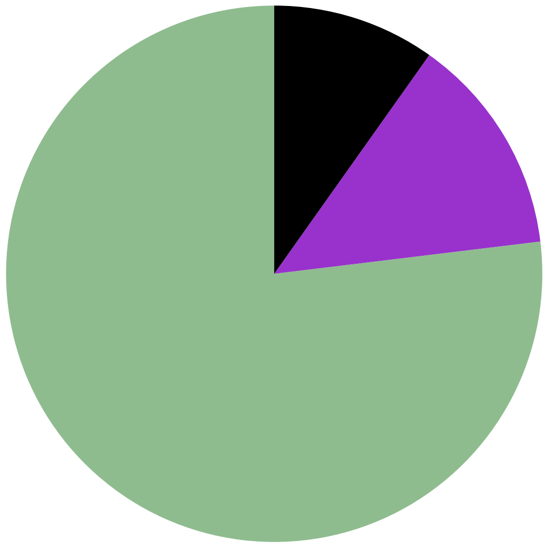

- Signature E, 76.92%
- Signature B, 13.26%
- Other Signatures Below Significance Threshold, 9.82%

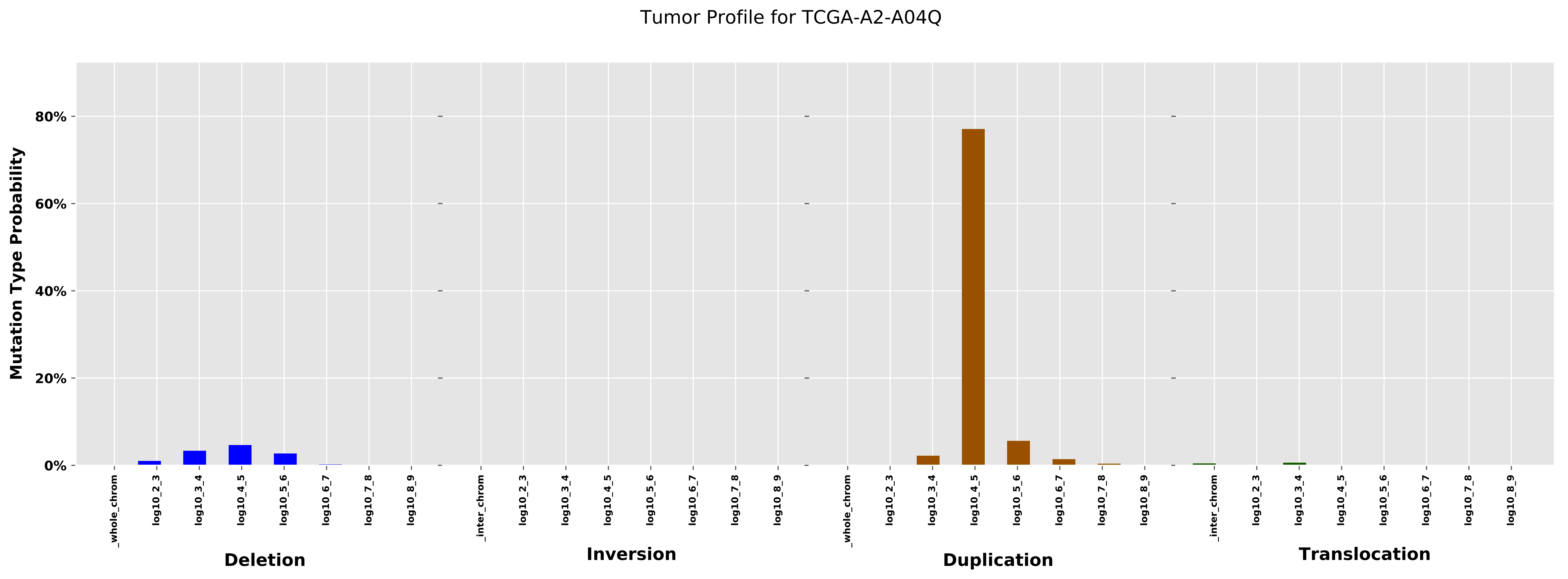

Tumor Profile for TCGA-A2-A04Q
Mutation Type Probability
80%
60%
40%
20%
0%
_whole_chrom
log10_2_3
log10_3_4
log10_4_5
log10_5_6
log10_6_7
log10_7_8
log10_8_9
_inter_chrom
Deletion
Inversion
Duplication
Translocation

Cancer processes Weights for TCGA-BH-A0B3

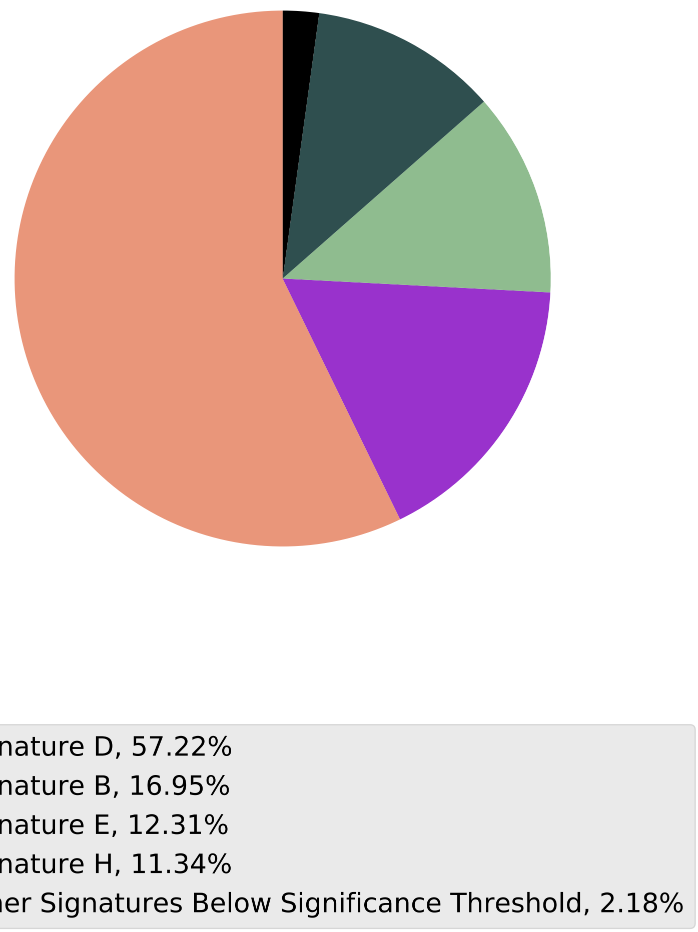

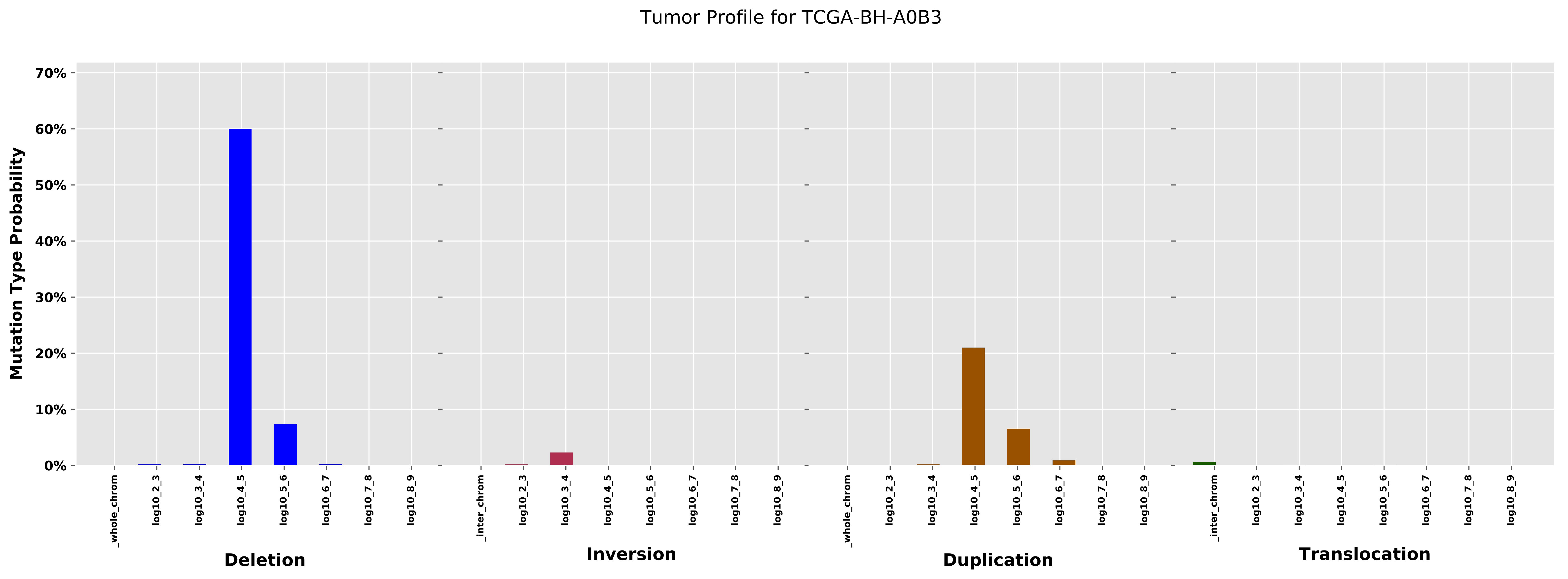

Tumor Profile for TCGA-BH-A0B3
Mutation Type Probability
70%
60%
50%
40%
30%
20%
10%
0%
_whole_chrom
log10_2_3
log10_3_4
log10_4_5
log10_5_6
log10_6_7
log10_7_8
log10_8_9
Deletion
_inter_chrom
Inversion
Duplication
Translocation

Cancer processes Weights for TCGA-BH-A18U

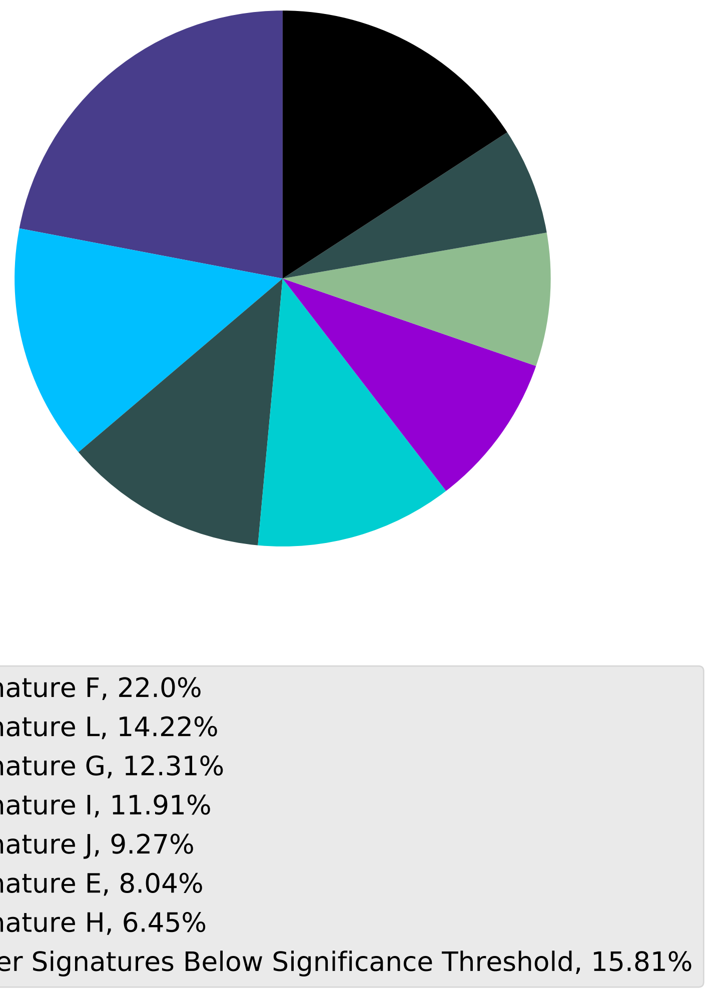

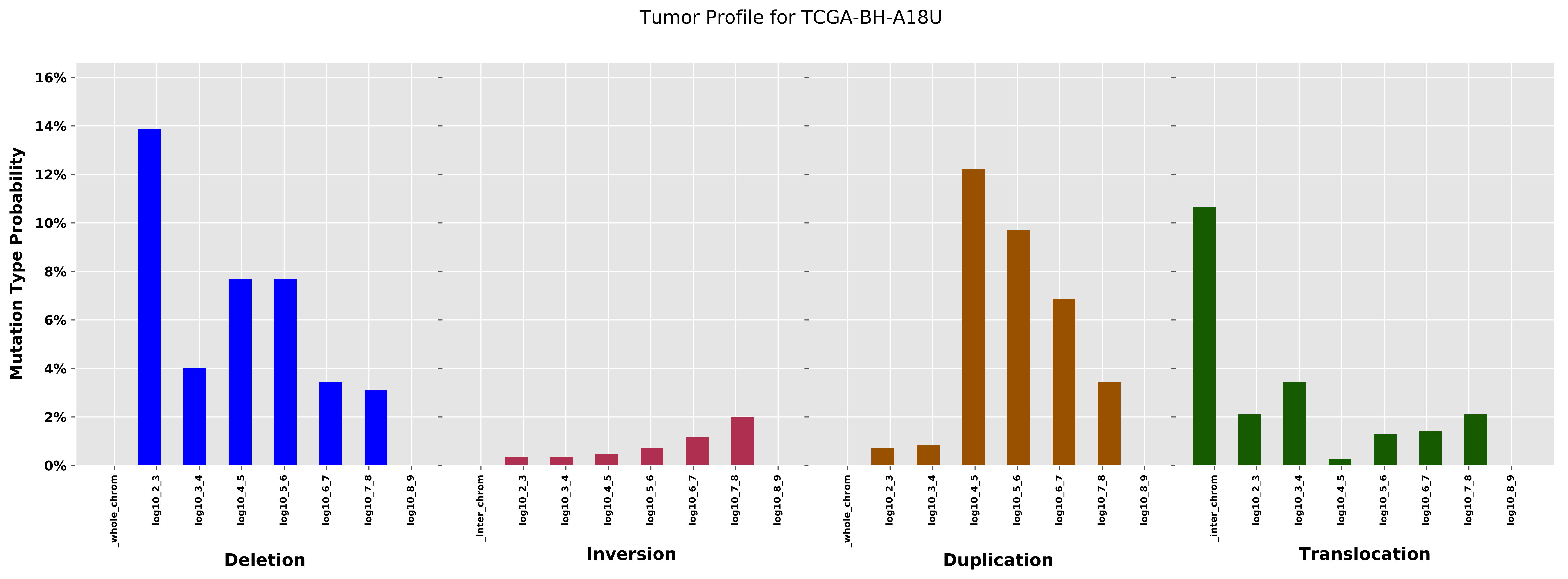
Tumor Profile for TCGA-BH-A18U
Mutation Type Probability
0%
2%
4%
6%
8%
10%
12%
14%
16%
_whole_chrom
log10_2_3
log10_3_4
log10_4_5
log10_5_6
log10_6_7
log10_7_8
log10_8_9
_inter_chrom
Deletion
Inversion
Duplication
Translocation

Cancer processes Weights for TCGA-GI-A2C9

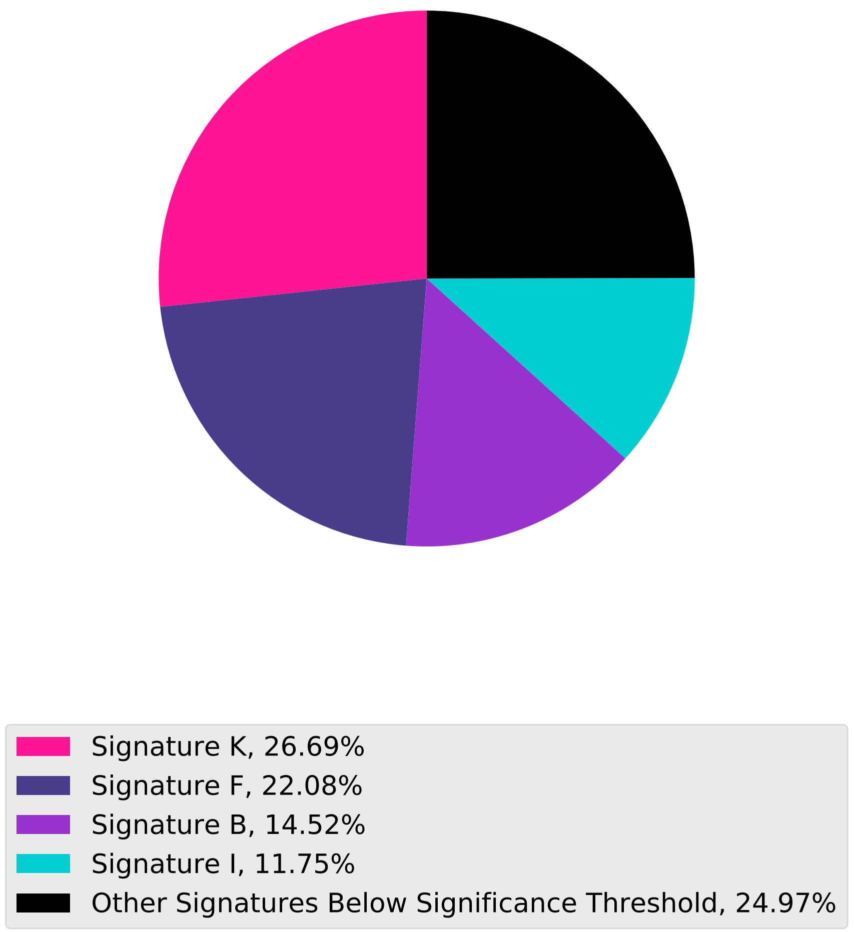

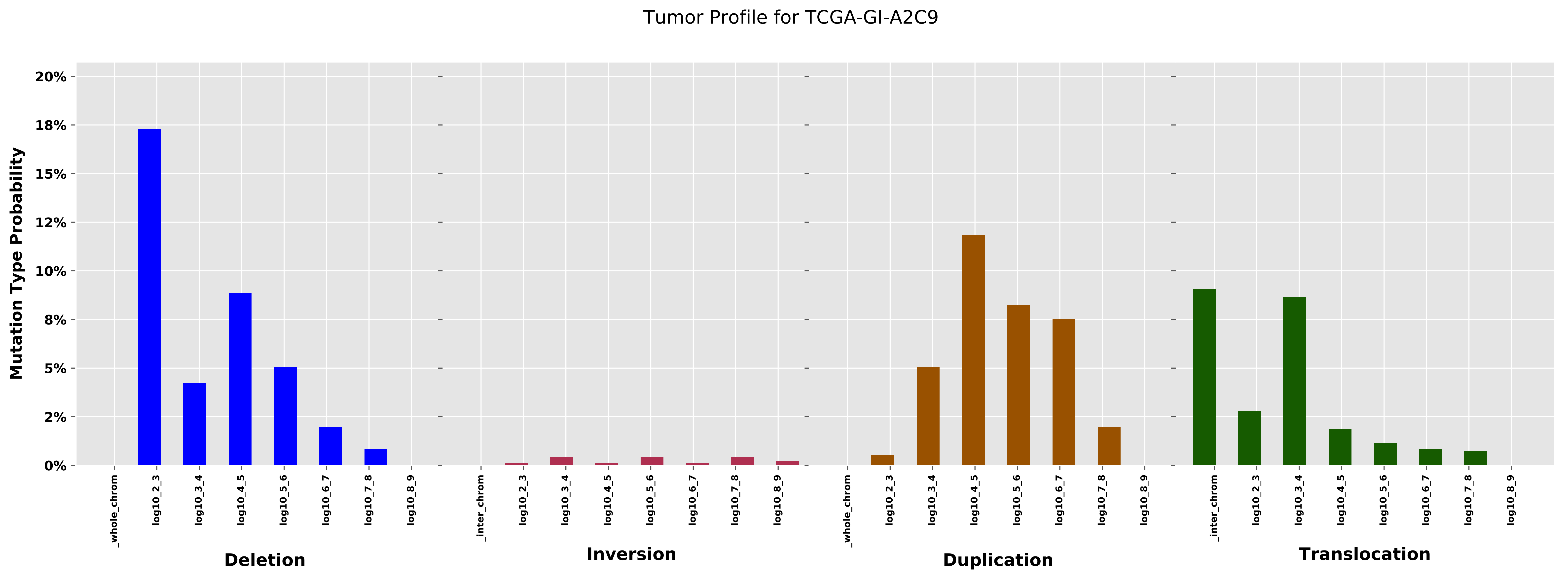

Tumor Profile for TCGA-GI-A2C9
Mutation Type Probability
20%
18%
15%
12%
10%
8%
5%
2%
0%
_whole_chrom
log10_2_3
log10_3_4
log10_4_5
log10_5_6
log10_6_7
log10_7_8
log10_8_9
_inter_chrom
Deletion
Inversion
Duplication
Translocation

Cancer processes Weights for TCGA-BH-A0DT

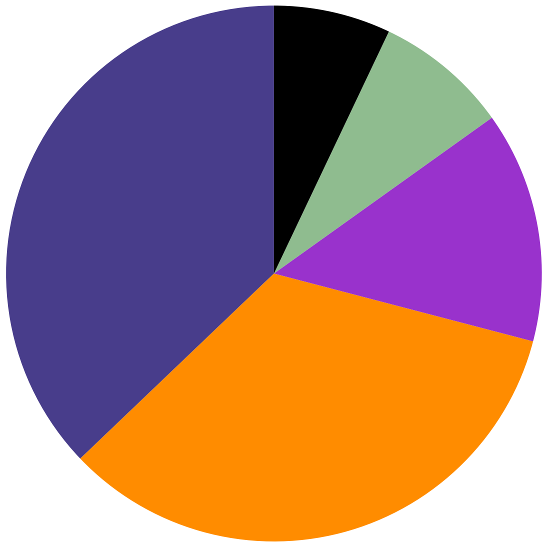

- Signature F, 37.14%
- Signature A, 33.79%
- Signature B, 13.95%
- Signature E, 8.06%
- Other Signatures Below Significance Threshold, 7.06%

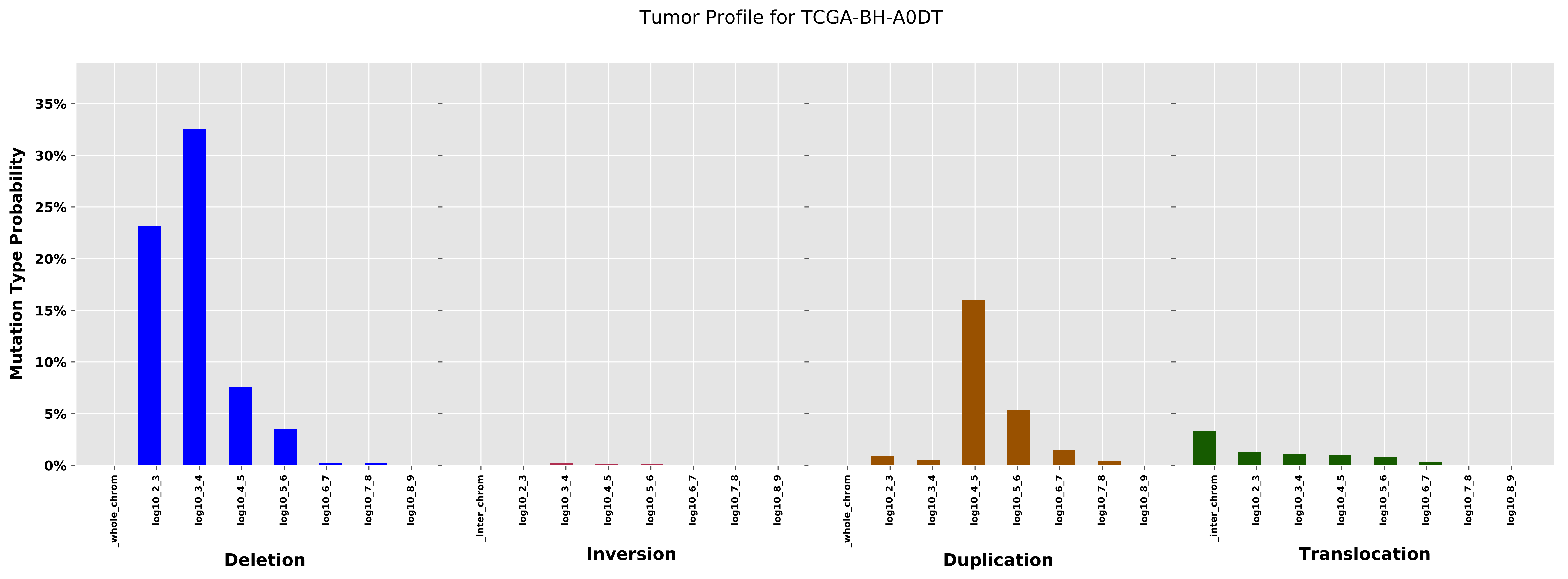
Tumor Profile for TCGA-BH-A0DT
Mutation Type Probability
35%
30%
25%
20%
15%
10%
5%
0%
_whole_chrom
log10_2_3
log10_3_4
log10_4_5
log10_5_6
log10_6_7
log10_7_8
log10_8_9
Deletion
_inter_chrom
log10_2_3
log10_3_4
log10_4_5
log10_5_6
log10_6_7
log10_7_8
log10_8_9
Inversion
_whole_chrom
log10_2_3
log10_3_4
log10_4_5
log10_5_6
log10_6_7
log10_7_8
log10_8_9
Duplication
_inter_chrom
log10_2_3
log10_3_4
log10_4_5
log10_5_6
log10_6_7
log10_7_8
log10_8_9
Translocation

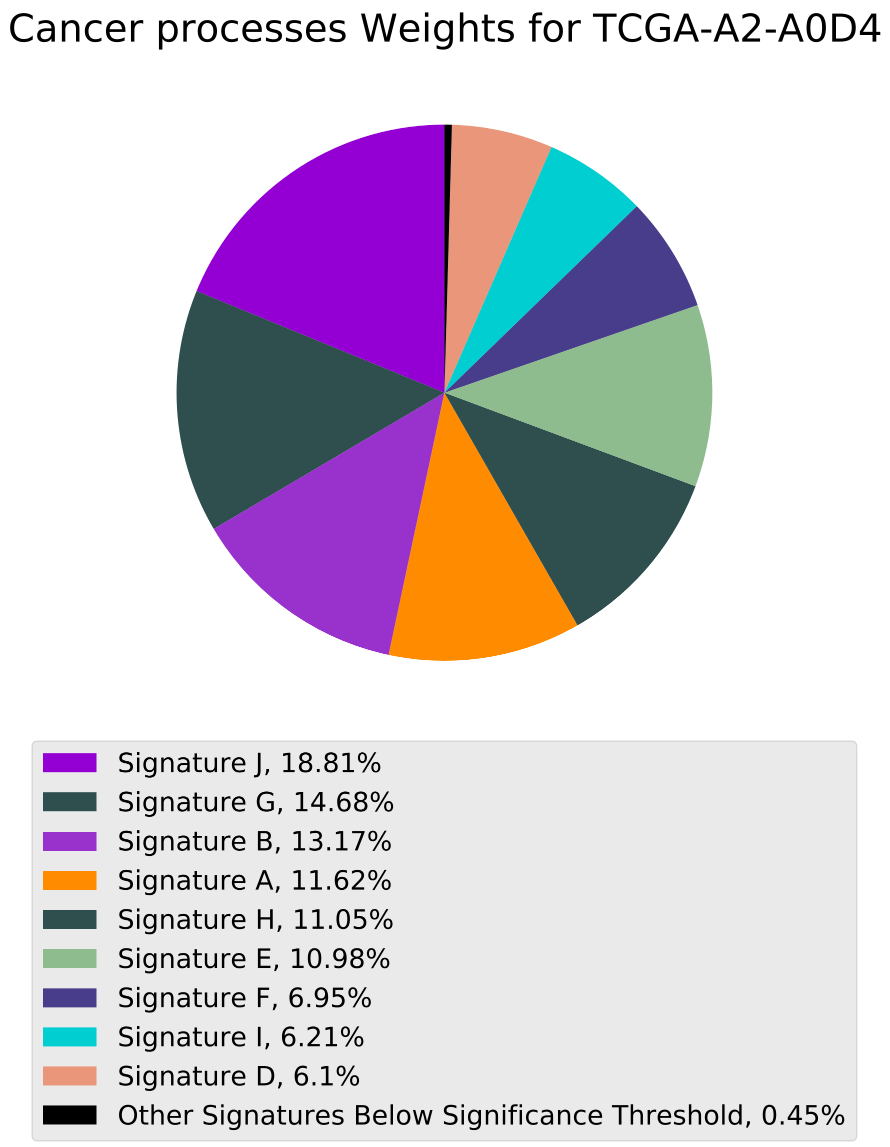

Cancer processes Weights for TCGA-A2-A0D4
Signature J, 18.81%
Signature G, 14.68%
Signature B, 13.17%
Signature A, 11.62%
Signature H, 11.05%
Signature E, 10.98%
Signature F, 6.95%
Signature I, 6.21%
Signature D, 6.1%
Other Signatures Below Significance Threshold, 0.45%

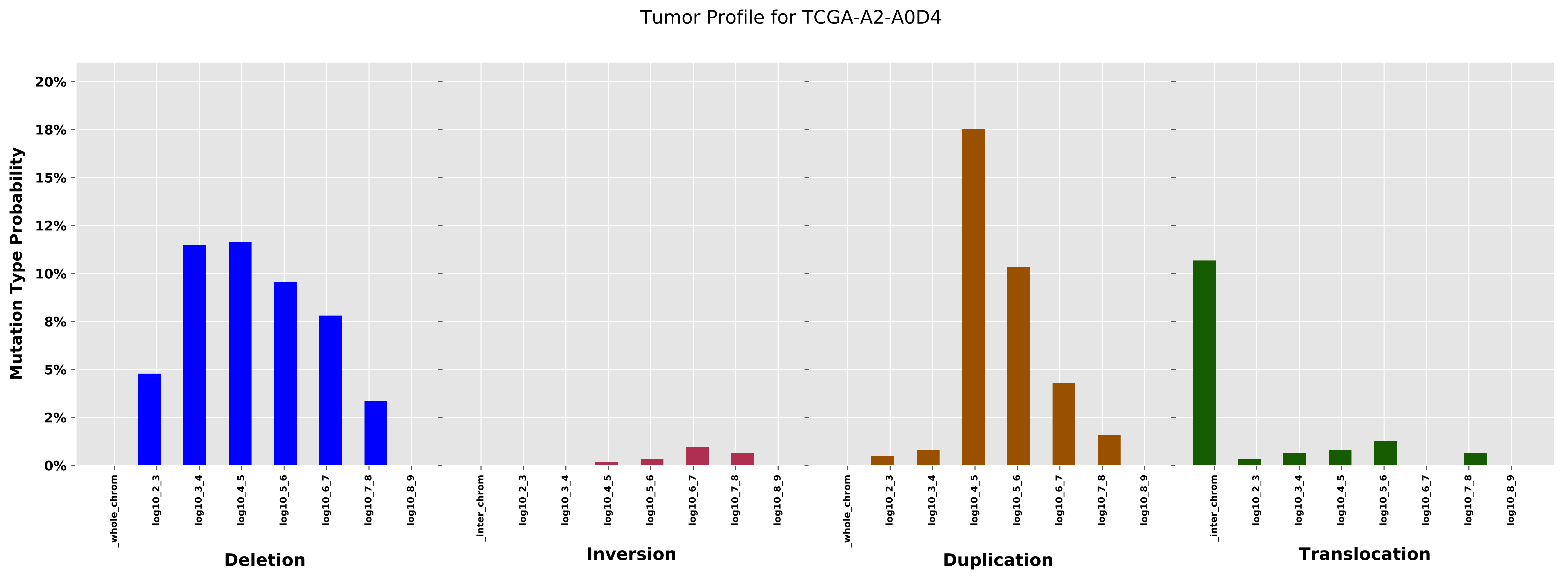
Tumor Profile for TCGA-A2-A0D4
Mutation Type Probability
20%
18%
15%
12%
10%
8%
5%
2%
0%
_whole_chrom
log10_2_3
log10_3_4
log10_4_5
log10_5_6
log10_6_7
log10_7_8
log10_8_9
_inter_chrom
Deletion
Inversion
Duplication
Translocation

Cancer processes Weights for TCGA-EW-A1P8

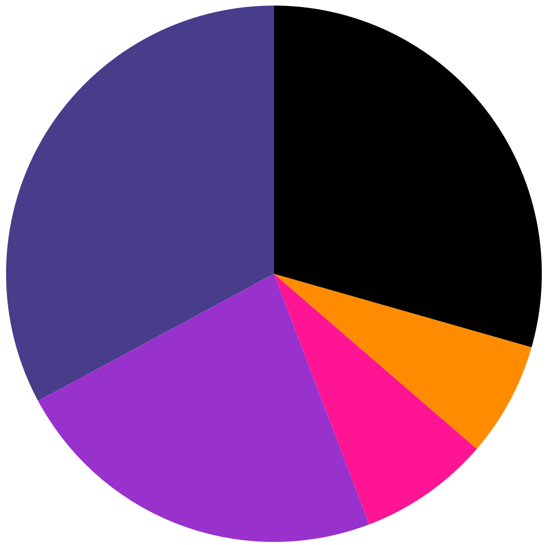

Signature F, 32.86%
Signature B, 22.93%
Signature K, 7.89%
Signature A, 6.9%
Other Signatures Below Significance Threshold, 29.42%

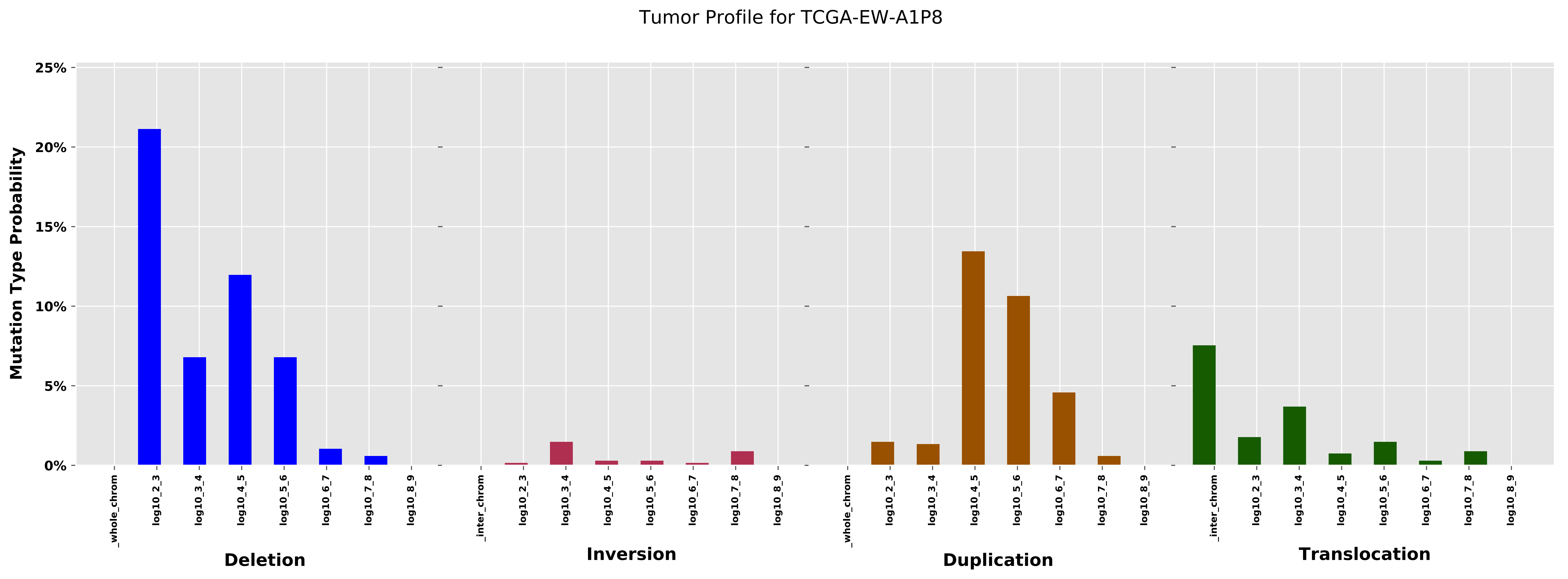
Tumor Profile for TCGA-EW-A1P8
Mutation Type Probability
25%
20%
15%
10%
5%
0%
_whole_chrom
log10_2_3
log10_3_4
log10_4_5
log10_5_6
log10_6_7
log10_7_8
log10_8_9
_inter_chrom
Deletion
Inversion
Duplication
Translocation

Cancer processes Weights for TCGA-A6-2683

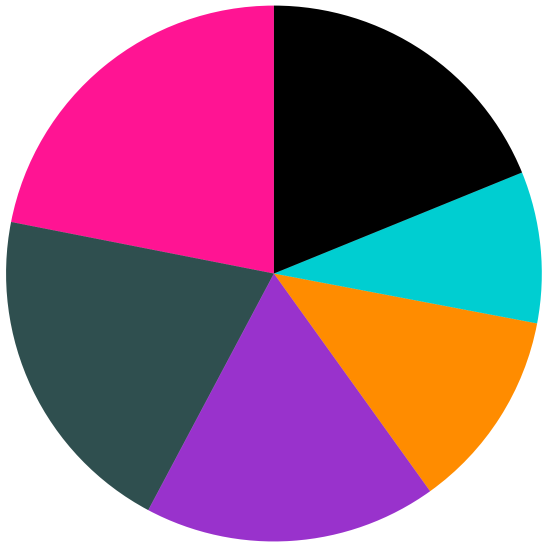

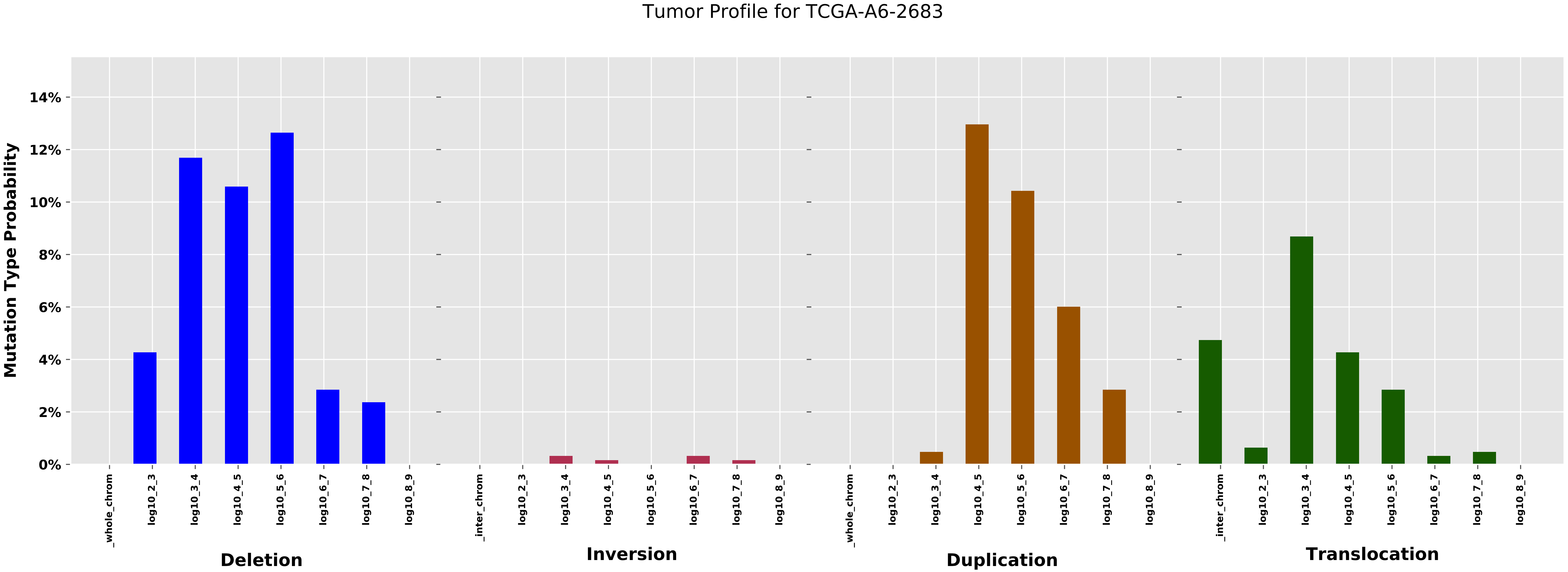
Tumor Profile for TCGA-A6-2683
Mutation Type Probability
0%
2%
4%
6%
8%
10%
12%
14%
_whole_chrom
log10_2_3
log10_3_4
log10_4_5
log10_5_6
log10_6_7
log10_7_8
log10_8_9
_inter_chrom
Deletion
Inversion
Duplication
Translocation

Cancer processes Weights for TCGA-AC-A2BK

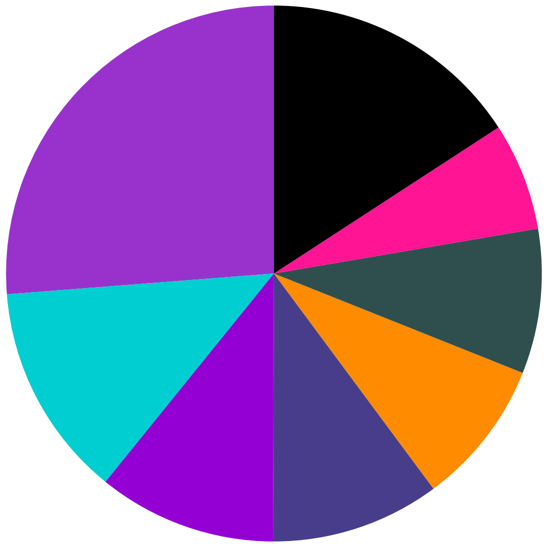

- Signature B, 26.23%
- Signature I, 12.94%
- Signature J, 10.77%
- Signature F, 10.21%
- Signature A, 8.8%
- Signature G, 8.73%
- Signature K, 6.49%
- Other Signatures Below Significance Threshold, 15.82%

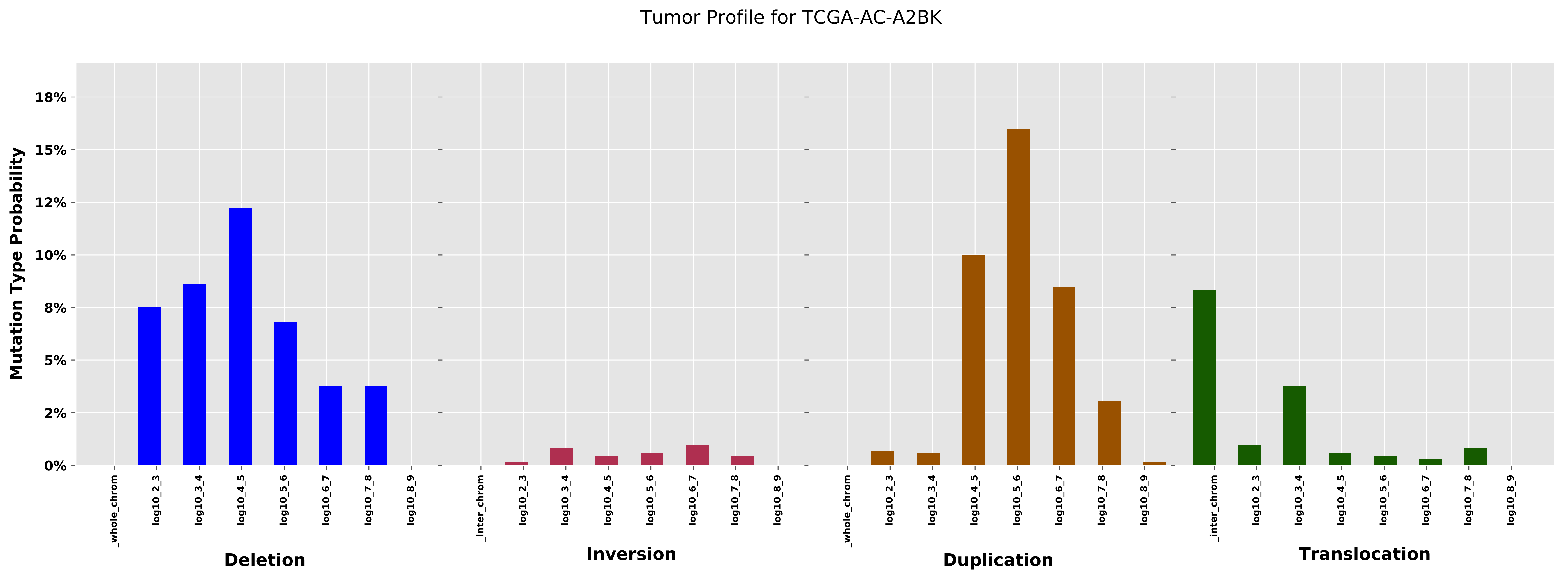

Tumor Profile for TCGA-AC-A2BK
Mutation Type Probability
18%
15%
12%
10%
8%
5%
2%
0%
_whole_chrom
log10_2_3
log10_3_4
log10_4_5
log10_5_6
log10_6_7
log10_7_8
log10_8_9
Deletion
_inter_chrom
Inversion
Duplication
Translocation

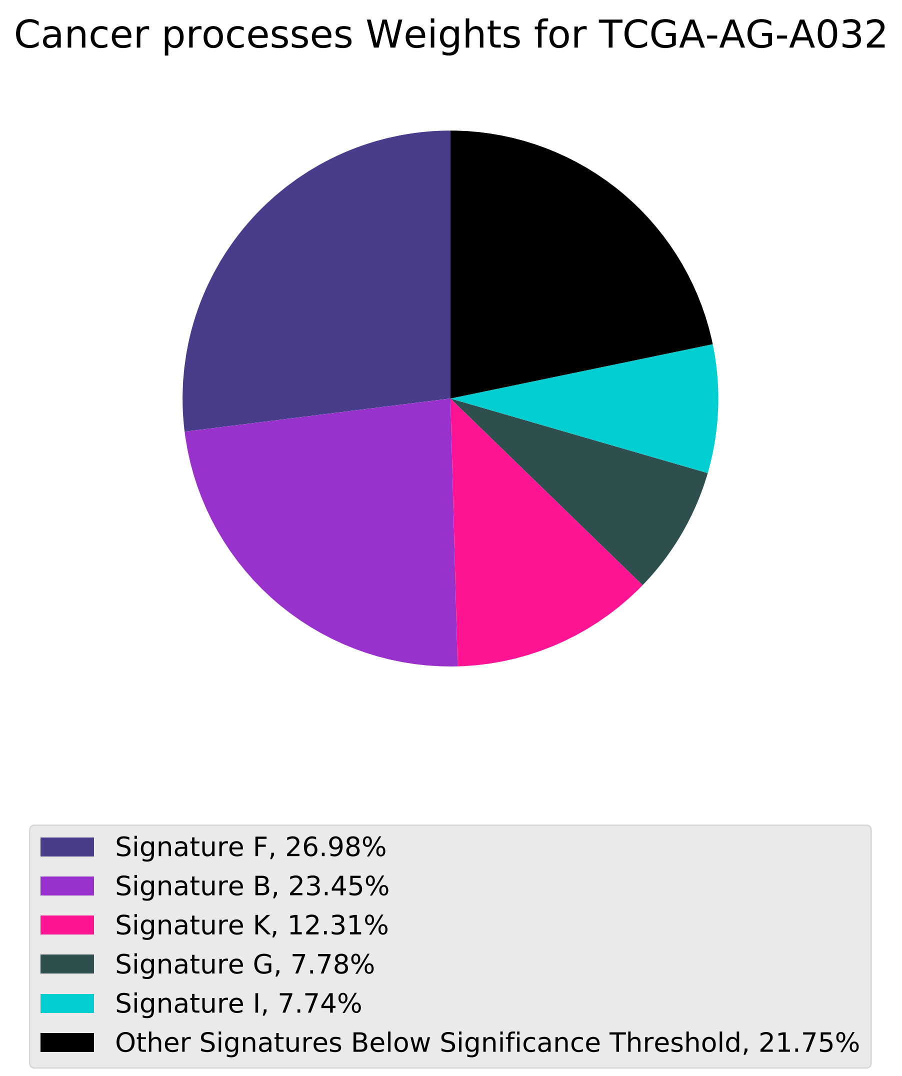
Cancer processes Weights for TCGA-AG-A032
Signature F, 26.98%
Signature B, 23.45%
Signature K, 12.31%
Signature G, 7.78%
Signature I, 7.74%
Other Signatures Below Significance Threshold, 21.75%

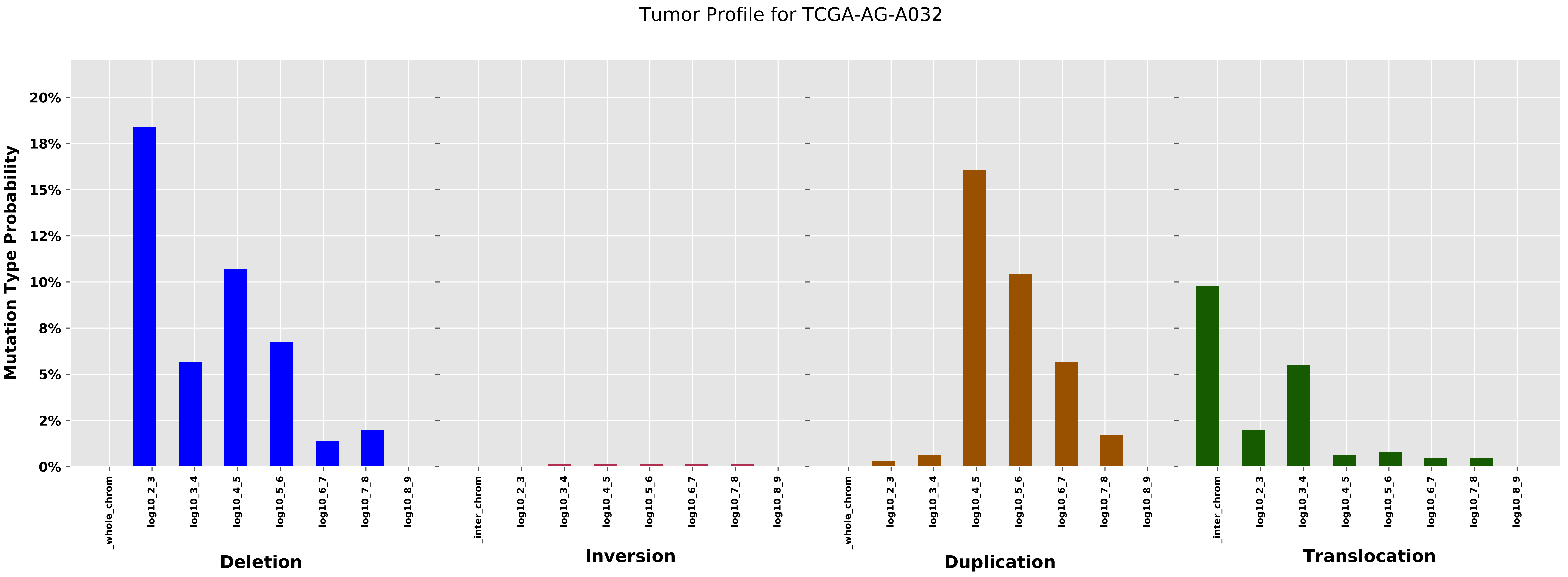
Tumor Profile for TCGA-AG-A032
Mutation Type Probability
20%
18%
15%
12%
10%
8%
5%
2%
0%
_whole_chrom
log10_2_3
log10_3_4
log10_4_5
log10_5_6
log10_6_7
log10_7_8
log10_8_9
_inter_chrom
Deletion
Inversion
Duplication
Translocation

Cancer processes Weights for TCGA-B6-A0I2

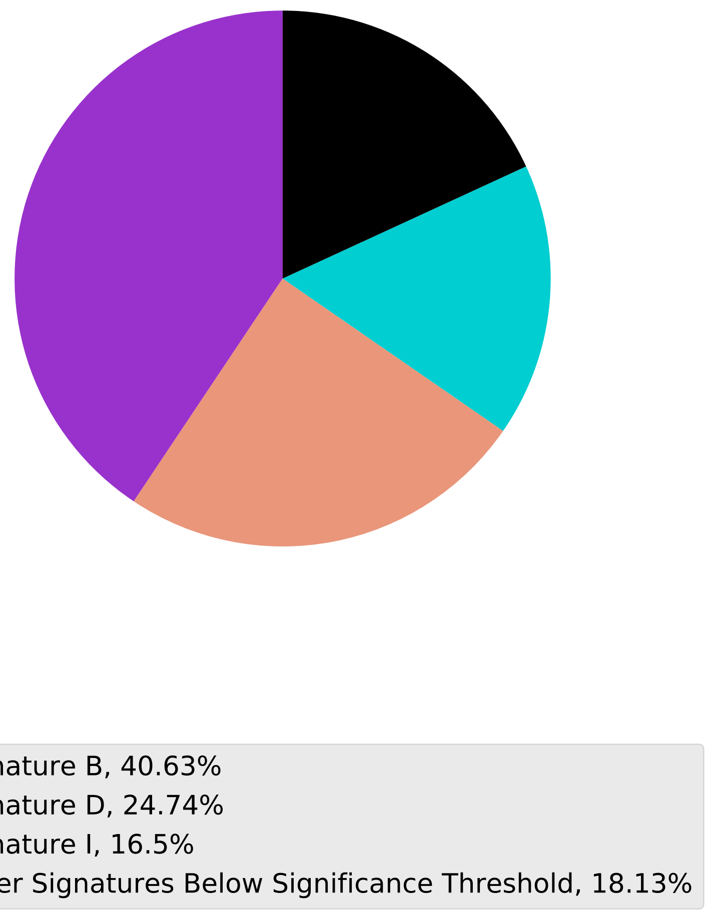

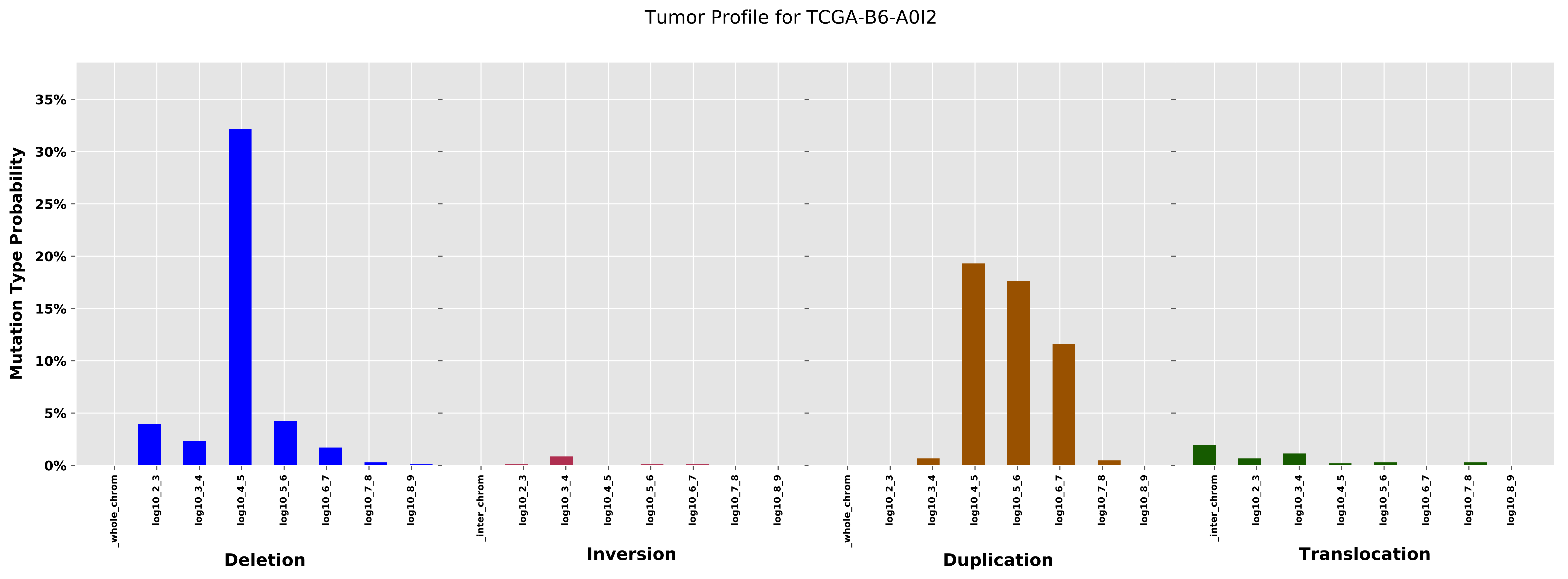
Tumor Profile for TCGA-B6-A0I2
Mutation Type Probability
0%
5%
10%
15%
20%
25%
30%
35%
_whole_chrom
log10_2_3
log10_3_4
log10_4_5
log10_5_6
log10_6_7
log10_7_8
log10_8_9
_inter_chrom
Deletion
Inversion
Duplication
Translocation

Cancer processes Weights for TCGA-A8-A094

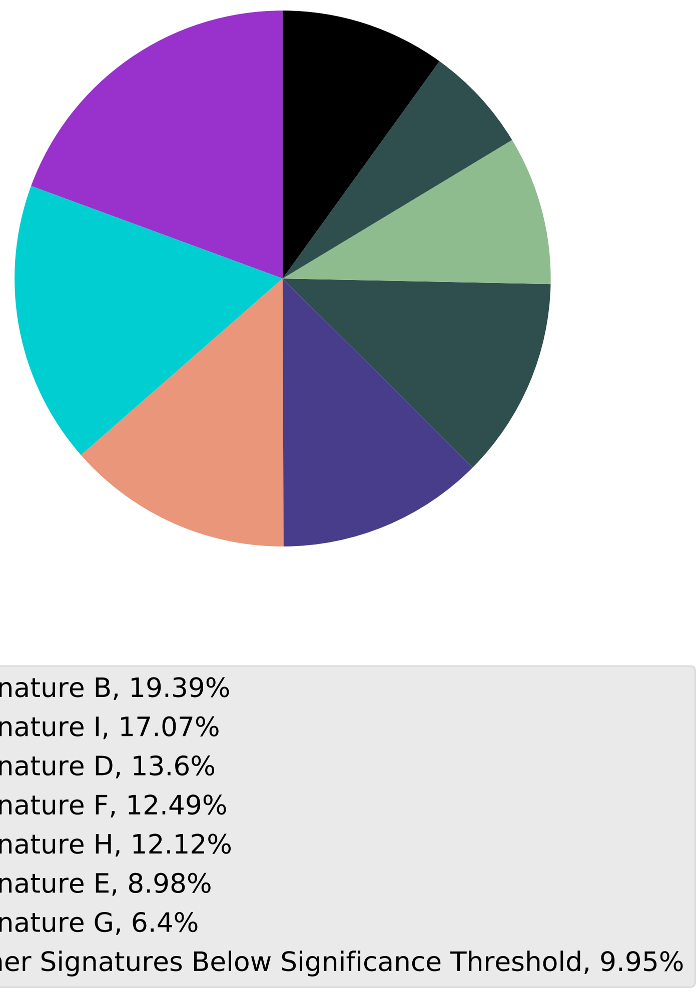

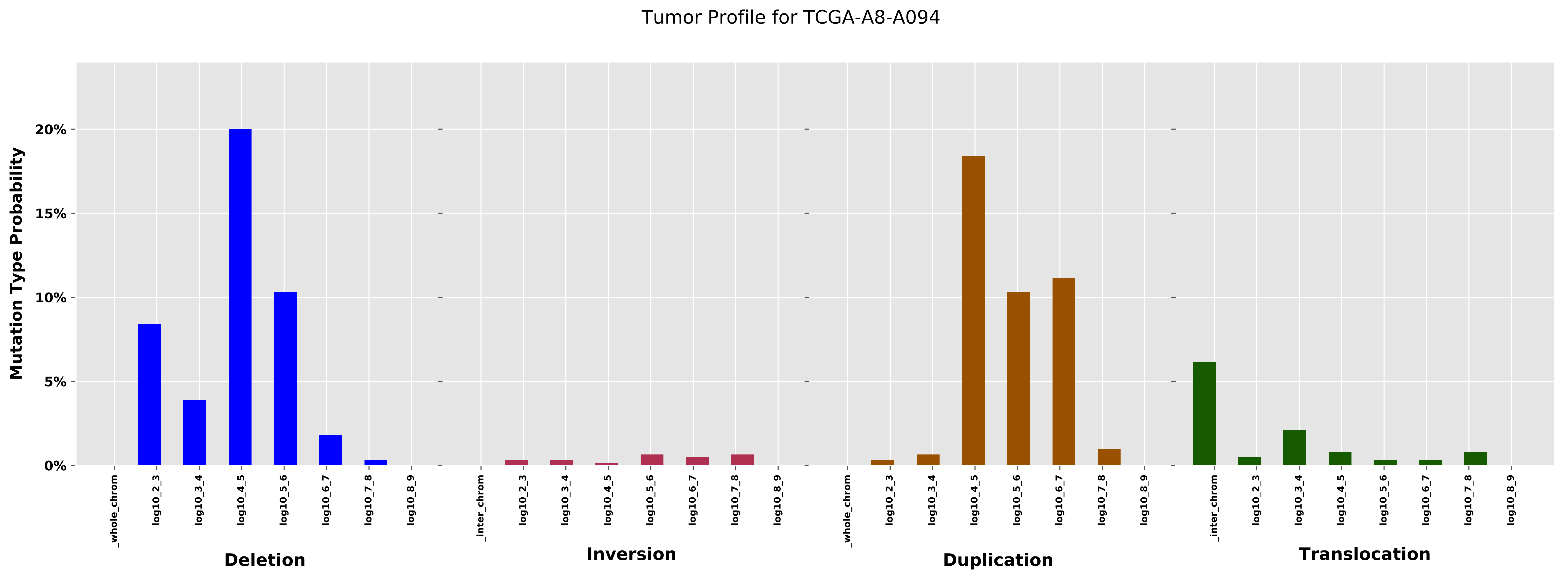
Tumor Profile for TCGA-A8-A094
Mutation Type Probability
0%
5%
10%
15%
20%
_whole_chrom
log10_2_3
log10_3_4
log10_4_5
log10_5_6
log10_6_7
log10_7_8
log10_8_9
_inter_chrom
Deletion
Inversion
Duplication
Translocation

Cancer processes Weights for TCGA-AA-3994

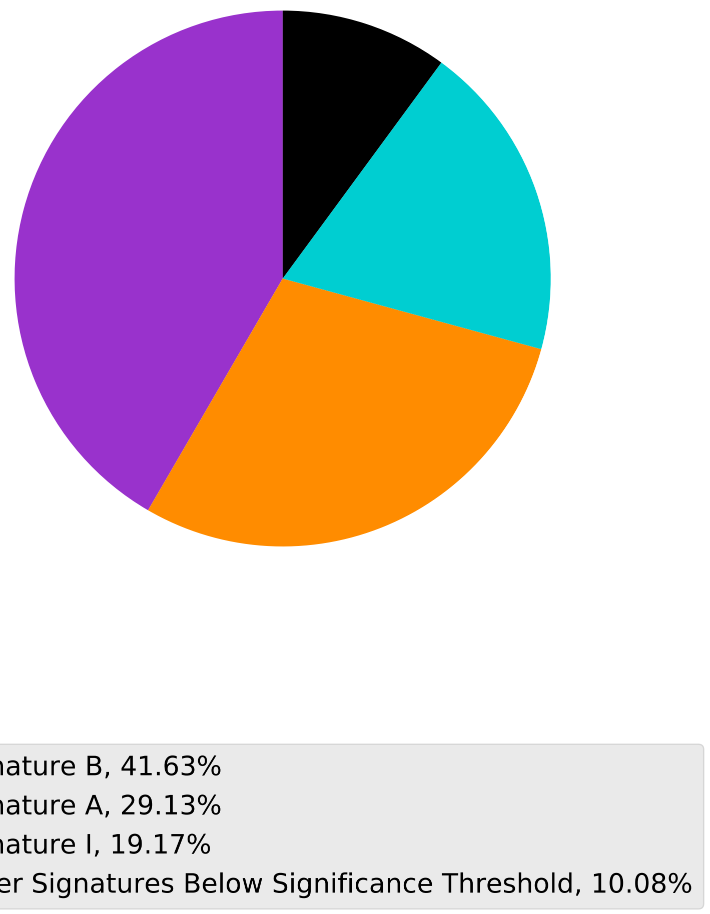

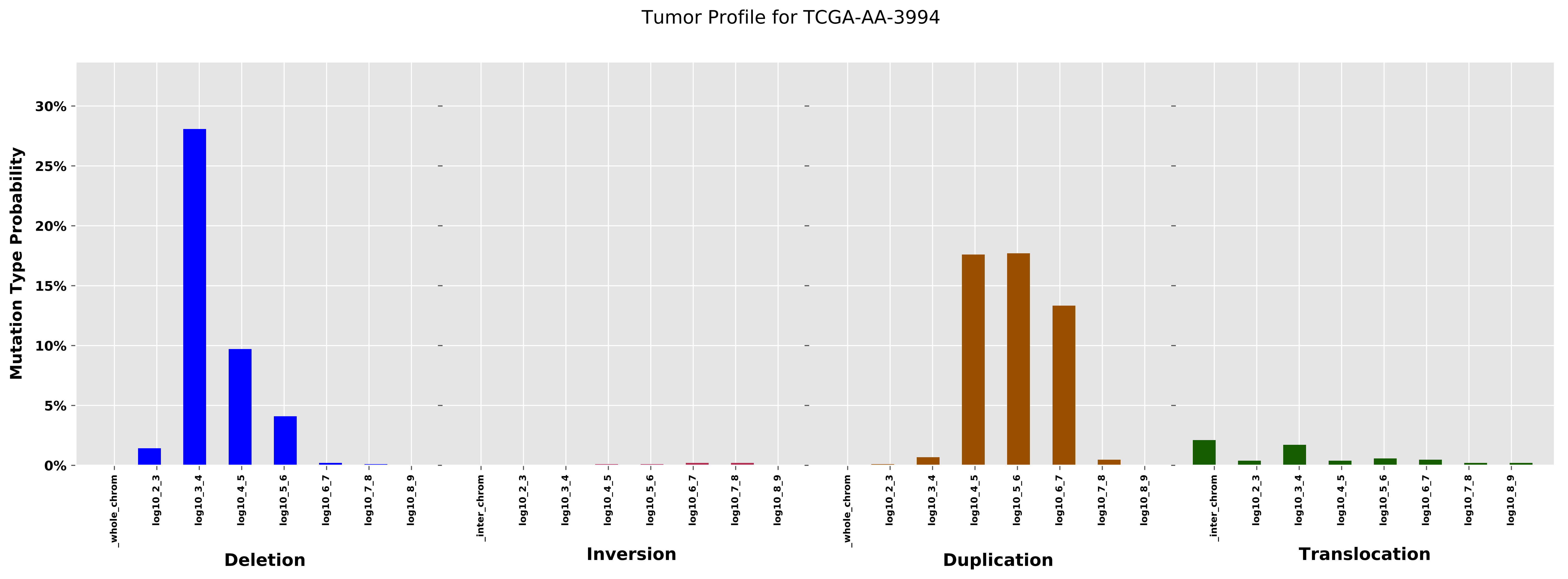
Tumor Profile for TCGA-AA-3994
Mutation Type Probability
30%
25%
20%
15%
10%
5%
0%
_whole_chrom
log10_2_3
log10_3_4
log10_4_5
log10_5_6
log10_6_7
log10_7_8
log10_8_9
_inter_chrom
Deletion
Inversion
Duplication
Translocation

#### Cancer processes Weights for TCGA-AG-3574

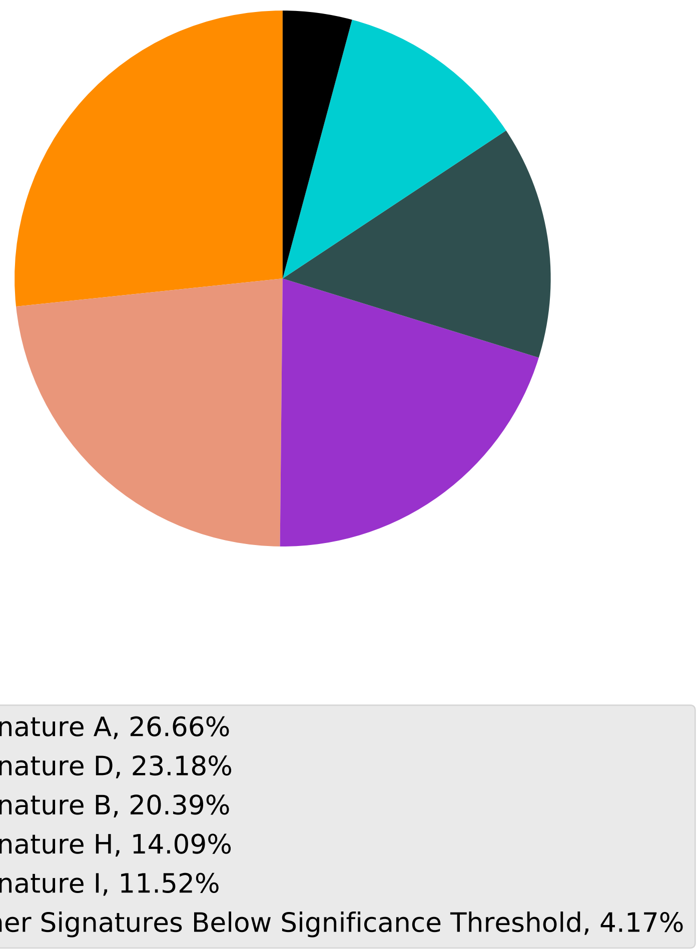

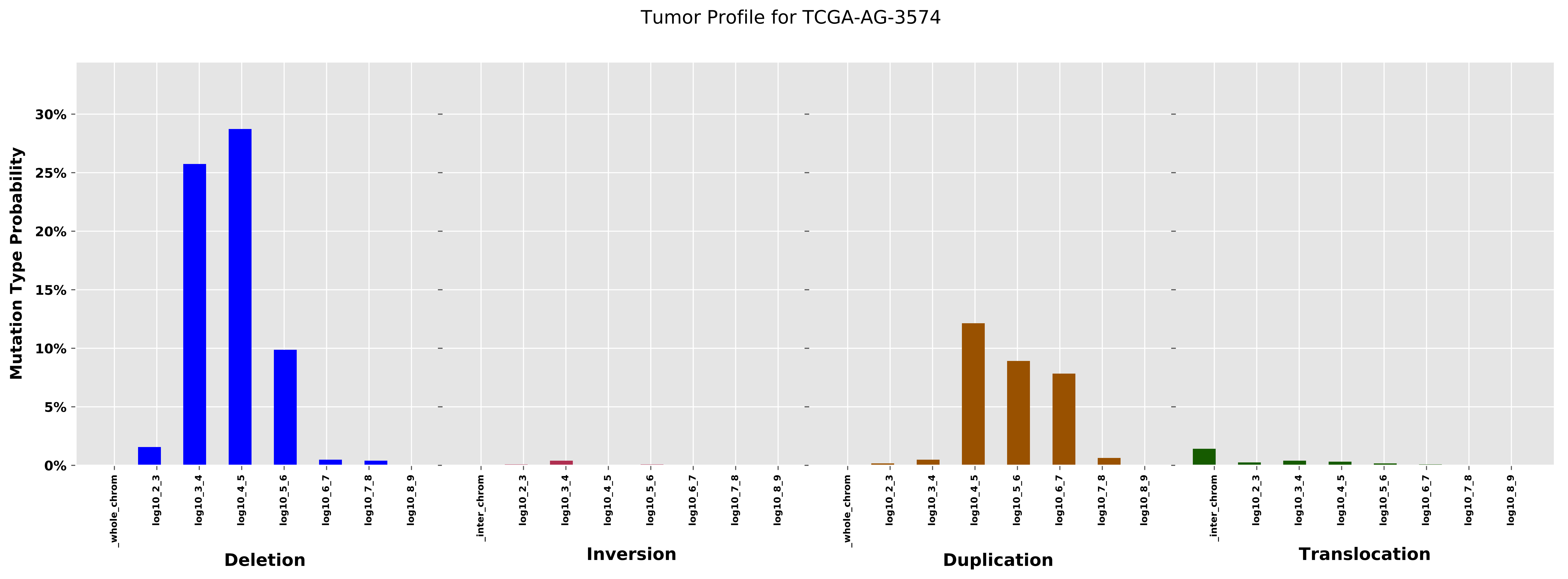
Tumor Profile for TCGA-AG-3574
Mutation Type Probability
0%
5%
10%
15%
20%
25%
30%
_whole_chrom
log10_2_3
log10_3_4
log10_4_5
log10_5_6
log10_6_7
log10_7_8
log10_8_9
_inter_chrom
Deletion
Inversion
Duplication
Translocation

Cancer processes Weights for TCGA-AO-A03L

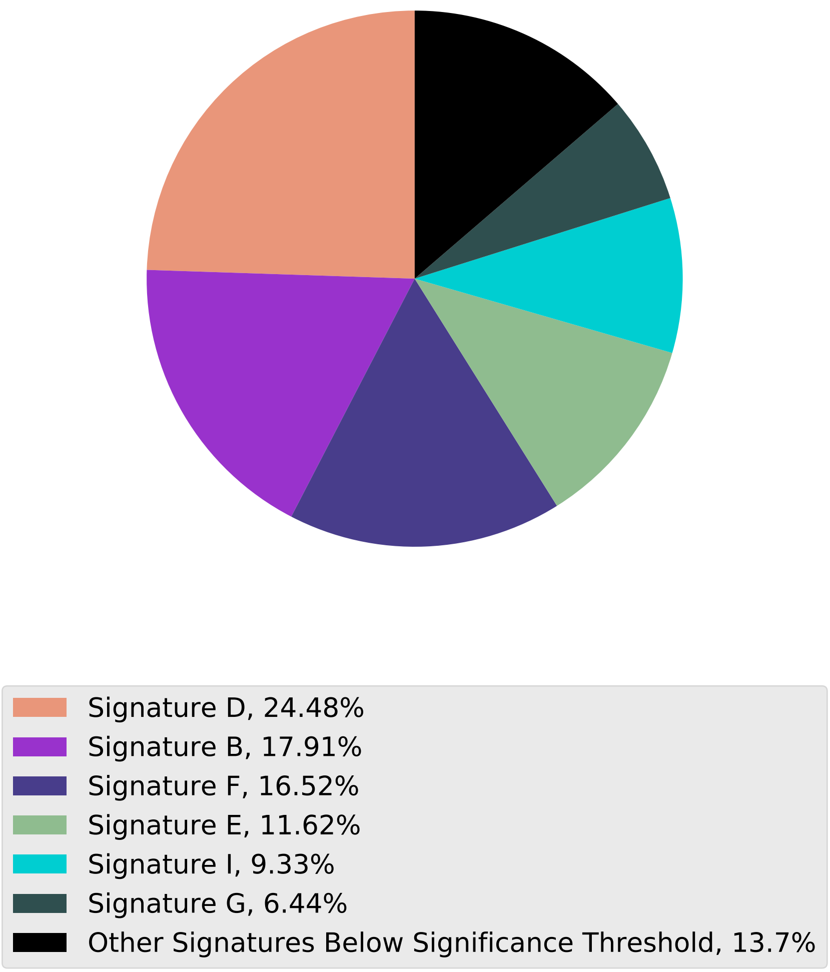

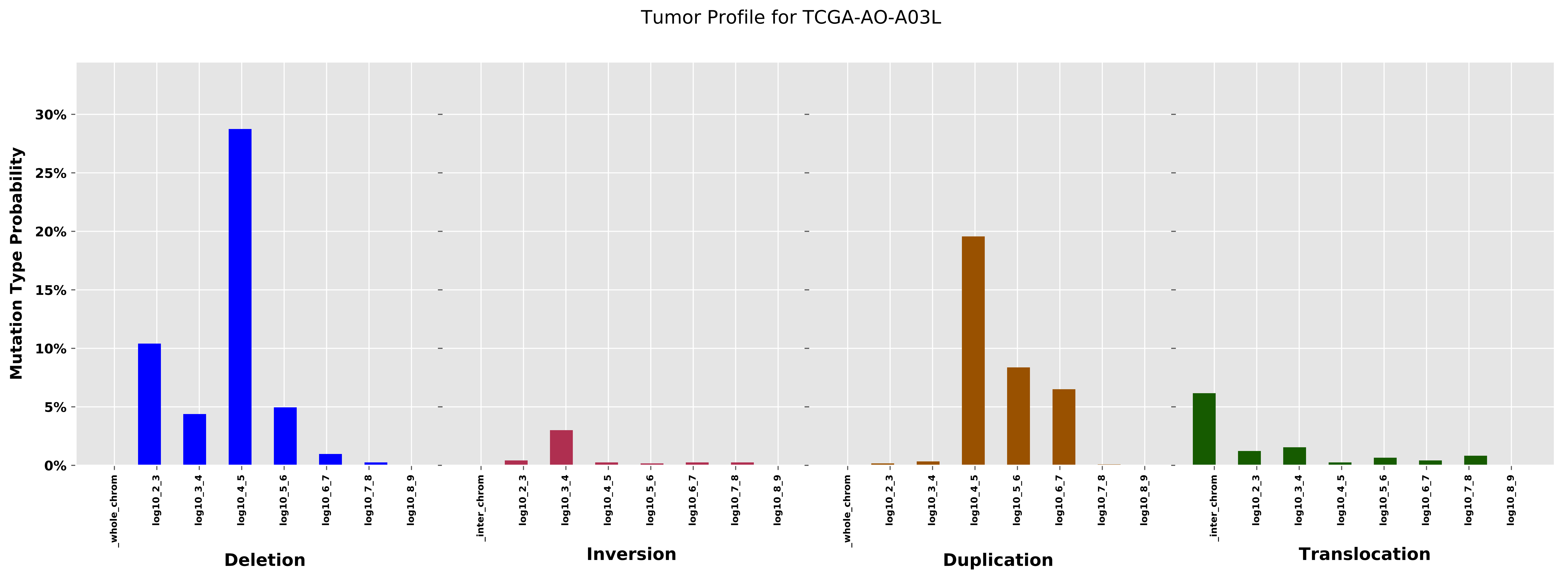
Tumor Profile for TCGA-AO-A03L
Mutation Type Probability
30%
25%
20%
15%
10%
5%
0%
_whole_chrom
log10_2_3
log10_3_4
log10_4_5
log10_5_6
log10_6_7
log10_7_8
log10_8_9
Deletion
_inter_chrom
log10_2_3
log10_3_4
log10_4_5
log10_5_6
log10_6_7
log10_7_8
log10_8_9
Inversion
_whole_chrom
log10_2_3
log10_3_4
log10_4_5
log10_5_6
log10_6_7
log10_7_8
log10_8_9
Duplication
_inter_chrom
log10_2_3
log10_3_4
log10_4_5
log10_5_6
log10_6_7
log10_7_8
log10_8_9
Translocation

Cancer processes Weights for TCGA-A6-6141

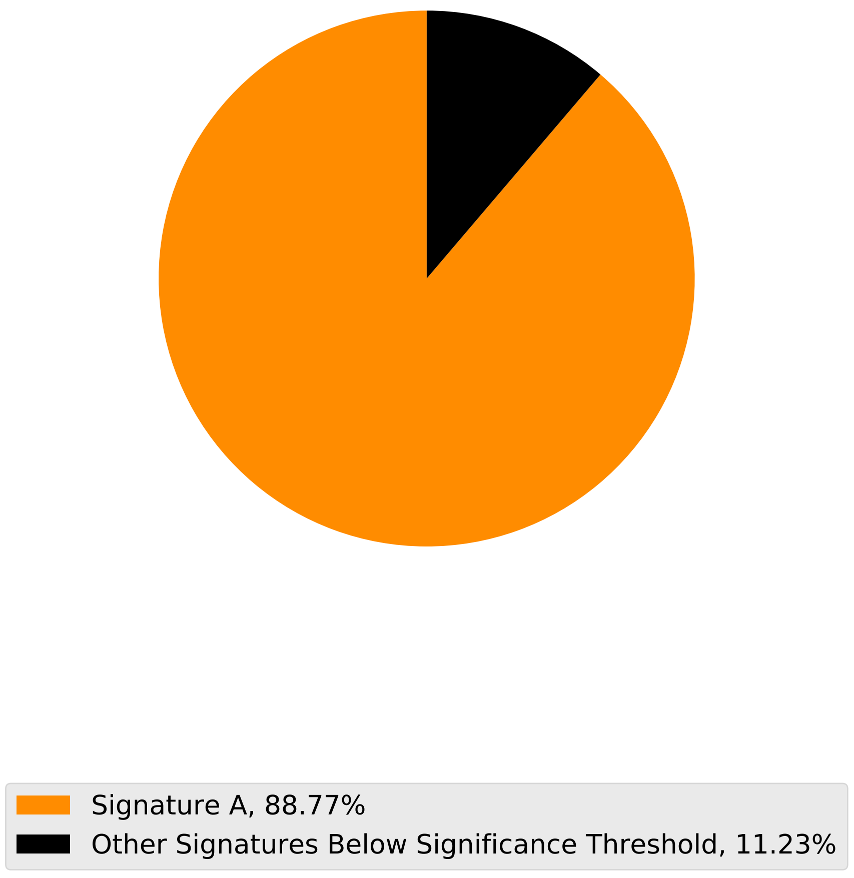

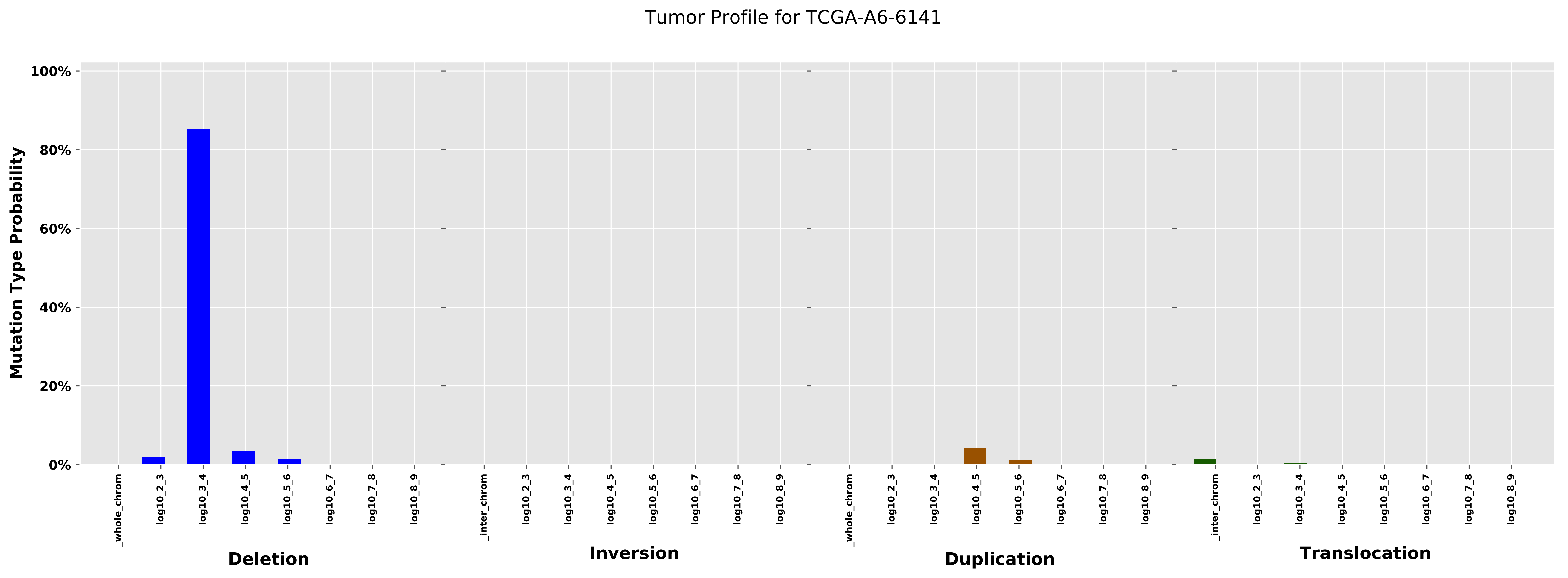
Tumor Profile for TCGA-A6-6141
Mutation Type Probability
100%
80%
60%
40%
20%
0%
_whole_chrom
log10_2_3
log10_3_4
log10_4_5
log10_5_6
log10_6_7
log10_7_8
log10_8_9
_inter_chrom
Deletion
Inversion
Duplication
Translocation

#### Cancer processes Weights for TCGA-AA-A02O

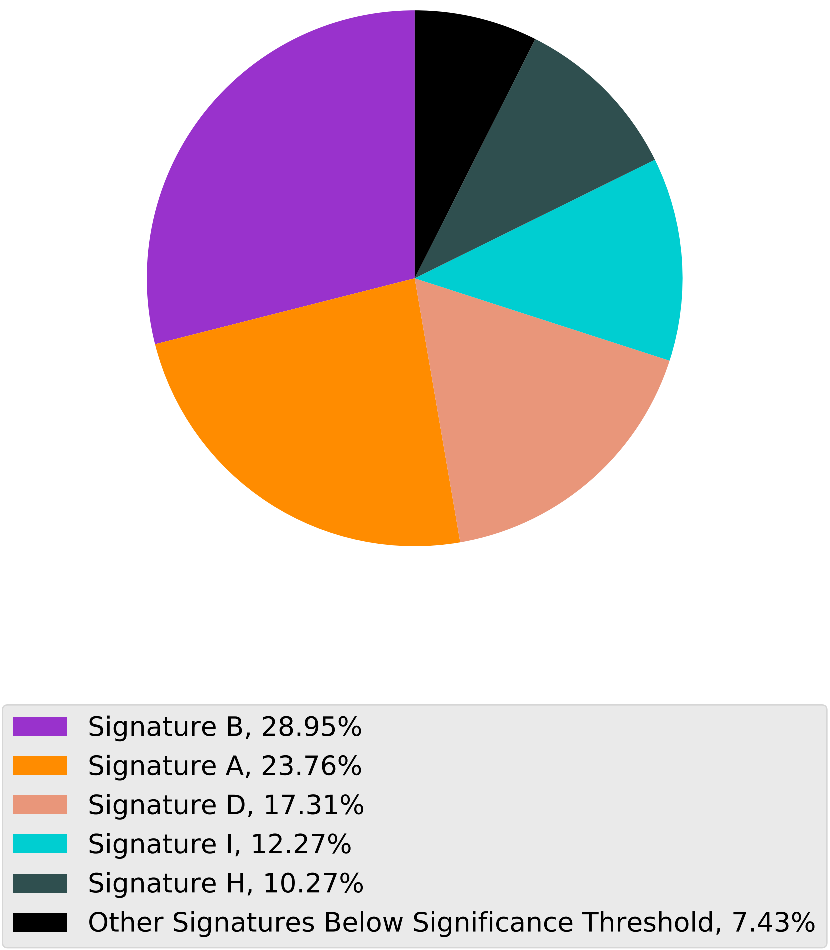

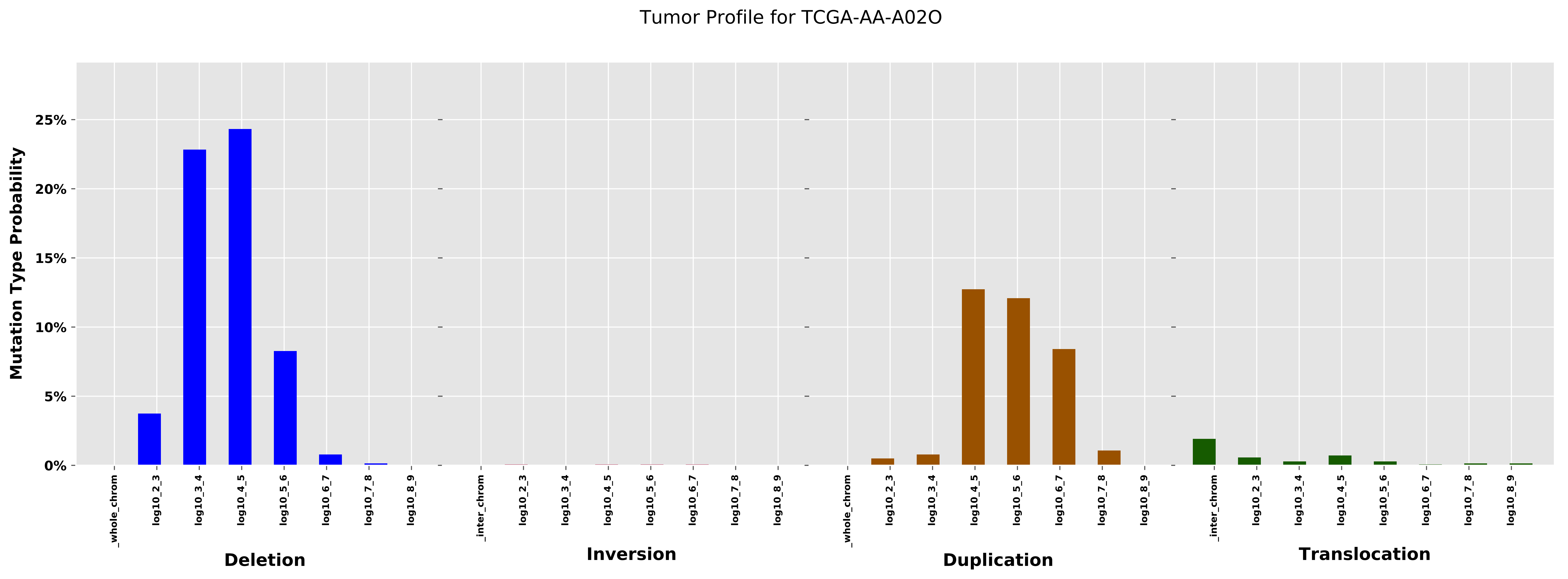

Tumor Profile for TCGA-AA-A02O
Mutation Type Probability
25%
20%
15%
10%
5%
0%
_whole_chrom
log10_2_3
log10_3_4
log10_4_5
log10_5_6
log10_6_7
log10_7_8
log10_8_9
Deletion
_inter_chrom
log10_2_3
log10_3_4
log10_4_5
log10_5_6
log10_6_7
log10_7_8
log10_8_9
Inversion
_whole_chrom
log10_2_3
log10_3_4
log10_4_5
log10_5_6
log10_6_7
log10_7_8
log10_8_9
Duplication
_inter_chrom
log10_2_3
log10_3_4
log10_4_5
log10_5_6
log10_6_7
log10_7_8
log10_8_9
Translocation

Cancer processes Weights for TCGA-AG-3885

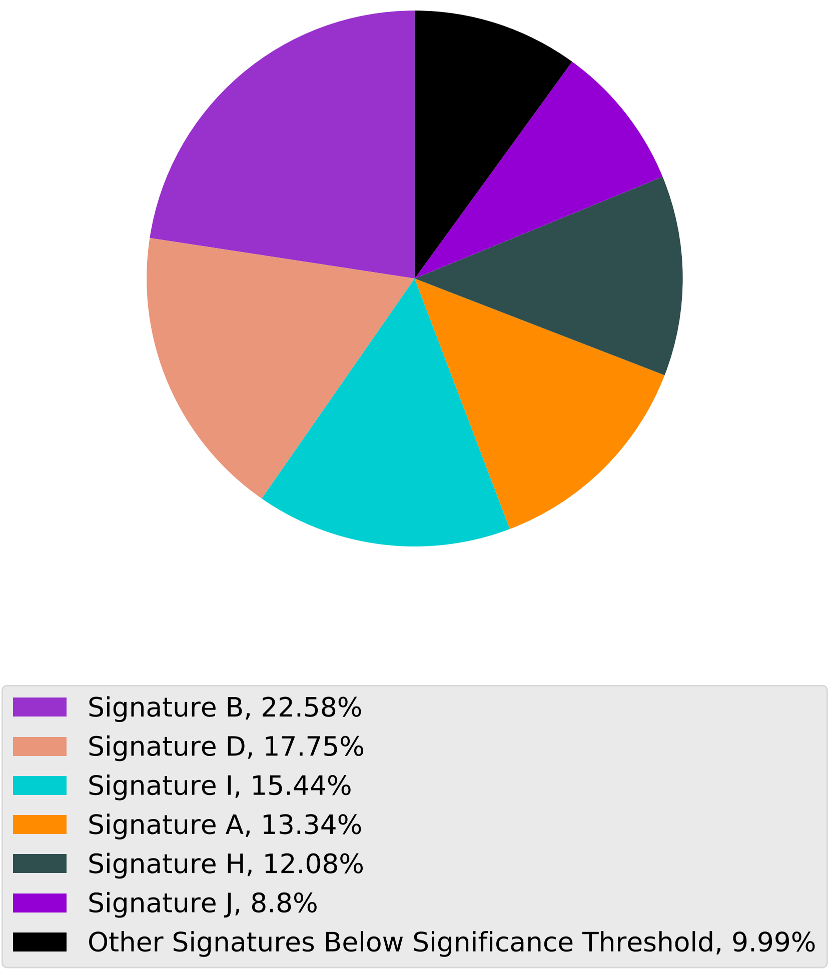

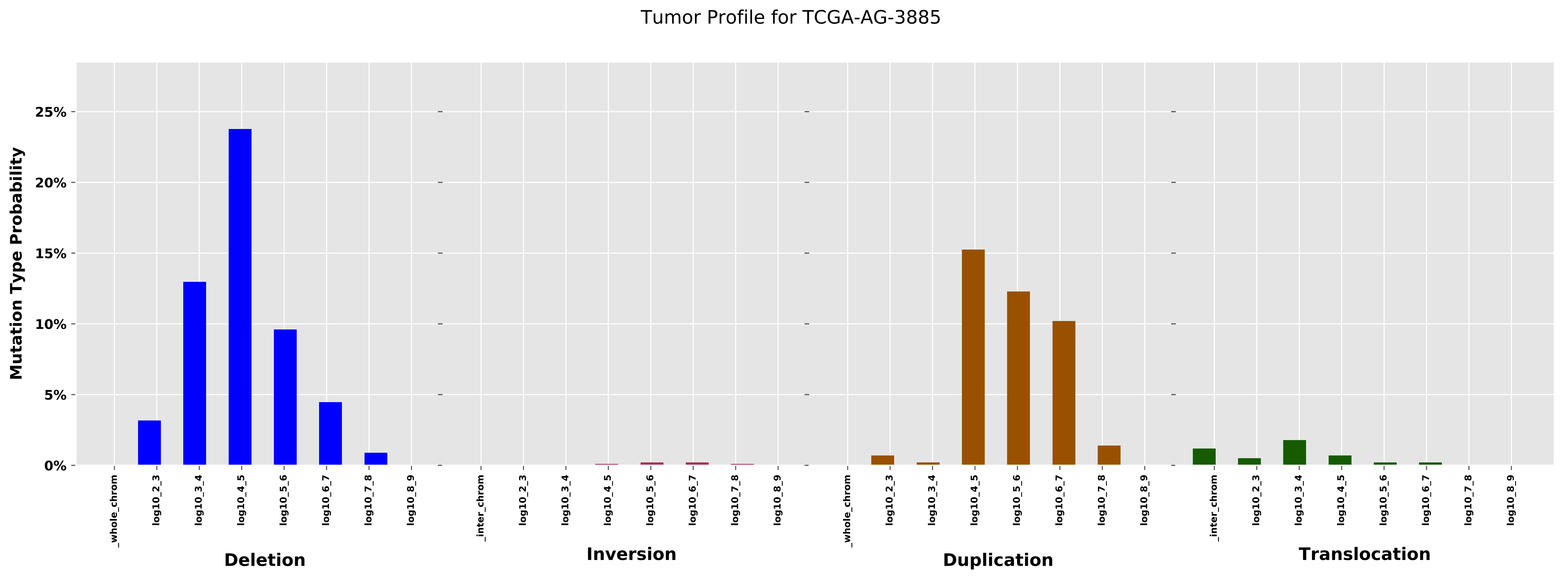
Tumor Profile for TCGA-AG-3885
Mutation Type Probability
25%
20%
15%
10%
5%
0%
_whole_chrom
log10_2_3
log10_3_4
log10_4_5
log10_5_6
log10_6_7
log10_7_8
log10_8_9
Deletion
_inter_chrom
Inversion
Duplication
Translocation

Cancer processes Weights for TCGA-AA-A01T

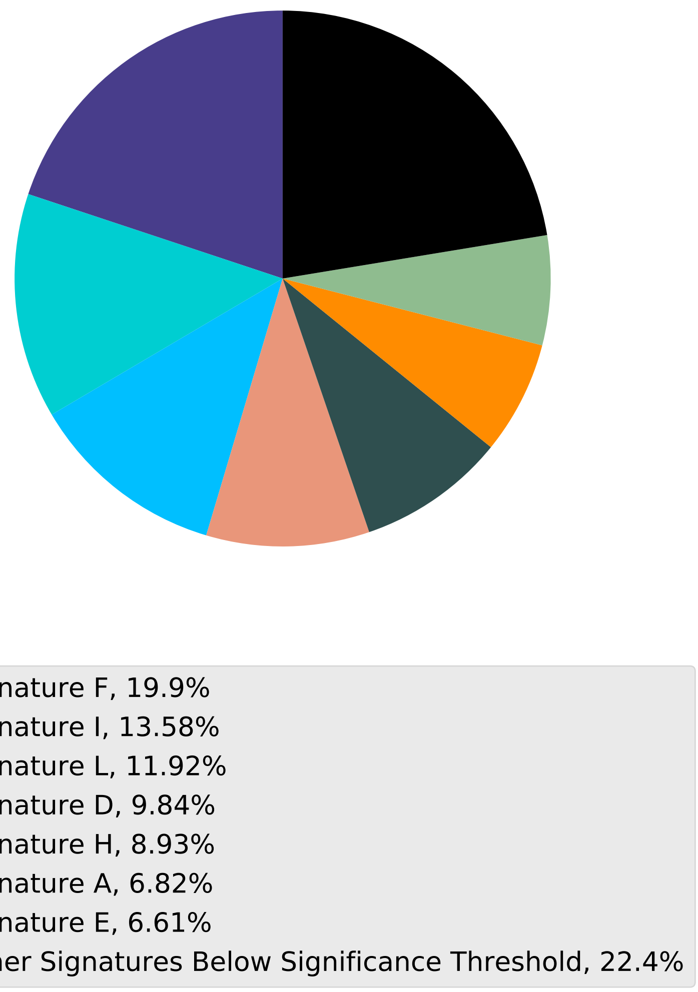

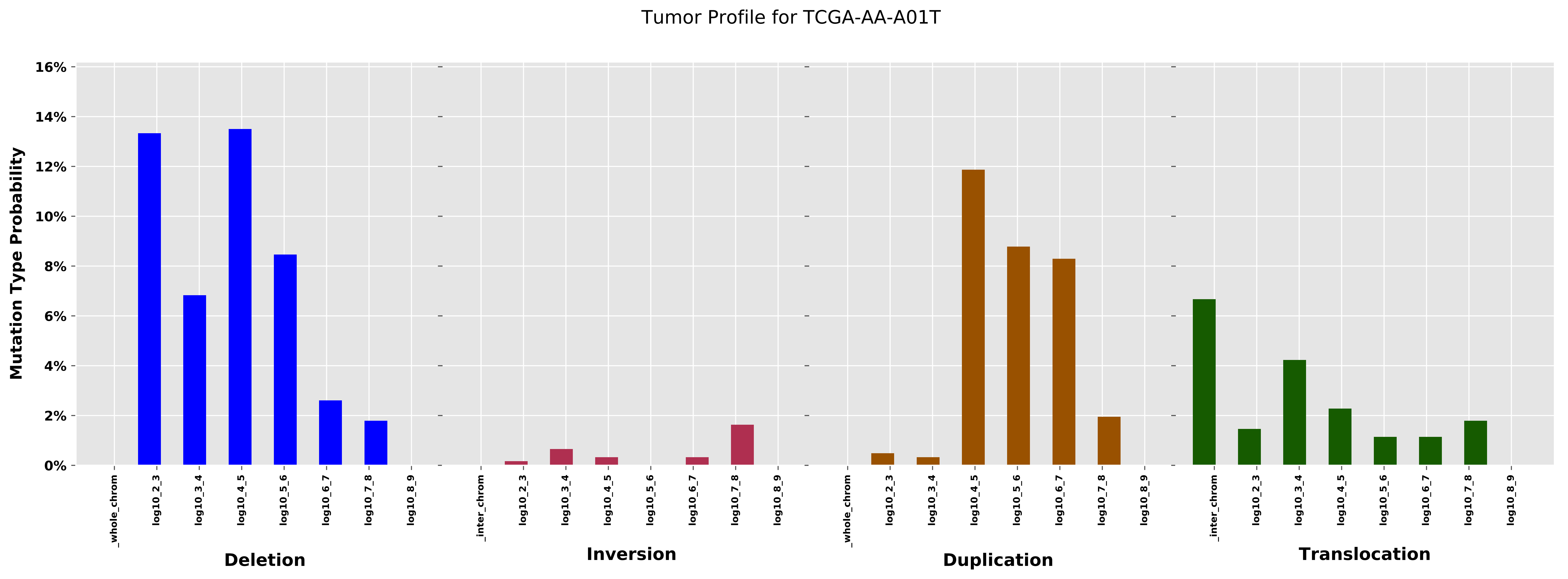

Tumor Profile for TCGA-AA-A01T
Mutation Type Probability
16%
14%
12%
10%
8%
6%
4%
2%
0%
_whole_chrom
log10_2_3
log10_3_4
log10_4_5
log10_5_6
log10_6_7
log10_7_8
log10_8_9
Deletion
_inter_chrom
log10_2_3
log10_3_4
log10_4_5
log10_5_6
log10_6_7
log10_7_8
log10_8_9
Inversion
_whole_chrom
log10_2_3
log10_3_4
log10_4_5
log10_5_6
log10_6_7
log10_7_8
log10_8_9
Duplication
_inter_chrom
log10_2_3
log10_3_4
log10_4_5
log10_5_6
log10_6_7
log10_7_8
log10_8_9
Translocation

Cancer processes Weights for TCGA-A8-A07I

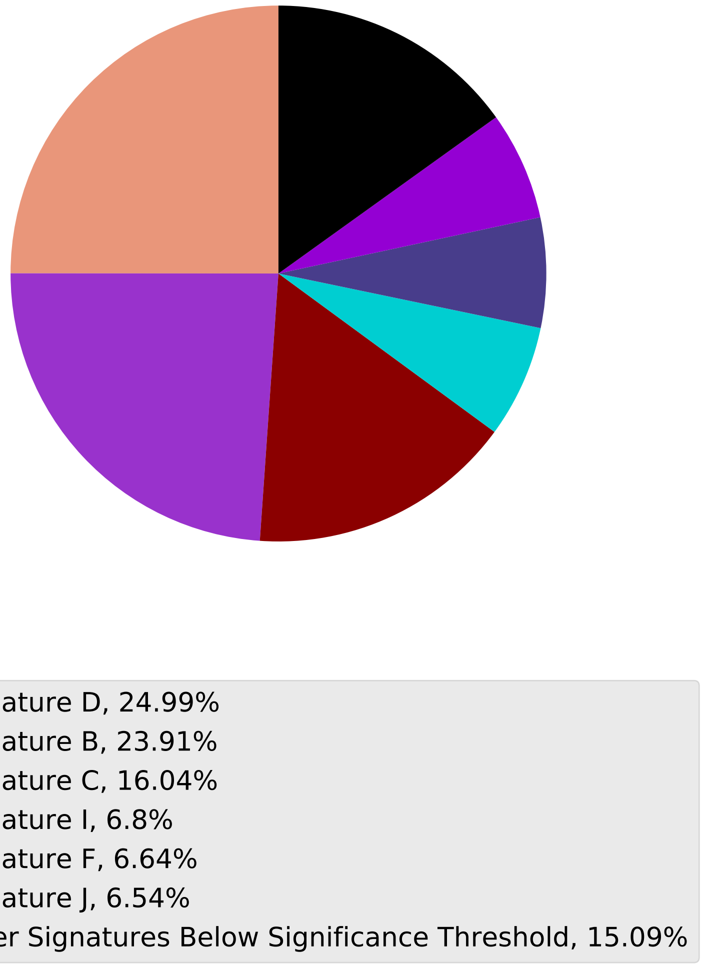

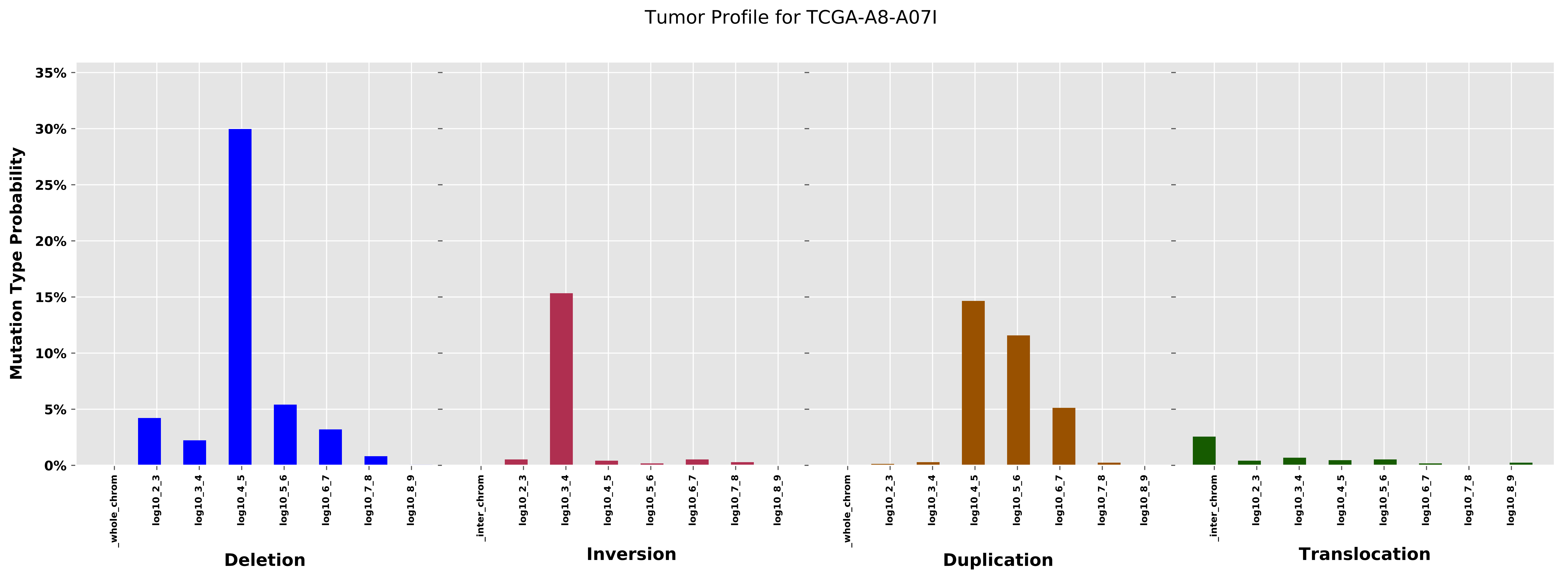
Tumor Profile for TCGA-A8-A07I
Mutation Type Probability
35%
30%
25%
20%
15%
10%
5%
0%
_whole_chrom
log10_2_3
log10_3_4
log10_4_5
log10_5_6
log10_6_7
log10_7_8
log10_8_9
Deletion
_inter_chrom
log10_2_3
log10_3_4
log10_4_5
log10_5_6
log10_6_7
log10_7_8
log10_8_9
Inversion
_whole_chrom
log10_2_3
log10_3_4
log10_4_5
log10_5_6
log10_6_7
log10_7_8
log10_8_9
Duplication
_inter_chrom
log10_2_3
log10_3_4
log10_4_5
log10_5_6
log10_6_7
log10_7_8
log10_8_9
Translocation

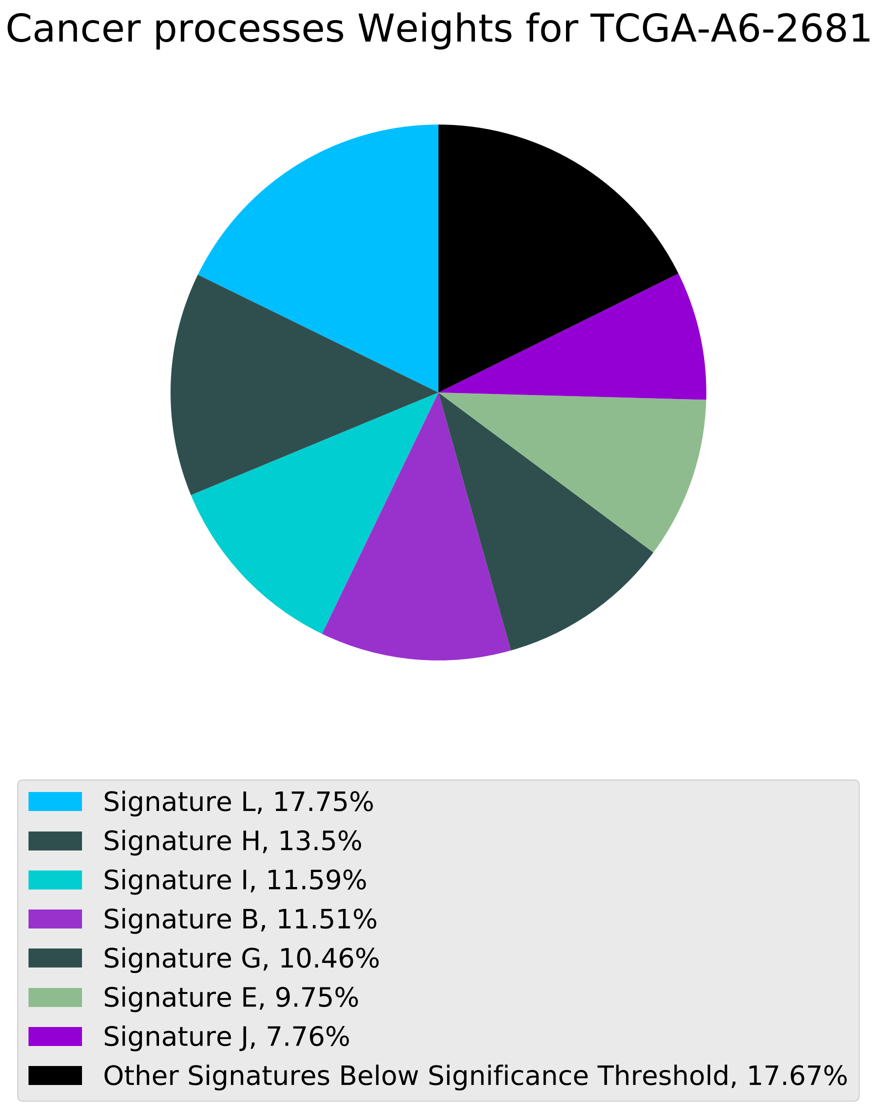

Cancer processes Weights for TCGA-A6-2681
Signature L, 17.75%
Signature H, 13.5%
Signature I, 11.59%
Signature B, 11.51%
Signature G, 10.46%
Signature E, 9.75%
Signature J, 7.76%
Other Signatures Below Significance Threshold, 17.67%

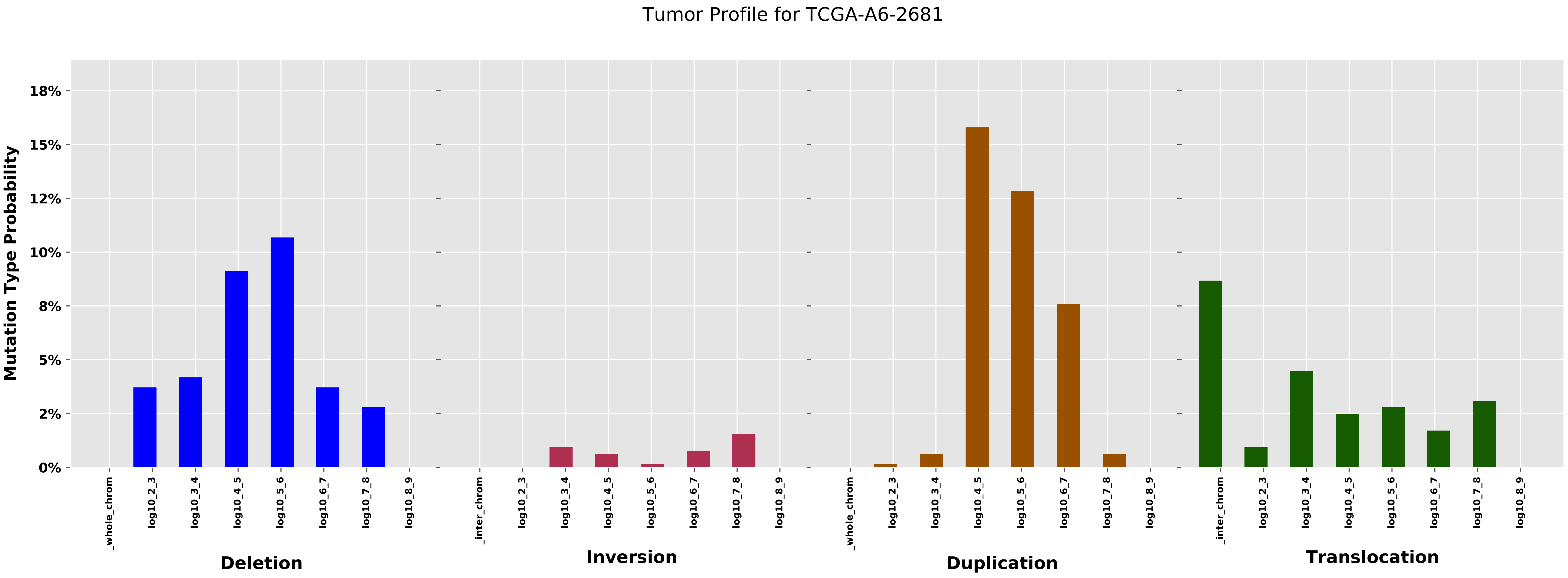

Tumor Profile for TCGA-A6-2681
Mutation Type Probability
0%
2%
5%
8%
10%
12%
15%
18%
_whole_chrom
log10_2_3
log10_3_4
log10_4_5
log10_5_6
log10_6_7
log10_7_8
log10_8_9
Deletion
_inter_chrom
Inversion
Duplication
Translocation

Cancer processes Weights for TCGA-AO-A124

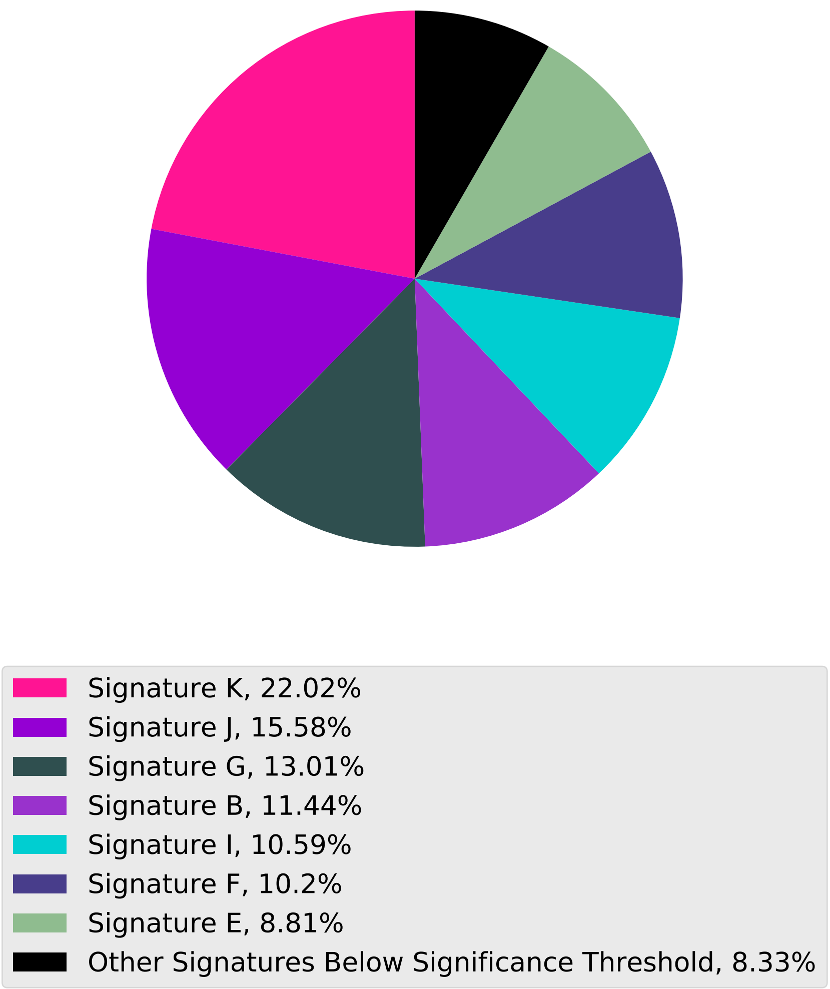

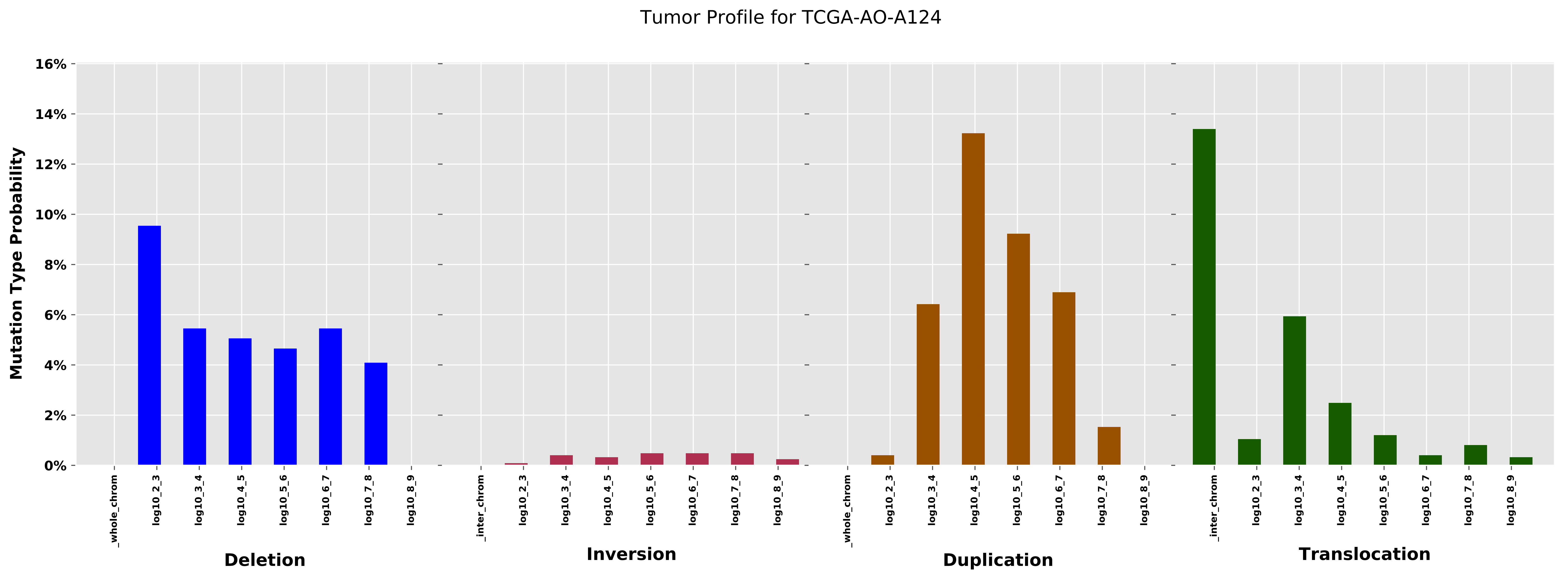
Tumor Profile for TCGA-AO-A124
Mutation Type Probability
16%
14%
12%
10%
8%
6%
4%
2%
0%
_whole_chrom
log10_2_3
log10_3_4
log10_4_5
log10_5_6
log10_6_7
log10_7_8
log10_8_9
_inter_chrom
Deletion
Inversion
Duplication
Translocation

Cancer processes Weights for TCGA-A2-A259

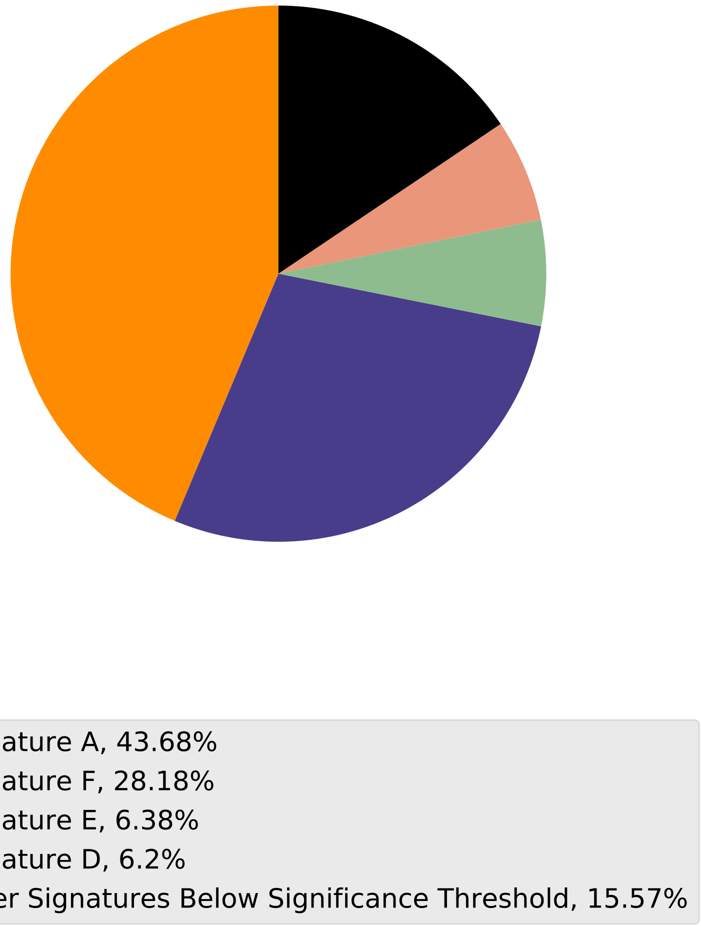

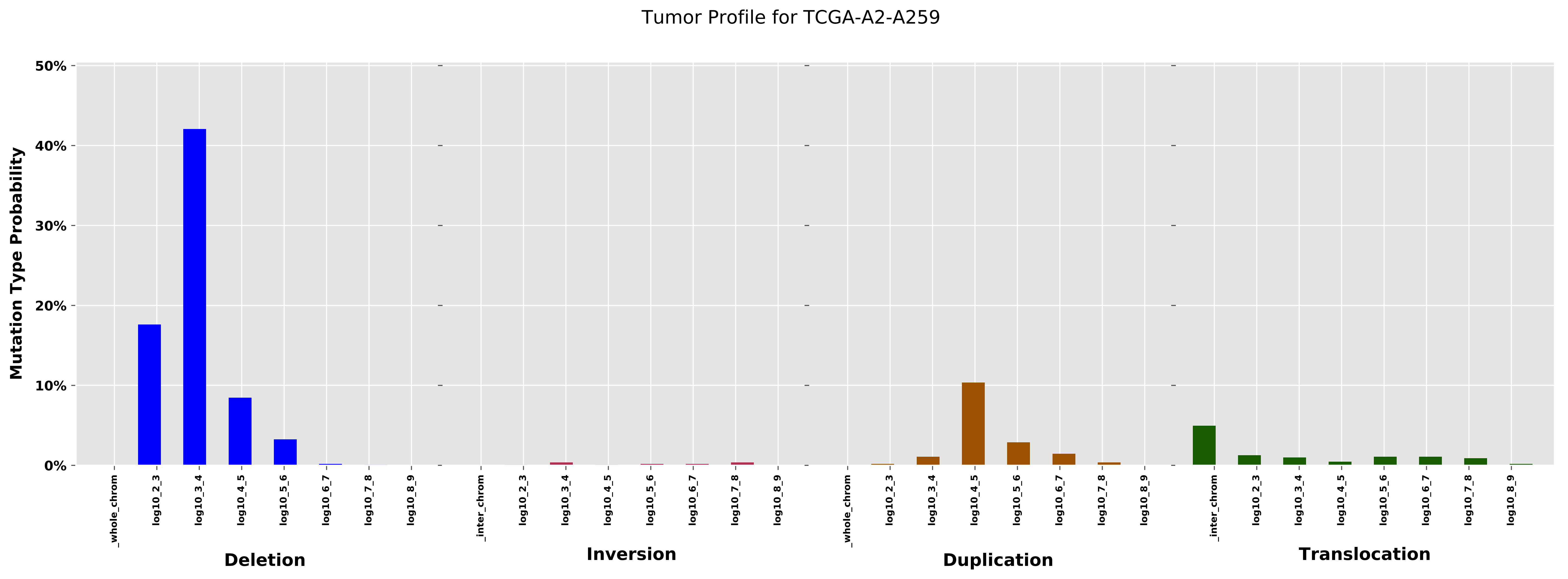

Tumor Profile for TCGA-A2-A259
Mutation Type Probability
50%
40%
30%
20%
10%
0%
_whole_chrom
log10_2_3
log10_3_4
log10_4_5
log10_5_6
log10_6_7
log10_7_8
log10_8_9
Deletion
_inter_chrom
log10_2_3
log10_3_4
log10_4_5
log10_5_6
log10_6_7
log10_7_8
log10_8_9
Inversion
_whole_chrom
log10_2_3
log10_3_4
log10_4_5
log10_5_6
log10_6_7
log10_7_8
log10_8_9
Duplication
_inter_chrom
log10_2_3
log10_3_4
log10_4_5
log10_5_6
log10_6_7
log10_7_8
log10_8_9
Translocation

Cancer processes Weights for TCGA-BH-A0DG

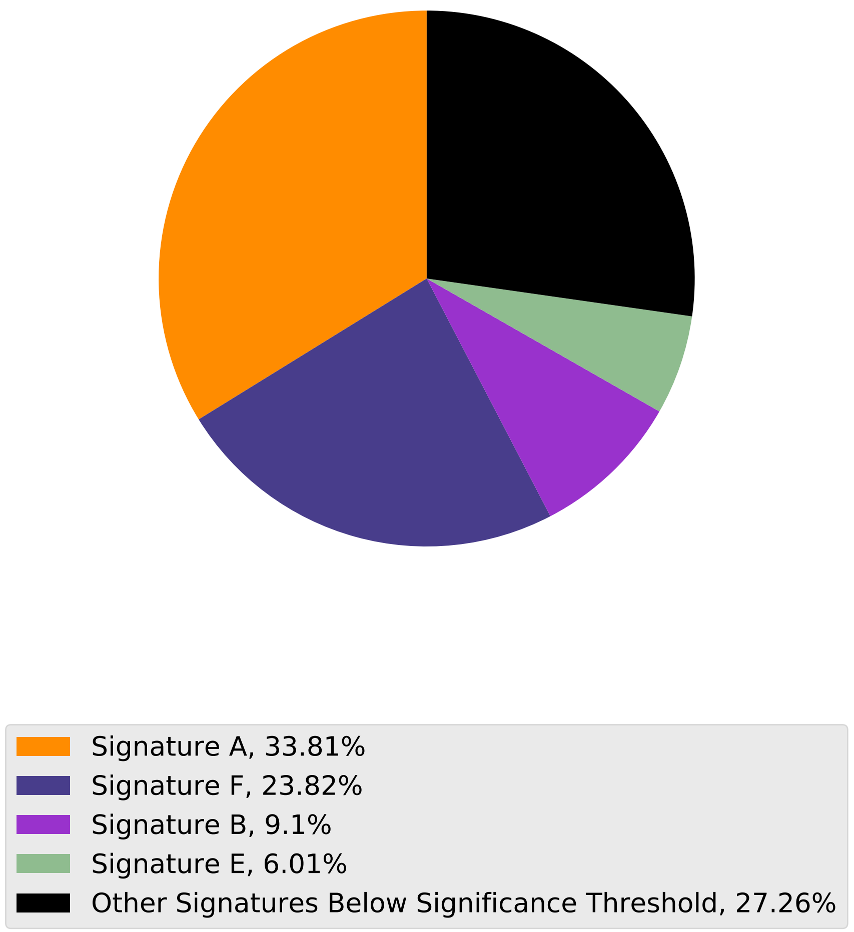

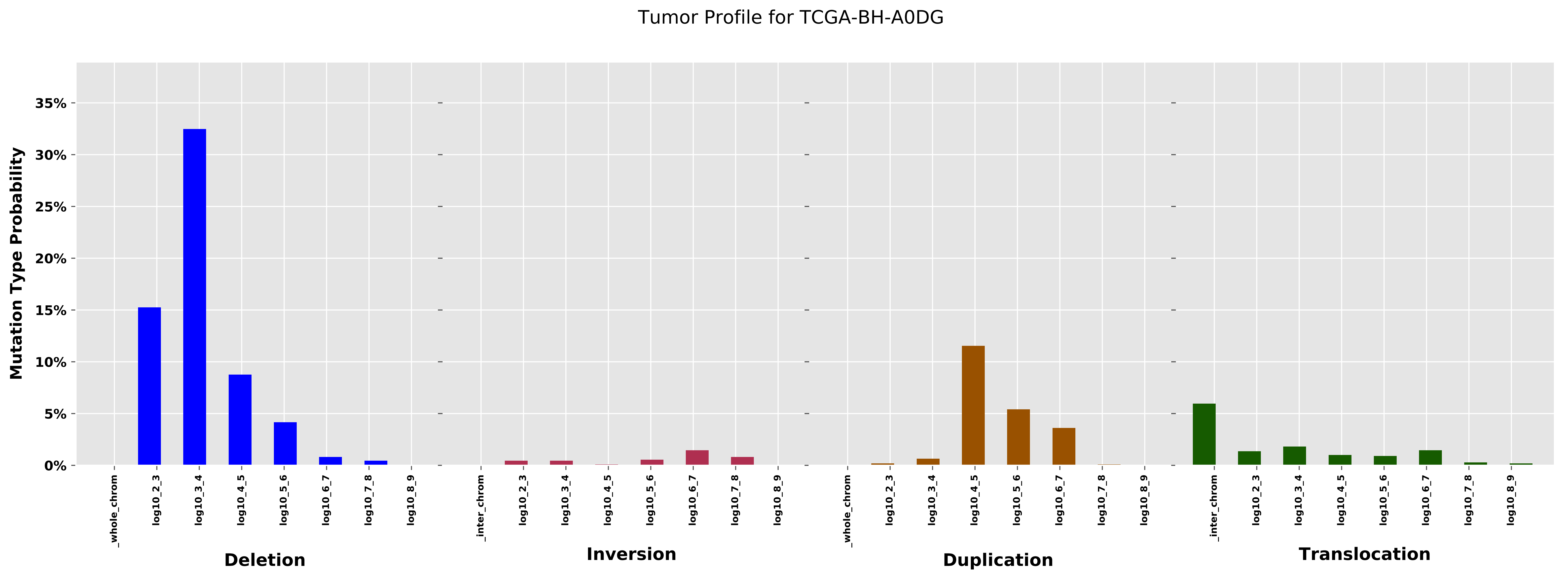
Tumor Profile for TCGA-BH-A0DG
Mutation Type Probability
0%
5%
10%
15%
20%
25%
30%
35%
_whole_chrom
log10_2_3
log10_3_4
log10_4_5
log10_5_6
log10_6_7
log10_7_8
log10_8_9
_inter_chrom
Deletion
Inversion
Duplication
Translocation

Cancer processes Weights for TCGA-A2-A3KC

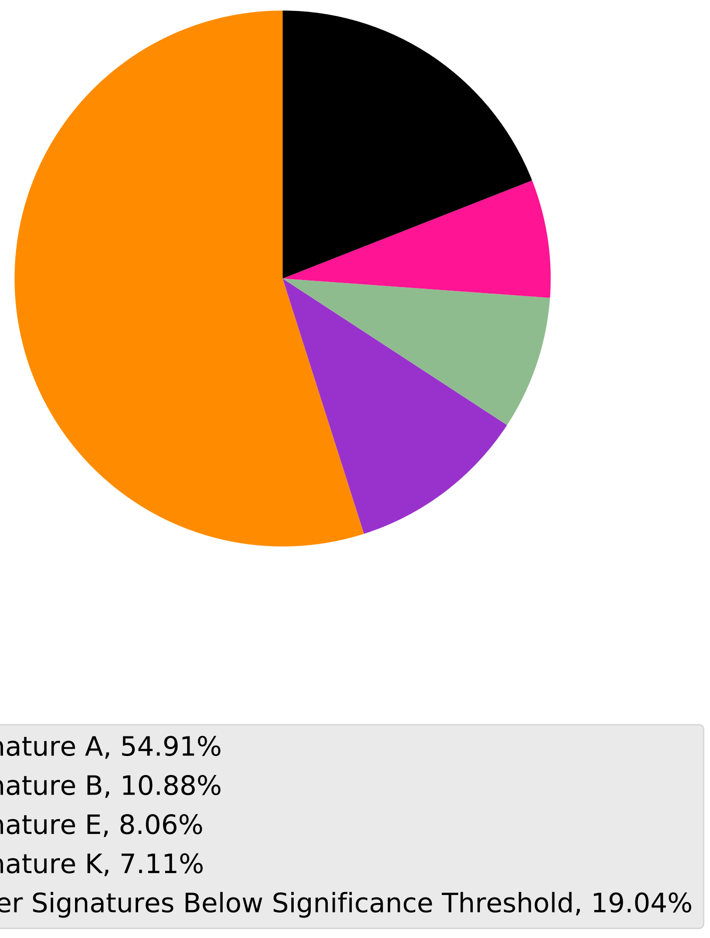

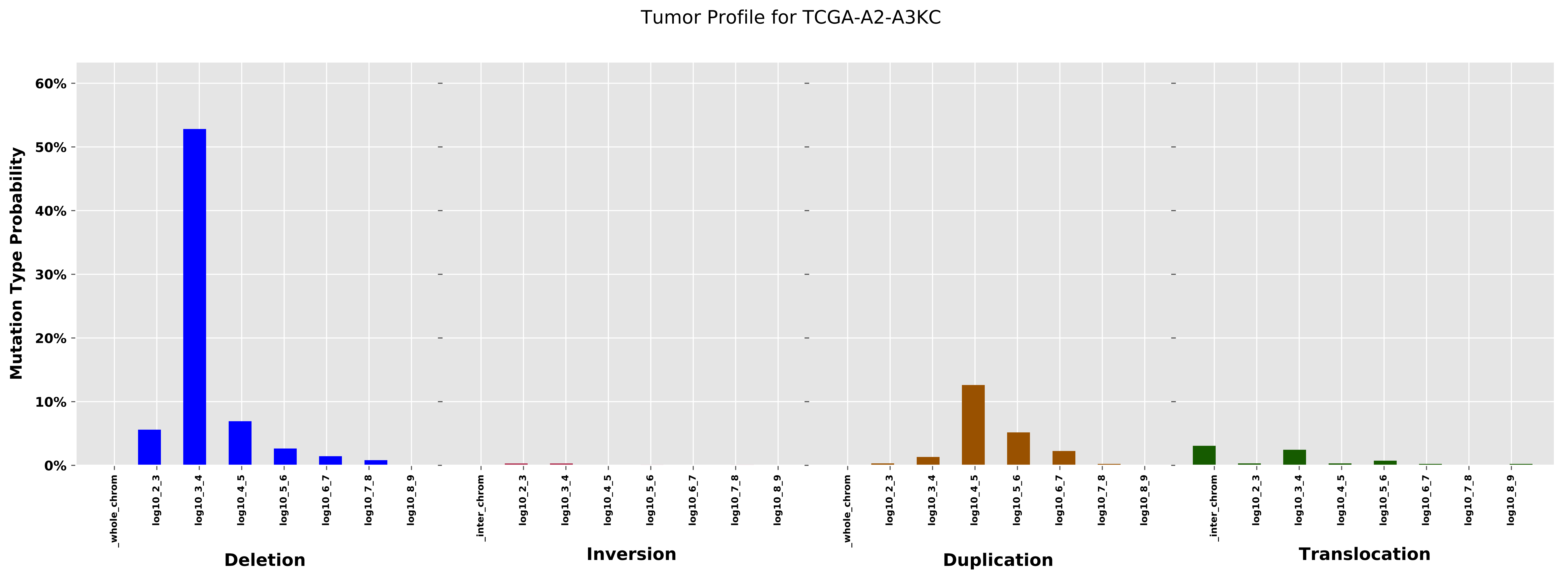

Tumor Profile for TCGA-A2-A3KC
Mutation Type Probability
60%
50%
40%
30%
20%
10%
0%
_whole_chrom
log10_2_3
log10_3_4
log10_4_5
log10_5_6
log10_6_7
log10_7_8
log10_8_9
_inter_chrom
Deletion
Inversion
Duplication
Translocation

Cancer processes Weights for TCGA-AA-3514

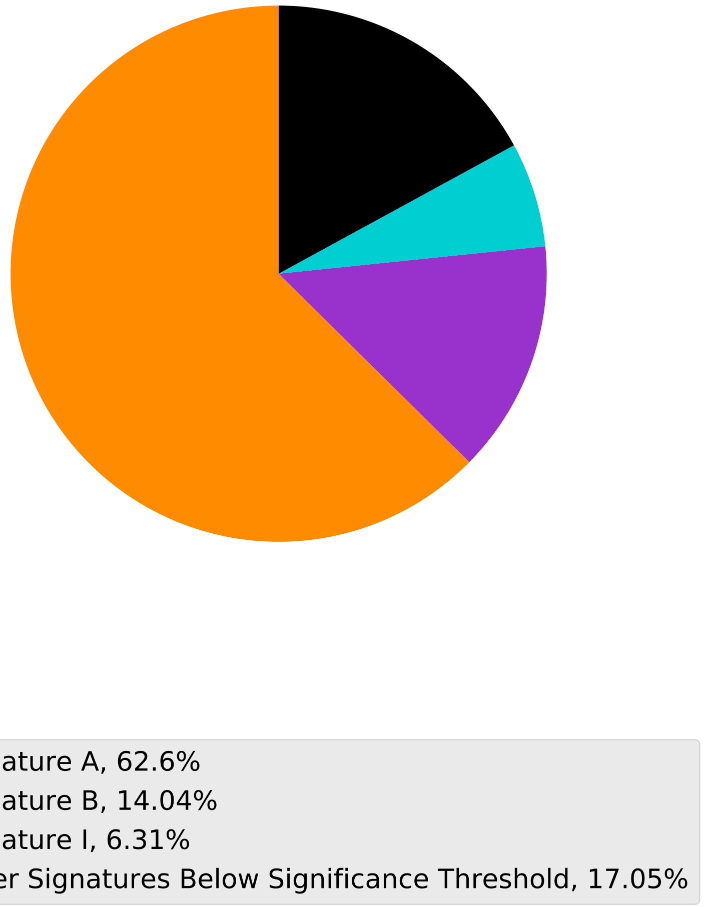

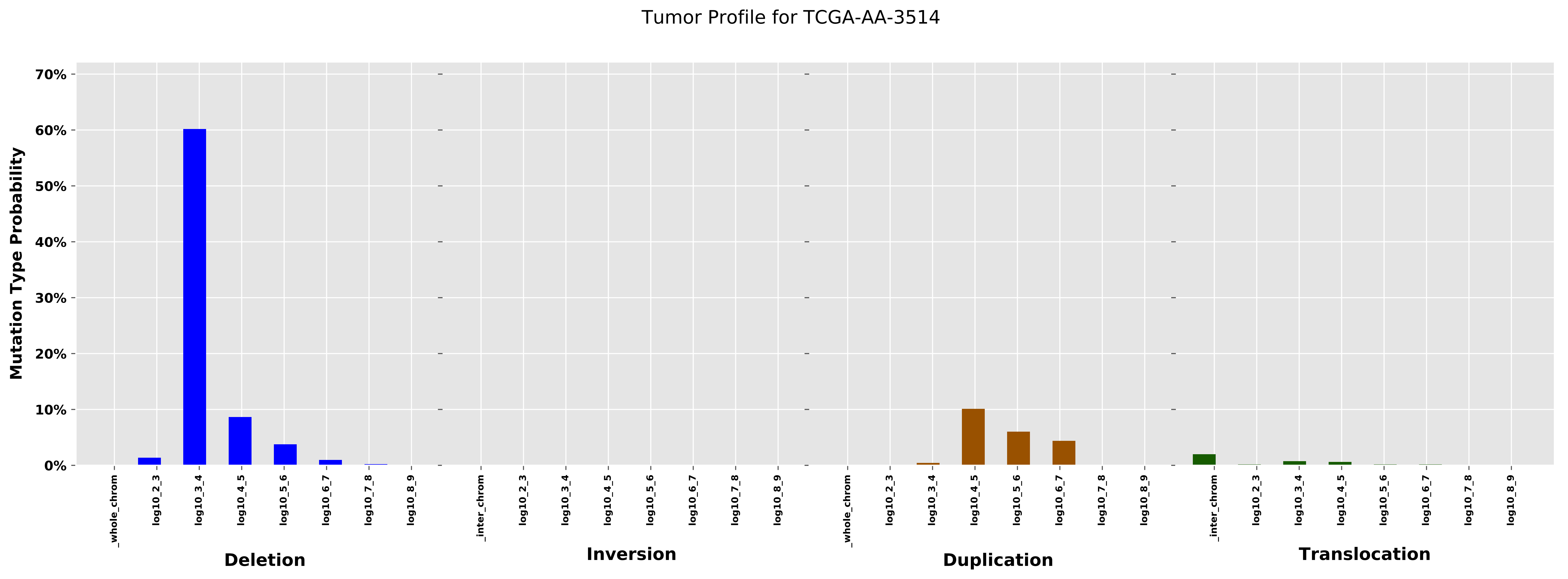
Tumor Profile for TCGA-AA-3514
Mutation Type Probability
70%
60%
50%
40%
30%
20%
10%
0%
_whole_chrom
log10_2_3
log10_3_4
log10_4_5
log10_5_6
log10_6_7
log10_7_8
log10_8_9
_inter_chrom
Deletion
Inversion
Duplication
Translocation

#### Cancer processes Weights for TCGA-AN-A0XR

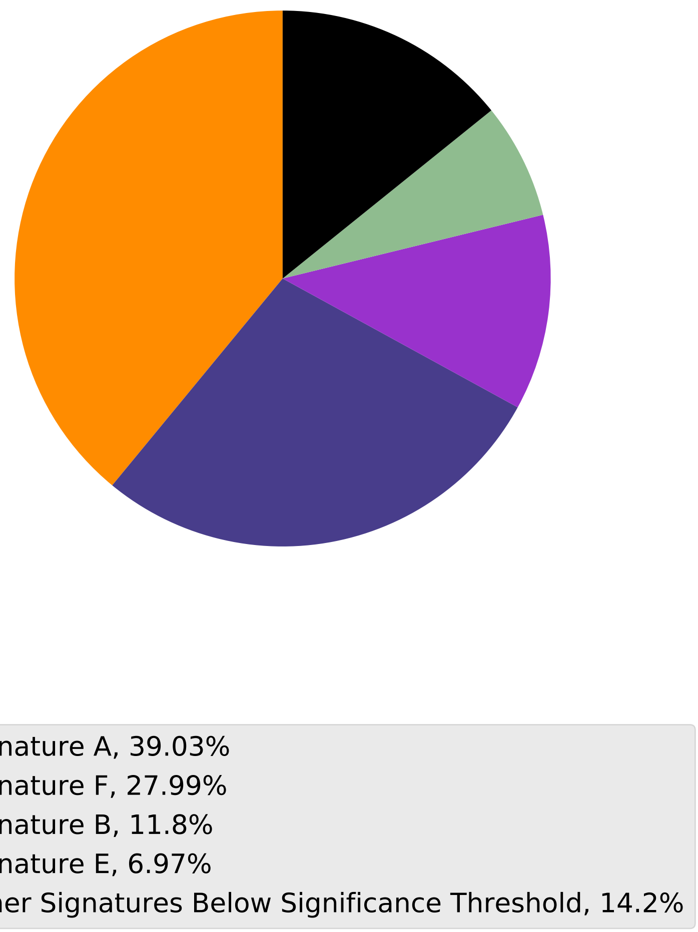

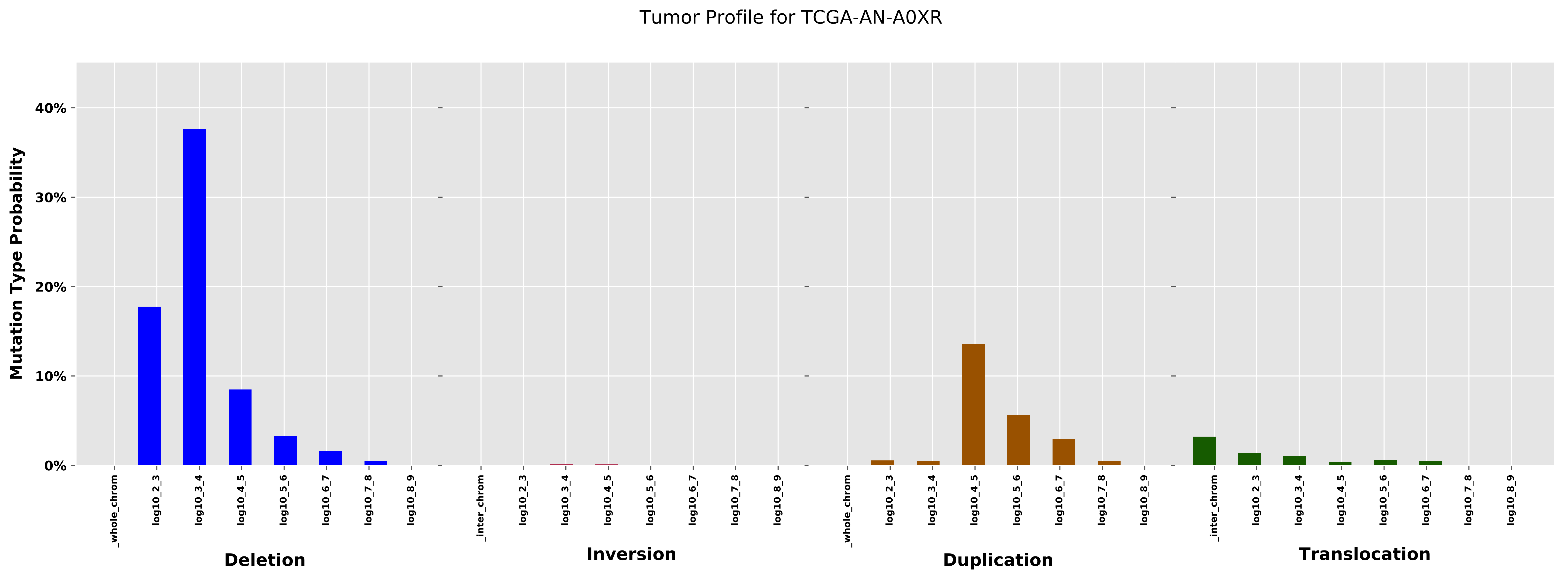
Tumor Profile for TCGA-AN-A0XR
Mutation Type Probability
40%
30%
20%
10%
0%
_whole_chrom
log10_2_3
log10_3_4
log10_4_5
log10_5_6
log10_6_7
log10_7_8
log10_8_9
Deletion
_inter_chrom
log10_2_3
log10_3_4
log10_4_5
log10_5_6
log10_6_7
log10_7_8
log10_8_9
Inversion
_whole_chrom
log10_2_3
log10_3_4
log10_4_5
log10_5_6
log10_6_7
log10_7_8
log10_8_9
Duplication
_inter_chrom
log10_2_3
log10_3_4
log10_4_5
log10_5_6
log10_6_7
log10_7_8
log10_8_9
Translocation

Cancer processes Weights for TCGA-BH-A0EA

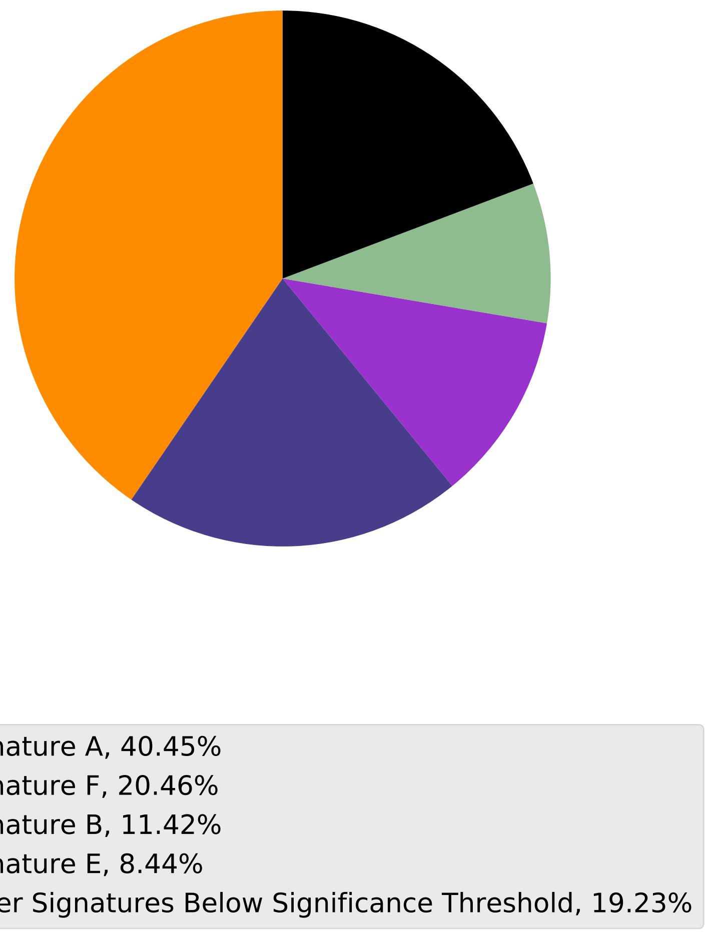

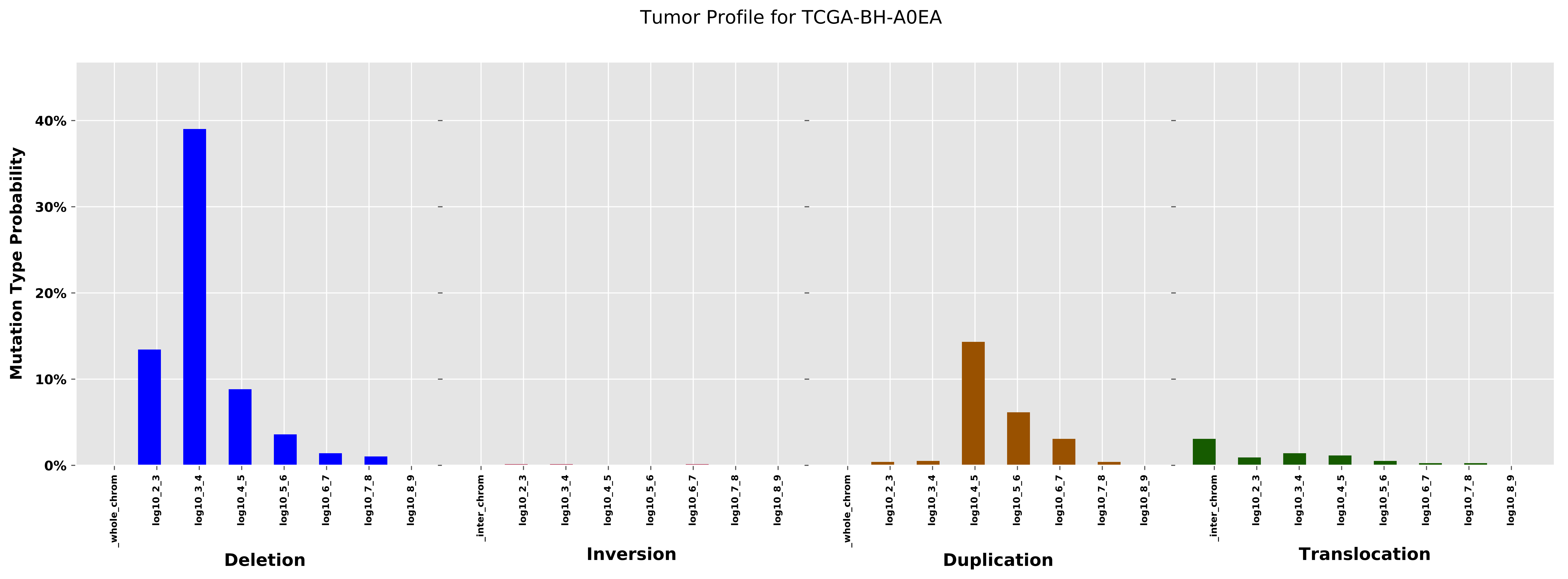

Tumor Profile for TCGA-BH-A0EA
Mutation Type Probability
40%
30%
20%
10%
0%
_whole_chrom
log10_2_3
log10_3_4
log10_4_5
log10_5_6
log10_6_7
log10_7_8
log10_8_9
_inter_chrom
Deletion
Inversion
Duplication
Translocation

Cancer processes Weights for TCGA-AR-A256

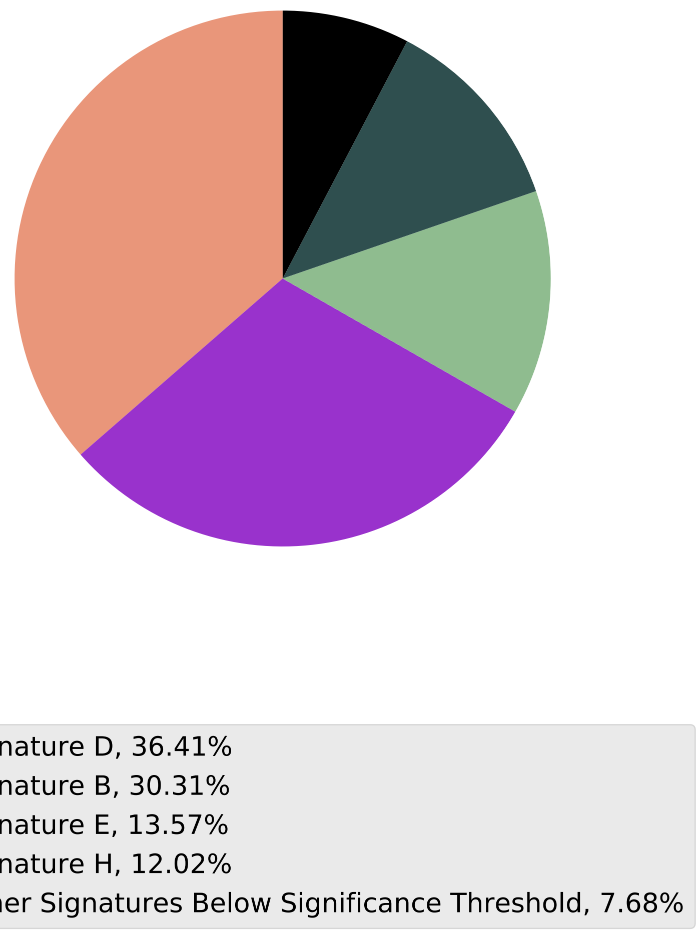

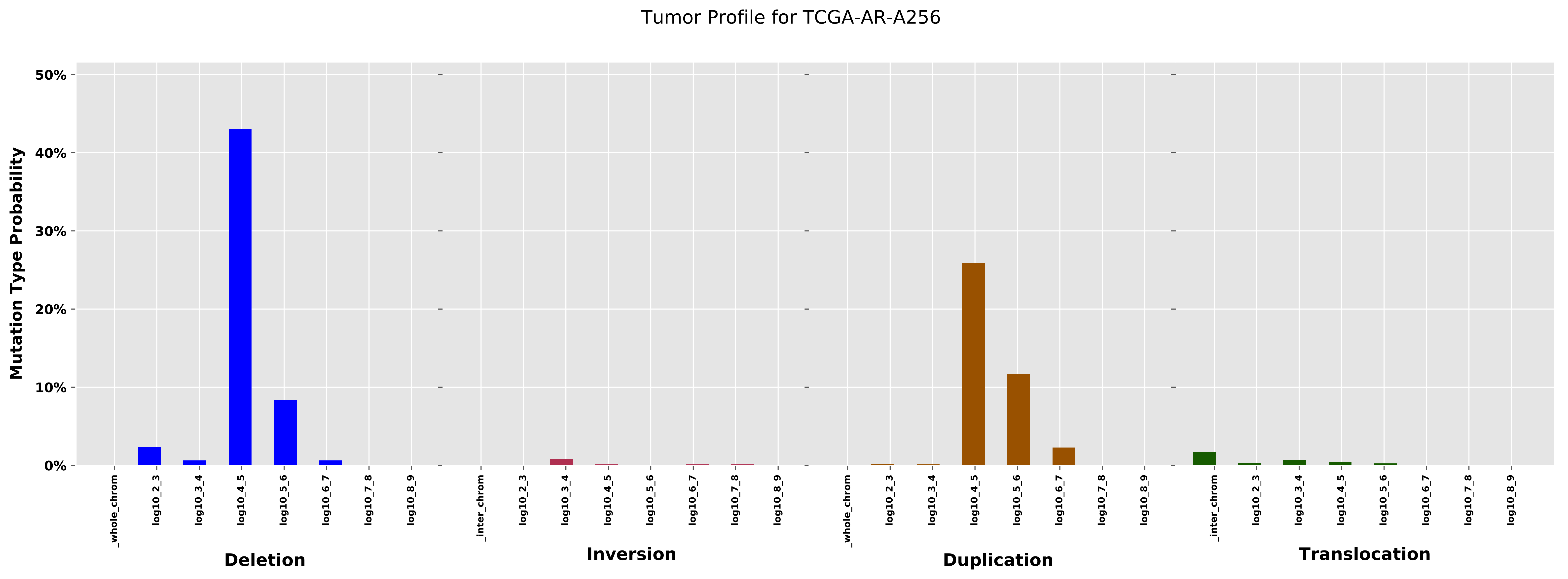

Tumor Profile for TCGA-AR-A256
Mutation Type Probability
50%
40%
30%
20%
10%
0%
_whole_chrom
log10_2_3
log10_3_4
log10_4_5
log10_5_6
log10_6_7
log10_7_8
log10_8_9
Deletion
_inter_chrom
log10_2_3
log10_3_4
log10_4_5
log10_5_6
log10_6_7
log10_7_8
log10_8_9
Inversion
_whole_chrom
log10_2_3
log10_3_4
log10_4_5
log10_5_6
log10_6_7
log10_7_8
log10_8_9
Duplication
_inter_chrom
log10_2_3
log10_3_4
log10_4_5
log10_5_6
log10_6_7
log10_7_8
log10_8_9
Translocation

Cancer processes Weights for TCGA-A8-A09X

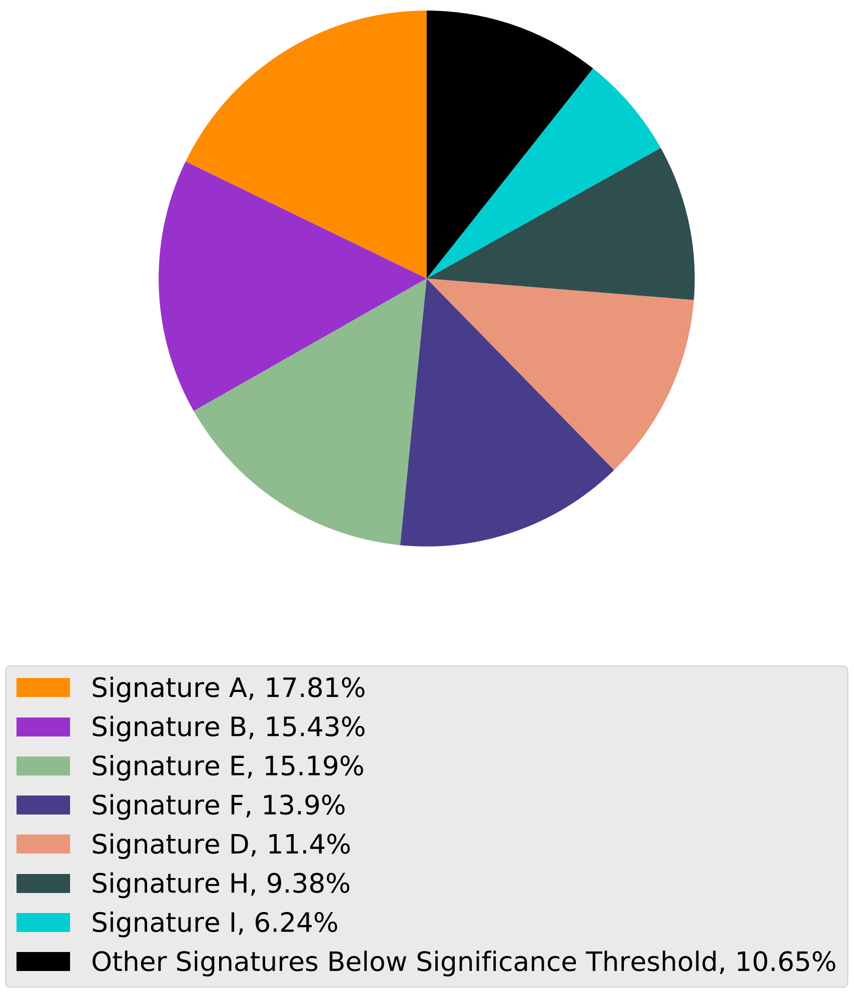

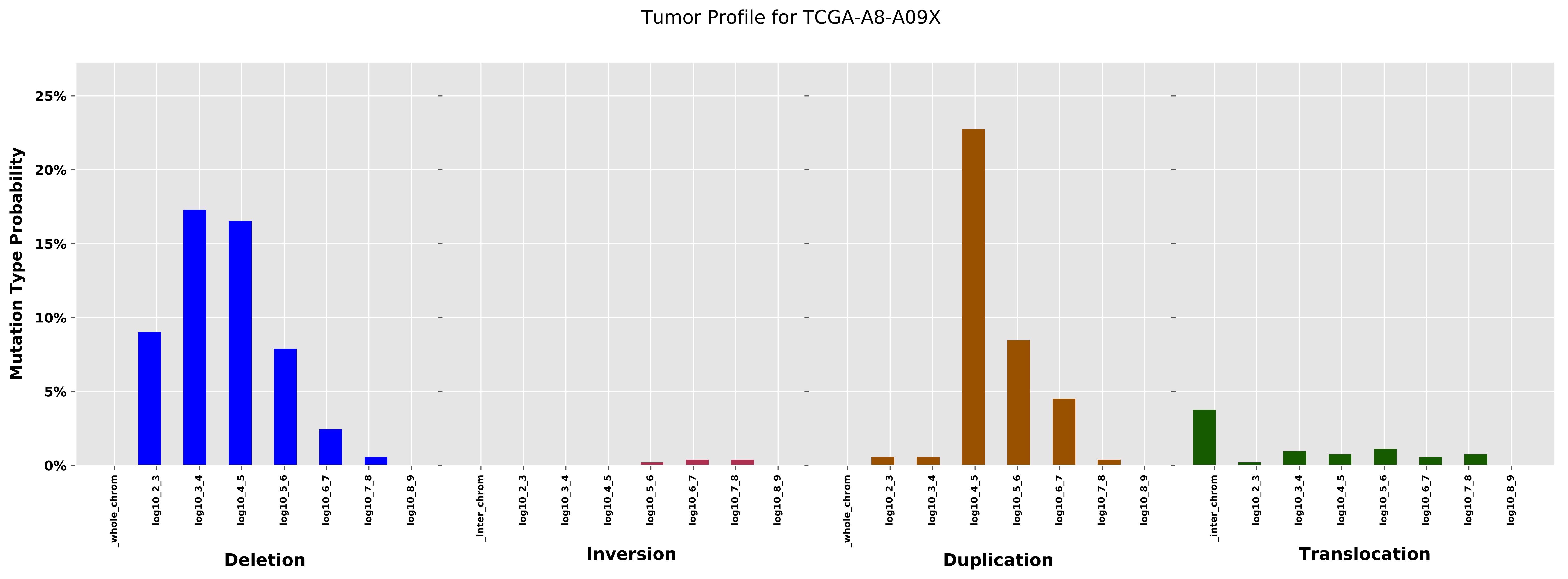
Tumor Profile for TCGA-A8-A09X
Mutation Type Probability
25%
20%
15%
10%
5%
0%
_whole_chrom
log10_2_3
log10_3_4
log10_4_5
log10_5_6
log10_6_7
log10_7_8
log10_8_9
Deletion
_inter_chrom
Inversion
Duplication
Translocation

Cancer processes Weights for TCGA-AA-A01V

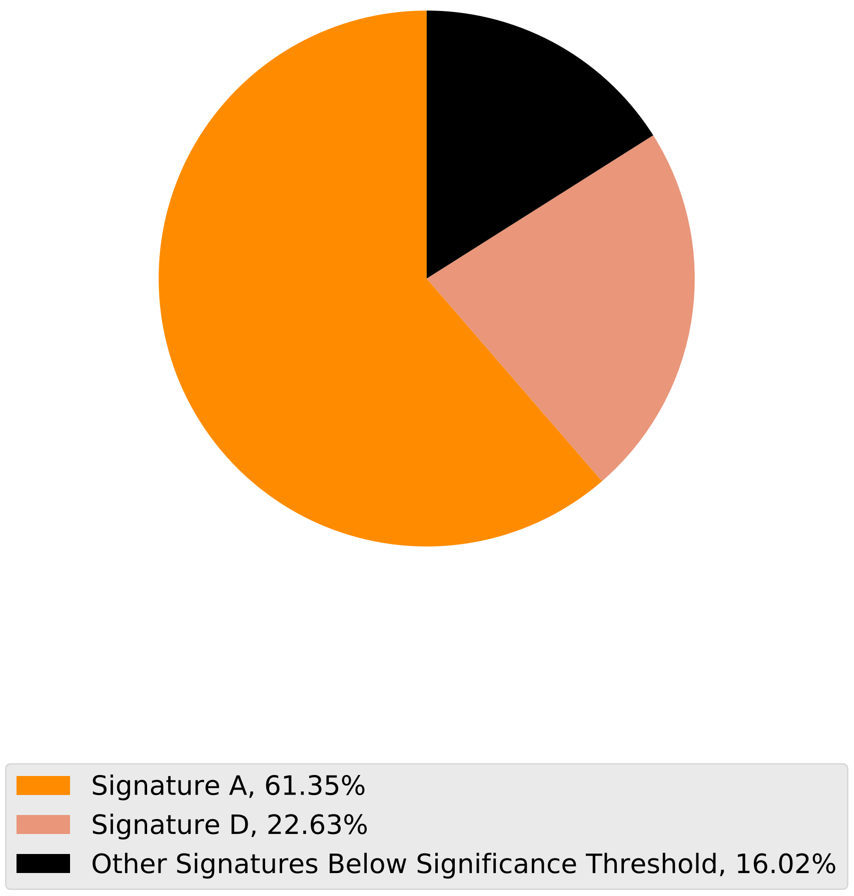

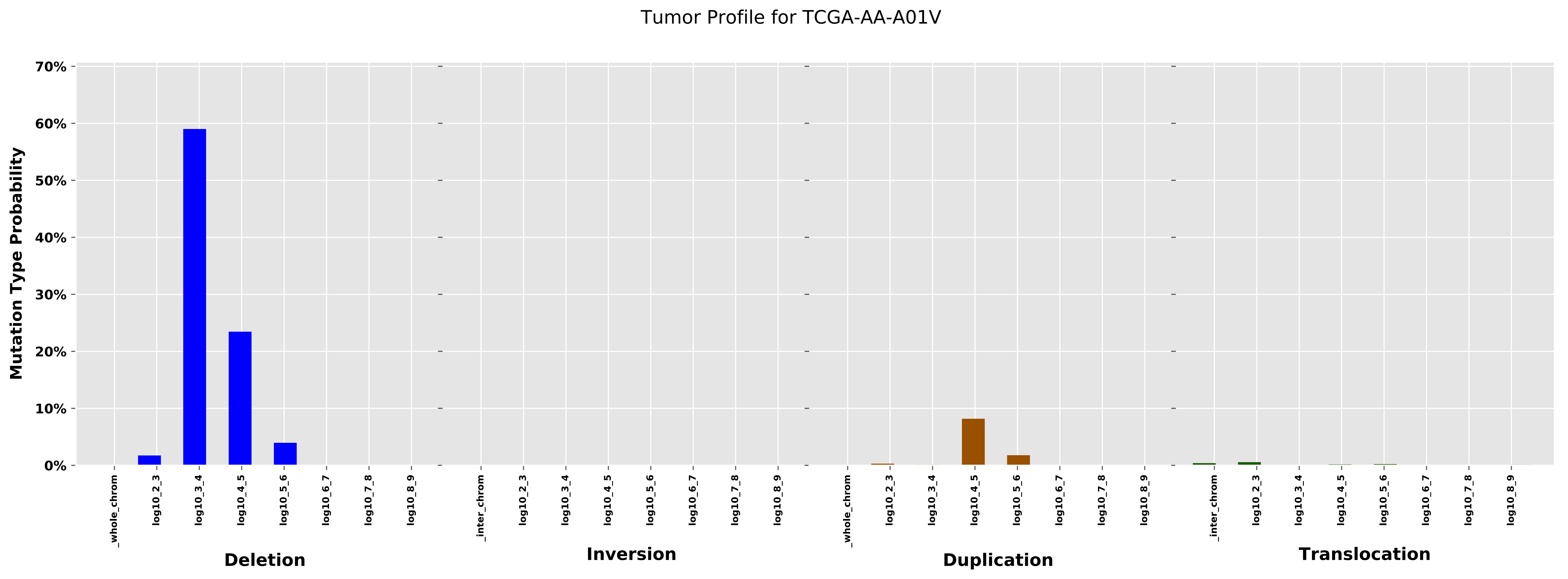
Tumor Profile for TCGA-AA-A01V
Mutation Type Probability
70%
60%
50%
40%
30%
20%
10%
0%
_whole_chrom
log10_2_3
log10_3_4
log10_4_5
log10_5_6
log10_6_7
log10_7_8
log10_8_9
Deletion
_inter_chrom
log10_2_3
log10_3_4
log10_4_5
log10_5_6
log10_6_7
log10_7_8
log10_8_9
Inversion
_whole_chrom
log10_2_3
log10_3_4
log10_4_5
log10_5_6
log10_6_7
log10_7_8
log10_8_9
Duplication
_inter_chrom
log10_2_3
log10_3_4
log10_4_5
log10_5_6
log10_6_7
log10_7_8
log10_8_9
Translocation

Cancer processes Weights for TCGA-A2-A04T

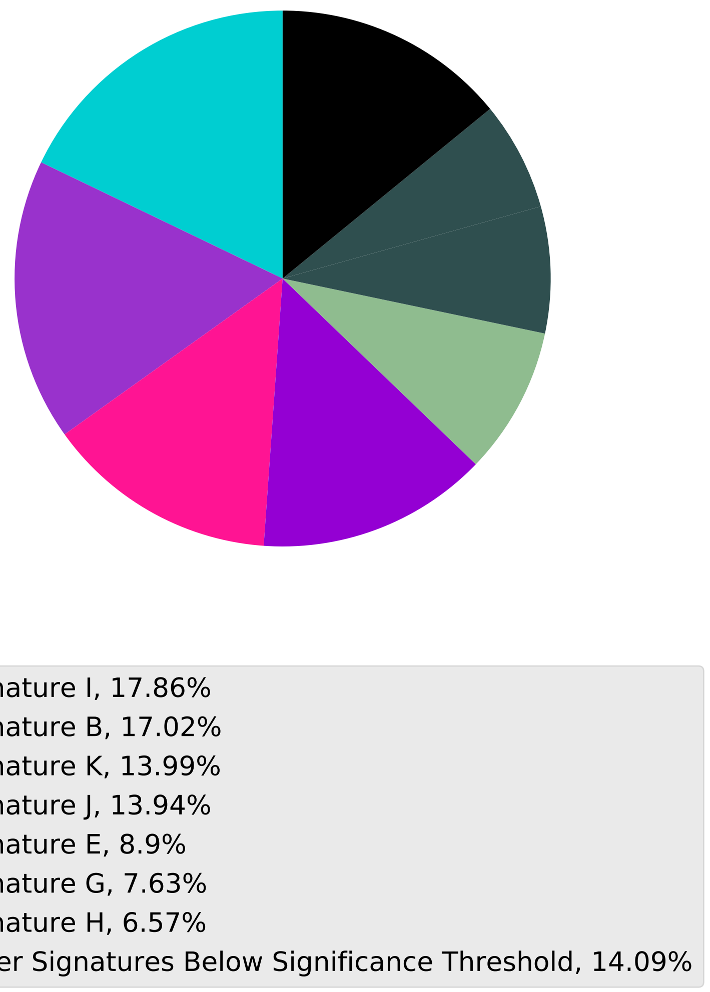

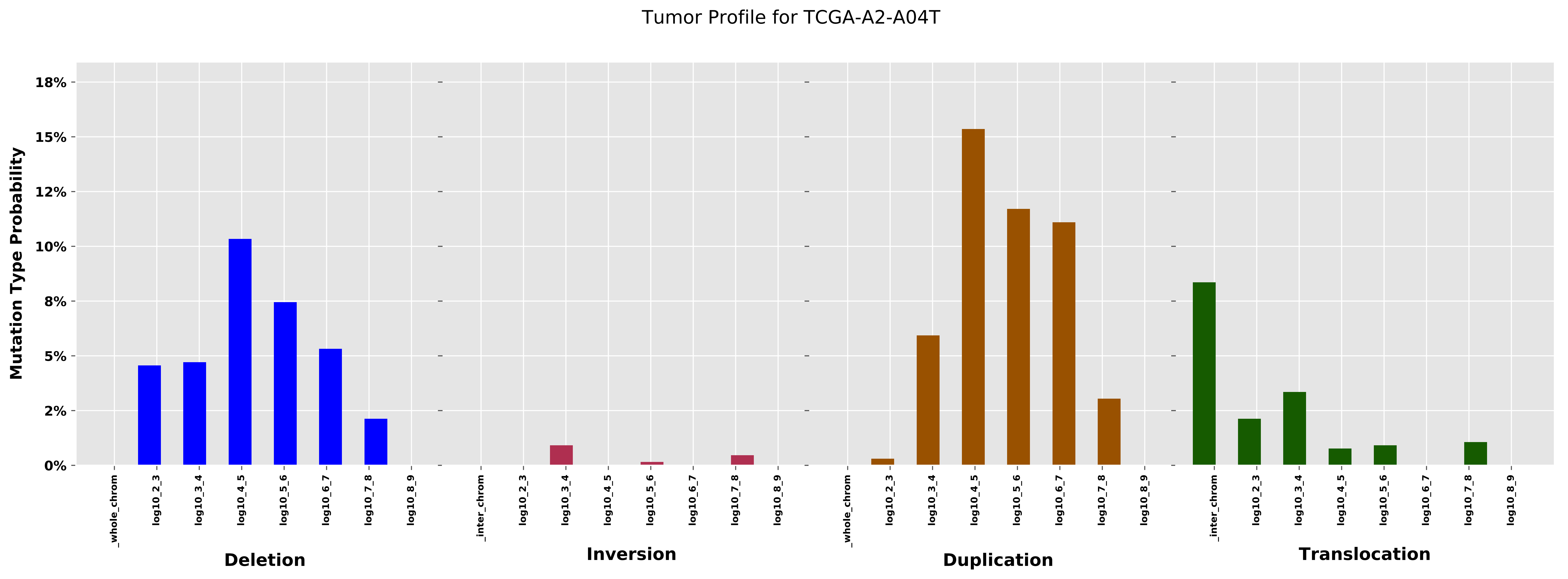

Tumor Profile for TCGA-A2-A04T
Mutation Type Probability
18%
15%
12%
10%
8%
5%
2%
0%
_whole_chrom
log10_2_3
log10_3_4
log10_4_5
log10_5_6
log10_6_7
log10_7_8
log10_8_9
_inter_chrom
Deletion
Inversion
Duplication
Translocation

Cancer processes Weights for TCGA-BH-A0H6

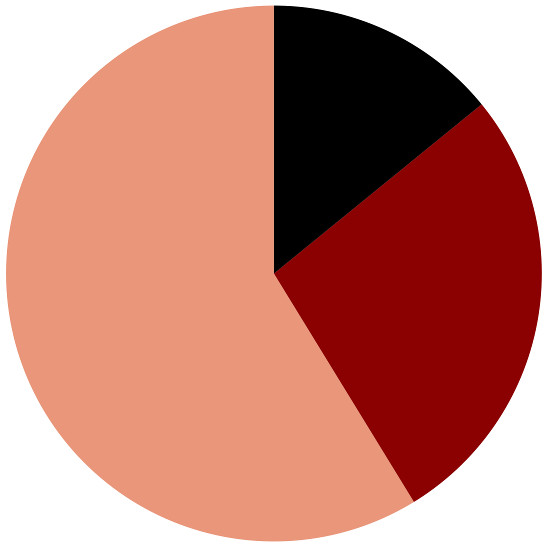

Signature D, 58.75%
Signature C, 27.12%
Other Signatures Below Significance Threshold, 14.12%

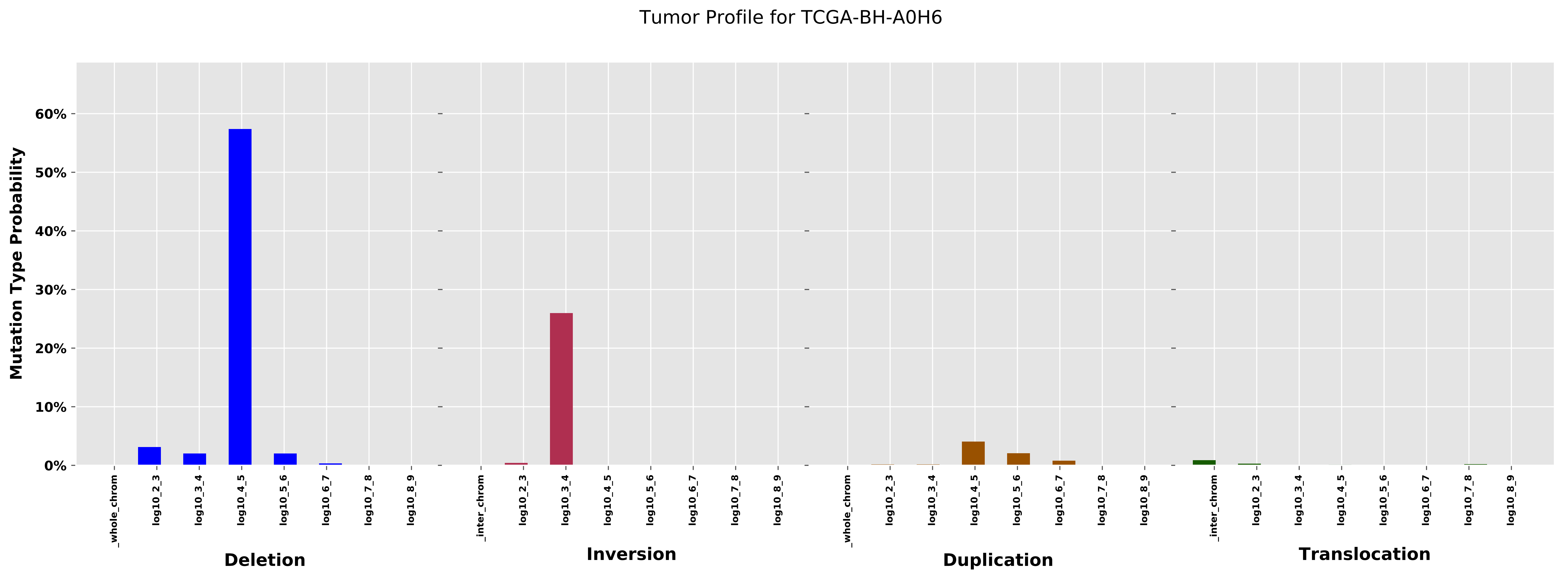

Tumor Profile for TCGA-BH-A0H6
Mutation Type Probability
60%
50%
40%
30%
20%
10%
0%
_whole_chrom
log10_2_3
log10_3_4
log10_4_5
log10_5_6
log10_6_7
log10_7_8
log10_8_9
_inter_chrom
Deletion
Inversion
Duplication
Translocation

Cancer processes Weights for TCGA-D8-A27F

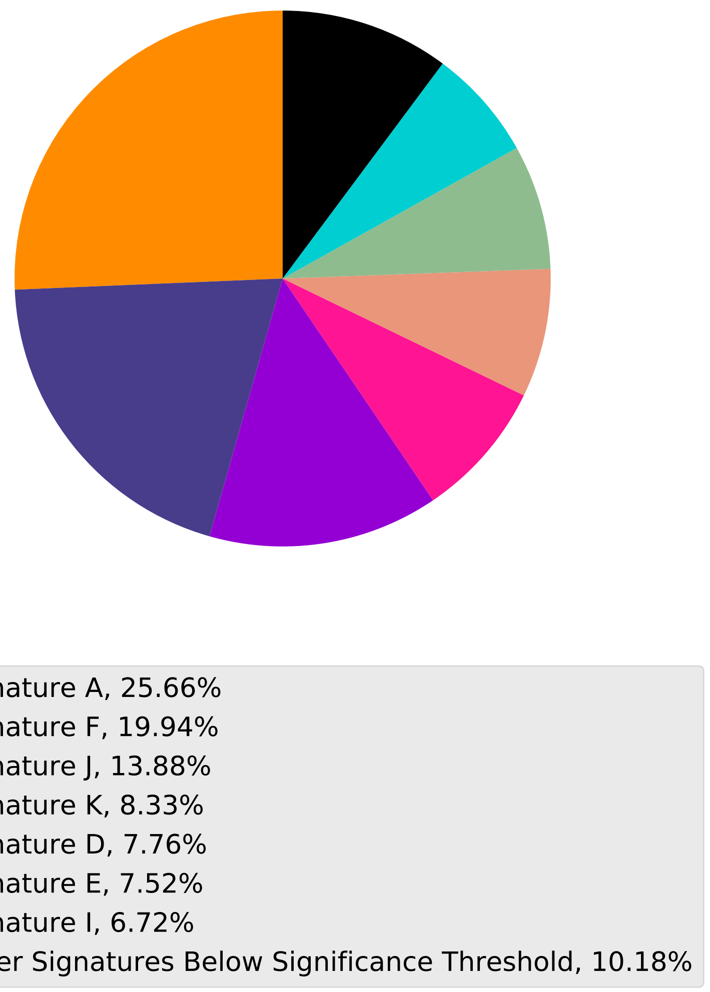

Tumor Profile for TCGA-D8-A27F

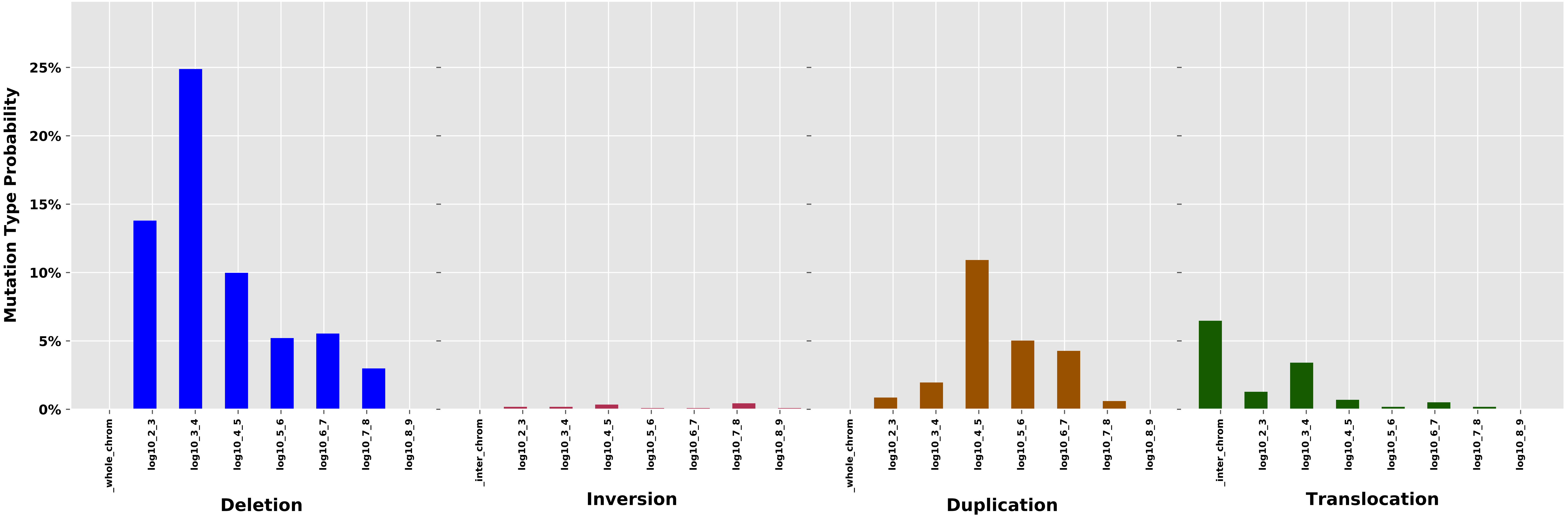

Cancer processes Weights for TCGA-AG-3727

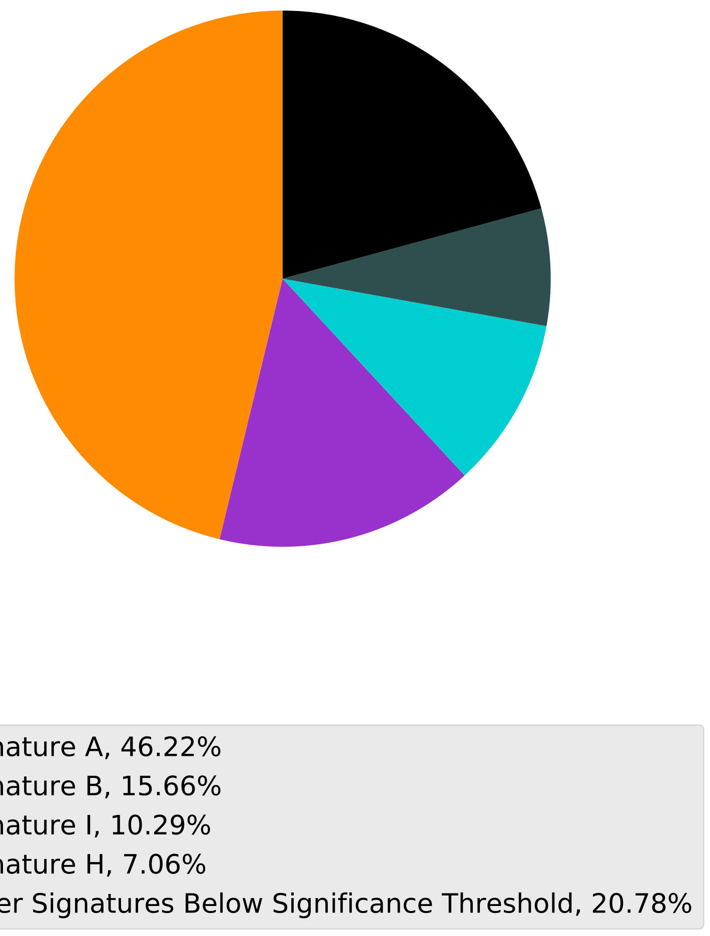

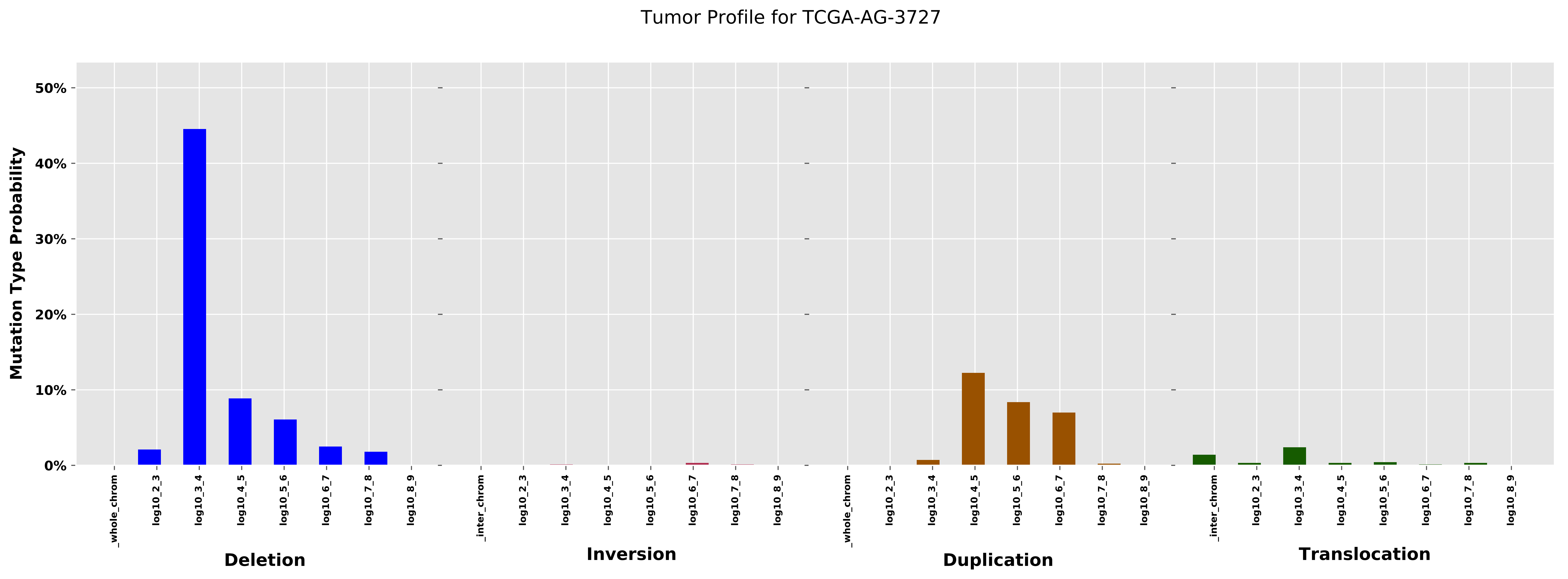

Tumor Profile for TCGA-AG-3727
Mutation Type Probability
50%
40%
30%
20%
10%
0%
_whole_chrom
log10_2_3
log10_3_4
log10_4_5
log10_5_6
log10_6_7
log10_7_8
log10_8_9
Deletion
_inter_chrom
log10_2_3
log10_3_4
log10_4_5
log10_5_6
log10_6_7
log10_7_8
log10_8_9
Inversion
_whole_chrom
log10_2_3
log10_3_4
log10_4_5
log10_5_6
log10_6_7
log10_7_8
log10_8_9
Duplication
_inter_chrom
log10_2_3
log10_3_4
log10_4_5
log10_5_6
log10_6_7
log10_7_8
log10_8_9
Translocation

Cancer processes Weights for TCGA-A8-A09I

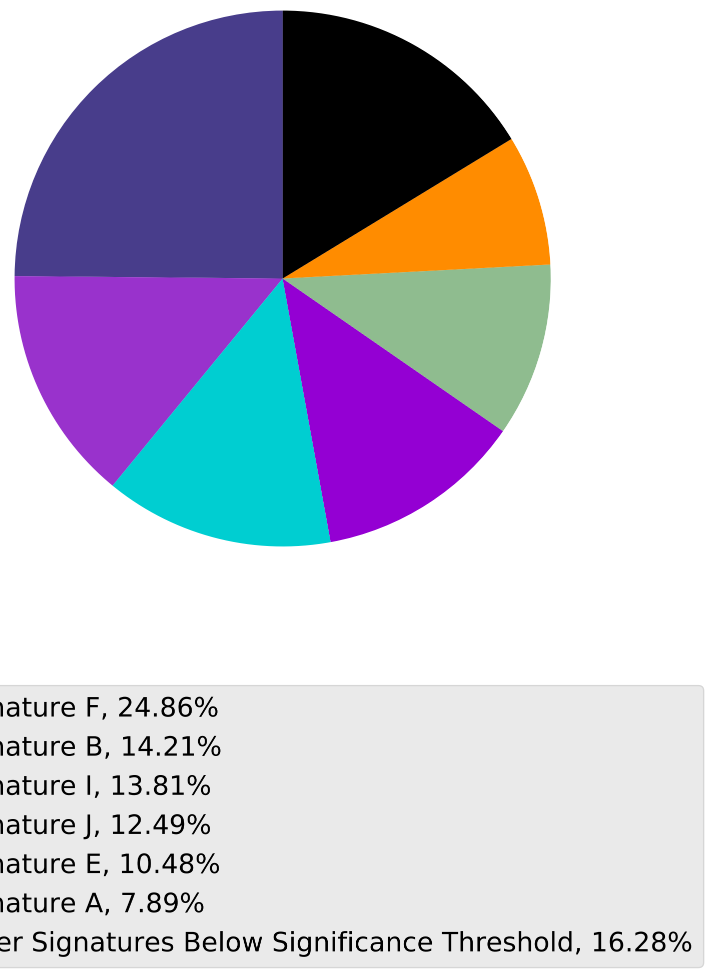

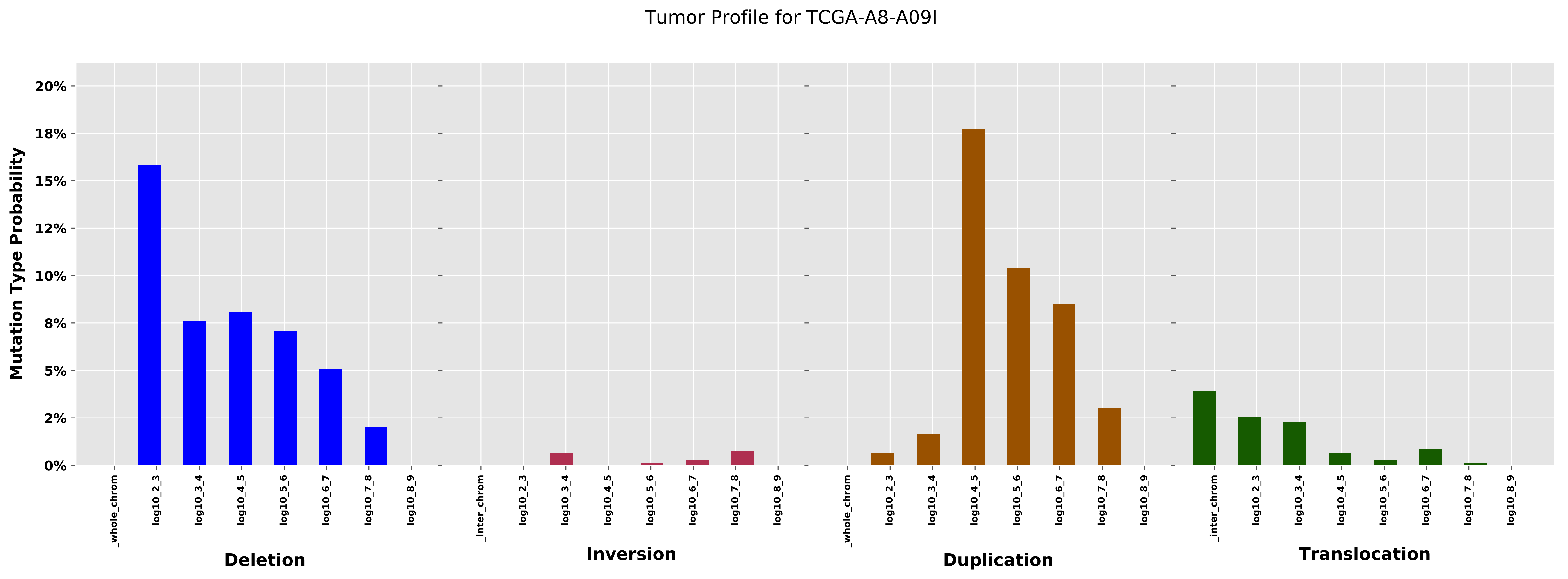

Tumor Profile for TCGA-A8-A09I
Mutation Type Probability
20%
18%
15%
12%
10%
8%
5%
2%
0%
_whole_chrom
log10_2_3
log10_3_4
log10_4_5
log10_5_6
log10_6_7
log10_7_8
log10_8_9
_inter_chrom
Deletion
Inversion
Duplication
Translocation

Cancer processes Weights for TCGA-AA-3664

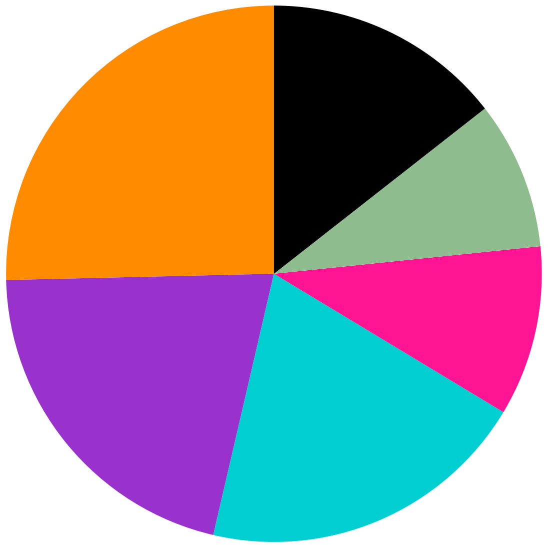

Signature A, 25.37%
Signature B, 20.99%
Signature I, 20.01%
Signature K, 10.27%
Signature E, 8.93%
Other Signatures Below Significance Threshold, 14.43%

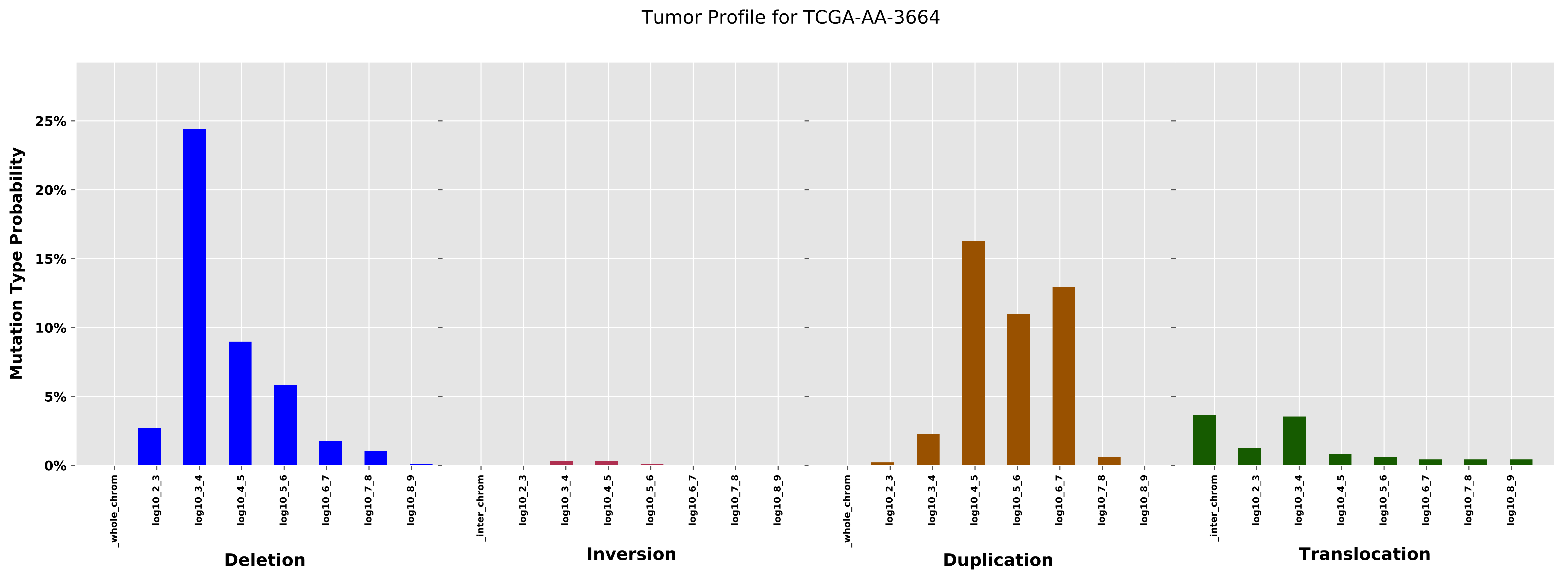

Tumor Profile for TCGA-AA-3664
Mutation Type Probability
25%
20%
15%
10%
5%
0%
_whole_chrom
log10_2_3
log10_3_4
log10_4_5
log10_5_6
log10_6_7
log10_7_8
log10_8_9
_inter_chrom
Deletion
Inversion
Duplication
Translocation

Cancer processes Weights for TCGA-E2-A1LL

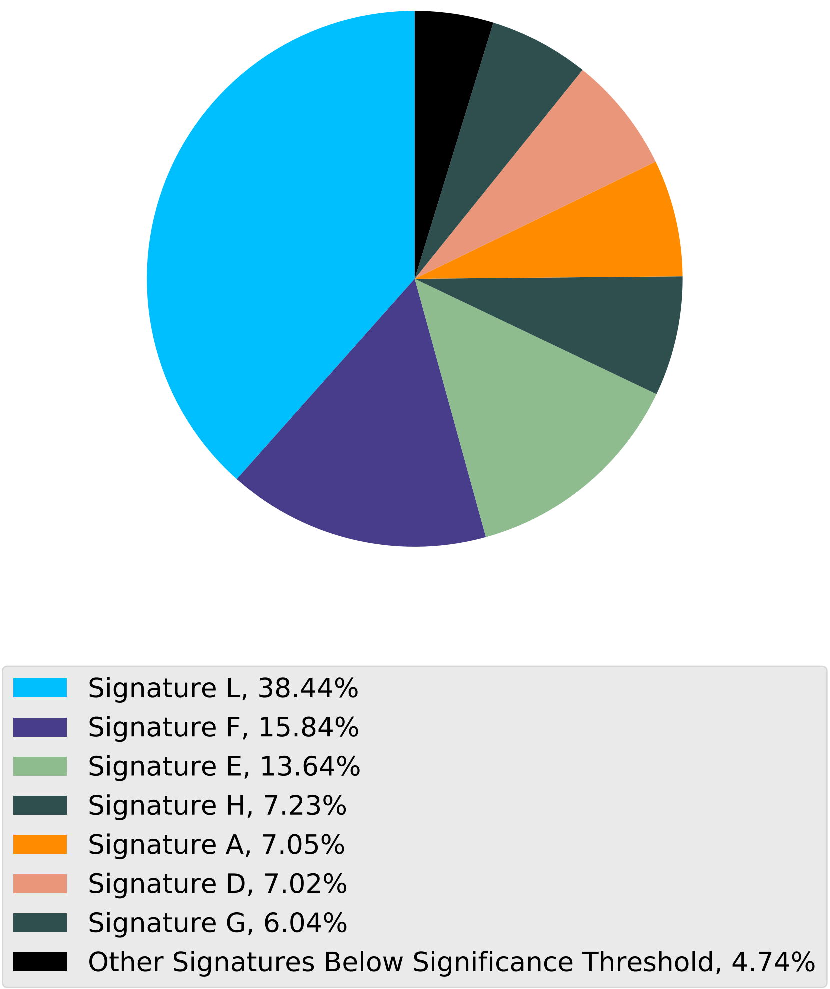

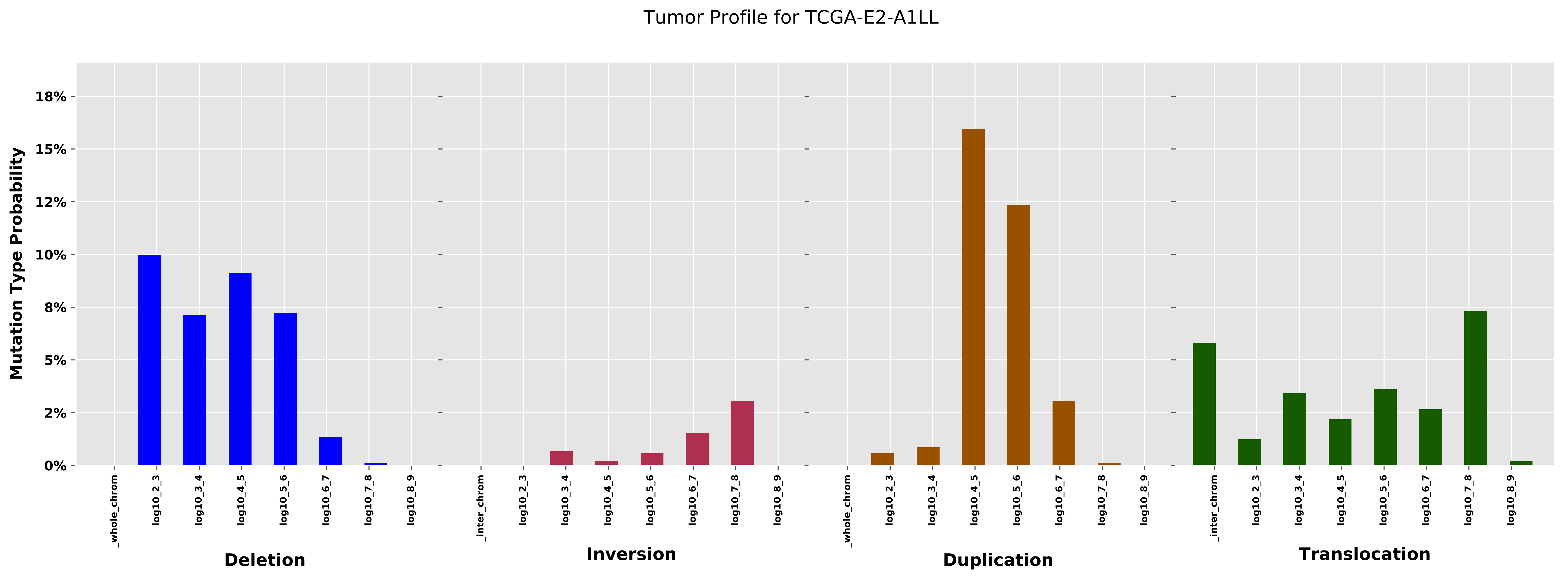

Tumor Profile for TCGA-E2-A1LL
Mutation Type Probability
18%
15%
12%
10%
8%
5%
2%
0%
_whole_chrom
log10_2_3
log10_3_4
log10_4_5
log10_5_6
log10_6_7
log10_7_8
log10_8_9
_inter_chrom
Deletion
Inversion
Duplication
Translocation

Cancer processes Weights for TCGA-A1-A0SM

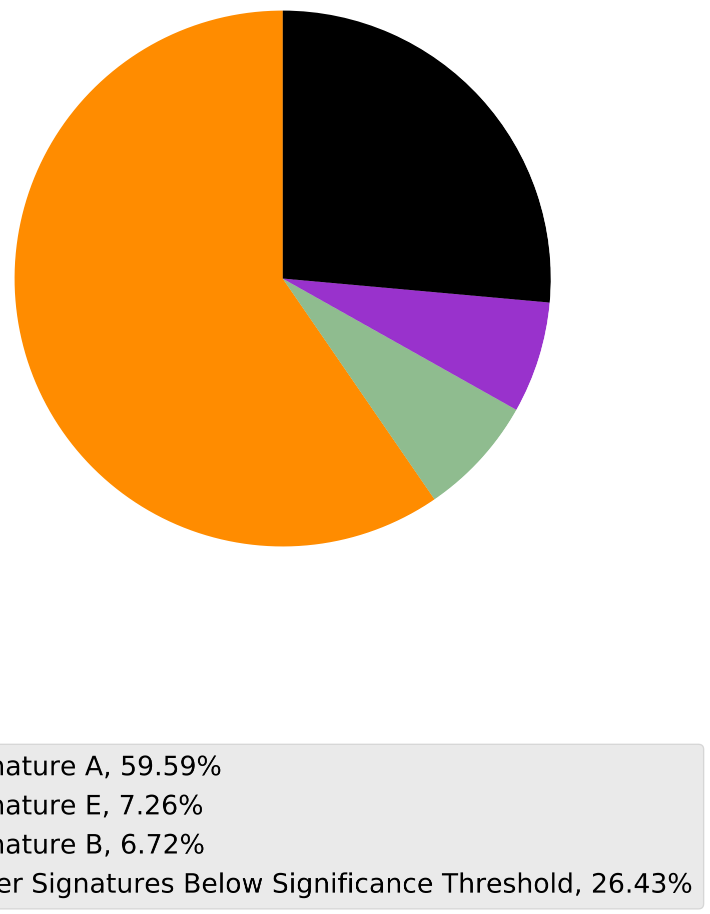

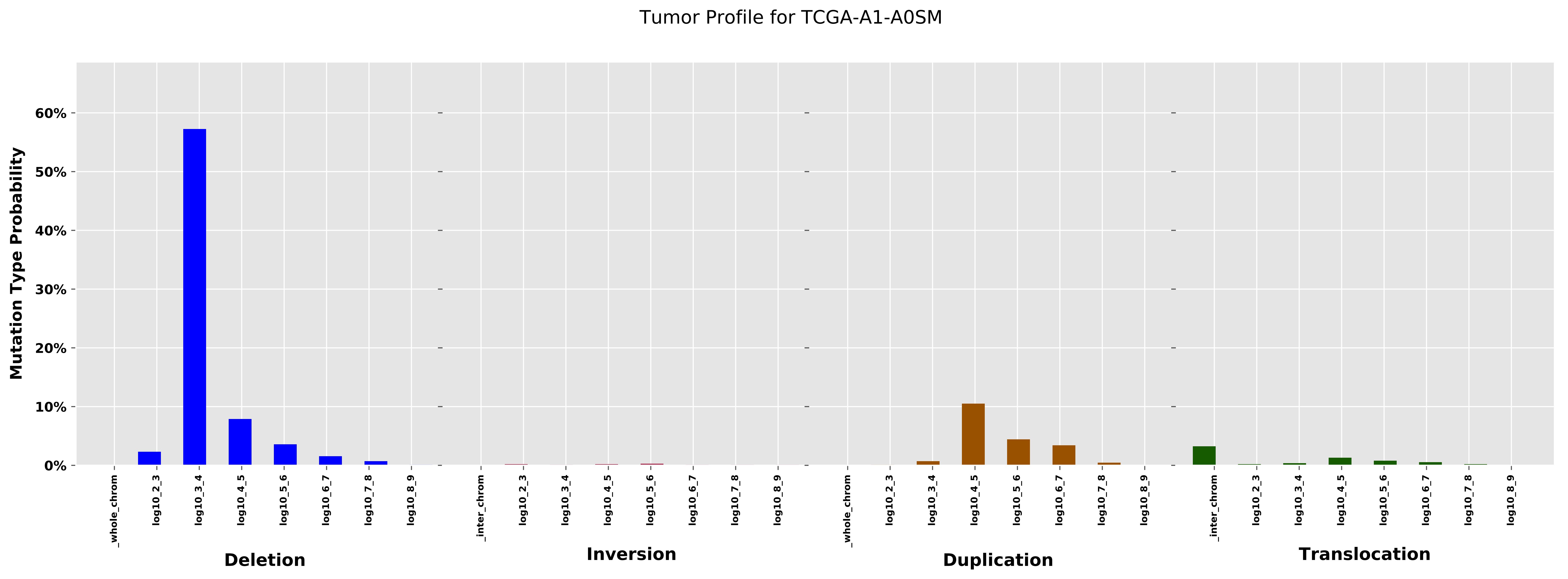
Tumor Profile for TCGA-A1-A0SM
Mutation Type Probability
60%
50%
40%
30%
20%
10%
0%
_whole_chrom
log10_2_3
log10_3_4
log10_4_5
log10_5_6
log10_6_7
log10_7_8
log10_8_9
_inter_chrom
Deletion
Inversion
Duplication
Translocation

Cancer processes Weights for TCGA-C8-A130

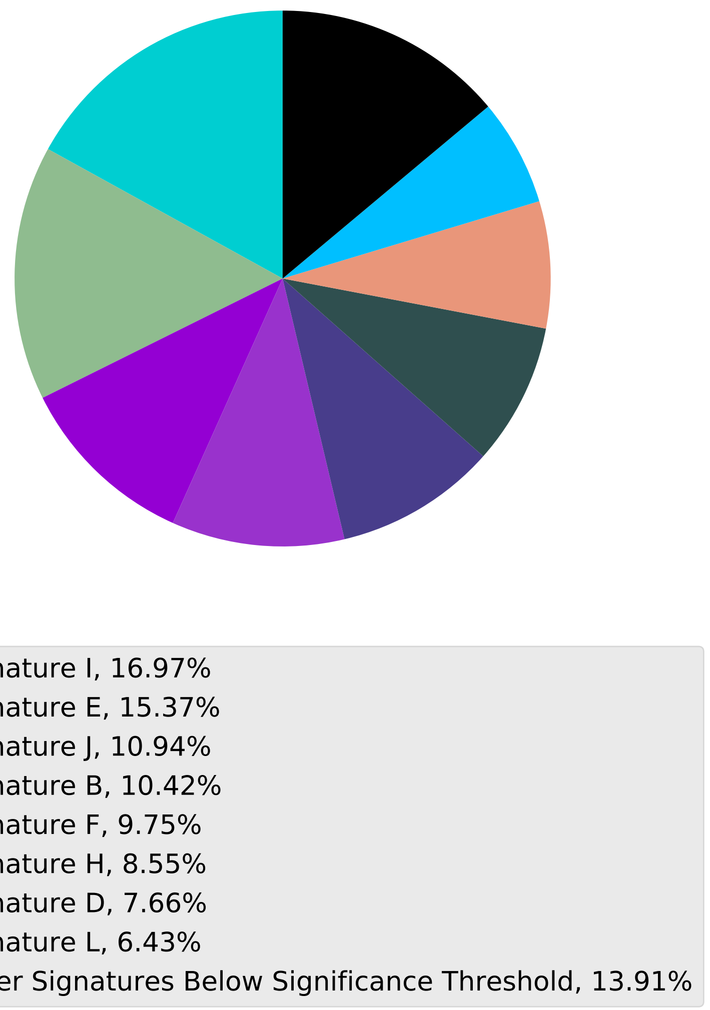

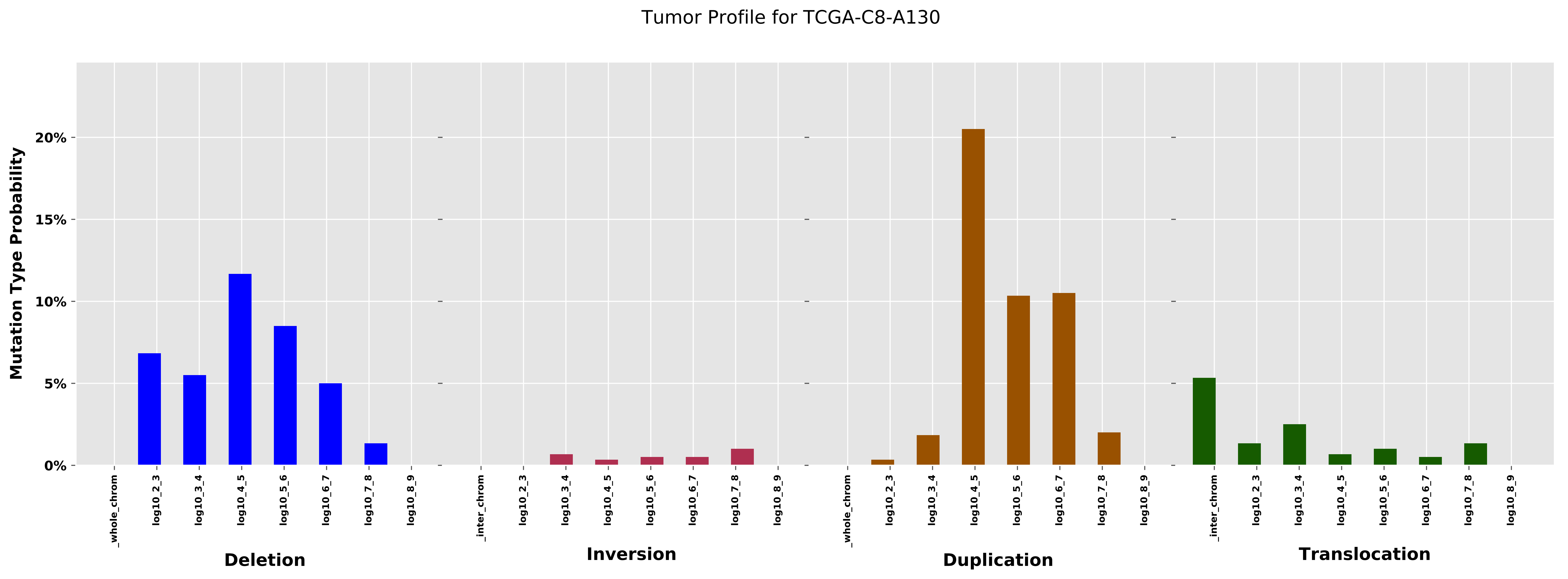

Tumor Profile for TCGA-C8-A130
Mutation Type Probability
20%
15%
10%
5%
0%
_whole_chrom
log10_2_3
log10_3_4
log10_4_5
log10_5_6
log10_6_7
log10_7_8
log10_8_9
_inter_chrom
Deletion
Inversion
Duplication
Translocation

Cancer processes Weights for TCGA-BH-A0H0

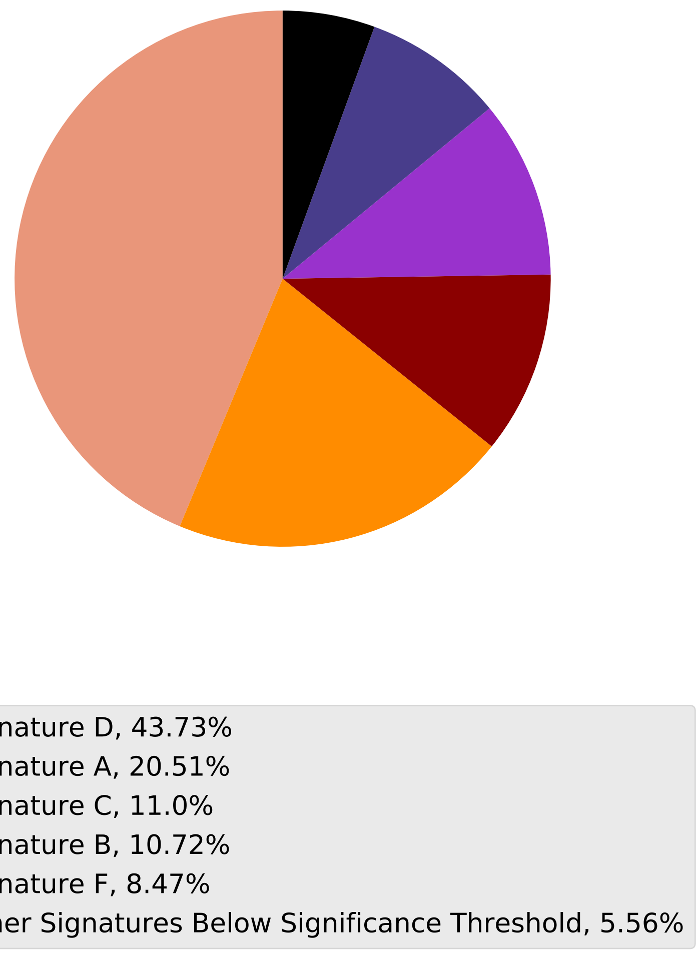

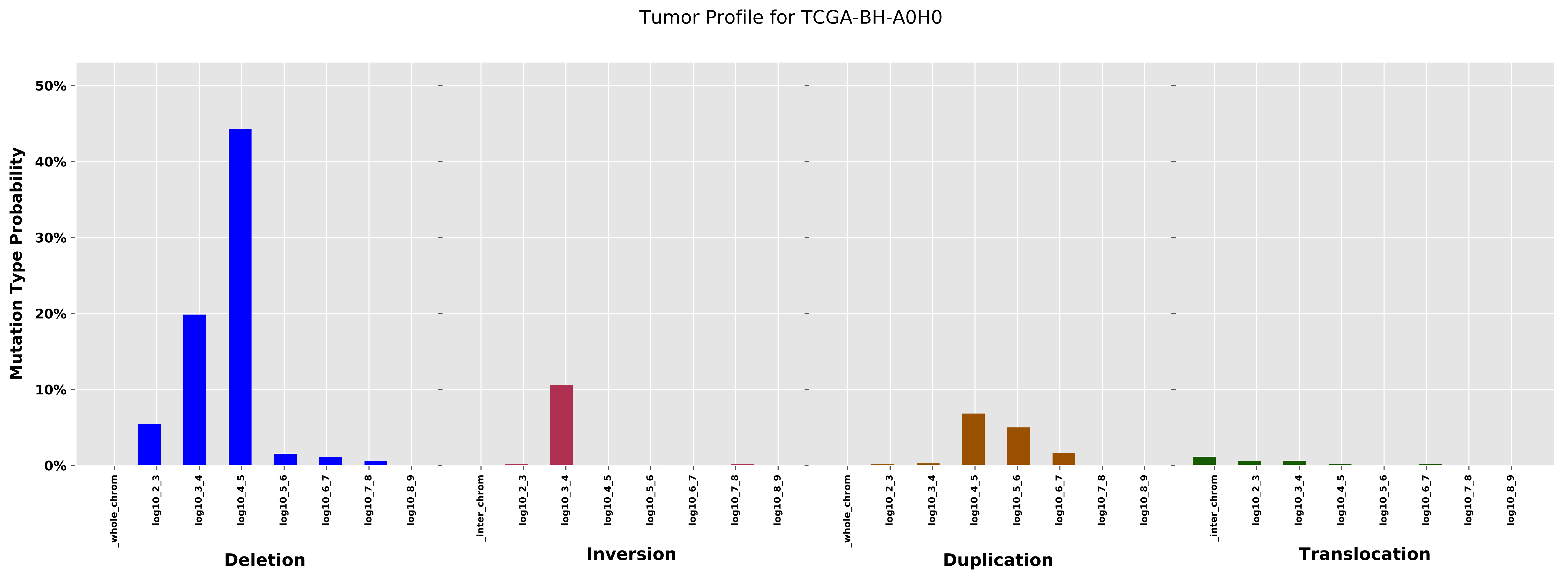

Tumor Profile for TCGA-BH-A0H0
Mutation Type Probability
50%
40%
30%
20%
10%
0%
_whole_chrom
log10_2_3
log10_3_4
log10_4_5
log10_5_6
log10_6_7
log10_7_8
log10_8_9
_inter_chrom
Deletion
Inversion
Duplication
Translocation

Cancer processes Weights for TCGA-EW-A1J5

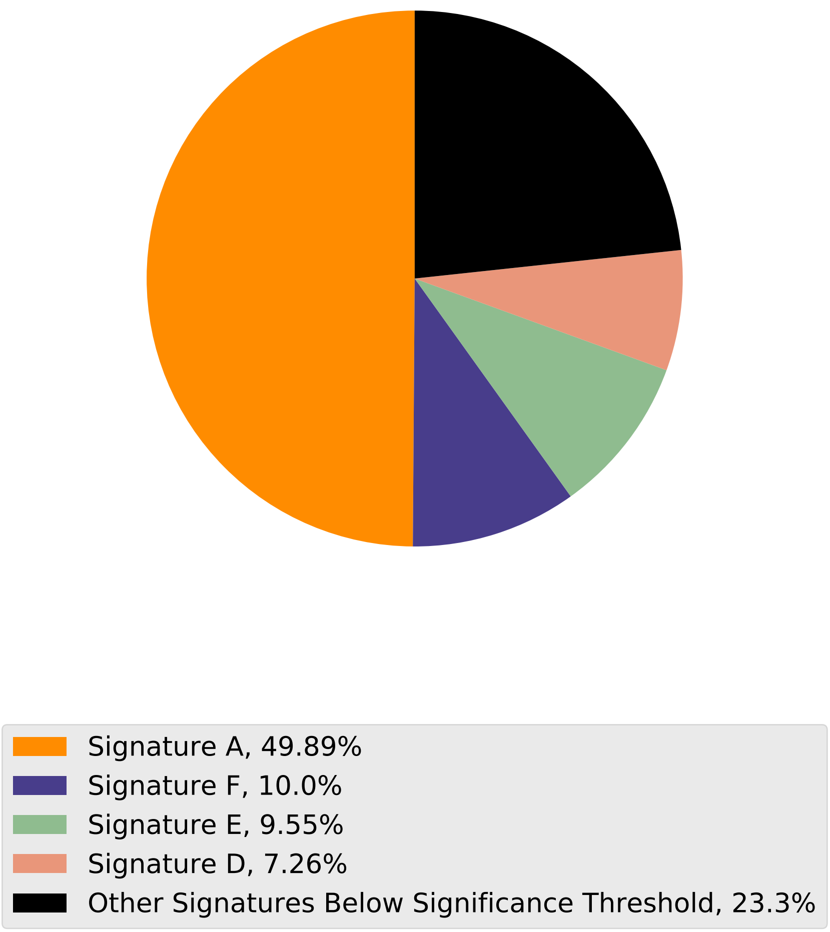

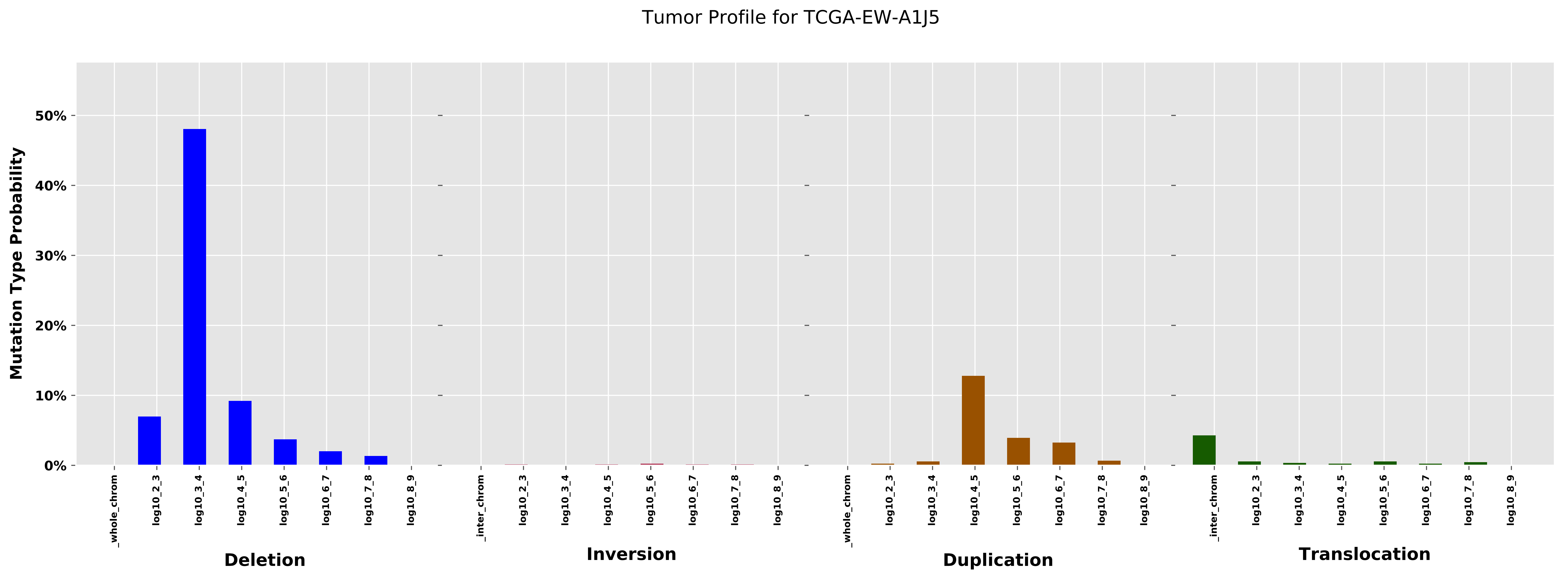

Tumor Profile for TCGA-EW-A1J5
Mutation Type Probability
0%
10%
20%
30%
40%
50%
_whole_chrom
log10_2_3
log10_3_4
log10_4_5
log10_5_6
log10_6_7
log10_7_8
log10_8_9
_inter_chrom
Deletion
Inversion
Duplication
Translocation

Cancer processes Weights for TCGA-EW-A1PC

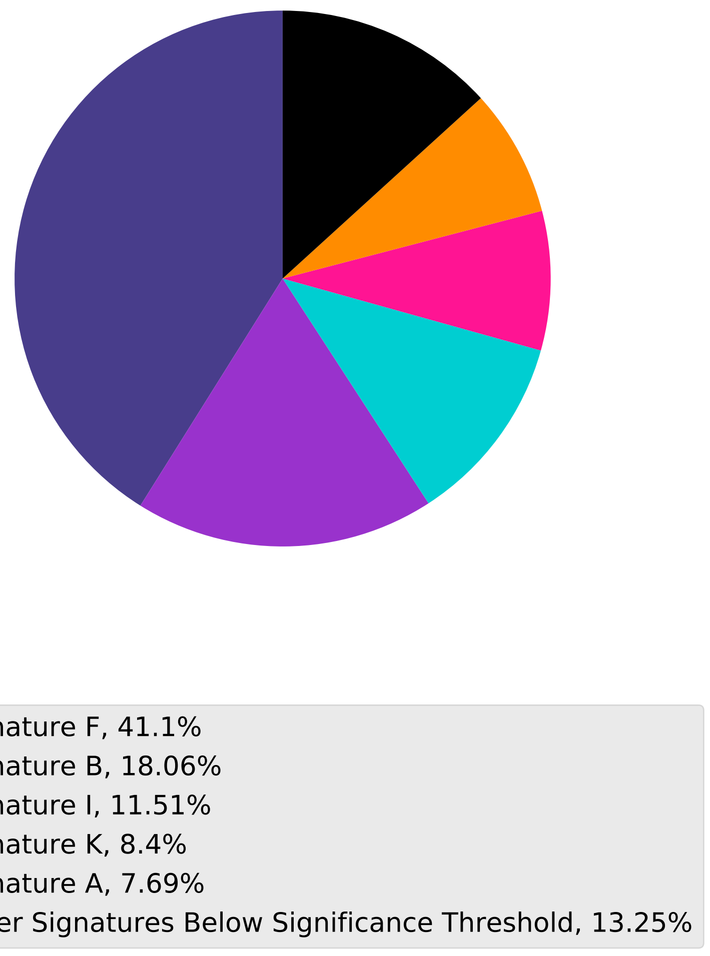

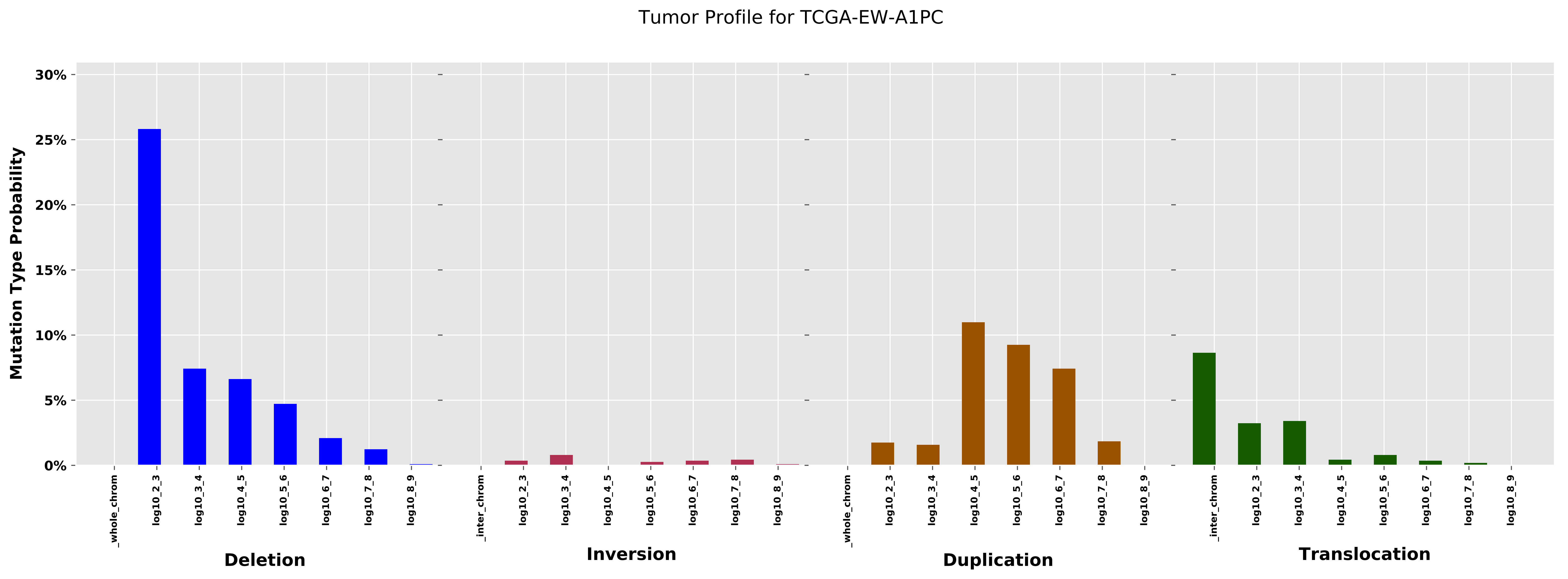
Tumor Profile for TCGA-EW-A1PC
Mutation Type Probability
30%
25%
20%
15%
10%
5%
0%
_whole_chrom
log10_2_3
log10_3_4
log10_4_5
log10_5_6
log10_6_7
log10_7_8
log10_8_9
_inter_chrom
Deletion
Inversion
Duplication
Translocation

Cancer processes Weights for TCGA-BH-A0E0

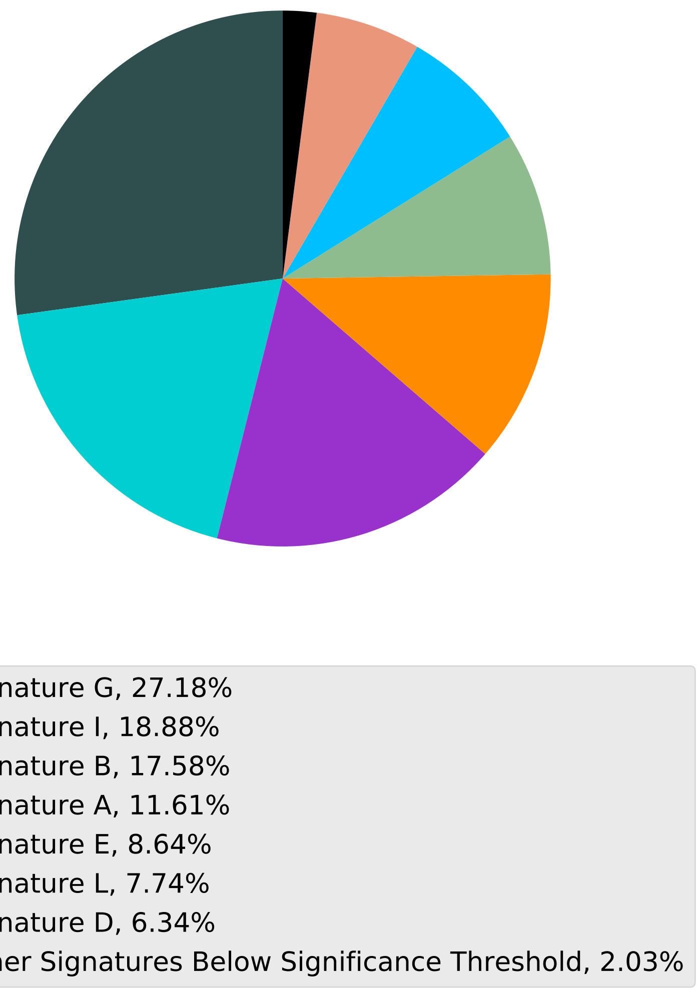

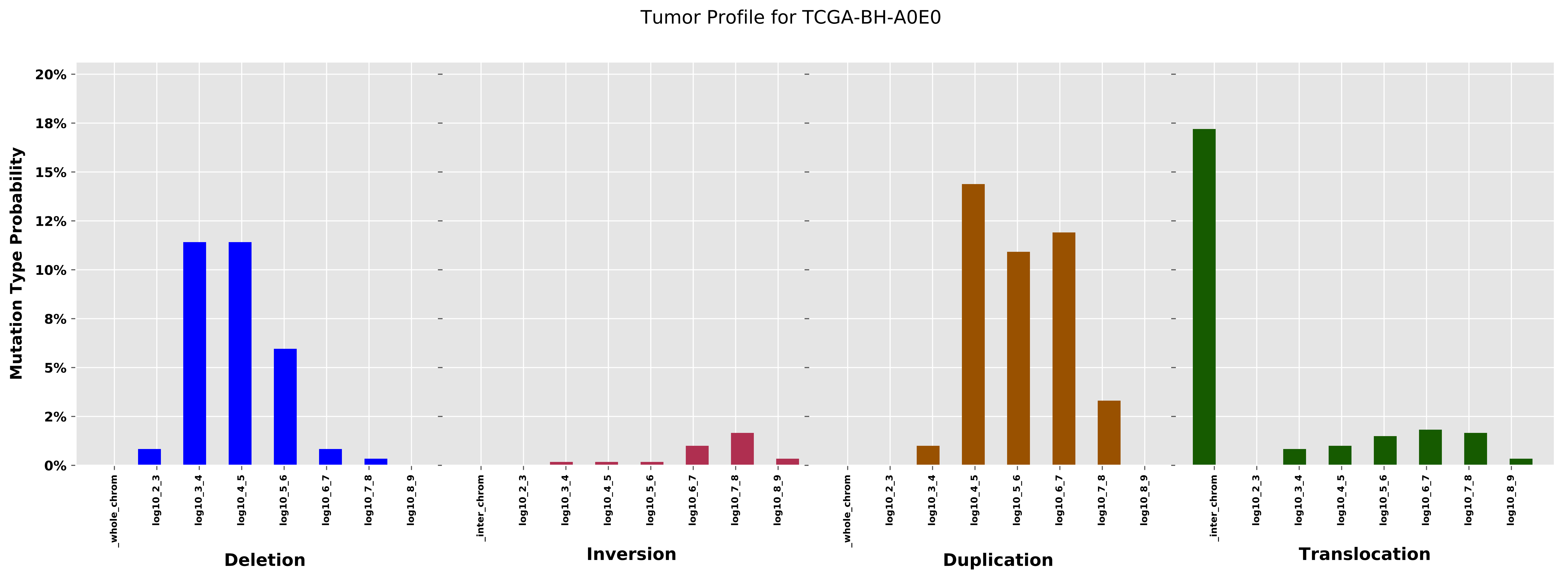
Tumor Profile for TCGA-BH-A0E0
Mutation Type Probability
20%
18%
15%
12%
10%
8%
5%
2%
0%
_whole_chrom
log10_2_3
log10_3_4
log10_4_5
log10_5_6
log10_6_7
log10_7_8
log10_8_9
Deletion
_inter_chrom
log10_2_3
log10_3_4
log10_4_5
log10_5_6
log10_6_7
log10_7_8
log10_8_9
Inversion
_whole_chrom
log10_2_3
log10_3_4
log10_4_5
log10_5_6
log10_6_7
log10_7_8
log10_8_9
Duplication
_inter_chrom
log10_2_3
log10_3_4
log10_4_5
log10_5_6
log10_6_7
log10_7_8
log10_8_9
Translocation

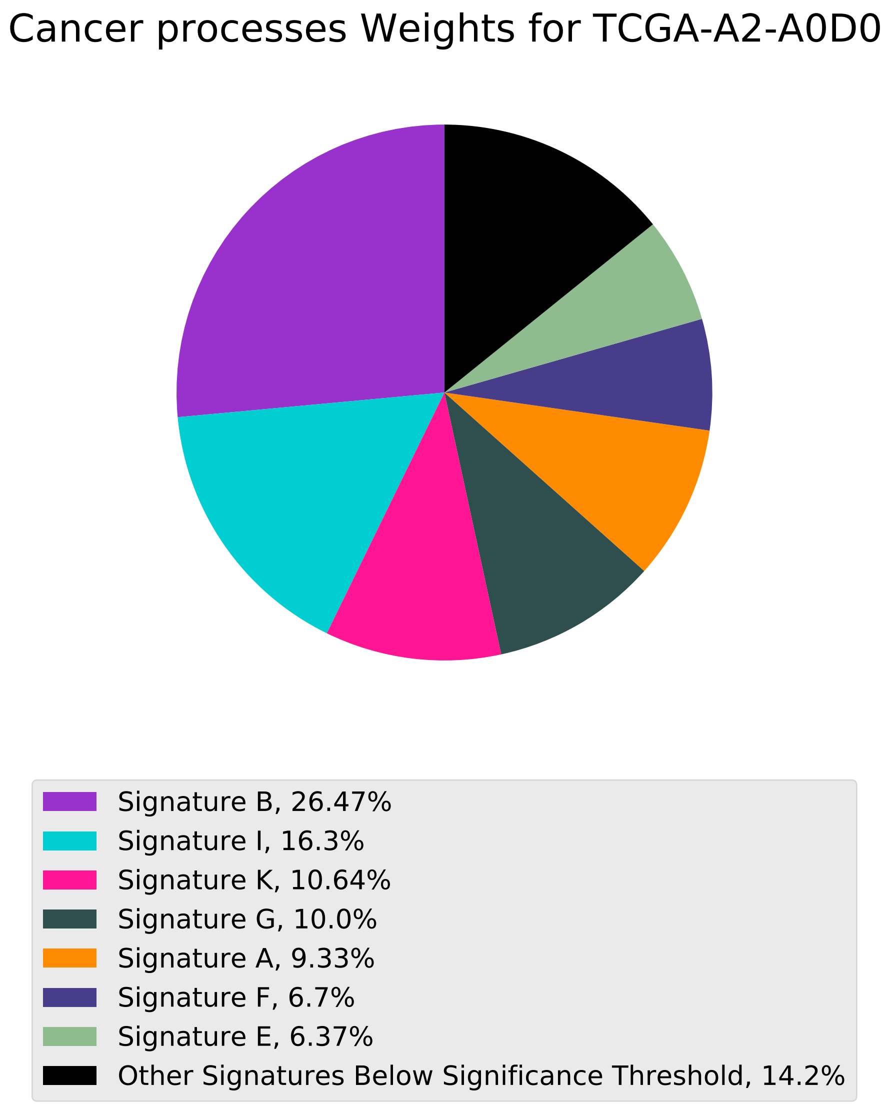

Cancer processes Weights for TCGA-A2-A0D0
Signature B, 26.47%
Signature I, 16.3%
Signature K, 10.64%
Signature G, 10.0%
Signature A, 9.33%
Signature F, 6.7%
Signature E, 6.37%
Other Signatures Below Significance Threshold, 14.2%

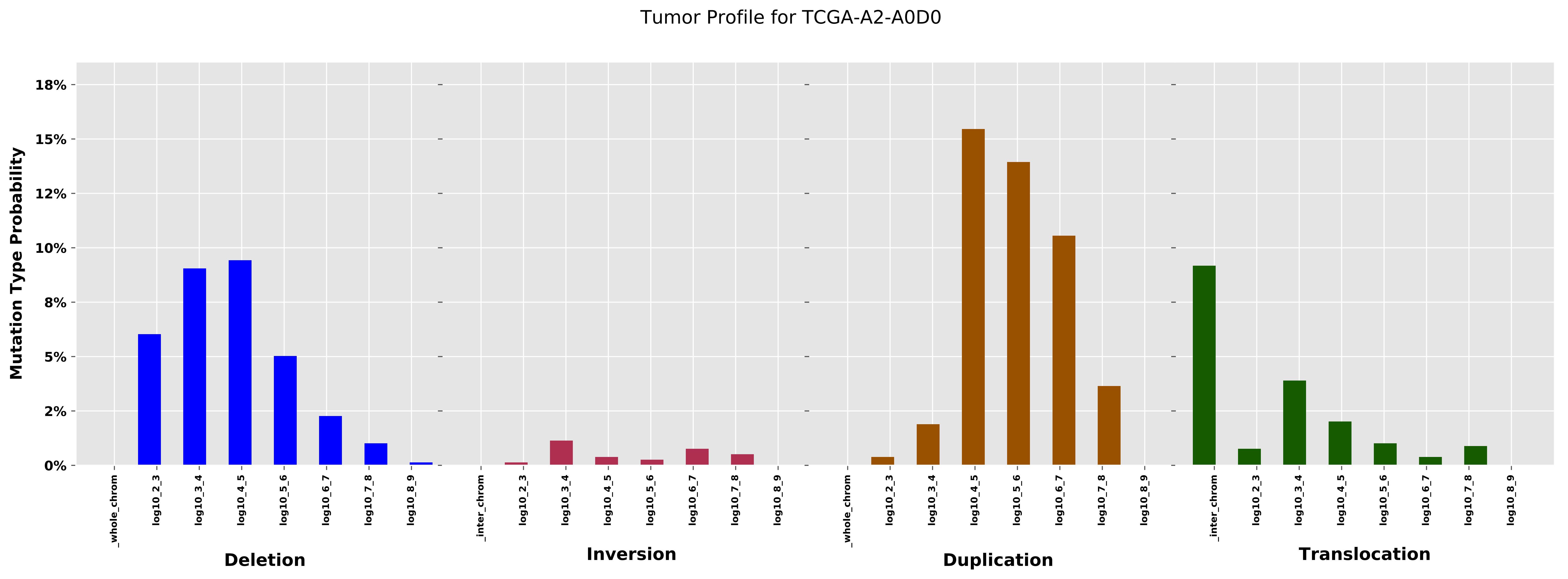
Tumor Profile for TCGA-A2-A0D0
Mutation Type Probability
18%
15%
12%
10%
8%
5%
2%
0%
_whole_chrom
log10_2_3
log10_3_4
log10_4_5
log10_5_6
log10_6_7
log10_7_8
log10_8_9
Deletion
_inter_chrom
Inversion
Duplication
Translocation

Cancer processes Weights for TCGA-AA-3956

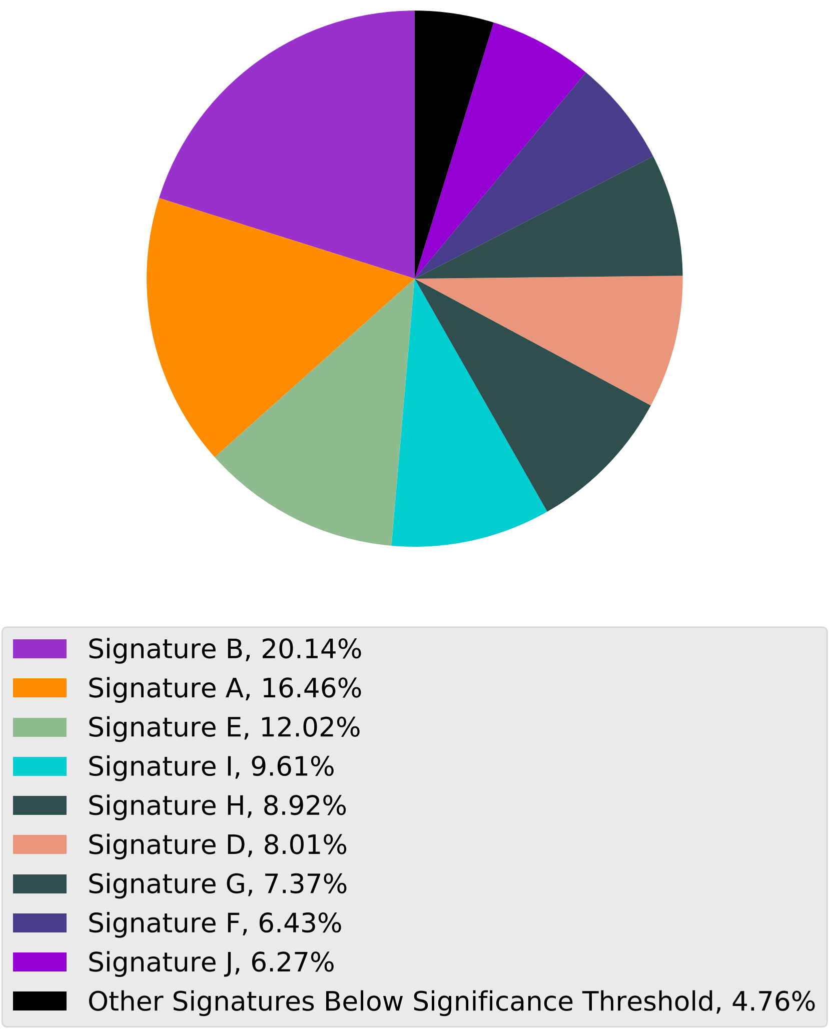

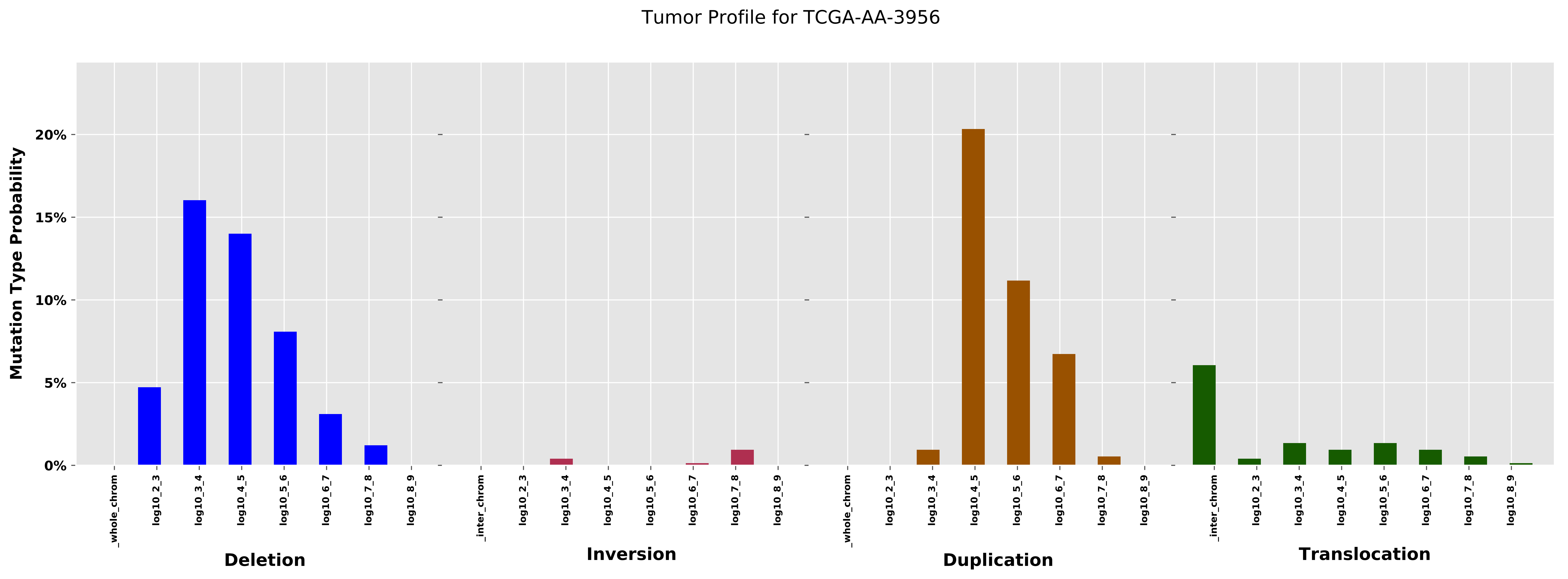
Tumor Profile for TCGA-AA-3956
Mutation Type Probability
20%
15%
10%
5%
0%
_whole_chrom
log10_2_3
log10_3_4
log10_4_5
log10_5_6
log10_6_7
log10_7_8
log10_8_9
Deletion
_inter_chrom
log10_2_3
log10_3_4
log10_4_5
log10_5_6
log10_6_7
log10_7_8
log10_8_9
Inversion
_whole_chrom
log10_2_3
log10_3_4
log10_4_5
log10_5_6
log10_6_7
log10_7_8
log10_8_9
Duplication
_inter_chrom
log10_2_3
log10_3_4
log10_4_5
log10_5_6
log10_6_7
log10_7_8
log10_8_9
Translocation

Cancer processes Weights for TCGA-AR-A0TX

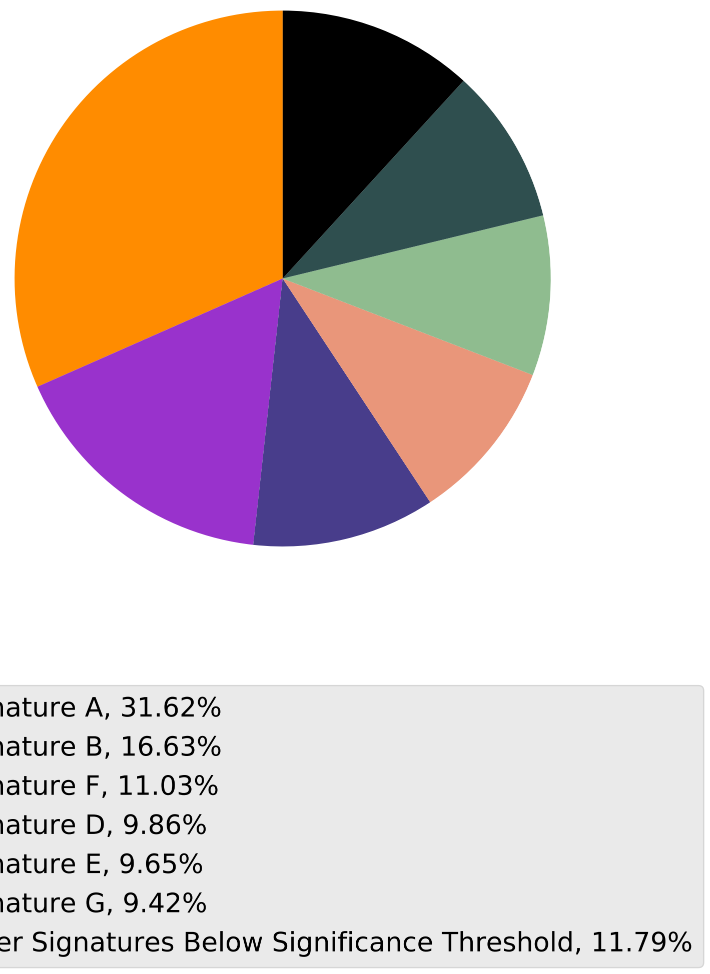

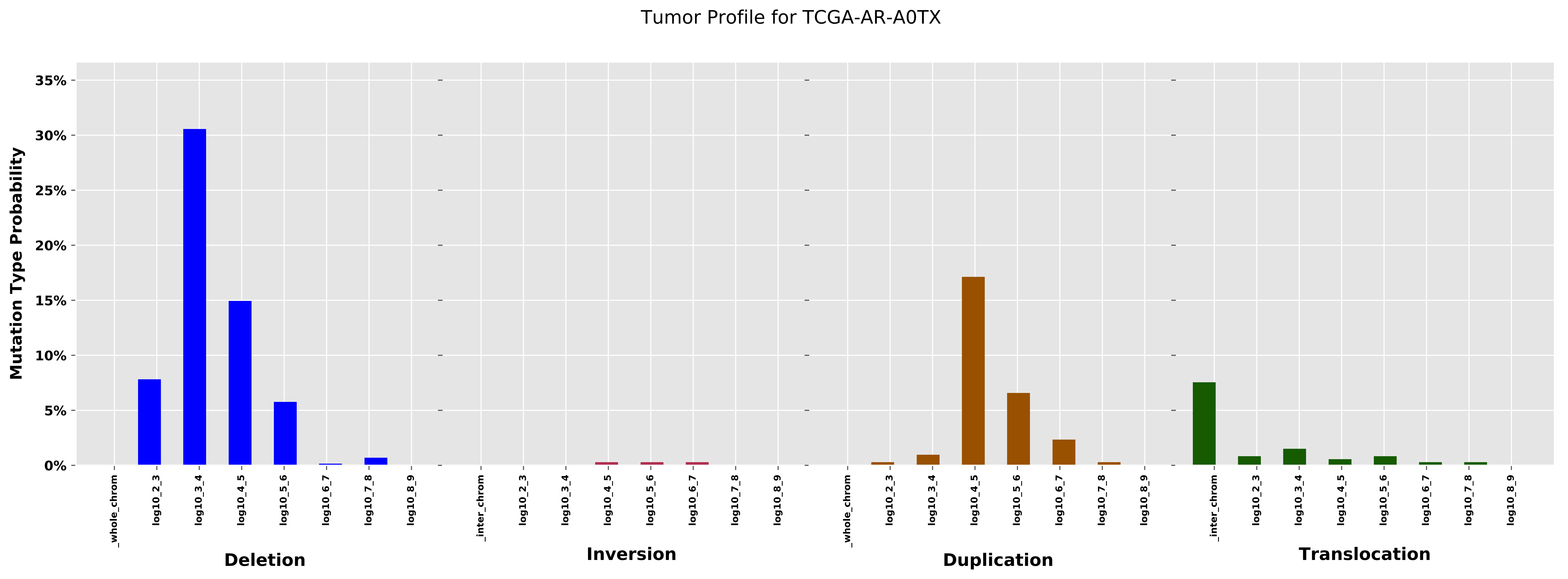

Tumor Profile for TCGA-AR-A0TX
Mutation Type Probability
35%
30%
25%
20%
15%
10%
5%
0%
_whole_chrom
log10_2_3
log10_3_4
log10_4_5
log10_5_6
log10_6_7
log10_7_8
log10_8_9
Deletion
_inter_chrom
log10_2_3
log10_3_4
log10_4_5
log10_5_6
log10_6_7
log10_7_8
log10_8_9
Inversion
_whole_chrom
log10_2_3
log10_3_4
log10_4_5
log10_5_6
log10_6_7
log10_7_8
log10_8_9
Duplication
_inter_chrom
log10_2_3
log10_3_4
log10_4_5
log10_5_6
log10_6_7
log10_7_8
log10_8_9
Translocation

Cancer processes Weights for TCGA-EI-6917

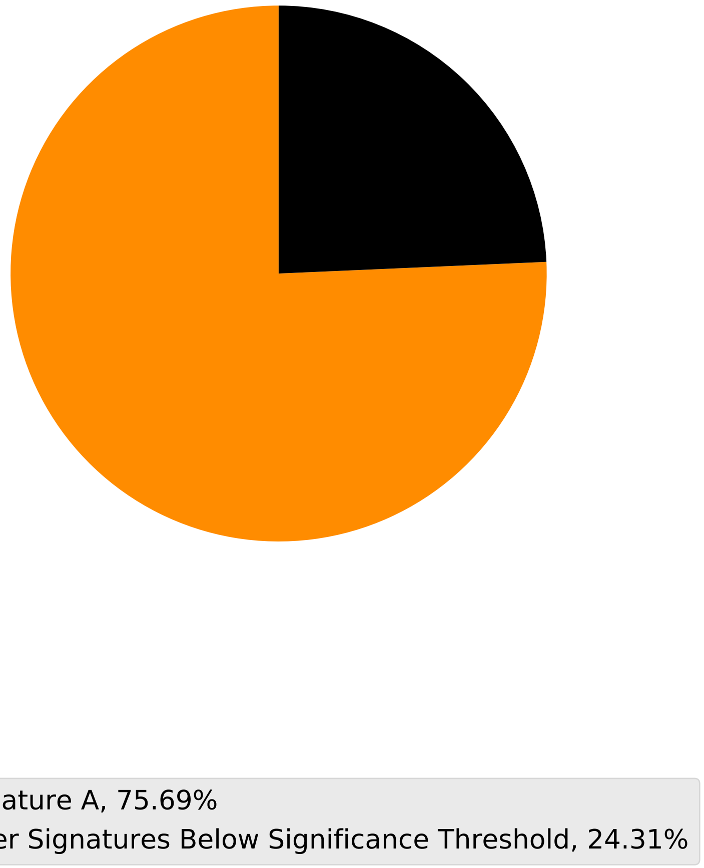

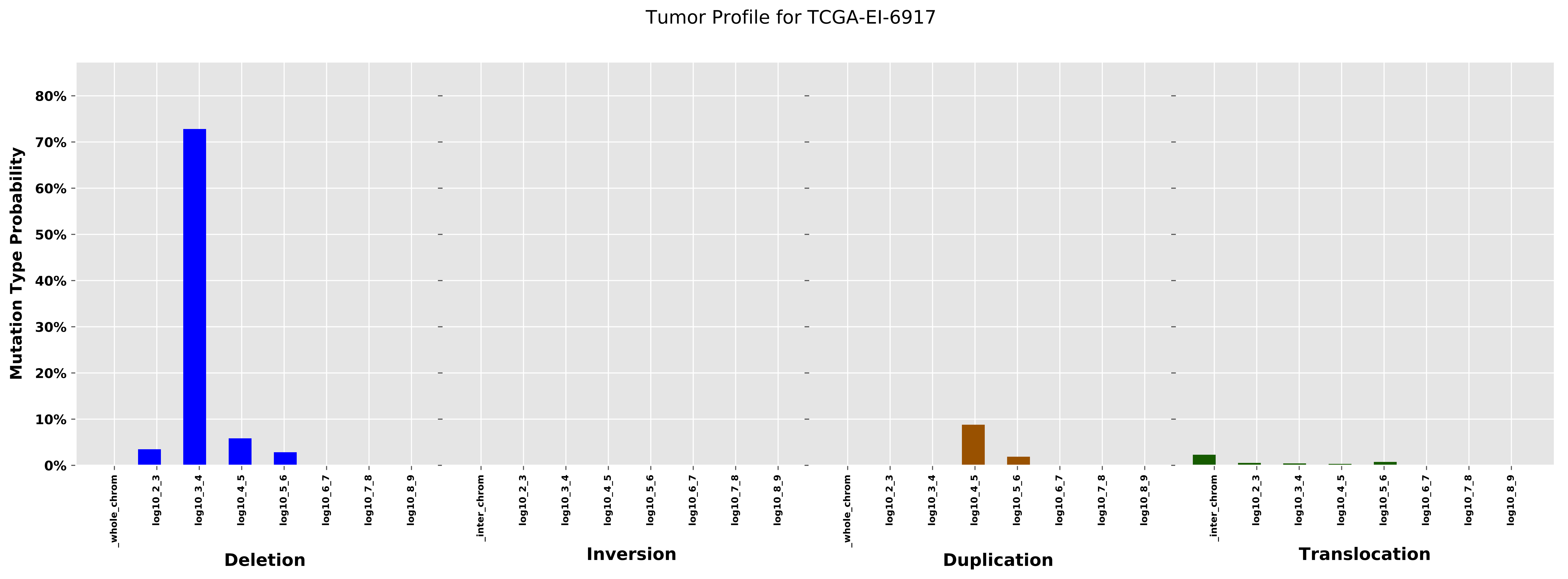

Tumor Profile for TCGA-EI-6917
Mutation Type Probability
80%
70%
60%
50%
40%
30%
20%
10%
0%
_whole_chrom
log10_2_3
log10_3_4
log10_4_5
log10_5_6
log10_6_7
log10_7_8
log10_8_9
_inter_chrom
Deletion
Inversion
Duplication
Translocation

#### Cancer processes Weights for TCGA-AA-A01X

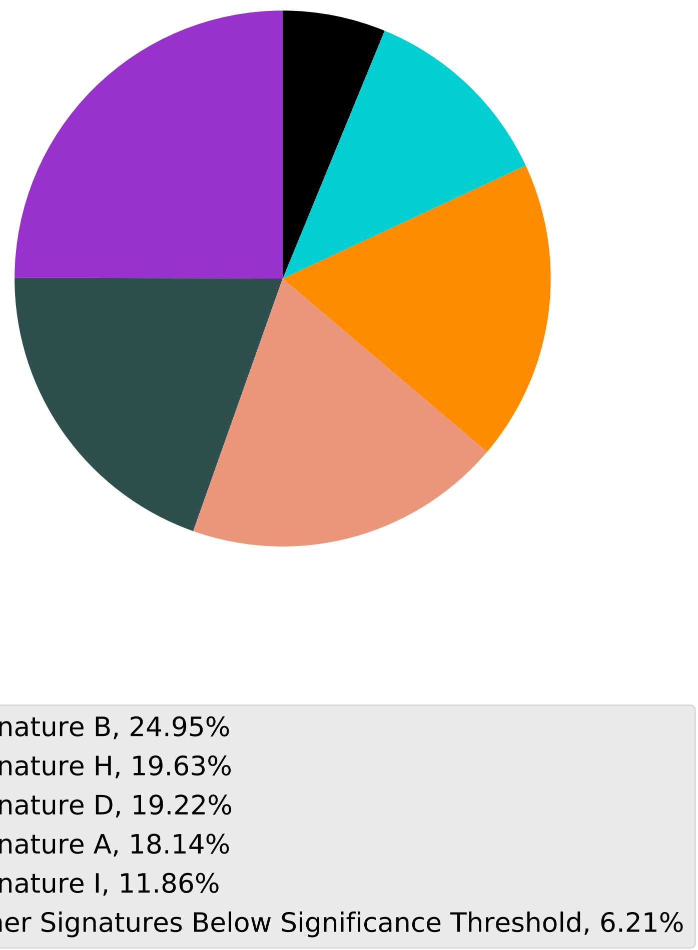

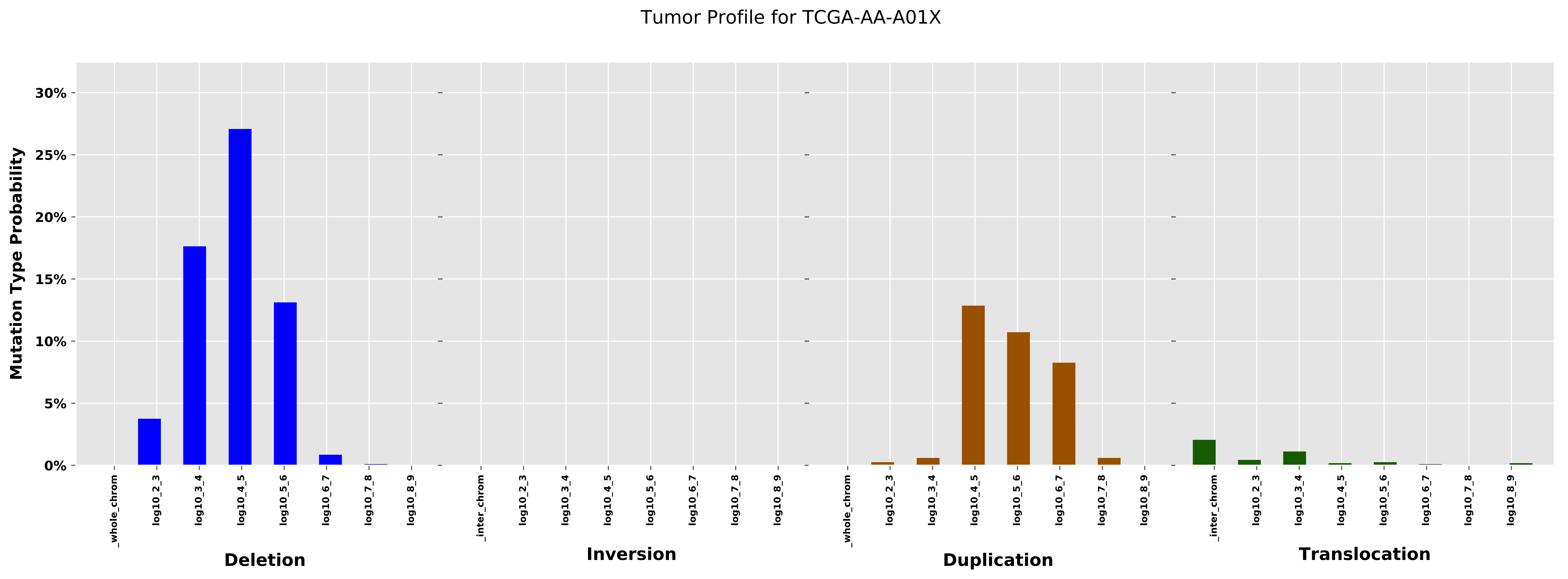
Tumor Profile for TCGA-AA-A01X
Mutation Type Probability
30%
25%
20%
15%
10%
5%
0%
_whole_chrom
log10_2_3
log10_3_4
log10_4_5
log10_5_6
log10_6_7
log10_7_8
log10_8_9
_inter_chrom
Deletion
Inversion
Duplication
Translocation

Cancer processes Weights for TCGA-E2-A15H

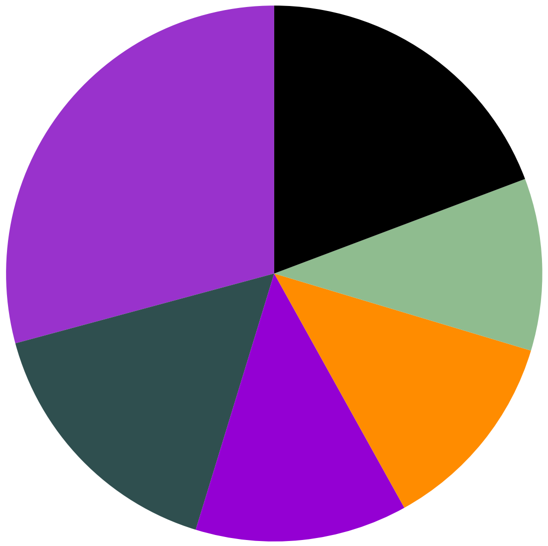

- Signature B, 29.2%
- Signature H, 16.08%
- Signature J, 12.79%
- Signature A, 12.29%
- Signature E, 10.39%
- Other Signatures Below Significance Threshold, 19.26%

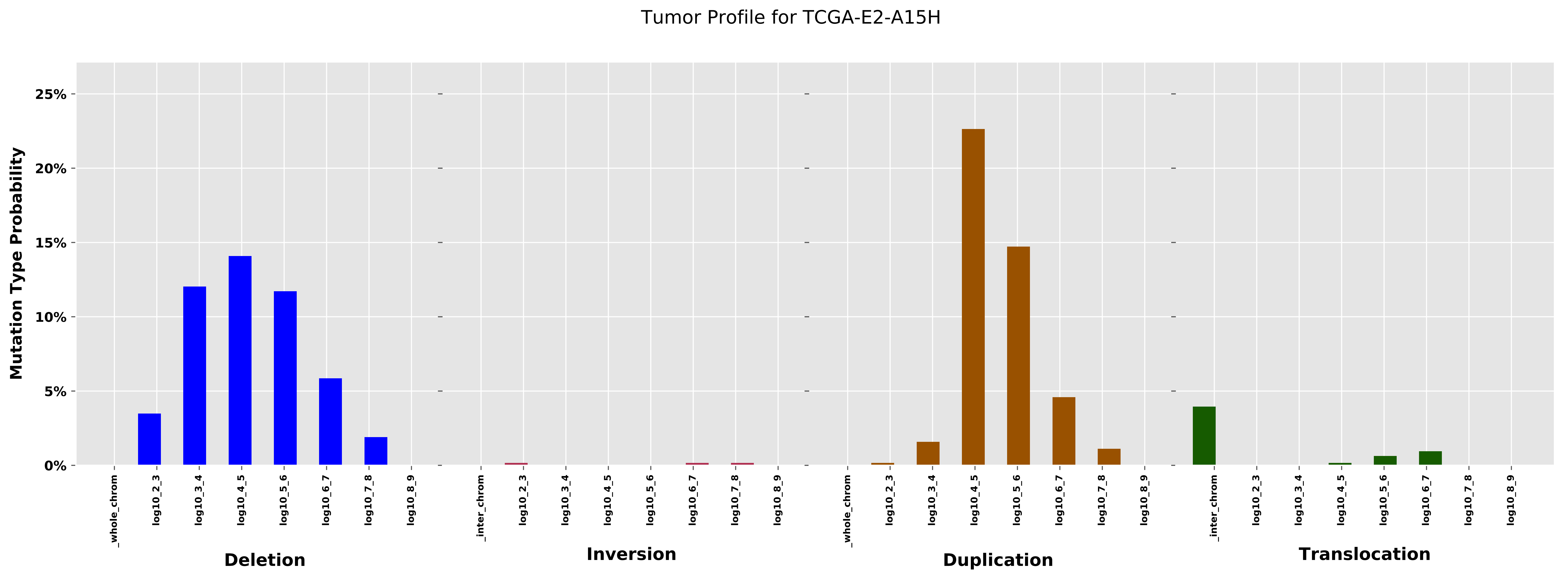
Tumor Profile for TCGA-E2-A15H
Mutation Type Probability
0%
5%
10%
15%
20%
25%
_whole_chrom
log10_2_3
log10_3_4
log10_4_5
log10_5_6
log10_6_7
log10_7_8
log10_8_9
_inter_chrom
Deletion
Inversion
Duplication
Translocation

Cancer processes Weights for TCGA-E9-A1NH

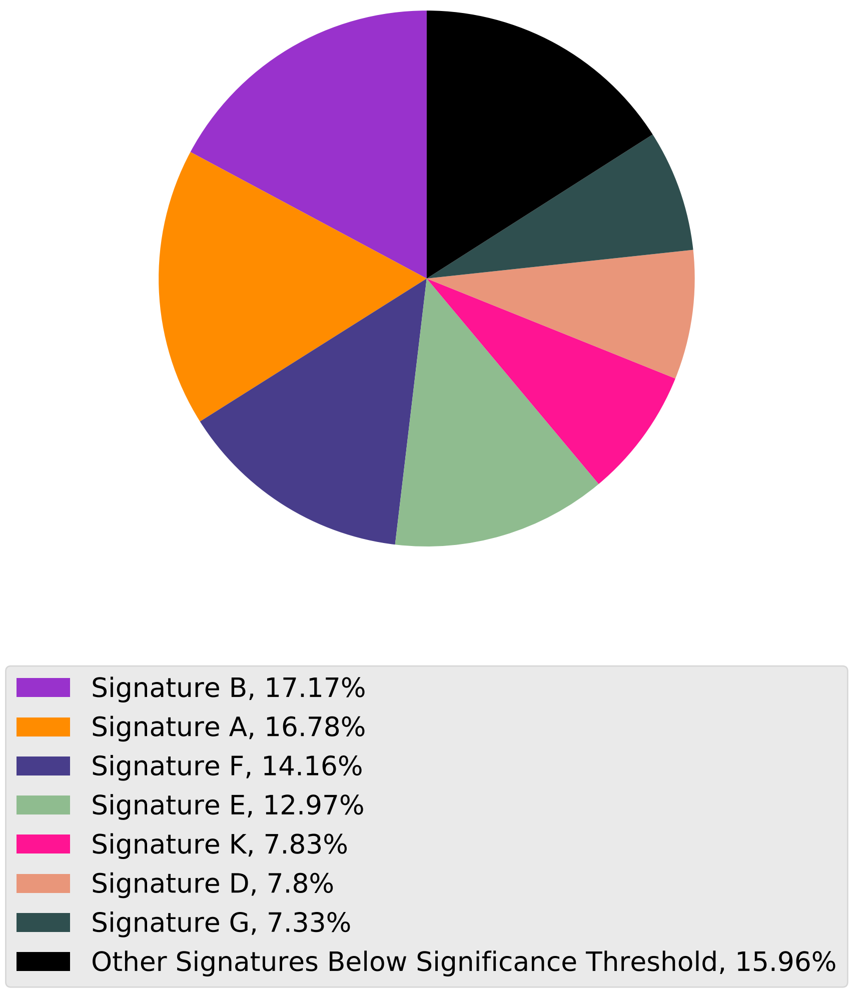

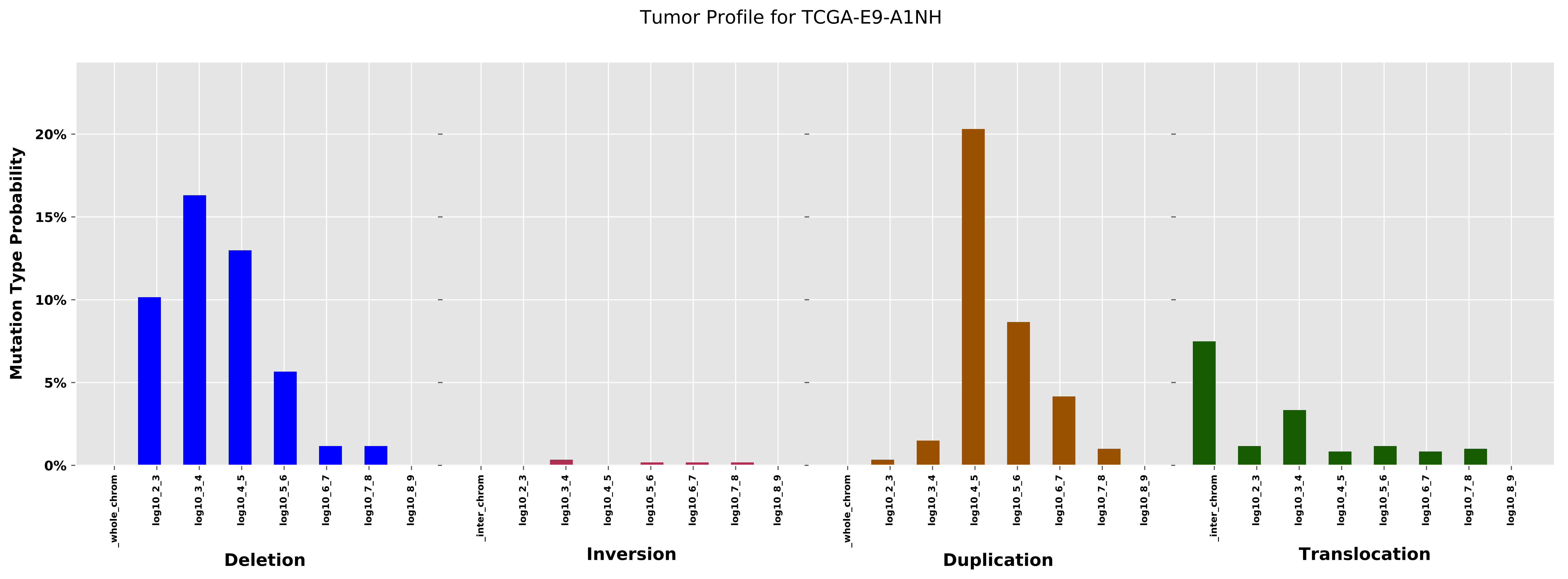

Tumor Profile for TCGA-E9-A1NH
Mutation Type Probability
0%
5%
10%
15%
20%
_whole_chrom
log10_2_3
log10_3_4
log10_4_5
log10_5_6
log10_6_7
log10_7_8
log10_8_9
Deletion
_inter_chrom
log10_2_3
log10_3_4
log10_4_5
log10_5_6
log10_6_7
log10_7_8
log10_8_9
Inversion
_whole_chrom
log10_2_3
log10_3_4
log10_4_5
log10_5_6
log10_6_7
log10_7_8
log10_8_9
Duplication
_inter_chrom
log10_2_3
log10_3_4
log10_4_5
log10_5_6
log10_6_7
log10_7_8
log10_8_9
Translocation

Cancer processes Weights for TCGA-AA-A01R

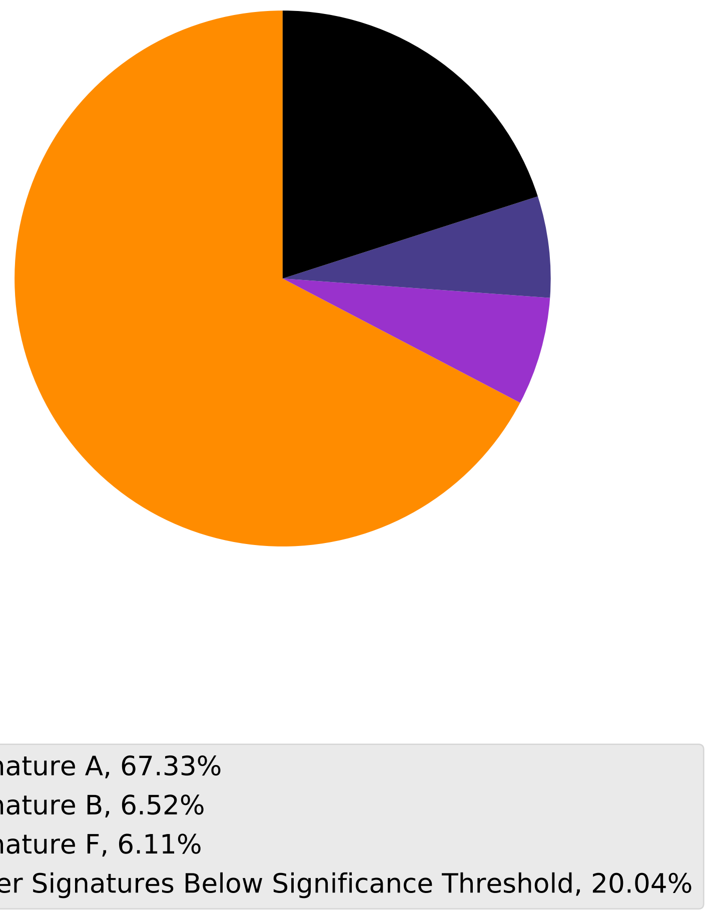

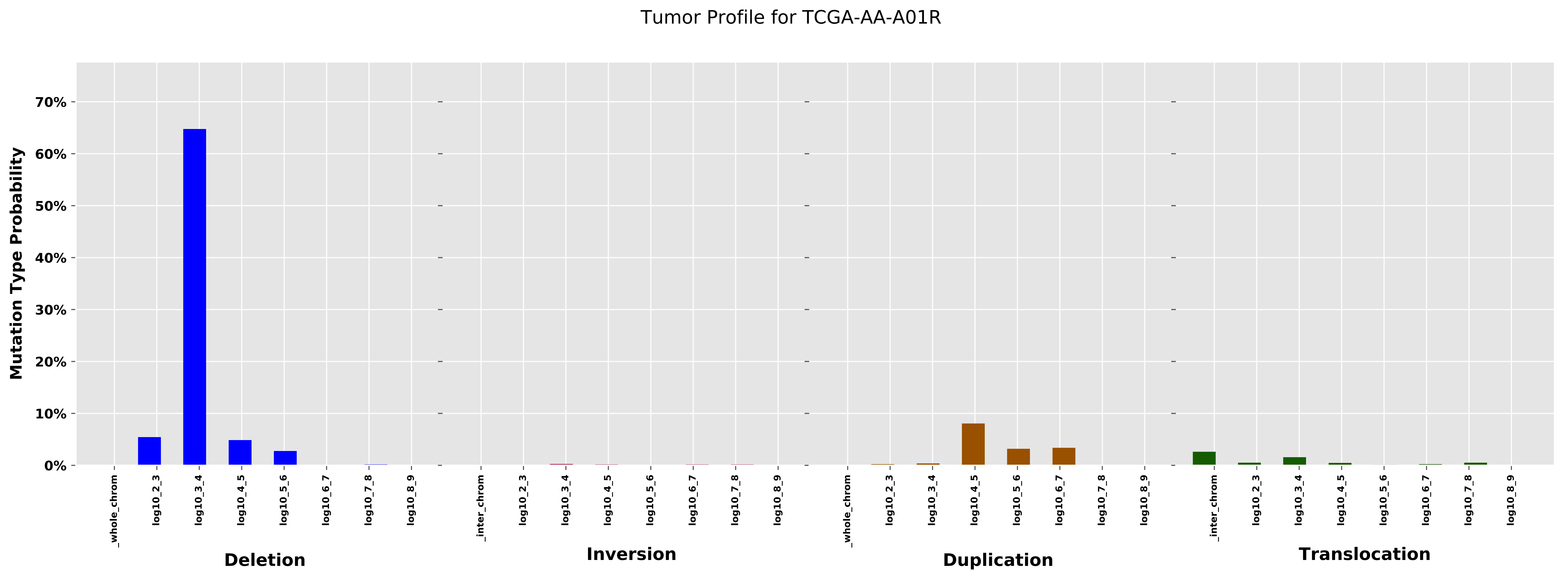

Tumor Profile for TCGA-AA-A01R
Mutation Type Probability
70%
60%
50%
40%
30%
20%
10%
0%
_whole_chrom
log10_2_3
log10_3_4
log10_4_5
log10_5_6
log10_6_7
log10_7_8
log10_8_9
_inter_chrom
log10_2_3
log10_3_4
log10_4_5
log10_5_6
log10_6_7
log10_7_8
log10_8_9
_whole_chrom
log10_2_3
log10_3_4
log10_4_5
log10_5_6
log10_6_7
log10_7_8
log10_8_9
_inter_chrom
log10_2_3
log10_3_4
log10_4_5
log10_5_6
log10_6_7
log10_7_8
log10_8_9
Deletion
Inversion
Duplication
Translocation

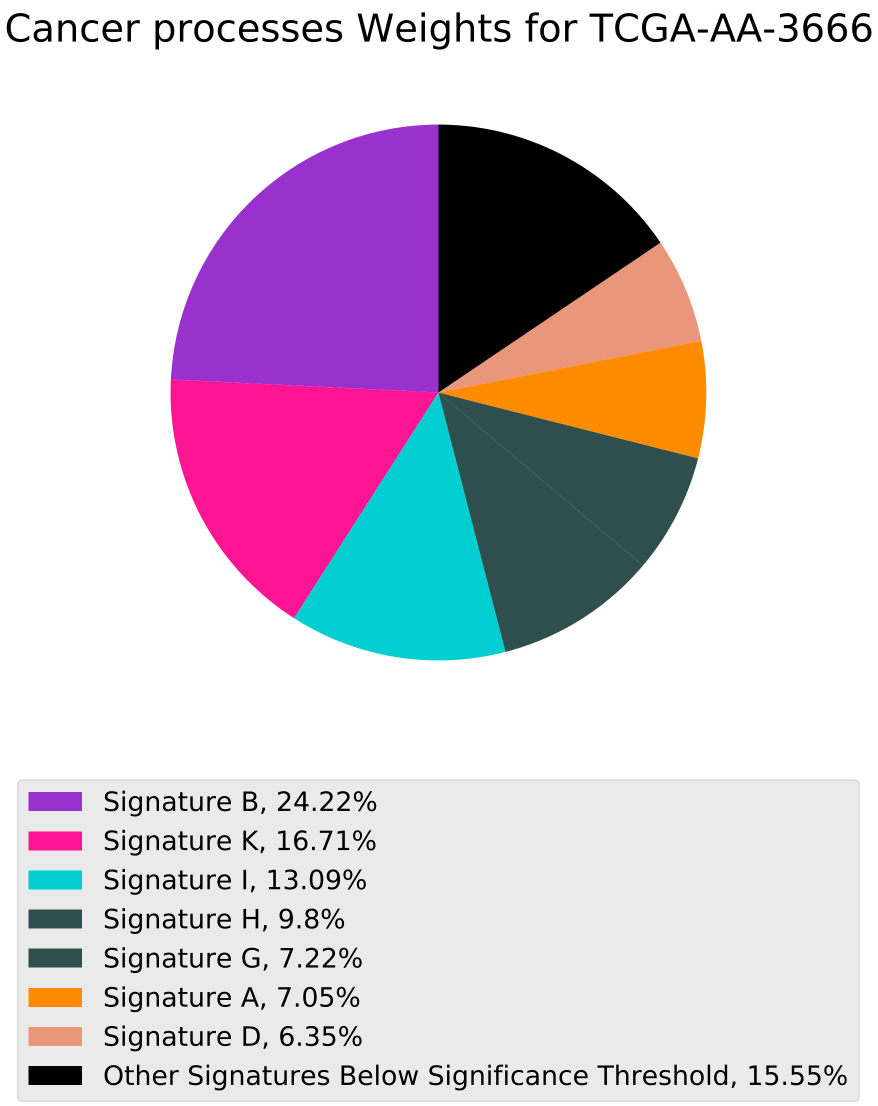
Cancer processes Weights for TCGA-AA-3666
Signature B, 24.22%
Signature K, 16.71%
Signature I, 13.09%
Signature H, 9.8%
Signature G, 7.22%
Signature A, 7.05%
Signature D, 6.35%
Other Signatures Below Significance Threshold, 15.55%

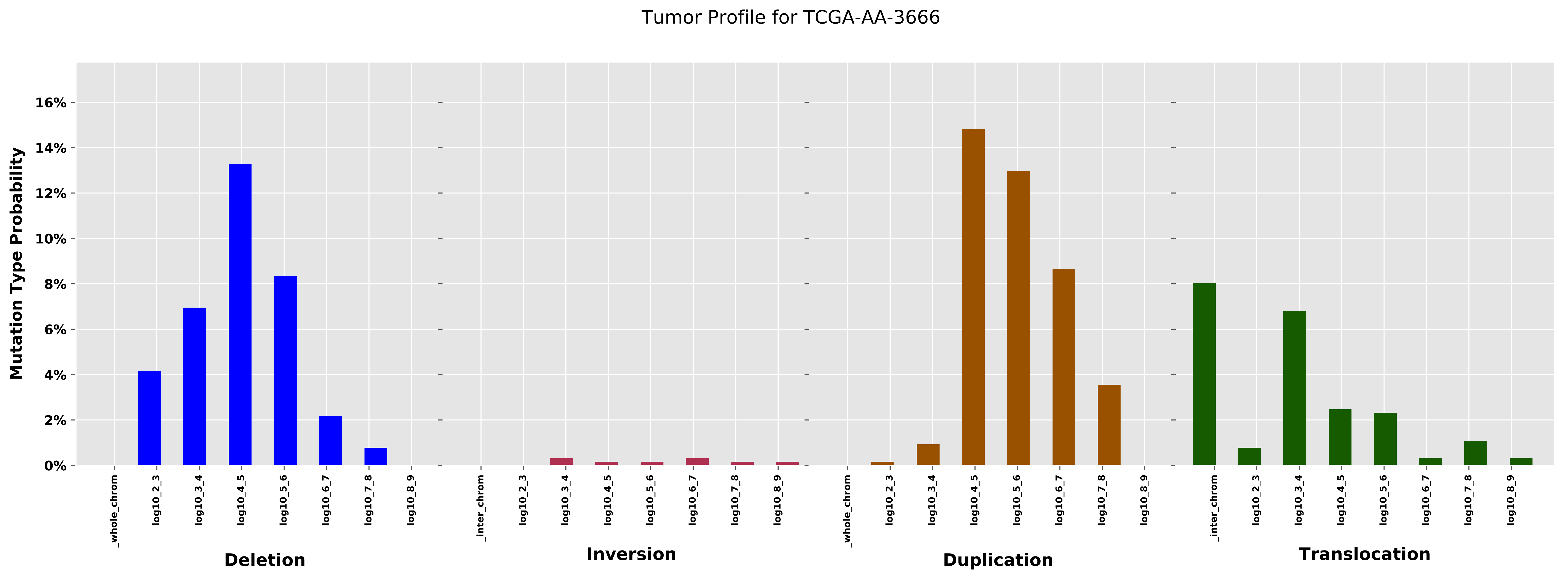

Tumor Profile for TCGA-AA-3666
Mutation Type Probability
0%
2%
4%
6%
8%
10%
12%
14%
16%
_whole_chrom
log10_2_3
log10_3_4
log10_4_5
log10_5_6
log10_6_7
log10_7_8
log10_8_9
_inter_chrom
Deletion
Inversion
Duplication
Translocation

Cancer processes Weights for TCGA-AR-A2LK

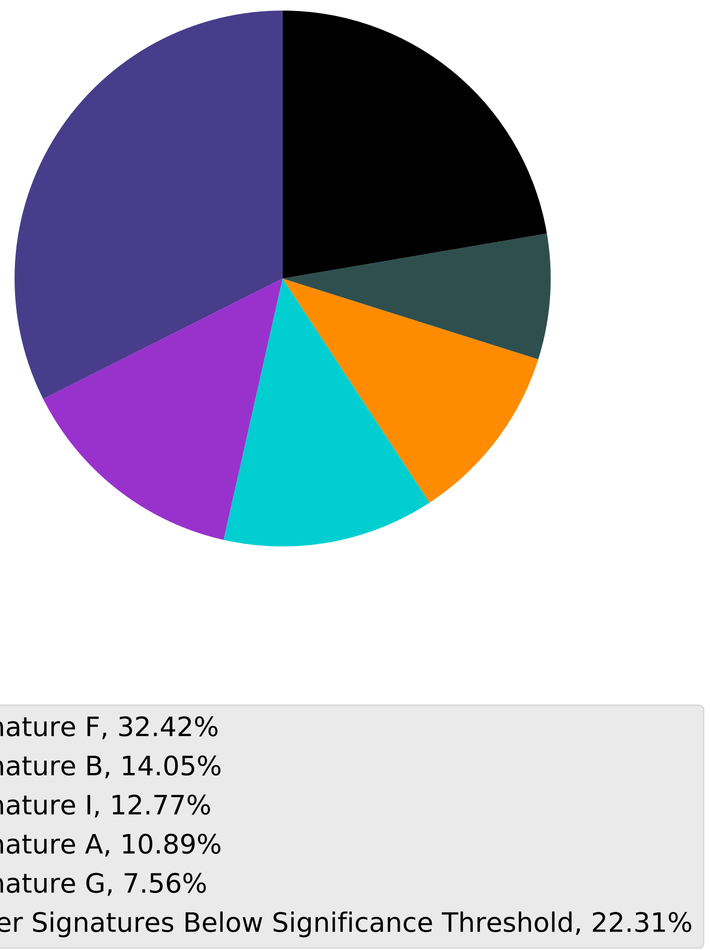

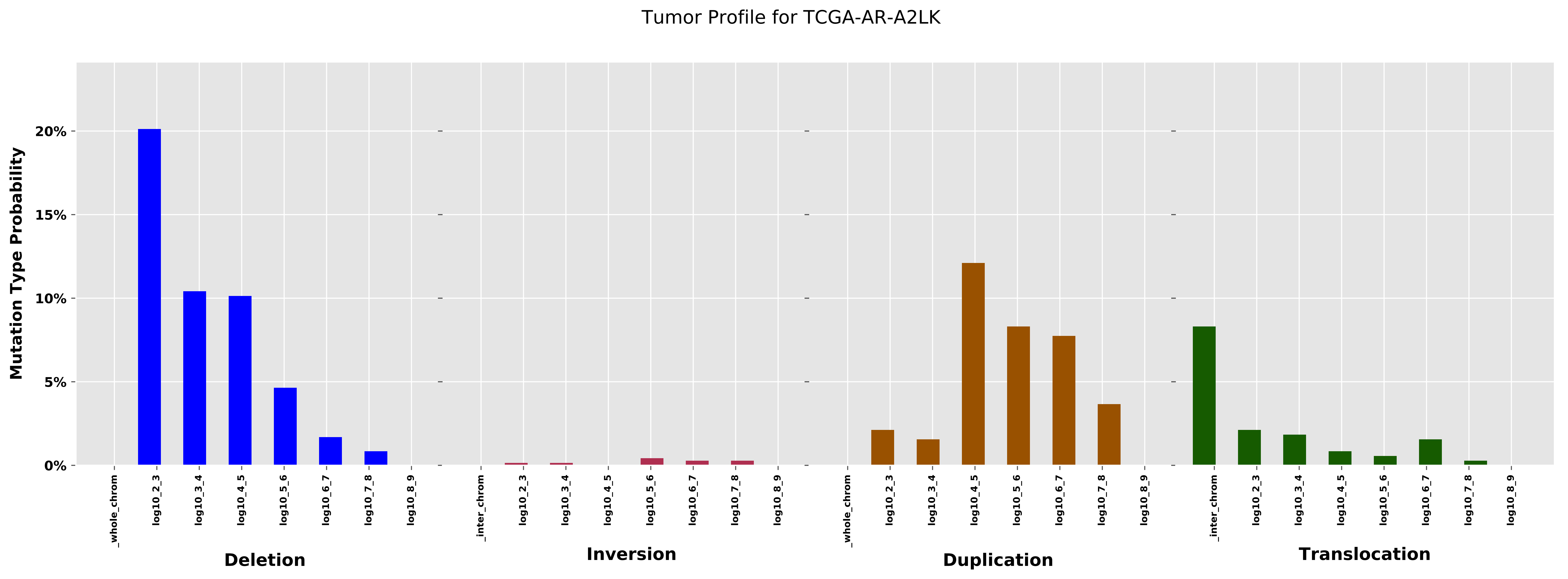

Tumor Profile for TCGA-AR-A2LK
Mutation Type Probability
20%
15%
10%
5%
0%
_whole_chrom
log10_2_3
log10_3_4
log10_4_5
log10_5_6
log10_6_7
log10_7_8
log10_8_9
_inter_chrom
Deletion
Inversion
Duplication
Translocation

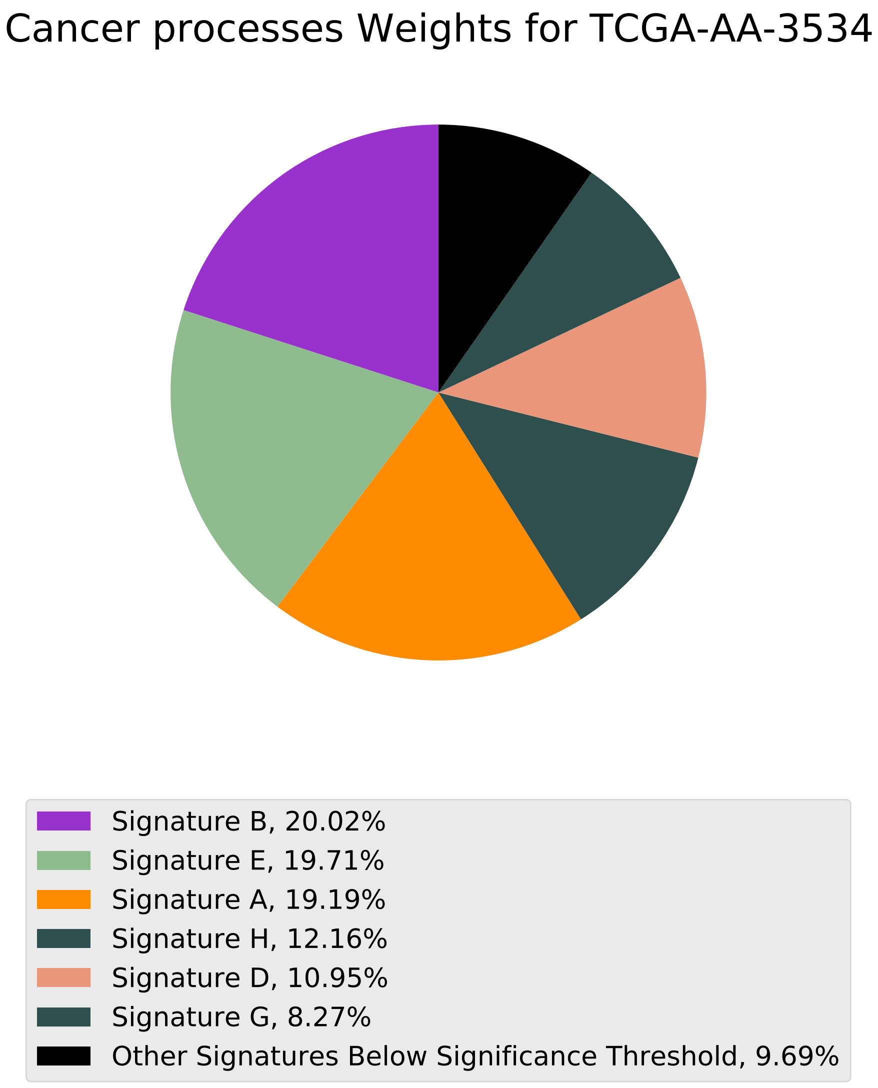
Cancer processes Weights for TCGA-AA-3534
Signature B, 20.02%
Signature E, 19.71%
Signature A, 19.19%
Signature H, 12.16%
Signature D, 10.95%
Signature G, 8.27%
Other Signatures Below Significance Threshold, 9.69%

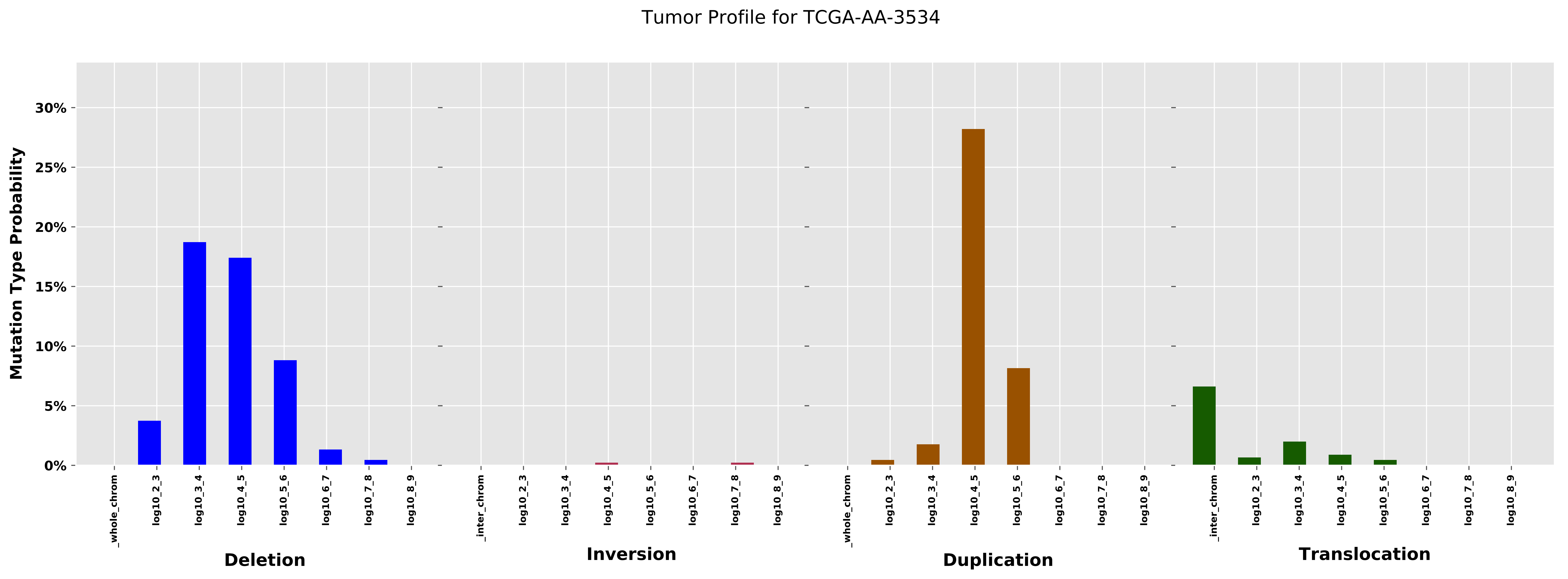

Tumor Profile for TCGA-AA-3534
Mutation Type Probability
30%
25%
20%
15%
10%
5%
0%
_whole_chrom
log10_2_3
log10_3_4
log10_4_5
log10_5_6
log10_6_7
log10_7_8
log10_8_9
Deletion
_inter_chrom
log10_2_3
log10_3_4
log10_4_5
log10_5_6
log10_6_7
log10_7_8
log10_8_9
Inversion
_whole_chrom
log10_2_3
log10_3_4
log10_4_5
log10_5_6
log10_6_7
log10_7_8
log10_8_9
Duplication
_inter_chrom
log10_2_3
log10_3_4
log10_4_5
log10_5_6
log10_6_7
log10_7_8
log10_8_9
Translocation

Cancer processes Weights for TCGA-A2-A25B

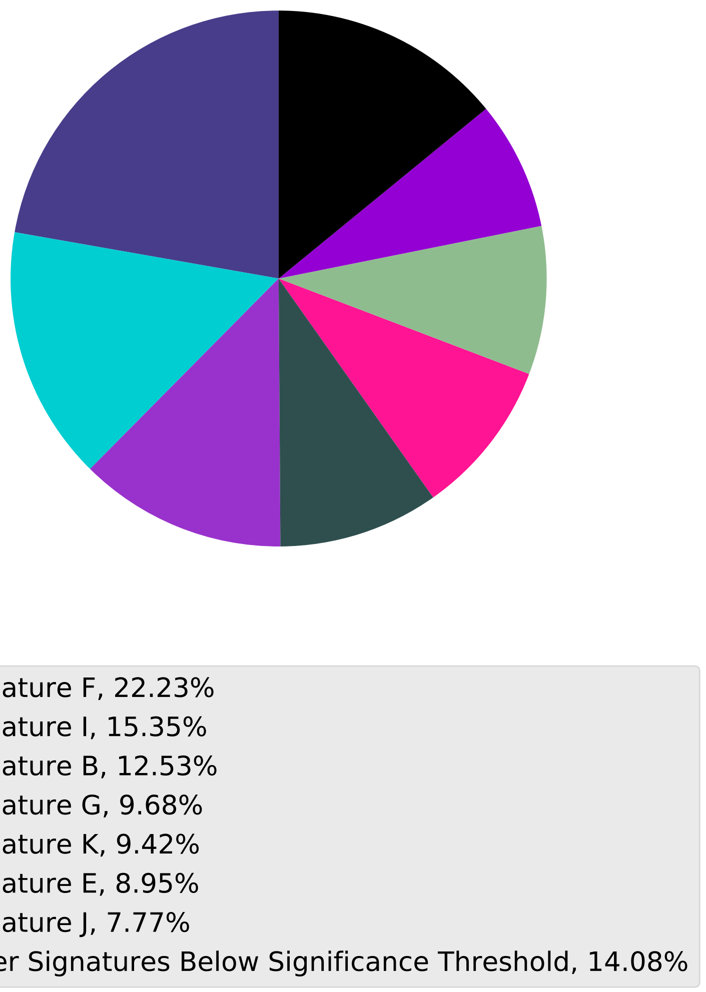

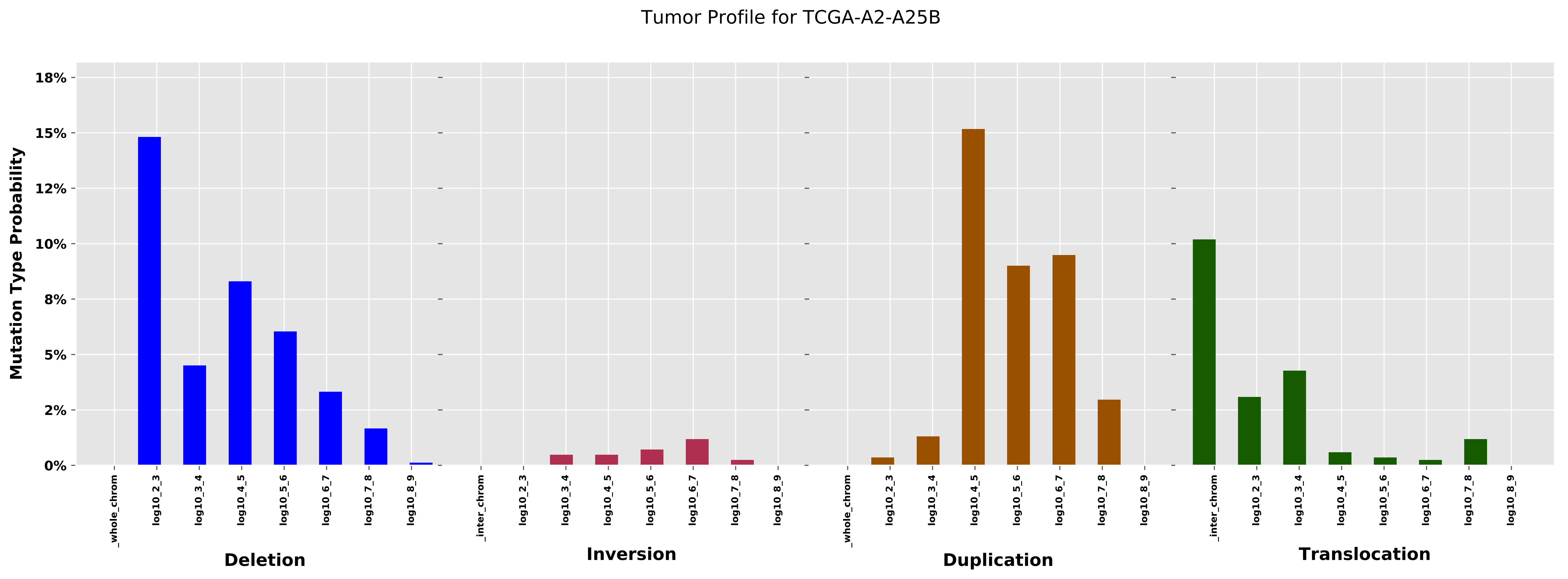
Tumor Profile for TCGA-A2-A25B
Mutation Type Probability
18%
15%
12%
10%
8%
5%
2%
0%
_whole_chrom
log10_2_3
log10_3_4
log10_4_5
log10_5_6
log10_6_7
log10_7_8
log10_8_9
_inter_chrom
Deletion
Inversion
Duplication
Translocation

Cancer processes Weights for TCGA-AG-4008

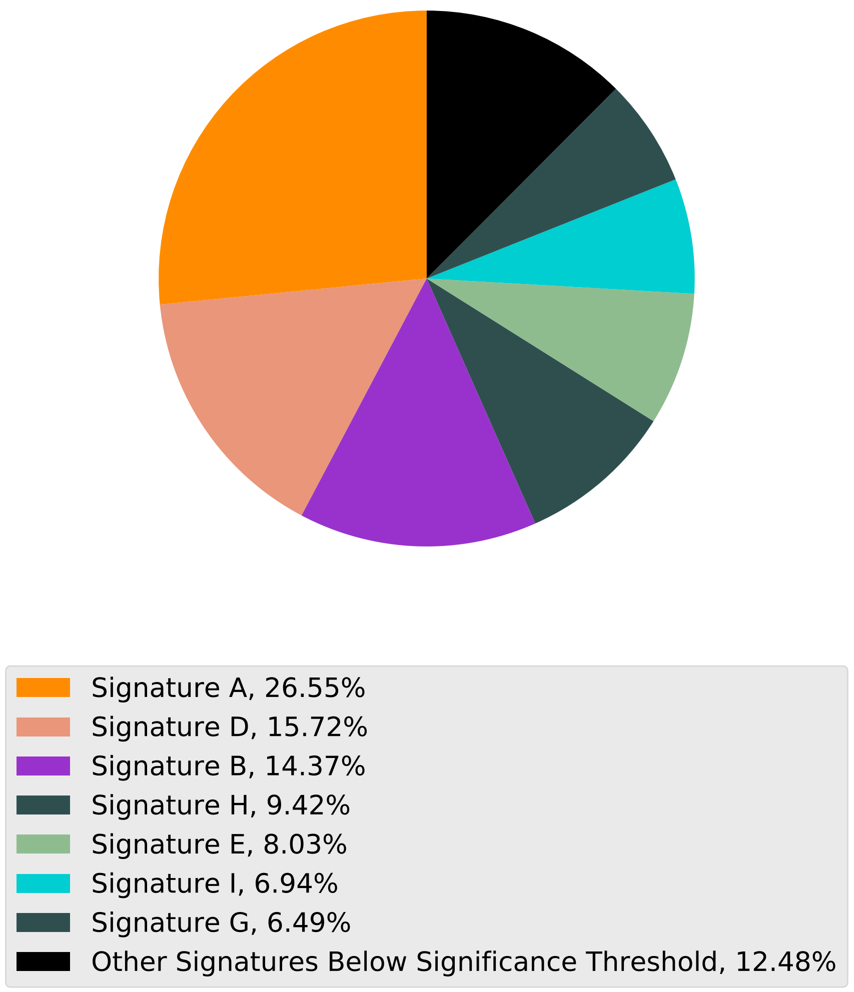

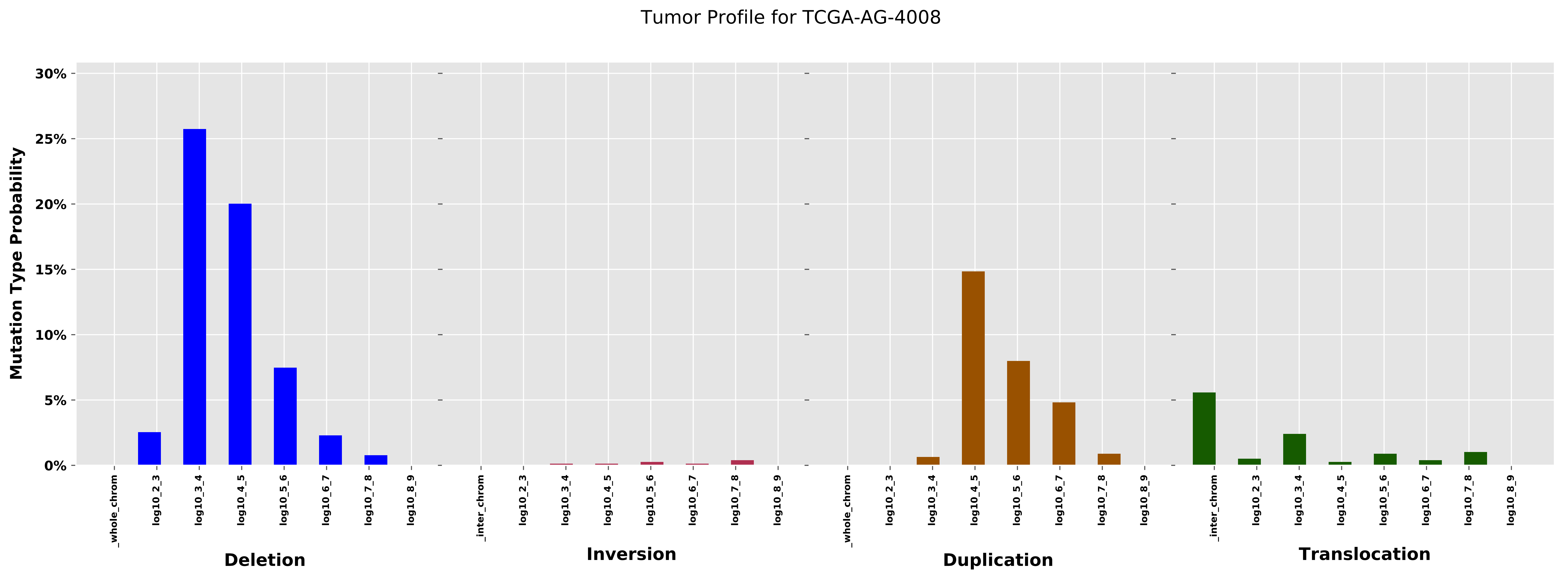

Tumor Profile for TCGA-AG-4008
Mutation Type Probability
30%
25%
20%
15%
10%
5%
0%
_whole_chrom
log10_2_3
log10_3_4
log10_4_5
log10_5_6
log10_6_7
log10_7_8
log10_8_9
Deletion
_inter_chrom
log10_2_3
log10_3_4
log10_4_5
log10_5_6
log10_6_7
log10_7_8
log10_8_9
Inversion
_whole_chrom
log10_2_3
log10_3_4
log10_4_5
log10_5_6
log10_6_7
log10_7_8
log10_8_9
Duplication
_inter_chrom
log10_2_3
log10_3_4
log10_4_5
log10_5_6
log10_6_7
log10_7_8
log10_8_9
Translocation

Cancer processes Weights for TCGA-A7-A26G

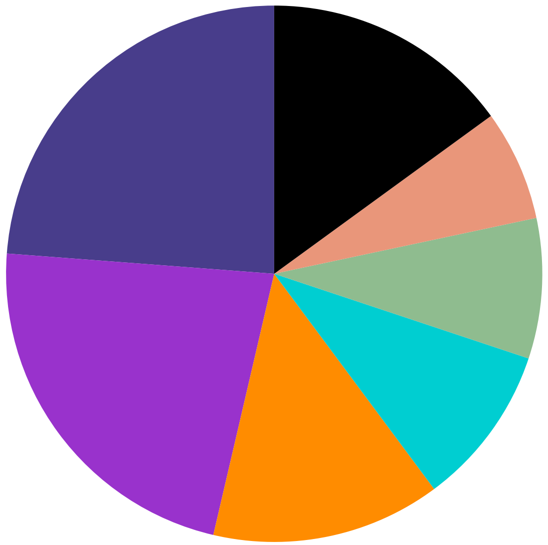

- Signature F, 23.8%
- Signature B, 22.56%
- Signature A, 13.82%
- Signature I, 9.71%
- Signature E, 8.46%
- Signature D, 6.67%
- Other Signatures Below Significance Threshold, 14.98%

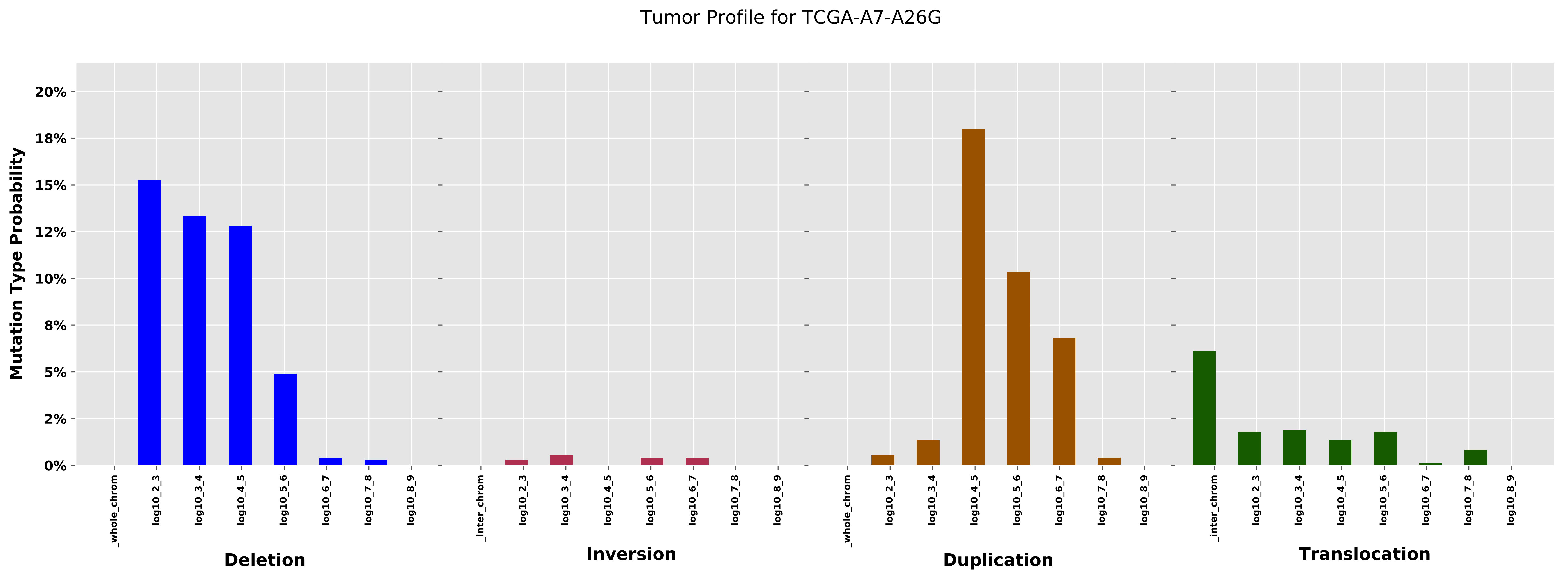
Tumor Profile for TCGA-A7-A26G
Mutation Type Probability
0%
2%
5%
8%
10%
12%
15%
18%
20%
_whole_chrom
log10_2_3
log10_3_4
log10_4_5
log10_5_6
log10_6_7
log10_7_8
log10_8_9
_inter_chrom
Deletion
Inversion
Duplication
Translocation

Cancer processes Weights for TCGA-AR-A24Z

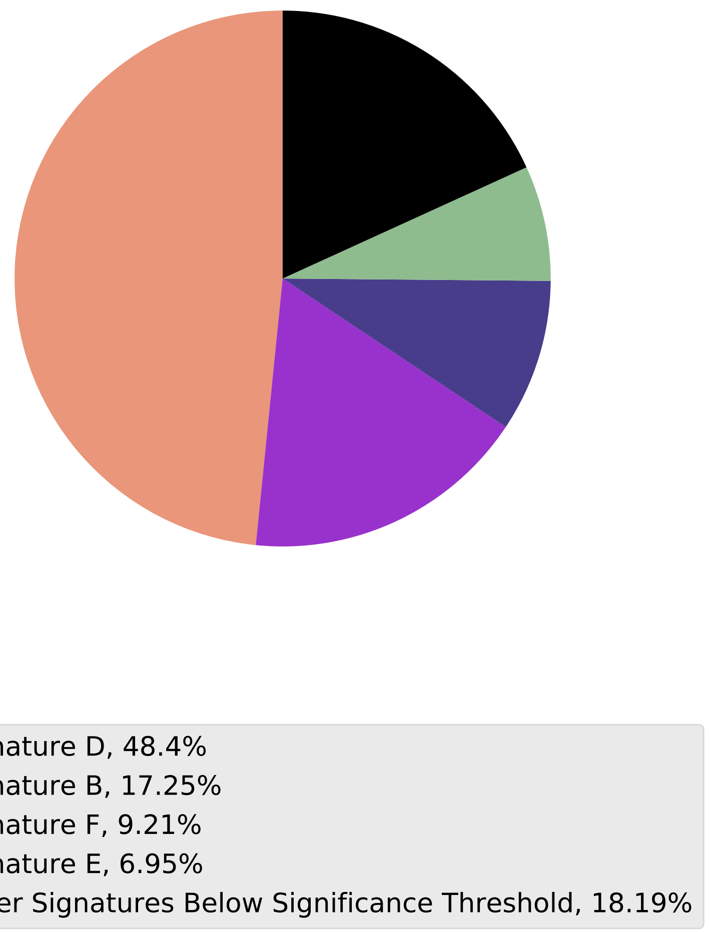

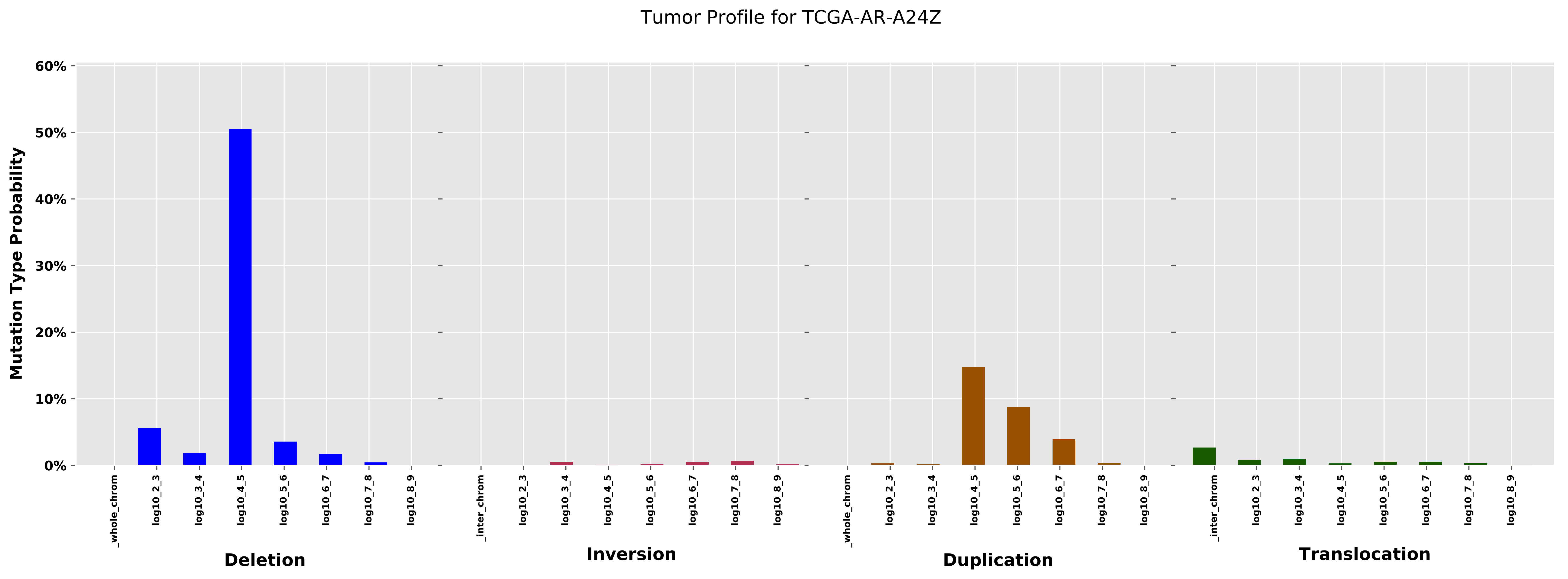

Tumor Profile for TCGA-AR-A24Z
Mutation Type Probability
60%
50%
40%
30%
20%
10%
0%
_whole_chrom
log10_2_3
log10_3_4
log10_4_5
log10_5_6
log10_6_7
log10_7_8
log10_8_9
Deletion
_inter_chrom
log10_2_3
log10_3_4
log10_4_5
log10_5_6
log10_6_7
log10_7_8
log10_8_9
Inversion
_whole_chrom
log10_2_3
log10_3_4
log10_4_5
log10_5_6
log10_6_7
log10_7_8
log10_8_9
Duplication
_inter_chrom
log10_2_3
log10_3_4
log10_4_5
log10_5_6
log10_6_7
log10_7_8
log10_8_9
Translocation

Cancer processes Weights for TCGA-A2-A04P

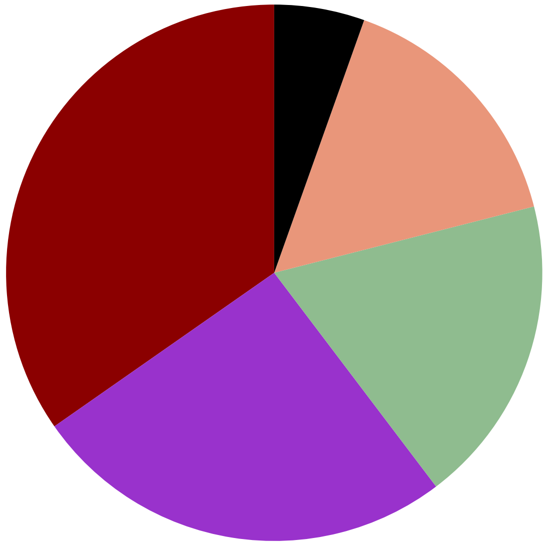

- Signature C, 34.71%
- Signature B, 25.61%
- Signature E, 18.66%
- Signature D, 15.58%
- Other Signatures Below Significance Threshold, 5.45%

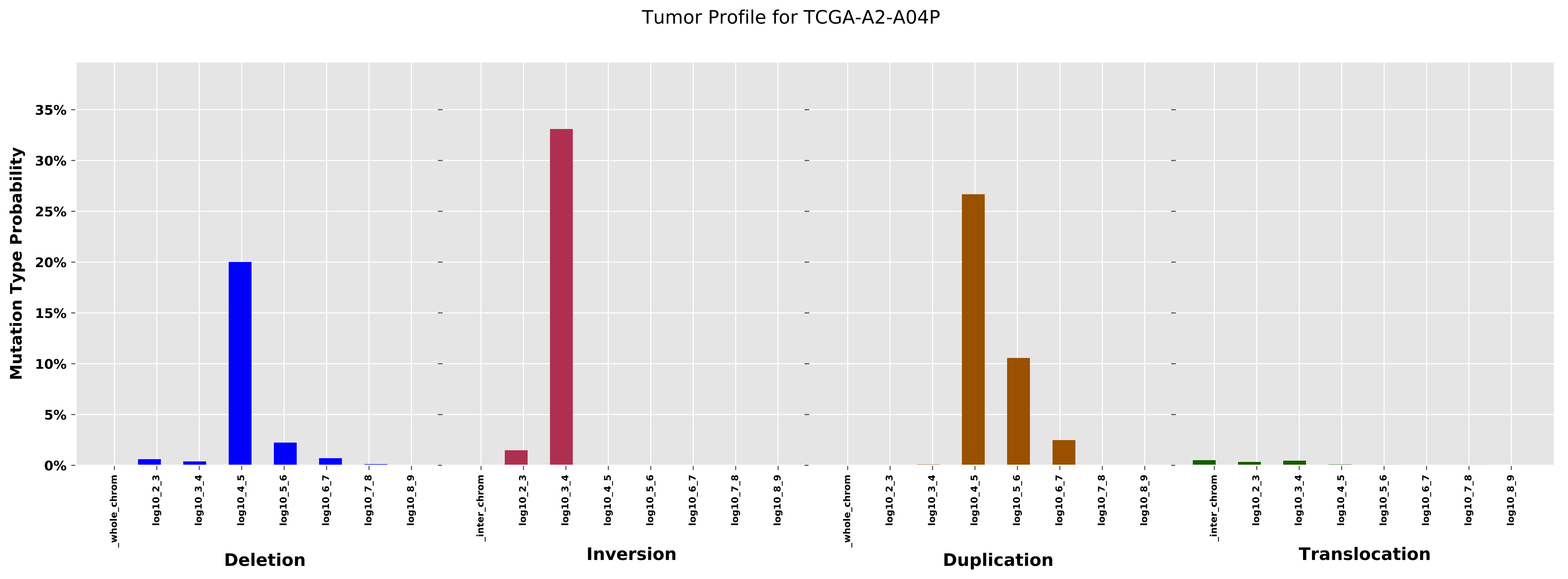

Tumor Profile for TCGA-A2-A04P
Mutation Type Probability
35%
30%
25%
20%
15%
10%
5%
0%
_whole_chrom
log10_2_3
log10_3_4
log10_4_5
log10_5_6
log10_6_7
log10_7_8
log10_8_9
_inter_chrom
Deletion
Inversion
Duplication
Translocation

Cancer processes Weights for TCGA-CA-6718

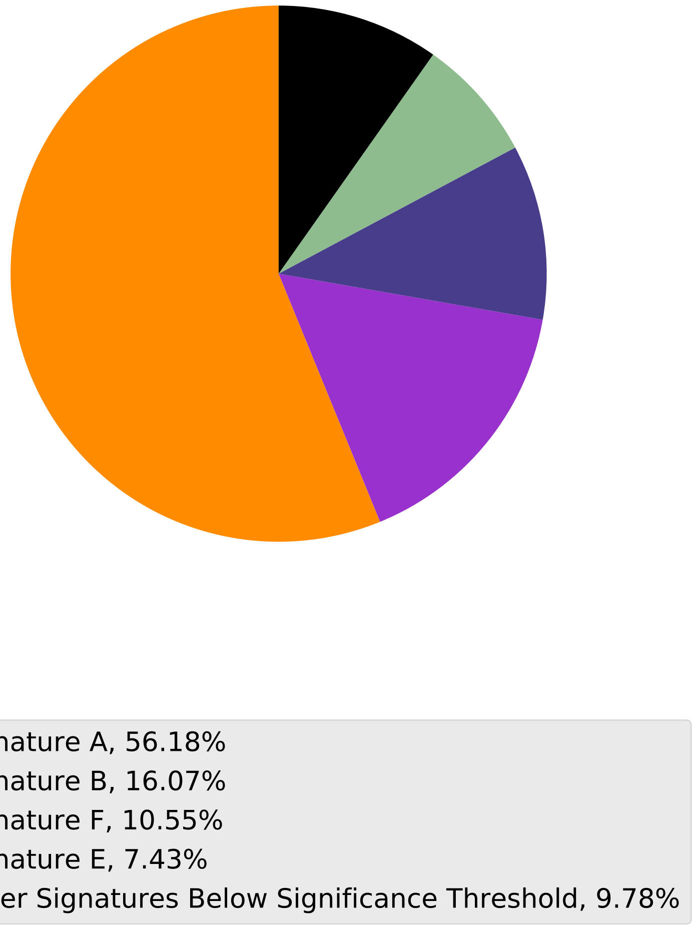

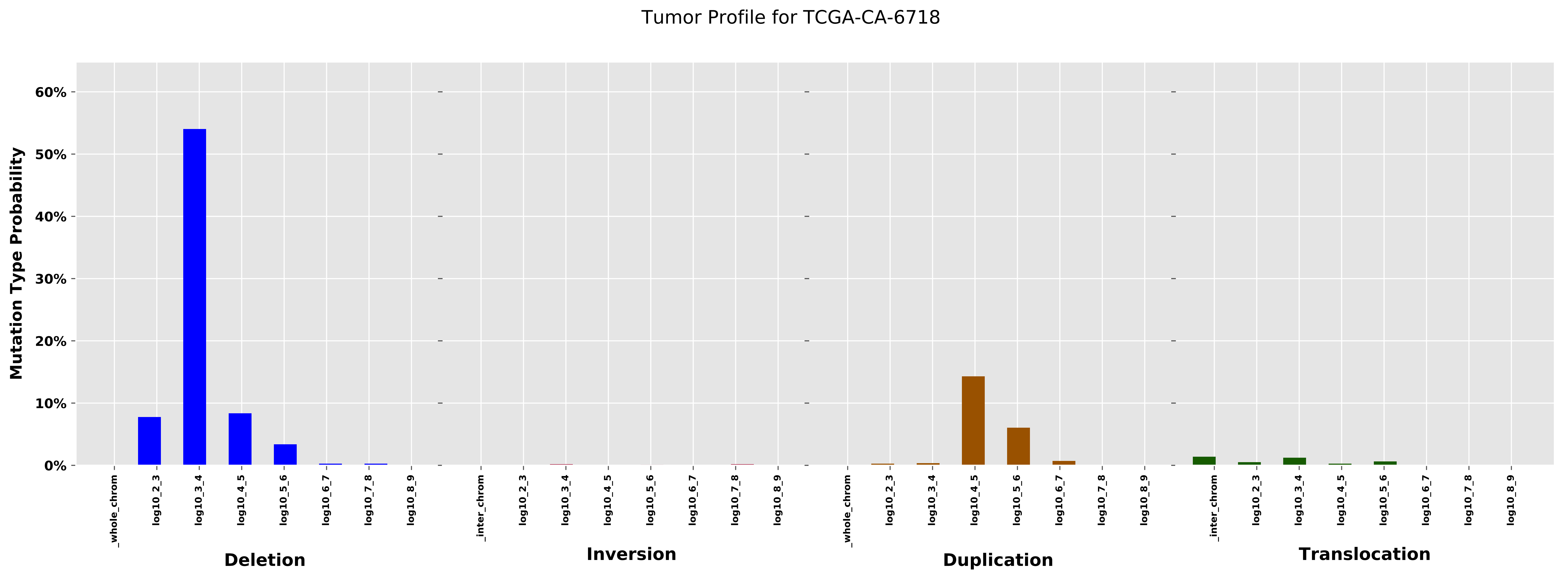
Tumor Profile for TCGA-CA-6718
Mutation Type Probability
0%
10%
20%
30%
40%
50%
60%
_whole_chrom
log10_2_3
log10_3_4
log10_4_5
log10_5_6
log10_6_7
log10_7_8
log10_8_9
_inter_chrom
Deletion
Inversion
Duplication
Translocation

Cancer processes Weights for TCGA-BH-A0AV

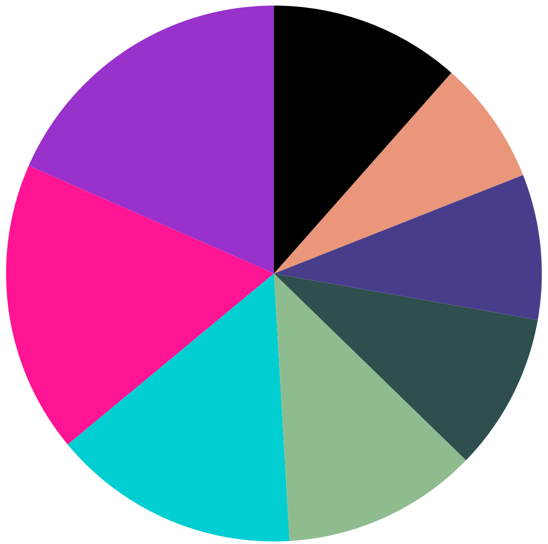

| |
|---|
| Signature B, 18.41% |
| Signature K, 17.61% |
| Signature I, 14.91% |
| Signature E, 11.8% |
| Signature G, 9.5% |
| Signature F, 8.77% |
| Signature D, 7.48% |
| Other Signatures Below Significance Threshold, 11.52% |

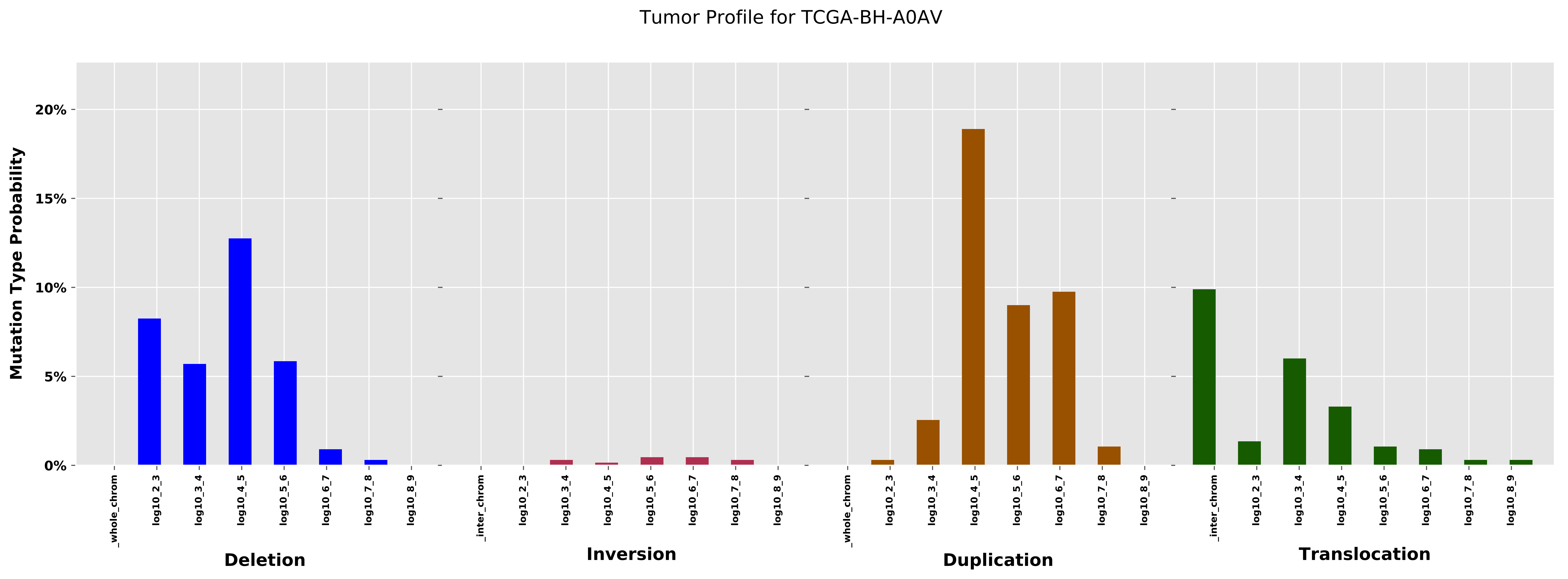
Tumor Profile for TCGA-BH-A0AV
Mutation Type Probability
20%
15%
10%
5%
0%
_whole_chrom
log10_2_3
log10_3_4
log10_4_5
log10_5_6
log10_6_7
log10_7_8
log10_8_9
_inter_chrom
Deletion
Inversion
Duplication
Translocation

Cancer processes Weights for TCGA-A2-A0CM

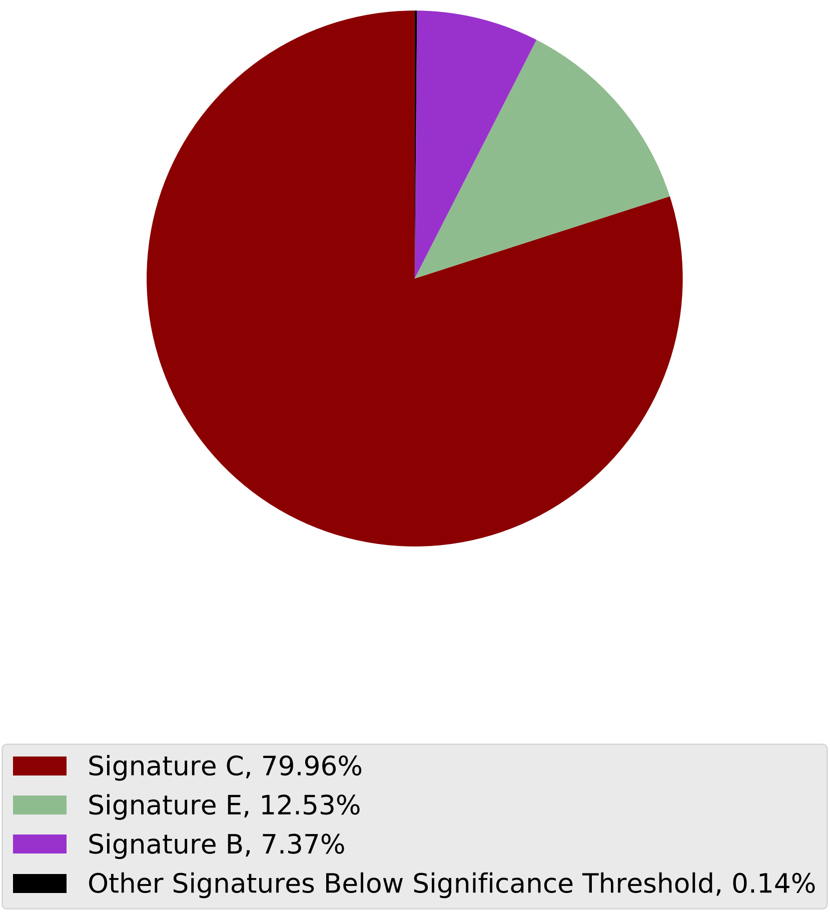

Tumor Profile for TCGA-A2-A0CM

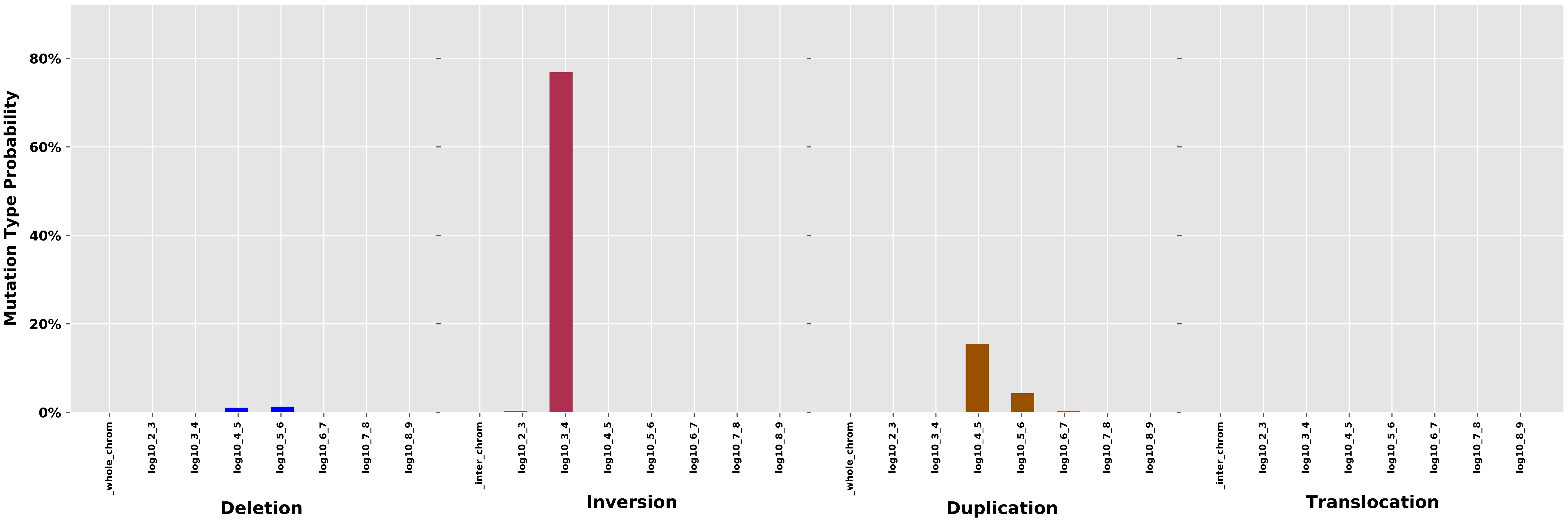

Cancer processes Weights for TCGA-AN-A04D

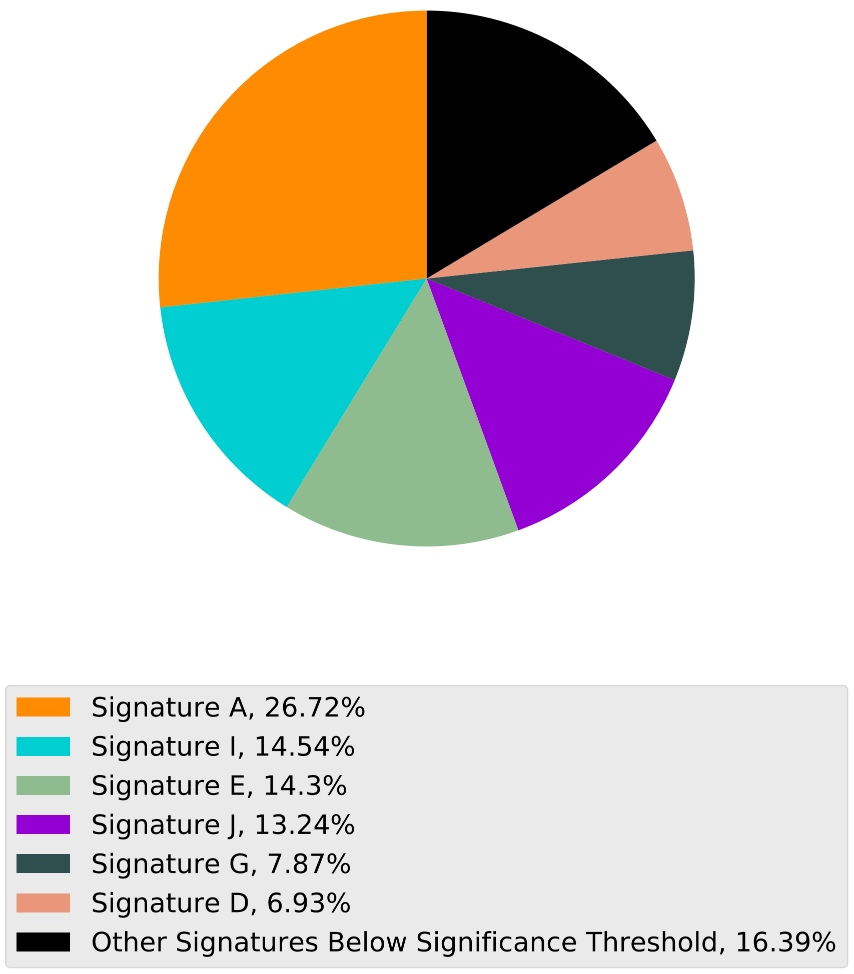

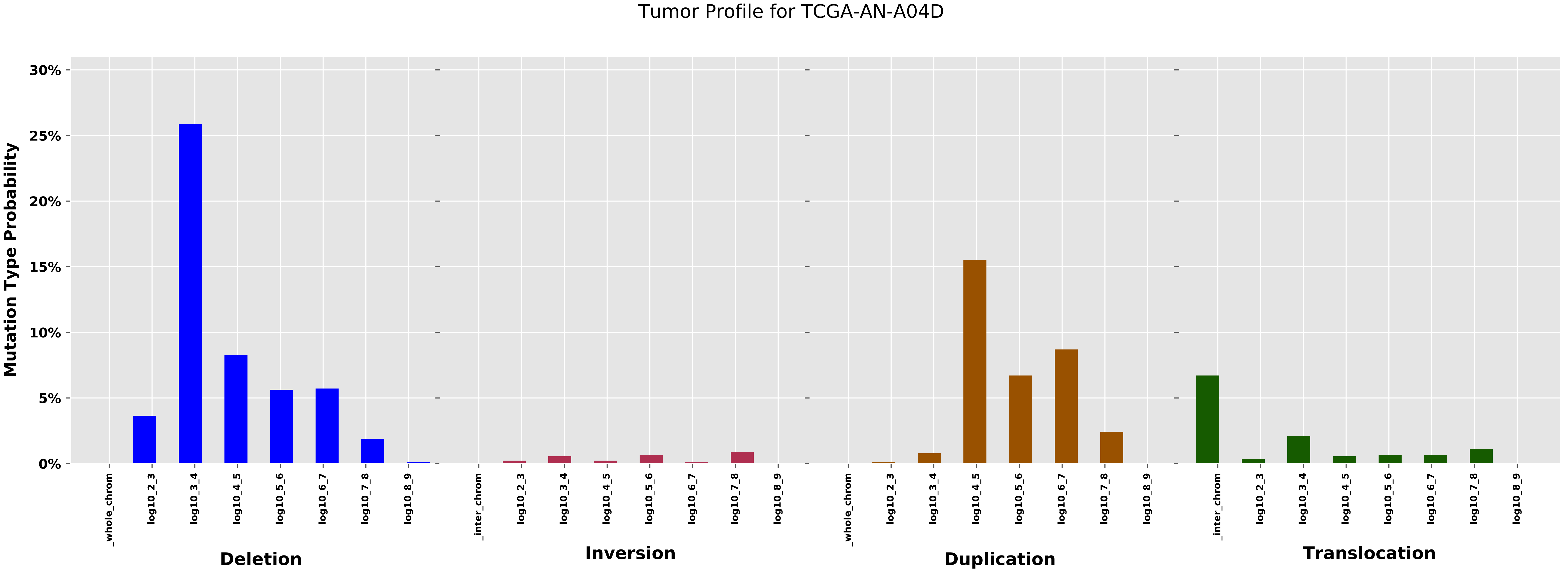
Tumor Profile for TCGA-AN-A04D
Mutation Type Probability
30%
25%
20%
15%
10%
5%
0%
_whole_chrom
log10_2_3
log10_3_4
log10_4_5
log10_5_6
log10_6_7
log10_7_8
log10_8_9
_inter_chrom
Deletion
Inversion
Duplication
Translocation

### Cancer processes Weights for TCGA-A8-A092

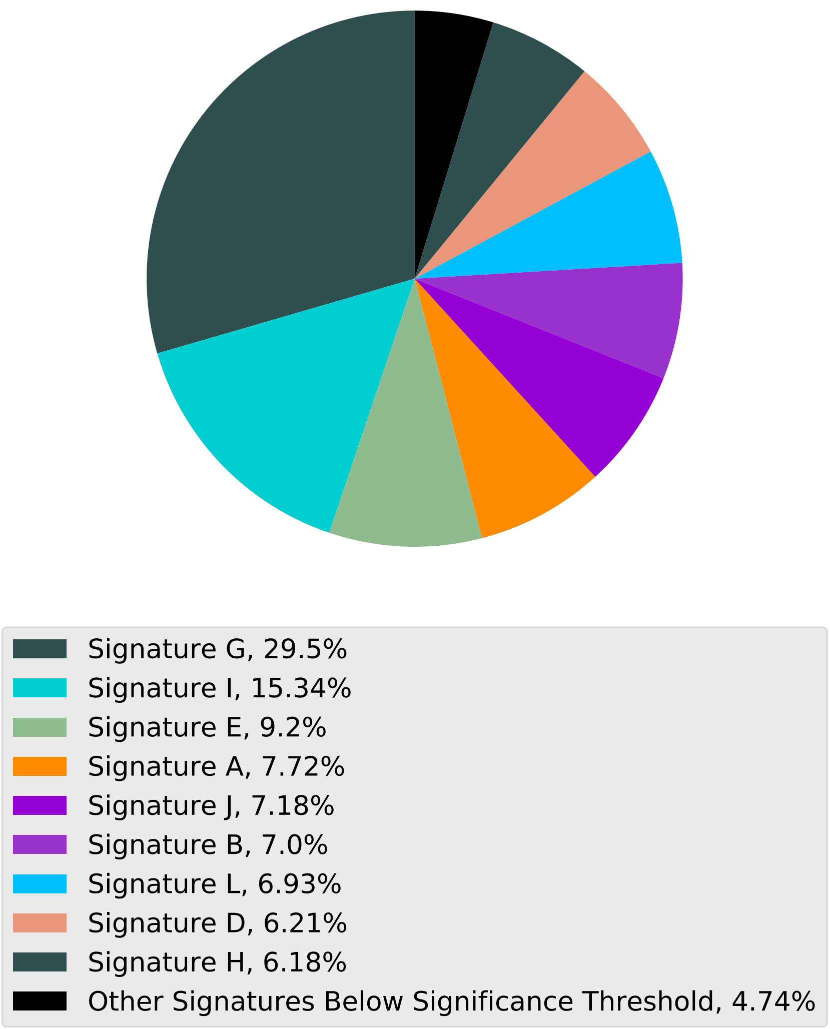

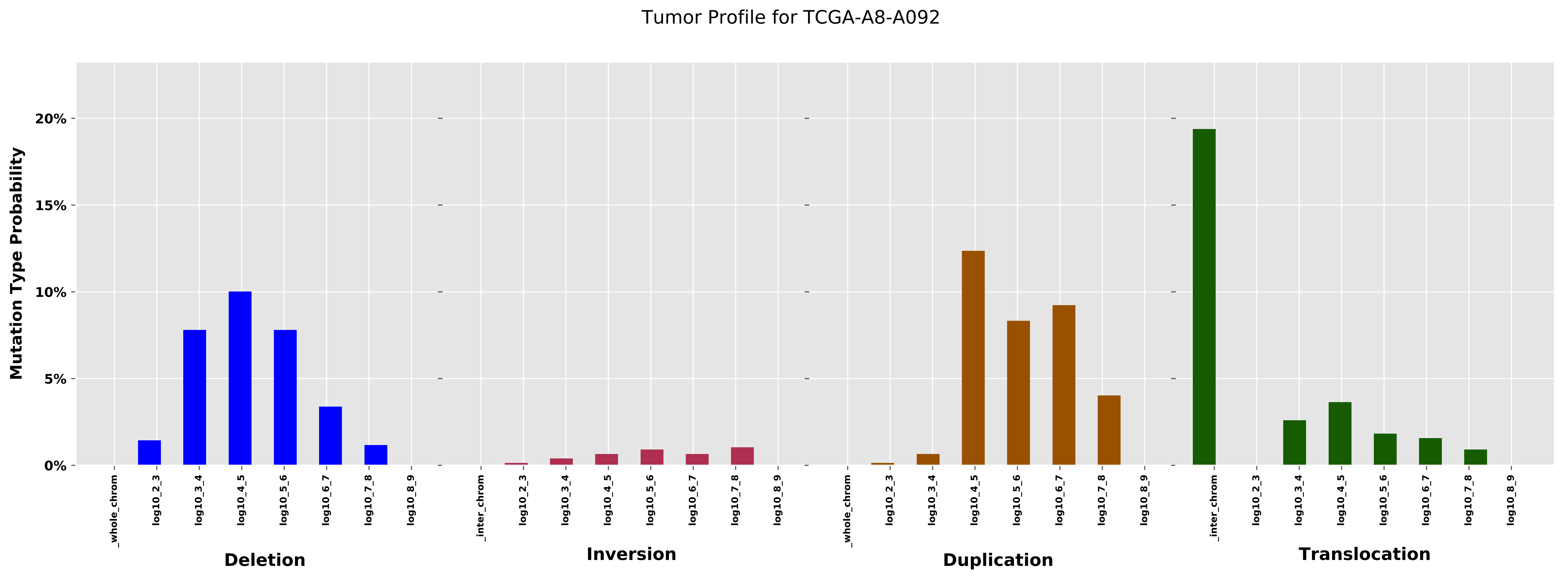

Tumor Profile for TCGA-A8-A092
Mutation Type Probability
20%
15%
10%
5%
0%
_whole_chrom
log10_2_3
log10_3_4
log10_4_5
log10_5_6
log10_6_7
log10_7_8
log10_8_9
_inter_chrom
Deletion
Inversion
Duplication
Translocation

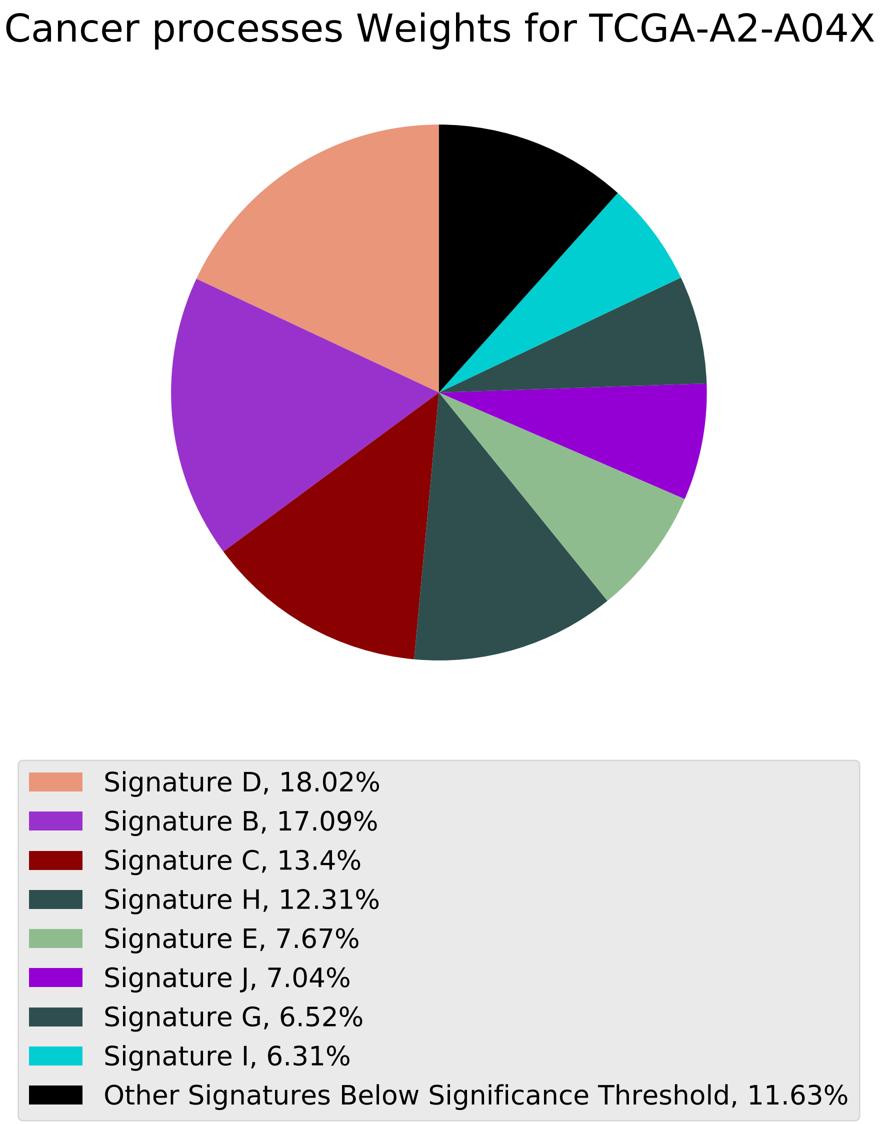
Cancer processes Weights for TCGA-A2-A04X
Signature D, 18.02%
Signature B, 17.09%
Signature C, 13.4%
Signature H, 12.31%
Signature E, 7.67%
Signature J, 7.04%
Signature G, 6.52%
Signature I, 6.31%
Other Signatures Below Significance Threshold, 11.63%

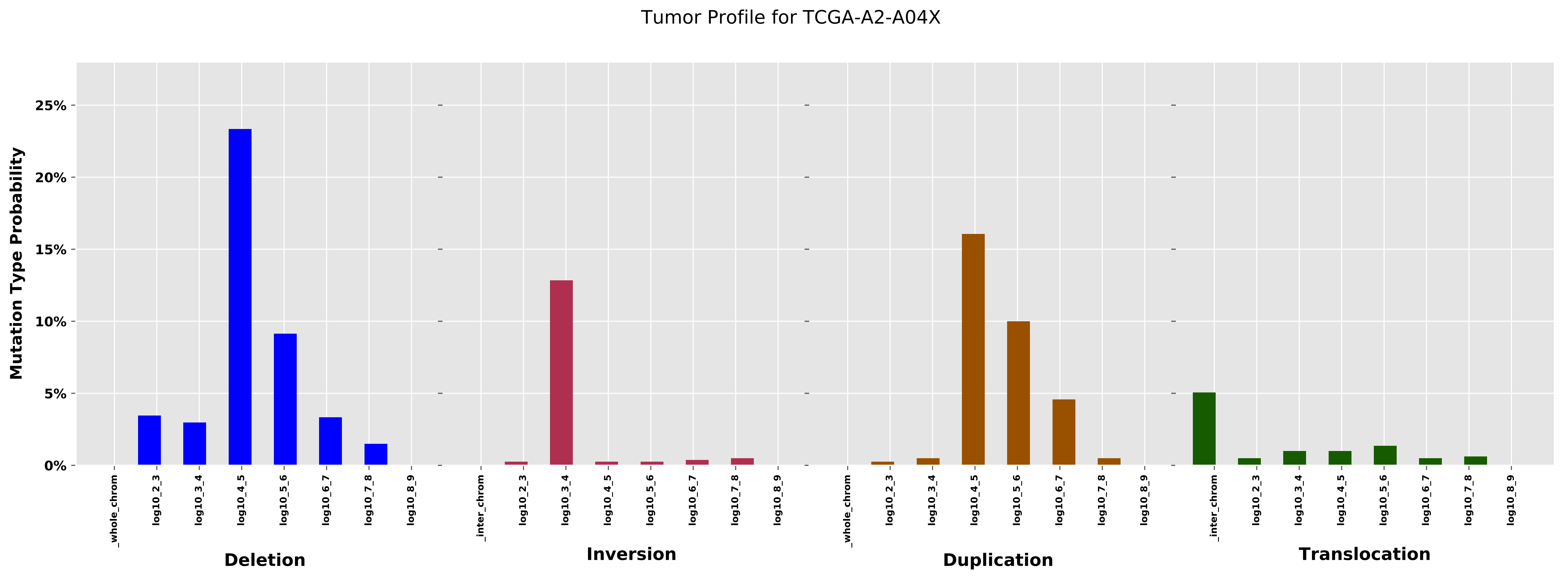
Tumor Profile for TCGA-A2-A04X
Mutation Type Probability
25%
20%
15%
10%
5%
0%
_whole_chrom
log10_2_3
log10_3_4
log10_4_5
log10_5_6
log10_6_7
log10_7_8
log10_8_9
_inter_chrom
Deletion
Inversion
Duplication
Translocation

Cancer processes Weights for TCGA-A2-A0D2

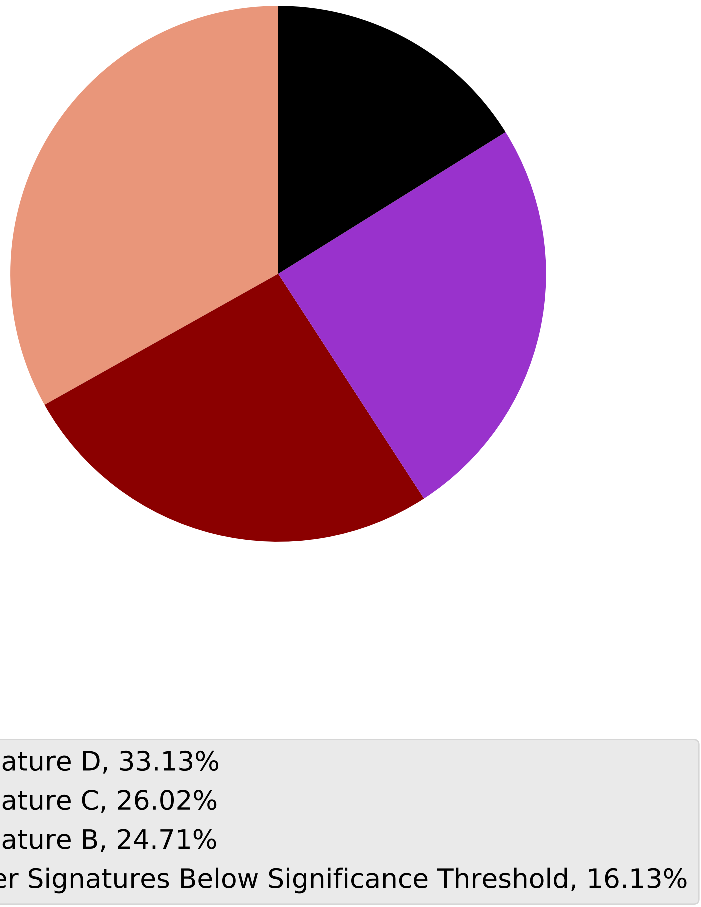

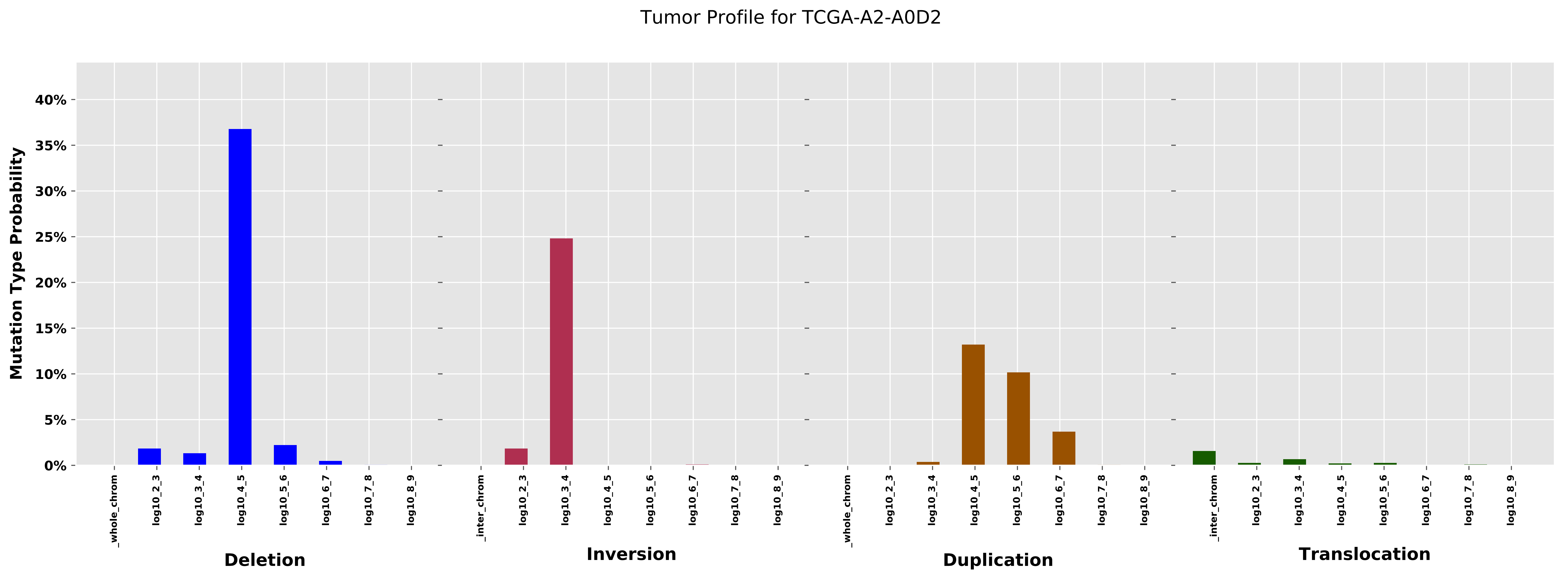

Tumor Profile for TCGA-A2-A0D2
Mutation Type Probability
40%
35%
30%
25%
20%
15%
10%
5%
0%
_whole_chrom
log10_2_3
log10_3_4
log10_4_5
log10_5_6
log10_6_7
log10_7_8
log10_8_9
Deletion
_inter_chrom
Inversion
Duplication
Translocation

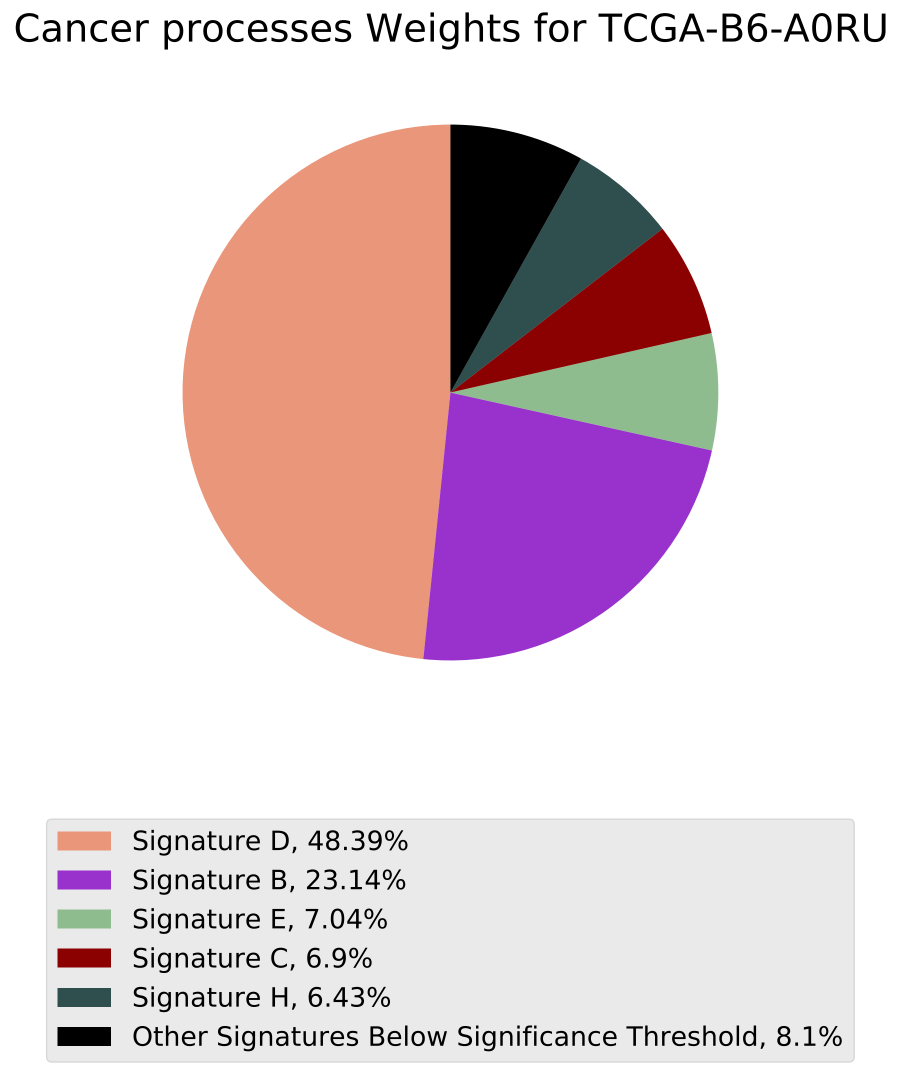
Cancer processes Weights for TCGA-B6-A0RU
Signature D, 48.39%
Signature B, 23.14%
Signature E, 7.04%
Signature C, 6.9%
Signature H, 6.43%
Other Signatures Below Significance Threshold, 8.1%

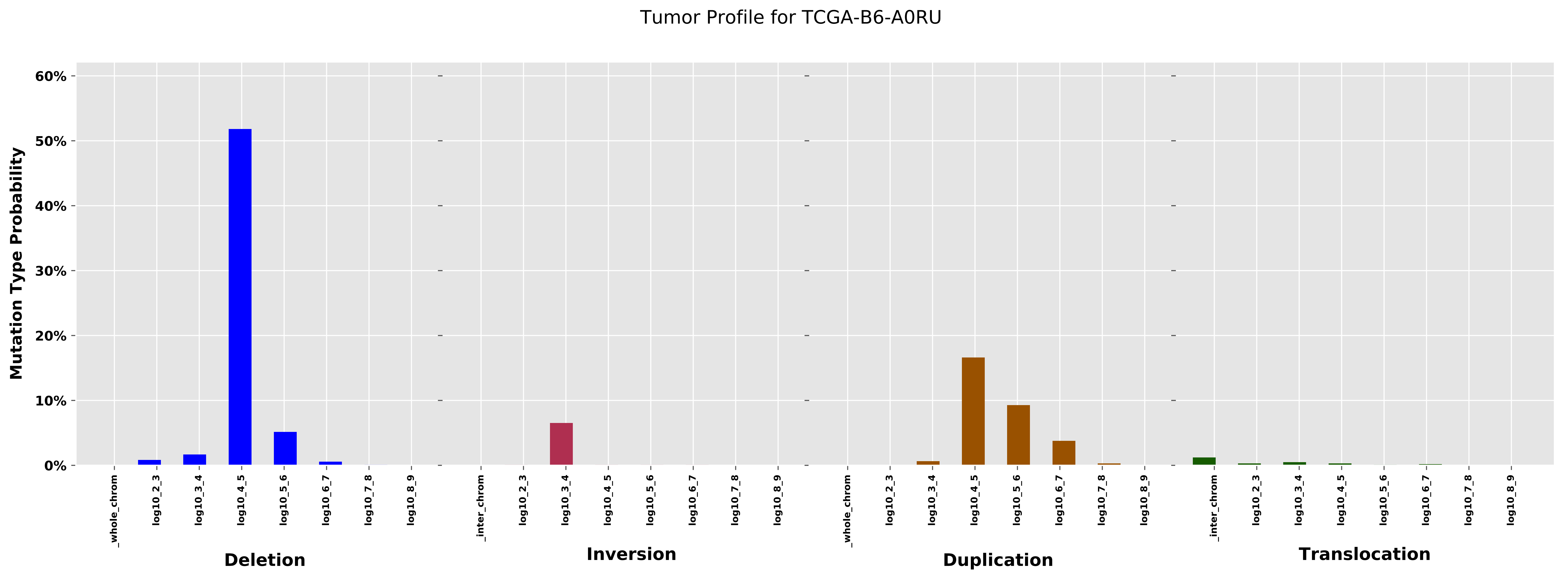

Tumor Profile for TCGA-B6-A0RU
Mutation Type Probability
60%
50%
40%
30%
20%
10%
0%
_whole_chrom
log10_2_3
log10_3_4
log10_4_5
log10_5_6
log10_6_7
log10_7_8
log10_8_9
_inter_chrom
Deletion
Inversion
Duplication
Translocation
