## Additional File 7 for "pyCancerSig: subclassifying human cancer with comprehensive single nucleotide, structural and microsatellite mutational signature deconstruction from whole genome sequencing"

### Figure 1: Combined mutational profiles of samples with highest SVB versus samples with lowest SVB

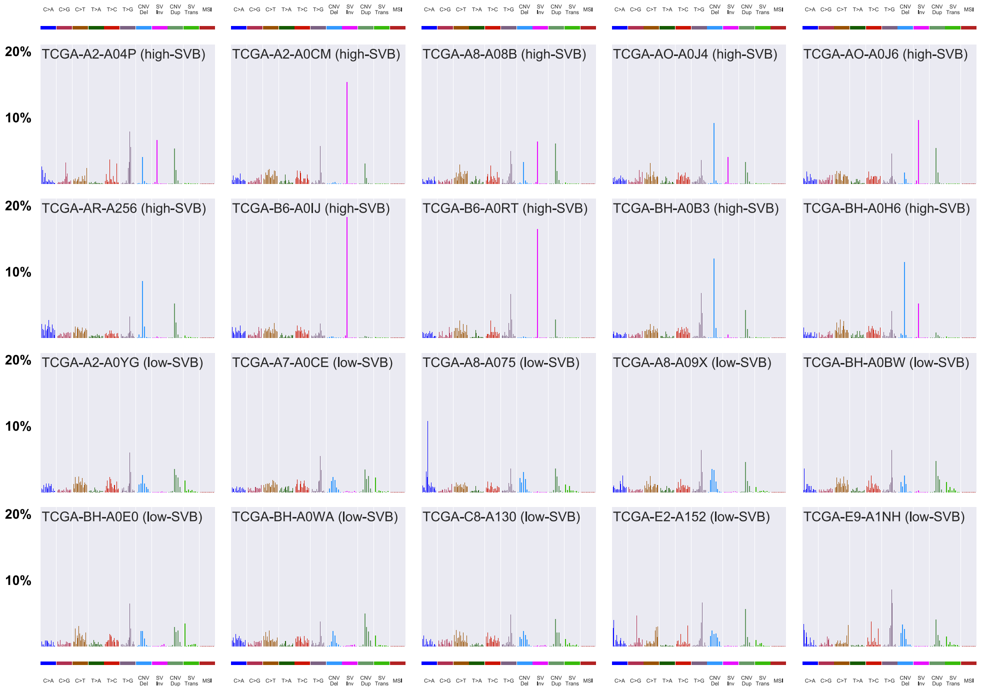


The samples in the first two rows have the highest SVB and the samples in the last two rows have the lowest SVB

### Figure 2: SV-only mutational profiles of samples with highest SVB versus samples with lowest SVB

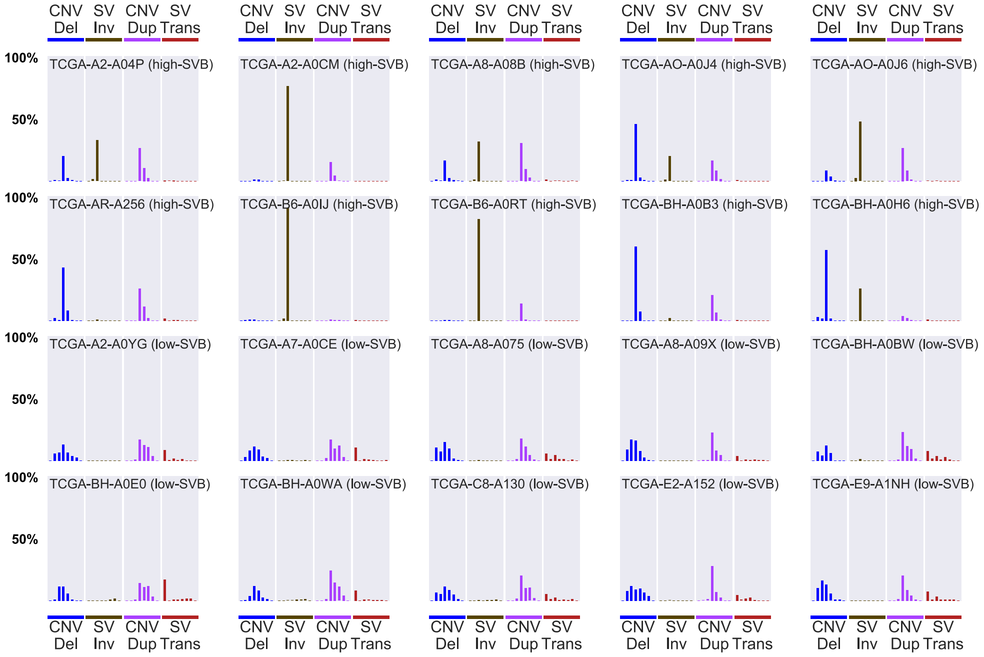


The samples in the first two rows have the highest SVB and the samples in the last two rows have the lowest SVB
